# The Genomic Basis of Electric Signal Diversity

**DOI:** 10.64898/2026.01.14.699493

**Authors:** Jason R. Gallant, Sophie Picq, Lauren Koenig, Nestor Ngoua Aba’a, Franck Nzigou, Hans Kevin Mipounga

## Abstract

Behavioral diversity is a striking result of species diversity, shaped by intertwined mechanisms, developmental pathways, and evolutionary histories. Yet the genetic underpinnings of behavioral evolution remain obscure, even though they are key to understanding how organisms adapt and diversify. Weakly electric African elephantfishes provide a rare opportunity to dissect the genetics of behavior because their electric communication signals are quantifiable, stereotyped, and directly tied to identifiable cellular mechanisms. Among elephantfishes, electric organ discharges (EODs) vary widely across species and populations, shaping mate choice, social interactions, and reproductive isolation. A key axis of this diversity is the presence or absence of an initial phase in the EOD waveform—a discrete, repeated evolutionary transition whose genetic basis has remained unresolved. Here we show, using whole-genome resequencing, population-genomic analyses, transcriptomics, and histology from 305 specimens, that biphasic EODs in *Paramormyrops kingsleyae* have evolved repeatedly in association with different genomic loci in different populations. All associated regions are noncoding, some show signatures of selection, and several are linked to differences in electric organ gene expression or protein localization, consistent with cis-regulatory change. Some associated haplotypes also occur in triphasic populations or across *Paramormyrops*, suggesting that part of this variation is old. These repeated transitions followed different genetic routes, suggesting that the developmental program underlying electric signaling contains multiple points of regulatory sensitivity. Distinct genetic changes can therefore produce the same behavioral phenotype, enabling repeated evolutionary transitions even within a single species. More broadly, our results show how developmental architecture shapes the evolutionary pathways available to behavior, helping explain why communication signals in elephantfishes—and in other radiations—diversify so rapidly.

---

Species radiations are defined by divergence, yet the same phenotype often arises repeatedly among their constituent lineages ^1–5^. That repetition is what makes them powerful for genetics: repeated origins provide multiple avenues to distinguish the loci underlying phenotypic diversity from background genetic variation^6–8^. Studies of the genetic basis of convergent evolution have contributed to an emerging perspective that adaptive traits often arise from selection on standing ancestral genetic variation rather than *de-novo* mutations^9^ and that the same or similar genetic substrates are repeatedly used in the evolution of convergent traits^10,11^. However, the generalizability of these findings remains uncertain^12^, as many studies focus primarily on morphological traits, particularly coloration and pigmentation. A central question is therefore whether repeated origins of a trait, especially within a species, reuse the same genomic regions or follow different genetic routes.

While the study of convergently evolved morphological traits has provided critical insights into the genetics of adaptation, much less is known about the genetic architecture of physiological and behavioral traits, despite their pivotal roles in shaping evolutionary and ecological dynamics in adaptive radiations^1,13^. Behavioral traits, especially those linked to prezygotic isolation (i.e. courtship signals), are central to species diversification and often exhibit striking patterns of convergence in adaptive radiations^5,14–17^, highlighting the role of behavioral isolation in speciation^18^. Understanding the genetic basis of courtship behavior may provide key insights into why certain groups undergo explosive radiations while others do not^19^. Despite this potential, the genetic basis of behavioral traits remains understudied, partly due to the perceived complexity of these traits, which often involve intricate anatomical and physiological underpinnings^4,8,20,21^.

African elephant fishes (Teleostei: Mormyridae) offer an exceptional system for exploring these questions. Mormyrids emit brief pulses of electricity, termed electric organ discharges (EODs), which they use for electrolocation, navigation, and communication^22^. EODs represent a diverse, easily quantifiable behavior^23^ directly relevant to speciation and adaptive evolution^16,24,25^. Mormyrid EOD waveforms are species-specific and stereotyped rather than learned, arising from anatomical and physiological variation in the electric organ^26^. EOD waveforms vary primarily in duration, polarity, and the number of phases^27–30^, and playback studies demonstrate these specific temporal features are critical for species recognition^25,31–33^. EOD duration is also seasonally sexually dimorphic in many species: mature reproductive males produce EODs considerably longer than the species-typical waveform, produced by females year-round and males outside the breeding season^28,34–36^. Nearly all mormyrid EODs can be categorized as either triphasic (TP) or biphasic (BP) (Fig. 1A), and the anatomical basis for these differences is well established: TP individuals possess electrocytes in their electric organ with penetrating stalks and anterior innervation, while BP individuals have non-penetrating stalks with posterior innervation^26,27,37,38^.

**Fig. 1.**
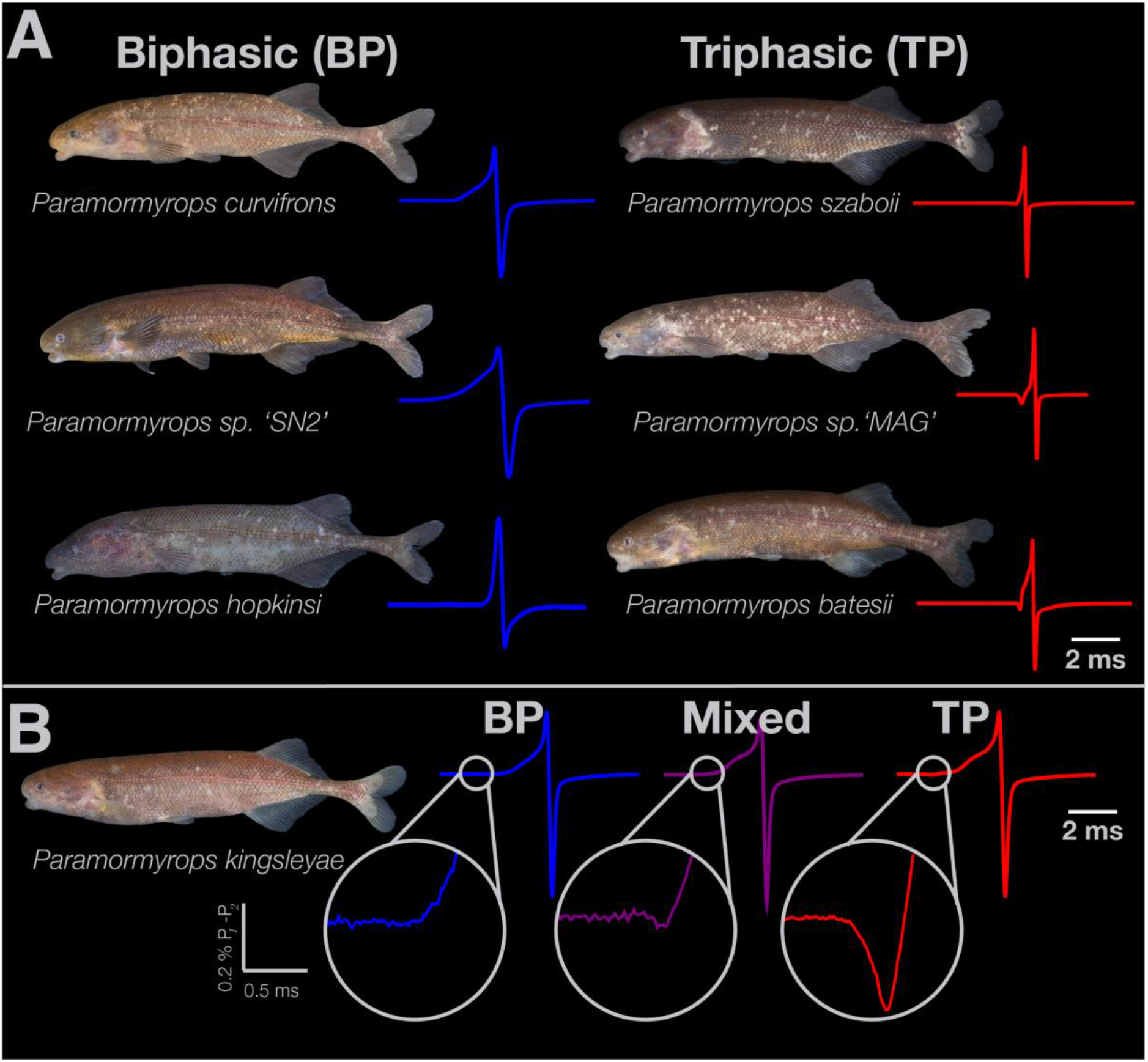
Electric signal diversity within the *Paramormyrops*. **A**, Representative *Paramormyrops* species showing variation in electric organ discharge (EOD) duration and number of phases, classified as biphasic (BP, blue traces) or triphasic (TP, red). **B,** Across Gabon, *P. kingsleyae* populations segregate BP (blue), TP (red), and ‘mixed’ (purple) EOD phenotypes; mixed individuals produce small triphasic components and possess mosaics of penetrating, anteriorly innervated electrocytes and non-penetrating, posteriorly innervated electrocytes.

While research has begun to identify expression correlates of EOD variation^39–43^, the mutational basis of EOD differences remains unknown, limiting our understanding of how electric signal evolution has progressed in this group. In this study, we leveraged EOD diversity among the rapidly diverged *Paramormyrops* to determine the genomic architecture of species signaling behavior in mormyrids. *Paramormyrops* has undergone a geographically restricted radiation of over 20 morphologically cryptic species in the river systems of West-Central Africa over the past 5–6 million years^28,29,44,45^. We recently described a polymorphism for TP and BP EOD waveforms (Fig. 1B) in the species *P. kingsleyae,* a geographically widespread member of this genus^27,32^. While most populations are fully fixed for either TP or BP waveforms, we identified a third waveform class (“mixed”) in a region in southern Gabon (Fig. 2), characterized anatomically by a mix of penetrating and non-penetrating electrocytes within a single electric organ, which earlier work interpreted as possible hybridization ^27,32^. Behavioral experiments demonstrated that *P. kingsleyae* can distinguish between BP and TP waveforms ^16^, emphasizing the potential for these traits to drive and potentially reinforce reproductive isolation^16^.

**Fig. 2.**
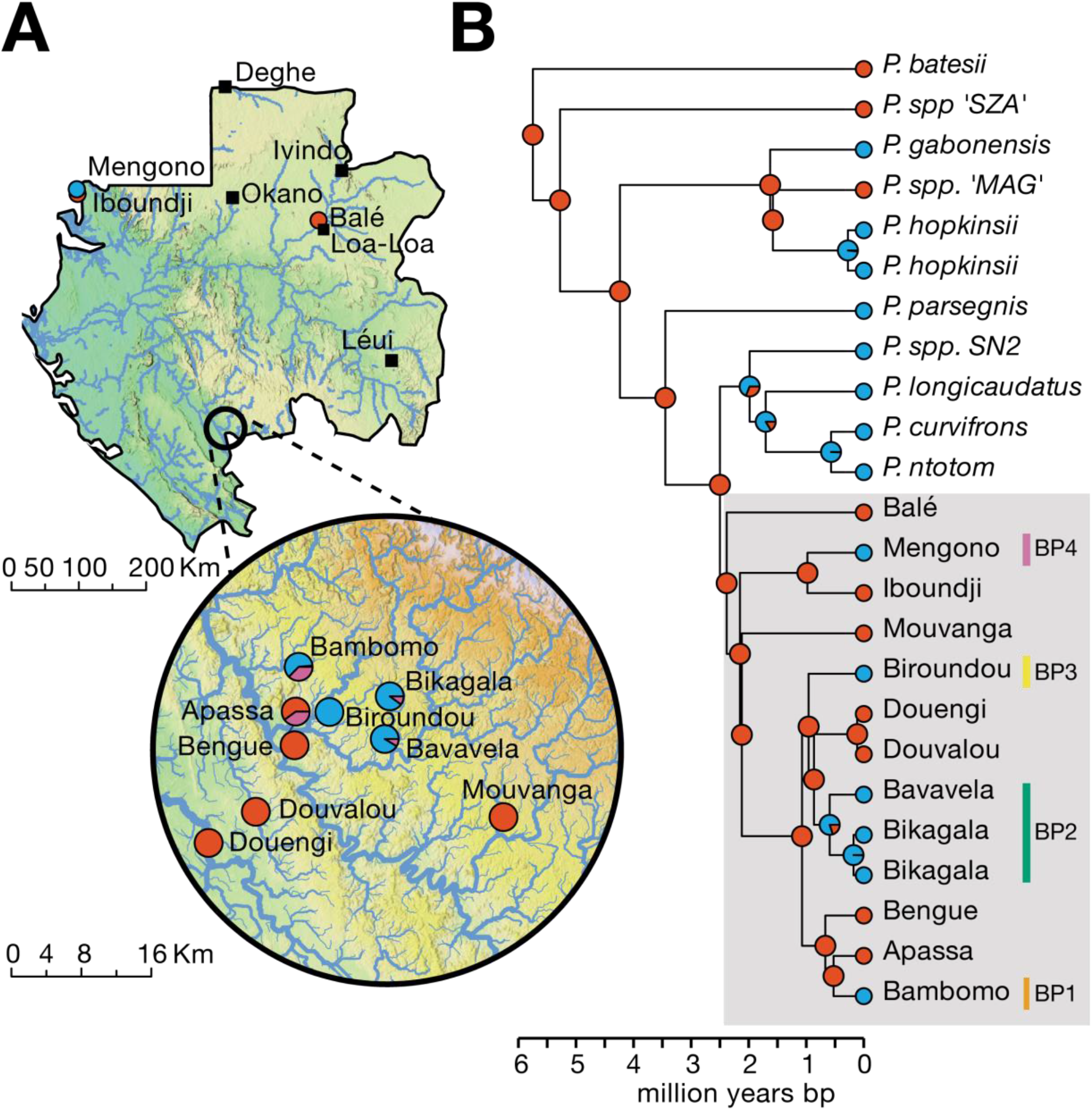
A. Geographic distribution and repeated origins of biphasic EODs. **A**, Sampling localities for *P. kingsleyae* in Gabon, West-Central Africa, with pie charts indicating the proportion of BP (blue), TP (red), and mixed (purple) EOD phenotypes at each site; black squares mark localities of other *Paramormyrops* species. A full list of specimens, along with their museum accession numbers, is provided in Supplementary Data S1. **B,** Time-calibrated species tree for *Paramormyrops* based on 25,137 maximum-likelihood gene trees, with ancestral-state reconstruction of P0 presence (TP, red) versus absence (BP, blue); pie charts at nodes show posterior probabilities. Triphasic EODs are inferred as ancestral for the genus (∼5.75 Myr ago), with eight independent origins of BP EODs (median 8, 95% HPD interval 8–10), four among other *Paramormyrops* species and four within the *P. kingsleyae* clade (∼2.5 Myr; all nodes local posterior probability >0.9999).

## New Genomic Resources Enable High-Resolution Analyses in *Paramormyrops*

We began by assembling a new chromosome-level reference genome for *Paramormyrops kingsleyae* using a single biphasic male specimen from Bambomo Creek from a combination of PacBio CCS reads and Dovetail Omni-C reads (see Materials and Methods). The final assembly measured 993 MB, with a scaffold N_50_ of 35 MB and 95.29% complete single-copy BUSCOs, representing a significant improvement in contiguity compared to our previous efforts for this species^46^. Notably, 99% of the assembled sequences were assigned to 25 autosomes (2_n_=50), consistent with karyotypes described for other *Paramormyrops* species^47^. The assembly was deposited in NCBI GenBank with accession number JBLIQM000000000 (BioProject PRJNA1218457) and annotated by the NCBI Eukaryotic Genome Annotation Pipeline (RefSeq annotation GCF_048594095.1-RS_2025_03; 24,872 protein-coding genes; see Materials and Methods).

Next, we collected EOD recordings and fin clips from 305 specimens, representing various *Paramormyrops* species and *P. kingsleyae* populations (Figs. 1, 2, Supplementary Data S1, S2). For each specimen, we extracted DNA from fin clips and performed paired-end whole genome sequencing on an Illumina HiSeq 2500 (2 × 125 or 2 × 150 bp reads; mean sequencing depth 20.0×, range 5.2–35.6×; mean aligned read depth 10.8×). Reads are available from the NCBI Sequence Read Archive under BioProject PRJNA1218444 (per-sample accessions in Supplementary Data S1). Sequenced reads were aligned to the newly assembled *P. kingsleyae* reference genome, followed by joint variant discovery and calling, yielding 51,220,152 single nucleotide polymorphisms (SNPs) with a minor allele frequency (MAF) > 0.05. Although the reference genome derives from a BP individual, read depth, the fraction of sites without reads and heterozygote allele balance were similar in BP and TP individuals, including in highly variable non-coding regions and at the GWAS peaks (Table S3; see Materials and Methods). Nucleotide diversity (π) and Tajima’s D statistic was calculated for each non-overlapping 5-kb windows across the genome for each population, while pairwise F_st_ statistics were computed for each non-overlapping 5-kb window for each pairwise population and species comparison. This analysis revealed strong population structure among *P. kingsleyae* populations, but with genome-wide F_st_ values significantly lower than those observed in pairwise comparisons between distinct *Paramormyrops* species (Fig. S1).

## Biphasic EODs Have Repeatedly Evolved Across Taxonomic Scales

Our next step was to quantify the number of independent origins of biphasic EOD waveforms across mormyrids, within the genus *Paramormyrops* and among populations of *P. kingsleyae*. Using a recent time-calibrated phylogenetic tree of Osteoglossiformes^44^, we extracted a subtree containing 106 species within the Mormyroidea where phenotypic data was publicly available (Supplementary Data S3). Using a maximum-likelihood approach, we estimated a character transition matrix based on a directional Mk model of EOD waveform evolution and generated 1,000 replicate stochastic character maps for the mormyrid phylogeny, to obtain posterior probability distributions for each ancestral node in the mormyrid phylogeny (Fig. S2). Based on this analysis, we estimate within the mormyrids that there was a single origin of biphasic waveforms from monophasic waveforms (median 1, 95% HPD interval 1-2); 10 origins of triphasic EODs from biphasic ancestors (median 10, 95% HPD interval 6-16); and 18 origins of biphasic EODs from triphasic ancestors (median 18, 95% HPD interval 11-22).

We then assessed the number of independent origins of biphasic EOD waveforms among *Paramormyrops* species and between *P. kingsleyae* populations. Drawing on our whole-genome resequencing data, we constructed 25,137 maximum-likelihood gene trees from 22 taxa and then estimated a species tree for *Paramormyrops* under the multi-species coalescent from these individual gene trees (Fig. S3). We then applied a penalized likelihood approach to obtain an ultrametric, time calibrated phylogenetic tree, using previously estimated divergence times^44^. Again, we used maximum likelihood to estimate a character transition matrix based on a directional Mk model of EOD waveform evolution. Using this model, we generated 1,000 replicate stochastic character maps for the *Paramormyrops* phylogeny to obtain posterior probability distribution for each ancestral node in the *Paramormyrops* phylogeny (Fig. 2B). Based on this analysis, we estimated that TP EODs are the ancestral state for the *Paramormyrops* genus, with 8 independent origins of biphasic EODs from triphasic ancestors (median 8, 95% HPD interval 8-10) including 4 origins of BP EODs among populations of *P. kingsleyae* (Fig. 2B).

Together, our phylogenomic analyses paint a striking picture of multiple, independent origins of BP EODs across the Mormyridae and within *Paramormyrops*, a trait with important consequence in species recognition. Finally, we show that EOD variation among populations of *P. kingsleyae* is a microcosm of this overall pattern, indicating a possible four independent origins of BP EODs (Fig. 2B; BP1-4).

## Distinct Genetic Loci Underlie Independent Origins of BP EODs in *P. kingsleyae*

Recognizing that the genome-wide phylogenomic patterns described above might not accurately reflect the underlying history of causative alleles, we leveraged the relatively low genome-wide differentiation (Fig. S1) observed between TP populations of *P. kingsleyae* in southern Gabon and three adjacent BP populations of *P. kingsleyae* to determine whether BP EODs originated independently among *P. kingsleyae* using a genome-wide association study (GWAS).

Our GWAS used a linear mixed model approach to account for relatedness, and compared BP and TP *P. kingsleyae* from all populations (Fig. 2B grey box; n=89 TP, n=109 BP individuals of 240 genotyped; 42 mixed individuals excluded, see below), and identified six narrow genomic regions (peaks 1-6) that were significantly associated with BP EODs (Fig. 3, Table S1). Remarkably, three of these regions (Peaks 1,4, and 5) exhibited genotypes that predicted BP phenotypes in Bambomo Creek (median McFadden’s pseudo R^2^ 0.73–0.94; Fig. 3C) but not in Bikagala, Bavavela or Biroundou Creeks (median McFadden’s pseudo R^2^ ≤ 0.34; Fig. 3C). Peak 1 occurs in an intergenic region near *LOC111844745*, an uncharacterized gene that is at most a putative homolog of human *C6orf132* (Fig. 3B) and exhibits a single fixed haplotypic difference (Fig. S4) between BP in Bambomo Creek and geographically proximate TP individuals. Peak 4 lies in an intergenic region between *cntnap5* and *gypc* (Fig. 3B) and displays at least four distinct haplotypes (Fig. S4) among individuals from Apassa, Bambomo, and Bengue Creeks. Similarly, Peak 5 occurs within the introns of the gene *sptbn4b* (Fig. 3B) and, also exhibits multiple haplotypes (Fig. S4). In Bambomo Creek, both Peak 4 and Peak 5 BP-associated alleles consistently occurred in one or two copies, suggesting that the causative BP allele is dominant relative to the TP-associated allele (Fig. S4). For Peaks 1, 4, and 5 the BP associated allele was only found in Bambomo Creek and not in other biphasic *P. kingsleyae* populations (i.e. Bikagala, Bavavela, and Biroundou Creeks), whereas the Bambomo Peak 1 haplotype occurs in Biroundou Creek (8 of 16 individuals), where it does not account for BP EODs. No BP-associated haplotypes for peaks 1,4, and 5 were found in other *Paramormyrops* species exhibiting biphasic EODs (Fig. S4).

**Fig. 3.**
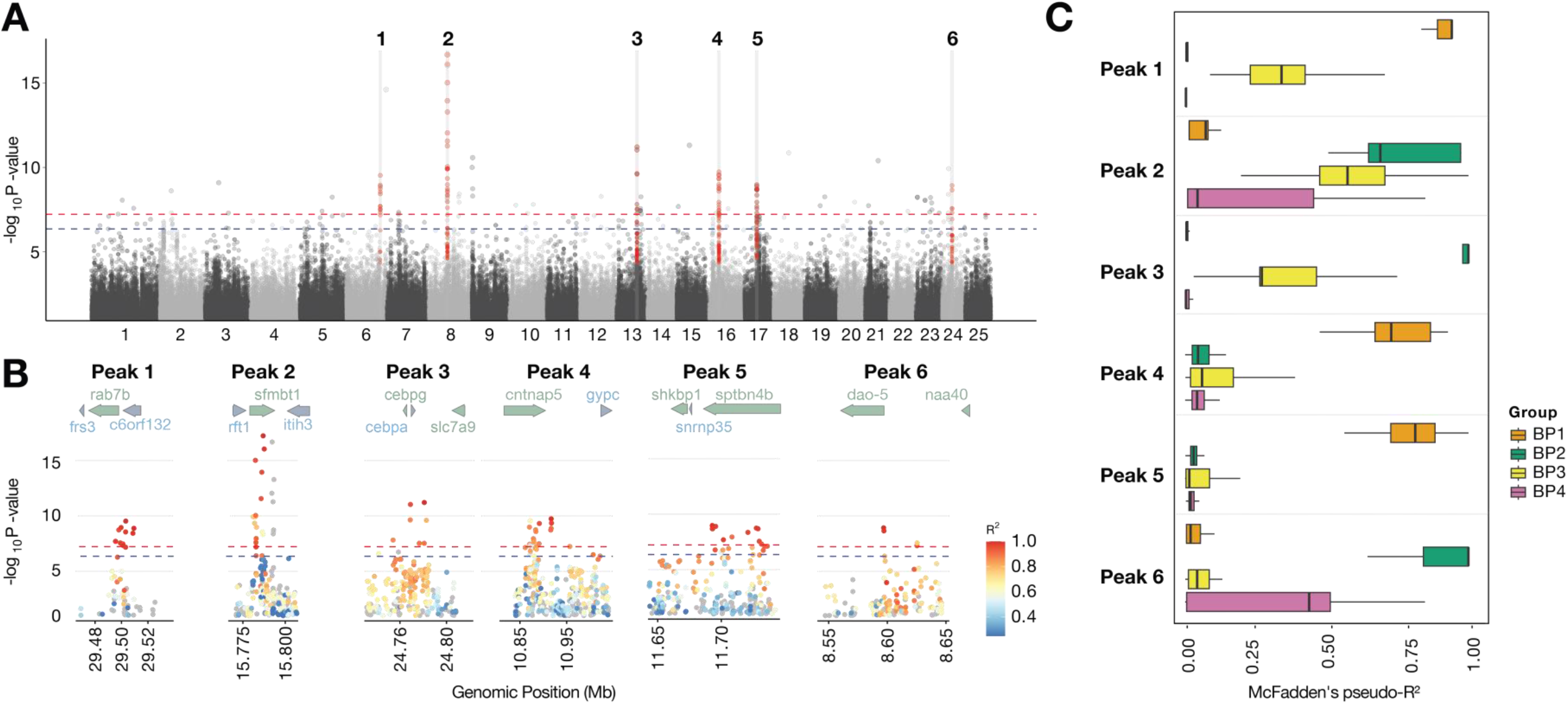
Genome-wide association mapping of biphasic versus triphasic EODs. **A**, Manhattan plot from a genome-wide association study comparing BP and TP individuals across all *P. kingsleyae* populations, revealing six narrow peaks strongly associated with BP EODs. **B,** Regional association plots for each peak; SNPs are coloured by linkage disequilibrium (R²) with the index SNP, and annotated genes are shown above each panel. **C,** McFadden’s pseudo-R² values for SNPs in high linkage disequilibrium (R² ≥ 0.6) with each index SNP, partitioned by BP lineage (BP1, Bambomo Creek; BP2, Bikagala and Bavavela Creeks; BP3, Biroundou Creek; BP4, Mengono Creek), summarizing the phenotypic variance explained by each peak in each group.

The remaining three GWAS peaks (Peaks 2, 3, and 6) predicted BP phenotypes in Bikagala, Bavavela or Biroundou Creeks (median McFadden’s pseudo R^2^≥ 0.5) but not in Bambomo Creek (median McFadden’s pseudo R^2^ ∼ 0.0, Fig. 3C). Peak 2 occurs between the genes *sfmbt1* and *itih3* (Fig. 3B), and exhibits long homozygous stretches of non-reference genotypes in Bikagala, Bavavela and Biroundou Creeks (Fig. S4). Peak 3 spans *cebpg* and *cebpa*, with its index SNP in the intergenic region 26 kb downstream of *cebpa* (Fig. 3B) and exhibits a long homozygous stretch in Bikagala and Bavavela creeks, but a distinct haplotype in Biroundou Creek. Peak 6 lies in the intergenic region between *dao-5* and *naa40* (Fig. 3B), and exhibits a long homozygous non-reference stretch found, among *P. kingsleyae*, only in Bikagala and Bavavela creeks (Fig. S4). For Peaks 2,3 and 6 the BP associated allele was only found in Bikagala, Bavavela, or in Biroundou creeks, but not in other BP *P. kingsleyae* populations (i.e. Bambomo Creek) or in other *Paramormyrops* species exhibiting BP EODs (Fig. S4), though we note that. *P.* sp. ’SN2’ carries a related haplotype with the BP allele at most SNPs but differs from every Bikagala/Bavavela copy at a block of SNPs within the peak, and *P. curvifrons* and *P. ntotom* carry the BP allele at about two-thirds of SNPs within the peak, interspersed with TP alleles (Fig. S4).

Phenotypically ‘mixed’ individuals were excluded from the GWAS but were genotyped at all six peaks using phased haplotypes (Table S2; Fig. S4; Supplemental Note). Surprisingly, mixed individuals carried the haplotypes typical of their own population: in Bambomo, 23 of 25 carried BP haplotypes at Peaks 1, 4 and 5; in Bikagala and Bavavela, all 5 carried BP haplotypes at Peaks 2, 3 and 6; but in Apassa, mixed individuals resembled TP individuals. They were also not a heterozygote class: in Bambomo, heterozygosity at Peak 4 or 5 was as common among mixed individuals (12 of 25) as among BP individuals (18 of 41; Fisher’s exact p = 0.80), consistent with dominance of the BP alleles. The two individuals with histologically confirmed mosaic electric organs came one from each background. A mixed Bambomo individual was homozygous for BP haplotypes at Peaks 1, 4 and 5, whereas the mixed Apassa individual carried one copy of the Bambomo Peak 1 haplotype but no BP haplotype at any of the other peaks.

Apassa mixed individuals had 10% or greater Bambomo-related ancestry (Fig. S8), spread across most chromosomes rather than distinct loci. This pattern is about what one Bambomo great-grandparent (12.5%) or great-great-grandparent (6.25%) would contribute: in contrast a first-generation hybrid or a first backcross (5–50% expected). A genome-wide scan within Bambomo found no locus associated with the mixed phenotype, although it had ≥80% power only for common variants (minor allele frequency ≥0.15) with per-allele odds ratios of 20–50.

## Introgression Does Not Explain the Repeated Evolution of BP EODs

Given the pervasive gene-tree/species-tree discordance in our phylogenomic analysis (Fig. S3), BP alleles could have been shared among lineages through hybridization and introgression, or inherited from a polymorphic common ancestor (incomplete lineage sorting), either among *Paramormyrops* species or among *P. kingsleyae* lineages. We tested for introgression at each GWAS peak with ABBA-BABA statistics (Patterson’s *D* and the f₄-ratio) and with the windowed statistic f_dM_, computed in Dsuite (Figs. S5, S6; Table S4; Supplementary Data S7; see Materials and Methods). Each test asked whether a BP lineage shares more derived alleles with another BP lineage than does its triphasic sister population, or more with other biphasic *Paramormyrops* species than does triphasic *P. kingsleyae* from Mouvanga, and compared each peak with the genome-wide value for the same test. The Bambomo lineage (BP1; Peaks 1, 4 and 5) showed no excess allele sharing with BP2 or BP3 at any peak. BP lineages shared excess derived alleles with other biphasic *Paramormyrops* at several peaks, but triphasic *P. kingsleyae* showed the same excess at the same peaks (Fig. S5), so this reflects the shared ancestry of these regions rather than transfer of BP alleles.

The only BP-specific signal was between BP2 and BP3 at Peak 3 (*D* = 0.18, *Z* = 2.5; *f*_dM_ above the 98th percentile of equal-sized genomic windows). At Peak 2, only 2 of 16 Biroundou fish carry the BP2 haplotype. The other Biroundou BP copies there, and all of them at Peak 3, are distinct haplotypes (Fig. S4). Divergence between BP2 and BP3 at these two peaks, scaled by divergence from the outgroup, is not reduced relative to the rest of the genome (relative node depth at the 75th–87th percentile). We therefore find no evidence that BP2 and BP3 exchanged these haplotypes recently but cannot tell whether they were shared through older gene flow or inherited from variation that predates the divergence of BP2, BP3 and their triphasic neighbors.

Taken together, these results indicate that populations of *P. kingsleyae* separated by as little as 10 km of river evolved biphasic EODs through changes at different loci, one set in Bambomo and another in Bikagala, Bavavela and Biroundou (Fig. 3), within the last 0.5–2 million years (Fig. 2).

## BP-Associated Loci Exhibit Mixed Signatures of Selection

A BP allele could have reached high frequency through selection or through drift, and we tested each in turn. We first asked whether BP-associated loci carry signatures of recent positive selection, using three statistics: haplotype length (Rsb), distortion of the site-frequency spectrum (the SweepFinder2 composite likelihood ratio, CLR) and loss of diversity (π in the BP lineage relative to π in the TP lineage). Each peak was tested in the lineage or lineages in which its BP allele arose, against the triphasic reference (BP1 vs TP1; BP2 and BP3 vs TP2). Each statistic was summarized as its most extreme window within 100 kb of the index SNP. It was then expressed as a percentile among comparable non-peak regions (see Materials and Methods). We considered a peak supported when at least two of the three statistics reached the 95^th^ percentile, and strong when one of them also reached the 99th.

Two of the six peaks were supported, both in the Bikagala and Bavavela lineage (BP2) (Fig. S7; Table S5). At Peak 6 (*naa40*), the CLR and the diversity ratio reached the 98.0th and 99.8th percentiles (empirical p = 0.007). At Peak 3 (*cebpg/cebpa*), Rsb reached the 96.7th percentile and the diversity ratio was lower than in every matched region (p = 0.019). None of the three Bambomo peaks was supported (p = 0.06–0.11), nor was Peak 2 (p = 0.056). Peak 4 came closest, with Rsb and CLR at the 92.0th and 92.9th percentiles but a diversity ratio at the 85.6th. These tests therefore support recent selection at two loci in one lineage and are inconclusive at the other four. A sweep that is old or soft, or one in a region of high recombination, may leave no trace detectable at these sample sizes. Peaks 4 and 5 carry multiple BP haplotypes, Peak 4 BP haplotypes also segregate in triphasic populations, and the BP alleles at both appear partly dominant; a sweep under these conditions leaves a weaker footprint than the hard-sweep model these statistics detect best.

We next asked whether the allele-frequency differences at BP-associated loci were unusual under neutral demographic histories. For each BP population and its TP reference, we fitted demographic models to the joint site-frequency spectrum and estimated the number of SNPs expected genome-wide to be at least as differentiated as each GWAS index SNP (Table S6). At Peaks 2–6, differences of the observed magnitude were readily produced under the fitted histories, indicating that extreme differentiation at these individual SNPs does not, by itself, require selection. Peak 1 in Bambomo was the exception: its index SNP differed in frequency by 0.99 between Bambomo and Apassa + Bengue, was more differentiated than any SNP outside the GWAS peaks, and was exceptionally unlikely under the fitted neutral histories (E = 0.04– 0.32 expected SNPs). However, allowing local reductions in effective population size increased this expectation to approximately four, leaving drift a plausible explanation. We also considered whether the clustering of highly differentiated SNPs within Peak 1 provided stronger evidence against drift. Although the region was unusual, three regions elsewhere in the genome exhibited denser clusters of comparably differentiated SNPs (empirical p = 0.04–0.10). Moreover, neutral simulations failed to reproduce the observed linkage disequilibrium in Bambomo, preventing a reliable model-based test of regional clustering (see Materials and Methods). Thus, although Peak 1 exhibits exceptional differentiation, we cannot confidently distinguish selection from drift at this locus.

Together, these analyses provide evidence consistent with recent positive selection at two BP-associated loci in the Bikagala–Bavavela lineage, but do not establish the evolutionary processes responsible for differentiation at the remaining loci. Our results therefore qualify the earlier proposal that geographic isolation and drift alone explain EOD diversification in *P. kingsleyae*^27,32^ : selection appears to have contributed to at least one origin of BP, while drift remains a plausible, but unconfirmed, explanation at other loci.

## BP-Associated Alleles Reshape Gene Expression and Protein Localization

To more directly connect BP-associated loci to electric-organ phenotypes, we sequenced RNA from the adult electric organs of 39 individuals from 8 TP and BP populations (Bambomo, Apassa, Bengue, Bikagala, Biroundou, Douengi, Douvalou and Mouvanga Creeks; paired-end Illumina reads of 2 × 100 or 2 × 150 bp). Thirty-one libraries are new (NCBI Sequence Read Archive BioProject PRJNA1218444) and eight were published previously (BioProjects PRJNA417677 and PRJNA573805); per-sample accessions are in Supplementary Data S1. Reads were quantified at the gene level and analyzed in DESeq2 (see Materials and Methods). To increase power and focus on the loci implicated by our GWAS, we restricted formal differential expression tests to protein-coding genes whose annotated exons overlapped one of the six BP-associated peaks. For each peak, we contrasted expression using variance-stabilized expression values (VST) across all populations (Fig. 4).

**Fig 4.**
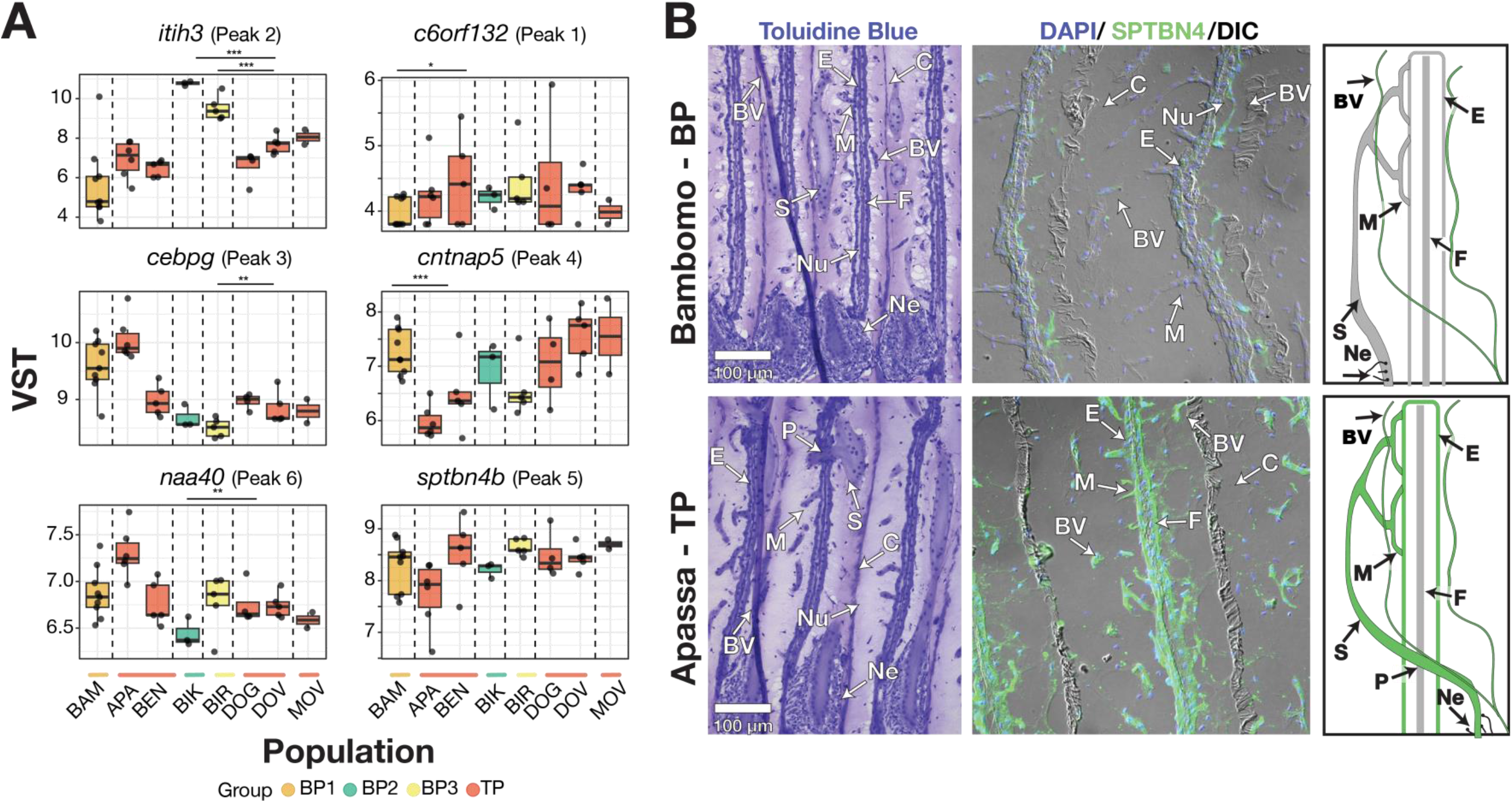
BP-associated alleles reshape gene expression and electric organ microanatomy. **A**, Variance-stabilized RNA-seq expression values for candidate genes within GWAS peaks in adult electric organs, grouped by BP lineages (BP1–BP3) and TP populations; boxplots show population-level expression differences, with *** p< 0.001; ** p<0.01 * p<0.05. **B,** Toluidine blue histology and SPTBN4 immunohistochemistry of adult electric organs from Bambomo (BP) and Apassa (TP) individuals; in TP tissue, SPTBN4 (green) labels stalks and microstalklets in penetrating stalks, which are absent from BP tissue. BV, blood vessel; S, stalk; Nu, nuclei; M, microstalklet; E, electrocyte; C, connective tissue; F, myofibril; Ne, nerve; P, penetration.

This targeted analysis revealed phenotypically associated expression shifts for a subset of genes within the peaks (Fig. 4A). Within peak 1, *LOC111844745* was barely expressed in adult electric organs (4 reads across 9 Bambomo and 38 across 11 Apassa and Bengue individuals), too few to test. The peak-4 gene *cntnap5* shows higher expression in Bambomo compared with TP1 populations (Fig. 4A). Among the loci associated with BP EODs in Bikagala, Bavavela and Biroundou, *itih3* exhibits strongly elevated expression in both BP2 and BP3 populations relative to their TP2 counterparts (Fig. 4A), whereas *cebpg* and *naa40* show more modest but significant shifts whose direction differs between BP2 and BP3 (Fig. 4A). Critically, the peak 5 gene, *sptbn4b* showed no consistent difference in expression between BP and TP groups, despite strong genetic associations (Fig. 4A).

To probe the cellular consequences of the BP-associated locus containing *sptbn4b*, we complemented the bulk RNA-seq analysis with immunohistochemistry for SPTBN4 protein in adult electric organs from representative TP and BP populations (Bambomo and Apassa Creek). SPTBN4 was prioritized for protein-level analysis because the *sptbn4b* locus shows one of the strongest and most consistent genetic associations, SPTBN4 is a key organizer of excitable membrane domains in other vertebrates, and a high-quality antibody that cross-reacts with mormyrid tissue was available (see *Materials and Methods*).

In TP electrocytes, SPTBN4 immunoreactivity is strong and highly polarized (Fig. 4B), forming continuous bands along the anterior and posterior membrane of the electrocyte and stalk regions, as well as in the surrounding vasculature. In the Bambomo (BP) individual, SPTBN4 labeling of these membranes is reduced but not absent, while labeling of the surrounding vasculature is maintained. Background-subtracted signal within the electrocyte band was 27– 75% of the Apassa level in each of four imaging sessions, and it varied among sections (one individual per population; Supplementary Methods). These protein-level differences occur in the absence of a consistent change in bulk sptbn4b transcript abundance, indicating that the BP-associated variation at this locus likely alters the spatial patterning or stability of SPTBN4 rather than its overall transcriptional level.

Together, the differential expression of several peak genes (e.g., *cntnap5*, *itih3*, *naa40*) and the altered subcellular localization of SPTBN4 at the *sptbn4b* locus are consistent with a primarily regulatory basis for BP-associated loci. All six GWAS peaks fall within intergenic or intronic regions, and the clearest expression changes involve altered transcription of genes rather than protein-coding substitutions. We emphasize that both RNA-seq and immunohistochemistry were conducted on fully differentiated adult electric organs; developmental studies in other mormyrids indicate that electrocyte penetrations are established early, when fish are ∼19 mm in length and ∼3 months of age^48^. Thus, while these differences persist into adulthood, our analysis likely misses the developmental window in which causal differences are strongest.

## BP-Associated Variants Alter Predicted Transcription Factor Binding Motifs

To investigate whether noncoding variation at BP-associated loci might affect cis-regulatory activity, we compared predicted transcription factor binding motifs between BP-and TP-associated alleles at three of our strongest GWAS peaks: Peaks 4 and 5 (BP1) and Peak 2 (BP2/BP3). We first tested whether these regions exhibited systematic differences in predicted binding potential. For 302 noncoding SNPs with −log10(p) > 3, we compared allele-specific predictions using 879 JASPAR vertebrate motifs against 10,000 sets of matched random SNPs (see Materials and Methods). Of 571 predicted allele-specific motif changes, 43% (247) were present only on the BP allele, compared with 49% among matched controls (95% range: 41–57%; p = 0.14). Neither individual peaks nor transcription factor families showed significant enrichment or consistent BP-biased motif changes (all peak p ≥ 0.14; all family q ≥ 0.36). Thus, BP-associated regions did not exhibit a systematic shift in predicted transcription factor binding potential, although individual variants could still have important regulatory effects.

We therefore used motif predictions to prioritize candidate regulatory variants within these regions. We identified 75 SNPs and indels in strong linkage disequilibrium with the GWAS index SNPs (r² ≥ 0.8), of which 39 altered at least one predicted motif by ten-fold or more between BP and TP haplotypes (best allele motif-match p < 10⁻⁴; Supplementary Data S4– S6). At Peak 5, within introns of *sptbn4b*, 26 of 47 linked variants altered predicted motifs. These included a SNP nearly perfectly linked to the index SNP (chr17:11,728,746; r² = 0.99) that introduces predicted C/EBP motifs on the BP allele, and two closely spaced SNPs (chr17:11,733,993 and 11,733,995; r² = 0.92) that produce a BP-specific BCL6/BCL6B motif. At Peak 4, 10 of 20 linked variants altered predicted motifs, including the intergenic index SNP (chr16:10,916,883), which introduces predicted IRF motifs on the BP allele. At Peak 2, three of eight linked variants altered motifs, including a SNP strongly linked to the index SNP (chr8:15,787,600; r² = 0.99) that produces a predicted BP-specific ZFP14 motif.

These variants provide candidate cis-regulatory mechanisms at loci where we independently observed altered SPTBN4 protein localization (Peak 5) or differential expression of *cntnap5* and *itih3* (Peaks 4 and 2) in BP electric organs. Although motif predictions alone cannot establish altered transcription factor binding or regulatory activity, they identify specific noncoding variants for experimental validation and offer a potential mechanistic link between BP-associated haplotypes and electric organ differentiation.

## Discussion

The repeated evolution of biphasic electric organ discharges (EODs) in *Paramormyrops* illustrates how similar changes in a communication signal can arise through distinct genetic architectures. Phylogenetic reconstruction indicates multiple losses of the initial EOD phase (Fig. 2), and our association analyses identify different sets of genomic regions linked to this phenotype in Bambomo than in Bikagala, Bavavela and Biroundou (Fig. 3). These associated regions lie on distinct haplotype backgrounds. Neither comparisons with other *Paramormyrops* species nor tests for introgression support the spread of a single BP-associated haplotype across these lineages (Figs. S4–S6), although the Bambomo Peak 1 haplotype is also found in Biroundou. Thus, while the evolutionary histories of individual associated variants are not all resolved, the repeated appearance of BP signals cannot be explained simply by the reuse of one shared genetic substrate. The same conspicuous waveform change has arisen in association with different genomic regions in different populations.

This pattern contrasts with prominent examples of repeated evolution involving recurrent use of the same loci. In freshwater sticklebacks, for example, repeated phenotypic change has involved both standing alleles at *Eda* and independent deletions affecting the same *Pitx1* enhancer^49,50^. A recent synthesis likewise suggested that repeated evolution within species often draws on shared standing ancestral variation^9^. In *P. kingsleyae*, however, the major genomic associations differ between BP origins. This does not mean that all BP-associated mutations arose de novo: the BP-associated haplotypes at Peaks 1 and 5 have not been found in sampled TP populations or related species, but many individual alleles at Peak 2 are also common in other *Paramormyrops*, including TP species, and the Bambomo-associated Peak 4 haplotype segregates at low frequency in TP Apassa and Bengue populations. The age of particular mutations therefore remains uncertain, and some associated variation may be old. What distinguishes this system is not necessarily a predominance of new mutations, but the use of different loci—and different haplotype combinations—to reach a similar phenotypic outcome. This broadens the range of genetic routes through which repeated evolution within species may occur^9,51^.

The developmental basis of the BP–TP transition suggests why several different loci could influence the same waveform feature. Mormyrid electrocytes differentiate from skeletal muscle during larval development and undergo extensive remodeling as posterior stalks form and pass through specialized penetrations in the electrocyte membrane^52^. These penetrations are necessary for the initial phase of the adult EOD; BP electrocytes lack them^37,45^. The BP phenotype can therefore be understood as the loss of a particular developmental specialization, rather than as the evolution of an entirely new cell type or electric organ. Establishing penetrating stalks requires the coordination of cell polarity, membrane organization, cytoskeletal attachments and extracellular interactions. Several genes adjacent to BP-associated regions have plausible roles in these processes, including genes associated with membrane–cytoskeleton interactions, cell adhesion and extracellular matrix organization. Related scaffolding systems maintain sharply differentiated membrane domains in neurons and muscle^53–56^, and previous comparisons of electric organs with skeletal muscle, and of species with divergent waveforms, implicate similar cellular functions^41,42,57–61^. Disturbance at more than one point in this developmental machinery could plausibly prevent penetration formation.

The molecular evidence is consistent with, but does not yet establish, a regulatory mechanism for these changes. The associated variants occur in noncoding regions; genes near several peaks show phenotype-associated differences in adult electric-organ expression, and BP and TP haplotypes differ in predicted transcription-factor binding motifs. Moreover, SPTBN4 labeling differs between TP and BP electrocytes despite the absence of a consistent bulk-transcript difference at *sptbn4b* (Fig. 4). These findings suggest several possible routes by which noncoding changes might alter the abundance, timing or spatial organization of proteins required for stalk development. Yet our expression and immunohistochemical observations come from adult organs, whereas penetrations form early in development^48^. Reduced SPTBN4 labeling at stalk membranes, for example, might be a consequence of absent penetrations rather than their cause; predicted motifs likewise do not demonstrate altered transcription-factor occupancy. Because juvenile and larval material is unavailable for these populations, establishing causal relationships will require developmental sampling and functional tests. For now, the candidate loci identify plausible points of sensitivity in a shared developmental process, not a demonstrated regulatory pathway.

Mixed-waveform individuals reveal an additional layer of complexity in that process. Where examined, their electric organs contain mosaics of electrocytes with and without penetrating stalks^27,32^. These fish do not simply carry intermediate genotypes at the major GWAS loci: mixed individuals generally carry the BP-associated haplotypes of their population in Bambomo, Bikagala and Bavavela, but TP-associated haplotypes in Apassa. They are not enriched for heterozygotes, and none is inferred to be a first-generation hybrid or first backcross. Similarly, six of 35 TP fish from Apassa and Bengue each carry a single copy of the Bambomo BP-associated haplotype at Peak 4 (*cntnap5*), even though all Bambomo BP fish carry at least one copy. Thus, the presence of a BP-associated haplotype is not invariably sufficient to produce a fully BP waveform across genetic backgrounds.

One explanation is a developmental threshold operating at the level of individual electrocytes. Under this model, major BP-associated alleles shift the distribution of liability for penetration formation, while genetic modifiers, environmental influences or developmental stochasticity determine whether particular cells cross the threshold. A mixed organ would then contain electrocytes on both sides of that threshold. Alternatively, stalk penetration and other aspects of electrocyte organization, including innervation, may be influenced by partially distinct developmental processes; different combinations of changes could yield similar mixed waveforms. The present data cannot distinguish these possibilities. Testing these models will require phenotype-balanced collections from the same sites and years, quantitative histology of whole organs and, ideally, developmental observations linking genotype to the formation of individual electrocytes.

The genetic routes by which BP phenotypes originated should also be distinguished from the processes that brought associated alleles to high frequency. Two loci in the Bikagala– Bavavela lineage (Peaks 3 and 6) have localized signatures consistent with recent selective sweeps (Fig. S7). None of the Bambomo loci meets our sweep criteria, however, and drift under the inferred demographic histories cannot be excluded even at Peak 1, the most differentiated SNP in that genome (Table S6). The evidence therefore does not support a single explanation for the spread of BP-associated alleles across populations. Selection may have contributed to some origins, while drift remains plausible at others; importantly, the observation that different loci are associated with BP signals does not depend on which process increased their frequencies.

This distinction also helps reconcile our results with earlier interpretations of EOD variation in *P. kingsleyae*. The original biogeographic hypothesis emphasized drift in geographically isolated populations^27^, whereas later work documented stable allele frequencies over roughly a decade and little correspondence between signal differences and neutral genetic distance^32^. Neither observation decisively rules out historical drift at loci of large effect. Stability over recent generations does not describe effective population sizes when BP-associated alleles initially rose, and a lack of correlation between waveform and genome-wide genetic distance is not unexpected when a phenotype is influenced by a few large-effect loci. Our coalescent estimates indicate substantial historical reductions in effective population size in all four BP populations (Fig. S9), potentially increasing the efficacy of drift during the relevant period. Thus, the earlier drift hypothesis remains viable where sweep tests are inconclusive, while the evidence at Peaks 3 and 6 is inconsistent with an exclusively neutral explanation.

Together, these findings point to developmental architecture as a potential determinant of how readily electric signals evolve repeatedly. The formation of penetrating stalks is a specialized developmental process that depends on the coordinated organization of membranes, cytoskeletal scaffolds, and extracellular interactions. Establishing this architecture may require many components to function together, whereas disrupting any one of several components could prevent its formation. The developmental complexity of penetrating electrocytes may therefore create a broad mutational target^62^, providing multiple genetic routes to the loss of the initial EOD phase. This asymmetry offers a possible explanation for the repeated evolution of BP signals: although the developmental machinery required to produce a triphasic discharge is complex, there may be many comparatively accessible ways to disrupt it.

This perspective has broader implications for understanding how behavior evolves. Behavioral evolution is often considered in terms of changes to neural circuits, sensory processing, or the mechanisms that generate new patterns of motor activity. Yet behavioral outputs also depend on the anatomical and physiological machinery through which neural activity is expressed. In electric fish, the EOD is the summed electrical output of electrocytes, so changes in their architecture directly alter the signal the animal produces^23^, without necessarily requiring changes to the neural circuitry that activates them. The loss of a specialized cellular structure can thus generate a distinctive communication signal, illustrating how evolutionary novelty can arise through the disruption of an existing developmental program rather than the construction of a new one. Such changes may have consequences beyond communication: because the EOD also supplies the electric field used for active electrolocation^22^, loss of its initial phase may alter sensory performance, although this remains untested. More generally, the evolutionary accessibility of behavioral phenotypes may depend not only on the strength of selection acting on them, but also on how readily their underlying developmental and physiological machinery can be modified. In this sense, repeated behavioral evolution need not reflect the reuse of favored genetic variants or even the same genes, but the existence of multiple ways to modify a shared biological system. Our findings evoke the opening line of *Anna Karenina*: all triphasic electrocytes are alike; each biphasic electrocyte is biphasic in its own way.

## Acknowledgments

Permits to collect fishes in Gabon and export them for this study were granted by l’Institut de Recherche en Ecologie Tropicale, l’Institut de Recherches Agronomiques et Forestières, and the Centre National de la Recherche Scientifique et Technologique. We are grateful for the valuable assistance and logistical support we received from J. D. Mbega and Jean Hervé Mve Beh, as well as students working in these institutions. Additionally, we thank C. D. Hopkins, C. Dillman, and J.P. Sullivan for their assistance in identifying specimens used from the Cornell Museum collections. As part of this analysis, we received media from the Macaulay Library at the Cornell Lab of Ornithology.

## Author contributions

Conceptualization, Supervision, Funding Acquisition: JRG, Methodology: JRG, SP, LK, NN, FN, KHM, Formal Analysis: JRG, Writing – original draft: JRG, Writing – review & editing: JRG, SP, LK, NN, FN, KHM

## Data and materials availability

Whole-genome resequencing reads and new electric organ RNA-seq reads are in the NCBI Sequence Read Archive under BioProject PRJNA1218444. The reference genome assembly is in GenBank under accession JBLIQM000000000 (BioProject PRJNA1218457); its RefSeq assembly and annotation are GCF_048594095.1. Previously published electric organ RNA-seq reads are under BioProjects PRJNA417677 and PRJNA573805. Per-sample accessions are listed in Supplementary Data S1. All other data are available in the main text or the supplementary materials.

## Code availability

All code used for the analyses in this study is available at https://github.com/msuefishlab/pkings_eod_gwas.

## Supplementary Materials

Materials and Methods

Supplementary Note

Figs. S1 to S9

Tables S1 to S6

References

Data S1 to S7

## Materials and Methods

### Field collections

Mormyrid specimens were collected in Gabon, West Central Africa, during field expeditions spanning 1998 to 2023. A comprehensive catalog of these specimens, including metadata and voucher accession numbers for the MSU Museum and the Cornell Museum of Vertebrates is provided in Supplementary Data File S1. Specimens were collected using hand nets and electric fish detectors, or through controlled light applications of rotenone. In cases where rotenone was used, fish were immediately transferred the fish to fresh, aerated water to allow for complete recovery. *Paramormyrops kingsleyae* exhibits sexual dimorphism in electric organ discharge (EOD) signals during the breeding season, with mature males producing EODs that are 2-3 times longer than those of females and non-breeding males. Our study focuses exclusively on the species-typical waveform: the EOD that females produce throughout the year and that males produce outside the breeding season.

### EOD recordings

EODs were recorded within 24 hours of capture in 1-to 5-liter plastic containers filled with water from the collection site. Signals were detected using bipolar chloride-coated silver wire electrodes, amplified (bandwidth = 0.0001-50 kHz) with a differential bioamplifier (CWE, Inc., Ardmore, PA), and digitized at sampling rates ranging from 100 kHz to 1 MHz. Head positivity was plotted upward using either a Daqbook or WaveBook (IOTECH, Cleveland, OH) or a USB-powered A-D converter (National Instruments, Austin, TX).

Each recording maintained a 16-bit vertical resolution, and multiple EOD waveforms were collected per specimen. Following EOD recording, specimens were euthanized with an overdose of MS-222, and tissue samples were preserved in 95% ethanol. Each specimen was assigned a unique identification tag and fixed in 10% phosphate-buffered formalin (pH 7.2) for at least two weeks before being transferred to 70% ethanol for long-term storage. Specimens were deposited in the Cornell University Museum of Vertebrates or the MSU Museum. All methods adhered to protocols approved by MSU Campus Animal Resources and the Institutional Animal Care and Use Committee (IACUC).

### Analysis of electric signal variation

Our analysis of EODs has been previously described by ^1–3^. Because the focal phenotype, P0, can be obscured by noise in the recording setup, we modified our typical analysis approach: first, we manually inspected each specimen’s EOD recording for amplifier clipping and discarded any waveforms that appeared clipped. Next, we normalized each specimen’s EODs by its P1-P2 voltage. Next, we aligned each specimen’s EODs in time using the time of P1 as reference. Finally, we averaged each of these aligned EOD recordings to obtain a final EOD for subsequent analysis. For each averaged EOD waveform, we made 21 measurements from each recorded EOD waveform using a custom written program in MATLAB (Mathworks, Inc.: Natick, MA). For each waveform, we calculated amplitudes, times, and slopes at nine landmarks defined by peaks, zero crossings, first derivative peaks, and threshold crossings, and output a composite plot for manual inspection. All plots of EODs from field captured specimens are available as Supplemental Data S2. Code for all EOD analysis is available at https://github.com/msuefishlab/pkings_eod_gwas/tree/main/code/02_Phenotyping.

### Reference Genome

While a reference genome for *Paramormyrops kingsleyae* is already published, ^4^ we were motivated to improve this assembly to obtain improved contiguity to better localize genotype-phenotype associations. We sequenced and assembled a new, chromosome-level assembly from a single biphasic male specimen of *Paramormryops kingsleaye* captured from Bambomo Creek in 2023 (metadata available under NCBI BioProject: PRJNA1218457). Samples from the brain, liver and skeletal muscle tissues were flash frozen and stored in a liquid nitrogen vapor shipper (MVE, Inc.) moments after euthanasia, and transported to Michigan State University for storage. Tissues were stored at -80°C until DNA extraction. DNA extraction, library preparation, sequencing, and assembly were conducted by Dovetail Genomics. High-fidelity sequencing was performed using PacBio CCS technology to generate 88.2 gigabase-pairs of sequence data (∼110x coverage of the genome). These reads were assembled using Hifiasm v0.15.4-r347 ^5^ with default parameters. To ensure data integrity, the assembly was blasted against the NCBI *nr* database, and these results were filtered for contaminants using blobtools v1.1.1^6^. Finally, haplotigs and contig overlaps were eliminated using purge_dups v1.2.5 ^7^.

The resulting draft assembly was further scaffolded using two Dovetail Omni-C libraries. For each Dovetail Omni-C library, chromatin was fixed in place with formaldehyde in the nucleus and then extracted from provided tissue samples. Fixed chromatin was digested with DNAse I, chromatin ends were repaired and ligated to a biotinylated bridge adapter followed by proximity ligation of adapter containing ends. After proximity ligation, crosslinks were reversed and the DNA purified. Purified DNA was treated to remove biotin that was not internal to ligated fragments. Sequencing libraries were generated using NEBNext Ultra enzymes and Illumina-compatible adapters. Biotin-containing fragments were isolated using streptavidin beads before PCR enrichment of each library. The library was sequenced on an Illumina HiSeqX platform to produce approximately 30x sequence coverage. The HiRise pipeline ^5^ was then used to align the Omni-C reds to the draft assembly using bwa ^8^, identify potential misjoins and generate a chromosome-level scaffolded assembly.

This Whole Genome Shotgun project has been deposited at DDBJ/ENA/GenBank under the accession JBLIQM000000000. Raw PacBio sequencing reads and Dovetail OmniC library reads were accessioned under NCBI BioProject PRJNA1218457.

### Genome Annotation

The assembly was annotated by the NCBI Eukaryotic Genome Annotation Pipeline (EGAP v10.3), which released it as RefSeq annotation GCF_048594095.1-RS_2025_03 (27 March 2025) of RefSeq assembly GCF_048594095.1 (PKINGS_0.4). Gnomon predicted gene models from aligned RNA-seq reads and proteins. The RNA-seq evidence was 38 runs from 37 previously published *P. kingsleyae* samples from 14 adult tissues, including electric organ (BioProjects PRJNA417677 and PRJNA573805; 1.56 billion reads, 73% aligned with STAR). The protein evidence was RefSeq proteins of human and six teleosts and Actinopterygii proteins from GenBank, aligned with ProSplign. Non-coding RNAs were predicted with cmsearch and tRNAscan-SE. The annotation contains 42,039 genes: 24,872 protein-coding genes, 15,870 non-coding genes and 1,253 pseudogenes. BUSCO v5.7.1 (actinopterygii_odb10, 3,640 genes) scored the annotated gene set as 98.6% complete (95.1% single-copy, 3.5% duplicated), 0.7% fragmented and 0.7% missing. We used these gene models to identify genes within and near GWAS peaks and to quantify RNA-seq expression. The exception is the phylogenomic gene alignments, which used the earlier NCBI annotation (Release 100) transferred to this assembly with Liftoff (see Phylogenomic Analysis).

### Whole Genome Resequencing

To investigate genome-wide variation, we performed whole-genome resequencing on 305 specimens described in Supplementary Data S1. DNA was extracted from ethanol-preserved fin clips using either DNeasy Tissue Kits (Qiagen, Inc.) or AgenCourt DNAadvance kits (Beckman-Coulter, Inc.). Each specimen provided at least 200 ng of DNA in 50 µL of H₂O for library preparation.

Library construction and sequencing were carried out by the Michigan State University Research Technology Support Facility (RTSF) Genomics Core or Genewiz (Azenta Life Sciences, Inc.). Libraries were prepared using either the Rubicon Genomics ThruPLEX DNA-Seq or the Illumina TruSeq Nano DNA Library kits, following manufacturer protocols. Following quality control, pooled libraries were sequenced on an Illumina HiSeq 2500 (high-output mode) in 2 × 125 bp or 2 × 150 bp mode, targeting an average depth of 14x coverage per specimen. Realized mean sequencing depth was 20.0× (range 5.2–35.6×), computed as total sequenced bases divided by the 993-Mb assembly length. Mean aligned read depth was 10.8× (range 4.2–20.1×), computed as the mean GATK DP across the genome-background sites of the all-sites VCF (see Reference bias), with sites without reads counted as zero. Base calling was performed using Illumina Real Time Analysis (RTA), and output files were demultiplexed and converted to FASTQ format using Illumina Bcl2fastq.

The resulting sequence data have been deposited in the NCBI Sequence Read Archive (SRA) under BioProject PRJNA1218444, with individual accession numbers provided in Supplementary Data S1.

### Variant Calling

Genotyping was performed using Terra (http://tera.bio), which enables implementation of the Genome Analysis Toolkit (GATK) variant discovery and calling pipeline using Google Cloud infrastructure^9^. FASTQ files were converted to BAM files to preserve metadata and were then aligned using bwa ^8^ to our newly obtained *P. kingsleyae* reference genome. Next, duplicate reads from each sample were marked using Picard ^10^ to prevent consideration as independent evidence for variation. Next, GATK was deployed to jointly discover SNP and short insertion-deletion variants across all samples in GVCF mode. Finally, the resulting gvcf files were used to perform joint genotype calling in each sample.

In humans and other model organisms, GATK Best Practices ^9^ recommend that variants obtained are compared to a validated set of variants through a process called Variant Quality Score Recalibration (VQSR), which is a machine-learning guided technique for filtering technical artefacts from true variants. As no such validated set of SNPs exists for *P. kingsleyae*, we implemented ‘hard filtering’ which is a recommended alternative for non-model organisms. Beginning with recommended settings ^9^, we manually examined annotations across all GATK determined variants and determined appropriate thresholds:

**SNPS**: QD > 5; QUAL > 30; SQR > 3; FS > 60 MQ< 40, MQRankSum > -12.5, ReadPosRankSum < -8; <2 DP <100

**INDELS**: QD > 5, FS > 200; ReadPosRankSum > -20; <2 DP < 100; GQ>10

Applying these filtering criteria, we obtained a final dataset with 51,220,152 SNPs and 13,733,103 Insertion/Deletion polymorphisms < 50 bp.

#### Reference bias

The reference assembly derives from a BP male of the Bambomo (BP1) lineage. Because BP EODs evolved independently in different populations, it carries the BP-associated haplotype at Peaks 1, 4 and 5, the TP-associated haplotype at Peaks 3 and 6, and a mosaic at Peak 2 (the BP-associated allele at 21 of 45 diagnostic SNPs). Per-sample alignment files were not retained after joint genotyping, so we assessed mapping bias between morphs from the raw all-sites VCF (invariant and variant sites) for the 198 GWAS individuals (109 BP, 89 TP). For each individual we computed read depth relative to its own mean depth in a genome background, the percentage of sites without reads, and the mean fraction of reference reads at heterozygous biallelic SNPs with depth ≥ 10, which rises above 0.5 when reads carrying non-reference alleles fail to map. We computed these measures in the six GWAS peaks; in 1,000 randomly chosen 5-kb windows from the top 5% of mean nucleotide diversity across the eight core *P. kingsleyae* populations, containing no annotated gene and not overlapping a peak; and in a genome background of 2-kb blocks spaced every 50 kb across the 25 chromosomes. BP and TP individuals were compared with two-sided Wilcoxon rank-sum tests, overall and within the three geographically matched comparisons (BP1 vs. TP1, BP2 vs. TP2 and BP3 vs. TP2). TP individuals showed no mapping deficit (Table S3). In the high-diversity non-coding windows, TP and BP individuals had similar relative depth (0.92 vs. 0.91), percentage of sites without reads (19.1% vs. 17.7%; P = 0.44) and heterozygote reference fraction (0.611 vs. 0.611; P = 0.92). At Peaks 1, 4 and 5, where TP individuals carry the non-reference haplotype, TP and BP individuals of the BP1 vs. TP1 comparison differed by at most 0.012 in relative depth and 0.3 percentage points in sites without reads. The only marked difference was at Peak 2, where Bikagala and Bavavela BP individuals lacked reads at 13.4% of sites, against 3.6% in Douvalou and Douengi TP individuals; this deficit falls on BP rather than TP individuals. All code for this workflow and analysis are available at https://github.com/msuefishlab/pkings_eod_gwas/tree/main/code/01_Terra and https://github.com/msuefishlab/pkings_eod_gwas/tree/main/code/03_VCF_QC_Filter.

### Phylogenomic Analysis

We obtained whole gene alignments by obtaining NCBI annotations from the previous *P. kingsleyae* genome ^11^, and mapping these gene models to our new genome assembly using Liftoff^12^. Next, for each of the 305 genomes, we extracted the biallelic SNPs within the 25,622 annotated genes and applied them to the reference sequence, creating 25,622 whole gene alignments for 305 individuals. To facilitate computation, we randomly selected a representative genome from each species or, in the case of *P. kingsleyae,* each population (see Supplemental Data S1). After filtering out empty gene alignments, the final dataset consisted of 22 taxa with 25,137 genes and 368,469,371 sites. Next, we used IQ-TREE v 2.3.1 ^13^ to find the best-fitting substitution model for each gene and reconstruct the maximum likelihood tree for each gene. Next, we used ASTRAL-IV v1.18.4.5 ^14–16^ to estimate a species tree and local posterior probability branch supports. Finally, IQ-TREE was utilized to compute gene concordance factors (gCF) and site concordance factors (sCF) for each branch of the species tree to further estimate branch support ^17^. All code for this workflow and analysis are available at https://github.com/msuefishlab/pkings_eod_gwas/tree/main/code/04_Phylogenomics.

### Ancestral State Reconstruction - Mormyrids

We were first motivated to assess the number of independent origins of P0-absent EOD waveforms across all known mormyrid species. We started with a recent time-calibrated phylogenetic tree of Osteoglossiformes ^18^ and extracted the subtree containing 169 species within the superfamily Mormyroidea.

Next, we obtained representative EOD recording plots obtained from 106 of these species based on literature and records present in the Macaulay Library at the Cornell Lab of Ornithology (Cornell University, Ithaca NY). We visually inspected each EOD and determined whether the EOD was monophasic (MP), P0-absent (BP) or P0-present (TP). Assigned phenotypes for all species, as well as accession numbers and/or references for EODs used in this analysis, are in Supplemental File S3.

Based on these results, we pruned the mormyrid subtree to tips where phenotype was able to be assessed (n=106 taxa). We estimated transition rates between phenotypic states using a maximum likelihood approach using *Phytools* v. 2.4-4 ^19^ function fitMk. We fit several MK models ^20^ to the phenotypic and phylogenetic data, including the conventional equal rates (ER), symmetric rates (SYM) and all rates different (ARD) models, as well as an additional ‘directional’ model (DIR) which allowed for unidirectional change between MP -> BP and bidirectional change between BP and TP. Based on AIC criterion, the best fitting model was the DIR model (log likelihood=-55.33). Using the estimated character transition matrix from the DIR model, we generated 1,000 replicate stochastic character maps for the mormyrid phylogeny using phytools *make.simmap*, which allowed us to obtain a posterior probability distributions for each ancestral node in the mormyrid phylogeny. All code for this workflow and analysis are available at https://github.com/msuefishlab/pkings_eod_gwas/tree/main/code/04_Phylogenomics.

### Ancestral State Reconstruction – *Paramormyrops*

We next wanted to determine the number of independent origins of P0-absent EOD waveforms among the sampled *Paramormyrops* species and within *P. kingsleyae* populations. Starting with the ASTRAL-IV ML summary tree, we used *makeChronosCalib* in *ape* ^21^ to apply a penalized likelihood method ^22^ to obtain an ultrametric, time calibrated tree, using dates obtained from previous estimates ^18^ for 5 taxa (MRCA *Paramormyrops sp.* ‘SN2’*-Paramormyrops batesii:* **5.75 MY**; MRCA *Paramormyrops szaboii* -*Paramormyrops sp.* ‘MAG’ **5.27 MY**; MRCA *Paramormyrops sp.* SN2 – *Paramormyrops curvifrons* **1.70 MY;** MRCA *Paramormyrops gabonensis – Paramormyrops* sp. ‘MAG’ **1.6 MY**;MRCA *Paramormyrops curvifrons – Paramormyrops hopkinsii* **4.23 MY** ; *Paramormyrops longicaudatus – Paramormyrops sp.* ‘SN2’ **1.98 MY).** All specimens in our phylogenetic tree were assigned as either P0-absent (BP) or P0-present (TP). We again estimated transition rates between phenotypic states using *Phytools fitMK* by fitting various models including the conventional equal rates (ER), symmetric rates (SYM) and all rates different (ARD) models, as well as an additional ‘directional’ model (DIR) which allowed for unidirectional change between TP -> BP. In this case, all models were similar in their log likelihood, and we ultimately selected to proceed with a directional model for consistency with the mormyrid analysis described above. Using the estimated character transition matrix from the DIR model, we generated 1,000 replicate stochastic character maps for the *Paramormyrops* phylogeny using phytools *make.simmap*, which allowed us to obtain posterior probability distributions for each ancestral node in the *Paramormyrops* phylogeny. All code for this workflow and analysis are available at https://github.com/msuefishlab/pkings_eod_gwas/tree/main/code/04_Phylogenomics.

### Population Genetics Analysis

We used pixy ^23^ to calculate π for each population, and Weir and Cockerham’s F_st_ and d_xy_ for each pair of populations, in non-overlapping 5-kb windows from biallelic SNPs and invariant sites. We used vcftools ^24^ to calculate Tajima’s D and the number of variable sites in the same windows for each population from biallelic SNPs. All code for this workflow and analysis are available at https://github.com/msuefishlab/pkings_eod_gwas/tree/main/code/05_PopGen.

### Association Analysis

A critical challenge in genome-wide association studies (GWAS) is accounting for population structure and cryptic relatedness, both of which can lead to inflated p-values and false-positive associations (i.e. the erroneous conclusion that all loci significantly contribute to phenotype; which is unlikely), or the detection of “false positive” loci (i.e. the erroneous conclusion that a particular locus contributes to phenotypic variation but is in fact associated with ancestry). In *P. kingsleyae*, previous analysis of population structure ^3^, and our present population genetics analysis reveal substantial genetic differentiation among population throughout Gabon. To mitigate these confounding effects, we utilized the genome-wide efficient mixed model association model association algorithm GEMMA ^25^ for genotype-phenotype association testing.

Our GWAS focused on individuals from all populations of *P. kingsleyae* sampled (see Supplementary Data S1). We categorized individuals by EOD waveform as TP, BP, or mixed (see <u>Analysis of electric signal variation, above)</u>. Of the 240 *P. kingsleyae* individuals genotyped, we excluded the 42 mixed individuals, leaving 109 BP and 89 TP individuals.

We constructed a centered relationship matrix (-*gk 1*) using a linkage disequilibrium (LD) thinned subset of SNPs (≤10% missing data, minor allele frequency MAF >5%, and one randomly selected SNP per 20 kb; n = 37,331 SNPs). GWAS was performed using a univariate linear mixed model (-lmm 2), incorporating the relatedness matrix as a random effect and testing each SNP with a likelihood ratio test. The full genotypic matrix contained all biallelic SNPs meeting quality control thresholds (n = 5,898,721 SNPs).

To check that our conclusions did not depend on the choice of test, we repeated the association with GEMMA’s Wald test (-lmm 1) and, for a matched comparison, with the likelihood ratio test on the same input. Both runs used a statistically phased version of the same call set (6,630,666 biallelic SNPs), with the same relatedness matrix, phenotypes and 100 permuted phenotypes for thresholds. The two tests gave nearly identical per-SNP evidence (Spearman ρ = 1.00 across 3,429,189 SNPs passing the frequency filter; median difference in −log₁₀(p) = −0.002, and −0.05 among SNPs with −log₁₀(p) > 4), and both recovered all six peaks reported in the main text, with the same index SNP at each peak. At Peaks 2, 3, 4 and 6 this index SNP differed from that of the main analysis under both tests, reflecting the different SNP set rather than the test.

The tests differed mainly in their permutation thresholds (significant / suggestive −log₁₀(p): Wald 7.98 / 6.80; likelihood ratio test 7.25 / 6.52; main analysis 7.22 / 6.36). The Wald threshold is inflated because 10 of its 100 permutations gave unstable estimates (undefined standard errors), three of which were among the five most extreme permutation minima. Under the lower likelihood-ratio thresholds, the phased data set yielded three further loci (chr2: 4.33– 4.60 Mb, suggestive only; chr5: 7.19–7.43 Mb; chr7: 10.45–10.51 Mb). All three appear among the candidate regions of the main analysis but fail its peak definition, and the chr5 locus also fails it in the phased data once a symmetric minor-allele-frequency filter (MAF > 0.15) is applied. Finally, replacing the one-sided frequency filter used before peak calling (alternate-allele frequency > 0.15; see below) with this symmetric filter left the maximum −log₁₀(p) and the number of suggestive SNPs unchanged at all six peaks.

In non-model organisms, determining genome-wide significance thresholds is challenging due to the lack of a well-defined linkage structure. To address this, we employed GEMMA-wrapper ^26^ to perform 100 phenotype permutations per GWAS. The minimum p-value from each permutation was used to define significance thresholds: SNPs exceeding the 67th percentile were considered “suggestive,” and those exceeding the 95th percentile were considered “significant.”.

We visualized results using Manhattan plots, identifying clusters of significant SNPs forming association peaks. Before defining peaks, we excluded SNPs at which the alternate (non-reference) allele had a frequency ≤ 0.15 among the 198 analysed individuals. We merged SNPs with -log10(p) > 5 that lay within 75 kb of one another into candidate regions, and retained a region as a peak if it contained >5 SNPs exceeding the suggestive threshold. Peak boundaries (Table S1) are the outermost SNPs with -log10(p) > 5 in each peak. For the haplotype-based selection scans and the introgression statistics, we used extended windows around the same six peaks, with boundaries at the outermost SNPs with -log10(p) > 3 (merged within 75 kb). For each peak we:

1. Identified the most significant SNP (herein referred to as the *index SNP*)
2. Used PLINK v. 1.9 (*85*) to calculate squared genotype correlation values (R^2^) for all SNPs within 10,000 SNPs or 10 Mb of the index SNP (whichever was less).
3. Intersected peak coordinates using the R package *GenomicRanges* ^27^ with *P. kingsleyae* annotations ^28^ within or adjacent to peaks.
4. Extracted the genotypes of the ≤ 250 most highly associated SNPs for all 305 *Paramormyrops* samples using BCFtools ^29^.
5. For each peak we took the up to 250 most strongly associated SNPs within the extended peak window (−log10(p) > 3) obtained in (4) and plotted the phased haplotypes of every individual at those SNPs (Fig. S4). Raw unphased genotypes agreed with the phased genotypes at every called site.

At each SNP the BP allele was the majority allele of BP individuals from the populations in which the BP allele arose (Bambomo for Peaks 1, 4 and 5; Bikagala, Bavavela and Biroundou for Peaks 2 and 3; Bikagala and Bavavela for Peak 6), and the TP allele was the majority allele of Apassa, Bengue, Douvalou and Douengi TP individuals. Only SNPs at which the two differ were plotted (26, 85, 163, 145, 95 and 86 SNPs at Peaks 1–6). To ask whether a BP-like haplotype found outside those populations was the same haplotype, we compared it with every BP haplotype of those populations. We called it the same haplotype if it matched one exactly, or differed from the nearest at one or two SNPs where it carried an allele also found on BP haplotypes of those populations. We called it a distinct haplotype if it differed from every one at three or more SNPs, or carried a read-supported allele found on none of them. For comparison, BP haplotypes within Bambomo differ among themselves at this scale: at Peak 4, 35 of 78 Bambomo BP haplotypes have no identical copy among the others.

To assess the variance explained by highly associated SNPs (from 4 above) we filtered the genotypes of the ≤250 most highly associated SNPs at each peak to those in strong linkage disequilibrium (LD; R² > 0.6) with each index SNP for only the *P. kingsleyae* individuals. Next, we assigned each individual to a phenotypic group based on its position in our ancestral character reconstruction (TP for triphasic, and BP1, BP2, BP3, BP4 for independently derived biphasic lineages; see Fig. 2). For each SNP, we fit a univariate logistic regression comparing individuals in that biphasic group (coded as 1) against all triphasic individuals (coded as 0) using the R function glm(phenotype ∼ genotype, family=“binomial”) and computed McFadden’s pseudo-R² for each regression using the pscl function pR2. This approach yielded separate variance explained estimates for each SNP × biphasic lineage combination, allowing us to assess whether SNPs have consistent or lineage-specific effects on the biphasic phenotype (Fig. 3C). All code for this workflow and analysis are available at https://github.com/msuefishlab/pkings_eod_gwas/tree/main/code/06_Association and https://github.com/msuefishlab/pkings_eod_gwas/tree/main/code/08_Peak_Analysis.

### Analysis of Introgression

We tested whether BP alleles at the GWAS peaks were shared among lineages through introgression with ABBA-BABA statistics computed in Dsuite v0.5 r58^30^. Individuals were assigned to eight groups: BP1 (Bambomo, n = 41), TP1 (Apassa and Bengue, 35), BP2 (Bikagala and Bavavela, 48), TP2 (Douengi and Douvalou, 31), BP3 (Biroundou, 16), TP_OUT (triphasic *P. kingsleyae* from Mouvanga, 10), other biphasic *Paramormyrops* (BP_ANC: *P. curvifrons* 5, *P. hopkinsii* 1, *P. longicaudatus* 3, *P. ntotom* 4 and *P.* sp. ’SN2’ 4) and the outgroup (*P. szaboii*, 11). Trios were oriented on the species tree (((((BP1,TP1),((BP2,TP2),BP3)),TP_OUT),BP_ANC),outgroup).

### Patterson’s D and f₄-ratio

Dsuite Dtrios computed Patterson’s *D*^31,32^ and the f₄-ratio^33^ with block-jackknife standard errors (20 blocks), (i) genome-wide, combining all 25 chromosomes (51.1 million biallelic SNPs), and (ii) within each GWAS peak, using the extended peak intervals (SNPs with −log₁₀ *p* > 3 merged within 75 kb; 34–400 kb). We report six tests: each pair of BP lineages, with the triphasic sister of one lineage as the reference ((BP1, TP1; BP2), (BP1, TP1; BP3), (BP2, TP2; BP3)), and each BP lineage against BP_ANC with Mouvanga TP *P. kingsleyae* as the reference ((BPx, TP_OUT; BP_ANC)). Two controls replaced the BP lineage with TP1 or TP2. *D* was signed so that positive values indicate excess allele sharing between the BP lineage (or control) and the third group. *P* values were adjusted across all 48 peak tests (Benjamini–Hochberg). Because genome-wide *D* differs from zero for most trios, we also compared each peak with the genome-wide value for the same trio, (*D*_peak_ − *D*_genome_)/SE_peak_, and considered a peak to show excess sharing when *q* < 0.05 and *D*_peak_ exceeded *D*_genome_ by more than two standard errors (Fig. S5; Table S4; Supplementary Data S7).

### Windowed statistics

Dsuite Dinvestigate computed f_dM_^34^ in windows of 50 informative SNPs with a step of 25 SNPs along each chromosome, for the same six tests. Trios were oriented with P1 and P2 as sisters, so negative f_dM_ indicates excess P1–P3 sharing; values were multiplied by −1 so that positive values indicate sharing between the BP lineage and the third group. This orientation agrees with the sign of genome-wide *D* for each trio and with the reciprocal trio (BP3, TP2; BP1). For each peak we took the median f_dM_ of the windows overlapping the extended interval and compared it with (i) all windows outside the peaks and (ii) medians of non-overlapping blocks of the same number of consecutive windows outside the peaks (Fig. S6).

### Targeted test at Peak 6

*P. curvifrons*, *P. ntotom* and *P.* sp. ’SN2’ carry many of the BP2 alleles at Peak 6 (Fig. S4), although none carries the BP2 haplotype, and BP_ANC pools them with two species, *P. hopkinsii* and *P. longicaudatus*, that carry no BP-like haplotype at the peak. We therefore repeated the Peak 6 tests with these three species alone as the third group (CURV3; 13 individuals), with the other groups, the peak interval and the Dsuite version as above, oriented on the tree (((((BP1,TP1),((BP2,TP2),BP3)),TP_OUT),CURV3),outgroup). The groups, tests, windows and decision rule were fixed before the test was run, although the allele sharing had been seen. The primary test was (TP_OUT, BP2; CURV3), as for the BP_ANC tests; we also computed the sister form (TP2, BP2; CURV3) and four controls, (TP1, TP_OUT; CURV3), (TP2, TP_OUT; CURV3), (BP1, TP_OUT; CURV3) and (TP_OUT, BP3; CURV3). P values were adjusted across these six tests. The rule was that of the tests above, with the addition that the peak median f_dM_ exceed the 95th percentile of equal-sized background blocks; excess sharing also shown by a triphasic control would indicate shared ancestry rather than a BP-specific signal. The analysis first reproduced the published Peak 6 values of all trios exactly (BP2–BP_ANC: *D* = 0.197, 445 informative sites). In the primary test, *D* = 0.231 (*Z* = 3.83, q = 3.9 × 10^−4^; 472 informative sites) and the peak f_dM_ was at the 96.1st percentile of background blocks, but *D* exceeded the genome-wide value for the trio (0.135) by only 1.59 standard errors, so the rule was not met. No control showed excess sharing (*D* = −0.082 to 0.024; f_dM_ at the 7.9th–20.5th percentiles). The sister form met the rule (*D* = 0.385, 5.07 standard errors above the genome-wide value; f_dM_ at the 98.8th percentile). Both reference groups, TP2 and Mouvanga, were triphasic controls in the association analysis that detected Peak 6. The test with Mouvanga as the reference had been designated primary before the test was run, as for the BP_ANC tests, and we base our conclusion on it: at Peak 6 the sharing cannot be distinguished from shared ancestral variation (Supplementary Data S7).

### Divergence between BP2 and BP3

To ask whether haplotypes shared by BP2 and BP3 were exchanged recently, we computed *d*_XY_^35^ with pixy^23^ in 5-kb windows between BP2 and BP3, between TP2 and BP3, and between each lineage and the outgroup *P. szaboii*, pooling the per-population counts. Because the peaks are more divergent than the genome average for every pair of lineages, we used relative node depth^36^, RND = *d*_XY_(BP2, BP3) / mean(*d*_XY_(BP2, outgroup), *d*_XY_(BP3, outgroup)), which scales out local differences in mutation rate and diversity. Recent gene flow lowers RND between the lineages that exchanged alleles. We compared RND at Peaks 2, 3 and 6 with non-overlapping blocks of the same number of consecutive windows elsewhere in the genome. Code is available at https://github.com/msuefishlab/pkings_eod_gwas/tree/main/code/10_Introgression.

### Tests for Selection at GWAS Peaks

We tested the six GWAS peaks for signatures of recent positive selection in the lineages associated with their BP alleles: Peaks 1, 4 and 5 in Bambomo (BP1, n = 42), Peaks 2 and 3 in Bikagala and Bavavela (BP2, n = 48) and Biroundou (BP3, n = 16), and Peak 6 in BP2. The TP references were Apassa and Bengue (TP1, n = 35) for BP1 and Douvalou and Douengi (TP2, n = 31) for BP2 and BP3. SNPs were phased using SHAPEIT4^37^ without a genetic map or reference panel, and ancestral states were assigned from 12 outgroup individuals (11 *P. szaboii* and one *P. hopkinsii*).

We examined three signatures of a selective sweep: unusually long haplotypes (Rsb^38^, calculated with rehh^39^), distortion of the site-frequency spectrum (the SweepFinder2^40,41^ composite likelihood ratio, CLR), and reduced diversity in BP relative to TP populations (πBP/πTP, calculated with pixy^23^). Rsb was standardized by chromosome and averaged over overlapping 20-kb windows stepped every 5 kb; CLR was calculated on a 10-kb grid against each lineage’s genome-wide frequency spectrum; and diversity ratios were calculated from 5-kb estimates summarized in 20-kb windows. Within ±100 kb of each index SNP, we retained the maximum Rsb and CLR and the minimum diversity ratio. For peaks tested in both BP2 and BP3, we retained the more extreme lineage result, using the same procedure for background regions.

We assessed each statistic against 4,383 control regions spaced every 200 kb across the genome, excluding the GWAS peaks and 250-kb flanks. To account for variation in data quality, we matched controls to each peak by SNP density and callable sequence, relaxing the matching range until at least 1,000 regions were available. We did not match on BP–TP differentiation, which could itself be elevated by selection. Each peak received a percentile for each statistic. We called a peak supported when at least two statistics reached the 95th percentile, and strong when one also reached the 99th. The empirical p-value was the fraction of matched controls whose second-highest percentile was at least as large as the peak’s. We also examined CLR in the TP reference to assess whether the signal was specific to the BP lineage.

We checked the sensitivity of these calls to search-region and window size, the choice of background regions, sample-size differences, ancestral polarization, and alternative haplotype statistics. Each variation was evaluated against its corresponding control distribution (Table S5). Code is available at https://github.com/msuefishlab/pkings_eod_gwas/tree/main/code/11_Drift_vs_Selection/sweep_detection.

### Drift Test at GWAS Peaks

We next asked whether neutral demographic histories could generate the allele-frequency differences observed at the GWAS index SNPs. We considered 11 population–peak comparisons: Bambomo at Peaks 1, 4 and 5; Bikagala and Bavavela separately at Peaks 2, 3 and 6; and Biroundou at Peaks 2 and 3. Each BP population was compared with its TP reference as defined above. These tests used all sampled fish, including mixed-phenotype individuals, to avoid inflating differentiation by selecting samples on the basis of EOD phenotype.

We fitted two-population demographic models to the joint site-frequency spectrum using dadi 2.4.3^42^. The input was an unphased, all-sites VCF containing invariant positions, after excluding unsupported genotypes and SNPs with excess heterozygosity. SNPs were polarized where at least two of six outer-clade species (*P. szaboii*, *P. magnostipes*, *P. parsegnis*, *P. hopkinsii*, *P. gabonensis* and *P. vadamans*) had data and all were fixed for the same allele. The index SNPs of Peaks 2 and 6 could not be polarized, so those comparisons used folded spectra. We excluded exons, the six GWAS peaks and their 250-kb flanks, a chromosome 4 inversion-like region, and poorly called regions when fitting the models. We considered divergence without migration, divergence with symmetric or asymmetric migration, and isolation with migration, a founder bottleneck and exponential growth. Model uncertainty was assessed with 100 block-bootstrap spectra. We retained only converged models with acceptable ancestral-state estimates and a predicted abundance of strongly differentiated SNPs within threefold of that observed. In the three comparisons with Douvalou and Douengi, TP-derived near-fixed differences (frequency difference ≥ 0.95) exceeded the fitted models’ expectation 1.4–3-fold, which none of the models explains.

For each comparison, we calculated *E*, the number of callable SNPs expected genome-wide to show an allele-frequency difference at least as large as the index SNP, and compared it with the observed count outside the focal peak. To err toward drift, we reported the largest of the best model’s upper bootstrap estimate, the point estimates of other admissible models, and an estimate allowing local reductions in effective population size associated with background selection (Table S6). Large values of E mean that differentiation of this magnitude is compatible with drift; they do not demonstrate that the index allele rose in frequency by drift. When no model met the adequacy criteria, we reported the observed genomic count without assigning a neutral expectation.

At Peak 1, 95% of the estimate allowing local reductions in effective population size comes from 25 of 897 1-Mb regions elsewhere in the genome with low estimated effective size. None of those regions holds a SNP as differentiated as the Peak 1 index SNP, and the Peak 1 region itself has an average effective size. Because the index-SNP comparison does not account for clustering of linked variants, we also examined Peak 1 as a region. No SNP outside the GWAS peaks matched the Peak 1 index difference (Δ = 0.99), so we counted clusters of SNPs at least as differentiated as the Peak 4 index SNP (Δ ≥ 0.81), using overlapping ±50-kb neighborhoods (±175 kb as a check), and compared Peak 1 with non-peak clusters, including controls matched for SNP density and callability. We also attempted a model-based regional test by simulating linked variation^43^ under the fitted histories and additional histories allowing population contraction. These simulations did not reproduce the observed decay of linkage disequilibrium in Bambomo, so we did not assign a neutral probability to the Peak 1 region as a whole. Both regional analyses were exploratory. Code is available at https://github.com/msuefishlab/pkings_eod_gwas/tree/main/code/11_Drift_vs_Selection/demographic_modeling and https://github.com/msuefishlab/pkings_eod_gwas/tree/main/code/11_Drift_vs_Selection/simulation.

### Historical Effective Population Size

Finally, we reconstructed changes in effective population size for four BP and four TP populations using SMC++ 1.15.4^44^, with up to 20 individuals per population, five of which served as distinguished individuals (the first five in each sample list; for Bambomo, mixed-phenotype fish sampled in 2019). We used the unphased all-sites genotypes, masking poorly called sites, GWAS peaks and the chromosome 4 inversion-like region. Estimates used the folded spectrum and assumed a mutation rate of 1 × 10⁻⁸ per site per generation, with uncertainty from ten chromosome bootstraps. Tests on simulated demographic histories^43^ indicated that reductions in effective population size could be detected, but their timing and precise magnitude were less reliable. We therefore use these reconstructions to assess whether historical population size may have permitted stronger drift, rather than to date specific demographic events (Fig. S9). Code is available at https://github.com/msuefishlab/pkings_eod_gwas/tree/main/code/12_Historical_Ne.

### RNA Sequencing and Differential Expression Analysis

Electric organ tissue samples were collected immediately after euthanasia in the field and preserved in RNA-later at 4 °C. They were subsequently transported to Michigan State University and stored at -80°C until processing. RNA extraction and RNA-seq library preparation were performed by GeneWiz (Azenta Life Sciences, Inc., South Plainfield, NJ). The libraries were pooled and sequenced on an Illumina HiSeq 2500 in 2 × 150 bp paired-end mode (31 libraries; 15.5–25.4 million read pairs per library). Raw sequence reads were deposited in the NCBI Sequence Read Archive (SRA) under BioProject PRJNA1218444. We added eight electric organ libraries sequenced for earlier studies: three Bambomo individuals from the *P. kingsleyae* genome project^4^ (BioProject PRJNA417677; Illumina HiSeq 2500 or HiSeq 4000, 2 × 100 or 2 × 150 bp) and five Bikagala and Mouvanga individuals (BioProject PRJNA573805; Illumina HiSeq 2000, 2 × 100 bp). In total, we analysed 39 electric organ libraries from eight populations: Apassa (6), Bambomo (9), Bengue (5), Bikagala (3), Biroundou (5), Douengi (4), Douvalou (5) and Mouvanga (2). Run accessions for every library are in Supplementary Data S1.

To ensure high-quality sequencing data, we processed raw reads using Trimmomatic v0.39 ^45^. This involved removing library adaptors, filtering out low-quality reads, and discarding short reads. We followed the recommended settings from ^46^: 2:30:10 SLIDINGWINDOW:4:5 LEADING:5 TRAILING:5 MINLEN:25. Next, we aligned the trimmed reads from each specimen to the reference transcriptome ^28^ (NCBI RefSeq annotation GCF_048594095.1-RS_2025_03; see Genome Annotation) using the STAR aligner ^47^. Transcript-level expression estimates were obtained using rsem-calculate-expression ^48^.

We focused our analysis on genes located within or near GWAS peaks identified in the association analysis. Transcript-level RSEM expected counts were imported into R using tximport ^49^ and summarized to the gene level using a transcript-to-gene mapping derived from the NCBI RefSeq annotation (GCF_048594095.1-RS_2025_03). Samples were filtered to include only individuals from populations used in GWAS1 (APA, BAM, BEN, MOV, BIR, DOG, DOV, BIK). Gene annotations were intersected with GWAS peak coordinates (defined in output_data/06_Association/PKINGS_ALL_WOB_EXCLUDED/PKINGS_ALL_WOB_EXCL UDED_PEAKS.bed) to identify candidate genes. Genes overlapping peaks exactly or located within 25 kb of a peak were retained for downstream analysis.

For visualization and comparison across populations, gene-level counts were normalized using variance-stabilizing transformation (VST) implemented in DESeq2 ^50^, with genes having fewer than 10 total counts across all samples removed prior to transformation. Expression patterns of candidate genes were visualized using heatmaps clustered by GWAS peak and boxplots grouped by population and phenotype. All code for this workflow and analysis are available at https://github.com/msuefishlab/pkings_eod_gwas/tree/main/code/09_RNASeq.

### Immunohistochemistry

One individual from Apassa Creek (TP EOD) and one from Bambomo Creek (BP EOD) were transported live from Gabon to Michigan State University in 2023, where they were maintained under standard laboratory conditions (26 °C; conductivity = 500 µS; pH = 7.0; 12 h light : 12 h dark). Electric organ discharges (EODs) were verified immediately prior to euthanasia. Fish were euthanized using an overdose of MS-222, and electric organs and brains were rapidly dissected, immersed in OCT compound (Tissue-Tek), and placed on a stainless-steel freezing bar (Pathology Innovations, Inc.) maintained at −20 °C. Tissues were embedded in OCT blocks, frozen, and serially sectioned parasagittally at 15 µm. Sections were air-dried and stored at −80 °C until staining.

For immunohistochemistry, slides were brought to room temperature in a humidified chamber and circled with a liquid blocking pen. Sections were rehydrated with three short rinses in wash buffer (1× PBS, 10 mM Tris pH 7.5, 0.3% Triton X-100), then blocked for 1 h at room temperature in wash buffer containing 5% normal goat serum (NGS). Slides were rinsed three times briefly, followed by three 10-min washes in wash buffer in a Coplin jar.

Primary antibody f5 (mouse anti-SPTBN4; Santa Cruz Biotechnology) was applied at 1:50 and incubated for 1 h at room temperature in a humidified chamber. Slides were washed again (3 quick + 3 × 10 min) before incubation with Alexa Fluor 488–conjugated goat anti-mouse secondary antibody (1:200) for 1 h at room temperature. After another wash series (3 quick + 3 × 10 min), nuclei were counterstained with NucBlue (DAPI; Invitrogen) for 15 min, followed by a final 5-min wash in buffer. Slides were coverslipped with Vectashield mounting medium and imaged using epifluorescence microscopy.

Each session’s Apassa and Bambomo sections and their no-primary controls were imaged with one objective (HC PL FLUOTAR 20×/0.50 or HCX PL FLUOTAR 40×/0.75) and identical exposure, lamp intensity and gain on an 8-bit camera. At most 1.1% of pixels were saturated. To quantify labeling, we defined the electrocyte band from the DAPI channel alone: the smoothed nuclear density above its Otsu threshold, dilated by 4 µm. We then measured the mean SPTBN4 intensity within this band and subtracted the same measure from the population’s no-primary control. For each of four session-by-objective image sets (1–2 Apassa and 1–5 Bambomo fields per set), we report the Bambomo/Apassa ratio of field medians. Fields are technical replicates from one individual per population, so we did not test for a population difference. Code is available at https://github.com/msuefishlab/pkings_eod_gwas/tree/main/code/16_IHC.

### Transcription Factor Binding Site Analysis

#### Allele-specific motif predictions

We asked whether noncoding variants at GWAS Peaks 2, 4 and 5 systematically altered predicted transcription factor binding motifs on BP-versus TP-associated alleles. We selected 302 noncoding SNPs with −log₁₀(p) > 3 from these peak regions. For each SNP, BP and TP alleles were assigned according to their frequencies in BP and TP fish, excluding individuals with mixed EODs. SNPs within 32 bp of one another were grouped to account for motifs spanning multiple variants. For each SNP or group, we constructed paired reference-based sequences carrying the BP or TP alleles with 32 bp of flanking sequence; other positions retained the reference allele. Both sequences were scanned with FIMO^51^ (MEME Suite v5.5.8) against 879 nonredundant vertebrate motifs from JASPAR 2024^52^, using a genome-derived nucleotide background. A predicted motif was counted when a hit with p < 10⁻⁴ overlapped a tested SNP, and was classified as BP-only or TP-only when detected on just one sequence.

#### Tests for directional enrichment

We tested whether BP-only motif predictions were overrepresented relative to 10,000 matched control sets drawn from noncoding SNPs more than 100 kb from GWAS peaks. Single-SNP controls were matched for nucleotide substitution class, CpG context, local GC content, allele frequency and reference-allele orientation; groups of nearby SNPs were matched instead for number and spacing of variants and GC content. Each control set was scanned and scored in the same way as the peak SNPs. The primary statistic was the proportion of directional motif predictions that were BP-only, pooled across the three peaks. We calculated two-sided empirical p-values for this proportion overall and at each peak, and tested individual transcription factor families for directional bias, applying the Benjamini– Hochberg correction^53^ across families.

#### Prioritization of candidate variants

Separately, we examined SNPs and indels within the peak regions and their extended boundaries that were strongly linked to the index SNP (genotype-dosage r² ≥ 0.8; genotype call rate ≥ 0.8). For each variant, we compared the most common local BP- and TP-associated haplotypes, using phased SNP data where available and the most common local sequence for unphased indels. Sequences extending 32 bp on either side of the variant were scanned with FIMO. We retained variants for which an overlapping motif had p < 10⁻⁴ on at least one allele and differed by at least ten-fold in match p-value between alleles; when one allele had no hit at p ≤ 10⁻², we assigned p = 10⁻² to that allele for this comparison. These predictions were used to prioritize variants, not to test enrichment (Supplementary Data S4–S6).

#### Use of generative AI

ChatGPT (OpenAI) and Claude (Anthropic; Claude Opus 5 and Opus 5.5) were used to help write, debug and run analysis code, and to edit the manuscript text for clarity. The authors reviewed and validated all code and results and take full responsibility for the content.

---

**Fig. S1.**
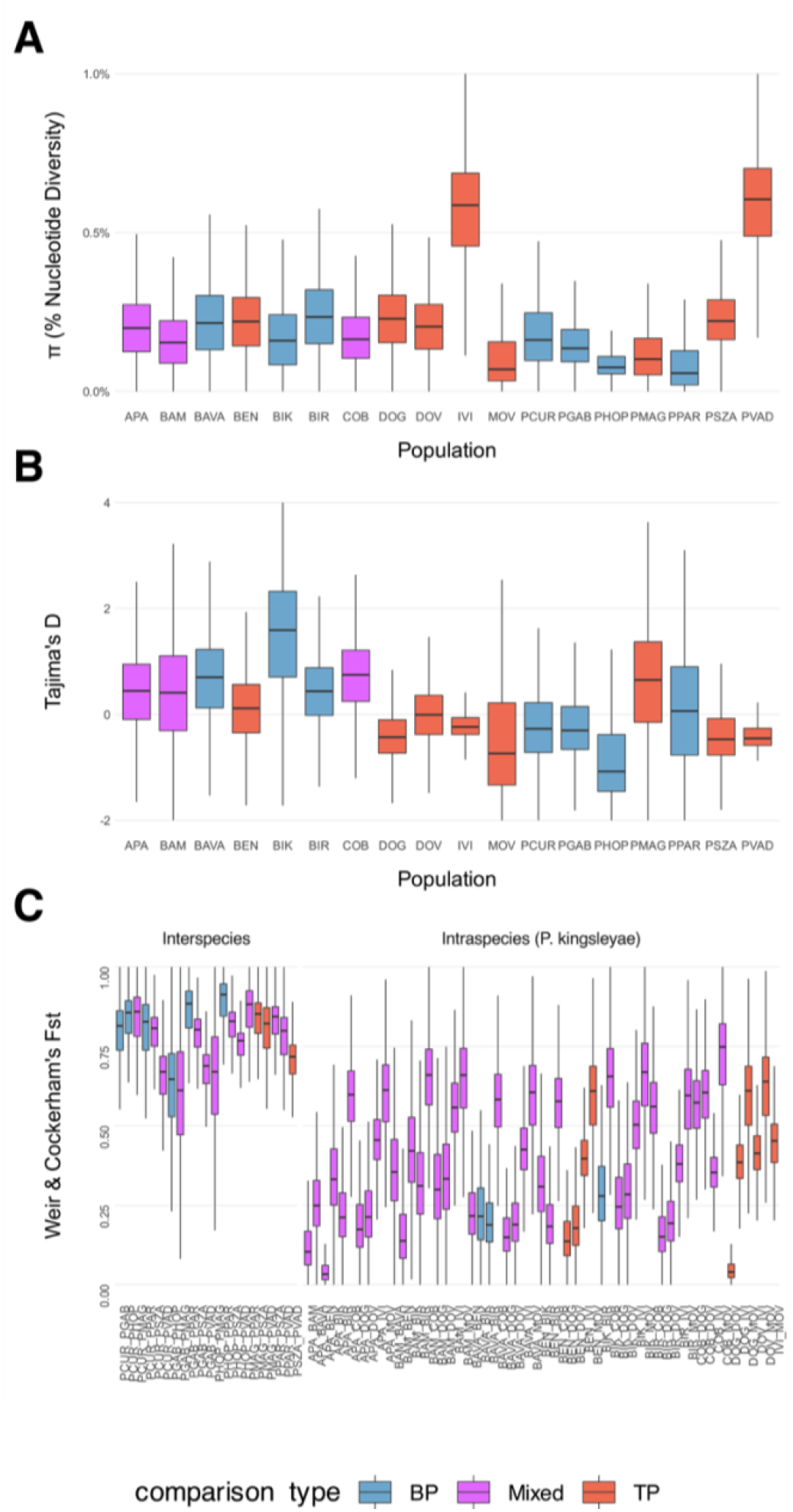
Summary population-genetic statistics across *Paramormyrops* populations. Nucleotide diversity (π), Tajima’s D, and pairwise F_ST_ were estimated in non-overlapping 5 kb windows for each population. **A,** π is broadly similar across *P. kingsleyae* populations and other *Paramormyrops* species, with no populations showing extreme loss of diversity. **B,** Genome-wide Tajima’s D is typically near zero, indicating that most populations do not show pervasive deviations from mutation–drift equilibrium at the scale of whole genomes. **C,**Windowed Weir and Cockerham F_ST_ estimates for all pairwise comparisons reveal strong genetic structure both among recognized *Paramormyrops* species and among populations within *P. kingsleyae*. Bars are coloured by phenotype category of the focal population: blue, biphasic (BP); red, triphasic (TP); purple, mixed.

**Fig. S2.**
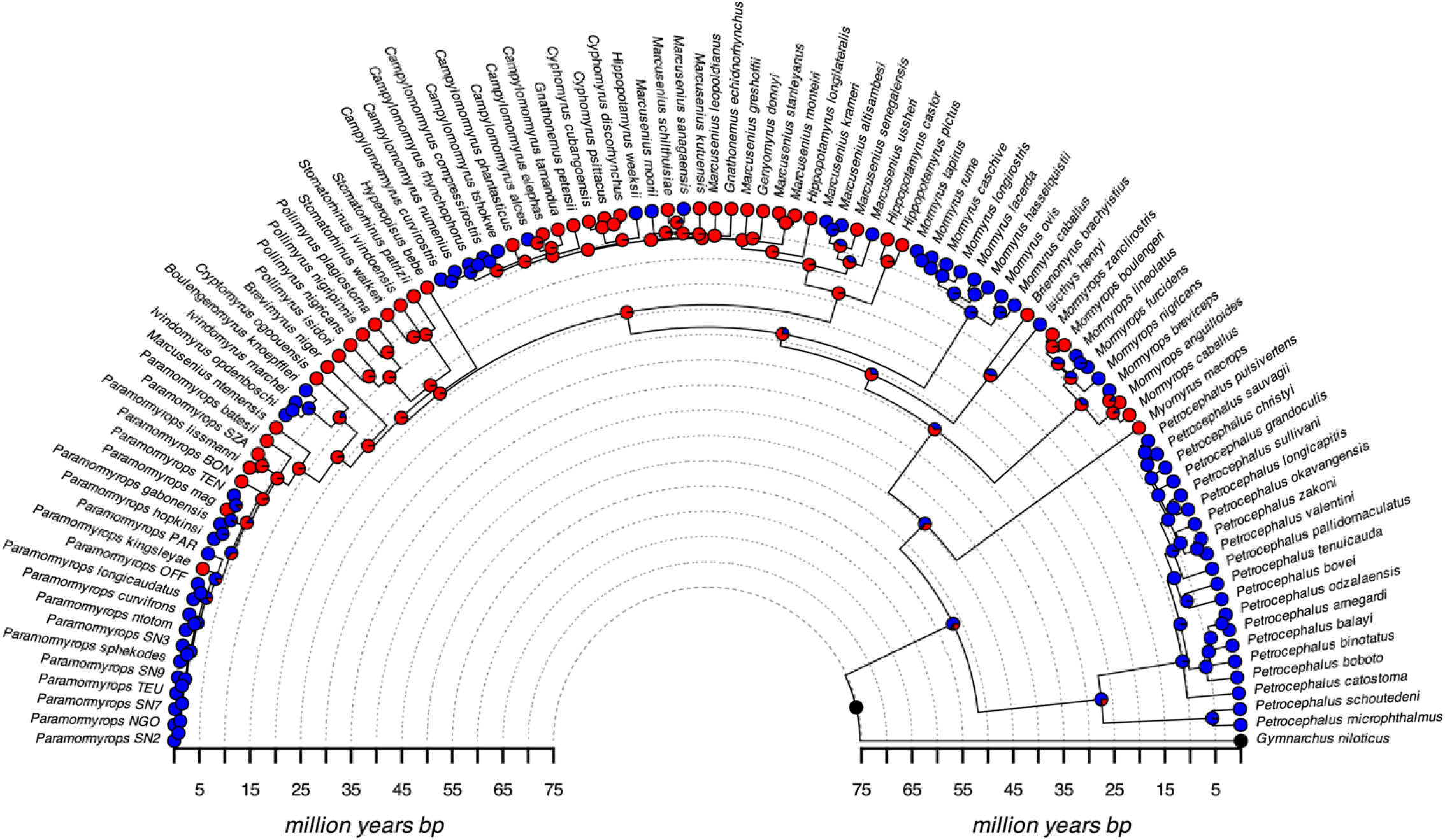
Ancestral reconstruction of P0 presence and absence across Mormyridae. A time-calibrated phylogeny of Mormyridae (pruned from a previously published tree) is annotated with ancestral-state reconstructions of the presence (P0-present, red) or absence (P0-absent, blue) of the small leading head-positive phase (P0) in the electric organ discharge (EOD). P0-absent EODs arose early in mormyrid evolution (∼50 million years ago), and the most likely origin of P0-present EODs is in the most recent common ancestor of *Mormyrops* and *Petrocephalus*. Across 1,000 stochastic character maps, we infer a median of 18 transitions from P0-present to P0-absent EODs (95% HPD interval 11–22) and 10 from P0-absent to P0-present EODs (6–16). Tip colors indicate observed phenotypes; pie charts at internal nodes show posterior probabilities of ancestral states. Time is shown in millions of years before present along the outer axis; see Materials and Methods for details of the phylogeny and reconstruction.

**Fig. S3.**
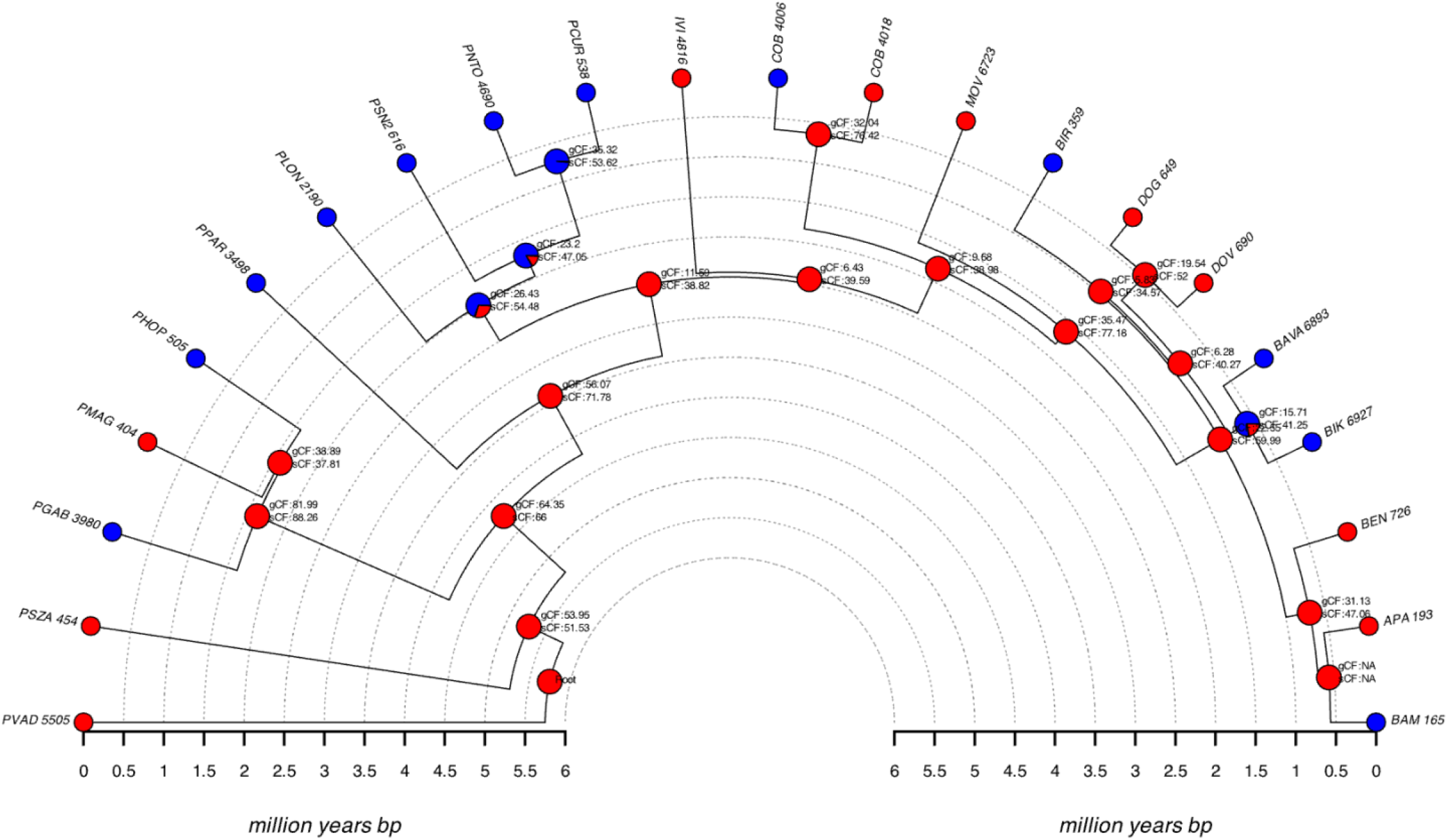
Gene-tree and site-pattern concordance across *Paramormyrops* lineages. Time-calibrated species tree for *Paramormyrops* and focal *P. kingsleyae* lineages, annotated with gene concordance factors (gCF) and site concordance factors (sCF) at internal nodes. Node symbols summarize the percentage of gene trees and site patterns supporting the species-tree topology at each node, providing a quantitative measure of gene-tree/species-tree concordance. Many internal branches show high sCF but modest gCF, indicating that while the dominant site pattern supports the species tree, individual gene trees are frequently discordant consistent with incomplete lineage sorting and complex population history rather than simple bifurcating divergence. Time is plotted in millions of years before present along the radial axis.

**Fig S4.**
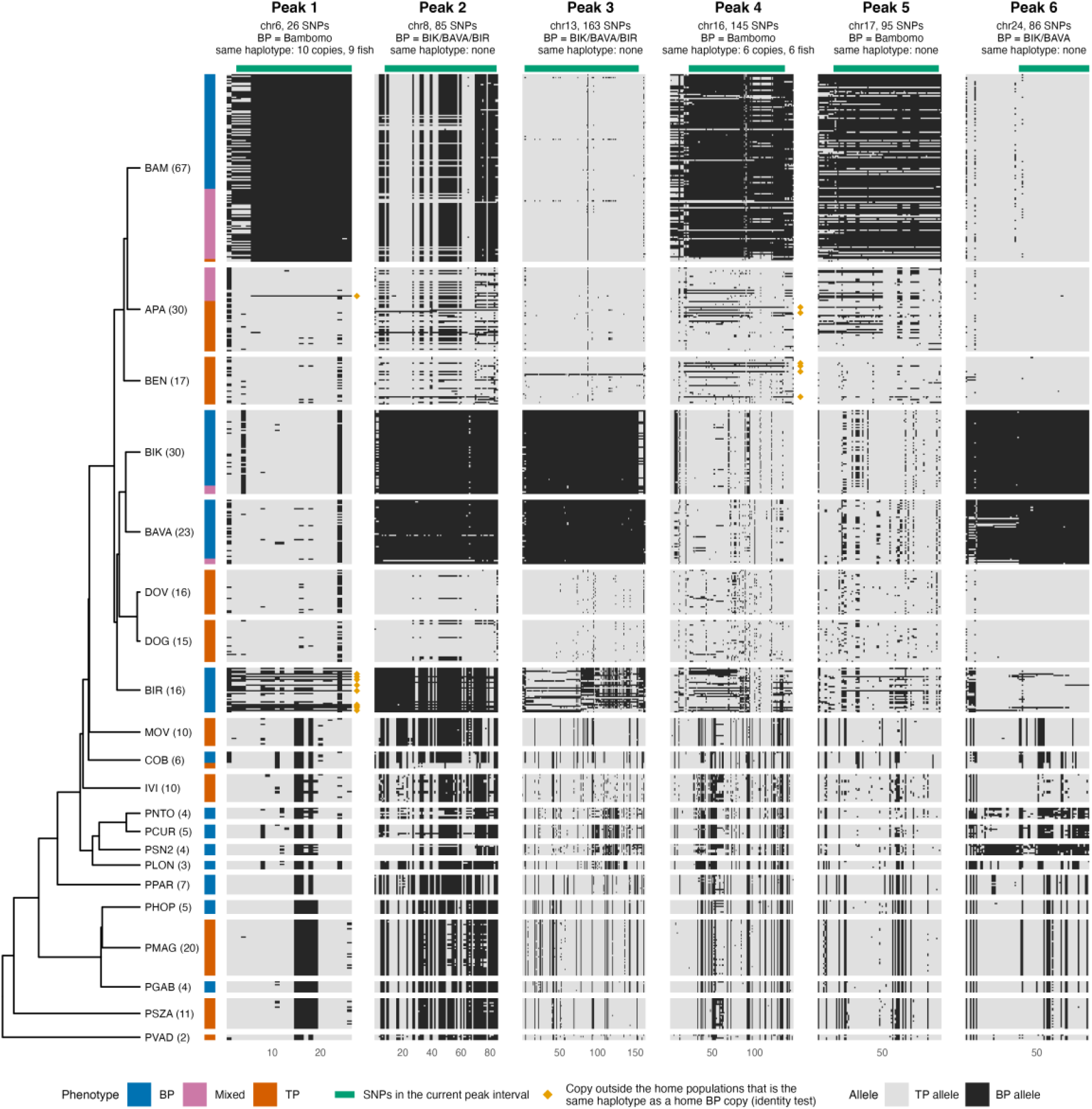
Haplotypes at the six GWAS peaks. Rows are phased haplotypes, two per individual, grouped by population and ordered as in the species tree (left; number of individuals in parentheses). The coloured strip gives each individual’s EOD phenotype, and within each individual the more BP-like haplotype is drawn on top. Columns are the up to 250 most strongly associated SNPs within each extended peak window (−log10(p) > 3; Methods) at which the BP and TP consensus haplotypes differ, in genome order. The BP allele is the consensus allele of BP individuals of the populations in which the BP allele arose (Bambomo for Peaks 1, 4 and 5; Bikagala, Bavavela and Biroundou for Peaks 2 and 3; Bikagala and Bavavela for Peak 6), and the TP allele that of Apassa, Bengue, Douvalou and Douengi TP individuals. Green bars mark SNPs inside each peak interval. Orange diamonds mark haplotypes outside those populations that are the same haplotype as one of their BP haplotypes (Methods).

**Fig. S5.**
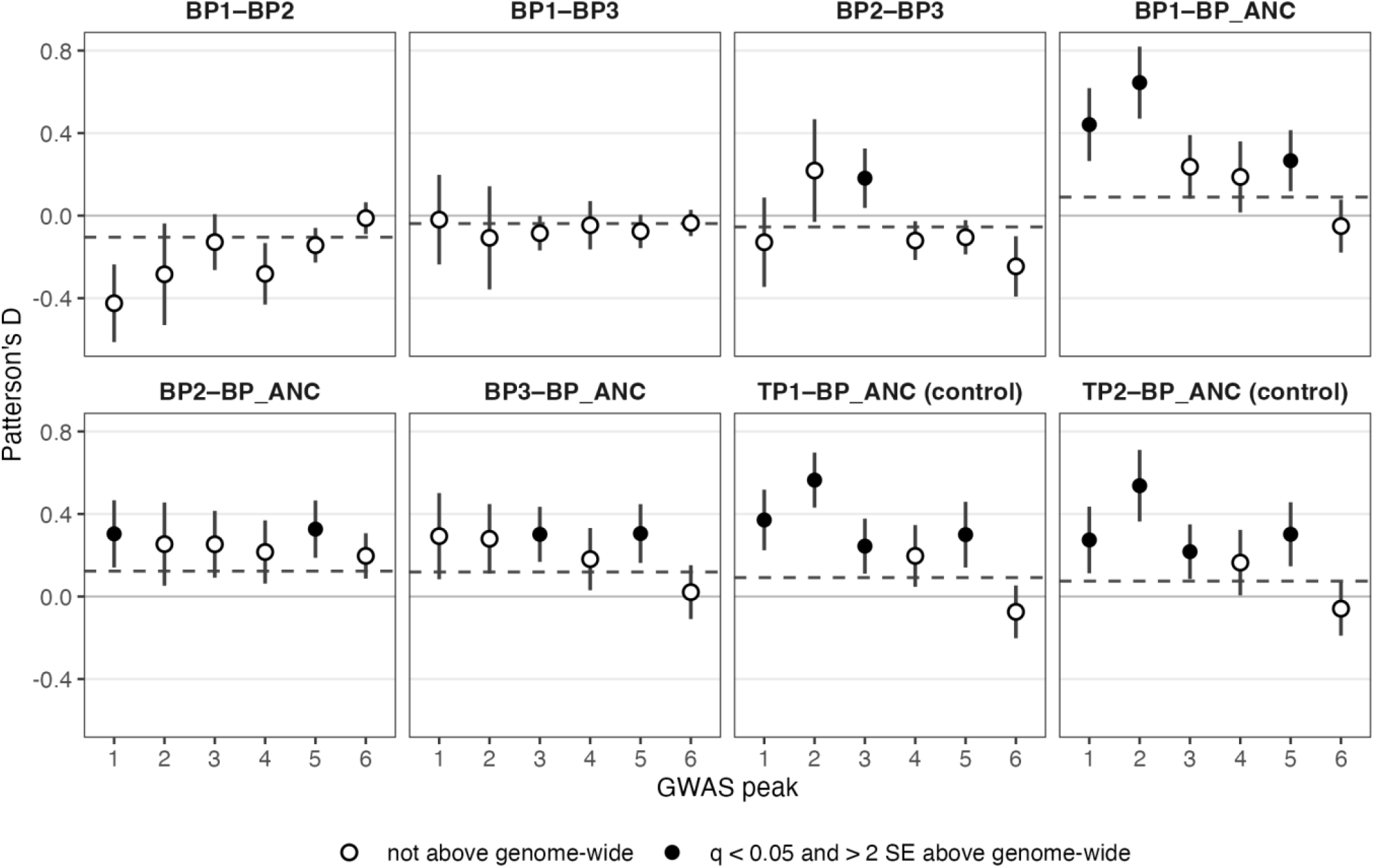
Patterson’s D at the six GWAS peaks. Each panel is one ABBA-BABA test; points are *D* within each extended peak interval with 95% block-jackknife intervals, and the dashed line is the genome-wide *D* for the same trio. *D* is signed so that positive values indicate excess allele sharing between the two named lineages relative to the reference: the triphasic sister for BP–BP tests, and triphasic *P. kingsleyae* from Mouvanga for tests against other biphasic *Paramormyrops* (BP_ANC). Filled points: *q* < 0.05 and more than two standard errors above the genome-wide value. BP1 shows no excess sharing with BP2 or BP3 at any peak. Excess sharing with BP_ANC at Peaks 1, 2, 3 and 5 is matched by triphasic *P. kingsleyae* (TP1 and TP2 controls), so it is not specific to BP lineages. The one BP-specific signal is BP2–BP3 at Peak 3, where both lineages carry the BP allele. Values are in Table S4 and Supplementary Data S7.

**Fig. S6.**
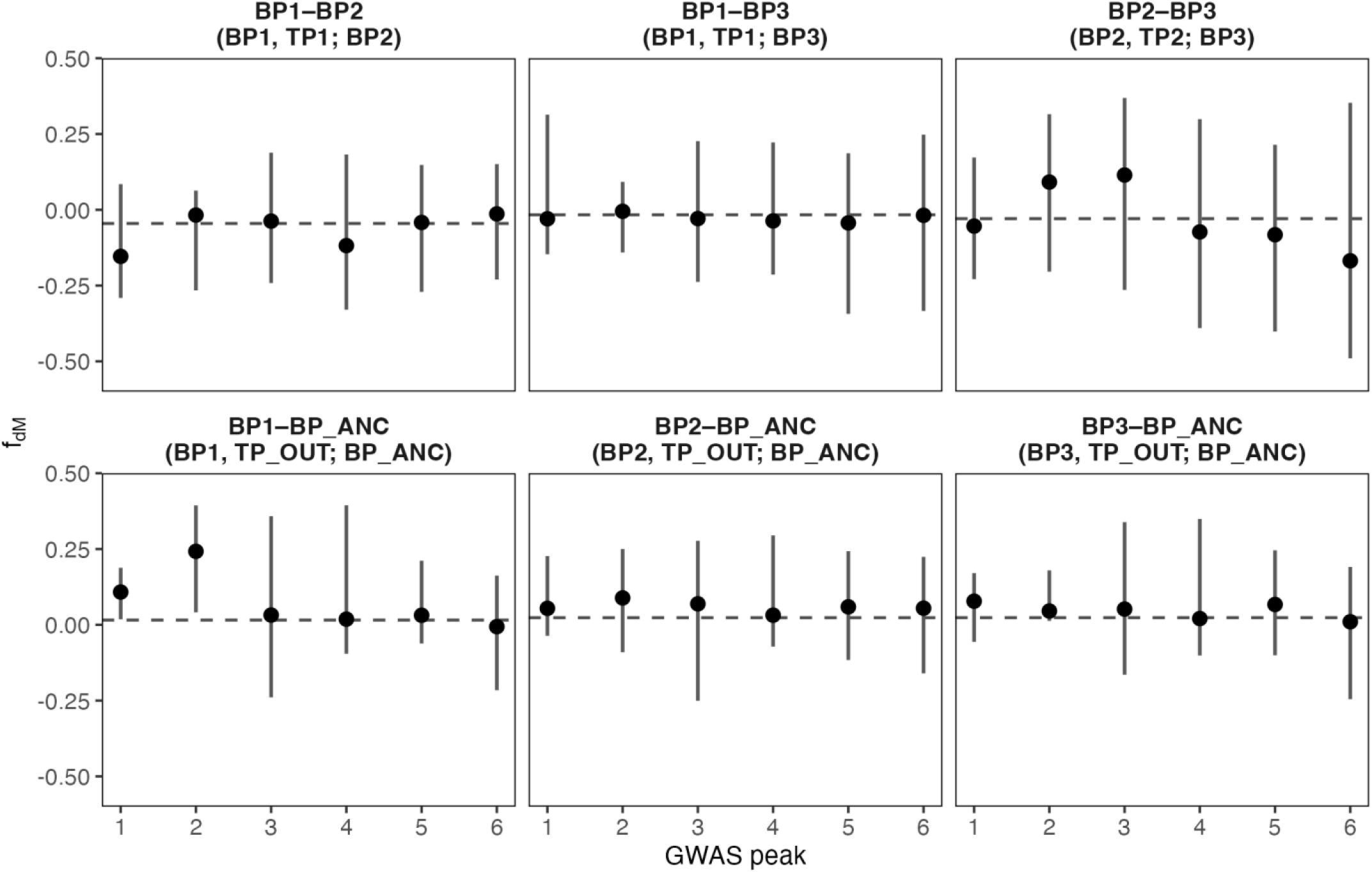
Windowed f_dM_ at the GWAS peaks. For each test (panel) and peak, the point is the median f_dM_ over windows of 50 informative SNPs overlapping the extended peak interval, and the bar spans the 2.5–97.5% quantiles of those windows. f_dM_ is signed so that positive values indicate sharing between the two named lineages; panel titles give the trio (P1, P2; P3). The dashed line is the genome-wide median outside all peaks. Peak medians lie between the 20th and 85th percentiles of background windows, except BP1–BP_ANC at Peak 2 (98th). Against medians of equal-sized blocks of background windows, BP1–BP_ANC at Peaks 1 and 2 and BP2–BP3 at Peak 3 exceed the 95th percentile; the first two are matched by triphasic *P. kingsleyae* in the *D* statistics (Fig. S5), and the third reflects the BP alleles that BP2 and BP3 share at Peak 3.

**Fig S7.**
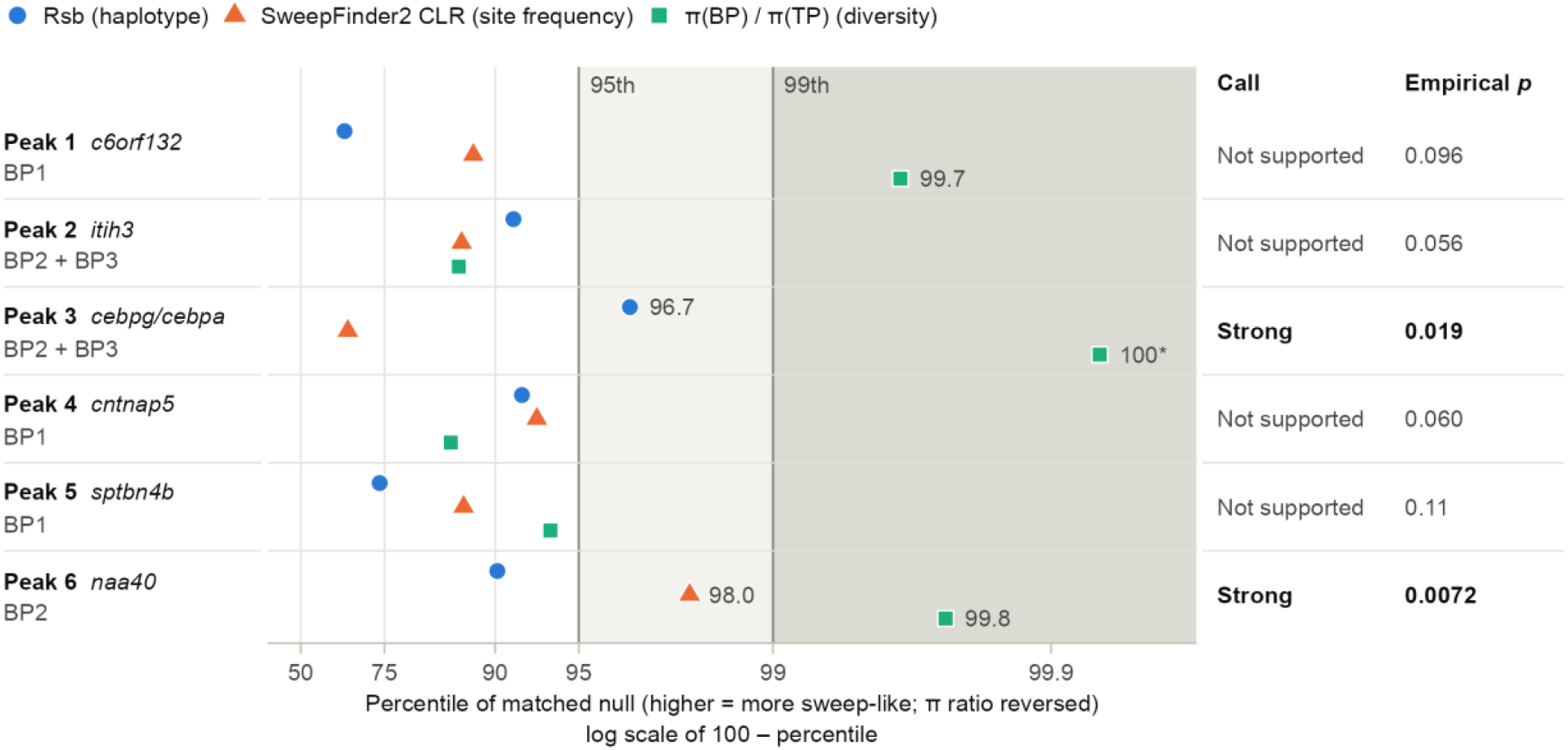
Calibrated tests for selective sweeps at the six GWAS peaks. Each peak was tested in the biphasic lineage in which its BP allele arose (BP1, Bambomo; BP2, Bikagala and Bavavela; BP3, Biroundou) against its triphasic reference (TP1, Apassa and Bengue; TP2, Douvalou and Douengi). Points give the percentile of each primary statistic among non-peak regions matched for SNP density and callable fraction (n = 1,074–1,518 per peak): Rsb between the BP lineage and its reference (haplotype length), the SweepFinder2 composite likelihood ratio (CLR; site-frequency spectrum) and π_BP_/π_TP_ (diversity; low values are sweep-like, so its percentile is reversed). Each statistic was summarised in the most extreme window within 100 kb of the index SNP, and the null regions were summarised in the same way. Where two lineages were tested (Peaks 2 and 3), the higher value is shown and the null takes the same maximum. The x axis is on a log scale of 100 − percentile; shading marks the 95th and 99th percentiles, and values are printed only at or above the 95th. A peak is supported when at least two statistics reach the 95th percentile, and strong when one of them also reaches the 99th. The empirical p is the fraction of matched null regions whose composite score (the second-highest of the three percentiles) is at least as high as the peak’s. Peak 1 is not supported despite a π ratio at the 99.7th percentile, because only one statistic reaches the 95th. *No matched null region had a lower π ratio than Peak 3 (n = 1,497). Applied to the null regions, the rule calls 1.3–2.9% supported and 0.5–1.7% strong. No correction across the six peaks is applied here, and sensitivity analyses are given in Table S5. In post hoc checks, a Benjamini–Hochberg correction^53^ across the six peaks leaves Peak 6 (q = 0.043; Peak 3, q = 0.056). With 40-kb windows centred on the index SNP, rather than chosen within ± 100 kb, only Peak 3 is called strong (p = 0.006). Peak 6 is not supported (p = 0.18), because its CLR and diversity-ratio signals lie 43–45 kb from the index SNP.

**Table S1.** GWAS peaks. Peaks correspond to those in Fig. 3A,B. Start and End are the outermost SNPs with −log10(p) > 5 in each peak (see Association Analysis). n SNPs gives the number of SNPs above the significance and suggestive permutation thresholds (−log10(p) > 7.22 and > 6.36) within these bounds. Genes are all annotated genes (NCBI RefSeq annotation GCF_048594095.1-RS_2025_03) that overlap the peak or lie within 25 kb of it (†, within 25 kb only). Symbol gives the gene name used in the main text; Description is the RefSeq description.

| Peak | Chr. | Start | End | n SNPs | Gene | Symbol | Description |
|---|---|---|---|---|---|---|---|
| 1 | chr6 | 29,492,938 | 29,509,385 | 12 / 13 | frs3 † | *frs3* | fibroblast growth factor receptor substrate 3 |
|  |  |  |  |  | LOC111844745 | *c6orf132* | uncharacterized LOC111844745 |
|  |  |  |  |  | LOC111844692 † | *elf3* | ETS-related transcription factor Elf-3-like |
| 2 | chr8 | 15,781,018 | 15,801,982 | 24 / 28 | rft1 † | *rft1* | RFT1 homolog |
|  |  |  |  |  | LOC111839880 | *sfmbt1* | scm-like with four MBT domains protein 1 |
|  |  |  |  |  | LOC111841399 | *itih3* | inter-alpha-trypsin inhibitor heavy chain H3-like |
| 3 | chr13 | 24,683,587 | 24,784,771 | 8 / 11 | LOC111843522 | — | nuclear factor of activated T-cells, cytoplasmic 3-like |
|  |  |  |  |  | LOC111843545 | — | FH1/FH2 domain-containing protein 1-like |
|  |  |  |  |  | LOC111843556 | — | 11-beta-hydroxysteroid dehydrogenase type 2-like |
|  |  |  |  |  | agrp | *agrp* | agouti related neuropeptide |
|  |  |  |  |  | LOC111843547 | *ctra* | chymotrypsin A-like |
|  |  |  |  |  | LOC111843498 | *cebpg* | CCAAT/enhancer-binding protein gamma-like |
|  |  |  |  |  | LOC111843507 | *cebpa* | CCAAT/enhancer-binding protein alpha-like |
|  |  |  |  |  | LOC111843506 † | *slc7a10* | asc-type amino acid transporter 1-like |
|  |  |  |  |  | LOC111843516 † | *rnf182* | E3 ubiquitin-protein ligase RNF182-like |
|  |  |  |  |  | LOC111843559 † | *slc7a9* | b(0,+)-type amino acid transporter 1-like |
| 4 | chr16 | 10,861,379 | 11,021,498 | 18 / 27 | LOC111845409 † | *cntnap5* | contactin-associated protein-like 5 |
|  |  |  |  |  | LOC111845410 | *gypc* | glycophorin-C-like |
|  |  |  |  |  | trnav-uac | *trnav-uac* | transfer RNA valine (anticodon UAC) |
|  |  |  |  |  | ndufa10 † | *ndufa10* | NADH:ubiquinone oxidoreductase subunit A10 |
| 5 | chr17 | 11,625,395 | 11,736,085 | 12 / 21 | foxa3 † | *foxa3* | forkhead box A3 |
|  |  |  |  |  | pld3 | *pld3* | phospholipase D family member 3 |
|  |  |  |  |  | LOC111844037 | — | mRNA decay activator protein ZFP36L1 |
|  |  |  |  |  | shkbp1 | *shkbp1* | SH3KBP1 binding protein 1 |
|  |  |  |  |  | snrnp35 | *snrnp35* | small nuclear ribonucleoprotein 35 (U11/U12) |
|  |  |  |  |  | sptbn4b | *sptbn4b* | spectrin, beta, non-erythrocytic 4b |
|  |  |  |  |  | blvrb † | *blvrb* | biliverdin reductase B |
|  |  |  |  |  | kcnk6 † | *kcnk6* | potassium channel, subfamily K, member 6 |
| 6 | chr24 | 8,596,847 | 8,660,182 | 5 / 7 | LOC111842864 † | *dao-5* | uncharacterized LOC111842864 |
|  |  |  |  |  | naa40 | *naa40* | N-alpha-acetyltransferase 40, NatD catalytic subunit |
|  |  |  |  |  | LOC111835120 | — | REST corepressor 2-like |
|  |  |  |  |  | LOC111835121 | — | uncharacterized LOC111835121 |
|  |  |  |  |  | mark2a † | *mark2a* | MAP/microtubule affinity-regulating kinase 2a |

**Data S1. (separate file) Listing of all specimens used in this analysis.** Excel formatted listing of all specimens used in this analysis, along with associated metadata, museum accession numbers (where available) and associated NCBI accession numbers for short reads.

**Data S2. (separate file) Plot of all EOD recordings used in this analysis.** A ZIP archive of PNG images corresponding to the EODs of individuals listed in Data File S1, where available.

**Data S3. (separate file) Listing of all publicly available EOD recordings used in this analysis.** Excel formatted listing of all EOD recordings used in this analysis, along with accession numbers or citations where available, as well as assigned phenotype.

**Data S4. (separate file) Candidate variants and predicted motif changes at Peak 4 (*cntnap5*). Tab-delimited. One row per variant in linkage disequilibrium with the index SNP (r² ≥ 0.8) and per predicted motif change (≥ ten-fold difference between alleles, best allele p < 10⁻⁴). Variants without a predicted change appear once, with empty motif fields. The columns give position, alleles and which allele is BP- or TP-associated; r² with the index SNP and genotype call rate; allele frequencies in BP and TP fish and the association −log10(p); annotation class, nearest gene and distance to its transcription start site; the frequency of the phased local haplotype used; and the motif ID, name and JASPAR family, the predicted p on each allele, the log10 fold difference and the allele on which the motif is present.**

**Data S5. (separate file) As Data S4, for Peak 5 (*sptbn4b*).**

**Data S6. (separate file) As Data S4, for Peak 2 (*itih3*).**

**Data S7. (separate file) ABBA-BABA statistics at the GWAS peaks. Patterson’s *D*, *Z*, *P* and *q* values, f₄-ratio, genome-wide *D* and informative-site counts for each test and peak (Dsuite Dtrios; see Analysis of Introgression). The last six rows are the targeted test at Peak 6 against *P. curvifrons*, *P. ntotom* and *P.* sp. ’SN2’ (CURV3), with q values adjusted across those six tests.**

## Supplemental Note: Genotypes of mixed-phenotype individuals

### Sample and phenotype

“Mixed” individuals have electric organs containing both penetrating, anteriorly innervated electrocytes and non-penetrating, posteriorly innervated electrocytes^1,3^. Their averaged EODs carry a small P0 phase (Fig. 1B; Data S2). We classified 42 individuals as mixed from their EODs: 25 from Bambomo, 12 from Apassa, 3 from Bikagala and 2 from Bavavela. These individuals were excluded from the GWAS. The EOD boundary between mixed and BP is imperfect. One Bambomo individual with histologically confirmed mosaic electrocytes (BAM_6494) produced an EOD indistinguishable from BP and was analysed as BP.

### Genotypes at the GWAS peaks

We defined canonical BP and TP haplotypes for each peak from phased genotypes of reference individuals. The BP reference was Bambomo BP individuals for Peaks 1, 4 and 5, Bikagala, Bavavela and Biroundou BP individuals for Peaks 2 and 3, and Bikagala and Bavavela BP individuals for Peak 6. The TP reference was TP individuals from Apassa, Bengue, Douvalou and Douengi. Mixed individuals were not used as references. At SNPs where the two reference consensus haplotypes differed, we assigned each phased haplotype of every individual to the closer consensus. This gave one diploid state per peak: BP homozygous, heterozygous or TP homozygous (Table S2).

In Bambomo, 23 of 25 mixed individuals carried a BP haplotype at all three Bambomo-origin peaks, as did all 41 BP individuals. The exceptions were BAM_169 (TP homozygous at Peak 4) and BAM_247 (TP homozygous at Peak 5). No BP individual lacked a BP haplotype at Peak 4 or 5 (0 of 41 vs 2 of 25 mixed; Fisher’s exact test, p = 0.14). All Bambomo individuals were BP homozygous at Peak 1. Mixed and BP individuals did not differ in heterozygosity at Peak 4 (4 of 25 vs 4 of 41; p = 0.47), at Peak 5 (10 of 25 vs 15 of 41; p = 0.80) or at either (12 of 25 vs 18 of 41; p = 0.80). Twelve mixed individuals (of 25) were BP homozygous at all three Bambomo peaks, against 23 of 41 BP individuals (p = 0.61).

All five Bikagala and Bavavela mixed individuals carried BP genotypes at Peaks 2, 3 and 6, like the BP individuals of those populations; one was heterozygous at Peak 2. Apassa mixed individuals resembled Apassa TP individuals at all six peaks. One of them (APA_6681) carried a single copy of the Bambomo BP haplotype at Peak 1, which no other Apassa individual carried.

Seven individuals had histologically confirmed electrocyte anatomy^1,3^.

- The two mosaic individuals carried the genotypes of their populations. BAM_6494 was BP homozygous at Peaks 1, 4 and 5. APA_6681 carried one copy of the Bambomo Peak 1 haplotype and no Bambomo or Bikagala/Bavavela BP haplotype at the other peaks.

- The only TP individual sampled in Bambomo (BAM_6597, histologically TP) was TP homozygous at Peaks 4 and 5 and BP homozygous at Peak 1.

- The histologically BP Bambomo individual (BAM_6497) was BP homozygous at Peaks 1, 4 and 5.

- The three histologically TP Apassa individuals were TP homozygous at every peak, except that APA_6675 carried one recombinant Peak 4 haplotype that matches no Bambomo haplotype (see Dominance).

### Dominance

In Bambomo, every BP individual carried at least one BP haplotype at both Peaks 4 and 5, and 18 of 41 were heterozygous at one or both, so the BP alleles behave as dominant (main text). If heterozygosity at these loci produced the mixed phenotype, mixed individuals would be enriched for heterozygotes. They are not (above), so the mixed phenotype is not a heterozygote class. Outside Bambomo, a single copy of a Bambomo BP haplotype at Peak 4 occurs in 2 of 18 Apassa and 4 of 17 Bengue TP individuals, always without a BP haplotype at Peak 5. One of these copies is identical to a sampled Bambomo haplotype; the other five differ from the nearest at one or two SNPs, at alleles that other Bambomo haplotypes carry. Six further Apassa and Bengue TP individuals carry recombinant Peak 4 haplotypes that a two-class assignment would call BP but that match no Bambomo haplotype. Peak 4 carries several haplotypes (Fig. S4) that our two-class assignment groups together, and one copy of a Bambomo BP haplotype at Peak 4 alone is not sufficient for a BP EOD.

### Ancestry

We estimated ancestry with ADMIXTURE on 40,758 LD-thinned SNPs (MAF ≥ 5%, ≤ 20% missing data, one SNP per 20 kb; 240 P. kingsleyae). Cross-validation error was lowest at K = 8 (0.2307), closely followed by K = 10 (0.2309) (Fig. S8). For each individual, we summed the ancestry assigned to clusters characteristic of populations with the opposite EOD phenotype.

- Mixed individuals from Bambomo, Bikagala and Bavavela had none (< 0.01%) at either K, like BP individuals from the same populations.

- Apassa mixed individuals carried a small Bambomo-cluster component at K = 8: median 7%, maximum 14%, against a median below 0.01% in Apassa TP individuals.

- At K = 10 that component was absent (maximum 0.8%). Apassa instead split between an Apassa cluster and a Bengue-like cluster.

No mixed individual carried more than 14% opposite-phenotype ancestry at either K. An F1 or F2 hybrid would carry about 50%, and a first-generation backcross about 25%. In an exploratory follow-up of the Apassa component, we estimated local ancestry along phased chromosomes and, as a check, projected unphased genotypes (depth ≥ 8, genotype quality ≥ 20) onto Bambomo and Apassa–Bengue allele frequencies. The Bambomo-related component was concentrated in the seven mixed individuals collected at one site in 2019. Their excess over 2009 Apassa TP individuals was 9–13 percentage points, detectable on 14–20 of the 25 chromosomes rather than in a few blocks, and not concentrated at the GWAS peaks; the two methods agreed genome-wide (r = 0.93). The three 2019 mixed individuals from two other sites, and the two collected in 2009, were near the TP baseline. The site’s excess is about what one Bambomo great-grandparent (12.5%) or great-great-grandparent (6.25%) would contribute; a Bambomo grandparent would contribute 25%, as in a first backcross. Tract lengths, which could separate such recent admixture from older gene flow, depend on phasing and were not used to date it. No TP individuals from that site were sequenced, so this excess cannot be separated from collection site, year or sequencing batch, and we do not interpret it as a cause of the mixed phenotype.

### Association scans

To avoid confounding phenotype with population, we scanned Bambomo alone: 25 mixed vs 41 BP individuals, GEMMA Wald test with the relatedness matrix, thresholds from 100 phenotype permutations. No SNP reached the suggestive threshold (top −log10 p = 6.29; suggestive 6.78; significant 7.69). Peaks 1, 4 and 5 showed no association (all p > 10⁻³).

To ask what this scan could detect, we planted one causal SNP, drawn from the observed Bambomo genotypes, and sampled 25 cases among the 66 individuals with weight OR^dosage. We tested each planted SNP with GEMMA and the observed relatedness matrix, and scored it against the permutation threshold (100 replicates per odds ratio and frequency bin).

- Even with error-free phenotypes, power reached 80% only for common variants with very large effects: per-allele odds ratio ≥ 20 at minor allele frequency 0.30–0.50 (85%), and odds ratio 50 at 0.15–0.30 (86%; 79% at odds ratio 20).

- For rarer variants (0.05–0.15), power was 29% even at an odds ratio of 50.

- At odds ratios ≤ 3, power was ≤ 3% in every bin. Misclassified EODs would lower power further.

The scan could therefore have detected only near-Mendelian loci for the mixed phenotype.

An earlier pooled scan compared all 42 mixed individuals with all BP individuals across populations (likelihood-ratio test, thresholds from 100 permutations). One peak exceeded the significance threshold, on chromosome 3 at 17.35–17.51 Mb (lead SNP chr3:17,486,697; −log10 p = 8.43 vs 8.20). Two exceeded the suggestive threshold (7.28): chromosome 20 at 4.69–5.15 Mb (8.11) and chromosome 24 at 11.83–12.15 Mb (7.50).

- The chromosome 3 peak partly reflects allele-frequency differences among populations. Its alternative allele is fixed or nearly fixed (≥ 97%) in the Bikagala, Bavavela and Biroundou BP populations and segregates in Apassa, Bengue and Bambomo (31–56%).

- Within Bambomo, the same allele is depleted in mixed individuals (0.08 vs 0.45 in BP individuals), but the region does not reach genome-wide significance in the within-Bambomo scan (best p = 1.3 × 10⁻⁵).

- Within Apassa, the difference is in the same direction but small (0.25 vs 0.36; Fisher’s exact p = 0.41).

### Interpretation

Mixed individuals carry the peak genotypes typical of their population: BP genotypes in Bambomo, Bikagala and Bavavela, and TP genotypes in Apassa. They are not enriched for heterozygotes at the GWAS loci, and none is a first-generation hybrid or first backcross. The mixed phenotype therefore arises on both genetic backgrounds, consistent with partial penetrance of electrocyte penetration at the level of individual electrocytes. Whether an individual is mixed depends on variation outside the six GWAS loci, or below the resolution of our sample. That variation may include loci of smaller effect, developmental noise or environmental effects. These results support, rather than challenge, the conclusion that the six GWAS peaks are the major loci underlying the BP/TP difference in the sampled populations.

**Table S2.** Genotypes at the six GWAS peaks in populations containing mixed individuals. Each cell gives the number of individuals that are TP homozygous / heterozygous / BP homozygous for the peak’s canonical haplotypes, from phased genotypes. Peaks 1, 4 and 5 are of Bambomo origin, Peaks 2 and 3 of Bikagala/Bavavela/Biroundou origin, and Peak 6 of Bikagala/Bavavela origin. The Bambomo TP individual (BAM_6597) is histologically TP. APA_223 was collected in Bambomo and is counted there, despite its ID prefix. Genotypes are two-class calls: each haplotype is assigned to whichever of the BP and TP consensus haplotypes it is nearer. A BP-class call does not mean the haplotype is the same as those of the BP populations; Fig. S4 and the Dominance section give the haplotype-level comparison.

| **Population** | **EOD** | **n** | **Peak 1** | **Peak 2** | **Peak 3** | **Peak 4** | **Peak 5** | **Peak 6** |
|---|---|---|---|---|---|---|---|---|
| Bambomo | BP | 41 | 0/0/41 | 41/0/0 | 41/0/0 | 0/4/37 | 0/15/26 | 41/0/0 |
| Bambomo | mixed | 25 | 0/0/25 | 25/0/0 | 25/0/0 | 1/4/20 | 1/10/14 | 25/0/0 |
| Bambomo | TP | 1 | 0/0/1 | 1/0/0 | 1/0/0 | 1/0/0 | 1/0/0 | 1/0/0 |
| Apassa | TP | 18 | 18/0/0 | 16/1/1 | 18/0/0 | 12/6/0 | 18/0/0 | 18/0/0 |
| Apassa | mixed | 12 | 11/1/0 | 11/1/0 | 12/0/0 | 10/2/0 | 12/0/0 | 12/0/0 |
| Bikagala | BP | 27 | 27/0/0 | 0/0/27 | 0/0/27 | 27/0/0 | 27/0/0 | 0/0/27 |
| Bikagala | mixed | 3 | 3/0/0 | 0/0/3 | 0/0/3 | 3/0/0 | 3/0/0 | 0/0/3 |
| Bavavela | BP | 21 | 21/0/0 | 0/0/21 | 0/0/21 | 21/0/0 | 21/0/0 | 0/0/21 |
| Bavavela | mixed | 2 | 2/0/0 | 0/1/1 | 0/0/2 | 2/0/0 | 2/0/0 | 0/0/2 |

**Table S3.**
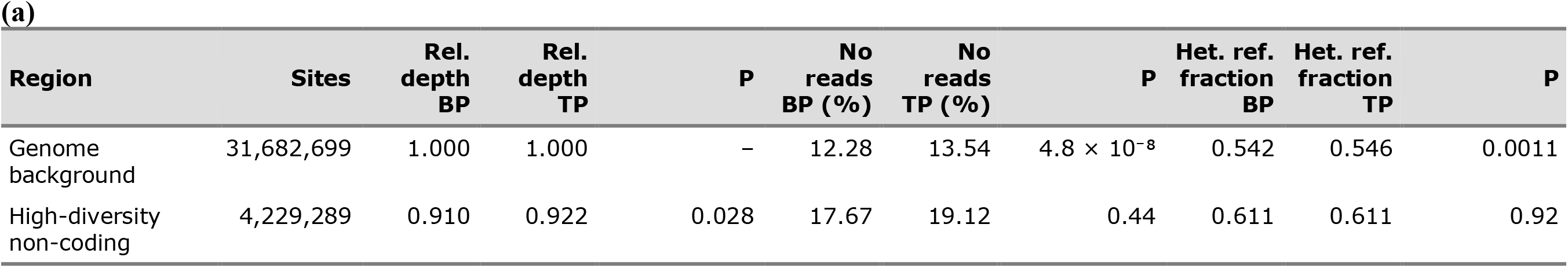
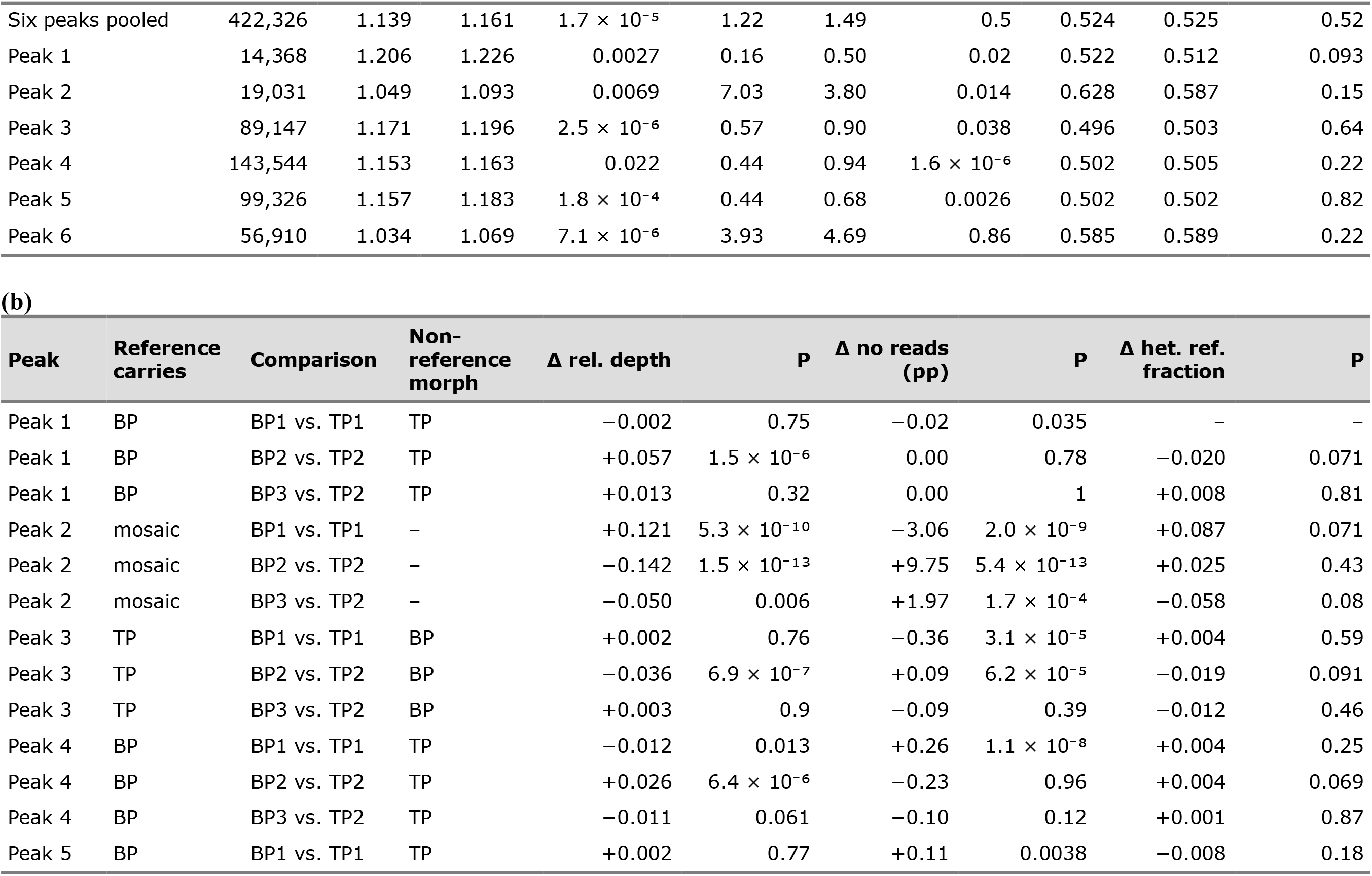
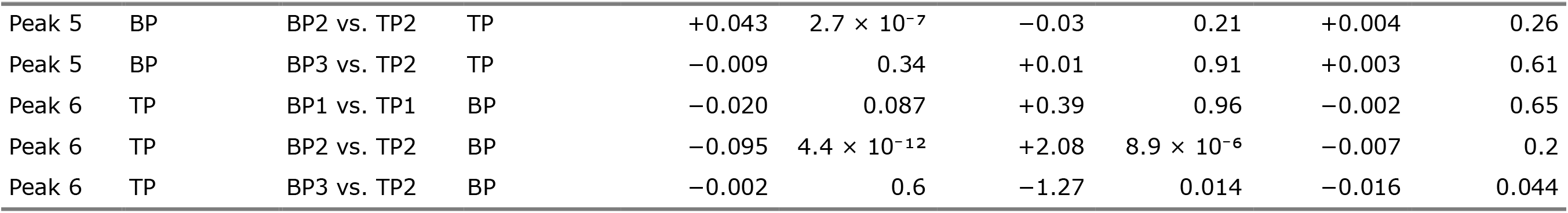
Mapping proxies for reference bias in BP and TP individuals. Values come from the raw all-sites VCF for the 198 GWAS individuals (109 BP, 89 TP). Rel. depth is mean read depth divided by the individual’s mean depth in the genome background; no reads is the percentage of sites without reads; het. ref. fraction is the mean fraction of reference reads at heterozygous biallelic SNPs with depth ≥ 10. P values are from two-sided Wilcoxon rank-sum tests between BP and TP individuals. (a) All GWAS individuals, by region. Genome background: 2-kb blocks every 50 kb; high-diversity non-coding: 1,000 5-kb windows from the top 5% of nucleotide diversity, without annotated genes. (b) Each peak within the three geographically matched comparisons: BP1 vs. TP1, Bambomo BP vs. Apassa and Bengue TP; BP2 vs. TP2, Bikagala and Bavavela BP vs. Douvalou and Douengi TP; BP3 vs. TP2, Biroundou BP vs. Douvalou and Douengi TP. Reference carries gives the phenotype-associated haplotype present in the reference assembly. Δ is the non-reference morph minus the reference-matching morph (at Peak 2, where the reference is a mosaic, BP minus TP); pp, percentage points. Reference bias would give a negative Δ rel. depth and positive Δ no reads and Δ het. ref. fraction. Dashes mark values that could not be computed (fewer than 20 heterozygous sites per individual).

| (a) |  |  |  |  |  |  |  |  |  |  |
|---|---|---|---|---|---|---|---|---|---|---|
| **Region** | **Sites** | **Rel. depth BP** | **Rel. depth TP** | **P** | **No reads BP (%)** | **No reads TP (%)** | **P** | **Het. ref. fraction BP** | **Het. ref. fraction TP** | **P** |
| Genome background | 31,682,699 | 1.000 | 1.000 | – | 12.28 | 13.54 | 4.8 × 10<sup>−8</sup> | 0.542 | 0.546 | 0.0011 |
| High-diversity non-coding | 4,229,289 | 0.910 | 0.922 | 0.028 | 17.67 | 19.12 | 0.44 | 0.611 | 0.611 | 0.92 |
| Six peaks pooled | 422,326 | 1.139 | 1.161 | 1.7 × 10<sup>−5</sup> | 1.22 | 1.49 | 0.5 | 0.524 | 0.525 | 0.52 |
| Peak 1 | 14,368 | 1.206 | 1.226 | 0.0027 | 0.16 | 0.50 | 0.02 | 0.522 | 0.512 | 0.093 |
| Peak 2 | 19,031 | 1.049 | 1.093 | 0.0069 | 7.03 | 3.80 | 0.014 | 0.628 | 0.587 | 0.15 |
| Peak 3 | 89,147 | 1.171 | 1.196 | 2.5 × 10<sup>−6</sup> | 0.57 | 0.90 | 0.038 | 0.496 | 0.503 | 0.64 |
| Peak 4 | 143,544 | 1.153 | 1.163 | 0.022 | 0.44 | 0.94 | 1.6 × 10<sup>−6</sup> | 0.502 | 0.505 | 0.22 |
| Peak 5 | 99,326 | 1.157 | 1.183 | 1.8 × 10<sup>−4</sup> | 0.44 | 0.68 | 0.0026 | 0.502 | 0.502 | 0.82 |
| Peak 6 | 56,910 | 1.034 | 1.069 | 7.1 × 10<sup>−6</sup> | 3.93 | 4.69 | 0.86 | 0.585 | 0.589 | 0.22 |
| (b) |  |  |  |  |  |  |  |  |  |  |
| **Peak** | **Reference carries** | **Comparison** | **Non-reference morph** | **Δ rel. depth** | **P** | **Δ no reads (pp)** | **P** | **Δ het. ref. fraction** | **P** |  |
| Peak 1 | BP | BP1 vs. TP1 | TP | −0.002 | 0.75 | −0.02 | 0.035 | – | – |  |
| Peak 1 | BP | BP2 vs. TP2 | TP | +0.057 | 1.5 × 10<sup>−6</sup> | 0.00 | 0.78 | −0.020 | 0.071 |  |
| Peak 1 | BP | BP3 vs. TP2 | TP | +0.013 | 0.32 | 0.00 | 1 | +0.008 | 0.81 |  |
| Peak 2 | mosaic | BP1 vs. TP1 | – | +0.121 | 5.3 × 10<sup>−10</sup> | −3.06 | 2.0 × 10<sup>−9</sup> | +0.087 | 0.071 |  |
| Peak 2 | mosaic | BP2 vs. TP2 | – | −0.142 | 1.5 × 10<sup>−13</sup> | +9.75 | 5.4 × 10<sup>−13</sup> | +0.025 | 0.43 |  |
| Peak 2 | mosaic | BP3 vs. TP2 | – | −0.050 | 0.006 | +1.97 | 1.7 × 10<sup>−4</sup> | −0.058 | 0.08 |  |
| Peak 3 | TP | BP1 vs. TP1 | BP | +0.002 | 0.76 | −0.36 | 3.1 × 10<sup>−5</sup> | +0.004 | 0.59 |  |
| Peak 3 | TP | BP2 vs. TP2 | BP | −0.036 | 6.9 × 10<sup>−7</sup> | +0.09 | 6.2 × 10<sup>−5</sup> | −0.019 | 0.091 |  |
| Peak 3 | TP | BP3 vs. TP2 | BP | +0.003 | 0.9 | −0.09 | 0.39 | −0.012 | 0.46 |  |
| Peak 4 | BP | BP1 vs. TP1 | TP | −0.012 | 0.013 | +0.26 | 1.1 × 10<sup>−8</sup> | +0.004 | 0.25 |  |
| Peak 4 | BP | BP2 vs. TP2 | TP | +0.026 | 6.4 × 10<sup>−6</sup> | −0.23 | 0.96 | +0.004 | 0.069 |  |
| Peak 4 | BP | BP3 vs. TP2 | TP | −0.011 | 0.061 | −0.10 | 0.12 | +0.001 | 0.87 |  |
| Peak 5 | BP | BP1 vs. TP1 | TP | +0.002 | 0.77 | +0.11 | 0.0038 | −0.008 | 0.18 |  |
| Peak 5 | BP | BP2 vs. TP2 | TP | +0.043 | 2.7 × 10<sup>−7</sup> | −0.03 | 0.21 | +0.004 | 0.26 |  |
| Peak 5 | BP | BP3 vs. TP2 | TP | −0.009 | 0.34 | +0.01 | 0.91 | +0.003 | 0.61 |  |
| Peak 6 | TP | BP1 vs. TP1 | BP | −0.020 | 0.087 | +0.39 | 0.96 | −0.002 | 0.65 |  |
| Peak 6 | TP | BP2 vs. TP2 | BP | −0.095 | 4.4 × 10<sup>−12</sup> | +2.08 | 8.9 × 10<sup>−6</sup> | −0.007 | 0.2 |  |
| Peak 6 | TP | BP3 vs. TP2 | BP | −0.002 | 0.6 | −1.27 | 0.014 | −0.016 | 0.044 |  |

**Table S4.** ABBA-BABA statistics at the GWAS peaks. Patterson’s *D* (*Z* in parentheses) within each extended peak interval, signed so that positive values indicate excess sharing between the named lineages, and genome-wide *D* for the same trio (all genome-wide |*Z*| ≥ 14). † *q* < 0.05 and more than two standard errors above the genome-wide value. BP_ANC, other biphasic *Paramormyrops*; the TP1 and TP2 rows are controls. f₄-ratios, *P* and *q* values and site counts are in Supplementary Data S7.

| Test | Peak 1 | Peak 2 | Peak 3 | Peak 4 | Peak 5 | Peak 6 | Genome-wide D |
|---|---|---|---|---|---|---|---|
| BP1–BP2 | -0.42 (-4.4) | -0.28 (-2.3) | -0.13 (-1.9) | -0.28 (-3.7) | -0.14 (-3.4) | -0.01 (-0.3) | -0.104 |
| BP1–BP3 | -0.02 (-0.2) | -0.11 (-0.8) | -0.09 (-2.0) | -0.05 (-0.8) | -0.08 (-1.8) | -0.04 (-1.1) | -0.038 |
| BP2–BP3 | -0.13 (-1.2) | 0.22 (1.7) | 0.18 (2.5)† | -0.12 (-2.5) | -0.10 (-2.5) | -0.25 (-3.3) | -0.055 |
| BP1–BP_ANC | 0.44 (4.9)† | 0.65 (7.2)† | 0.24 (3.0) | 0.19 (2.1) | 0.27 (3.5)† | -0.05 (-0.8) | 0.090 |
| BP2–BP_ANC | 0.30 (3.7)† | 0.25 (2.5) | 0.25 (3.1) | 0.22 (2.8) | 0.33 (4.6)† | 0.20 (3.5) | 0.123 |
| BP3–BP_ANC | 0.29 (2.8) | 0.28 (3.2) | 0.30 (4.4)† | 0.18 (2.4) | 0.30 (4.2)† | 0.02 (0.3) | 0.119 |
| TP1–BP_ANC (control) | 0.37 (4.9)† | 0.56 (8.3)† | 0.24 (3.6)† | 0.20 (2.6) | 0.30 (3.7)† | -0.07 (-1.1) | 0.092 |
| TP2–BP_ANC (control) | 0.27 (3.3)† | 0.54 (6.1)† | 0.22 (3.2)† | 0.16 (2.0) | 0.30 (3.8)† | -0.06 (-0.9) | 0.075 |

**Table S5.** Calibrated tests for selective sweeps at the GWAS peaks. Percentile of each primary statistic among non-peak regions matched for SNP density and callable fraction (n null), each summarised in the most extreme window within 100 kb of the index SNP (see Tests for Selection at GWAS Peaks). π_BP_/π_TP_ is reversed so that high percentiles are sweep-like. Where two lineages were tested, values are the maximum over both, as in the null. π ratio, π_BP_/π_TP_. TP CLR, SweepFinder2 CLR percentile in the matched triphasic reference. Composite, the second-highest of the three percentiles; empirical p, the fraction of matched null regions with a composite at least as high. Strong, the number of the eleven configurations (the headline and ten sensitivity analyses) in which the peak was called strong. The ten sensitivity analyses, each recomputed for peaks and null together, were a ± 50-kb search region; 40-kb windows; the unmatched null; a null also matched on BP–TP differentiation; a null stratified by a recombination proxy (mean inES of lineages outside the contrast); Rsb standardized genome-wide; all haplotype statistics after down-sampling every lineage to 16 individuals; the CLR on the folded spectrum; the extended peak interval in place of the fixed search region and window; and, as an exploratory analysis, XP-nSL^54^ in place of Rsb. Peak 3 is not supported with a ± 50-kb search region or with XP-nSL in place of Rsb; Peak 6 is supported but not strong over the extended peak interval; Peak 1 is supported, but not strong, only over the extended peak interval. *No matched null region had a lower π ratio. No correction across the six peaks is applied.

| **Peak** | **Gene** | **Lineages** | **Rsb** | **CLR** | **π ratio** | **TP CLR** | **Composite** | **Empirical p** | **Call** | **Strong** | **n null** |
|---|---|---|---|---|---|---|---|---|---|---|---|
| 1 | *LOC111844745* | BP1 | 65.2 | 88.0 | 99.7 | 50.5 | 88.0 | 0.096 | not supported | 0 of 11 | 1,436 |
| 2 | *itih3* | BP2 + BP3 | 91.4 | 86.8 | 86.5 | 95.4 | 86.8 | 0.056 | not supported | 0 of 11 | 1,115 |
| 3 | *cebpg/cebpa* | BP2 + BP3 | 96.7 | 66.1 | 100.0* | 23.8 | 96.7 | 0.019 | strong | 9 of 11 | 1,497 |
| 4 | *cntnap5* | BP1 | 92.0 | 92.9 | 85.6 | 56.9 | 92.0 | 0.060 | not supported | 0 of 11 | 1,074 |
| 5 | *sptbn4b* | BP1 | 74.0 | 87.0 | 93.7 | 69.8 | 87.0 | 0.107 | not supported | 0 of 11 | 1,518 |
| 6 | *naa40* | BP2 | 90.2 | 98.0 | 99.8 | 84.3 | 98.0 | 0.0072 | strong | 10 of 11 | 1,253 |

**Table S6.** Drift test at the GWAS index SNPs. For each BP population and peak (see Drift Test at GWAS Peaks): Δ, the BP–TP frequency difference of the derived allele at the index SNP (label-free analyses count differentiation in either direction); O, SNPs outside the unit’s own peak ± 250 kb that are at least as differentiated; E, the number expected genome-wide under the best admissible model (M2, split with asymmetric migration; M3, isolation with migration with a founder bottleneck and growth; f, folded spectrum); E 95% interval, from 100 block-bootstrap refits; E robust, the largest of the upper bootstrap bound, the point estimate under every admissible model and the estimate allowing local reductions in effective population size. Readings were fixed in advance: E robust ≤ 0.005, drift implausible across all units; 0.005–0.05, implausible for this unit; 0.05–1, unusual under drift; 1–20, drift makes a few such sites; > 20, drift readily makes this. By Markov’s inequality, the probability that drift produces at least one such SNP is at most E, whatever the linkage among sites. Where no model was admissible, O is reported.

| **Population** | **Peak** | **Analysis** | **Δ** | **O (non-peak)** | **E, best model** | **E 95% interval** | **E robust** | **Reading** |
|---|---|---|---|---|---|---|---|---|
| BAM | Peak 1 | polarized | 0.989 | 0 | 0.046 (M2) | 0.032–0.056 | 4.16 | drift makes a few [model-dependent] |
| BAM | Peak 4 | polarized | 0.808 | 347 | 690 (M2) | 576–773 | 1,169 | drift readily makes this |
| BAM | Peak 5 | polarized | 0.784 | 531 | 1,025 (M2) | 863–1,141 | 1,595 | drift readily makes this |
| BIK | Peak 2 | label-free | 1 | 11,709 | 4,787 (M3f) | 4,196–5,448 | 7,014 | drift readily makes this |
| BIK | Peak 3 | polarized | 1 | 6,300 | 3,864 (M3) | 3,214–4,380 | 7,527 | drift readily makes this |
| BIK | Peak 6 | label-free | 1 | 11,617 | 4,787 (M3f) | 4,196–5,448 | 7,014 | drift readily makes this |
| BAVA | Peak 2 | label-free | 0.957 | 2,577 | none admissible | — | — | no admissible model |
| BAVA | Peak 3 | polarized | 1 | 385 | 123 (M2) | 108–137 | 319 | drift readily makes this |
| BAVA | Peak 6 | label-free | 1 | 986 | none admissible | — | — | no admissible model |
| BIR | Peak 2 | label-free | 1 | 955 | none admissible | — | — | no admissible model |
| BIR | Peak 3 | polarized | 0.812 | 8,808 | 15,690 (M2) | 14,520–16,910 | 18,150 | drift readily makes this |

**Fig. S8.**
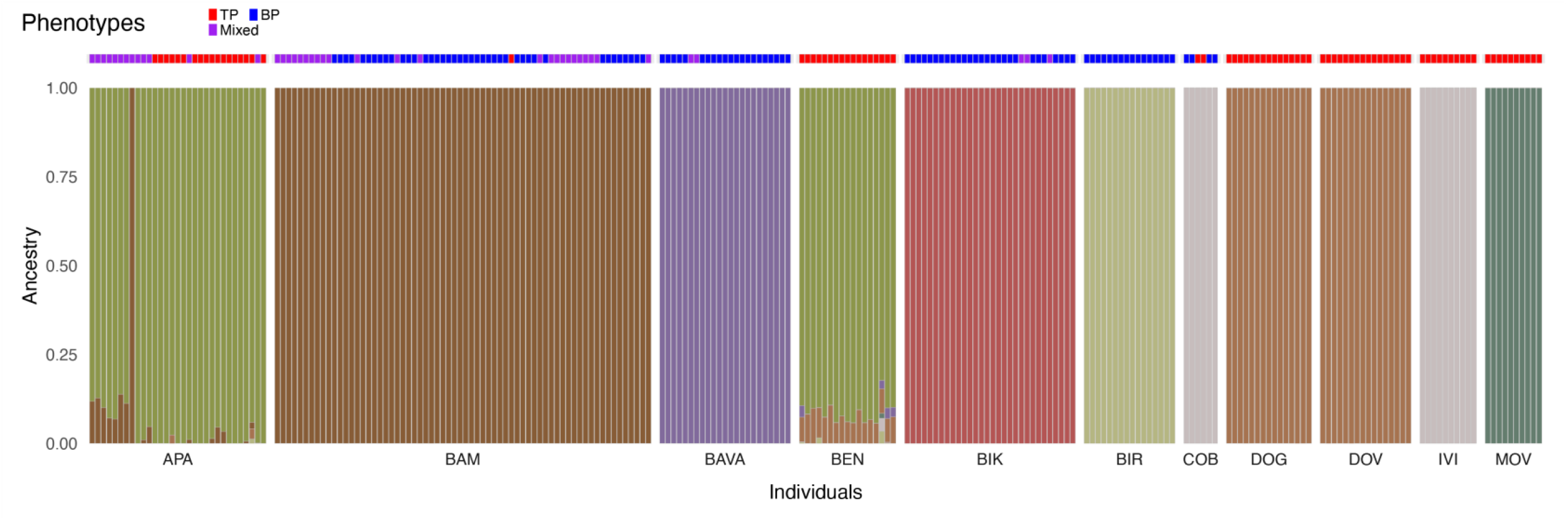
Population structure and mixed-phenotype individuals inferred with ADMIXTURE. Maximum-likelihood ancestry proportions (K = 8, the lowest cross-validation error; K = 10 was nearly tied) for 240 P. kingsleyae individuals genotyped at 40,758 LD-thinned SNPs. Each vertical bar is one individual, partitioned into coloured segments that give the proportion of ancestry assigned to each cluster. Individuals are grouped by population of origin, and phenotype is shown by the box beneath each bar (blue, BP; red, TP; purple, mixed). Most individuals derive nearly all of their ancestry from one cluster. Mixed individuals from Bambomo, Bikagala and Bavavela are indistinguishable from BP individuals of the same population. Mixed individuals from Apassa carry a small Bambomo-cluster component (median 7%, maximum 14%) that is absent at K = 10. Most of this component is carried by mixed individuals from one 2019 collection site (Supplemental Note). No mixed individual carries the 25–50% opposite-phenotype ancestry expected of first-generation hybrids or first backcrosses.

**Fig. S9.**
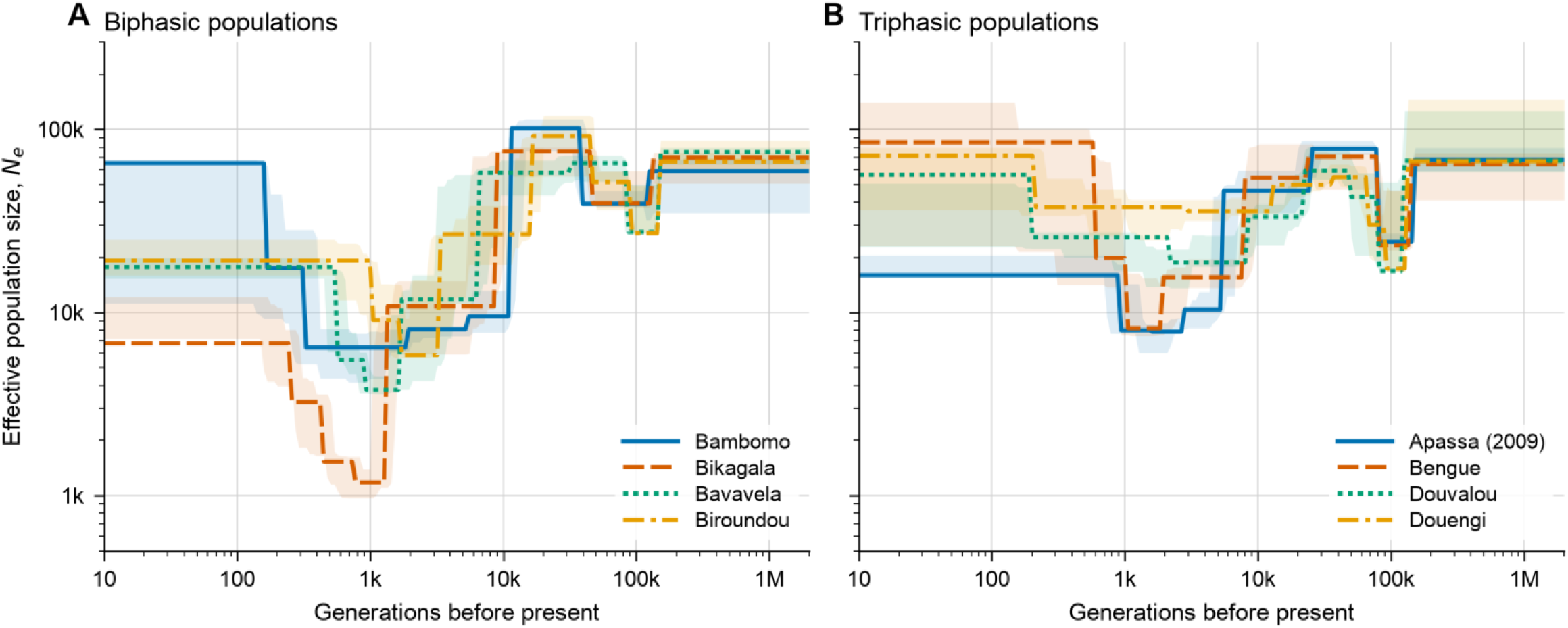
Historical effective population size. SMC++ estimates of effective population size (diploid individuals) through time in (A) the four BP populations and (B) the four TP populations, assuming a mutation rate of 1 × 10⁻⁸ per site per generation. Time is in generations before present, and both axes are logarithmic. Lines are point estimates and shaded bands the 2.5th–97.5th percentiles of 10 bootstraps over chromosomes. Colours identify populations within a panel only. Apassa is represented by its 2009 sample. Samples, masks and settings are given in Historical Effective Population Size. In simulations, SMC++ recovered reductions within roughly the last 1,000 generations but not the timing of older ones, and placed the minimum size about two-fold too low, so the reductions may be shallower than shown. We therefore interpret only the existence and approximate size of the reductions, not their timing or shape.

