## Supplemental Data S2 for "The Genomic Basis of Electric Signal Diversity": CombinedOutput.pdf

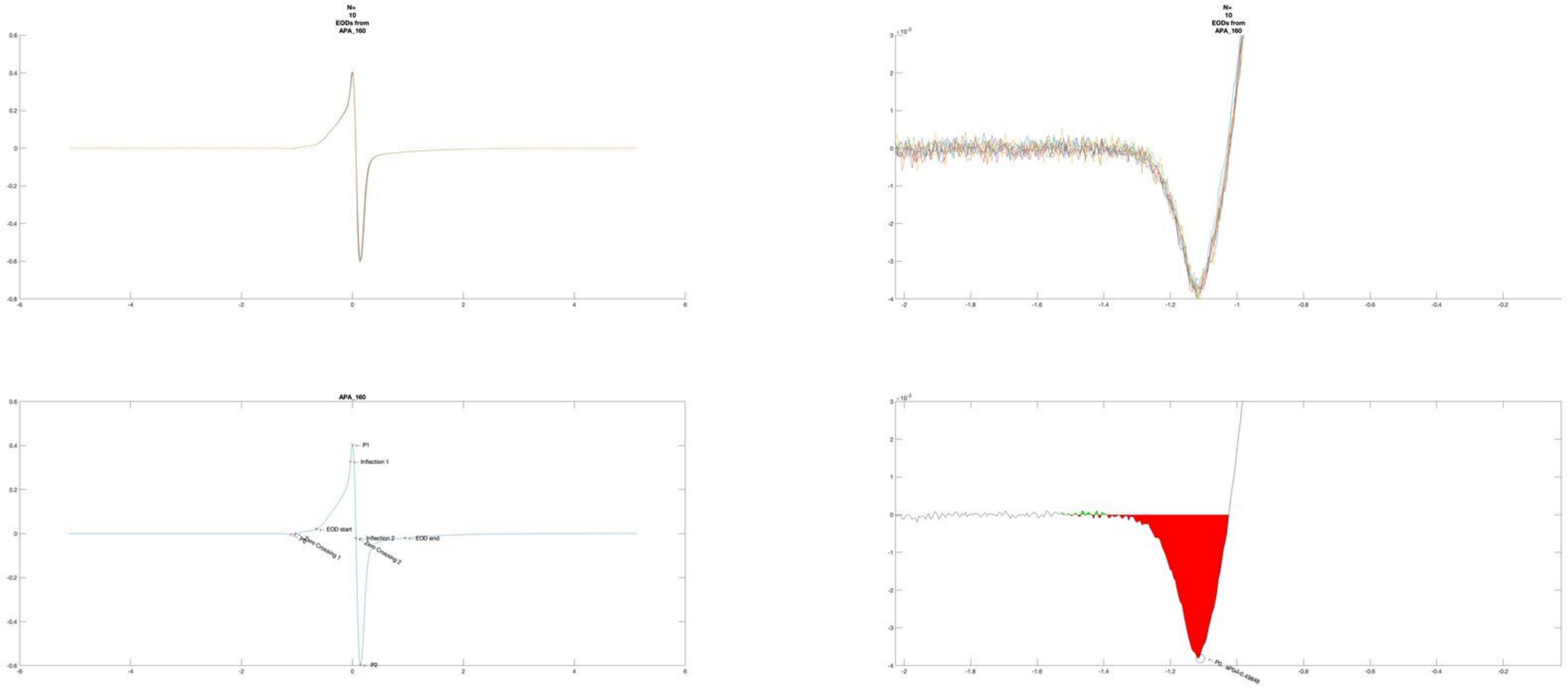
N=
10
EODs from
APA_160
N=
10
EODs from
APA_160
APA_160
P1
Inflection 1
EOD start
Inflection 2
EOD end
Zero Crossing 1
Zero Crossing 2
P2

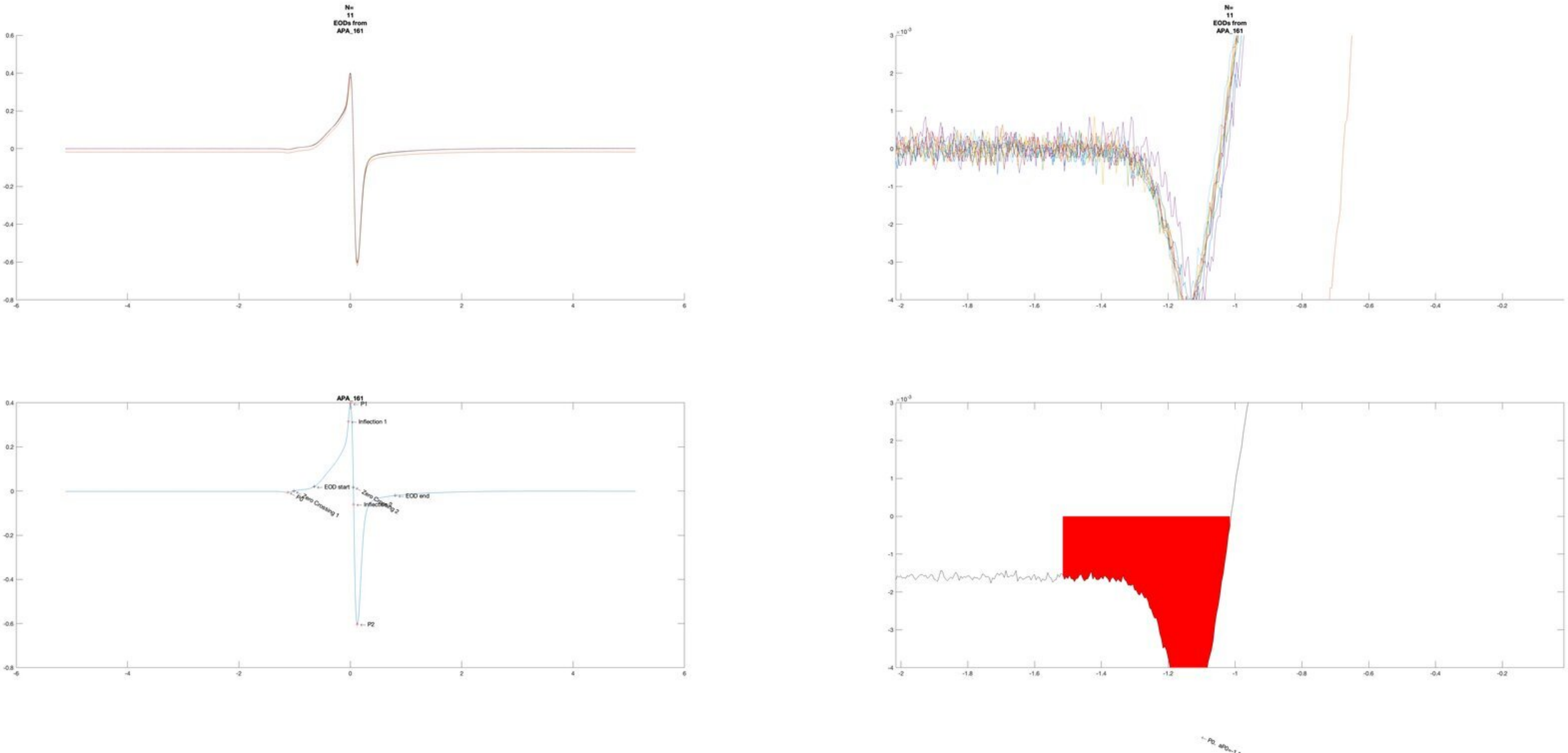
N=
11
EODs from
APA_161
N=
11
EODs from
APA_161
APA_161
P1
Inflection 1
EOD start
EOD end
Zero Crossing 1
P2

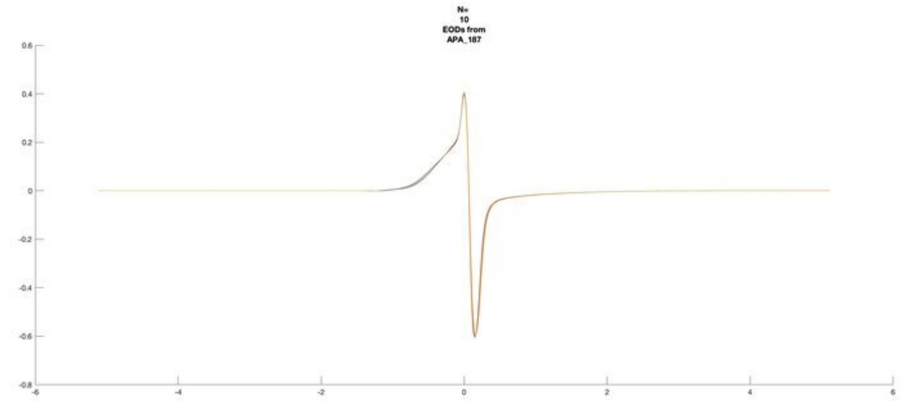
N=
10
EODs from
APA_187
0.6
0.4
0.2
0
-0.2
-0.4
-0.6
-0.8
-6
-4
-2
0
2
4
6

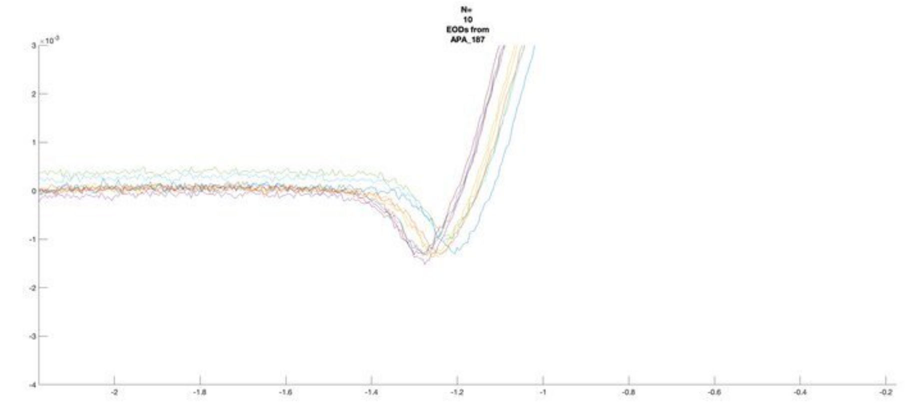
N=
10
EODs from
APA_187
×10⁻³
3
2
1
0
-1
-2
-3
-4
-2
-1.8
-1.6
-1.4
-1.2
-1
-0.8
-0.6
-0.4
-0.2

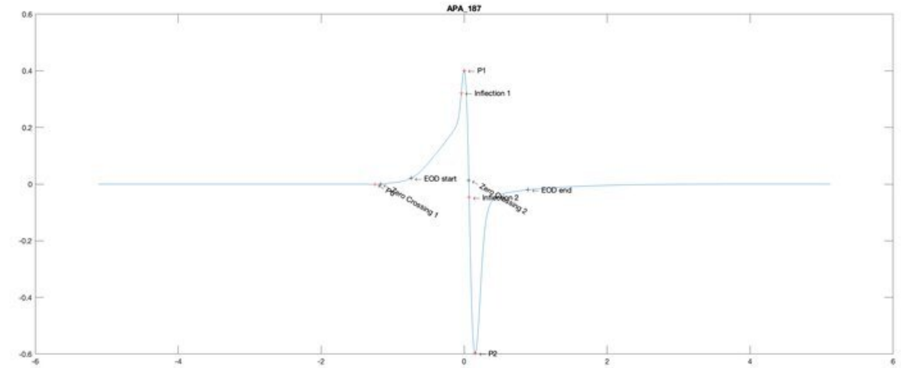
APA_187
P1
Inflection 1
EOD start
Zero Crossing 1
Zero Crossing 2
Inflection 2
EOD end
P2
0.6
0.4
0.2
0
-0.2
-0.4
-0.6
-6
-4
-2
0
2
4
6

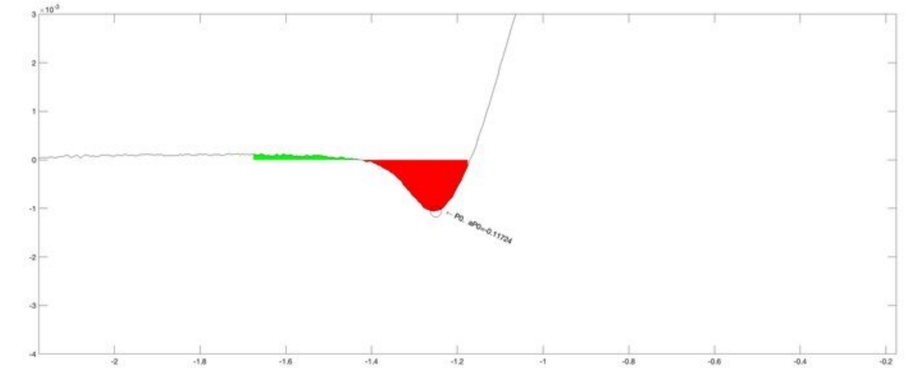
×10⁻³
P0, aP0=-0.11724
3
2
1
0
-1
-2
-3
-4
-2
-1.8
-1.6
-1.4
-1.2
-1
-0.8
-0.6
-0.4
-0.2

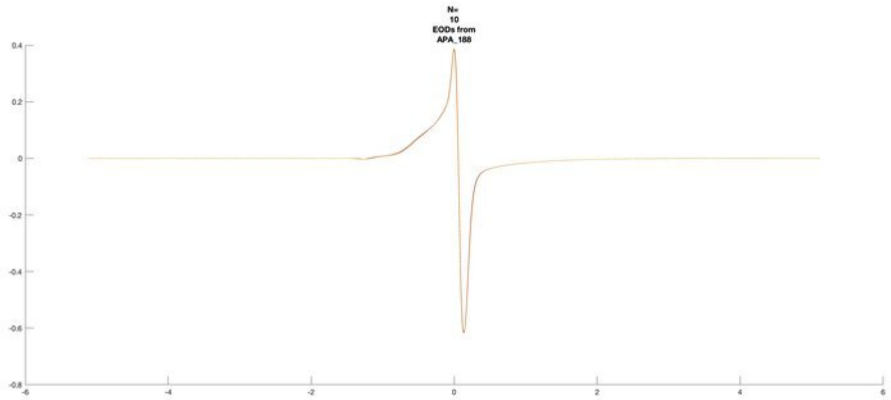
N=
10
EODs from
APA_188

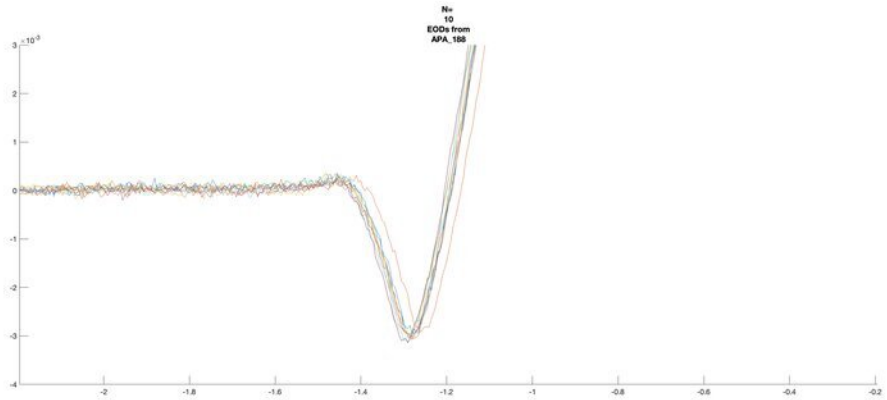
N=
10
EODs from
APA_188

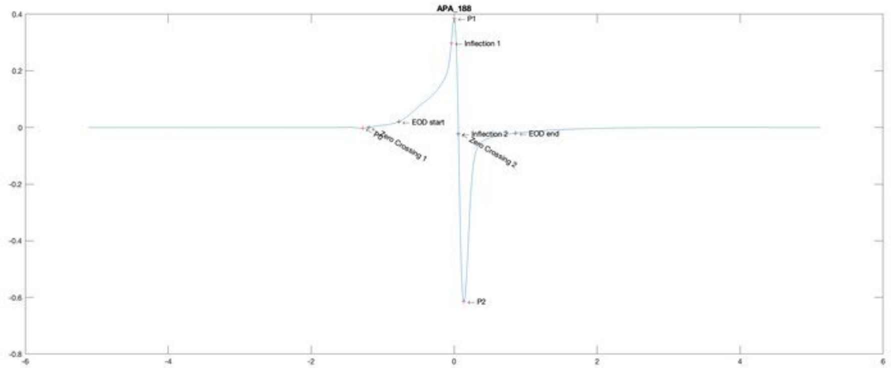
APA_188
P1
Inflection 1
EOD start
Zero Crossing 1
P0
Inflection 2
EOD end
Zero Crossing 2
P2

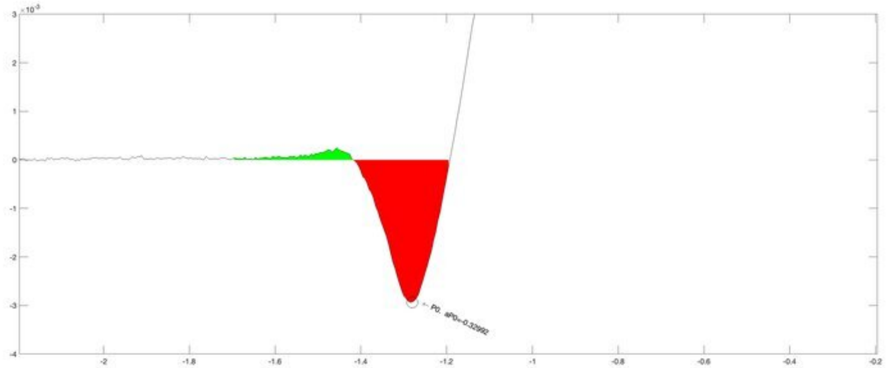
P0, aP0=-0.32992

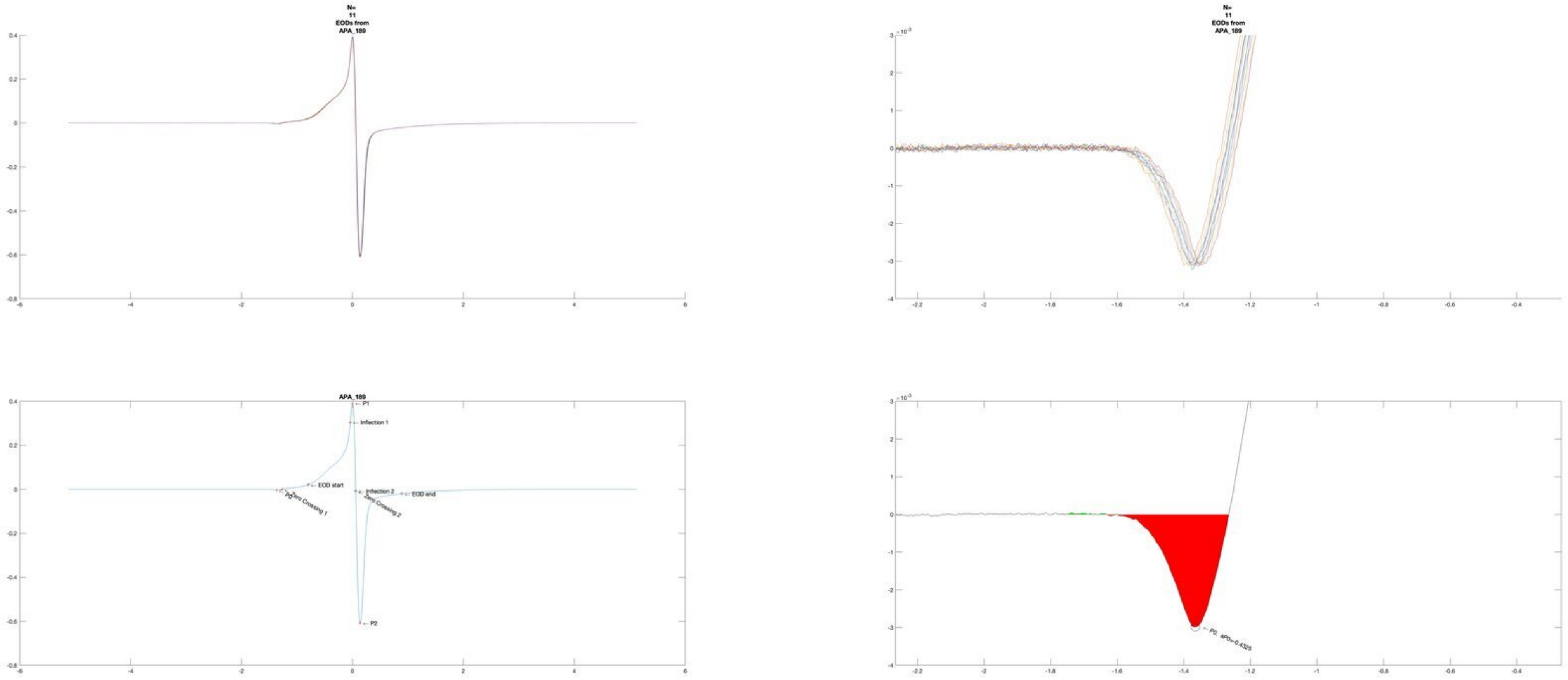
N=
11
EODs from
APA_189
N=
11
EODs from
APA_189
APA_189
P1
Inflection 1
EOD start
Inflection 2
EOD end
P0
Zero Crossing 1
Zero Crossing 2
P2
P0, aP0=-0.4325

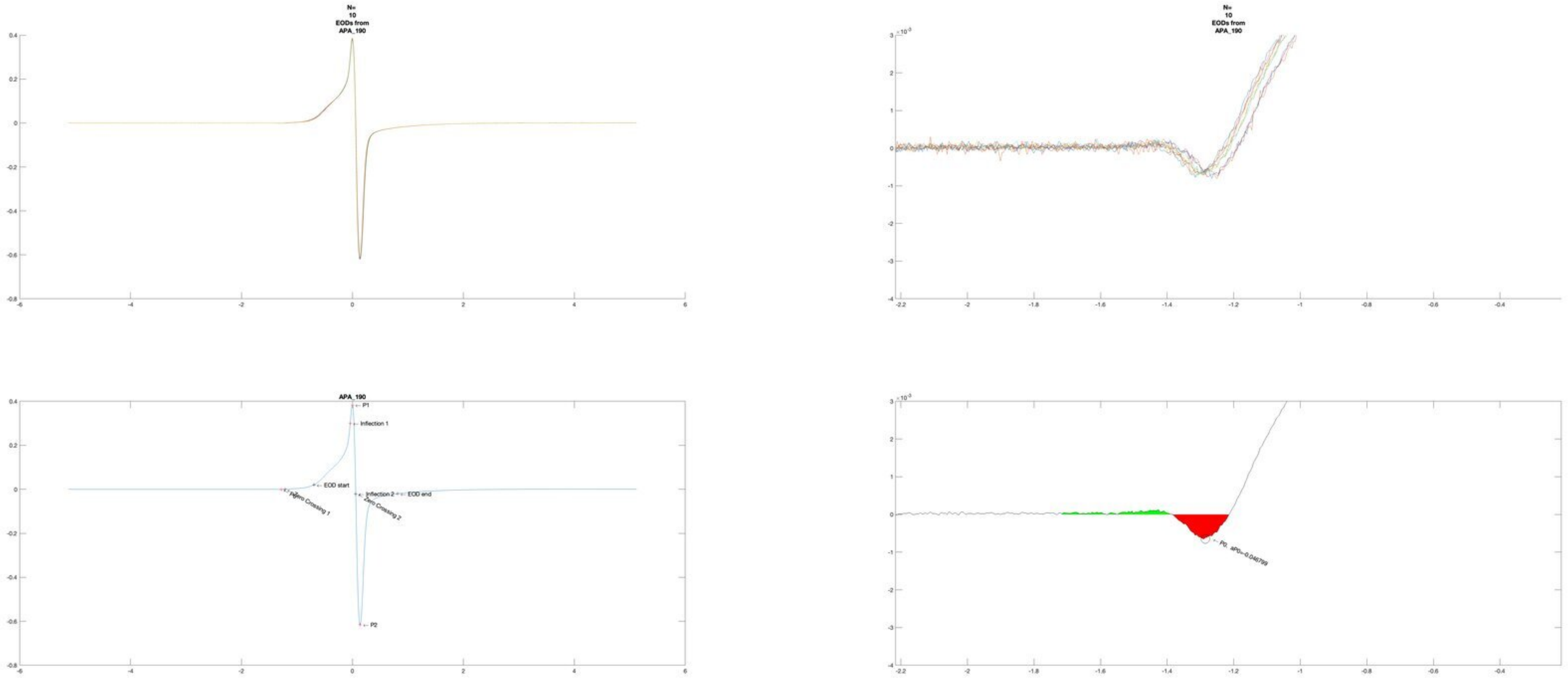
N=
10
EODs from
APA_190
N=
10
EODs from
APA_190
APA_190
P1
Inflection 1
EOD start
Inflection 2
EOD end
Zero Crossing 1
Zero Crossing 2
P2
P0, aP0=-0.046799

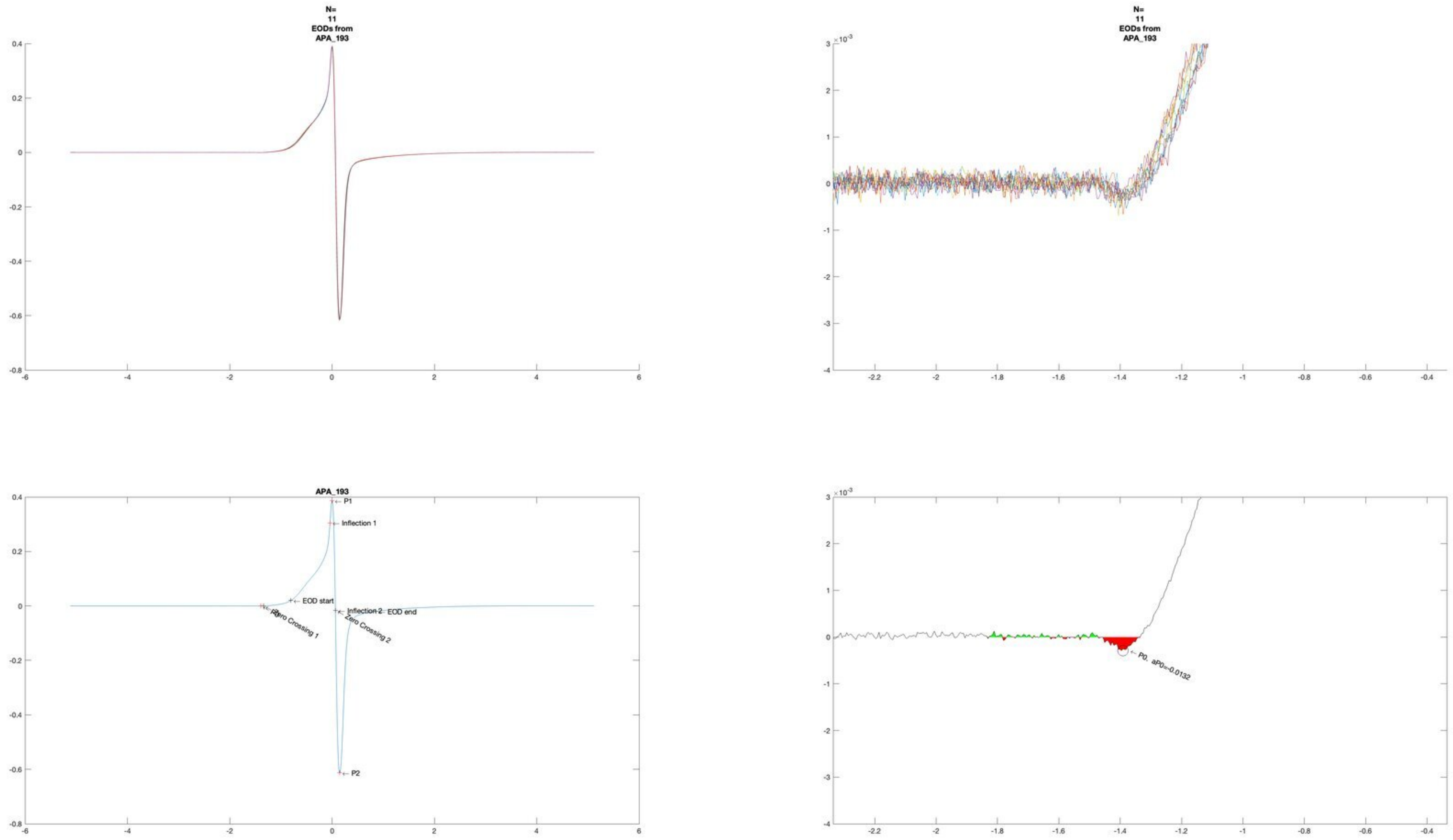
N=
11
EODs from
APA_193
N=
11
EODs from
APA_193
×10^-3
APA_193
P1
Inflection 1
EOD start
Inflection 2
EOD end
Zero Crossing 1
Zero Crossing 2
P2
×10^-3
P0. aP0=-0.0132

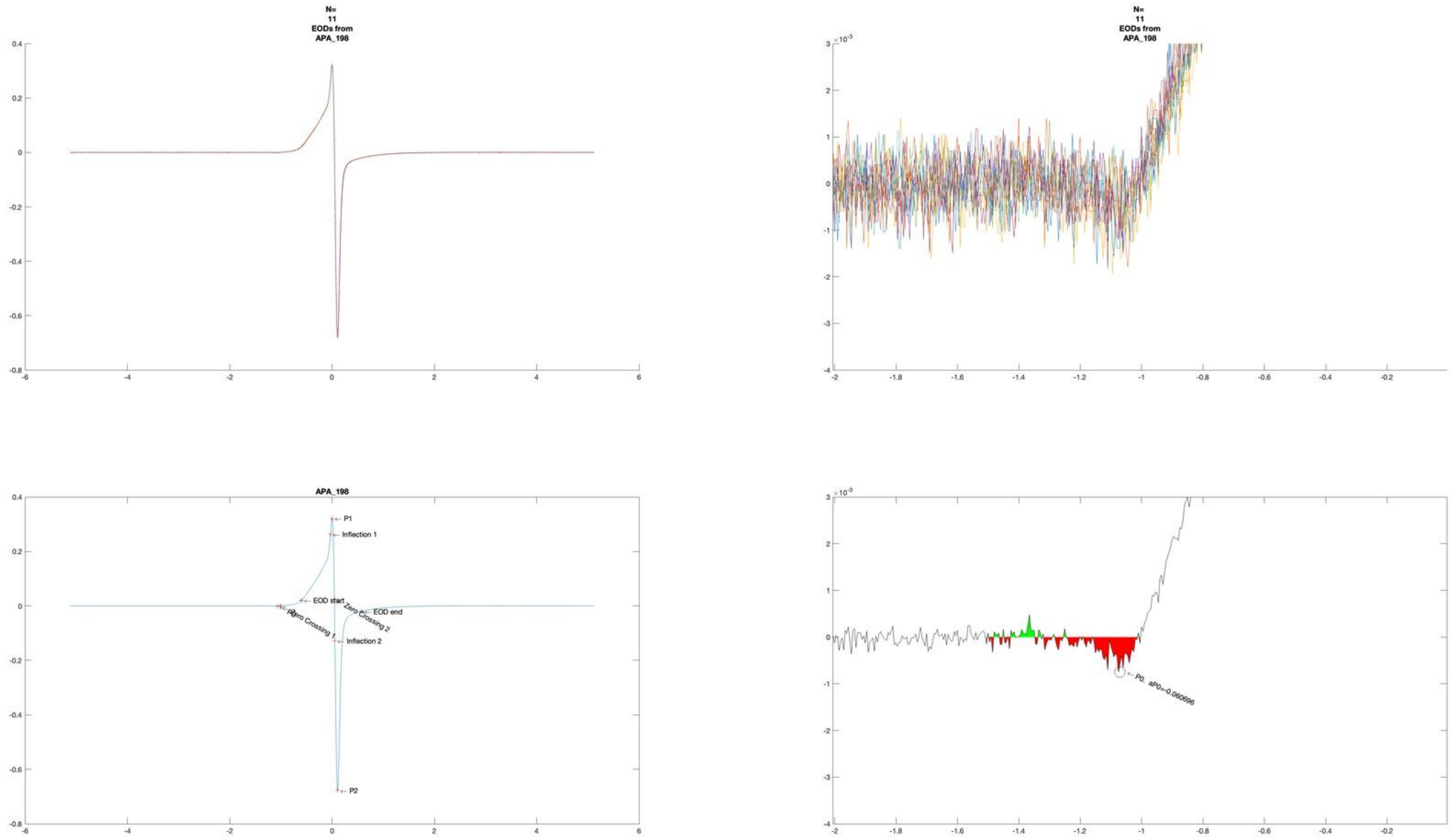
N=
11
EODs from
APA_198
APA_198
P1
Inflection 1
EOD start
Zero Crossing 2
EOD end
Zero Crossing 1
Inflection 2
P2
P0, aP0=-0.060696

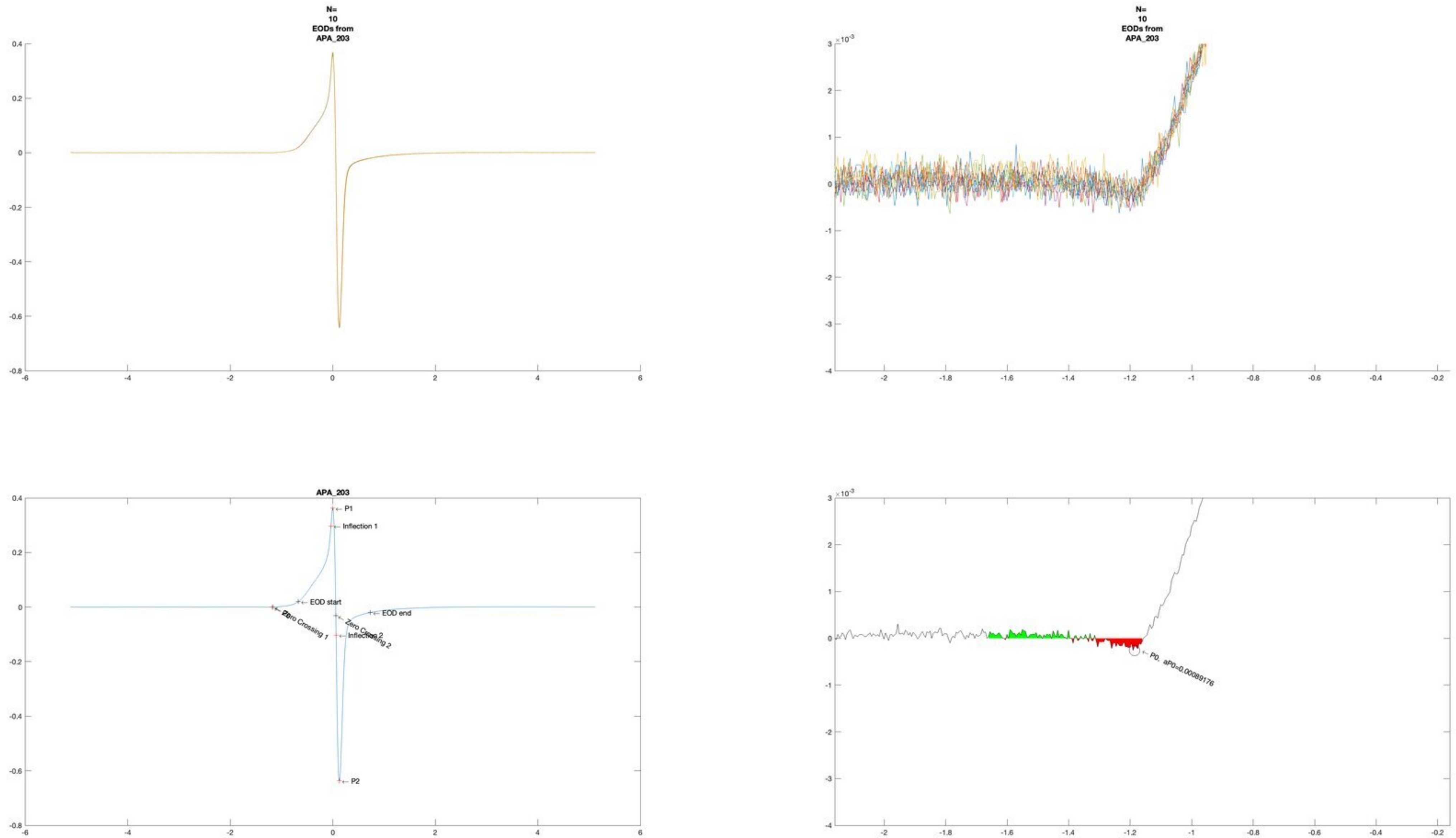
N=
10
EODs from
APA_203
N=
10
EODs from
APA_203
APA_203
P1
Inflection 1
EOD start
EOD end
Zero Crossing 1
Zero Crossing 2
Inflection 2
P2
P0, aP0=0.00089176

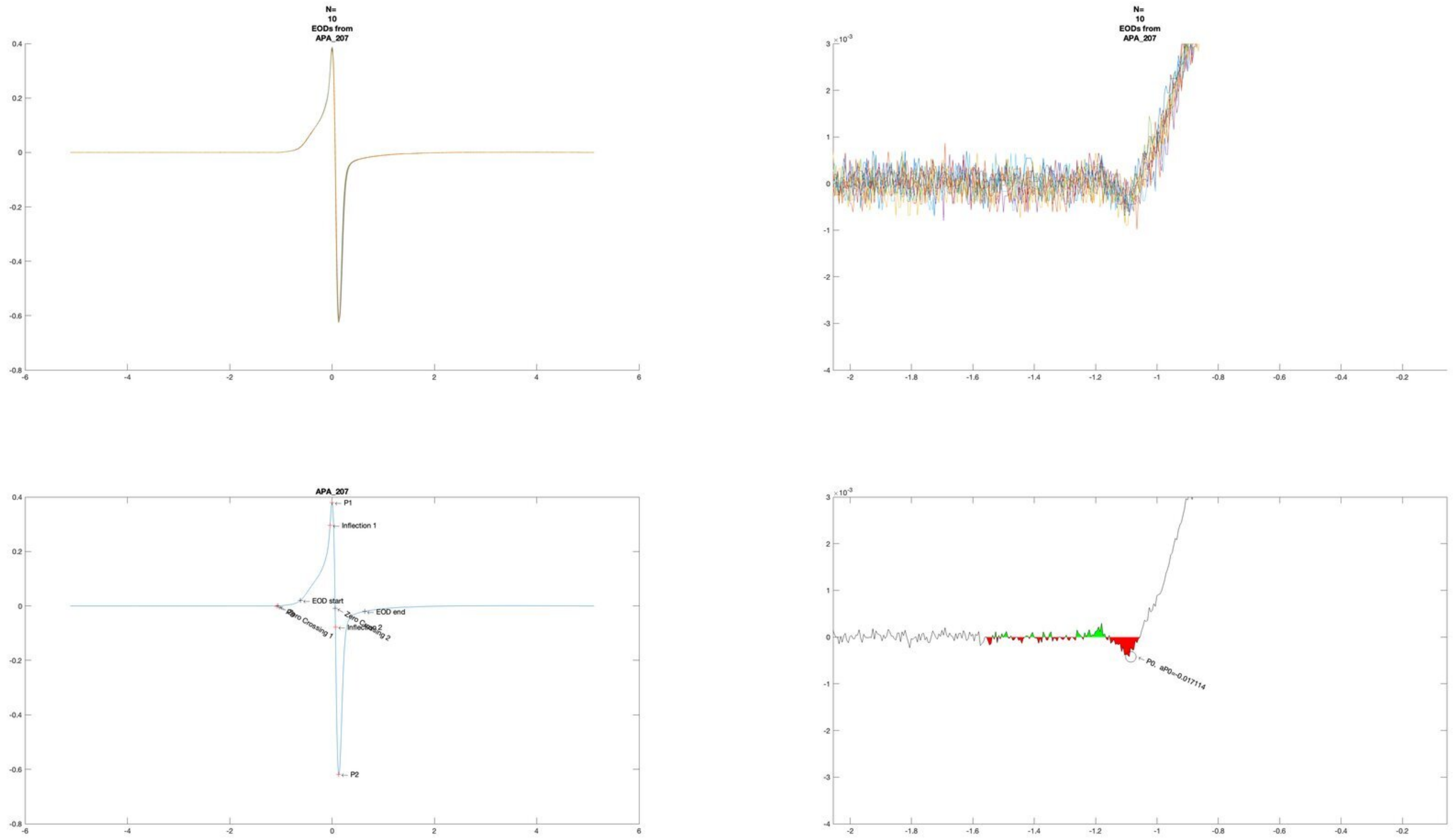
N=
10
EODs from
APA_207
N=
10
EODs from
APA_207
APA_207
P1
Inflection 1
EOD start
EOD end
Zero Crossing 1
Zero Crossing 2
Inflection 2
P2
P0, aP0=-0.017114

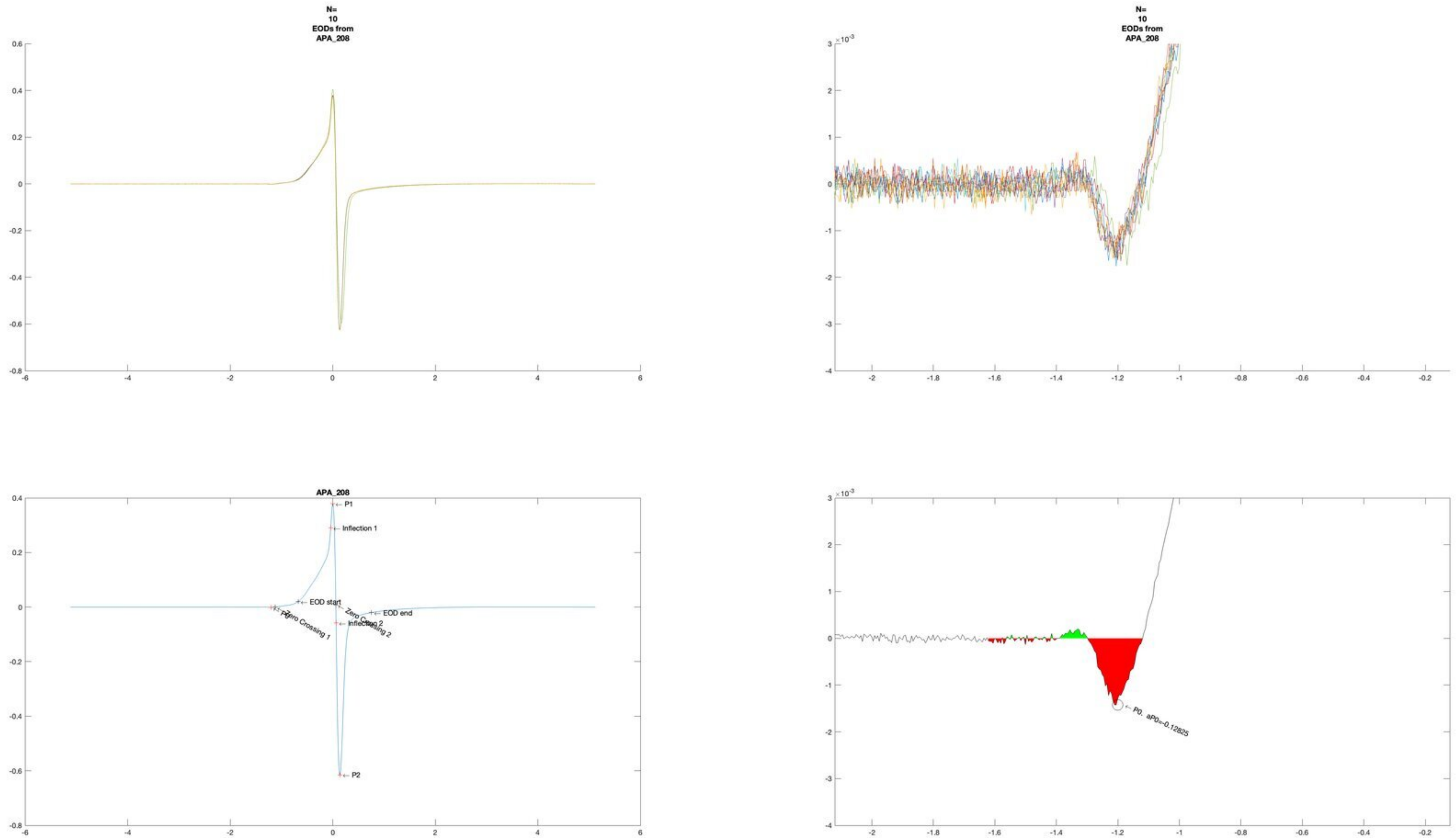
N=
10
EODs from
APA_208
N=
10
EODs from
APA_208
APA_208
P1
Inflection 1
EOD start
Zero Crossing 2
EOD end
P0
Zero Crossing 1
Inflection 2
P2
P0. aP0=-0.12825

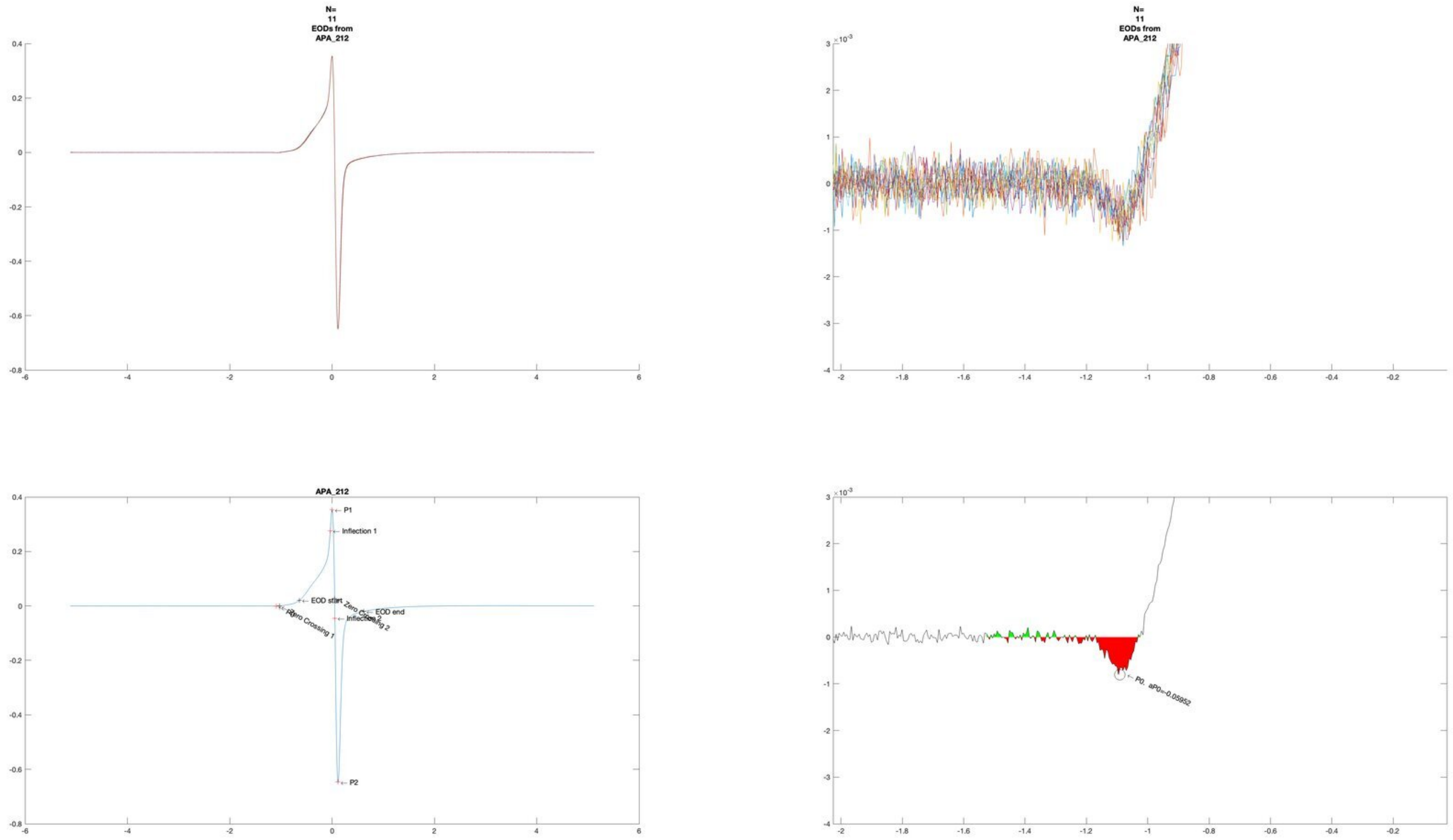
N=
11
EODs from
APA_212
N=
11
EODs from
APA_212
APA_212
P1
Inflection 1
EOD start
Zero Crossing 1
Zero Crossing 2
EOD end
Inflection 2
P2
P0, aP0=-0.05952

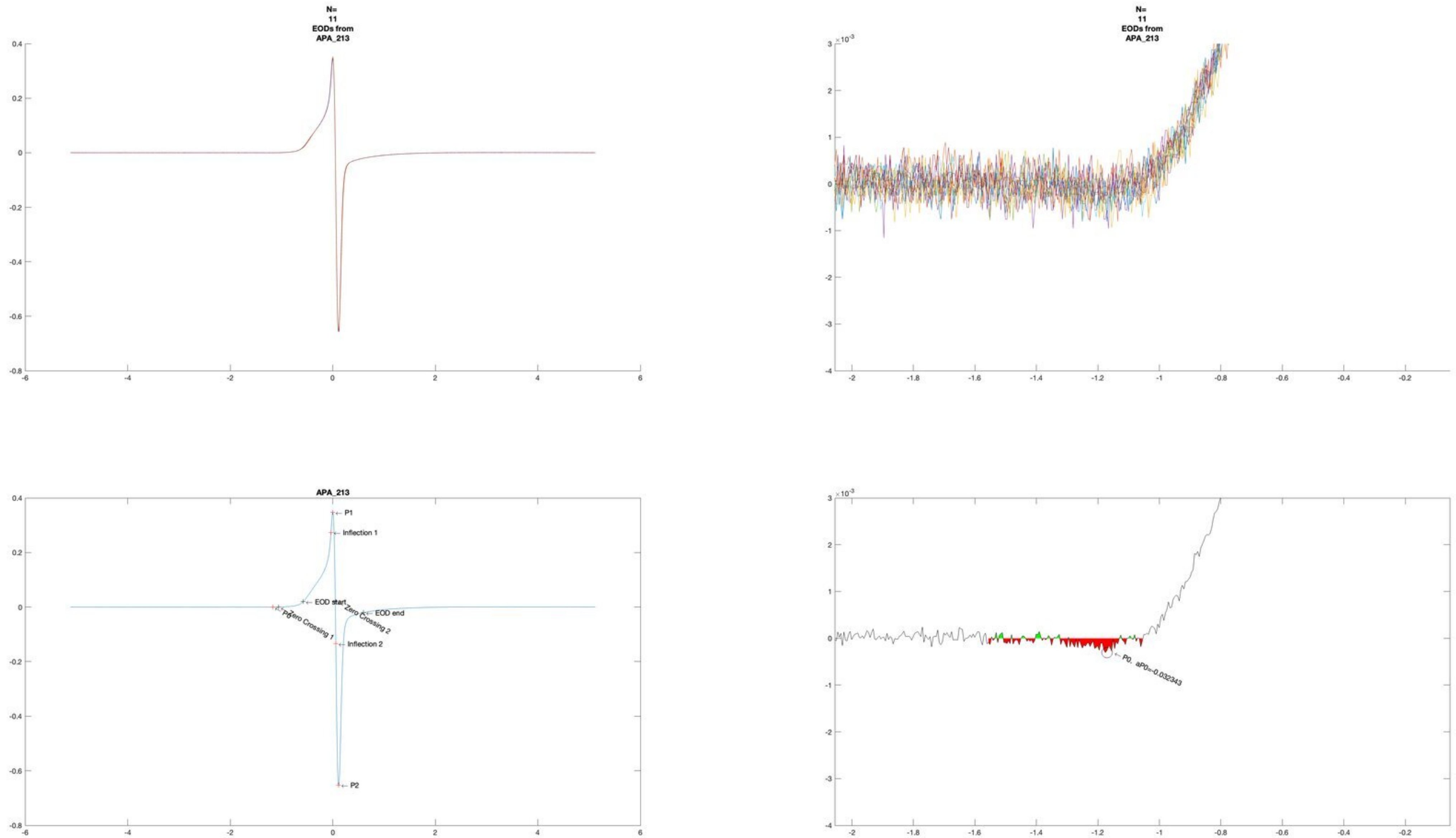
N=
11
EODs from
APA_213
N=
11
EODs from
APA_213
APA_213
P1
Inflection 1
EOD start
P0
Zero Crossing 1
Zero Crossing 2
EOD end
Inflection 2
P2
P0, aP0=-0.032343

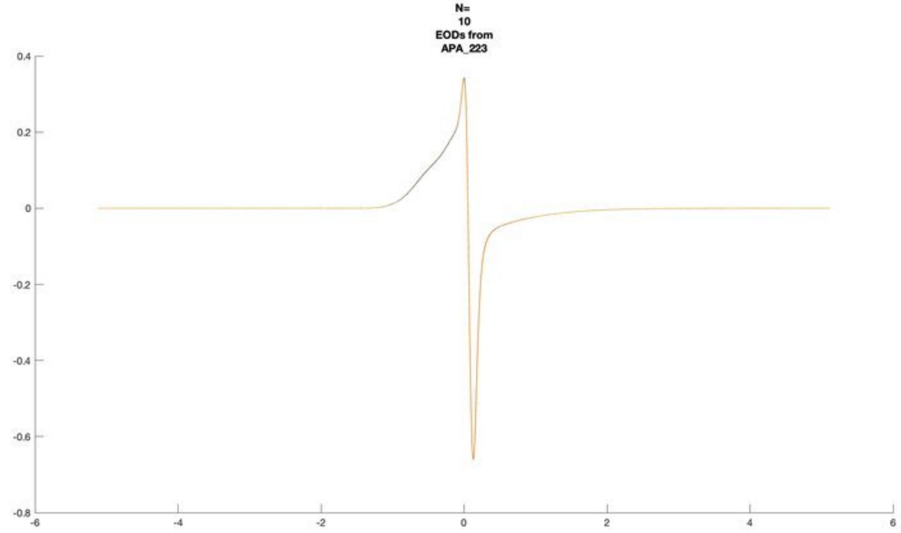
N=
10
EODs from
APA_223

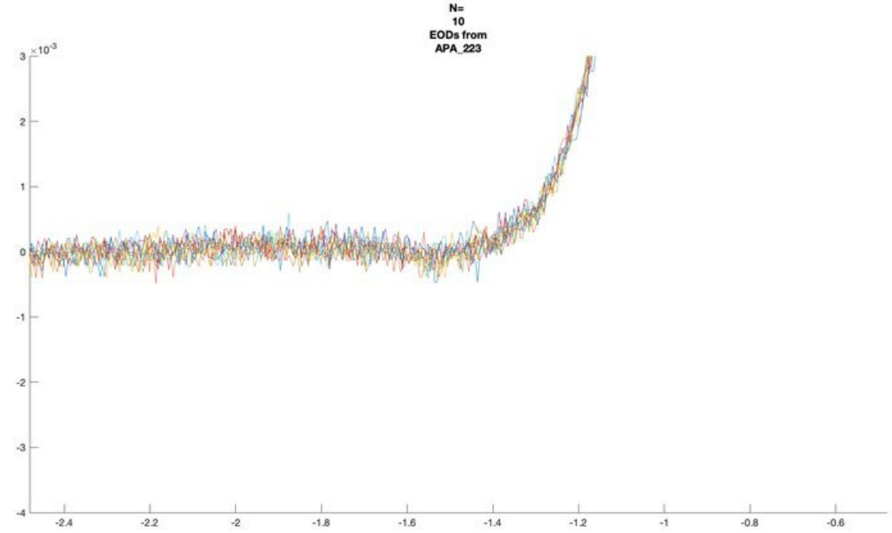
N=
10
EODs from
APA_223

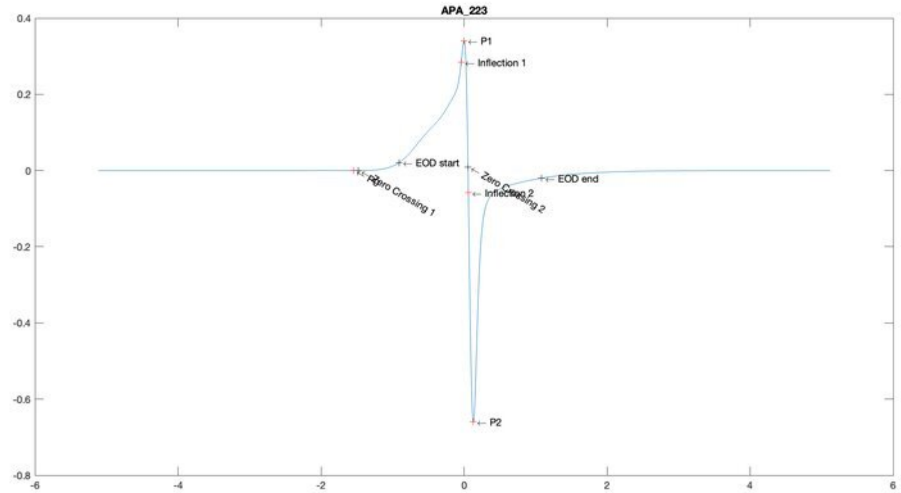
APA_223
P1
Inflection 1
EOD start
Zero Crossing 1
Zero Crossing 2
Inflection 2
EOD end
P2

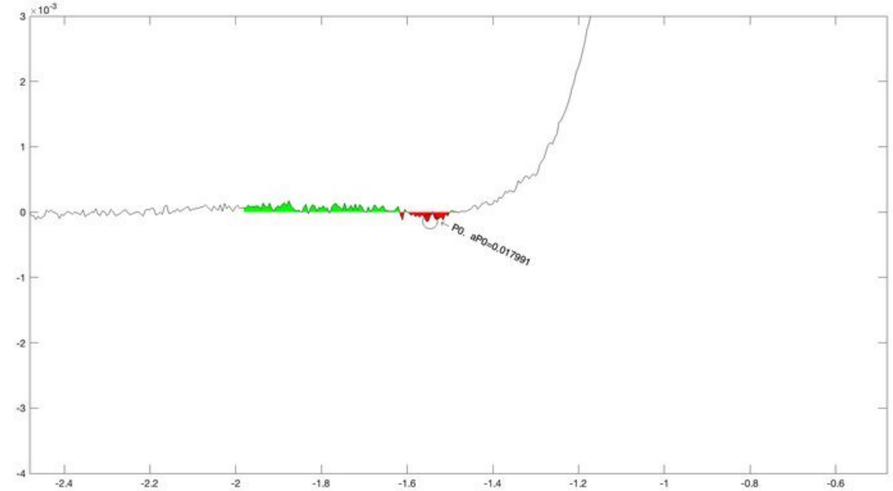
P0, aP0=0.017991

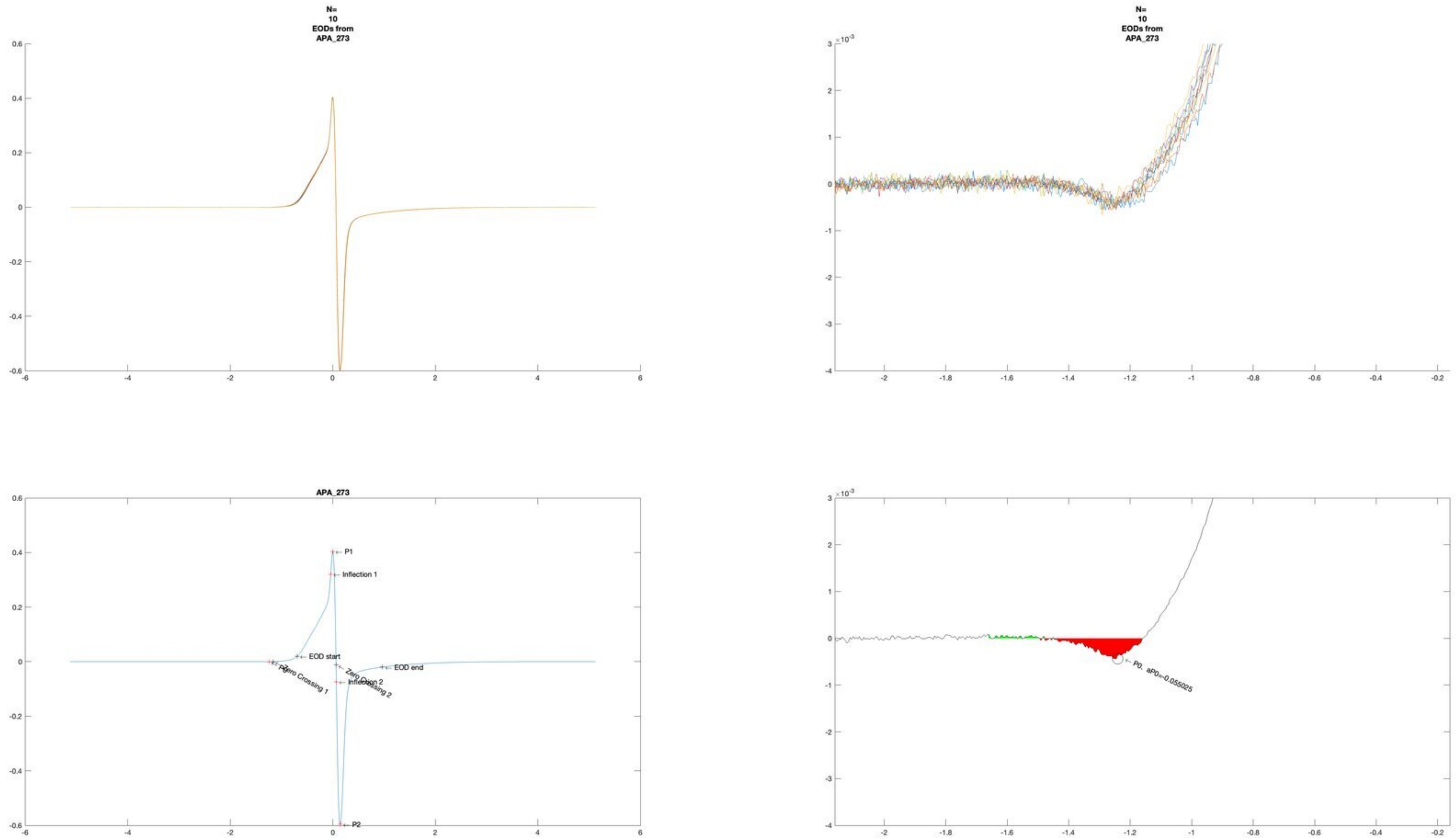

N=
10
EODs from
APA_273
N=
10
EODs from
APA_273
APA_273
P1
Inflection 1
EOD start
P0
Zero Crossing 1
Zero Crossing 2
Inflection 2
EOD end
P2
P0. aP0=-0.055025

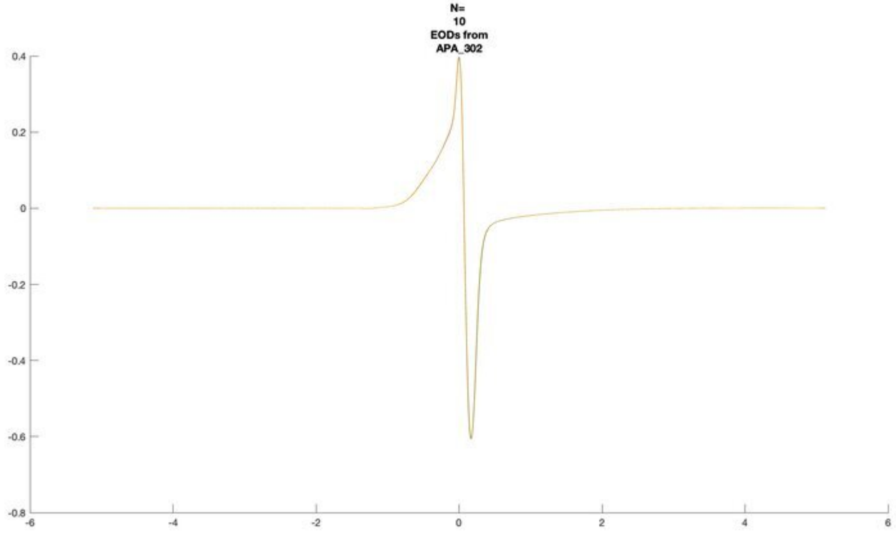
N=
10
EODs from
APA_302

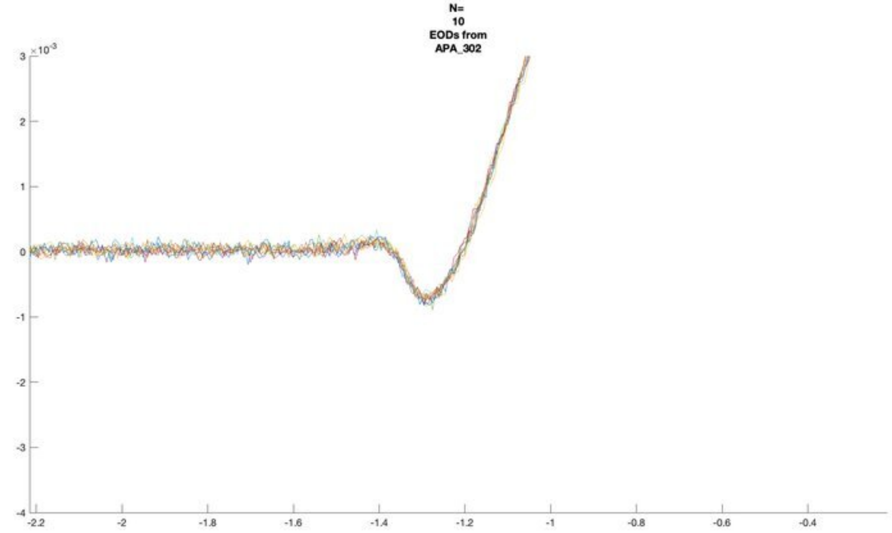
N=
10
EODs from
APA_302
×10^-3

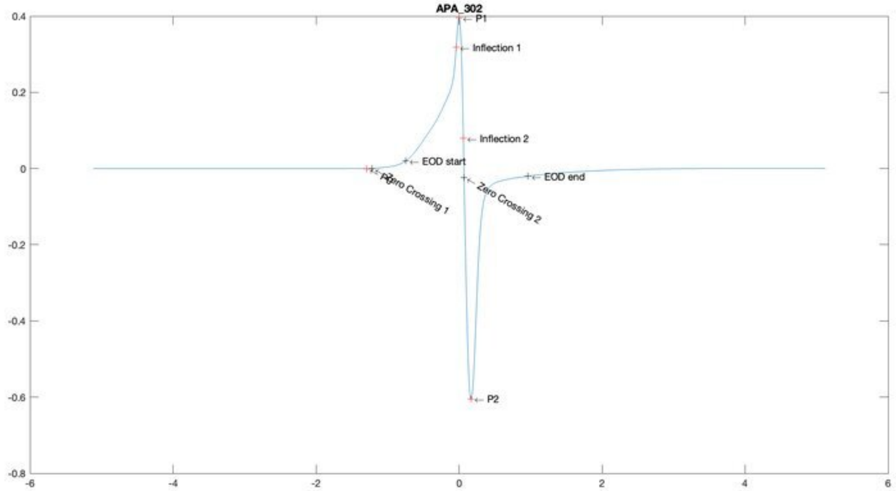
APA_302
P1
Inflection 1
Inflection 2
EOD start
EOD end
Zero Crossing 1
Zero Crossing 2
P2

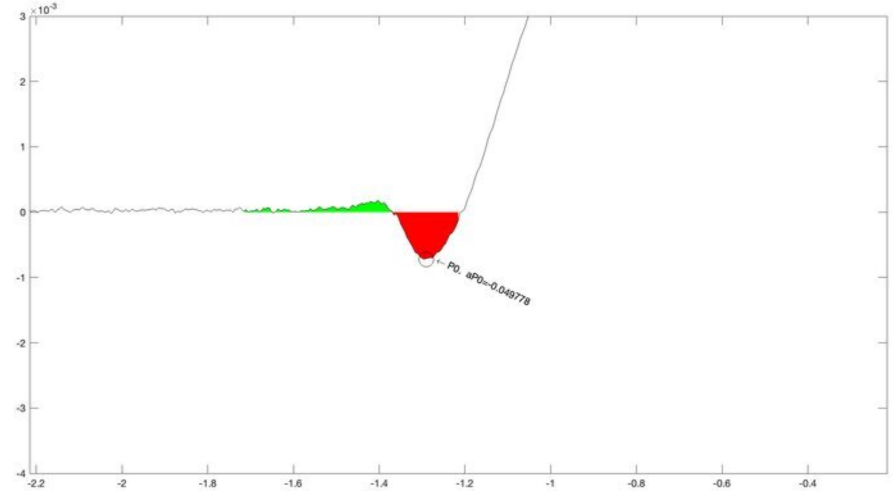
×10^-3
P0, aP0=-0.049778

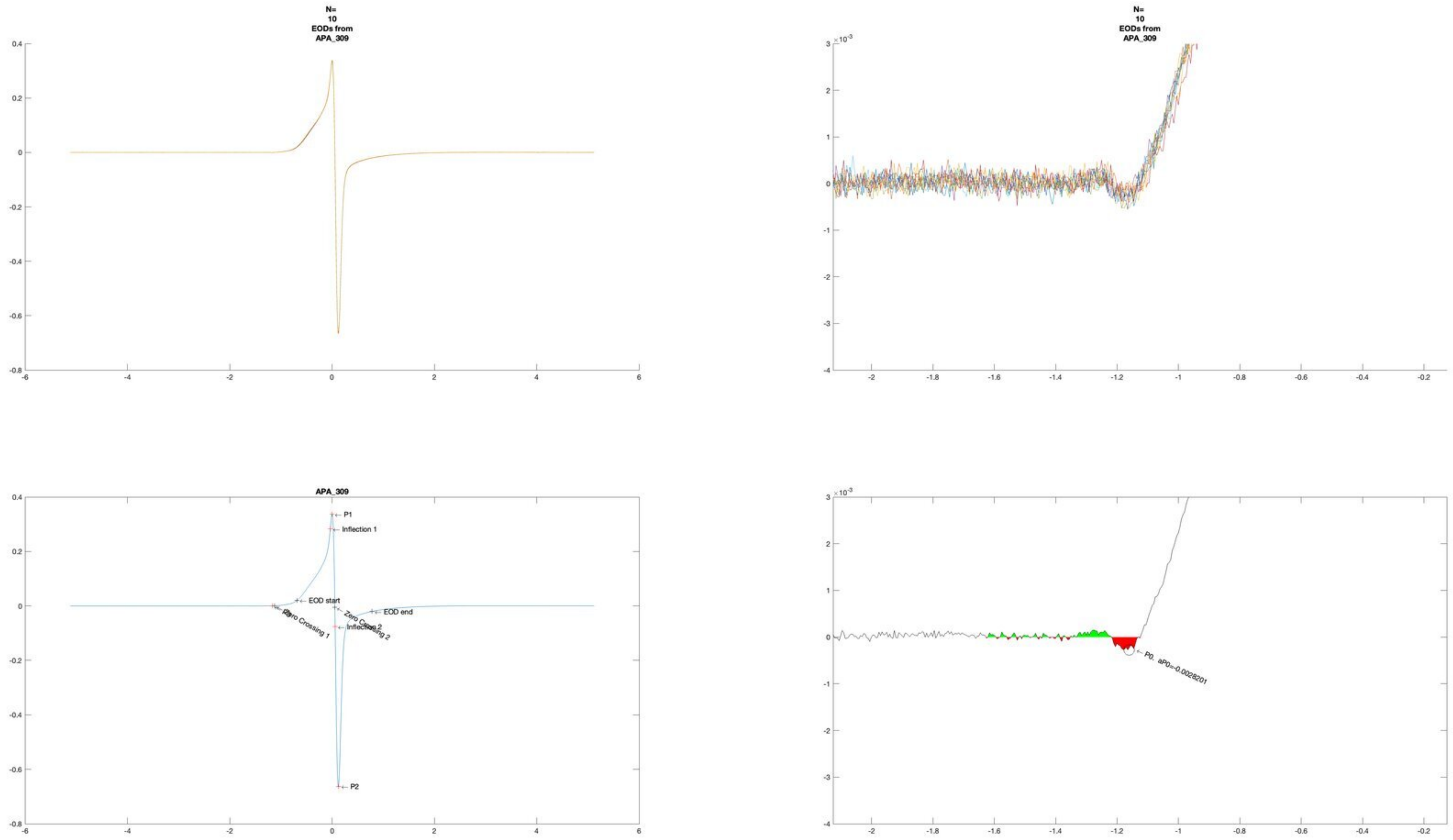
N=
10
EODs from
APA_309
N=
10
EODs from
APA_309
APA_309
P1
Inflection 1
EOD start
EOD end
Zero Crossing 1
Zero Crossing 2
Inflection 2
P2
P0, aP0=-0.0028201

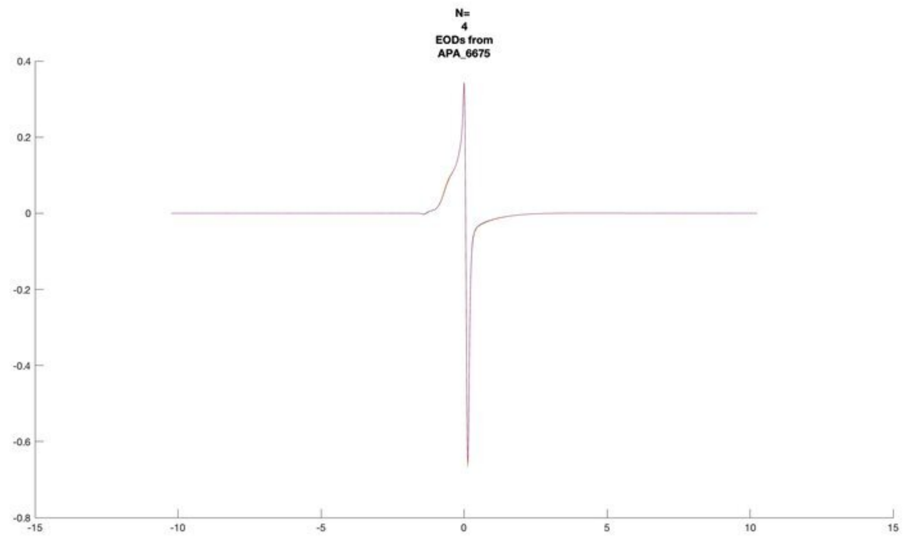
N=
4
EODs from
APA_6675

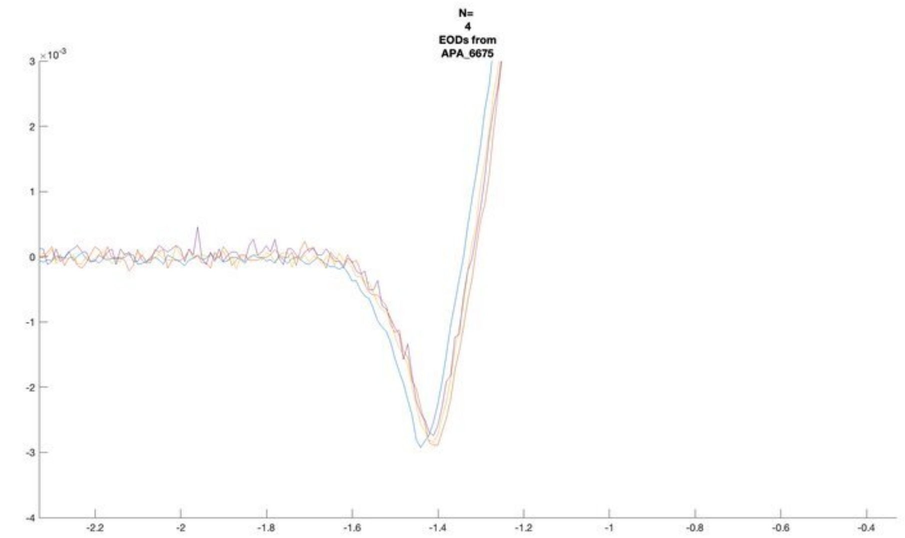
N=
4
EODs from
APA_6675
×10^-3

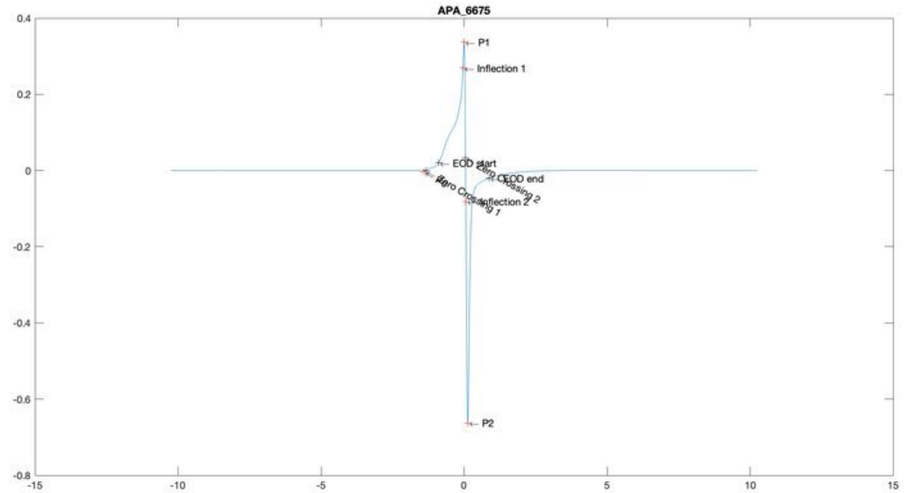
APA_6675
P1
Inflection 1
EOD start
EOD end
Inflection 2
P2

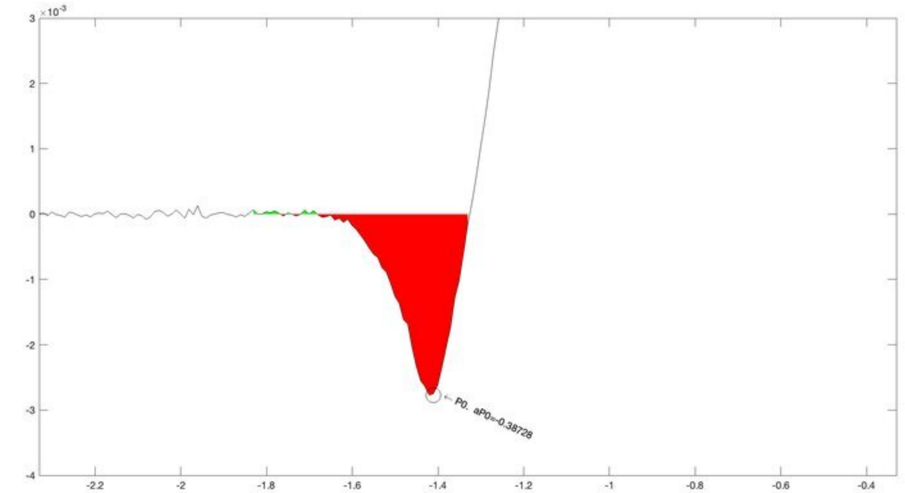
×10^-3
P0. aP0=-0.38728

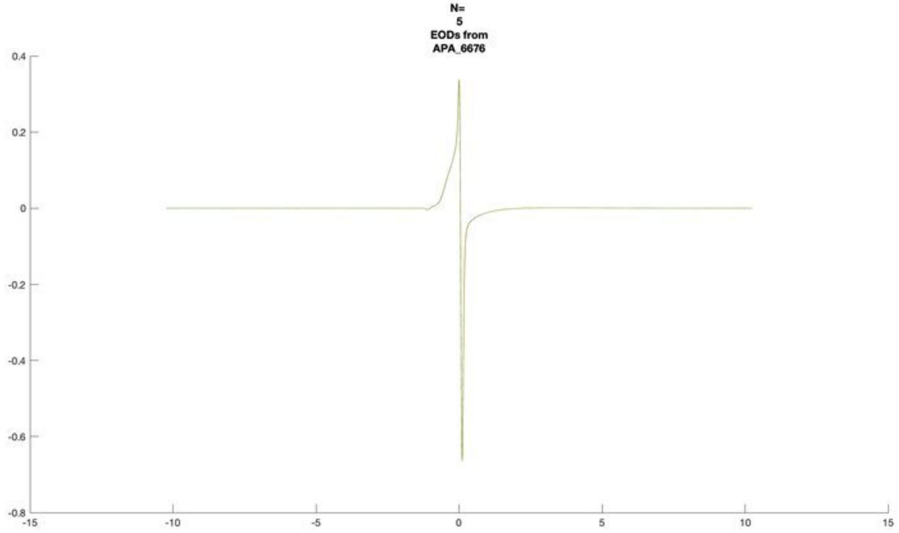
N=
5
EODs from
APA_6676

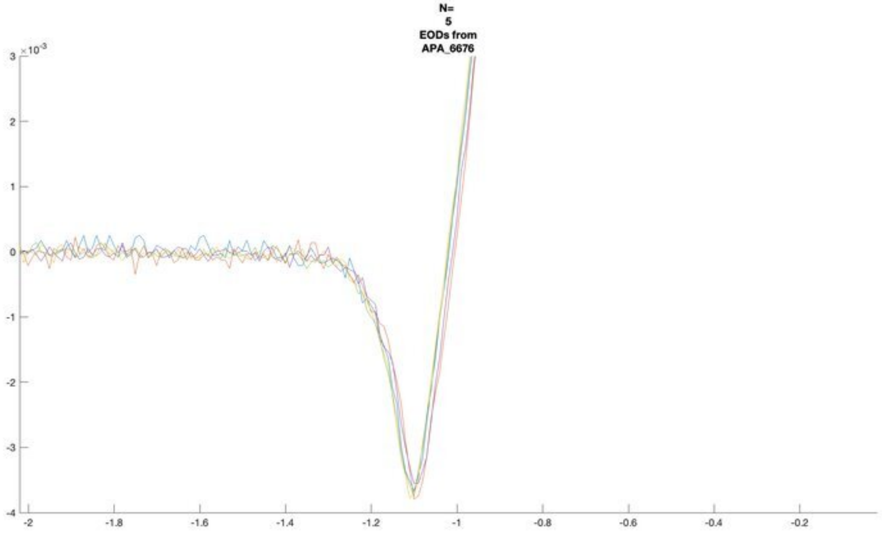
N=
5
EODs from
APA_6676
×10^-3

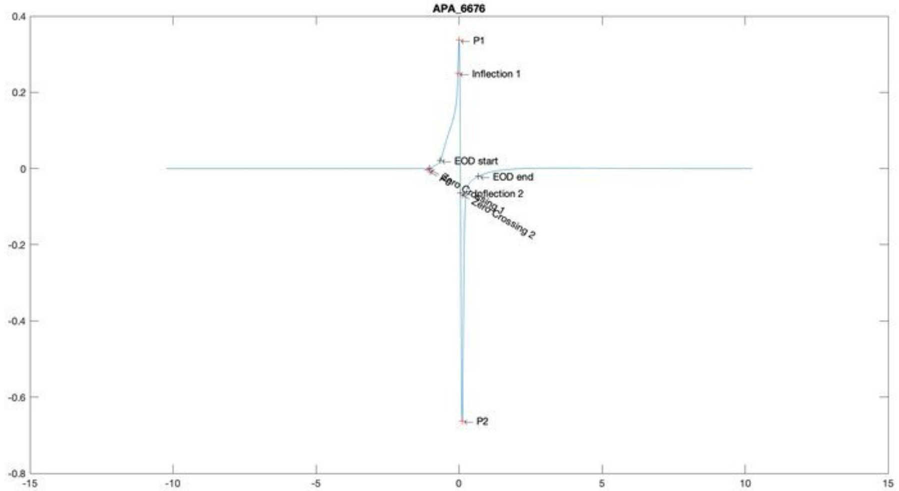
APA_6676
P1
Inflection 1
EOD start
EOD end
Inflection 2
Zero Crossing 2
P2

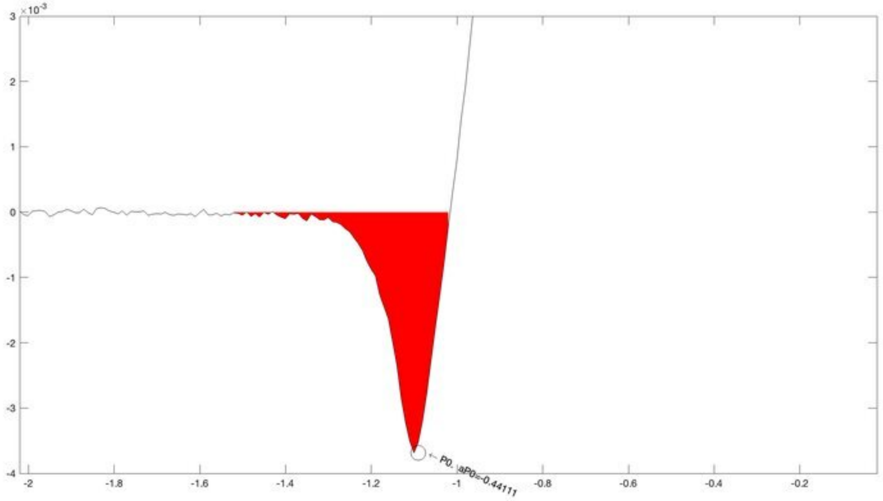
×10^-3
P0, aP0=-0.44111

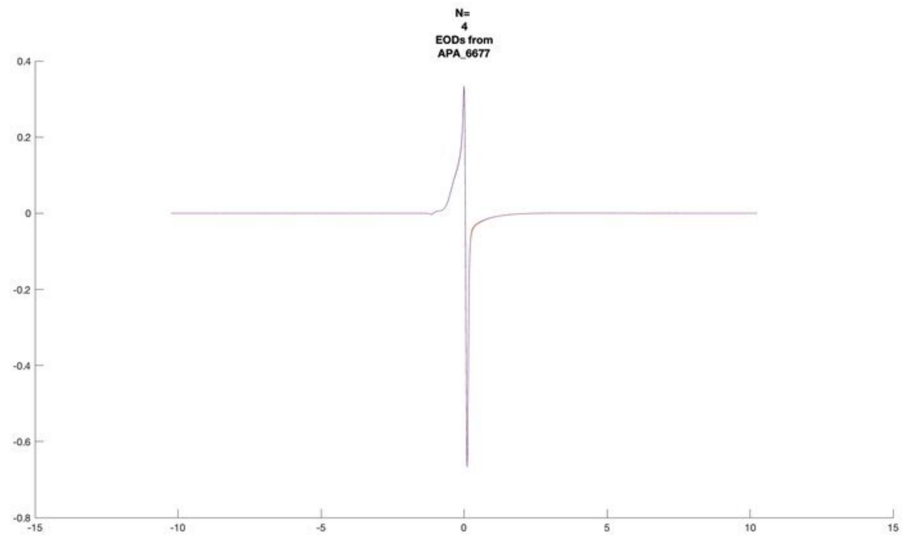
N=
4
EODs from
APA_6677

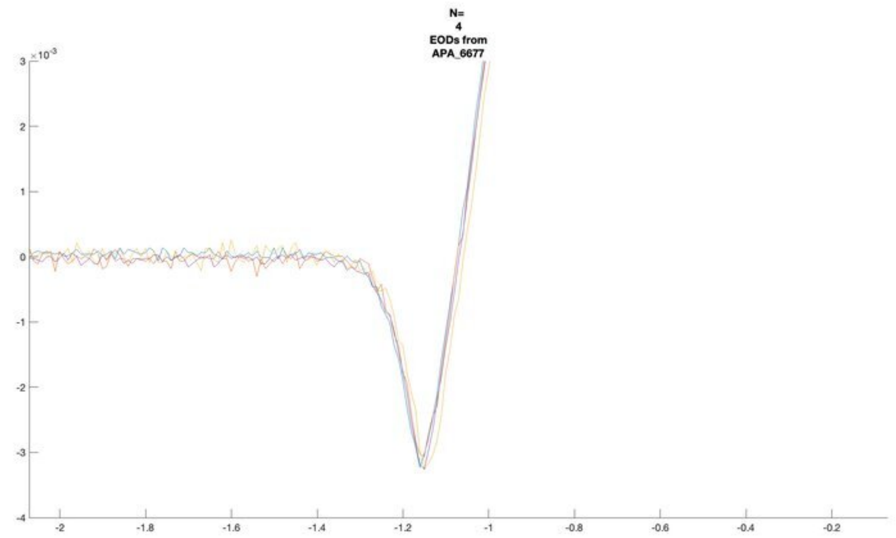
N=
4
EODs from
APA_6677

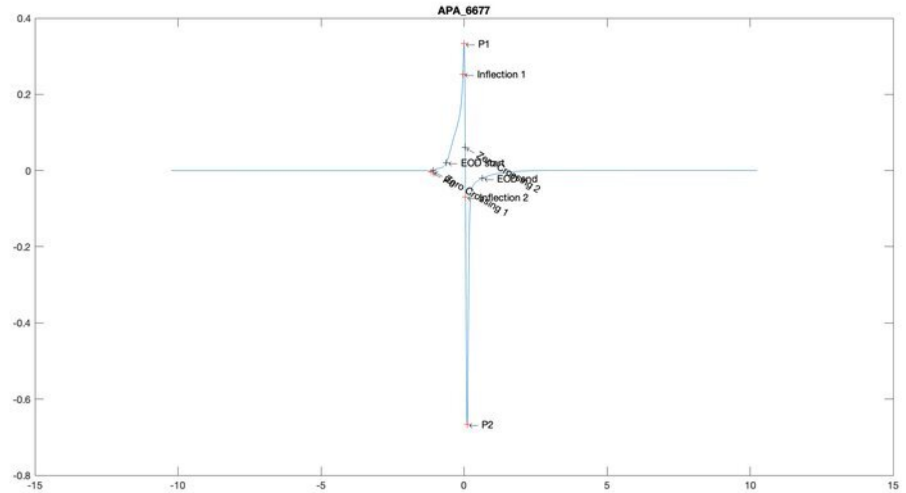
APA_6677
P1
Inflection 1
EOD start
P2

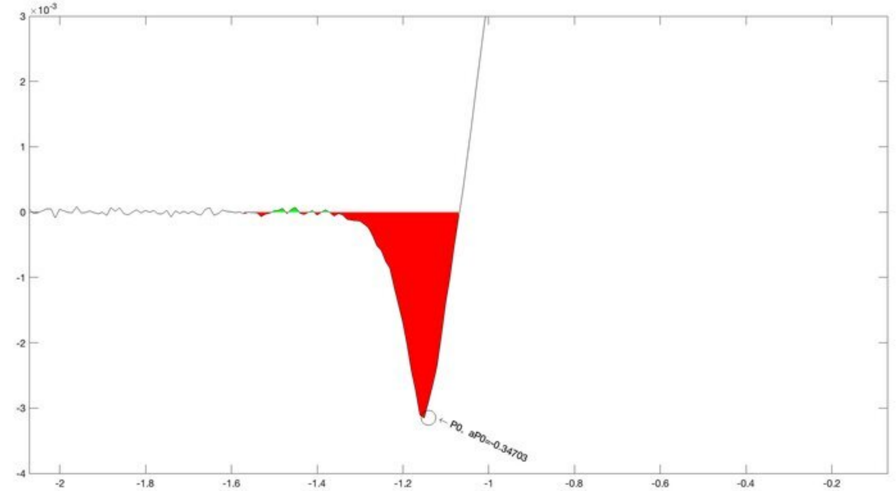
P0, aP0=-0.34703

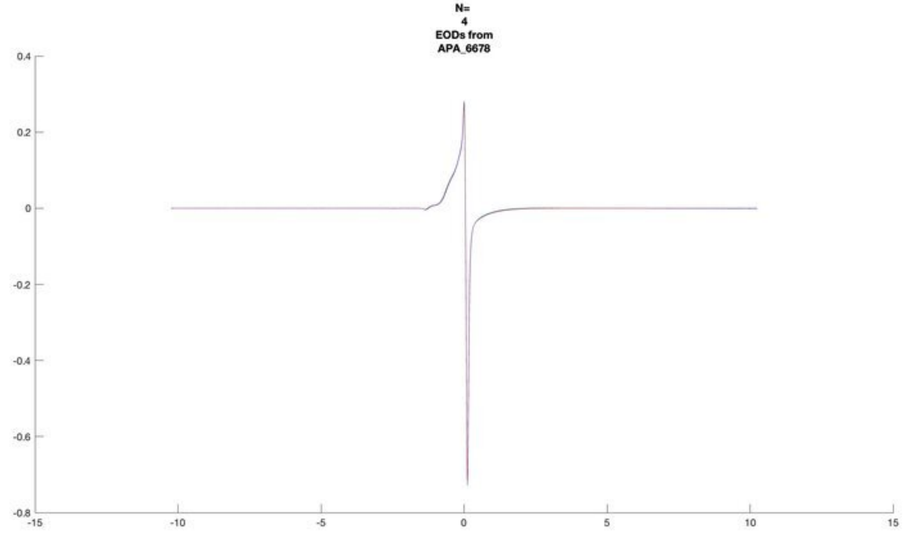
N=
4
EODs from
APA_6678

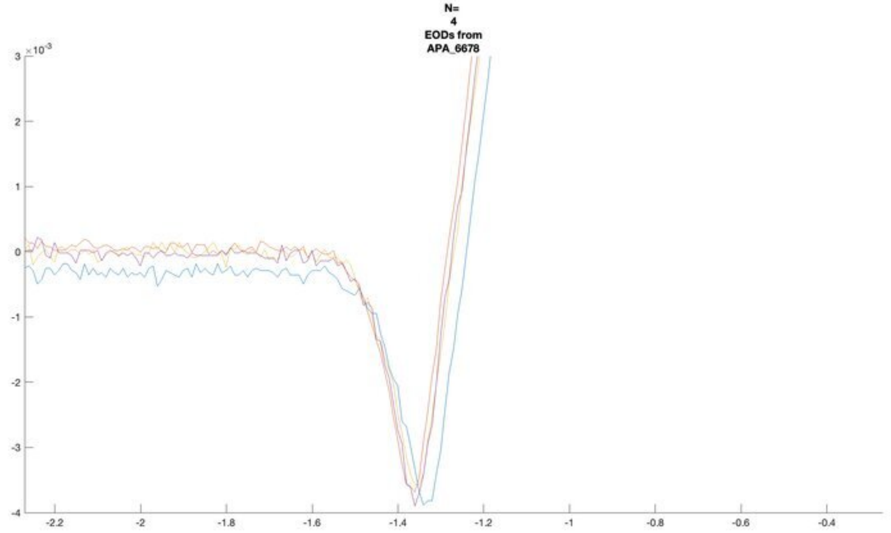
N=
4
EODs from
APA_6678
×10^-3

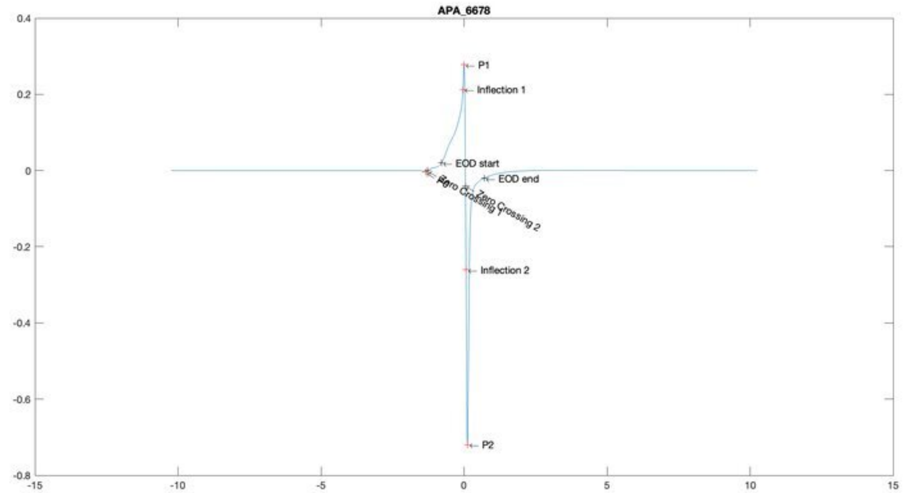
APA_6678
P1
Inflection 1
EOD start
EOD end
Zero Crossing 1
Zero Crossing 2
Inflection 2
P2

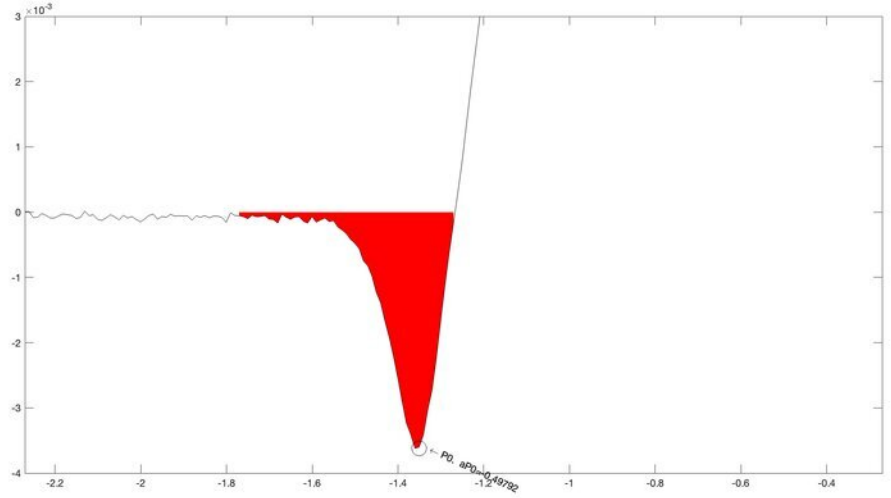
×10^-3
P0, aP0=-0.49792

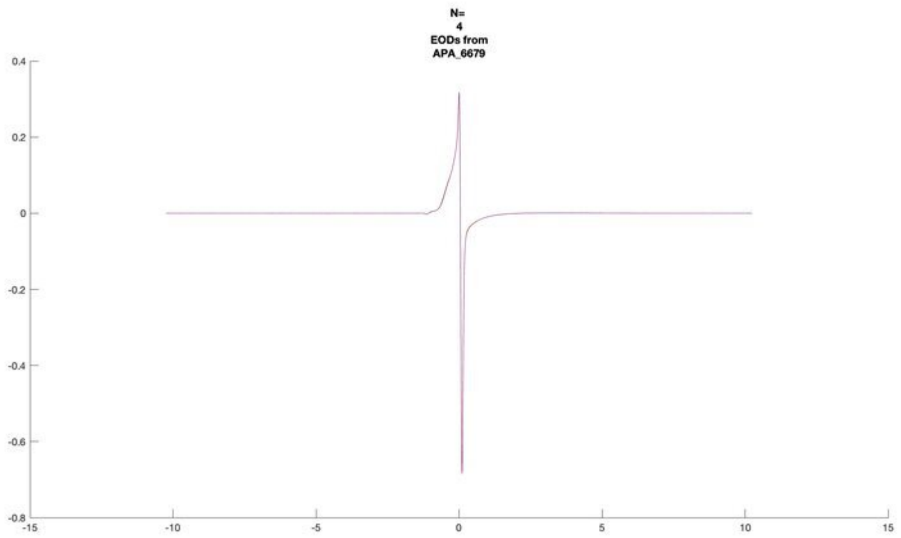
N=
4
EODs from
APA_6679

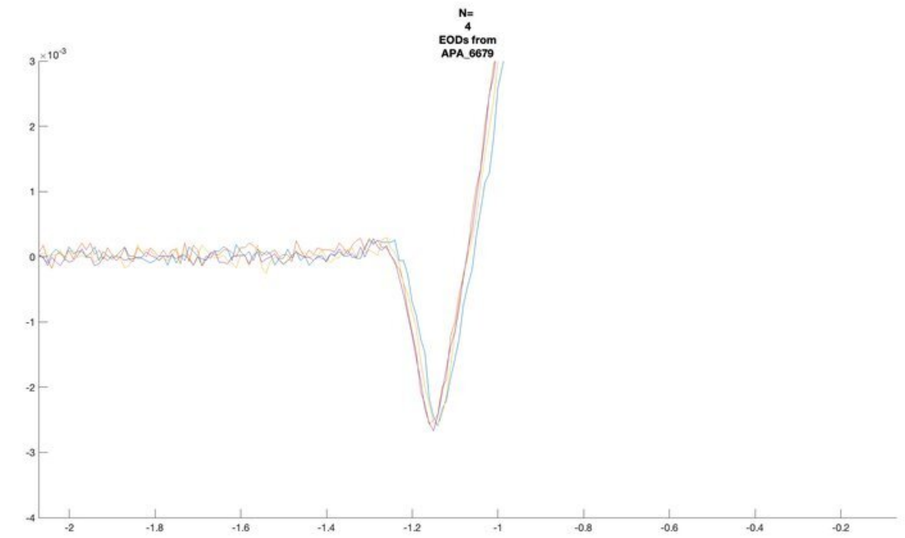
N=
4
EODs from
APA_6679

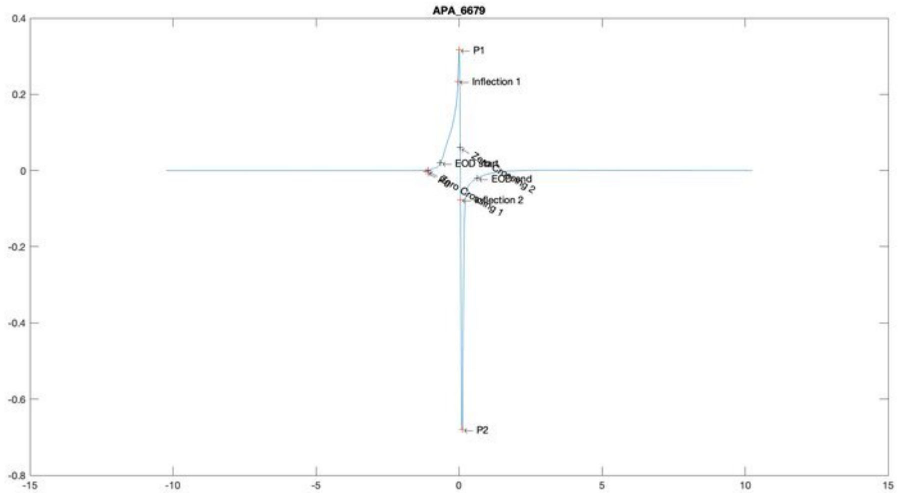
APA_6679
P1
Inflection 1
EOD start
EOD end
Inflection 2
P2

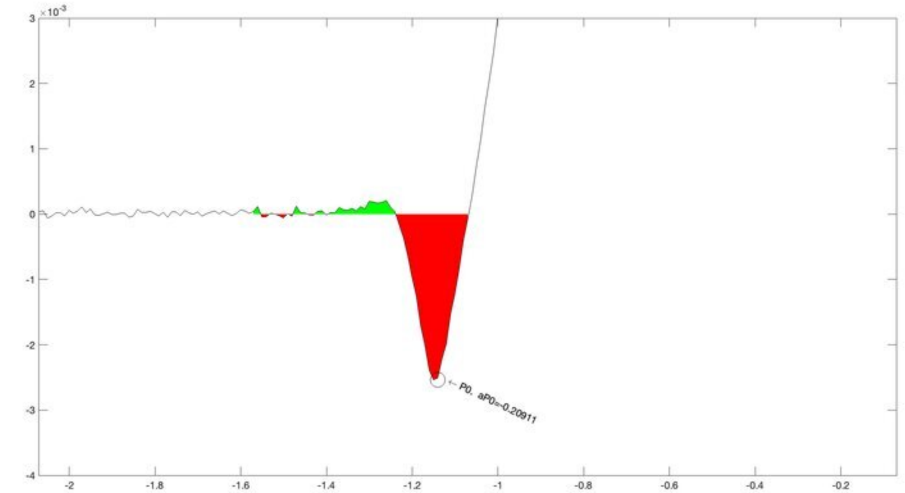
P0, aP0=-0.20911

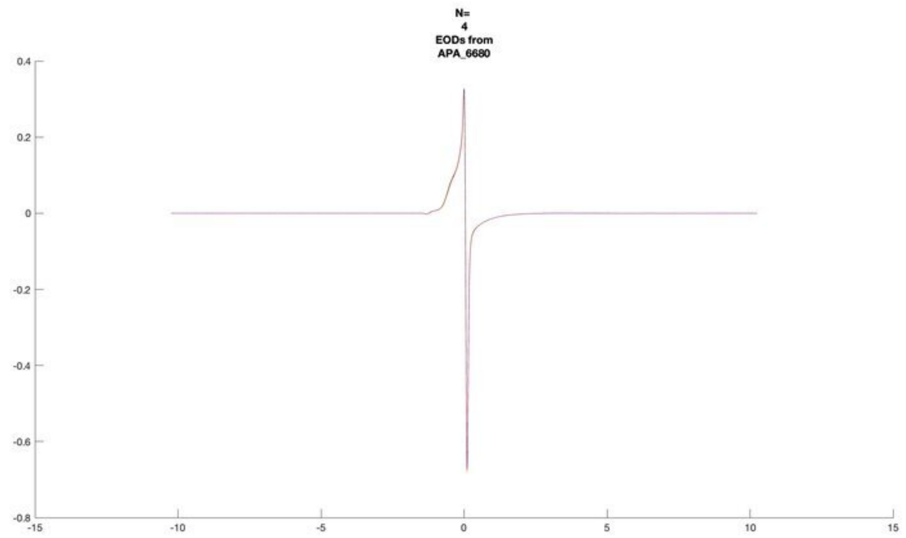
N=
4
EODs from
APA_6680

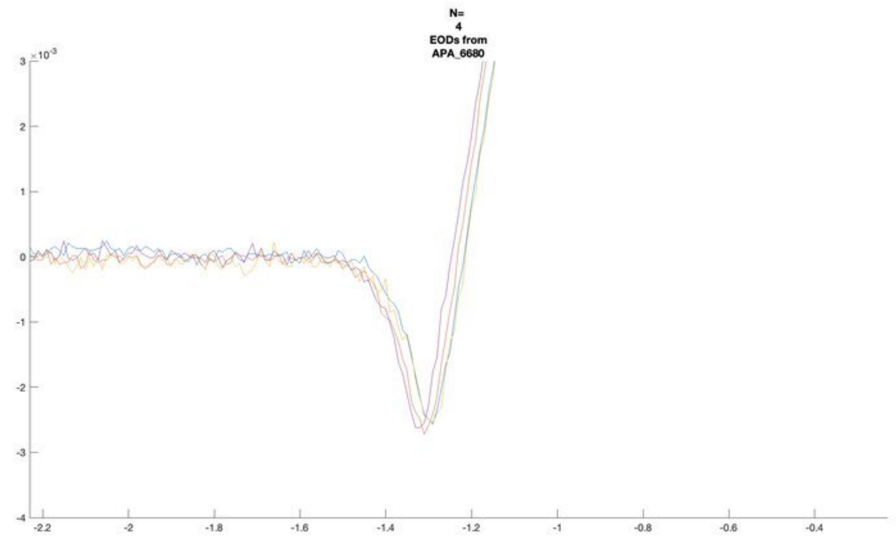
N=
4
EODs from
APA_6680

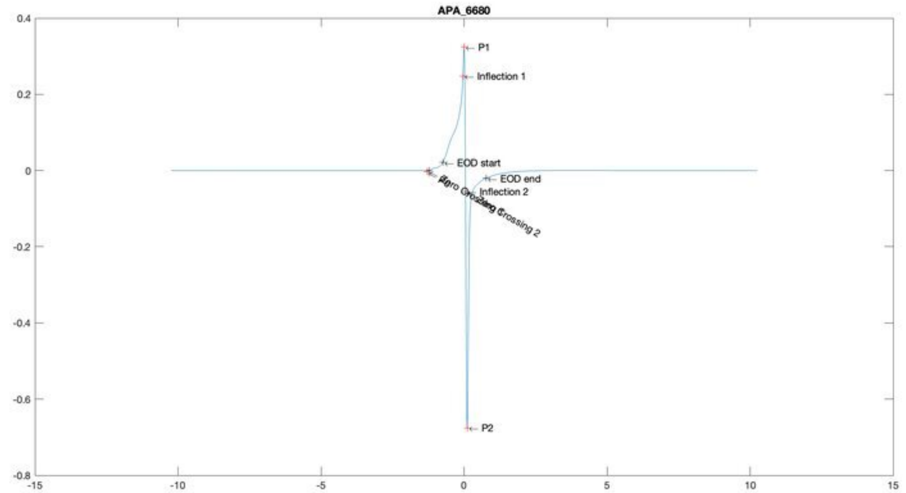
APA_6680
P1
Inflection 1
EOD start
EOD end
Inflection 2
P2

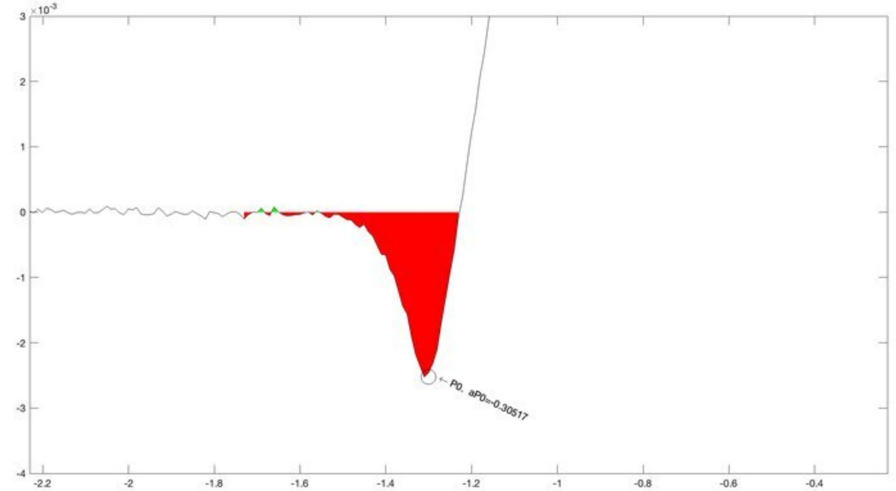
P0, aP0=-0.30517

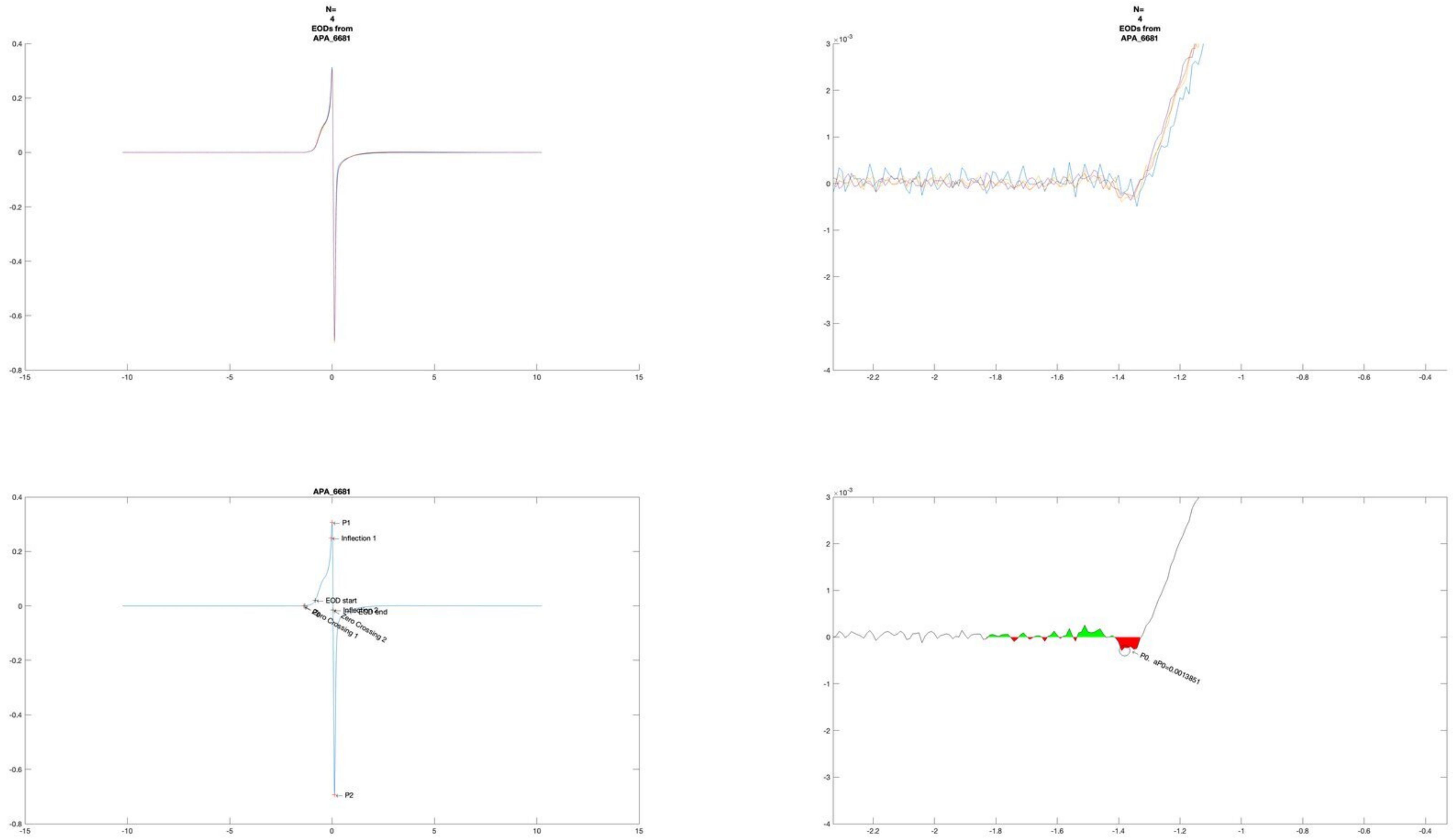
N=
4
EODs from
APA_6681
N=
4
EODs from
APA_6681
APA_6681
P1
Inflection 1
EOD start
Inflection 2
EOD end
Zero Crossing 1
Zero Crossing 2
P2
P0. aP0=0.0013851

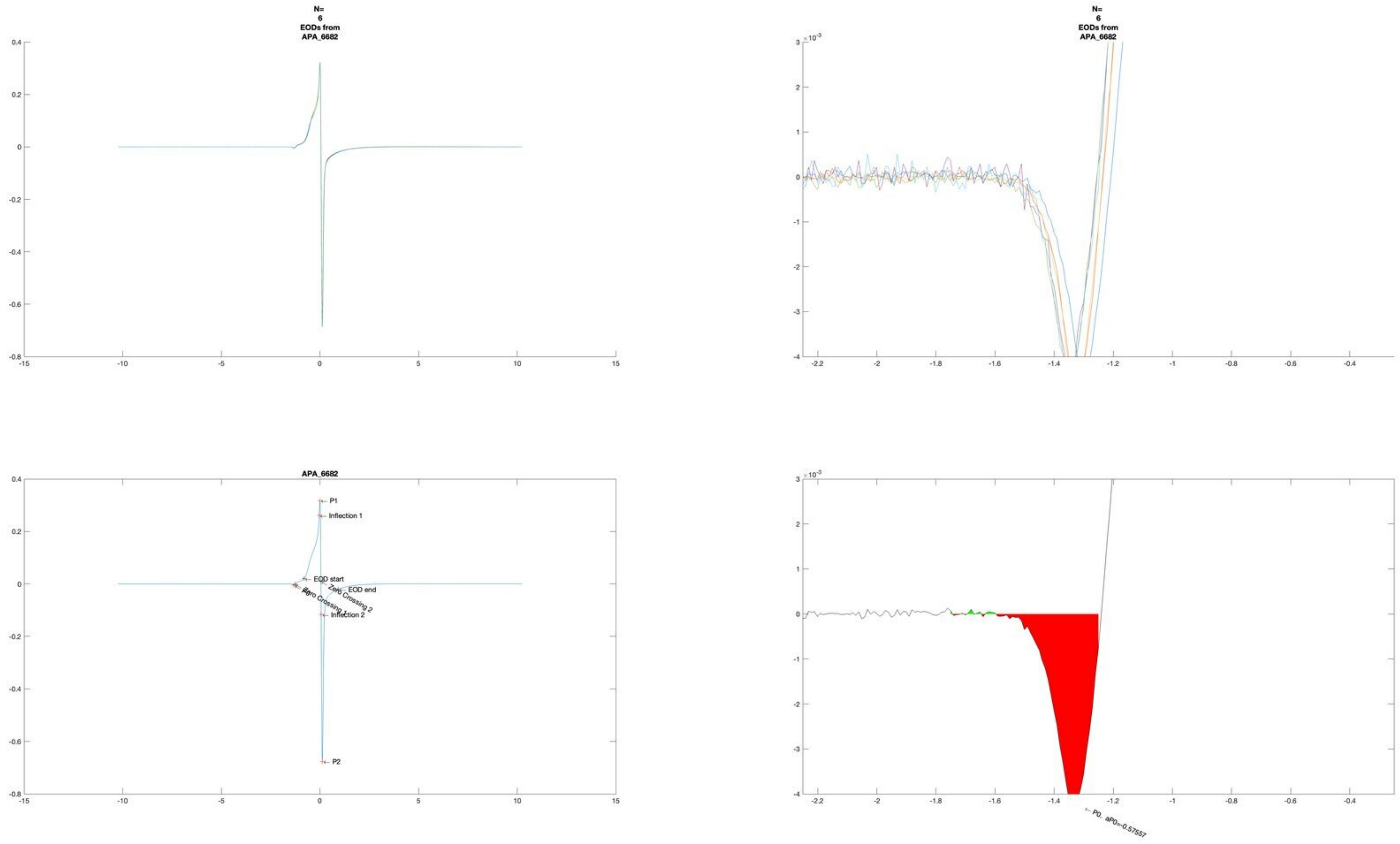
N=
6
EODs from
APA_6682
N=
6
EODs from
APA_6682
APA_6682
P1
Inflection 1
EOD start
EOD end
Zero Crossing 2
Zero Crossing 1
P0
Inflection 2
P2
P0, aP0=-0.57557

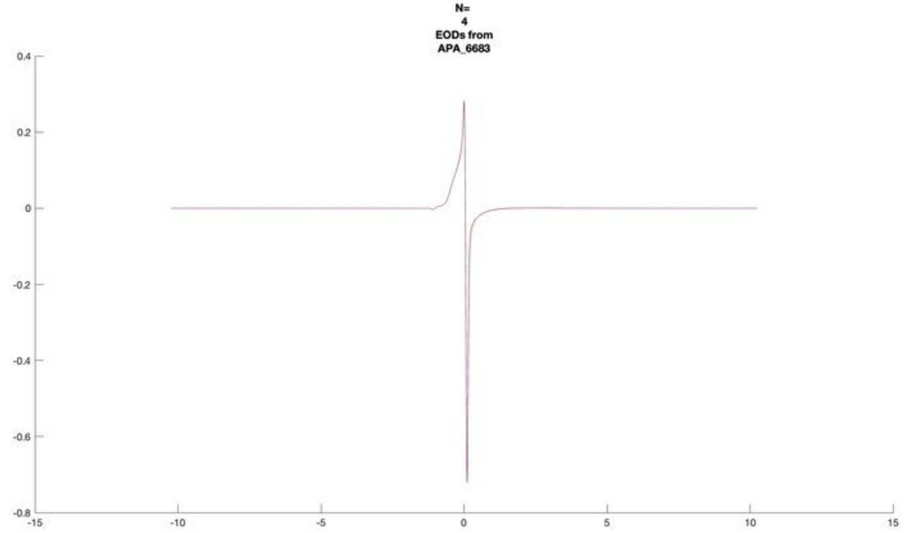
N=
4
EODs from
APA_6683

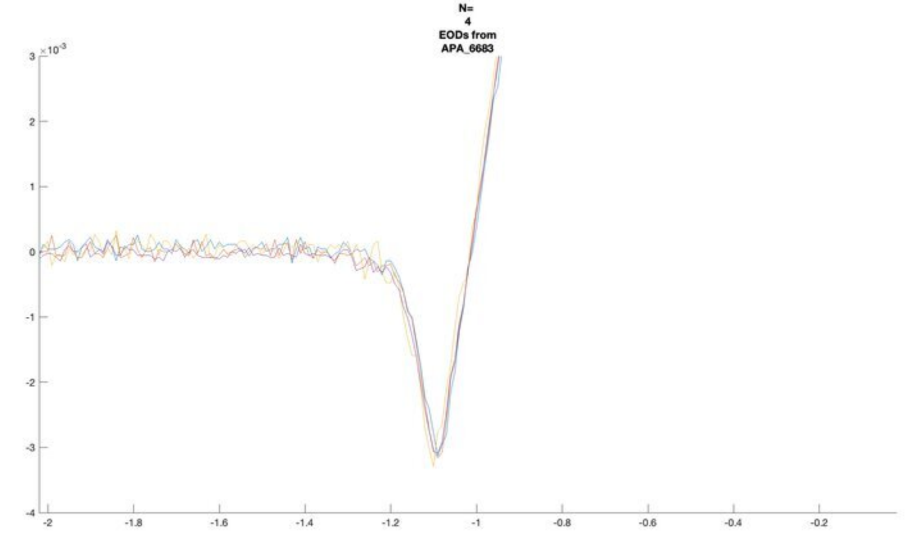
N=
4
EODs from
APA_6683

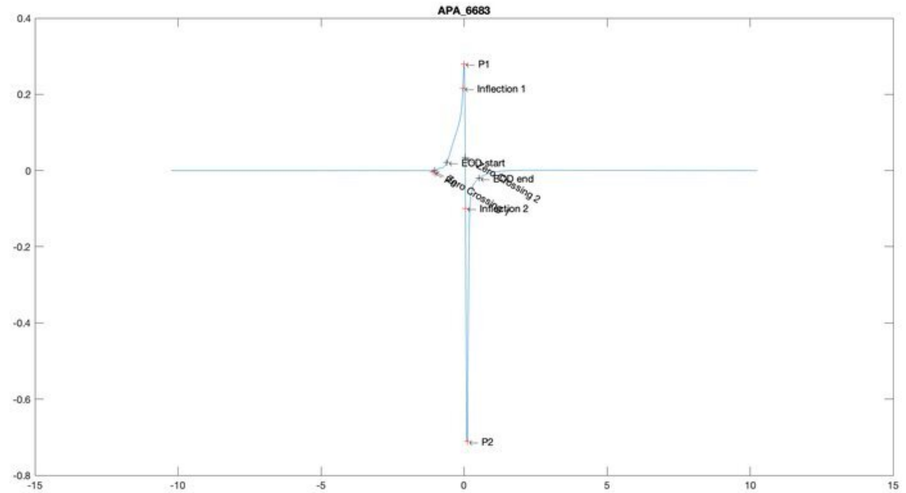
APA_6683
P1
Inflection 1
EOD start
EOD end
Inflection 2
P2

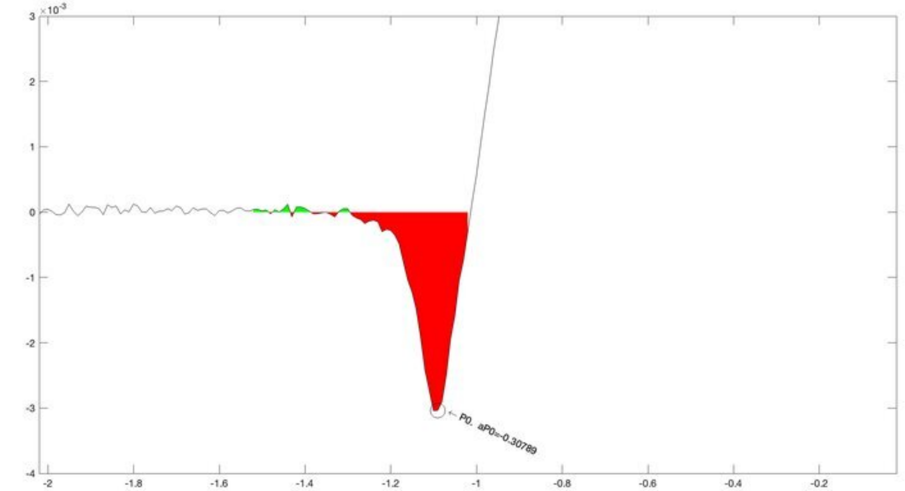
P0, aP0=-0.30789

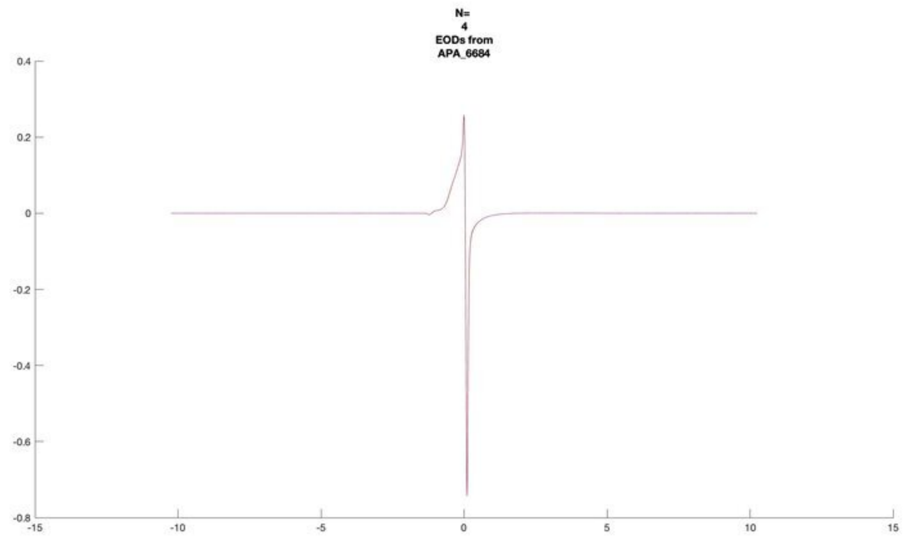
N=
4
EODs from
APA_6684

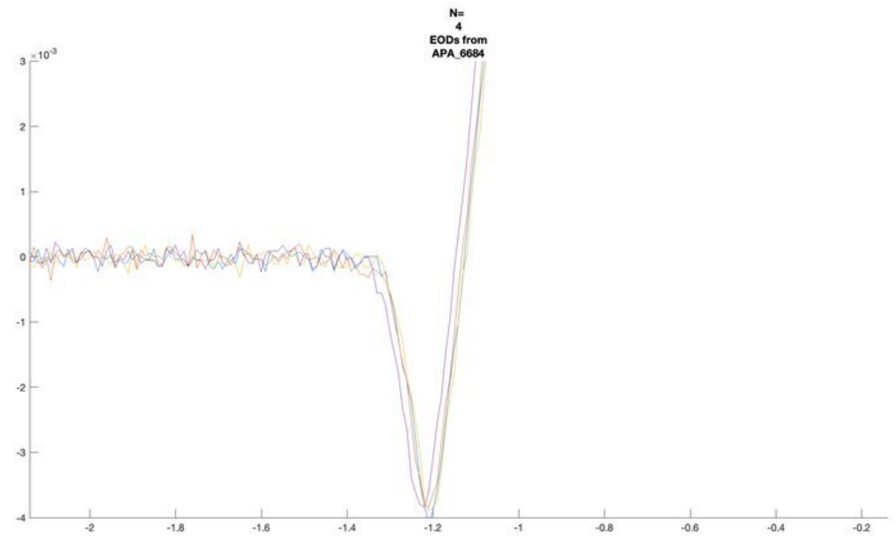
N=
4
EODs from
APA_6684

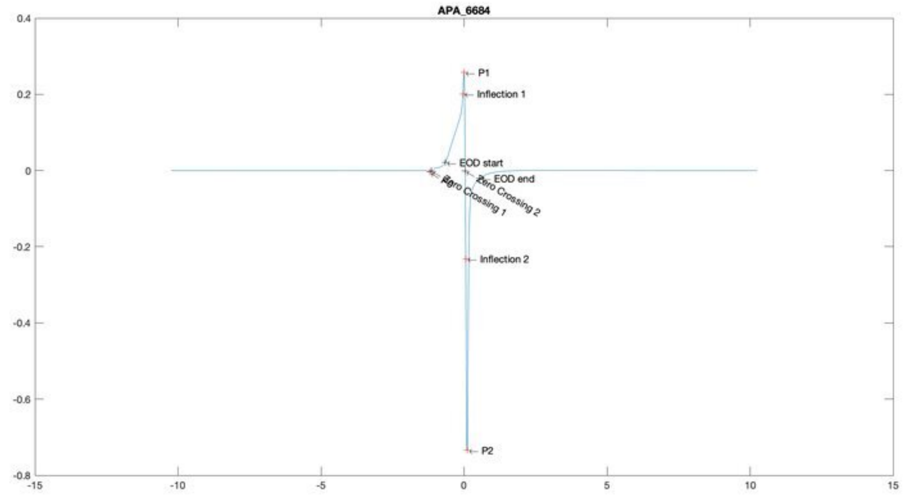
APA_6684
P1
Inflection 1
EOD start
EOD end
Zero Crossing 2
Zero Crossing 1
Inflection 2
P2

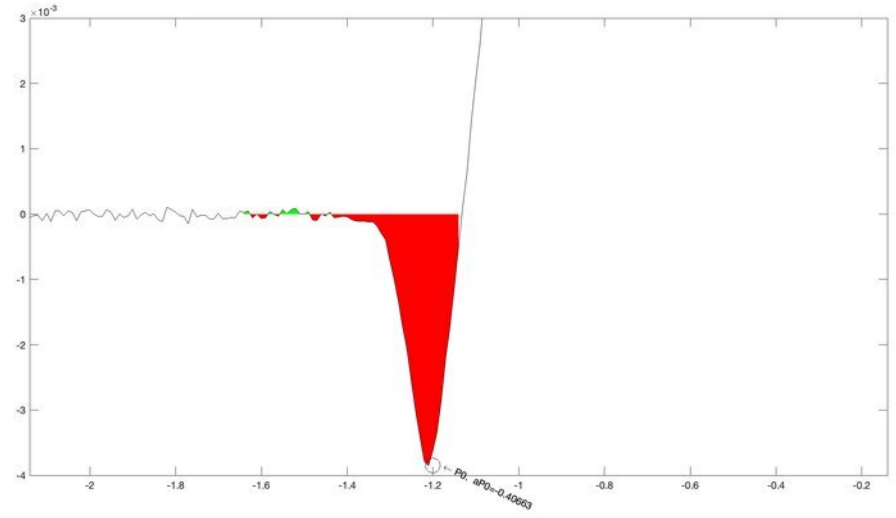
P0, aP0=-0.40663

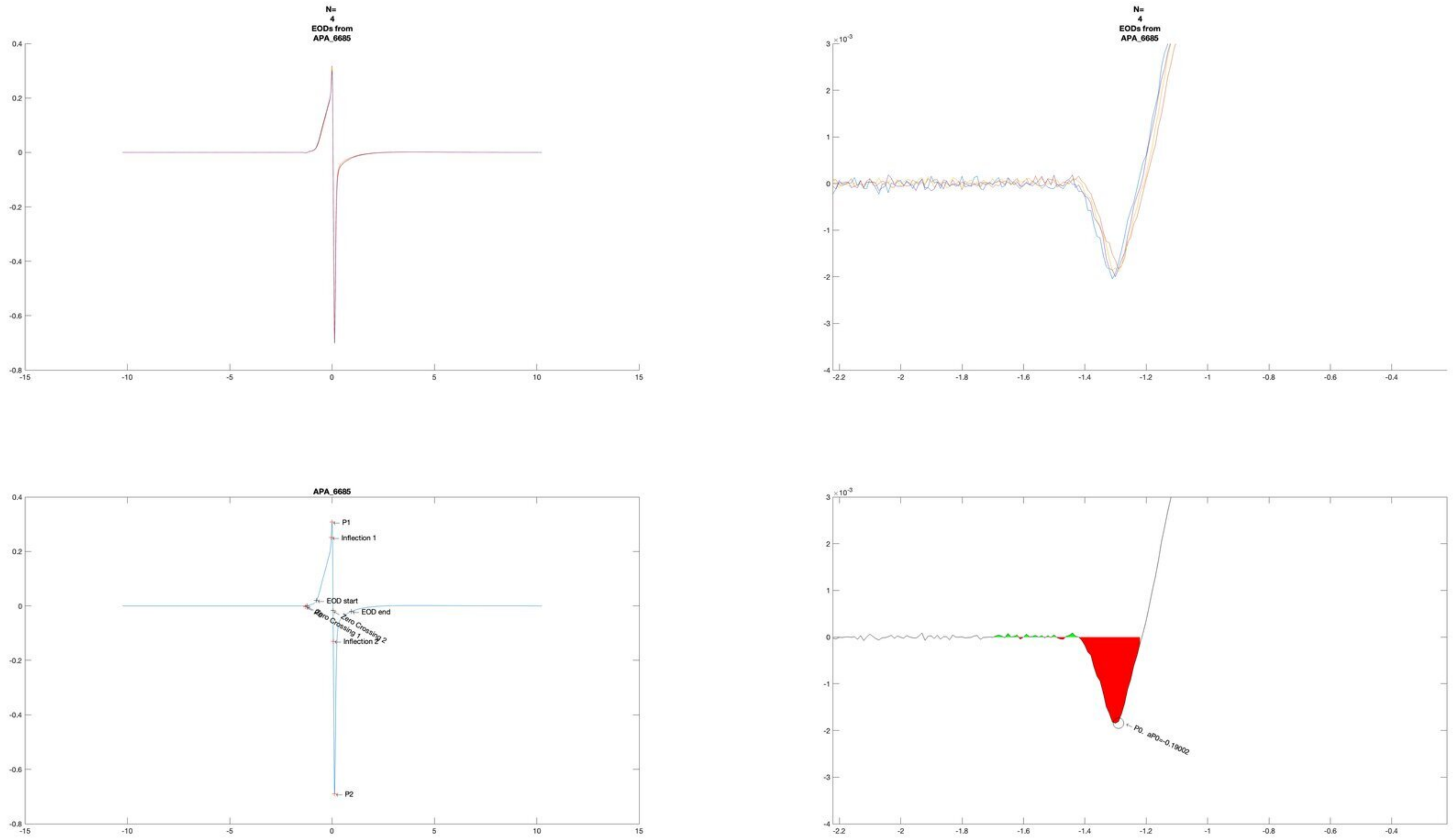
N=
4
EODs from
APA_6685
N=
4
EODs from
APA_6685
APA_6685
P1
Inflection 1
EOD start
EOD end
Zero Crossing 1
Zero Crossing 2
Inflection 2
P2
P0
P0, aP0=-0.19002
×10^-3

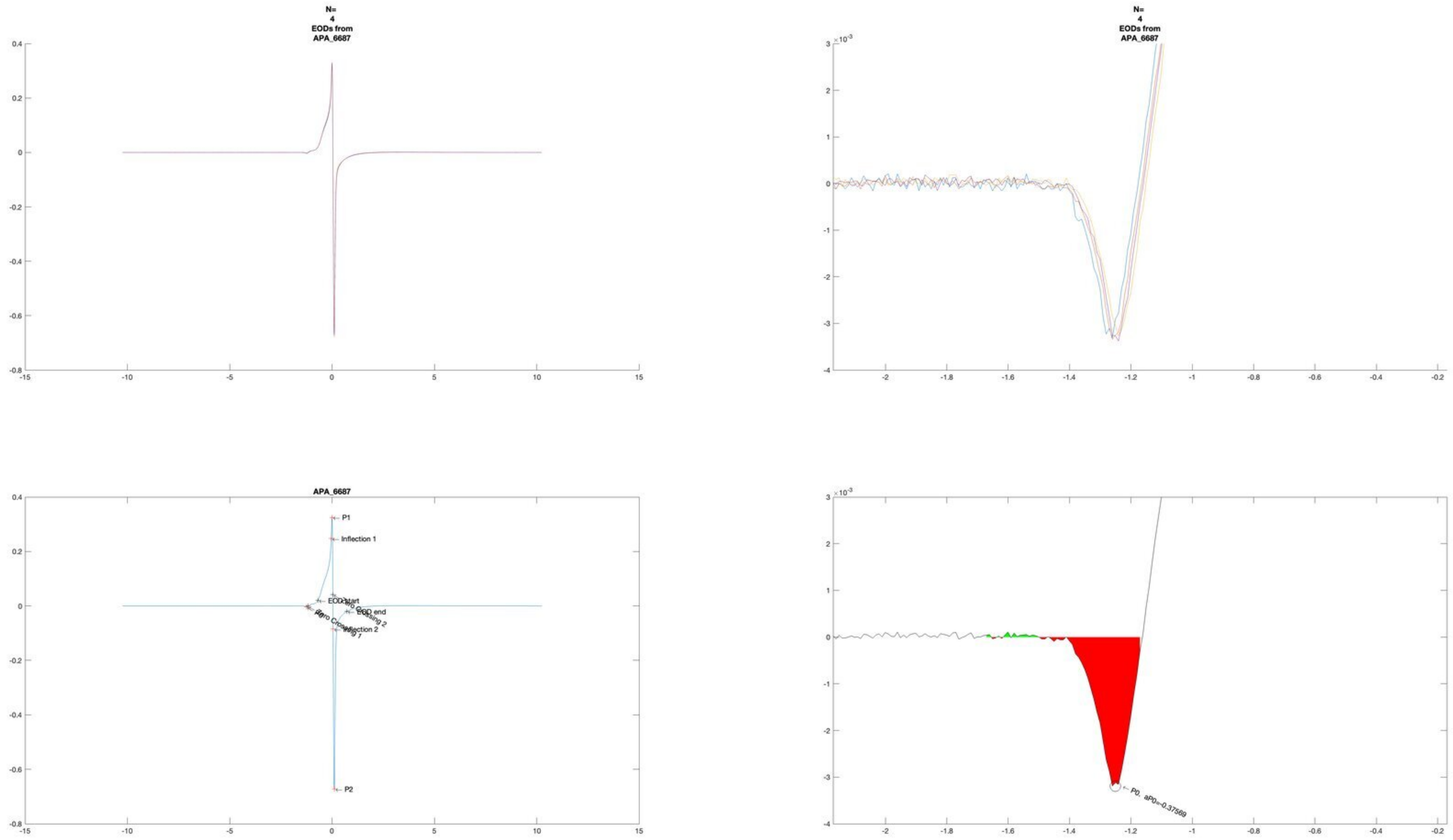
N=
4
EODs from
APA_6687
N=
4
EODs from
APA_6687
APA_6687
P1
Inflection 1
EOD start
EOD end
Inflection 2
P2
P0, aP0=-0.37569

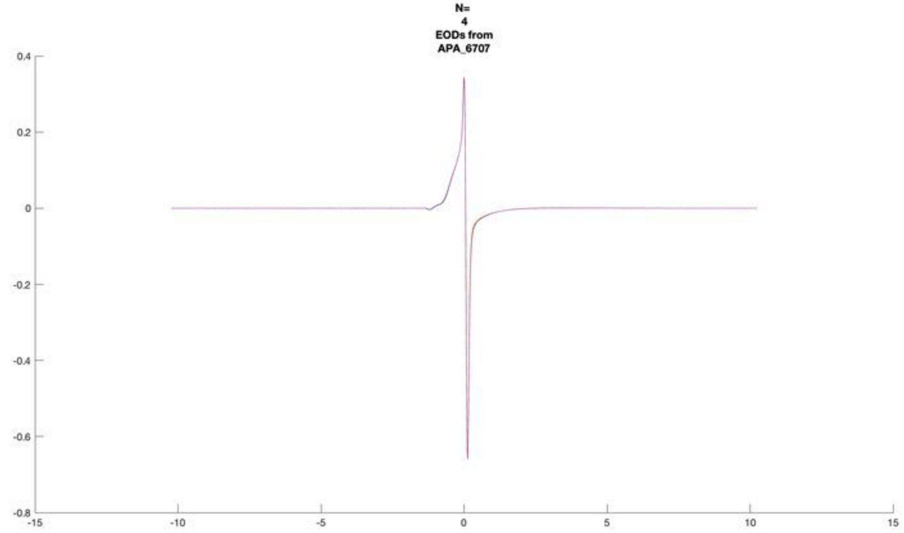
N=
4
EODs from
APA_6707

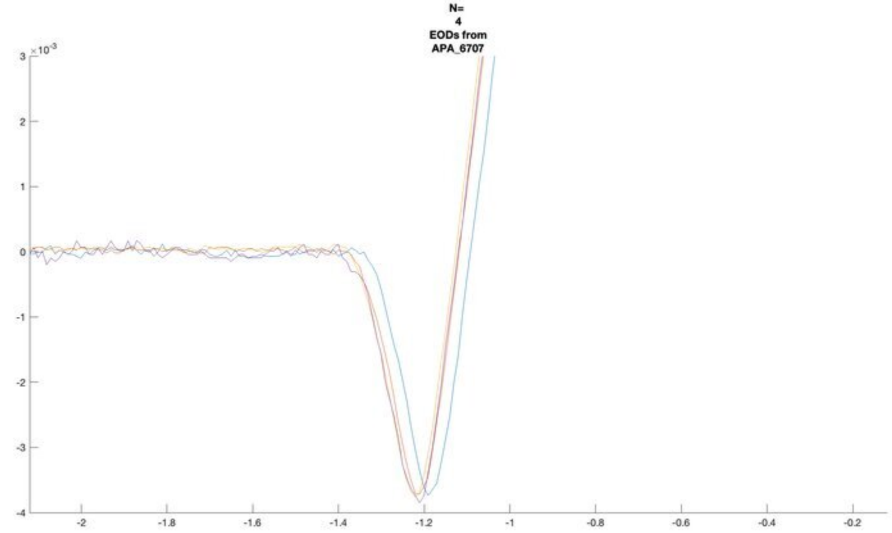
N=
4
EODs from
APA_6707
×10^-3

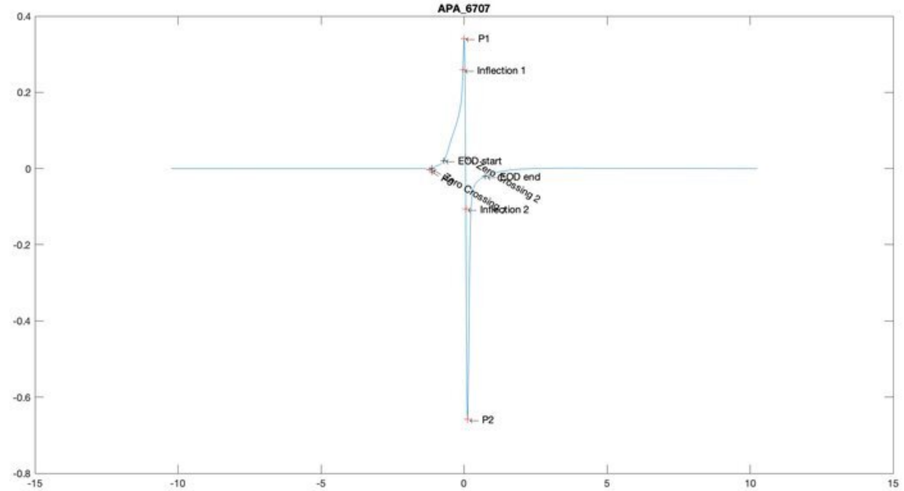
APA_6707
P1
Inflection 1
EOD start
EOD end
Zero Crossing 2
Inflection 2
P2

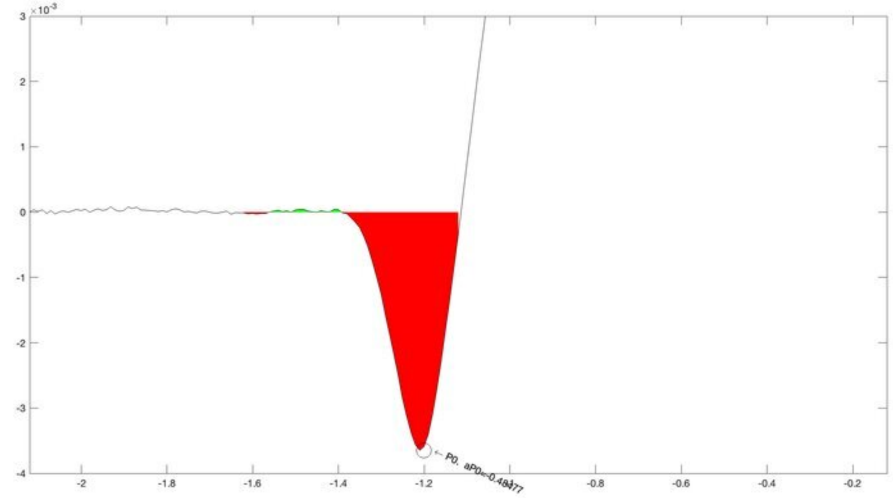
×10^-3

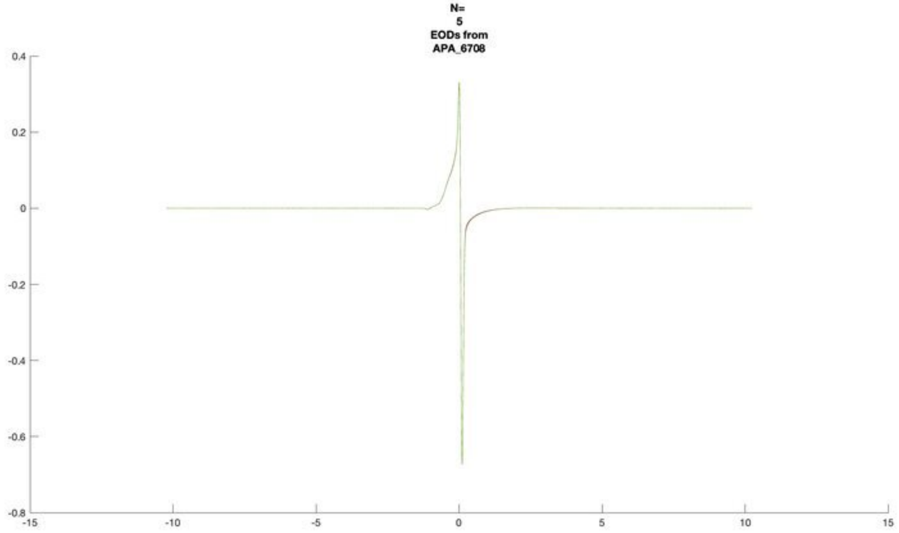
N=
5
EODs from
APA_6708

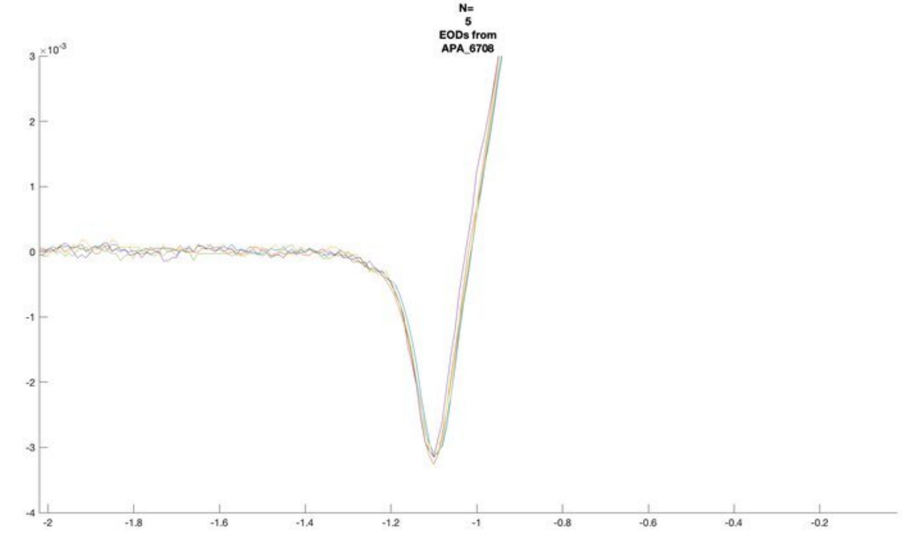
N=
5
EODs from
APA_6708
×10^-3

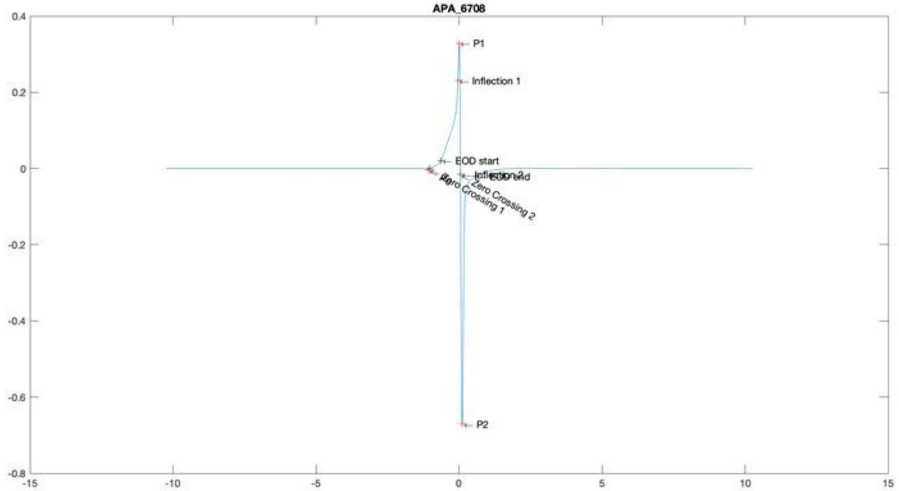
APA_6708
P1
Inflection 1
EOD start
Inflection 2
EOD end
Zero Crossing 2
Zero Crossing 1
P2

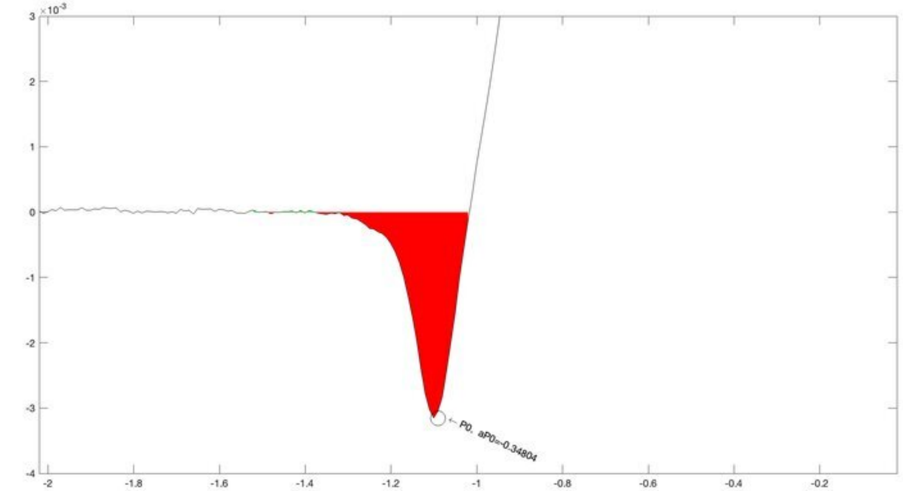
×10^-3
P0, aP0=-0.34804

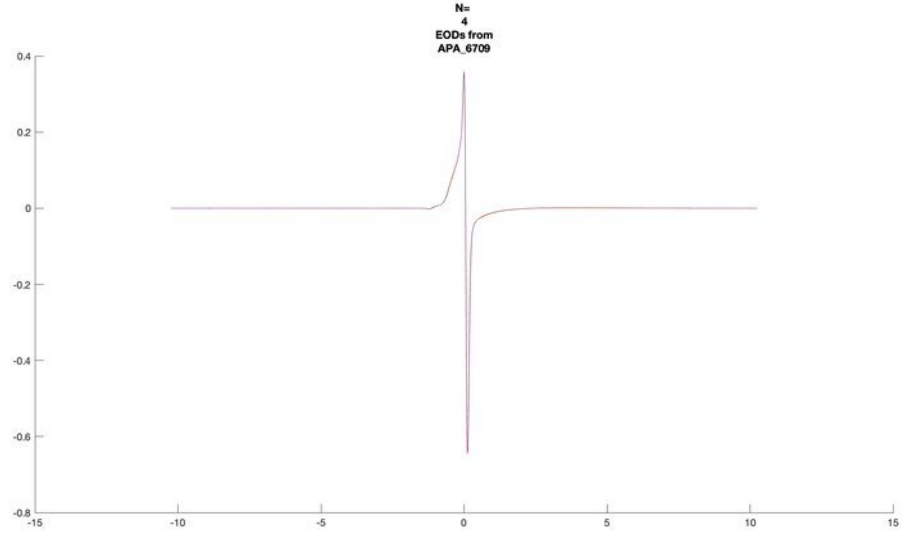
N=
4
EODs from
APA_6709

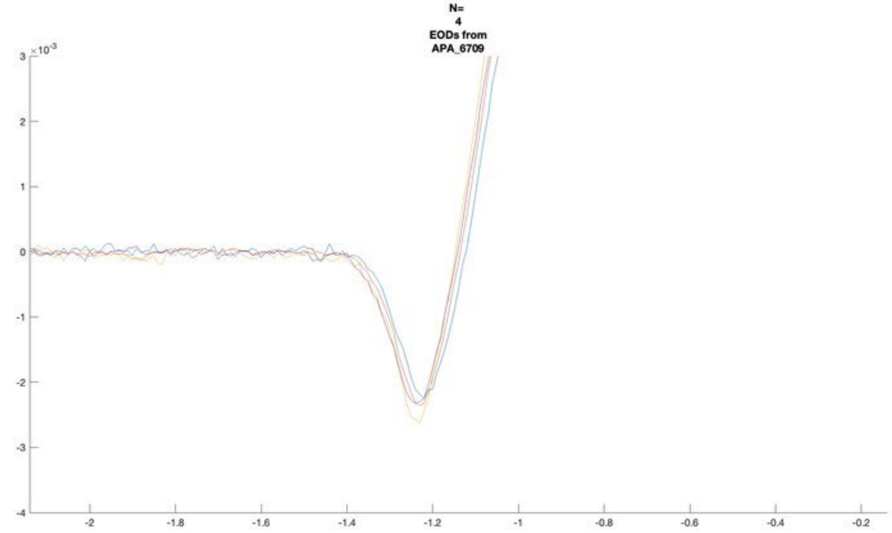
N=
4
EODs from
APA_6709
×10⁻³

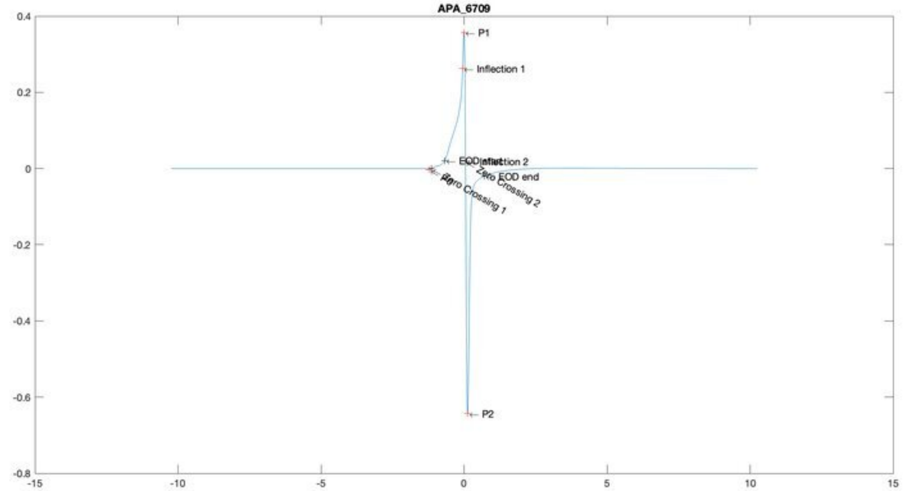
APA_6709
P1
Inflection 1
Inflection 2
EOD end
Zero Crossing 1
Zero Crossing 2
P2

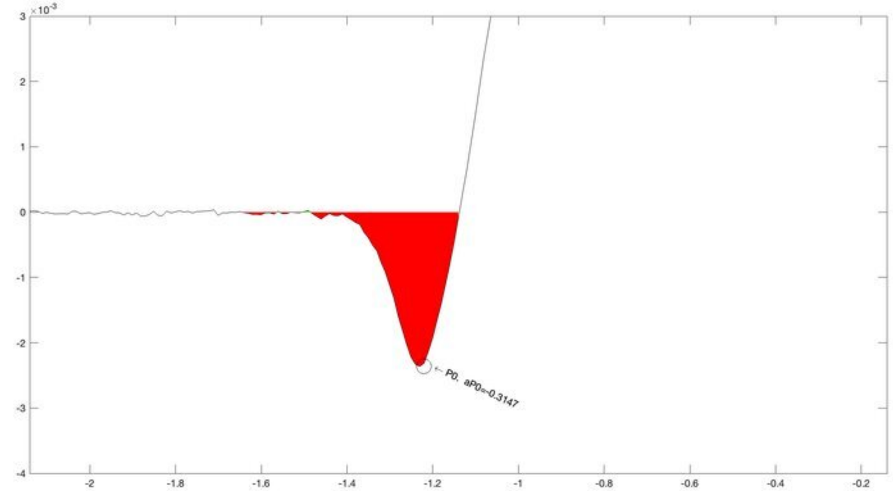
×10⁻³
P0. aP0=-0.3147

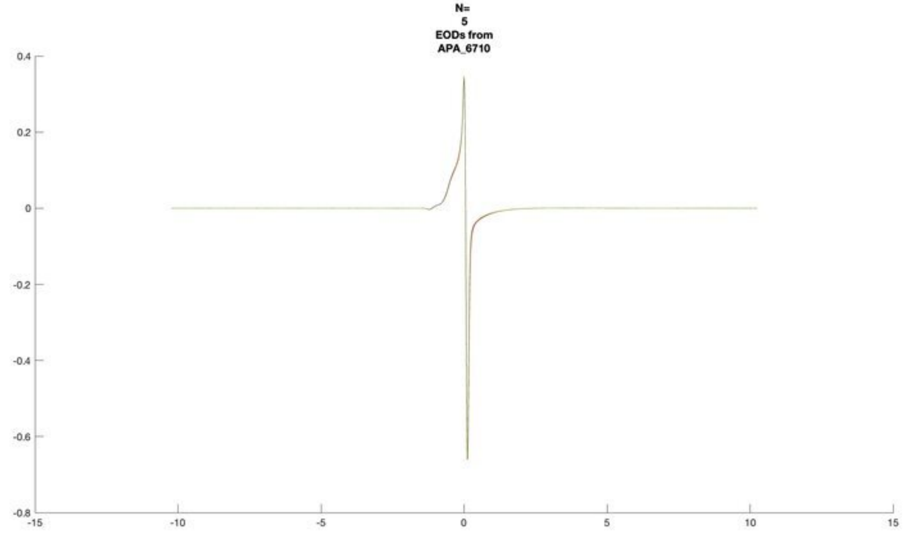
N=
5
EODs from
APA_6710

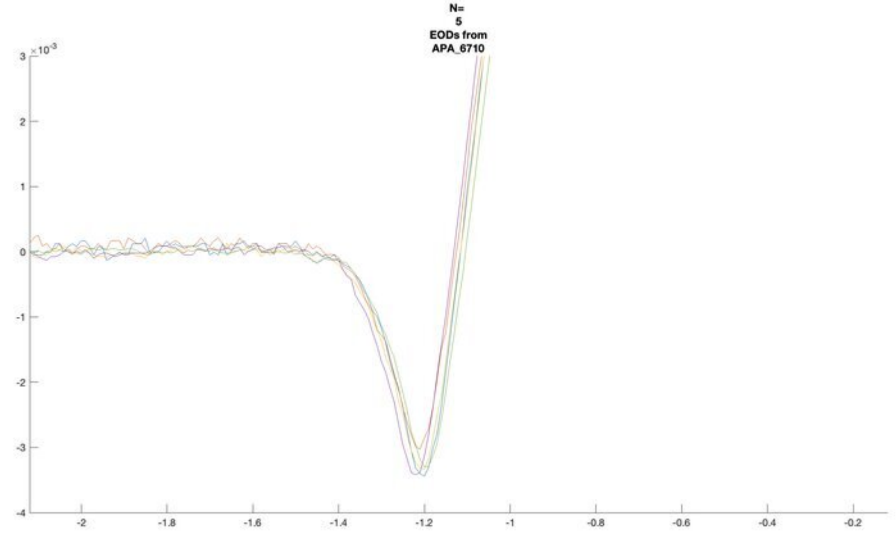
N=
5
EODs from
APA_6710

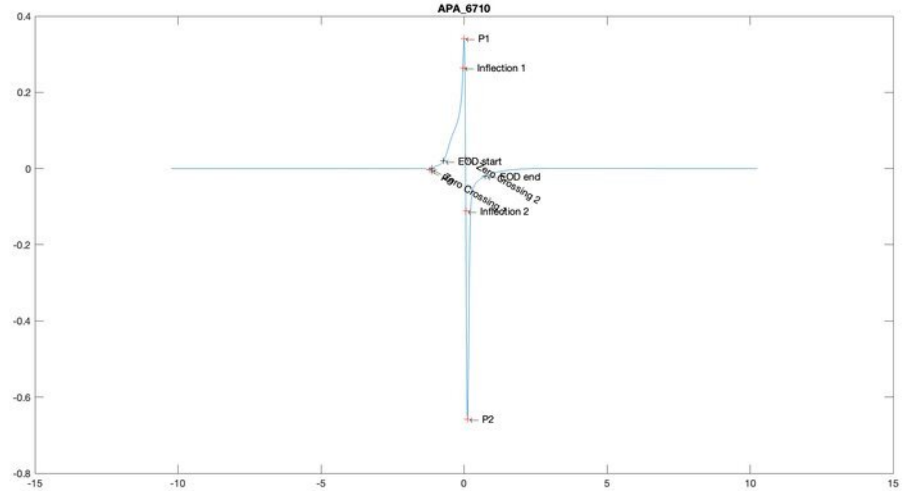
APA_6710
P1
Inflection 1
EOD start
EOD end
Inflection 2
P2

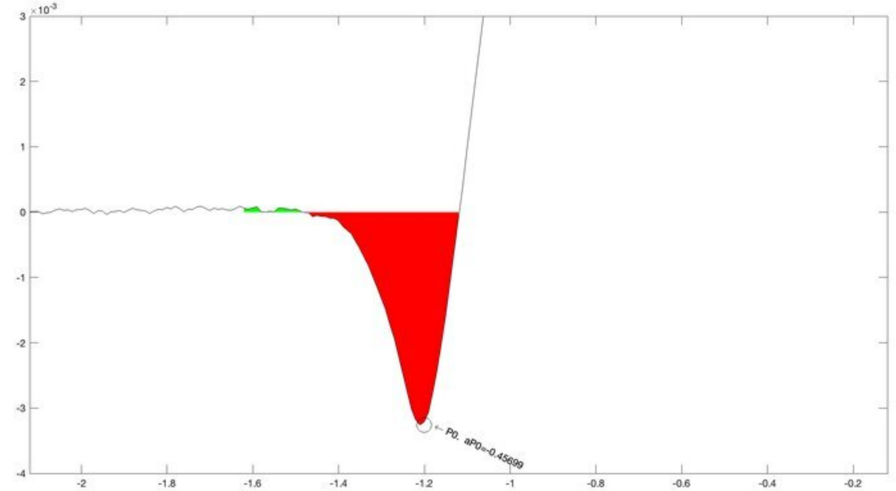
P0, aP0=-0.45699

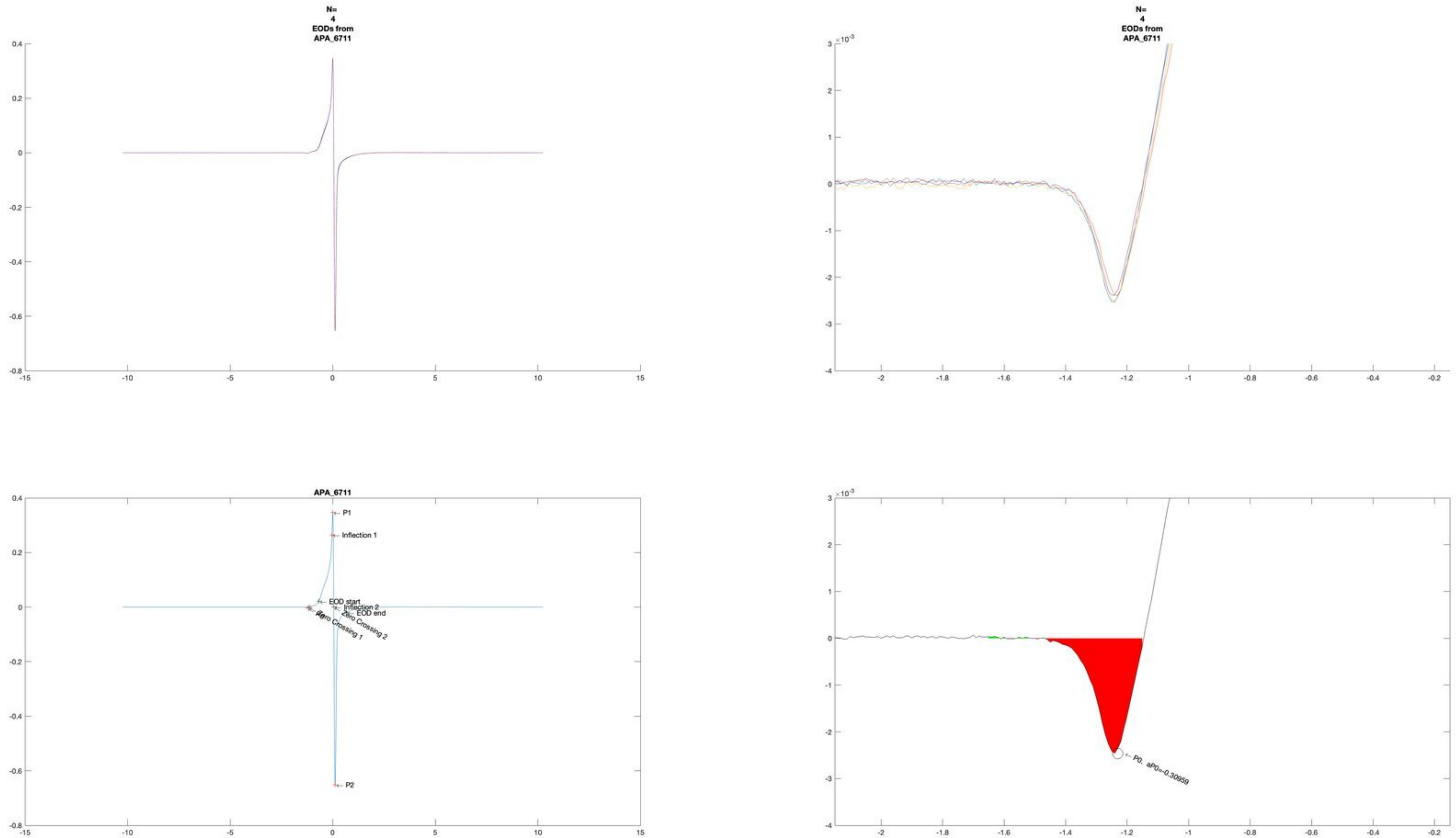
N=
4
EODs from
APA_6711
N=
4
EODs from
APA_6711
APA_6711
P1
Inflection 1
EOD start
Inflection 2
EOD end
Zero Crossing 2
Zero Crossing 1
P2
P0, aP0=-0.30959

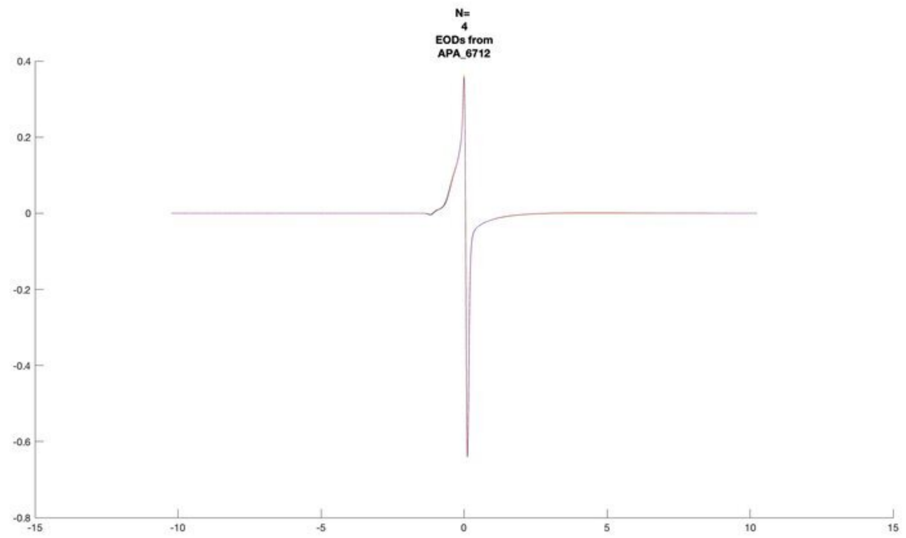
N=
4
EODs from
APA_6712

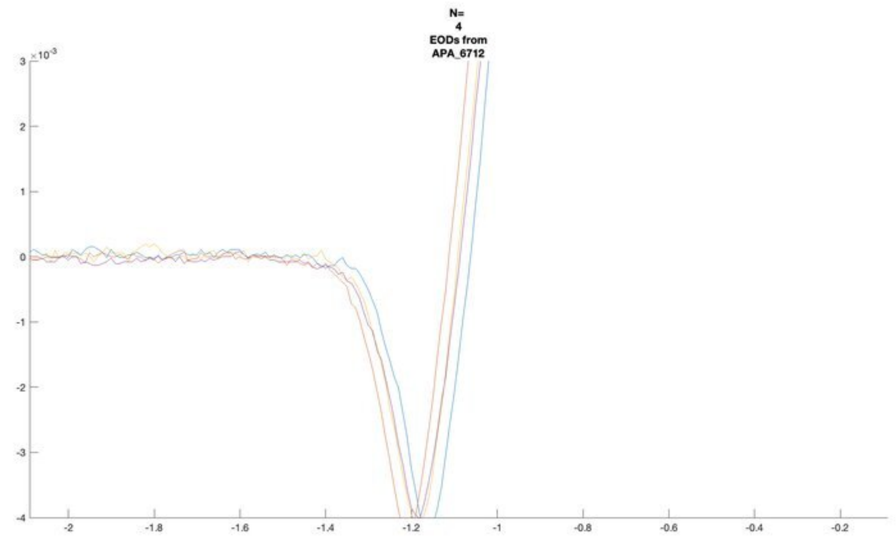
N=
4
EODs from
APA_6712

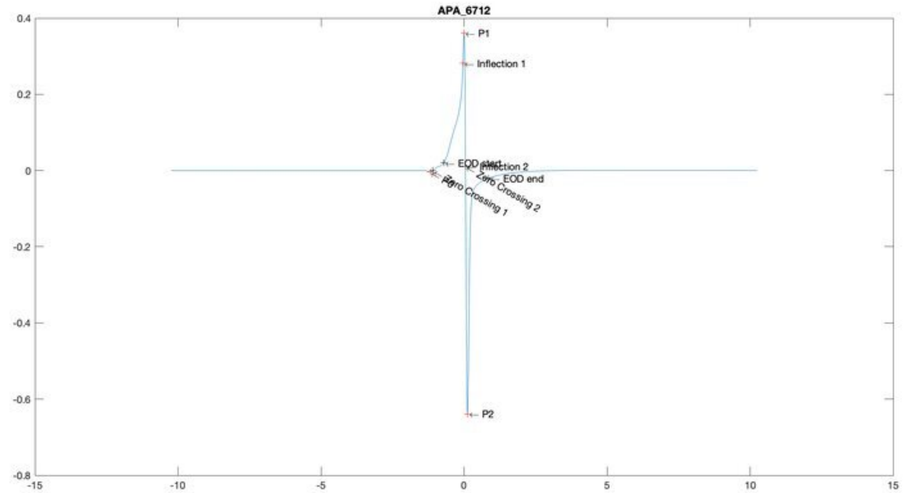
APA_6712
P1
Inflection 1
Inflection 2
EOD end
Zero Crossing 2
Zero Crossing 1
P2

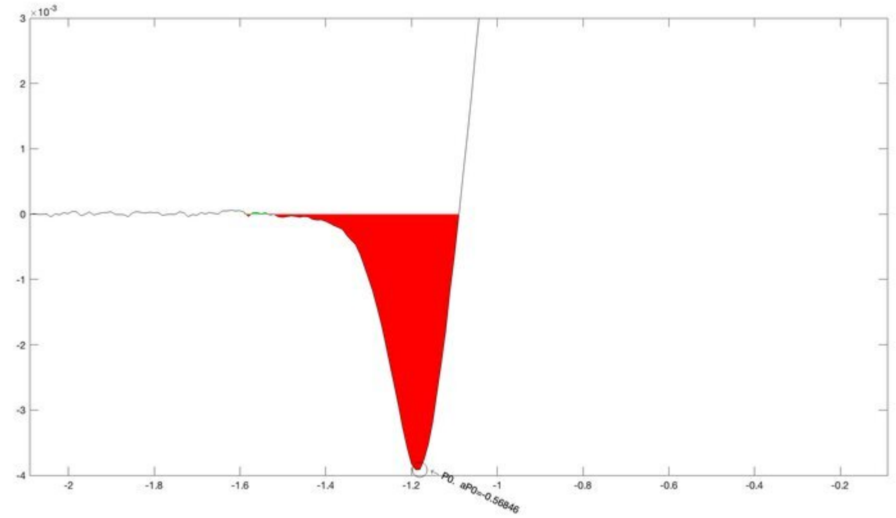
P0, aP0=-0.56846

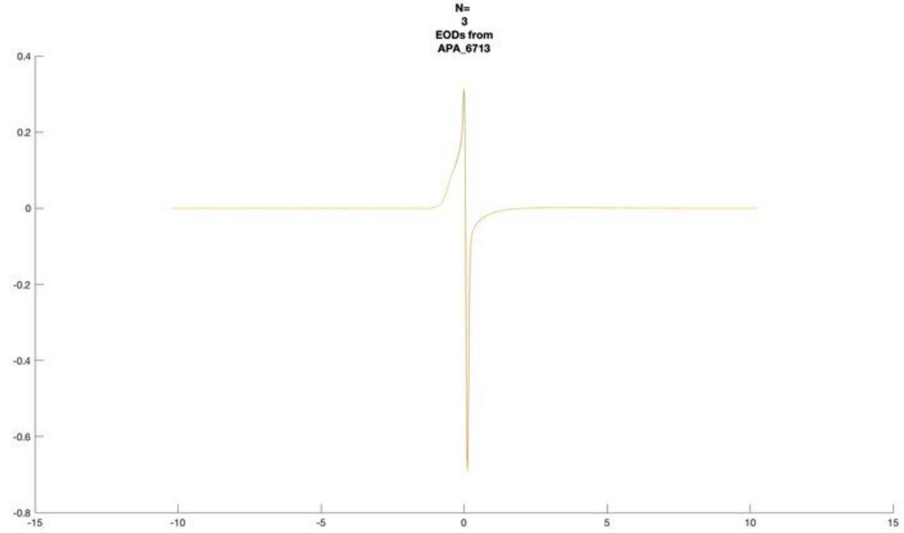
N=
3
EODs from
APA_6713
0.4
0.2
0
-0.2
-0.4
-0.6
-0.8
-15
-10
-5
0
5
10
15

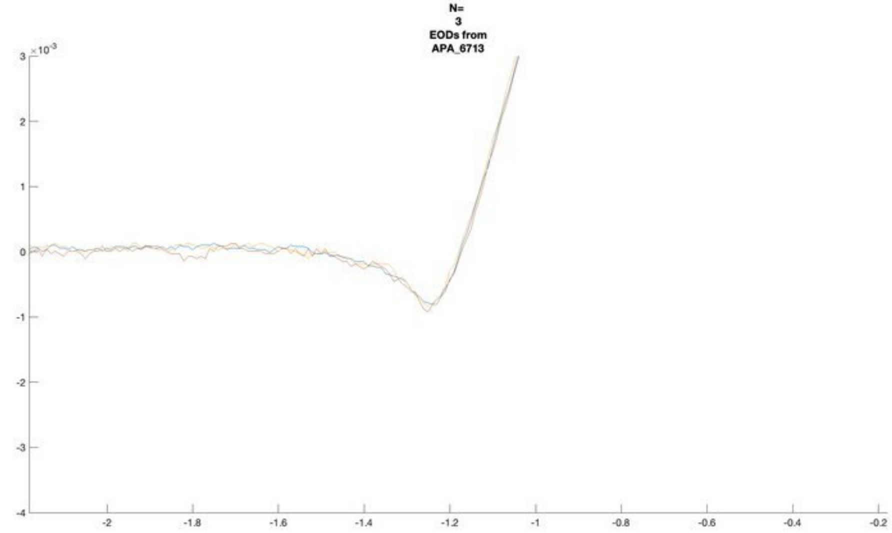
N=
3
EODs from
APA_6713
×10^-3
3
2
1
0
-1
-2
-3
-4
-2
-1.8
-1.6
-1.4
-1.2
-1
-0.8
-0.6
-0.4
-0.2

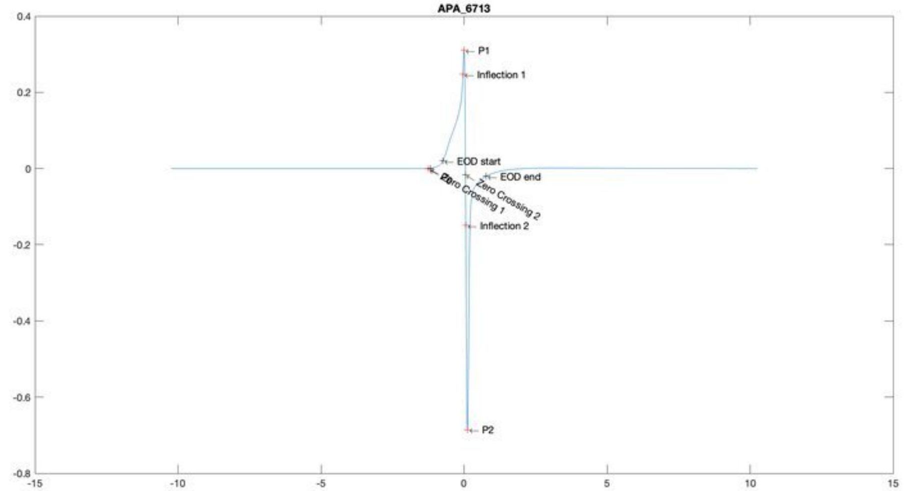
APA_6713
P1
Inflection 1
EOD start
EOD end
Zero Crossing 2
Zero Crossing 1
Inflection 2
P2
0.4
0.2
0
-0.2
-0.4
-0.6
-0.8
-15
-10
-5
0
5
10
15

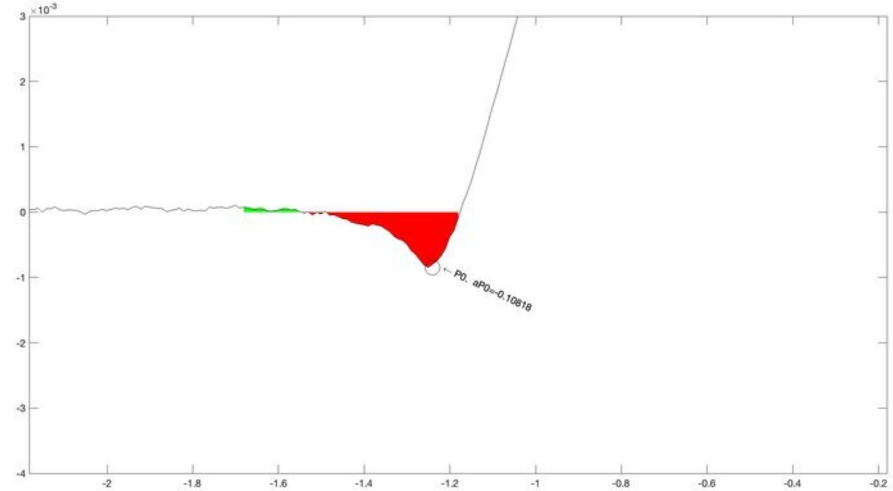
×10^-3
P0, aP0=-0.10818
3
2
1
0
-1
-2
-3
-4
-2
-1.8
-1.6
-1.4
-1.2
-1
-0.8
-0.6
-0.4
-0.2

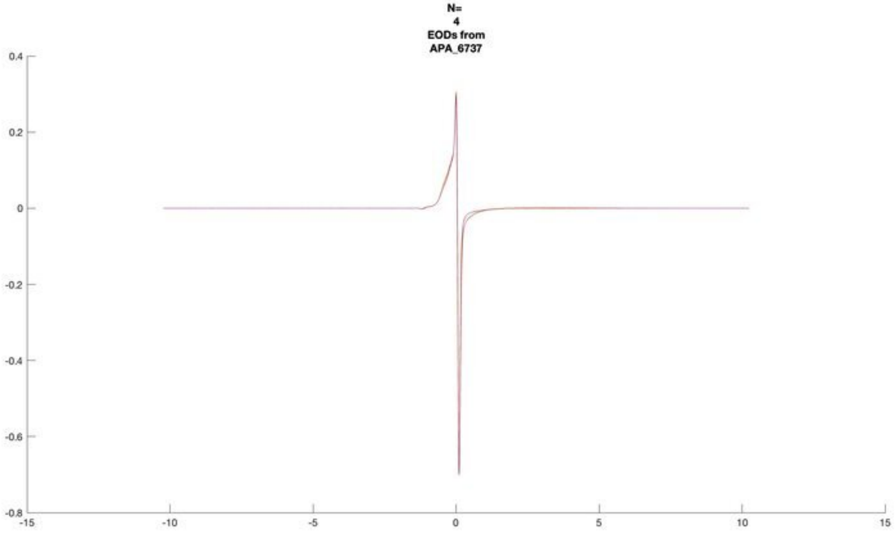
N=
4
EODs from
APA_6737

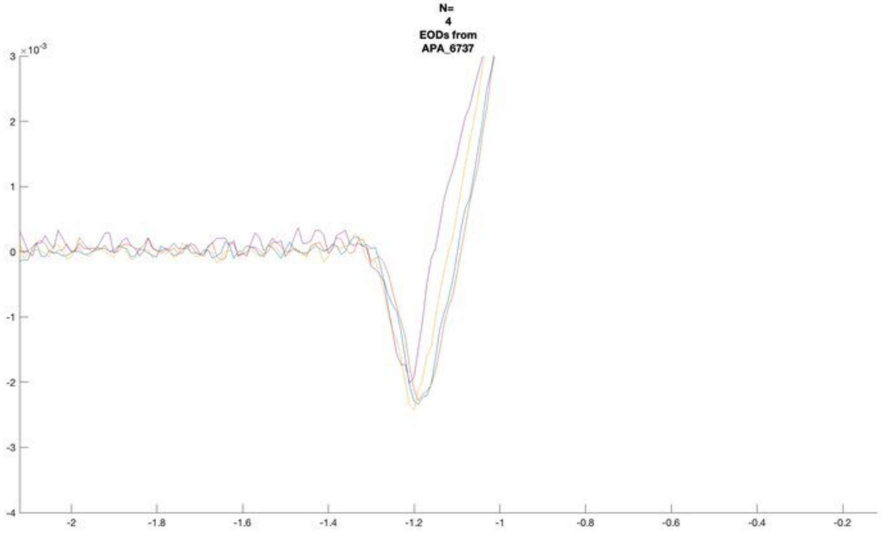
N=
4
EODs from
APA_6737
×10^-3

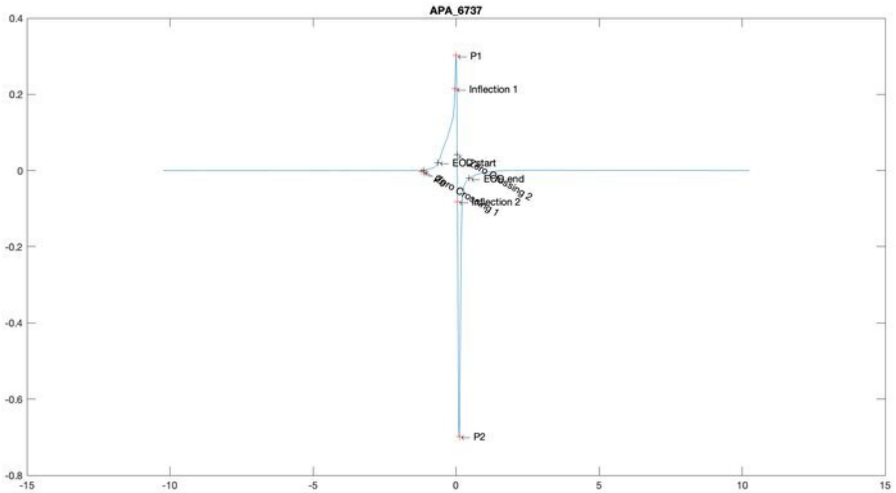
APA_6737
P1
Inflection 1
EOD start
EOD end
Inflection 2
P2

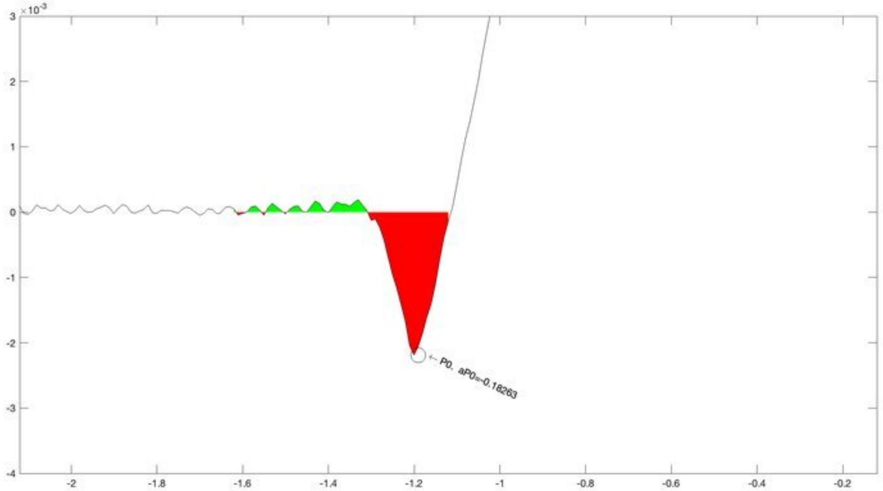
×10^-3
P0, aP0=-0.18263

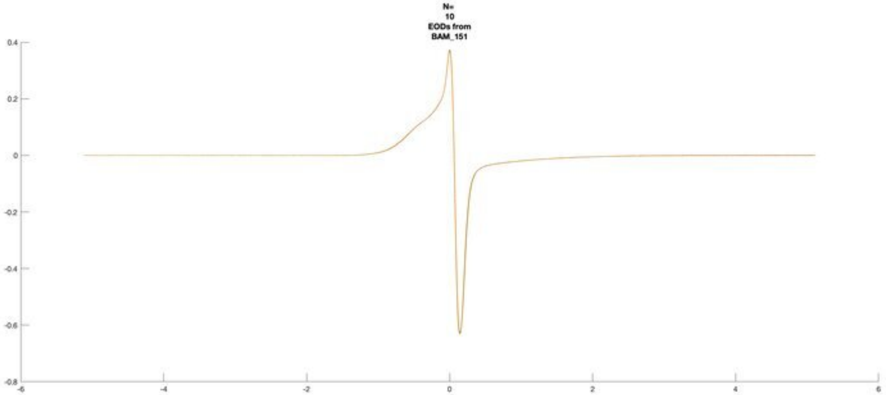
N=
10
EODs from
BAM_151

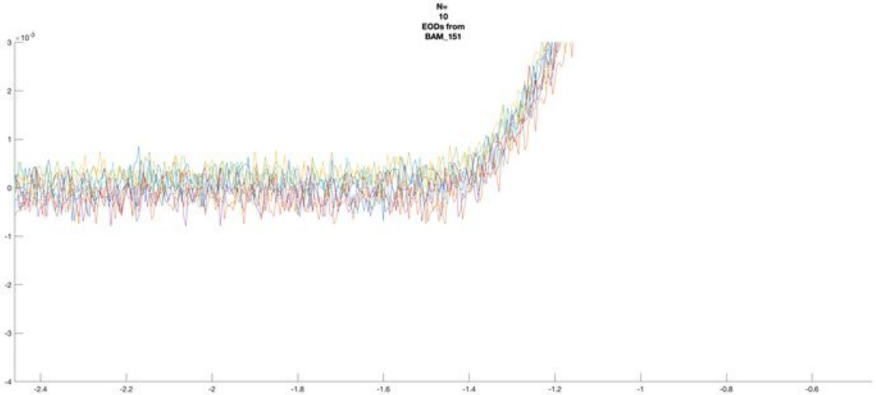
N=
10
EODs from
BAM_151

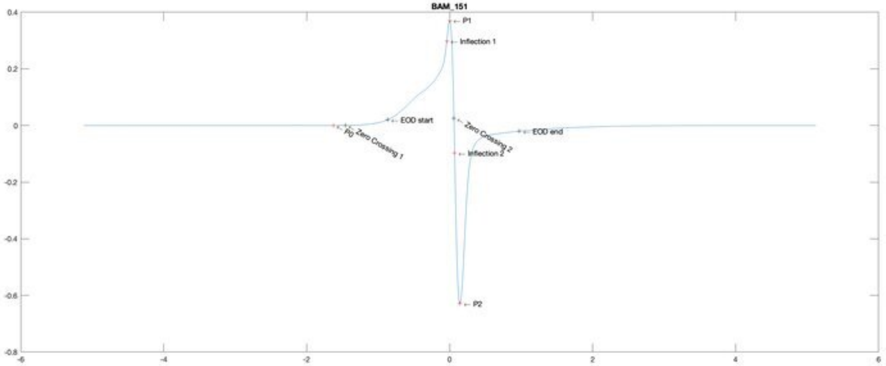
BAM_151
P1
Inflection 1
EOD start
P0
Zero Crossing 1
Zero Crossing 2
EOD end
Inflection 2
P2

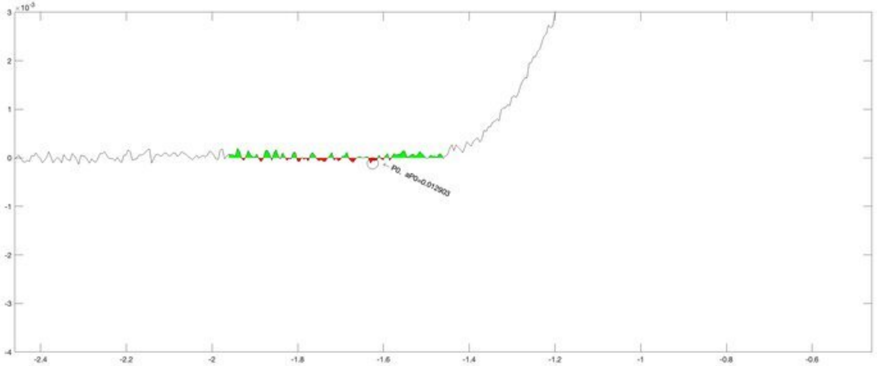
P0, aP0=0.012923

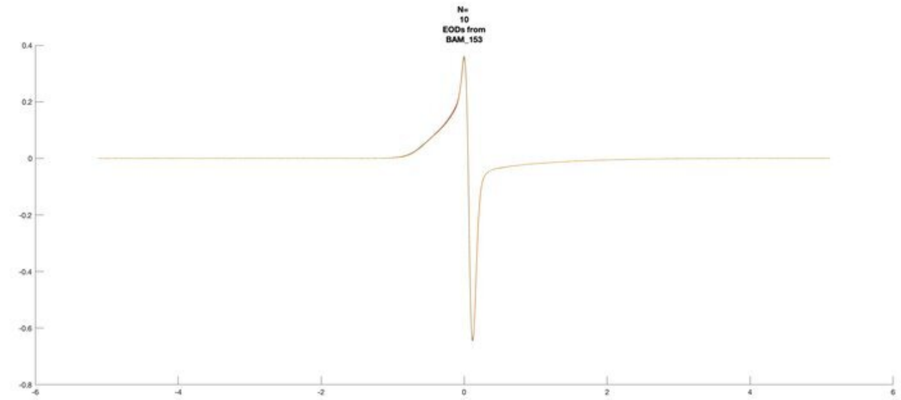
N=
10
EODs from
BAM_153

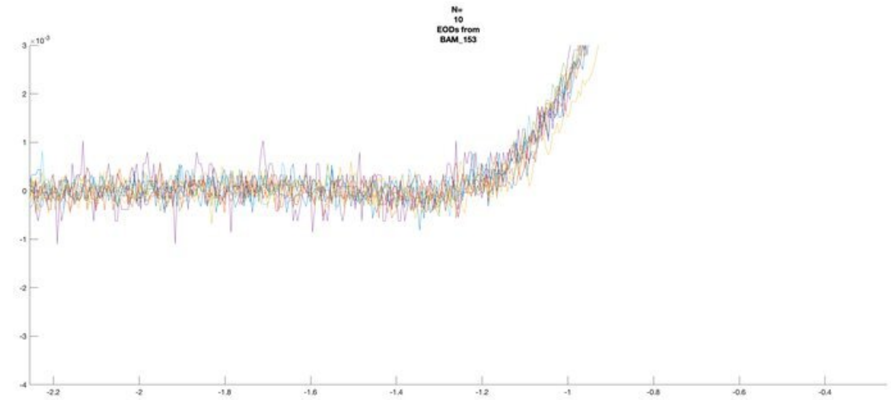
N=
10
EODs from
BAM_153

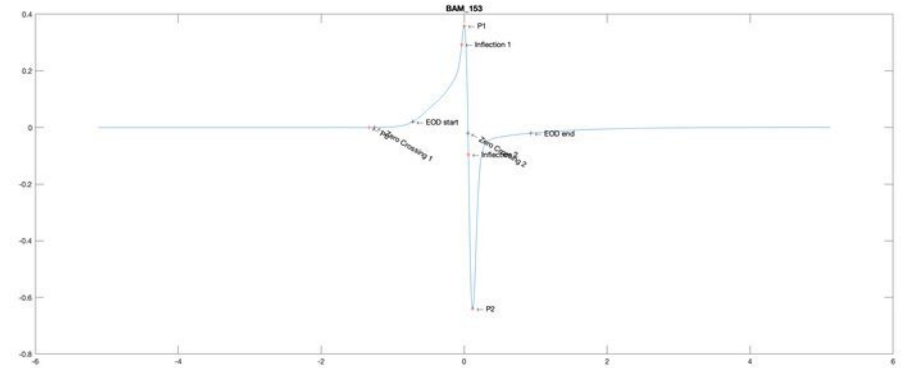
BAM_153
P1
Inflection 1
EOD start
Zero Crossing 1
Zero Crossing 2
Inflection 2
EOD end
P2

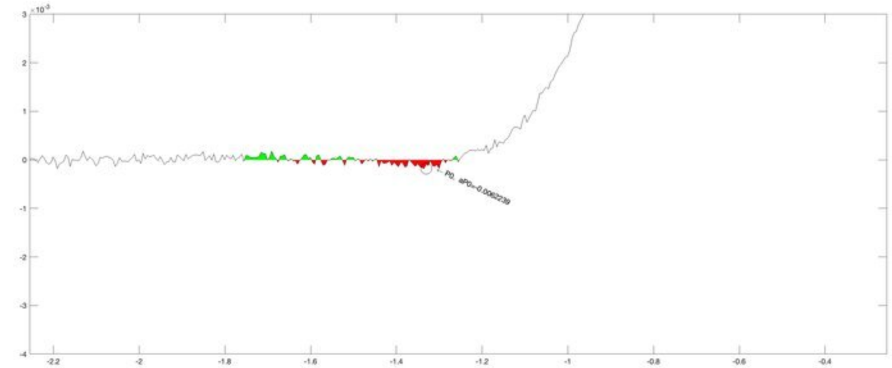
P0, aP0=-0.0062239

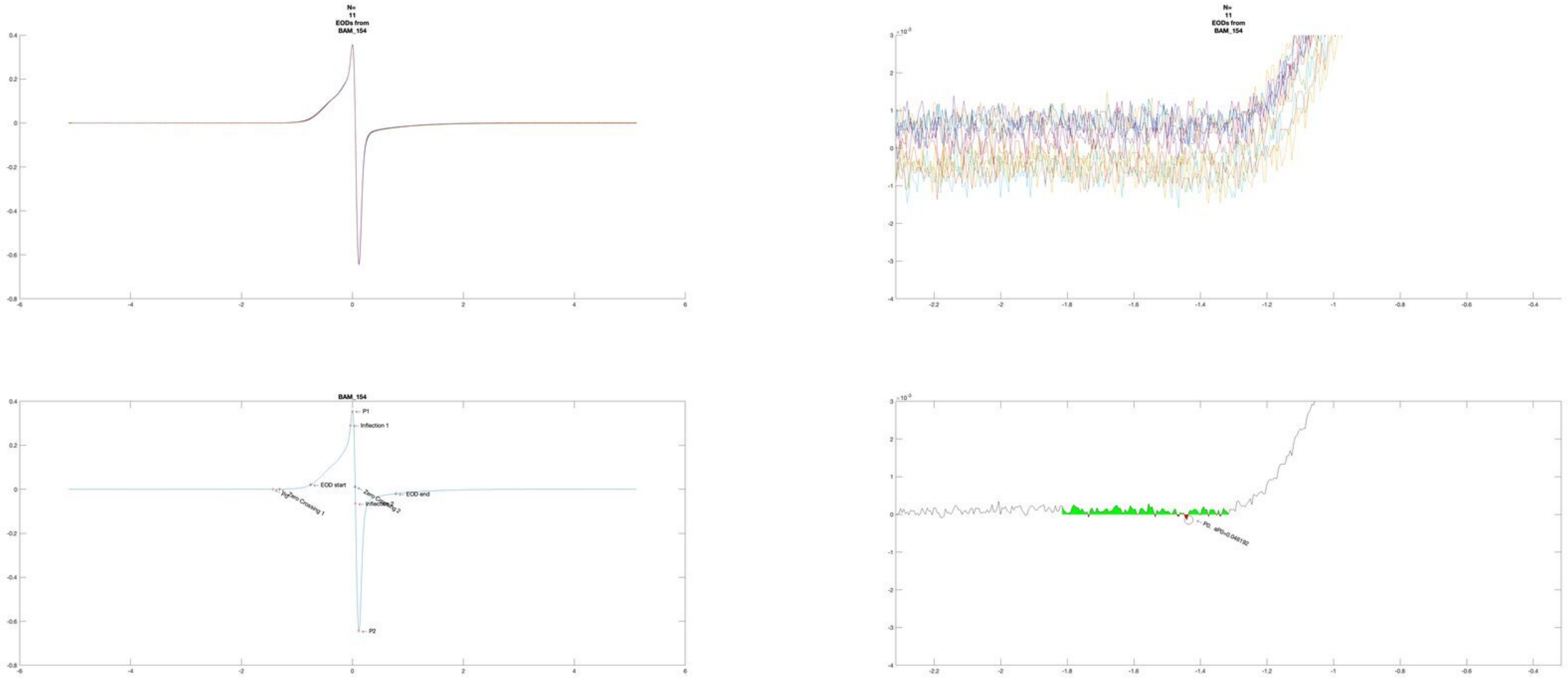

N=
11
EODs from
BAM_154
N=
11
EODs from
BAM_154
BAM_154
P1
Inflection 1
EOD start
EOD end
Zero Crossing 1
Zero Crossing 2
Inflection 2
P2
P0, aP0=0.046192

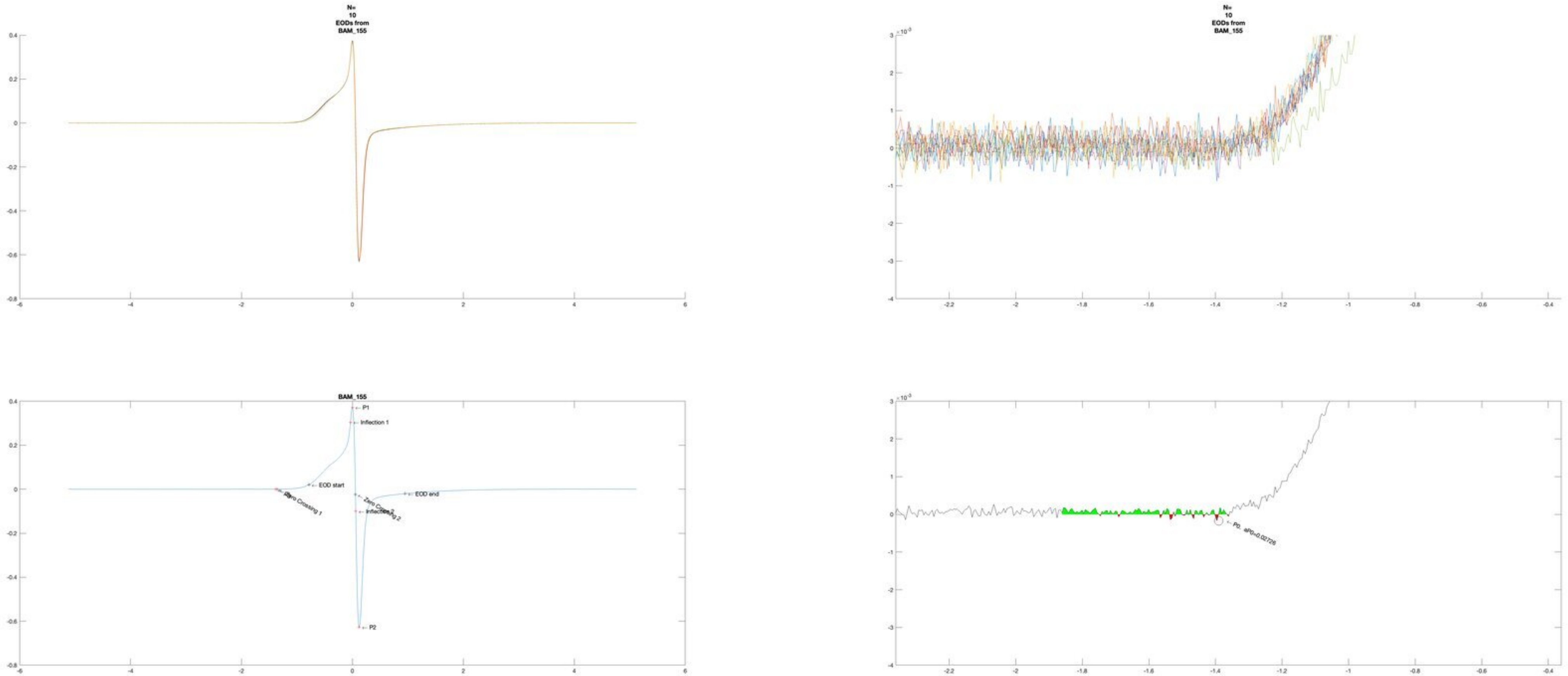
N=
10
EODs from
BAM_155
N=
10
EODs from
BAM_155
BAM_155
P1
Inflection 1
EOD start
Zero Crossing 1
Zero Crossing 2
Inflection 2
EOD end
P2
P0, aP0=0.02726

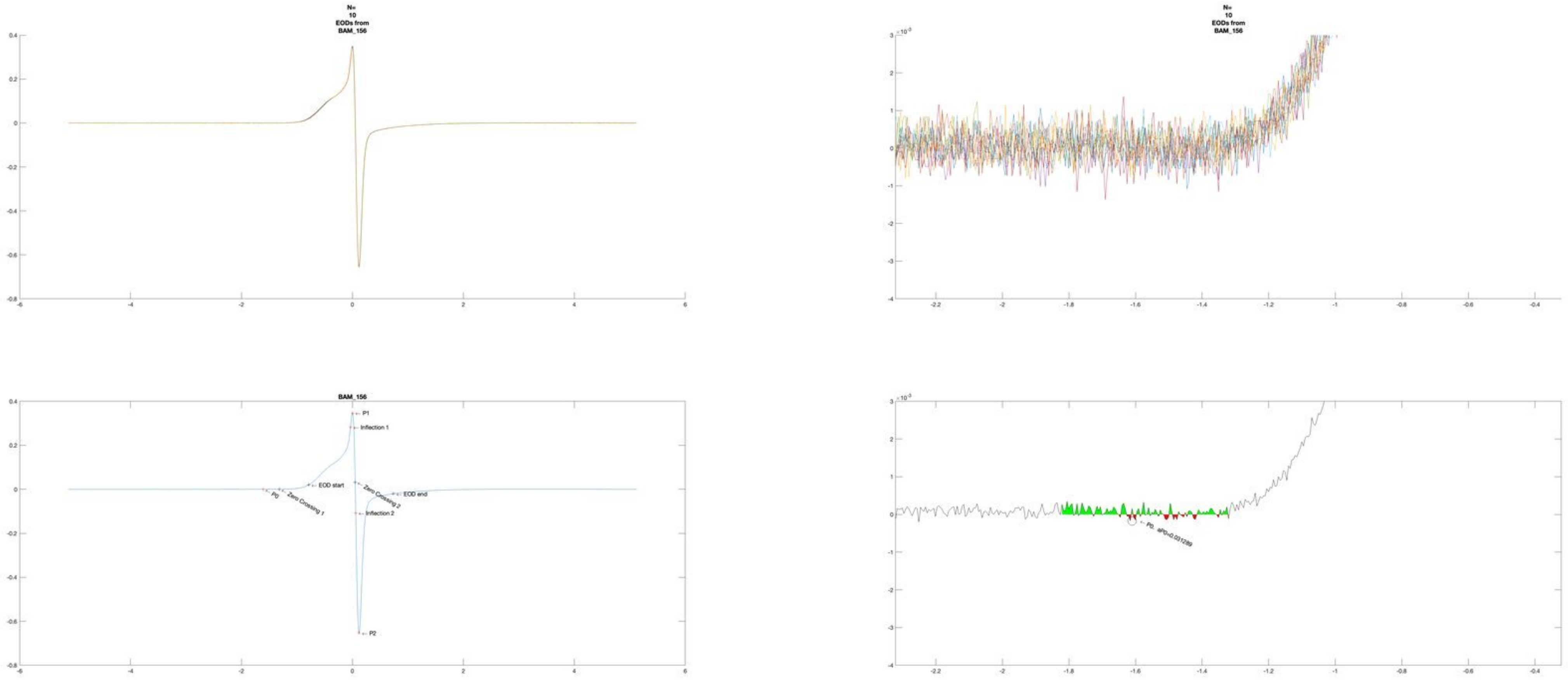
N=
10
EODs from
BAM_156
N=
10
EODs from
BAM_156
BAM_156
← P1
← Inflection 1
← EOD start
← P0
← Zero Crossing 1
← Zero Crossing 2
← EOD end
← Inflection 2
← P2
← P0, aP0=0.031289

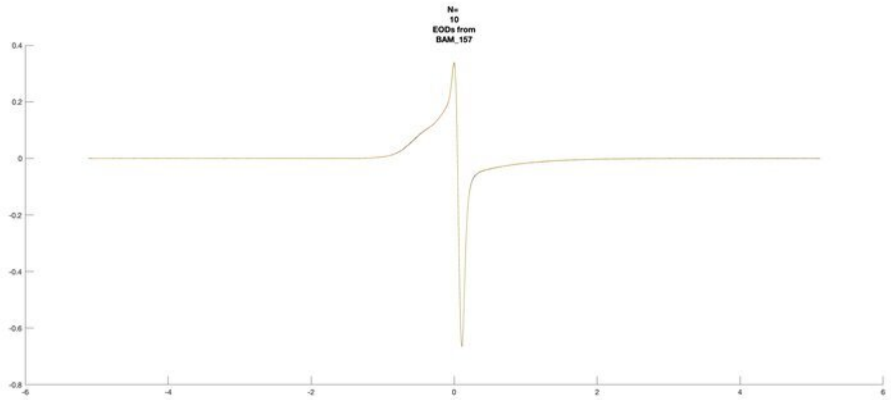
N=
10
EODs from
BAM_157

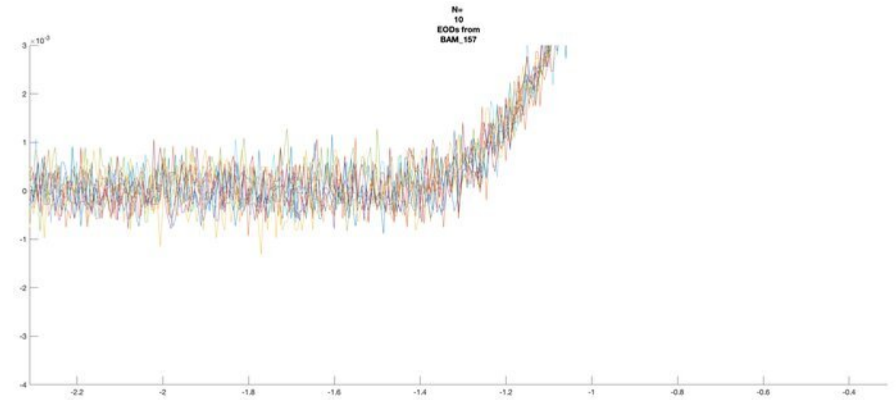
N=
10
EODs from
BAM_157

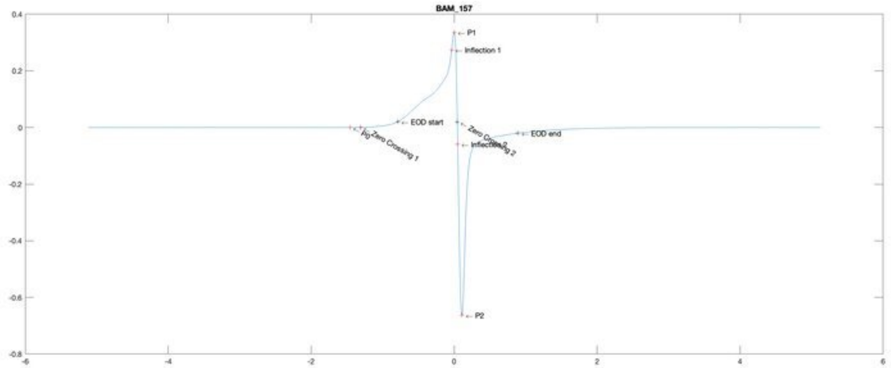
BAM_157
P1
Inflection 1
EOD start
Zero Crossing 1
EOD end
Zero Crossing 2
Inflection 2
P2

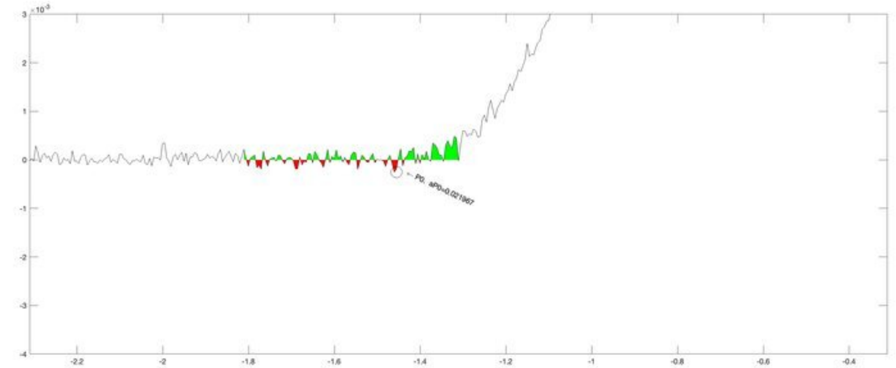
P0, aP0=0.021967

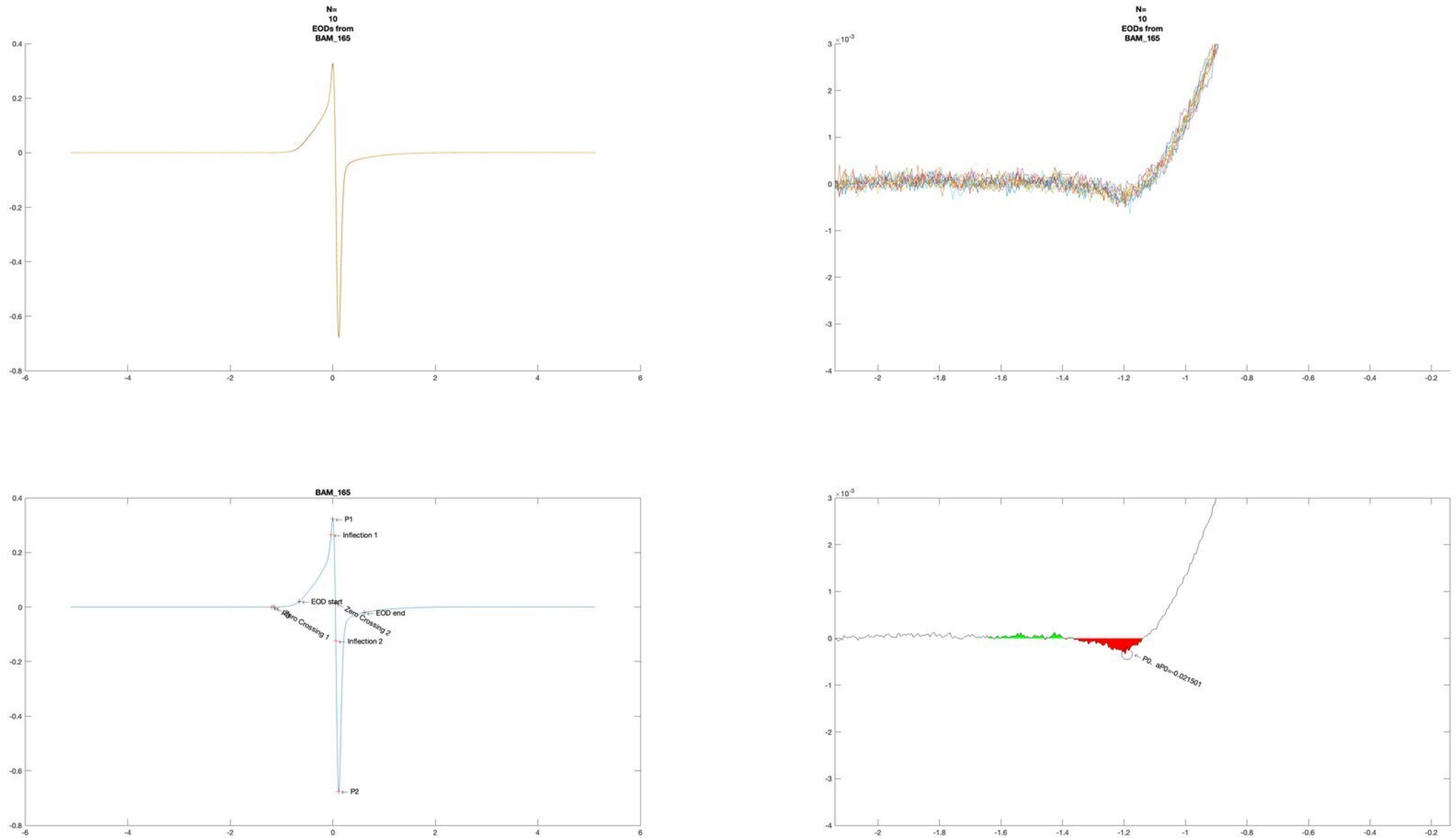
N=
10
EODs from
BAM_165
N=
10
EODs from
BAM_165
BAM_165
P1
Inflection 1
EOD start
Zero Crossing 2
EOD end
P0
Zero Crossing 1
Inflection 2
P2
P0. aP0=-0.021501

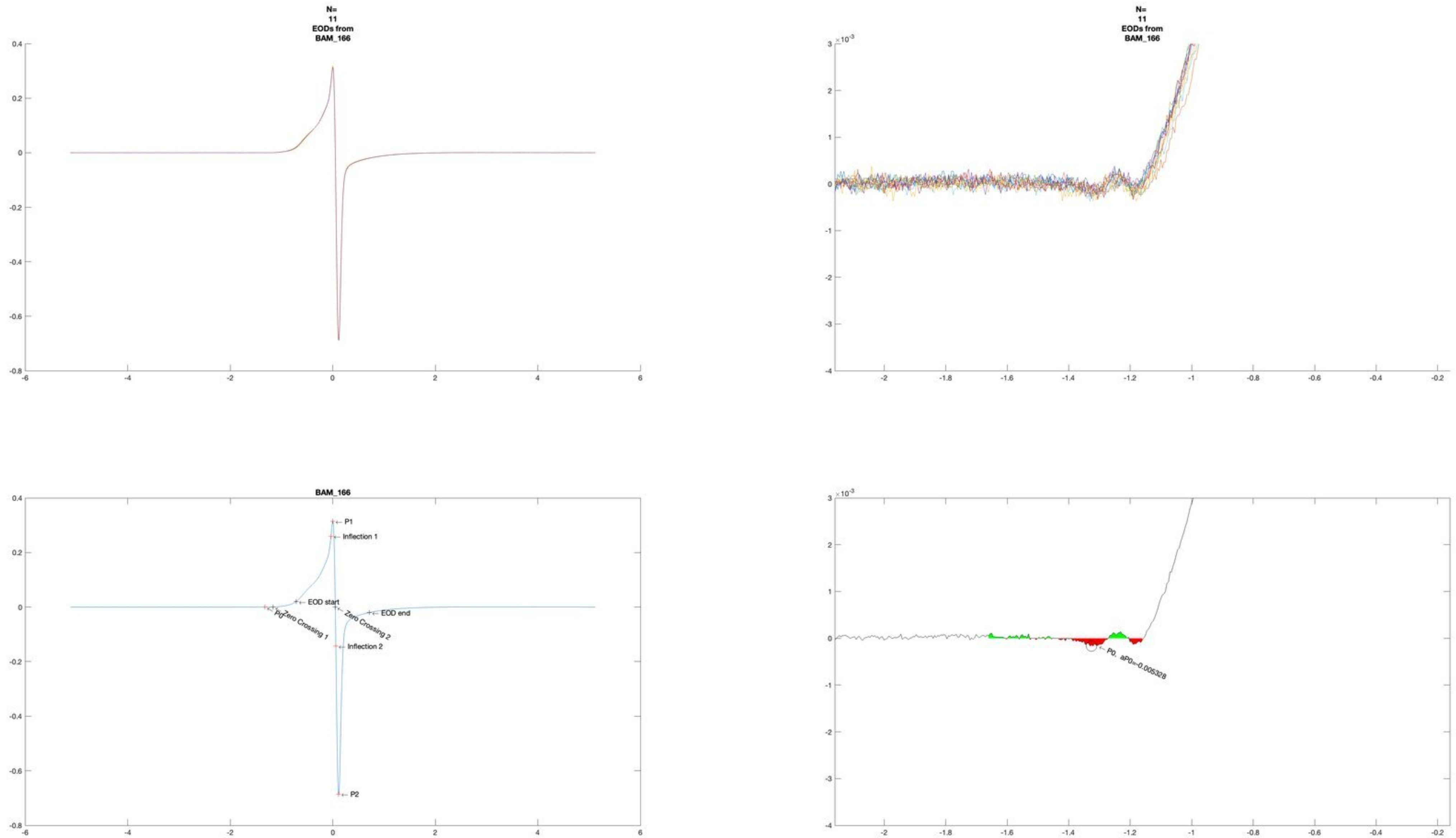
N=
11
EODs from
BAM_166
N=
11
EODs from
BAM_166
×10^-3
BAM_166
P1
Inflection 1
EOD start
P0
Zero Crossing 1
Zero Crossing 2
EOD end
Inflection 2
P2
×10^-3
P0, aP0=-0.005328

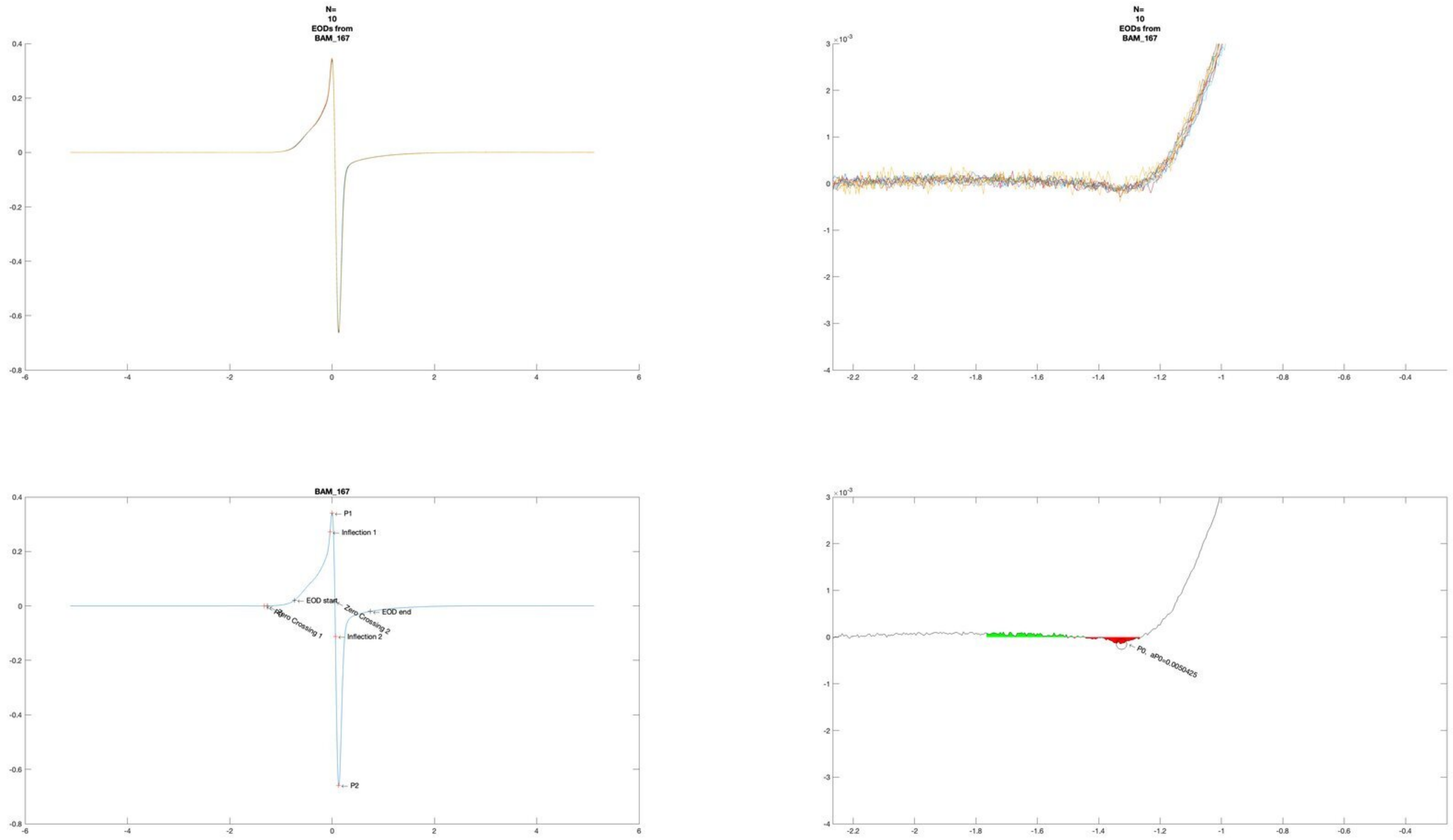
N=
10
EODs from
BAM_167
N=
10
EODs from
BAM_167
BAM_167
P1
Inflection 1
EOD start
Zero Crossing 2
EOD end
P0
Zero Crossing 1
Inflection 2
P2
P0, aP0=0.0050425

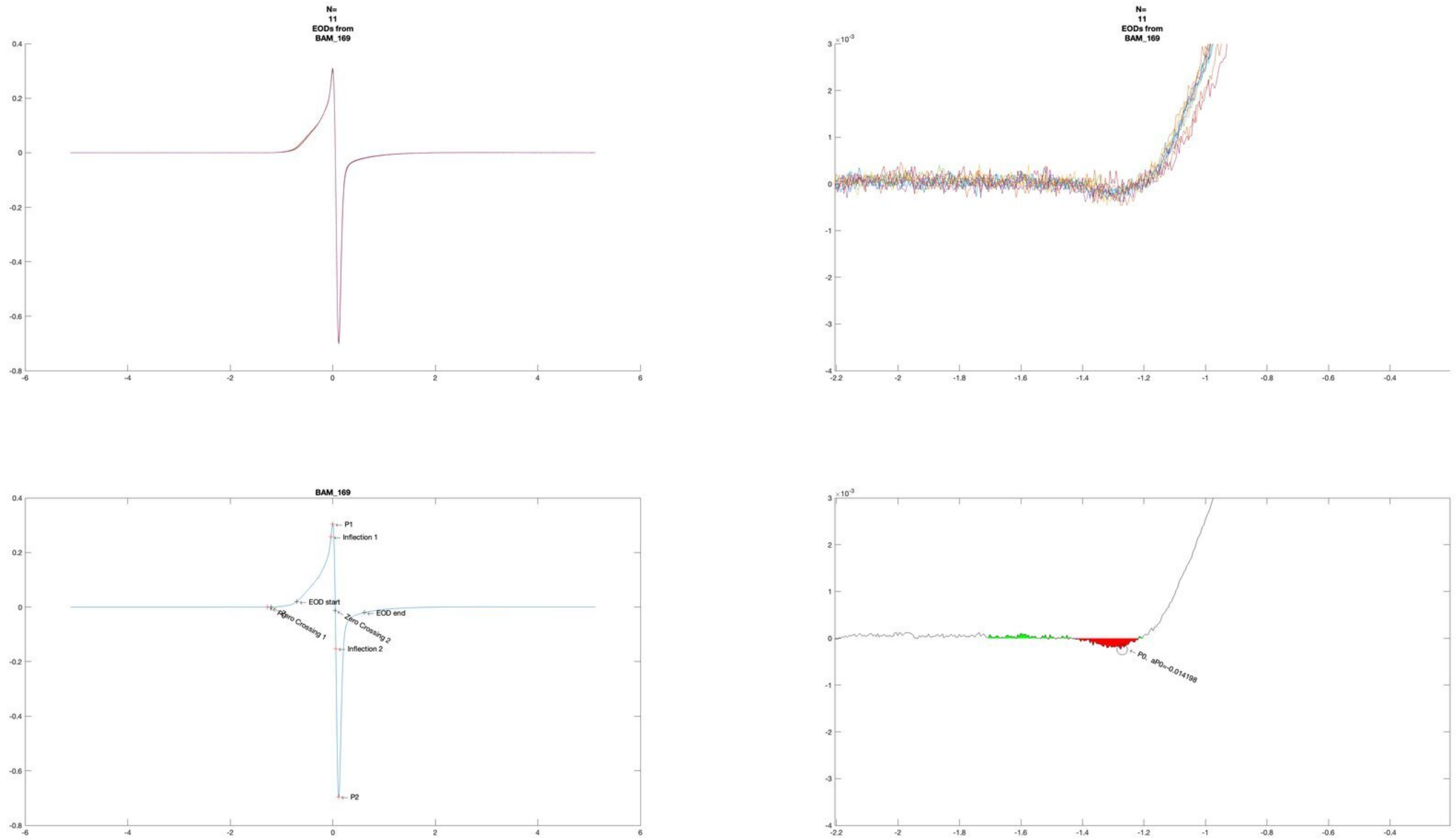
N=
11
EODs from
BAM_169
N=
11
EODs from
BAM_169
BAM_169
P1
Inflection 1
EOD start
P0
Zero Crossing 1
Zero Crossing 2
EOD end
Inflection 2
P2
P0, aP0=-0.014198

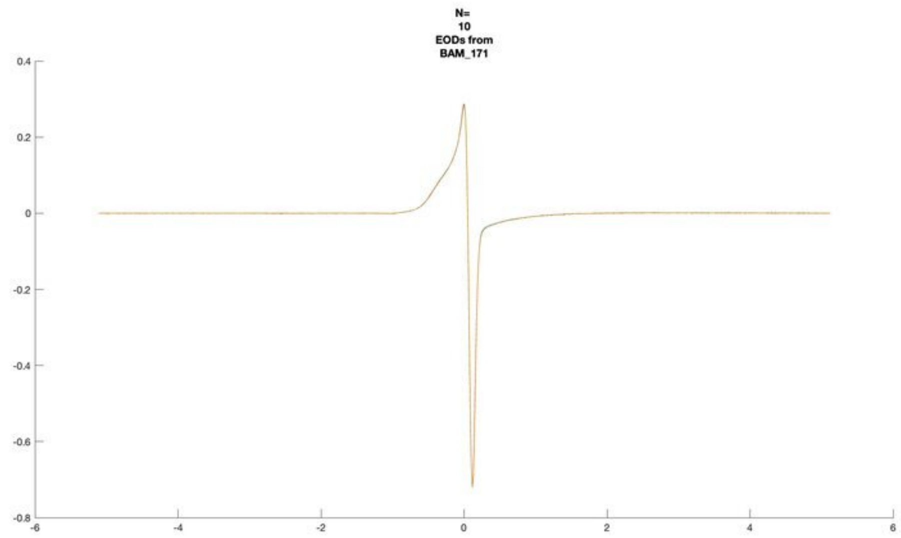
N=
10
EODs from
BAM_171

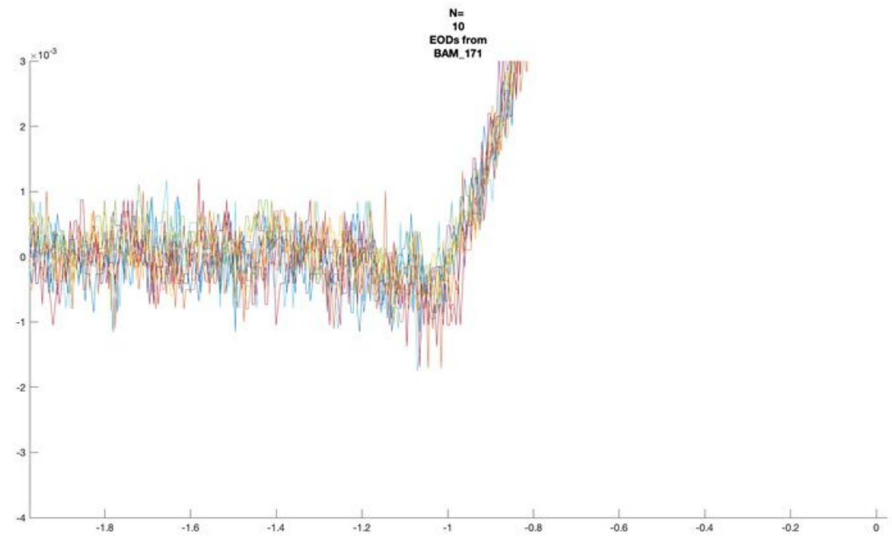
N=
10
EODs from
BAM_171
×10^-3

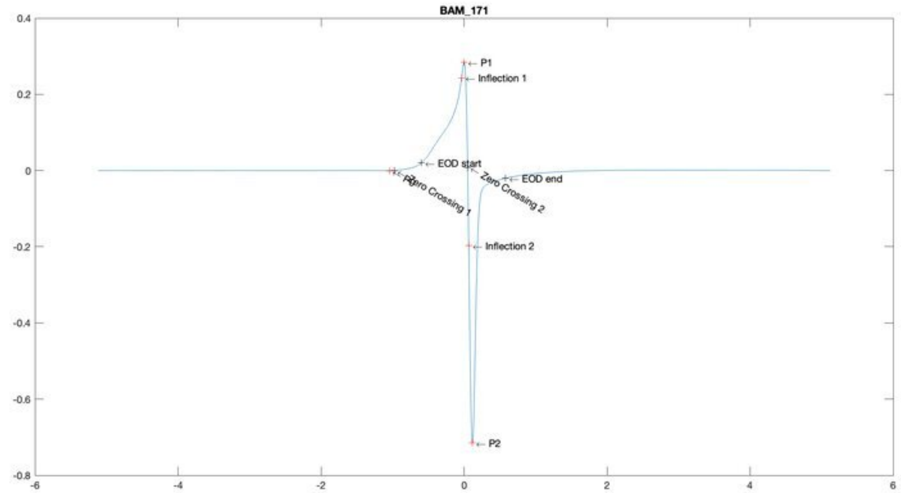
BAM_171
P1
Inflection 1
EOD start
Zero Crossing 1
Zero Crossing 2
EOD end
Inflection 2
P2

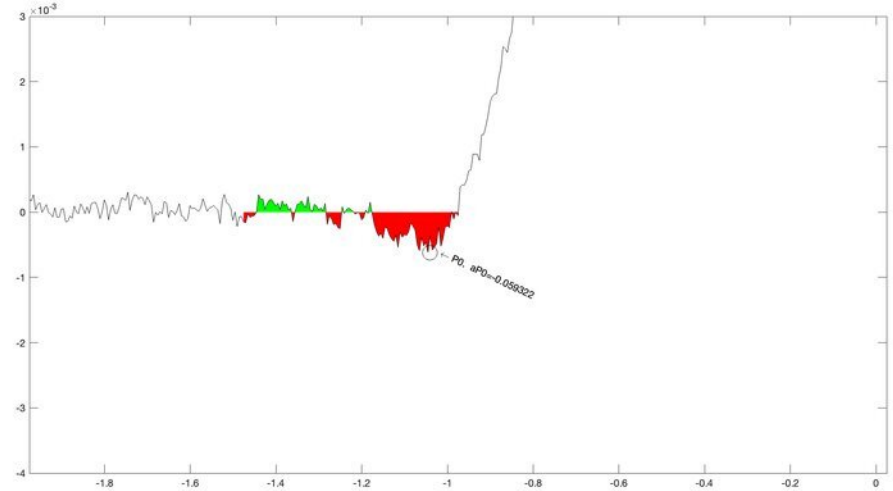
×10^-3
P0, aP0=-0.059322

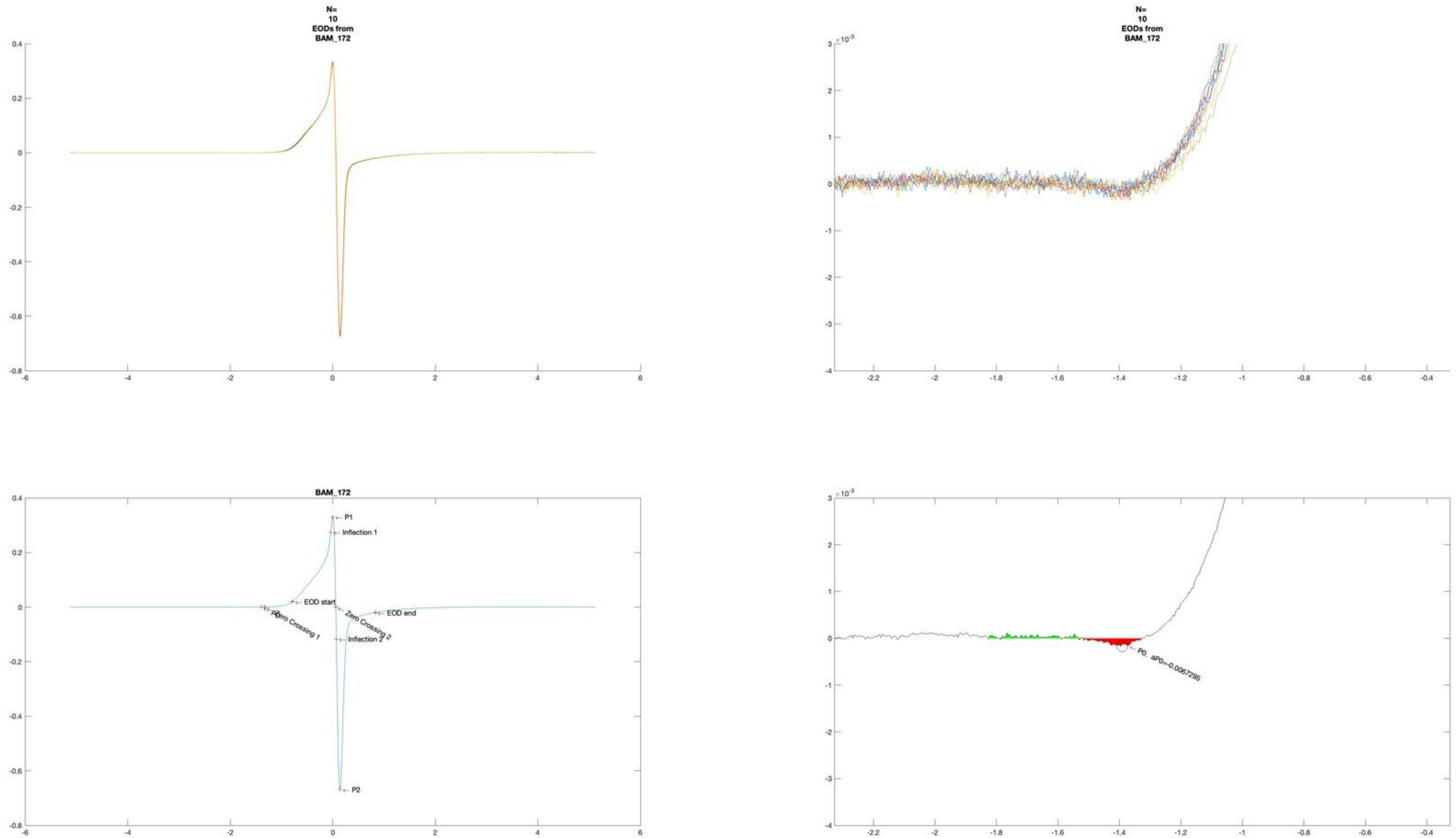
N=
10
EODs from
BAM_172
N=
10
EODs from
BAM_172
BAM_172
P1
Inflection 1
EOD start
Zero Crossing 1
Zero Crossing 2
EOD end
Inflection 2
P2
P0, aP0=-0.0067295

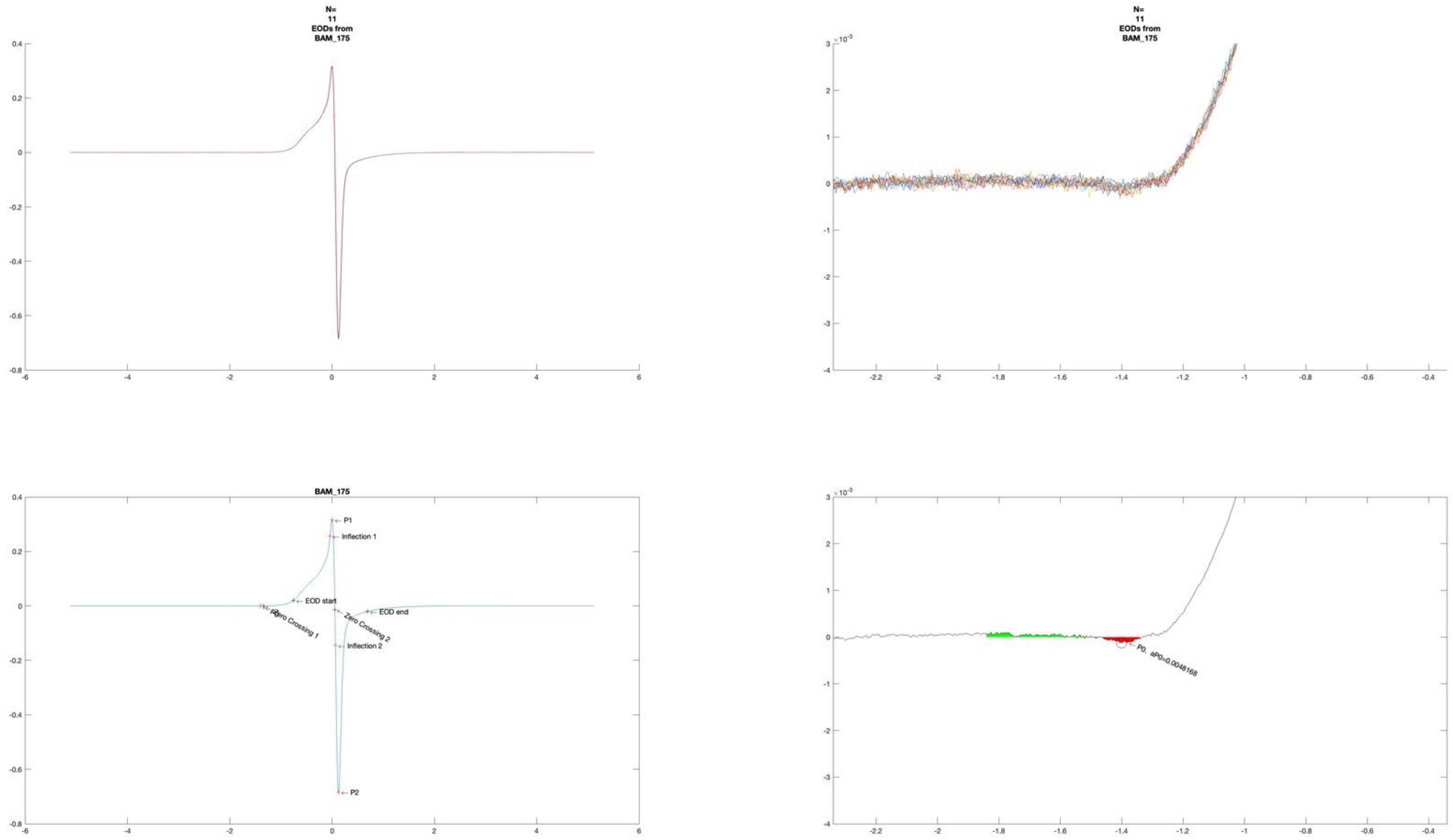
N=
11
EODs from
BAM_175
N=
11
EODs from
BAM_175
BAM_175
P1
Inflection 1
EOD start
EOD end
Zero Crossing 1
Zero Crossing 2
Inflection 2
P2
P0, aP0=0.0048168

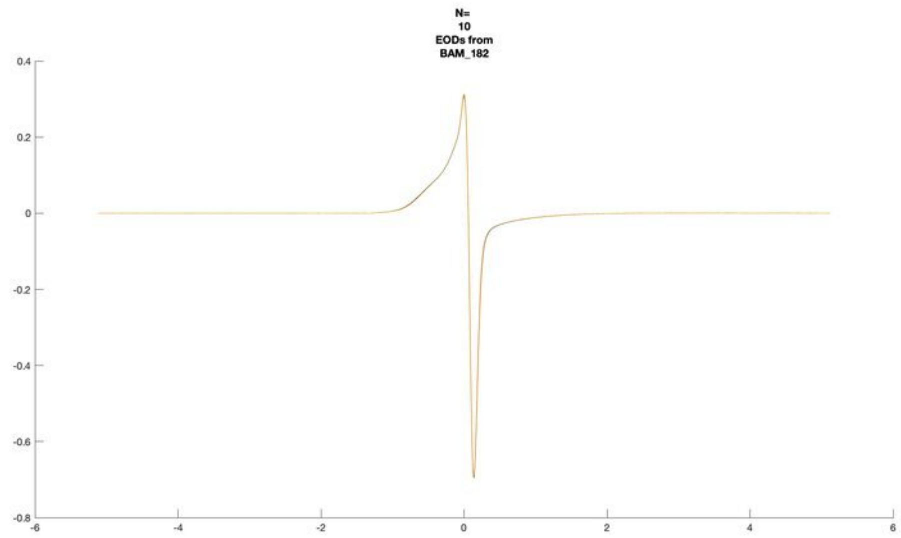
N=
10
EODs from
BAM_182

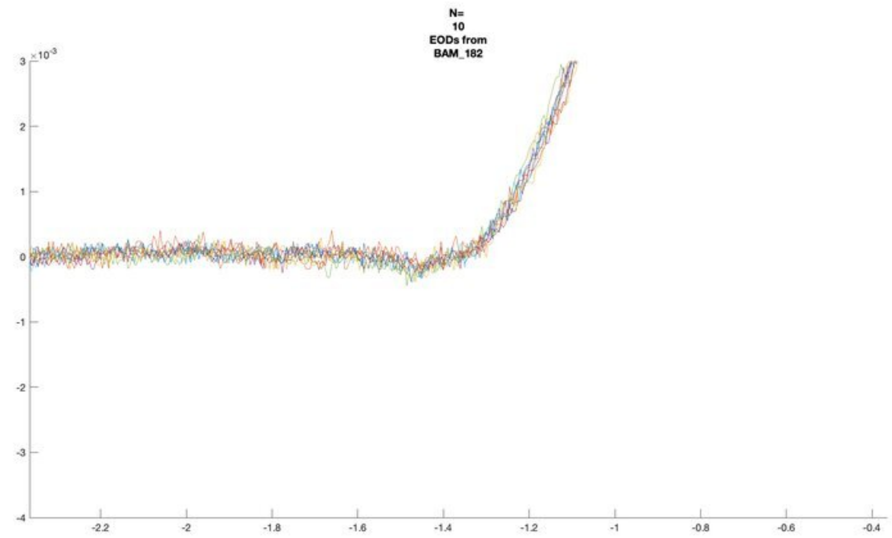
N=
10
EODs from
BAM_182
×10⁻³

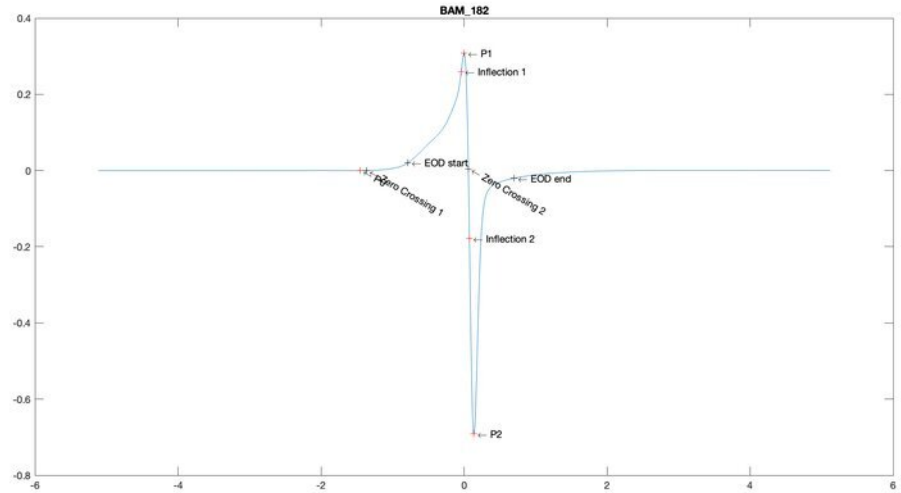
BAM_182
← P1
← Inflection 1
← EOD start
← P0
← Zero Crossing 1
← Zero Crossing 2
← EOD end
← Inflection 2
← P2

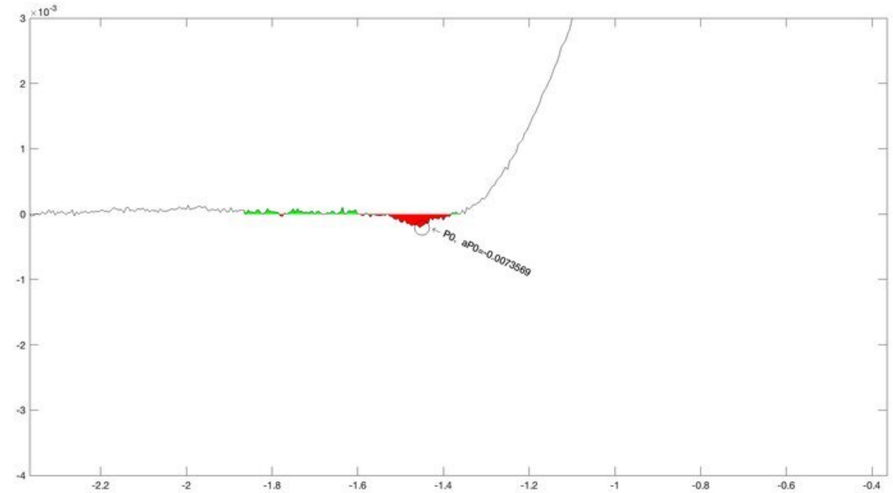
×10⁻³
← P0, aP0=-0.0073569

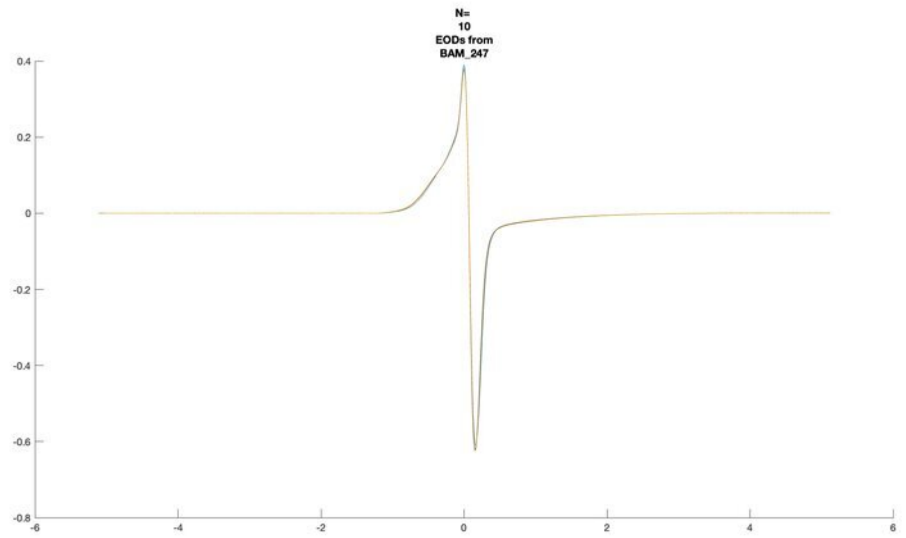
N=
10
EODs from
BAM_247

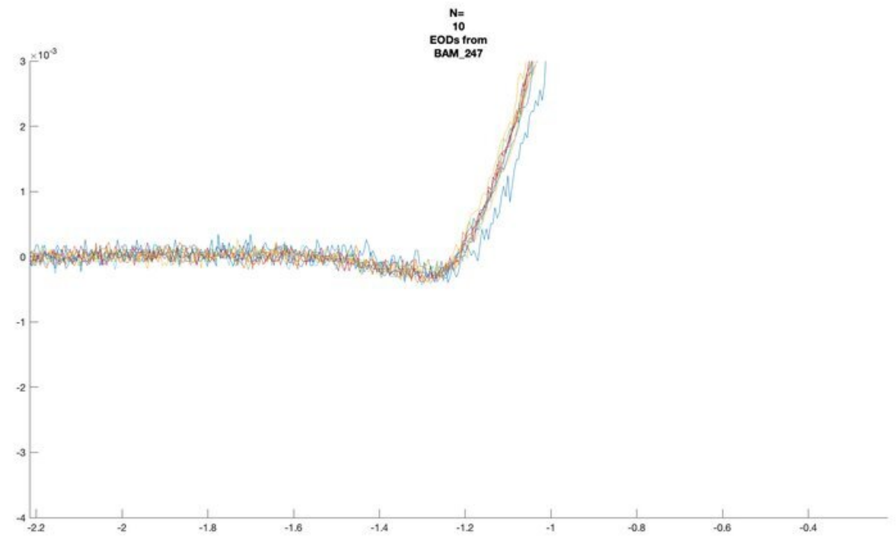
N=
10
EODs from
BAM_247

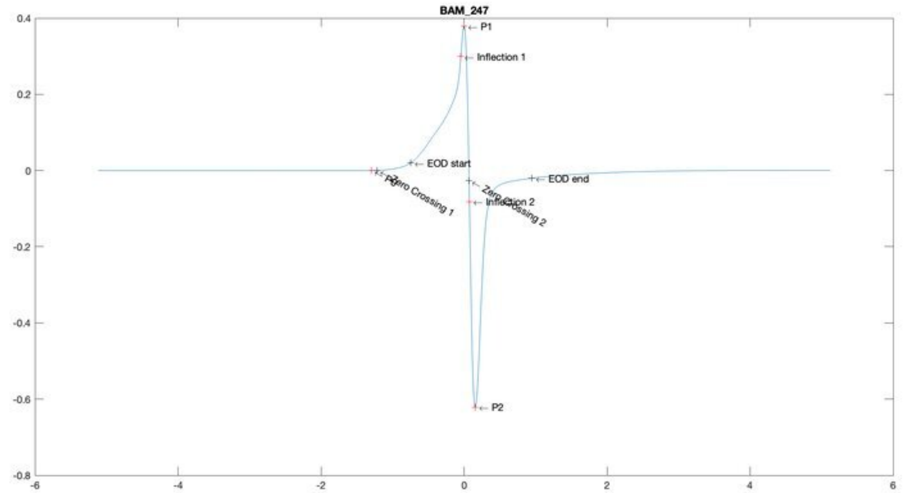
BAM_247
P1
Inflection 1
EOD start
EOD end
Zero Crossing 1
Zero Crossing 2
Inflection 2
P2

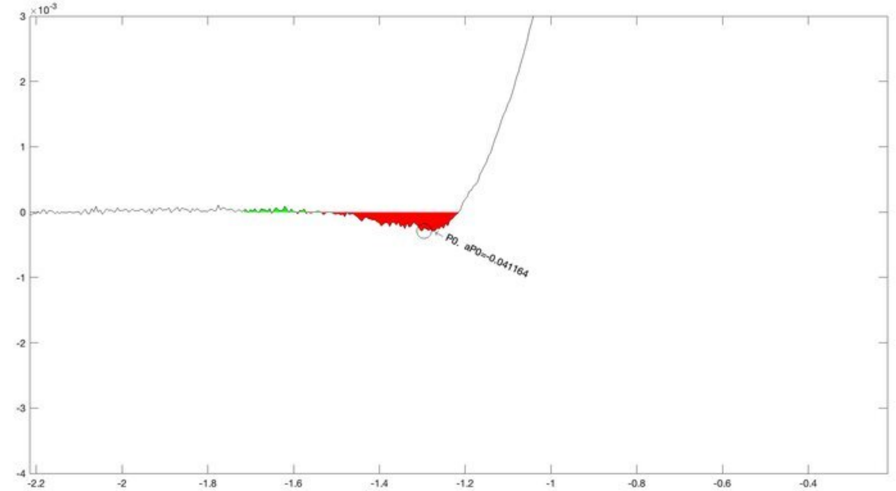
P0. aP0=-0.041164

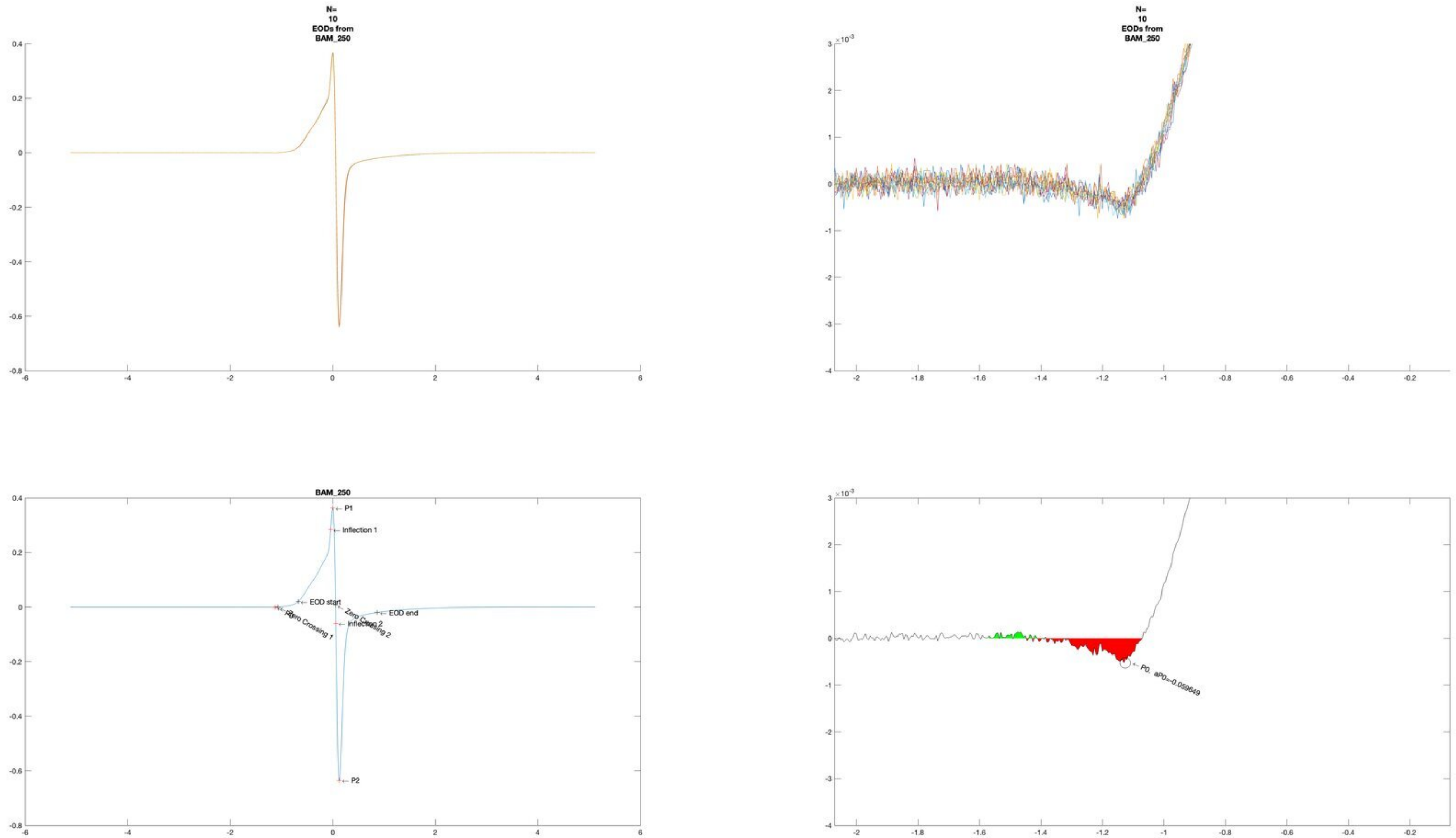
N=
10
EODs from
BAM_250
N=
10
EODs from
BAM_250
BAM_250
P1
Inflection 1
EOD start
EOD end
Zero Crossing 1
Zero Crossing 2
Inflection 2
P2
P0. aP0=-0.059649

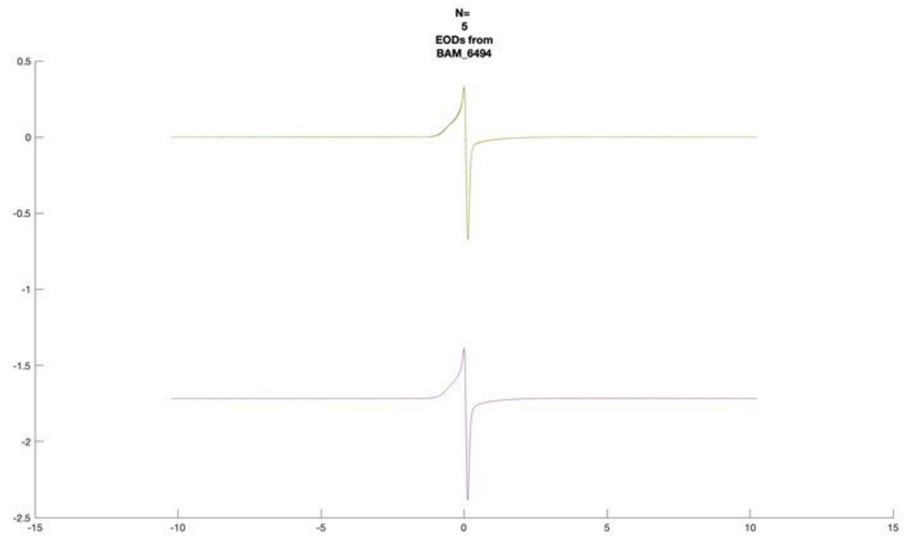
N=
5
EODs from
BAM_6494

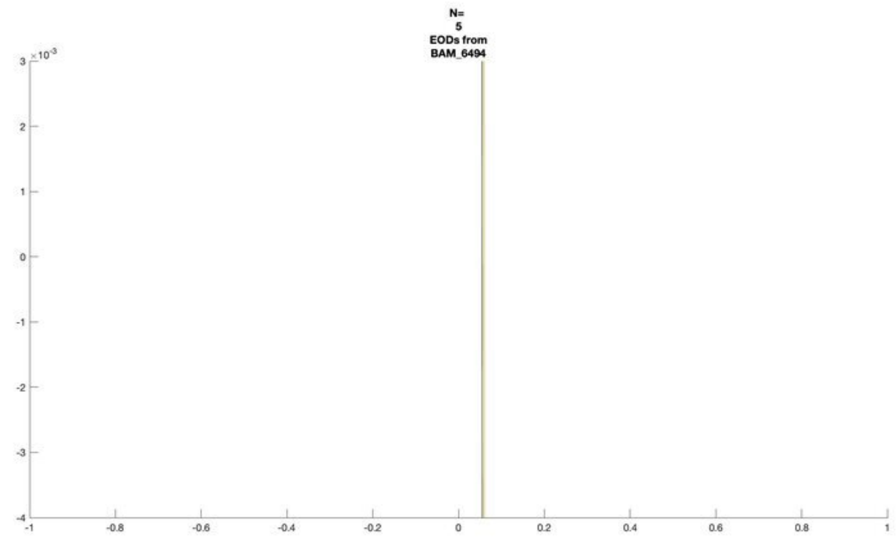
N=
5
EODs from
BAM_6494

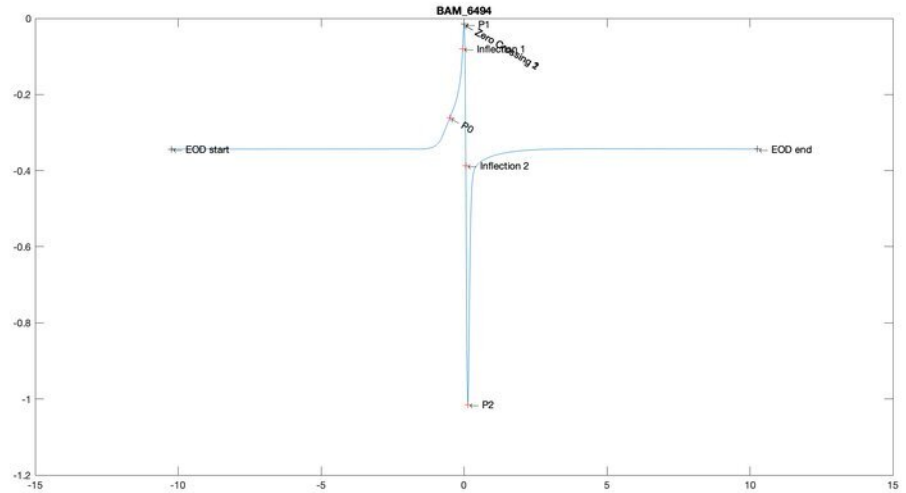
BAM_6494
P1
Zero Crossing 1
Inflection 1
Zero Crossing 2
P0
EOD start
EOD end
Inflection 2
P2

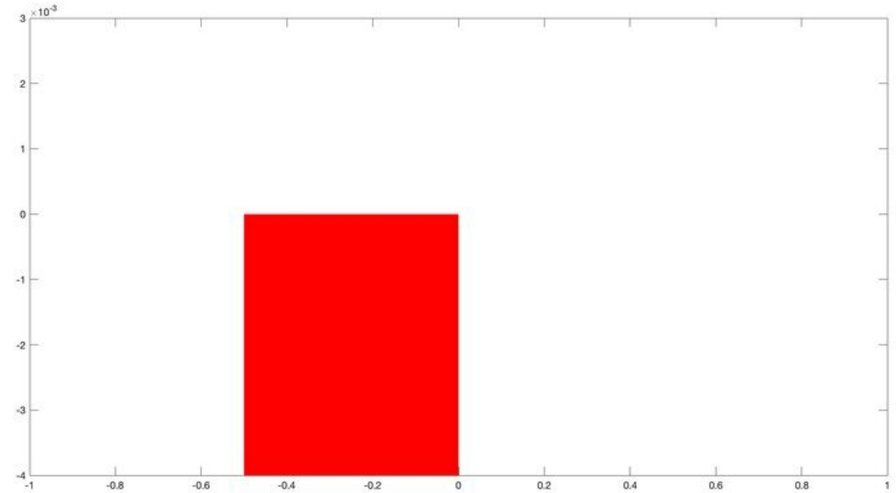

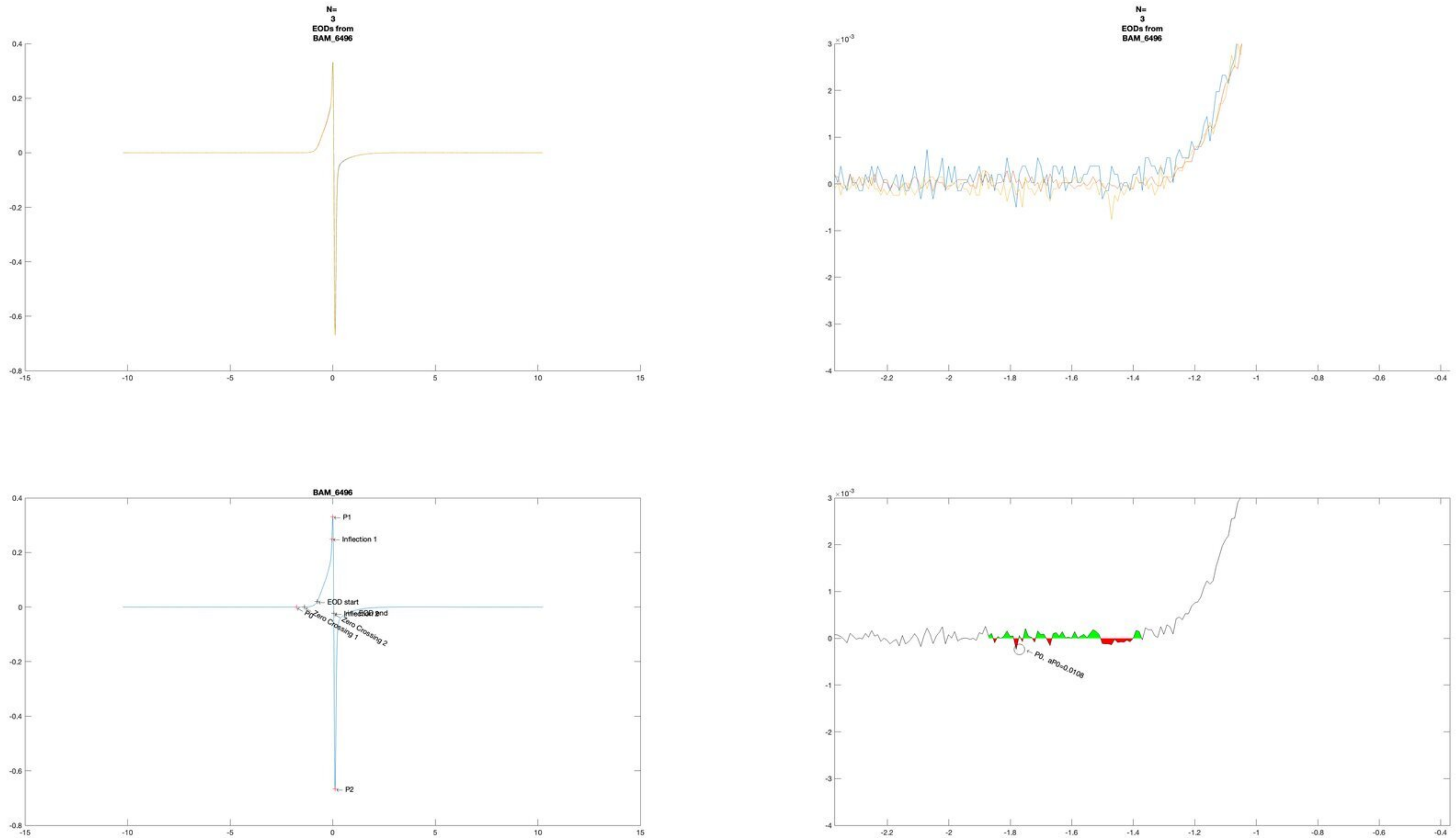
N=
3
EODs from
BAM_6496
N=
3
EODs from
BAM_6496
BAM_6496
P1
Inflection 1
EOD start
EOD end
Inflection 2
P0
Zero Crossing 1
Zero Crossing 2
P2
P0, aP0=0.0108

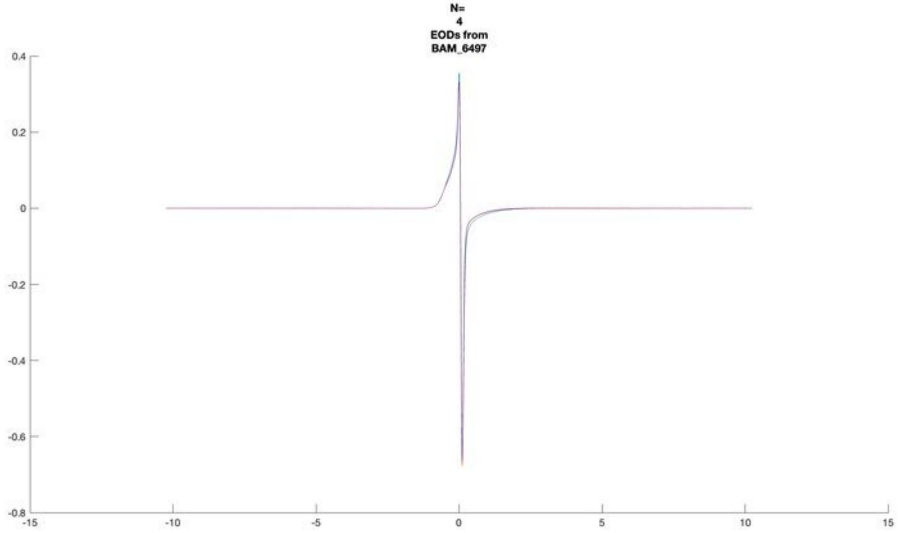
N=
4
EODs from
BAM_6497
0.4
0.2
0
-0.2
-0.4
-0.6
-0.8
-15
-10
-5
0
5
10
15

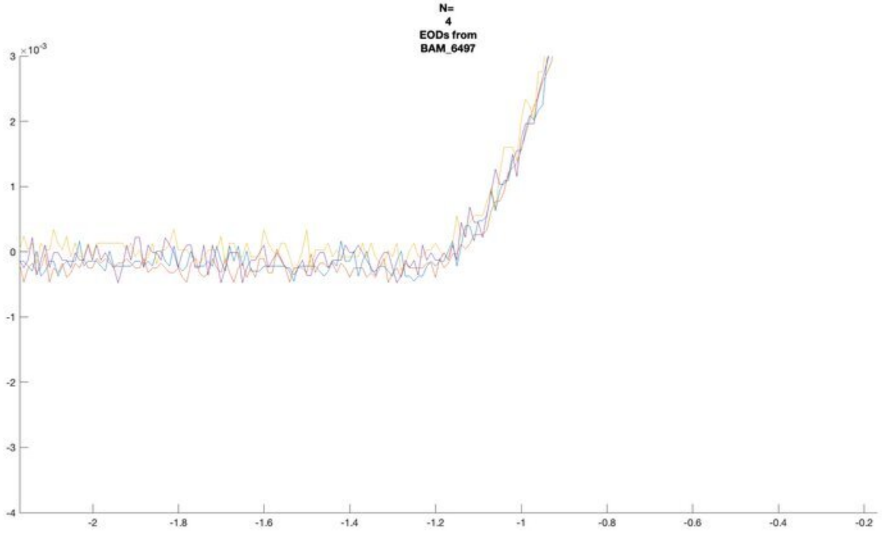
N=
4
EODs from
BAM_6497
×10^-3
3
2
1
0
-1
-2
-3
-4
-2
-1.8
-1.6
-1.4
-1.2
-1
-0.8
-0.6
-0.4
-0.2

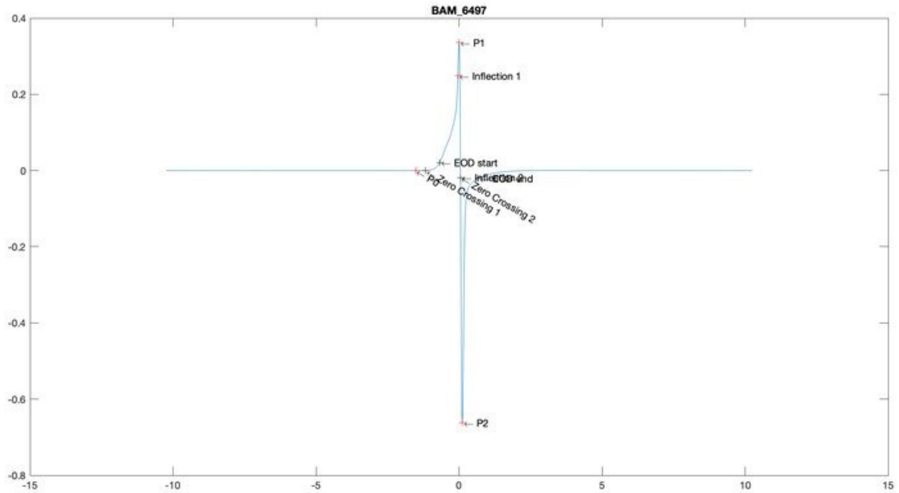
BAM_6497
P1
Inflection 1
EOD start
Inflection 2
EOD end
P0
Zero Crossing 1
Zero Crossing 2
P2
0.4
0.2
0
-0.2
-0.4
-0.6
-0.8
-15
-10
-5
0
5
10
15

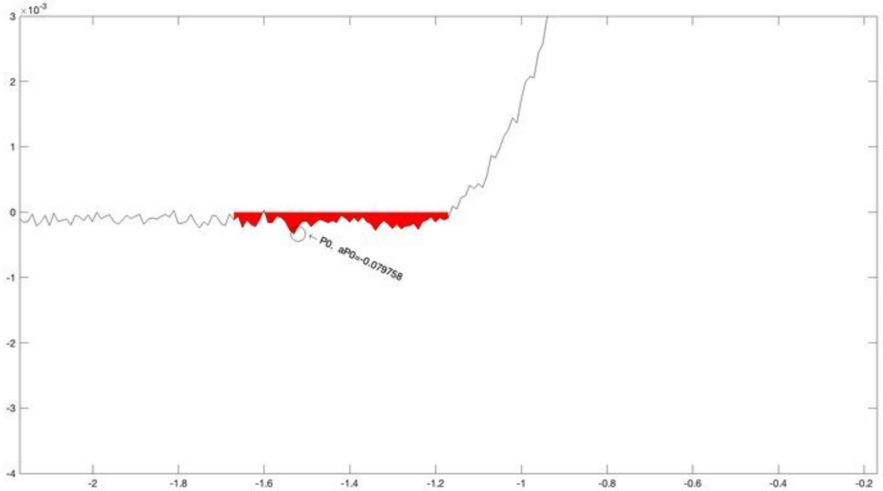
×10^-3
P0. aP0=-0.079758
3
2
1
0
-1
-2
-3
-4
-2
-1.8
-1.6
-1.4
-1.2
-1
-0.8
-0.6
-0.4
-0.2

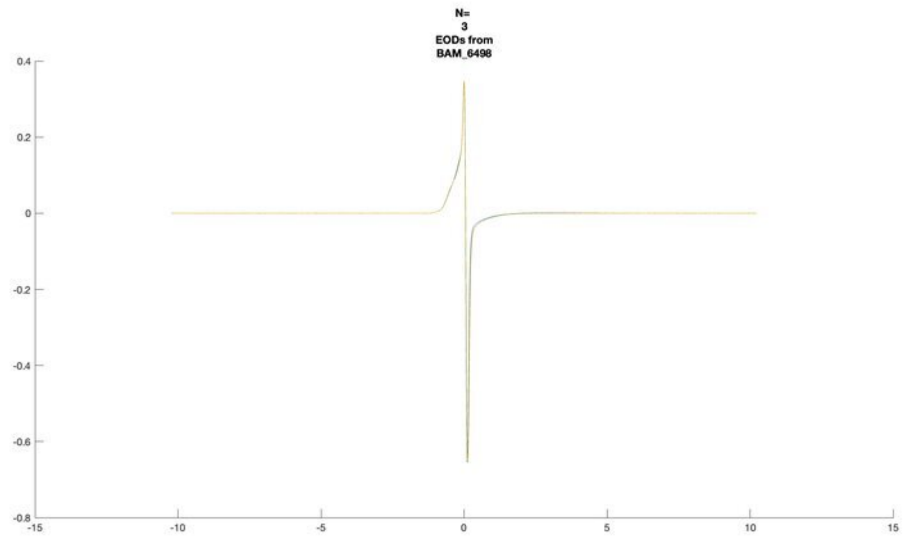
N=
3
EODs from
BAM_6498

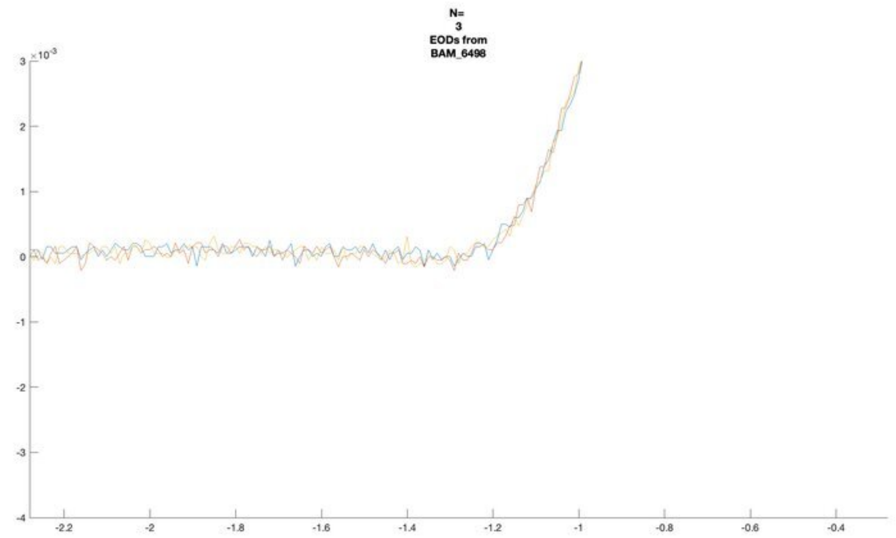
N=
3
EODs from
BAM_6498

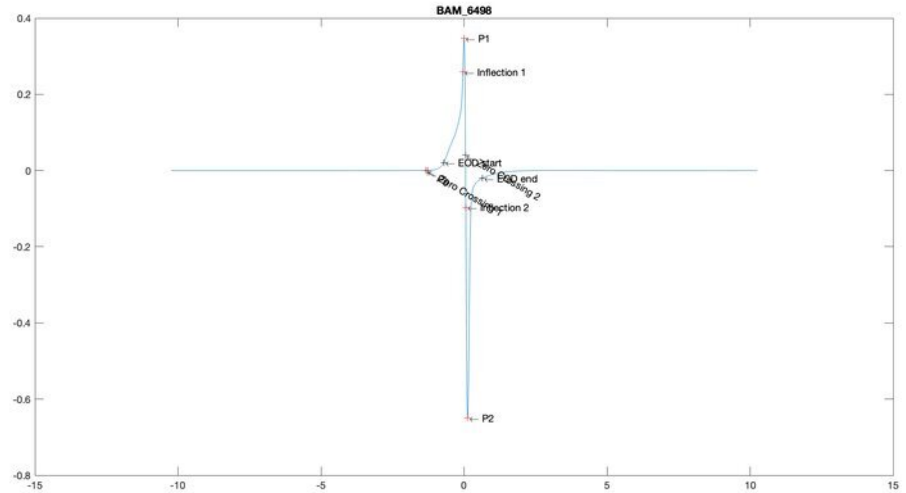
BAM_6498
P1
Inflection 1
EOD start
EOD end
Inflection 2
P2

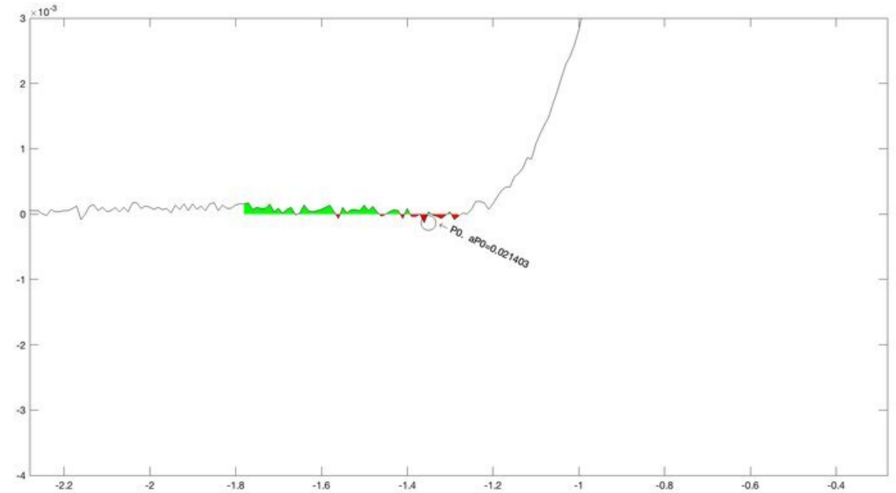
P0, aP0=0.021403

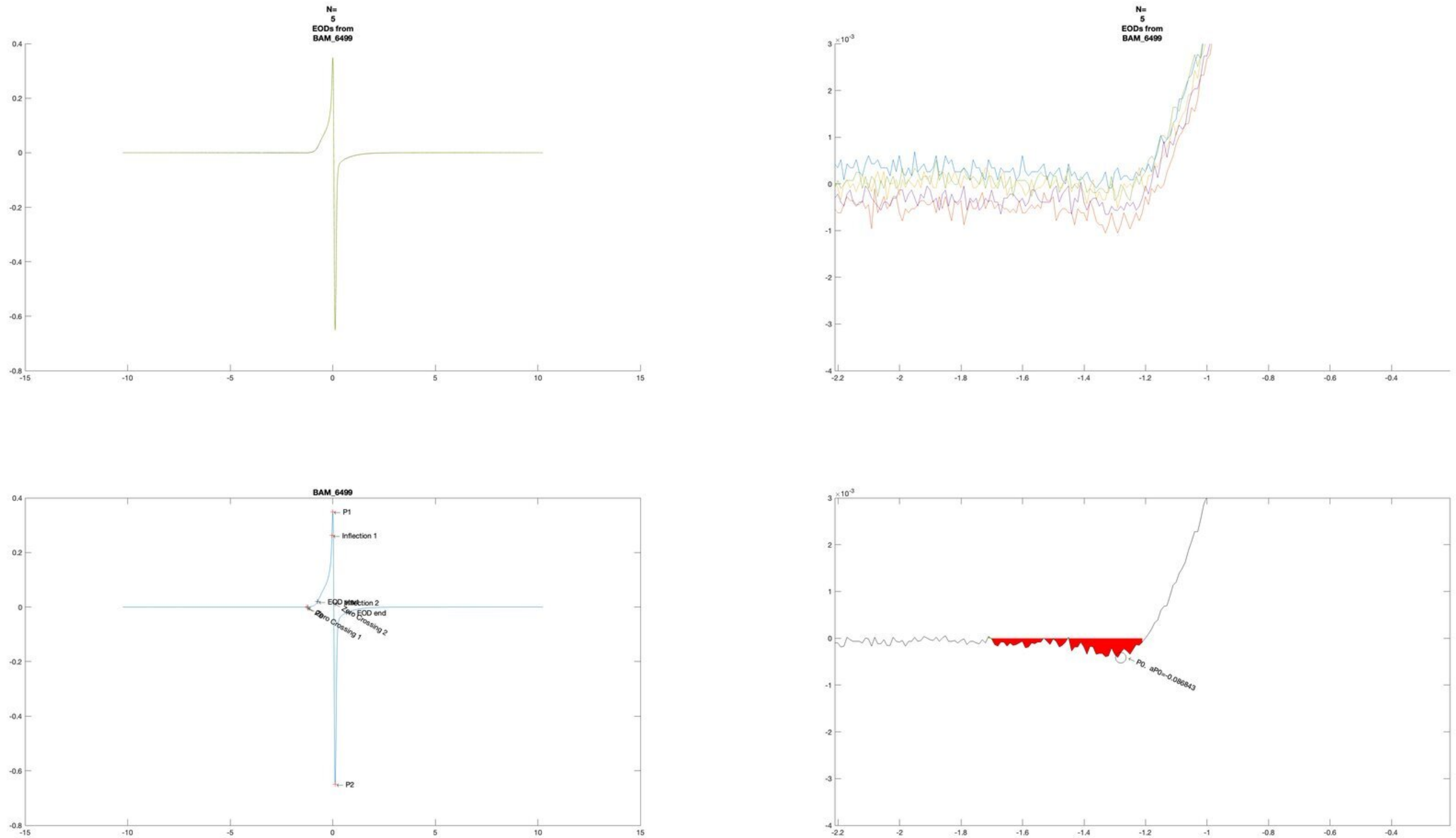
N=
5
EODs from
BAM_6499
N=
5
EODs from
BAM_6499
×10^-3
BAM_6499
P1
Inflection 1
EOD start
Inflection 2
EOD end
Zero Crossing 1
Zero Crossing 2
P2
×10^-3
P0, aP0=-0.086843

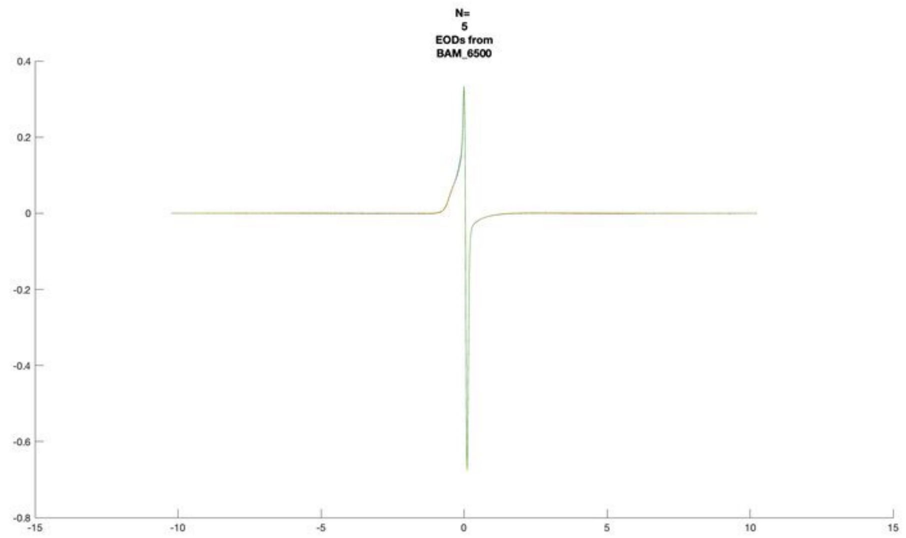
N=
5
EODs from
BAM_6500

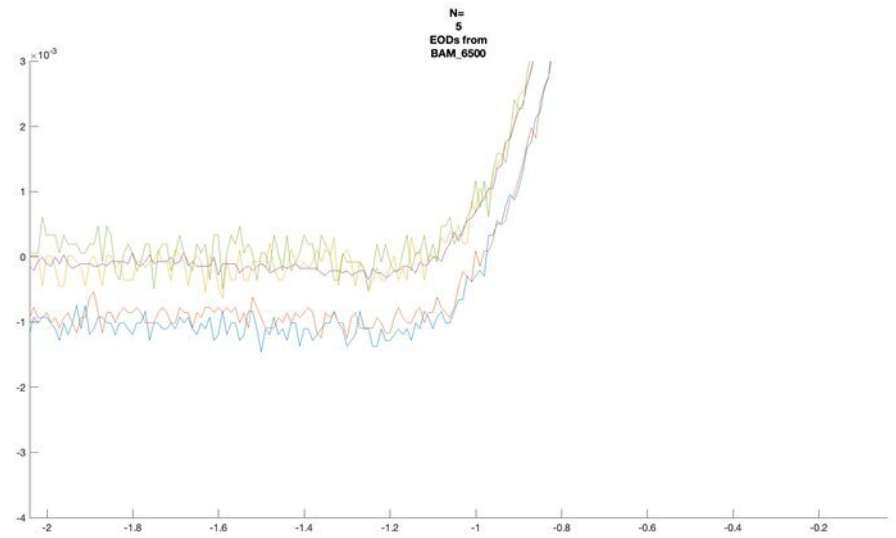
N=
5
EODs from
BAM_6500

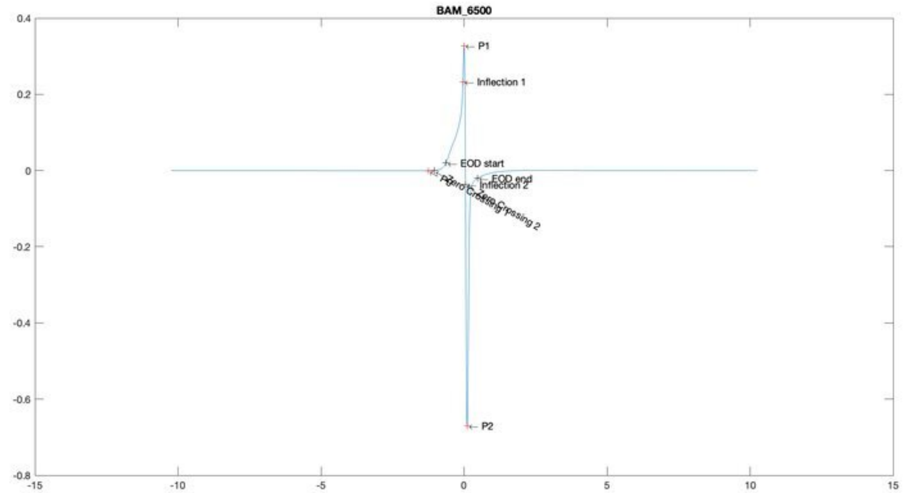
BAM_6500
P1
Inflection 1
EOD start
EOD end
Inflection 2
P2

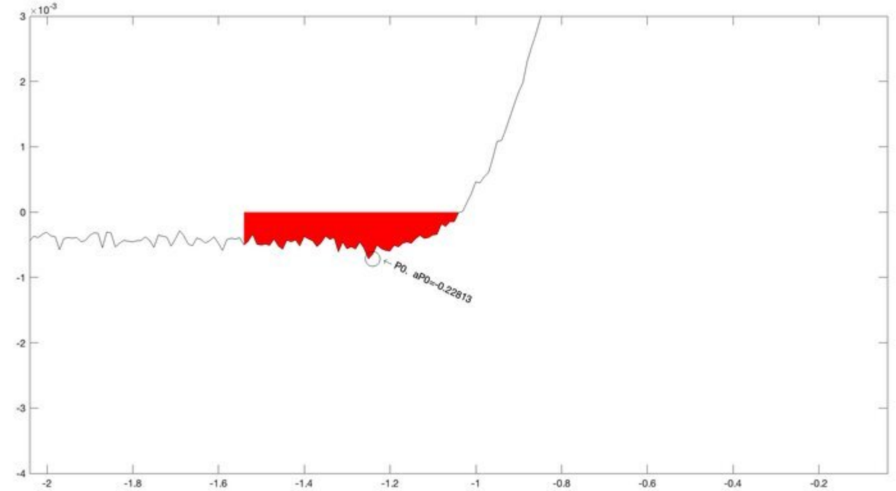
P0, aP0=-0.22813

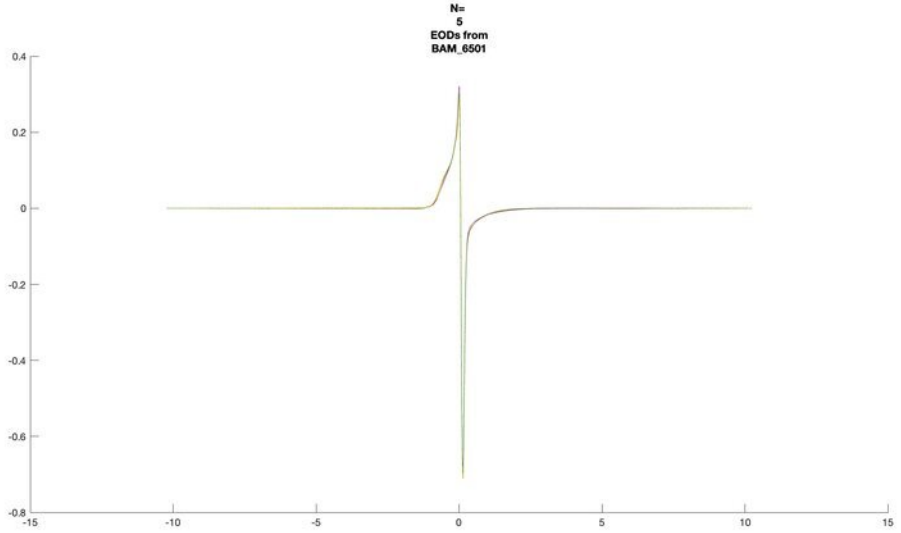
N=
5
EODs from
BAM_6501

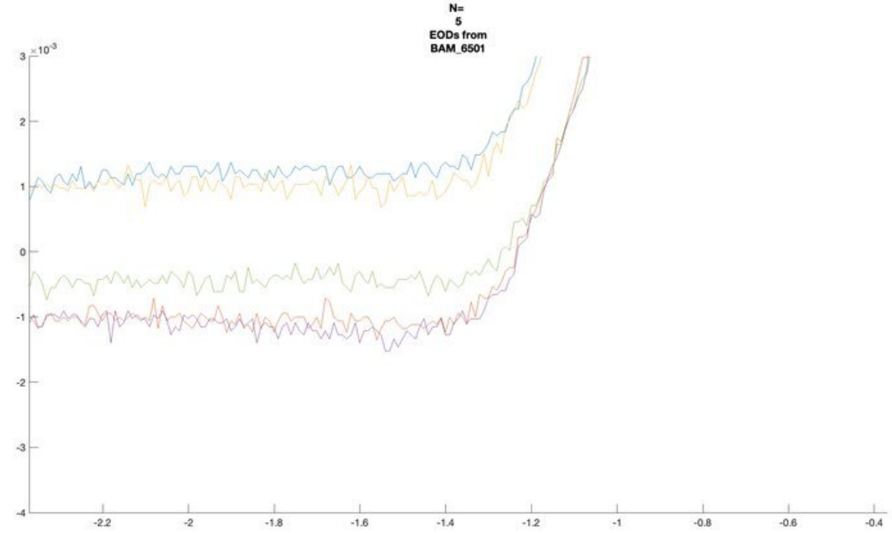
N=
5
EODs from
BAM_6501

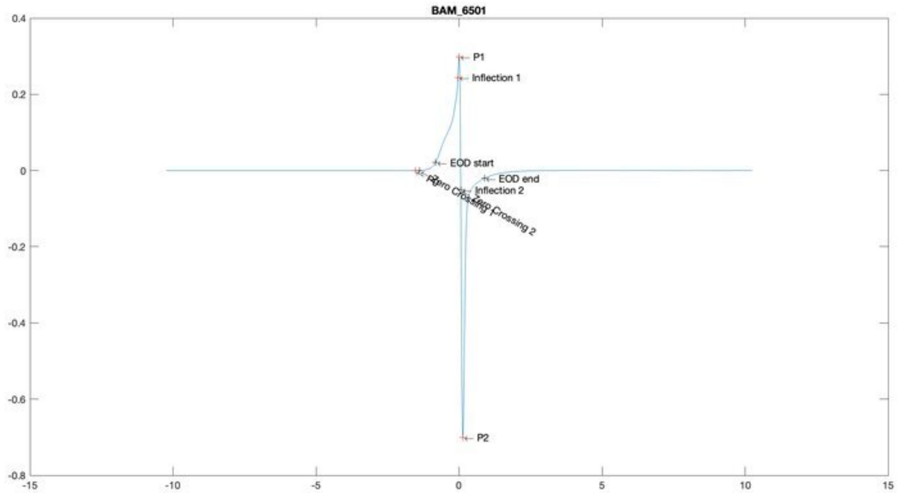
BAM_6501
P1
Inflection 1
EOD start
EOD end
Inflection 2
Zero Crossing 1
Zero Crossing 2
P2

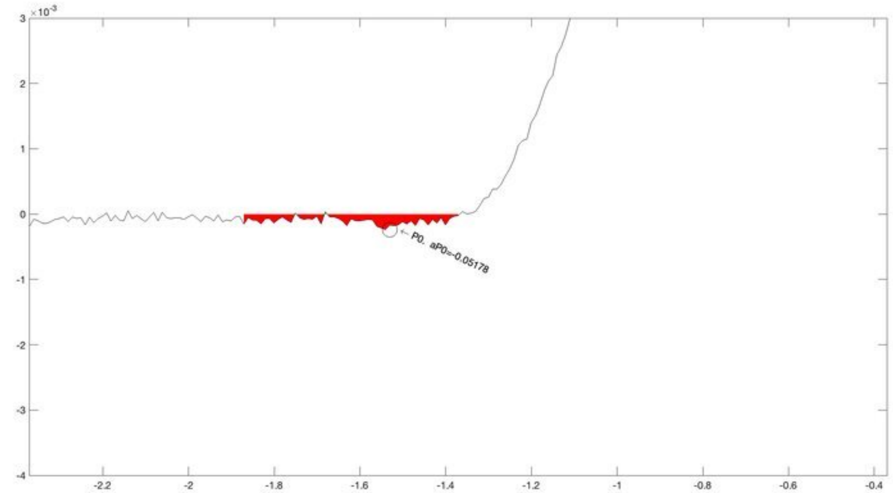
P0, aP0=-0.05178

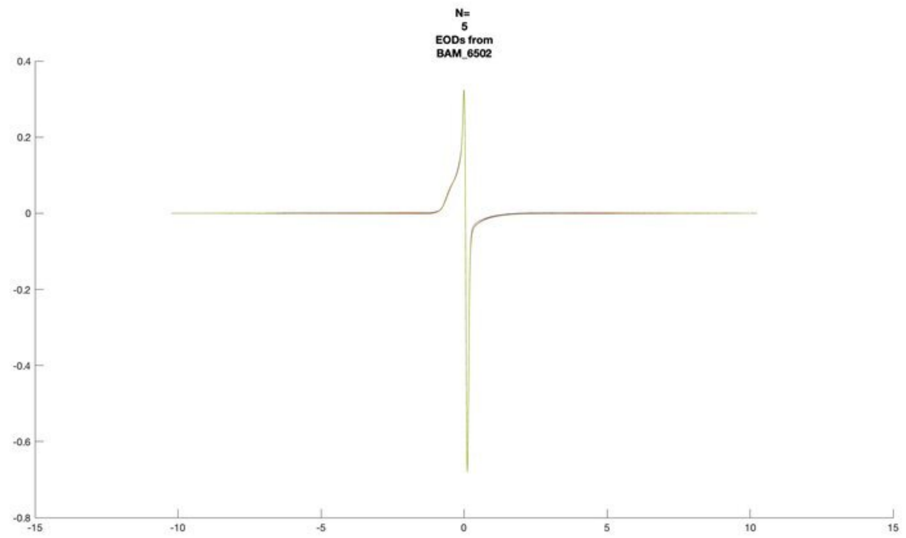
N=
5
EODs from
BAM_6502

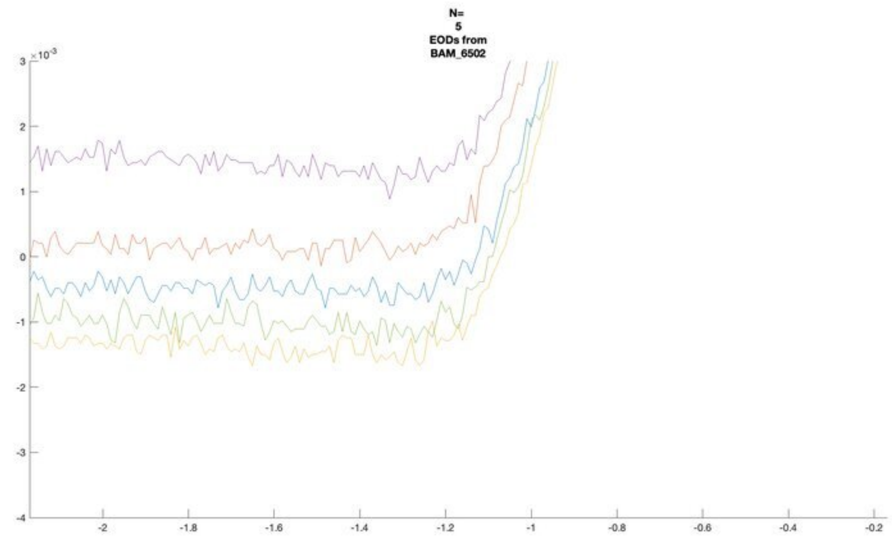
N=
5
EODs from
BAM_6502
×10^-3

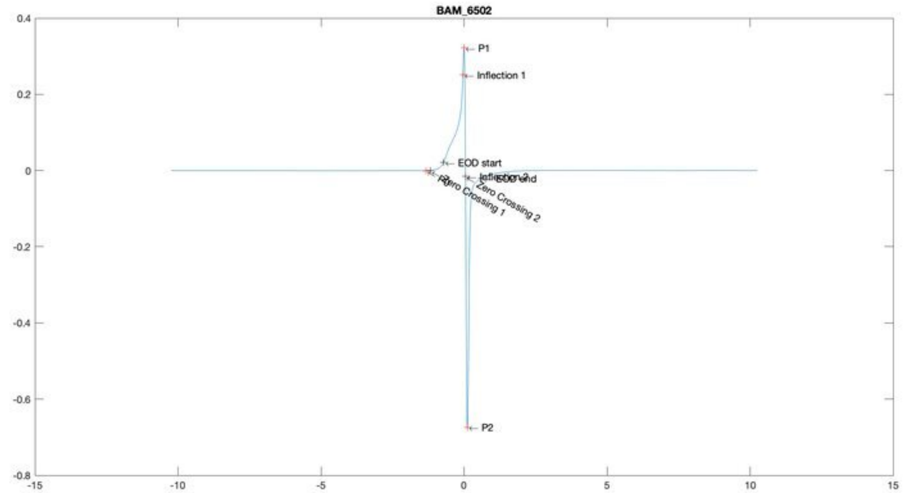
BAM_6502
P1
Inflection 1
EOD start
EOD end
Zero Crossing 1
Zero Crossing 2
P2

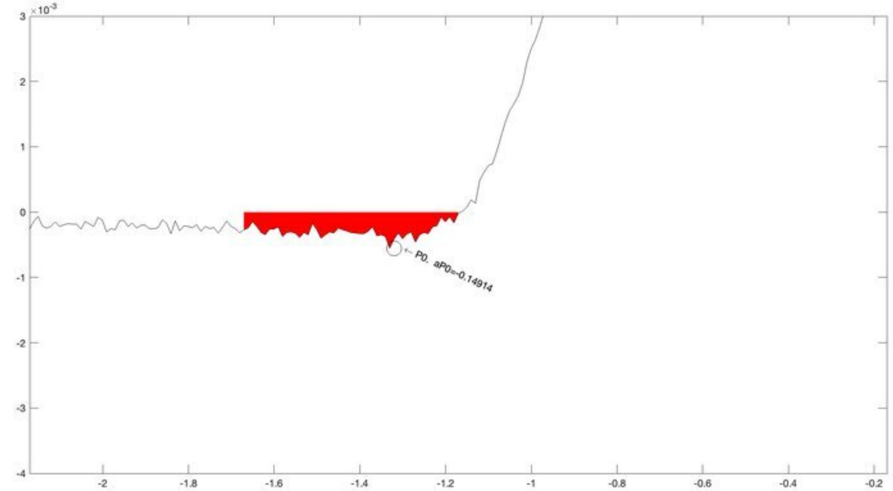
×10^-3
P0. aP0=-0.14914

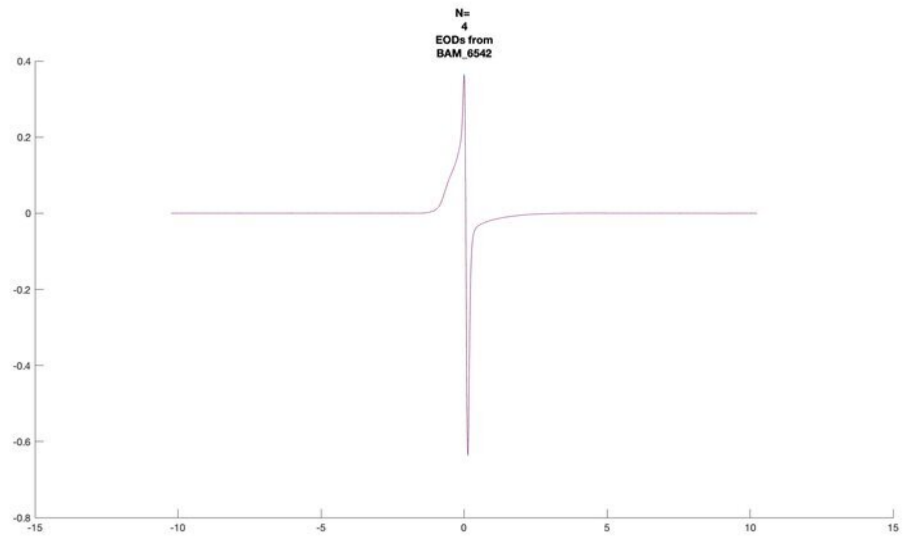
N=
4
EODs from
BAM_6542

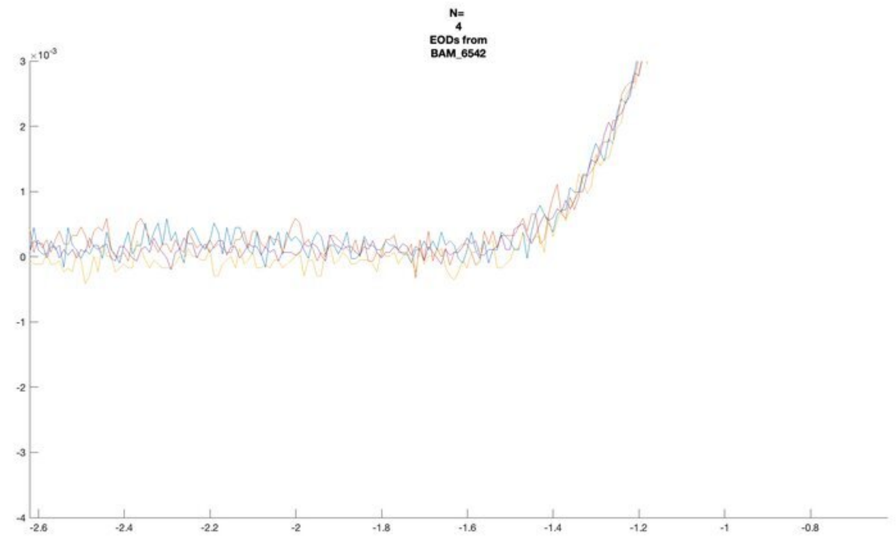
N=
4
EODs from
BAM_6542

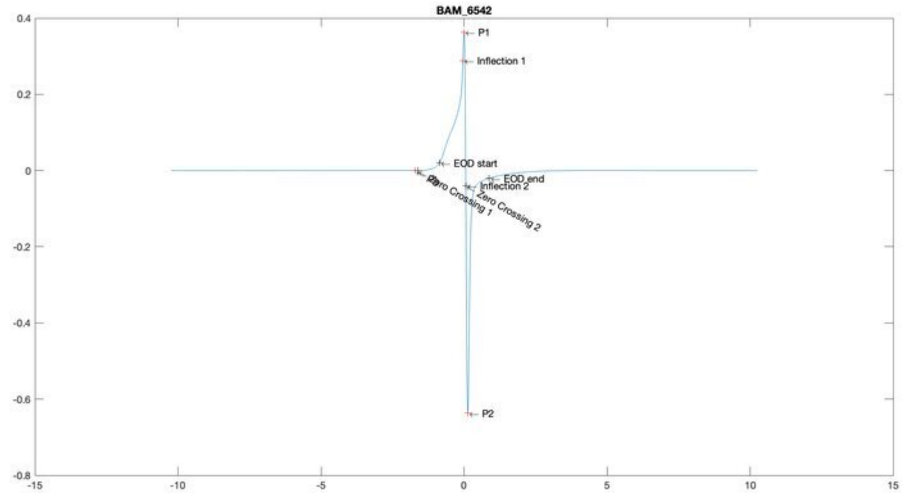
BAM_6542
P1
Inflection 1
EOD start
EOD_end
Inflection 2
Zero Crossing 1
Zero Crossing 2
P2

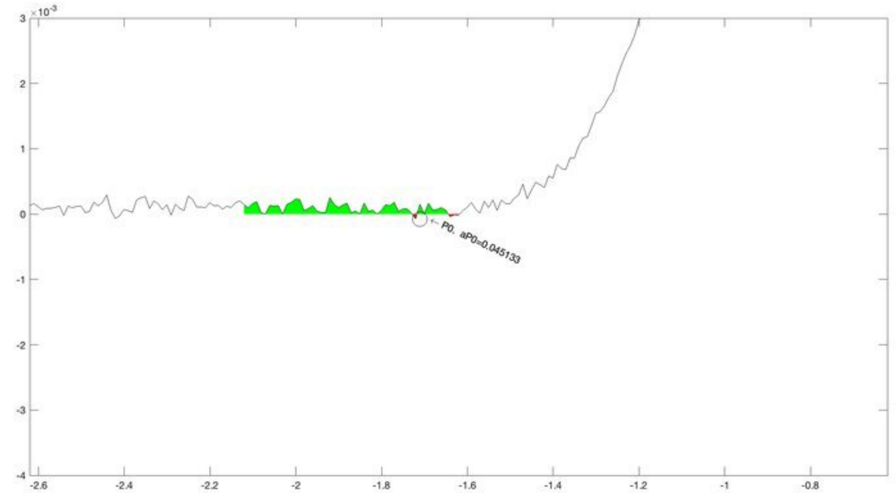
P0, aP0=0.045133

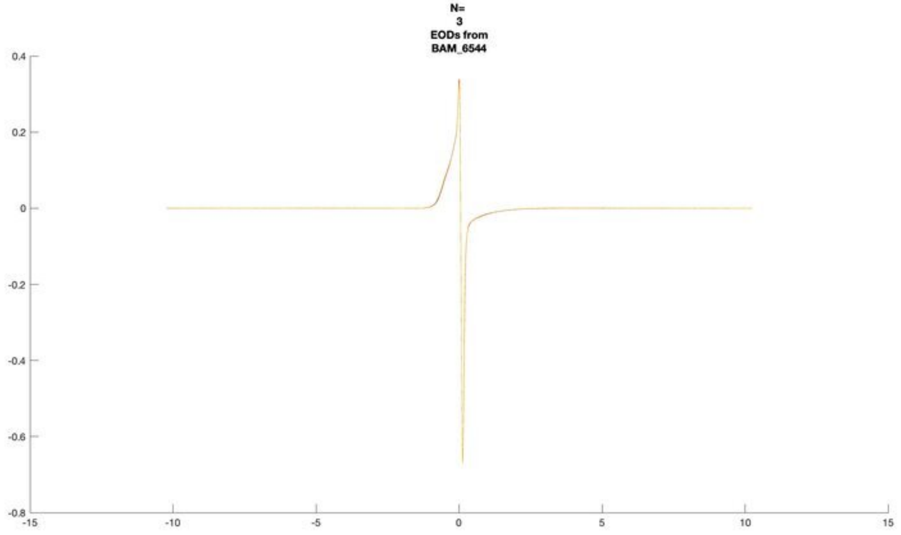
N=
3
EODs from
BAM_6544

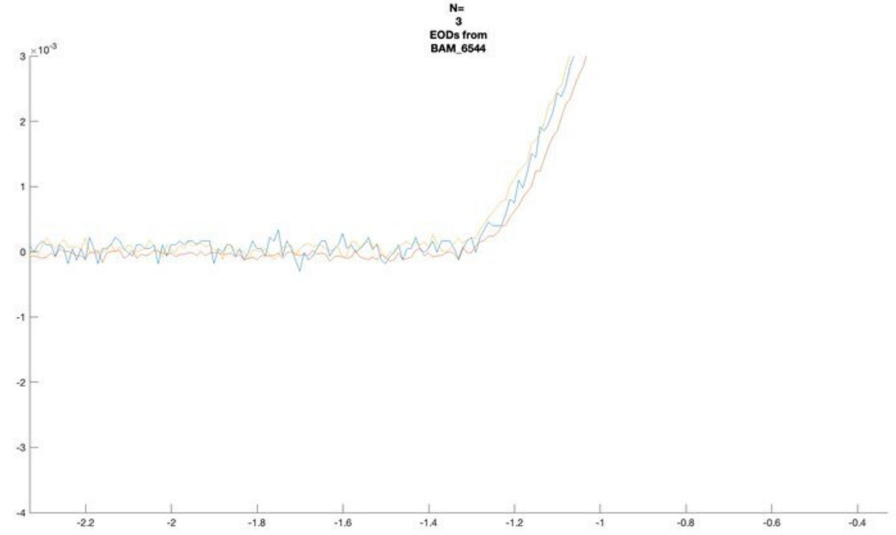
N=
3
EODs from
BAM_6544
×10^-3

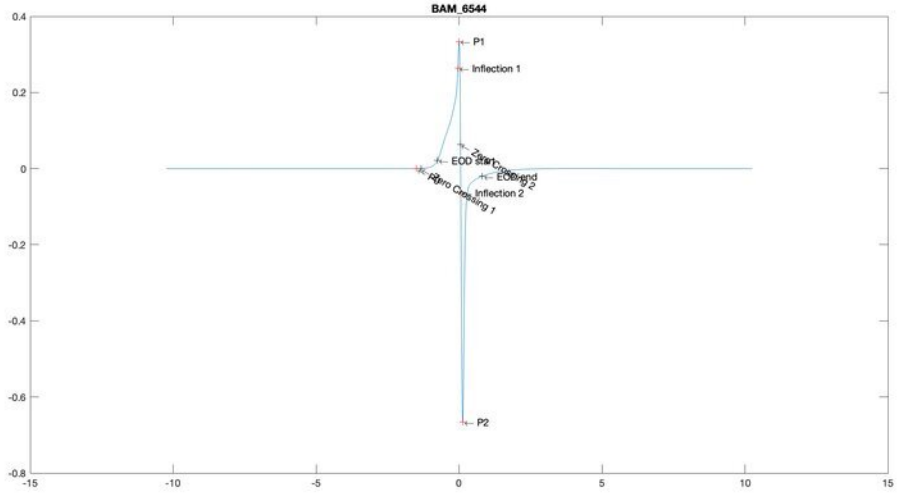
BAM_6544
P1
Inflection 1
EOD start
EOD end
Inflection 2
Zero Crossing 1
Zero Crossing 2
P2

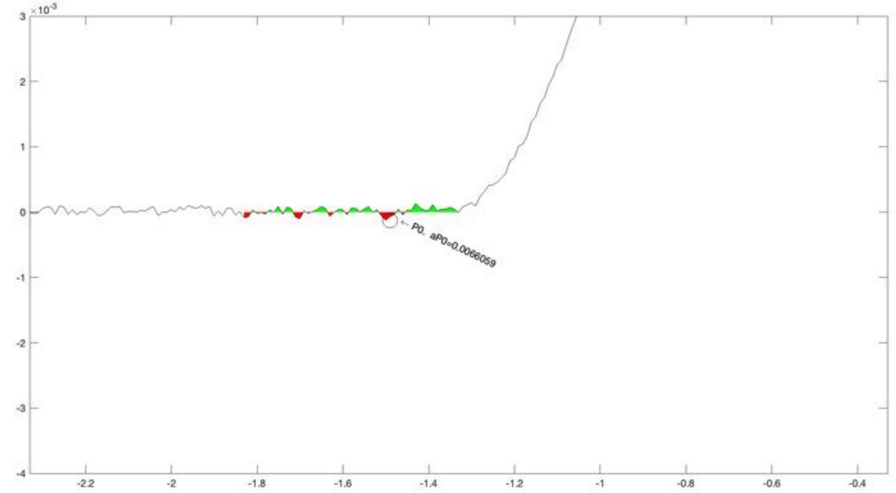
×10^-3
P0, aP0=0.0066059

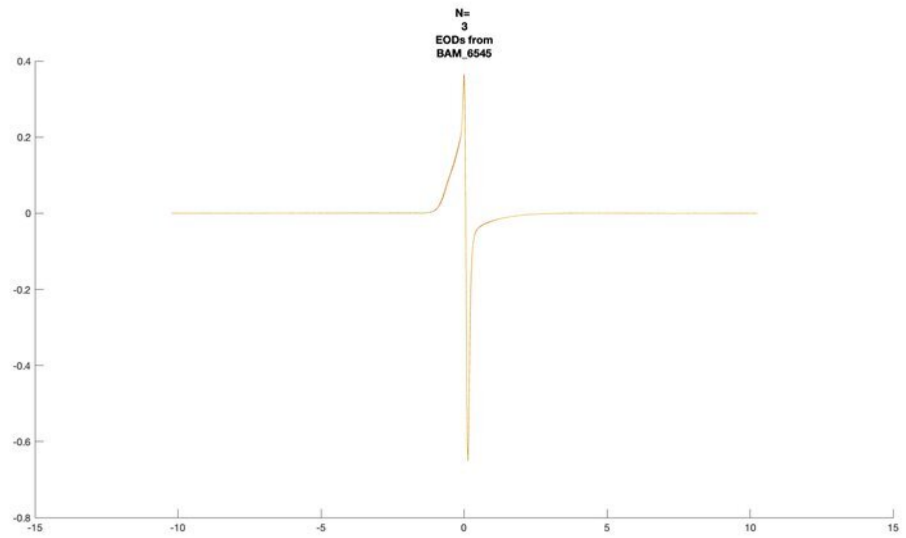
N=
3
EODs from
BAM_6545

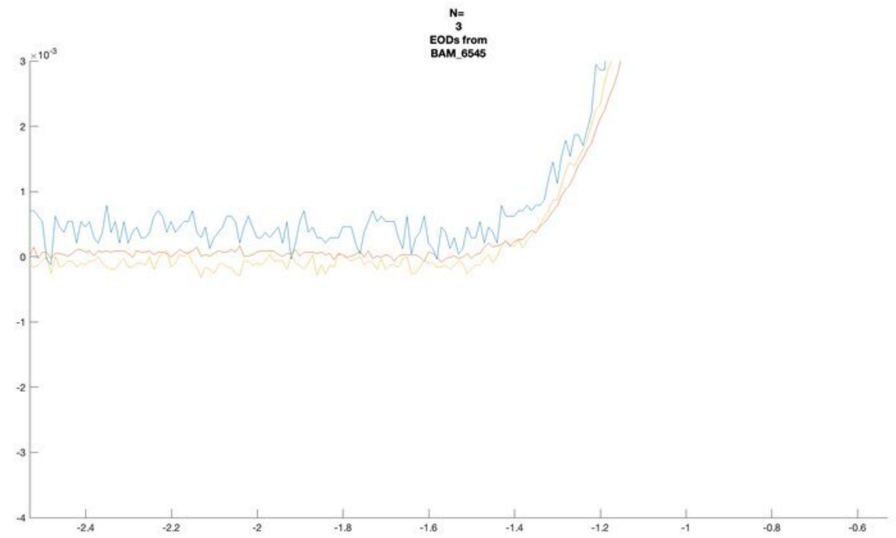
N=
3
EODs from
BAM_6545
×10^-3

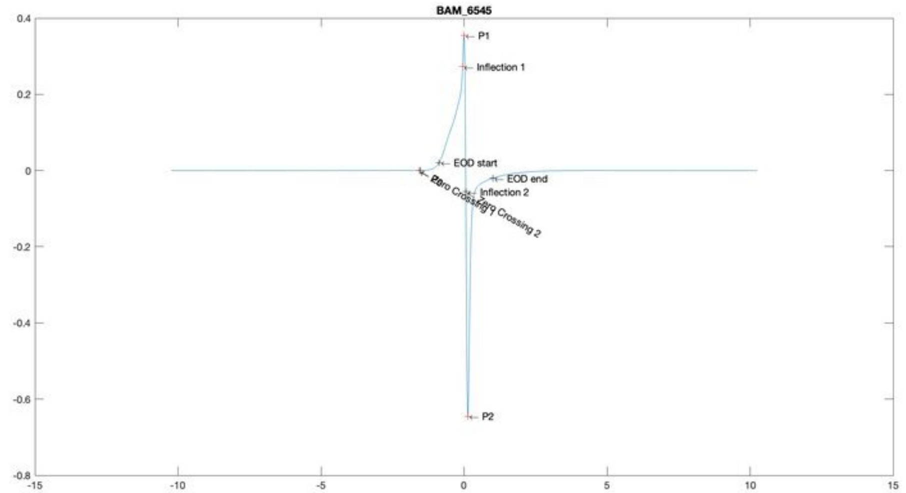
BAM_6545
P1
Inflection 1
EOD start
EOD end
Inflection 2
Zero Crossing 2
P2

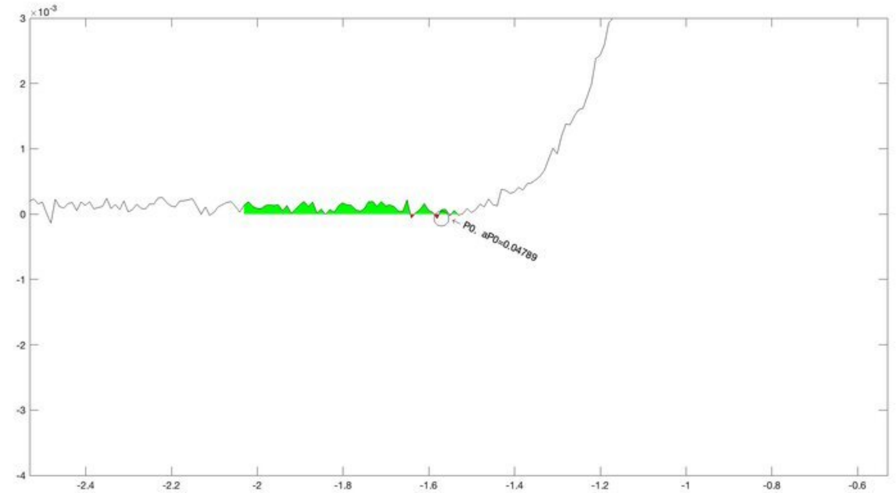
×10^-3
P0, aP0=0.04789

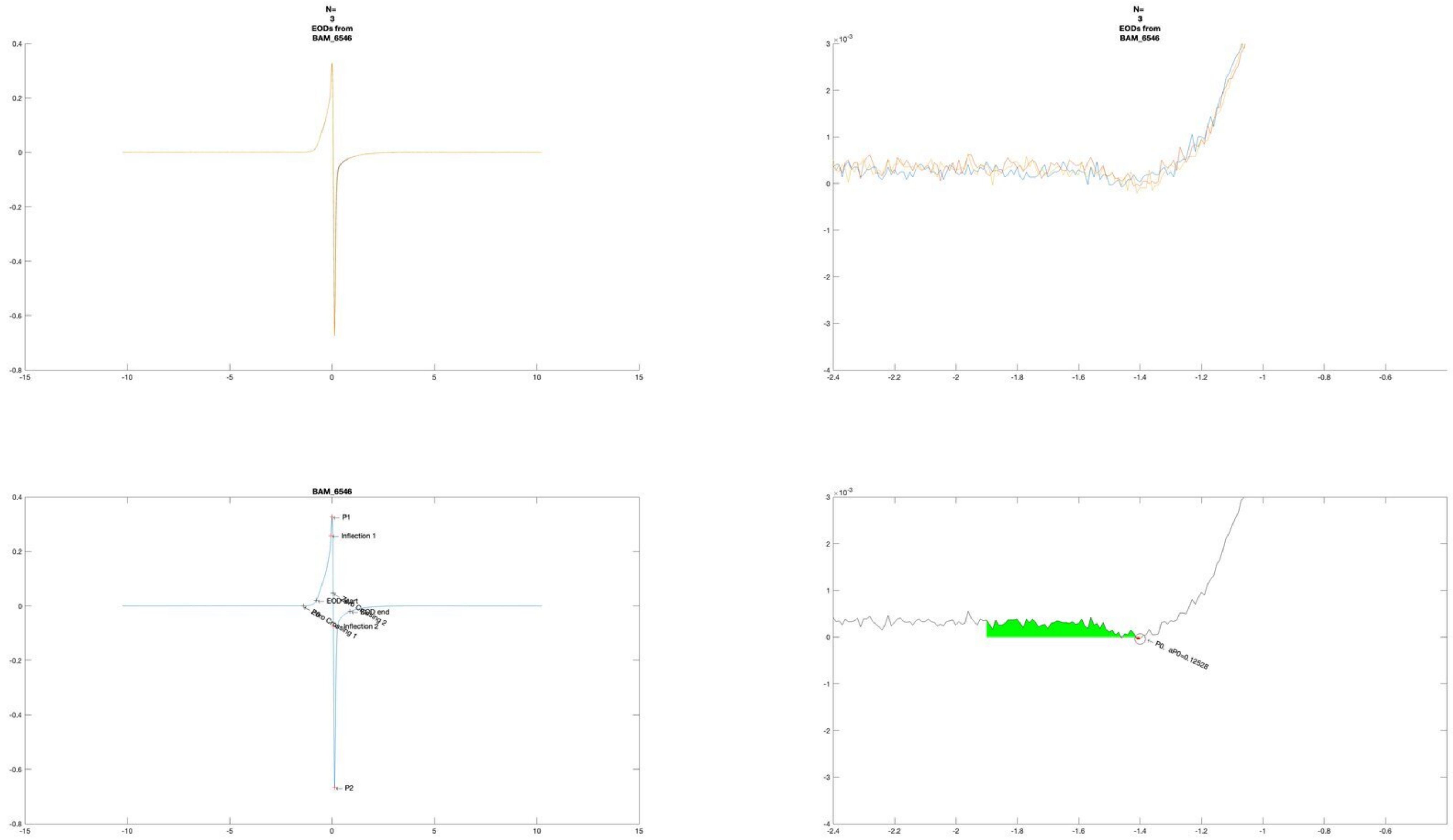
N=
3
EODs from
BAM_6546
N=
3
EODs from
BAM_6546
BAM_6546
P1
Inflection 1
EOD start
EOD end
Inflection 2
P2
P0, aP0=0.12528

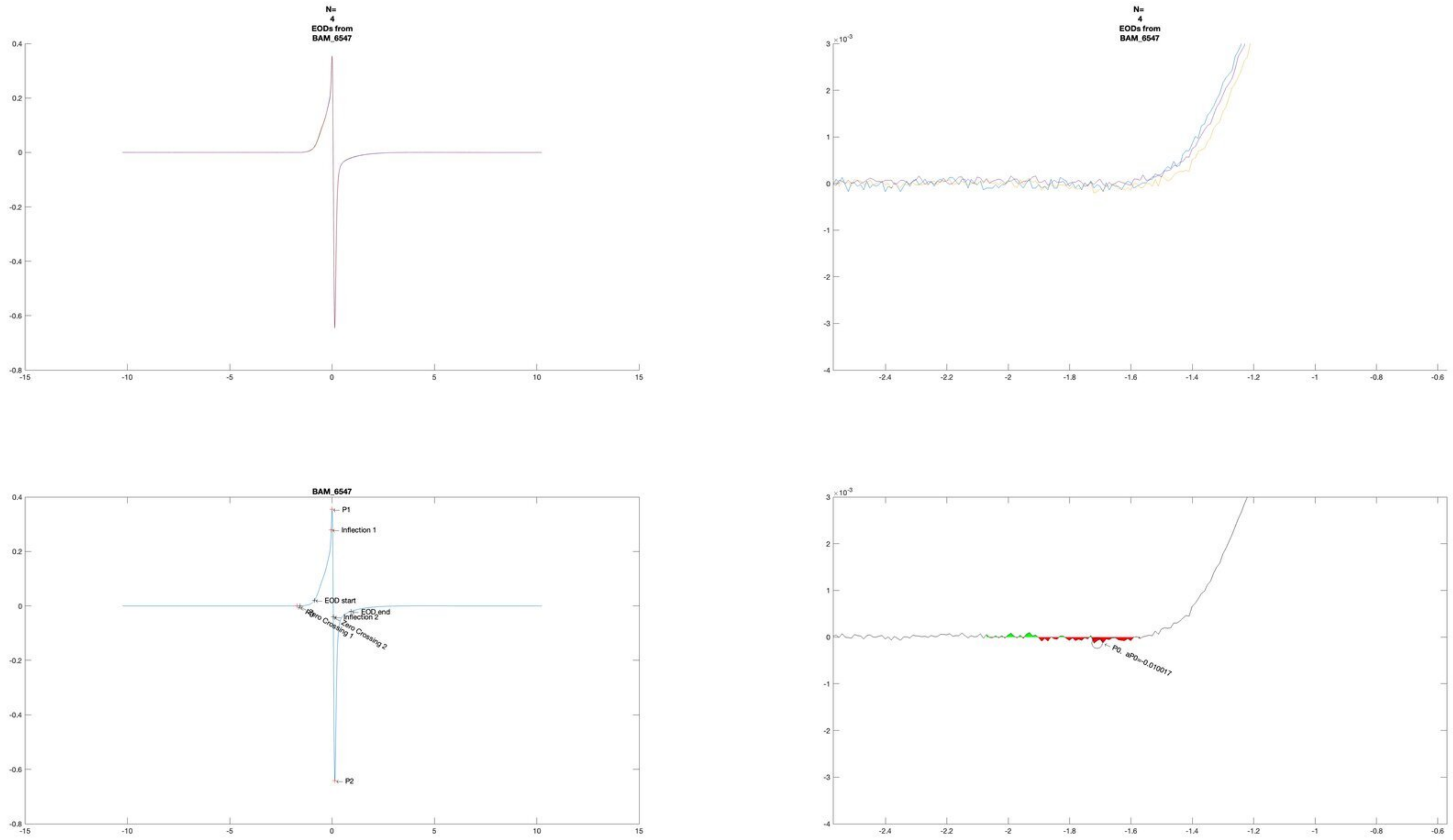

N=
4
EODs from
BAM_6547
N=
4
EODs from
BAM_6547
BAM_6547
P1
Inflection 1
EOD start
EOD end
Inflection 2
Zero Crossing 1
Zero Crossing 2
P2
P0, aP0=-0.010017

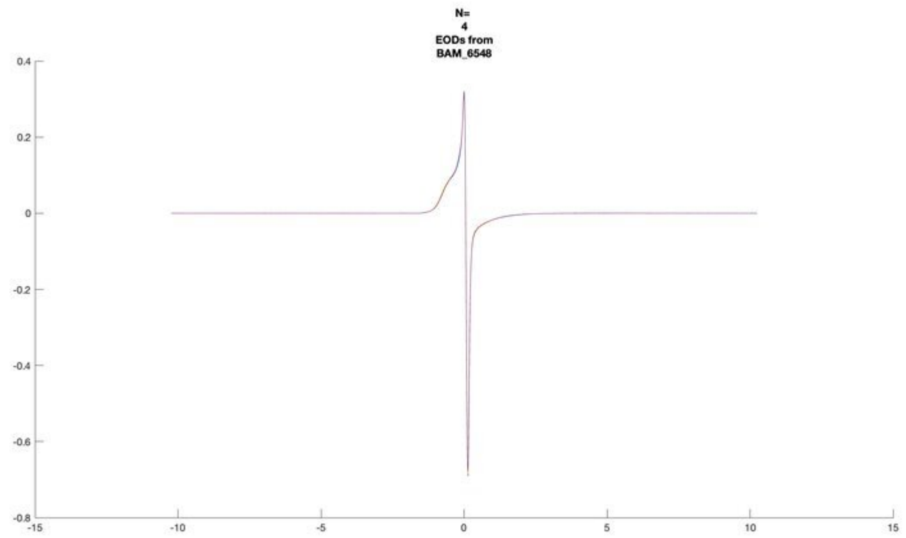
N=
4
EODs from
BAM_6548

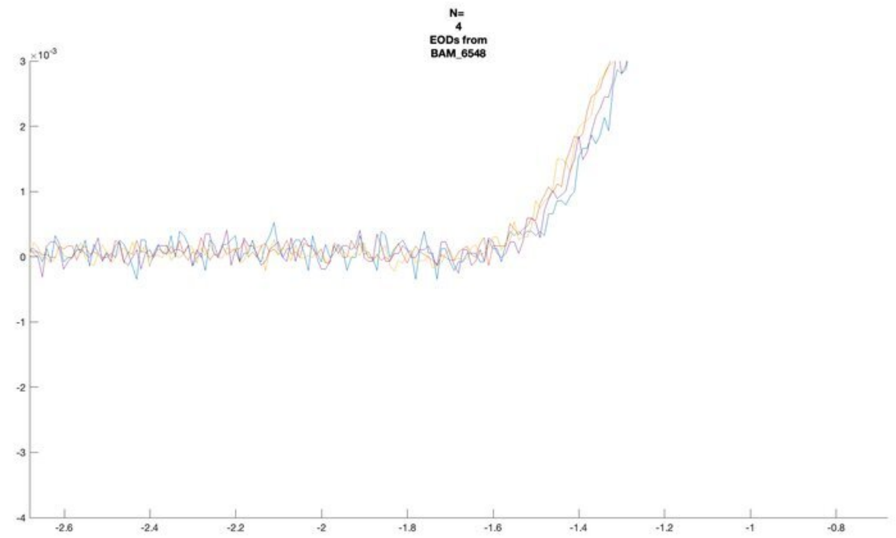
N=
4
EODs from
BAM_6548
×10^-3

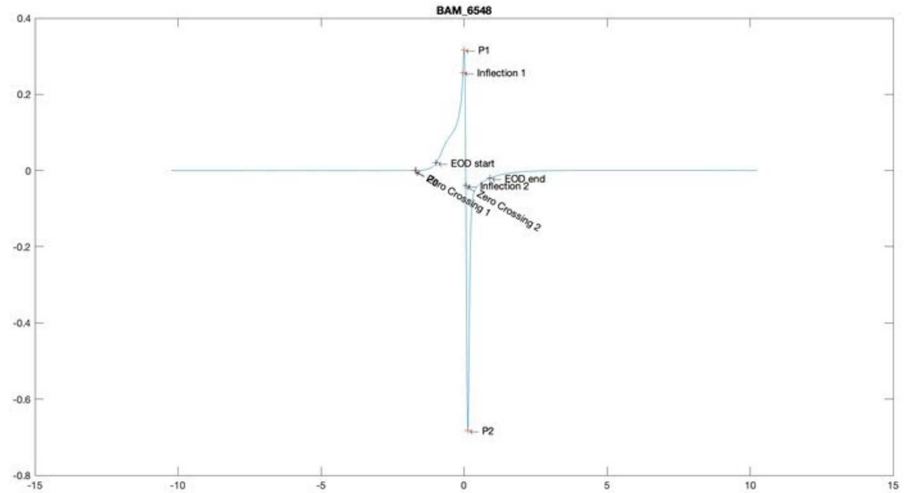
BAM_6548
P1
Inflection 1
EOD start
EOD_end
Inflection 2
Zero Crossing 1
Zero Crossing 2
P2

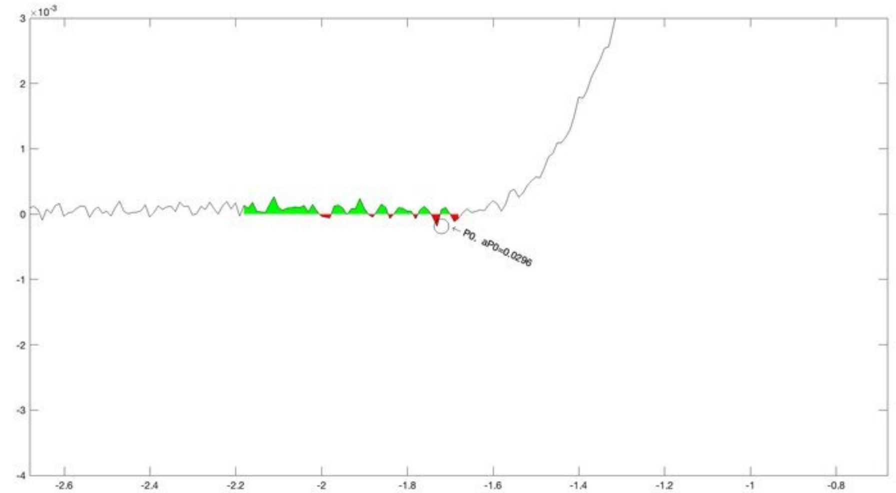
×10^-3
P0, aP0=0.0296

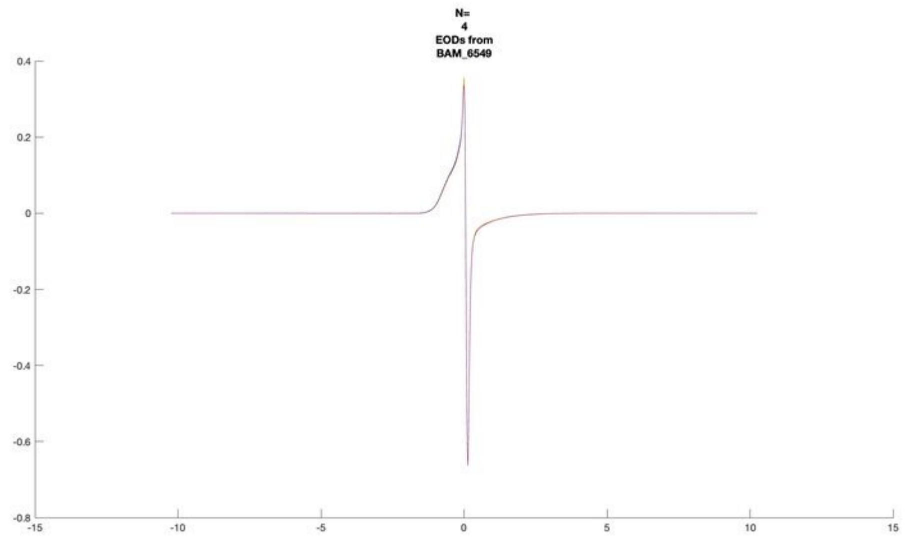
N=
4
EODs from
BAM_6549

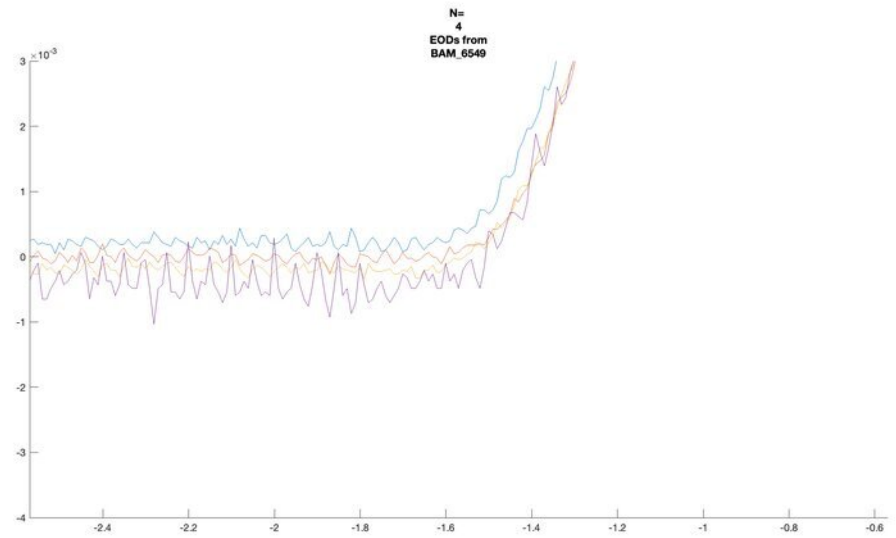
N=
4
EODs from
BAM_6549
×10^-3

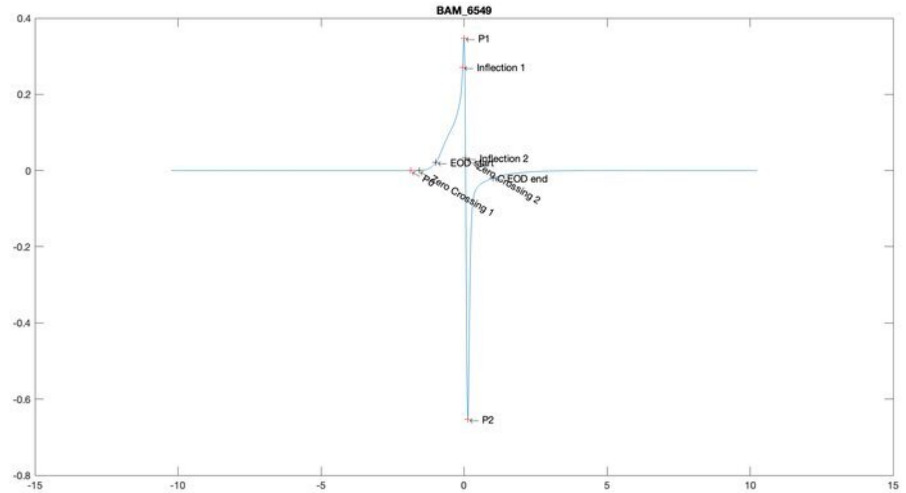
BAM_6549
P1
Inflection 1
Inflection 2
EOD start
EOD end
P0
Zero Crossing 1
Zero Crossing 2
P2

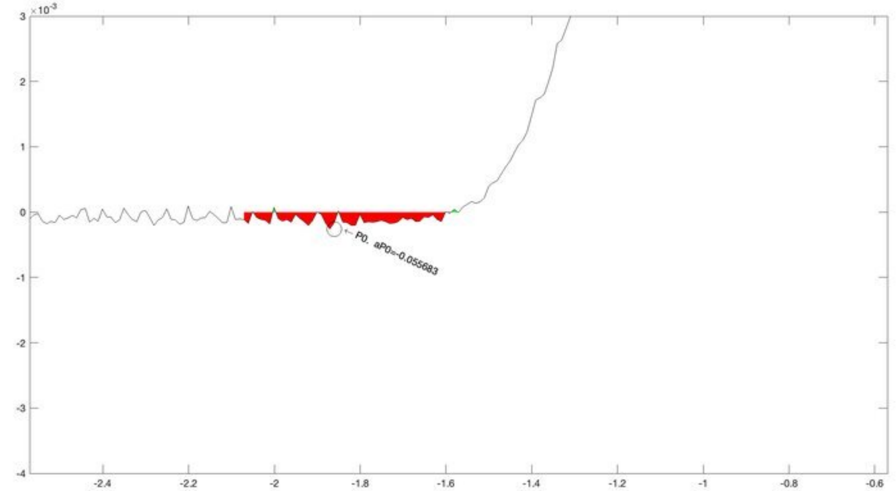
×10^-3
P0, aP0=-0.055683

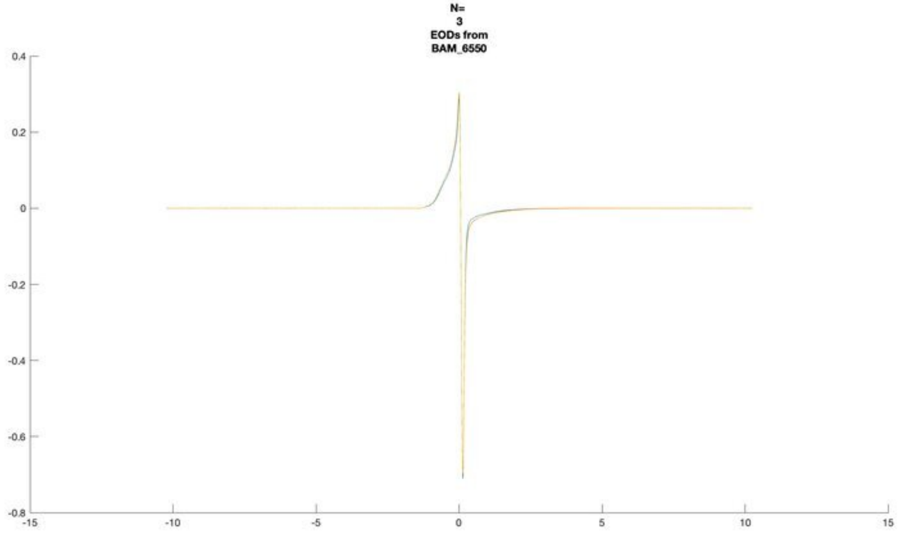
N=
3
EODs from
BAM_6550

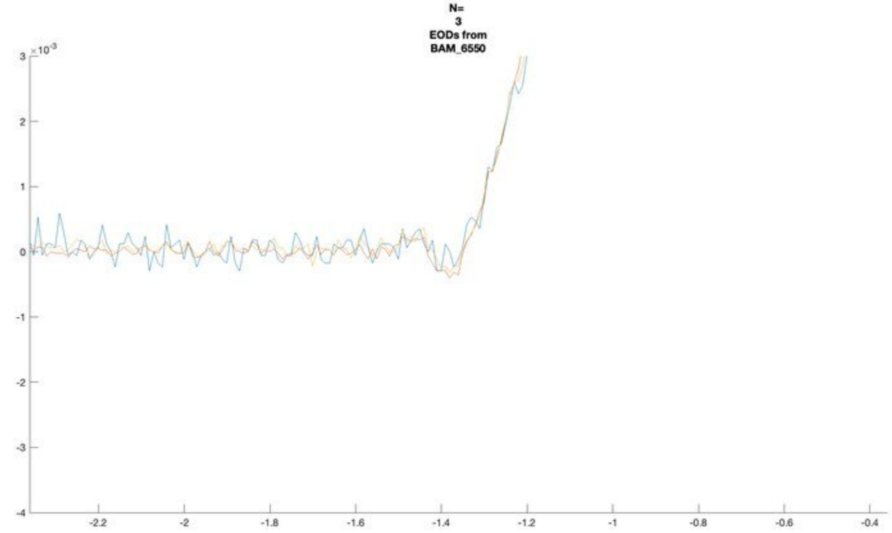
N=
3
EODs from
BAM_6550
×10^-3

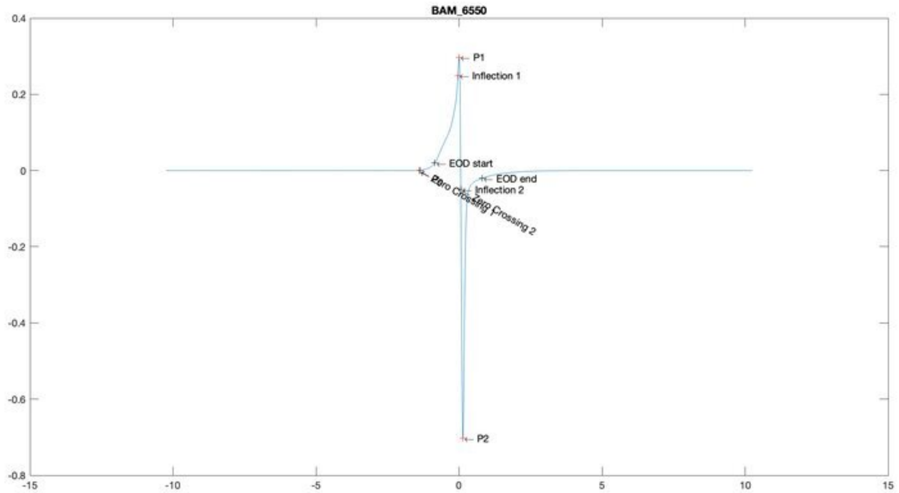
BAM_6550
P1
Inflection 1
EOD start
EOD end
Inflection 2
P2

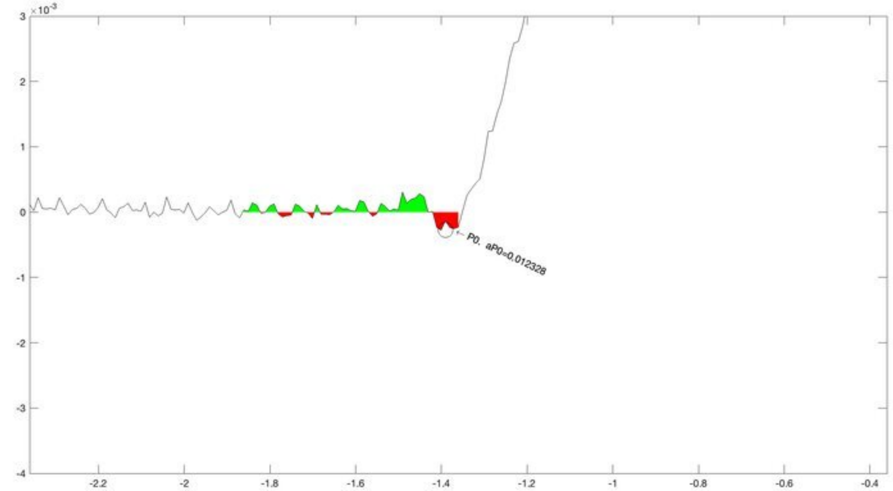
×10^-3
P0, aP0=0.012328

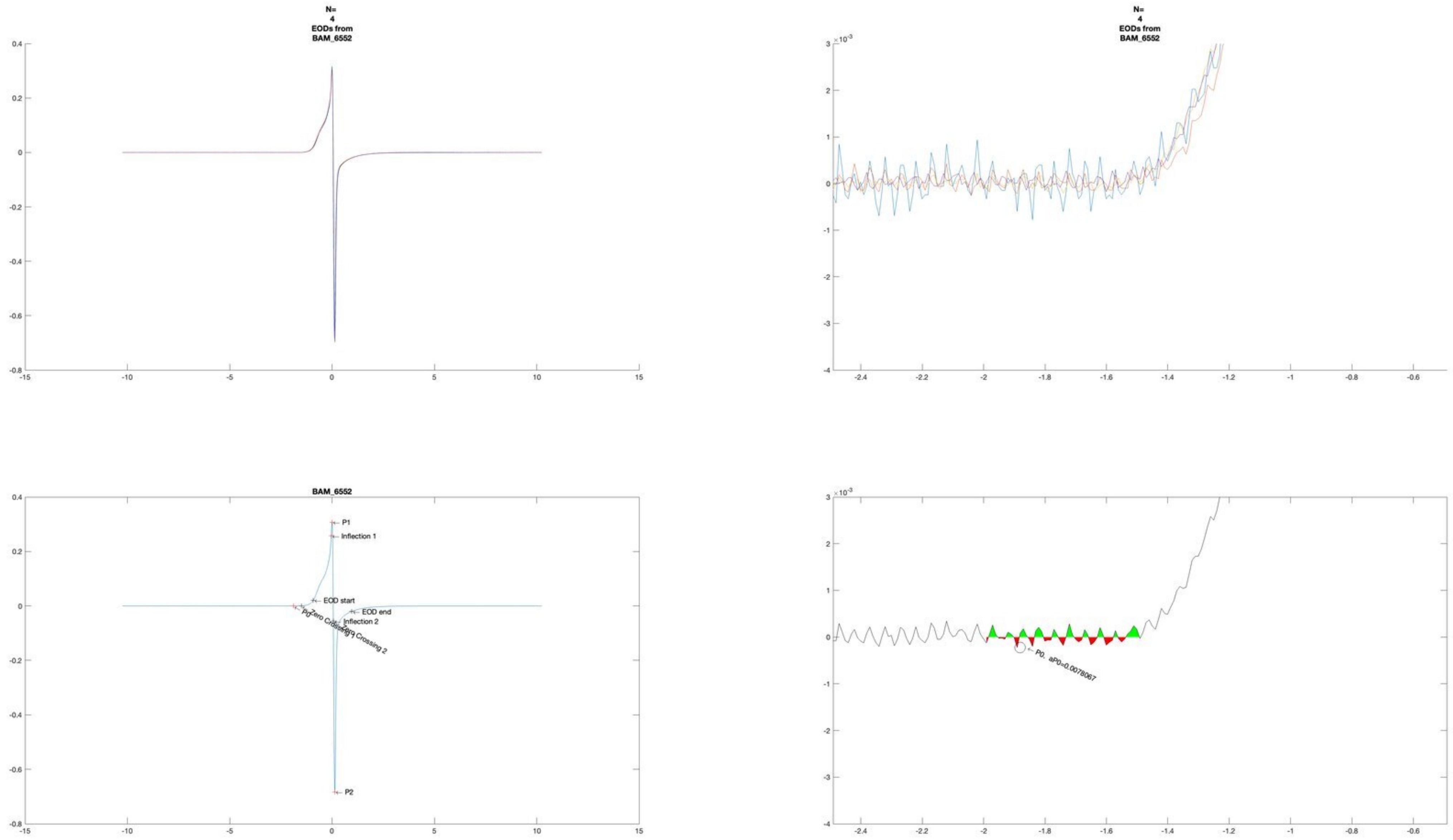

N=
4
EODs from
BAM_6552
N=
4
EODs from
BAM_6552
BAM_6552
P1
Inflection 1
EOD start
EOD end
Inflection 2
P0
Zero Crossing 1
Zero Crossing 2
P2
P0, aP0=0.0078067

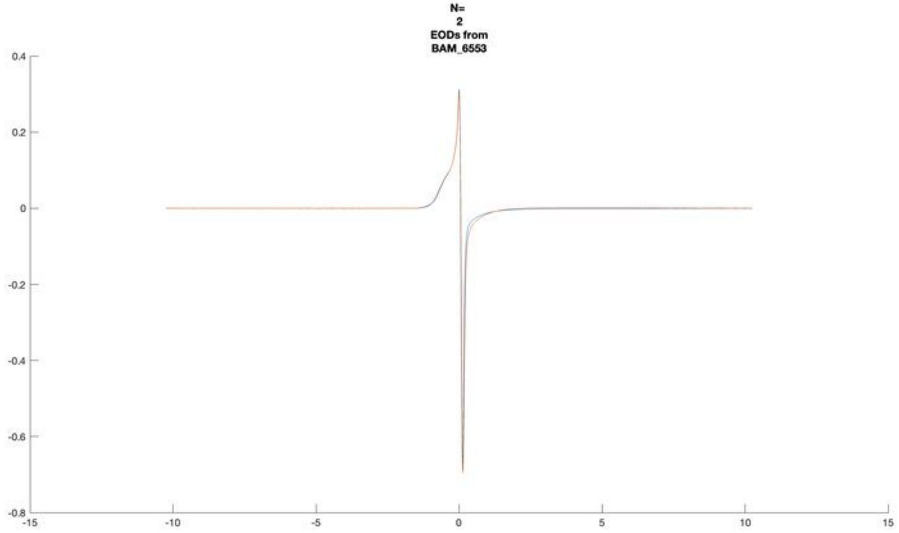
N=
2
EODs from
BAM_6553

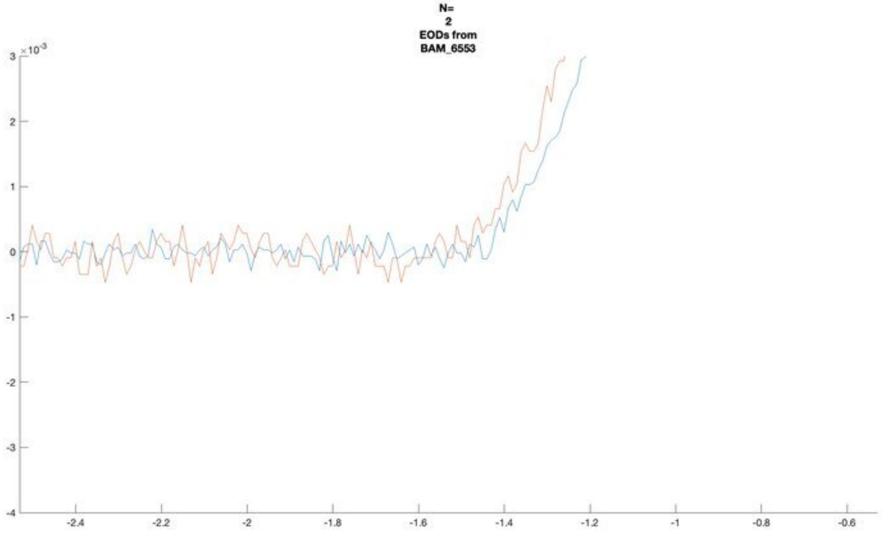
N=
2
EODs from
BAM_6553

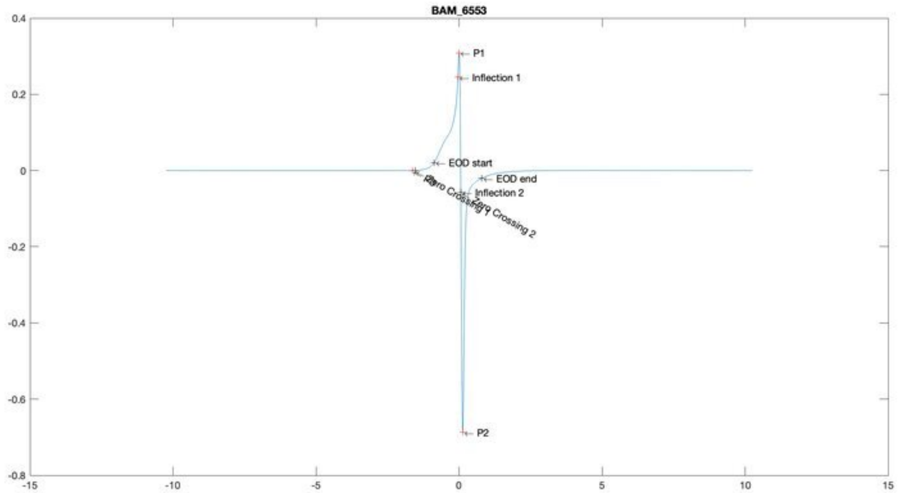
BAM_6553
P1
Inflection 1
EOD start
EOD end
Inflection 2
Zero Crossing 1
Zero Crossing 2
P2

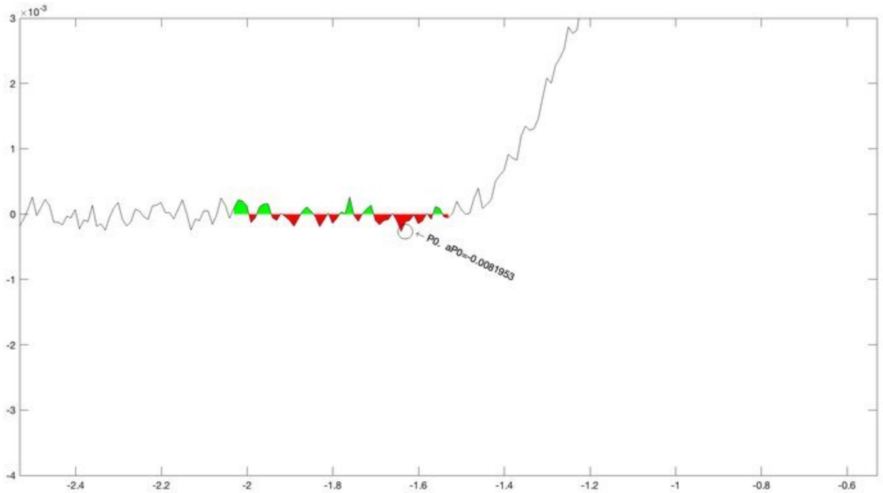
P0, aP0=-0.0081953

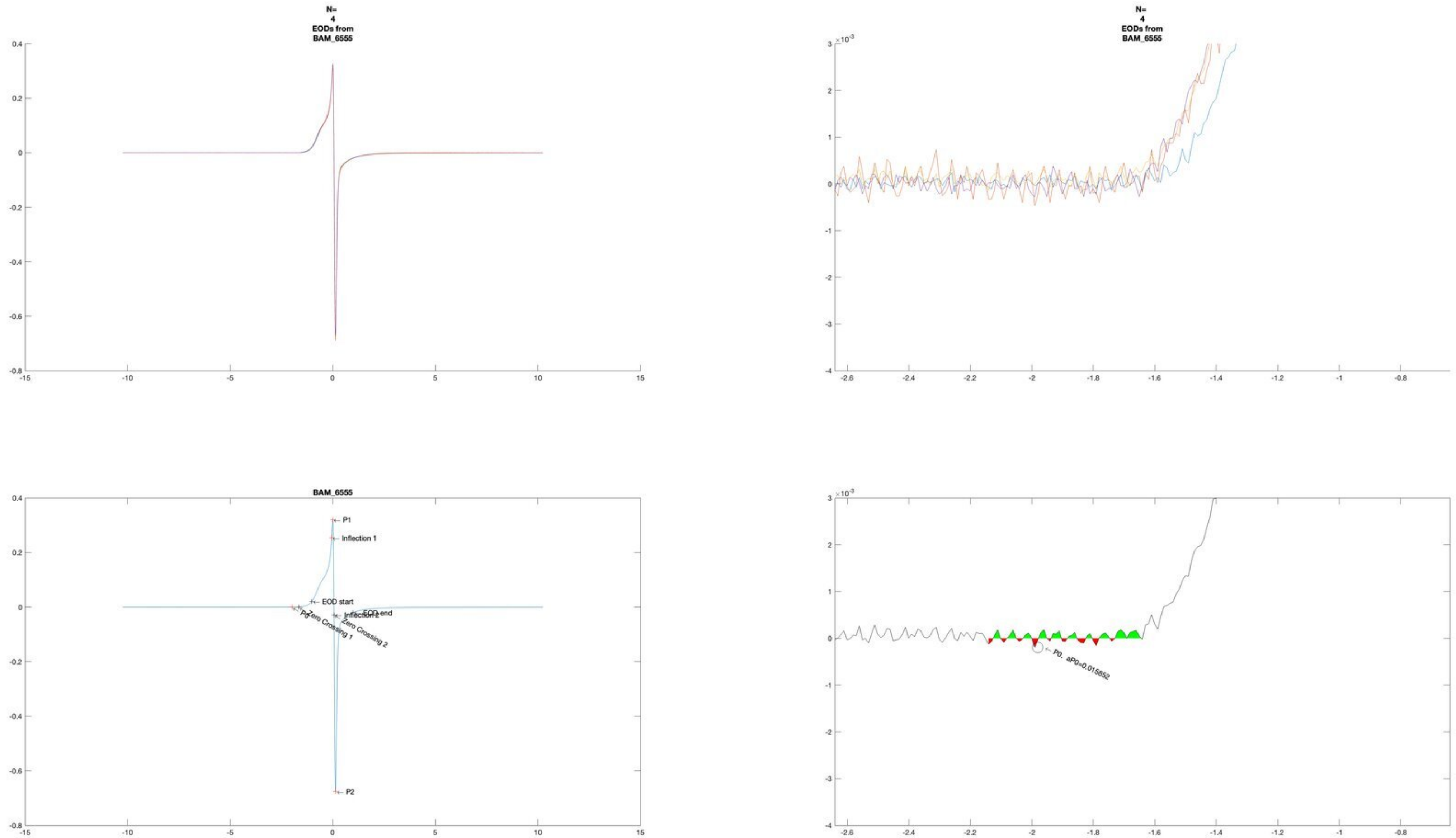
N=
4
EODs from
BAM_6555
N=
4
EODs from
BAM_6555
BAM_6555
P1
Inflection 1
EOD start
EOD end
Inflection 2
P0
Zero Crossing 1
Zero Crossing 2
P2
P0. aP0=0.015852

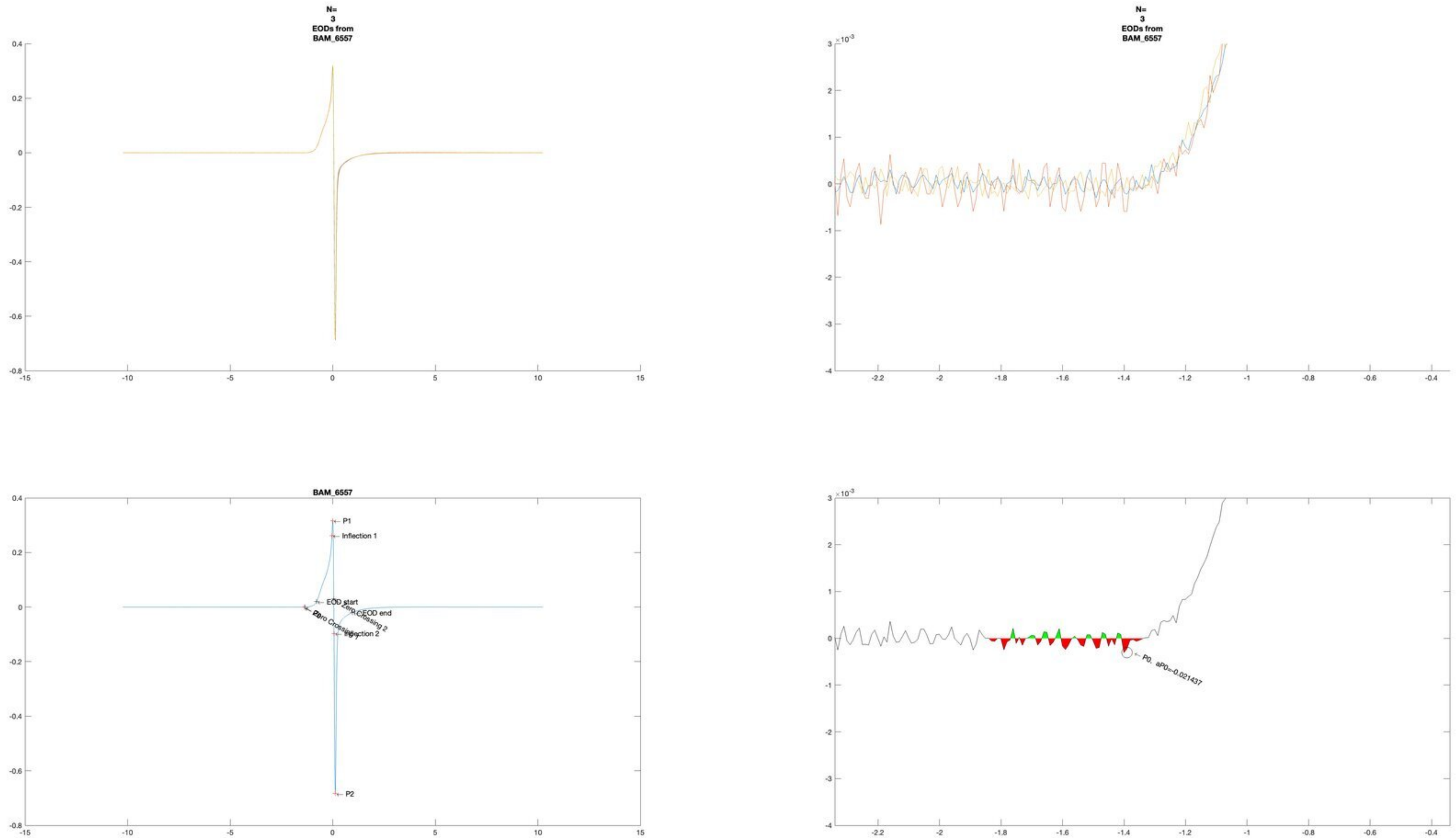
N=
3
EODs from
BAM_6557
N=
3
EODs from
BAM_6557
BAM_6557
P1
Inflection 1
EOD start
EOD end
Zero Crossing 2
Inflection 2
P2
P0. aP0=-0.021437

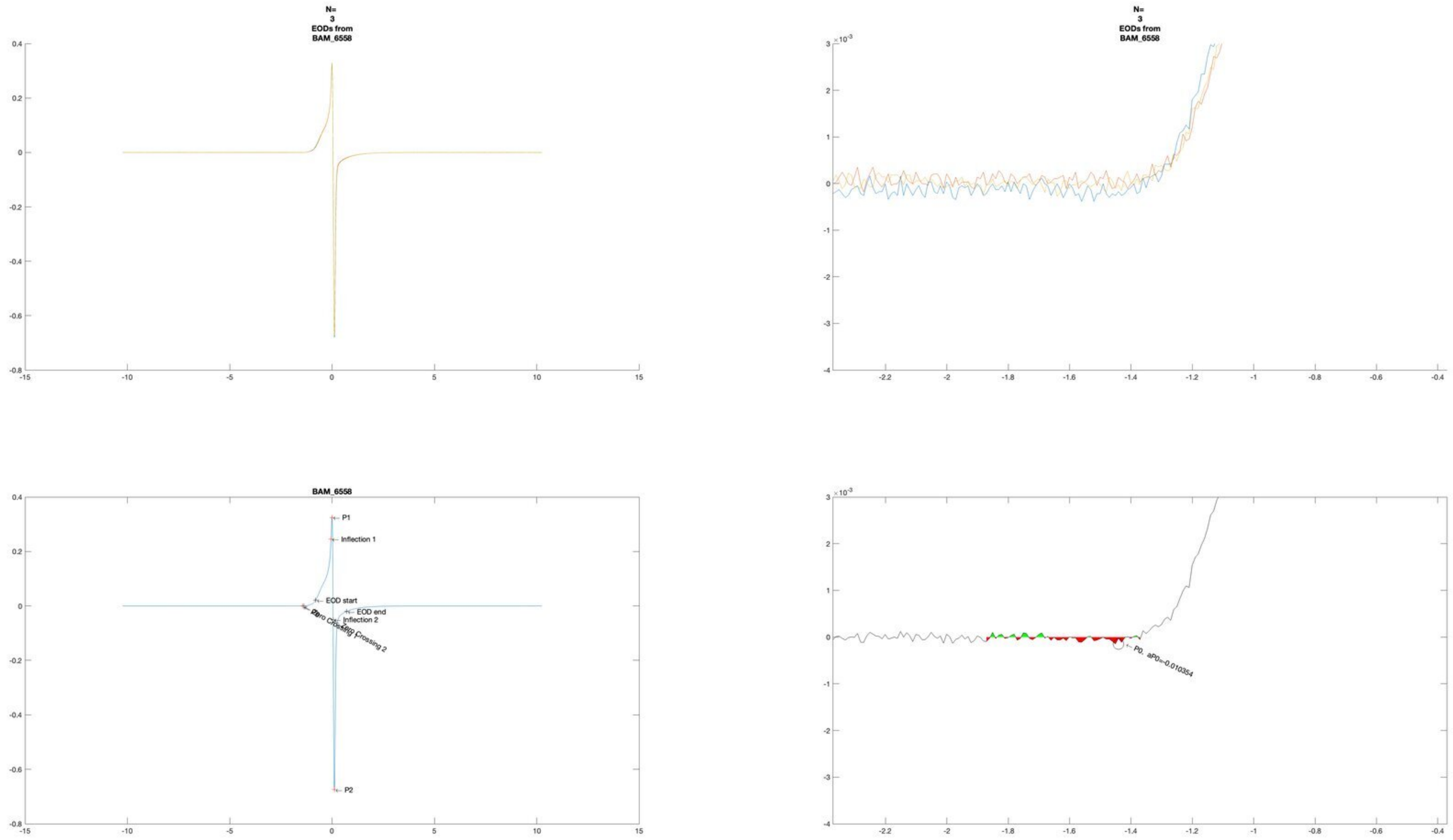
N=
3
EODs from
BAM_6558
N=
3
EODs from
BAM_6558
BAM_6558
P1
Inflection 1
EOD start
EOD end
Inflection 2
Zero Crossing 1
Zero Crossing 2
P2
P0, aP0=-0.010354

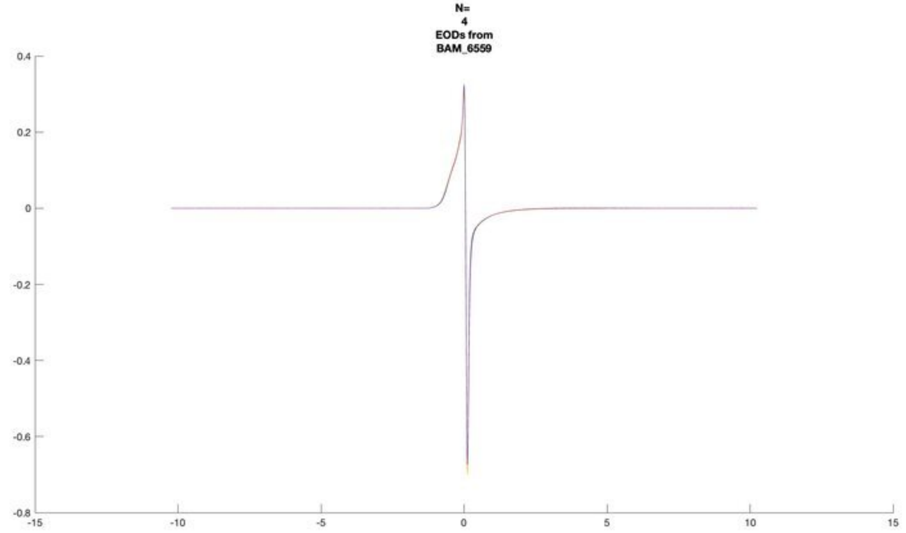
N=
4
EODs from
BAM_6559

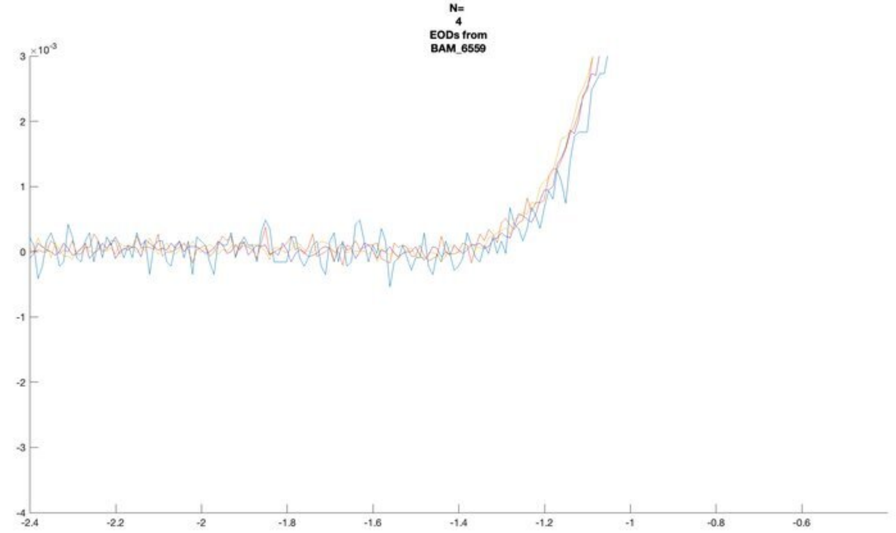
N=
4
EODs from
BAM_6559
×10^-3

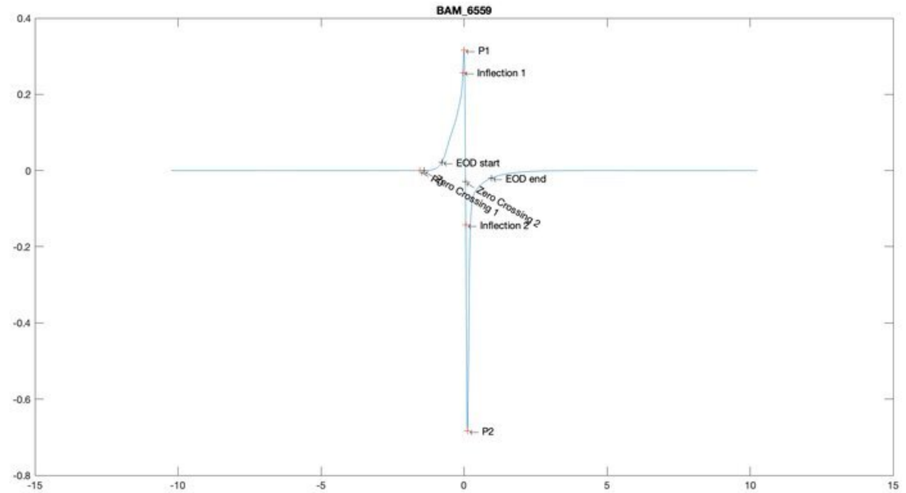
BAM_6559
P1
Inflection 1
EOD start
EOD end
Zero Crossing 1
Zero Crossing 2
Inflection 2
P2

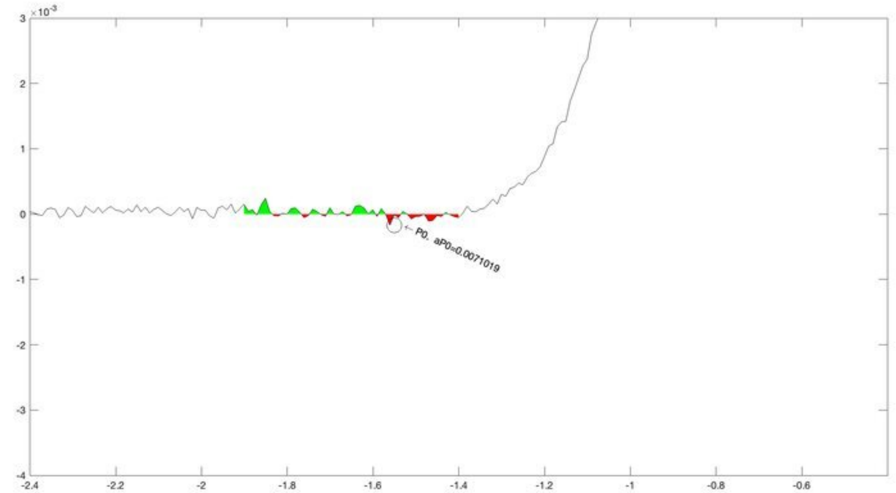
×10^-3
P0, aP0=0.0071019

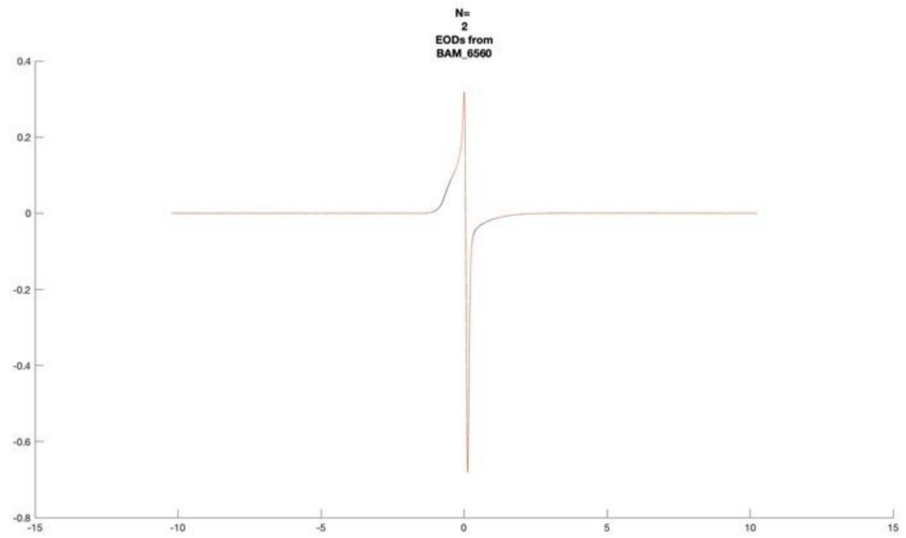
N=
2
EODs from
BAM_6560

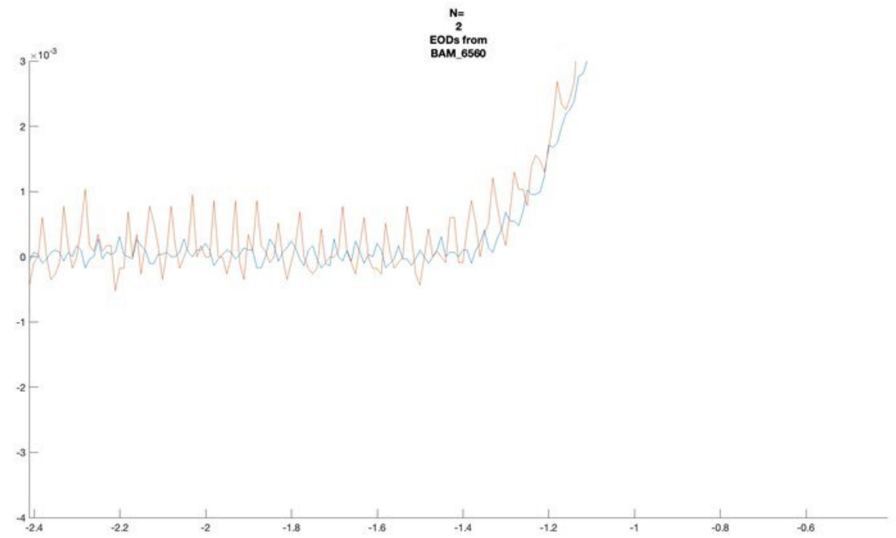
N=
2
EODs from
BAM_6560
$\times10^{-3}$

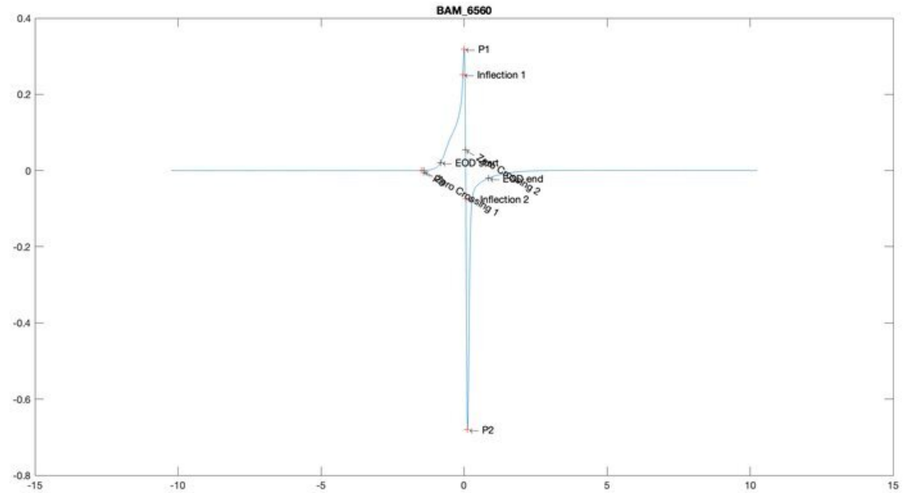
BAM_6560
P1
Inflection 1
EOD start
Zero Crossing 2
EOD end
Inflection 2
Zero Crossing 1
P2

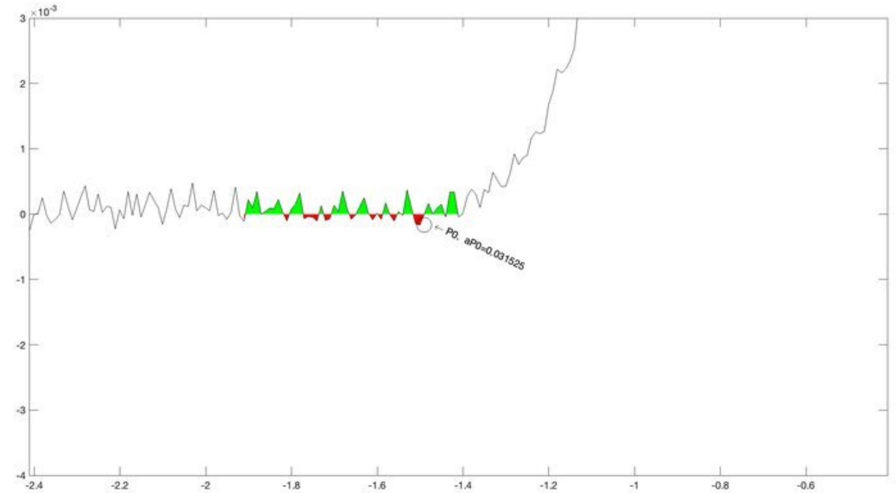
$\times10^{-3}$
P0, aP0=0.031525

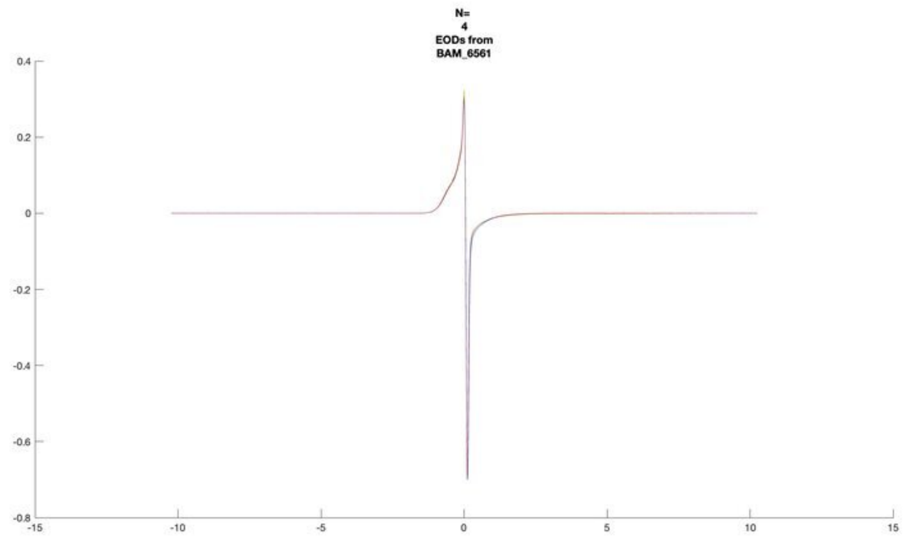
N=
4
EODs from
BAM_6561

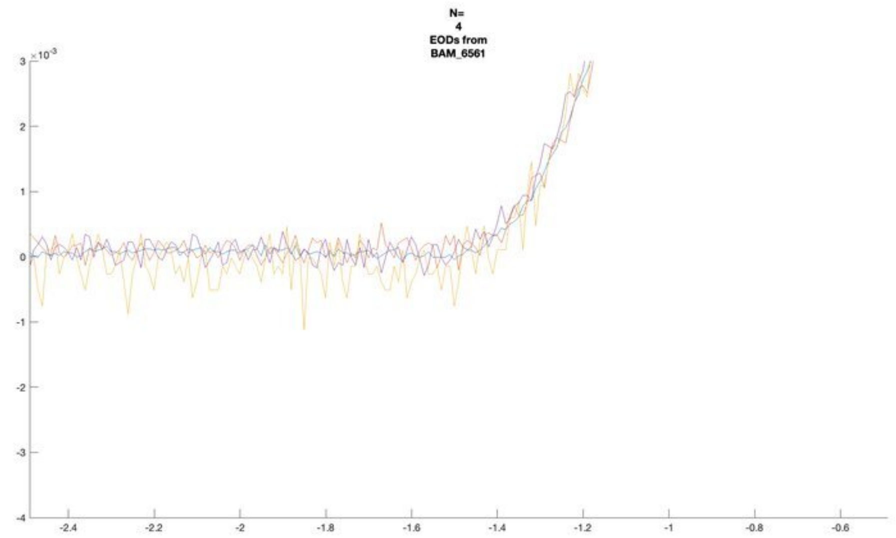
N=
4
EODs from
BAM_6561

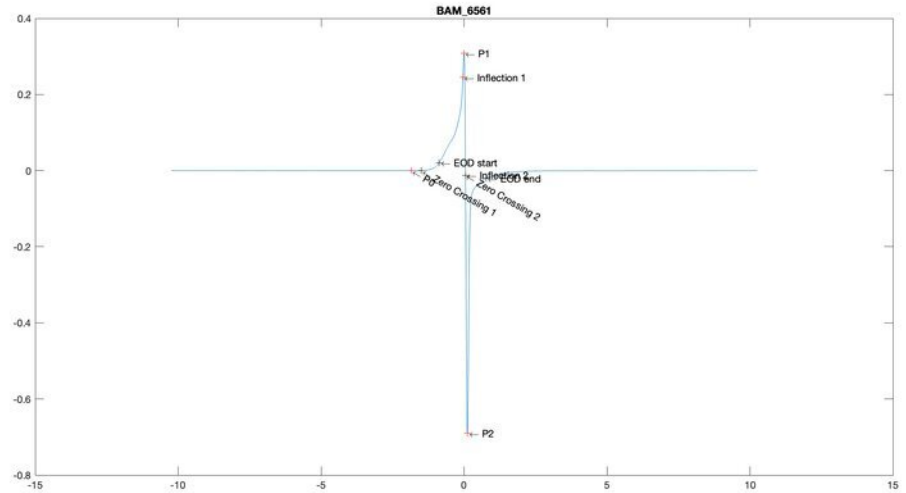
BAM_6561
P1
Inflection 1
EOD start
Inflection 2
EOD end
P0
Zero Crossing 1
Zero Crossing 2
P2

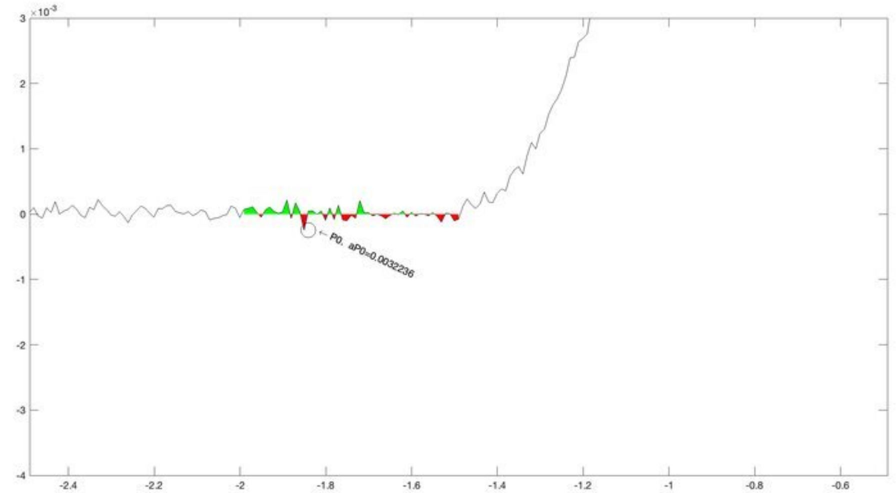
P0, aP0=0.0032236

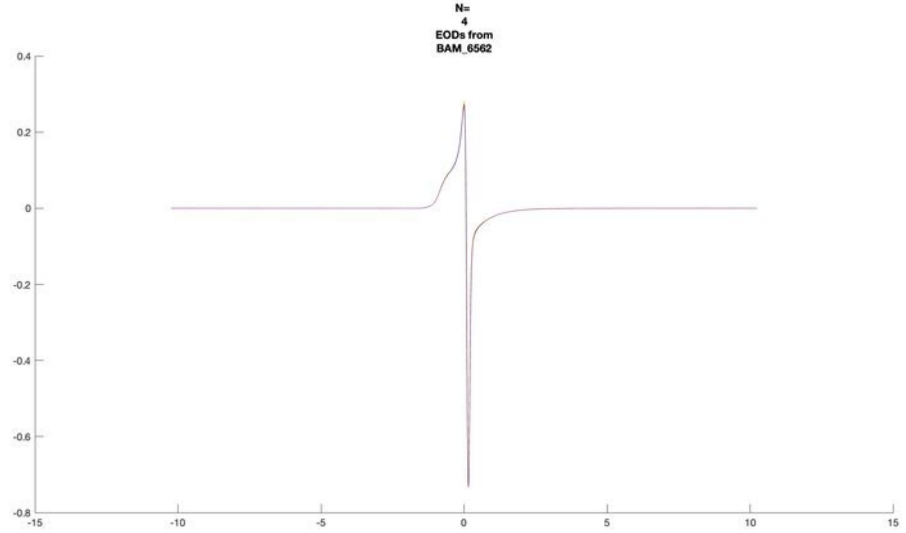
N=
4
EODs from
BAM_6562

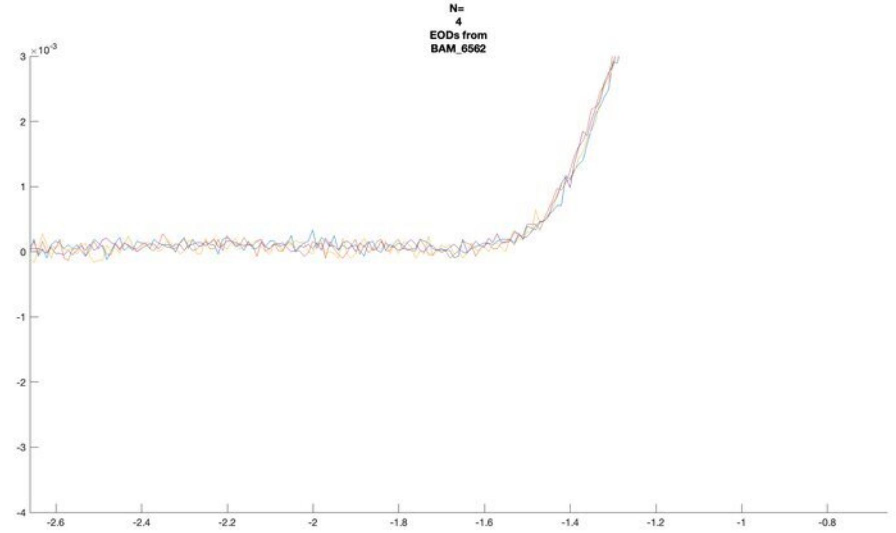
N=
4
EODs from
BAM_6562

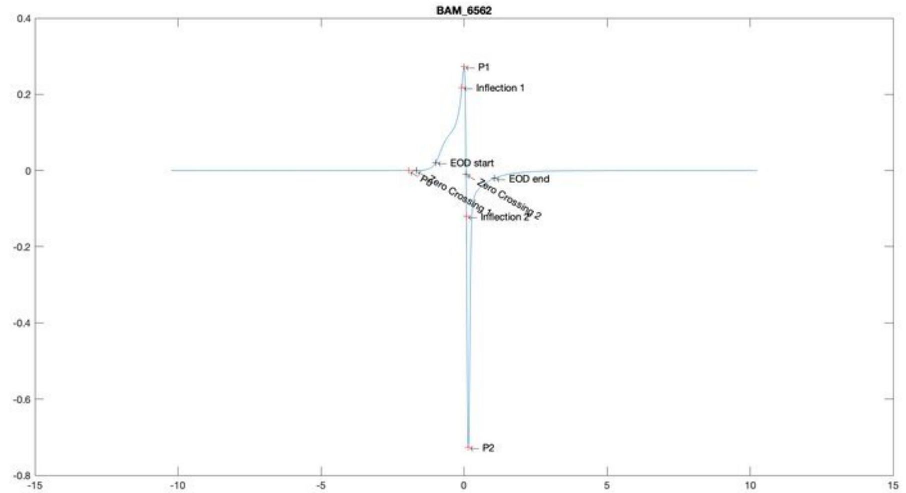
BAM_6562
P1
Inflection 1
EOD start
P0
Zero Crossing 1
Zero Crossing 2
EOD end
Inflection 2
P2

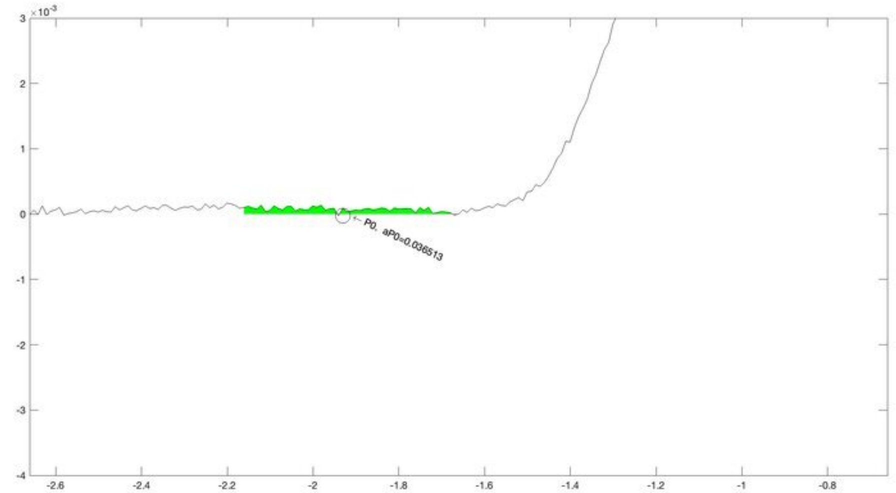
P0, aP0=0.036513

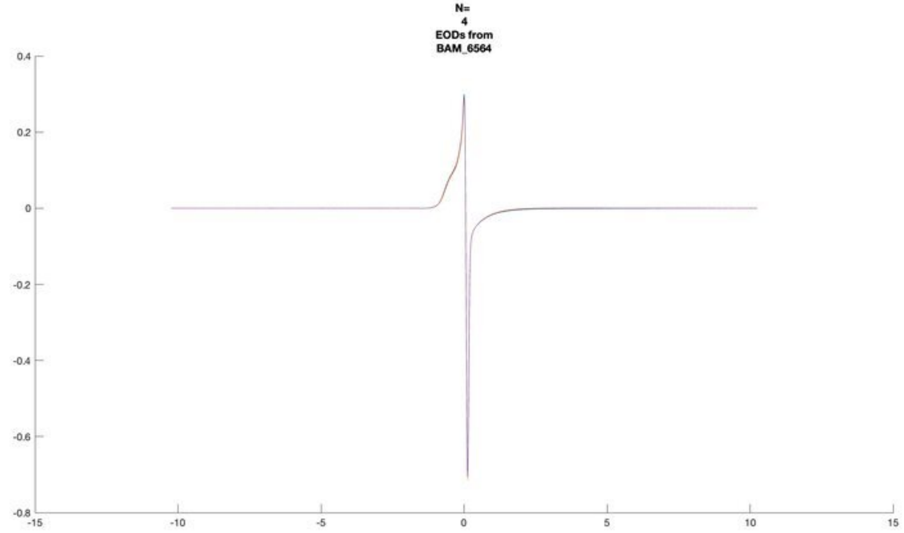
N=
4
EODs from
BAM_6564

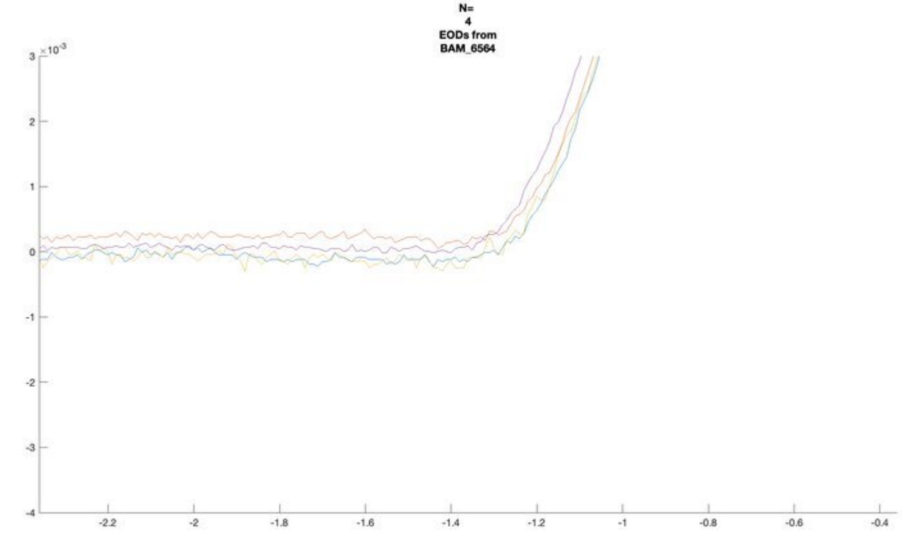
N=
4
EODs from
BAM_6564

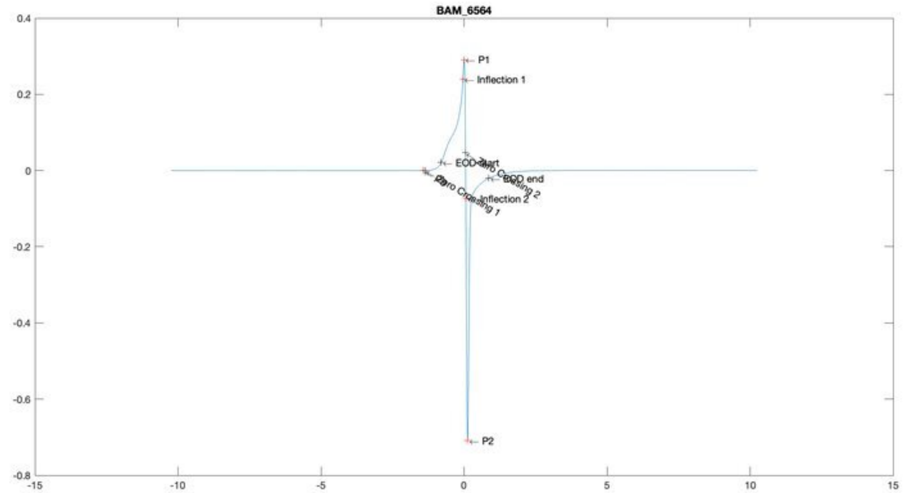
BAM_6564
P1
Inflection 1
EOD start
Zero Crossing 2
EOD end
Inflection 2
Zero Crossing 1
P2

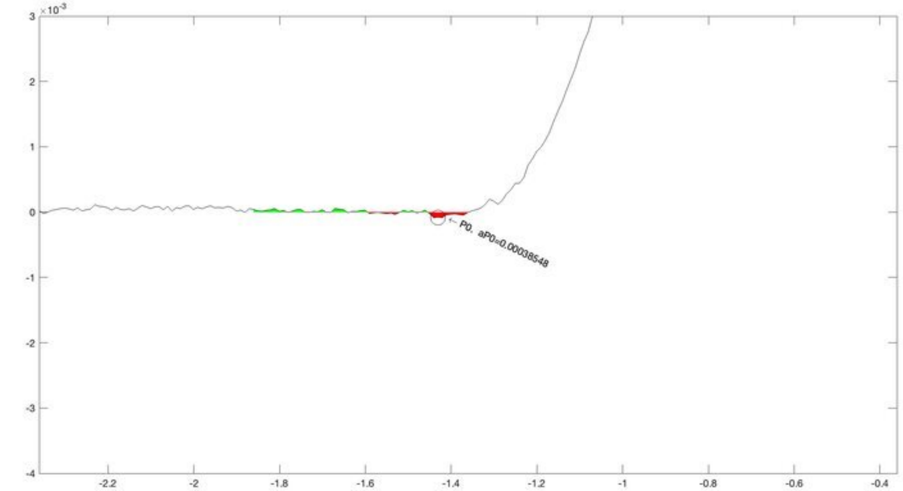
P0, aP0=0.00038548

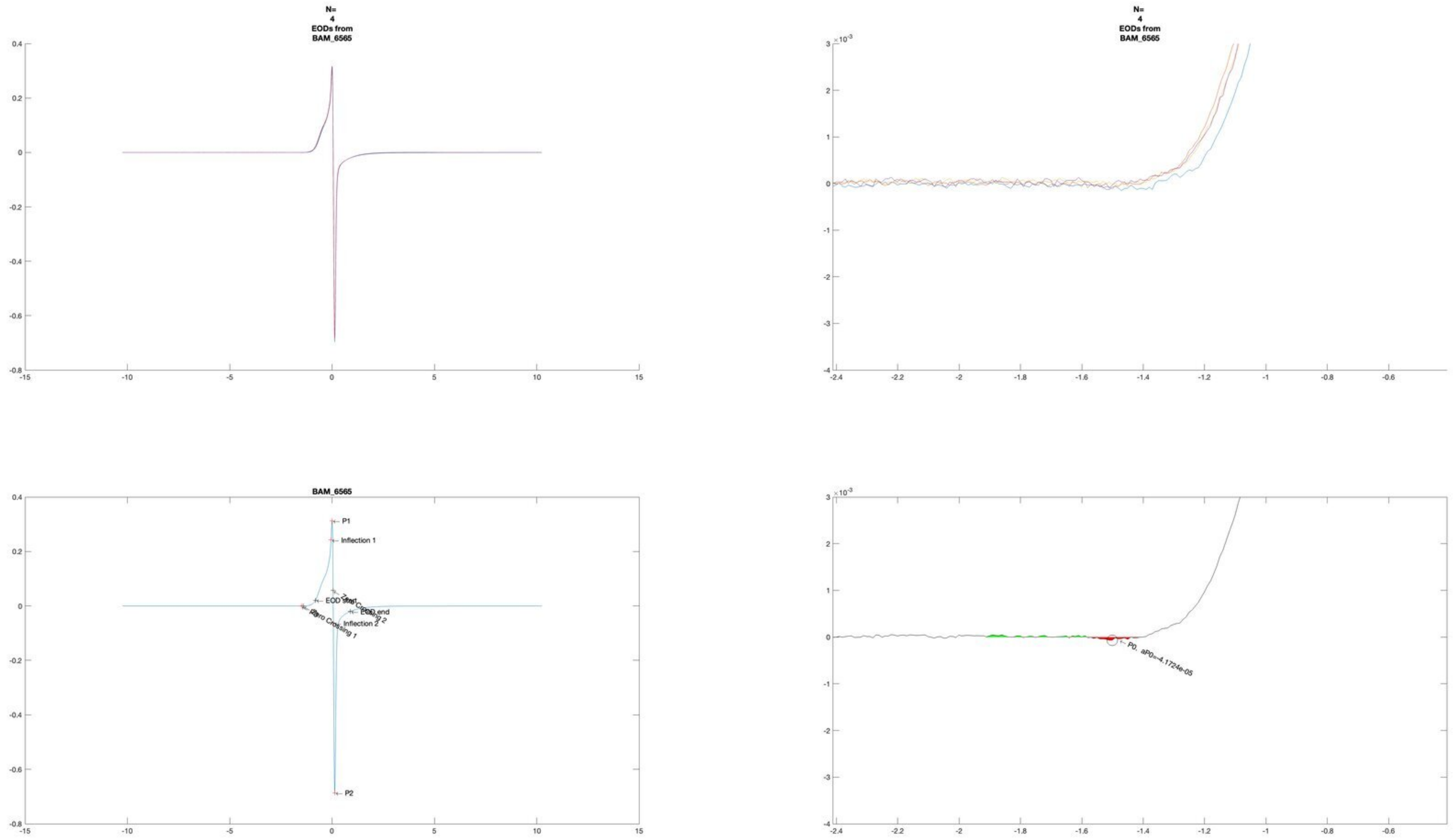

N=
4
EODs from
BAM_6565
BAM_6565
P1
Inflection 1
EOD start
Zero Crossing 1
Zero Crossing 2
EOD end
Inflection 2
P2
P0, aP0=-4.1724e-05

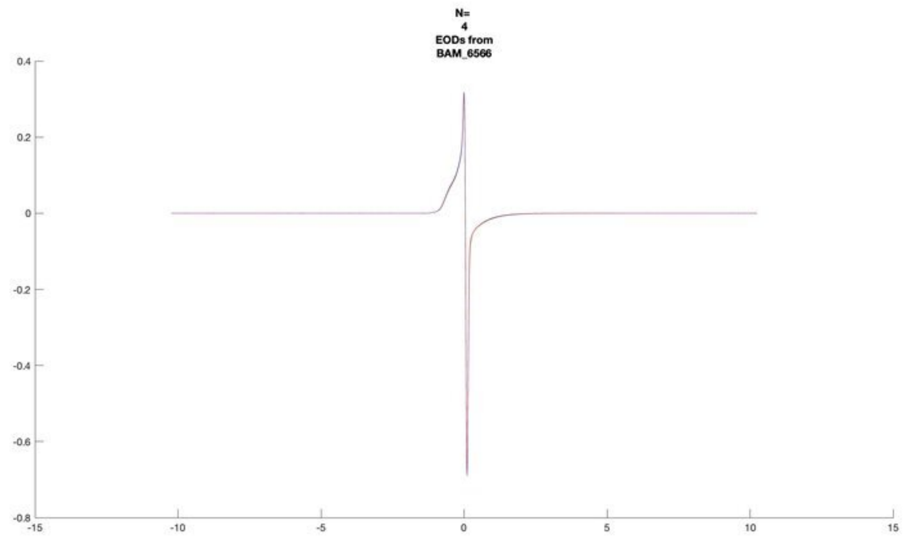
N=
4
EODs from
BAM_6566

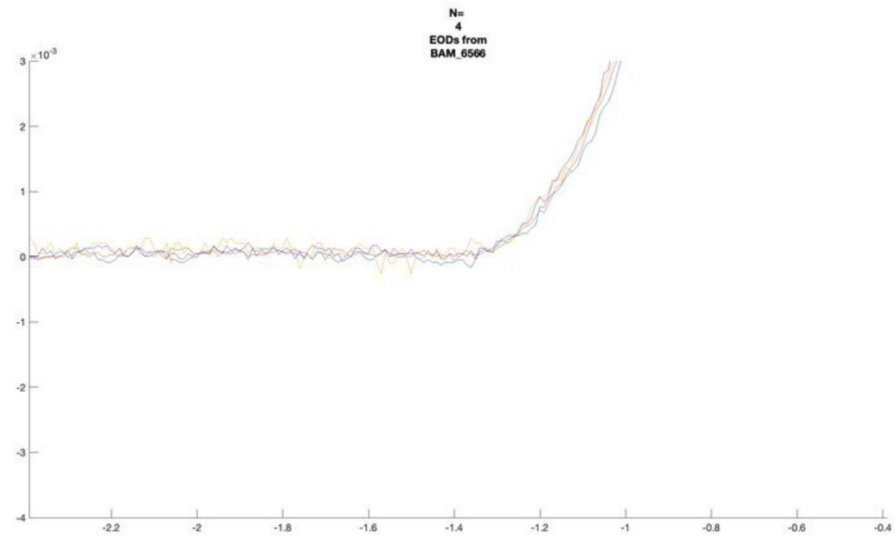
N=
4
EODs from
BAM_6566

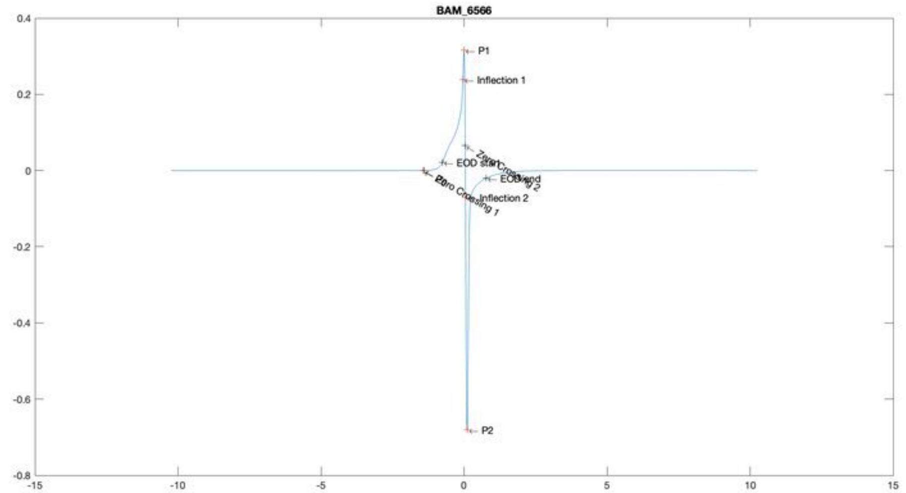
BAM_6566
P1
Inflection 1
EOD start
Zero Crossing 2
EOD end
Inflection 2
P0
Zero Crossing 1
P2

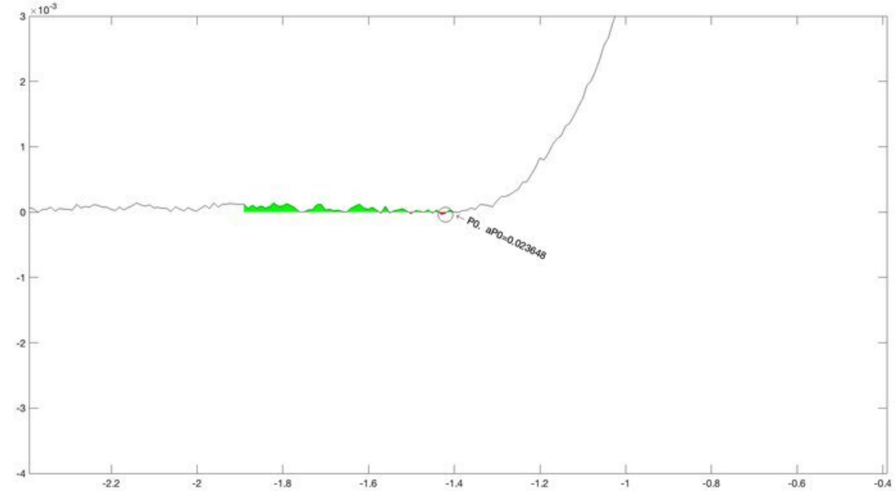
P0, aP0=0.023648

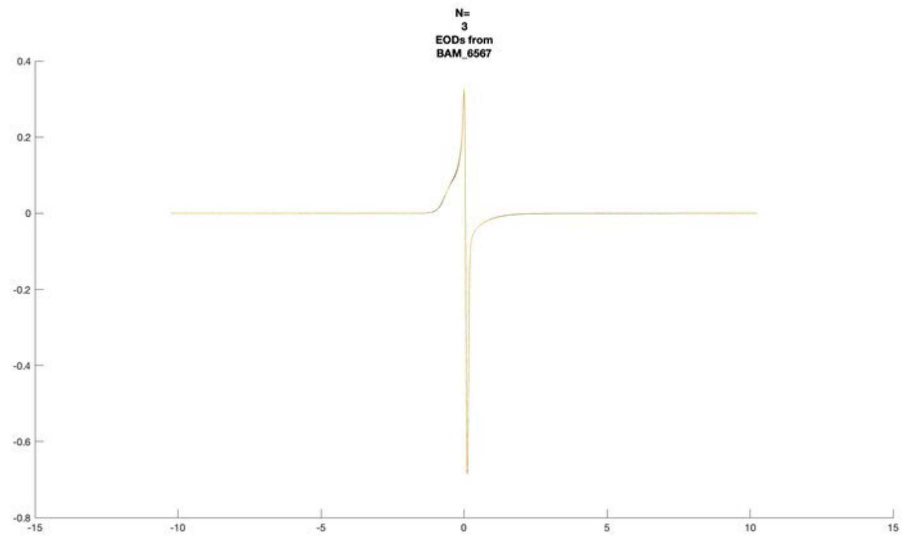
N=
3
EODs from
BAM_6567

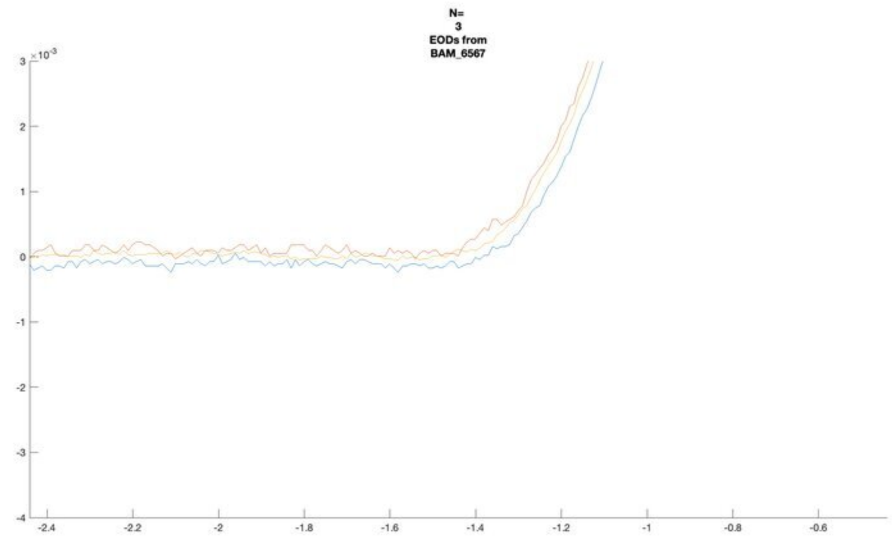
N=
3
EODs from
BAM_6567

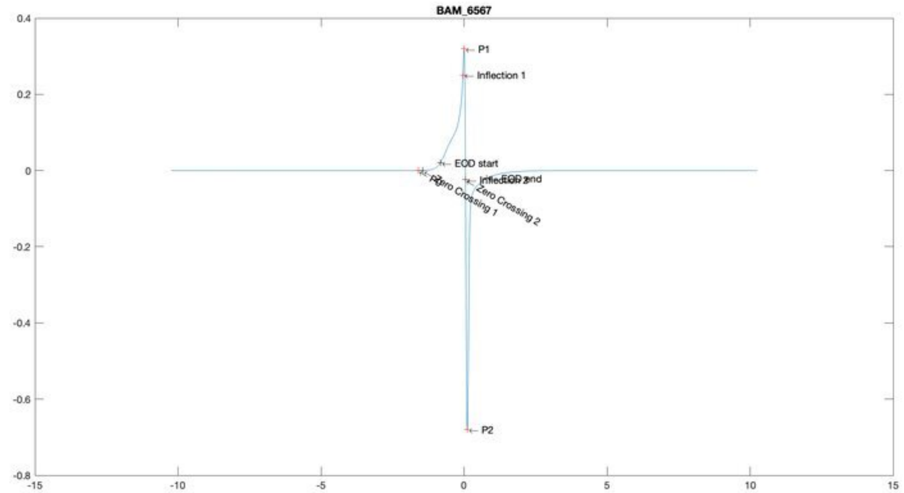
BAM_6567
P1
Inflection 1
EOD start
EOD end
Zero Crossing 1
Zero Crossing 2
P2

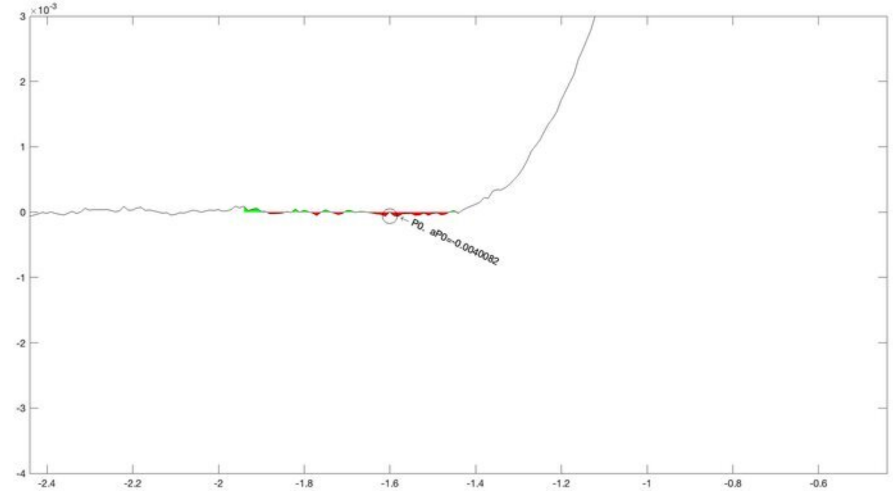
P0, aP0=-0.0040082

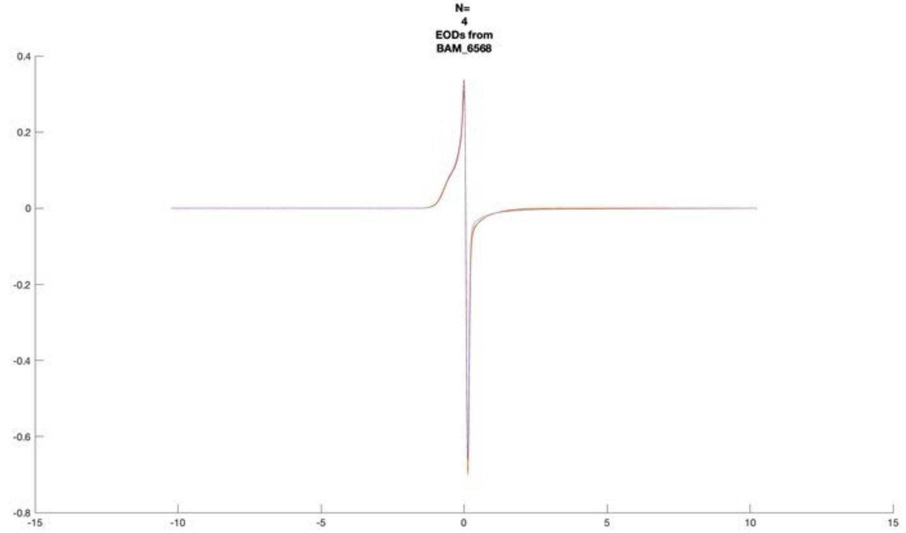
N=
4
EODs from
BAM_6568

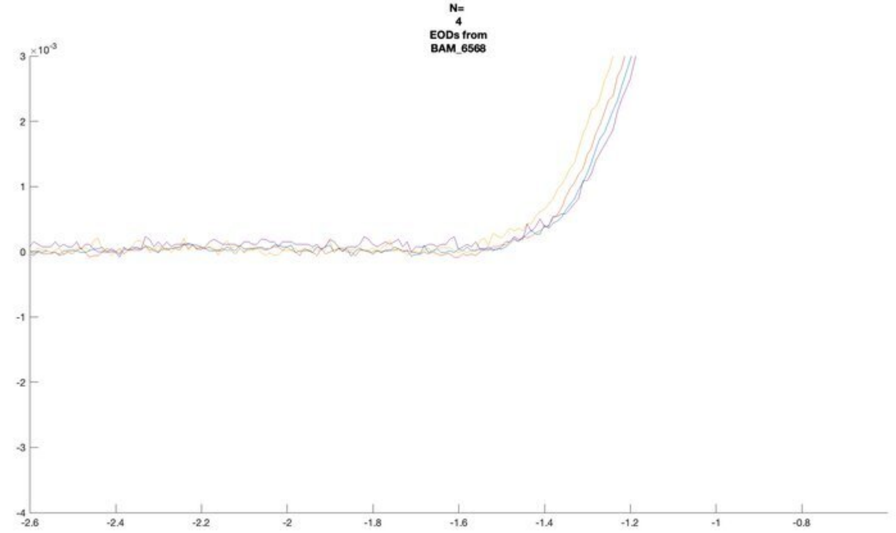
N=
4
EODs from
BAM_6568
×10⁻³

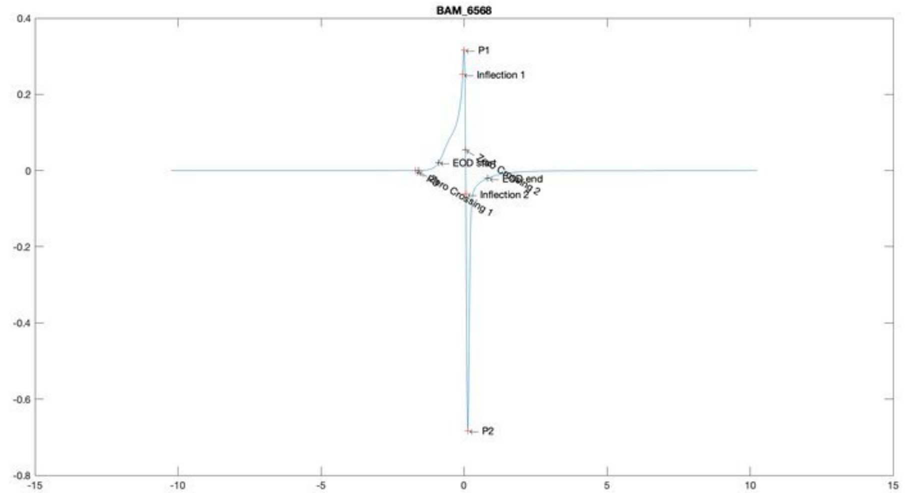
BAM_6568
P1
Inflection 1
EOD start
Zero Crossing 2
EOD end
Inflection 2
Zero Crossing 1
P2

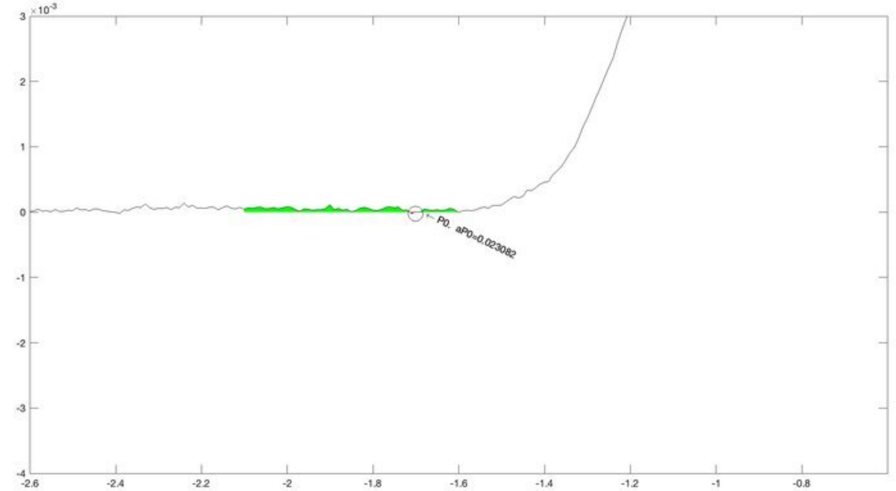
×10⁻³
P0, aP0=0.023082

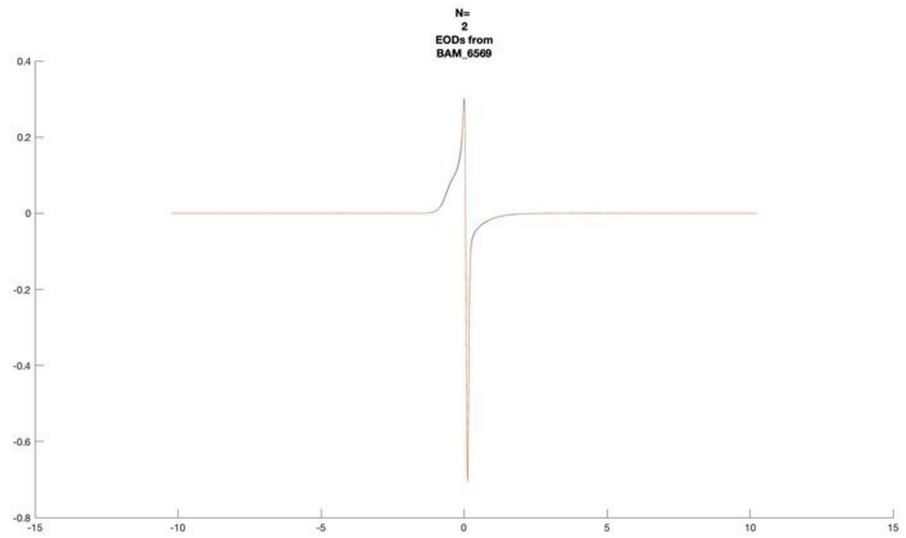
N=
2
EODs from
BAM_6569

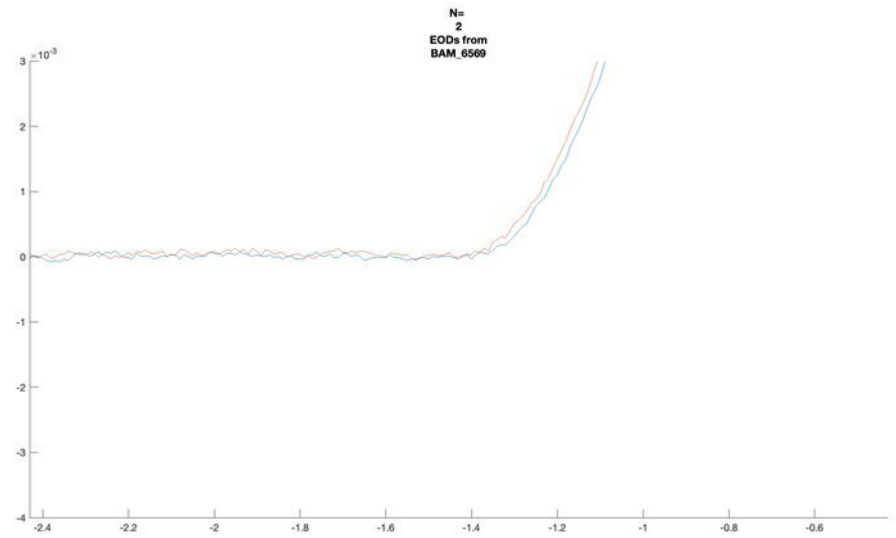
N=
2
EODs from
BAM_6569
×10^-3

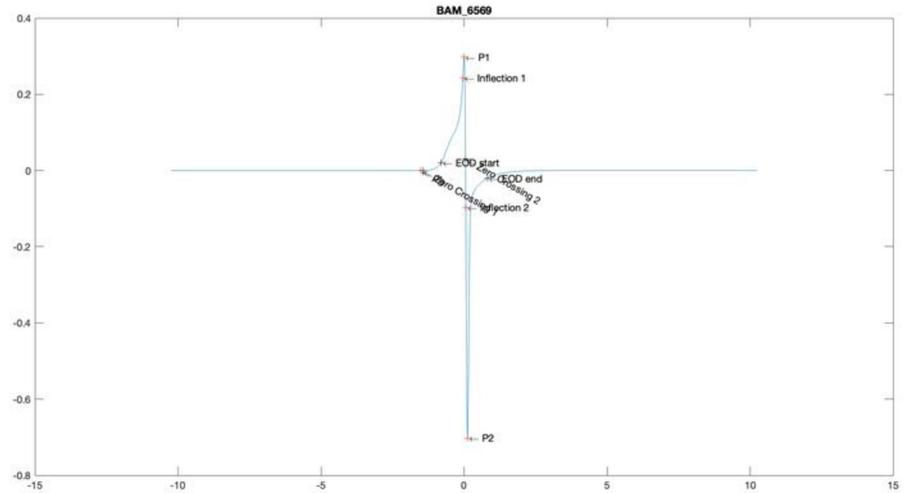
BAM_6569
P1
Inflection 1
EOD start
EOD end
P2

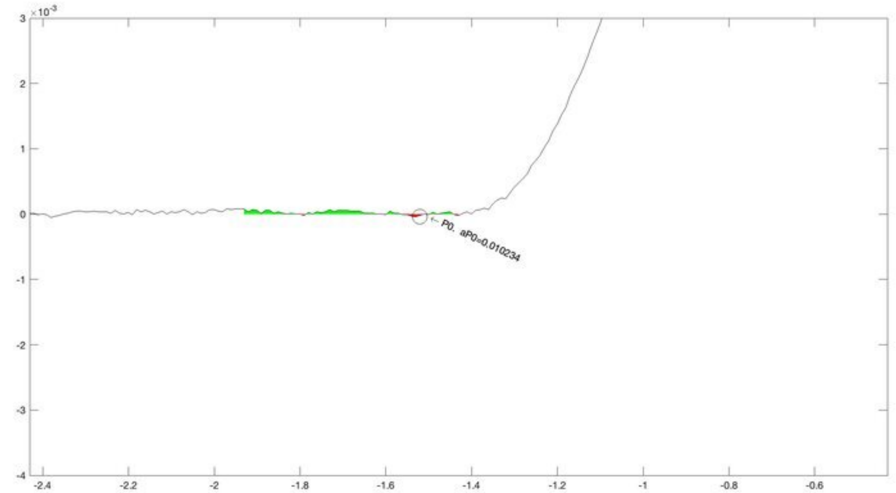
×10^-3
P0, aP0=0.010234

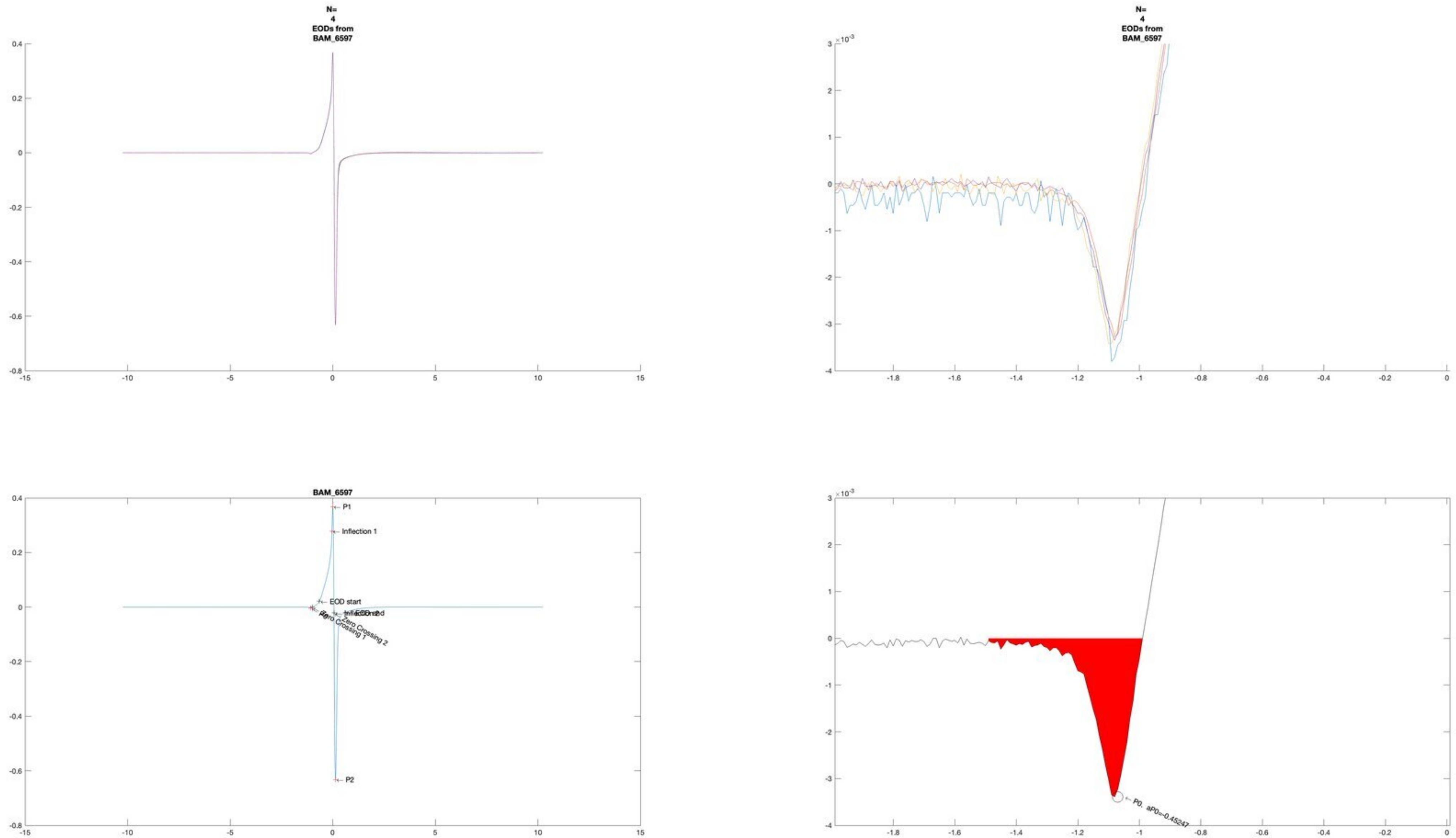
N=
4
EODs from
BAM_6597
N=
4
EODs from
BAM_6597
BAM_6597
P1
Inflection 1
EOD start
Zero Crossing 1
Zero Crossing 2
P2
P0, aP0=-0.45247

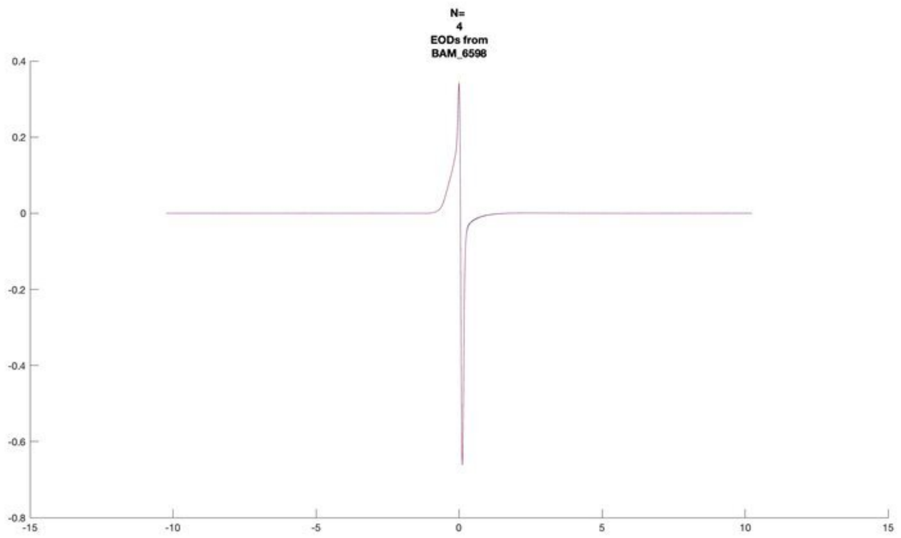
N=
4
EODs from
BAM_6598
0.4
0.2
0
-0.2
-0.4
-0.6
-0.8
-15
-10
-5
0
5
10
15

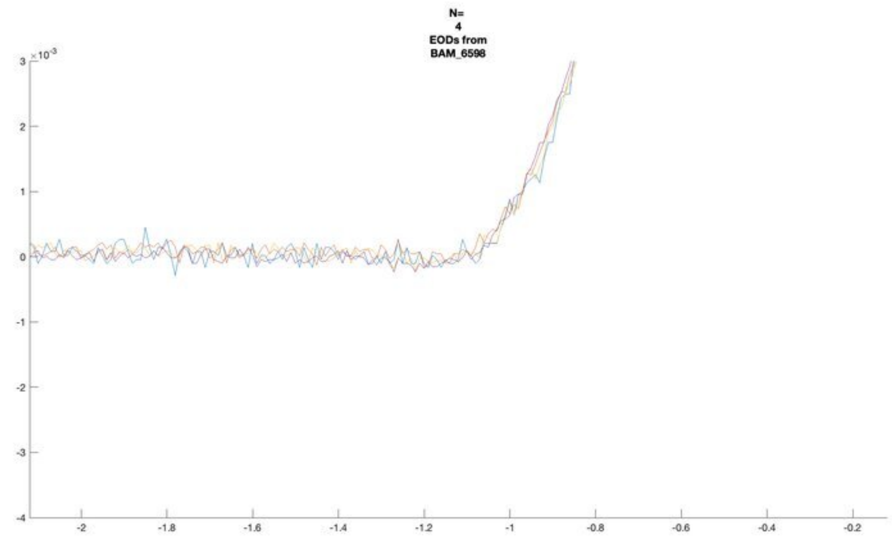
N=
4
EODs from
BAM_6598
×10^-3
3
2
1
0
-1
-2
-3
-4
-2
-1.8
-1.6
-1.4
-1.2
-1
-0.8
-0.6
-0.4
-0.2

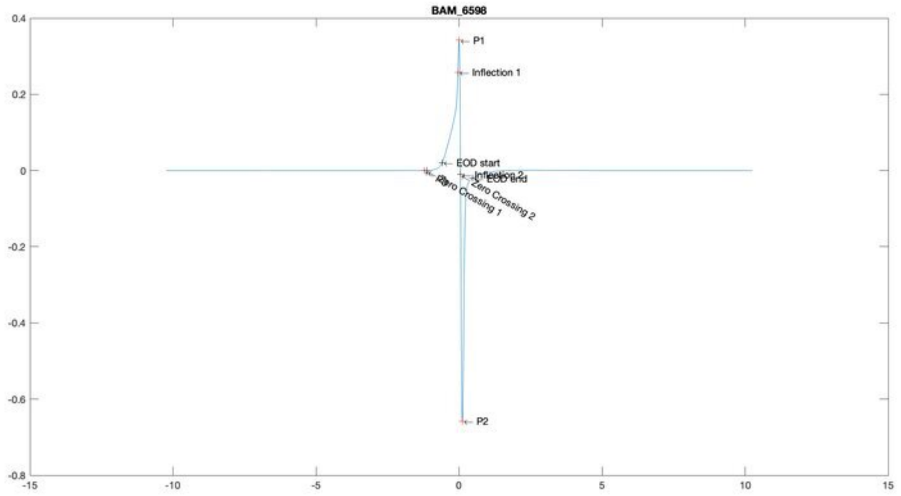
BAM_6598
P1
Inflection 1
EOD start
Inflection 2
EOD end
Zero Crossing 2
Zero Crossing 1
P2
0.4
0.2
0
-0.2
-0.4
-0.6
-0.8
-15
-10
-5
0
5
10
15

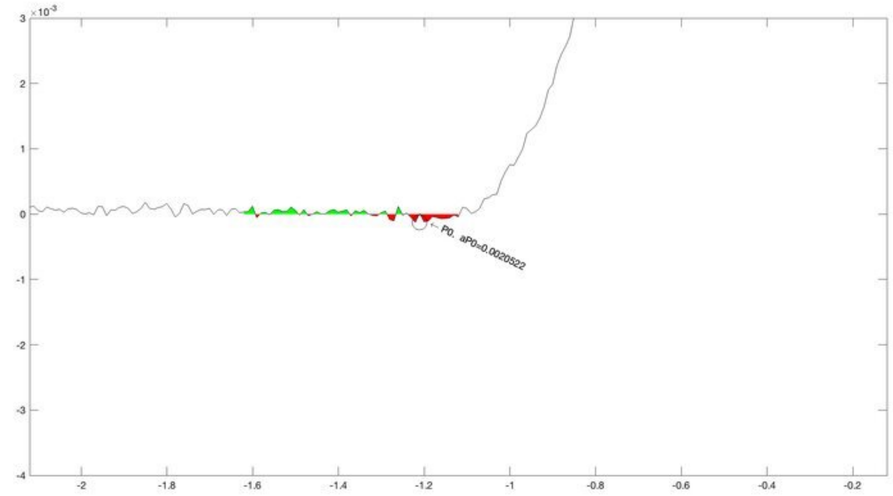
×10^-3
P0, aP0=0.0020522
3
2
1
0
-1
-2
-3
-4
-2
-1.8
-1.6
-1.4
-1.2
-1
-0.8
-0.6
-0.4
-0.2

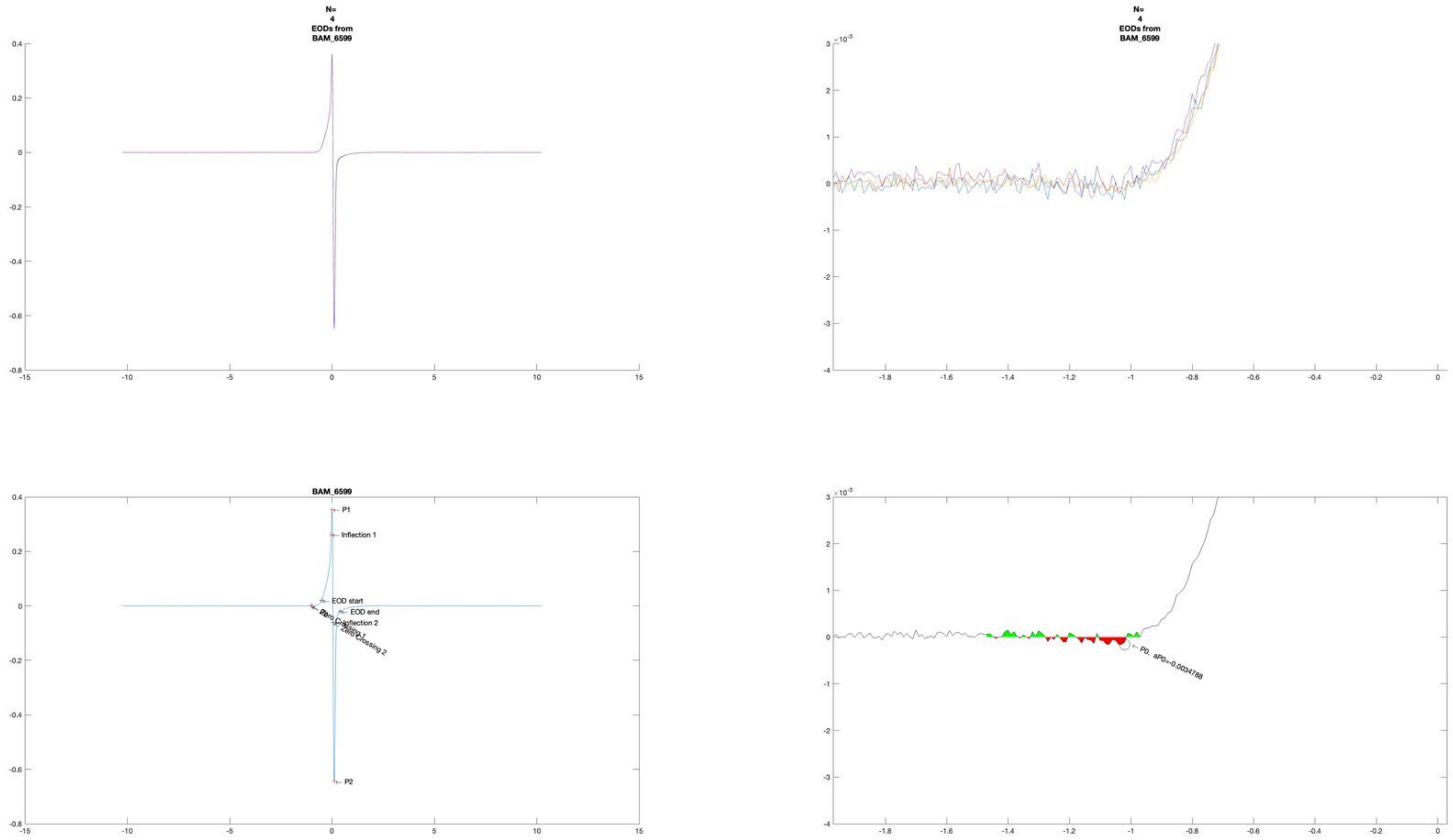
N=
4
EODs from
BAM_6599
N=
4
EODs from
BAM_6599
BAM_6599
P1
Inflection 1
EOD start
EOD end
Inflection 2
Zero Crossing 1
Zero Crossing 2
P0. aP0=-0.0034788

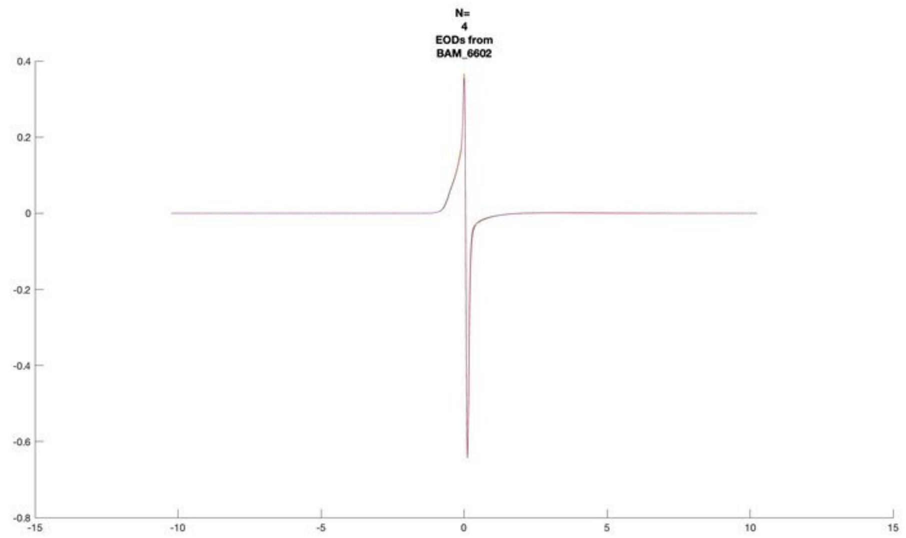
N=
4
EODs from
BAM_6602

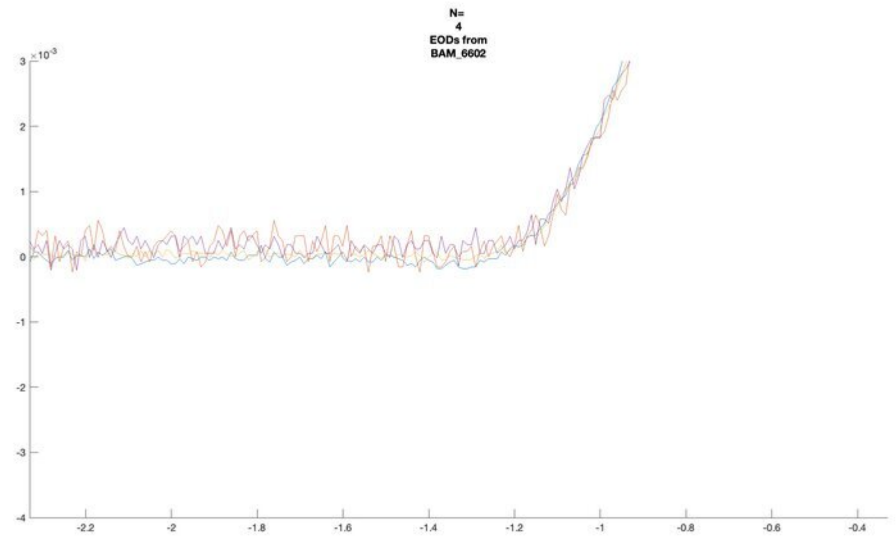
N=
4
EODs from
BAM_6602

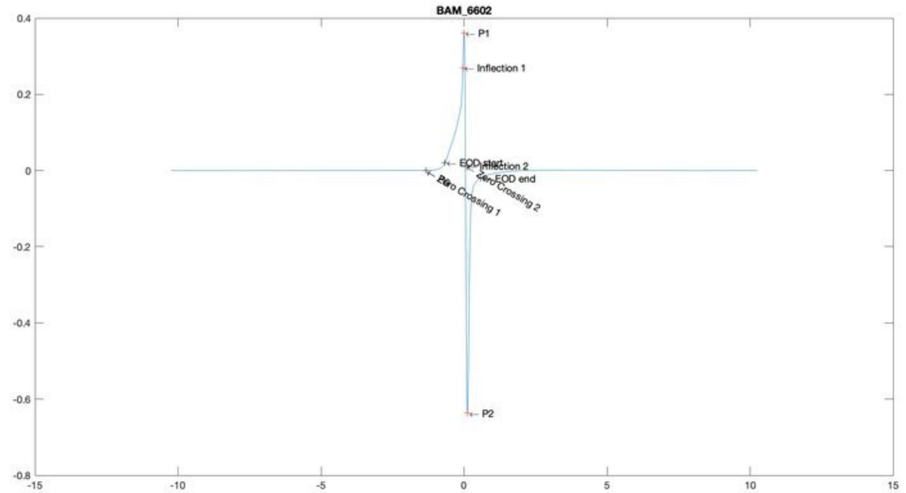
BAM_6602
P1
Inflection 1
EOD start
Inflection 2
EOD end
Zero Crossing 1
Zero Crossing 2
P2

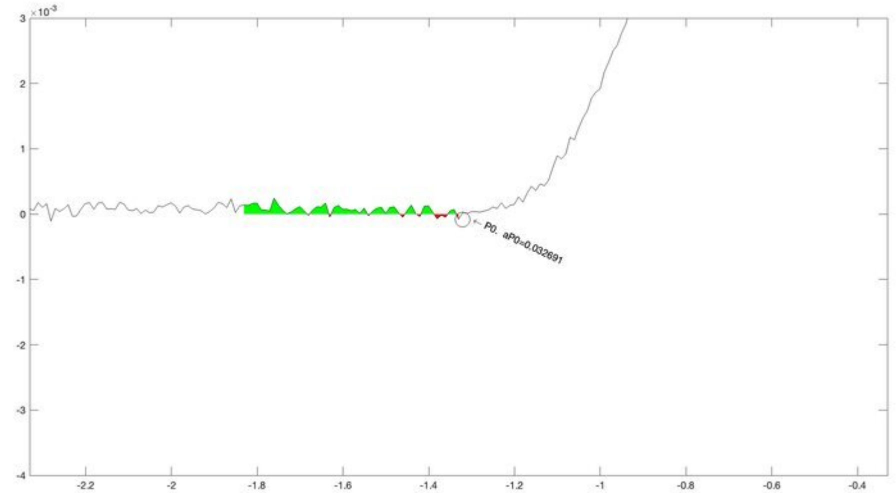
P0, aP0=0.032691

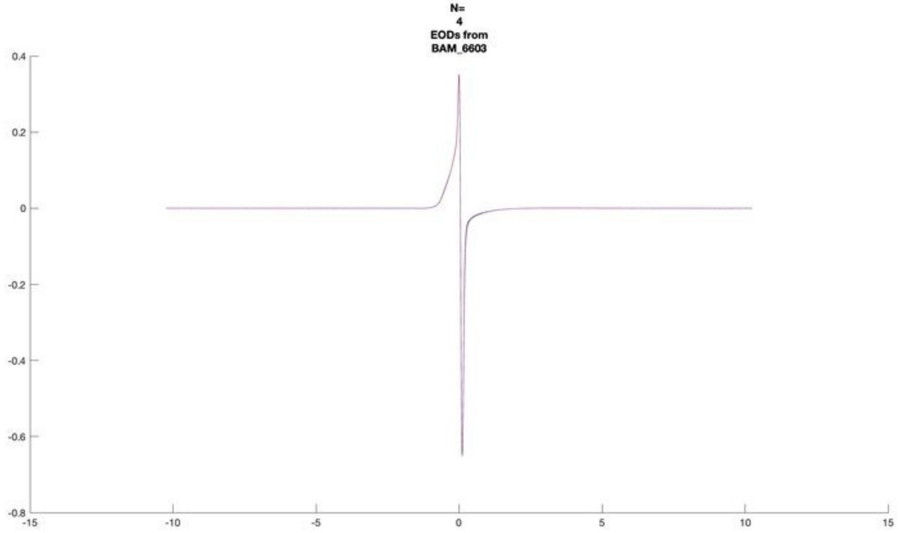
N=
4
EODs from
BAM_6603

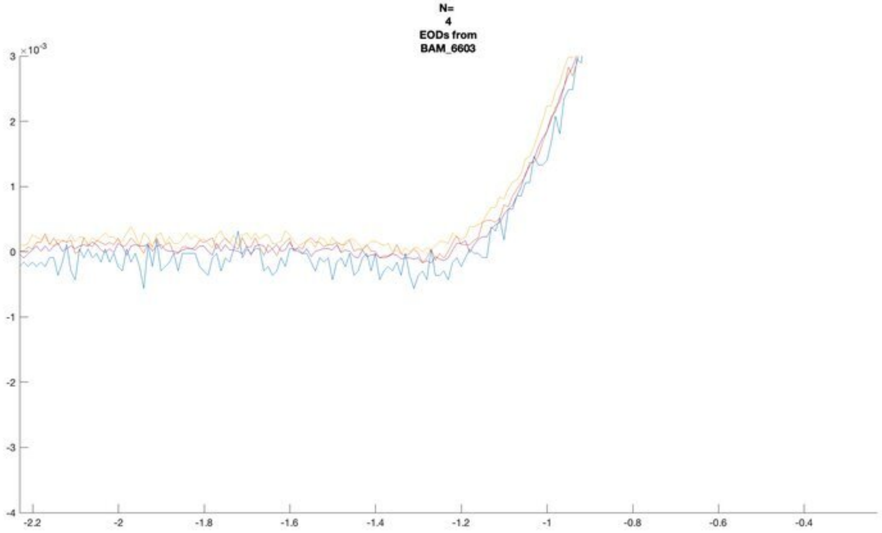
N=
4
EODs from
BAM_6603

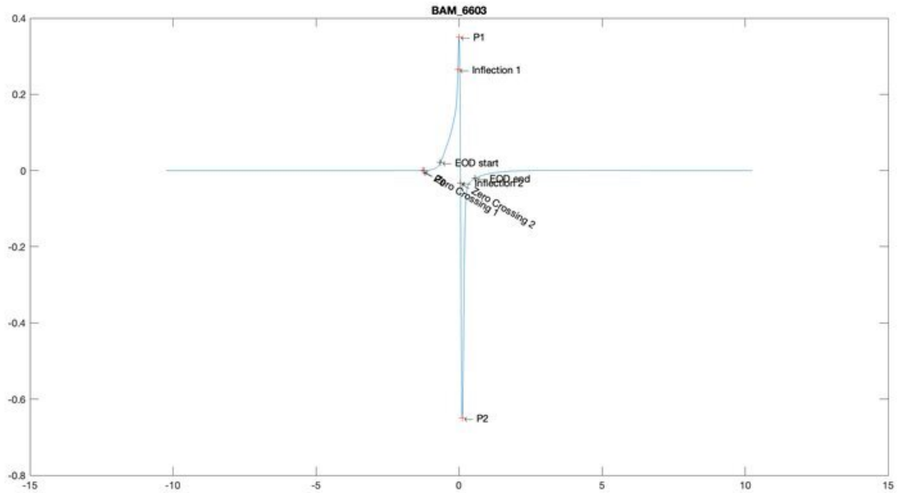
BAM_6603
P1
Inflection 1
EOD start
EOD end
Inflection 2
Zero Crossing 1
Zero Crossing 2
P0
P2

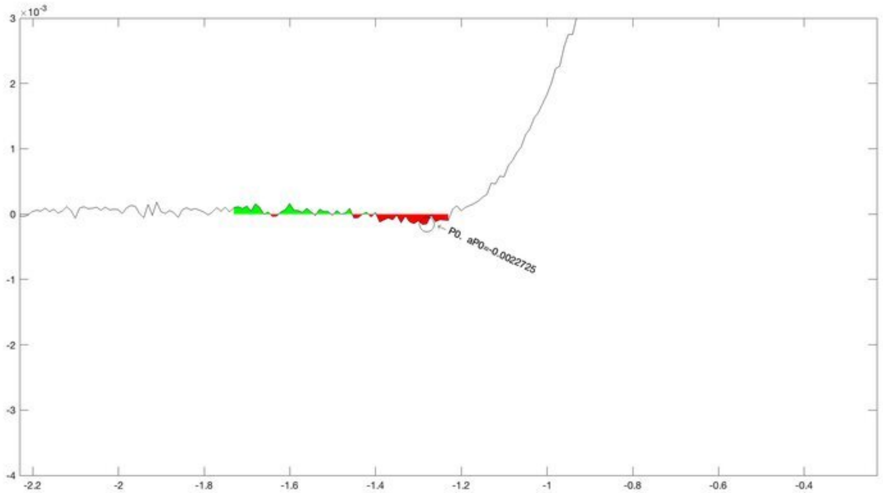
P0. aP0=-0.0022725

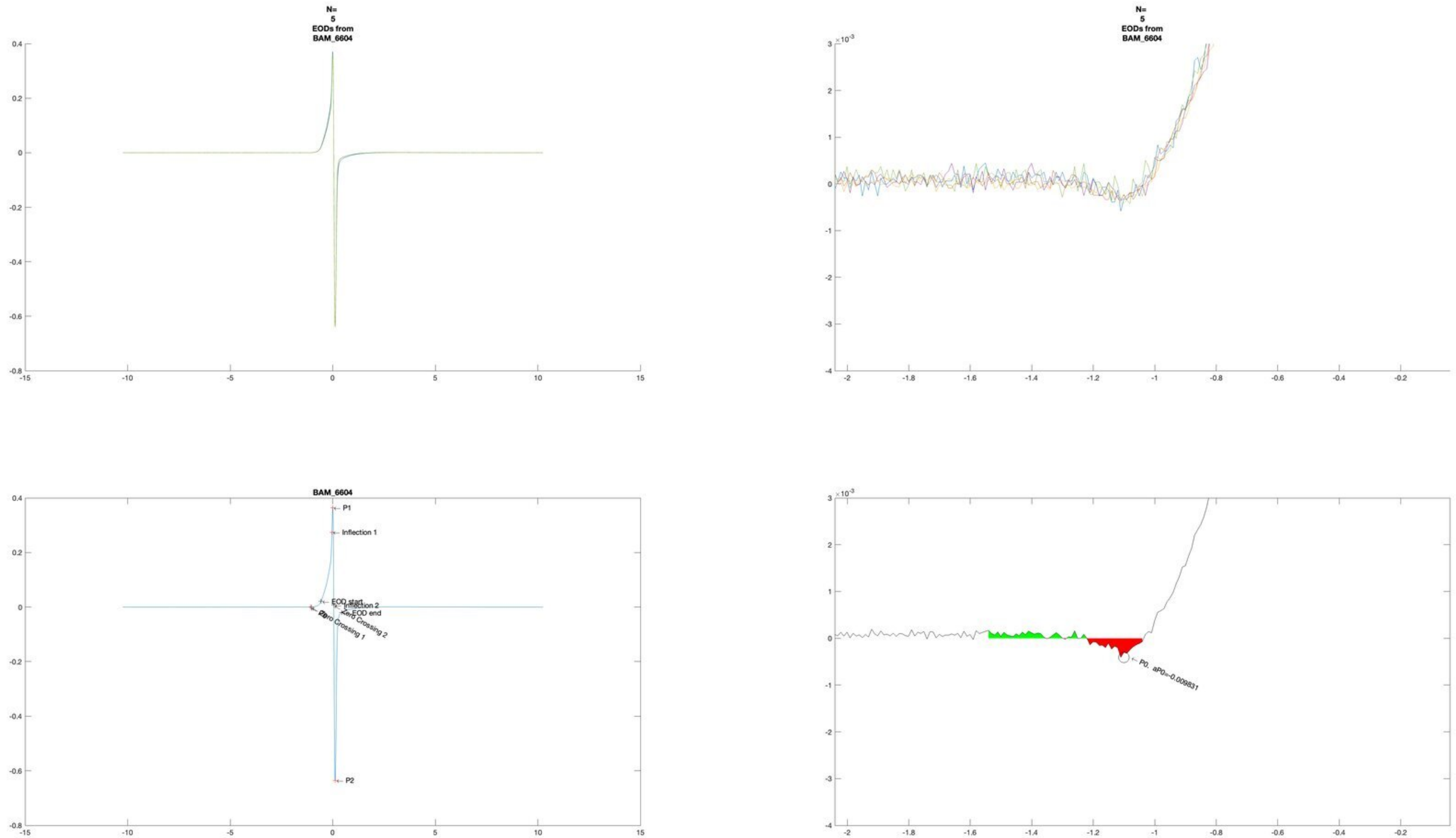
N=
5
EODs from
BAM_6604
N=
5
EODs from
BAM_6604
BAM_6604
P1
Inflection 1
EOD start
Inflection 2
EOD end
Zero Crossing 1
Zero Crossing 2
P2
P0, aP0=-0.009831

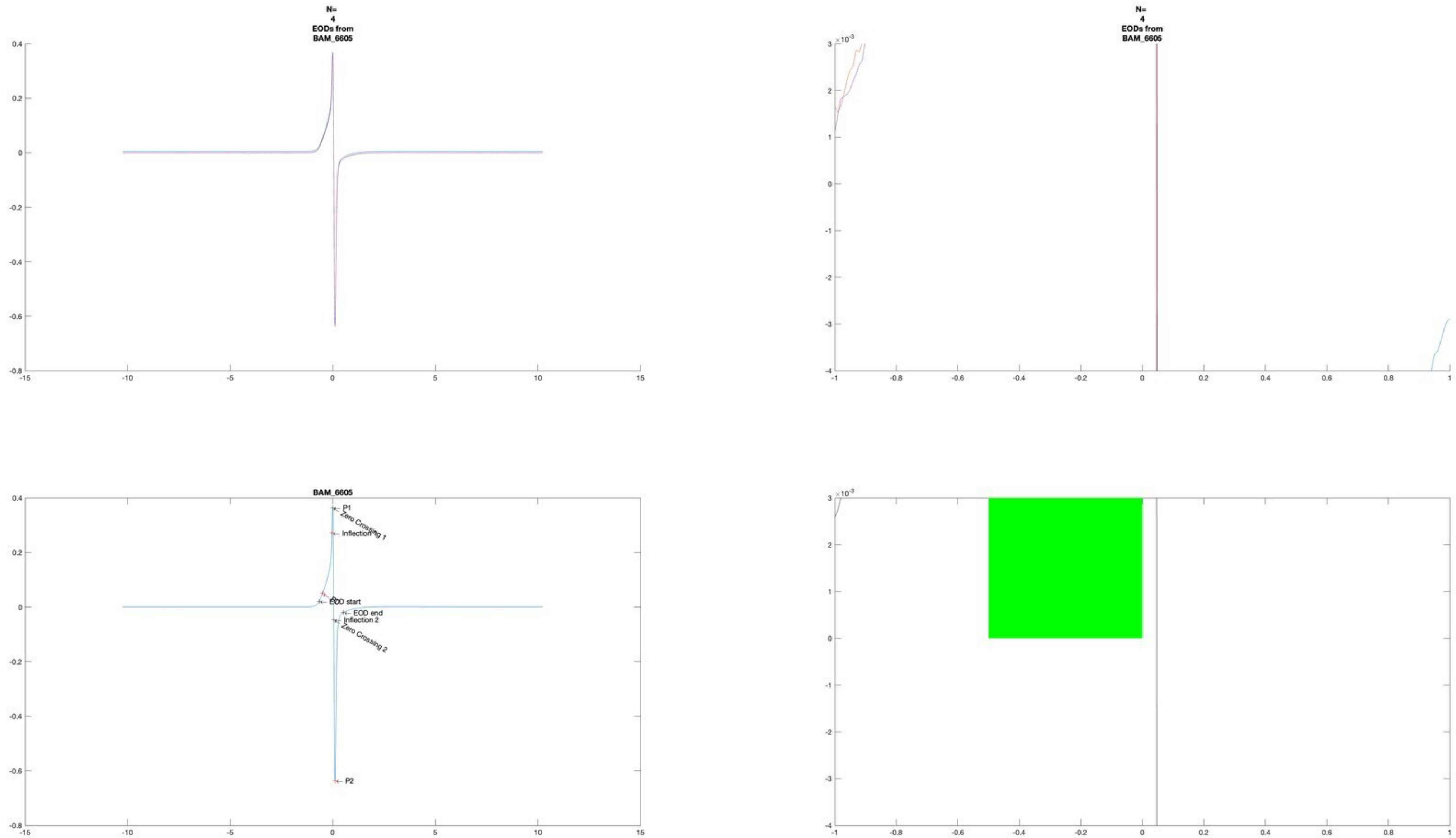

N=
4
EODs from
BAM_6605
N=
4
EODs from
BAM_6605
BAM_6605
P1
Zero Crossing 1
Inflection 1
EOD start
EOD end
Inflection 2
Zero Crossing 2
P2

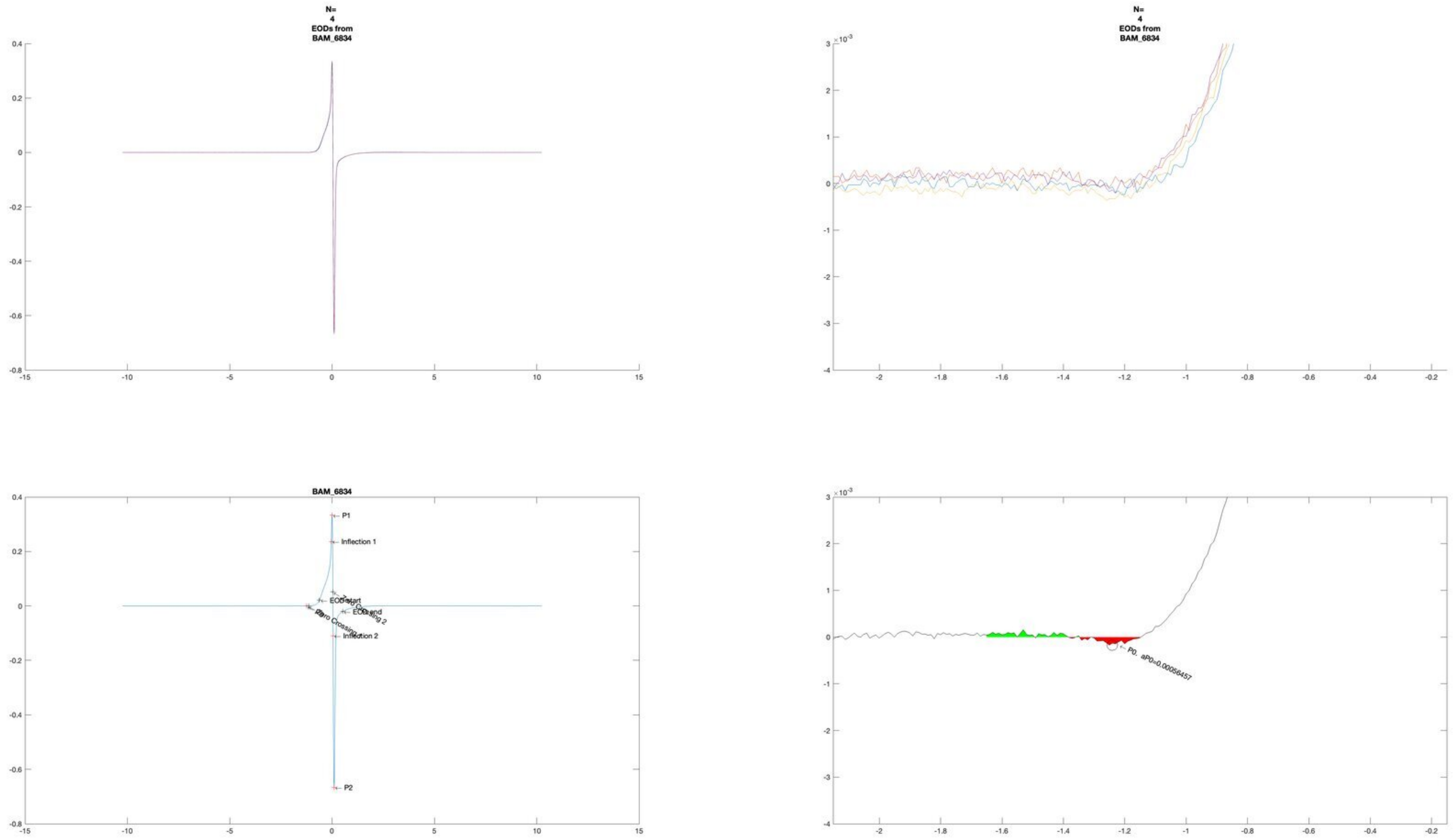

N=
4
EODs from
BAM_6834
N=
4
EODs from
BAM_6834
BAM_6834
P1
Inflection 1
EOD start
EOD end
Zero Crossing
Zero Crossing 2
Inflection 2
P2
P0, aP0=0.00056457

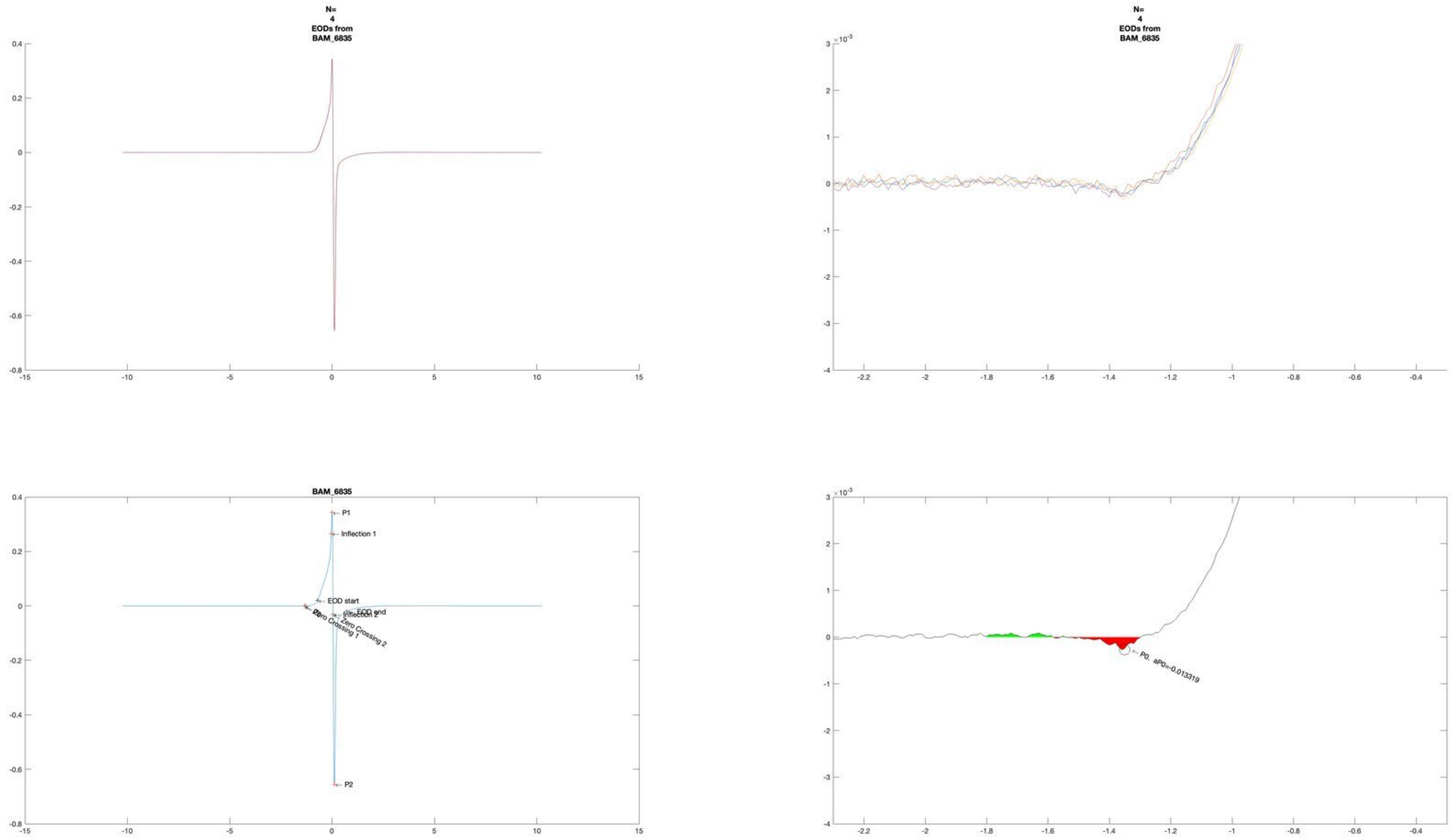

N=
4
EODs from
BAM_6835
N=
4
EODs from
BAM_6835
BAM_6835
P1
Inflection 1
EOD start
EOD end
Inflection 2
Zero Crossing 1
Zero Crossing 2
P2
P0, aP0=-0.013319

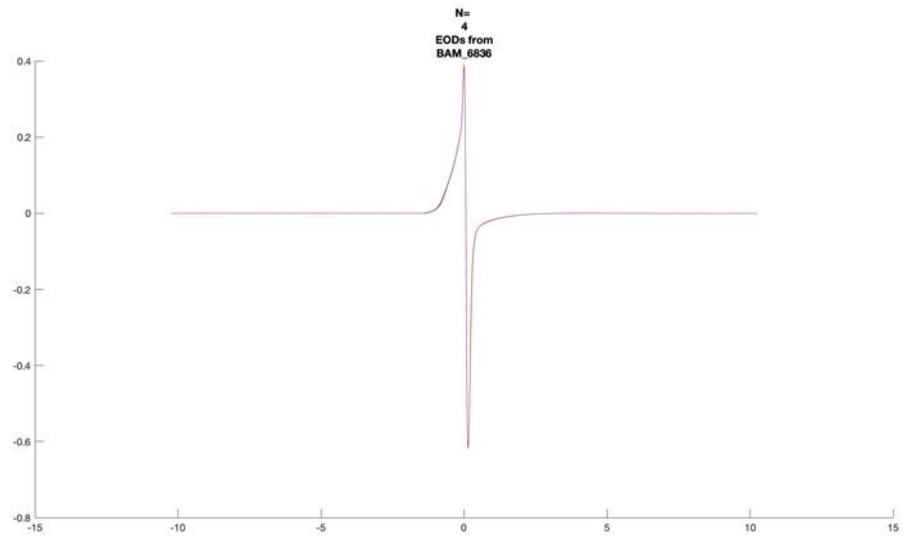
N=
4
EODs from
BAM_6836

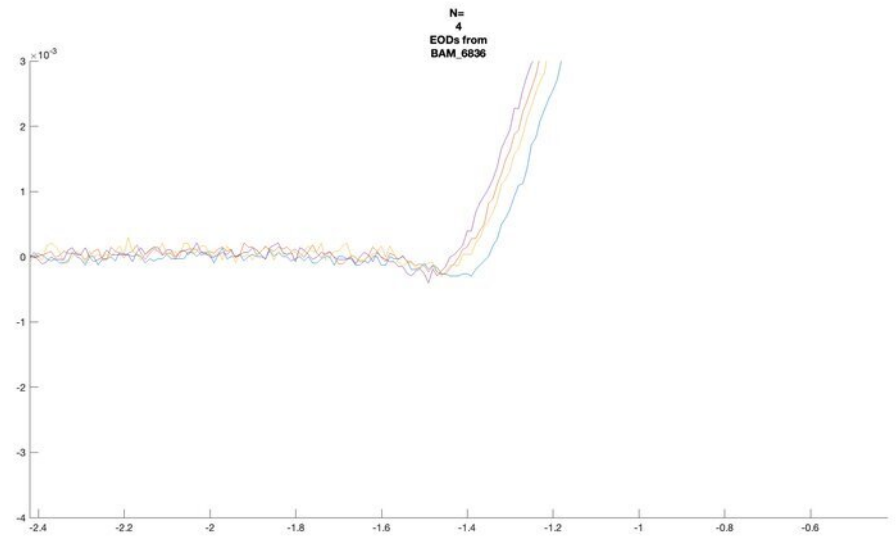
N=
4
EODs from
BAM_6836

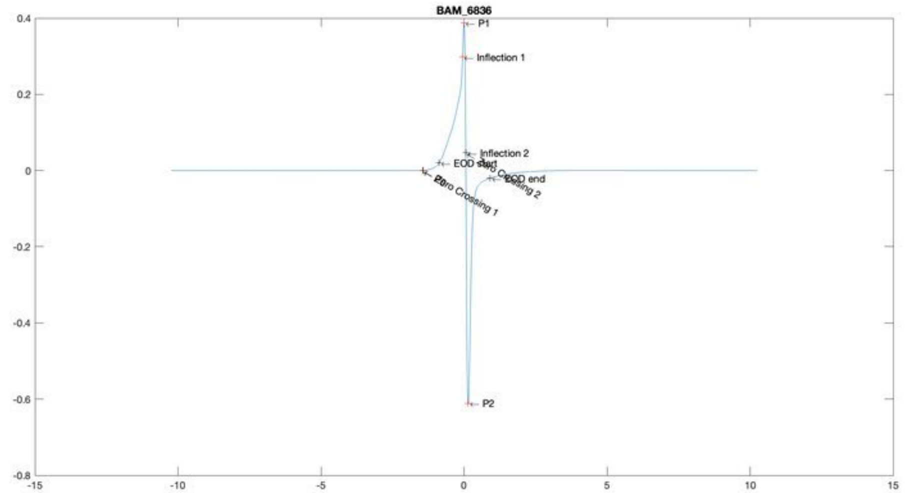
BAM_6836
P1
Inflection 1
Inflection 2
EOD start
EOD end
Zero Crossing 1
Zero Crossing 2
P2

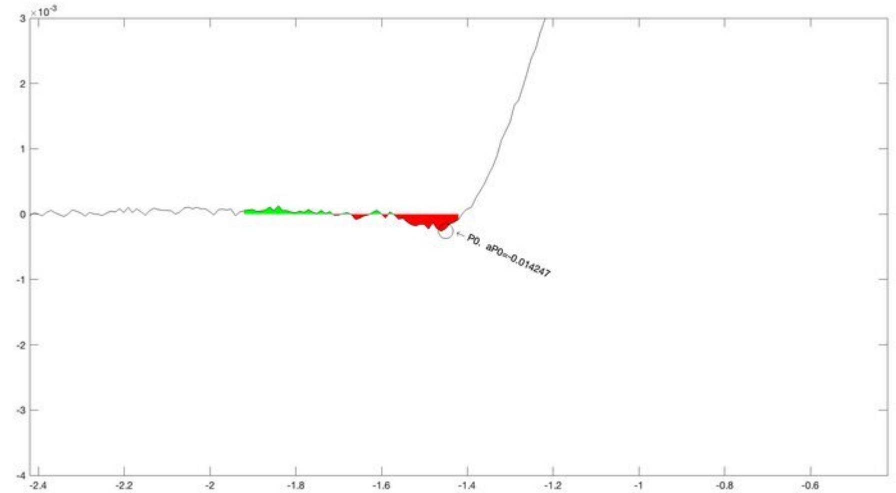
P0, aP0=-0.014247

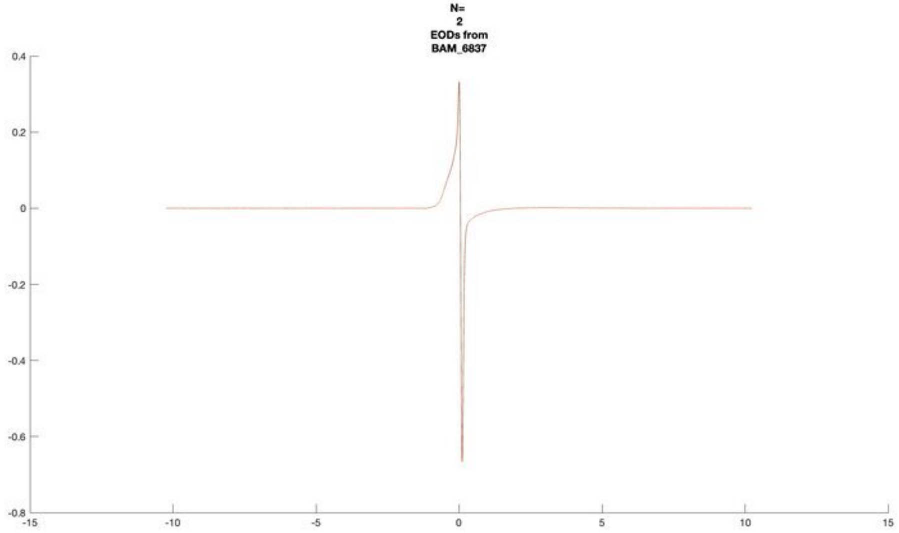
N=
2
EODs from
BAM_6837

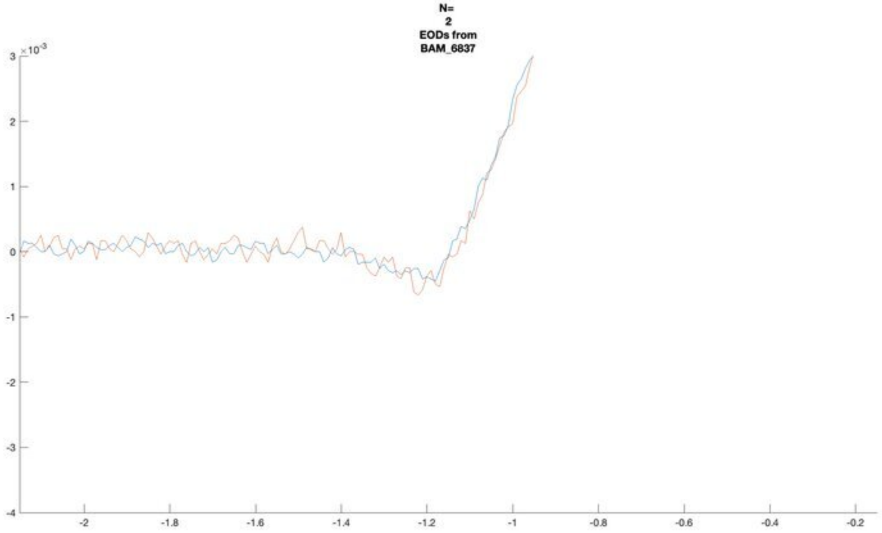
N=
2
EODs from
BAM_6837

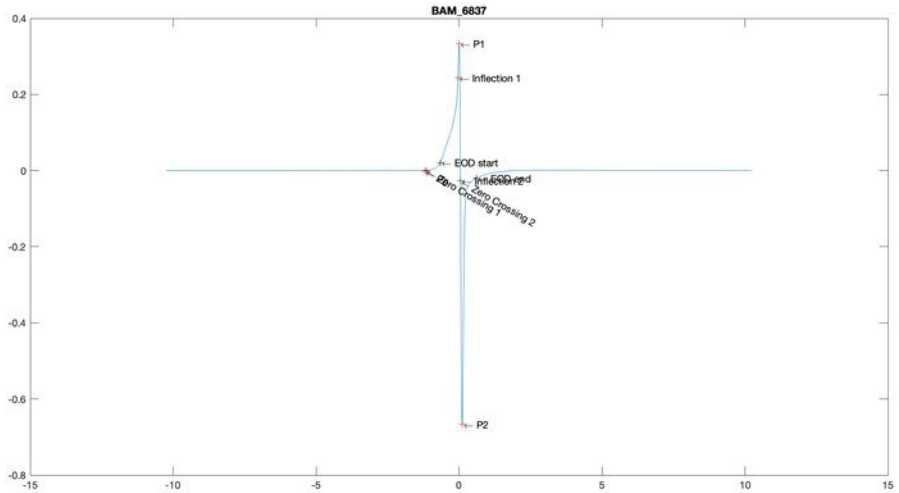
BAM_6837
P1
Inflection 1
EOD start
EOD end
Inflection 2
Zero Crossing 1
Zero Crossing 2
P2

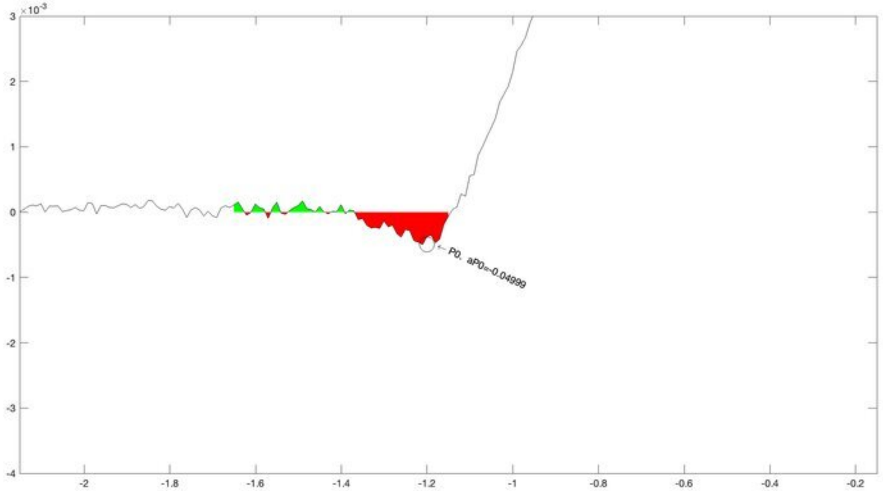
P0, aP0=-0.04999

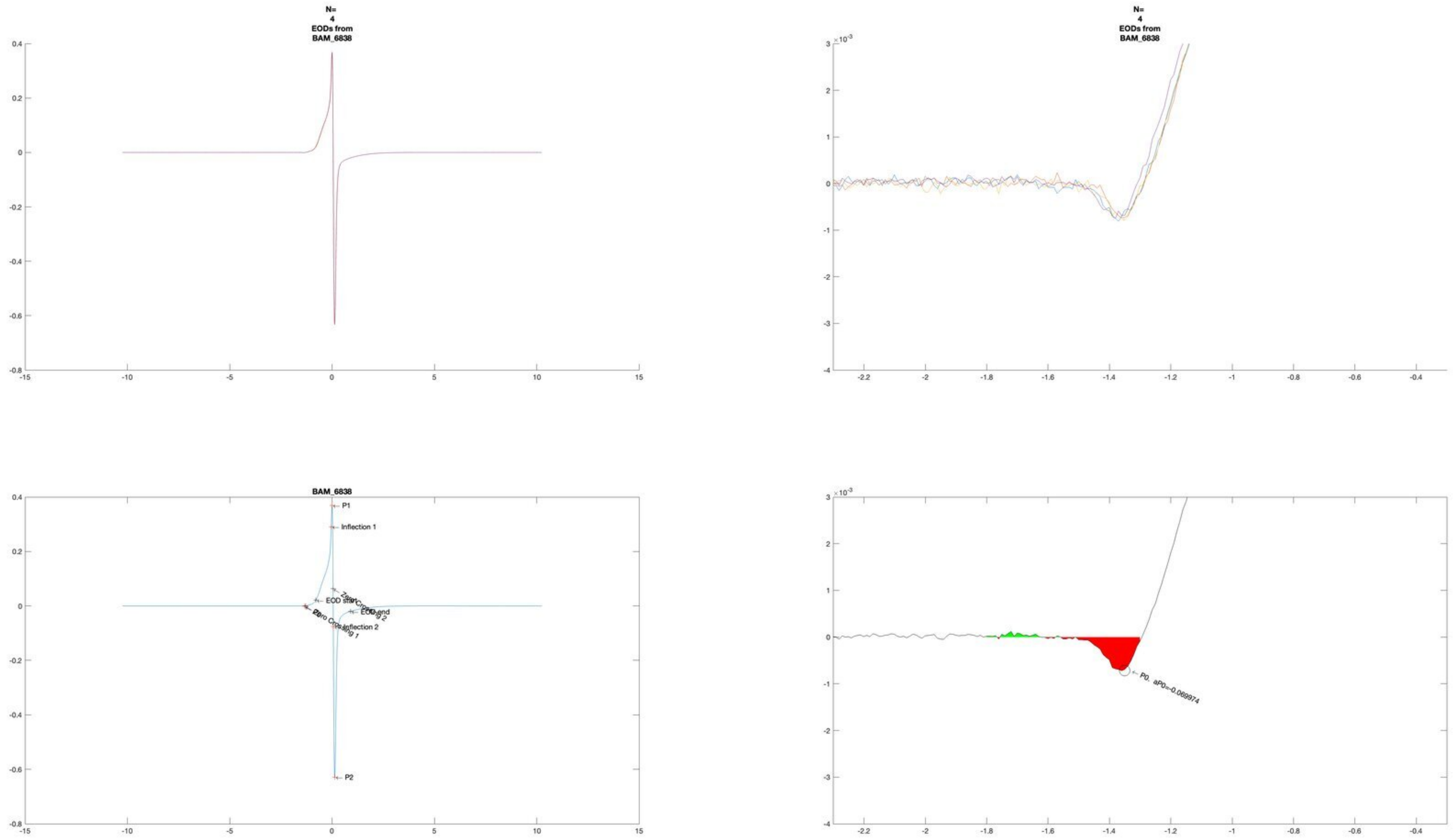
N=
4
EODs from
BAM_6838
N=
4
EODs from
BAM_6838
BAM_6838
P1
Inflection 1
EOD start
EOD end
Zero Crossing 1
Inflection 2
P2
P0. aP0=-0.069974

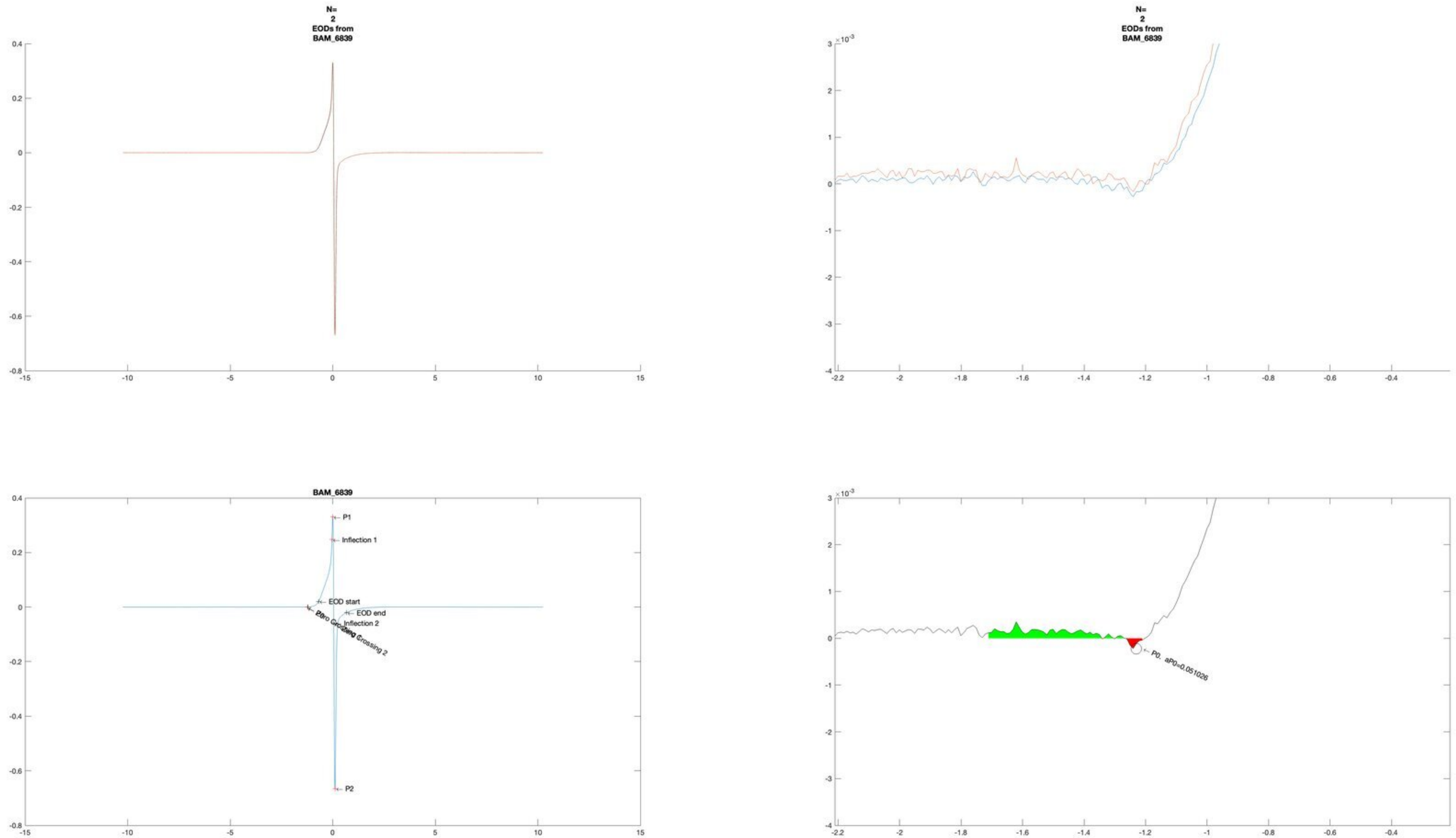
N=
2
EODs from
BAM_6839
N=
2
EODs from
BAM_6839
BAM_6839
P1
Inflection 1
EOD start
EOD end
Inflection 2
Zero Crossing 2
P2
P0, aP0=0.051026

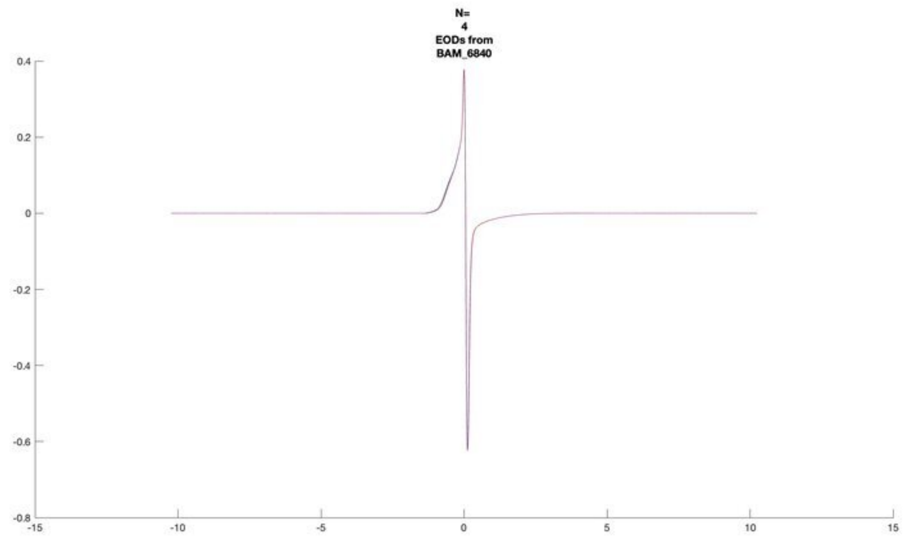
N=
4
EODs from
BAM_6840

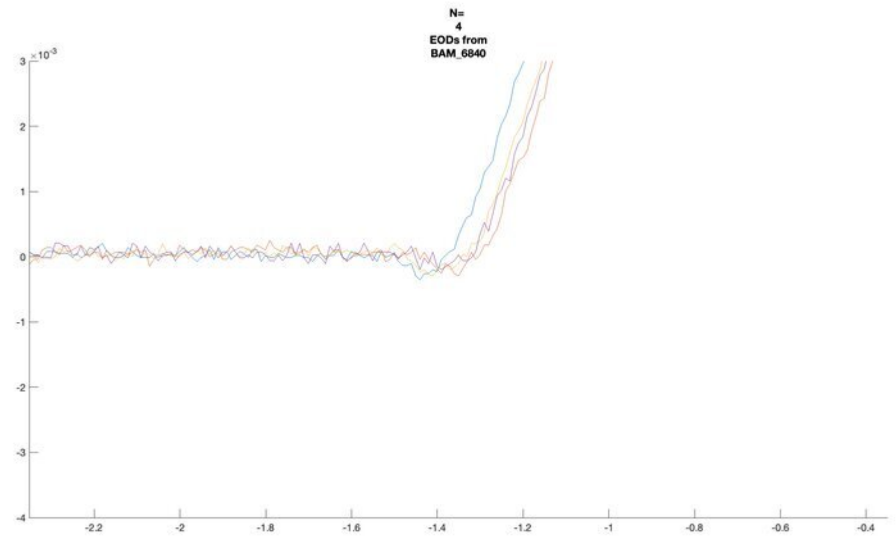
N=
4
EODs from
BAM_6840

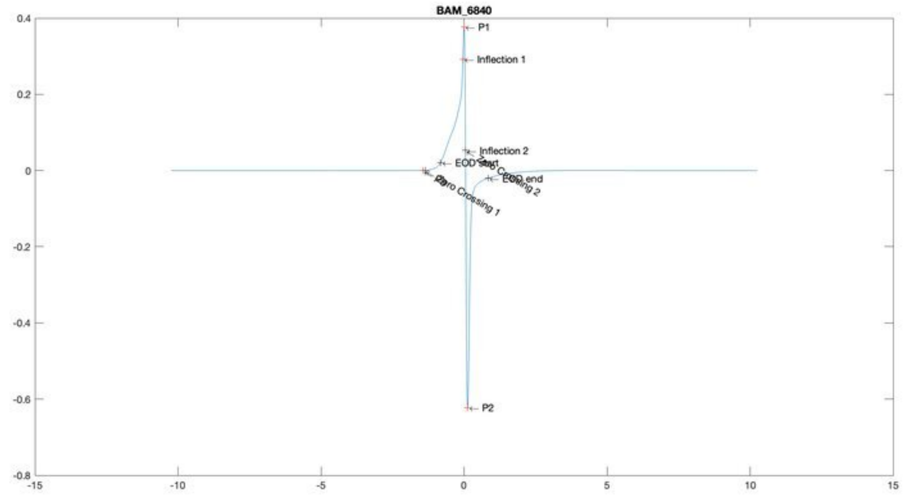
BAM_6840
P1
Inflection 1
Inflection 2
P2

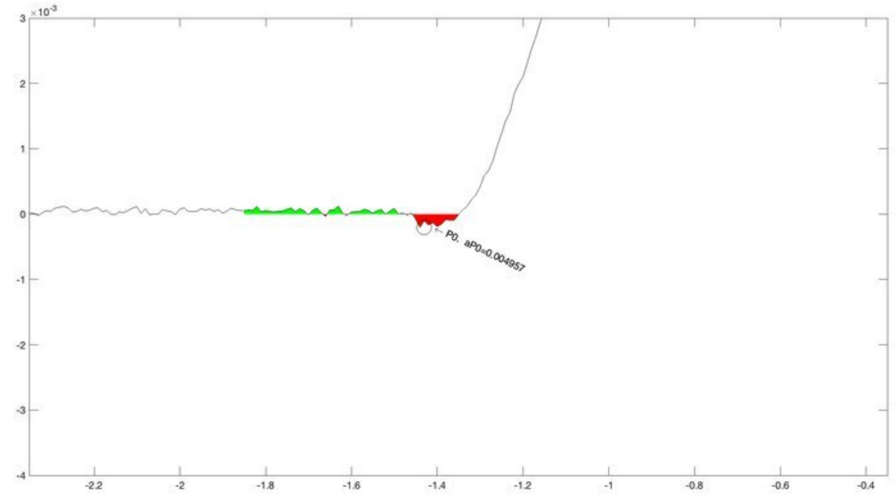
P0, aP0=0.004957

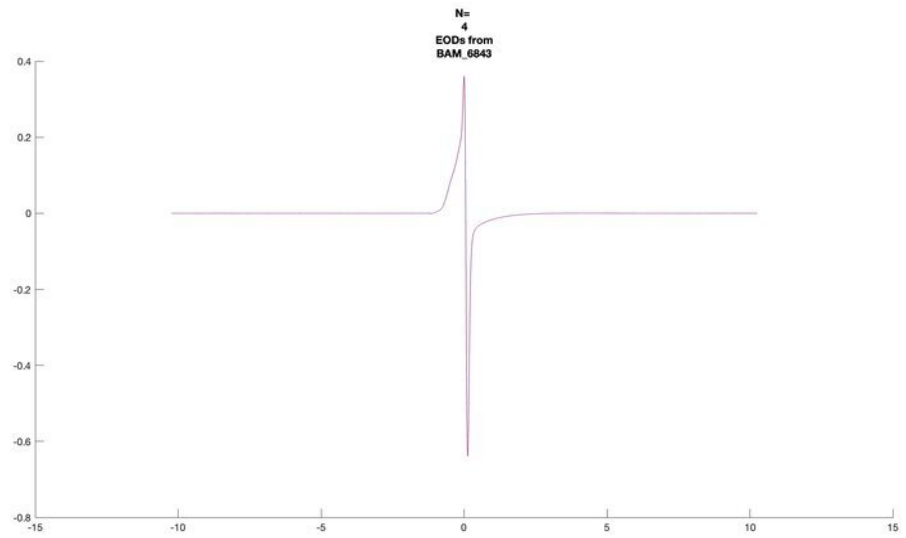
N=
4
EODs from
BAM_6843
0.4
0.2
0
-0.2
-0.4
-0.6
-0.8
-15
-10
-5
0
5
10
15

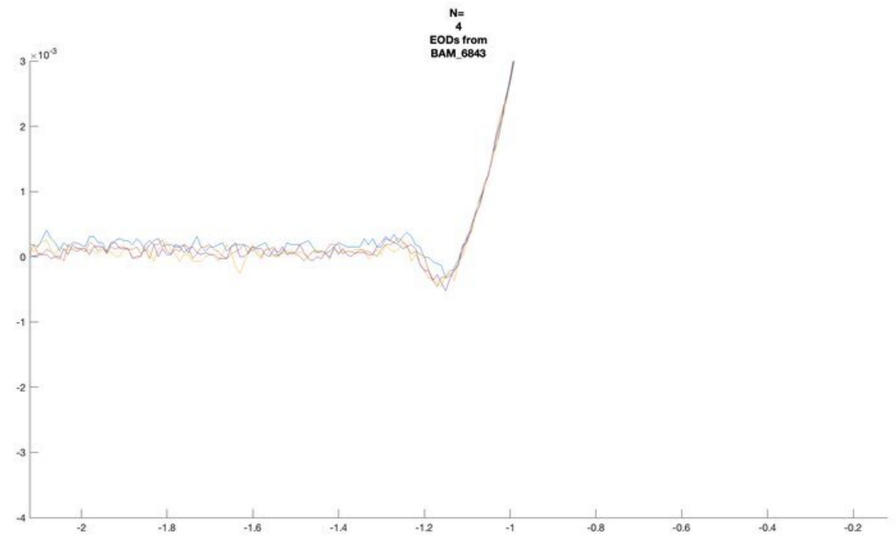
N=
4
EODs from
BAM_6843
×10⁻³
3
2
1
0
-1
-2
-3
-4
-2
-1.8
-1.6
-1.4
-1.2
-1
-0.8
-0.6
-0.4
-0.2

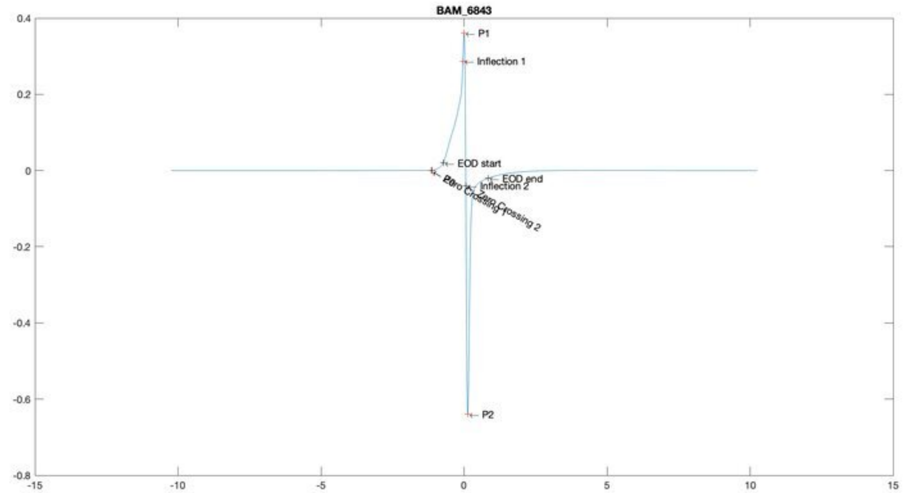
BAM_6843
P1
Inflection 1
EOD start
EOD end
Inflection 2
P0
Zero Crossing 1
Zero Crossing 2
P2
0.4
0.2
0
-0.2
-0.4
-0.6
-0.8
-15
-10
-5
0
5
10
15

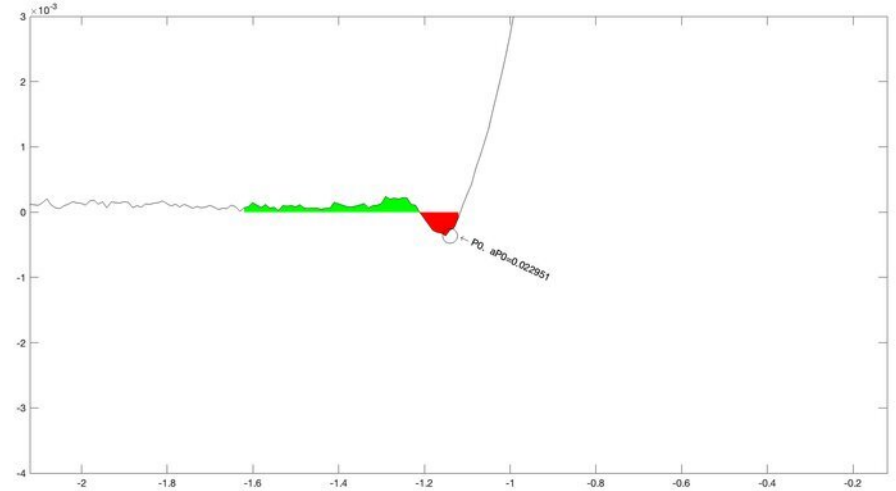
×10⁻³
P0. aP0=0.022951
3
2
1
0
-1
-2
-3
-4
-2
-1.8
-1.6
-1.4
-1.2
-1
-0.8
-0.6
-0.4
-0.2

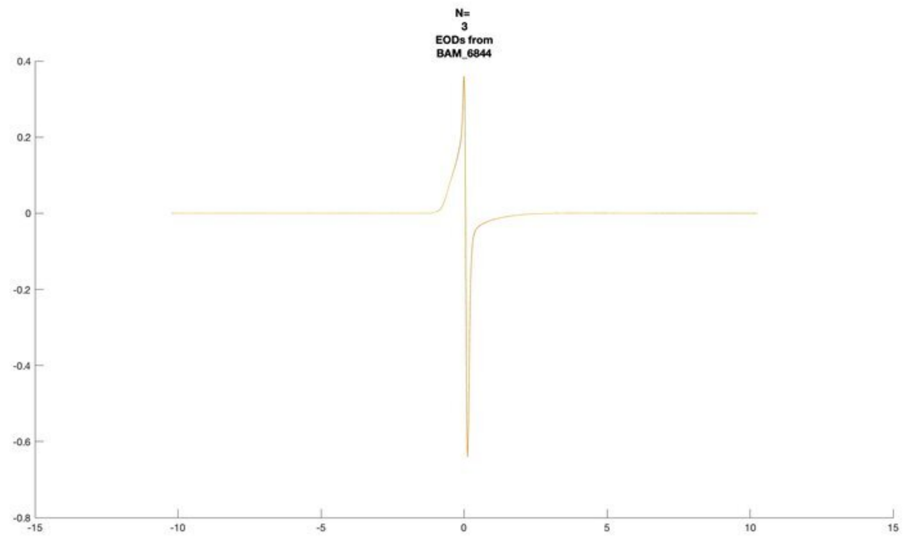
N=
3
EODs from
BAM_6844

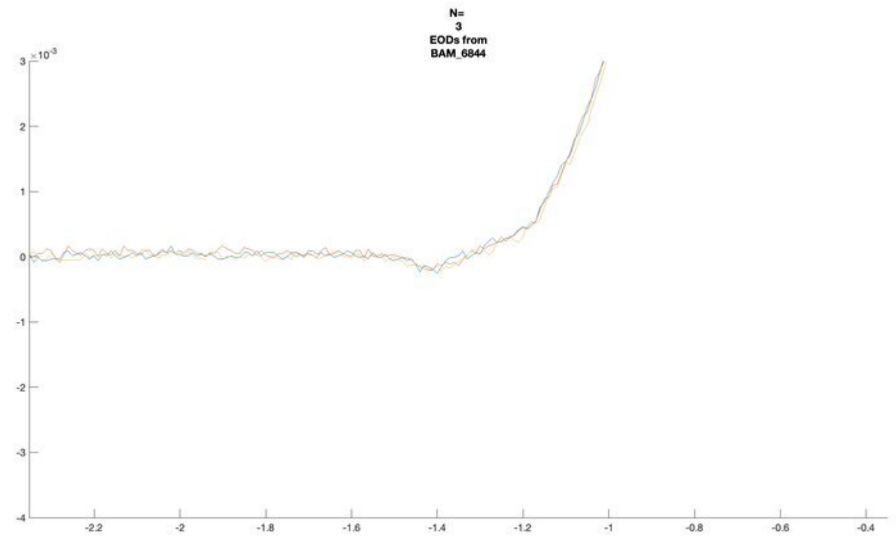
N=
3
EODs from
BAM_6844

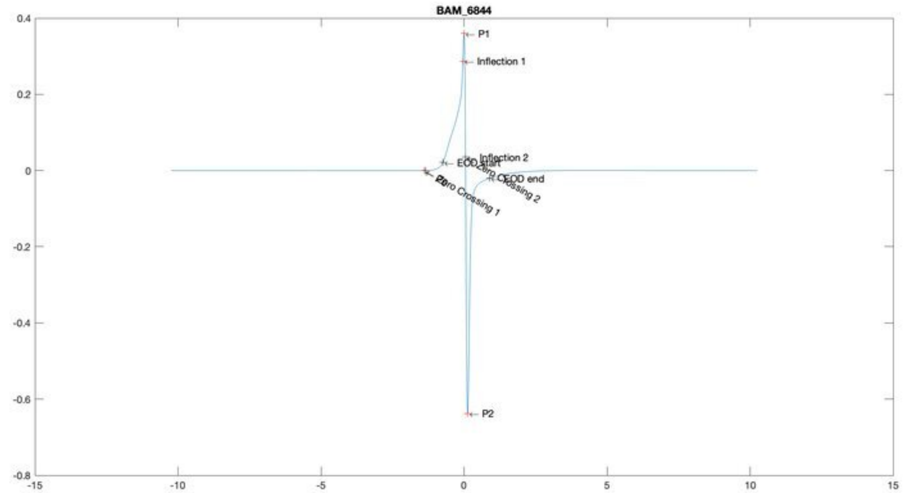
BAM_6844
P1
Inflection 1
Inflection 2
EOD start
Zero Crossing 1
EOD end
Zero Crossing 2
P2

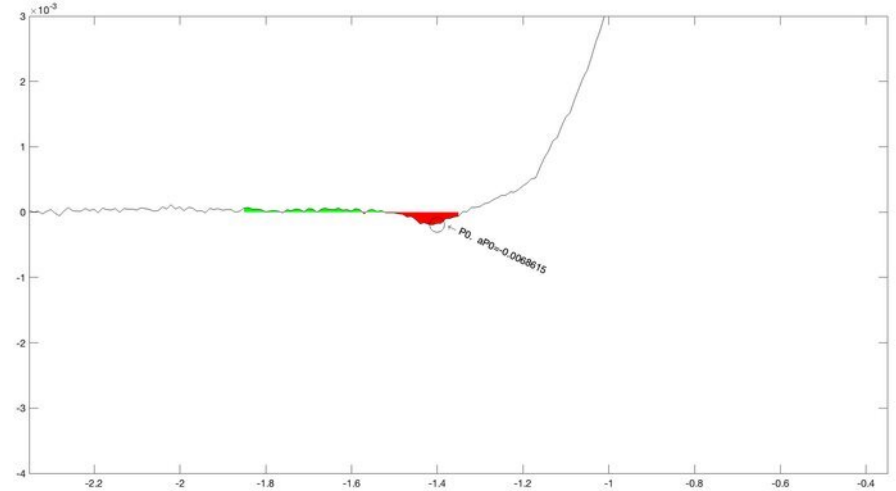
P0, aP0=-0.0068615

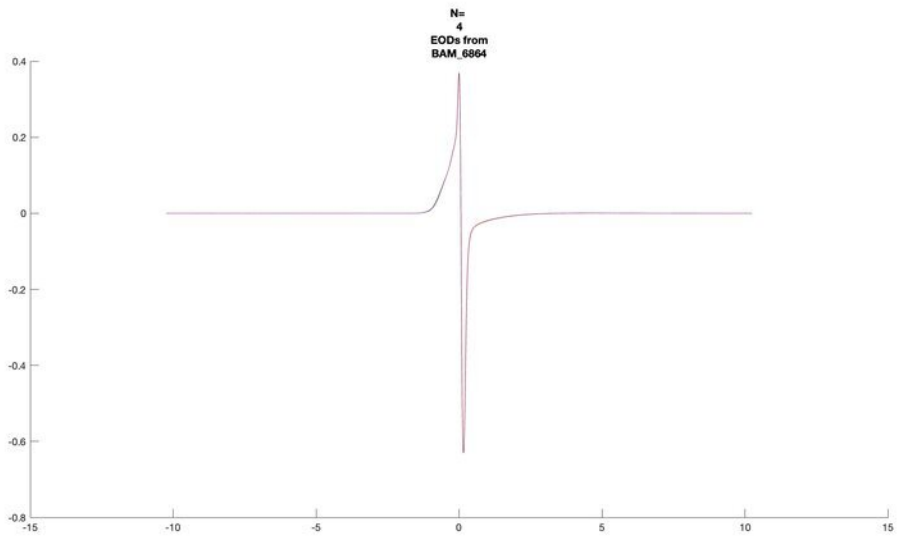
N=
4
EODs from
BAM_6864

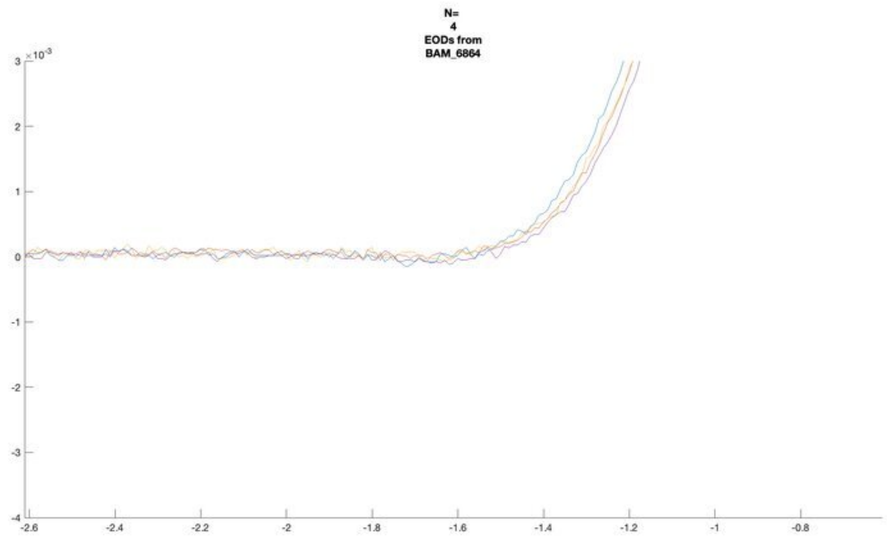
N=
4
EODs from
BAM_6864

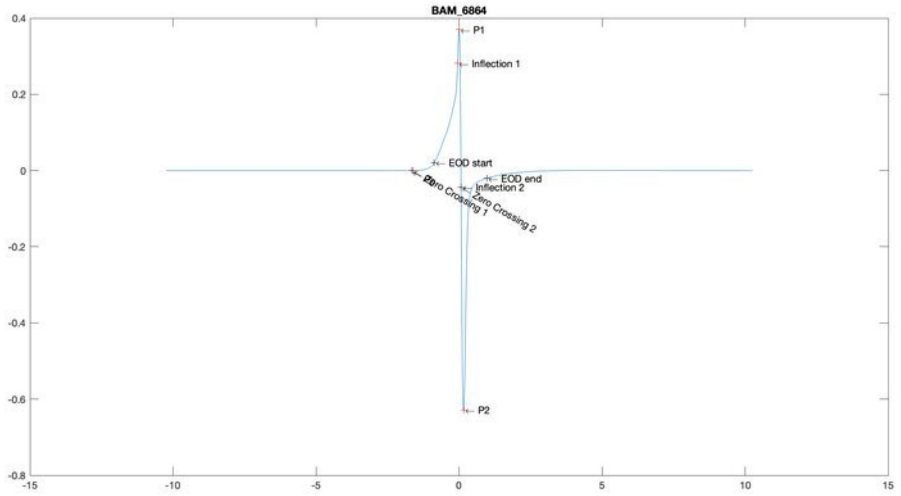
BAM_6864
P1
Inflection 1
EOD start
EOD end
Inflection 2
Zero Crossing 1
Zero Crossing 2
P2

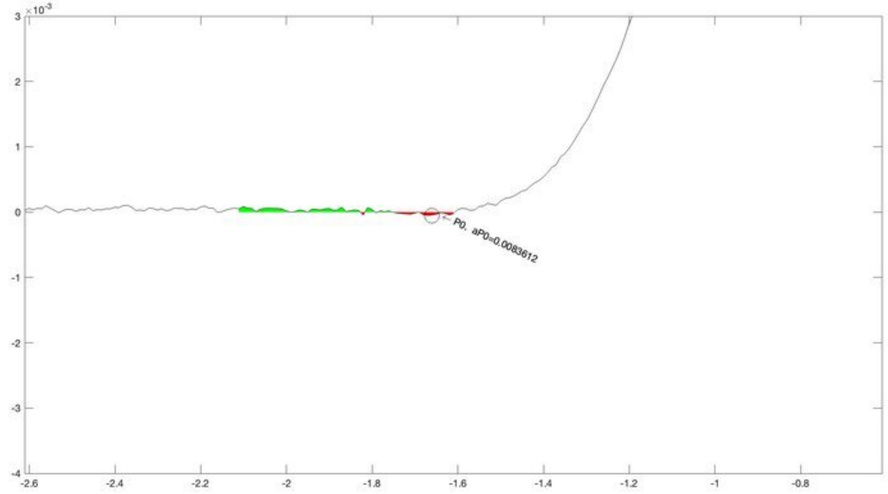
P0, aP0=0.0083612

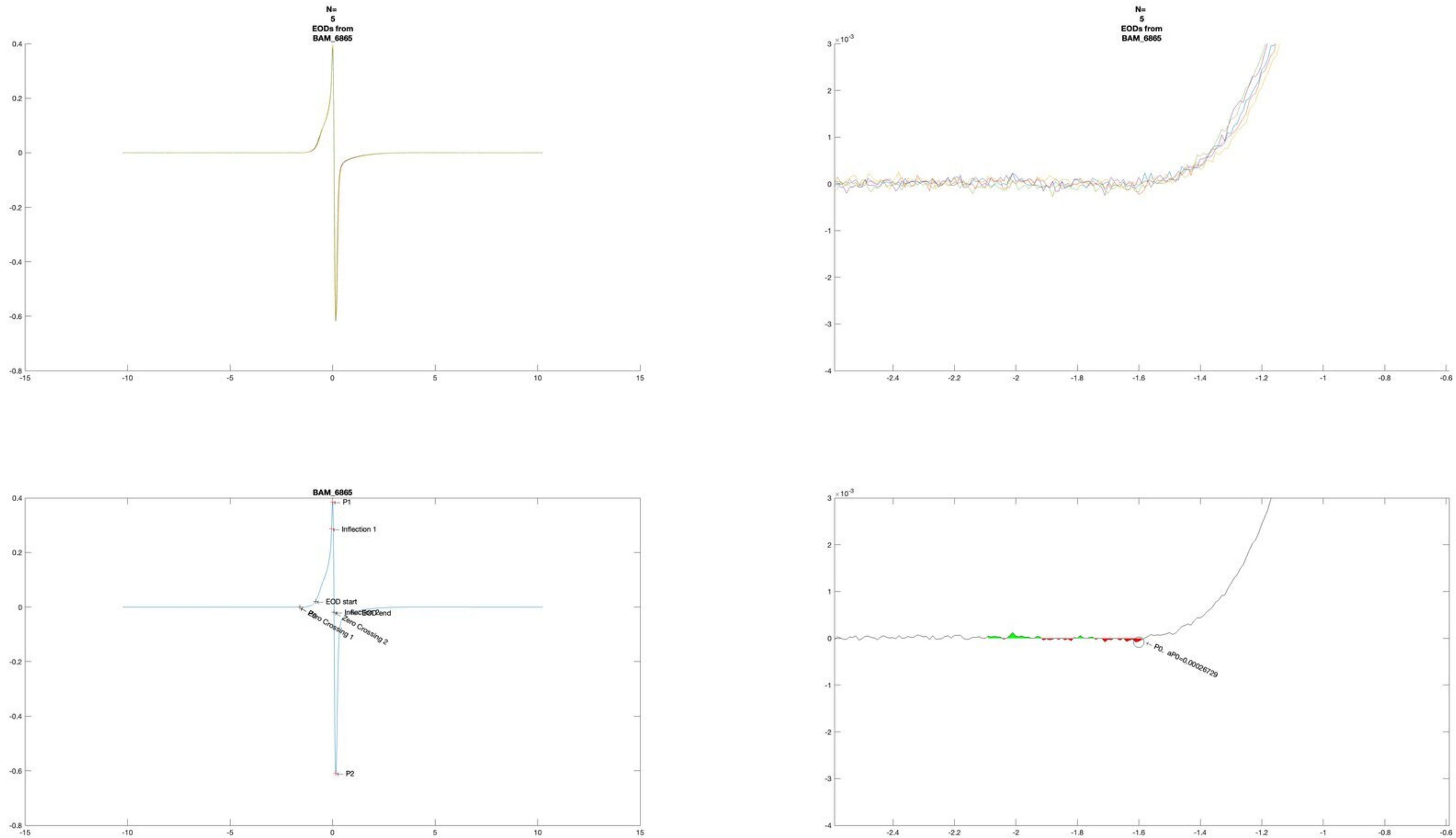

N=
5
EODs from
BAM_6865
N=
5
EODs from
BAM_6865
BAM_6865
P1
Inflection 1
EOD start
Zero Crossing 1
Zero Crossing 2
EOD end
P2
P0, aP0=0.00026729

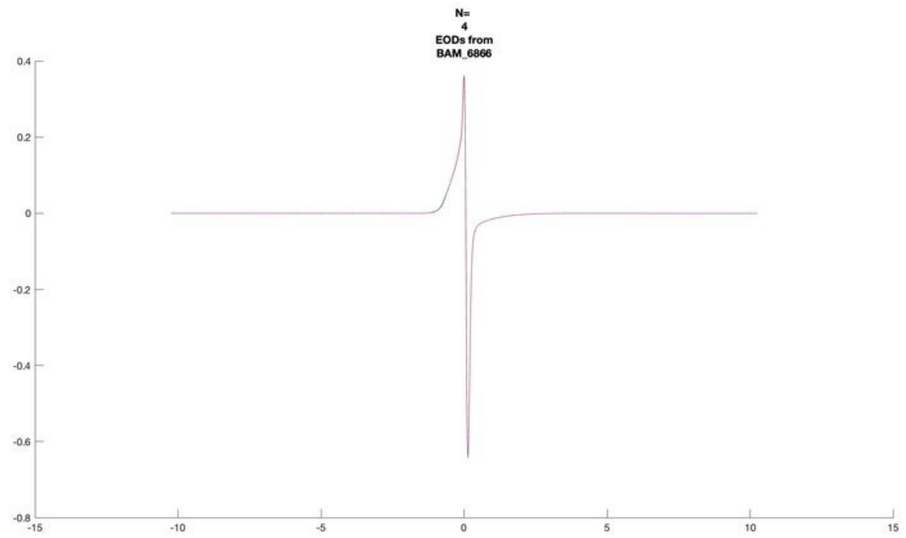
N=
4
EODs from
BAM_6866
0.4
0.2
0
-0.2
-0.4
-0.6
-0.8
-15
-10
-5
0
5
10
15

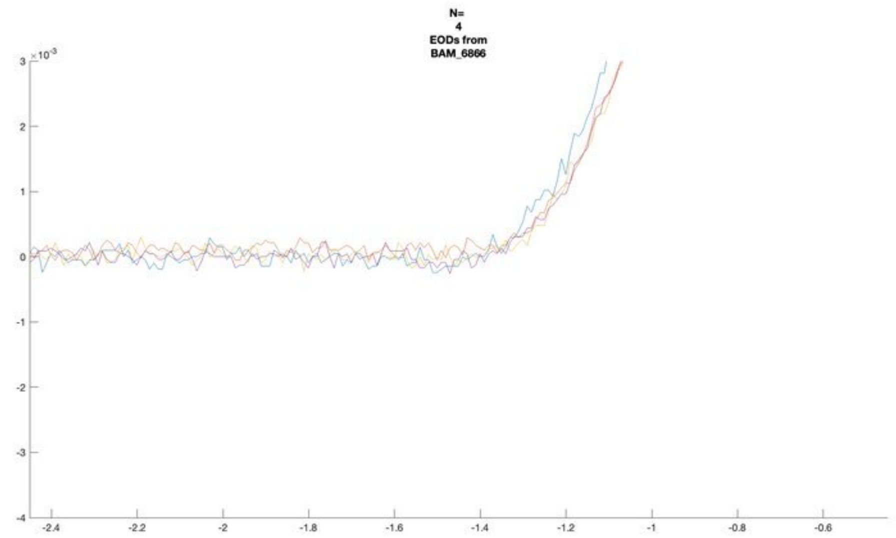
N=
4
EODs from
BAM_6866
×10^-3
3
2
1
0
-1
-2
-3
-4
-2.4
-2.2
-2
-1.8
-1.6
-1.4
-1.2
-1
-0.8
-0.6

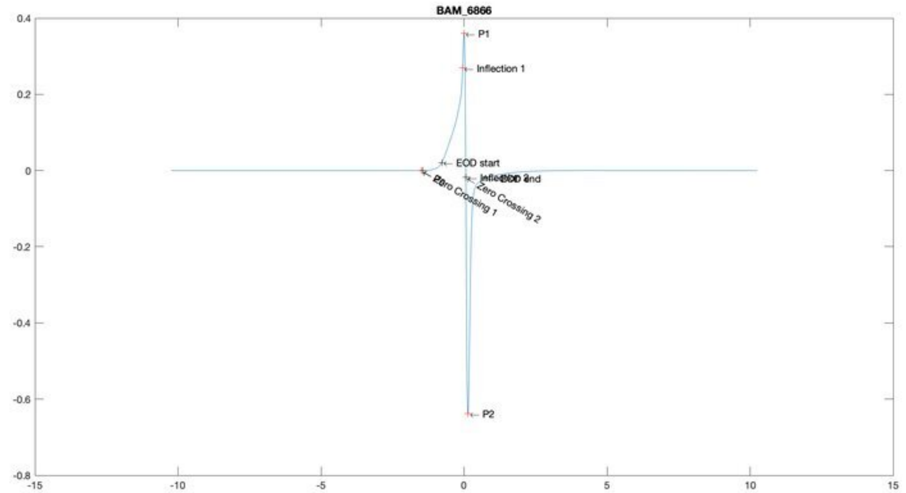
BAM_6866
P1
Inflection 1
EOD start
EOD end
Zero Crossing 1
Zero Crossing 2
P2
0.4
0.2
0
-0.2
-0.4
-0.6
-0.8
-15
-10
-5
0
5
10
15

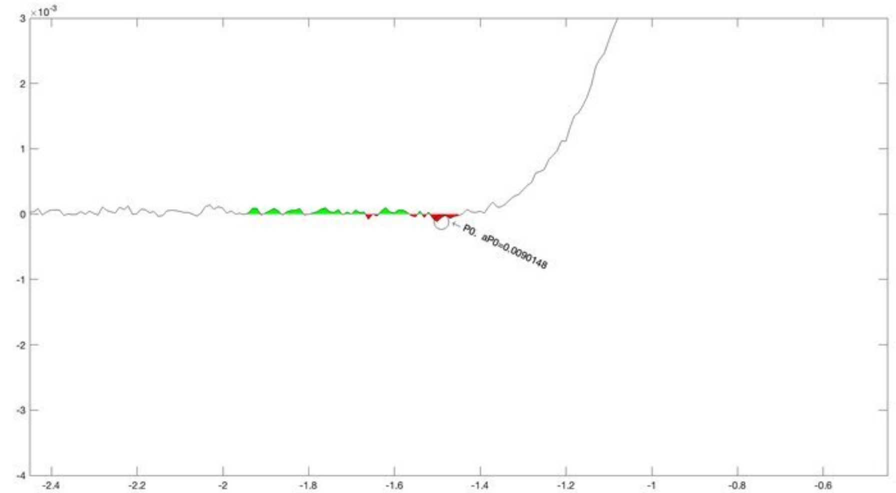
P0, aP0=0.0090148
×10^-3
3
2
1
0
-1
-2
-3
-4
-2.4
-2.2
-2
-1.8
-1.6
-1.4
-1.2
-1
-0.8
-0.6

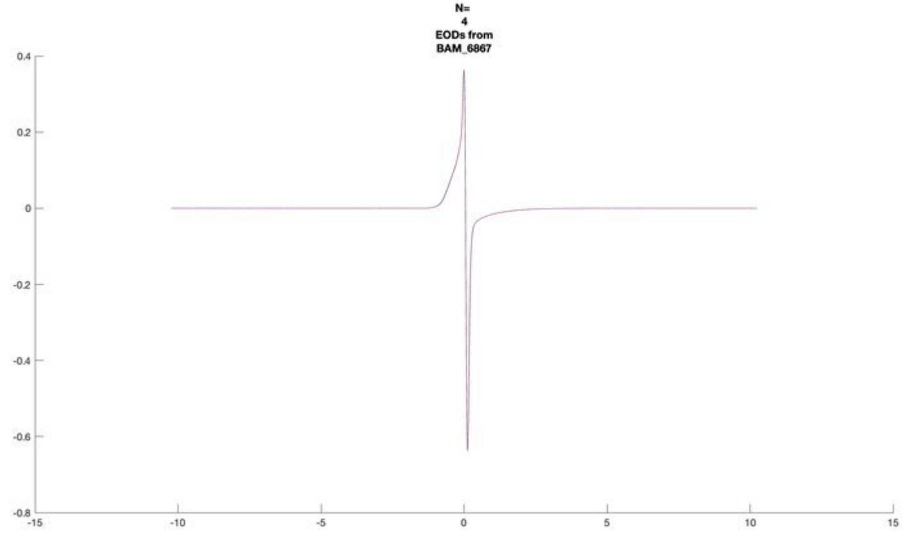
N=
4
EODs from
BAM_6867

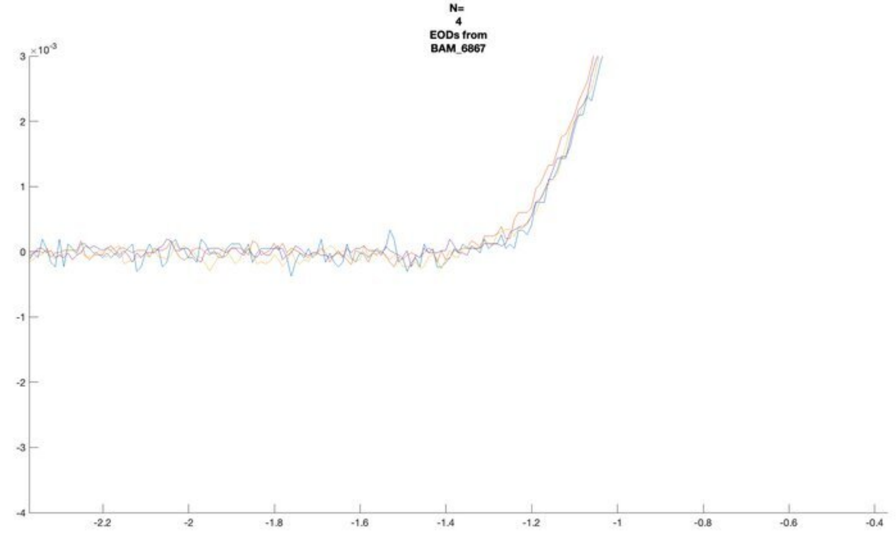
N=
4
EODs from
BAM_6867

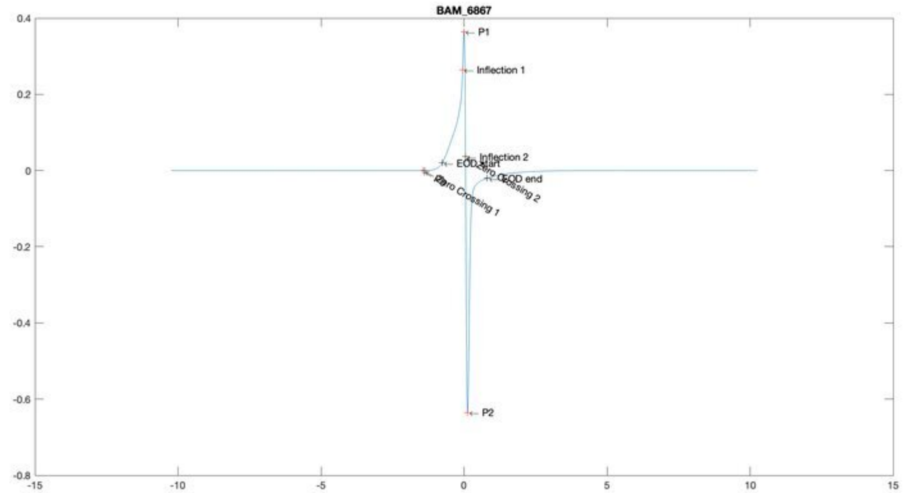
BAM_6867
P1
Inflection 1
Inflection 2
EOD end
P2

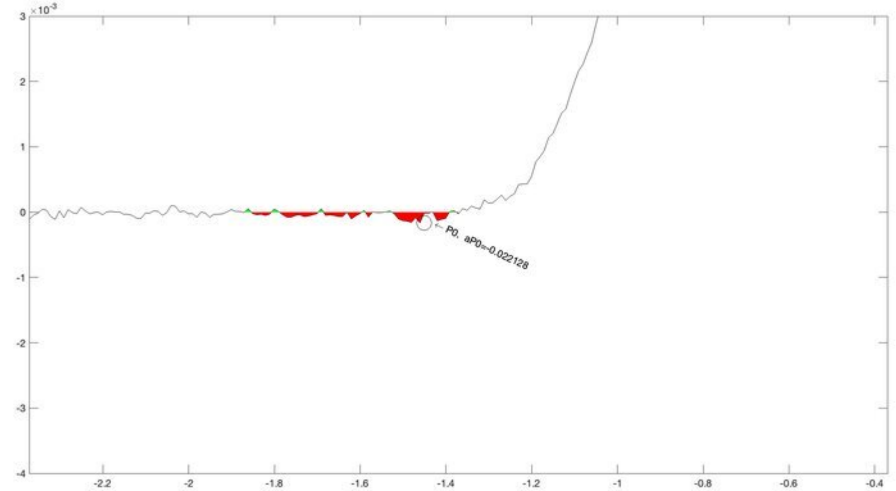
P0, aP0=-0.022128

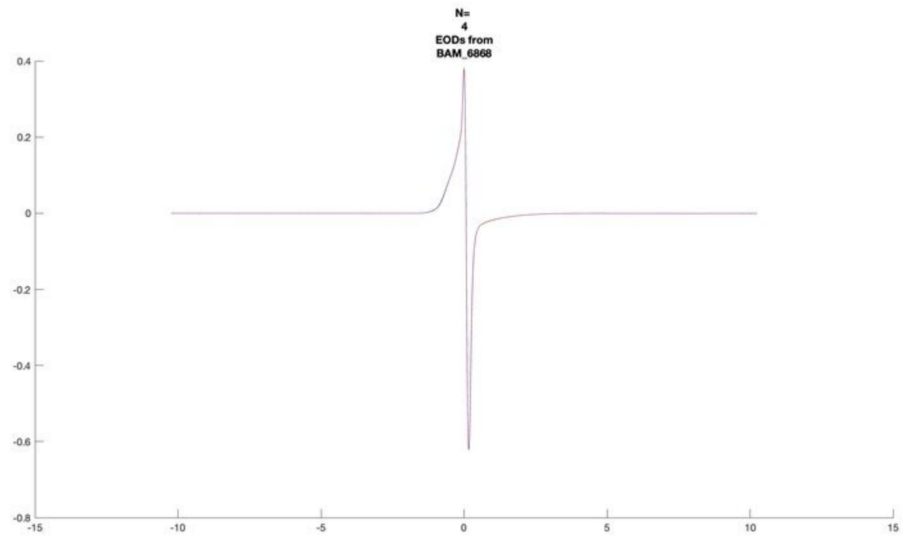
N=
4
EODs from
BAM_6868

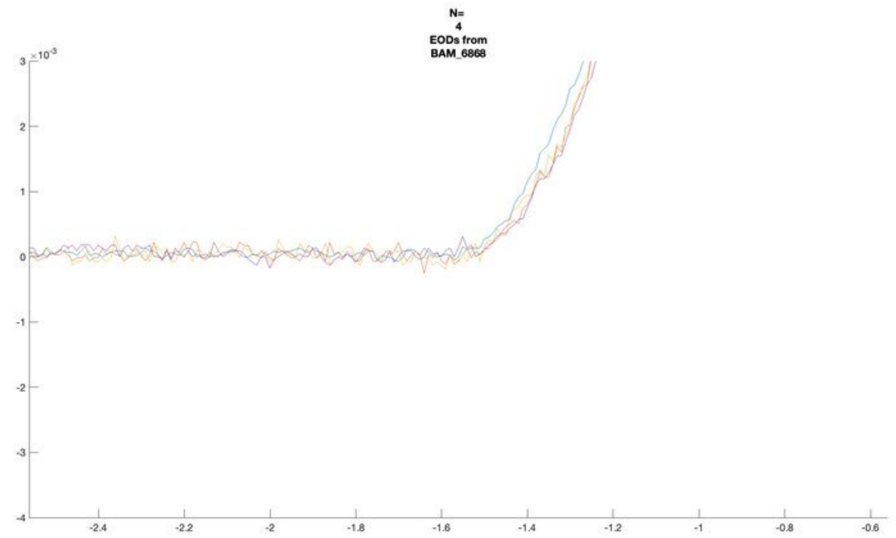
N=
4
EODs from
BAM_6868

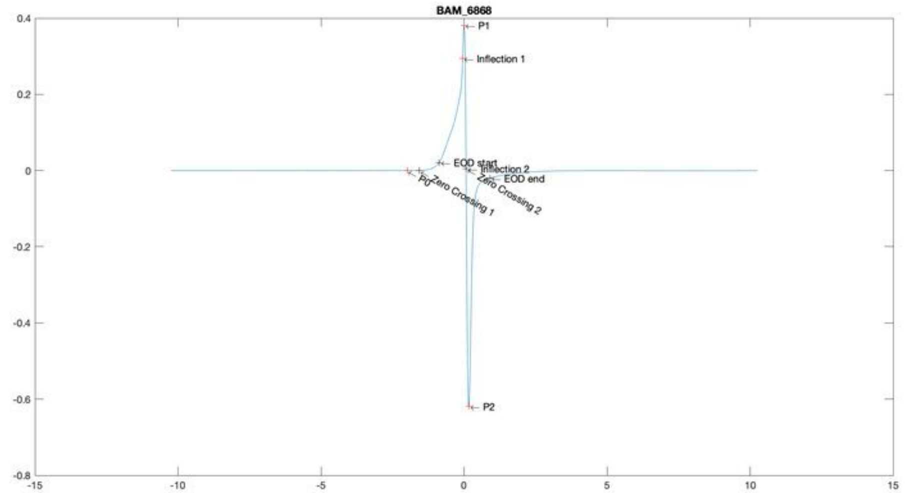
BAM_6868
P1
Inflection 1
EOD start
Inflection 2
EOD end
P0
Zero Crossing 1
Zero Crossing 2
P2

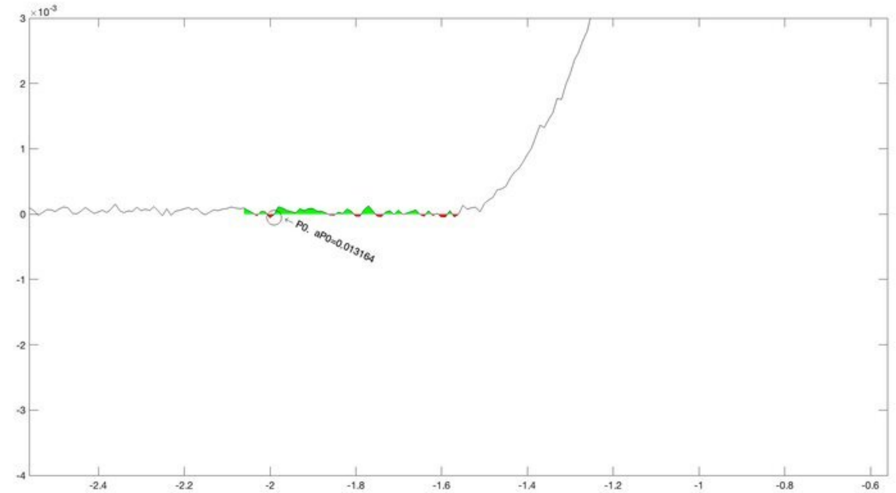
P0, aP0=0.013164

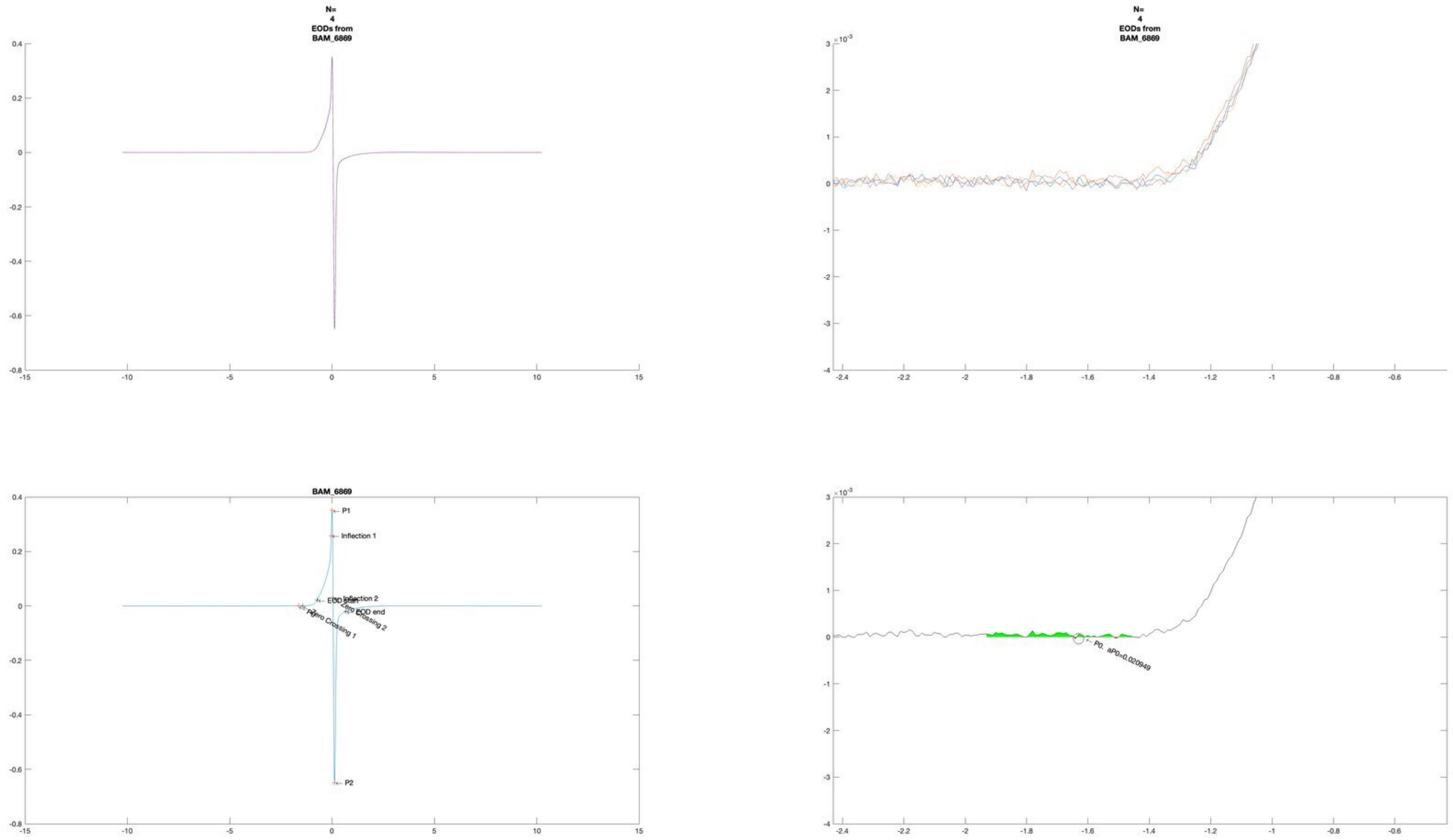
N=
4
EODs from
BAM_6869
N=
4
EODs from
BAM_6869
BAM_6869
P1
Inflection 1
Inflection 2
EOD start
EOD end
Zero Crossing 1
Zero Crossing 2
P2
P0, aP0=0.020949

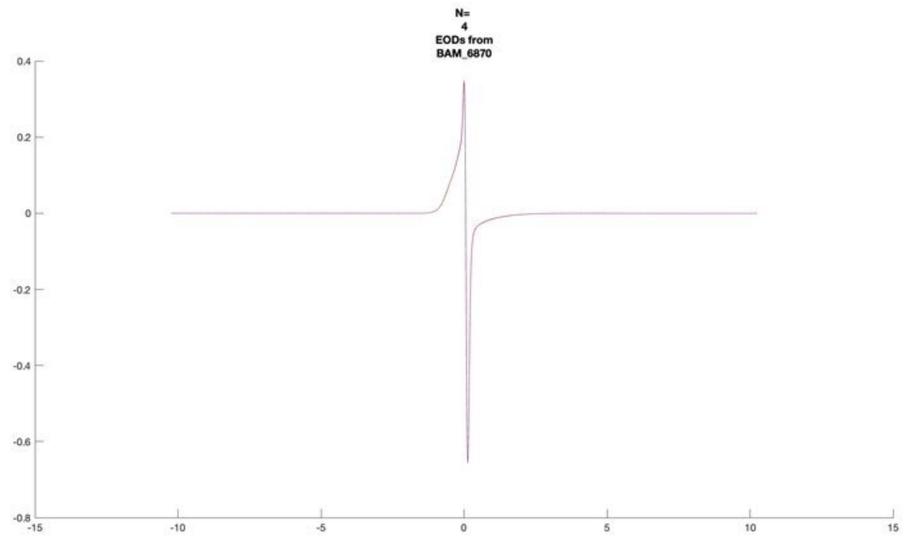
N=
4
EODs from
BAM_6870

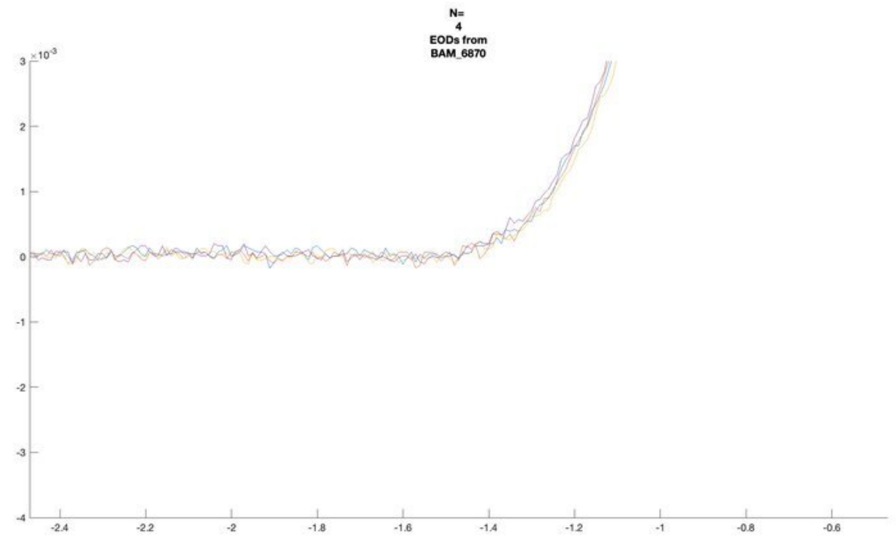
N=
4
EODs from
BAM_6870

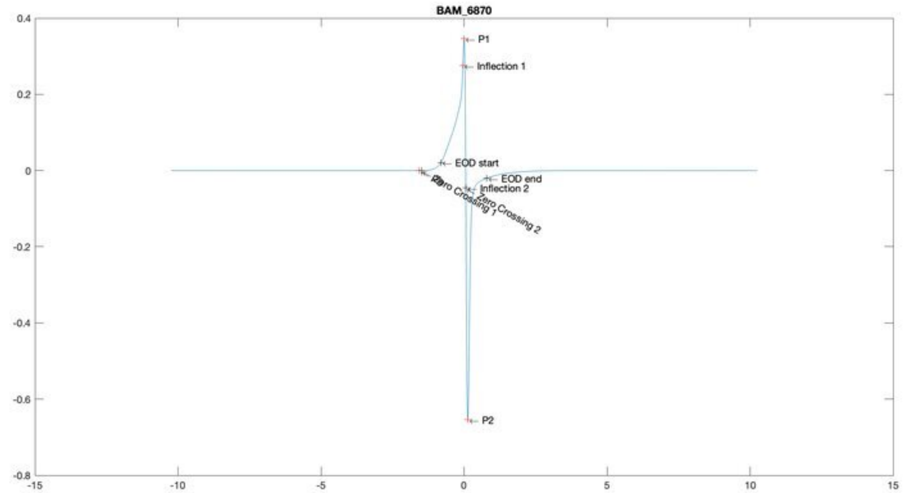
BAM_6870
P1
Inflection 1
EOD start
EOD end
Inflection 2
Zero Crossing 1
Zero Crossing 2
P2

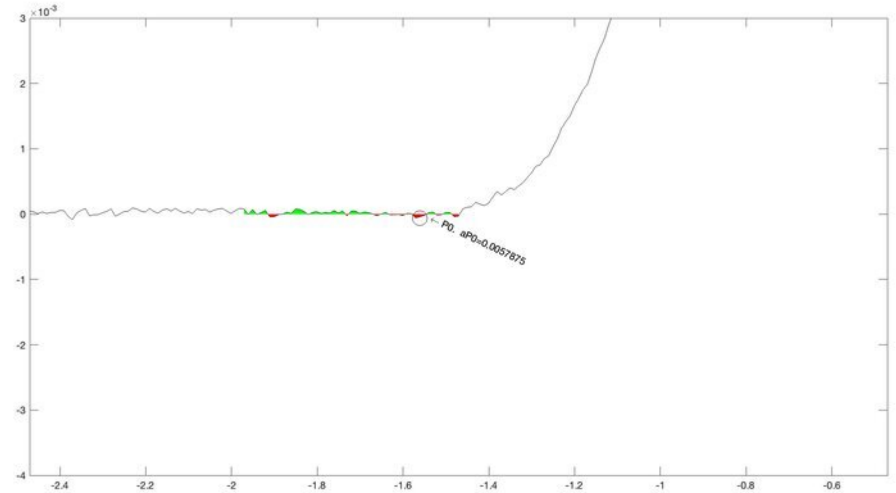
P0, aP0=0.0057875

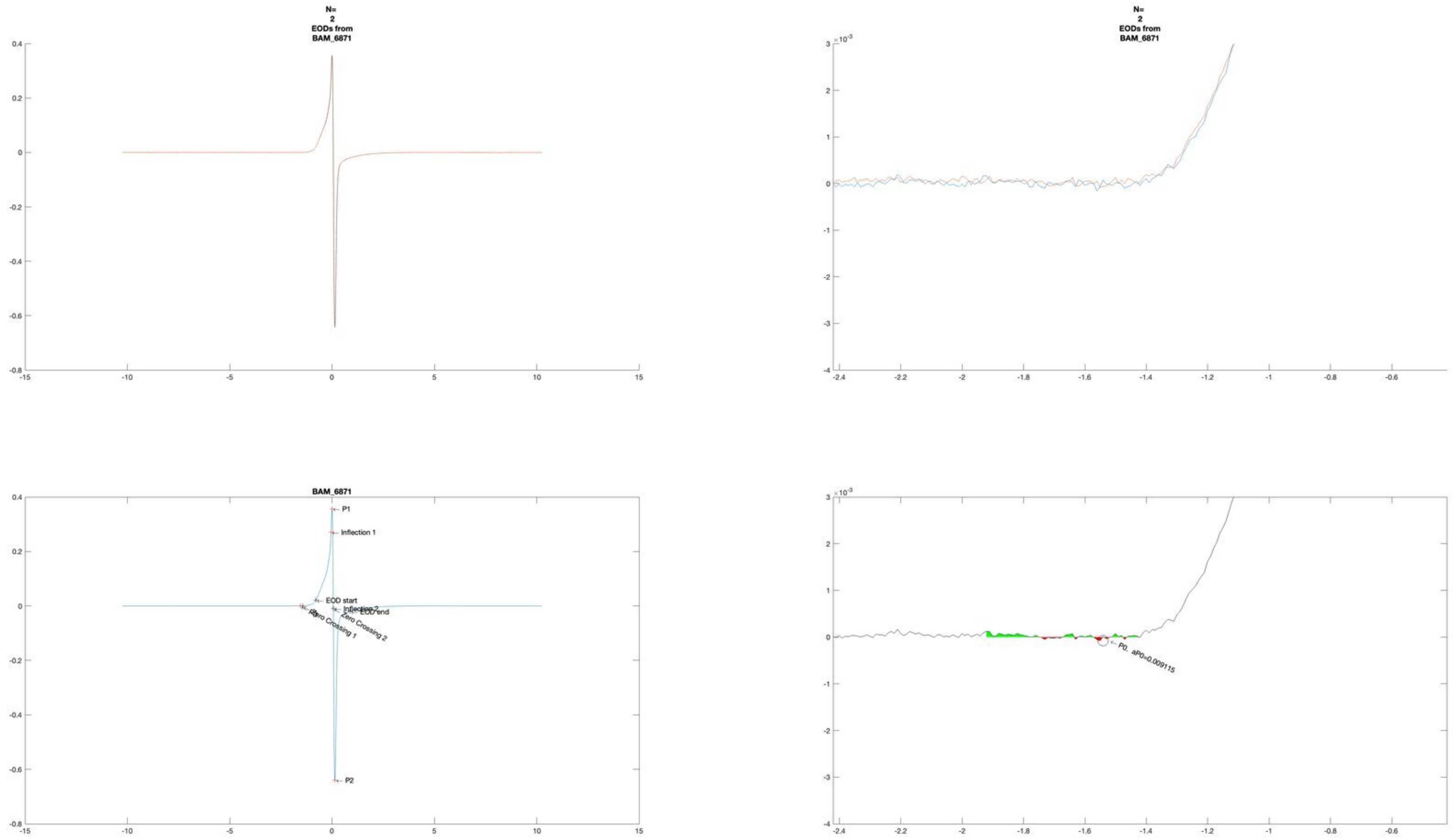
N=
2
EODs from
BAM_6871
N=
2
EODs from
BAM_6871
BAM_6871
P1
Inflection 1
EOD start
Inflection 2
EOD end
Zero Crossing 1
Zero Crossing 2
P2
P0, aP0=0.009115
×10^-3

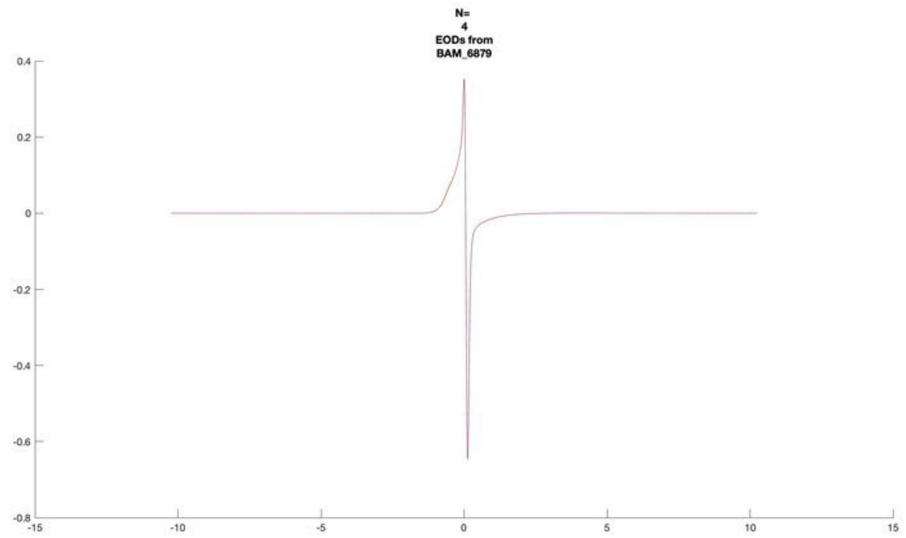
N=
4
EODs from
BAM_6879

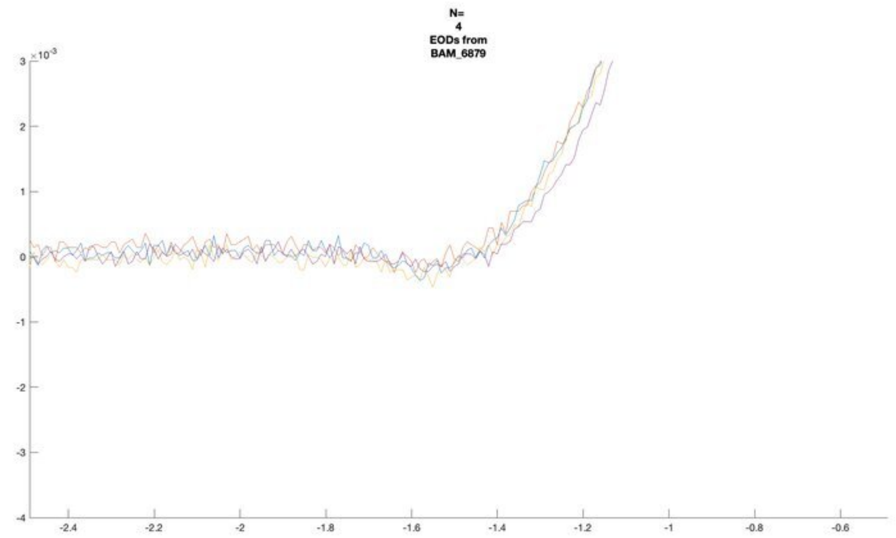
N=
4
EODs from
BAM_6879
×10^-3

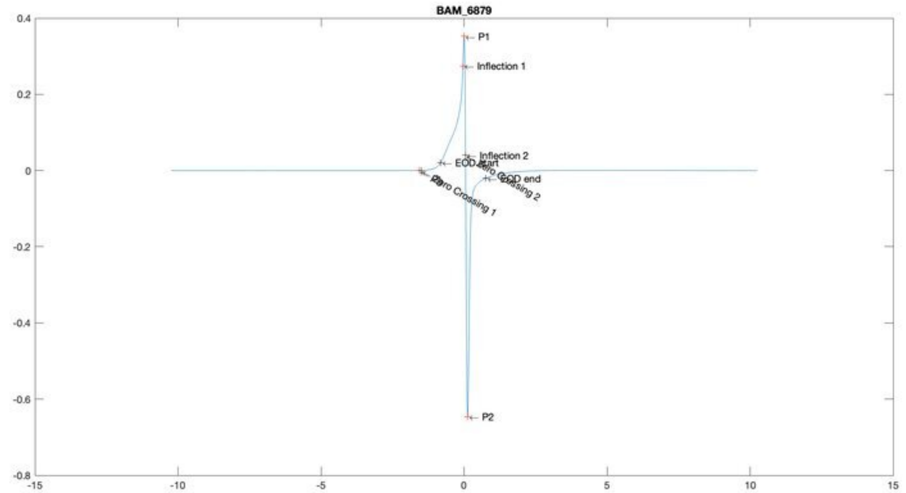
BAM_6879
P1
Inflection 1
Inflection 2
EOD end
P2

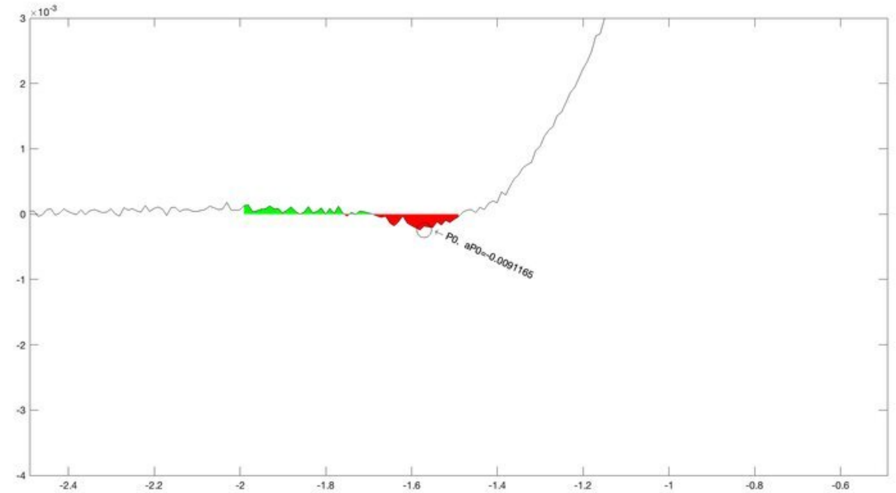
×10^-3
P0, aP0=-0.0091165

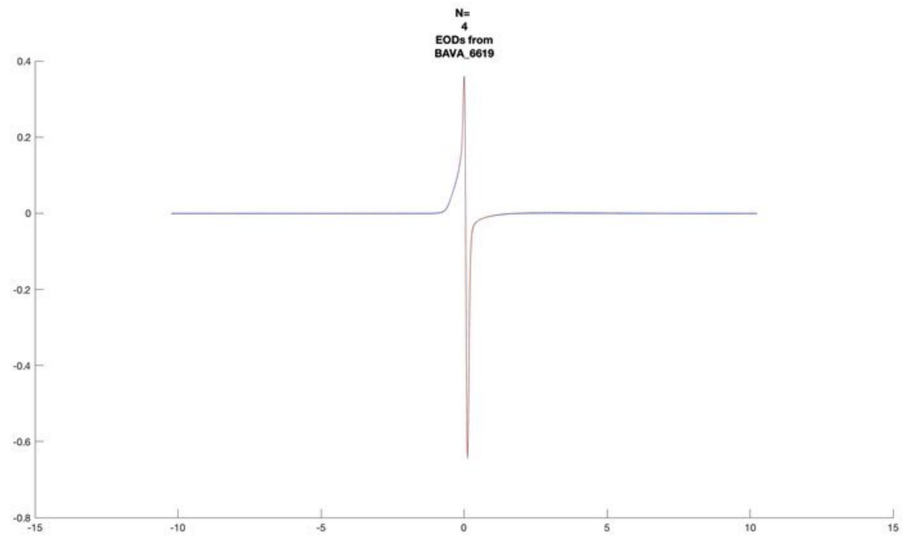
N=
4
EODs from
BAVA_6619

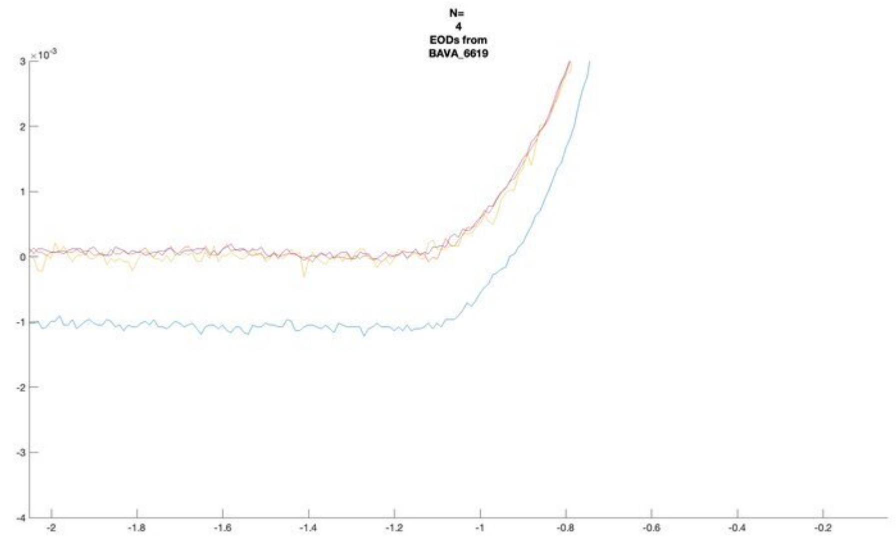
N=
4
EODs from
BAVA_6619

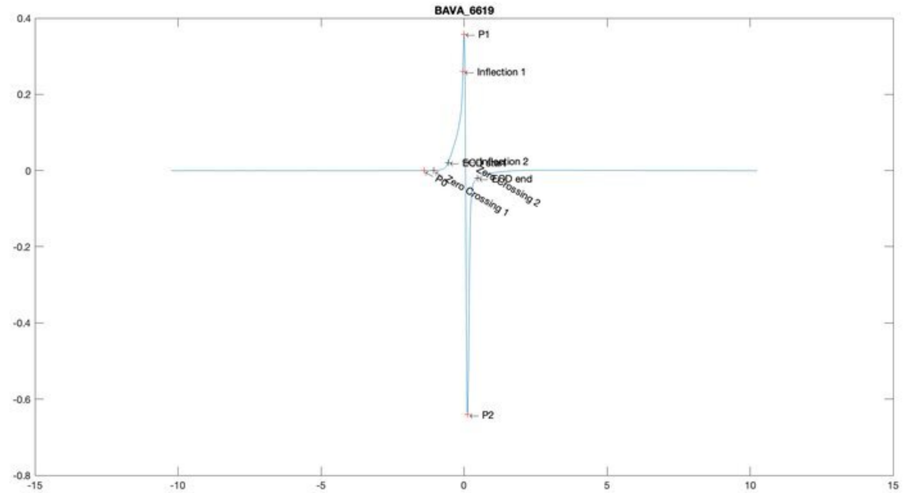
BAVA_6619
P1
Inflection 1
Inflection 2
EOD start
EOD end
P0
Zero Crossing 1
Zero Crossing 2
P2

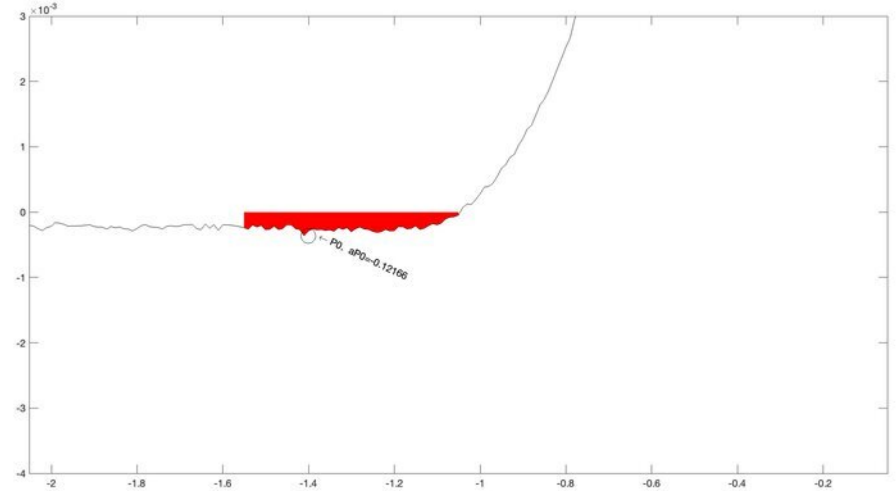
P0, aP0=-0.12166

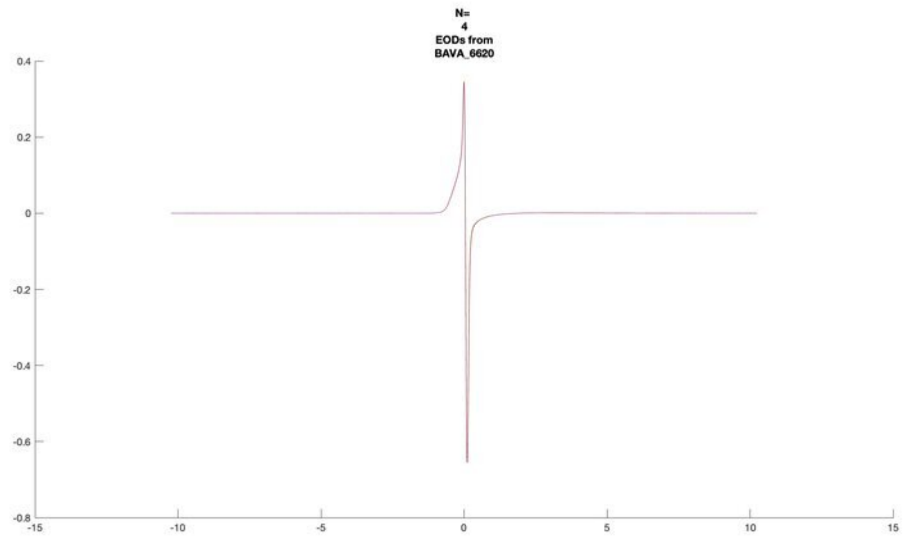
N=
4
EODs from
BAVA_6620

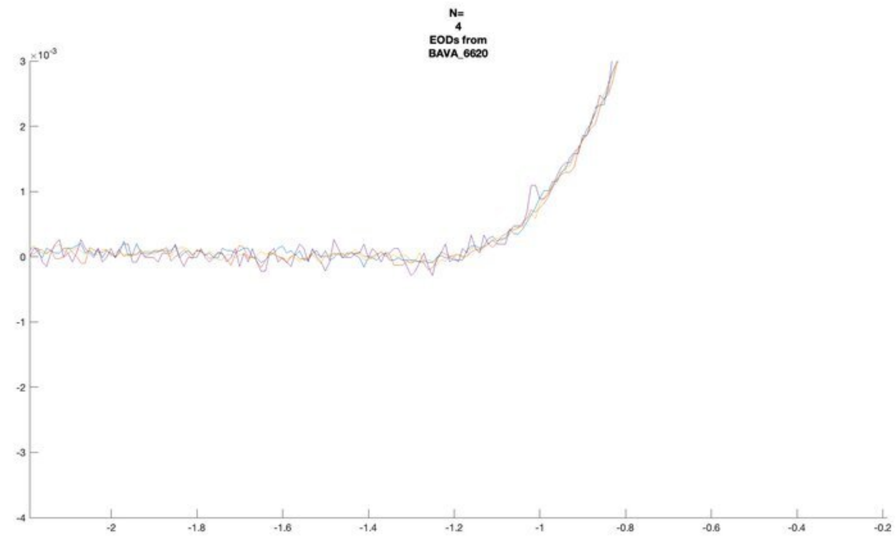
N=
4
EODs from
BAVA_6620
×10^-3

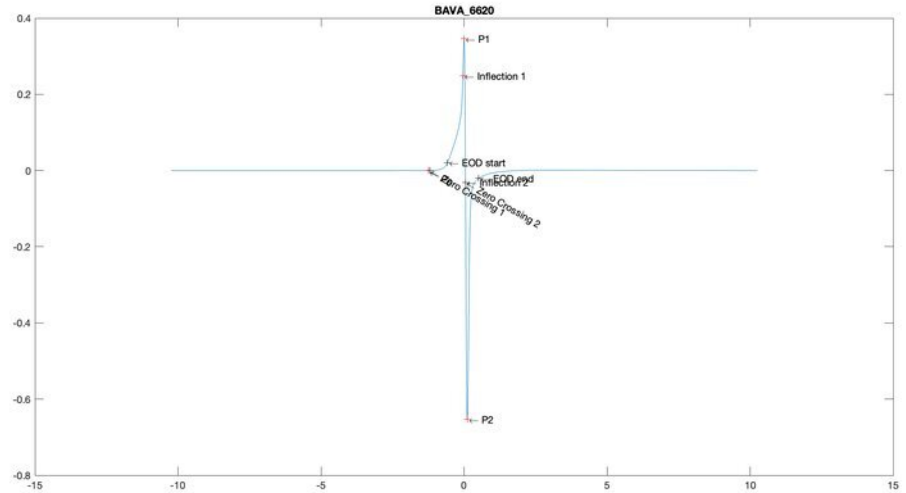
BAVA_6620
P1
Inflection 1
EOD start
EOD end
Inflection 2
Zero Crossing 1
Zero Crossing 2
P2

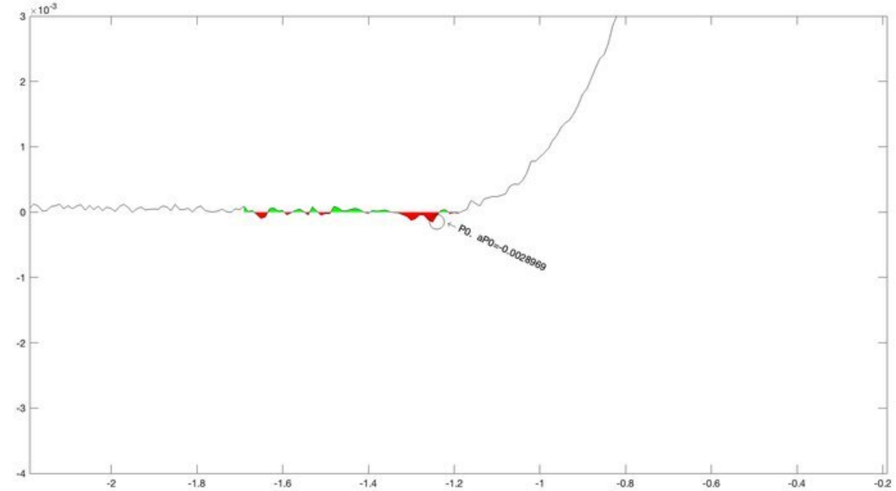
×10^-3
P0. aP0=-0.0028969

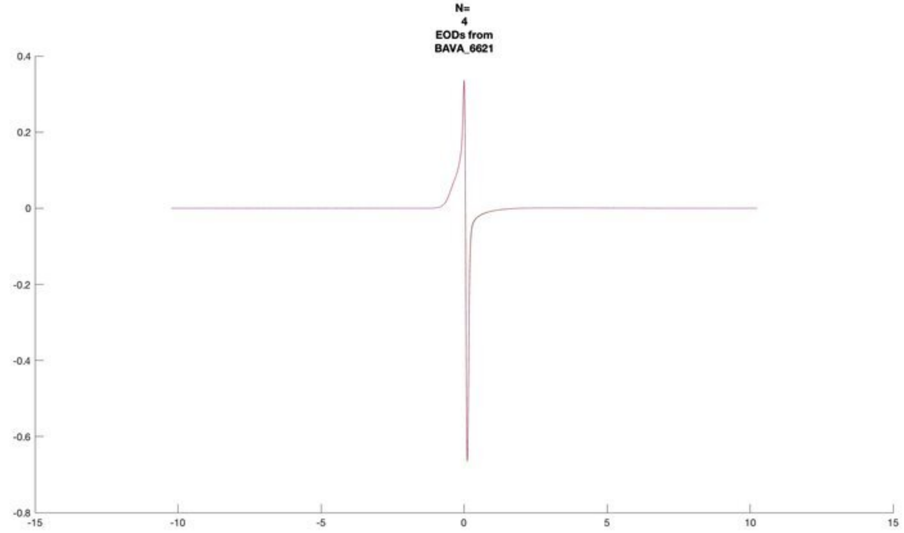
N=
4
EODs from
BAVA_6621

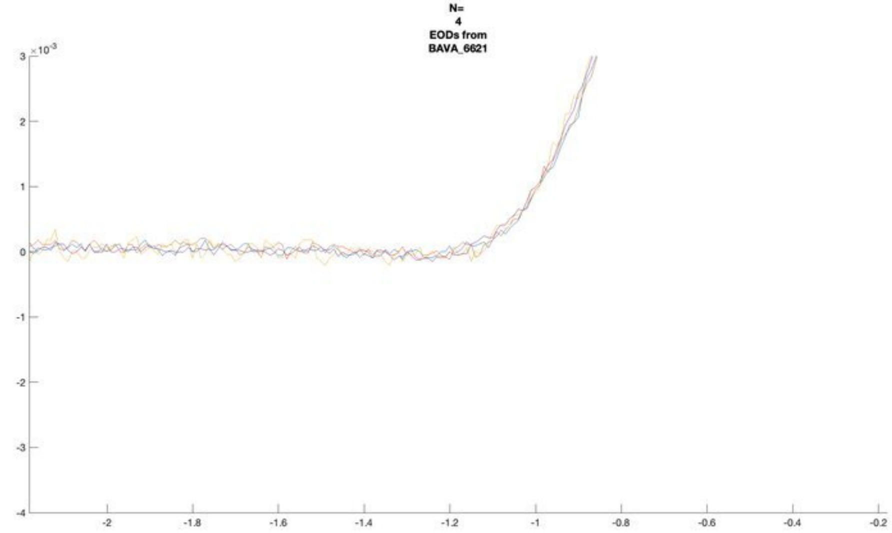
N=
4
EODs from
BAVA_6621
$\times 10^{-3}$

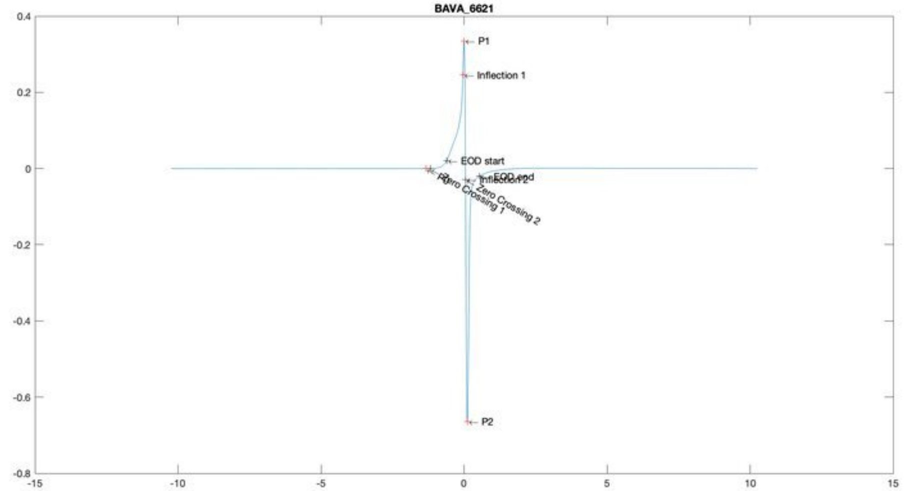
BAVA_6621
P1
Inflection 1
EOD start
EOD end
Inflection 2
Zero Crossing 2
Zero Crossing 1
P0

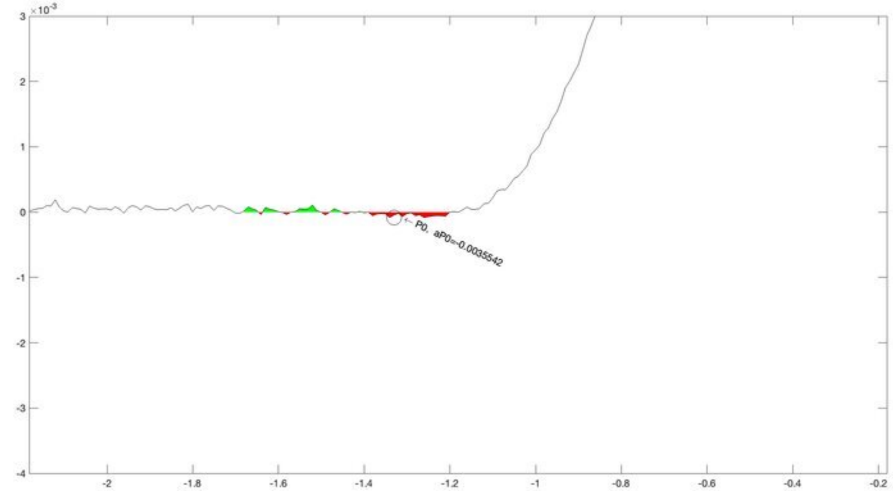
$\times 10^{-3}$
P0, aP0=-0.0035542

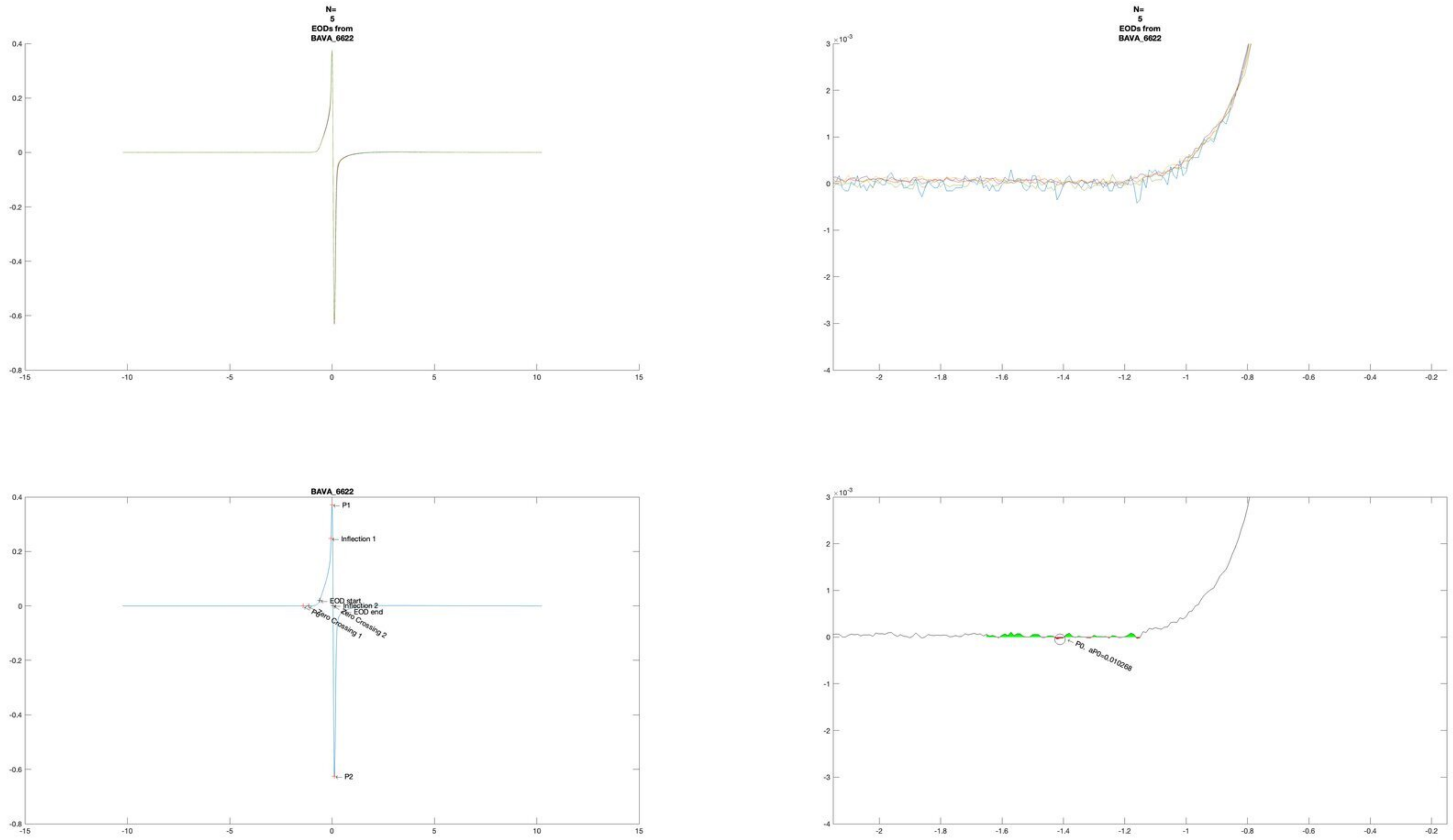
N=
5
EODs from
BAVA_6622
N=
5
EODs from
BAVA_6622
×10^-3
BAVA_6622
P1
Inflection 1
EOD start
Inflection 2
EOD end
Zero Crossing 2
P0
Zero Crossing 1
P2
×10^-3
P0, aP0=0.010268

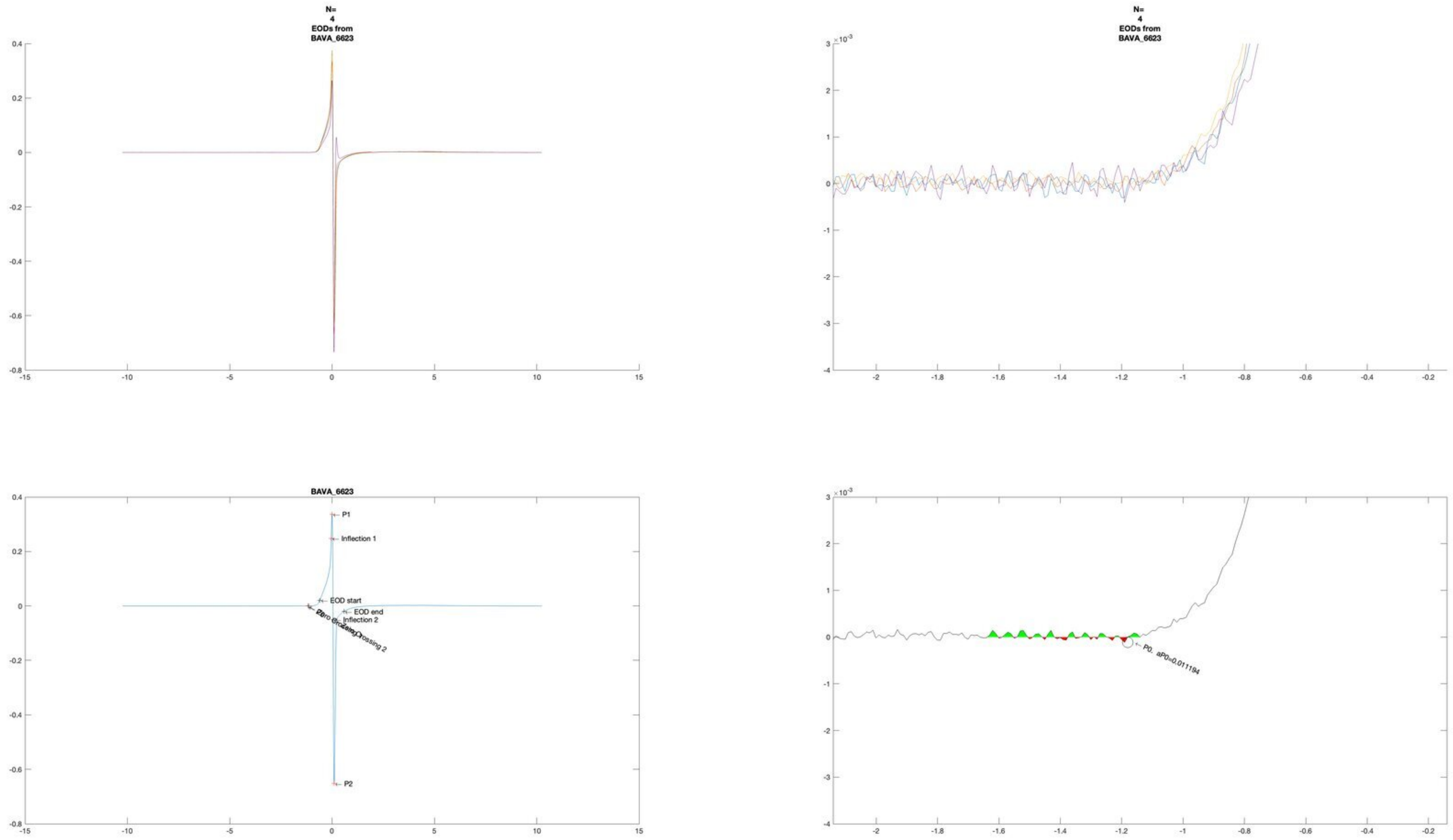
N=
4
EODs from
BAVA_6623
N=
4
EODs from
BAVA_6623
×10^-3
BAVA_6623
P1
Inflection 1
EOD start
EOD end
Inflection 2
Zero Crossing 2
P2
×10^-3
P0, aP0=0.011194

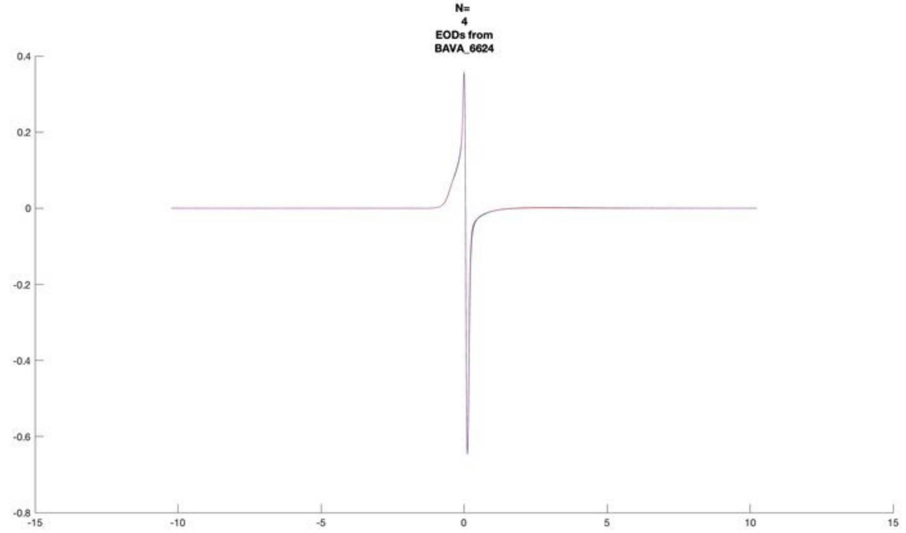
N=
4
EODs from
BAVA_6624

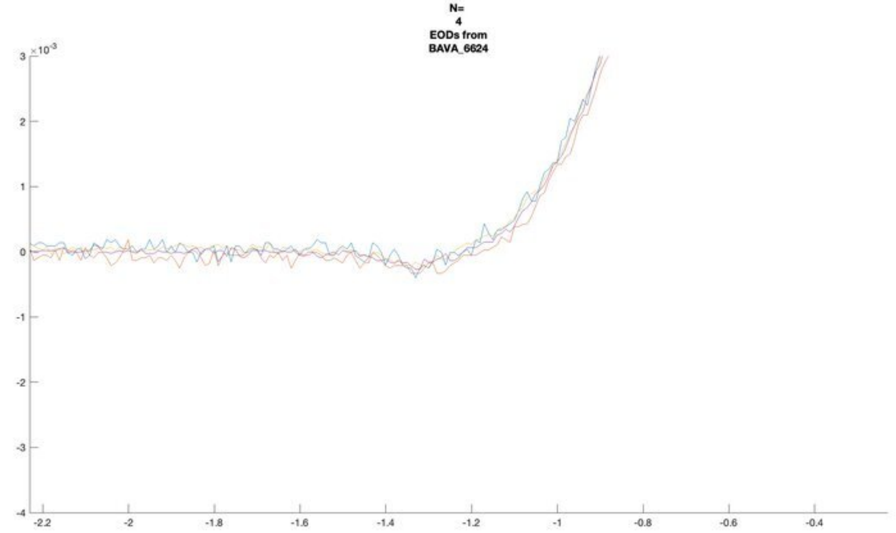
N=
4
EODs from
BAVA_6624
×10^-3

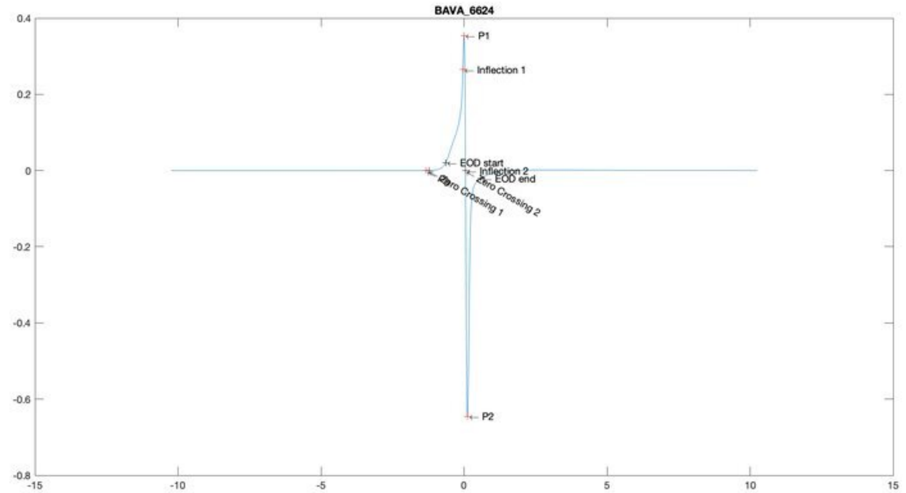
BAVA_6624
P1
Inflection 1
EOD start
Inflection 2
EOD end
Zero Crossing 2
Zero Crossing 1
P2

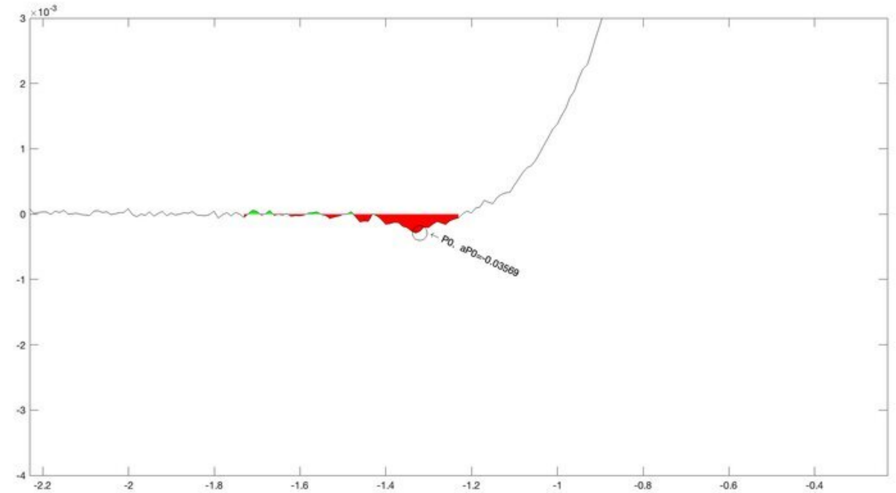
×10^-3
P0, aP0=-0.03569

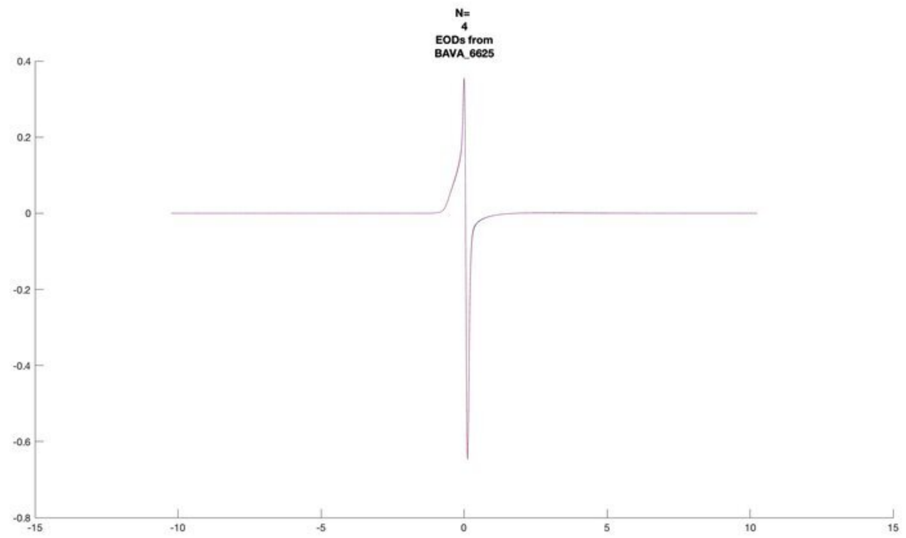
N=
4
EODs from
BAVA_6625

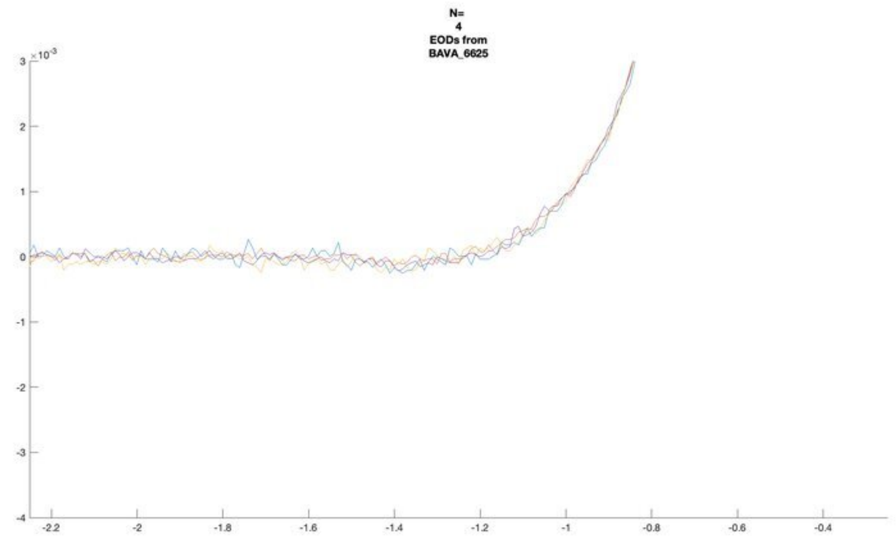
N=
4
EODs from
BAVA_6625

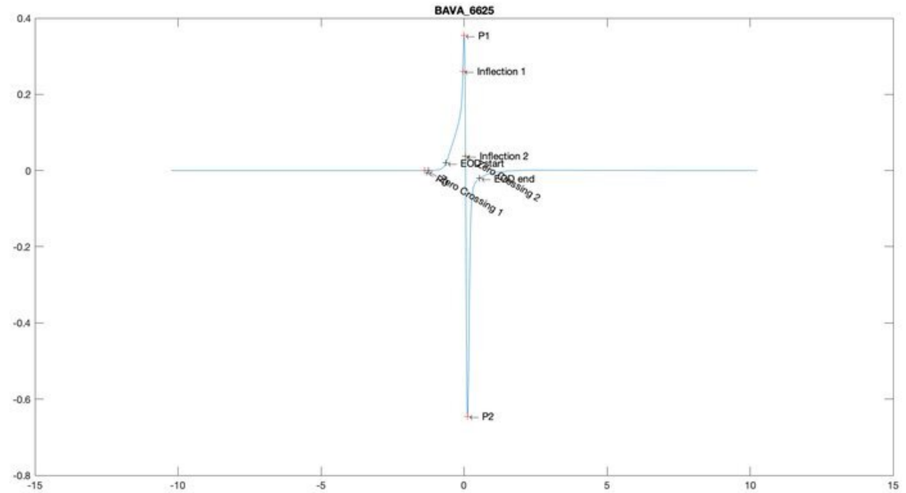
BAVA_6625
P1
Inflection 1
Inflection 2
EOD start
EOD end
Zero Crossing 1
Zero Crossing 2
P2

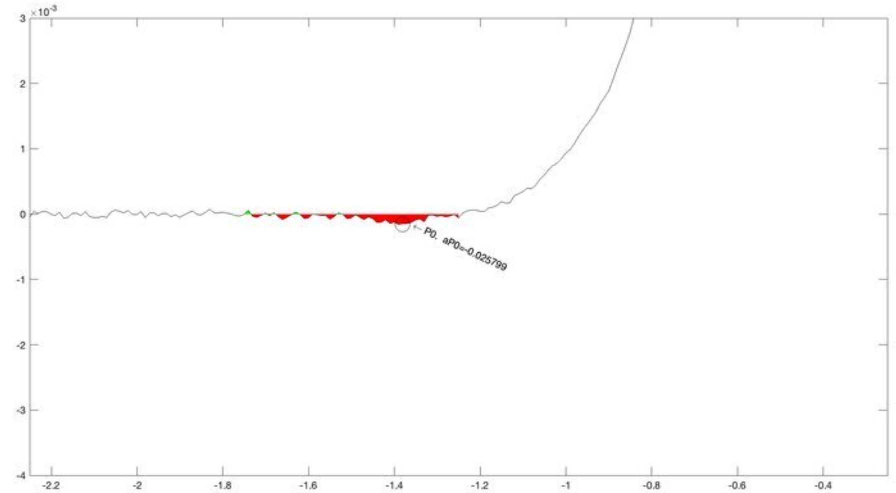
P0, aP0=-0.025799

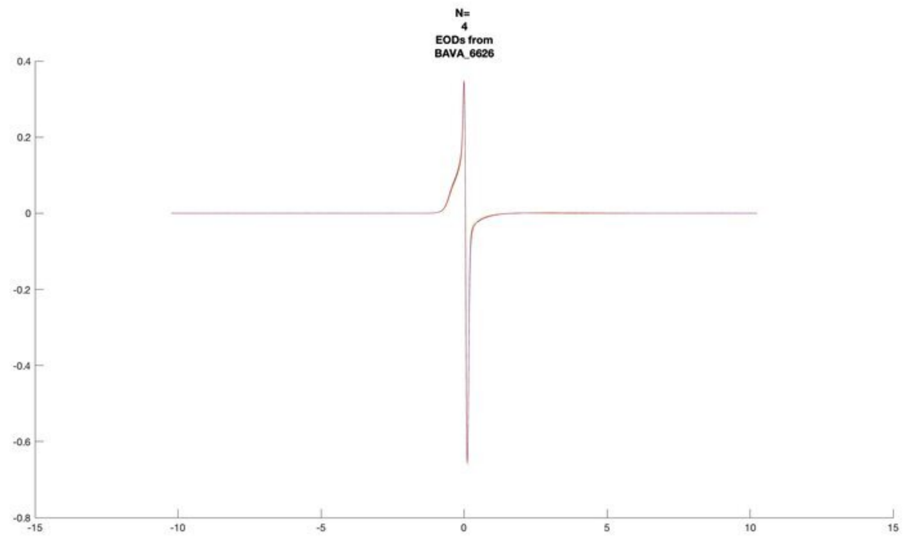
N=
4
EODs from
BAVA_6626

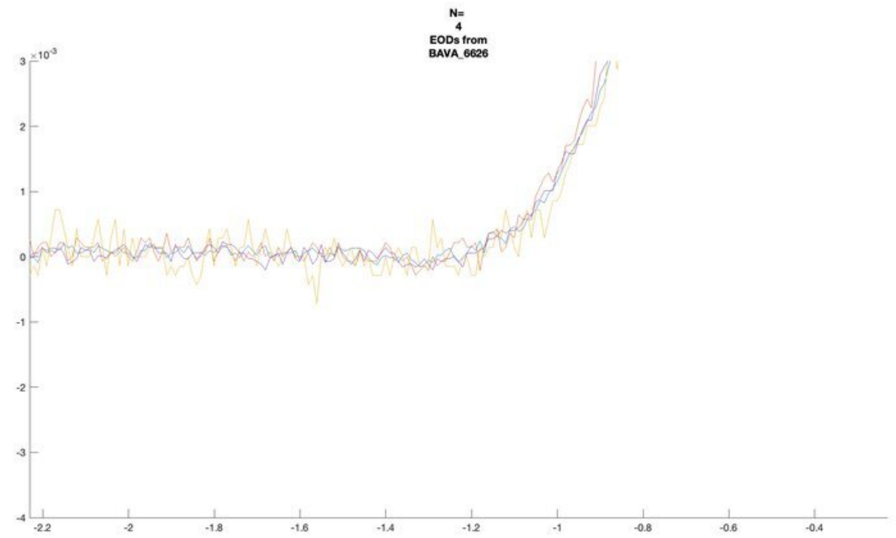
N=
4
EODs from
BAVA_6626

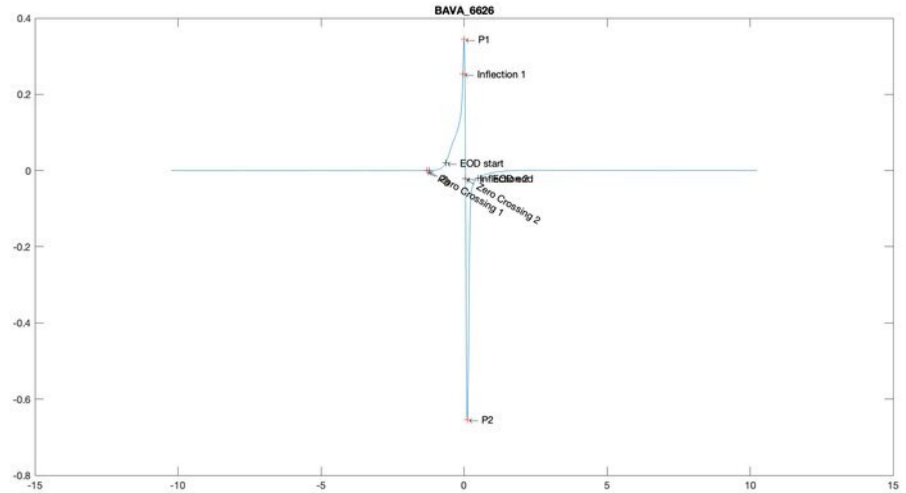
BAVA_6626
P1
Inflection 1
EOD start
Zero Crossing 1
Zero Crossing 2
P2

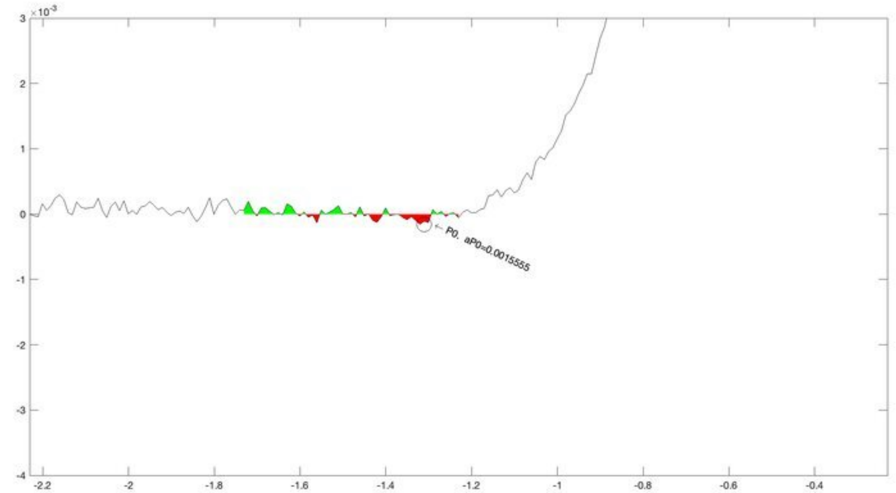
P0. aP0=0.0015555

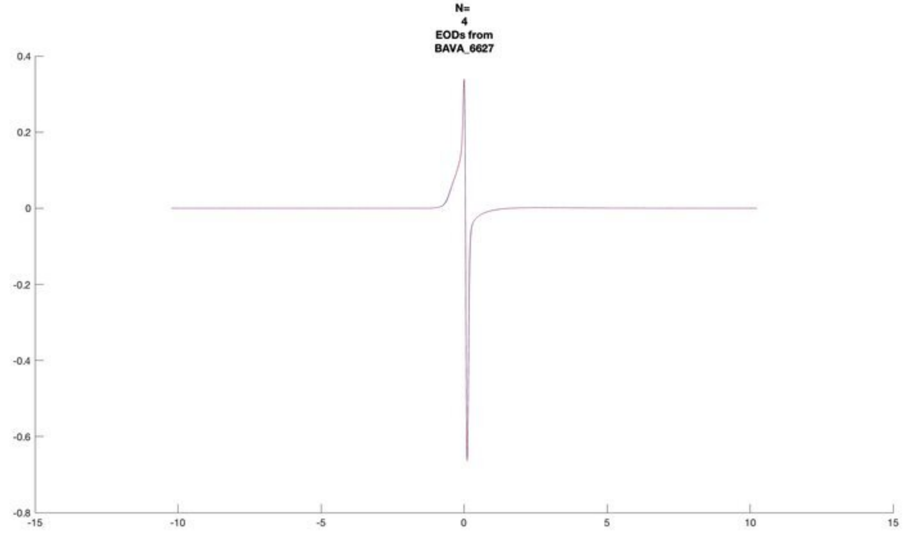
N=
4
EODs from
BAVA_6627

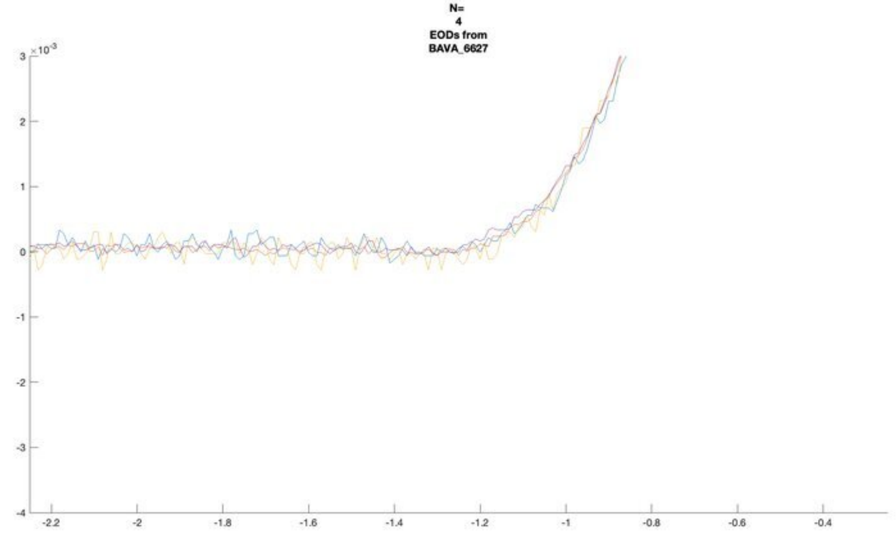
N=
4
EODs from
BAVA_6627

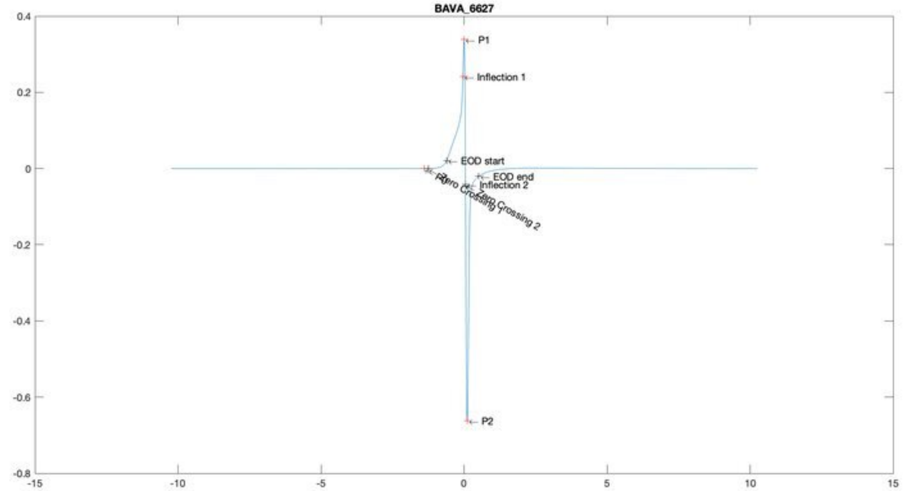
BAVA_6627
P1
Inflection 1
EOD start
EOD end
Inflection 2
Zero Crossing 2
P2

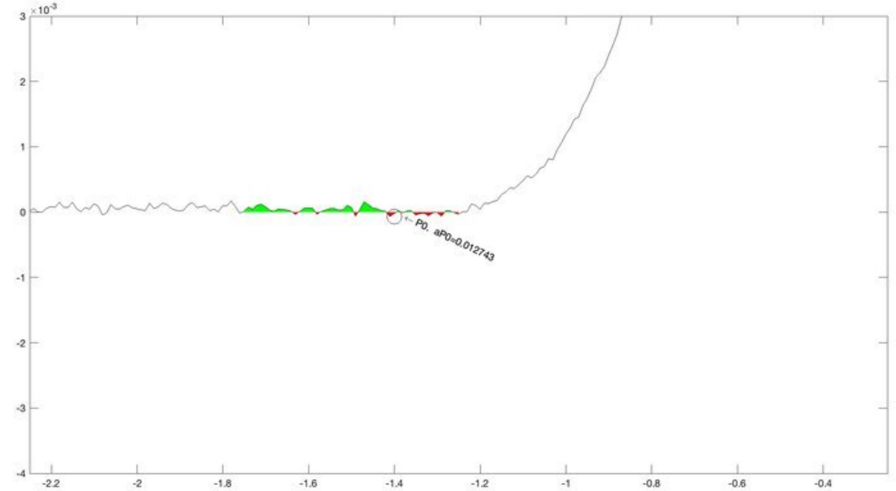
P0, aP0=0.012743

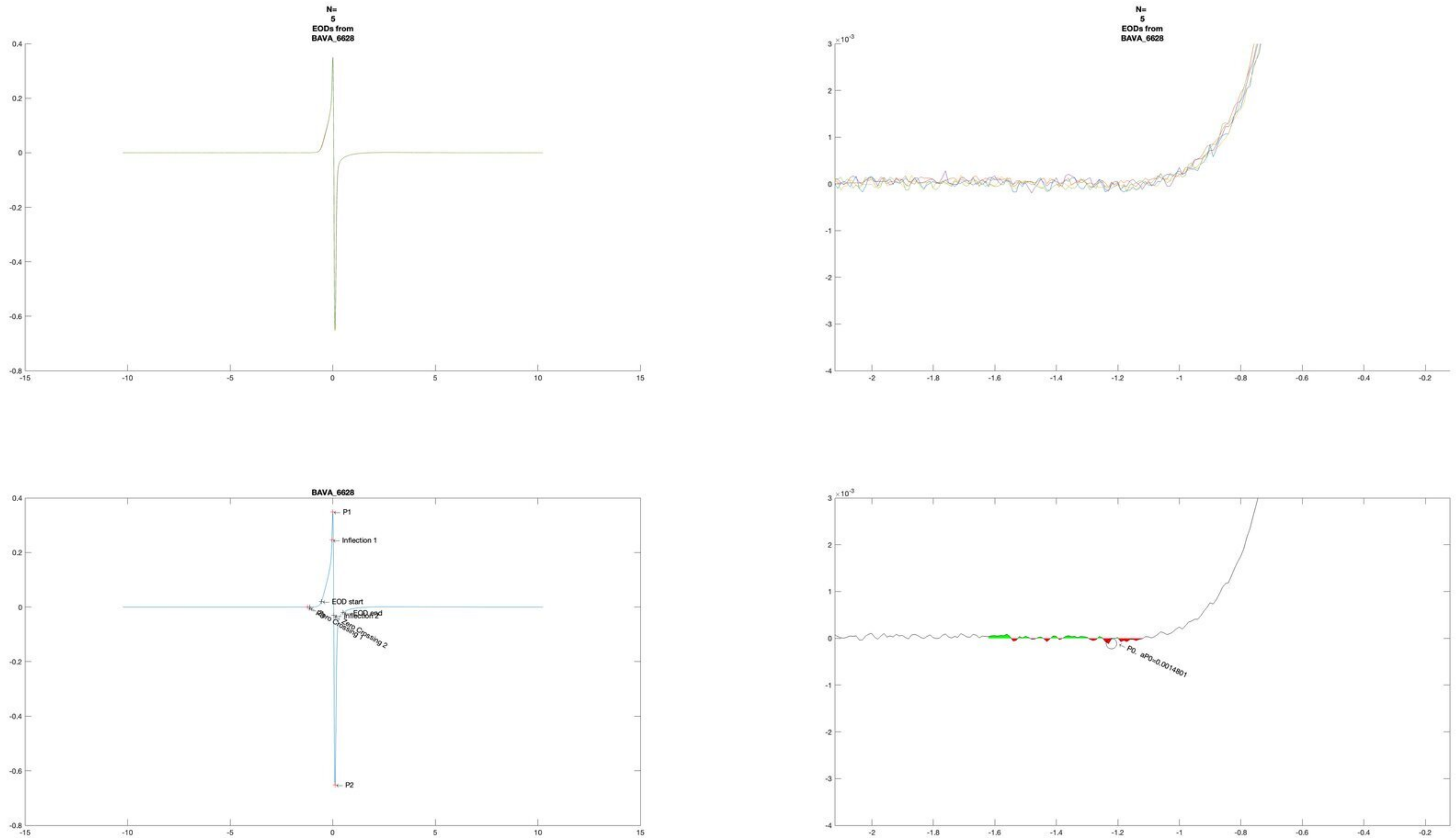
N=
5
EODs from
BAVA_6628
N=
5
EODs from
BAVA_6628
BAVA_6628
P1
Inflection 1
EOD start
EOD end
Inflection 2
Zero Crossing 1
Zero Crossing 2
P2
P0, aP0=0.0014801

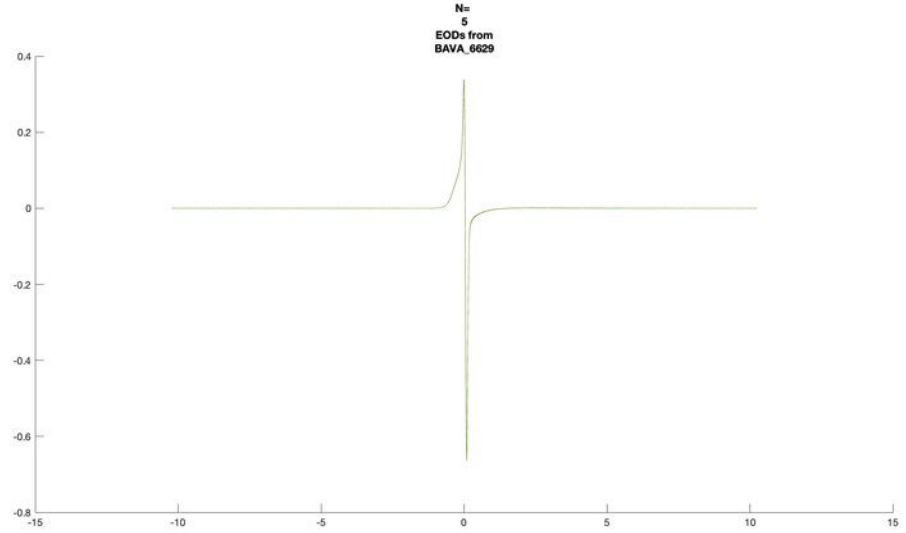
N=
5
EODs from
BAVA_6629

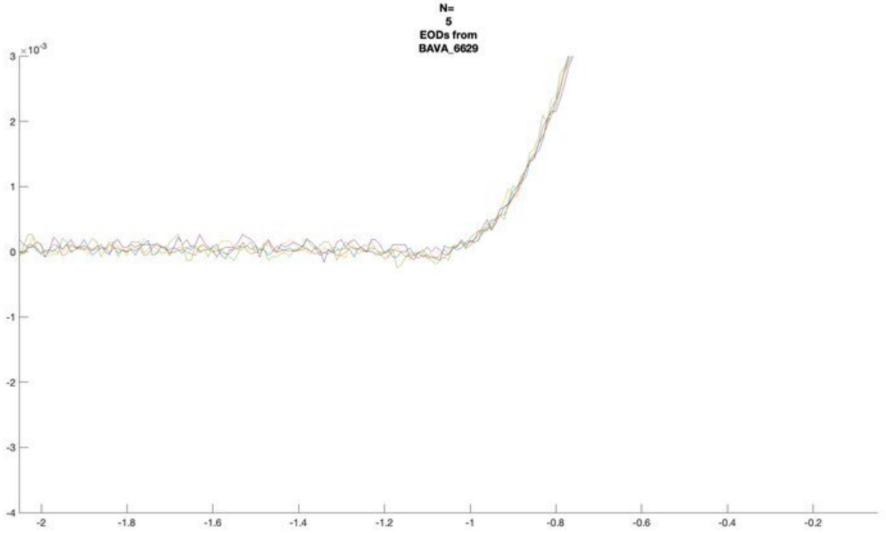
N=
5
EODs from
BAVA_6629

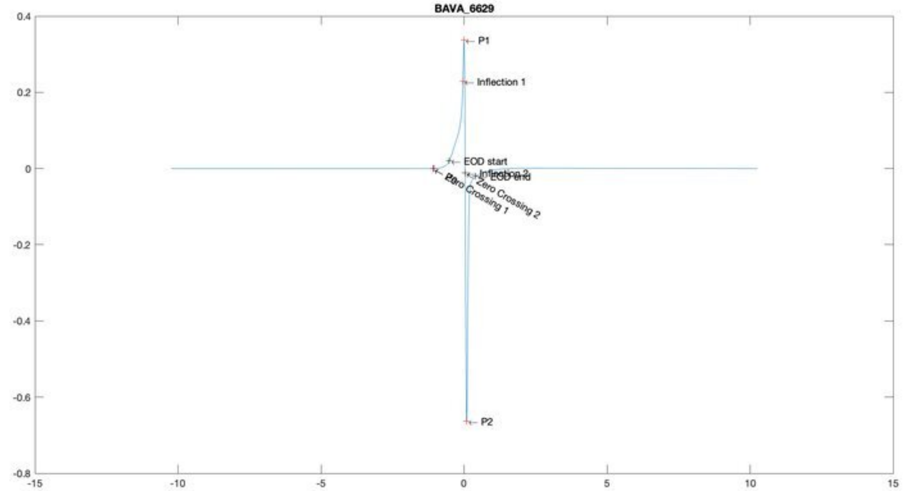
BAVA_6629
P1
Inflection 1
EOD start
Inflection 2
EOD end
Zero Crossing 1
Zero Crossing 2
P2

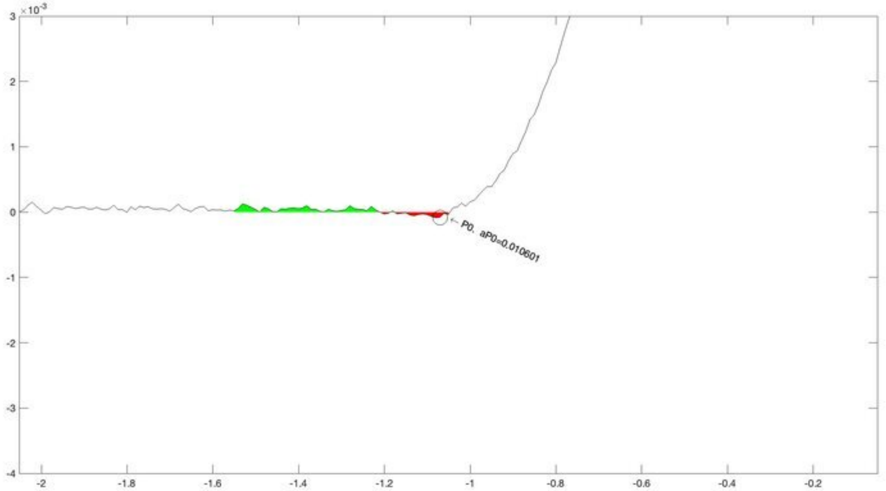
P0, aP0=0.010601

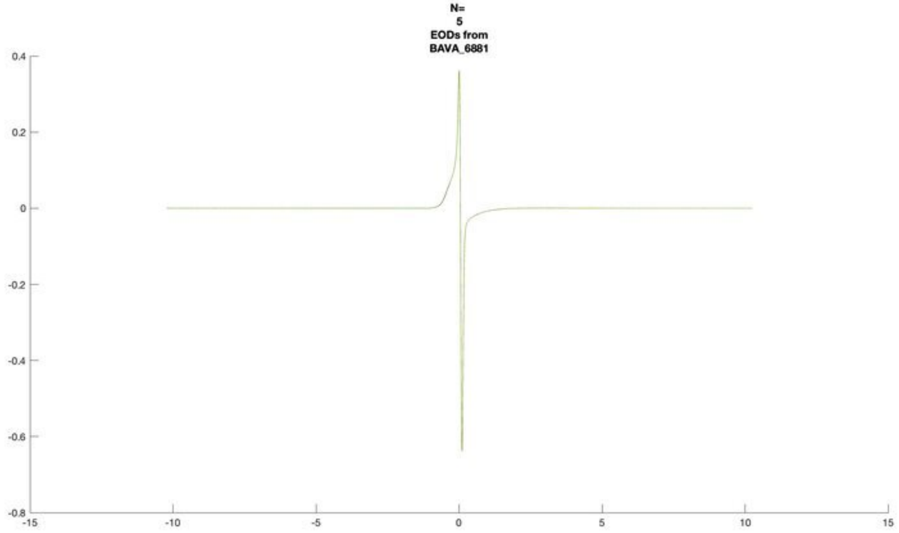
N=
5
EODs from
BAVA_6881

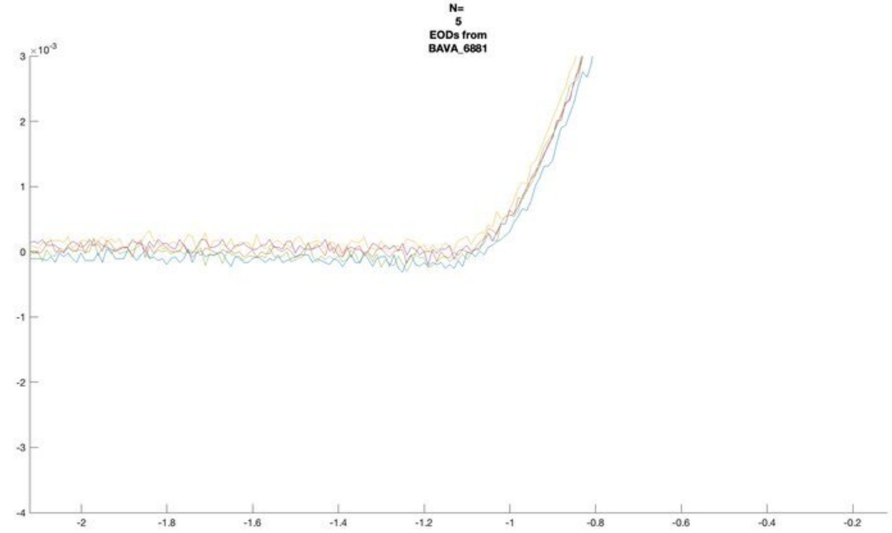
N=
5
EODs from
BAVA_6881
×10^-3

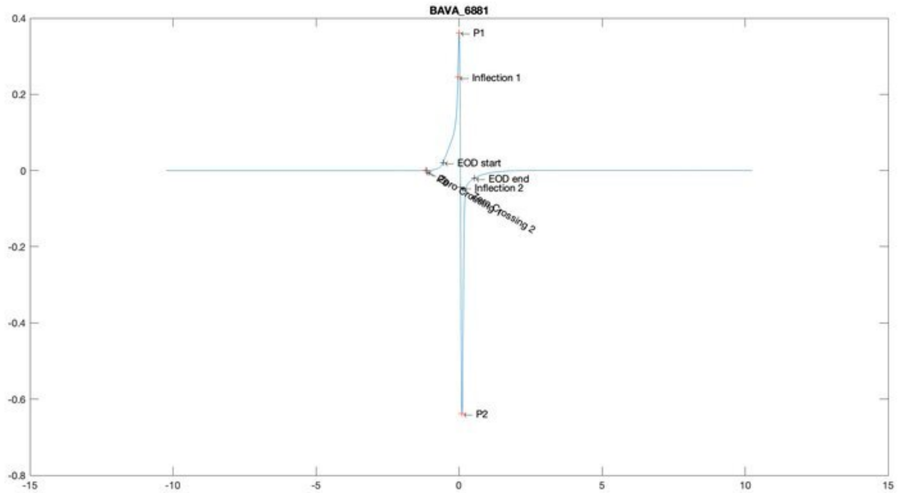
BAVA_6881
P1
Inflection 1
EOD start
EOD end
Inflection 2
Zero Crossing 2
P2

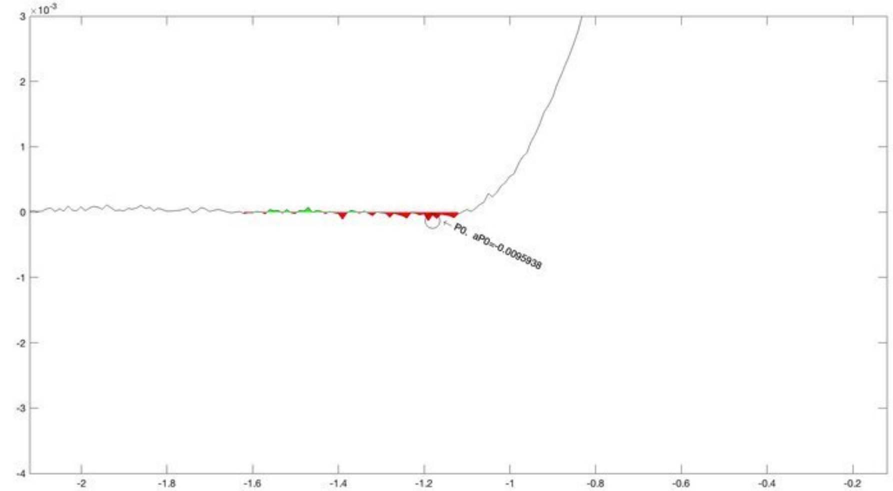
×10^-3
P0, aP0=-0.0095938

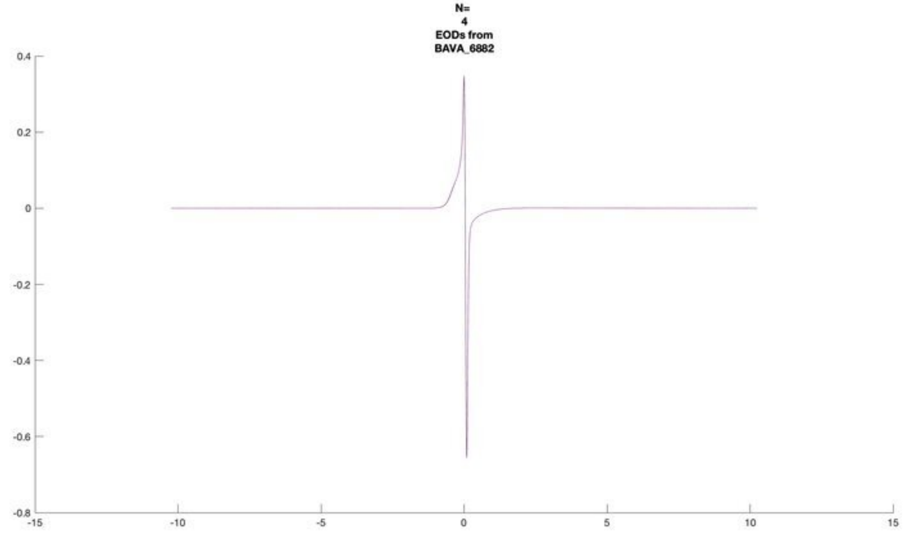
N=
4
EODs from
BAVA_6882

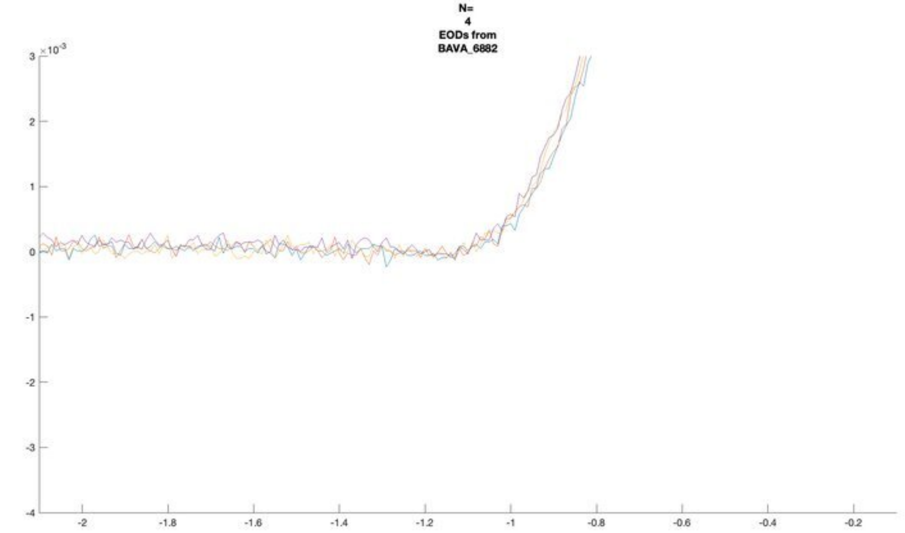
N=
4
EODs from
BAVA_6882
×10^-3

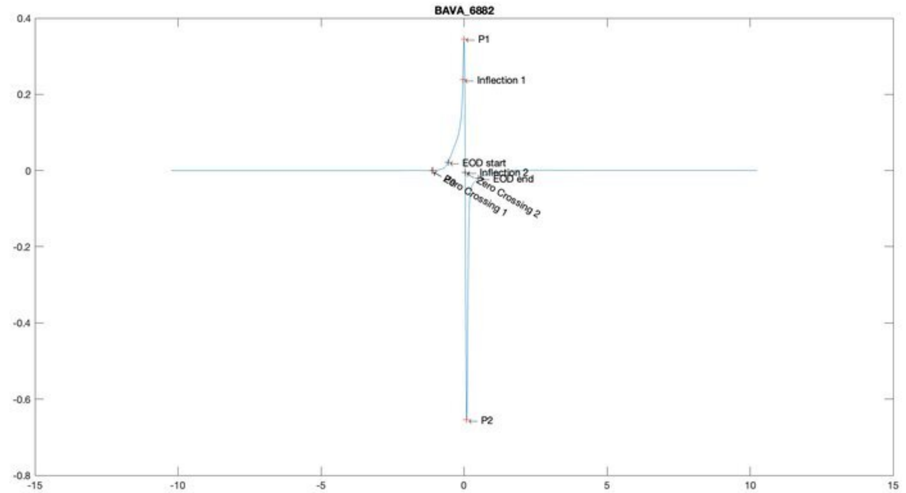
BAVA_6882
P1
Inflection 1
EOD start
Inflection 2
EOD end
Zero Crossing 2
Zero Crossing 1
P2

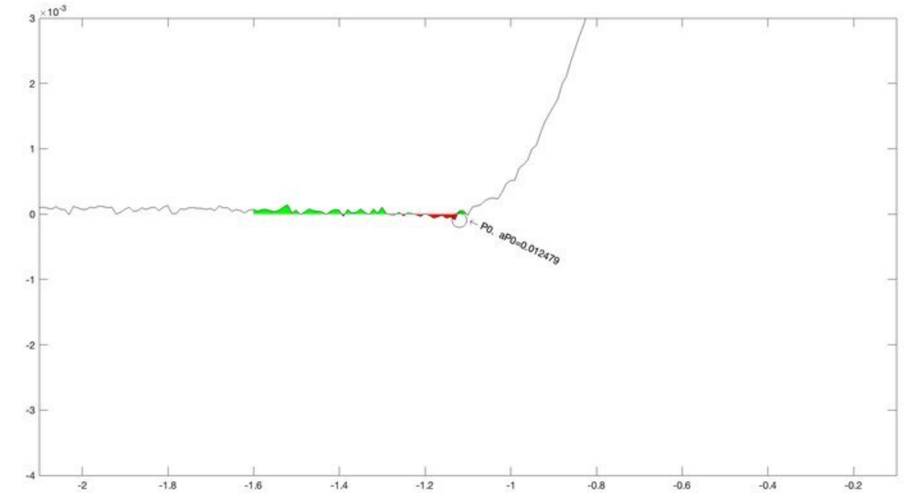
×10^-3
P0. aP0=0.012479

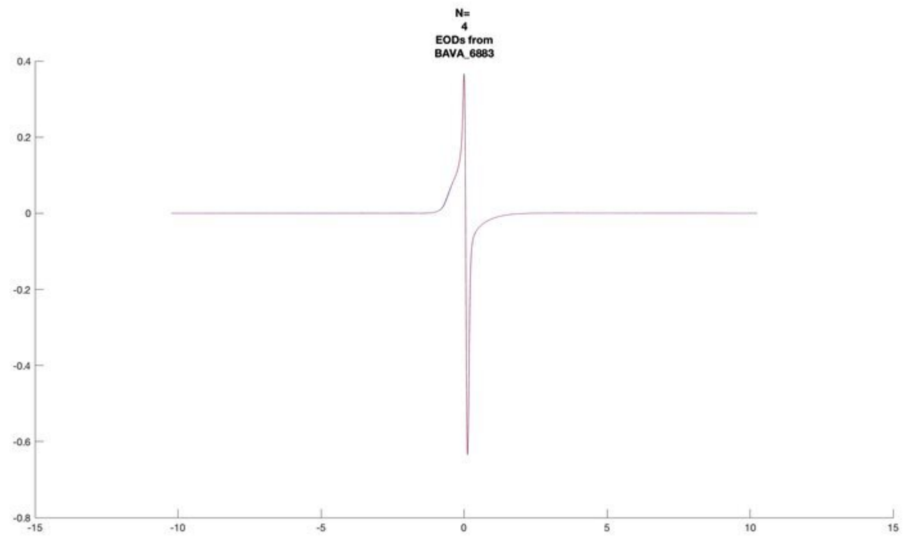
N=
4
EODs from
BAVA_6883

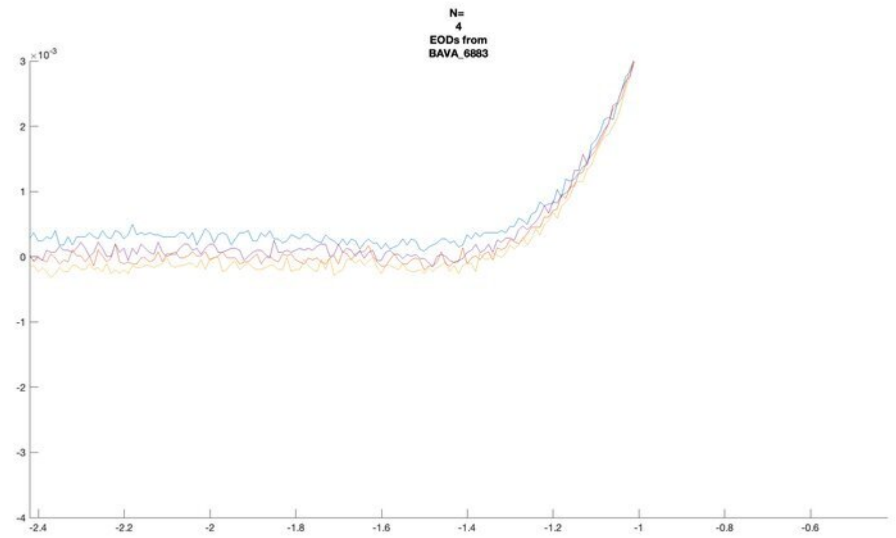
N=
4
EODs from
BAVA_6883

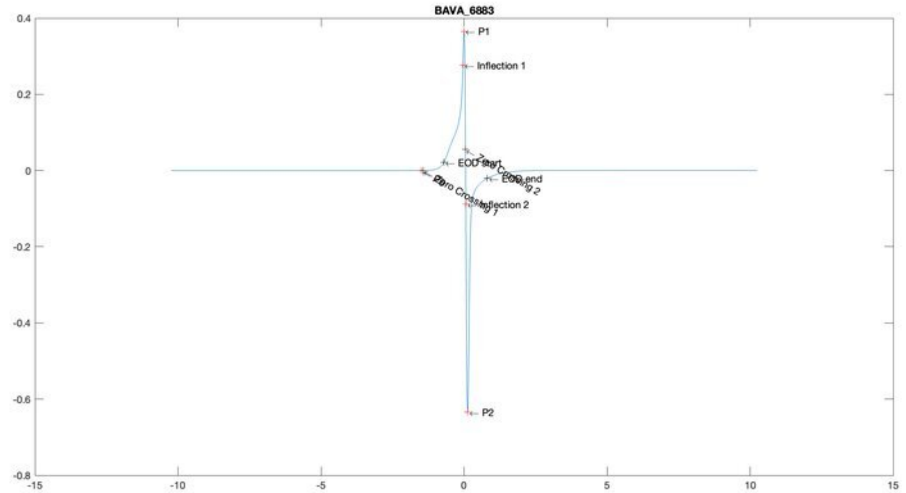
BAVA_6883
P1
Inflection 1
EOD end
P2

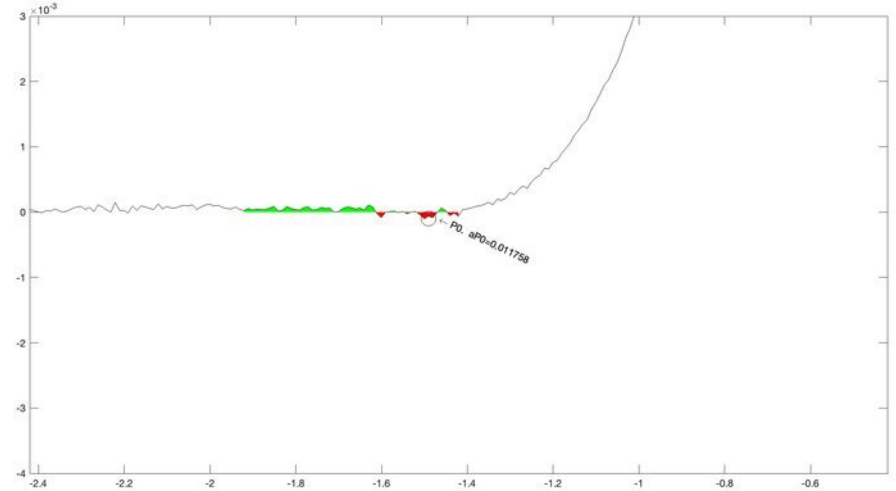
P0, aP0=0.011758

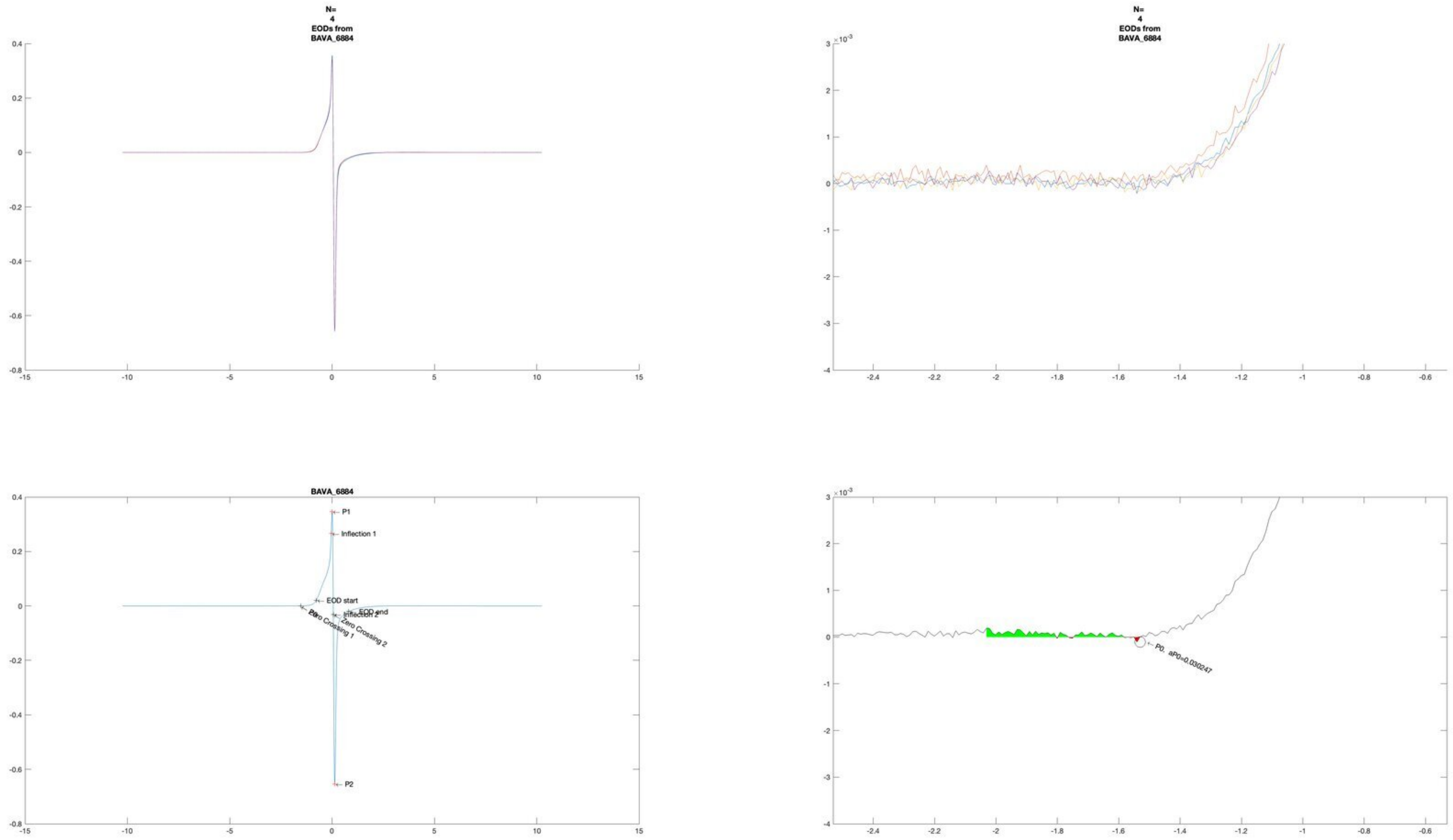

N=
4
EODs from
BAVA_6884
N=
4
EODs from
BAVA_6884
BAVA_6884
P1
Inflection 1
EOD start
EOD end
Inflection 2
Zero Crossing 1
Zero Crossing 2
P2
P0, aP0=0.030247

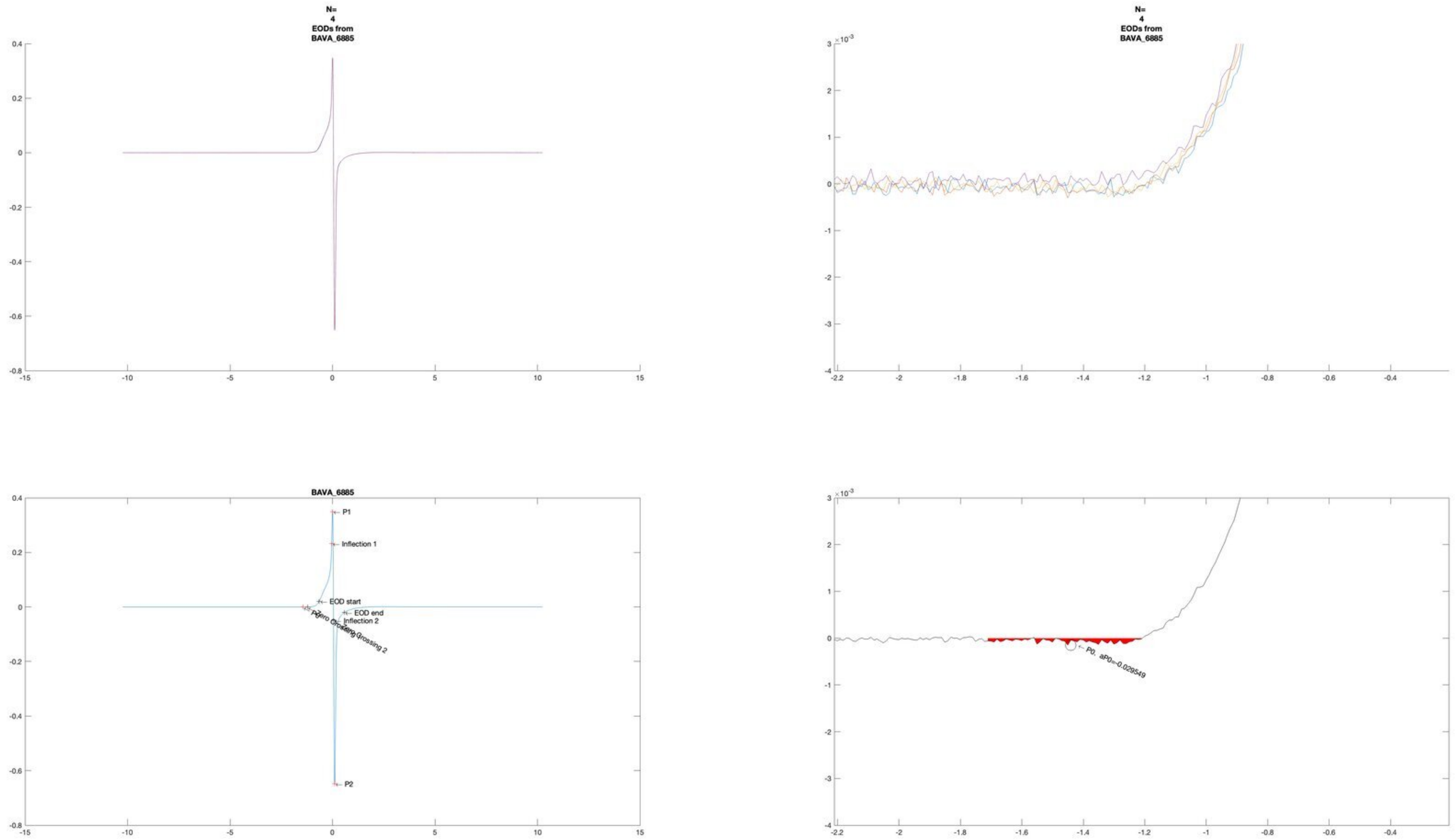
N=
4
EODs from
BAVA_6885
N=
4
EODs from
BAVA_6885
BAVA_6885
P1
Inflection 1
EOD start
EOD end
Inflection 2
Zero Crossing 2
P2
P0, aP0=-0.029549

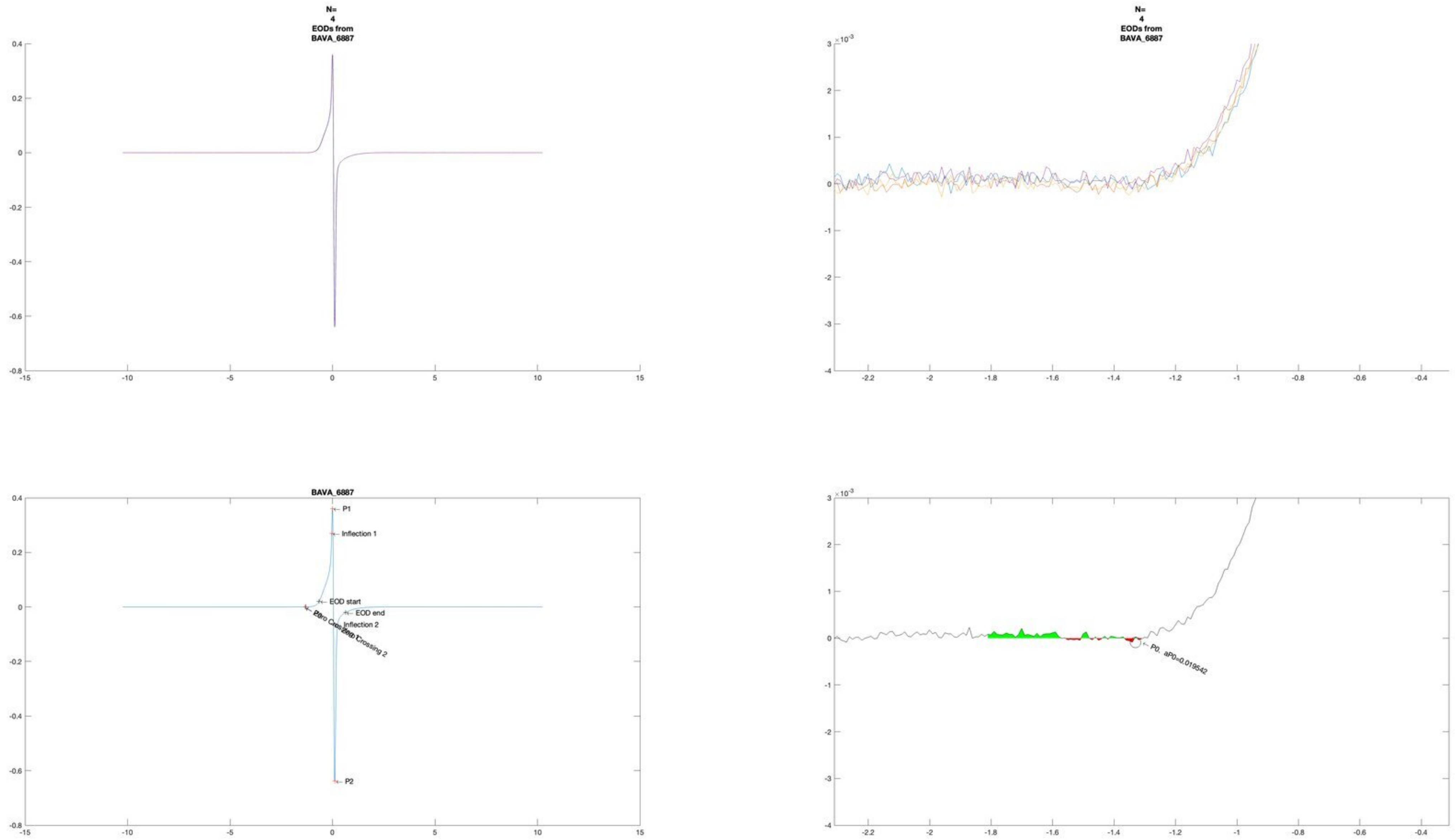
N=
4
EODs from
BAVA_6887
N=
4
EODs from
BAVA_6887
BAVA_6887
P1
Inflection 1
EOD start
EOD end
Inflection 2
Zero Crossing 2
P2
P0. aP0=0.019542

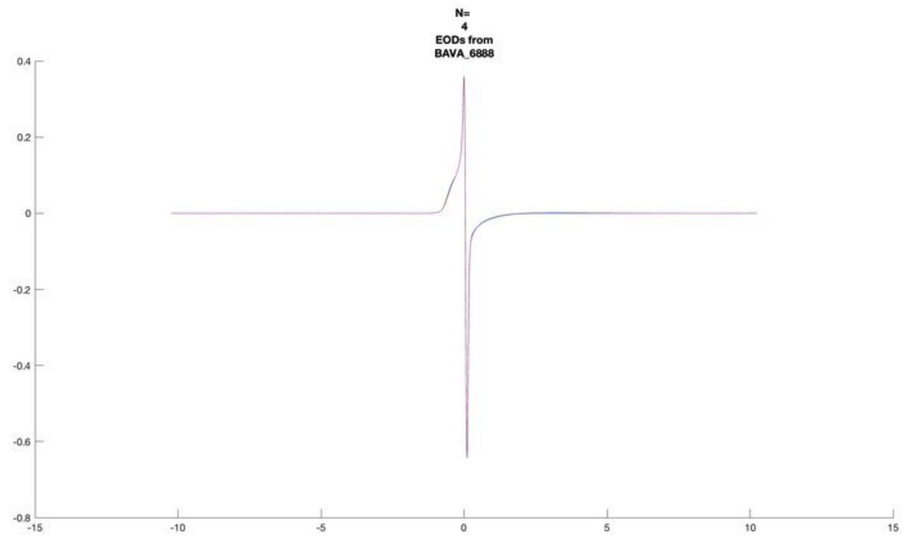
N=
4
EODs from
BAVA_6888

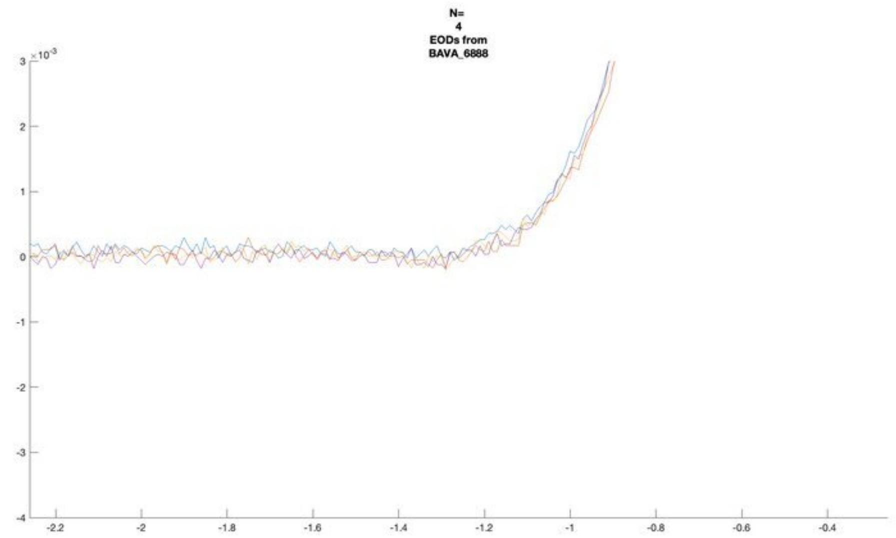
N=
4
EODs from
BAVA_6888
×10^-3

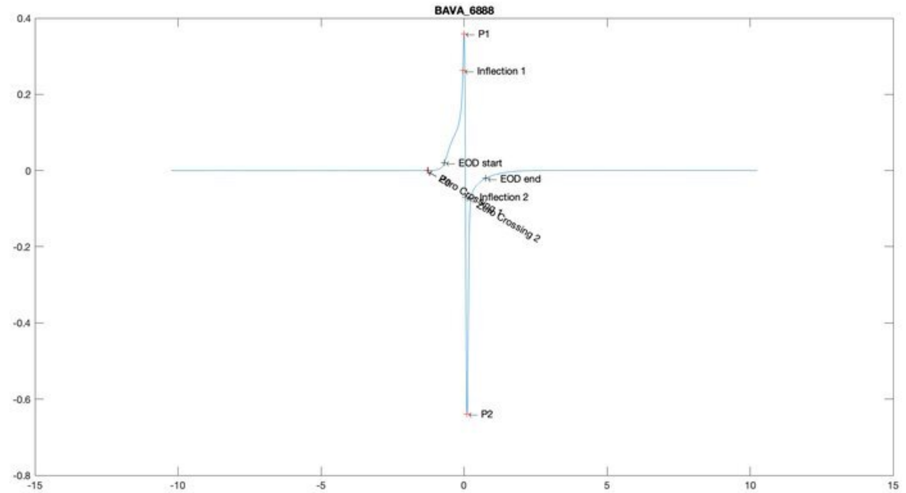
BAVA_6888
P1
Inflection 1
EOD start
EOD end
Inflection 2
Zero Crossing 2
P2

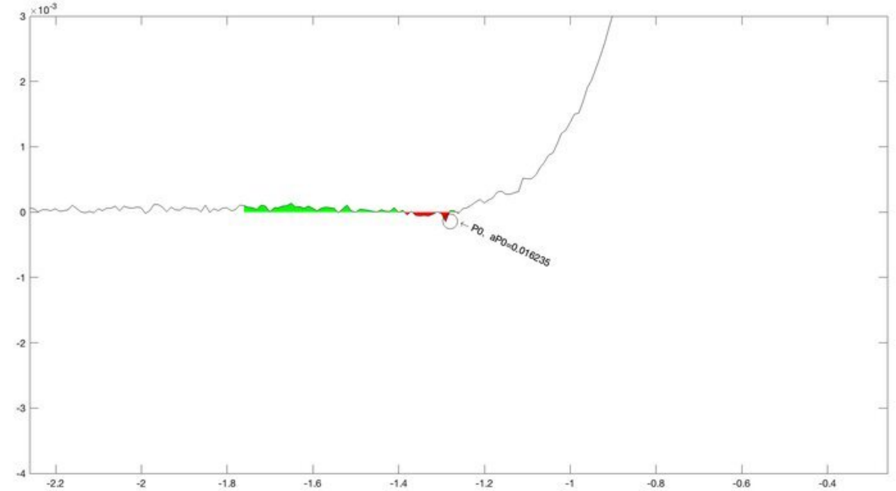
×10^-3
P0, aP0=0.016235

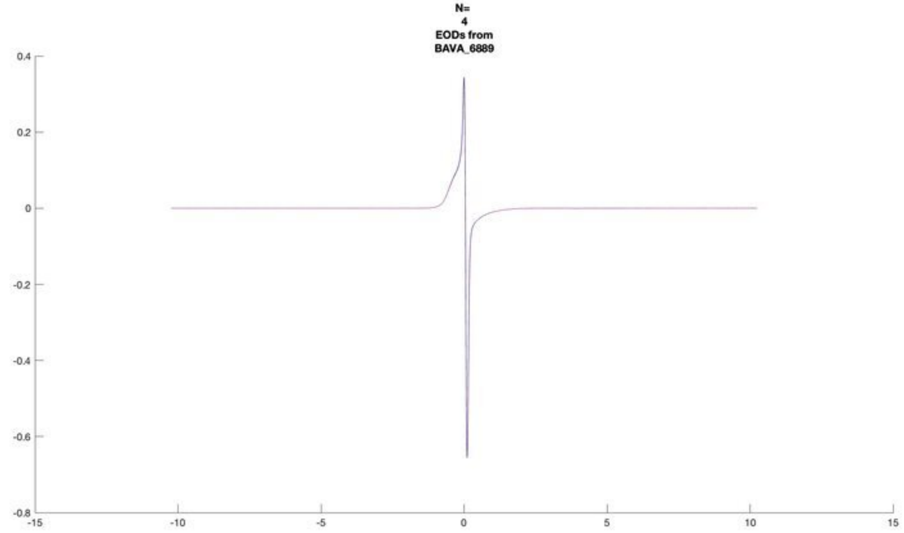
N=
4
EODs from
BAVA_6889

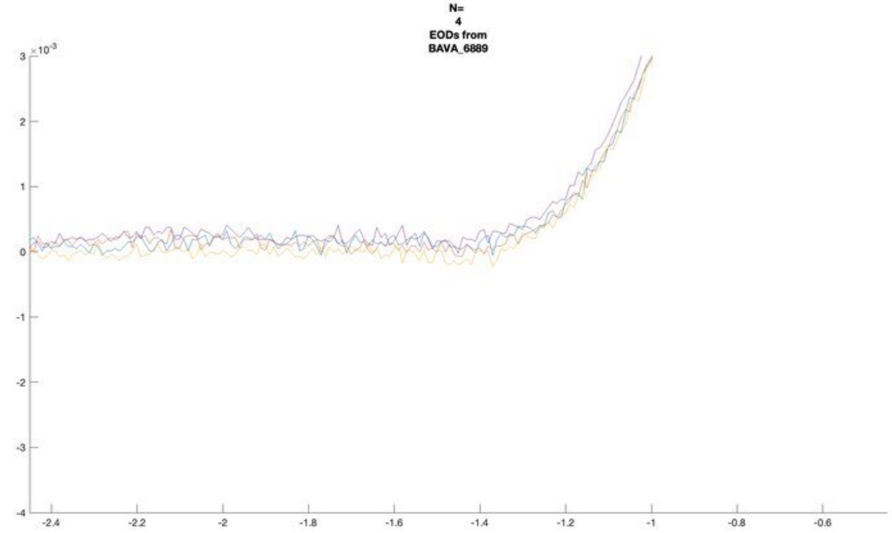
N=
4
EODs from
BAVA_6889

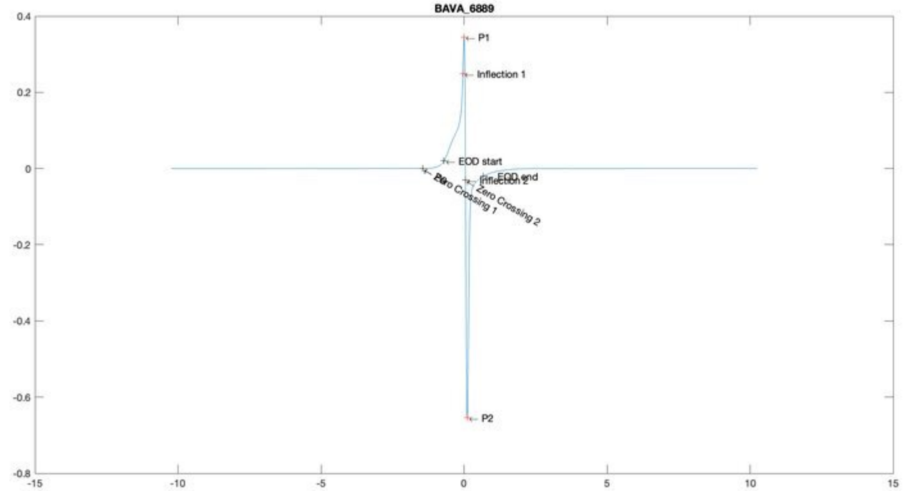
BAVA_6889
P1
Inflection 1
EOD start
EOD end
Inflection 2
P0
Zero Crossing 1
Zero Crossing 2
P2

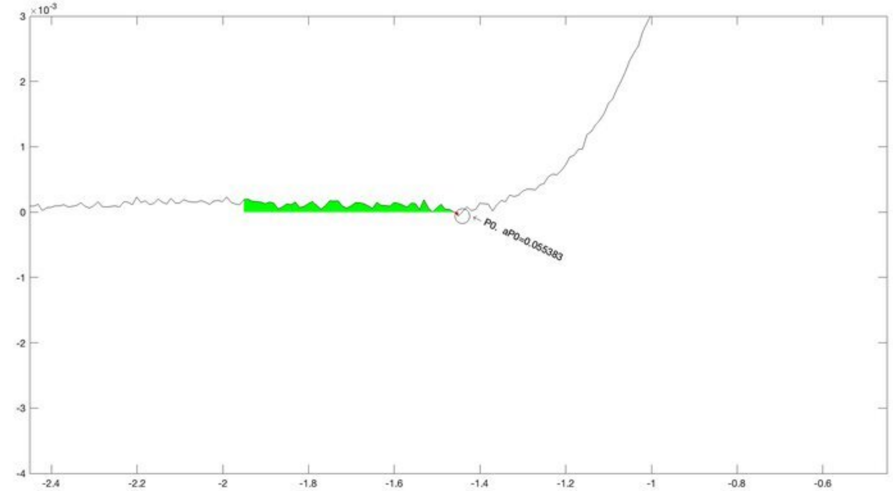
P0, aP0=0.055383

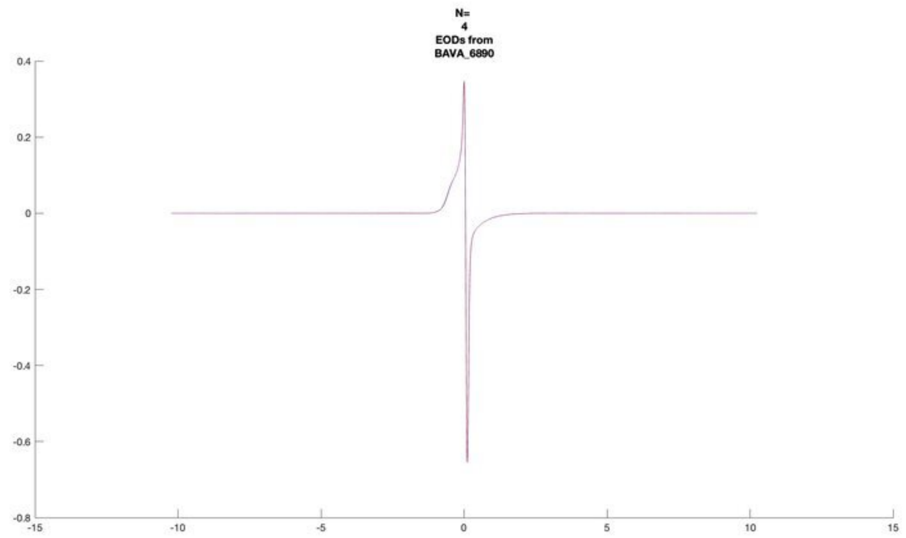

N=
4
EODs from
BAVA_6890


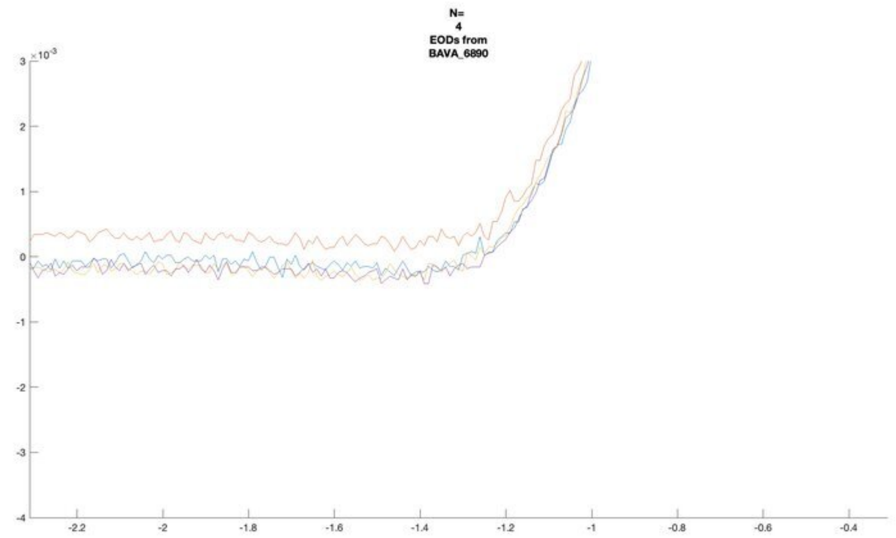

N=
4
EODs from
BAVA_6890


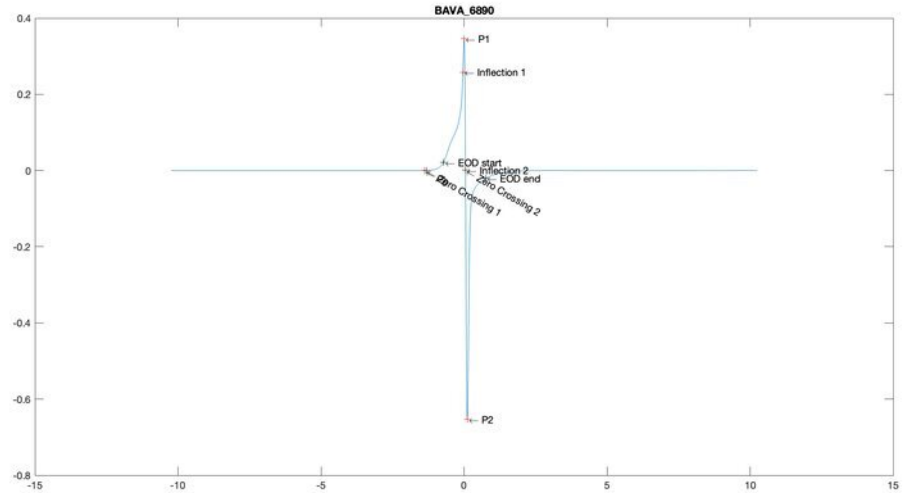

BAVA_6890
P1
Inflection 1
EOD start
Inflection 2
EOD end
Zero Crossing 1
Zero Crossing 2
P2


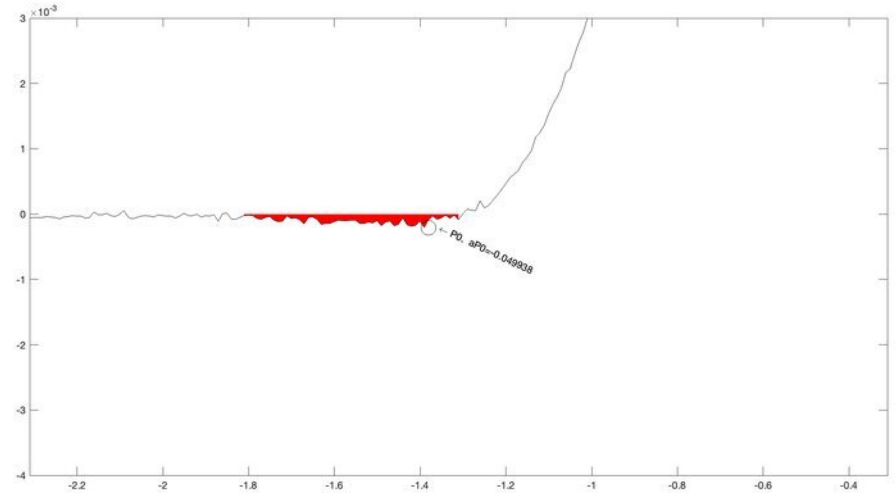

P0, aP0=-0.049938

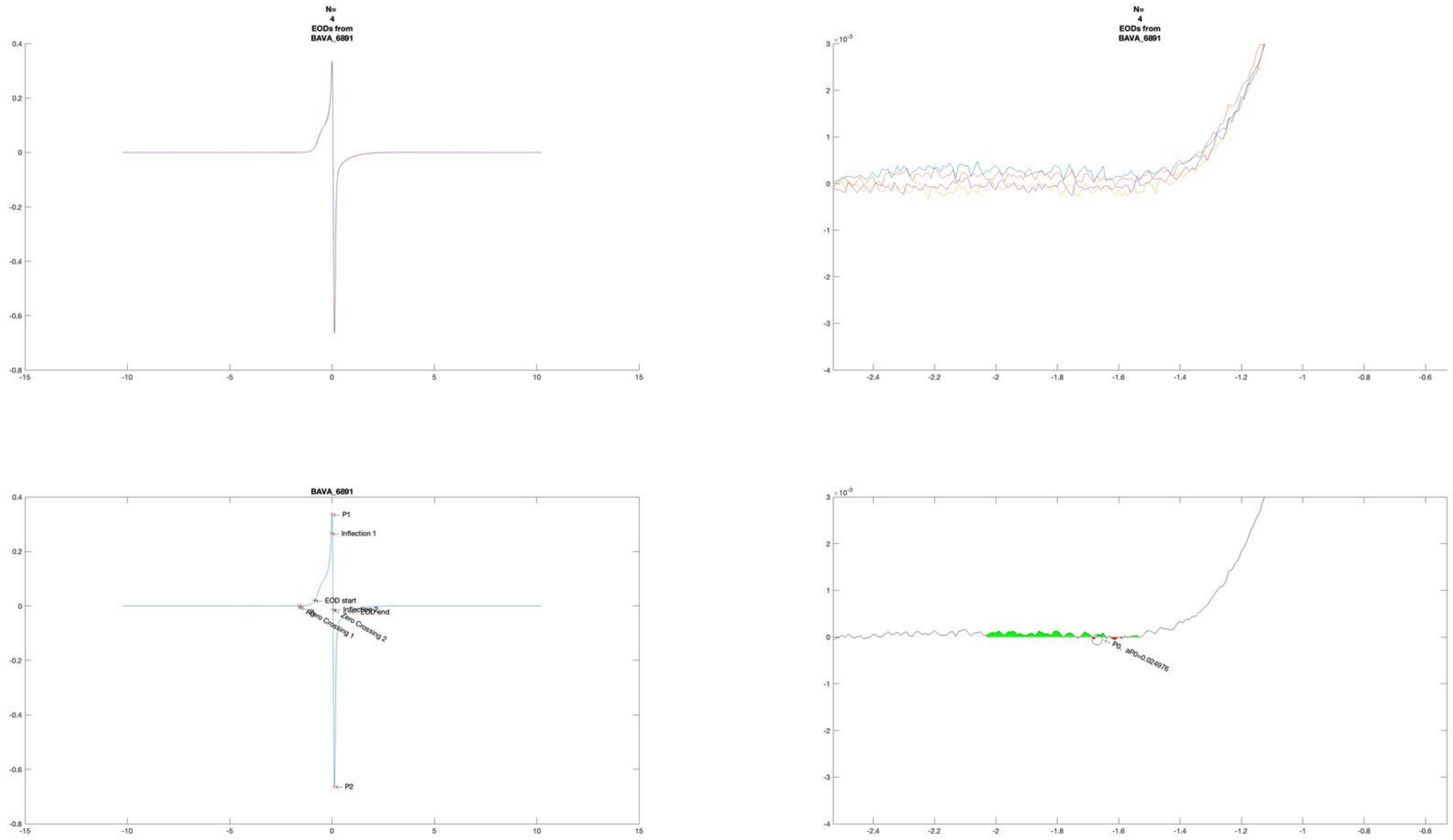
N=
4
EODs from
BAVA_6891
N=
4
EODs from
BAVA_6891
BAVA_6891
P1
Inflection 1
EOD start
Inflection 2
EOD end
Zero Crossing 1
Zero Crossing 2
P2
P0, aP0=0.024976

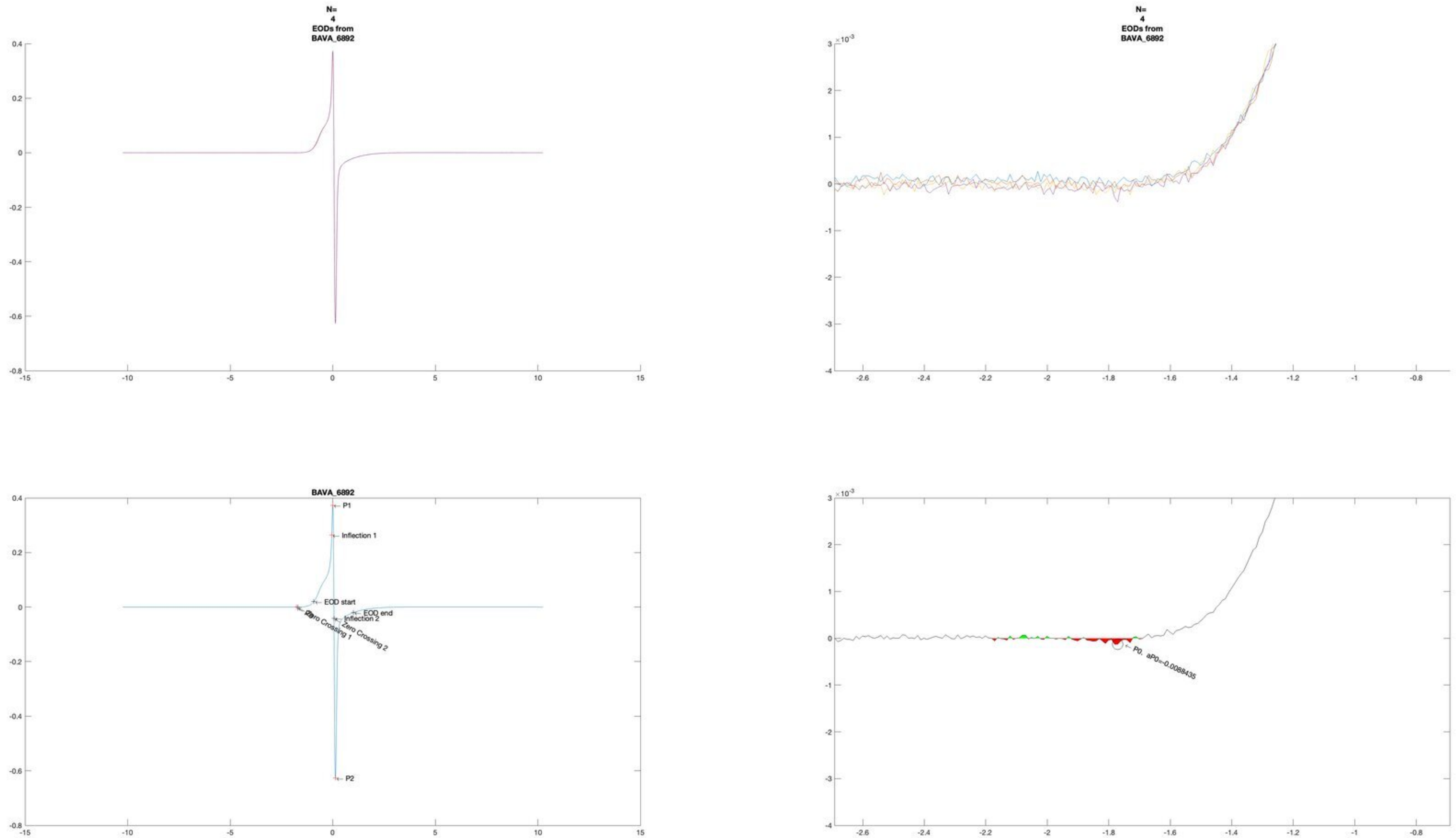
N=
4
EODs from
BAVA_6892
N=
4
EODs from
BAVA_6892
BAVA_6892
P1
Inflection 1
EOD start
EOD end
Inflection 2
Zero Crossing 1
Zero Crossing 2
P2
P0, aP0=-0.008435

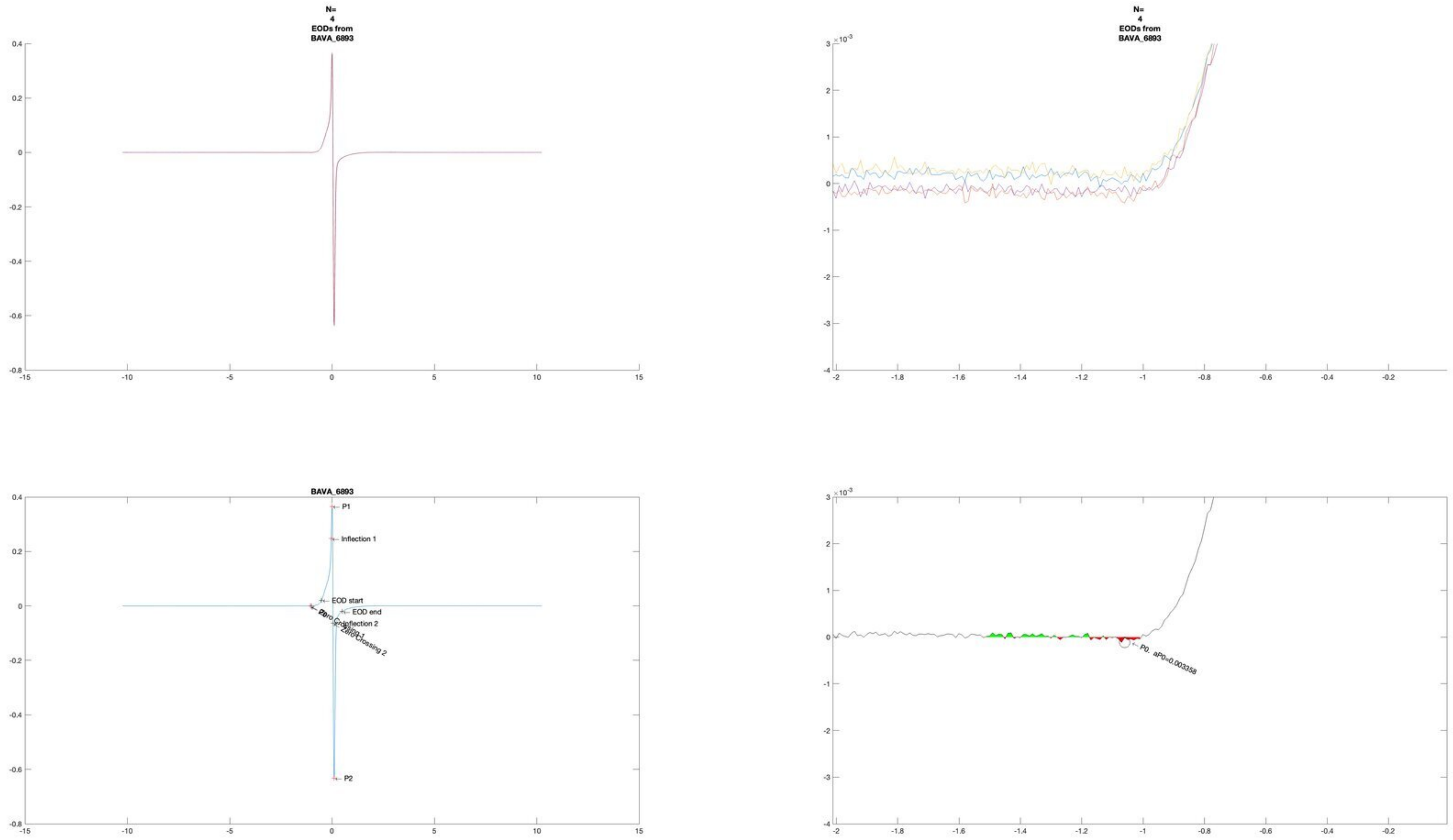
N=
4
EODs from
BAVA_6893
N=
4
EODs from
BAVA_6893
BAVA_6893
P1
Inflection 1
EOD start
EOD end
Inflection 2
Zero Crossing 2
P2
P0. aP0=0.003358

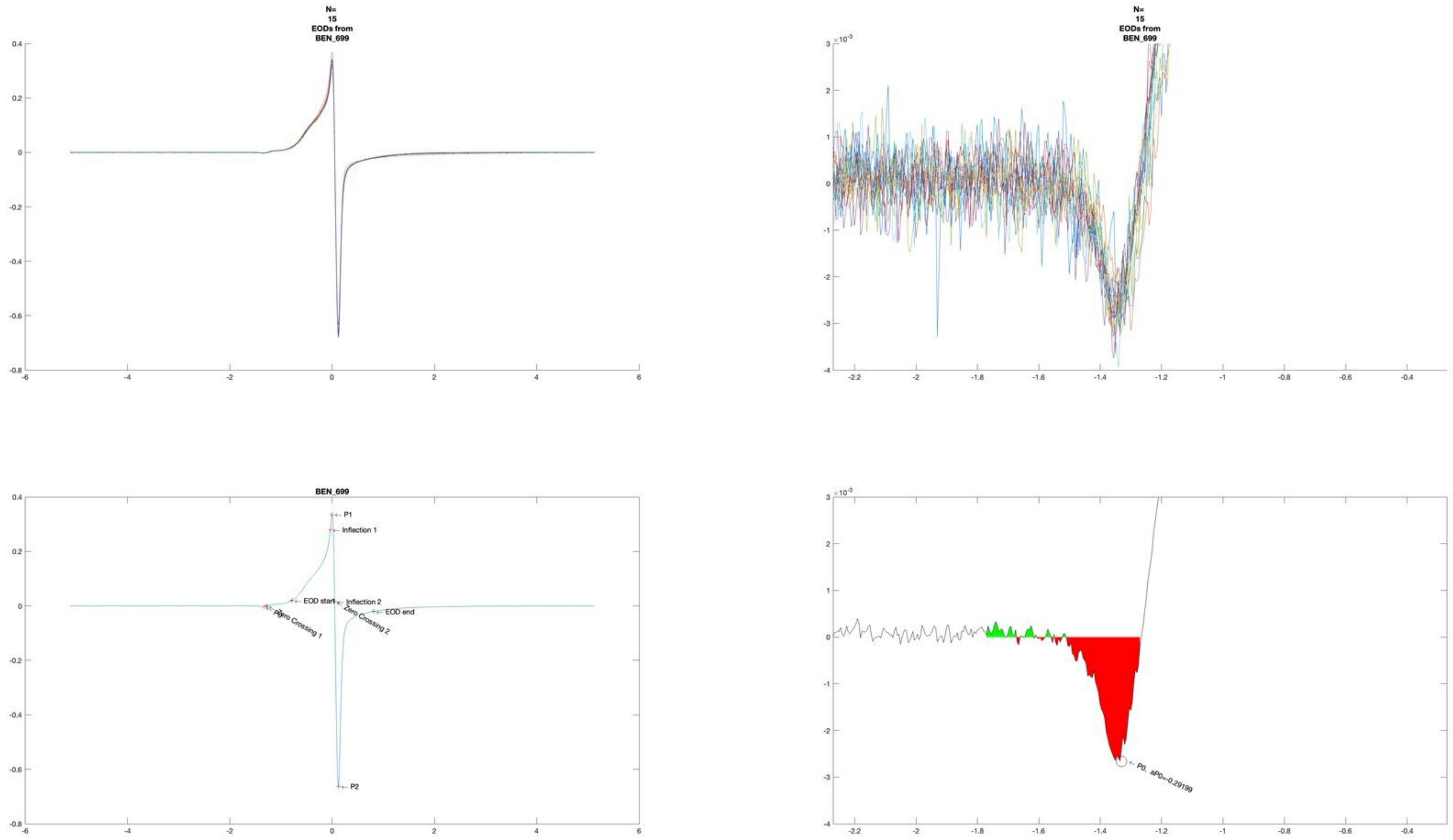

N=
15
EODs from
BEN_699
N=
15
EODs from
BEN_699
BEN_699
P1
Inflection 1
EOD start
Inflection 2
EOD end
P0
Zero Crossing 1
Zero Crossing 2
P2
P0, aP0=-0.29199

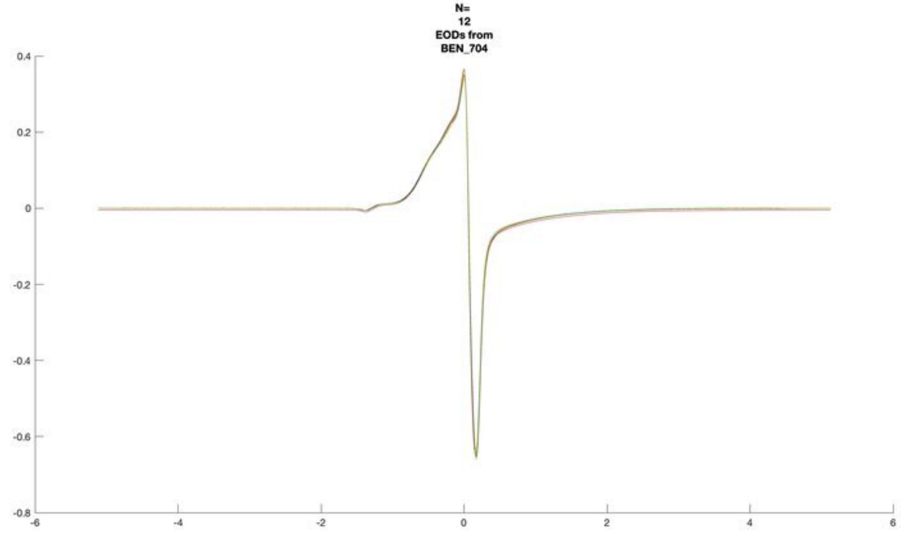
N=
12
EODs from
BEN_704

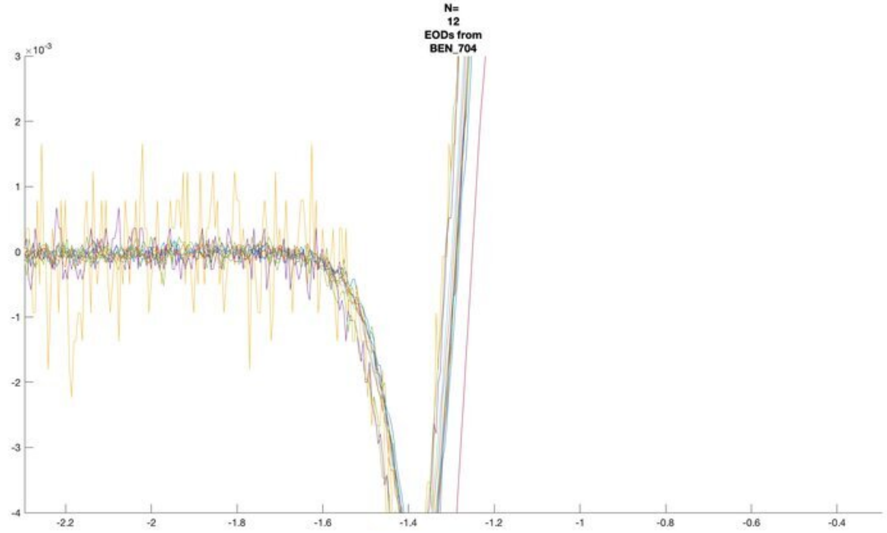
N=
12
EODs from
BEN_704
×10^-3

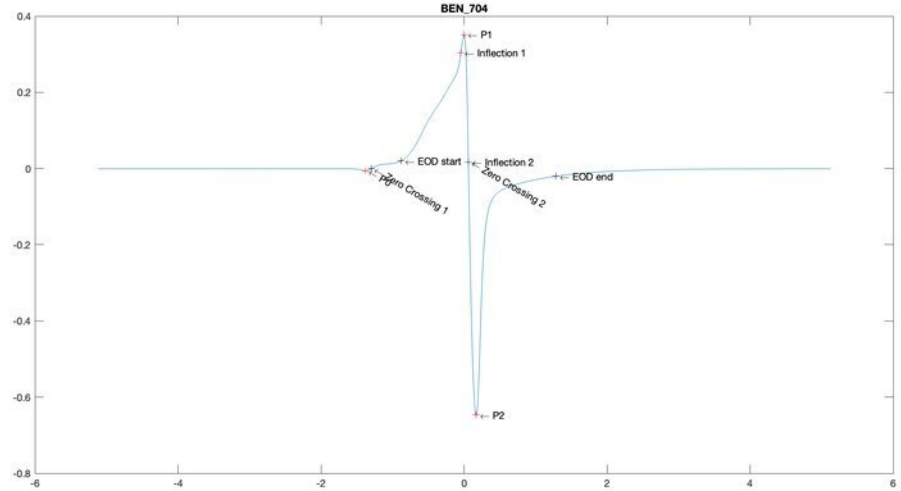
BEN_704
P1
Inflection 1
EOD start
Inflection 2
EOD end
P0
Zero Crossing 1
Zero Crossing 2
P2

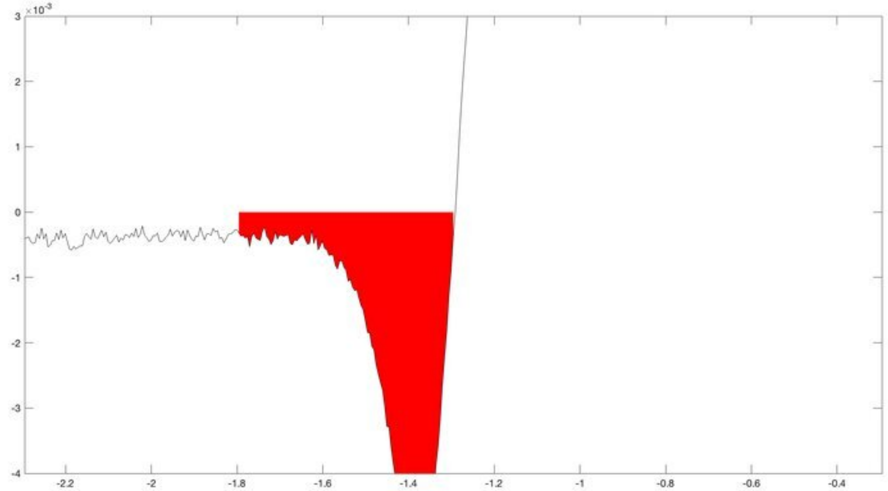
×10^-3

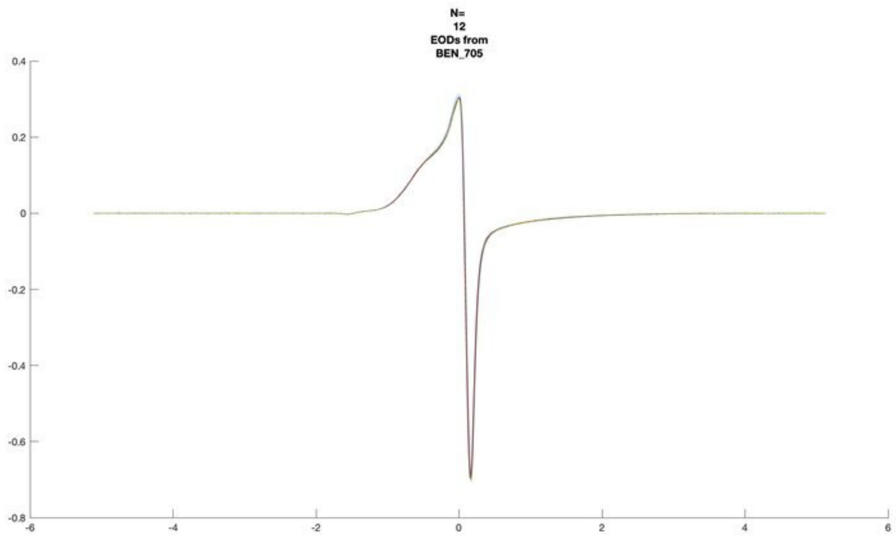
N=
12
EODs from
BEN_705

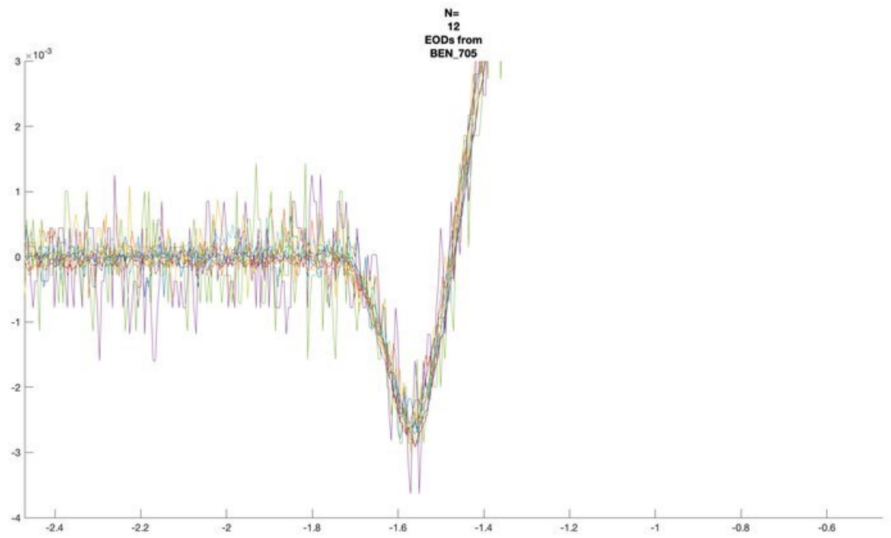
N=
12
EODs from
BEN_705
×10^-3

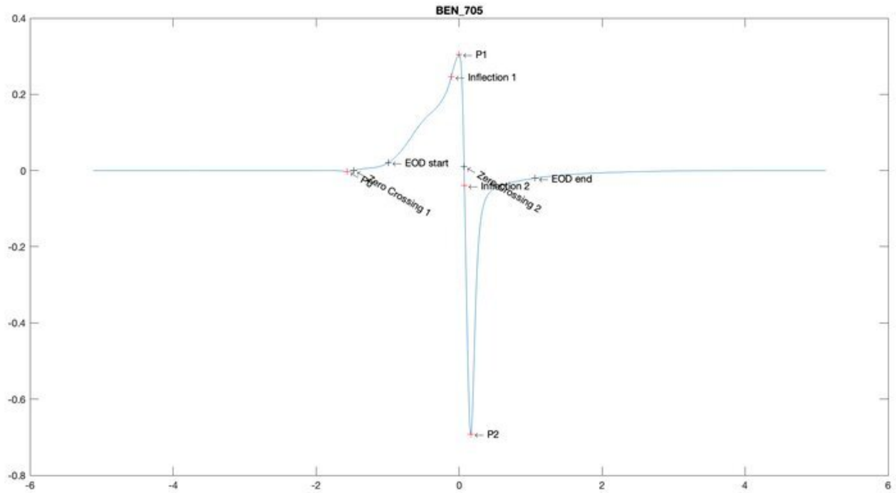
BEN_705
P1
Inflection 1
EOD start
P0
Zero Crossing 1
Zero Crossing 2
Inflection 2
EOD end
P2

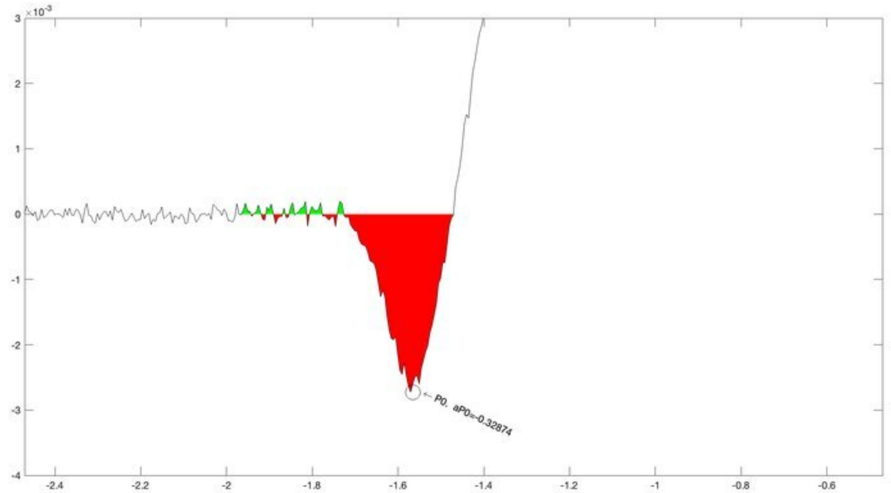
×10^-3
P0. aP0=-0.32874

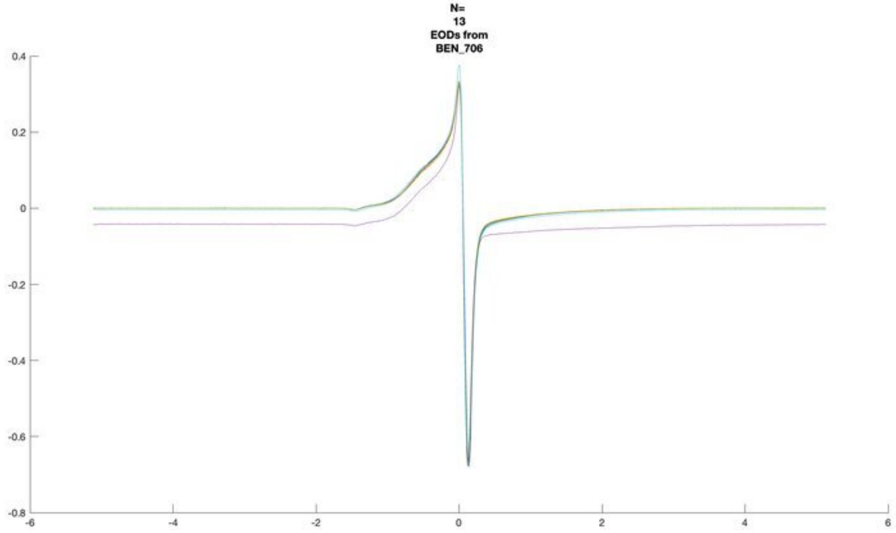
N=
13
EODs from
BEN_706

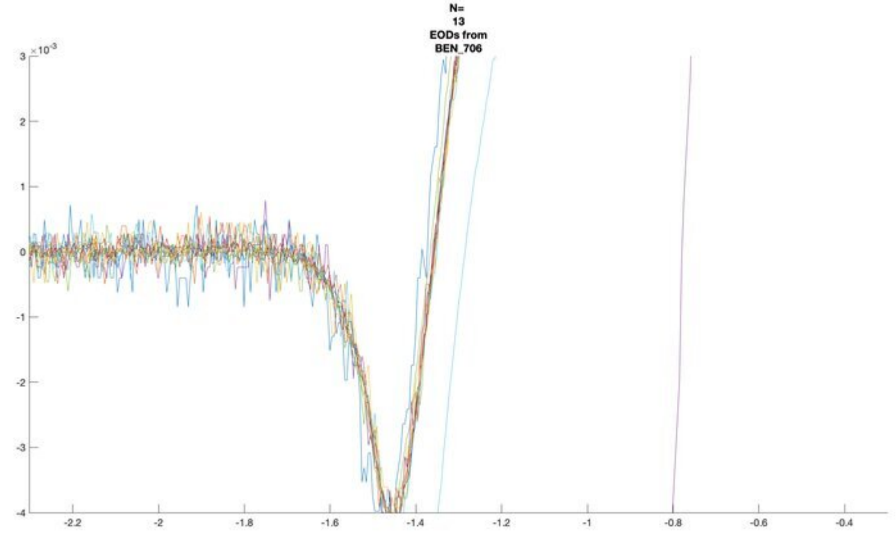
N=
13
EODs from
BEN_706

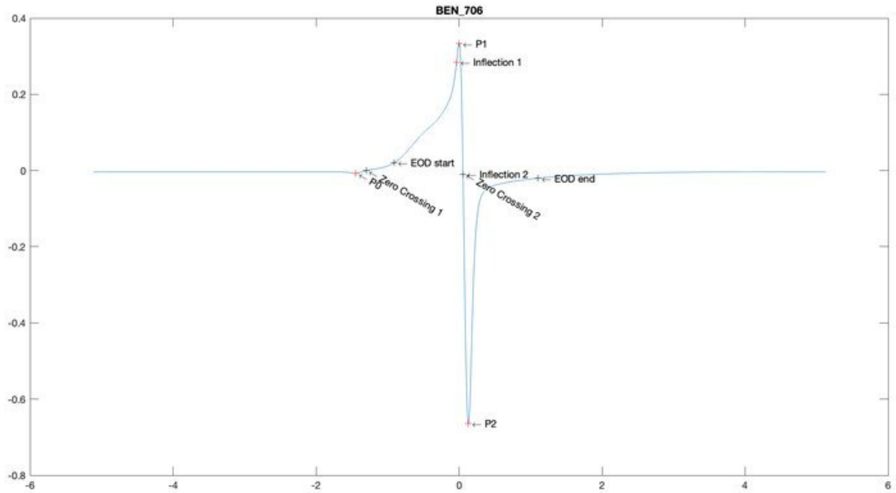
BEN_706
P1
Inflection 1
EOD start
P0
Zero Crossing 1
Inflection 2
Zero Crossing 2
EOD end
P2

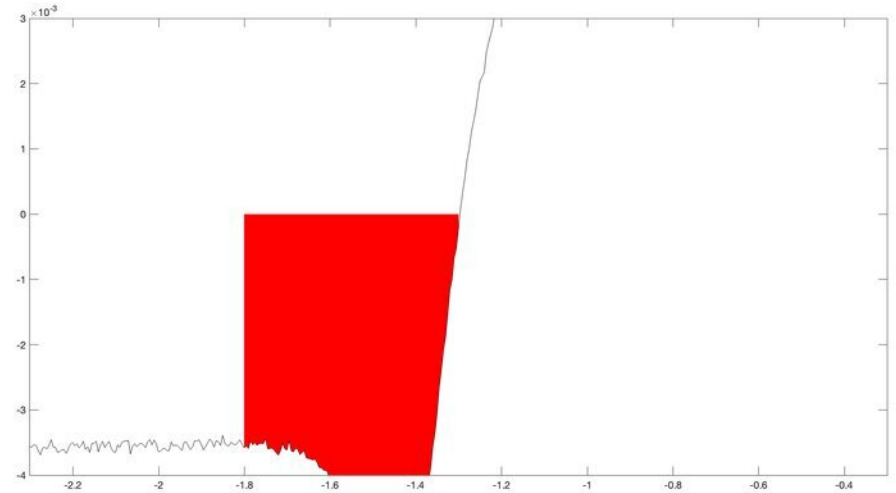

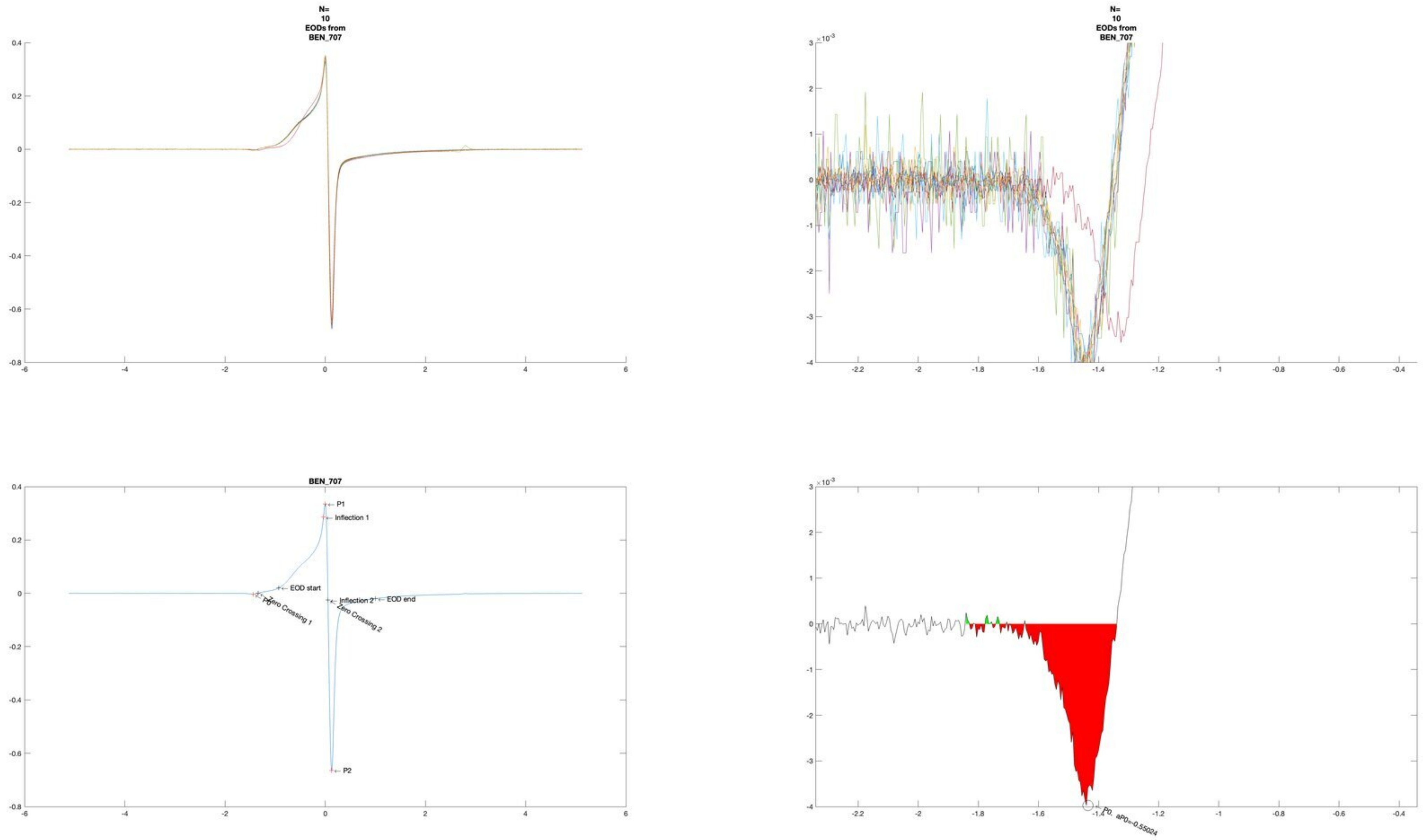
N=
10
EODs from
BEN_707
N=
10
EODs from
BEN_707
BEN_707
P1
Inflection 1
EOD start
P0
Zero Crossing 1
Inflection 2
Zero Crossing 2
EOD end
P2
P0, aP0=-0.55024

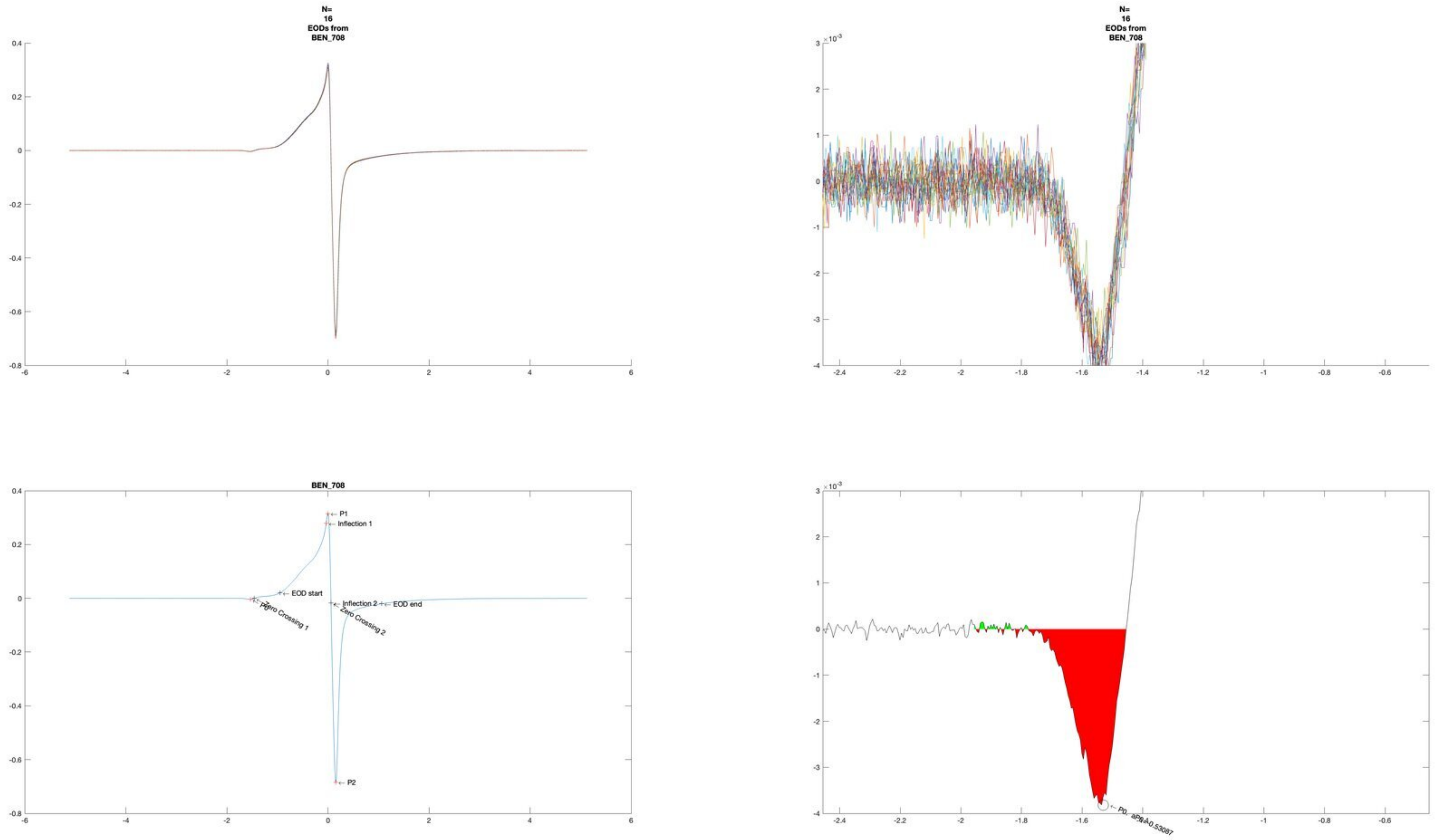
N=
16
EODs from
BEN_708
N=
16
EODs from
BEN_708
BEN_708
P1
Inflection 1
EOD start
Inflection 2
EOD end
Zero Crossing 1
Zero Crossing 2
P2

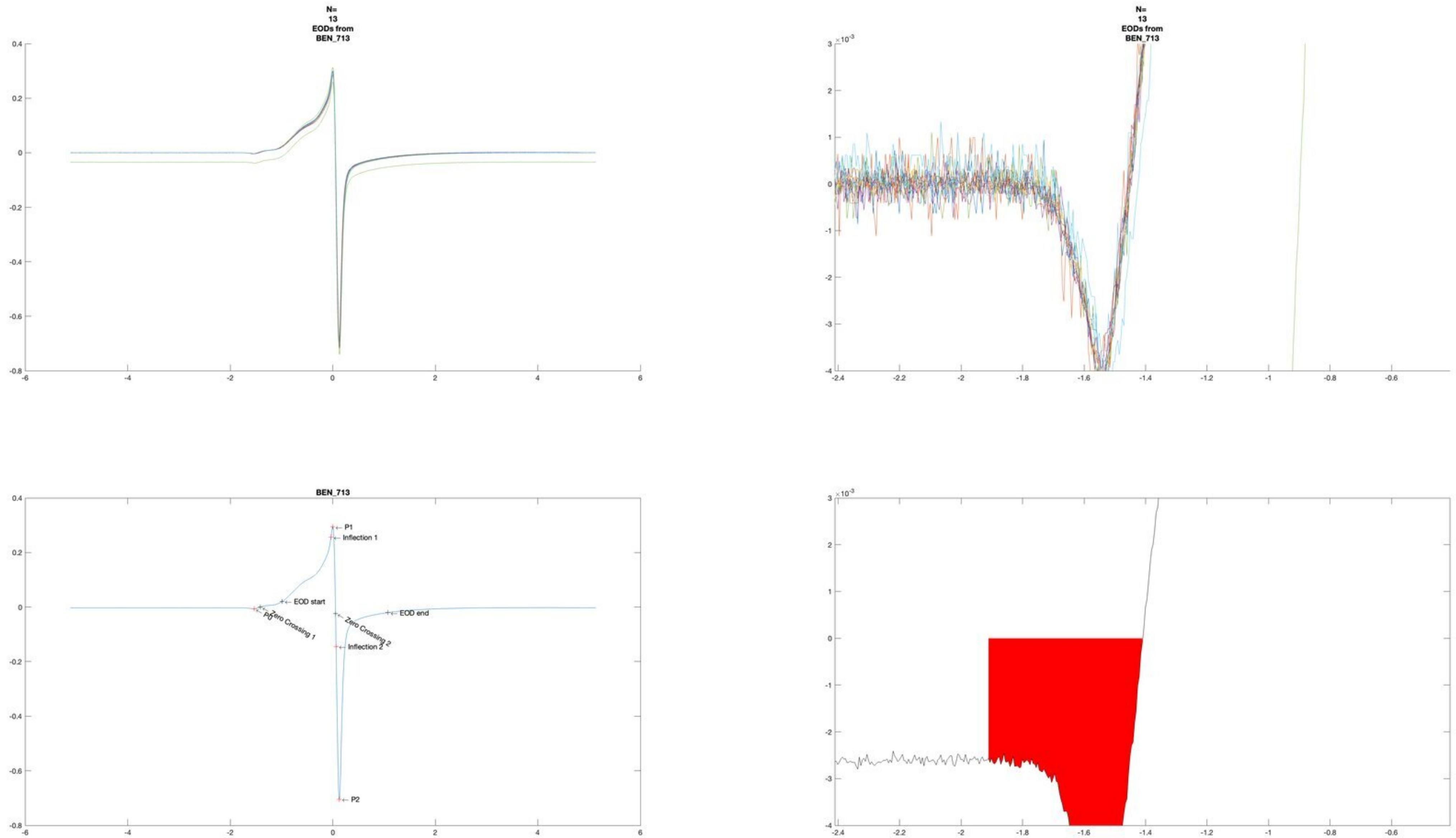
N=
13
EODs from
BEN_713
N=
13
EODs from
BEN_713
BEN_713
P1
Inflection 1
EOD start
P0
Zero Crossing 1
Zero Crossing 2
EOD end
Inflection 2
P2

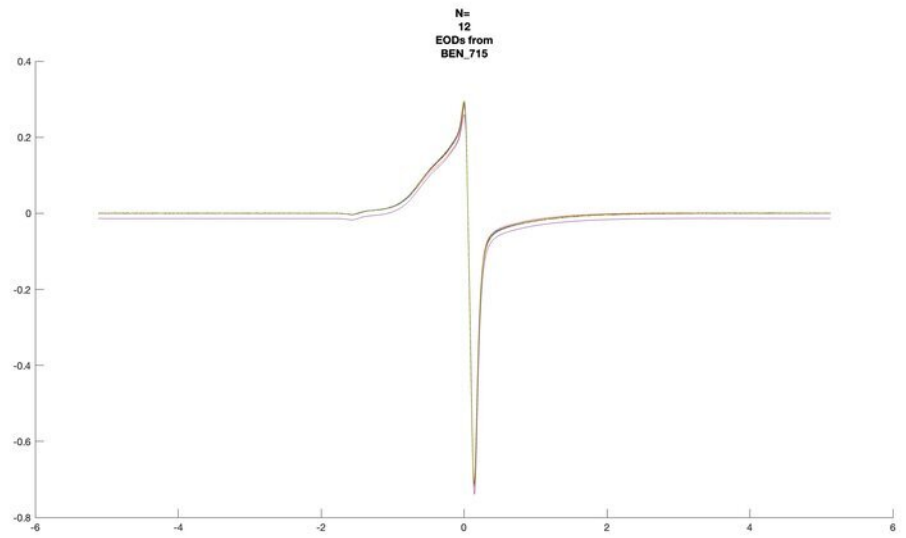
N=
12
EODs from
BEN_715

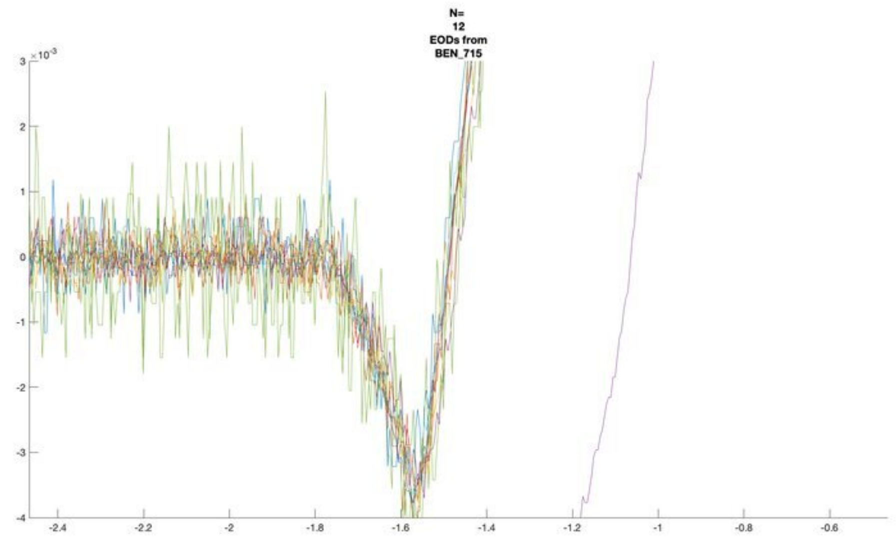
N=
12
EODs from
BEN_715

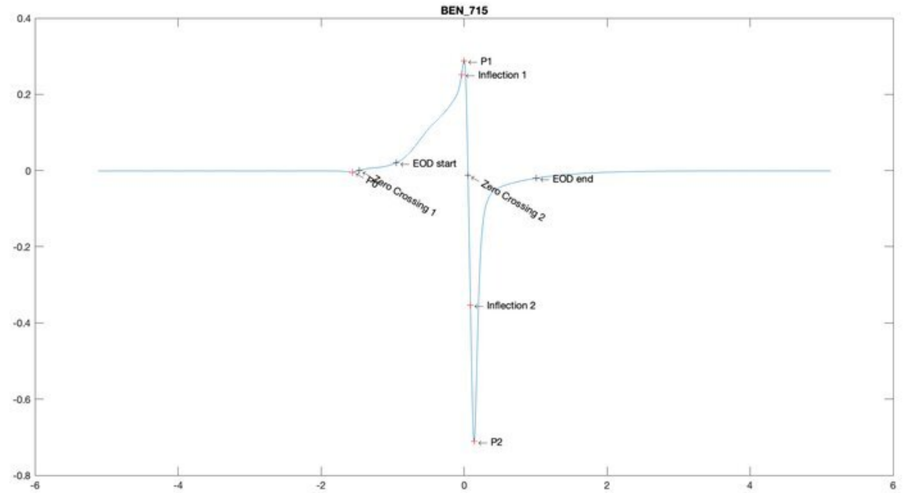
BEN_715
P1
Inflection 1
EOD start
P0
Zero Crossing 1
Zero Crossing 2
EOD end
Inflection 2
P2

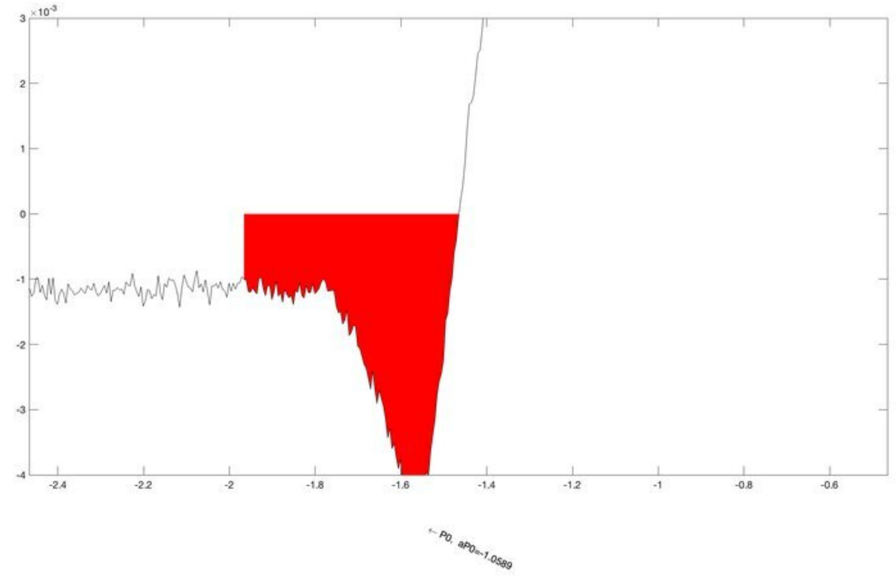
← P0, aP0=-1.0589

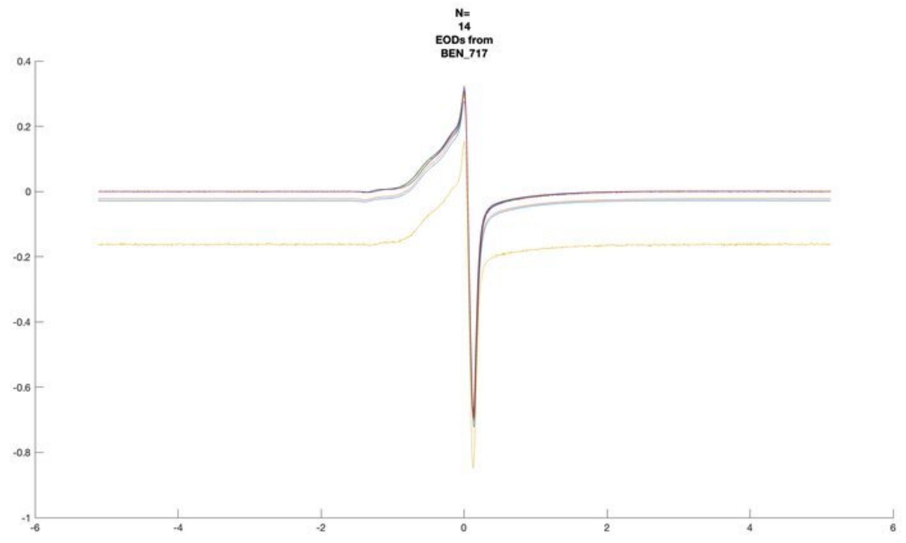
N=
14
EODs from
BEN_717

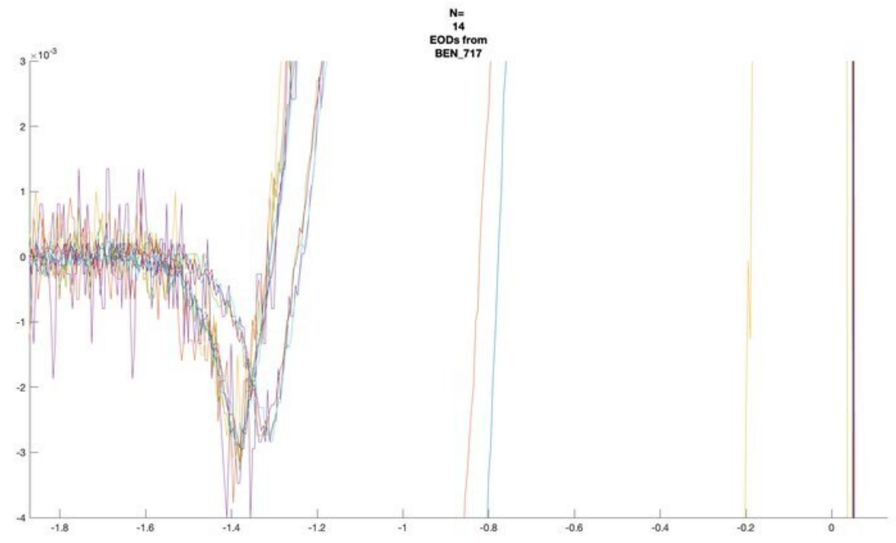
N=
14
EODs from
BEN_717

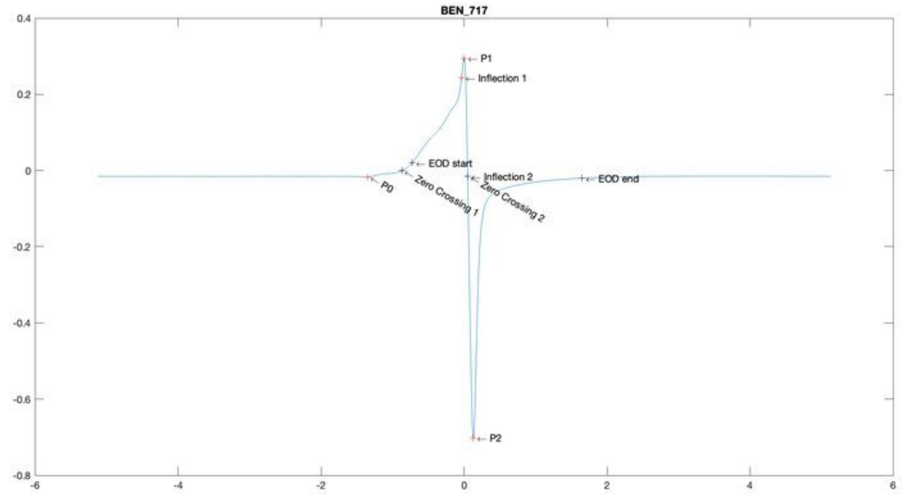
BEN_717
P1
Inflection 1
EOD start
Inflection 2
EOD end
P0
Zero Crossing 1
Zero Crossing 2
P2

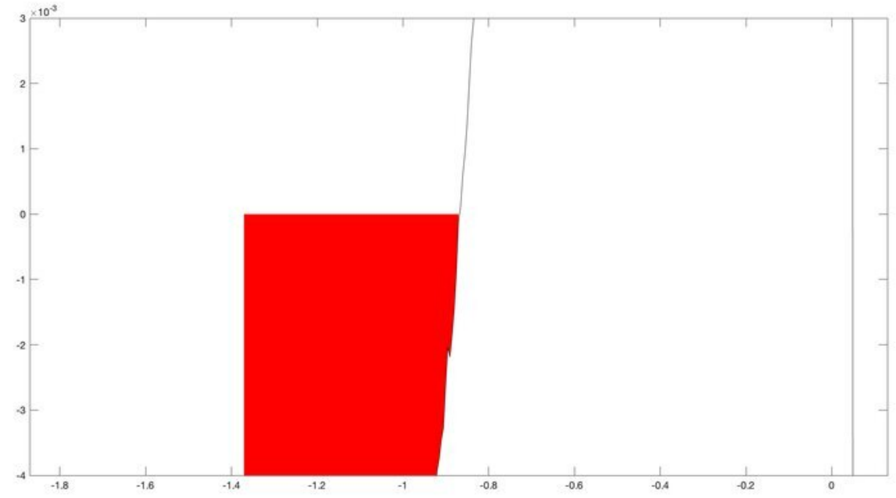

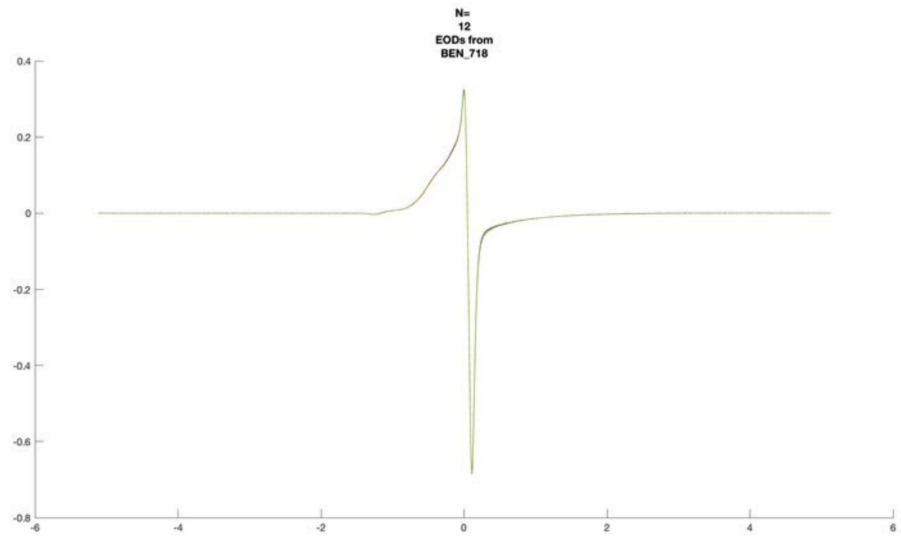
N=
12
EODs from
BEN_718

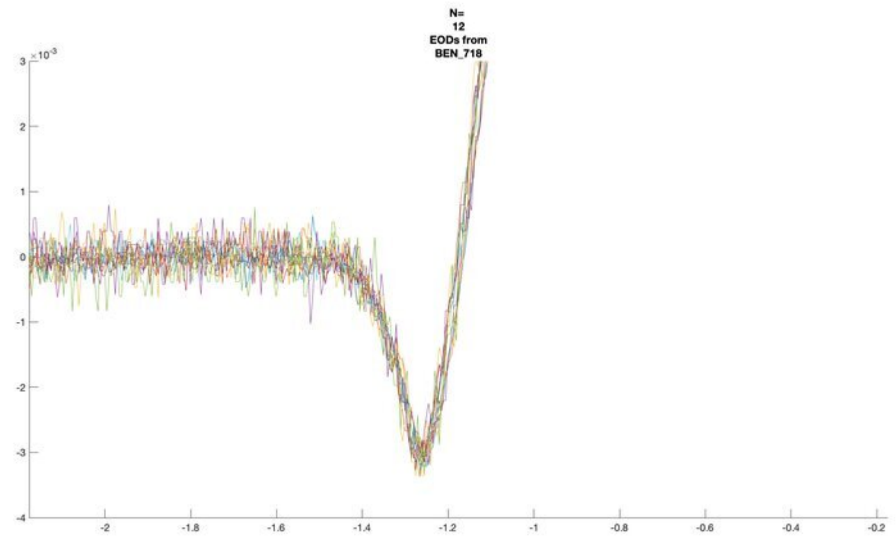
N=
12
EODs from
BEN_718
×10^-3

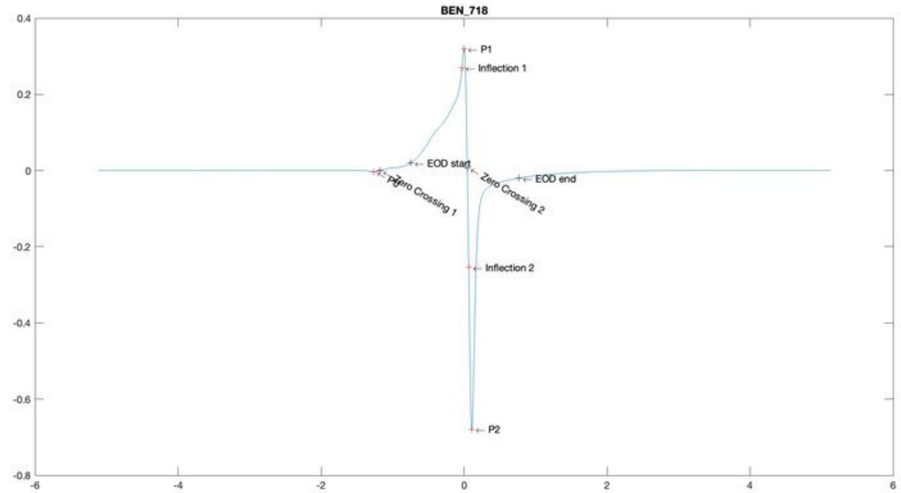
BEN_718
P1
Inflection 1
EOD start
P0
Zero Crossing 1
Zero Crossing 2
EOD end
Inflection 2
P2

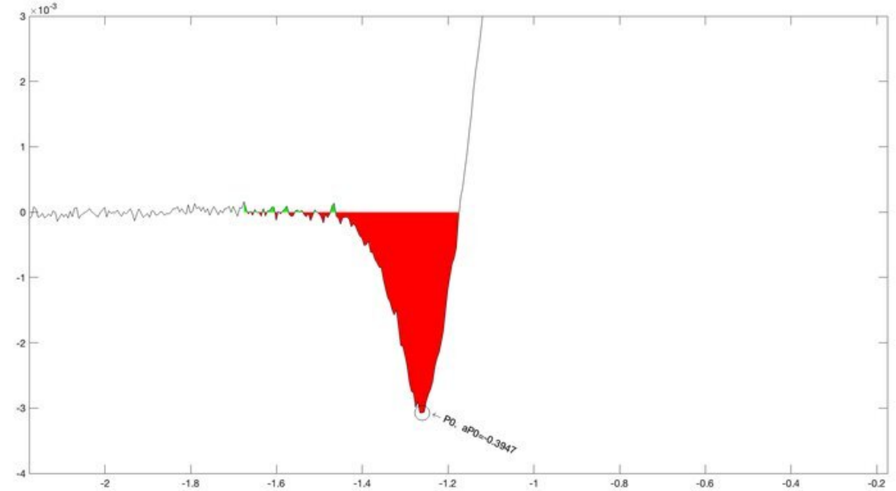
×10^-3
P0. aP0=-0.3947

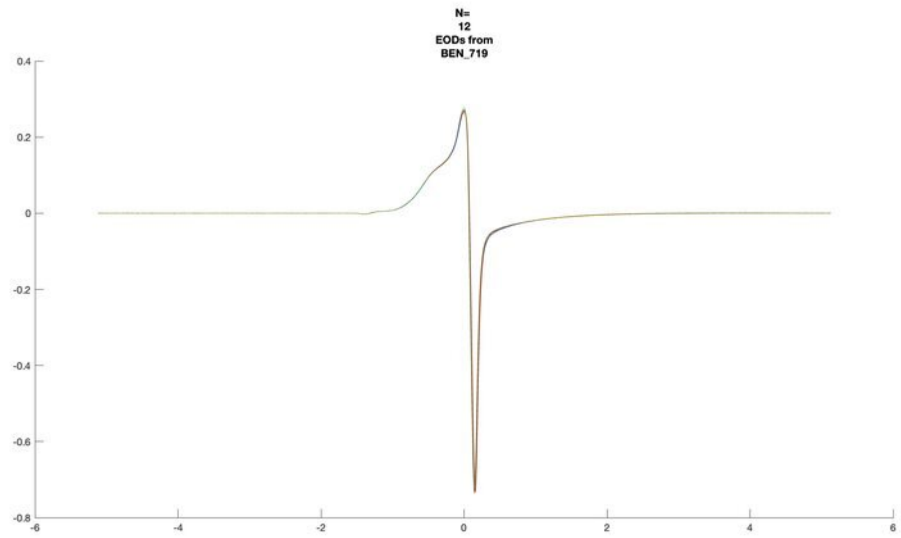
N=
12
EODs from
BEN_719

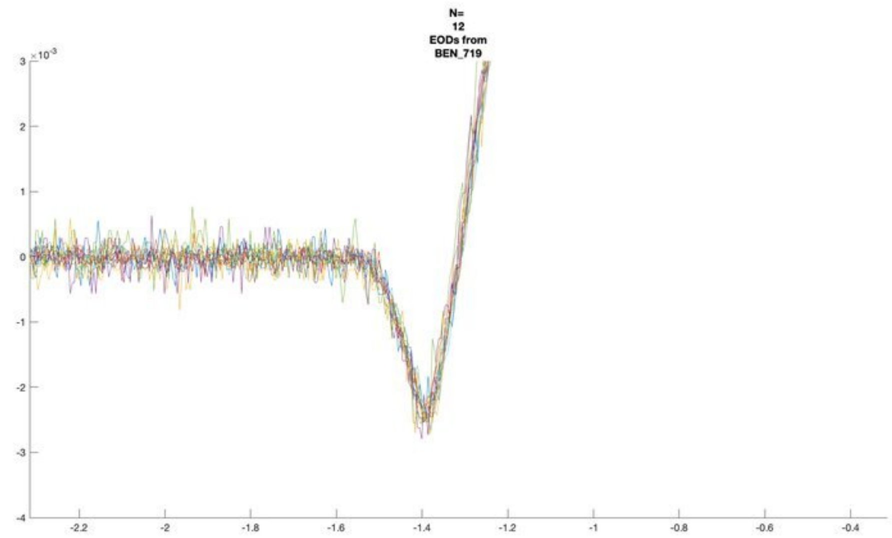
N=
12
EODs from
BEN_719
×10^-3

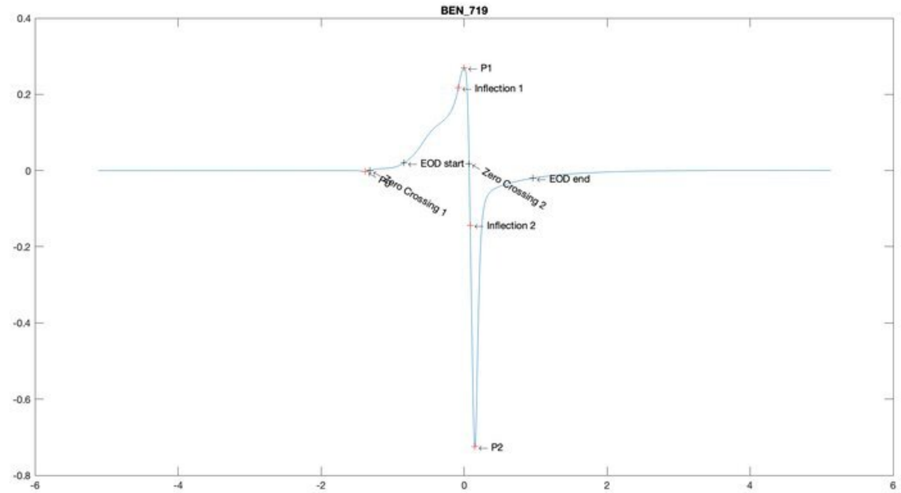
BEN_719
P1
Inflection 1
EOD start
Zero Crossing 1
Zero Crossing 2
EOD end
Inflection 2
P2

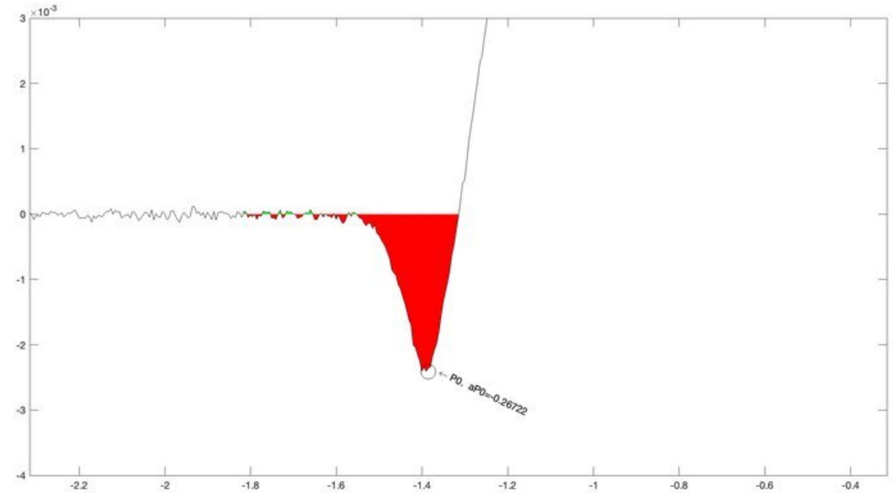
×10^-3
P0, aP0=-0.26722

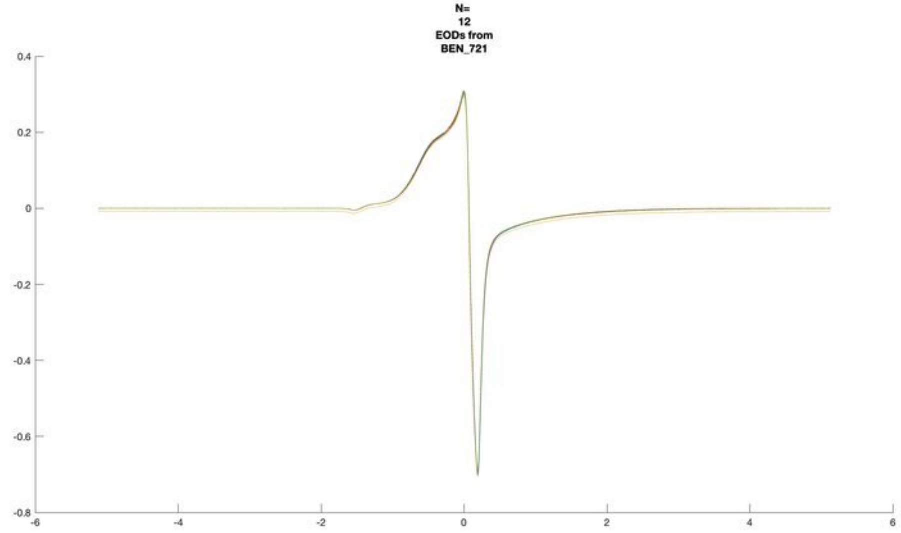
N=
12
EODs from
BEN_721

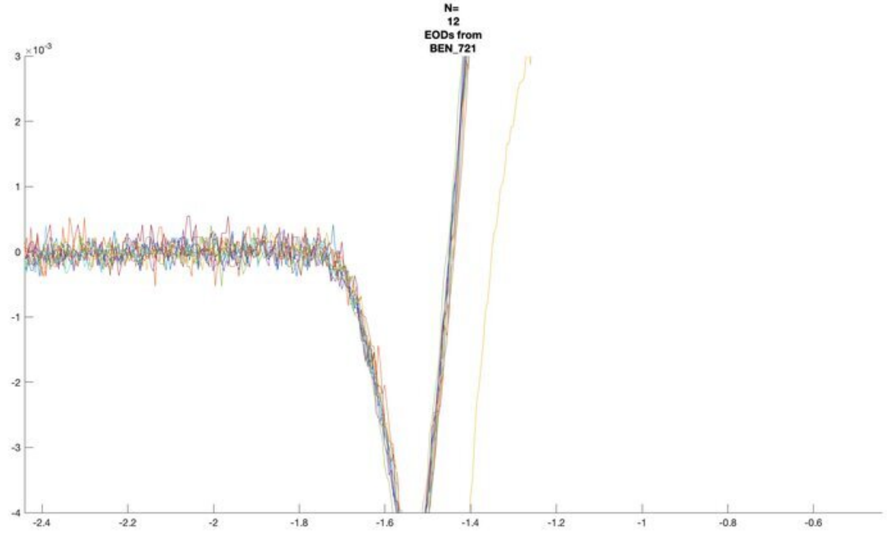
N=
12
EODs from
BEN_721
$\times 10^{-3}$

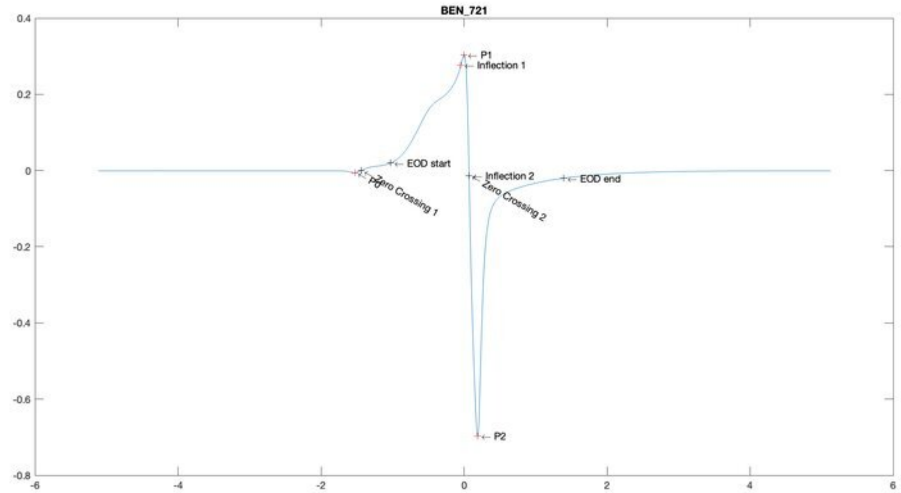
BEN_721
P1
Inflection 1
EOD start
P0
Zero Crossing 1
Inflection 2
Zero Crossing 2
EOD end
P2

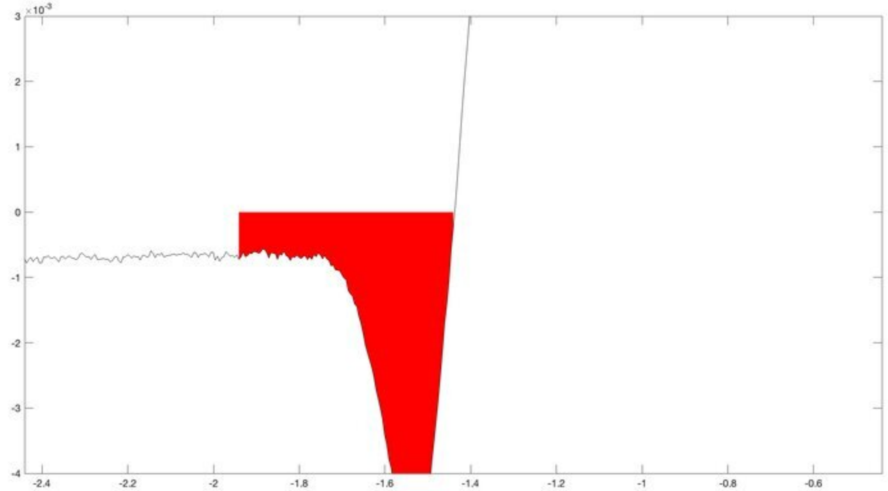
$\times 10^{-3}$

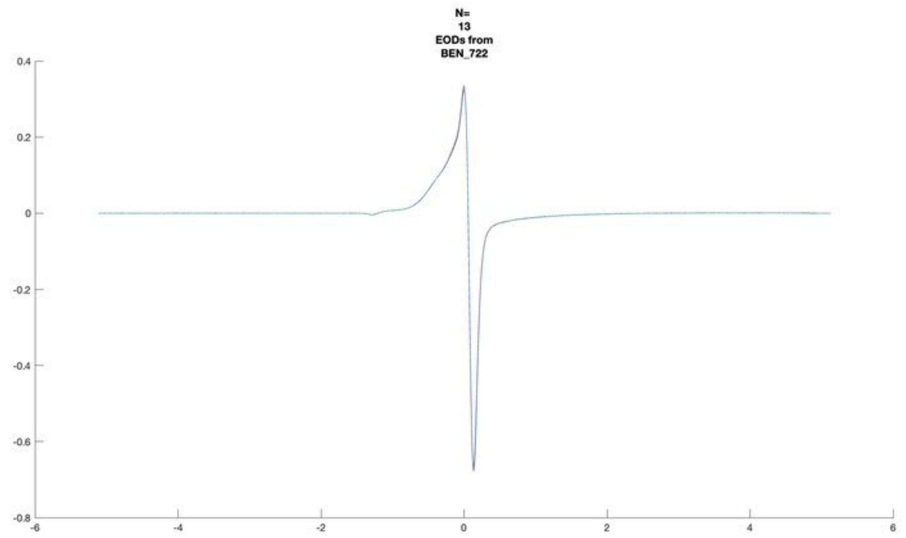
N=
13
EODs from
BEN_722

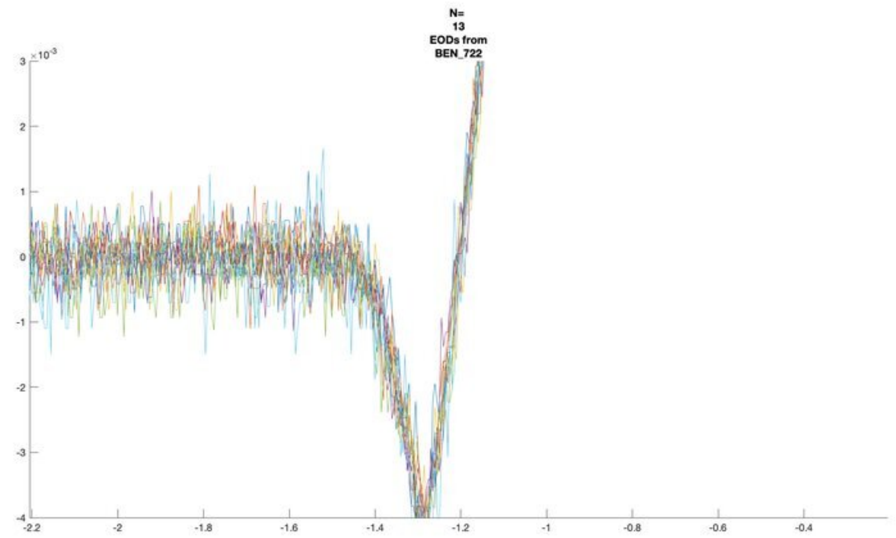
N=
13
EODs from
BEN_722
×10^-3

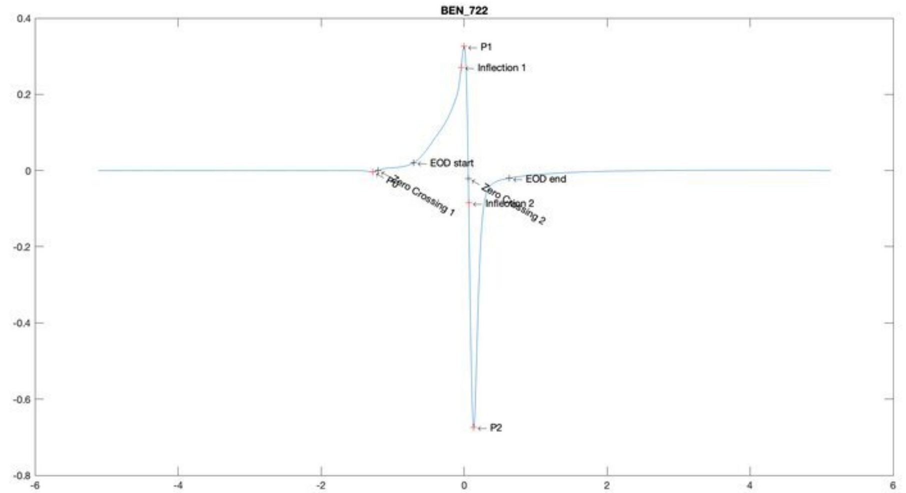
BEN_722
P1
Inflection 1
EOD start
EOD end
Zero Crossing 1
Zero Crossing 2
P2

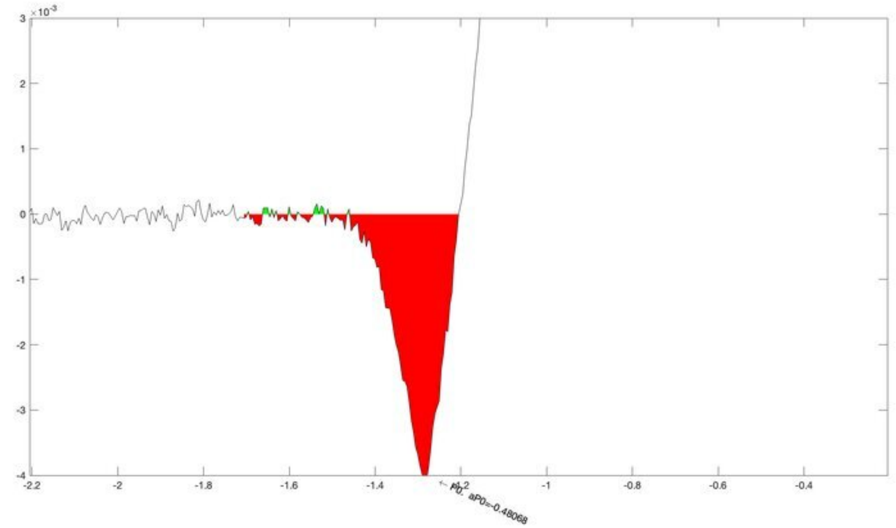
×10^-3
P0, aP0=-0.48068

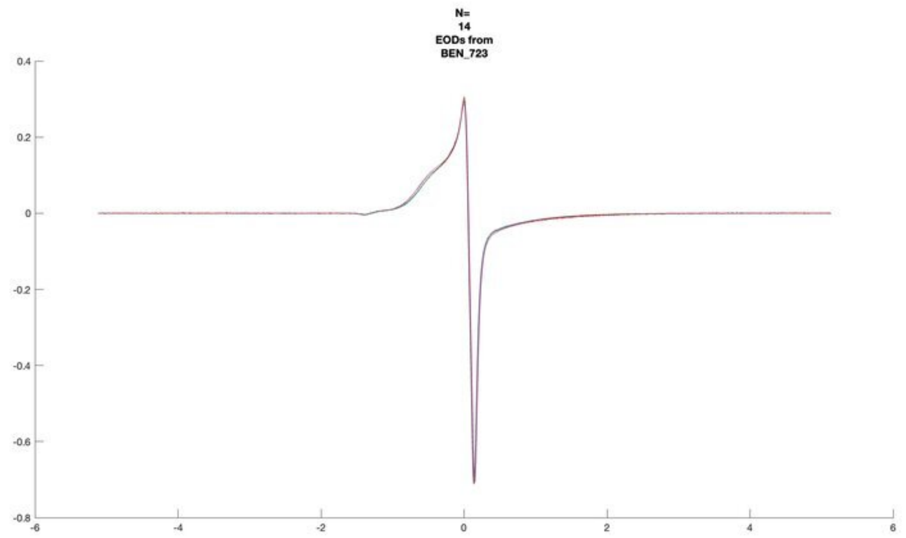
N=
14
EODs from
BEN_723

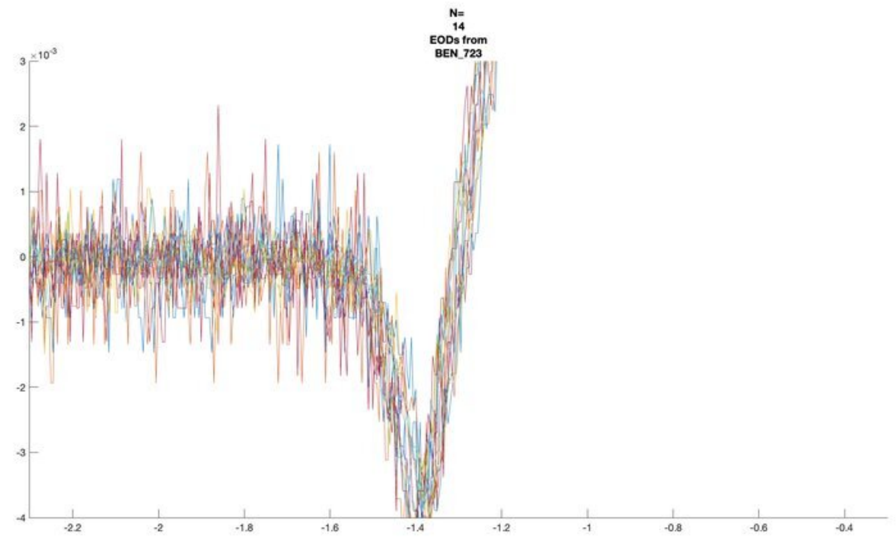
N=
14
EODs from
BEN_723
×10^-3

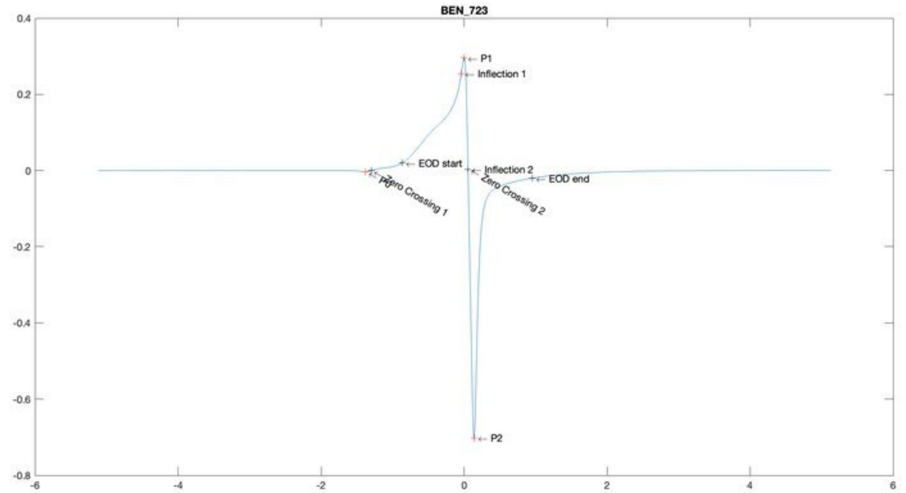
BEN_723
P1
Inflection 1
EOD start
Inflection 2
EOD end
P0
Zero Crossing 1
Zero Crossing 2
P2

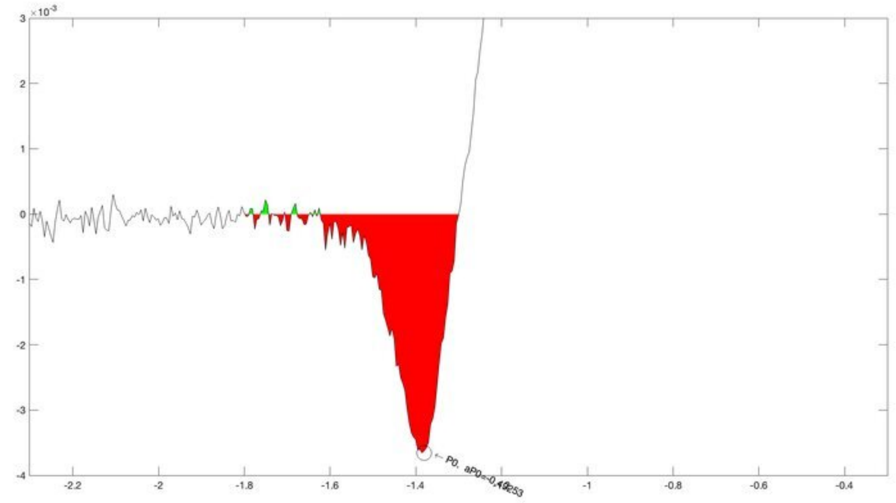
×10^-3
P0,

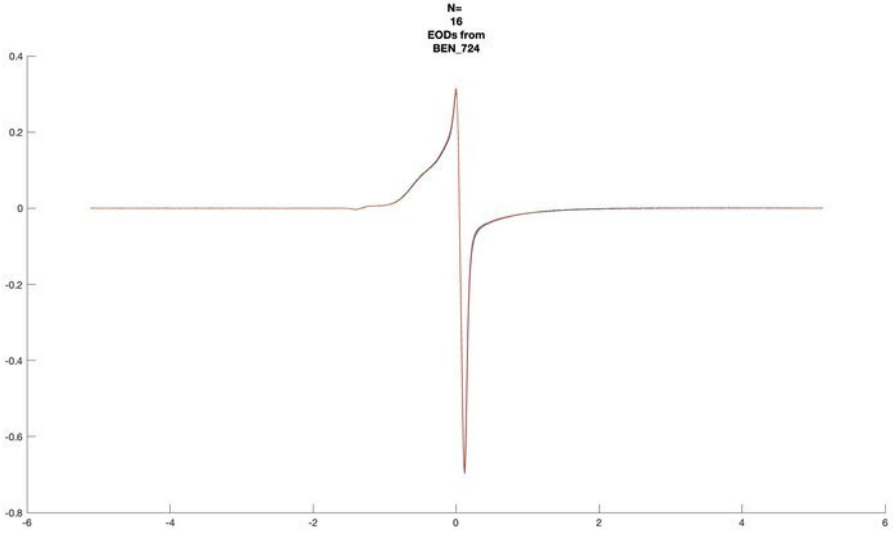
N=
16
EODs from
BEN_724

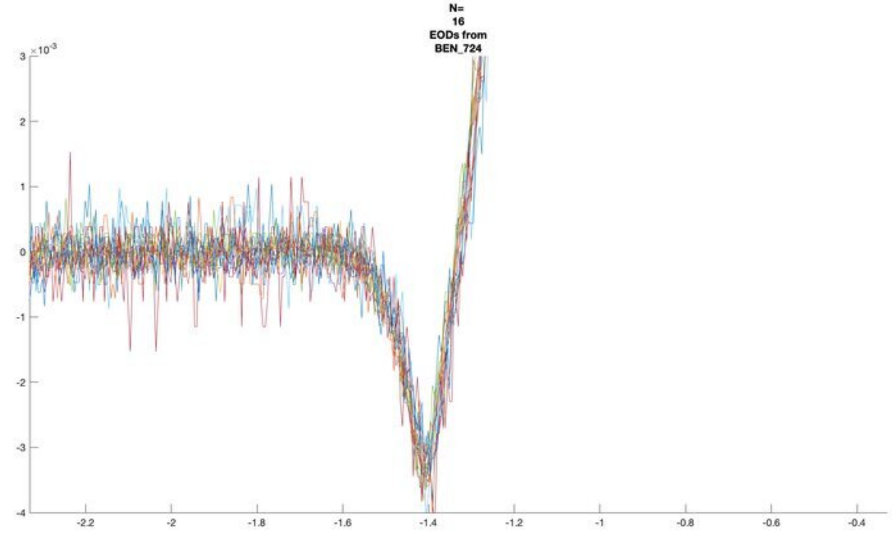
N=
16
EODs from
BEN_724
×10^-3

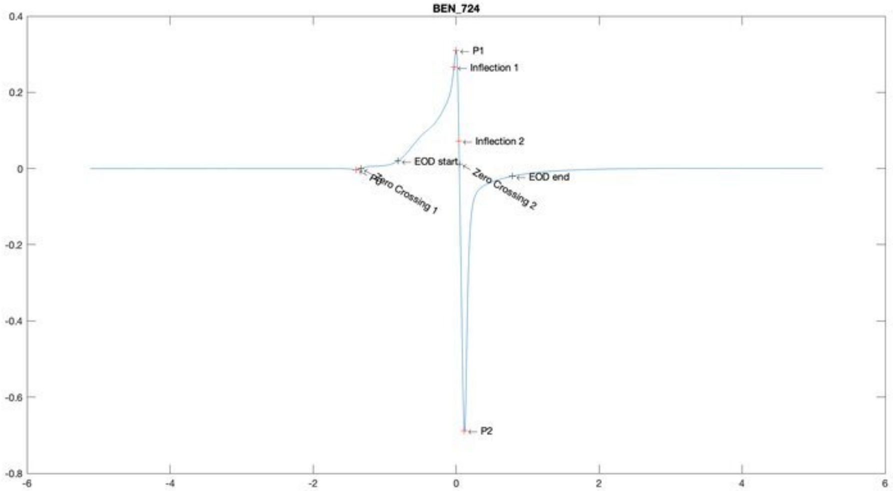
BEN_724
P1
Inflection 1
Inflection 2
EOD start
EOD end
P0
Zero Crossing 1
Zero Crossing 2
P2

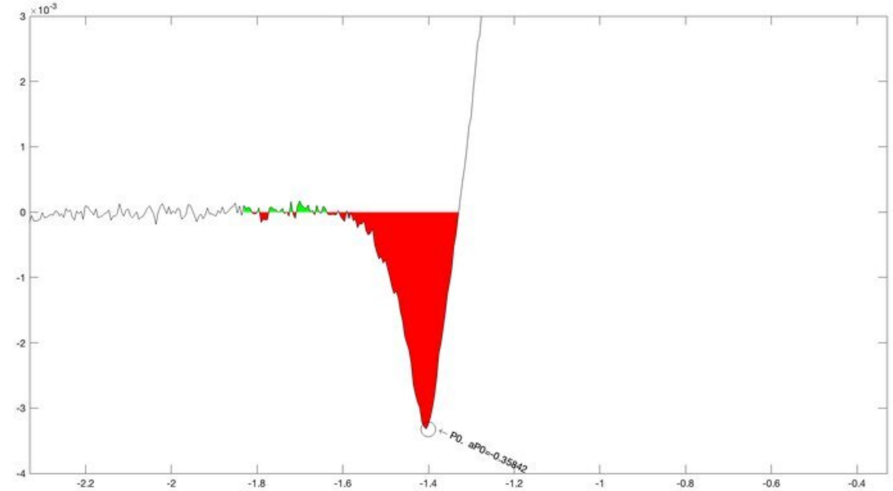
×10^-3
P0, aP0=-0.35842

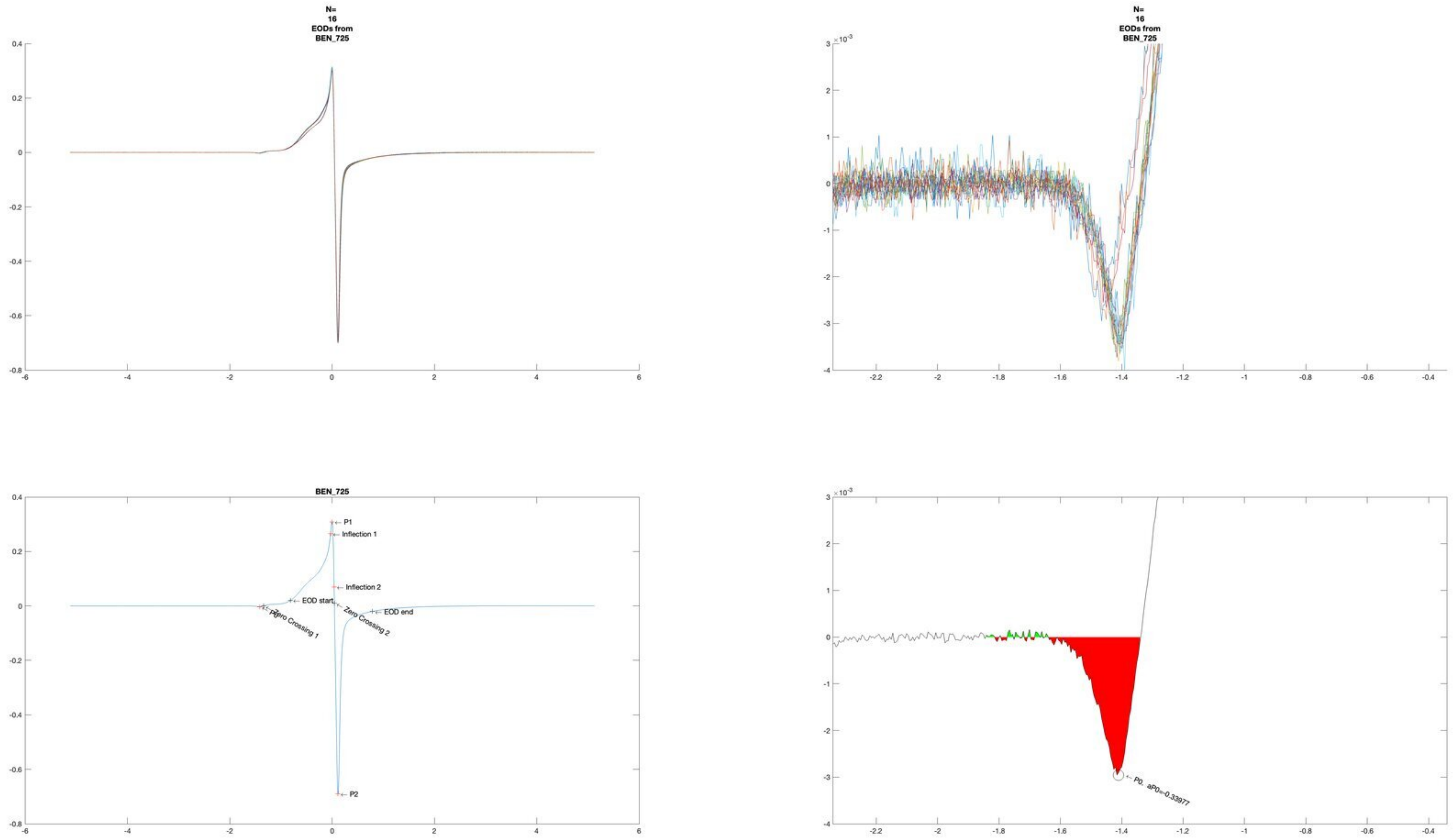

N=
16
EODs from
BEN_725
N=
16
EODs from
BEN_725
BEN_725
P1
Inflection 1
Inflection 2
EOD start
EOD end
P0
Zero Crossing 1
Zero Crossing 2
P2
P0, aP0=-0.33977

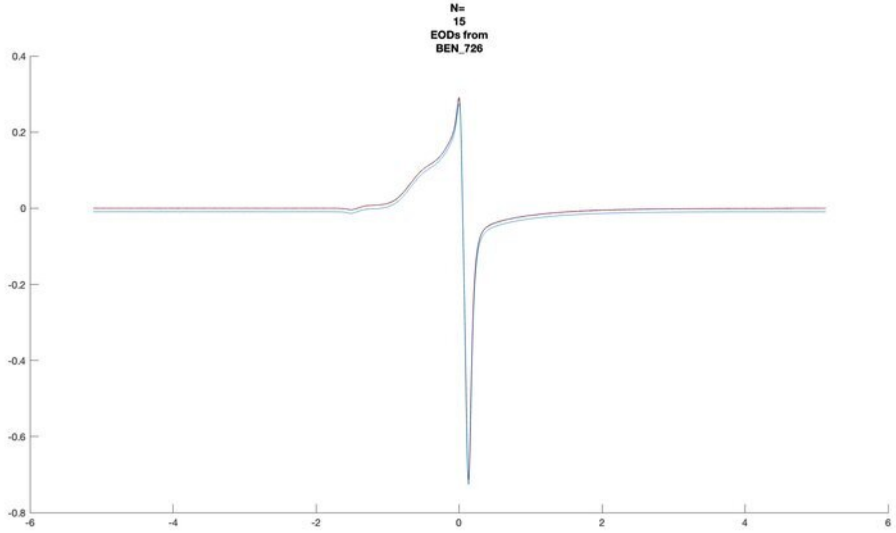
N=
15
EODs from
BEN_726

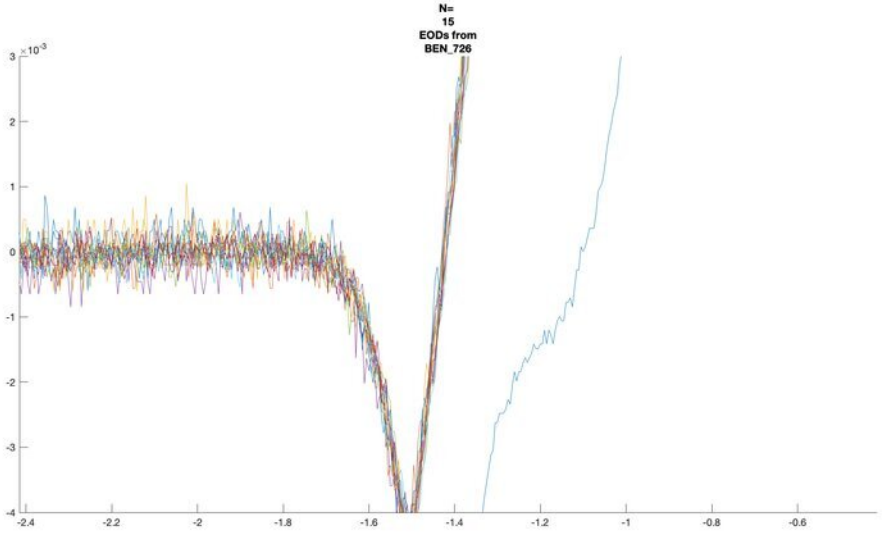
N=
15
EODs from
BEN_726

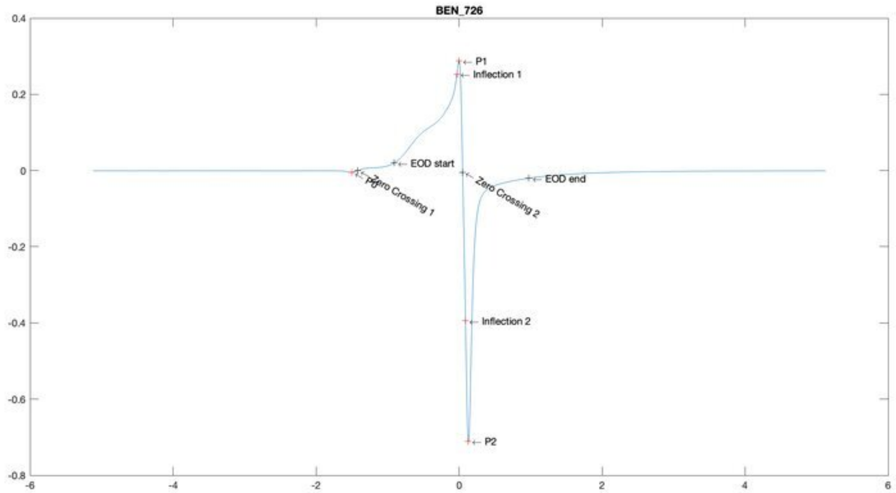
BEN_726
P1
Inflection 1
EOD start
P0
Zero Crossing 1
Zero Crossing 2
EOD end
Inflection 2
P2

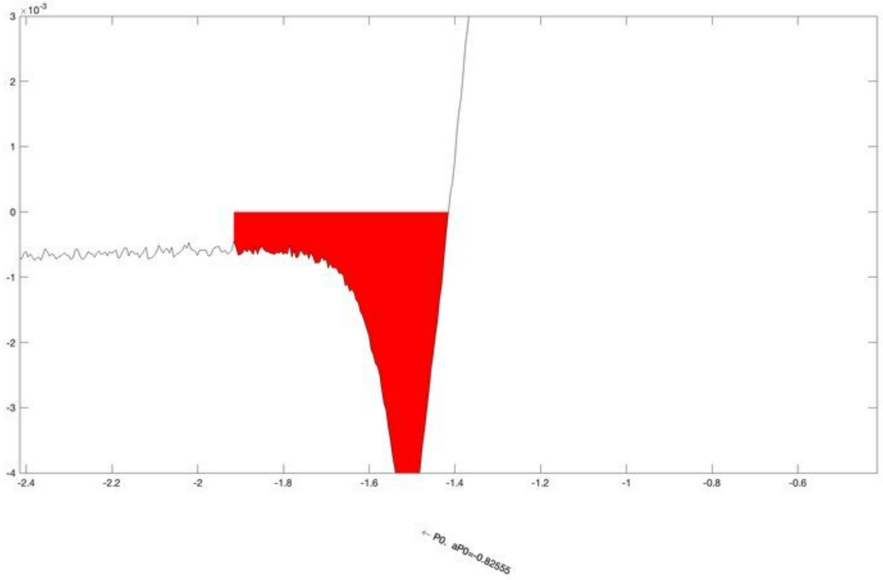
P0, aP0=-0.82555

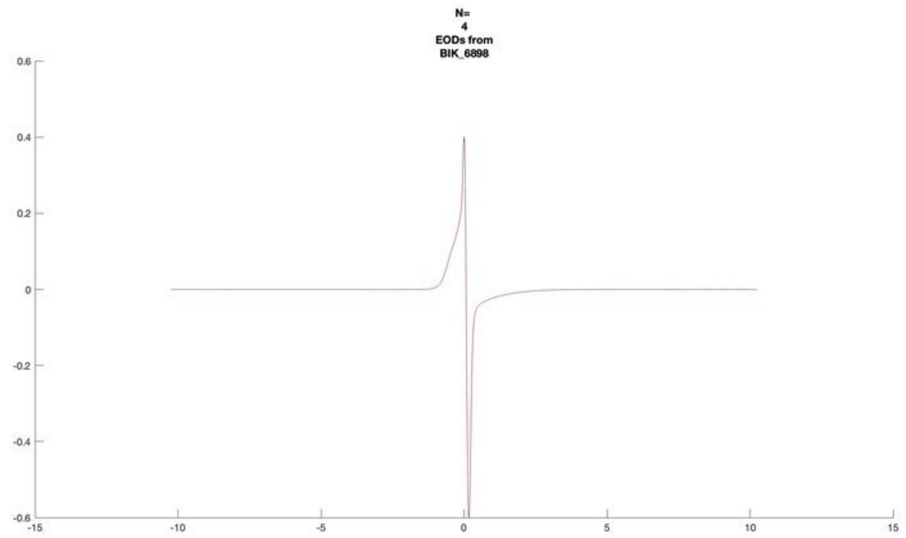
N=
4
EODs from
BIK_6898

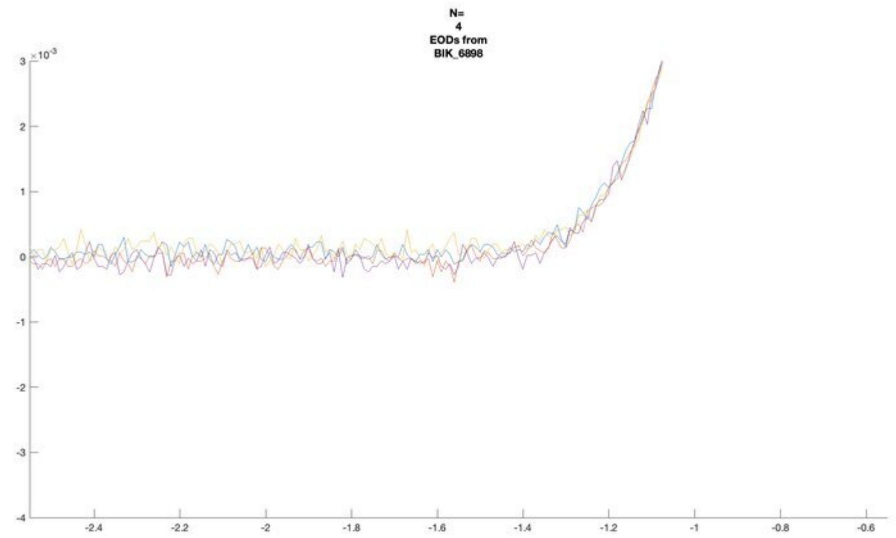
N=
4
EODs from
BIK_6898

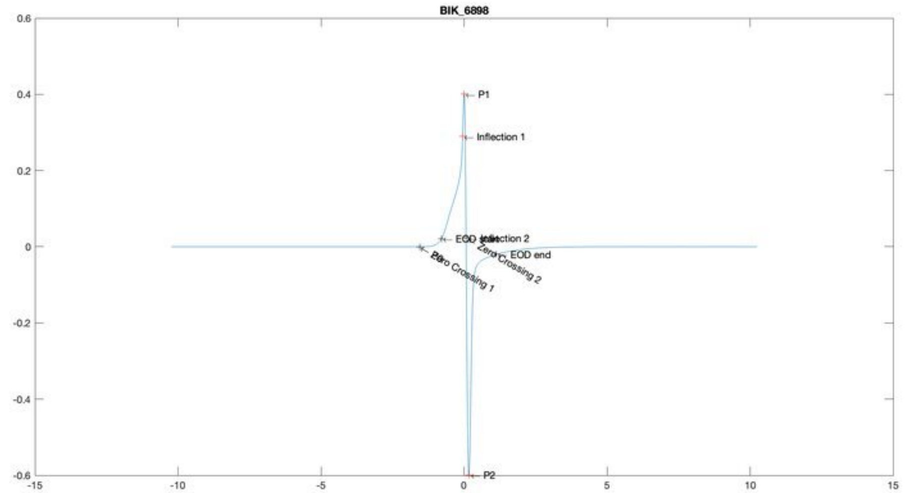
BIK_6898
P1
Inflection 1
Inflection 2
EOD end
Zero Crossing 1
Zero Crossing 2
P2

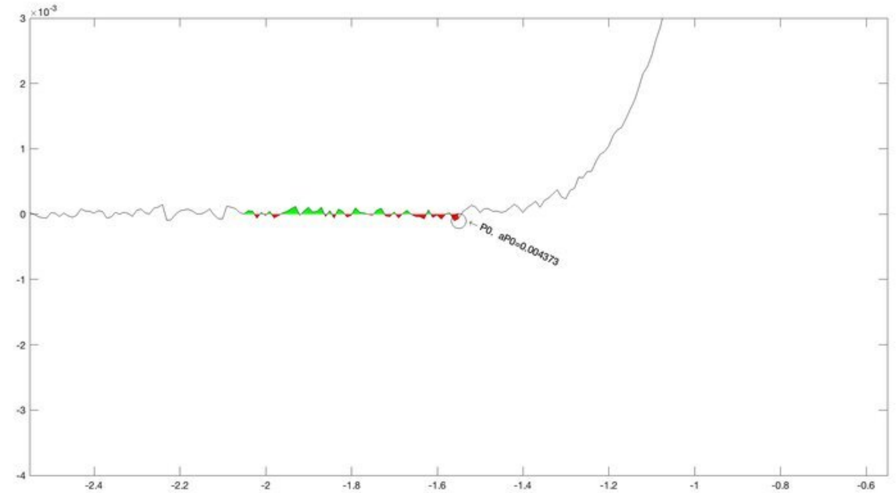
P0, aP0=0.004373

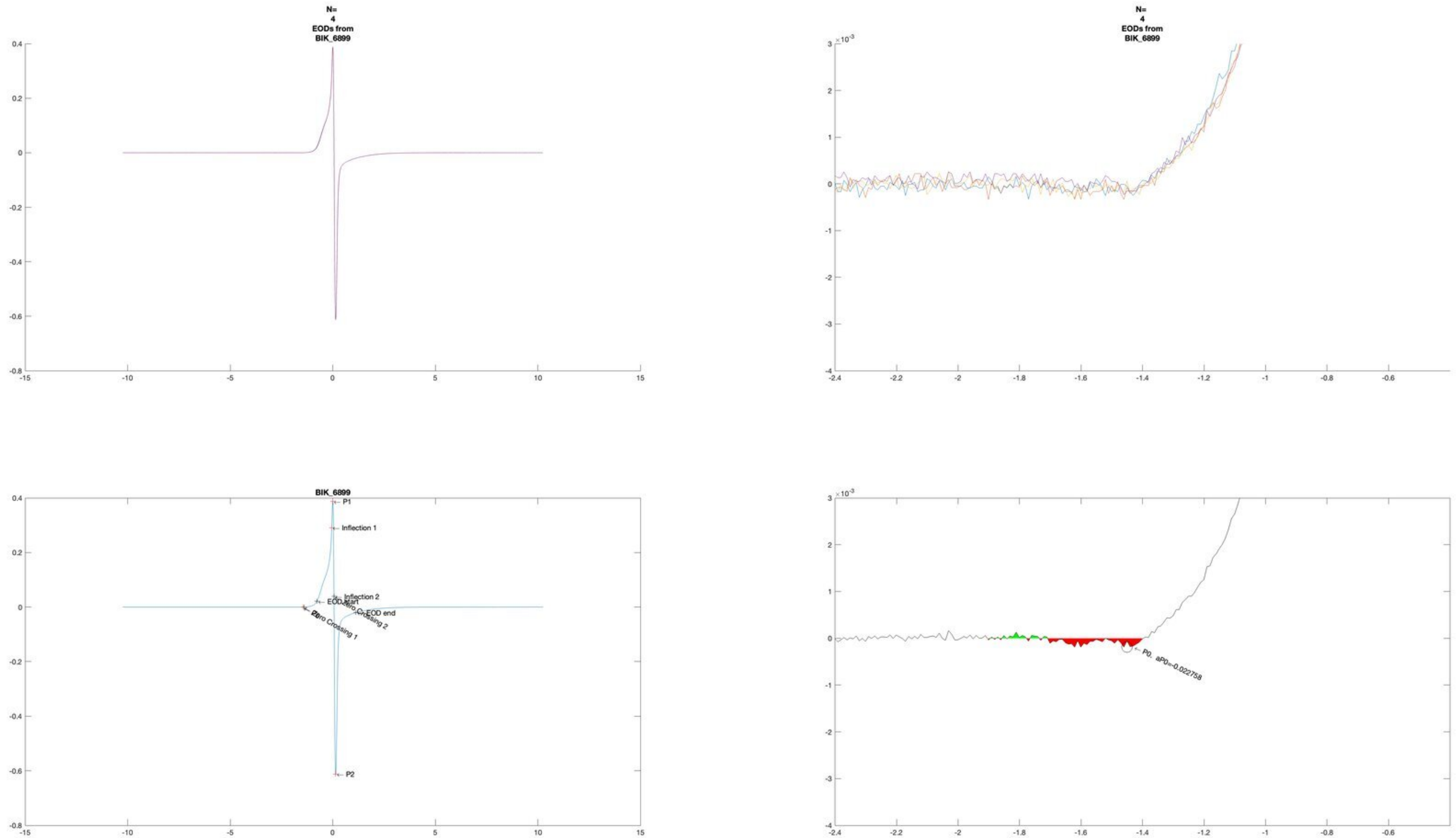
N=
4
EODs from
BIK_6899
N=
4
EODs from
BIK_6899
BIK_6899
P1
Inflection 1
Inflection 2
EOD start
EOD end
Zero Crossing 1
Zero Crossing 2
P2
P0, aP0=-0.022758

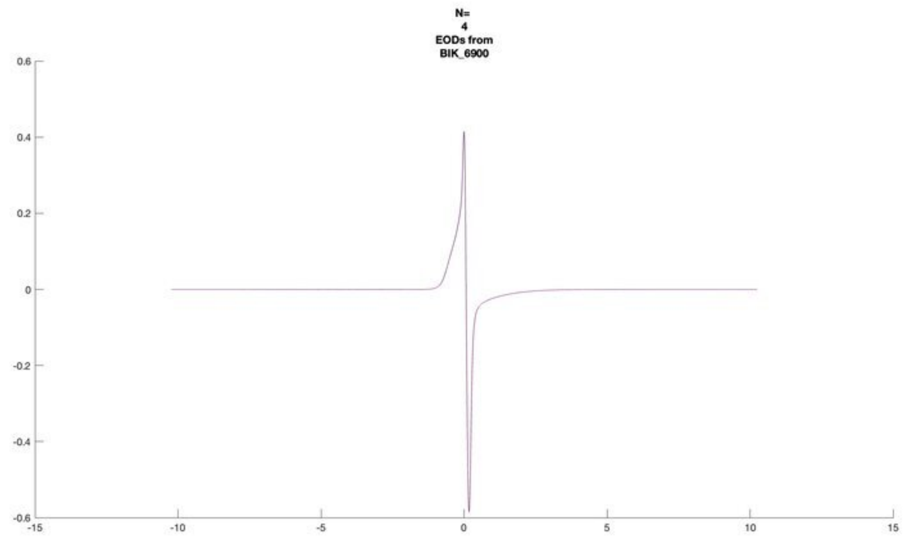
N=
4
EODs from
BIK_6900

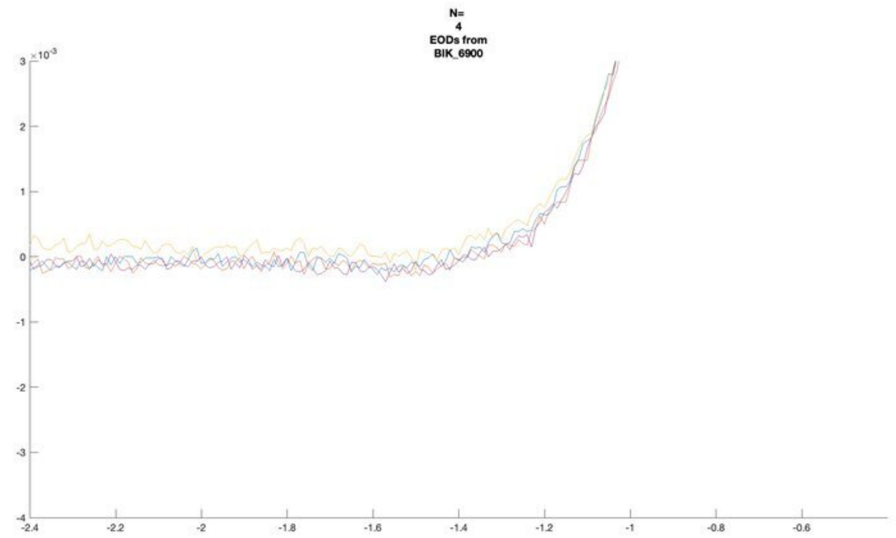
N=
4
EODs from
BIK_6900

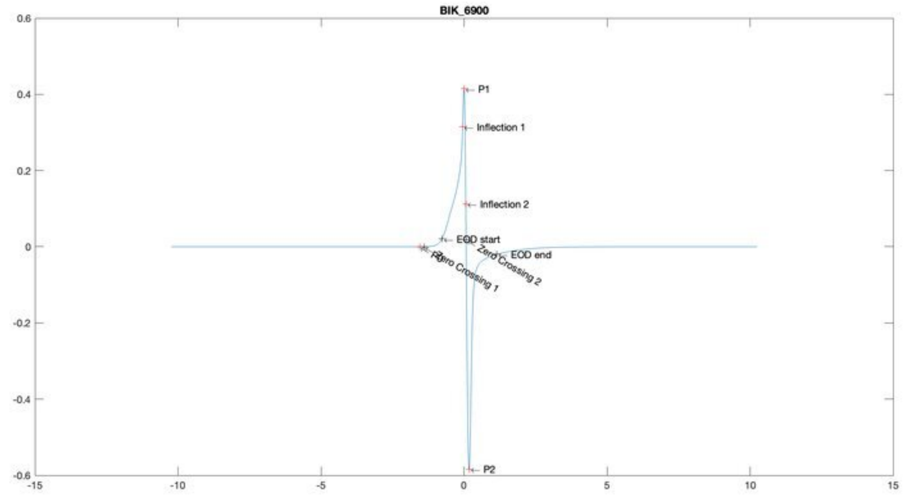
BIK_6900
P1
Inflection 1
Inflection 2
EOD start
EOD end
Zero Crossing 1
Zero Crossing 2
P2

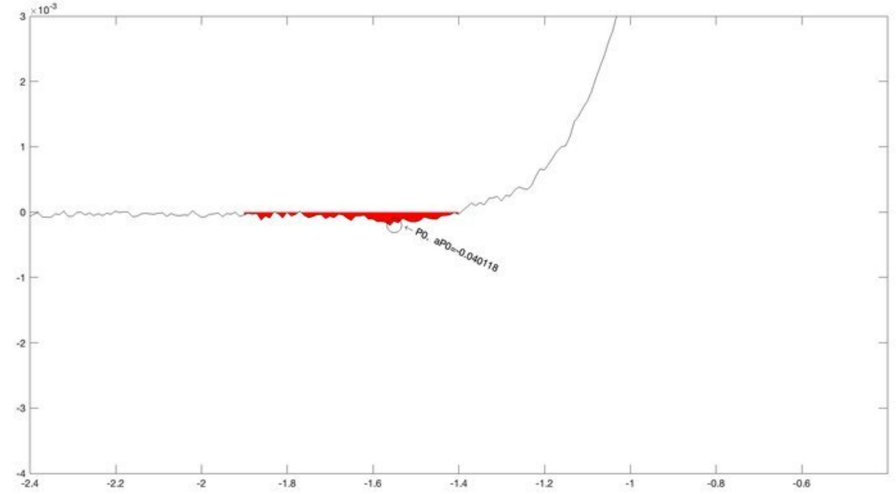
P0, aP0=-0.040118

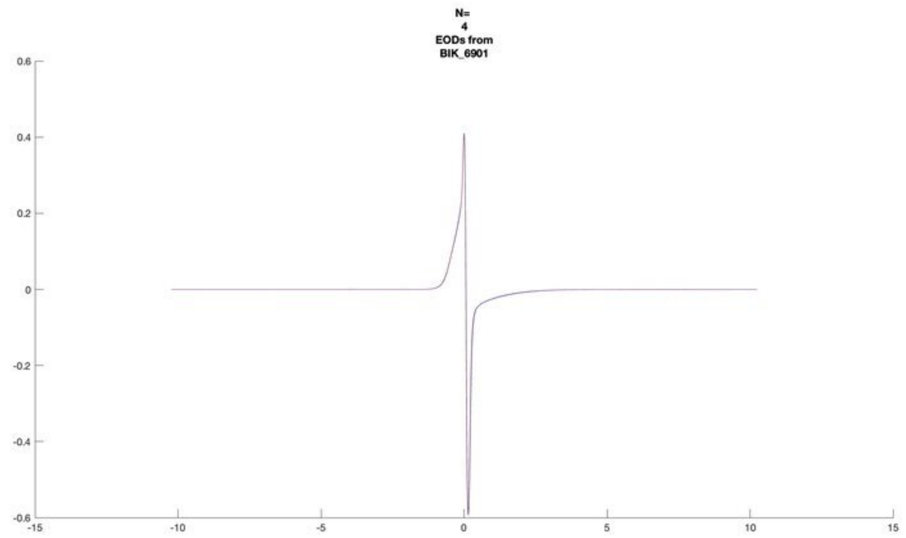
N=
4
EODs from
BIK_6901

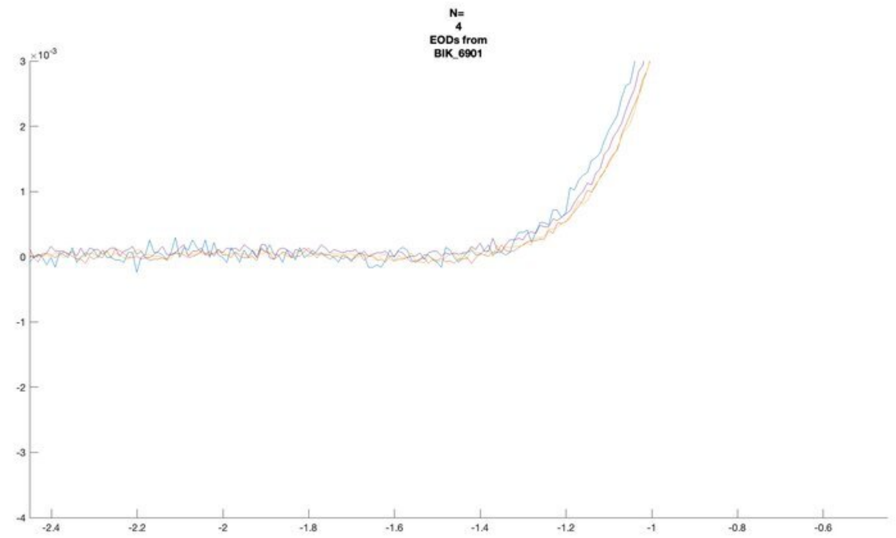
N=
4
EODs from
BIK_6901
×10^-3

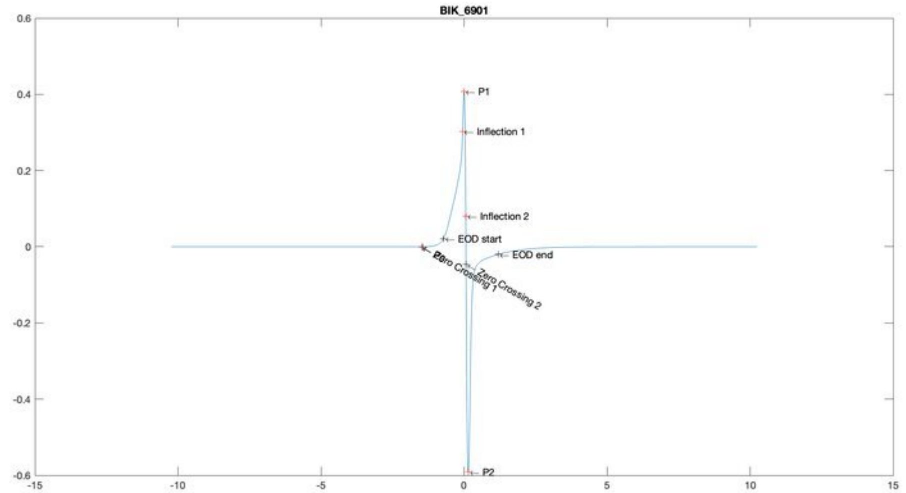
BIK_6901
P1
Inflection 1
Inflection 2
EOD start
EOD end
Zero Crossing 1
Zero Crossing 2
P2

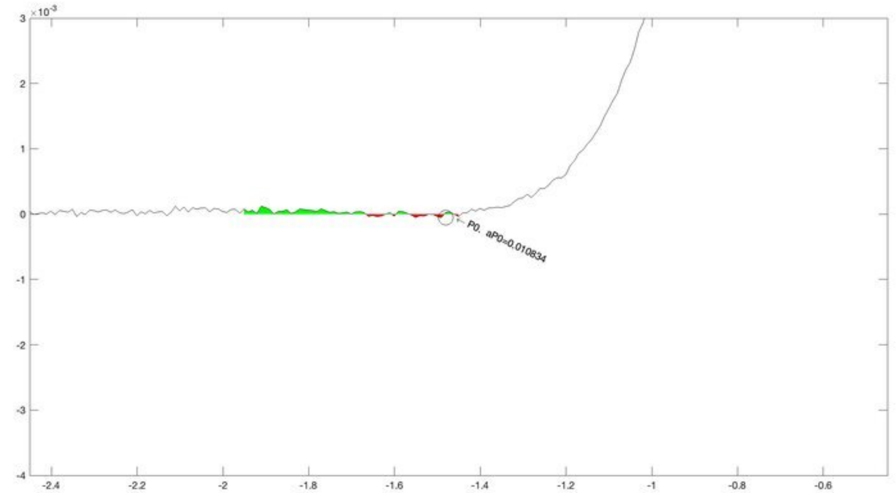
×10^-3
P0, aP0=0.010834

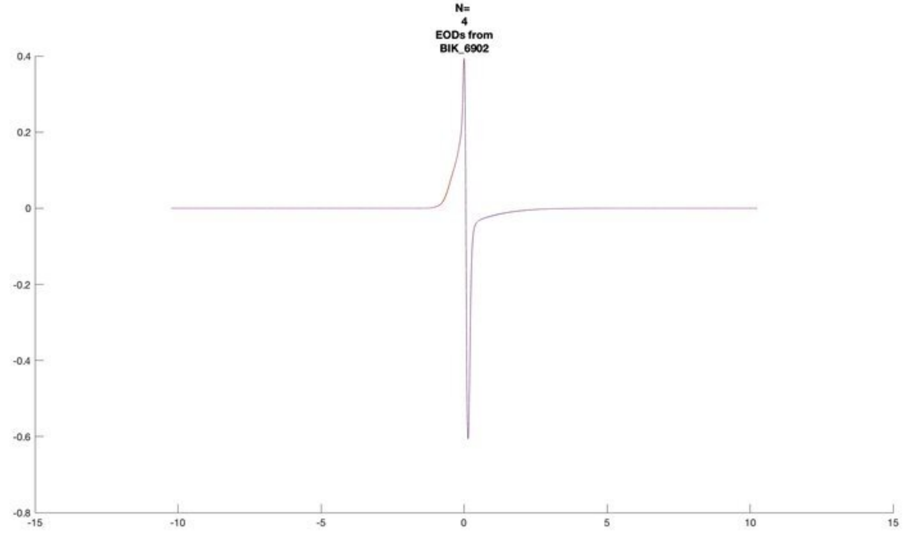
N=
4
EODs from
BIK_6902

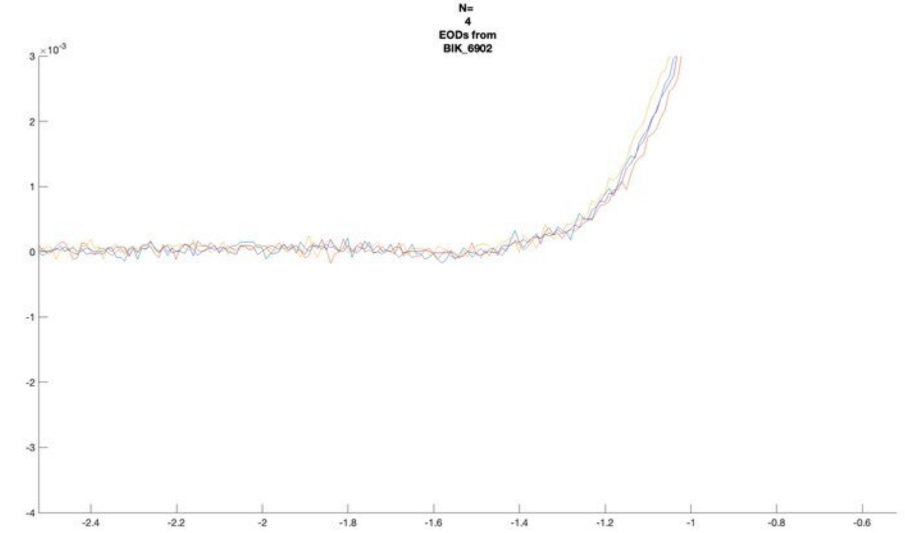
N=
4
EODs from
BIK_6902
$\times 10^{-3}$

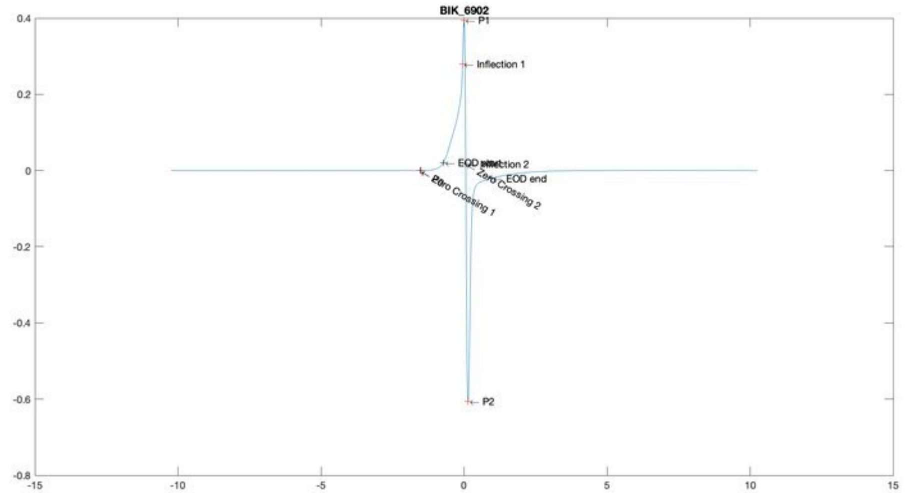
BIK_6902
P1
Inflection 1
EOD start
Inflection 2
EOD end
Zero Crossing 1
Zero Crossing 2
P2

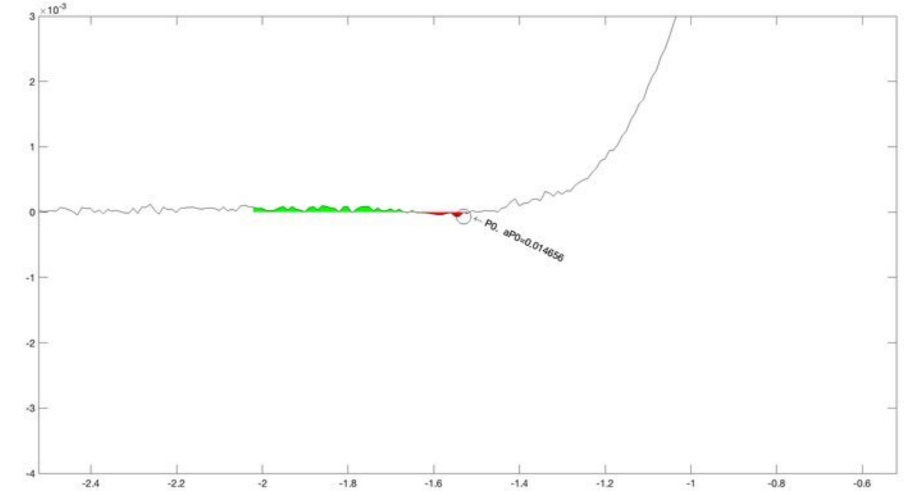
$\times 10^{-3}$
P0, aP0=0.014656

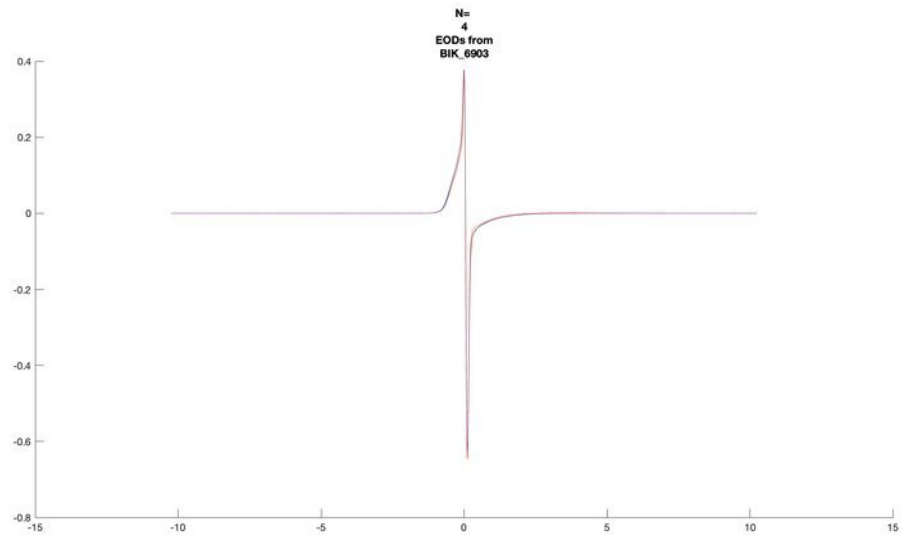
N=
4
EODs from
BIK_6903

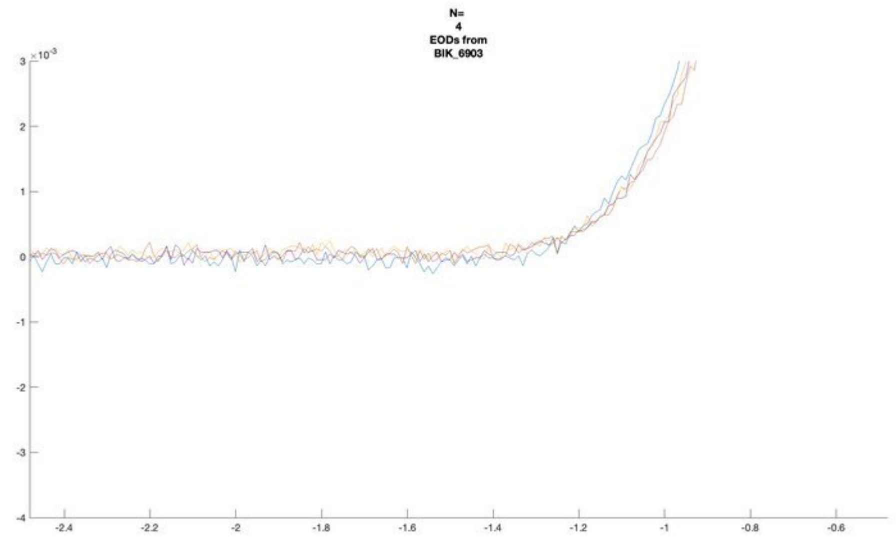
N=
4
EODs from
BIK_6903

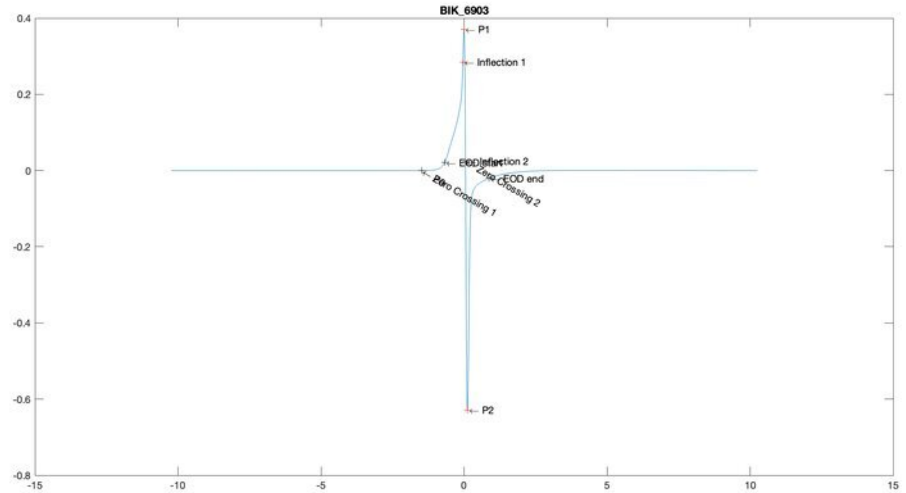
BIK_6903
P1
Inflection 1
Inflection 2
EOD end
Zero Crossing 1
Zero Crossing 2
P2

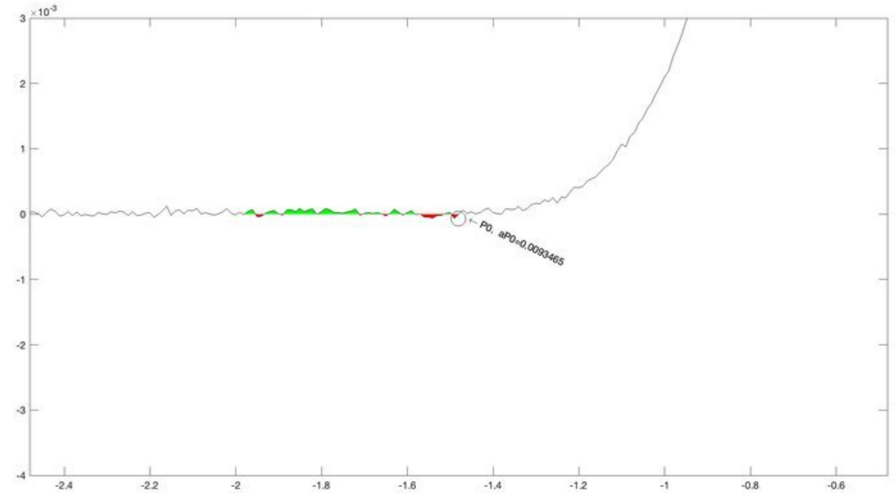
P0, aP0=0.0093465

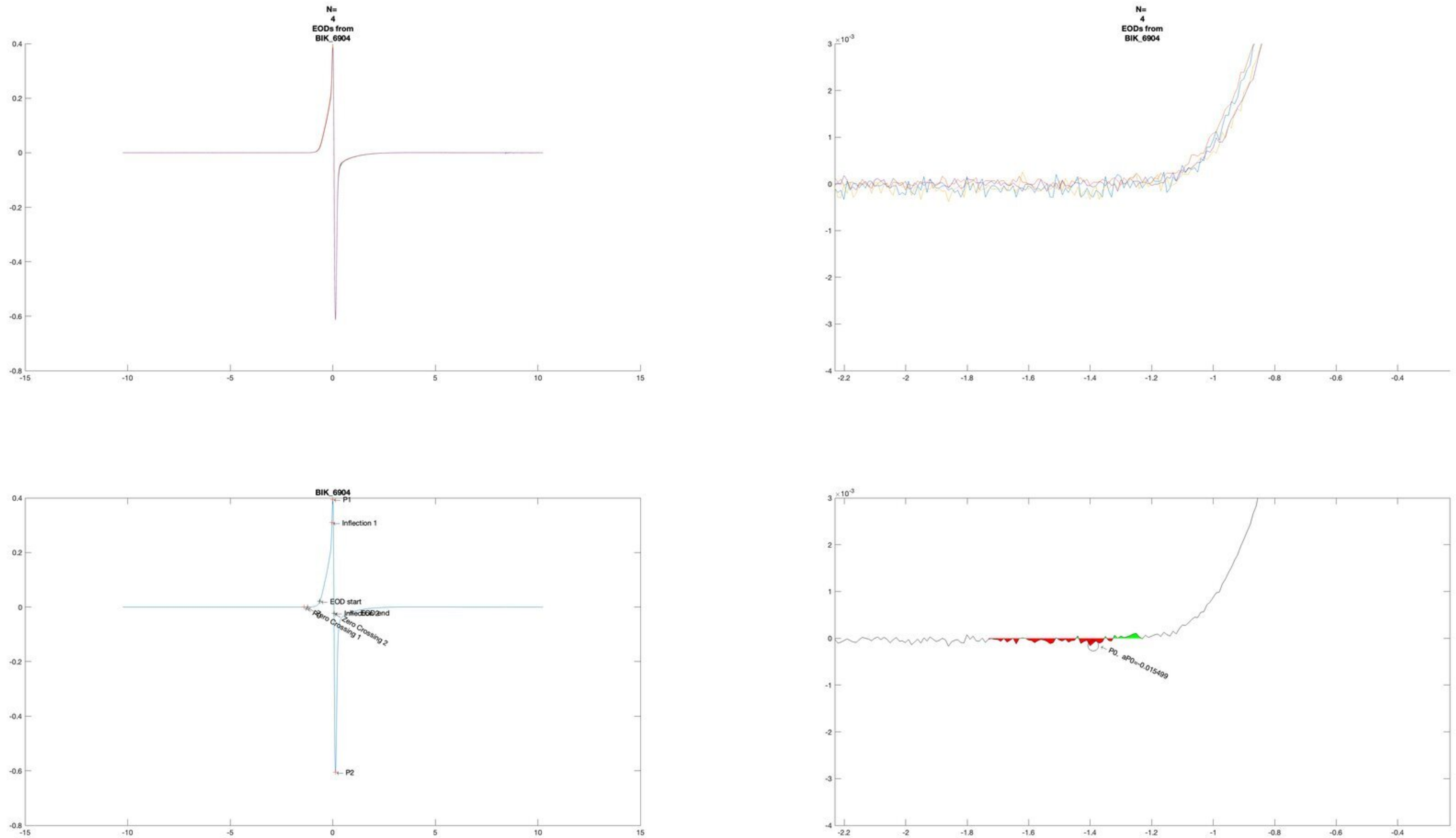
N=
4
EODs from
BIK_6904
N=
4
EODs from
BIK_6904
BIK_6904
P1
Inflection 1
EOD start
Inflection 2
EOD end
Zero Crossing 1
Zero Crossing 2
P2
P0, aP0=-0.015499

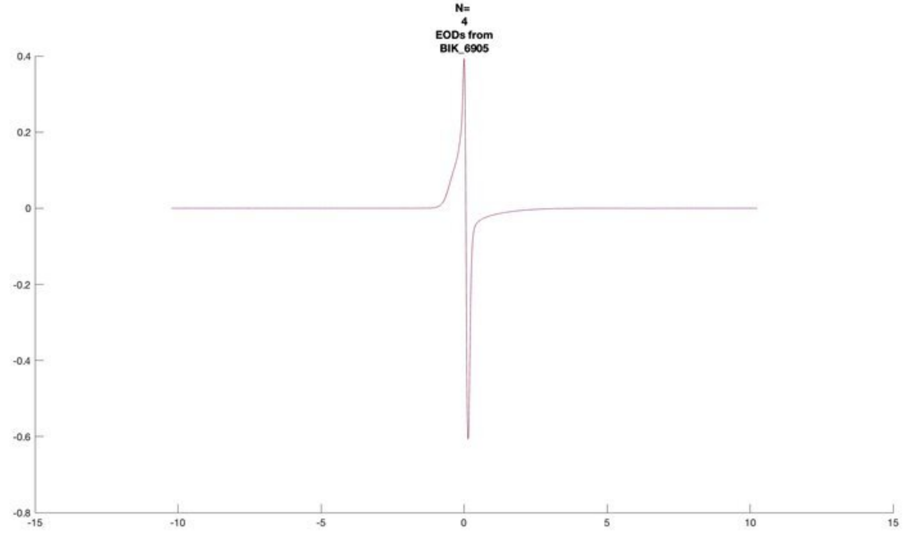
N=
4
EODs from
BIK_6905

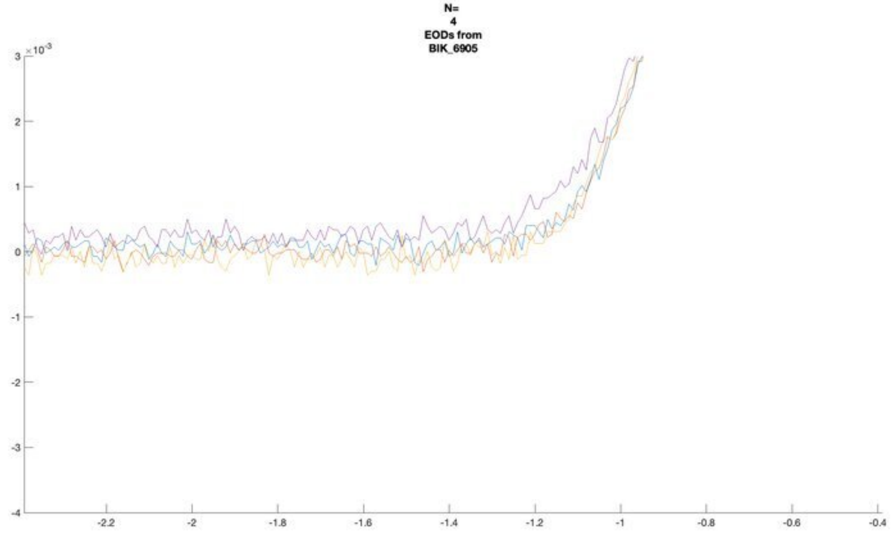
N=
4
EODs from
BIK_6905
×10^-3

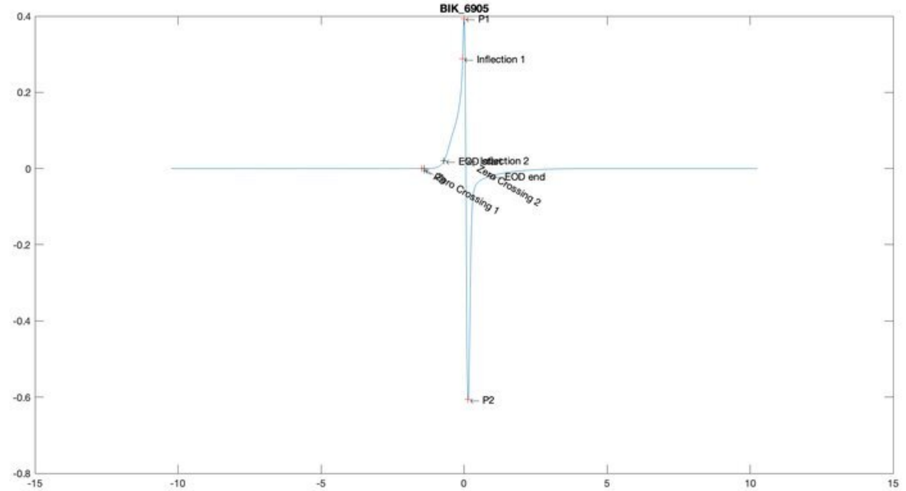
BIK_6905
P1
Inflection 1
Inflection 2
EOD start
EOD end
Zero Crossing 1
Zero Crossing 2
P2

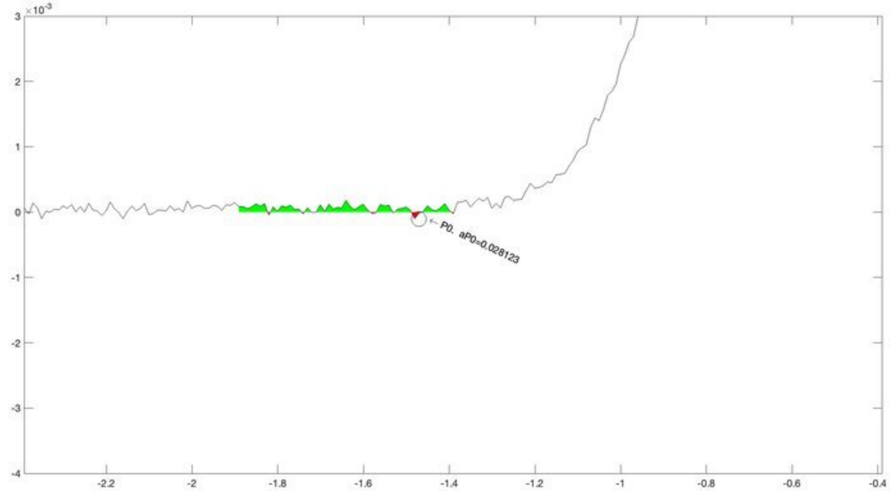
×10^-3
P0. aP0=0.028123

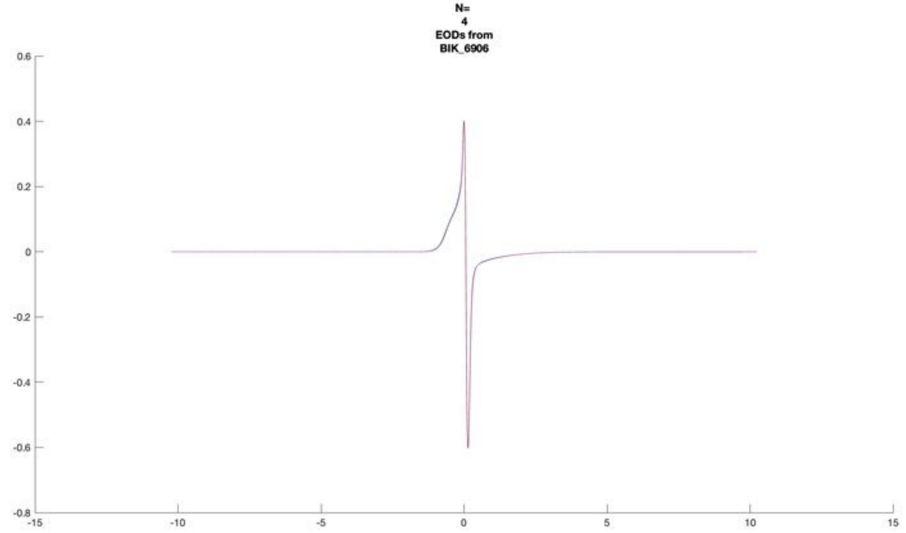
N=
4
EODs from
BIK_6906

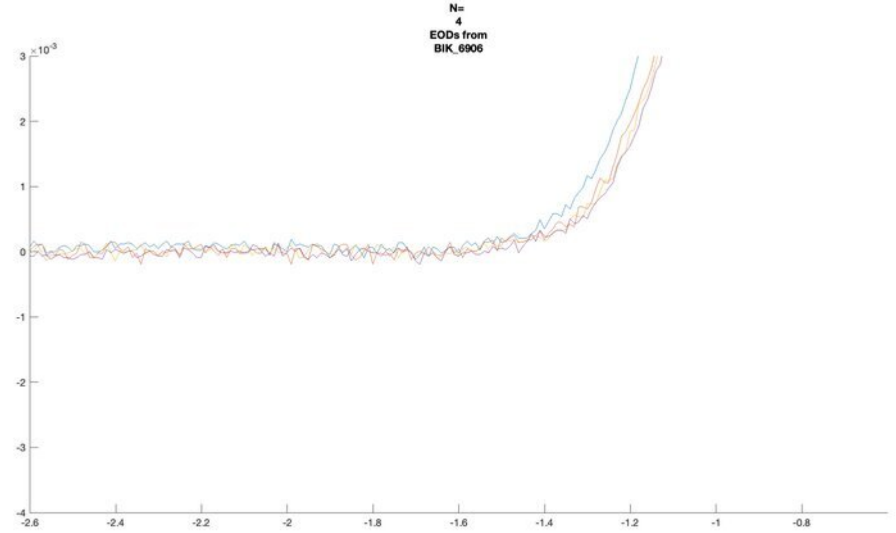
N=
4
EODs from
BIK_6906

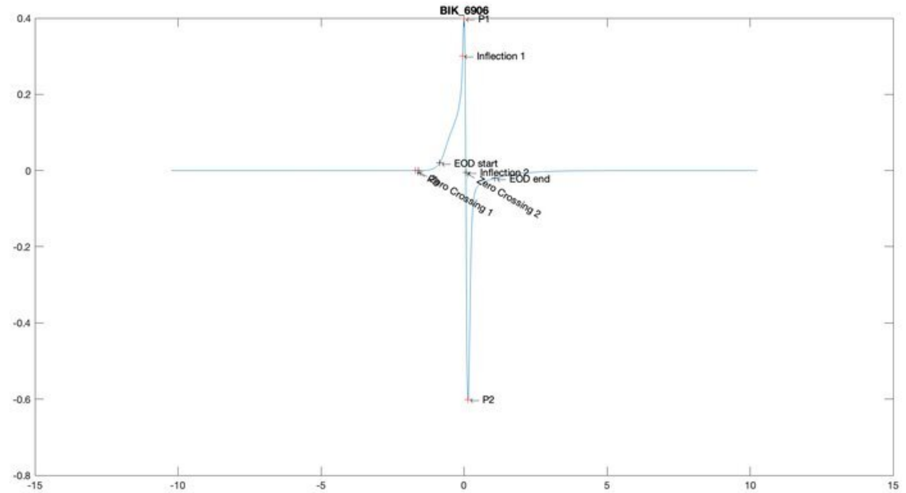
BIK_6906
P1
Inflection 1
EOD start
Inflection 2
EOD end
Zero Crossing 1
Zero Crossing 2
P2

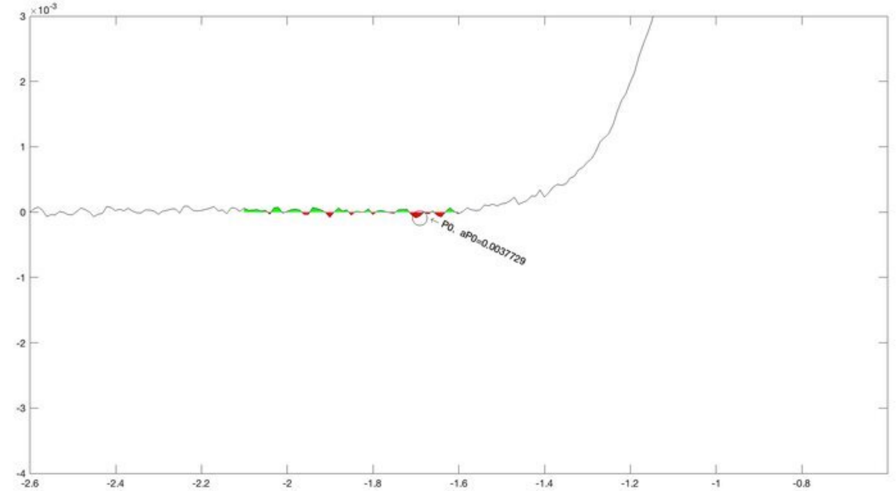
P0, aP0=0.0037729

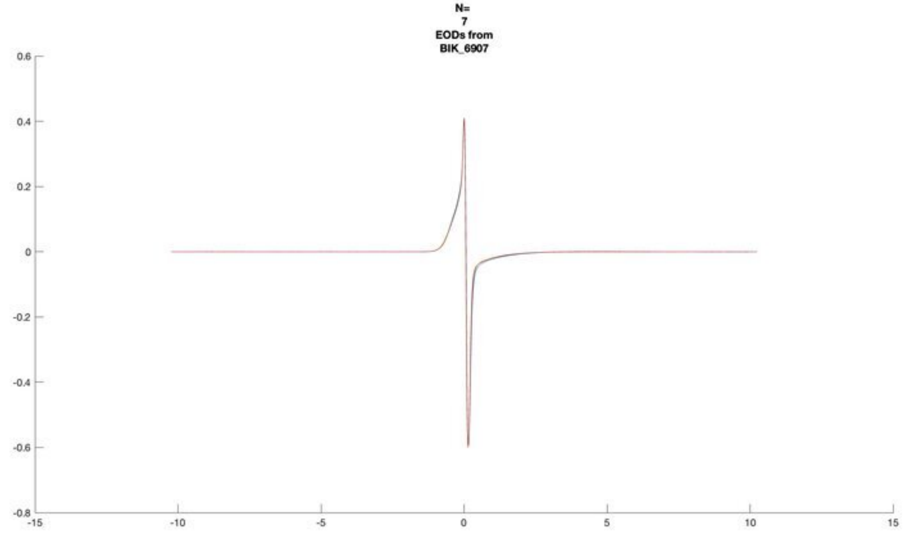
N=
7
EODs from
BIK_6907

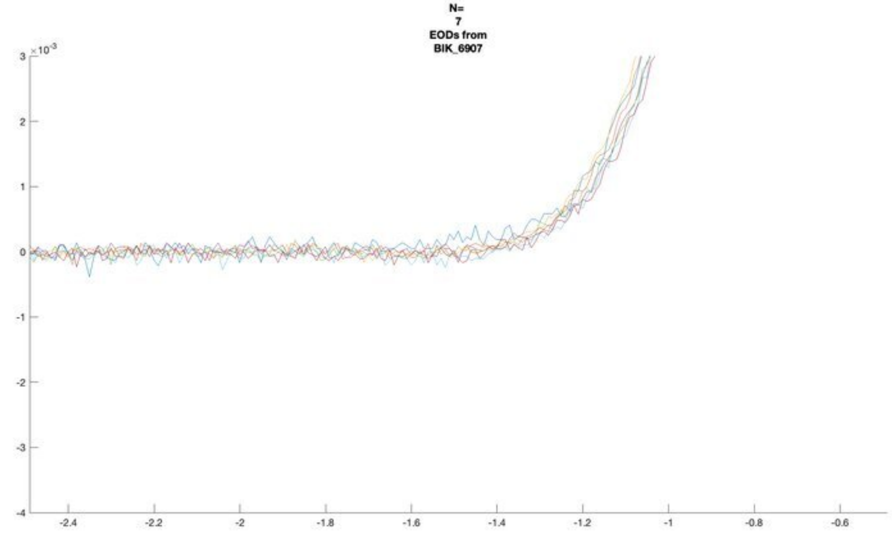
N=
7
EODs from
BIK_6907

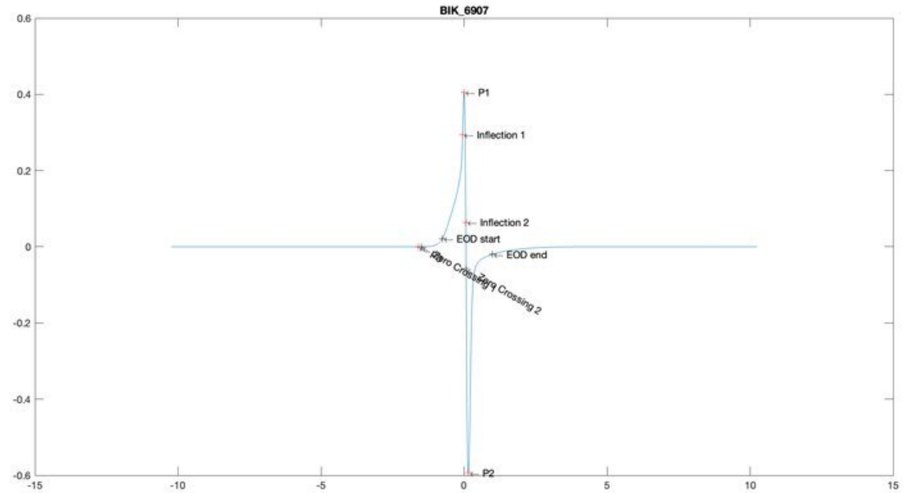
BIK_6907
P1
Inflection 1
Inflection 2
EOD start
EOD end
Zero Crossing 1
Zero Crossing 2
P2

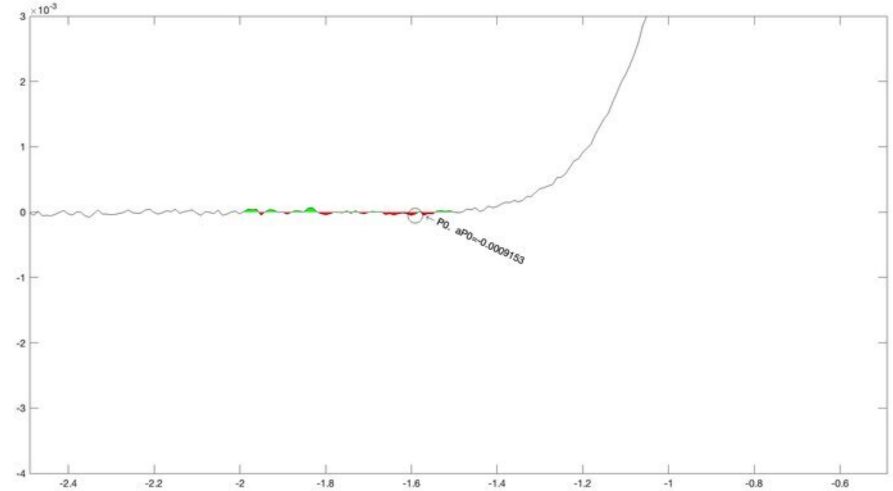
P0, aP0=-0.0009153

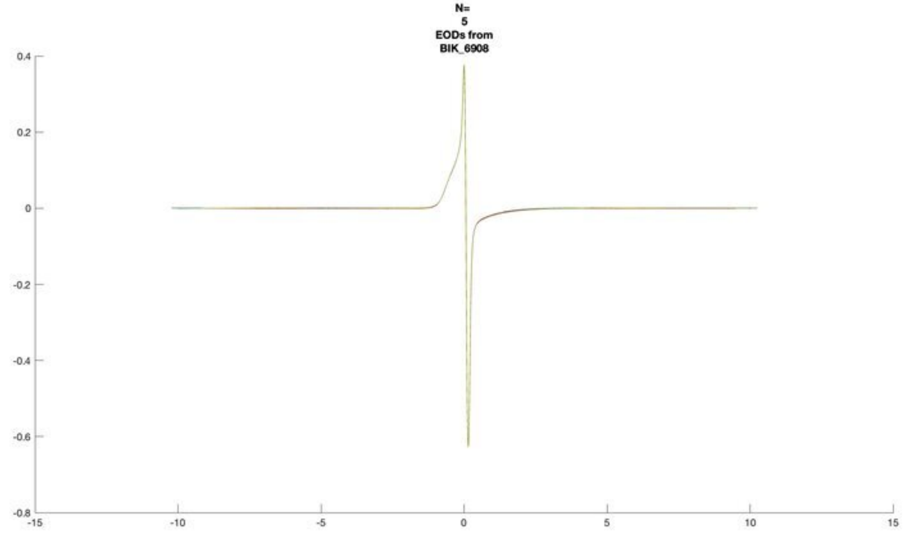
N=
5
EODs from
BIK_6908

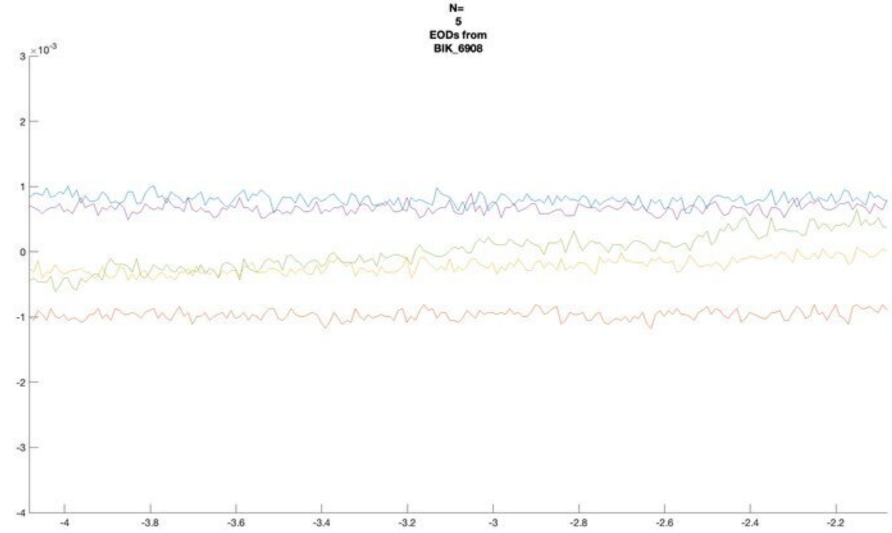
N=
5
EODs from
BIK_6908

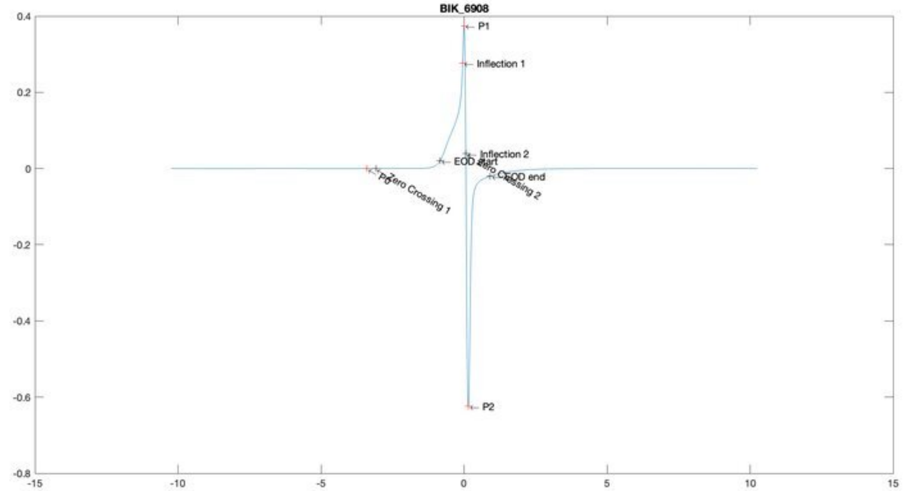
BIK_6908
P1
Inflection 1
Inflection 2
EOD start
EOD end
P0
Zero Crossing 1
Zero Crossing 2
P2

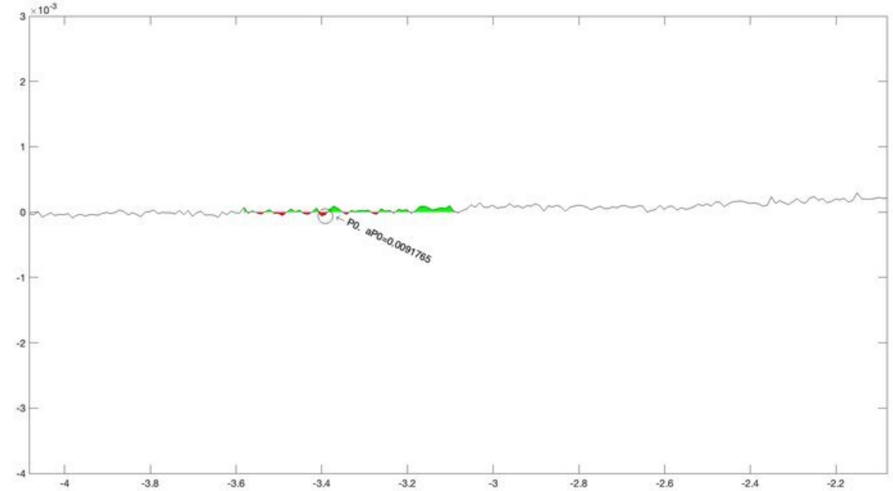
P0, aP0=0.0091765

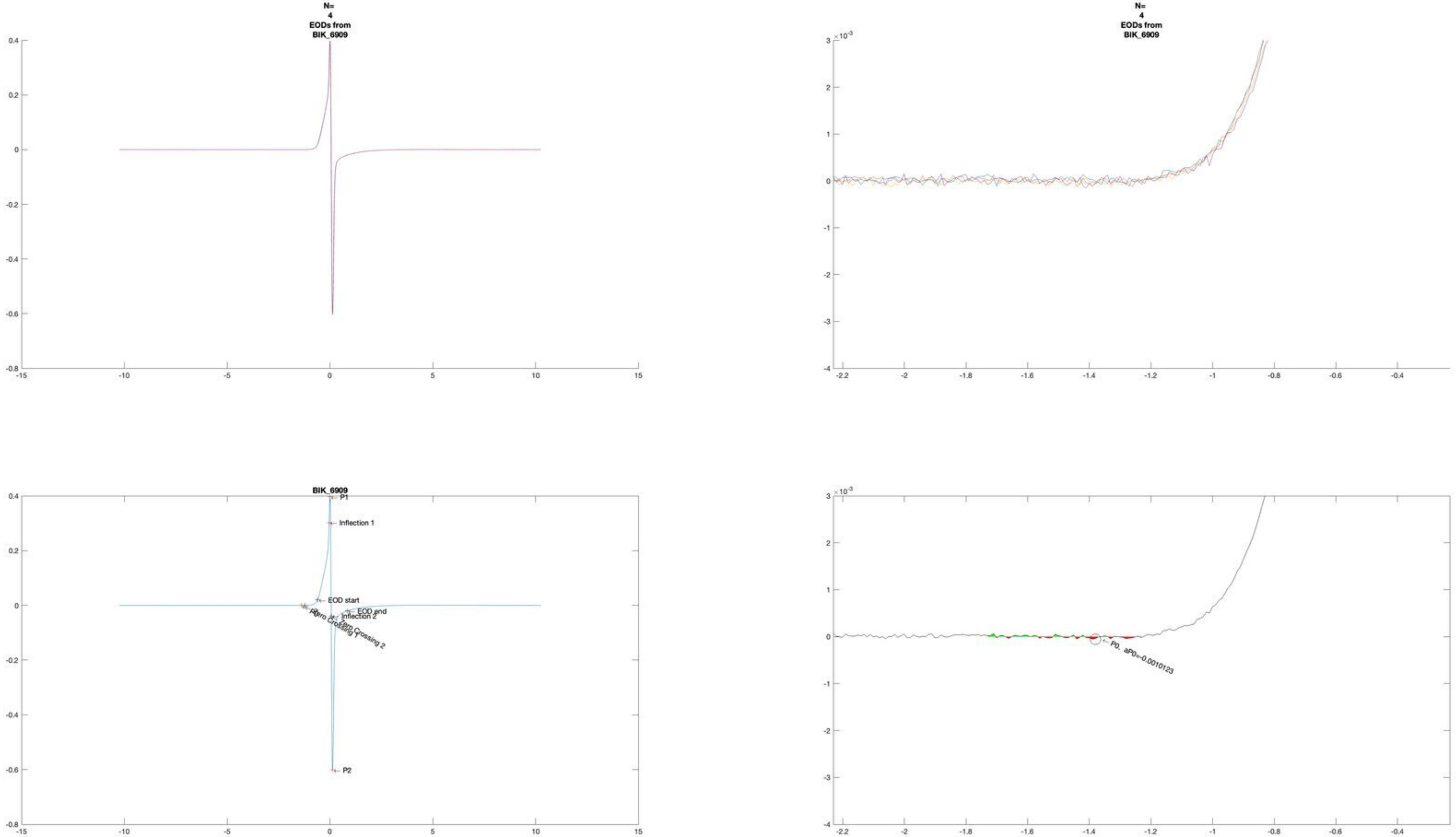
N=
4
EODs from
BIK_6909
BIK_6909
P1
Inflection 1
EOD start
EOD end
Inflection 2
Zero Crossing 1
Zero Crossing 2
P2
P0, aP0=-0.0010123

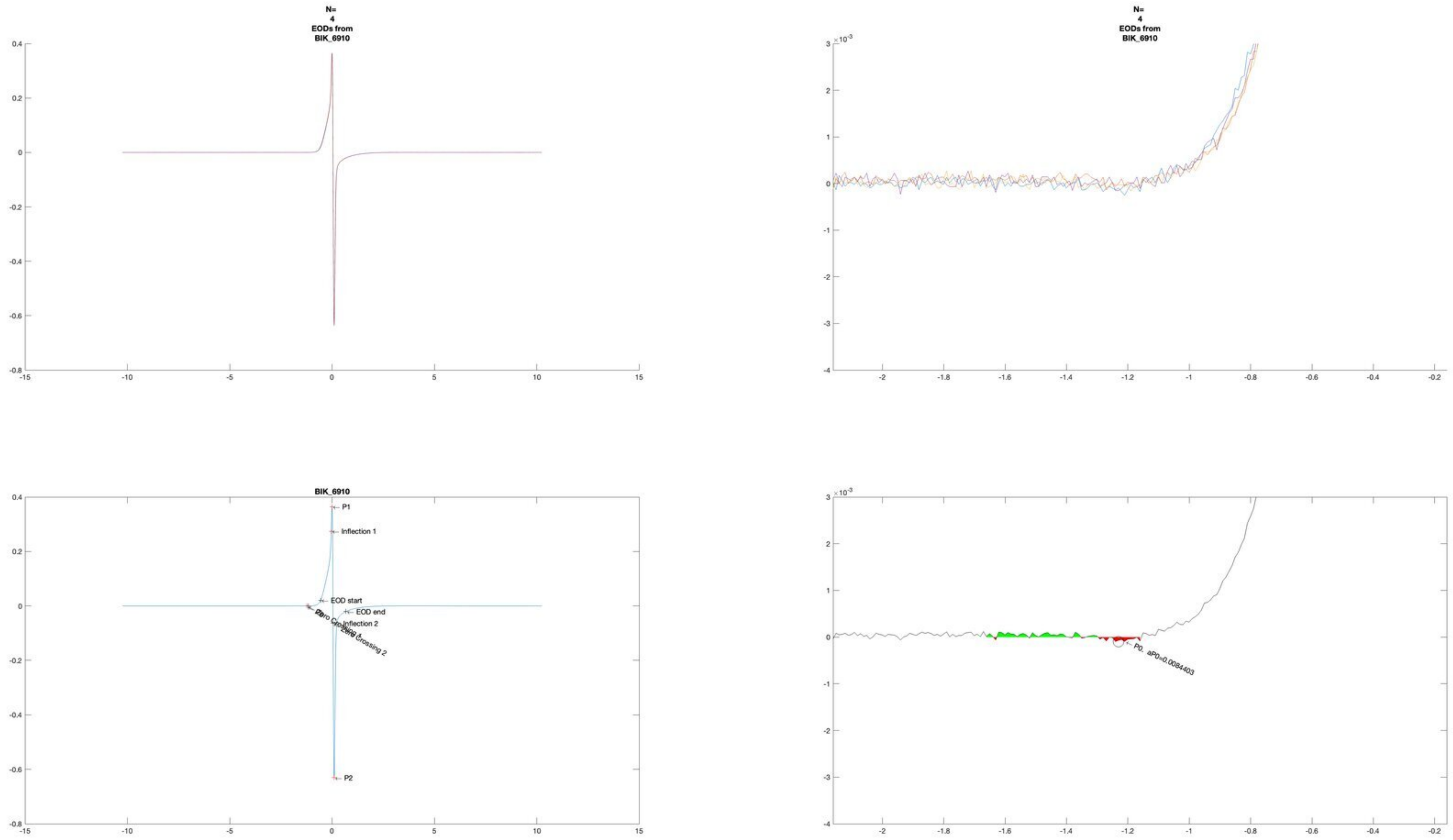
N=
4
EODs from
BIK_6910
N=
4
EODs from
BIK_6910
BIK_6910
P1
Inflection 1
EOD start
EOD end
Inflection 2
Zero Crossing 2
P2
P0, aP0=0.0084403

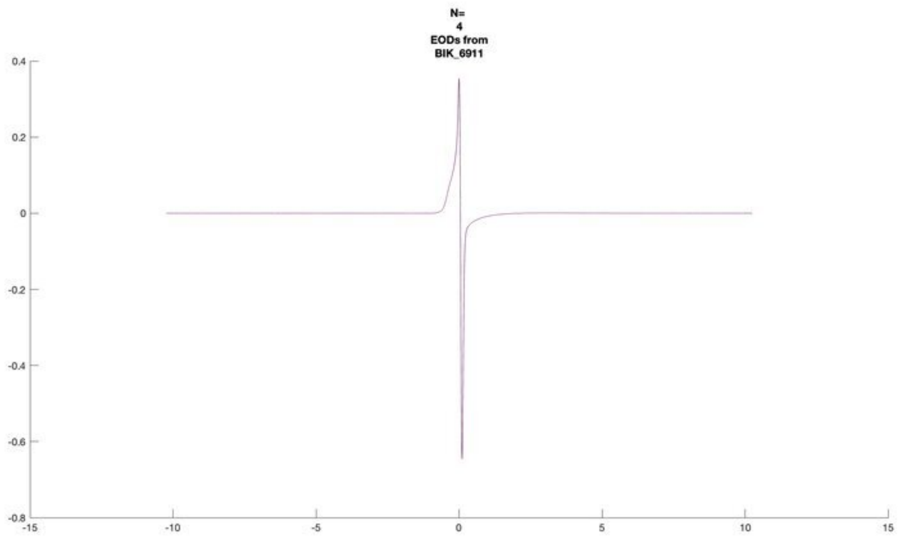

N=
4
EODs from
BIK_6911


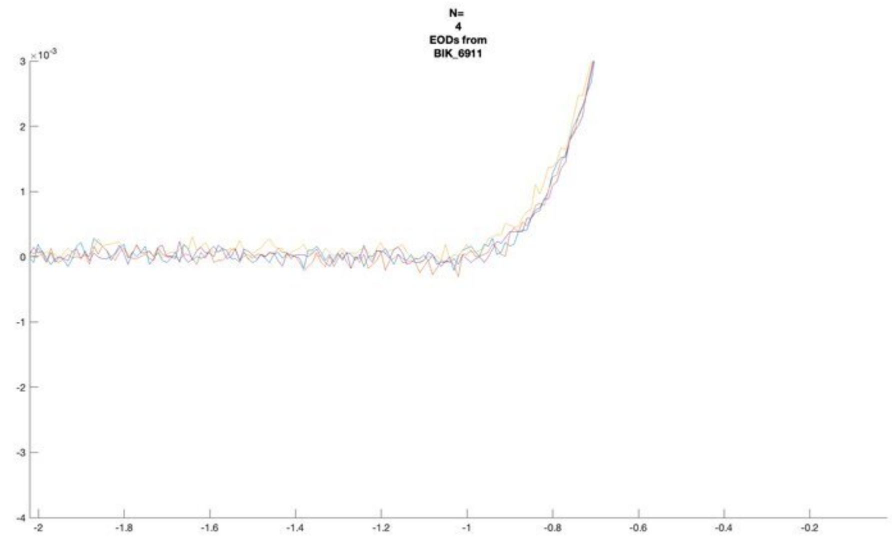

N=
4
EODs from
BIK_6911


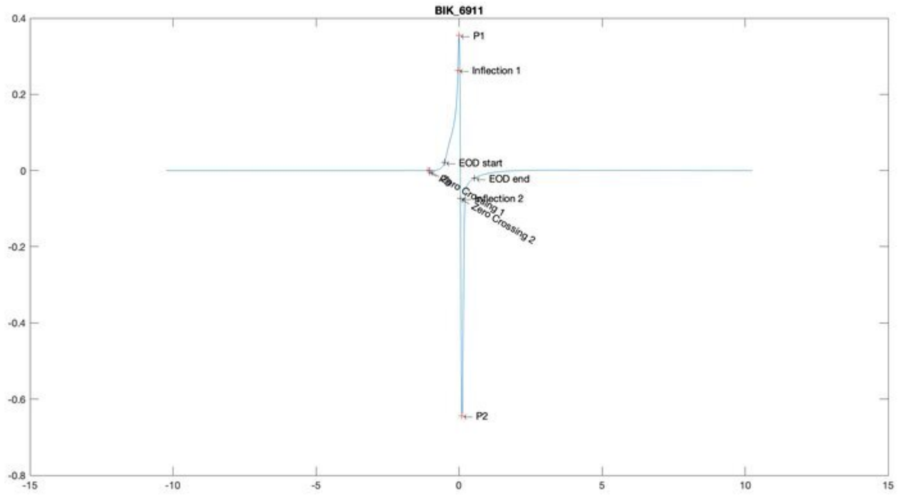

BIK_6911
P1
Inflection 1
EOD start
EOD end
Inflection 2
Zero Crossing 1
Zero Crossing 2
P2


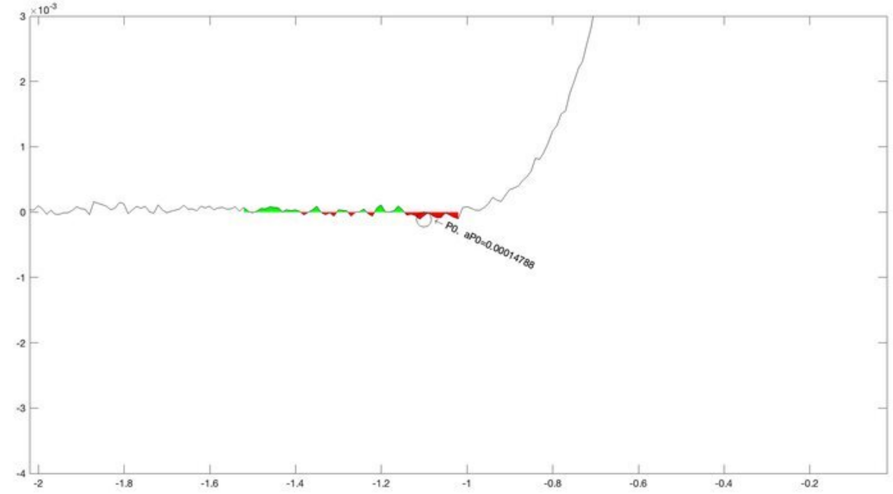

P0, aP0=0.00014788

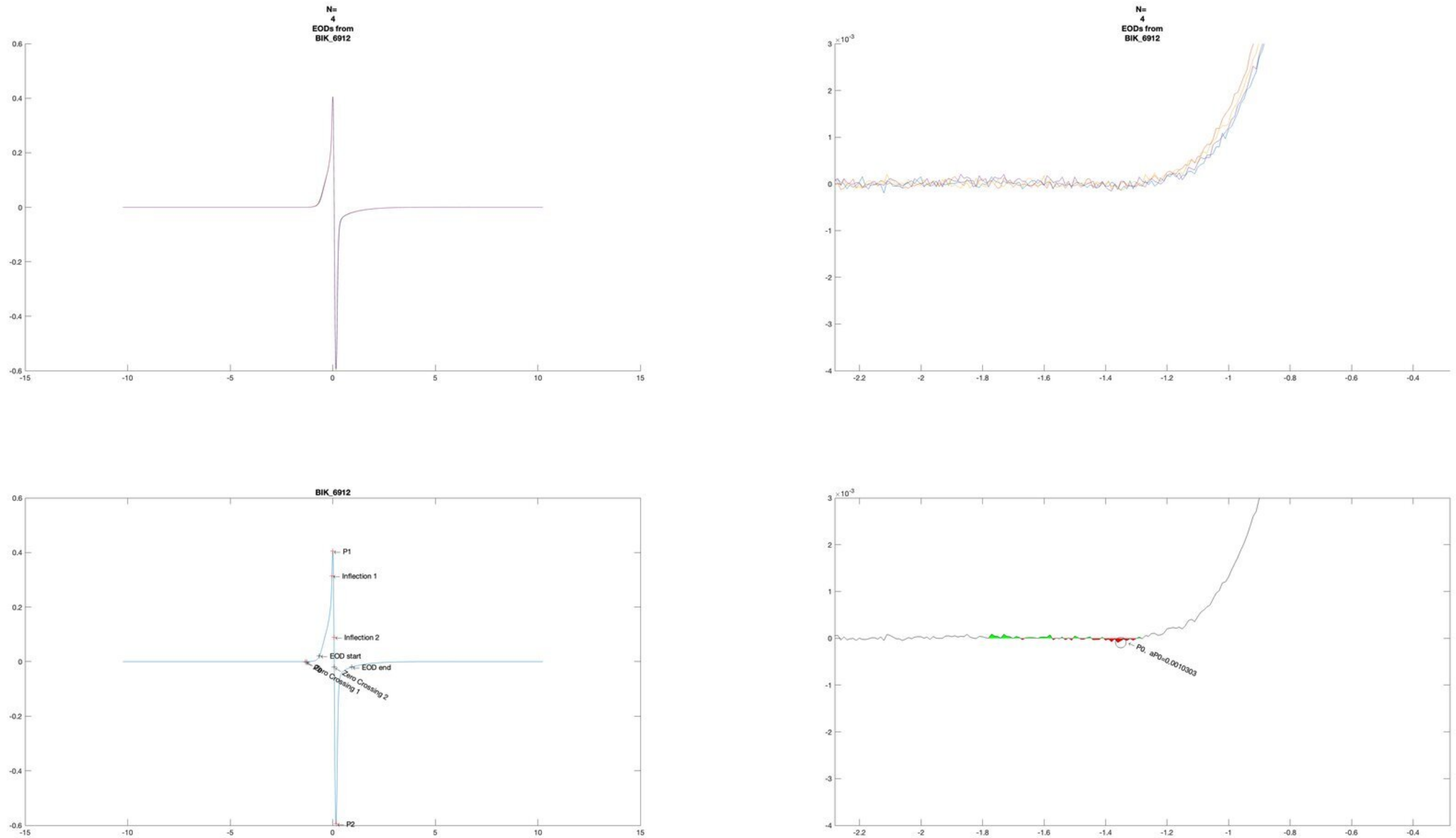
N=
4
EODs from
BIK_6912
N=
4
EODs from
BIK_6912
BIK_6912
P1
Inflection 1
Inflection 2
EOD start
Zero Crossing 1
Zero Crossing 2
EOD end
P2
P0, aP0=0.0010303

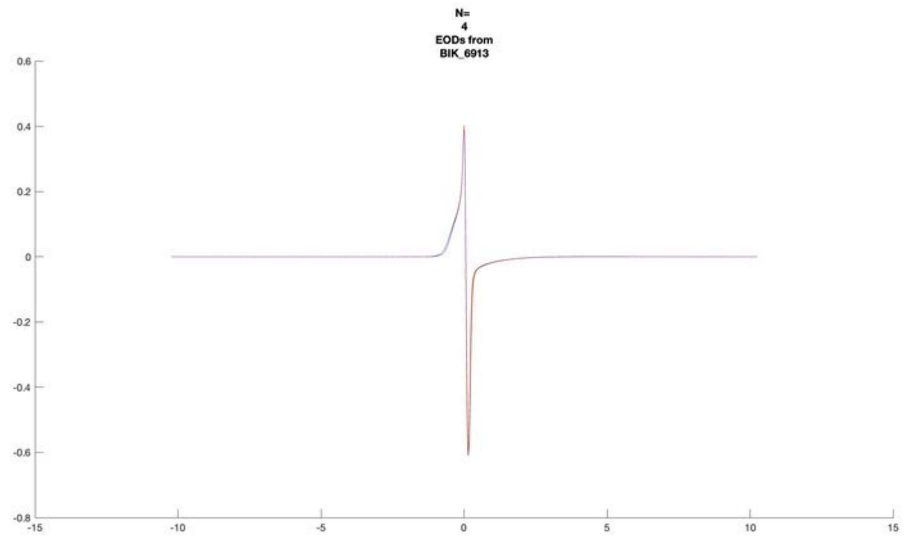
N=
4
EODs from
BIK_6913

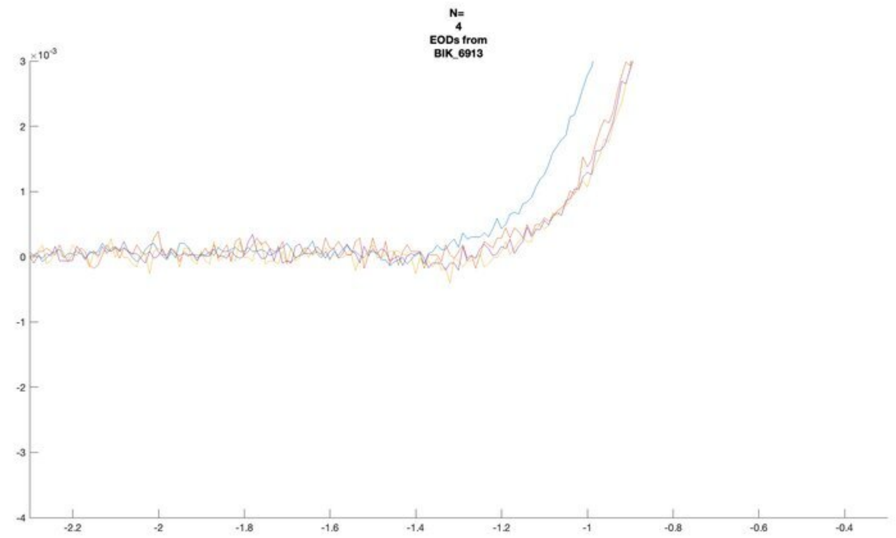
N=
4
EODs from
BIK_6913

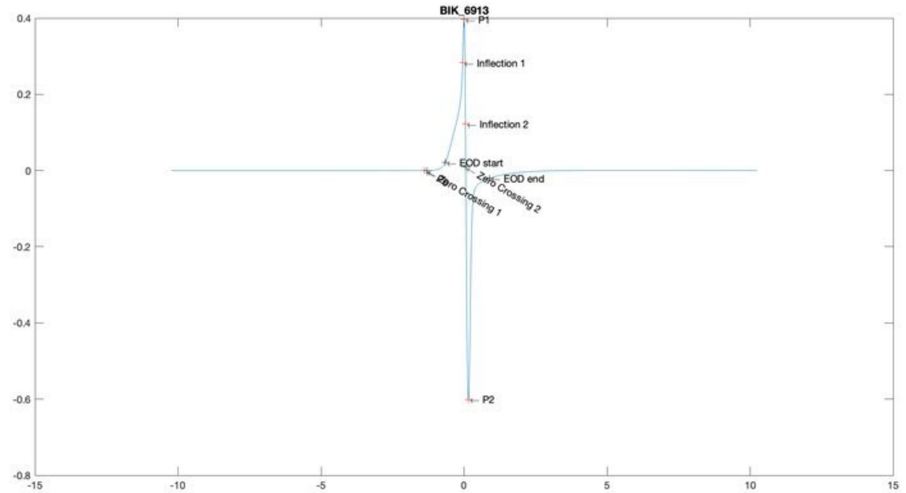
BIK_6913
P1
Inflection 1
Inflection 2
EOD start
EOD end
Zero Crossing 1
Zero Crossing 2
P2

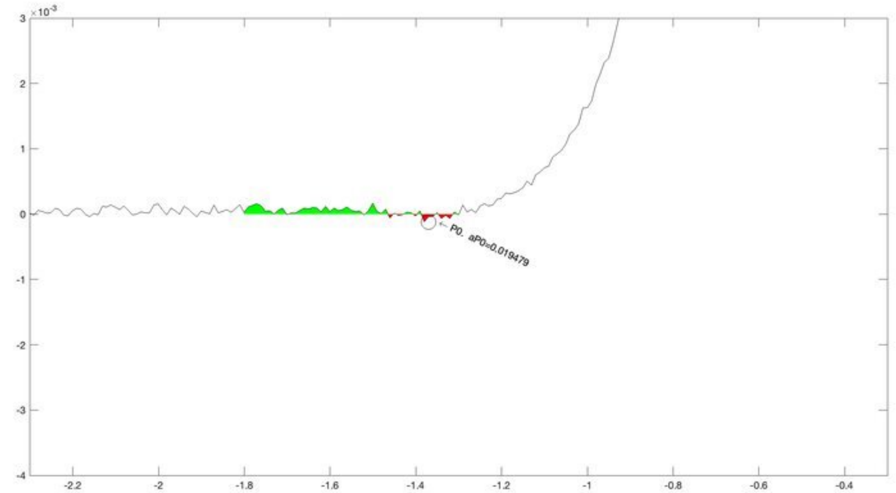
P0, aP0=0.019479

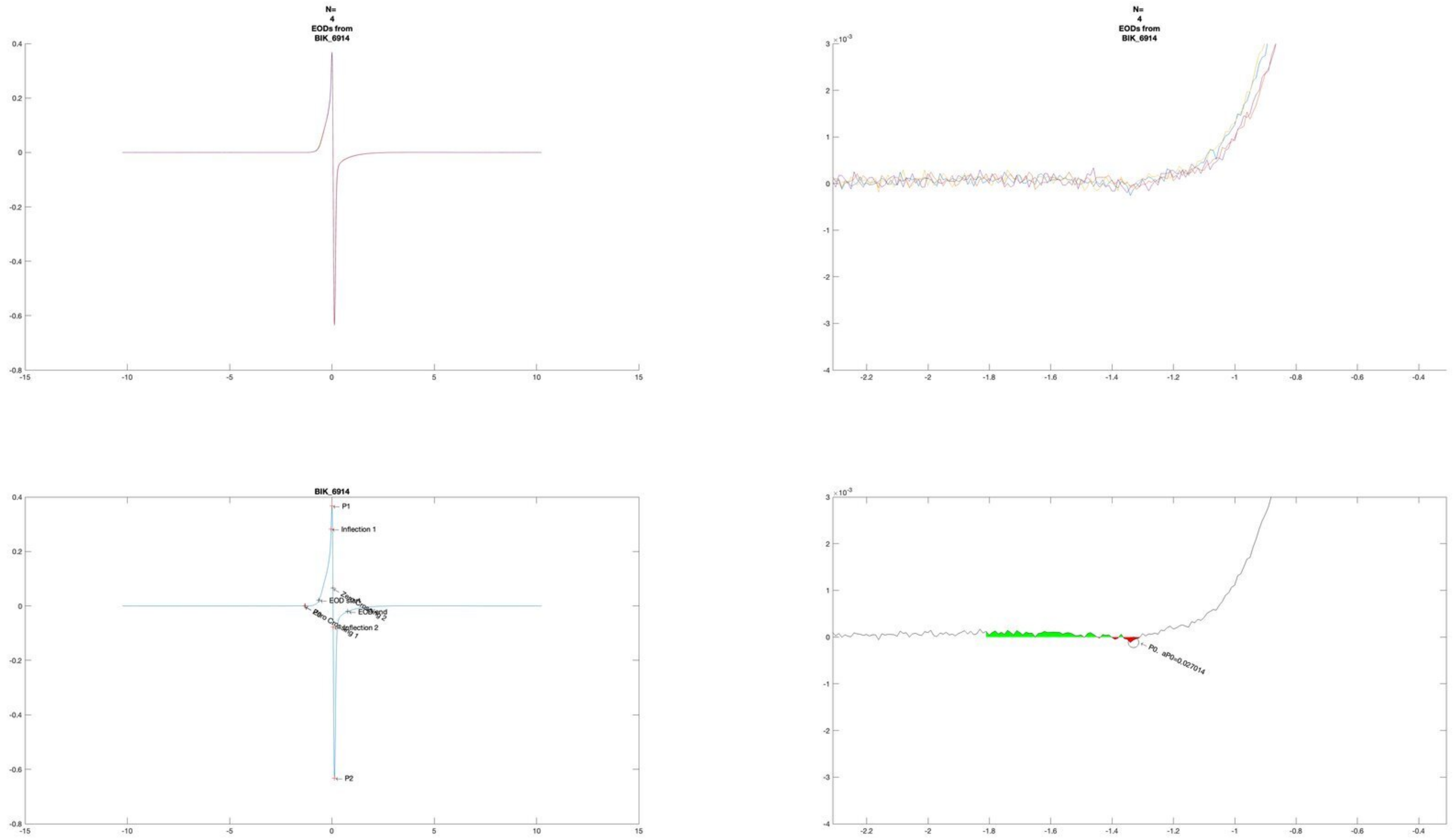
N=
4
EODs from
BIK_6914
N=
4
EODs from
BIK_6914
BIK_6914
P1
Inflection 1
EOD start
EOD end
Zero Crossing 1
Zero Crossing 2
Inflection 2
P0
P2
P0. aP0=0.027014

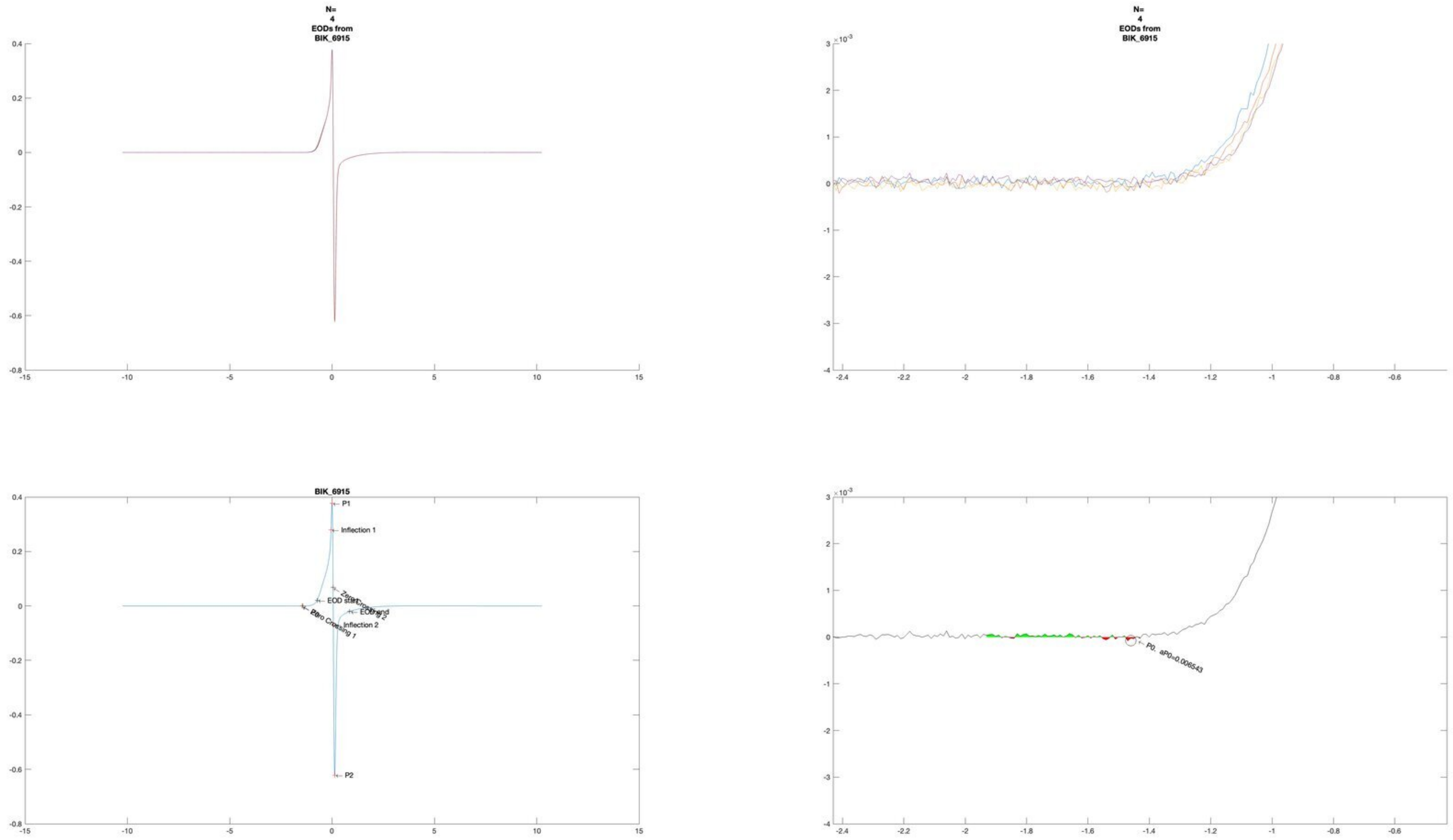
N=
4
EODs from
BIK_6915
N=
4
EODs from
BIK_6915
BIK_6915
P1
Inflection 1
EOD start
EOD end
Inflection 2
P2
P0, aP0=0.006543

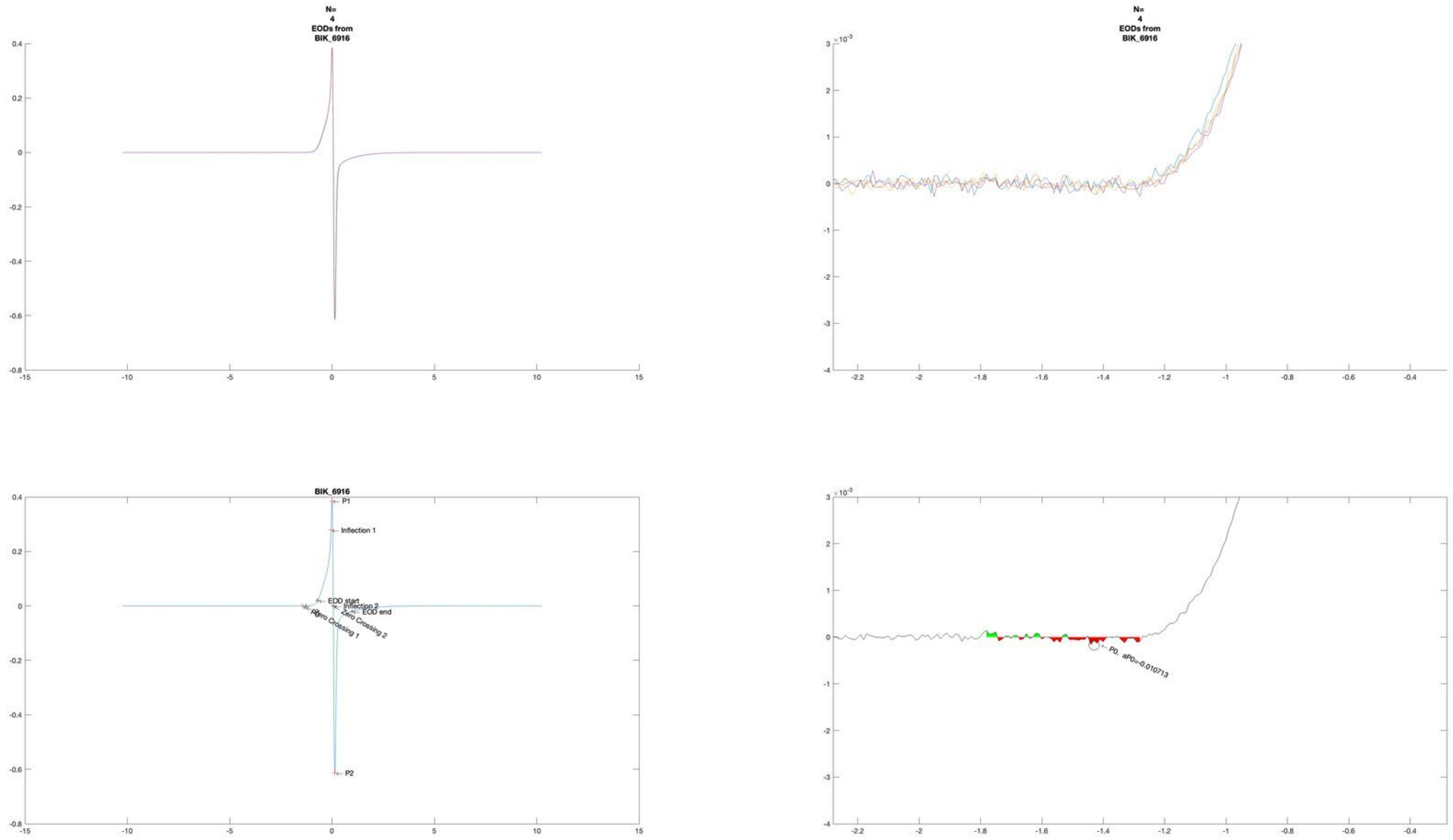
N=
4
EODs from
BIK_6916
N=
4
EODs from
BIK_6916
BIK_6916
P1
Inflection 1
EOD start
Inflection 2
EOD end
Zero Crossing 1
Zero Crossing 2
P2
P0, aP0=-0.010713

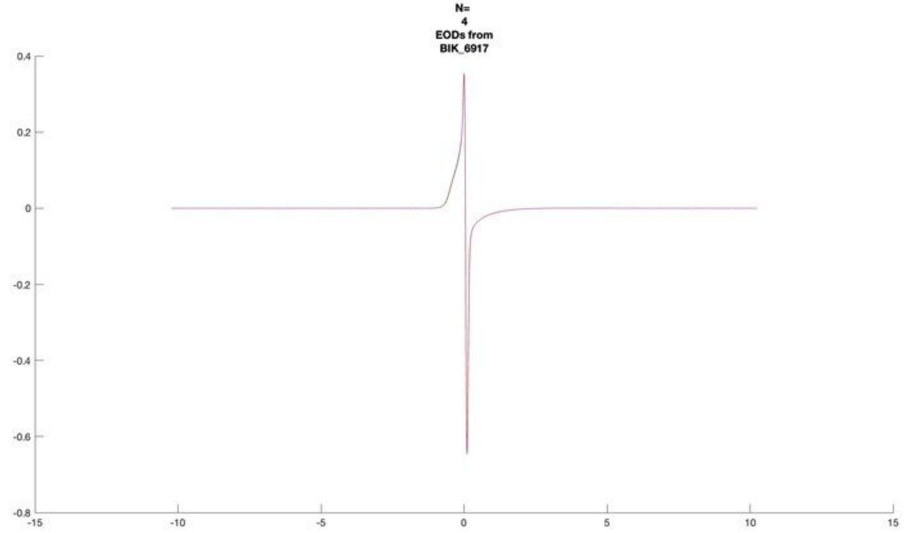
N=
4
EODs from
BIK_6917

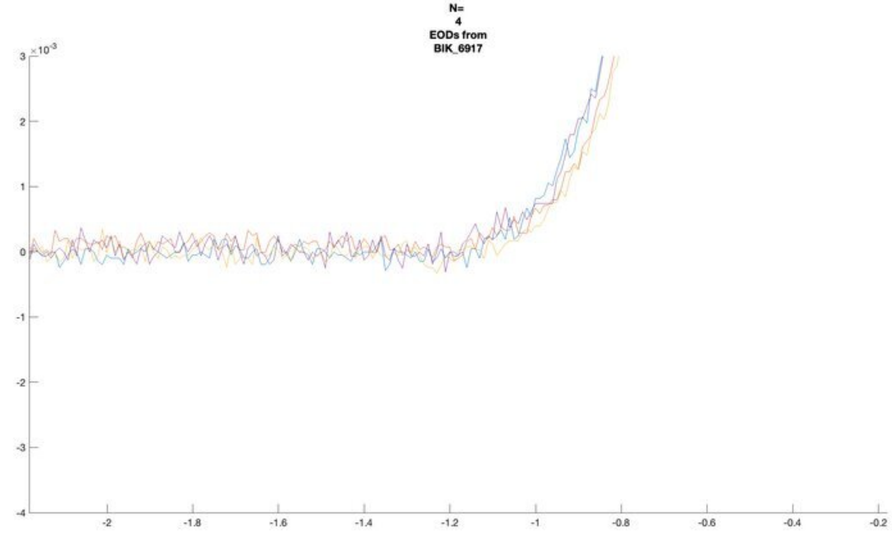
N=
4
EODs from
BIK_6917

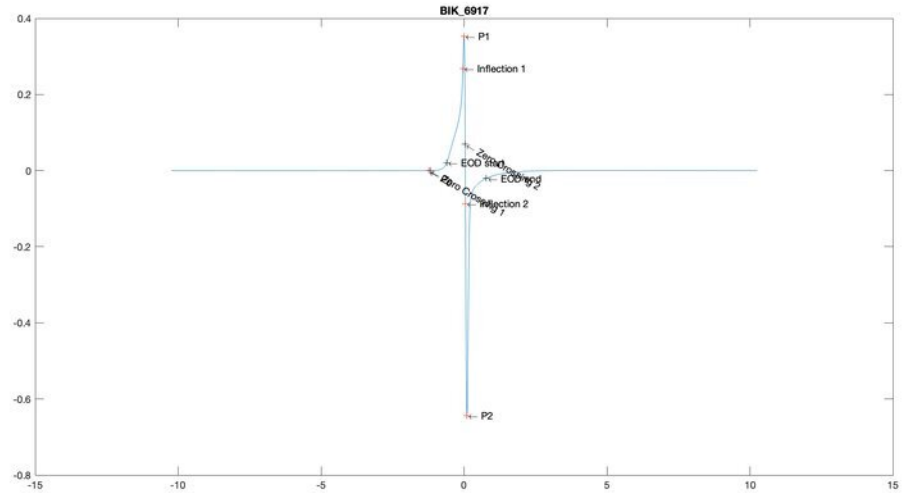
BIK_6917
P1
Inflection 1
EOD start
EOD end
Inflection 2
P2

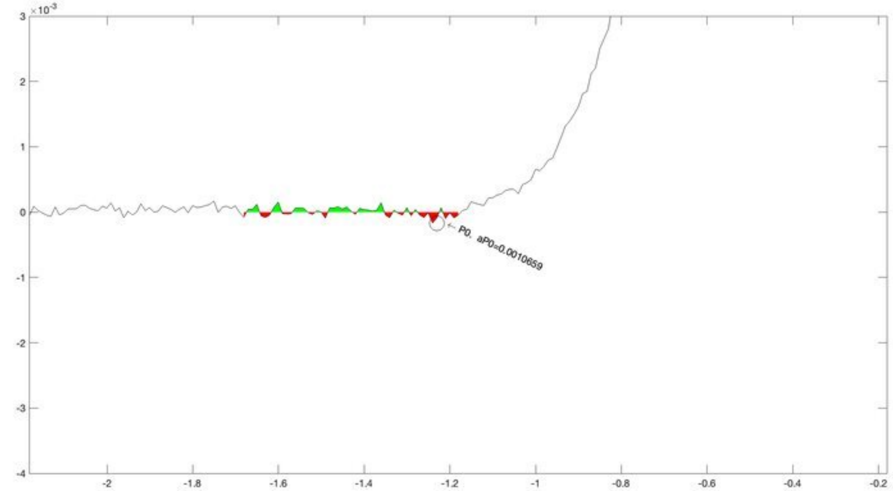
P0, aP0=0.0010659

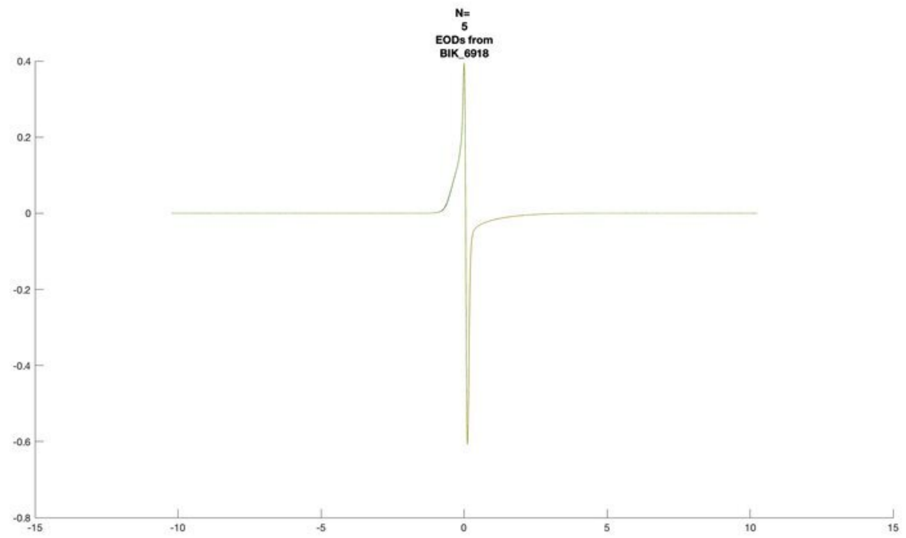
N=
5
EODs from
BIK_6918

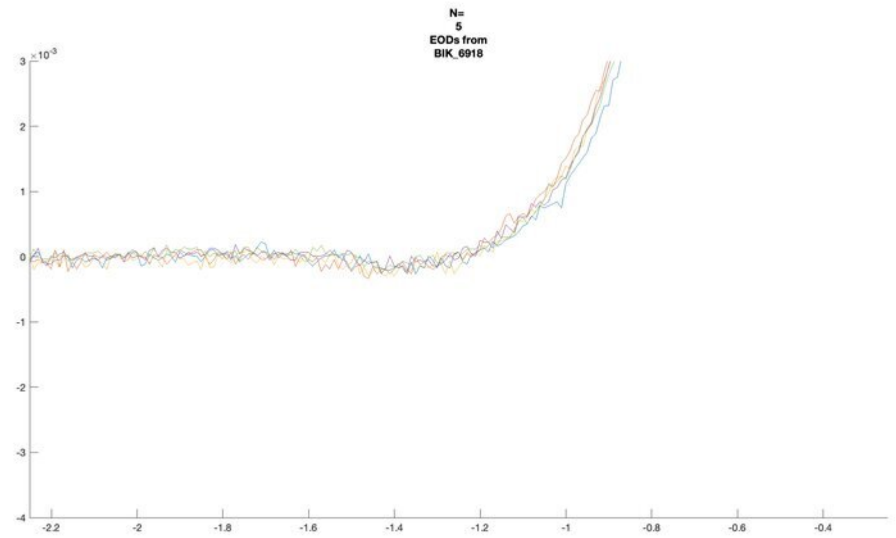
N=
5
EODs from
BIK_6918

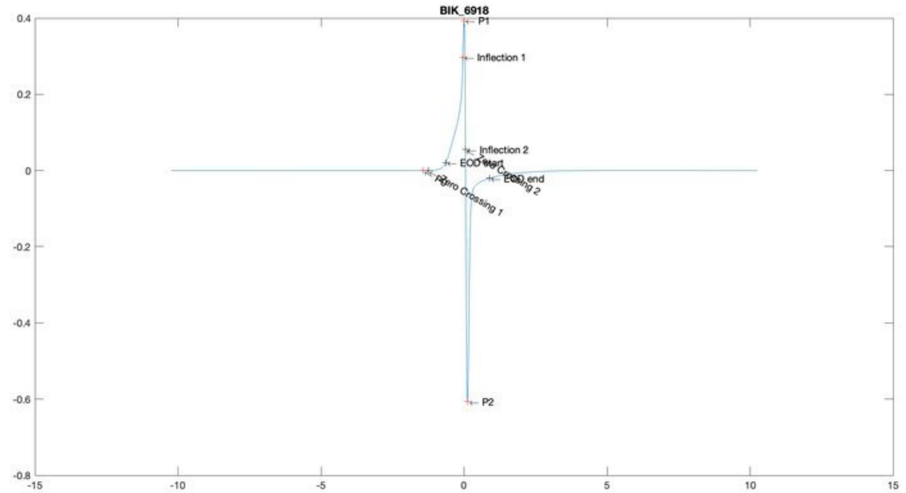
BIK_6918
P1
Inflection 1
Inflection 2
EOD start
EOD end
Zero Crossing 1
P2

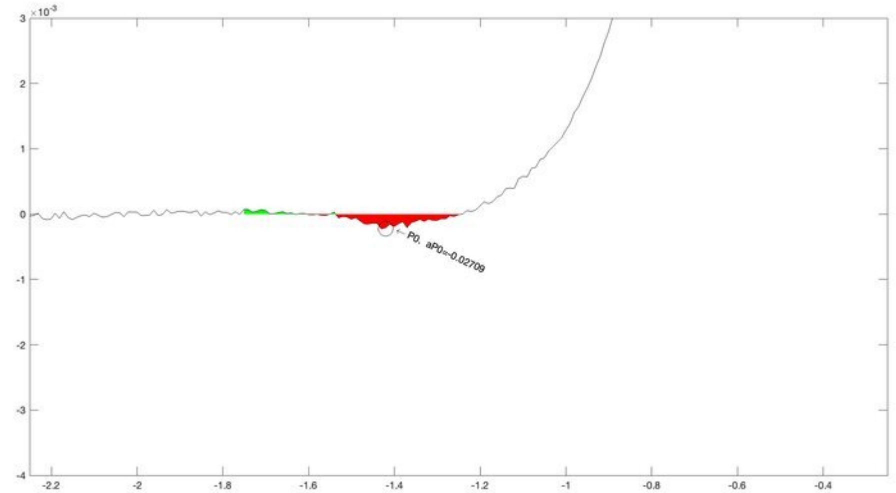
P0, aP0=-0.02709

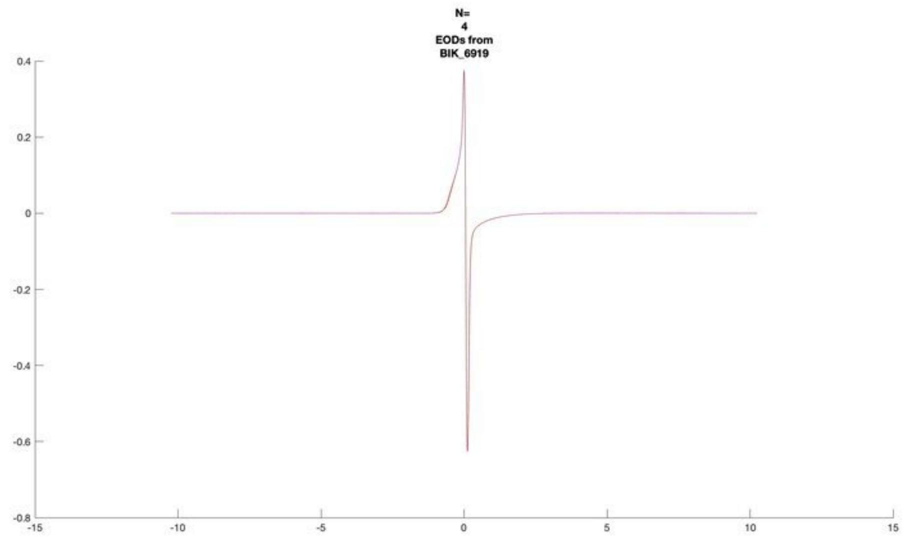
N=
4
EODs from
BIK_6919

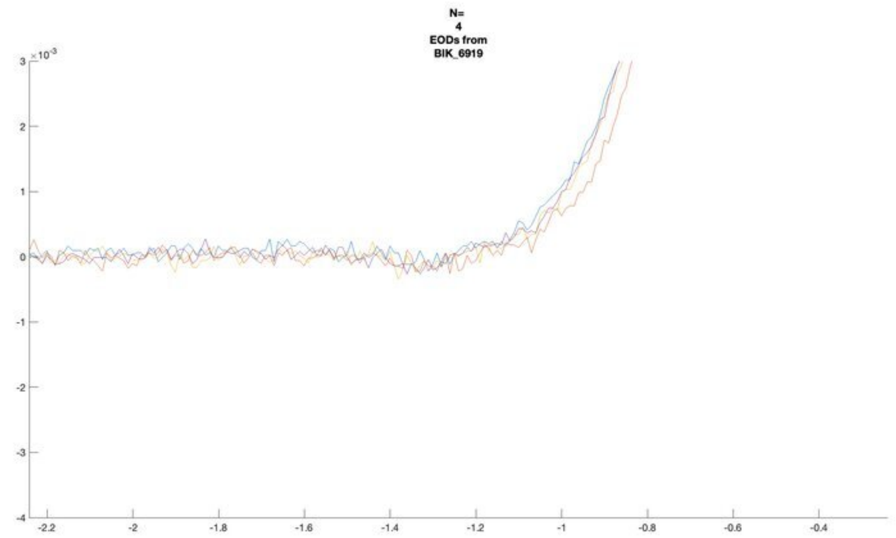
N=
4
EODs from
BIK_6919

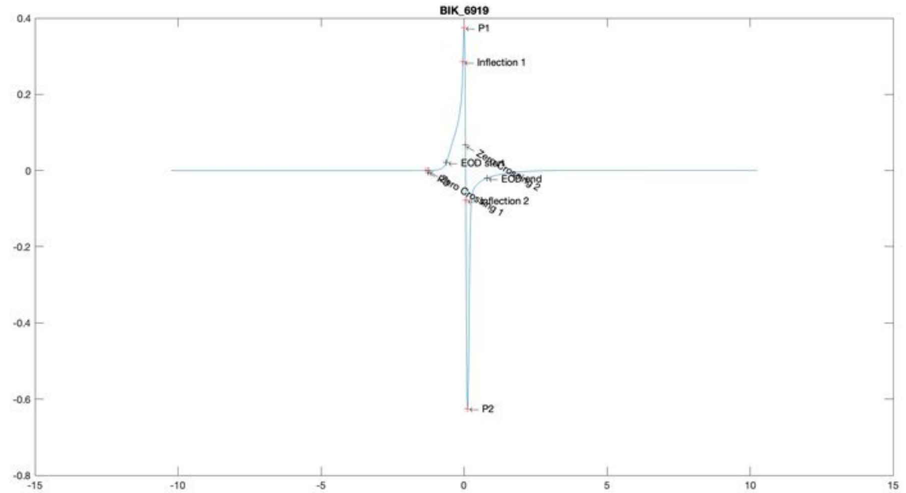
BIK_6919
P1
Inflection 1
EOD start
EOD end
Inflection 2
P2

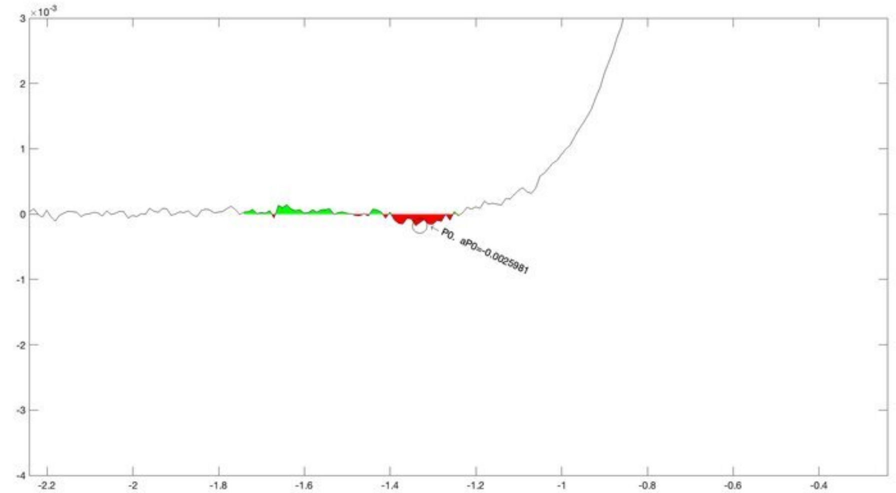
P0. aP0=-0.0025981

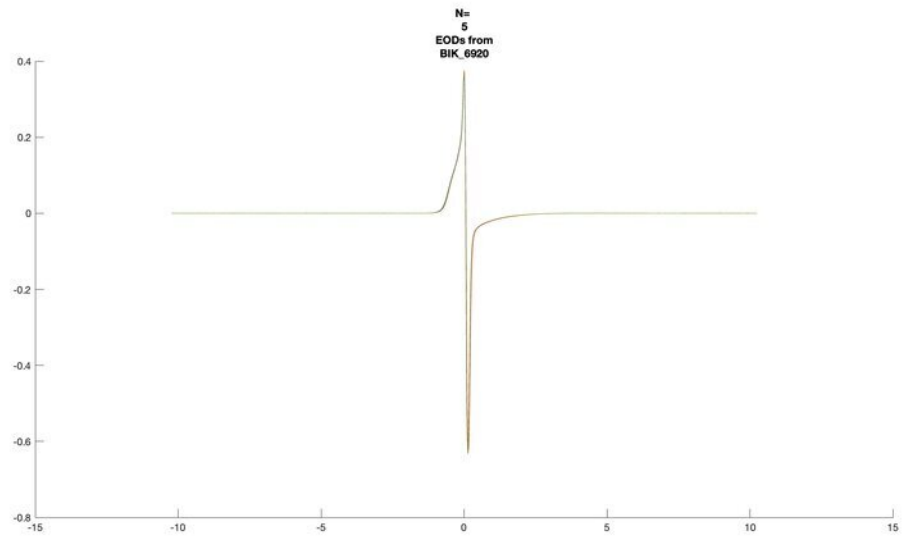
N=
5
EODs from
BIK_6920

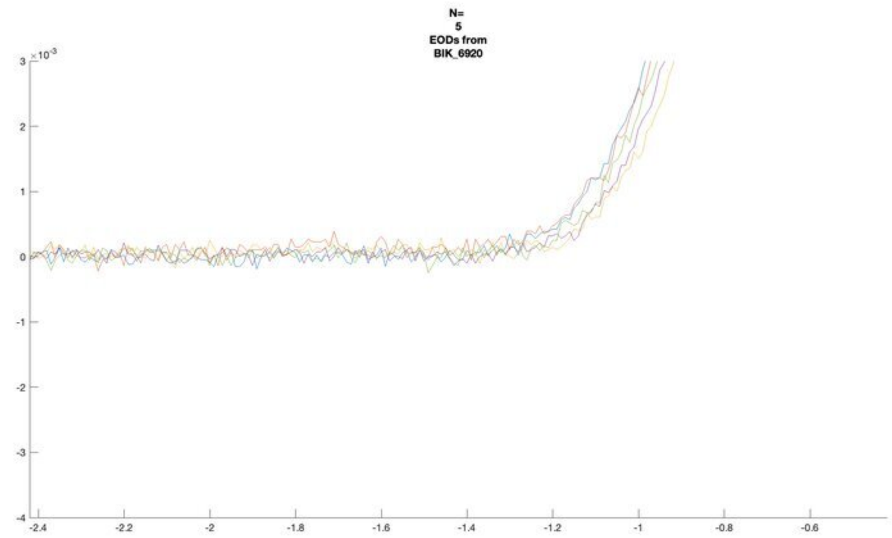
N=
5
EODs from
BIK_6920

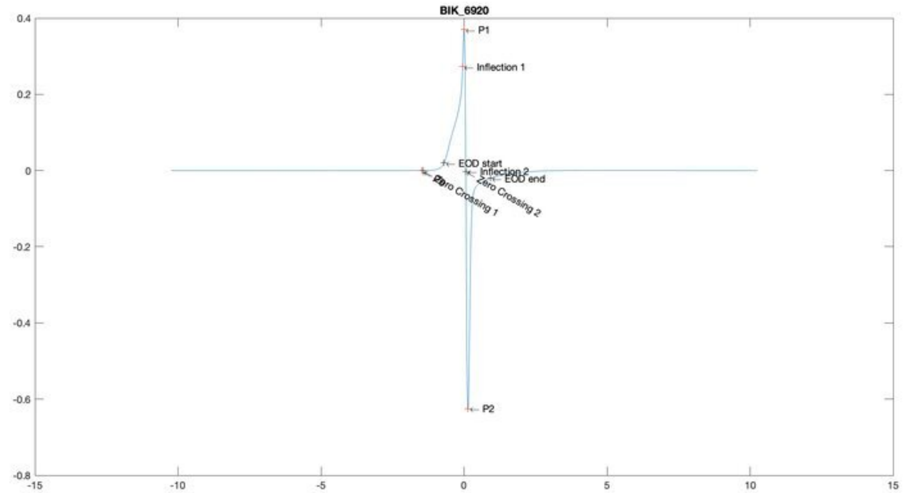
BIK_6920
P1
Inflection 1
EOD start
Inflection 2
EOD end
Zero Crossing 2
Zero Crossing 1
P2

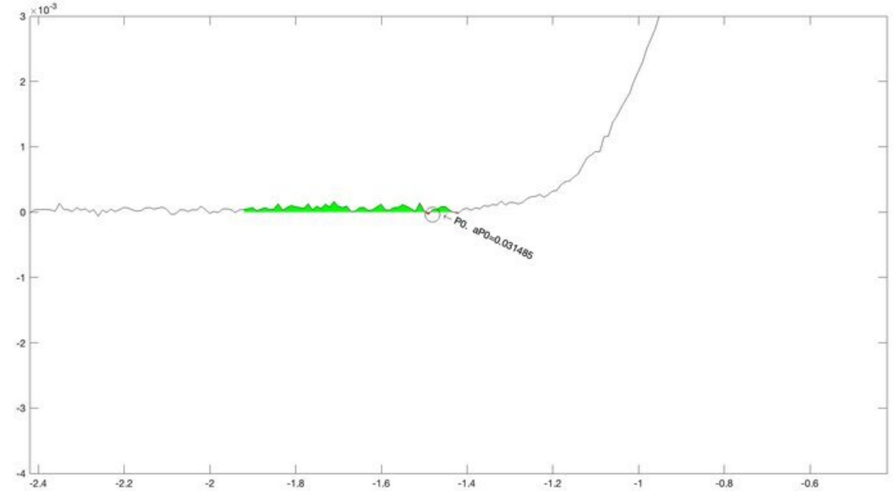
P0, aP0=0.031485

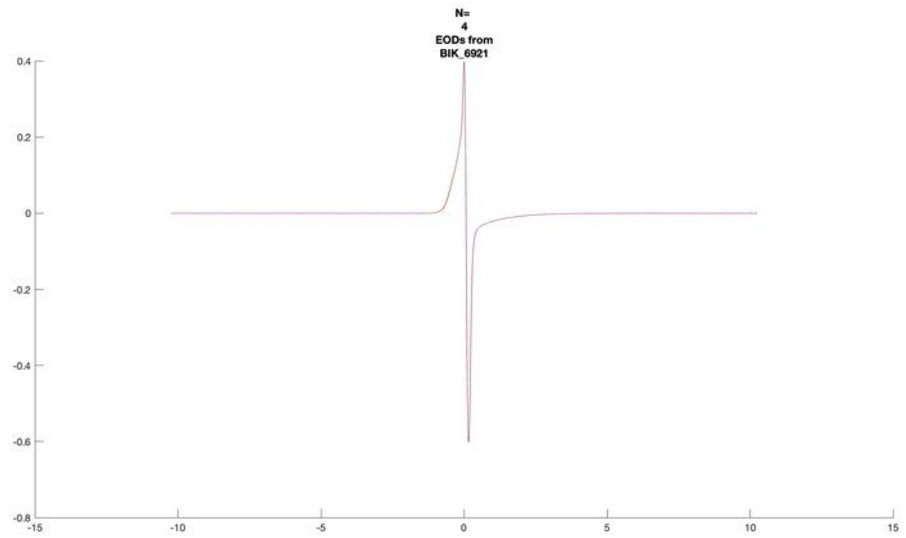
N=
4
EODs from
BIK_6921

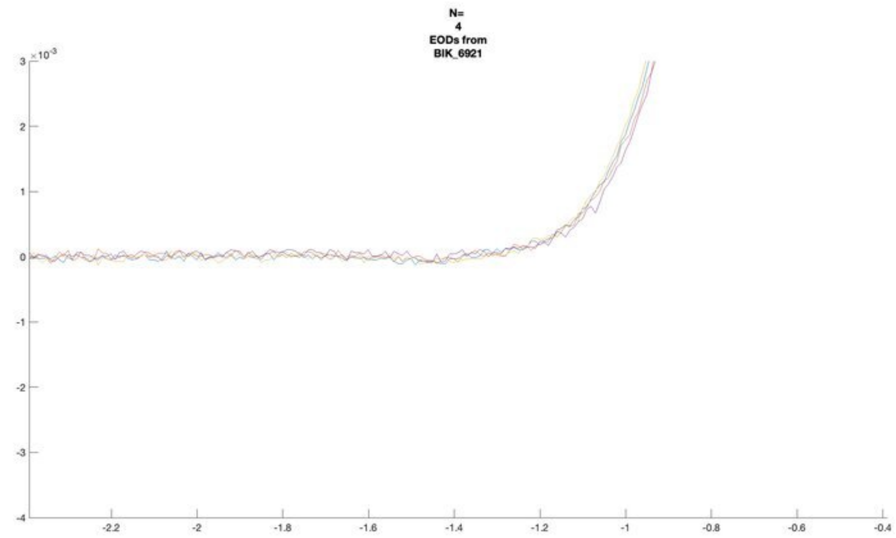
N=
4
EODs from
BIK_6921

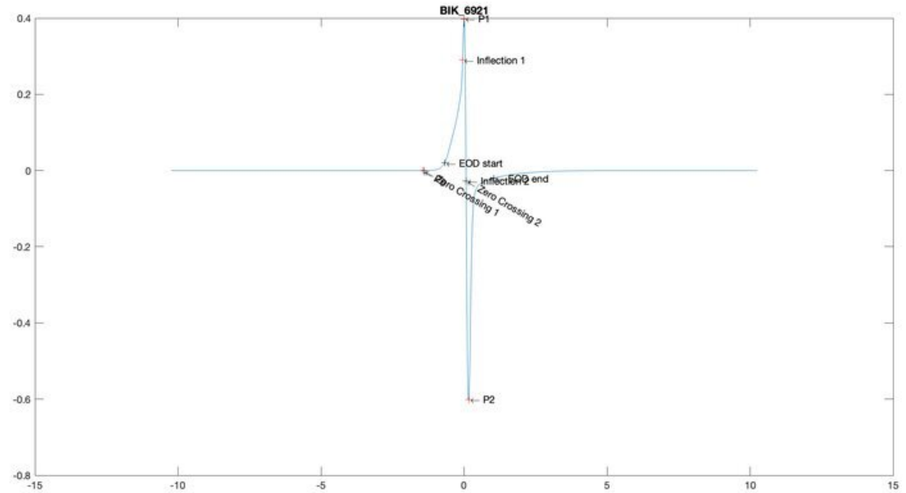
BIK_6921
P1
Inflection 1
EOD start
EOD end
Inflection 2
P0
Zero Crossing 1
Zero Crossing 2
P2

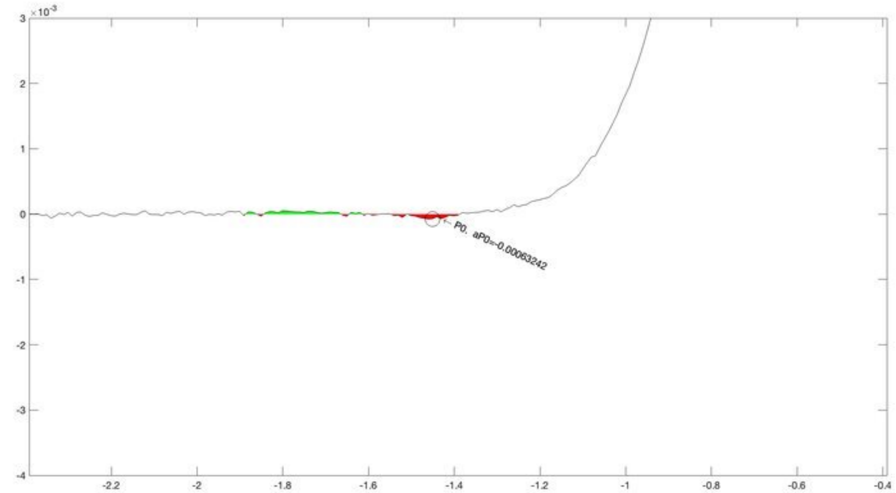
P0, aP0=-0.00063242

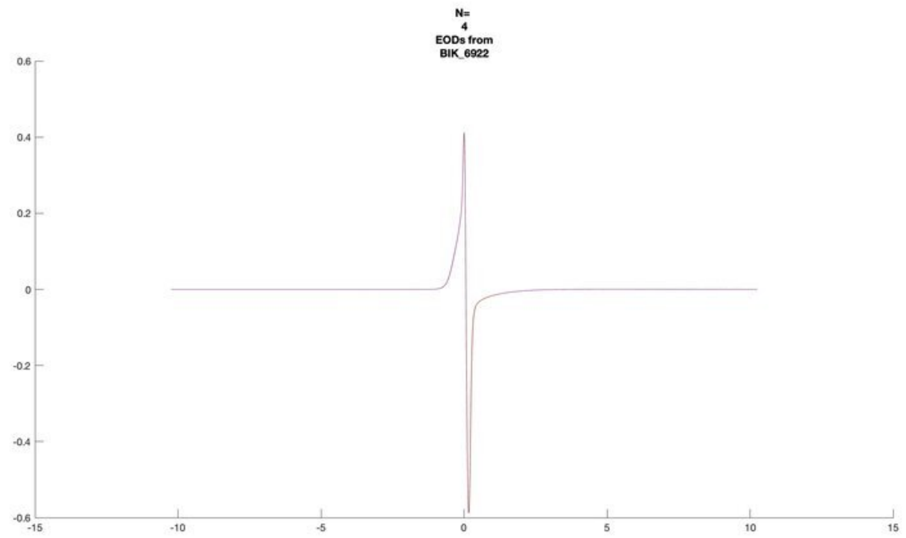
N=
4
EODs from
BIK_6922

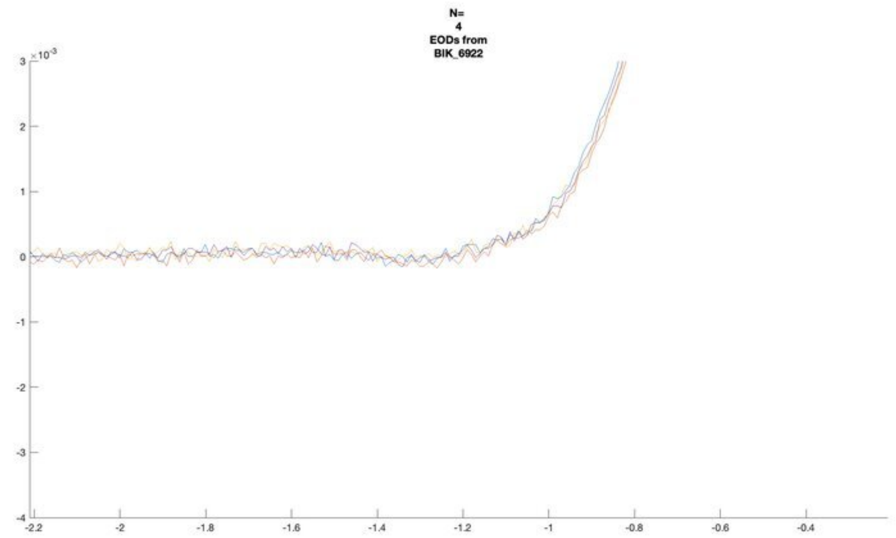
N=
4
EODs from
BIK_6922
×10^-3

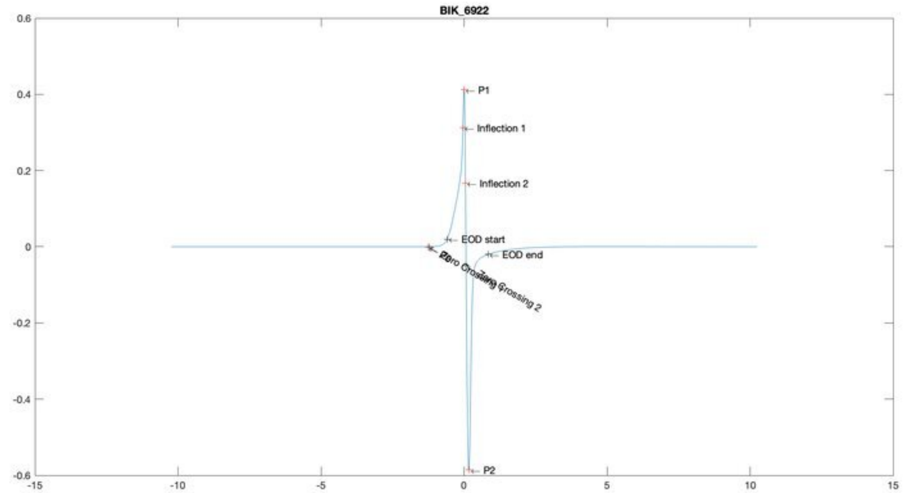
BIK_6922
P1
Inflection 1
Inflection 2
EOD start
EOD end
Zero Crossing 2
P2

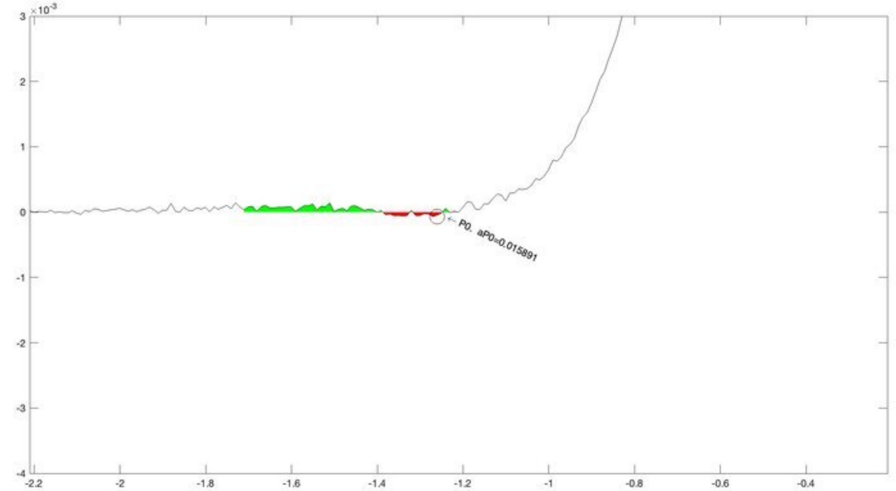
×10^-3
P0. aP0=0.015891

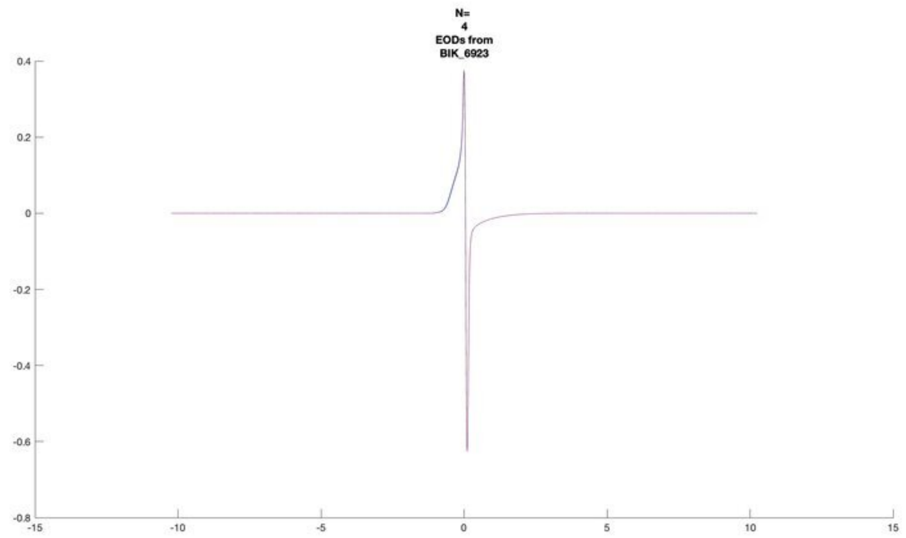
N=
4
EODs from
BIK_6923

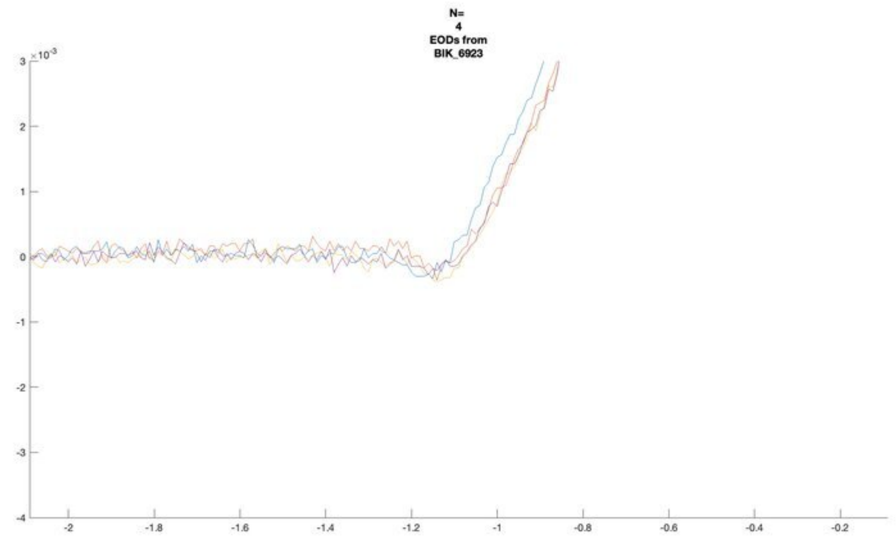
N=
4
EODs from
BIK_6923

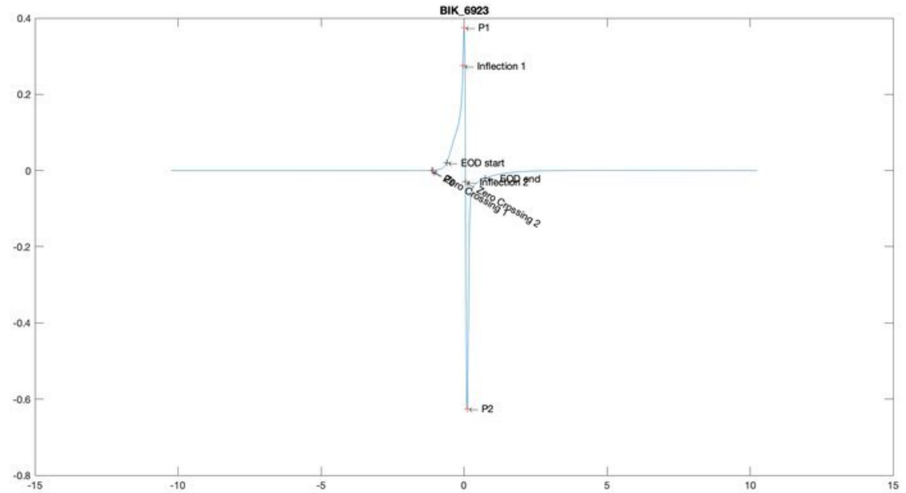
BIK_6923
P1
Inflection 1
EOD start
EOD end
Inflection 2
Zero Crossing 1
Zero Crossing 2
P2

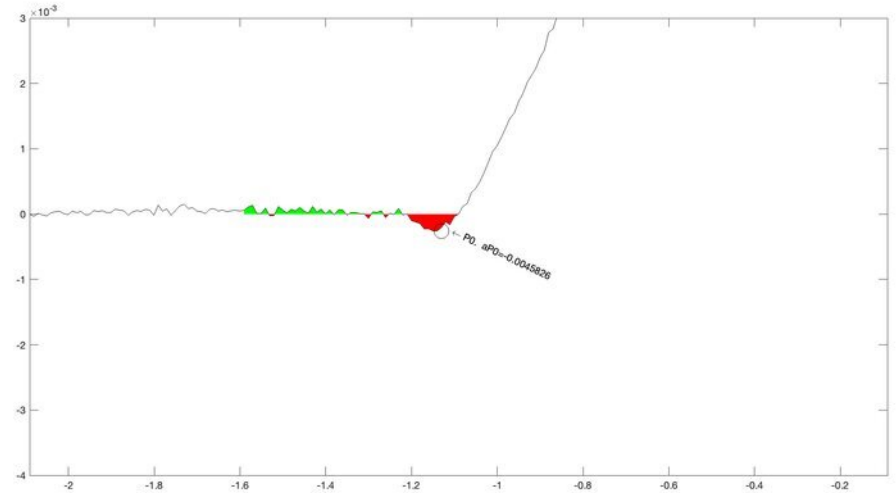
P0. aP0=-0.0045826

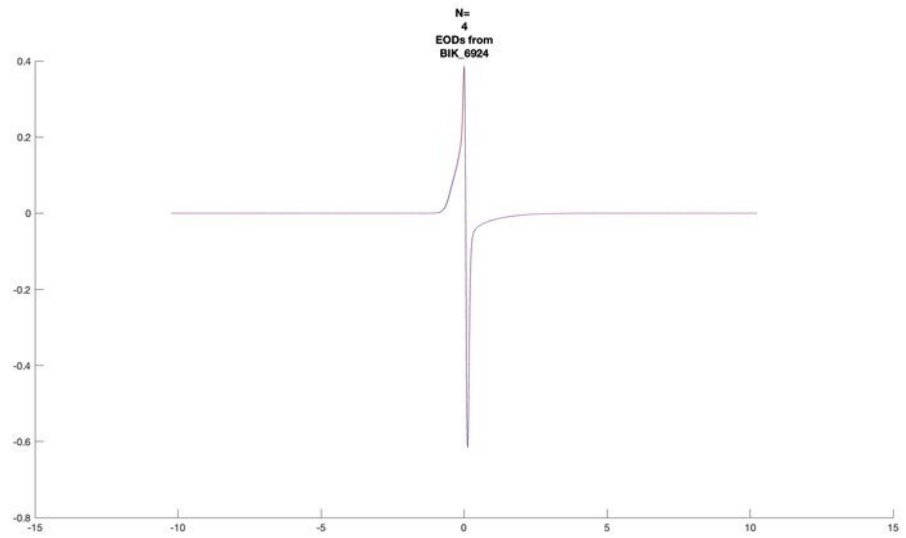
N=
4
EODs from
BIK_6924

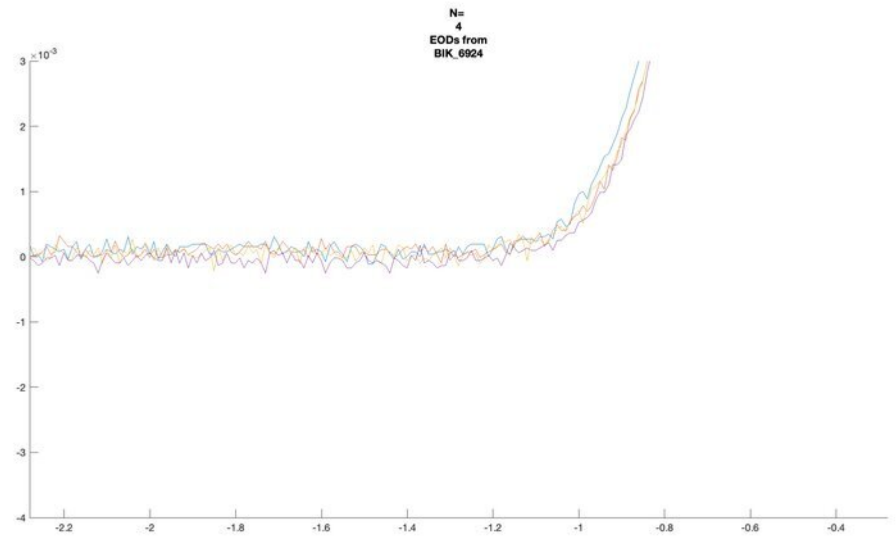
N=
4
EODs from
BIK_6924

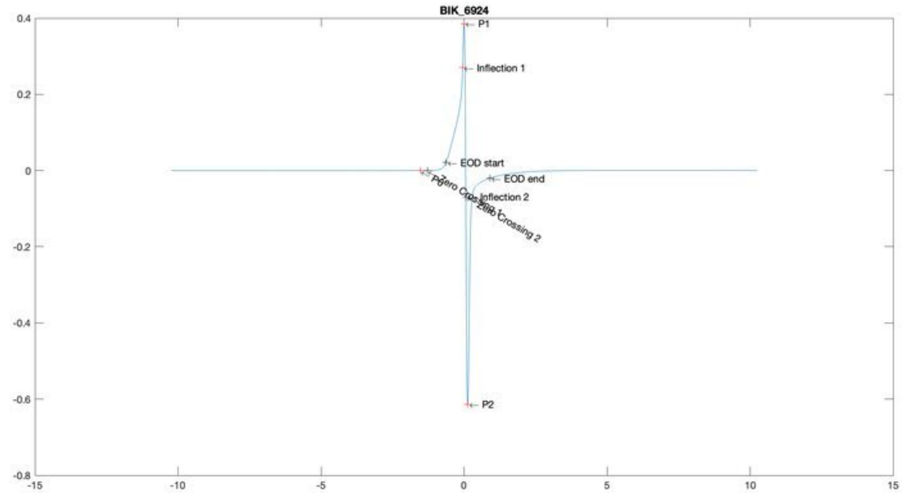
BIK_6924
P1
Inflection 1
EOD start
EOD end
Inflection 2
Zero Crossing 2
P2

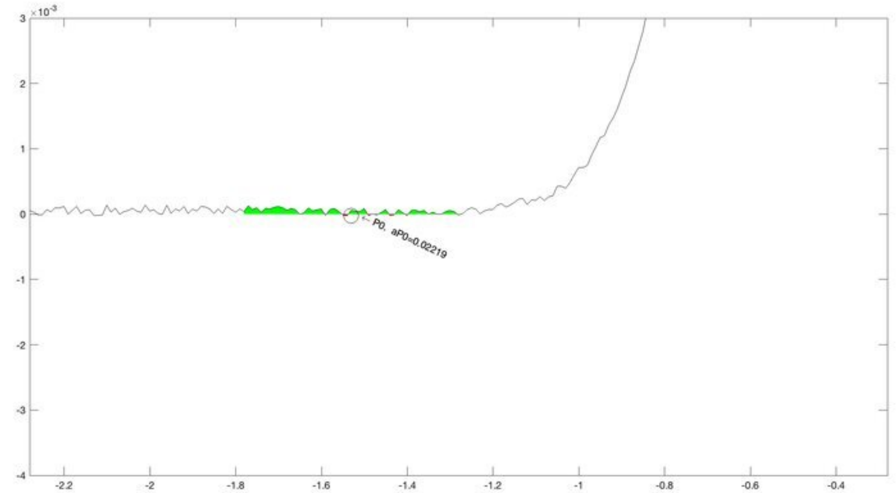
P0, aP0=0.02219

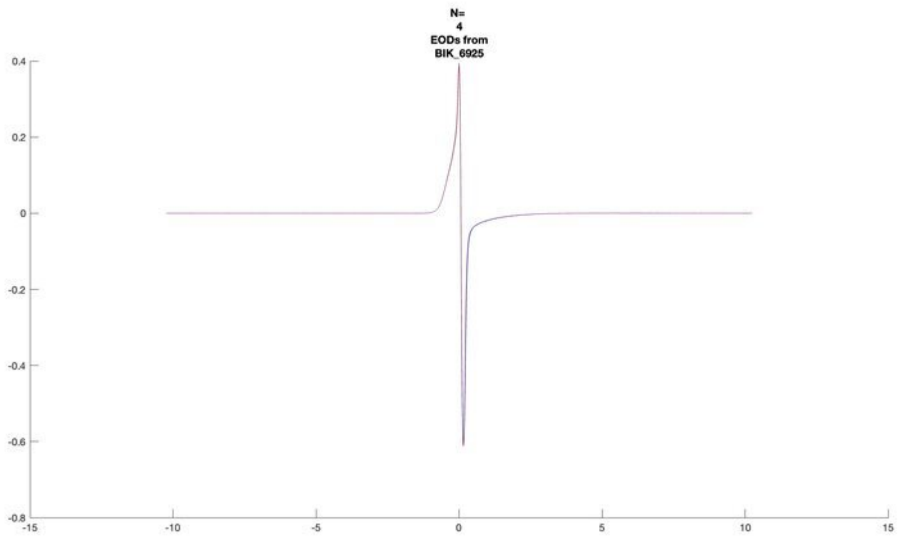
N=
4
EODs from
BIK_6925

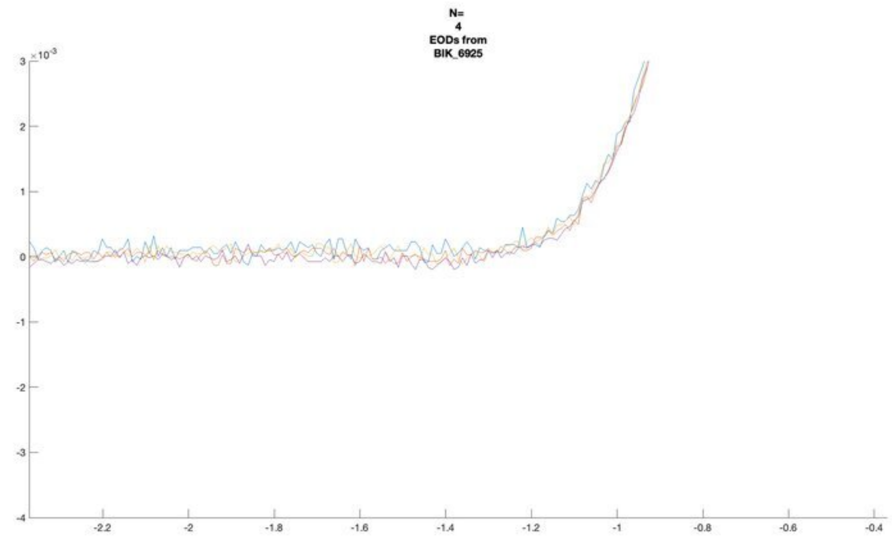
N=
4
EODs from
BIK_6925

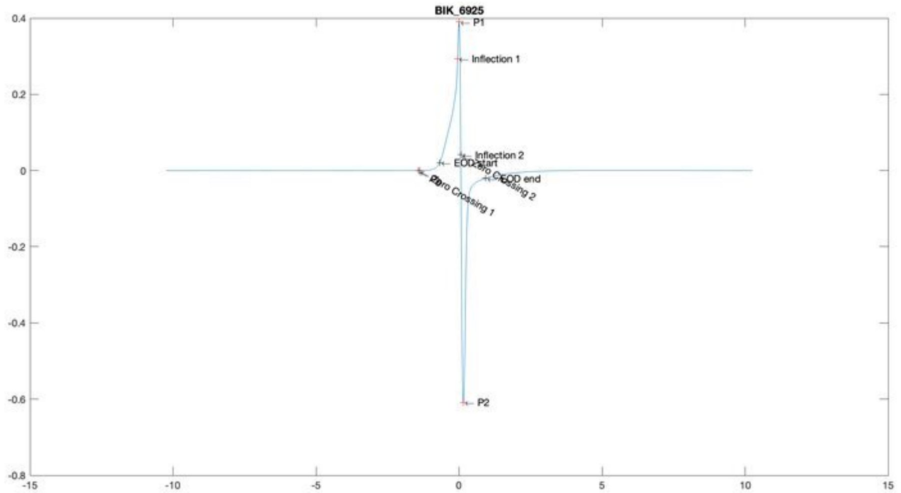
BIK_6925
P1
Inflection 1
Inflection 2
EOD start
EOD end
Zero Crossing 1
Zero Crossing 2
P2

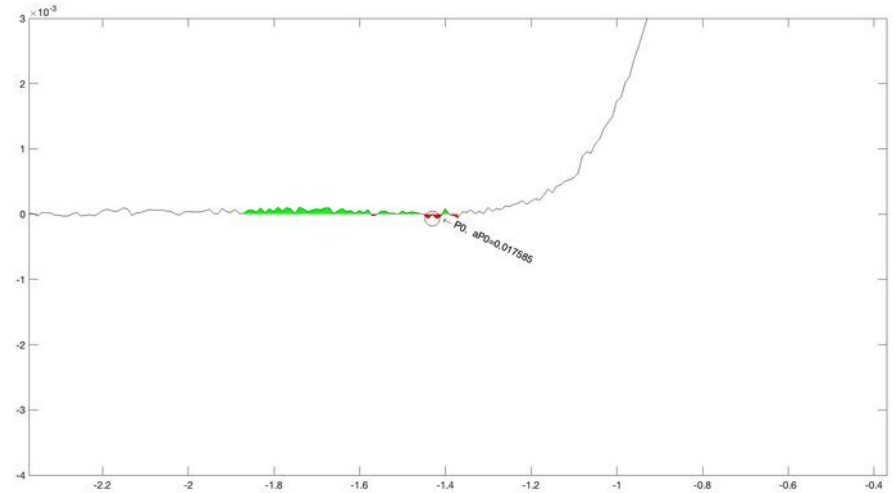
P0, aP0=0.017585

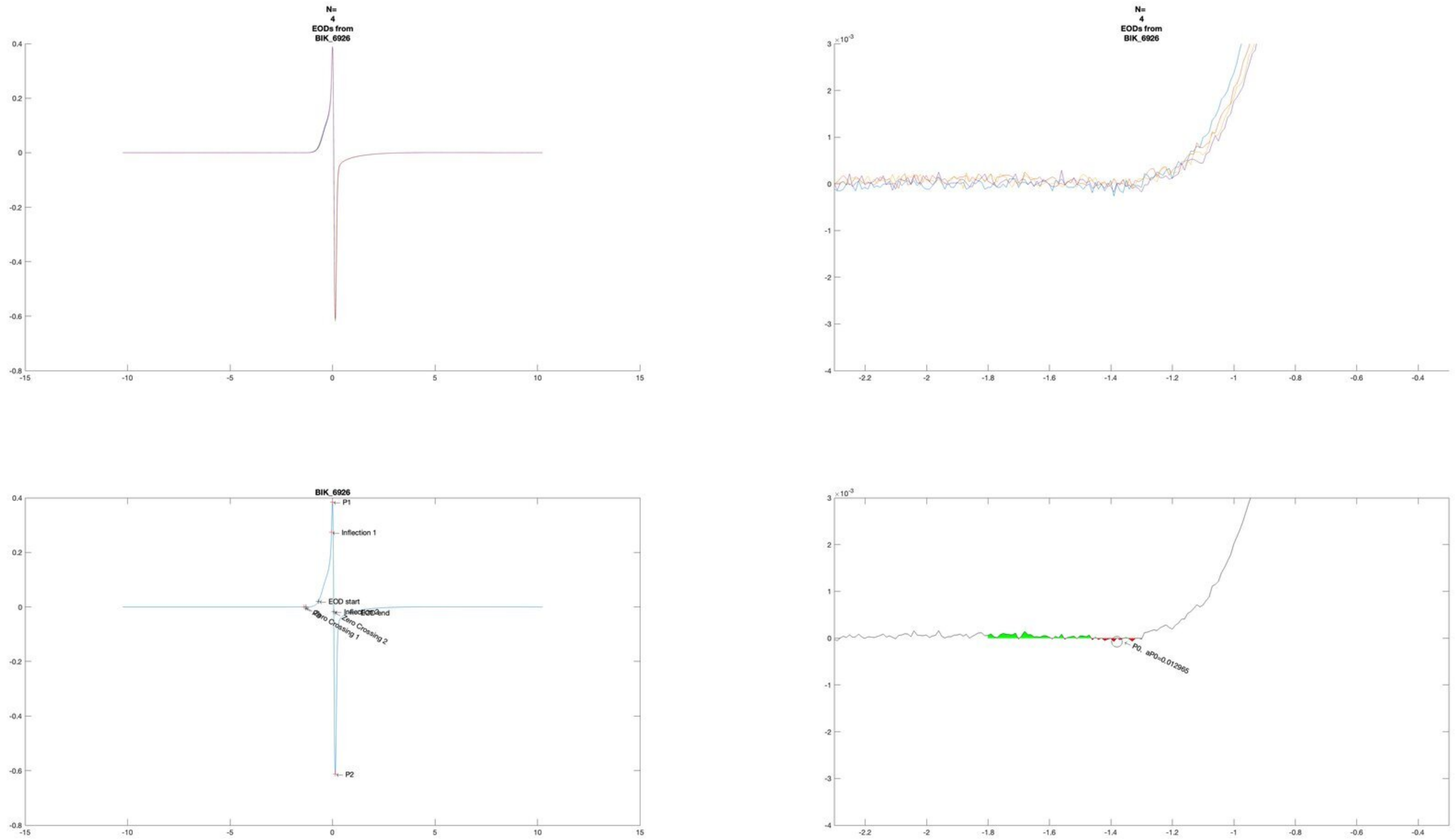
N=
4
EODs from
BIK_6926
N=
4
EODs from
BIK_6926
BIK_6926
P1
Inflection 1
EOD start
EOD end
Zero Crossing 1
Zero Crossing 2
P2
P0, aP0=0.012965

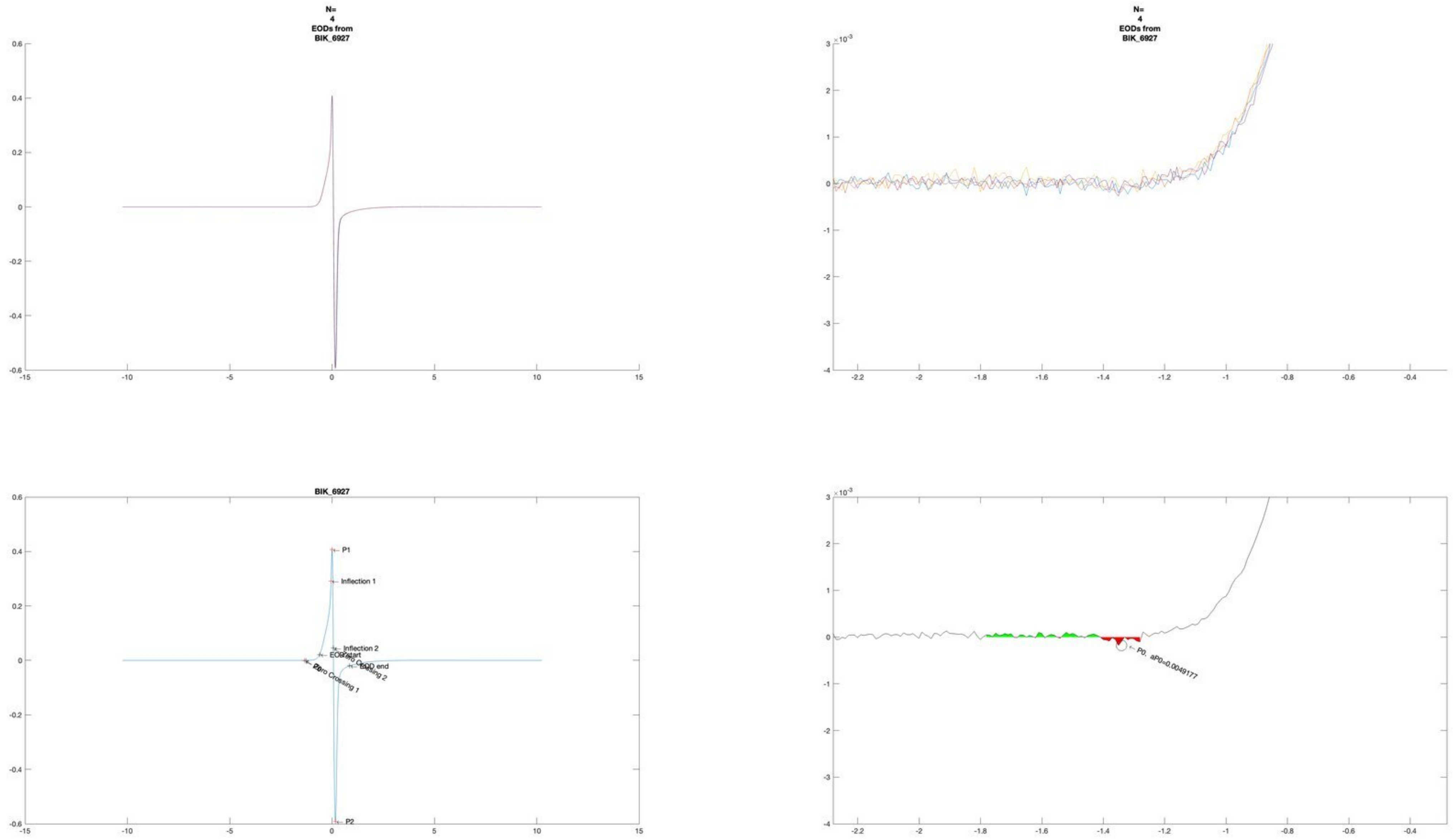
N=
4
EODs from
BIK_6927
N=
4
EODs from
BIK_6927
BIK_6927
P1
Inflection 1
Inflection 2
EOD start
EOD end
Zero Crossing 1
P2
P0. aP0=0.0049177

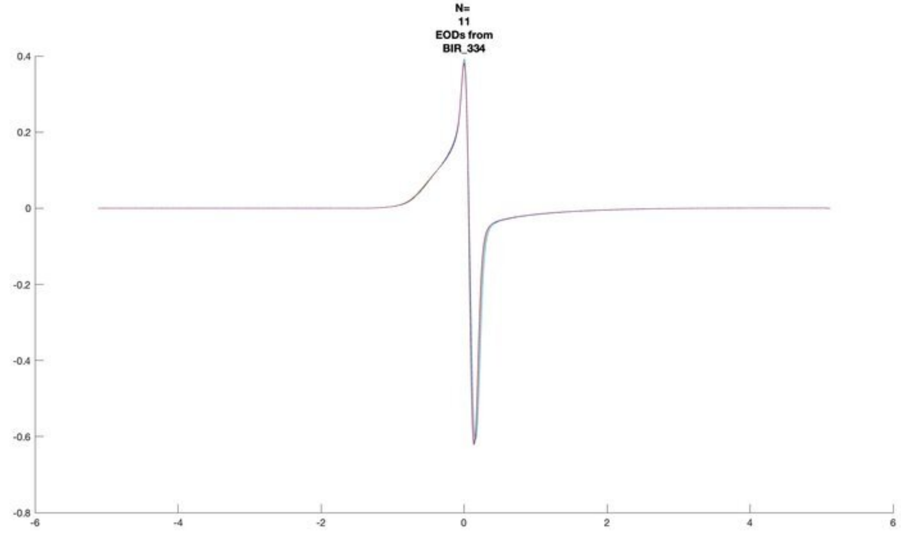
N=
11
EODs from
BIR_334

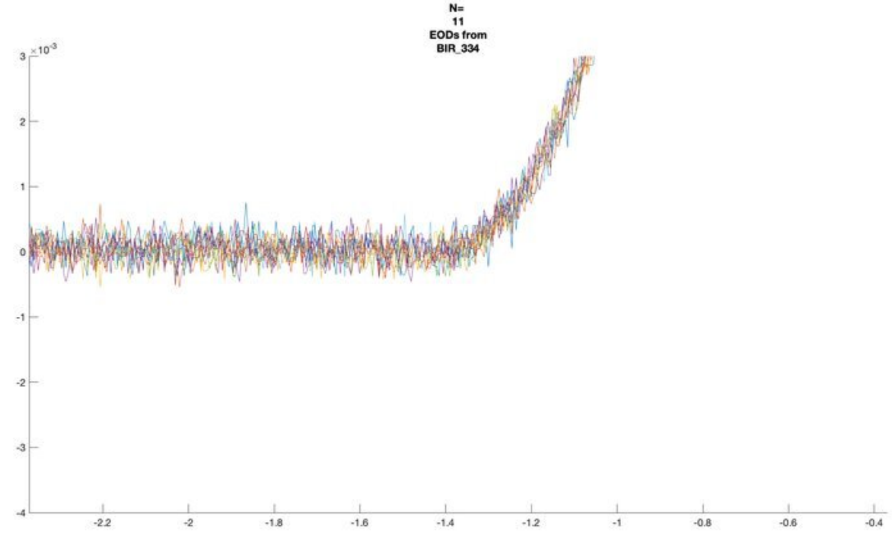
N=
11
EODs from
BIR_334

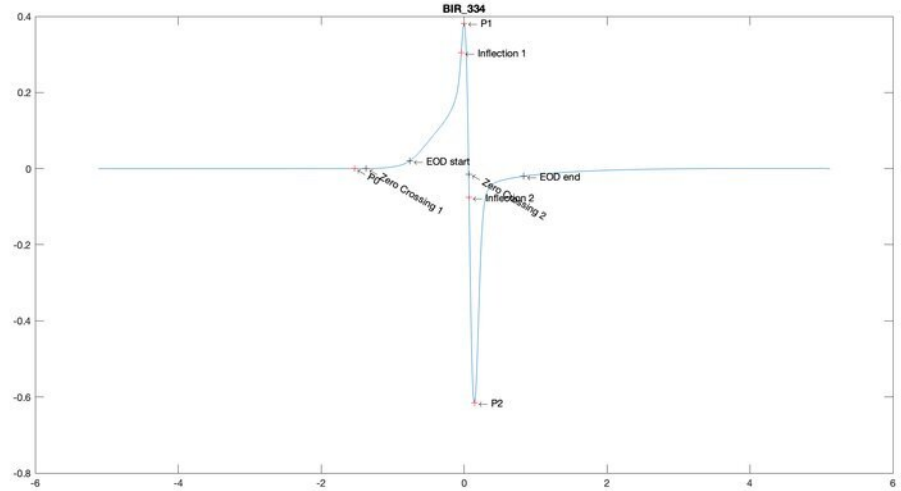
BIR_334
P1
Inflection 1
EOD start
EOD end
Zero Crossing 1
P0
Zero Crossing 2
Inflection 2
P2

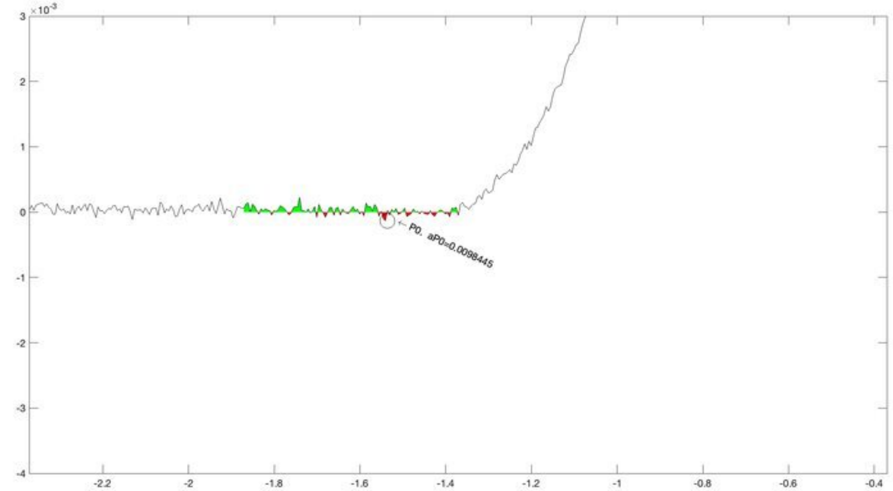
P0, aP0=0.0098445

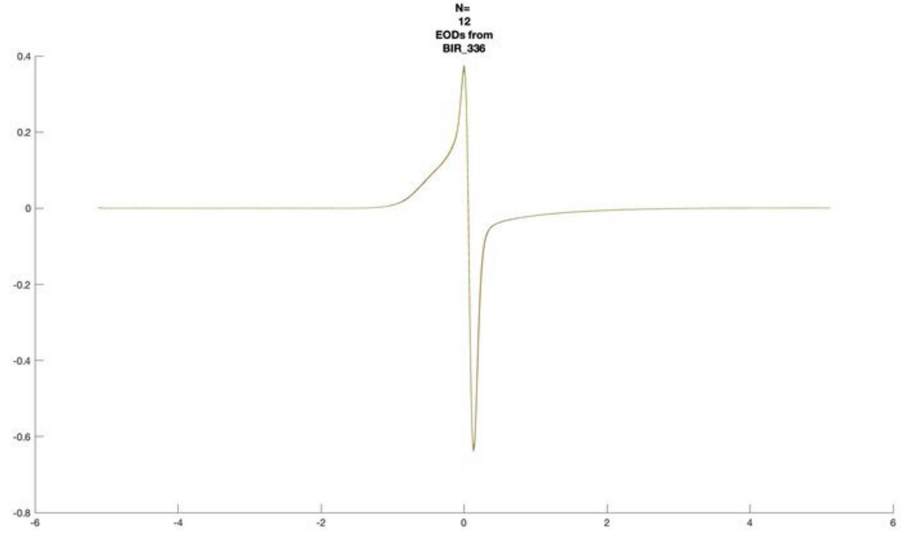
N=
12
EODs from
BIR_336
0.4
0.2
0
-0.2
-0.4
-0.6
-0.8
-6
-4
-2
0
2
4
6

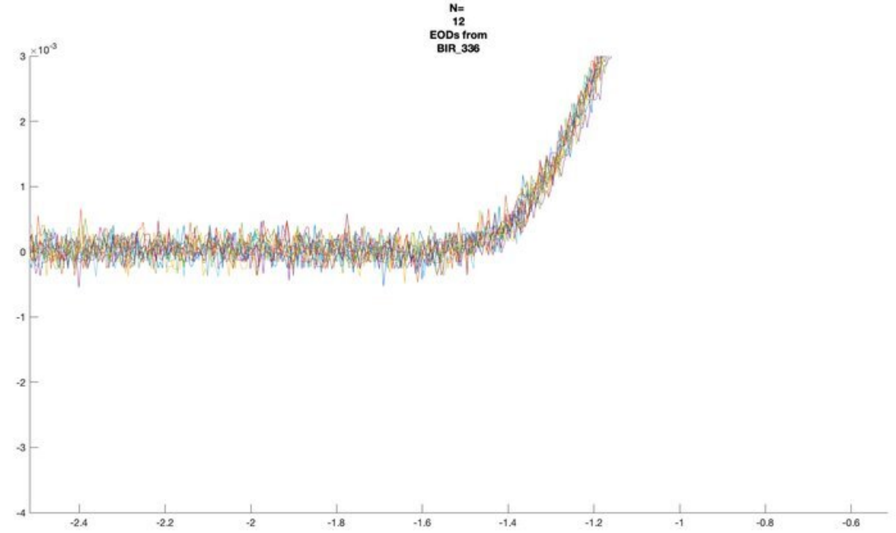
N=
12
EODs from
BIR_336
×10^-3
3
2
1
0
-1
-2
-3
-4
-2.4
-2.2
-2
-1.8
-1.6
-1.4
-1.2
-1
-0.8
-0.6

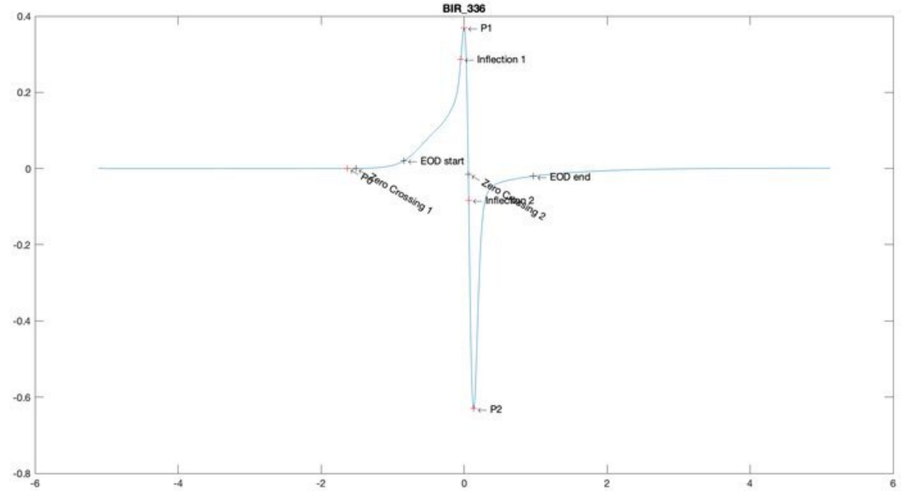
BIR_336
P1
Inflection 1
EOD start
EOD end
Zero Crossing 1
P0
Zero Crossing 2
Inflection 2
P2
0.4
0.2
0
-0.2
-0.4
-0.6
-0.8
-6
-4
-2
0
2
4
6

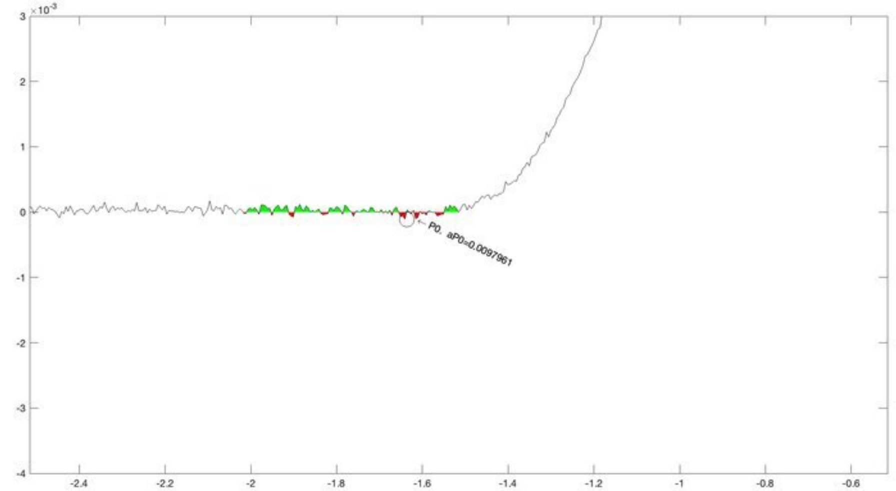
×10^-3
P0. aP0=0.0097961
3
2
1
0
-1
-2
-3
-4
-2.4
-2.2
-2
-1.8
-1.6
-1.4
-1.2
-1
-0.8
-0.6

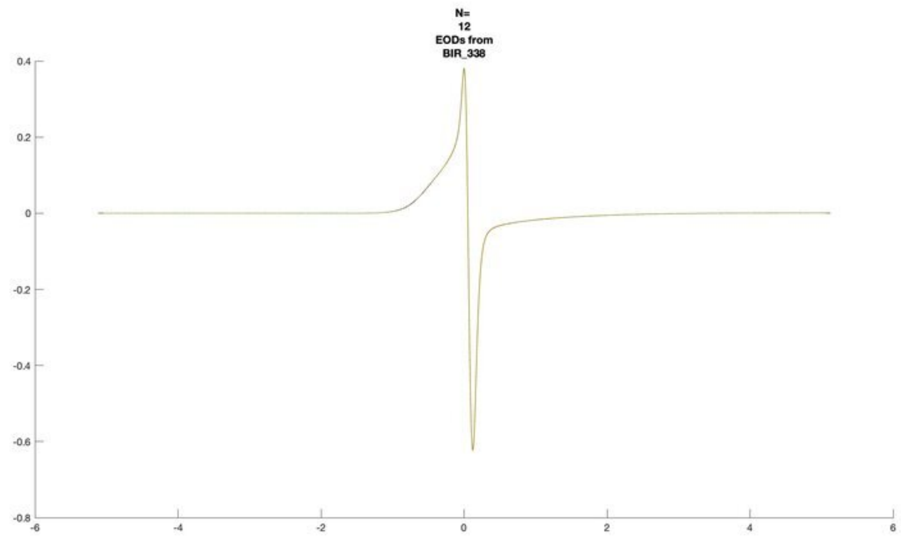
N=
12
EODs from
BIR_338

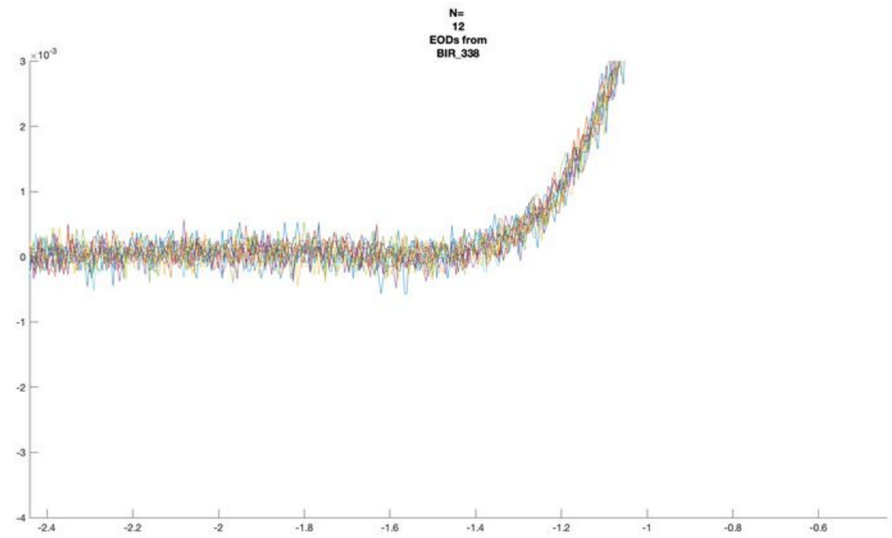
N=
12
EODs from
BIR_338
×10^-3

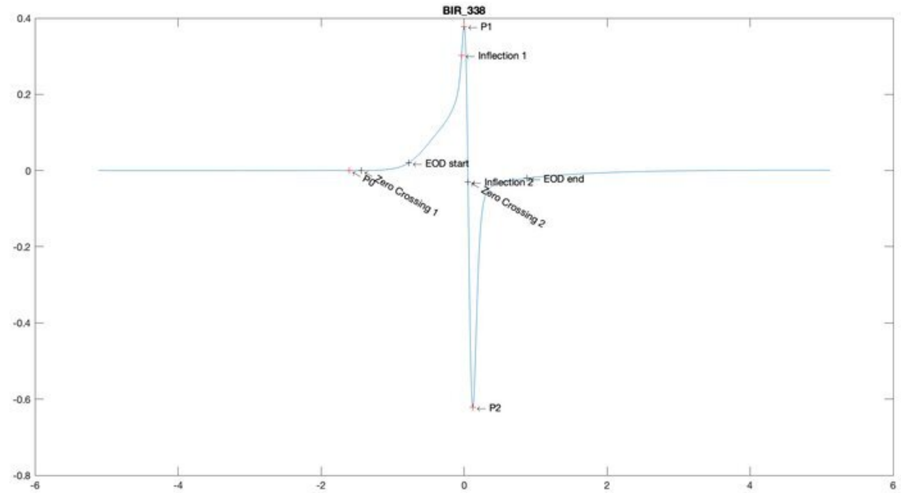
BIR_338
P1
Inflection 1
EOD start
P0
Zero Crossing 1
Inflection 2
Zero Crossing 2
EOD end
P2

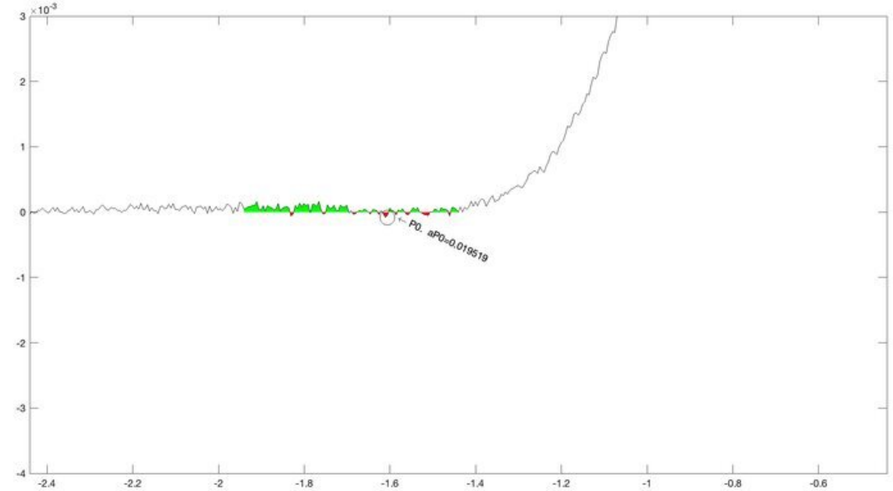
×10^-3
P0, aP0=0.019519

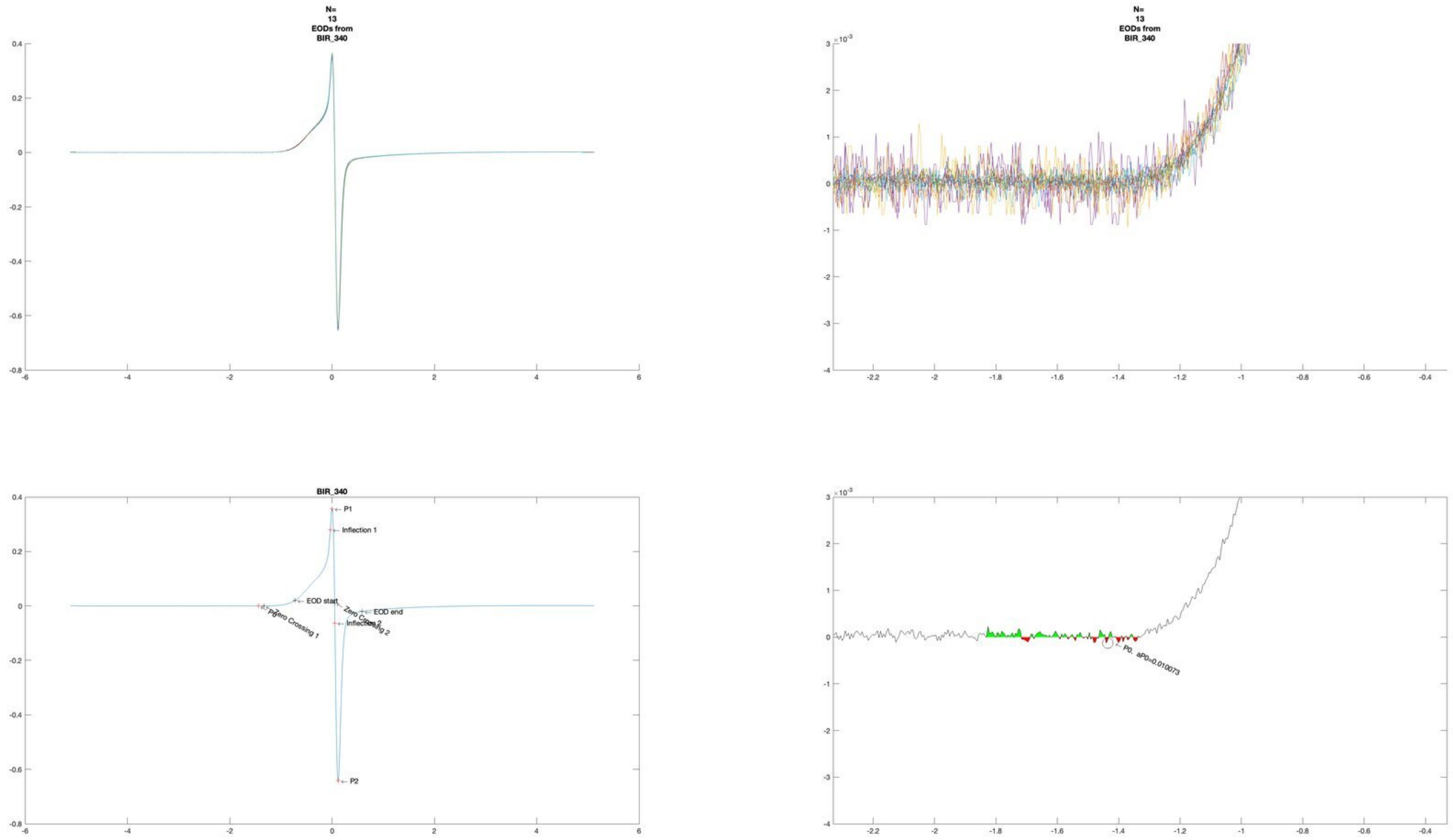
N=
13
EODs from
BIR_340
N=
13
EODs from
BIR_340
BIR_340
P1
Inflection 1
EOD start
EOD end
Zero Crossing 1
P0
P2
P0, aP0=0.010073

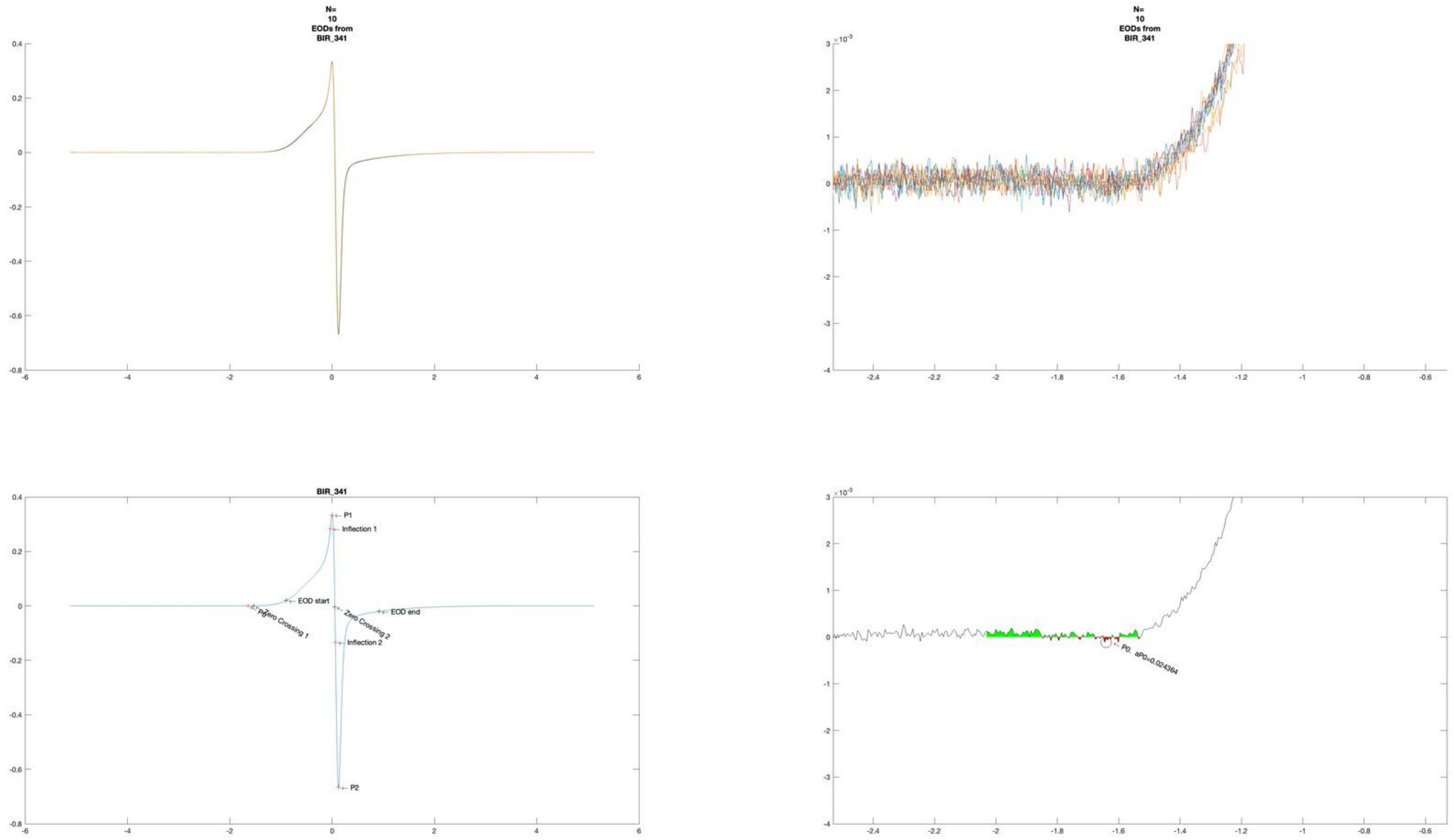
N=
10
EODs from
BIR_341
N=
10
EODs from
BIR_341
BIR_341
P1
Inflection 1
EOD start
P0
Zero Crossing 1
Zero Crossing 2
EOD end
Inflection 2
P2
P0. aP0=0.024364

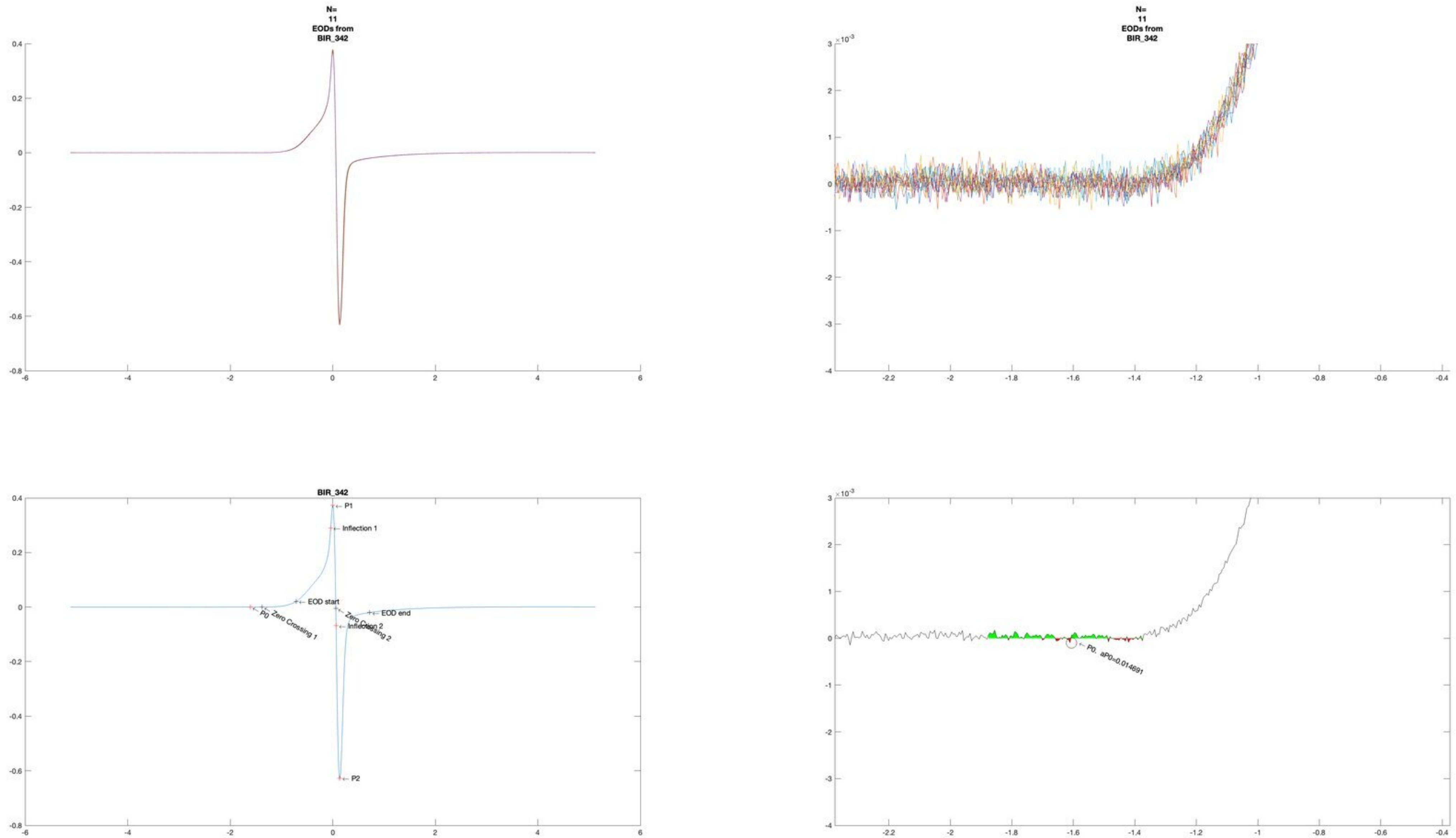

N=
11
EODs from
BIR_342
N=
11
EODs from
BIR_342
×10^-3
BIR_342
P1
Inflection 1
EOD start
P0
Zero Crossing 1
Zero Crossing 2
Inflection 2
EOD end
P2
×10^-3
P0, aP0=0.014691

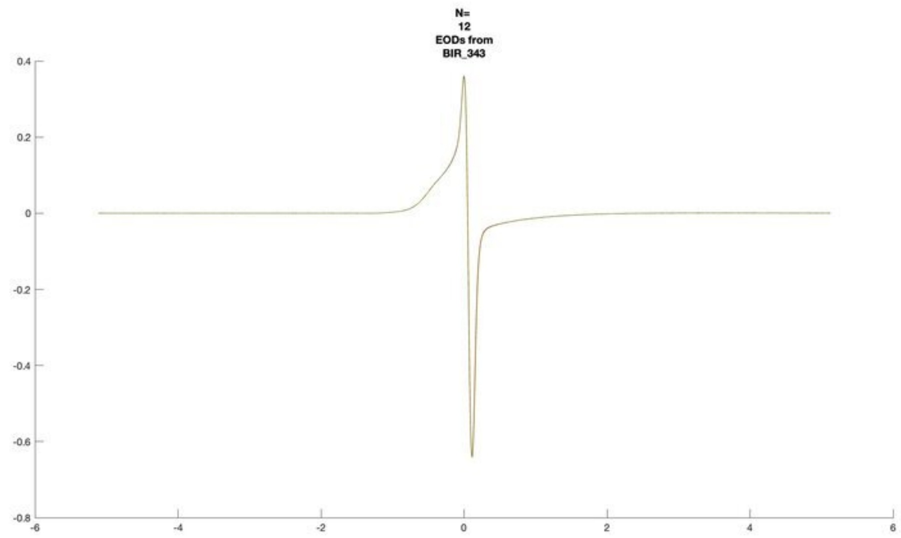
N=
12
EODs from
BIR_343

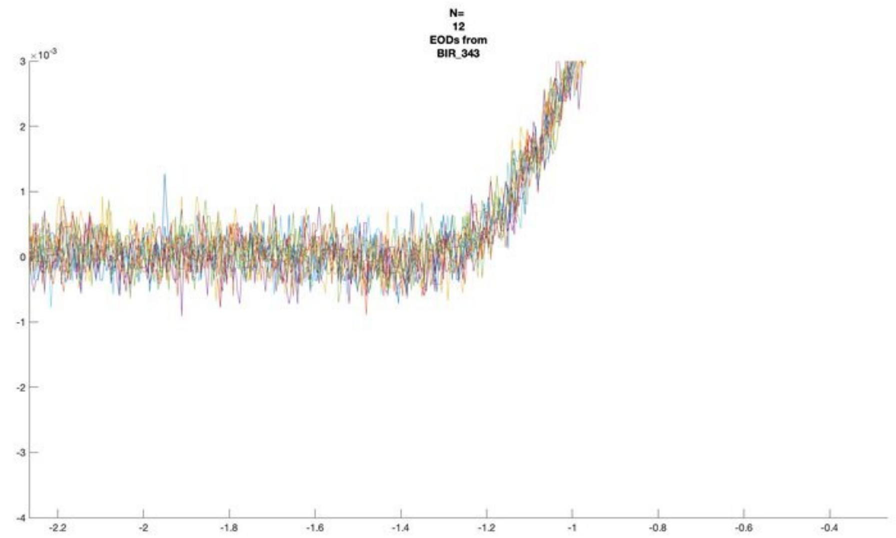
N=
12
EODs from
BIR_343

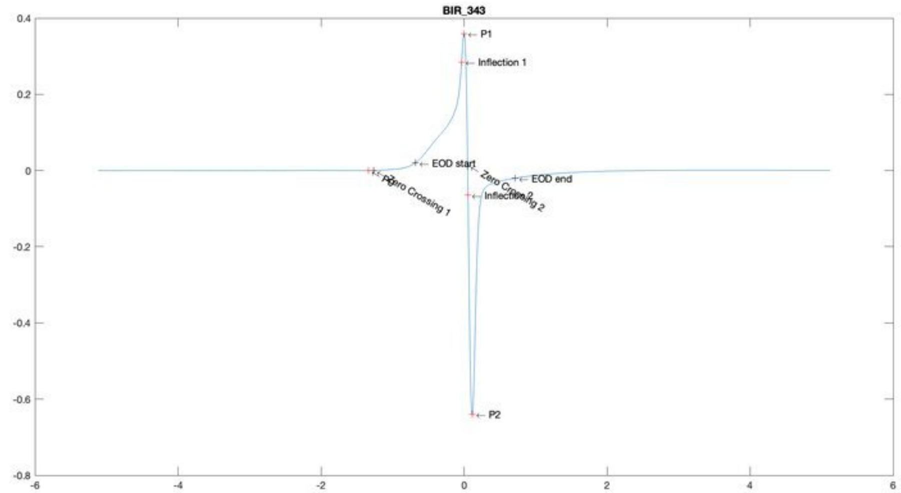
BIR_343
P1
Inflection 1
EOD start
EOD end
Zero Crossing 2
Inflection 2
Zero Crossing 1
P2

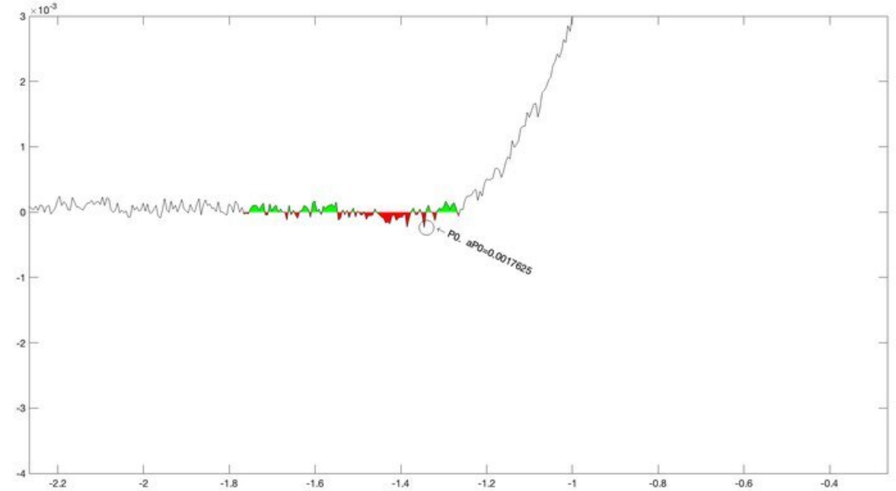
P0, aP0=0.0017625

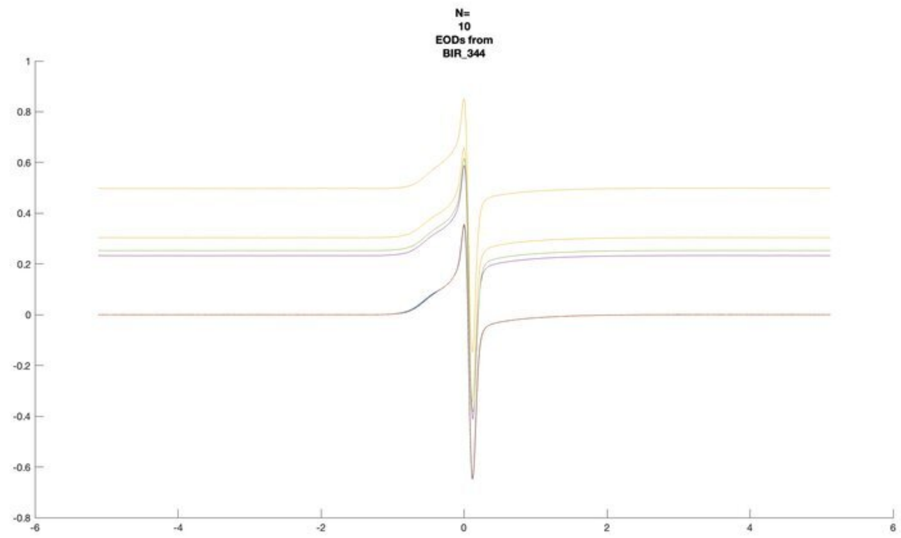
N=
10
EODs from
BIR_344

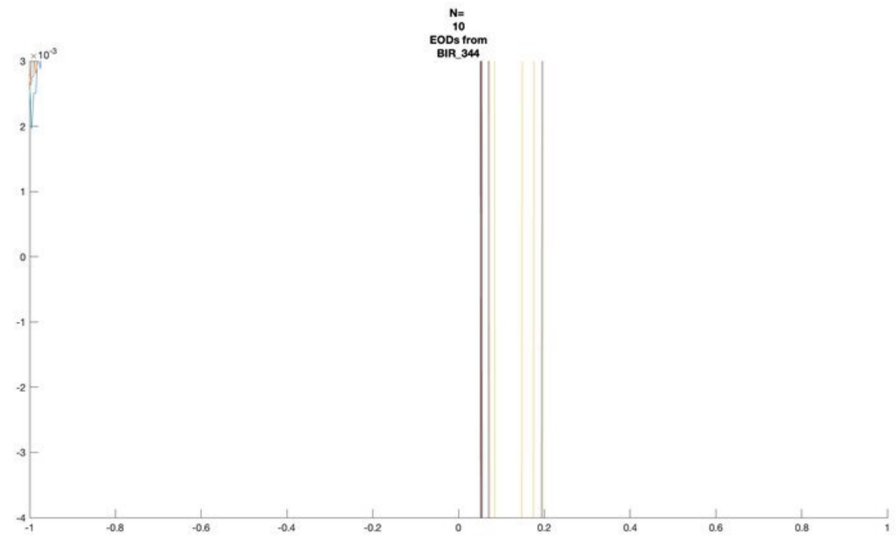
N=
10
EODs from
BIR_344

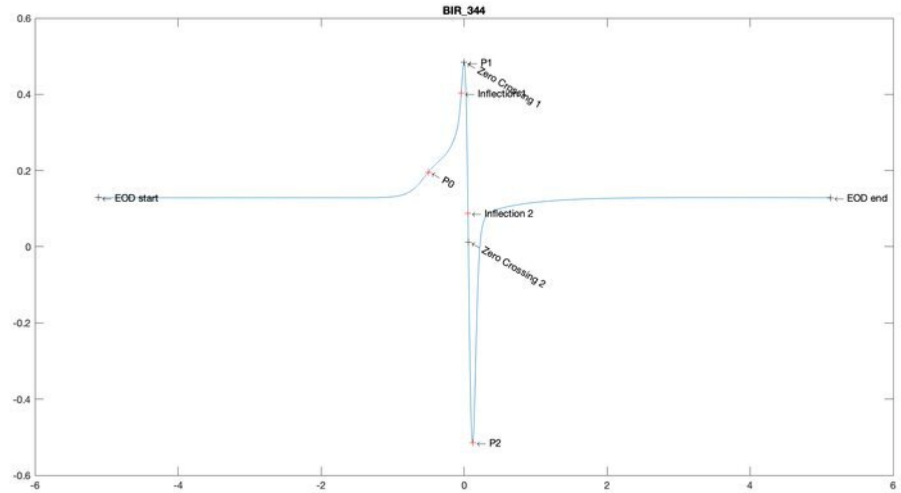
BIR_344
P1
Zero Crossing 1
Inflection 1
P0
EOD start
EOD end
Inflection 2
Zero Crossing 2
P2

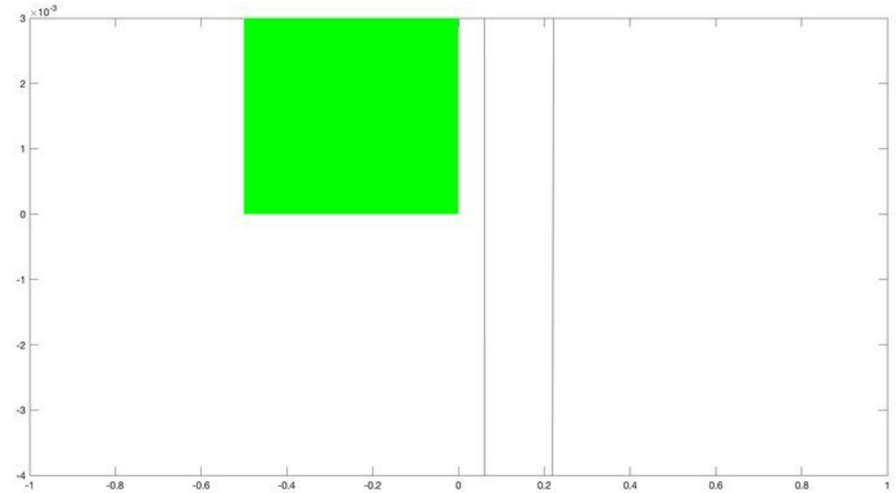

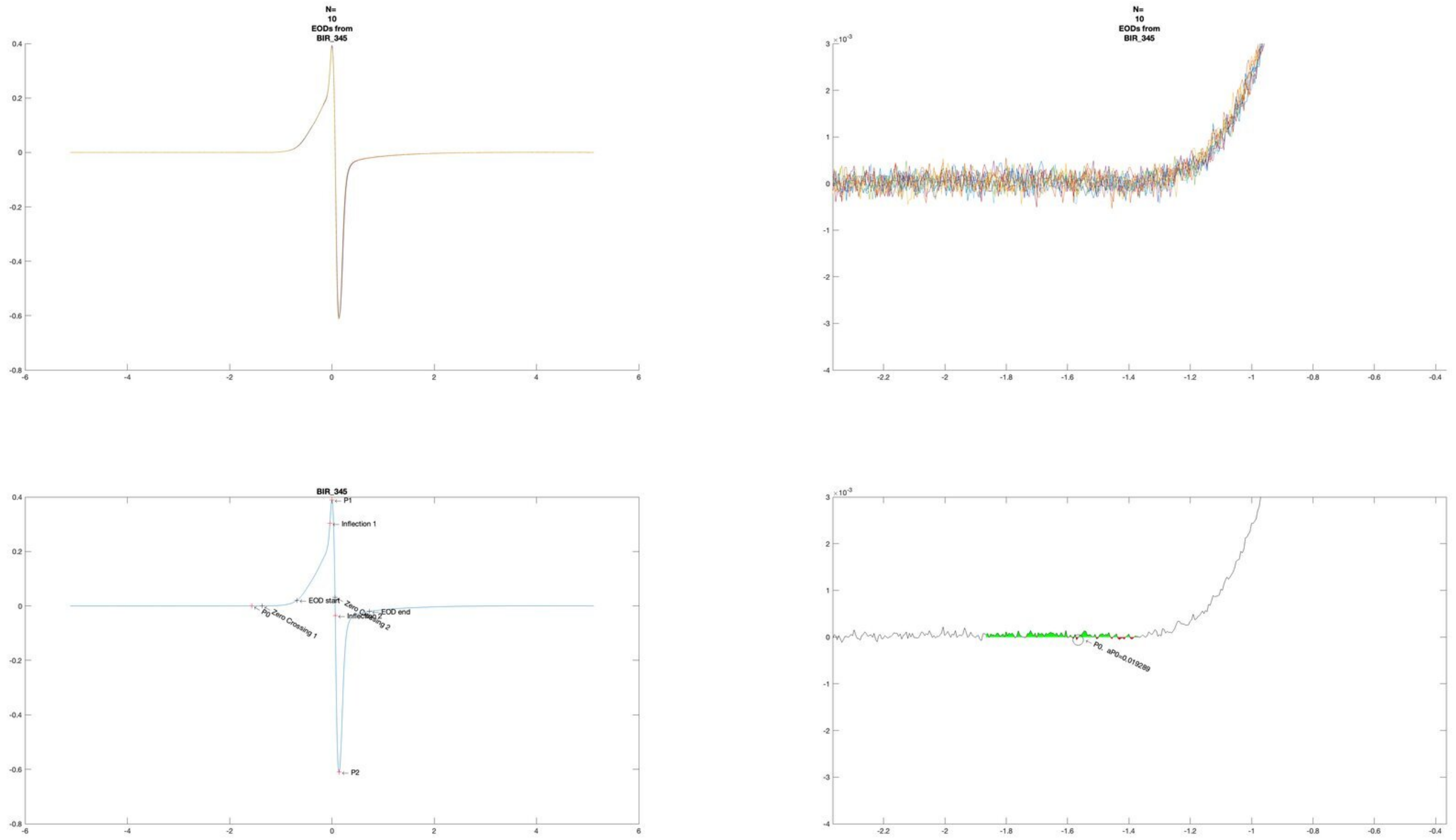
N=
10
EODs from
BIR_345
N=
10
EODs from
BIR_345
BIR_345
P1
Inflection 1
EOD start
P0
Zero Crossing 1
Inflection 2
Zero Crossing 2
EOD end
P2
P0, aP0=0.019289

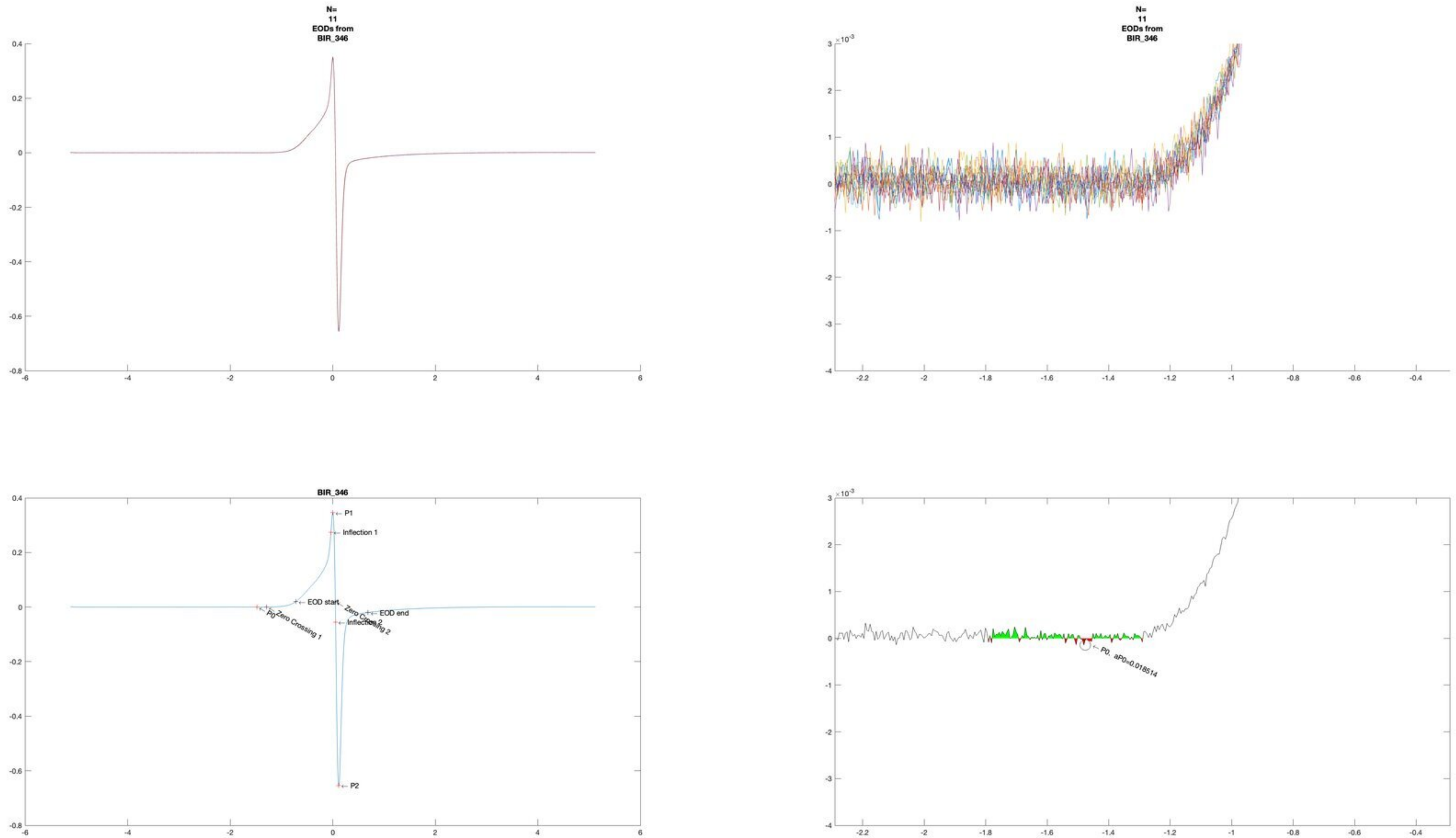
N=
11
EODs from
BIR_346
N=
11
EODs from
BIR_346
BIR_346
P1
Inflection 1
EOD start
Zero Crossing 2
EOD end
P0
Zero Crossing 1
Inflection 2
P2
P0. aP0=0.018514

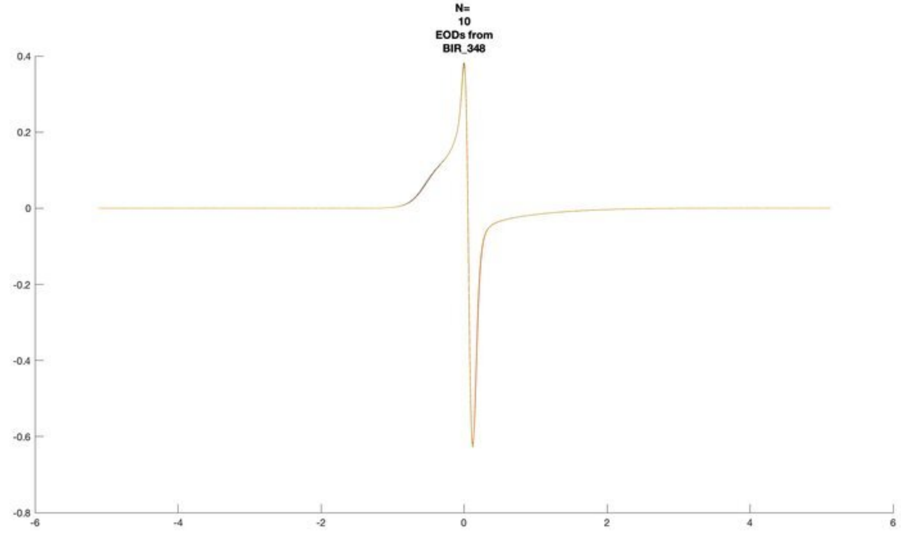
N=
10
EODs from
BIR_348

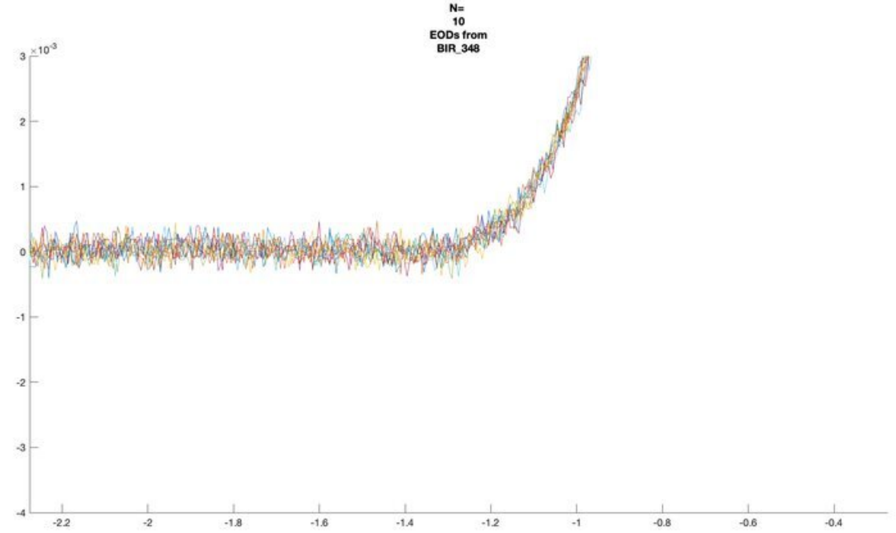
N=
10
EODs from
BIR_348
×10^-3

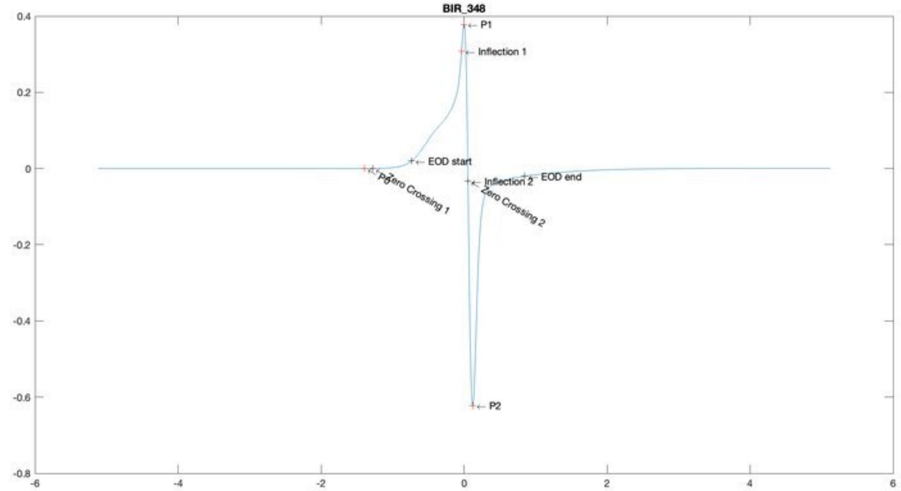
BIR_348
P1
Inflection 1
EOD start
P0
Zero Crossing 1
Inflection 2
Zero Crossing 2
EOD end
P2

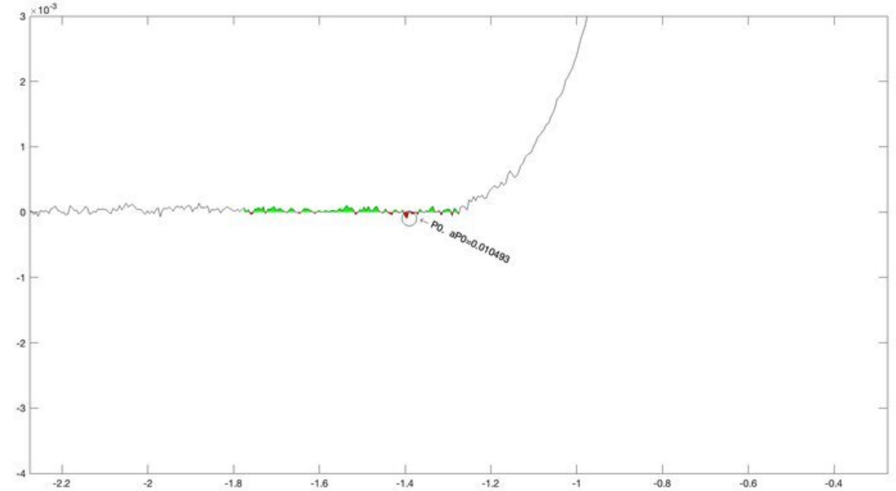
×10^-3
P0. aP0=0.010493

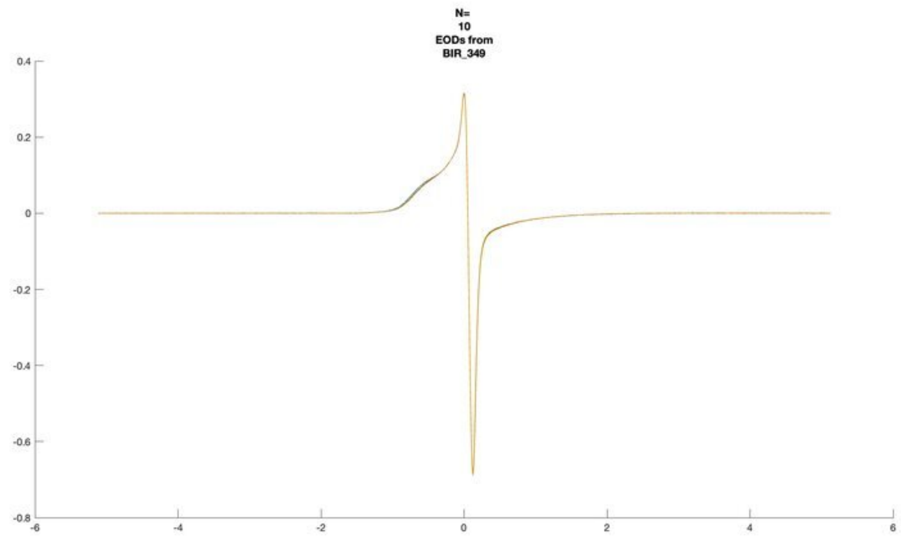
N=
10
EODs from
BIR_349
0.4
0.2
0
-0.2
-0.4
-0.6
-0.8
-6
-4
-2
0
2
4
6

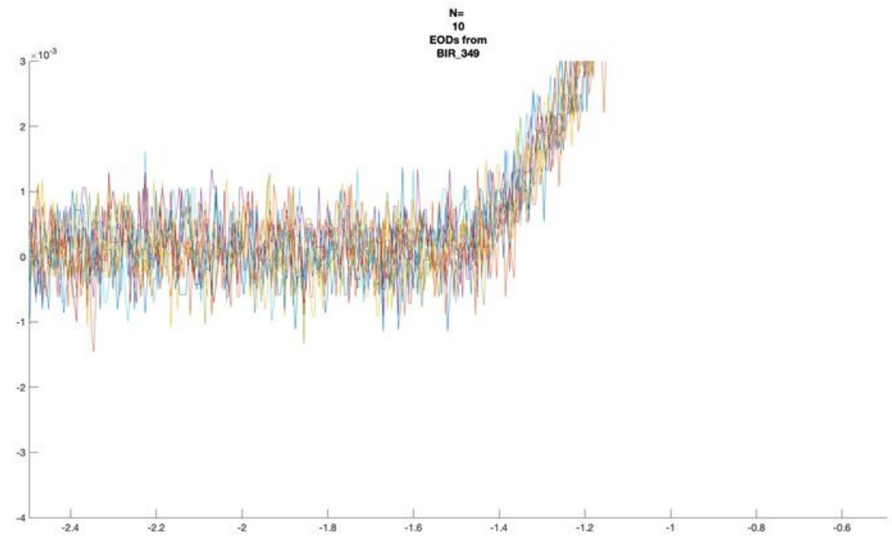
N=
10
EODs from
BIR_349
×10^-3
3
2
1
0
-1
-2
-3
-4
-2.4
-2.2
-2
-1.8
-1.6
-1.4
-1.2
-1
-0.8
-0.6

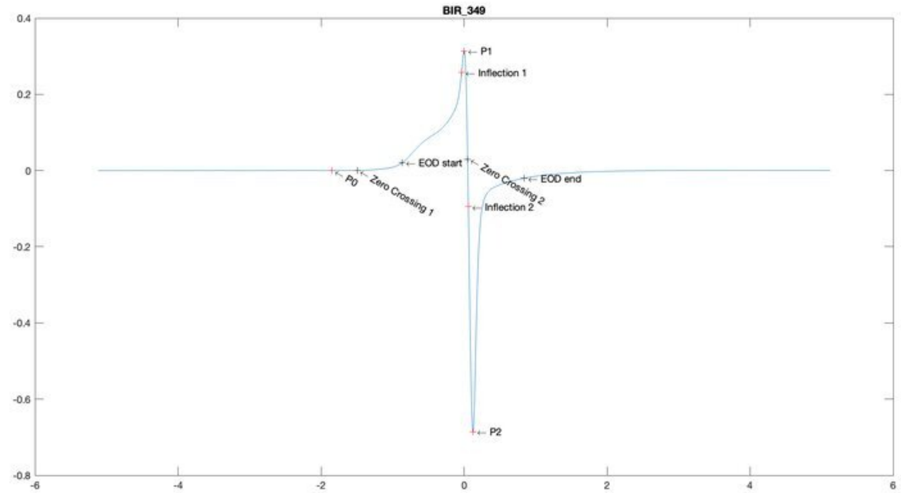
BIR_349
P1
Inflection 1
EOD start
Zero Crossing 2
EOD end
P0
Zero Crossing 1
Inflection 2
P2
0.4
0.2
0
-0.2
-0.4
-0.6
-0.8
-6
-4
-2
0
2
4
6

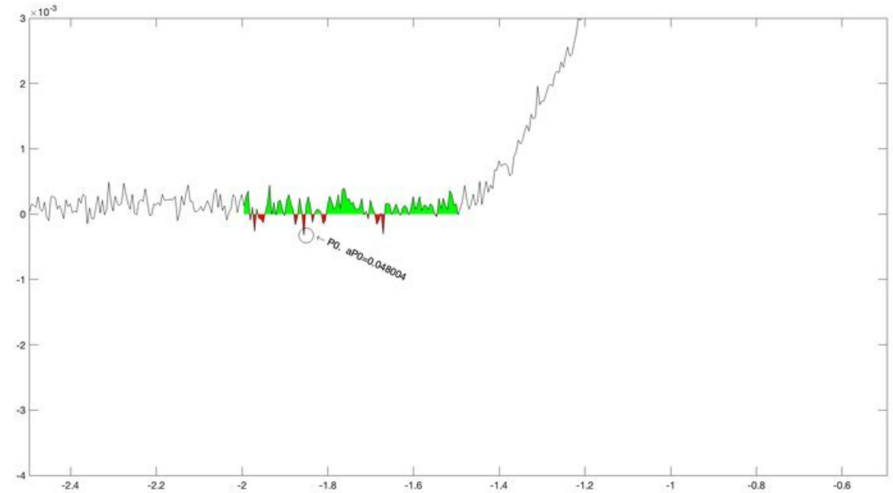
×10^-3
P0, aP0=0.048004
3
2
1
0
-1
-2
-3
-4
-2.4
-2.2
-2
-1.8
-1.6
-1.4
-1.2
-1
-0.8
-0.6

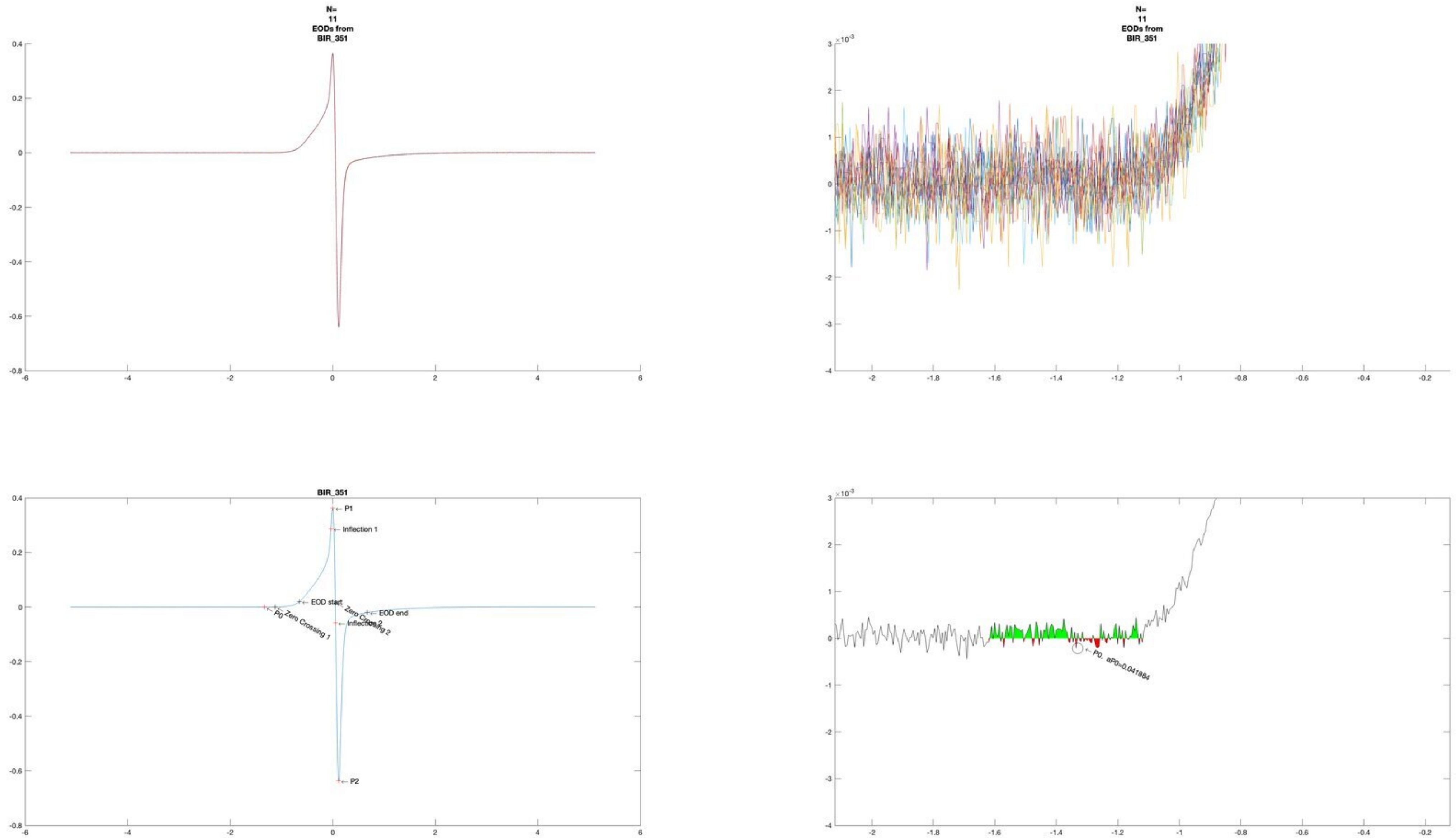
N=
11
EODs from
BIR_351
N=
11
EODs from
BIR_351
BIR_351
P1
Inflection 1
EOD start
Zero Crossing 1
P0
Zero Crossing 2
Inflection 2
EOD end
P2
P0, aP0=0.041884

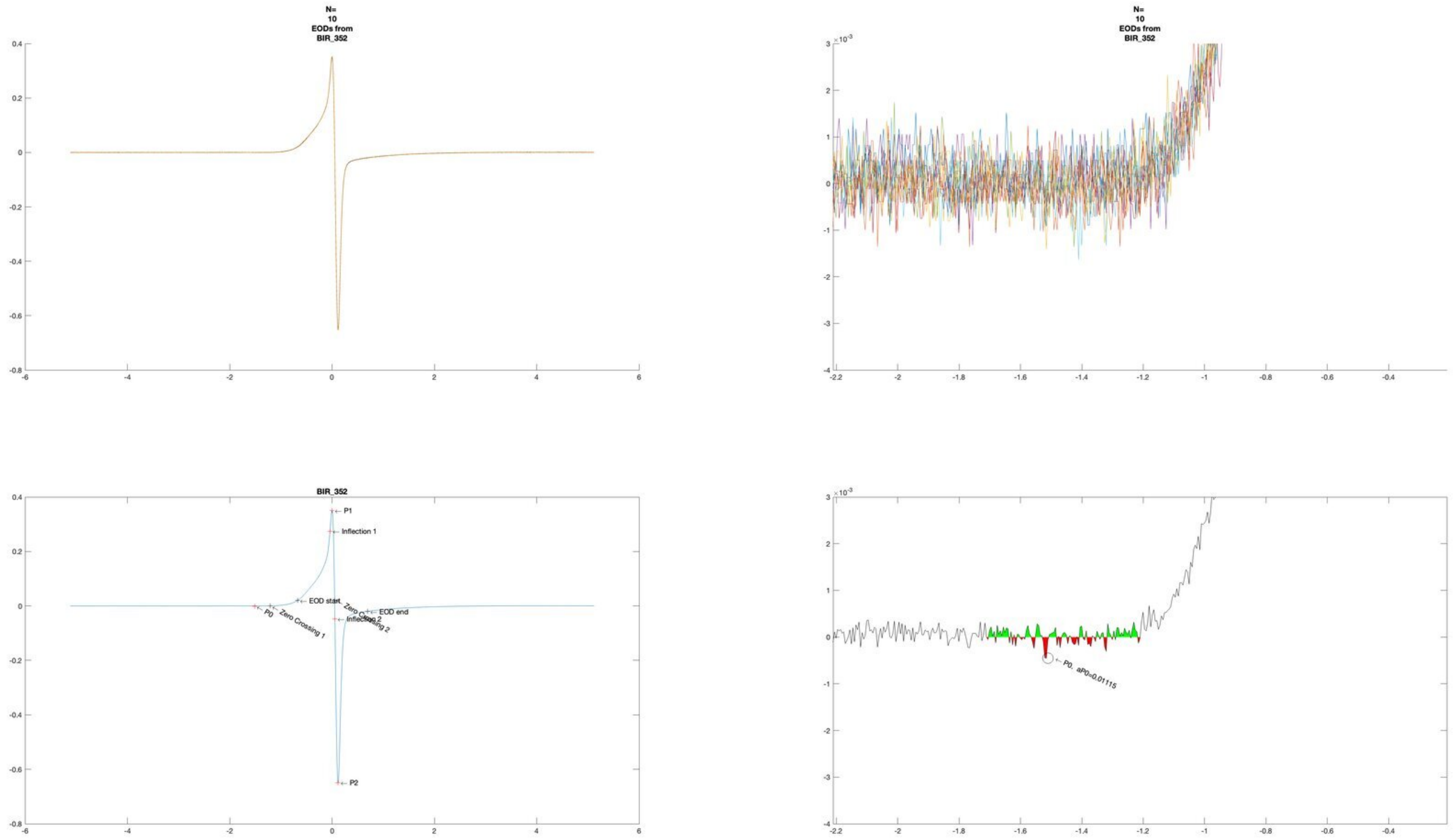
N=
10
EODs from
BIR_352
N=
10
EODs from
BIR_352
BIR_352
P1
Inflection 1
EOD start
Zero Crossing 2
Inflection 2
EOD end
P0
Zero Crossing 1
P2
P0, aP0=0.01115

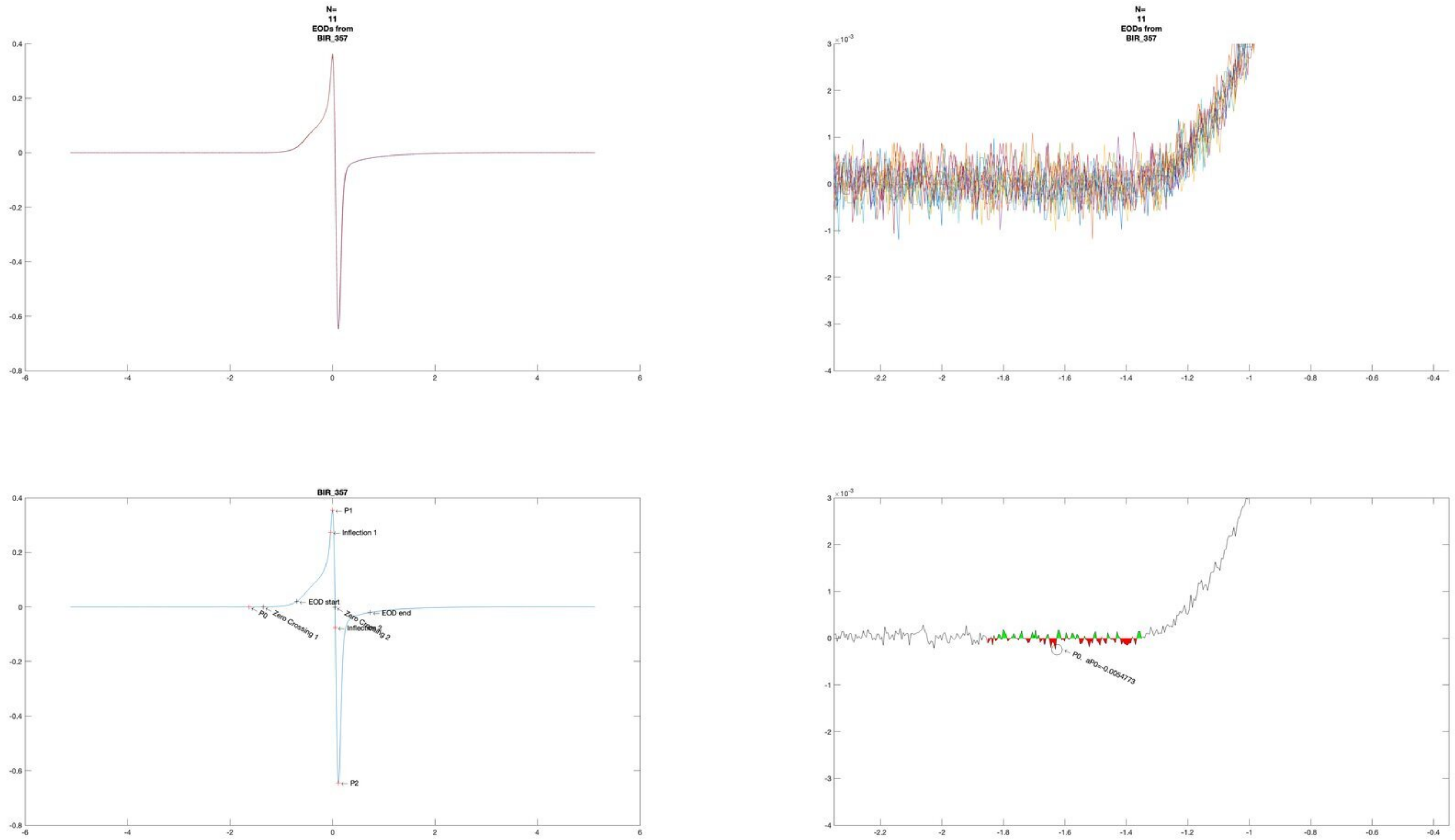
N=
11
EODs from
BIR_357
N=
11
EODs from
BIR_357
BIR_357
P1
Inflection 1
EOD start
EOD end
P0
Zero Crossing 1
Zero Crossing 2
Inflection 2
P2
P0, aP0=-0.0054773

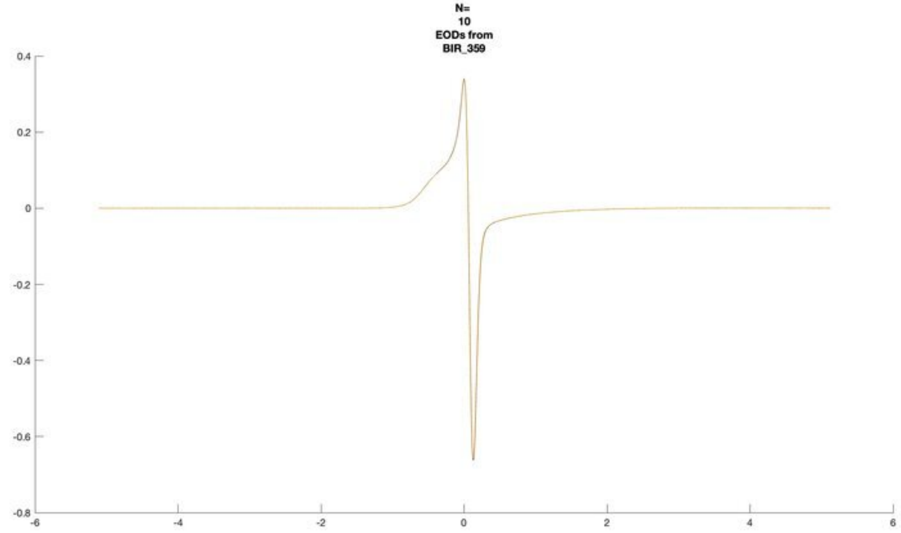
N=
10
EODs from
BIR_359

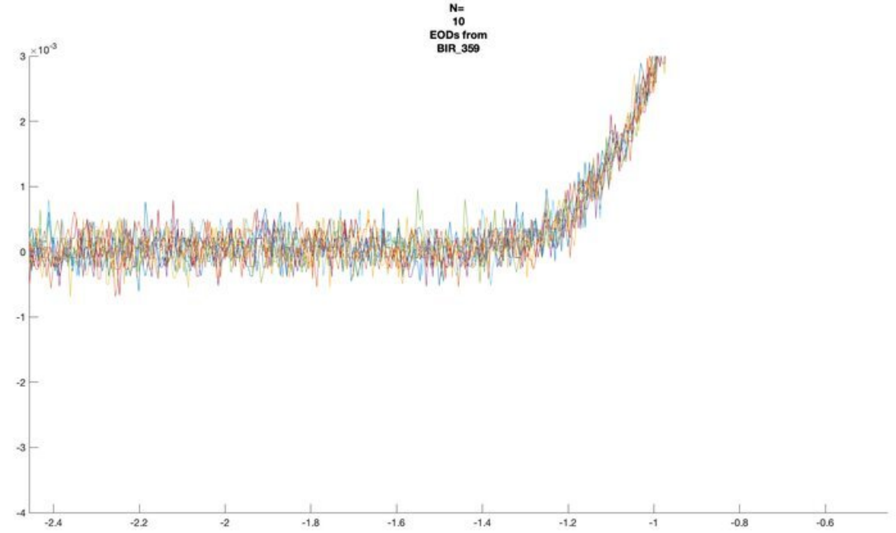
N=
10
EODs from
BIR_359
×10^-3

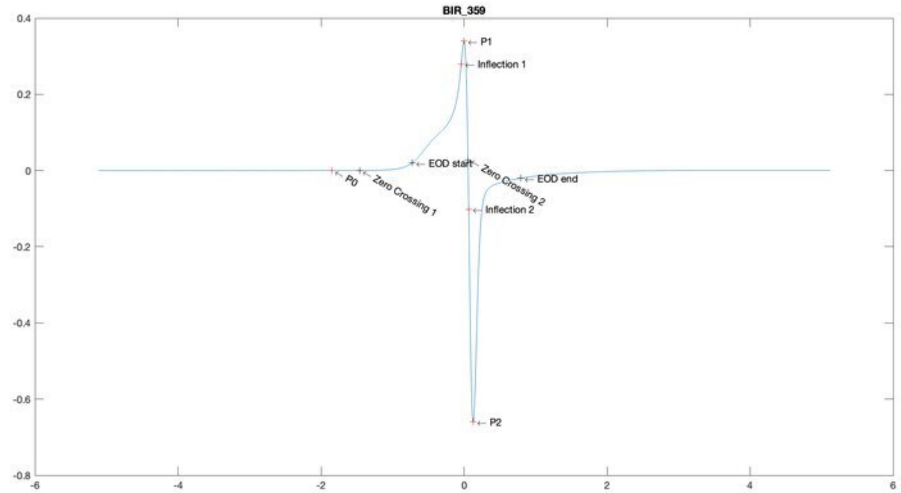
BIR_359
P1
Inflection 1
EOD start
Zero Crossing 2
EOD end
P0
Zero Crossing 1
Inflection 2
P2

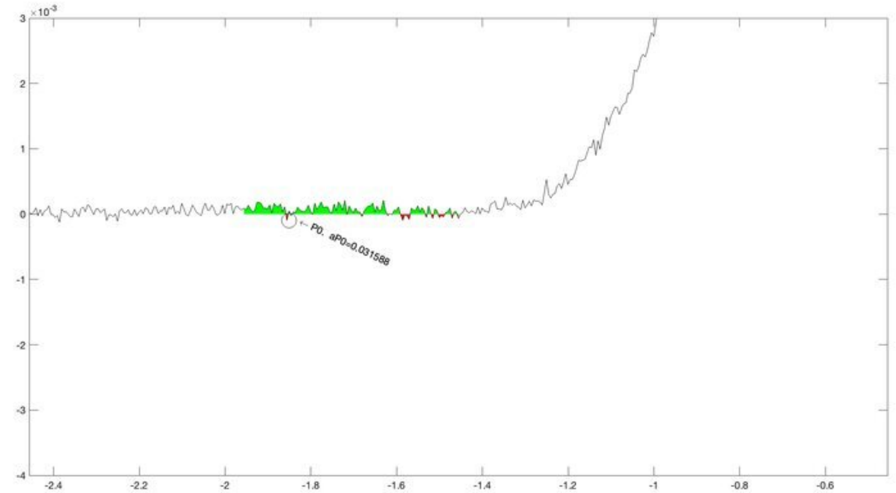
×10^-3
P0, aP0=0.031588

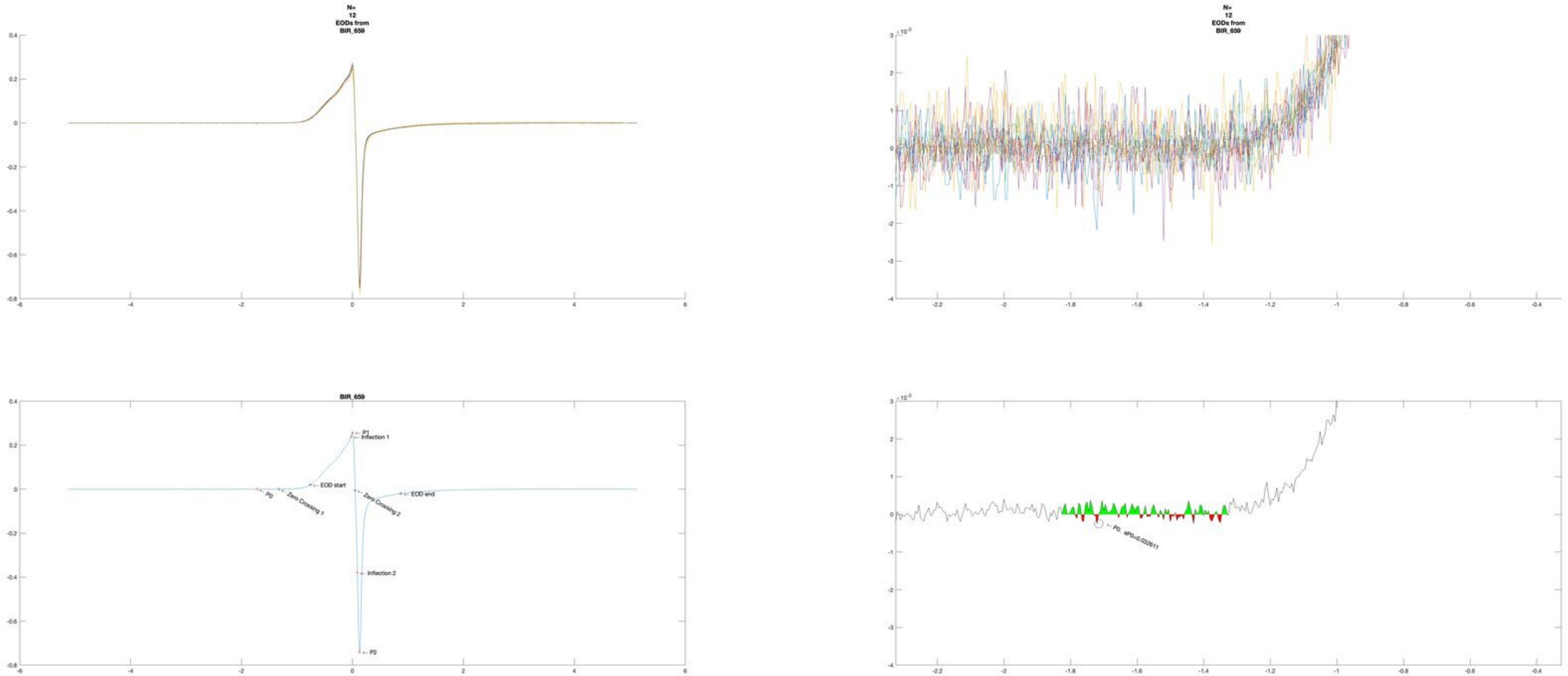

N=
12
EODs from
BIR_659
N=
12
EODs from
BIR_659
BIR_659
P1
Inflection 1
EOD start
P0
Zero Crossing 1
Zero Crossing 2
EOD end
Inflection 2
P2
P0, aP0=0.032611

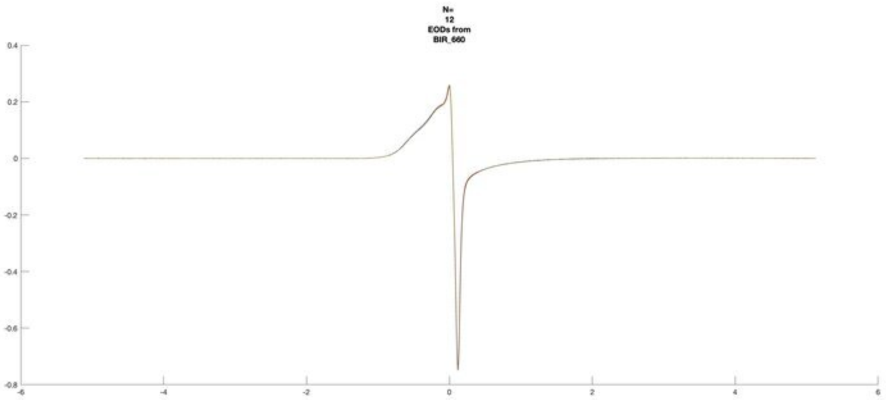

N=
12
EODs from
BIR_660
0.4
0.2
0
-0.2
-0.4
-0.6
-0.8
-6
-4
-2
0
2
4
6


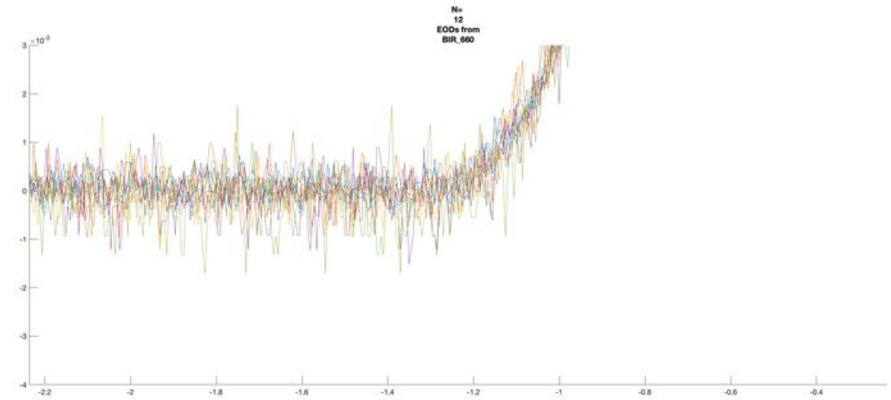

N=
12
EODs from
BIR_660
×10⁻³
3
2
1
0
-1
-2
-3
-4
-2.2
-2
-1.8
-1.6
-1.4
-1.2
-1
-0.8
-0.6
-0.4


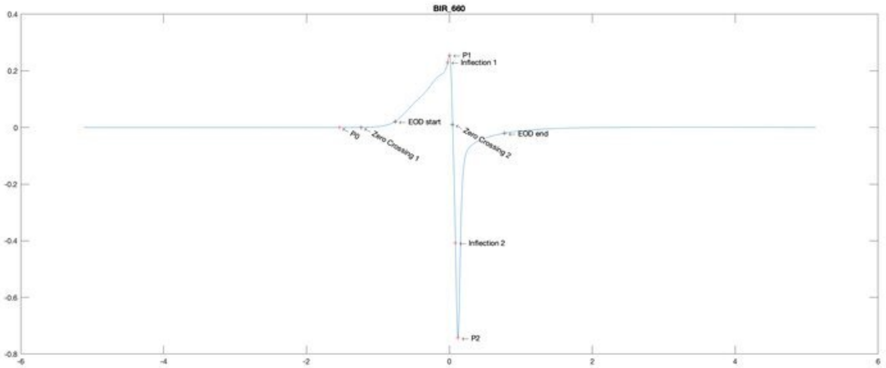

BIR_660
P1
Inflection 1
EOD start
P0
Zero Crossing 1
Zero Crossing 2
EOD end
Inflection 2
P2
0.4
0.2
0
-0.2
-0.4
-0.6
-0.8
-6
-4
-2
0
2
4
6


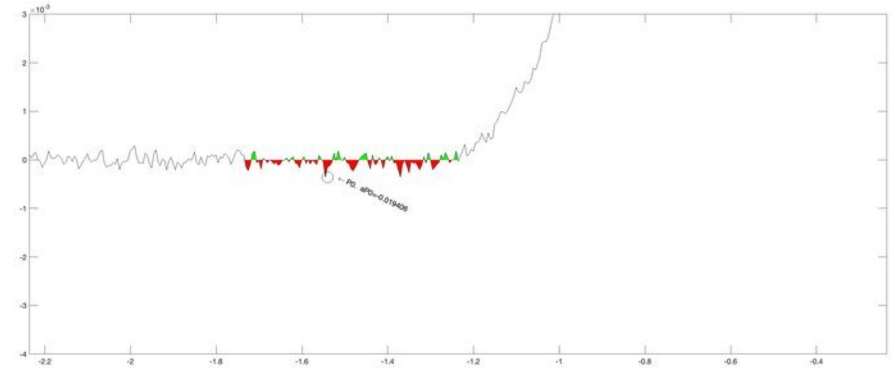

×10⁻³
P0, aP0=-0.019406
3
2
1
0
-1
-2
-3
-4
-2.2
-2
-1.8
-1.6
-1.4
-1.2
-1
-0.8
-0.6
-0.4

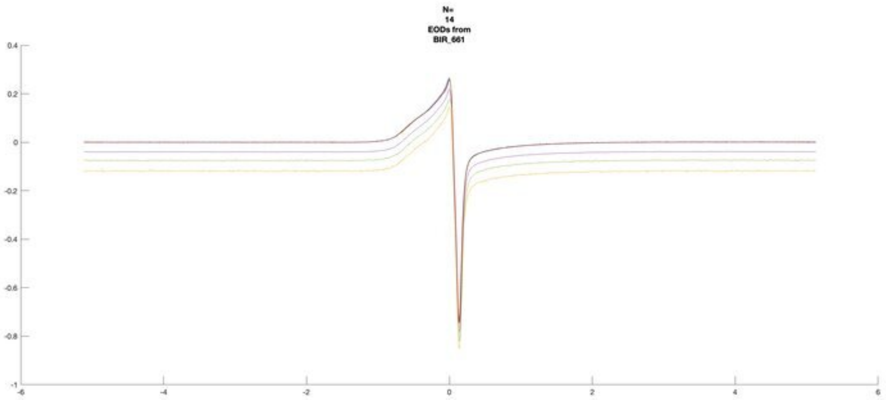
N=
14
EODs from
BIR_661

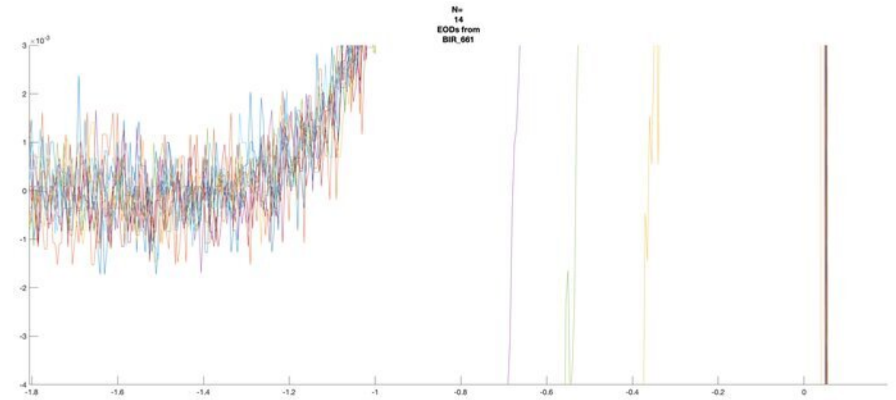
N=
14
EODs from
BIR_661

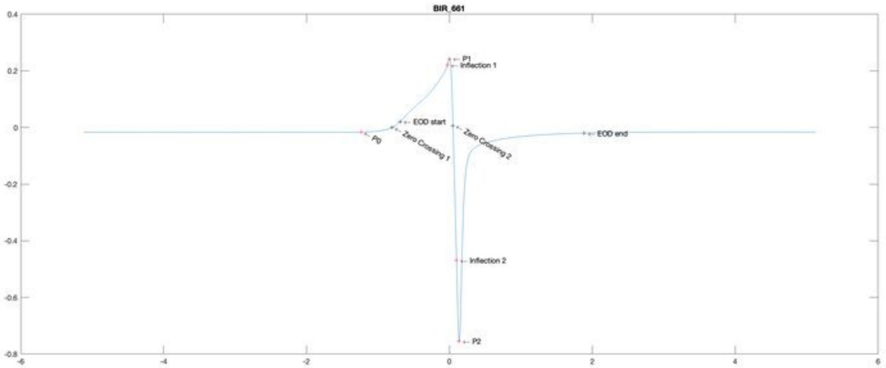
BIR_661
P1
Inflection 1
EOD start
P0
Zero Crossing 1
Zero Crossing 2
EOD end
Inflection 2
P2

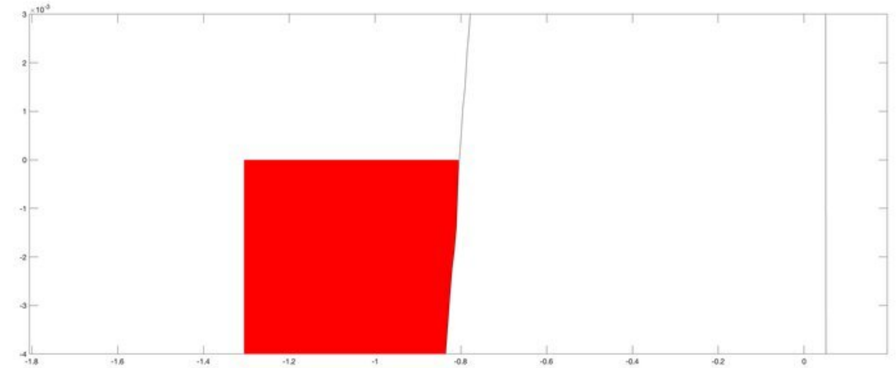

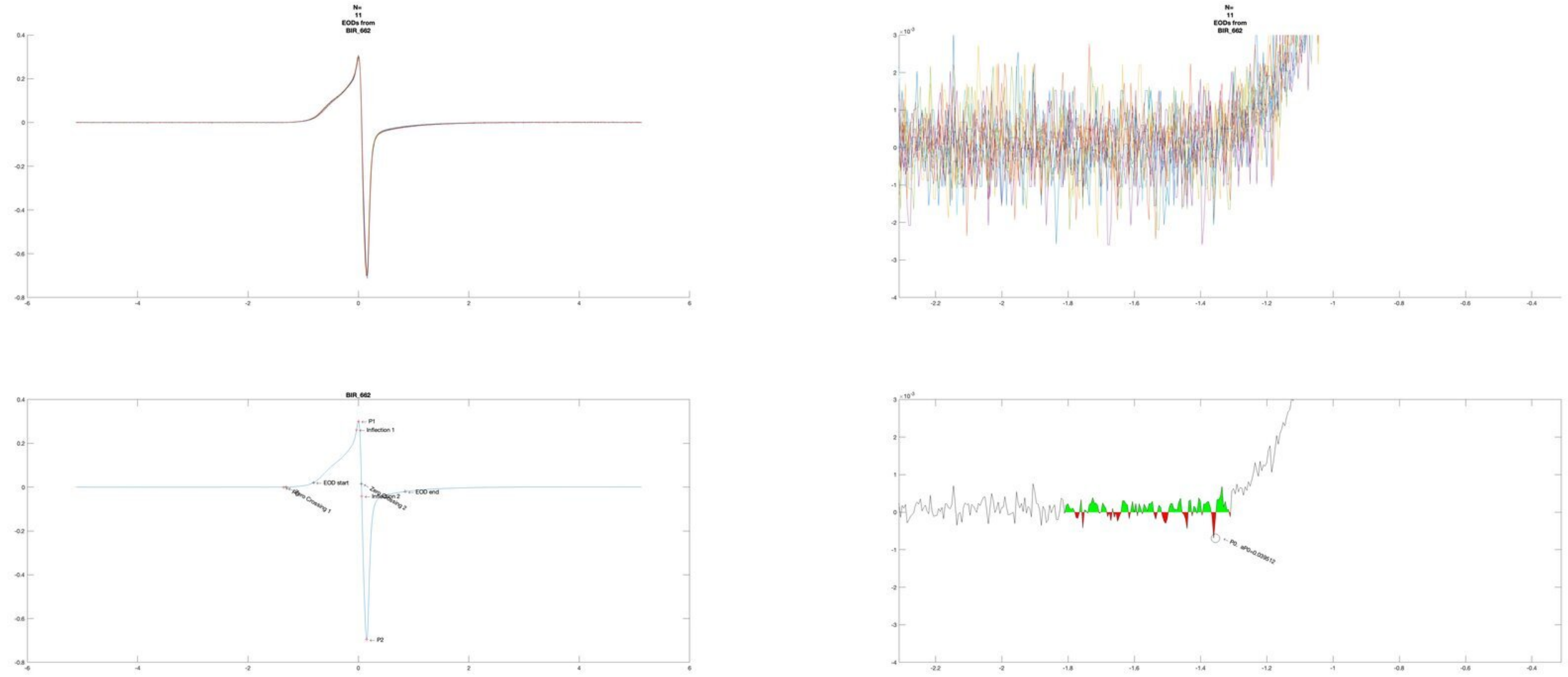

N=
11
EODs from
BIR_662
BIR_662
P1
Inflection 1
EOD start
Zero Crossing 1
Zero Crossing 2
Inflection 2
EOD end
P2
P0

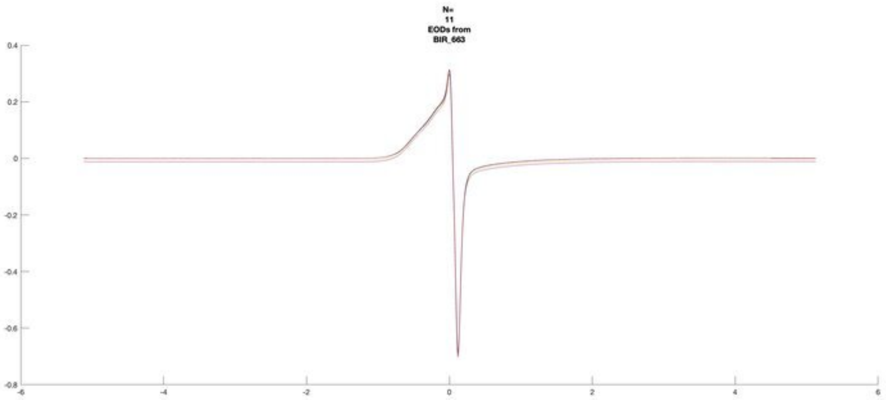
N=
11
EODs from
BIR_663

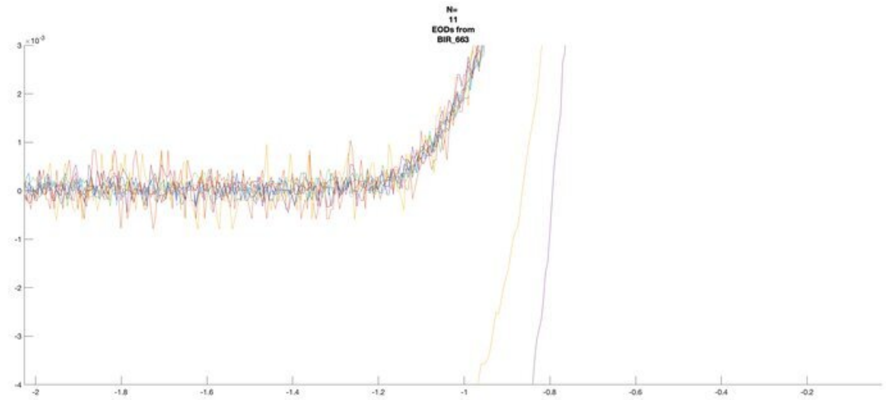
N=
11
EODs from
BIR_663

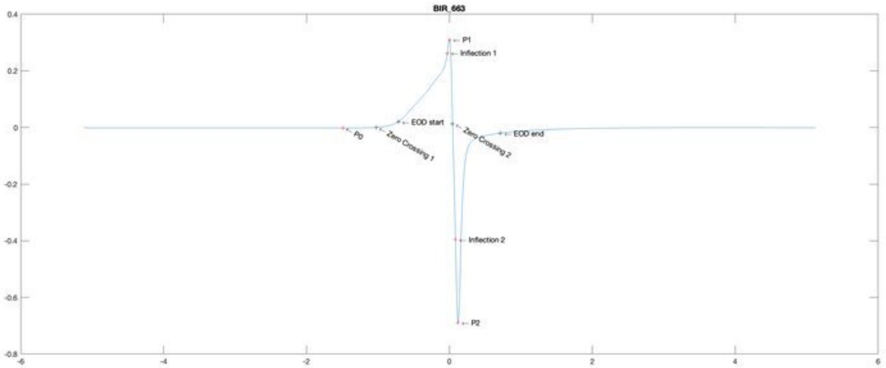
BIR_663
P1
Inflection 1
EOD start
P0
Zero Crossing 1
Zero Crossing 2
EOD end
Inflection 2
P2

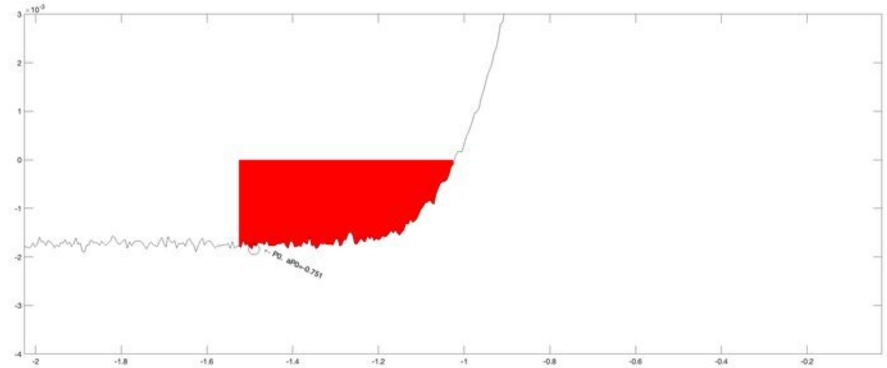
P0, aP0=-0.751

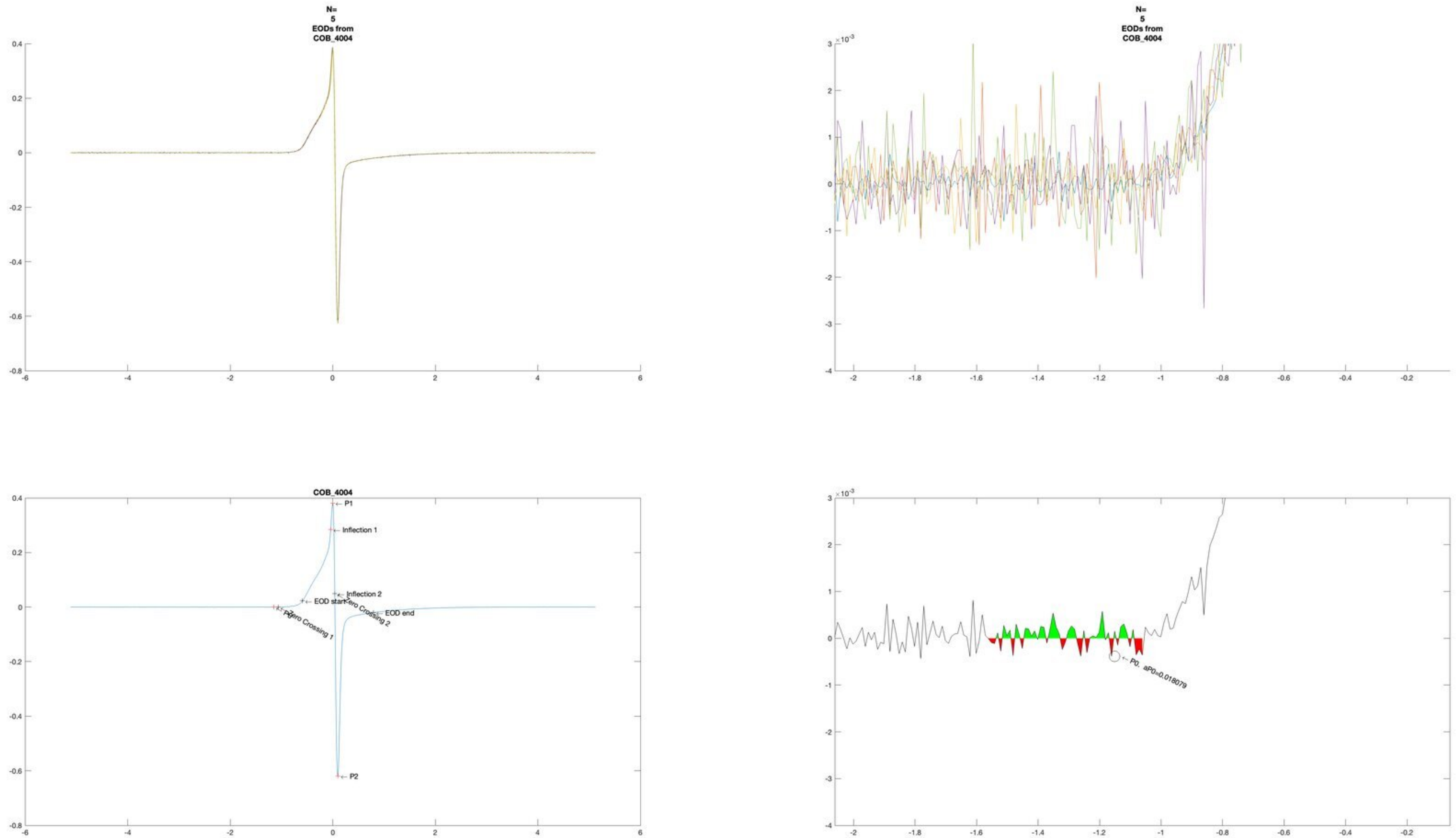
N=
5
EODs from
COB_4004
N=
5
EODs from
COB_4004
COB_4004
P1
Inflection 1
Inflection 2
EOD start
Zero Crossing 2
EOD end
Zero Crossing 1
P2
P0, aP0=0.018079

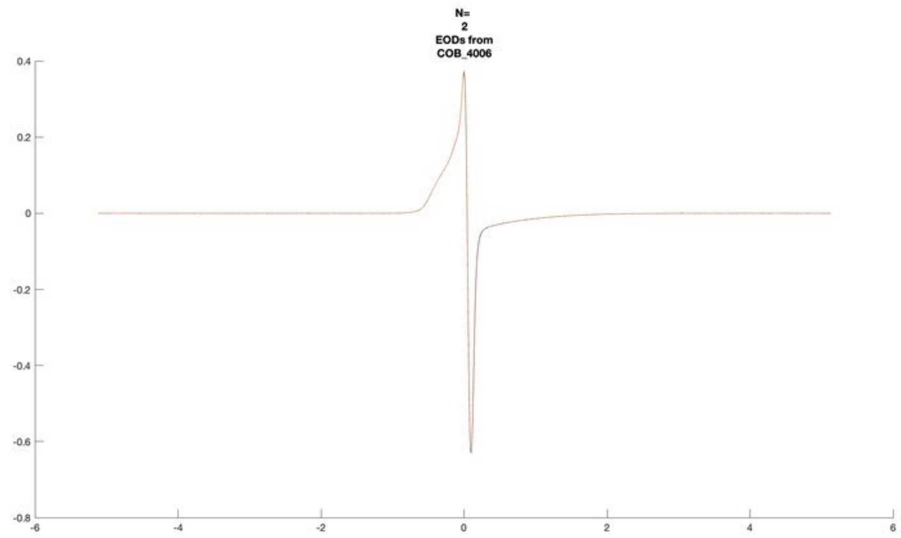
N=
2
EODs from
COB_4006

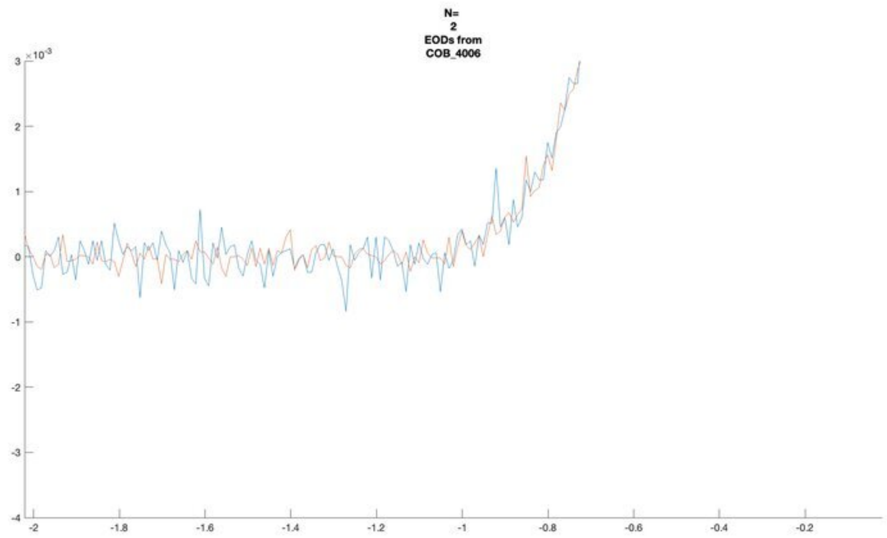
N=
2
EODs from
COB_4006

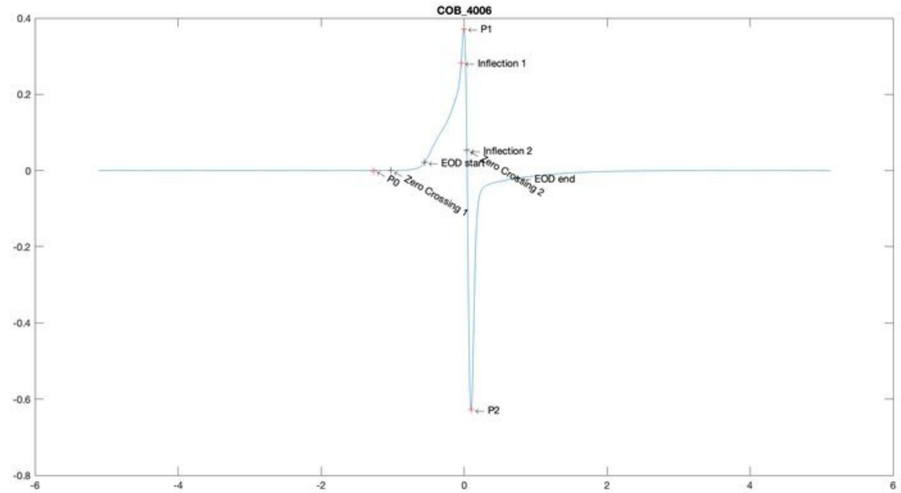
COB_4006
P1
Inflection 1
Inflection 2
EOD start
Zero Crossing 2
EOD end
P0
Zero Crossing 1
P2

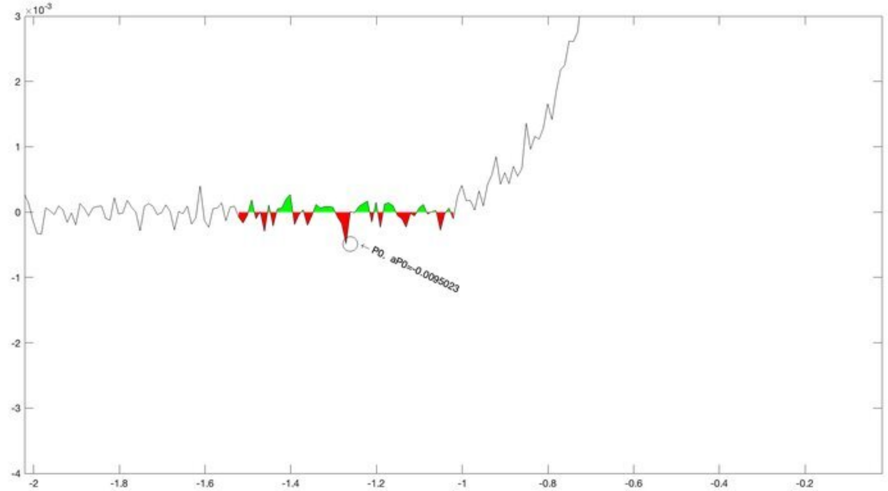
P0, aP0=-0.0095023

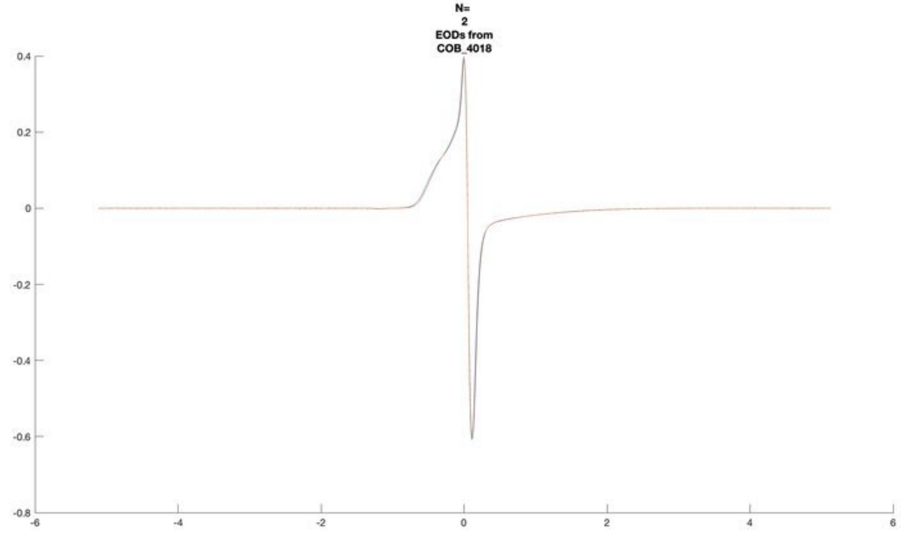
N=
2
EODs from
COB_4018

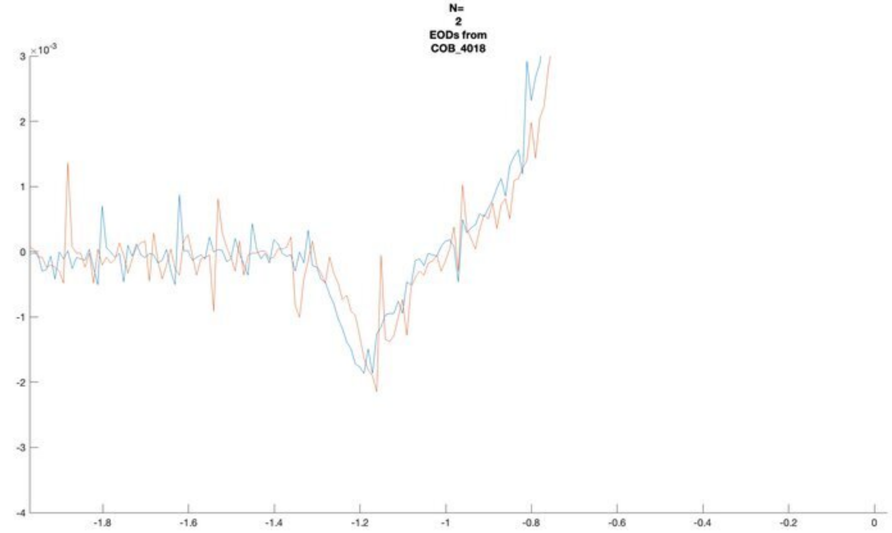
N=
2
EODs from
COB_4018
×10⁻³

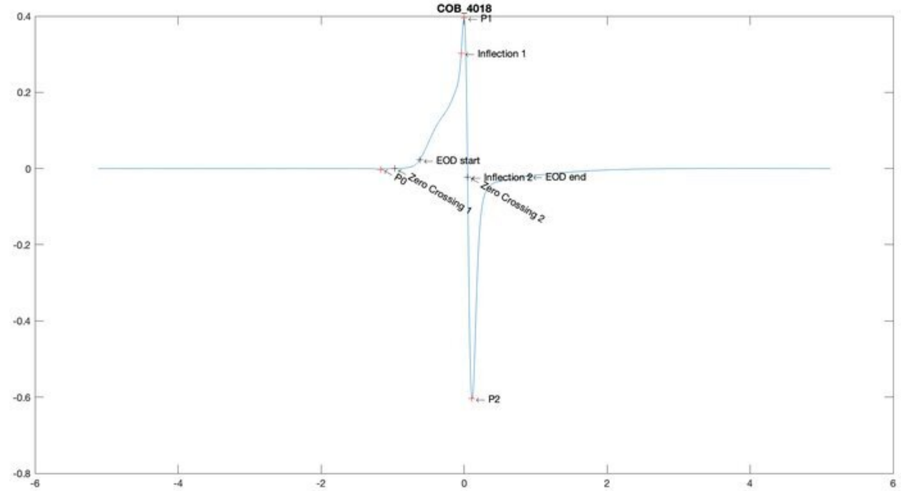
COB_4018
P1
Inflection 1
EOD start
Inflection 2
EOD end
P0
Zero Crossing 1
Zero Crossing 2
P2

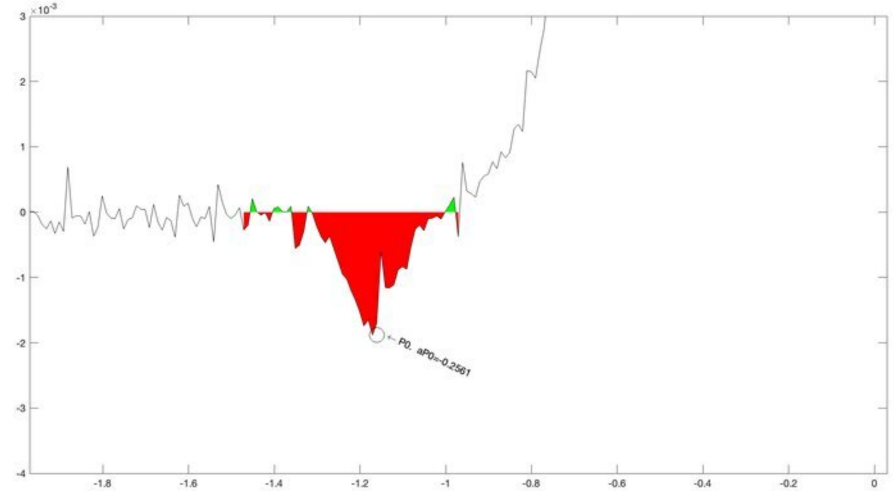
×10⁻³
P0. aP0=-0.2561

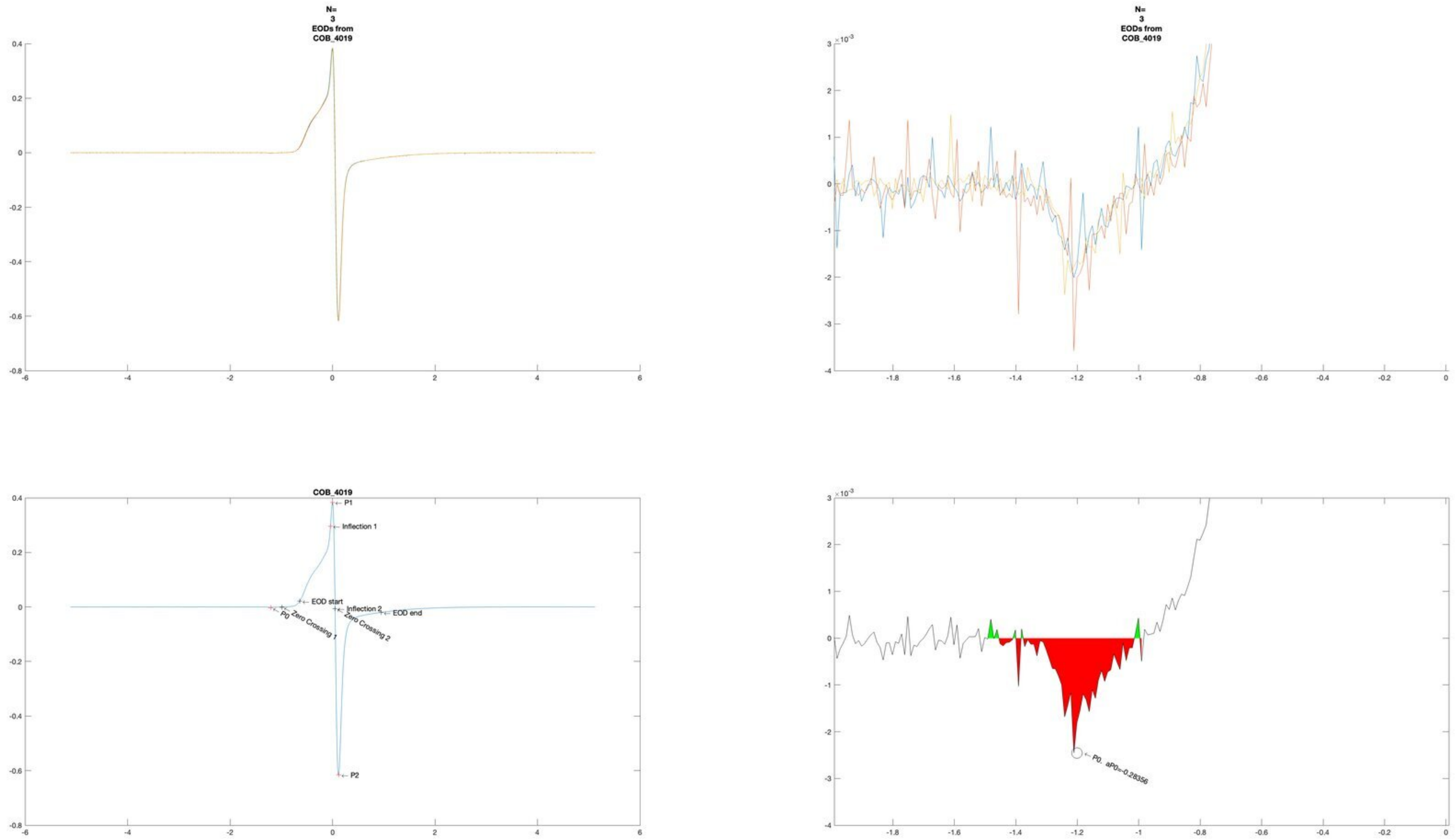
N=
3
EODs from
COB_4019
N=
3
EODs from
COB_4019
COB_4019
P1
Inflection 1
EOD start
Inflection 2
EOD end
P0
Zero Crossing 1
Zero Crossing 2
P2
P0. aP0=-0.28356

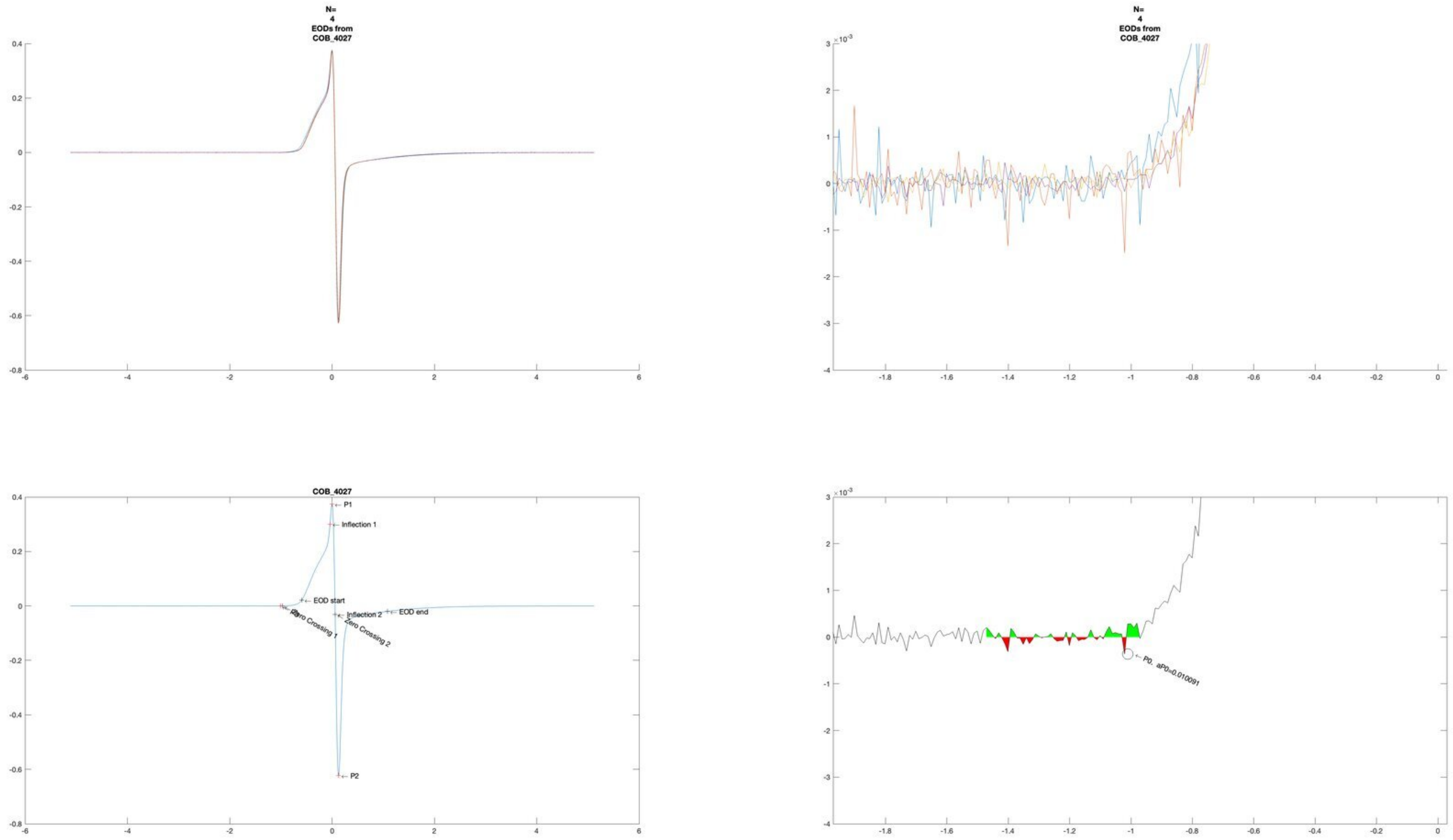
N=
4
EODs from
COB_4027
N=
4
EODs from
COB_4027
COB_4027
P1
Inflection 1
EOD start
Inflection 2
EOD end
Zero Crossing 1
Zero Crossing 2
P2
P0, aP0=0.010091

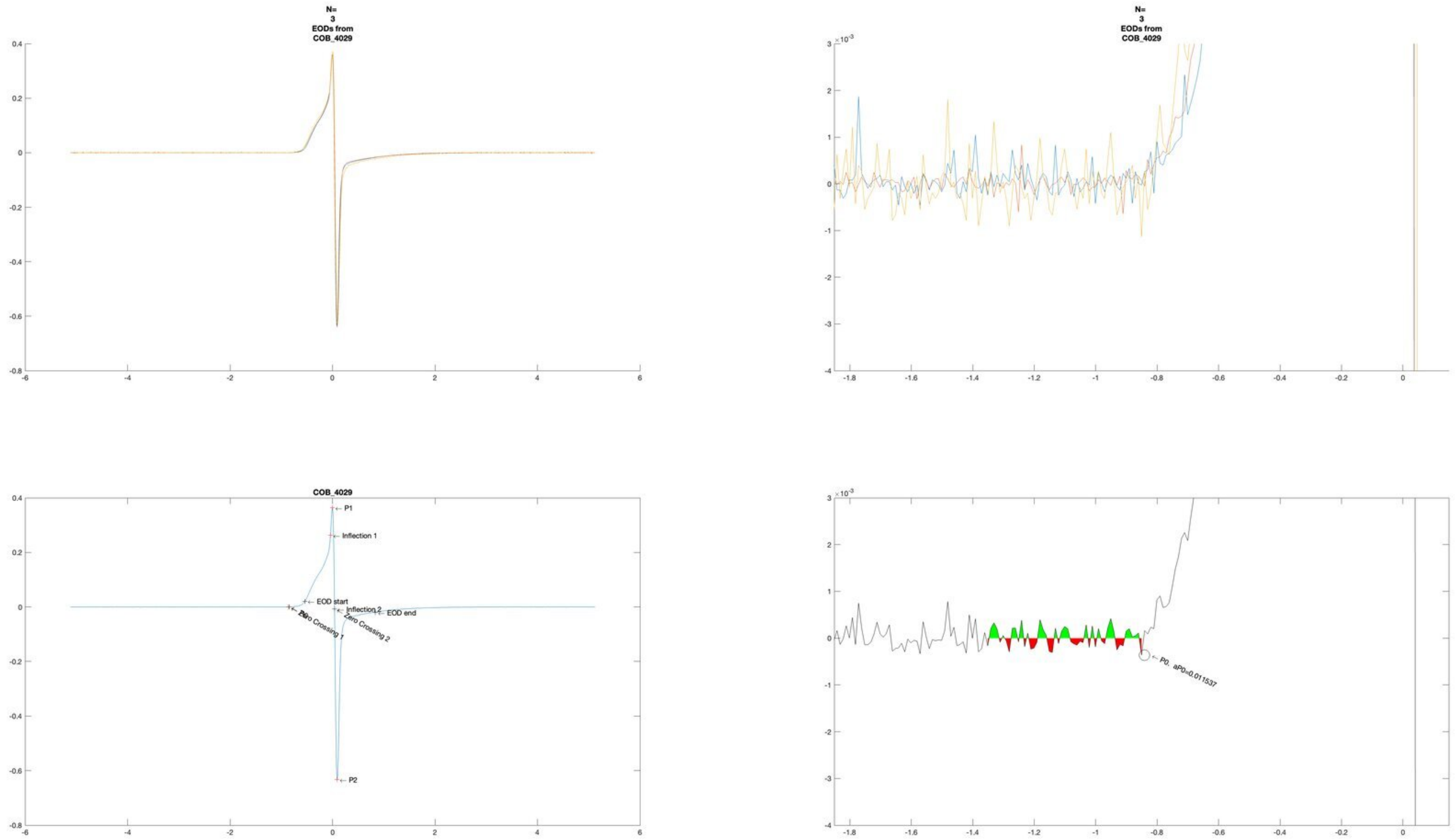
N=
3
EODs from
COB_4029
N=
3
EODs from
COB_4029
COB_4029
P1
Inflection 1
EOD start
Inflection 2
EOD end
Zero Crossing 1
Zero Crossing 2
P2
P0, aP0=0.011537

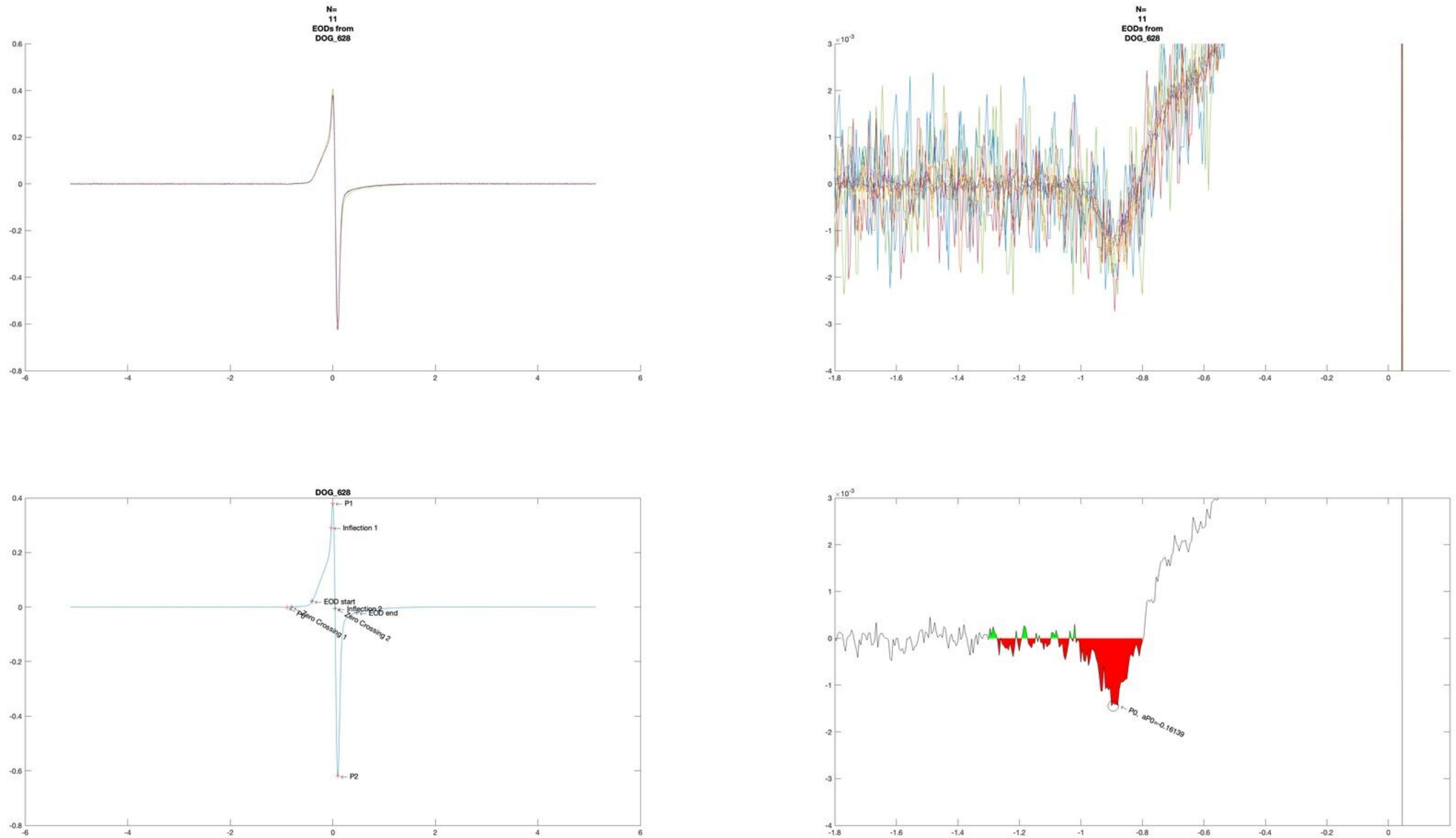
N=
11
EODs from
DOG_628
N=
11
EODs from
DOG_628
DOG_628
P1
Inflection 1
EOD start
Inflection 2
EOD end
Zero Crossing 1
Zero Crossing 2
P2
P0, aP0=-0.16139

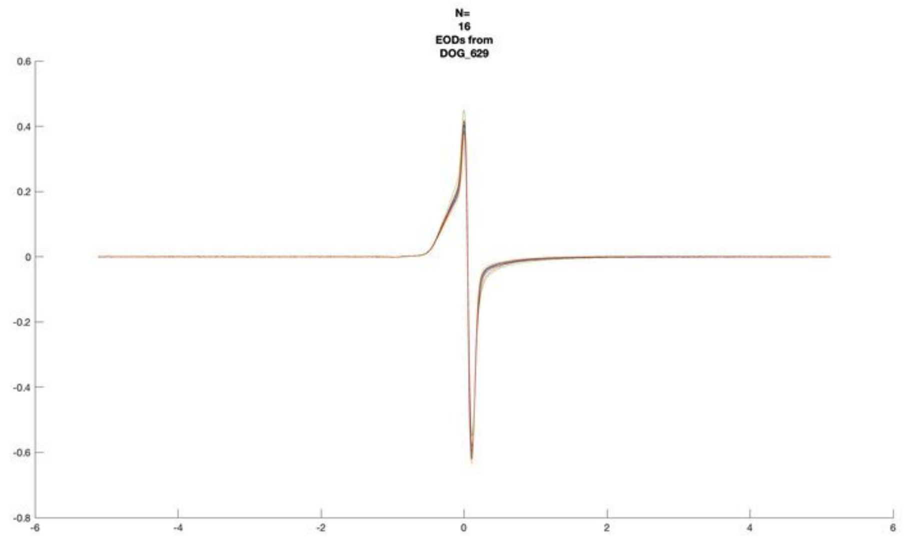
N=
16
EODs from
DOG_629

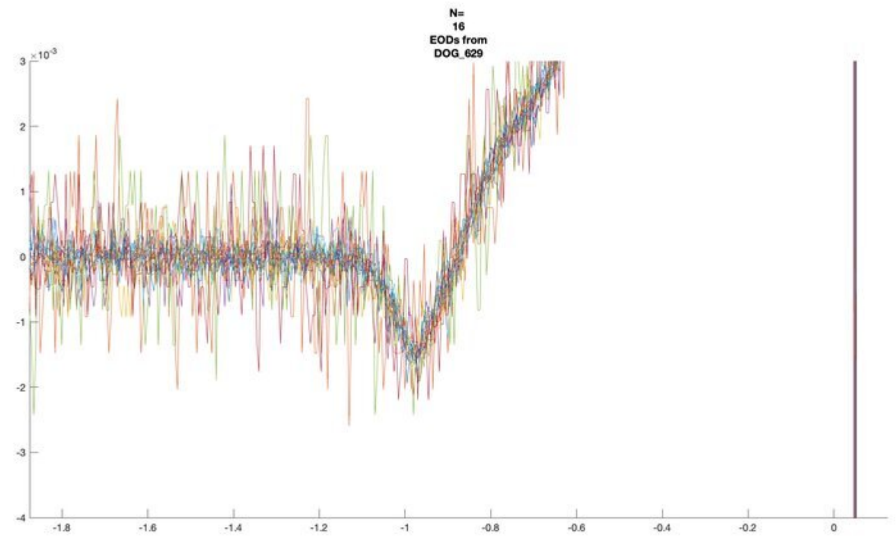
N=
16
EODs from
DOG_629
×10^-3

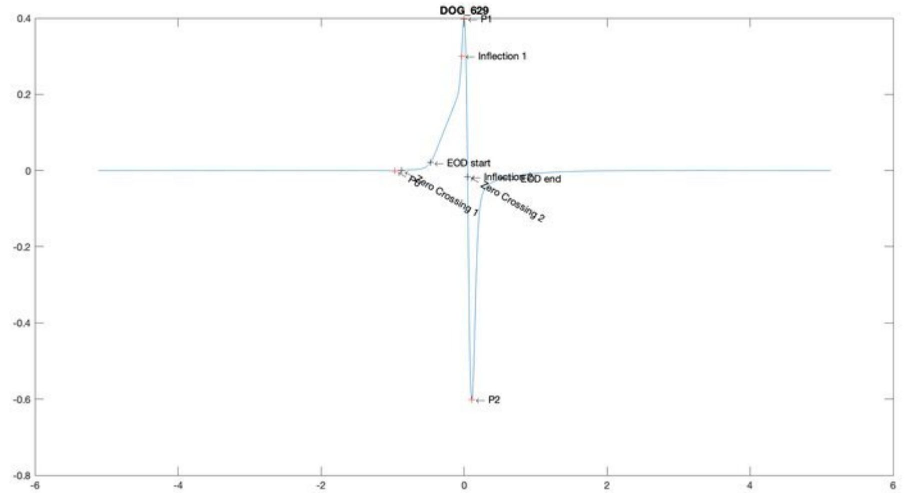
DOG_629
P1
Inflection 1
EOD start
EOD end
P0
Zero Crossing 1
Zero Crossing 2
P2

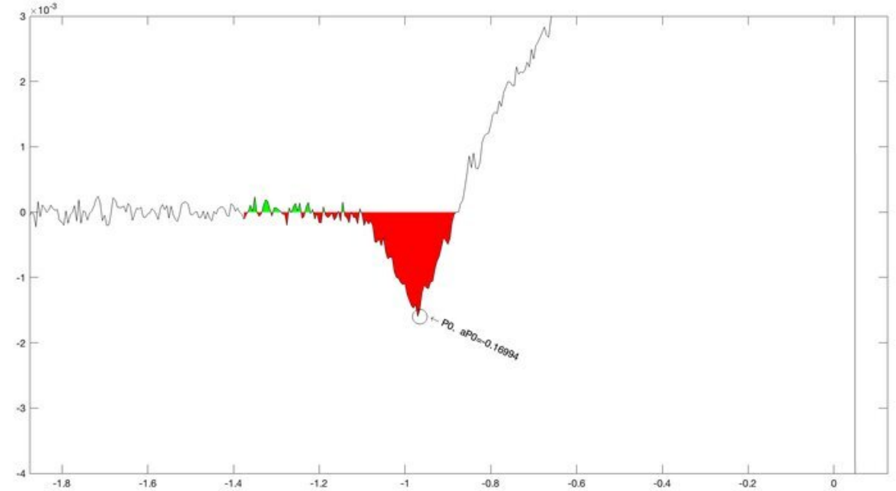
×10^-3
P0, aP0=-0.16994

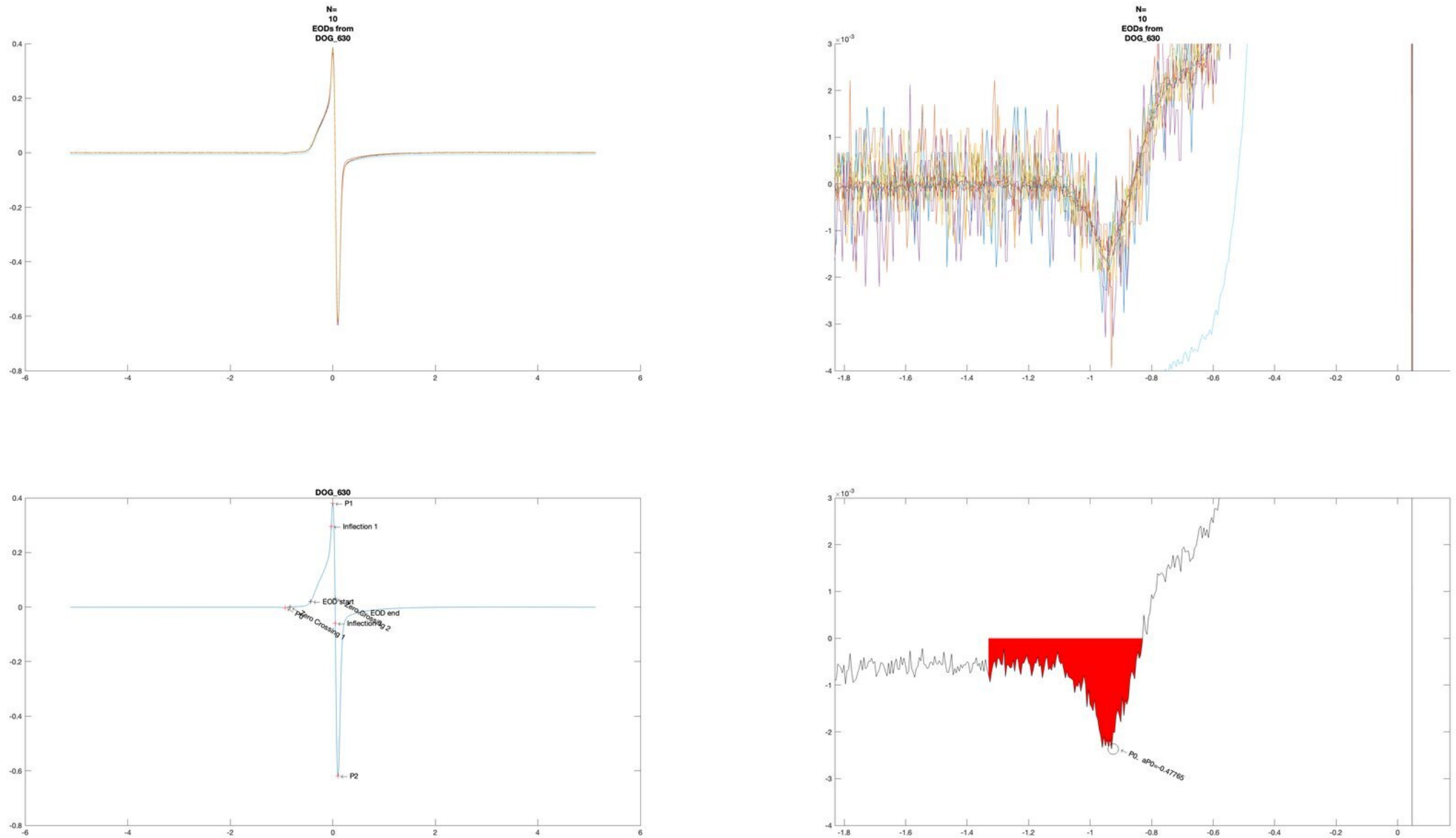
N=
10
EODs from
DOG_630
N=
10
EODs from
DOG_630
DOG_630
P1
Inflection 1
EOD start
EOD end
Zero Crossing 1
Zero Crossing 2
Inflection 2
P2
P0, aP0=-0.47765

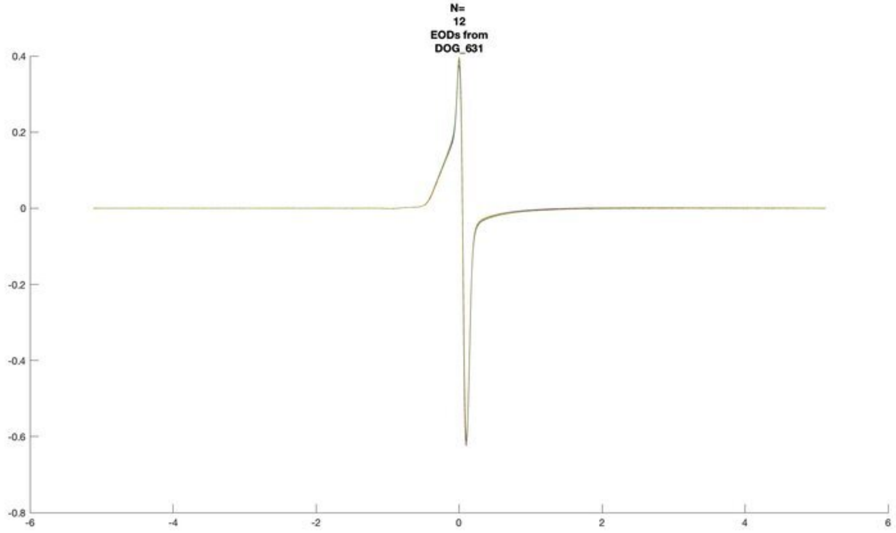
N=
12
EODs from
DOG_631

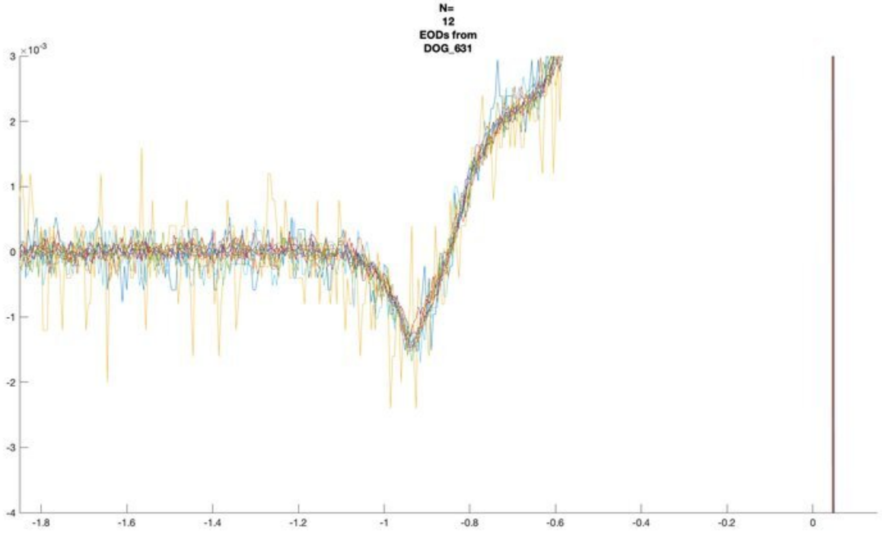
N=
12
EODs from
DOG_631

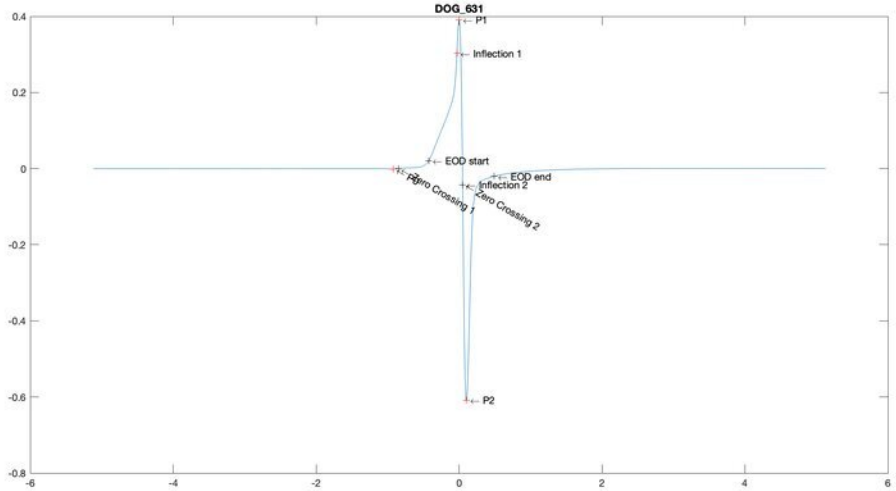
DOG_631
P1
Inflection 1
EOD start
EOD end
Inflection 2
Zero Crossing 1
Zero Crossing 2
P2

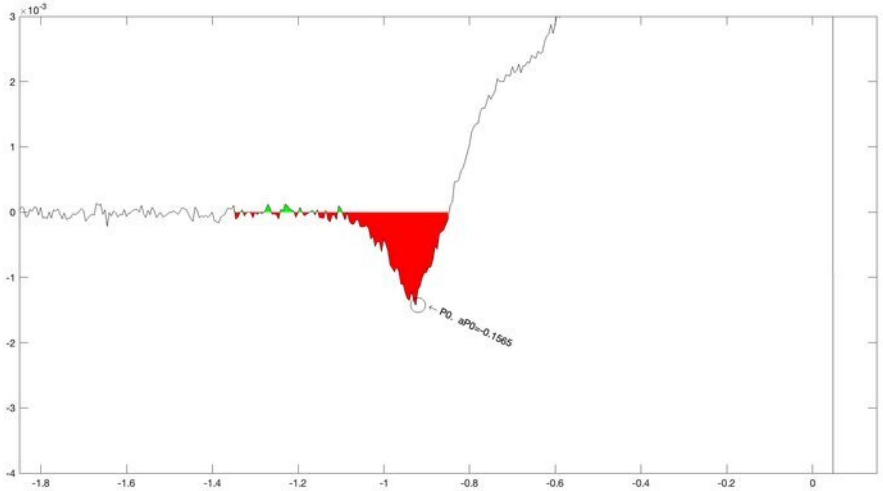
P0, aP0=-0.1565

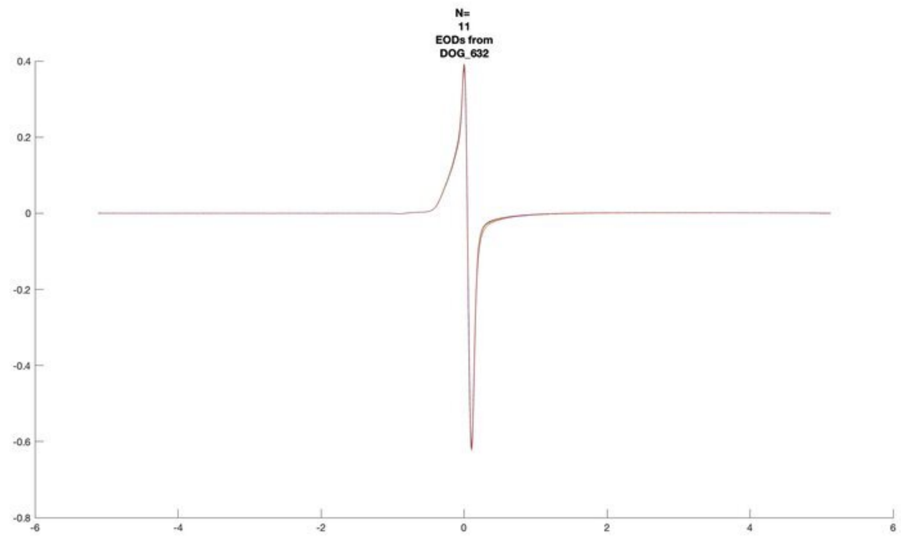
N=
11
EODs from
DOG_632

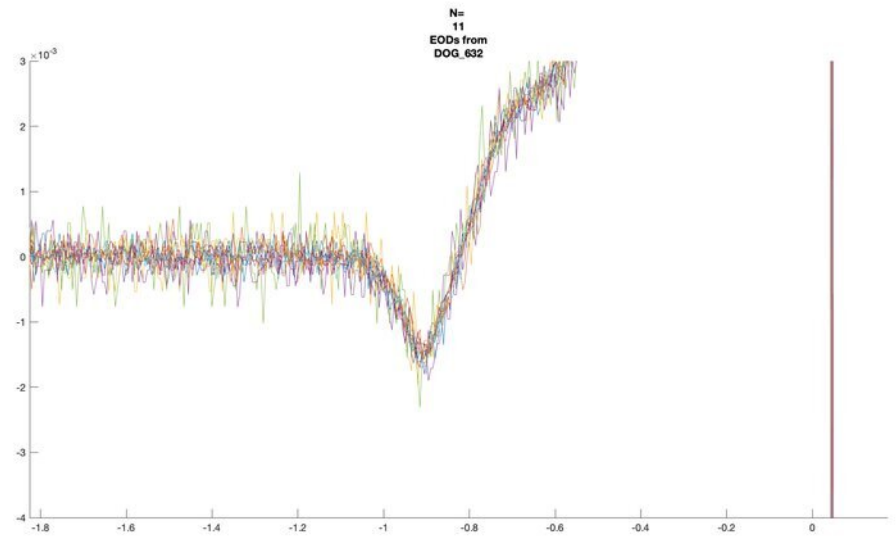
N=
11
EODs from
DOG_632

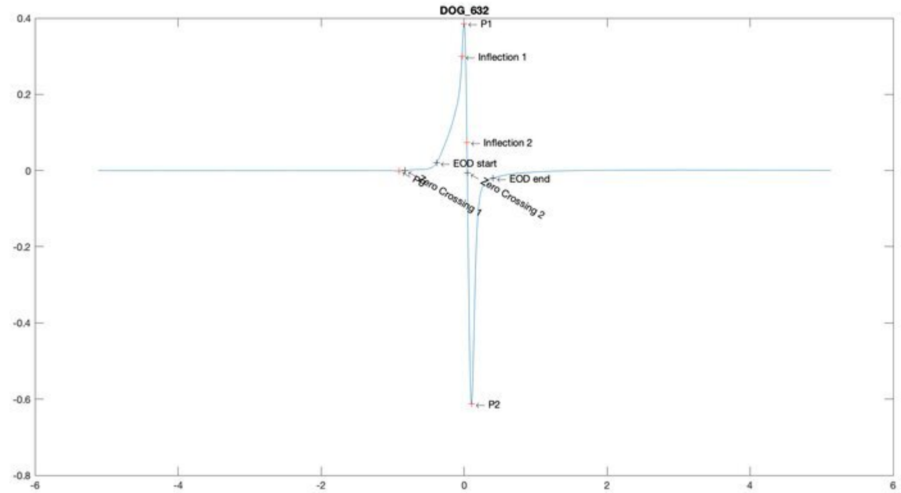
DOG_632
P1
Inflection 1
Inflection 2
EOD start
EOD end
Zero Crossing 1
Zero Crossing 2
P2

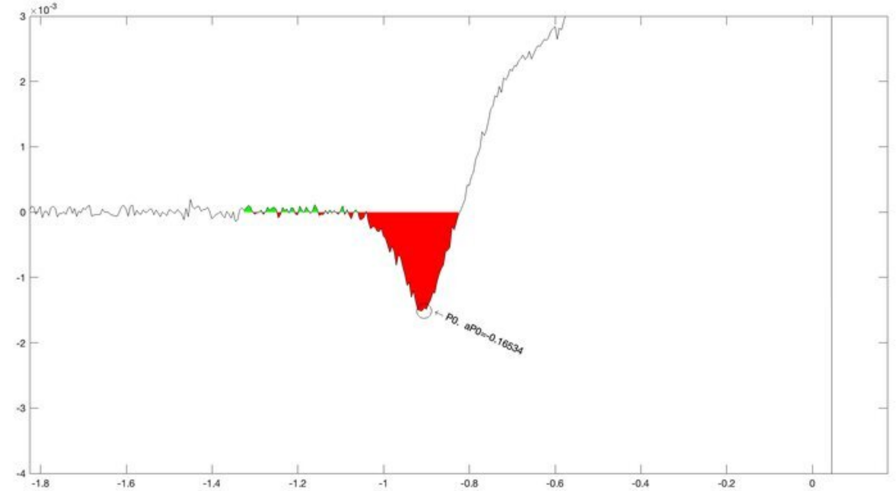
P0. aP0=-0.16534

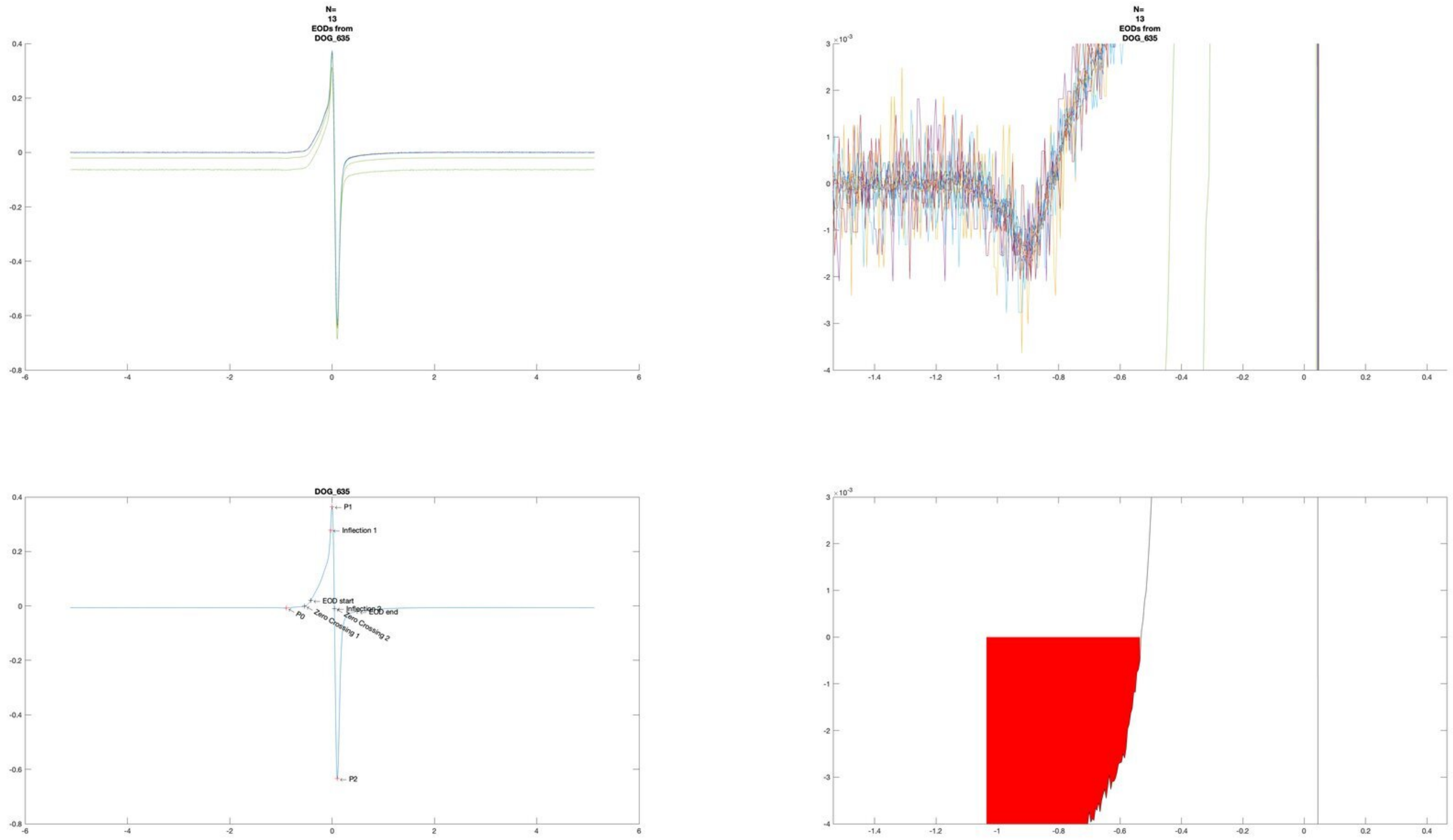
N=
13
EODs from
DOG_635
N=
13
EODs from
DOG_635
DOG_635
P1
Inflection 1
EOD start
Inflection 2
EOD end
P0
Zero Crossing 1
Zero Crossing 2
P2

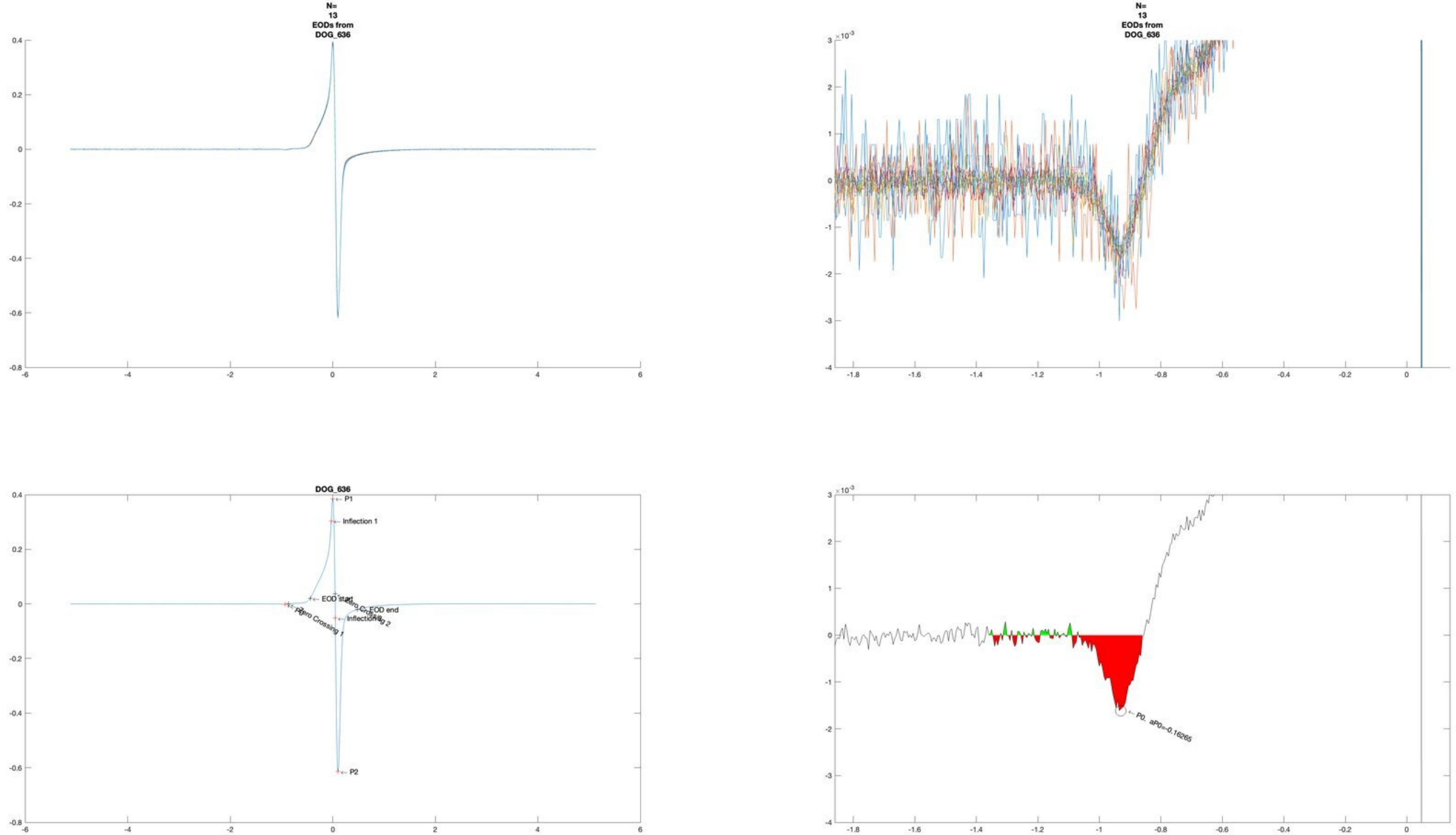

N=
13
EODs from
DOG_636
N=
13
EODs from
DOG_636
DOG_636
P1
Inflection 1
EOD start
EOD end
Zero Crossing 1
Zero Crossing 2
P2
P0. aP0=-0.16265

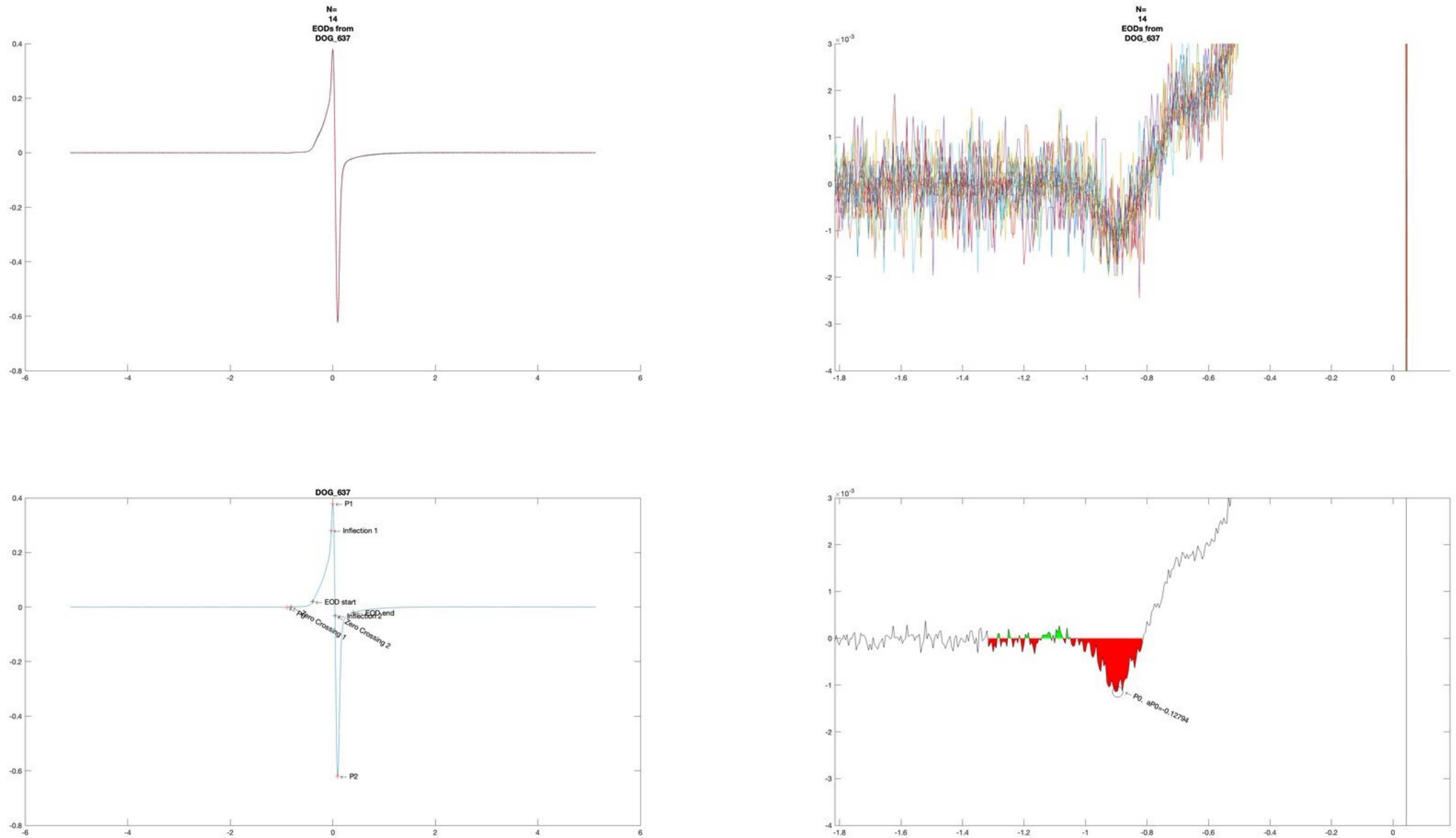
N=
14
EODs from
DOG_637
N=
14
EODs from
DOG_637
DOG_637
P1
Inflection 1
EOD start
EOD end
Inflection 2
Zero Crossing 1
Zero Crossing 2
P2
P0. aP0=-0.12794

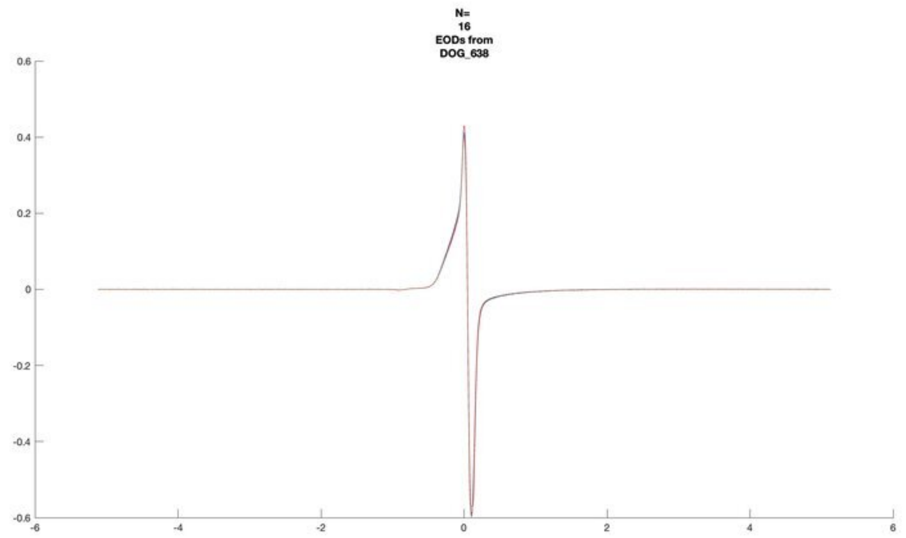

N=
16
EODs from
DOG_638


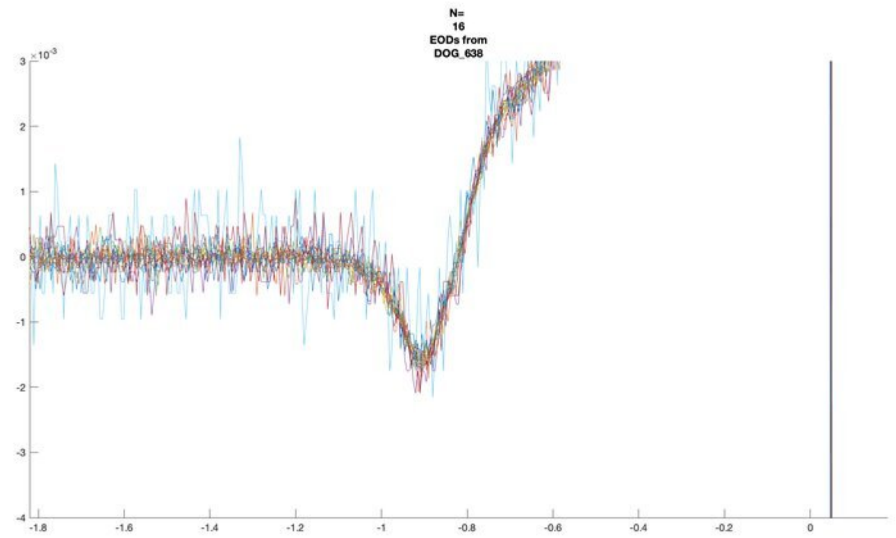

N=
16
EODs from
DOG_638


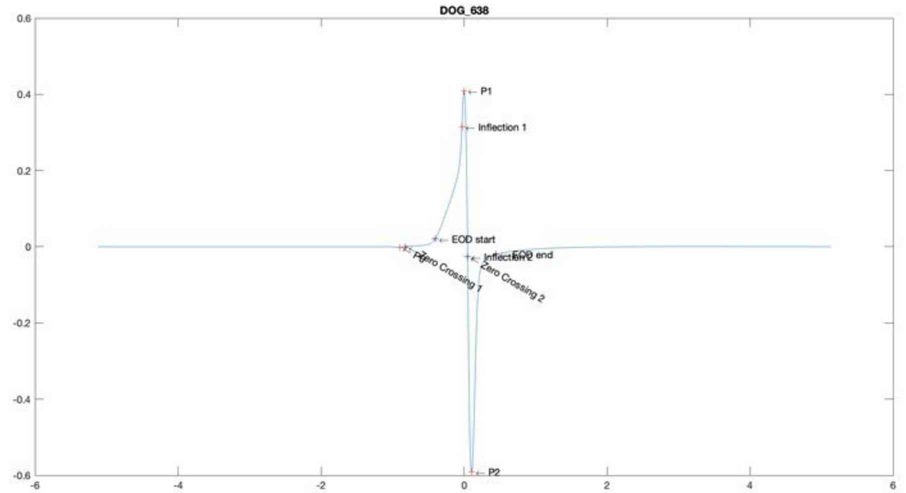

DOG_638
P1
Inflection 1
EOD start
EOD end
Zero Crossing 1
Zero Crossing 2
P2


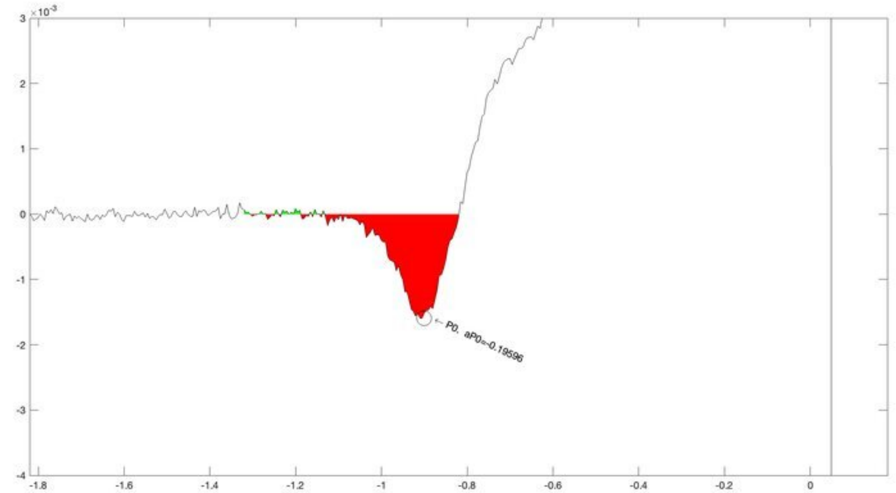

P0, aP0=-0.19596

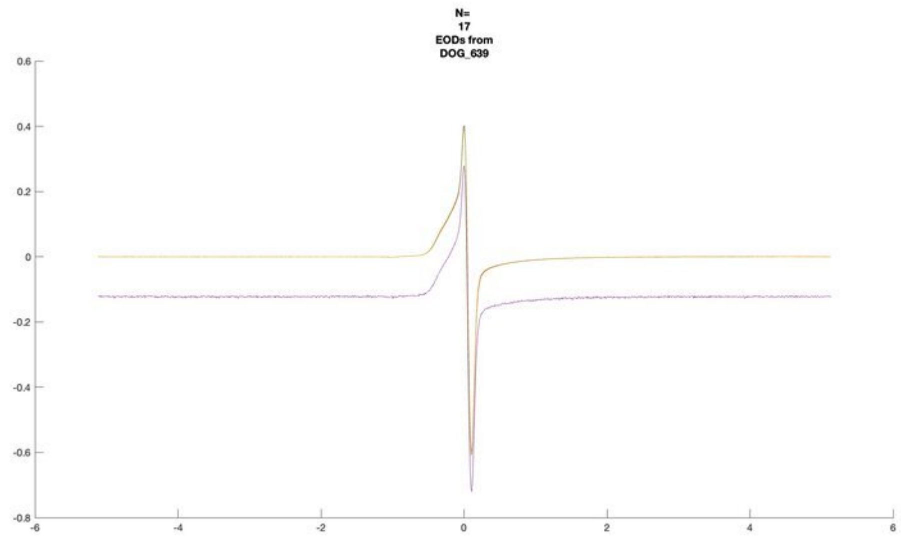
N=
17
EODs from
DOG_639

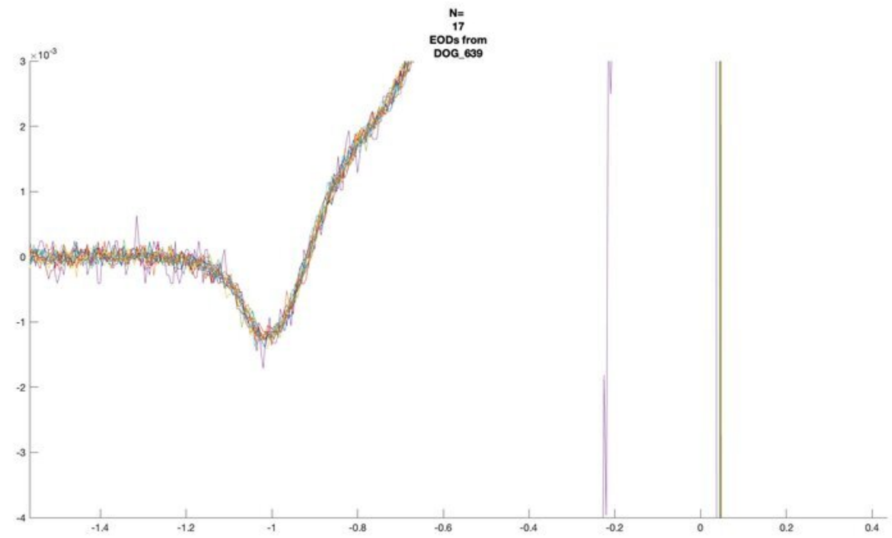
N=
17
EODs from
DOG_639

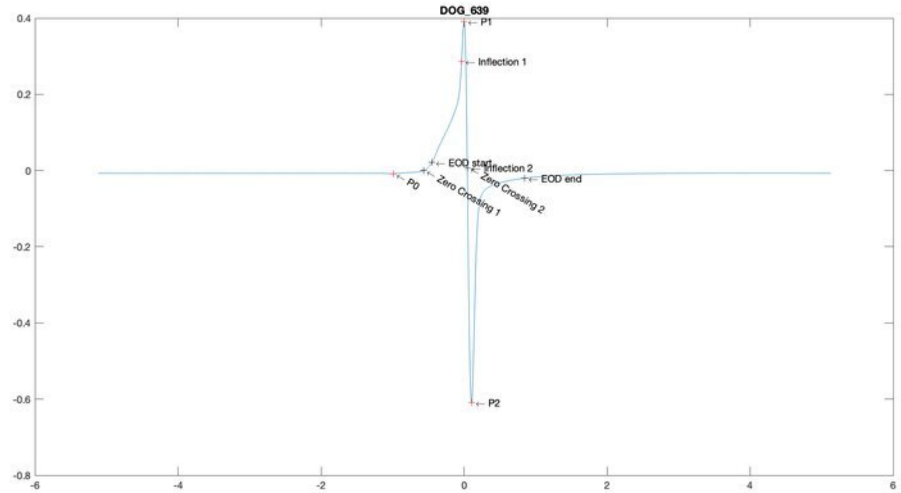
DOG_639
P1
Inflection 1
EOD start
Inflection 2
EOD end
P0
Zero Crossing 1
Zero Crossing 2
P2

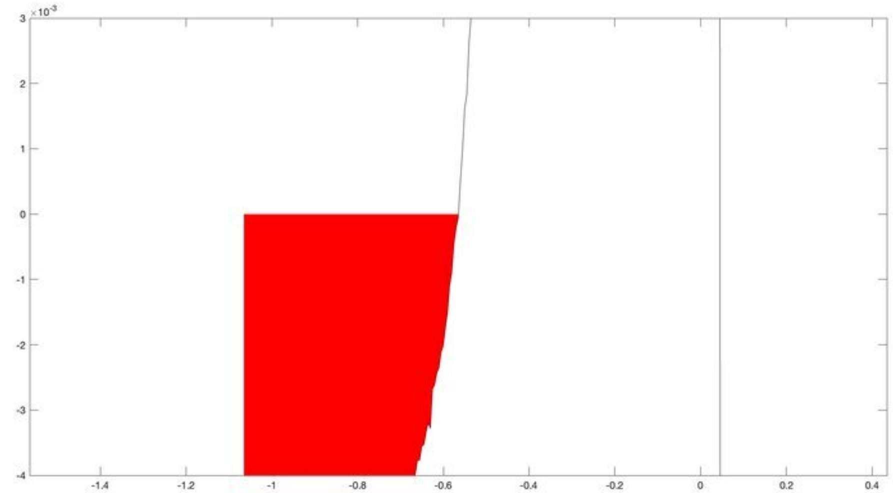

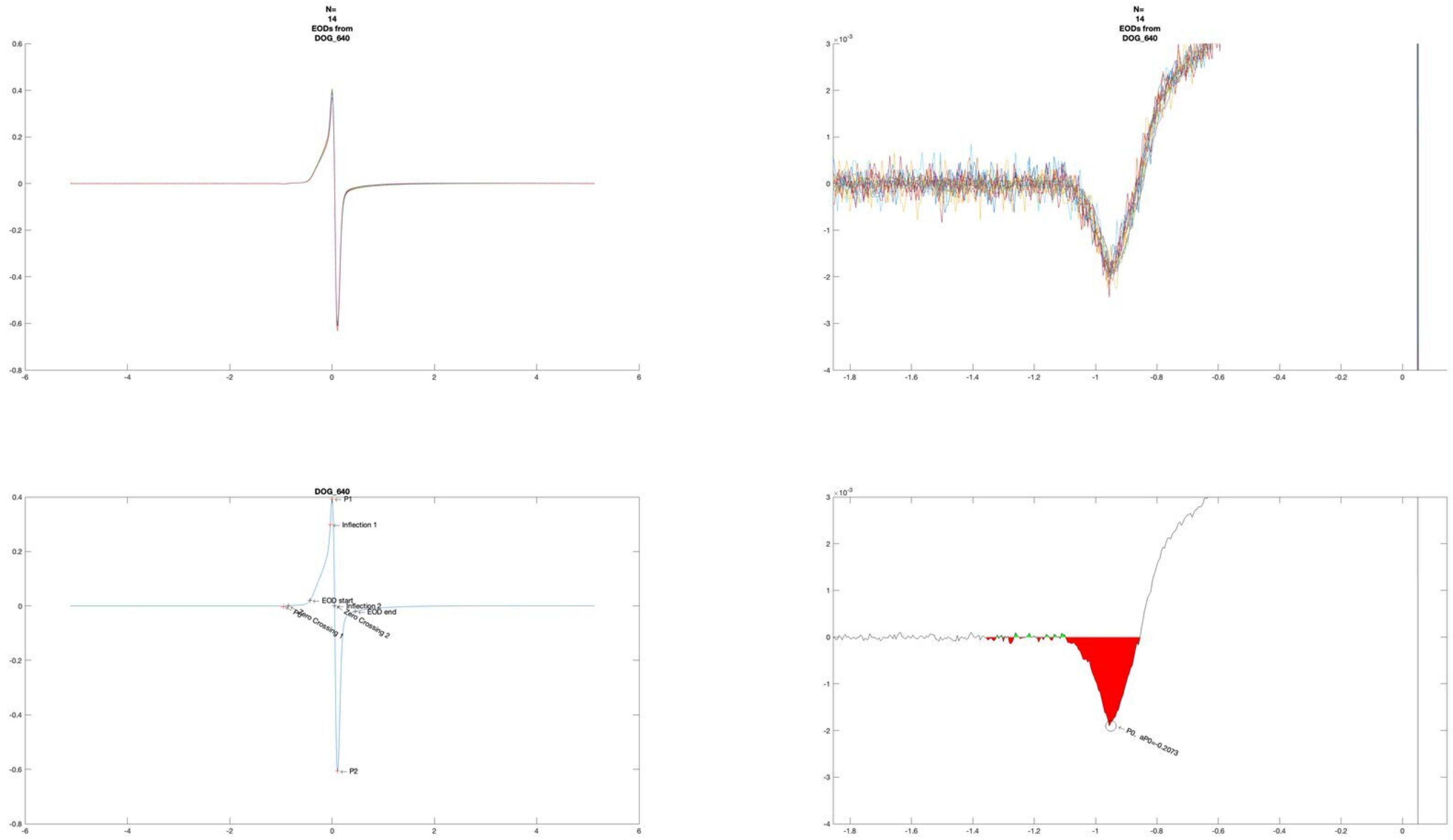

N=
14
EODs from
DOG_640
N=
14
EODs from
DOG_640
DOG_640
P1
Inflection 1
EOD start
Inflection 2
EOD end
Zero Crossing 1
Zero Crossing 2
P0
P2
P0, aP0=-0.2073

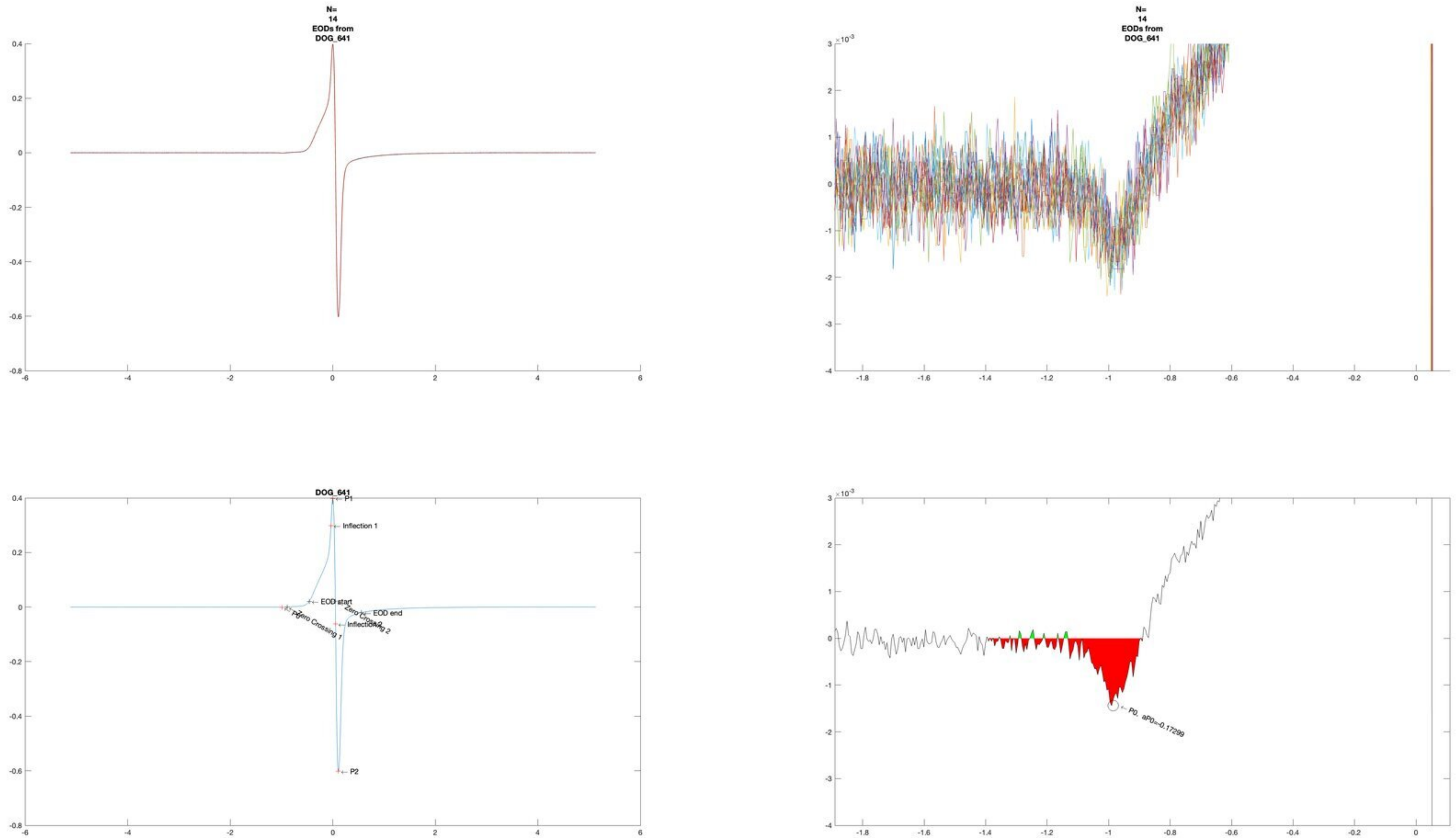
N=
14
EODs from
DOG_641
N=
14
EODs from
DOG_641
×10⁻³
DOG_641
P1
Inflection 1
EOD start
EOD end
P0
Zero Crossing 1
Zero Crossing 2
P2
×10⁻³
P0, aP0=-0.17299

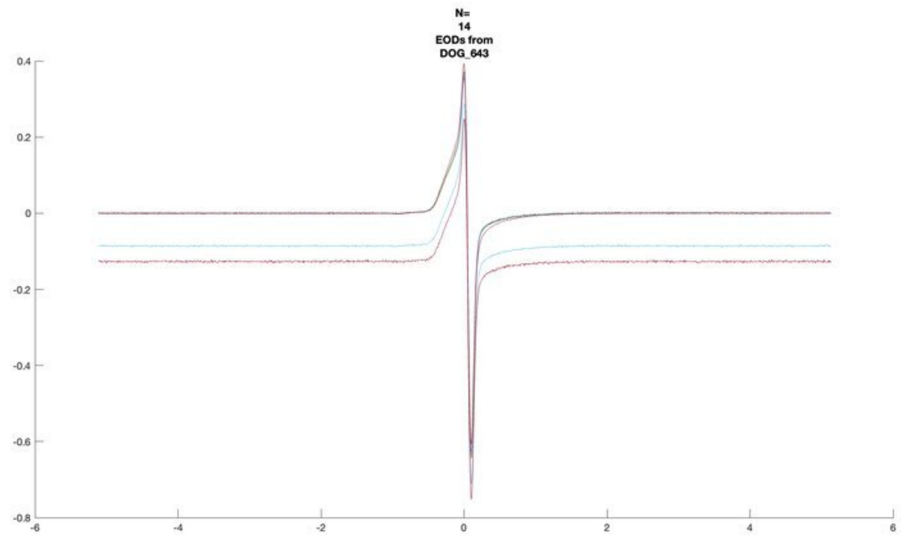
N=
14
EODs from
DOG_643

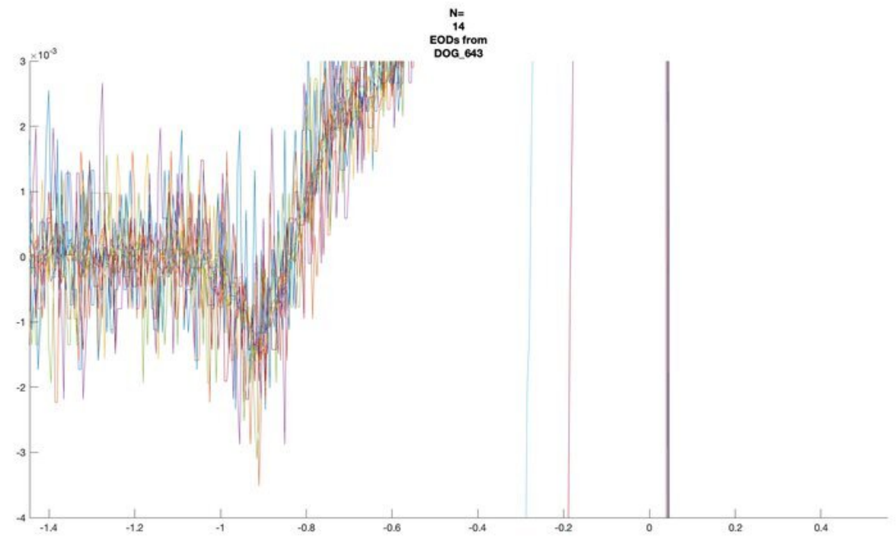
N=
14
EODs from
DOG_643

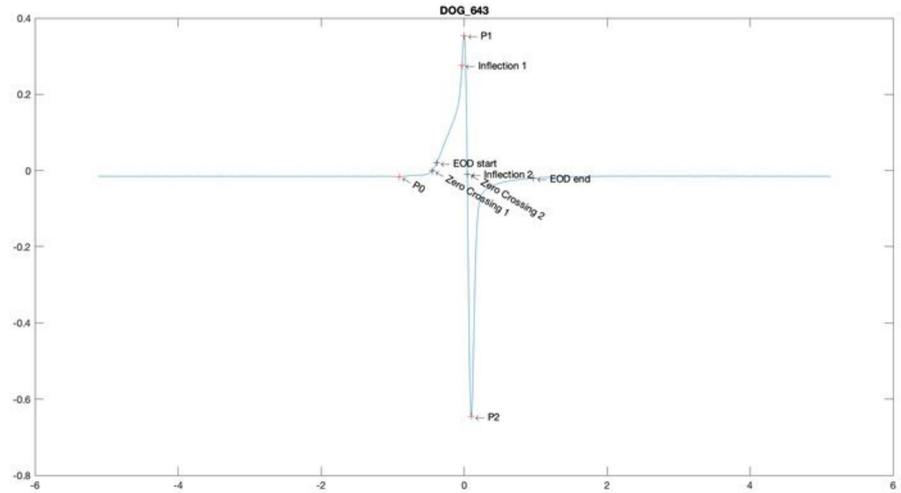
DOG_643
P1
Inflection 1
EOD start
Inflection 2
EOD end
P0
Zero Crossing 1
Zero Crossing 2
P2

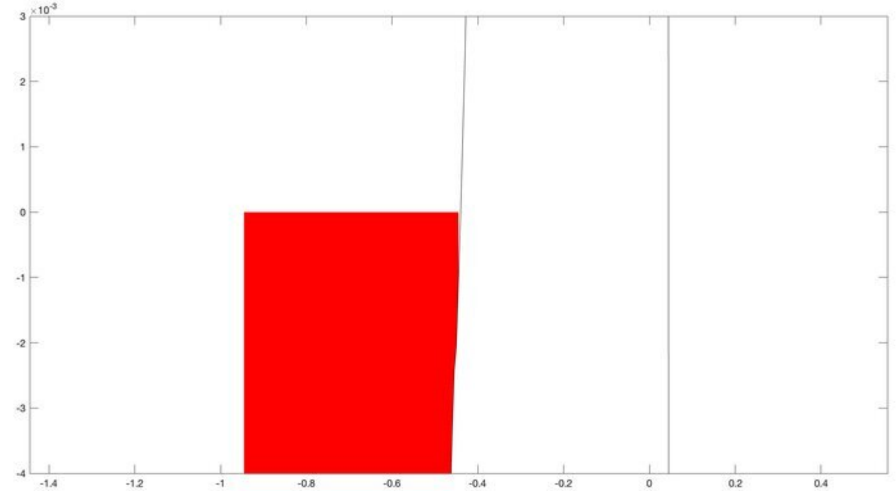

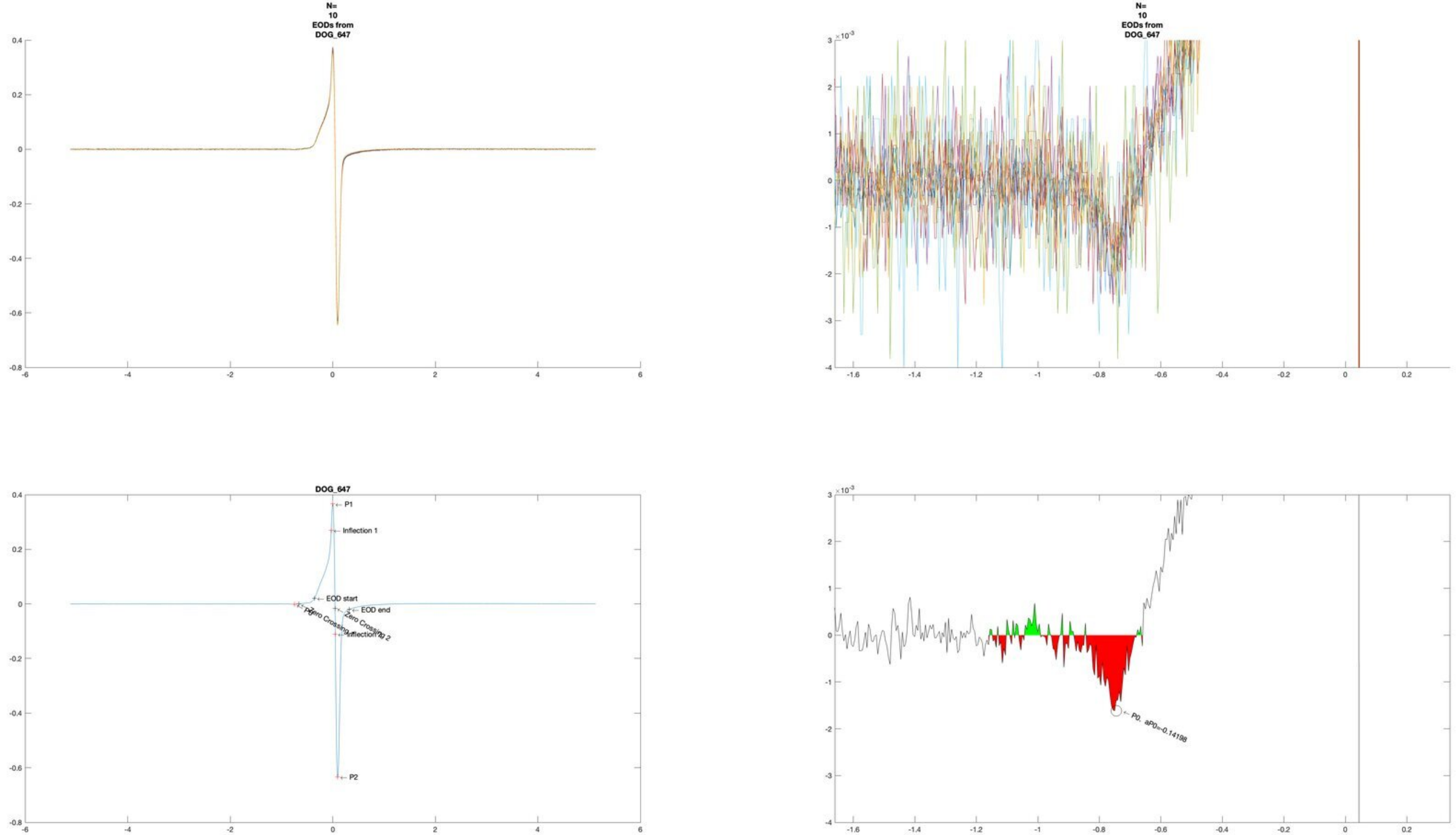

N=
10
EODs from
DOG_647
N=
10
EODs from
DOG_647
DOG_647
P1
Inflection 1
EOD start
EOD end
Zero Crossing 2
P0
Zero Crossing 1
Inflection 2
P2
P0. aP0=-0.14198

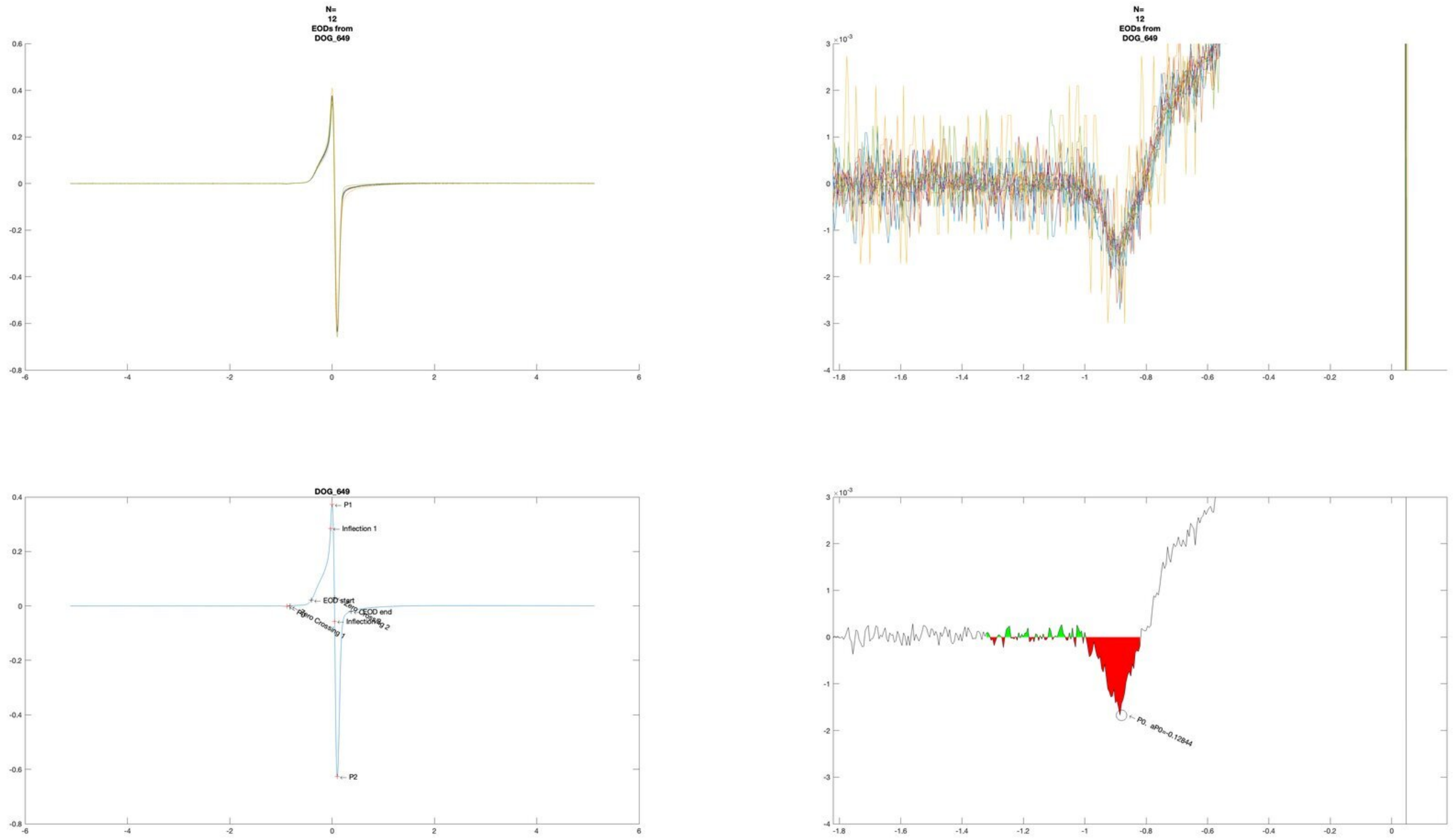
N=
12
EODs from
DOG_649
DOG_649
P1
Inflection 1
EOD start
EOD end
Zero Crossing 1
Zero Crossing 2
P2
P0, aP0=-0.12844

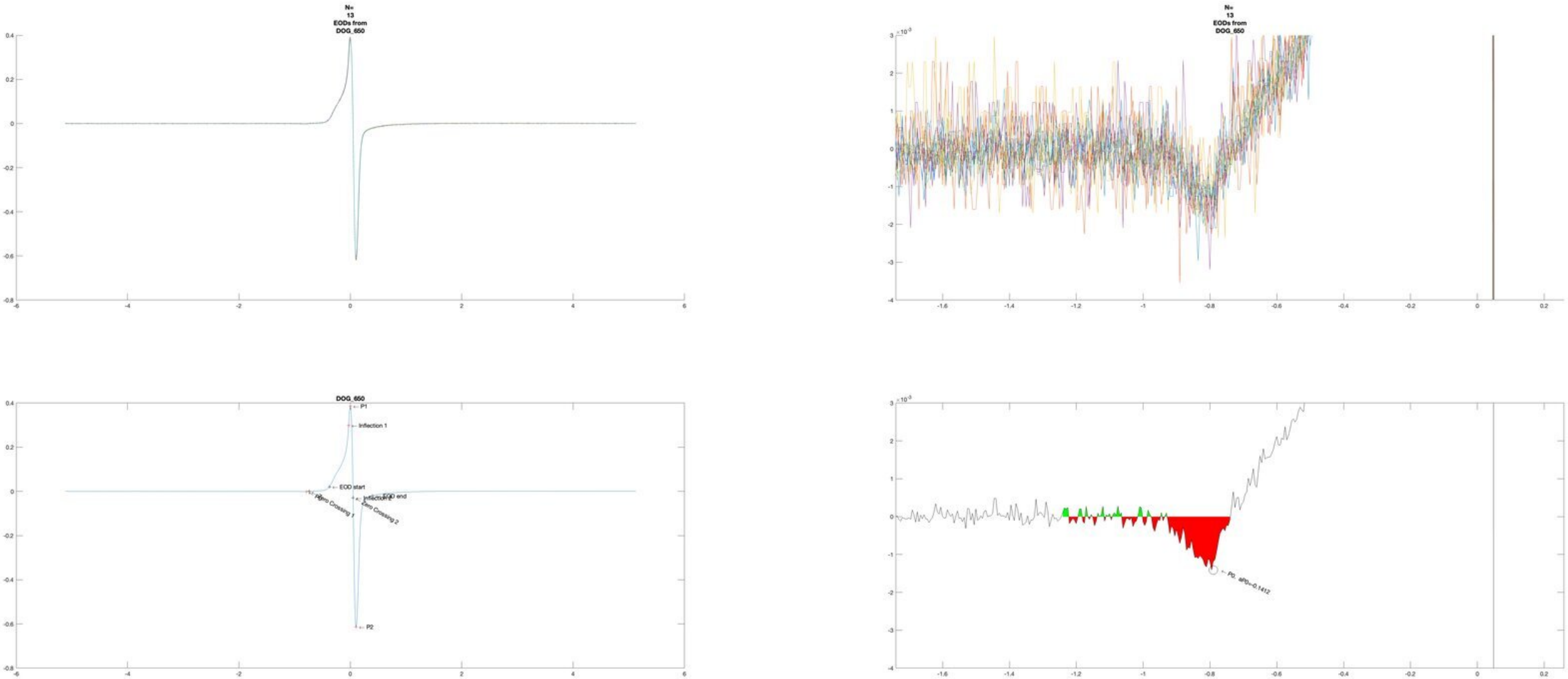
N= 13 EODs from DOG_650
N= 13 EODs from DOG_650
DOG_650
P1
Inflection 1
EOD start
EOD end
Inflection 2
Zero Crossing 1
Zero Crossing 2
P2
P0, aP0=-0.1412

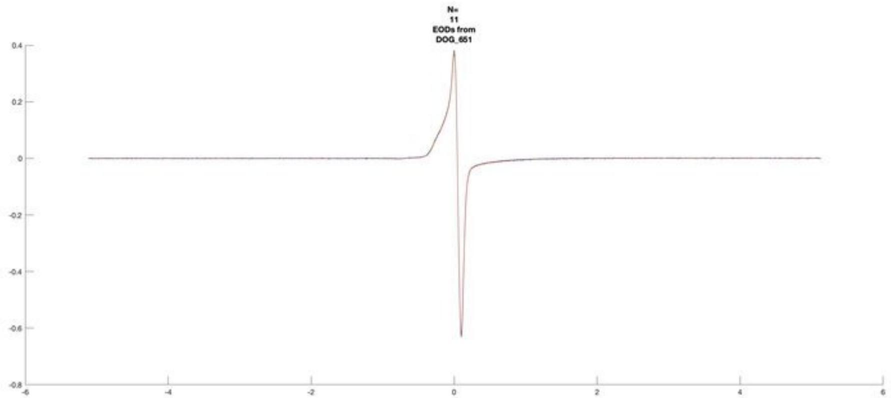
N=
11
EODs from
DOG_651

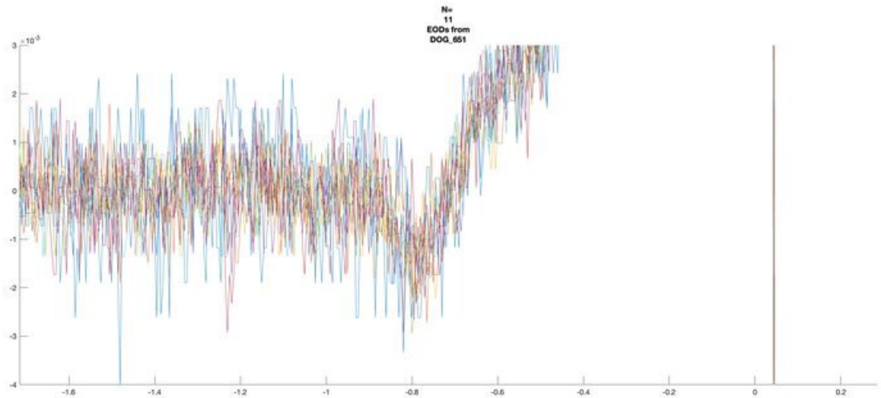
N=
11
EODs from
DOG_651

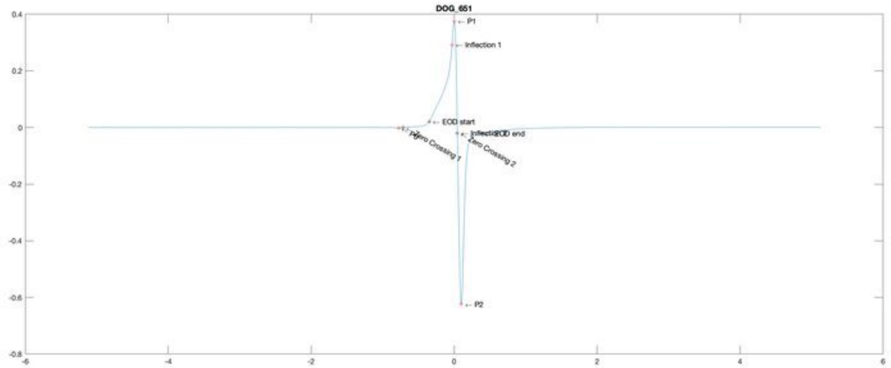
DOG_651
P1
Inflection 1
EOD start
EOD end
Zero Crossing 1
Zero Crossing 2
P2

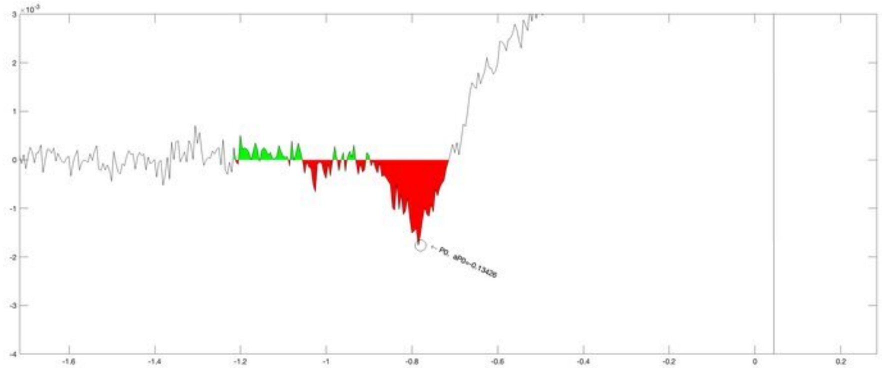
P0, aP0=-0.13426

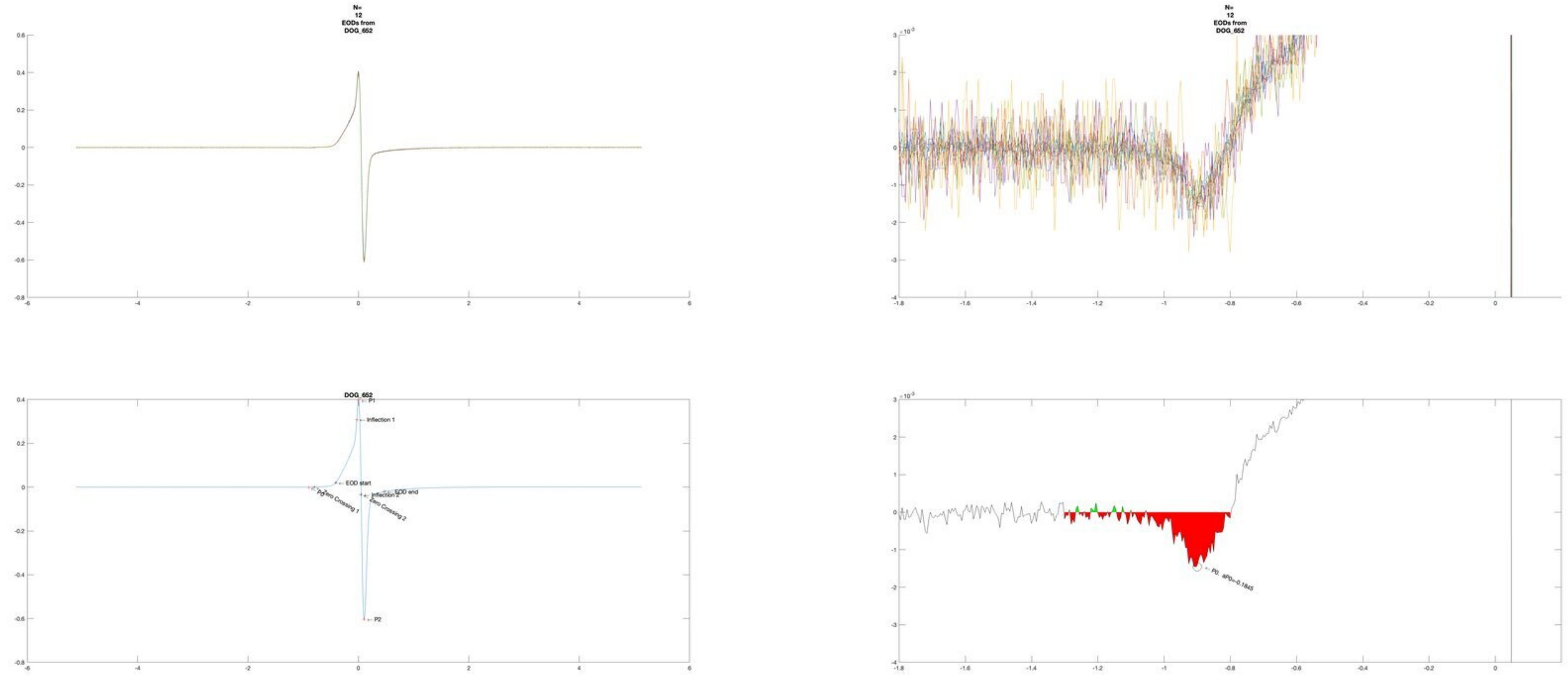

N=
12
EODs from
DOG_652
N=
12
EODs from
DOG_652
DOG_652
P1
Inflection 1
EOD start
EOD end
Inflection 2
Zero Crossing 1
Zero Crossing 2
P0
P2
P0, aP0=-0.1845

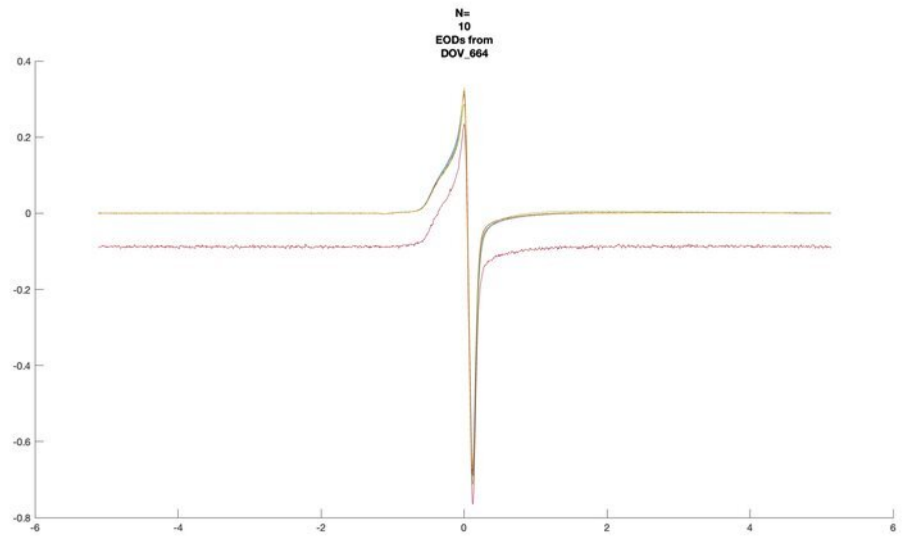
N=
10
EODs from
DOV_664

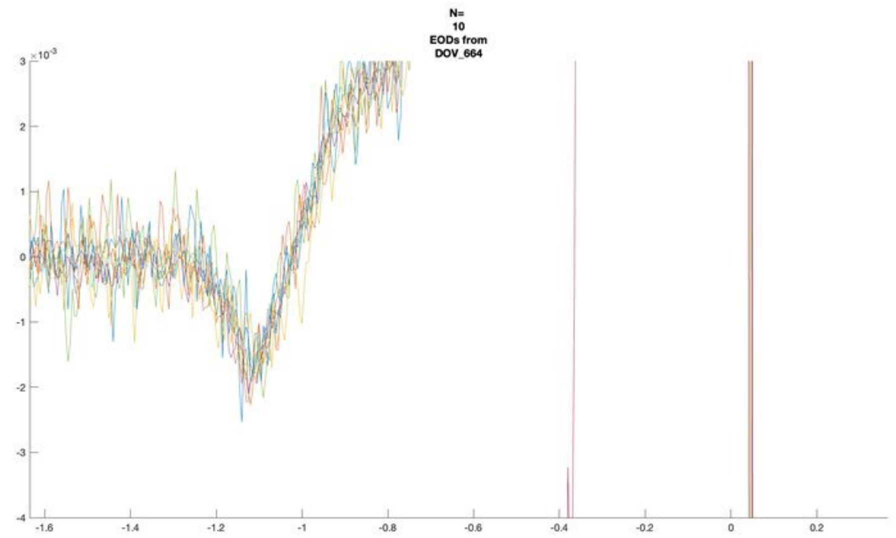
N=
10
EODs from
DOV_664

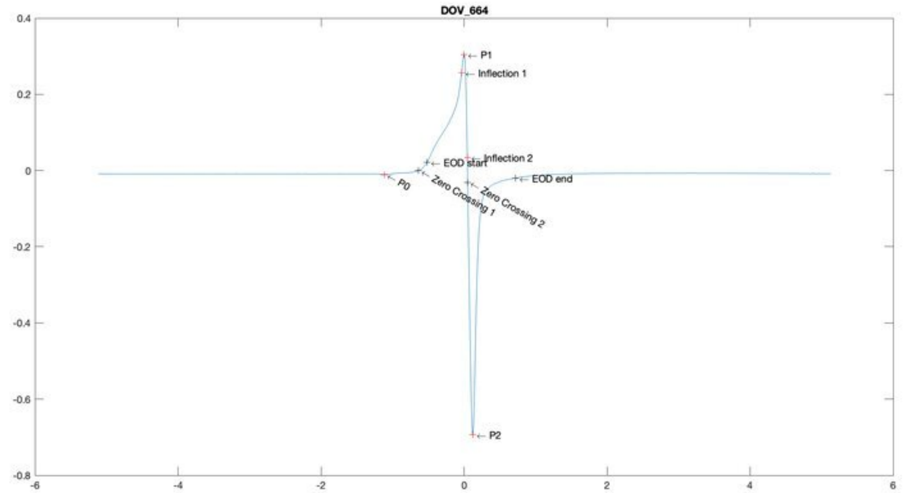
DOV_664
P1
Inflection 1
EOD start
Inflection 2
EOD end
P0
Zero Crossing 1
Zero Crossing 2
P2

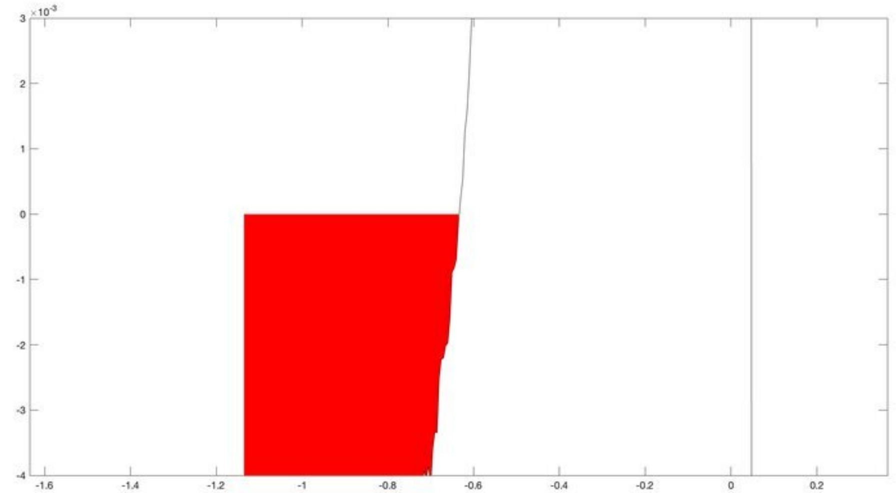

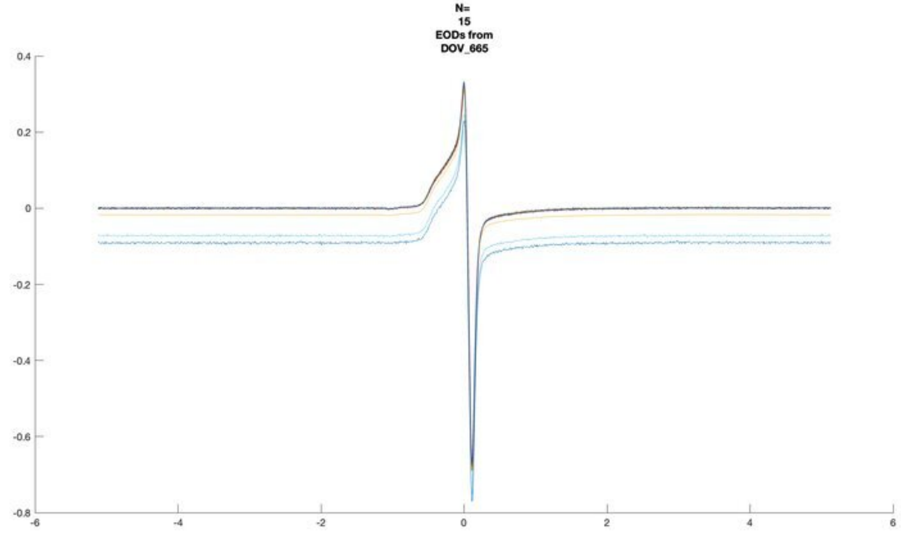
N=
15
EODs from
DOV_665

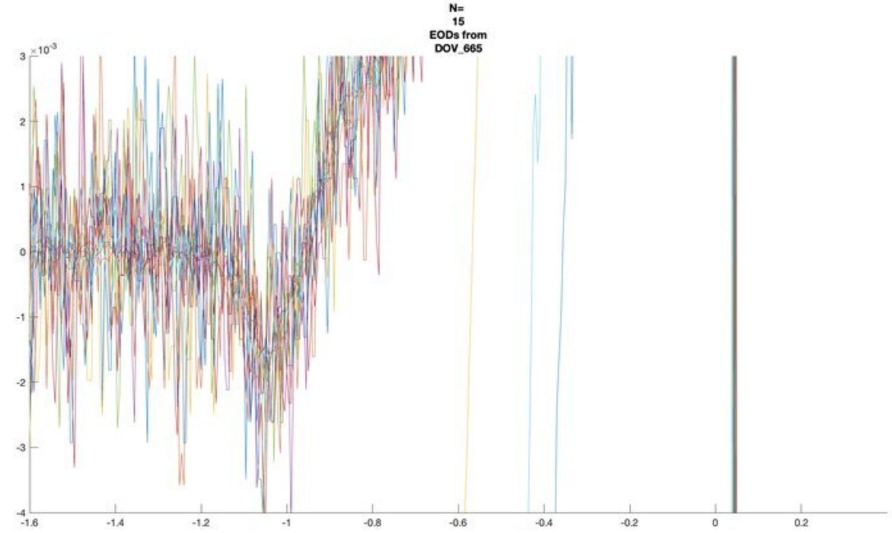
N=
15
EODs from
DOV_665

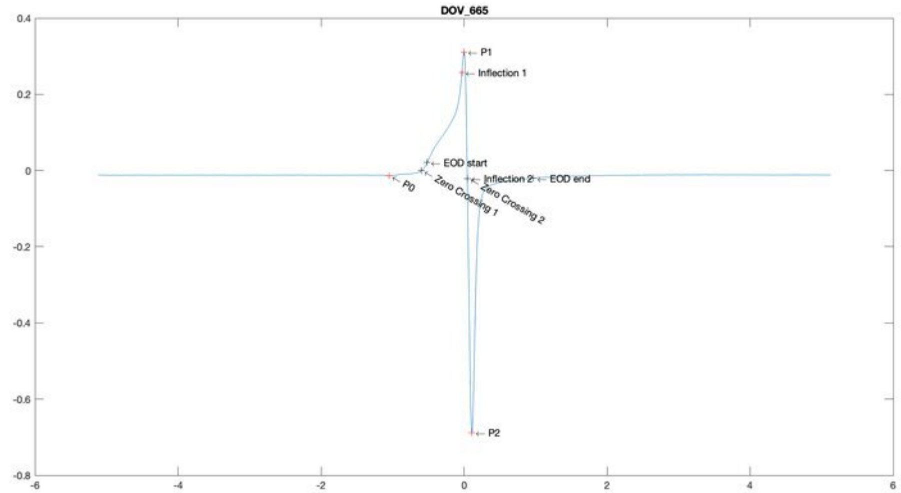
DOV_665
P1
Inflection 1
EOD start
P0
Zero Crossing 1
Inflection 2
EOD end
Zero Crossing 2
P2

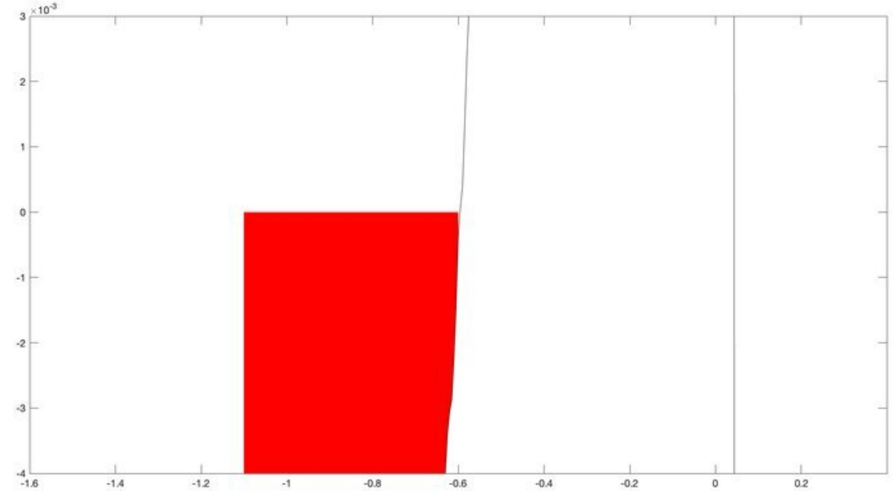

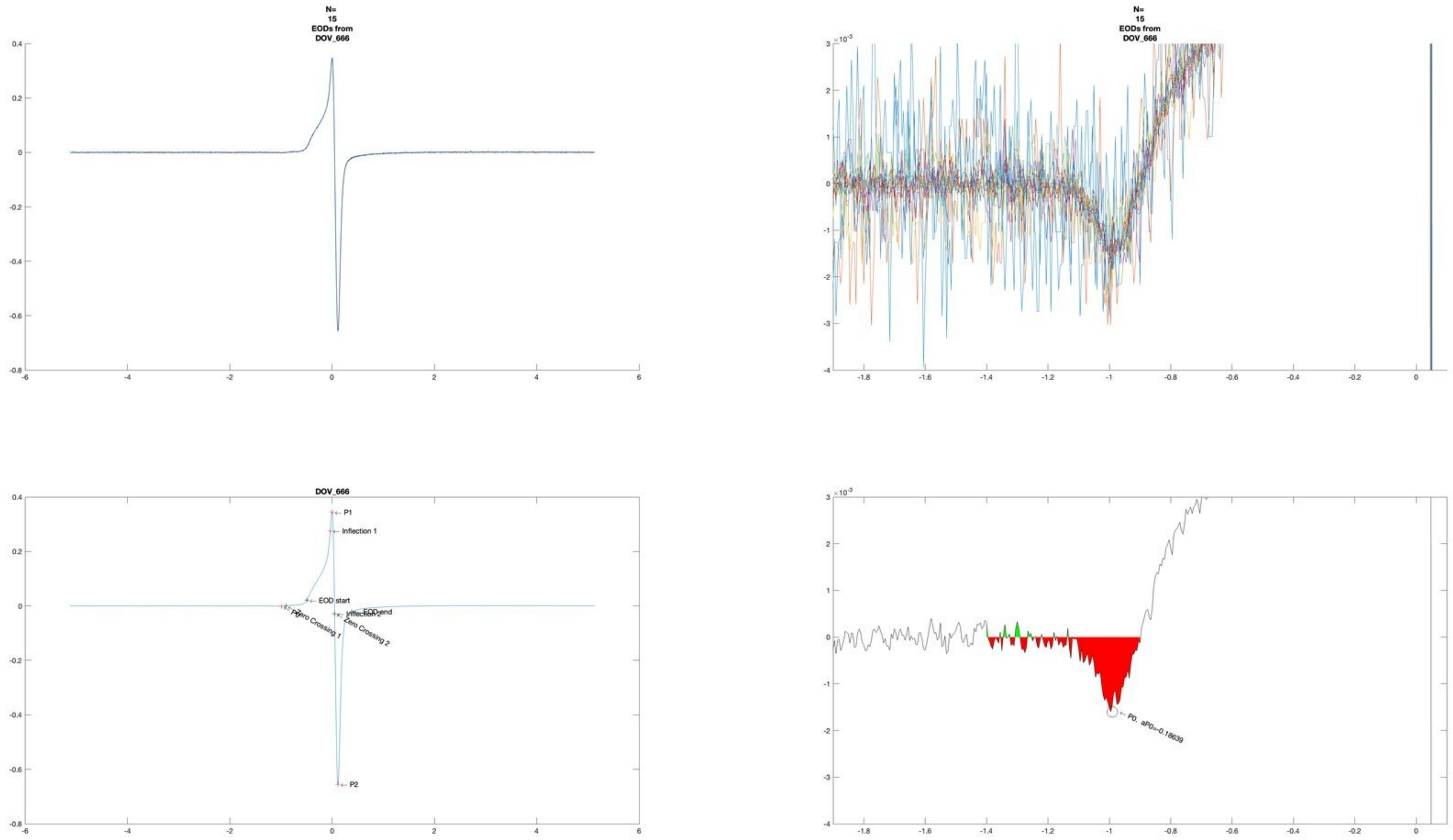
N=
15
EODs from
DOV_666
N=
15
EODs from
DOV_666
DOV_666
P1
Inflection 1
EOD start
EOD end
Inflection 2
P0
Zero Crossing 1
Zero Crossing 2
P2
P0, aP0=-0.18639

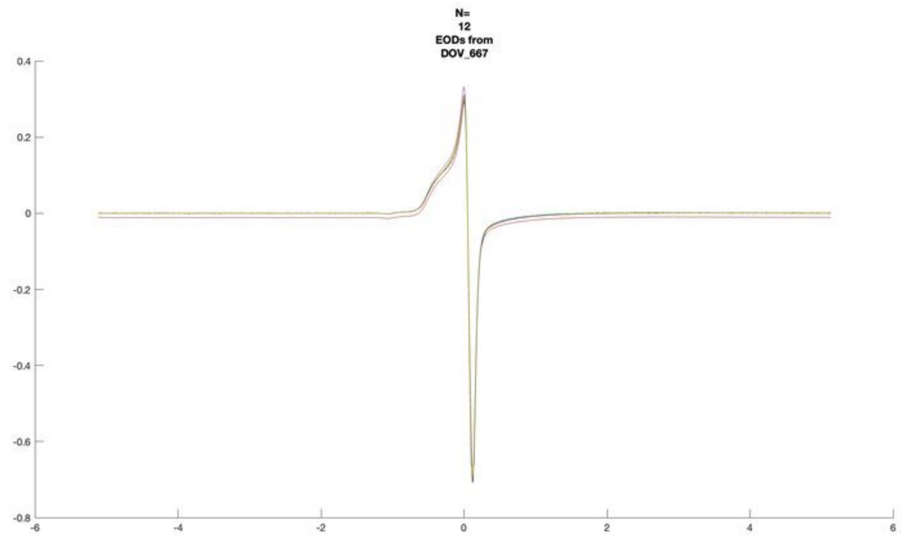
N=
12
EODs from
DOV_667

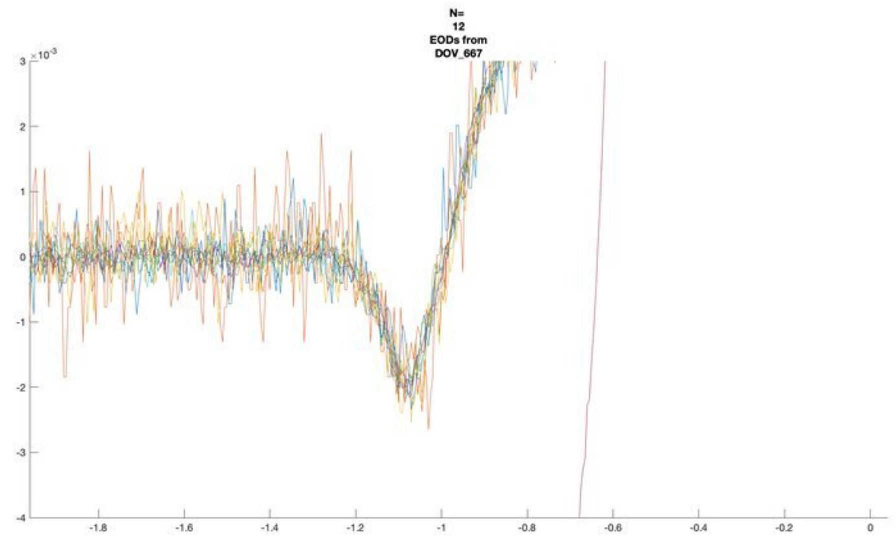
N=
12
EODs from
DOV_667

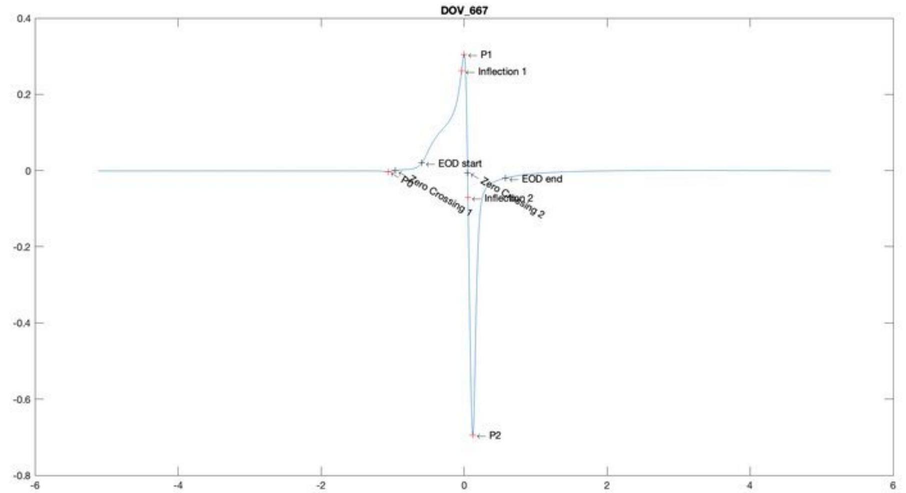
DOV_667
P1
Inflection 1
EOD start
EOD end
P0
Zero Crossing 1
Zero Crossing 2
Inflection 2
P2

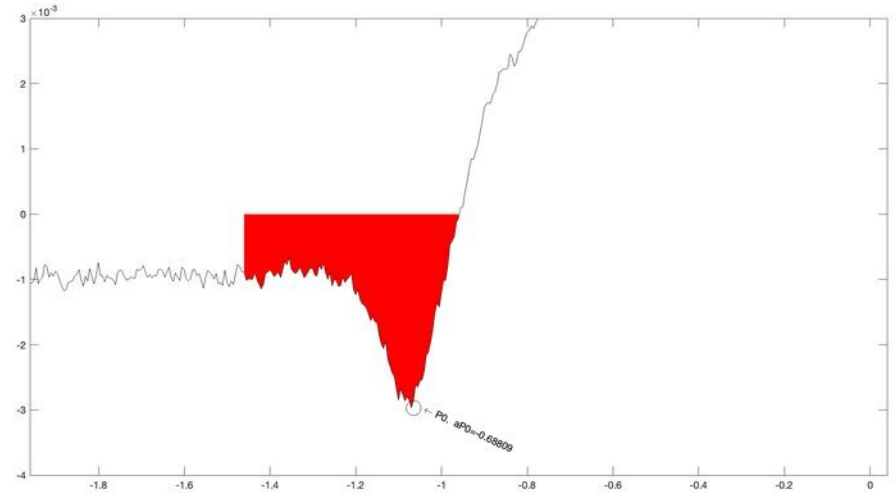
P0, aP0=-0.68809

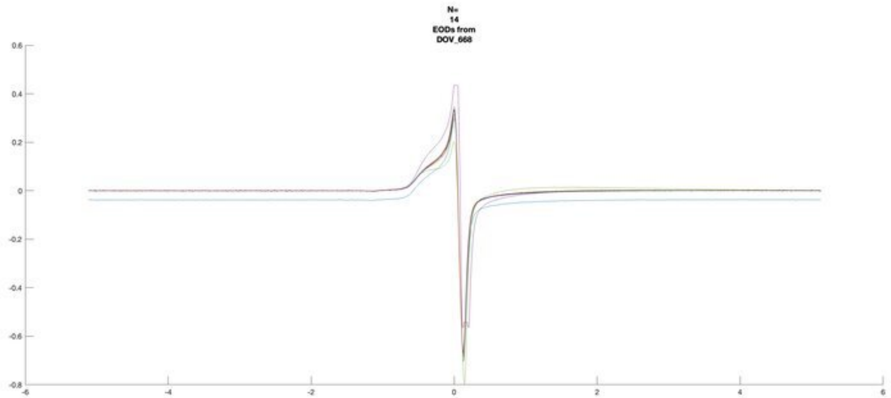
N=
14
EODs from
DOV_668

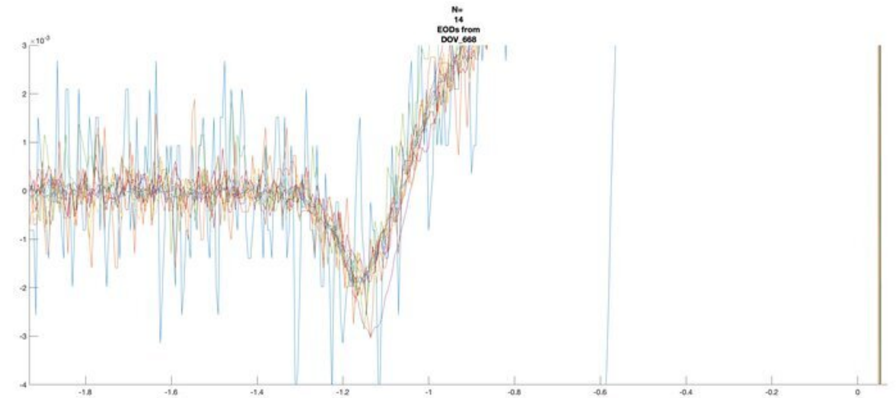
N=
14
EODs from
DOV_668

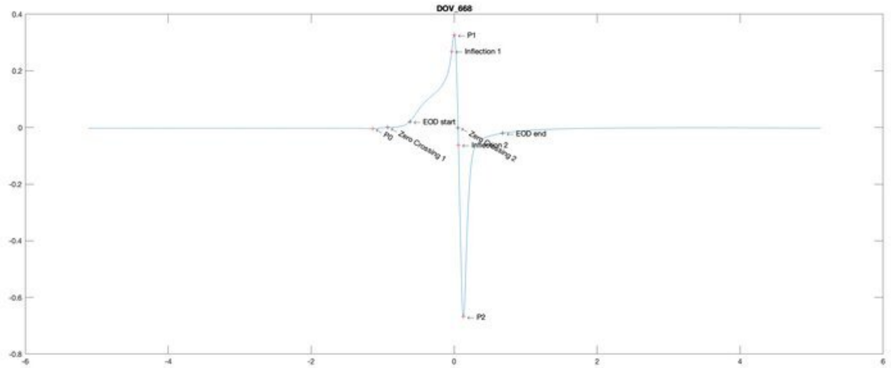
DOV_668
P1
Inflection 1
EOD start
P0
Zero Crossing 1
EOD end
P2

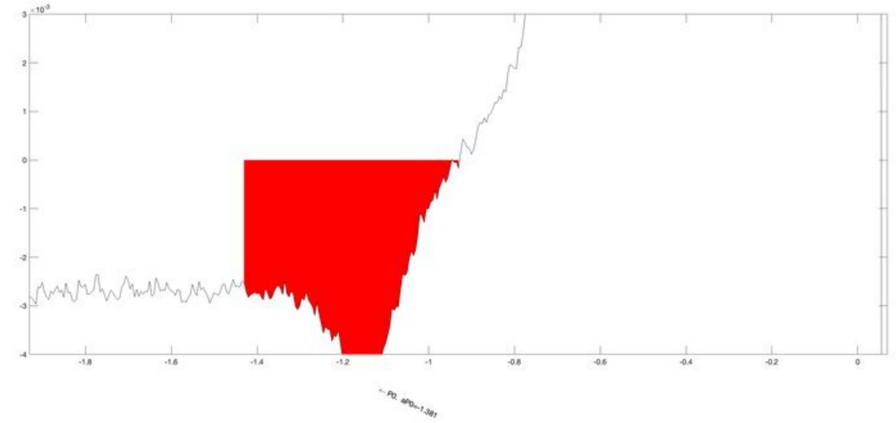
P0, aP0=-1.381

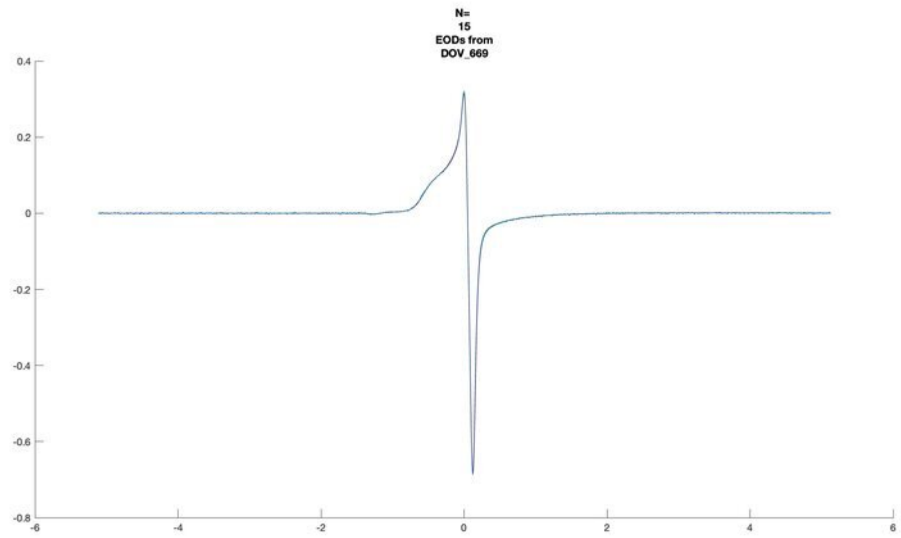
N=
15
EODs from
DOV_669

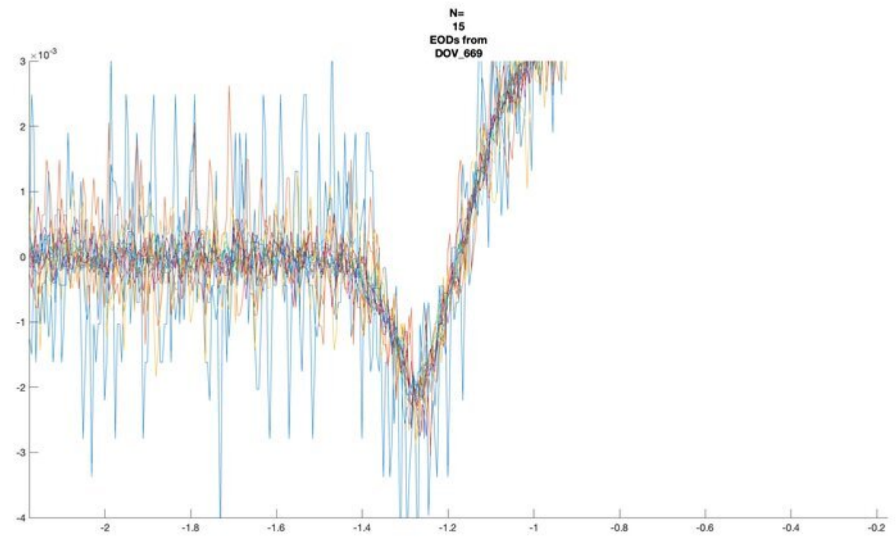
N=
15
EODs from
DOV_669

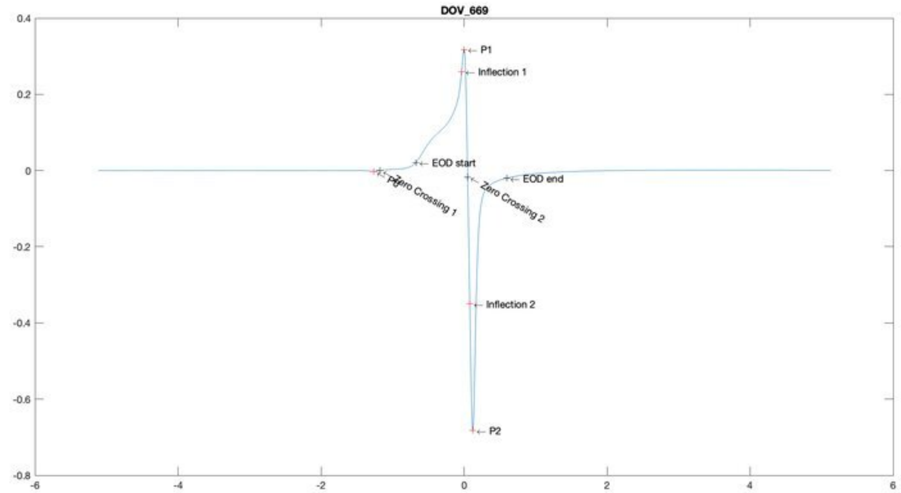
DOV_669
P1
Inflection 1
EOD start
EOD end
Zero Crossing 1
Zero Crossing 2
Inflection 2
P2

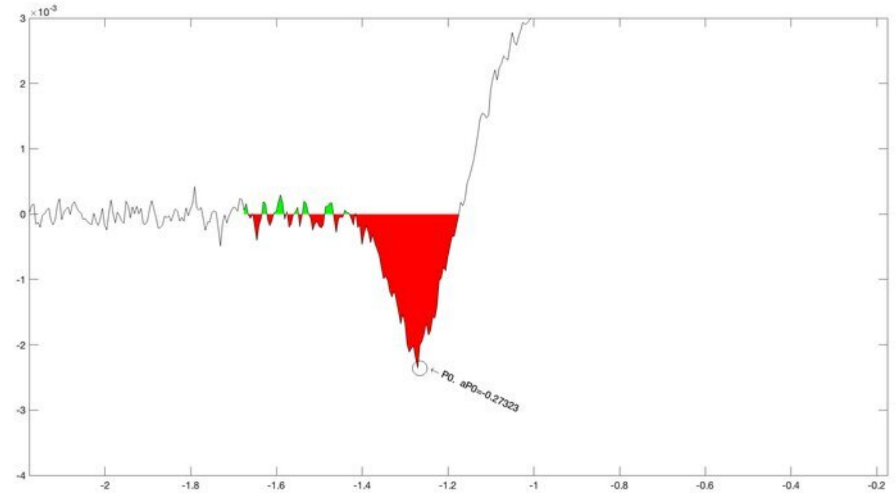
P0, aP0=-0.27323

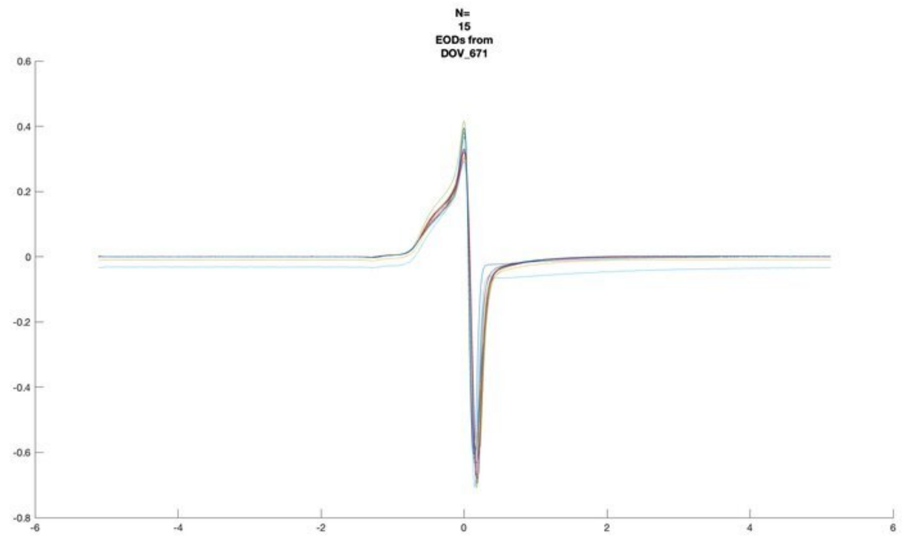
N=
15
EODs from
DOV_671

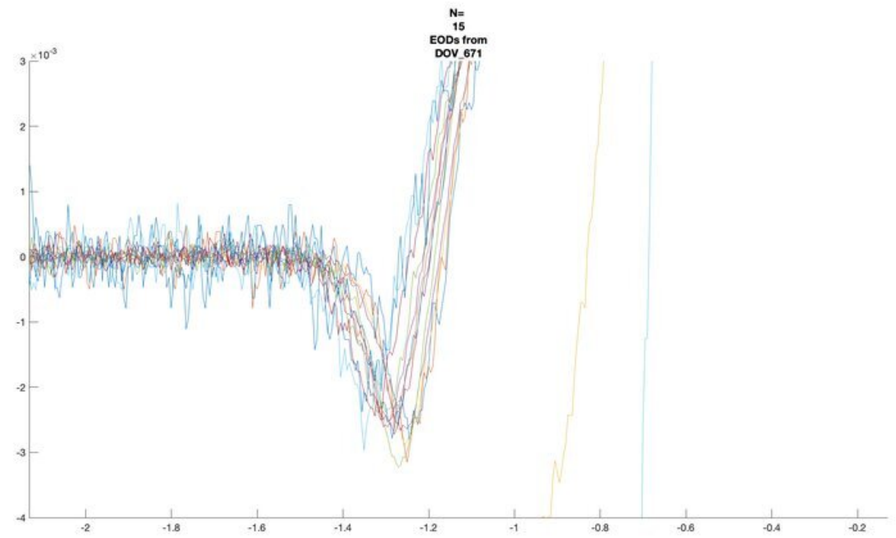
N=
15
EODs from
DOV_671

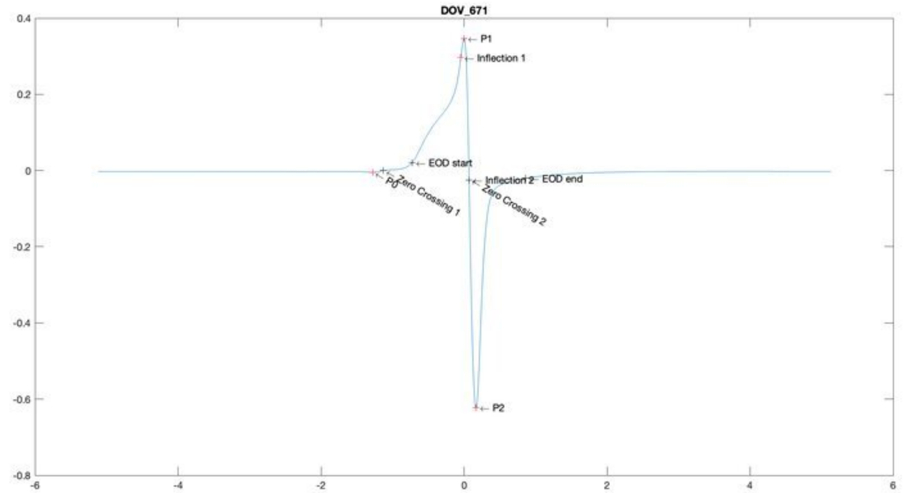
DOV_671
P1
Inflection 1
EOD start
P0
Zero Crossing 1
Inflection 2
EOD end
Zero Crossing 2
P2

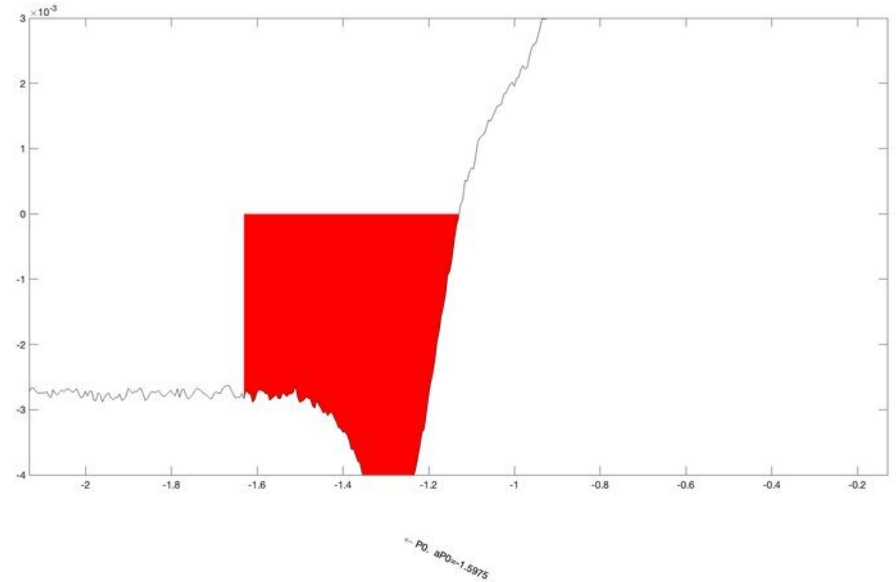
← P0, aP0=-1.5975

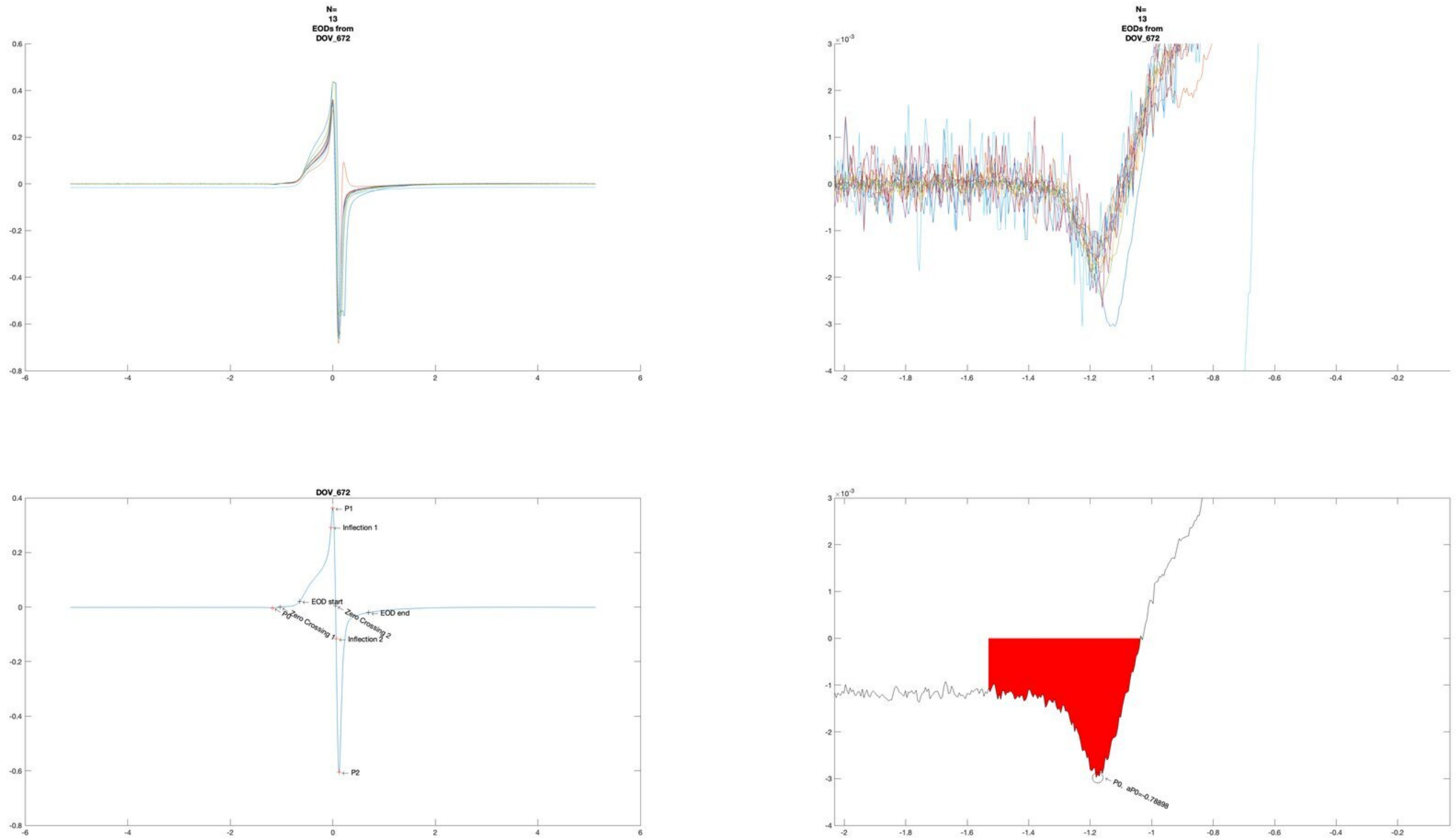
N=
13
EODs from
DOV_672
N=
13
EODs from
DOV_672
DOV_672
P1
Inflection 1
EOD start
P0
Zero Crossing 1
Zero Crossing 2
EOD end
Inflection 2
P2
P0. aP0=-0.78898

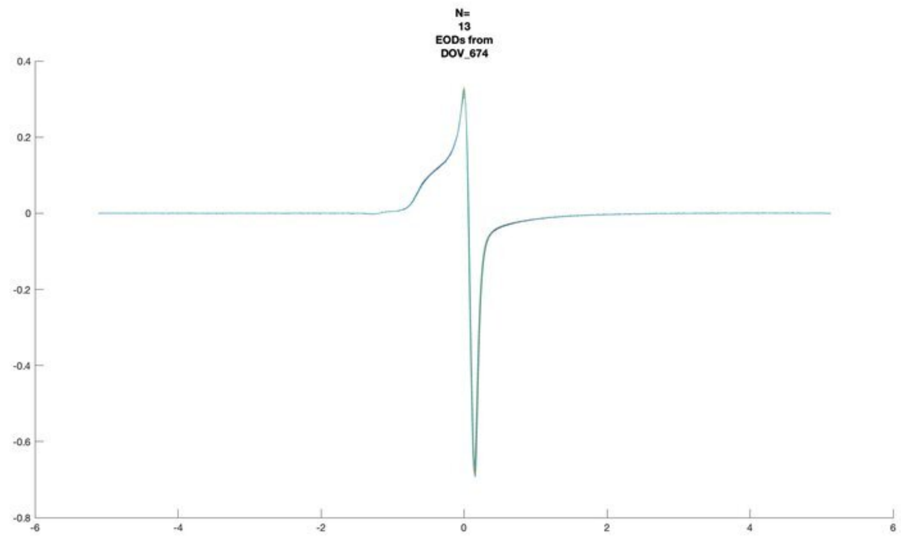
N=
13
EODs from
DOV_674

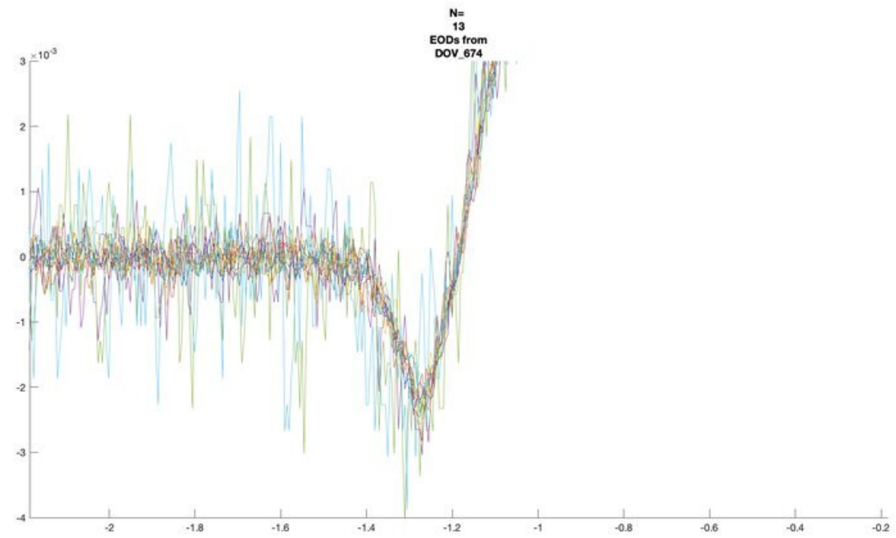
N=
13
EODs from
DOV_674
×10^-3

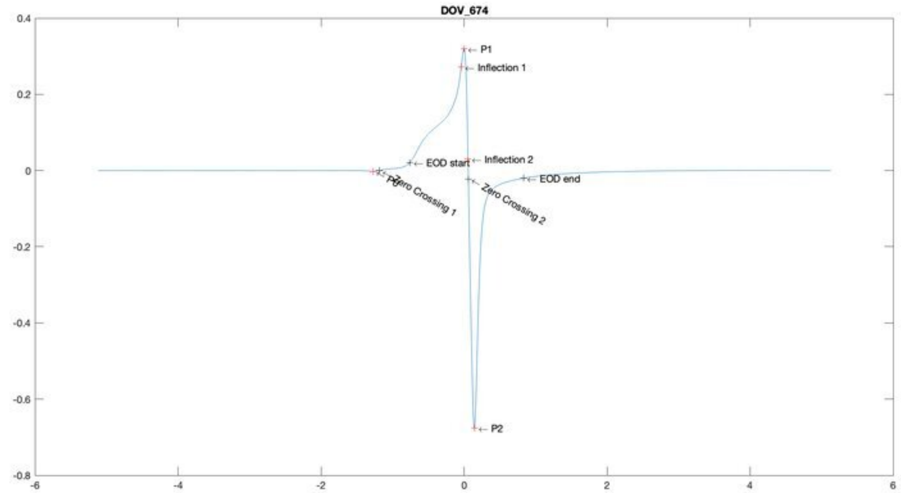
DOV_674
P1
Inflection 1
EOD start
Inflection 2
EOD end
P0
Zero Crossing 1
Zero Crossing 2
P2

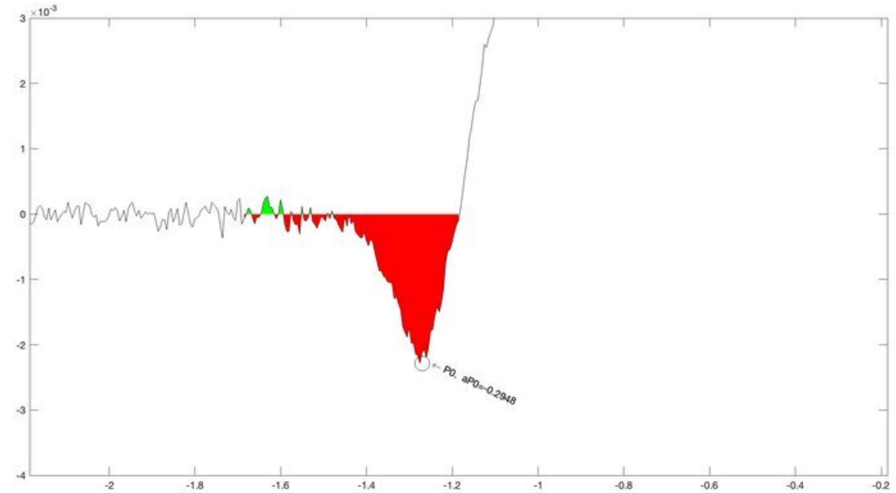
×10^-3
P0, aP0=-0.2948

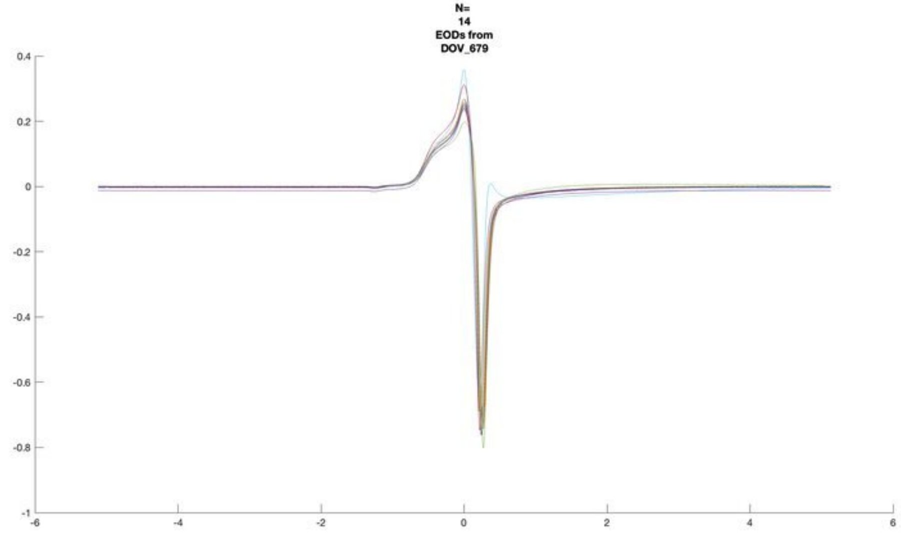
N=
14
EODs from
DOV_679

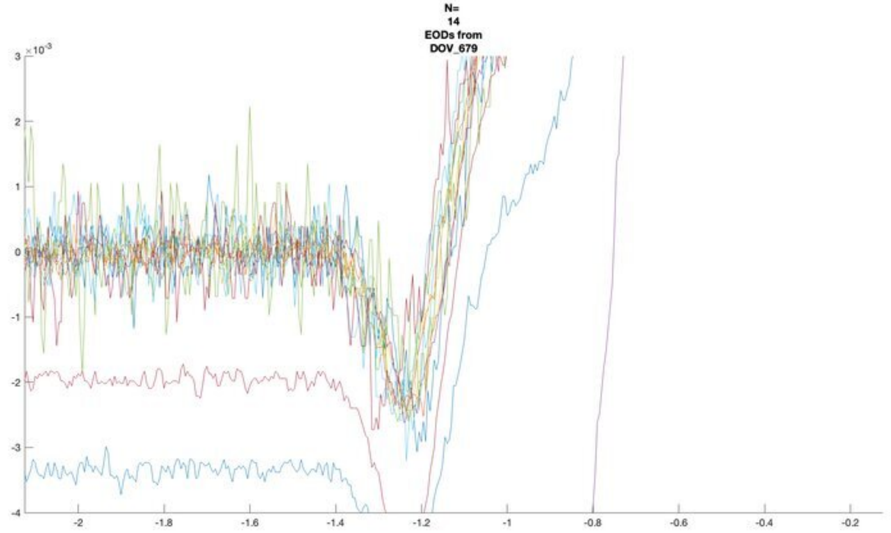
N=
14
EODs from
DOV_679

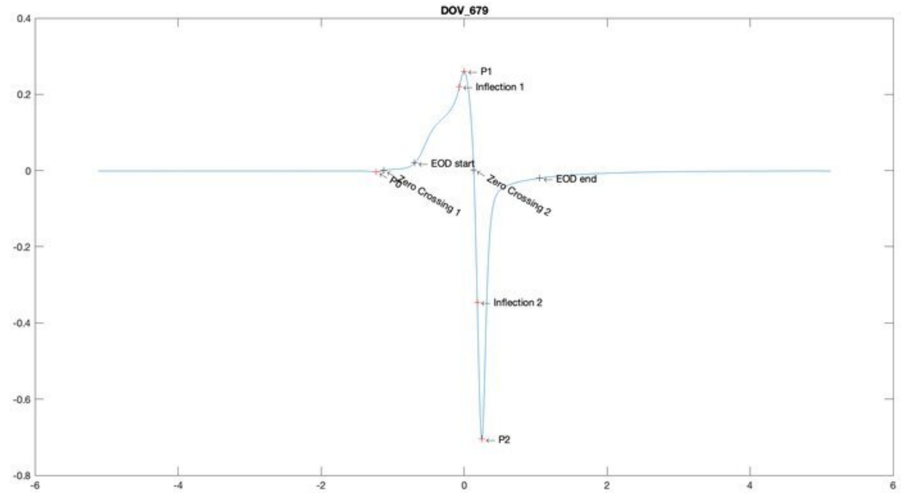
DOV_679
P1
Inflection 1
EOD start
P0
Zero Crossing 1
Zero Crossing 2
EOD end
Inflection 2
P2

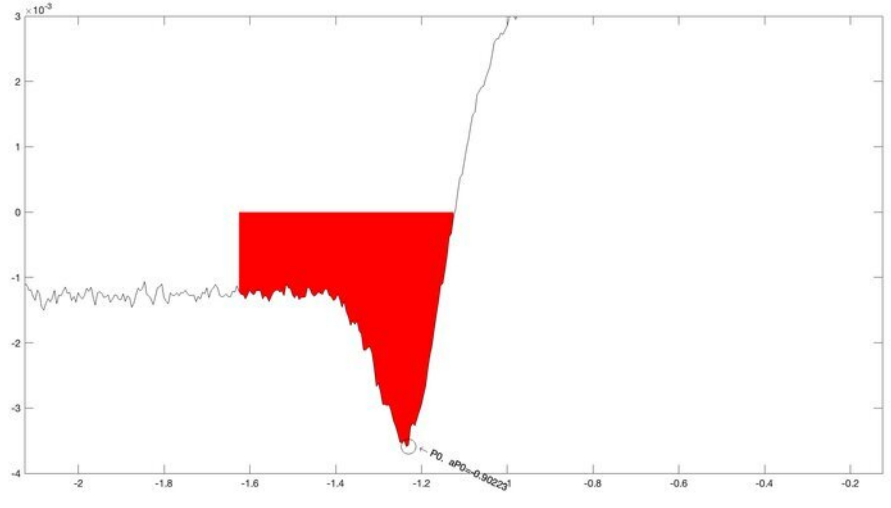
P0, aP0=-0.90223

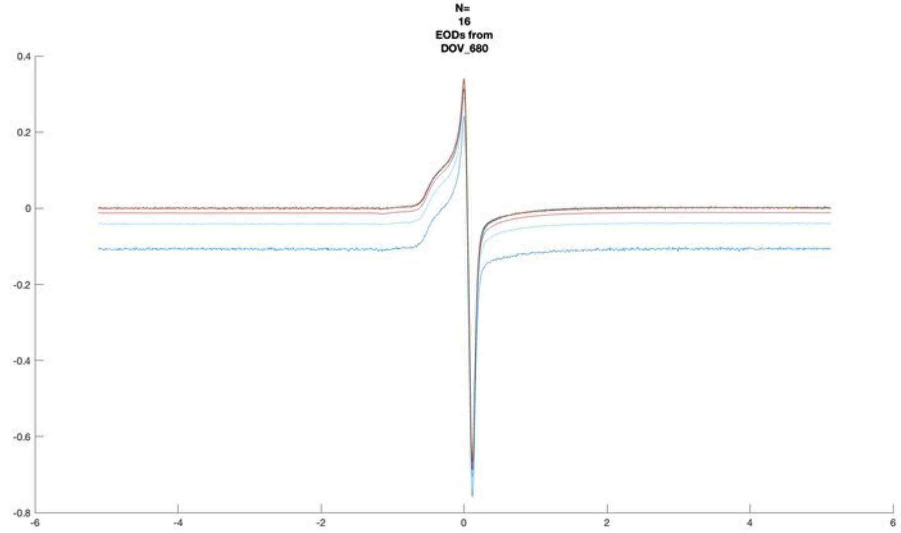
N=
16
EODs from
DOV_680
0.4
0.2
0
-0.2
-0.4
-0.6
-0.8
-6
-4
-2
0
2
4
6

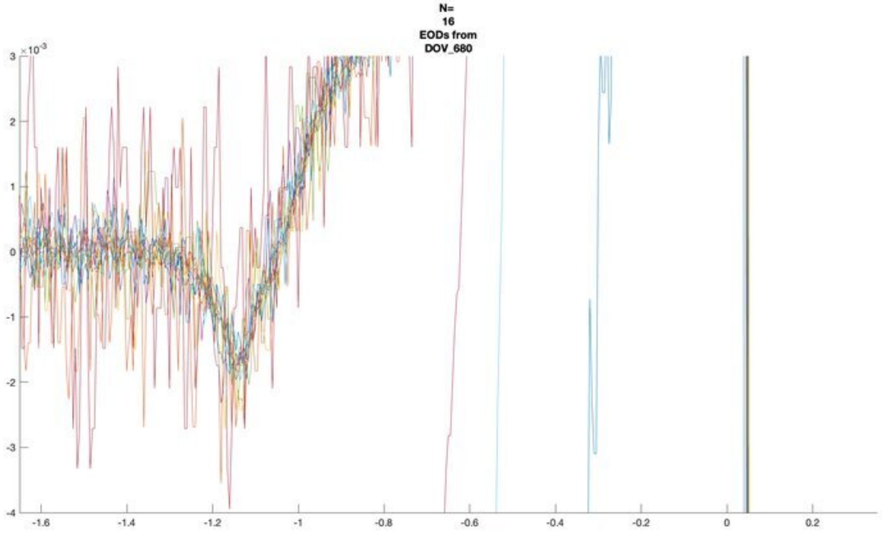
N=
16
EODs from
DOV_680
×10^-3
3
2
1
0
-1
-2
-3
-4
-1.6
-1.4
-1.2
-1
-0.8
-0.6
-0.4
-0.2
0
0.2

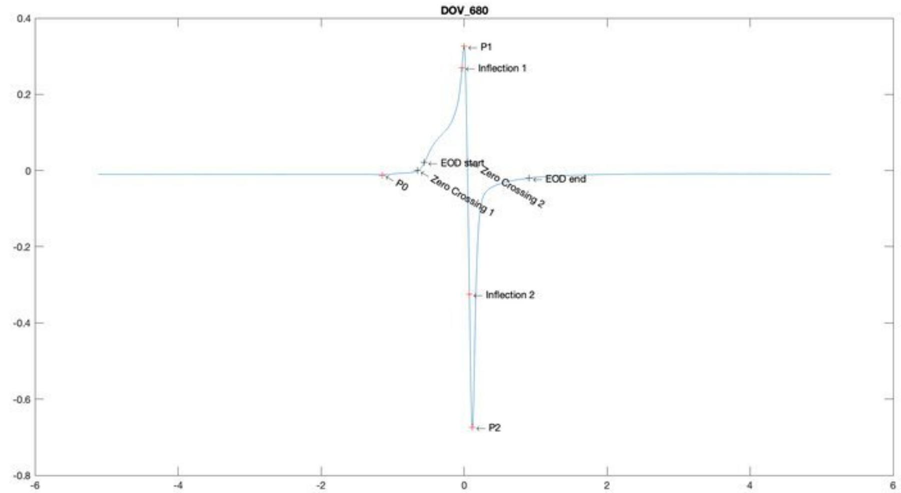
DOV_680
P1
Inflection 1
EOD start
Zero Crossing 2
EOD end
P0
Zero Crossing 1
Inflection 2
P2
0.4
0.2
0
-0.2
-0.4
-0.6
-0.8
-6
-4
-2
0
2
4
6

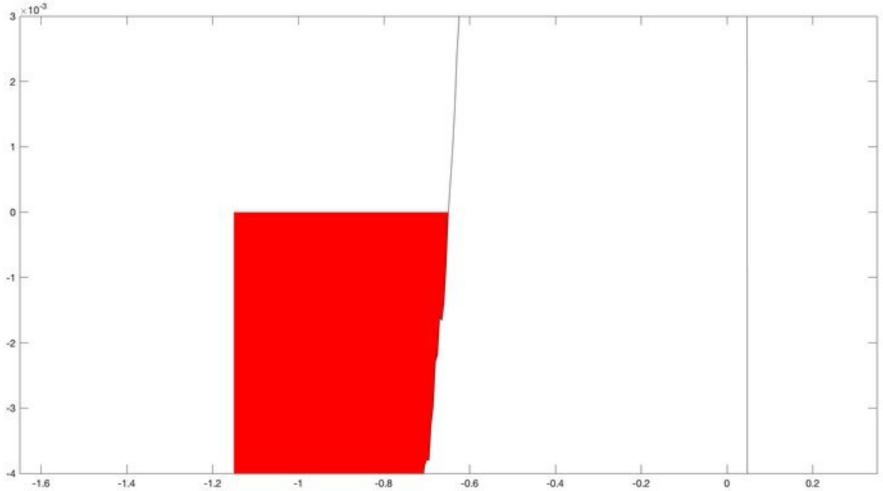
×10^-3
3
2
1
0
-1
-2
-3
-4
-1.6
-1.4
-1.2
-1
-0.8
-0.6
-0.4
-0.2
0
0.2

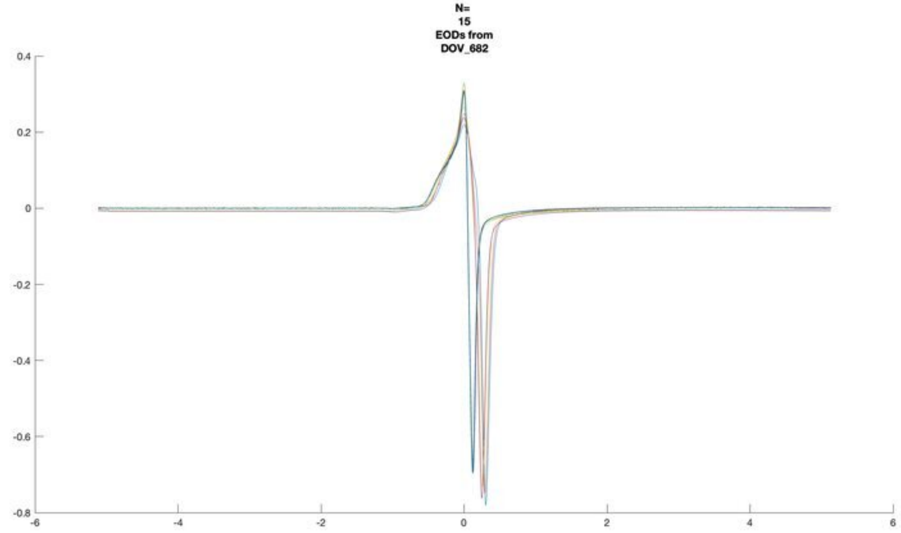
N=
15
EODs from
DOV_682

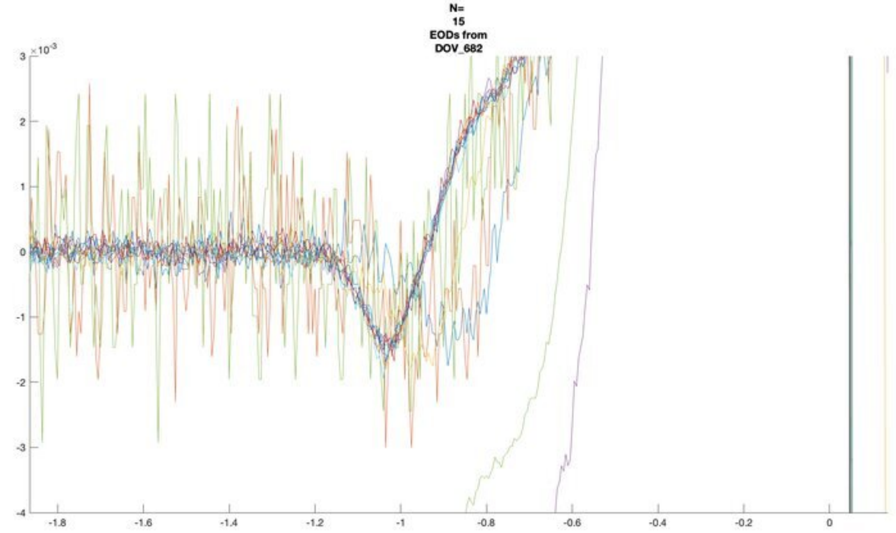
N=
15
EODs from
DOV_682

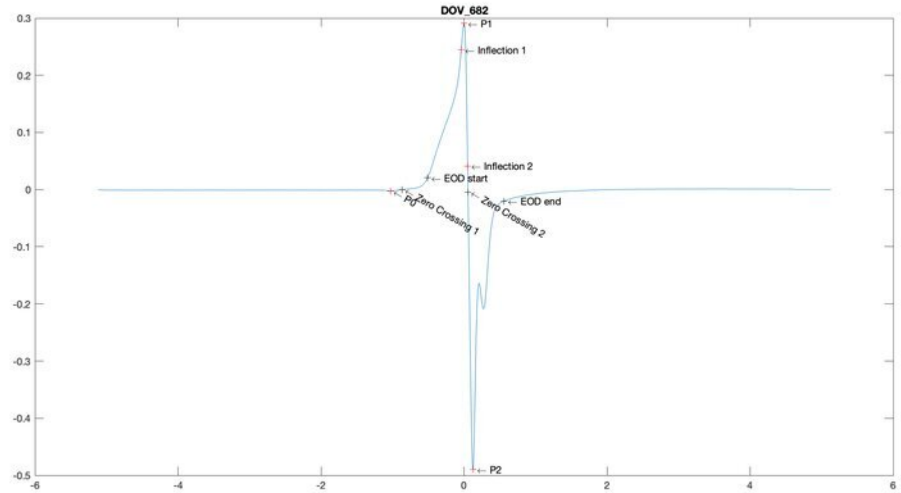
DOV_682
P1
Inflection 1
Inflection 2
EOD start
EOD end
P0
Zero Crossing 1
Zero Crossing 2
P2

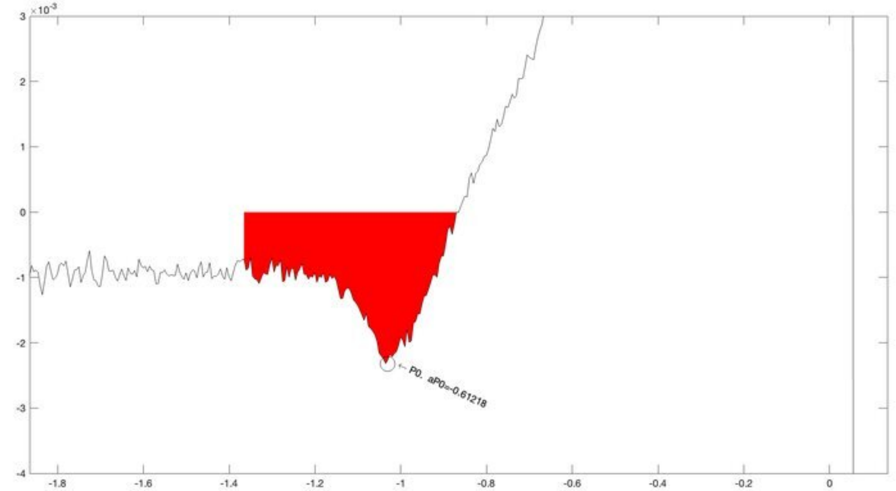
P0, aP0=-0.61218

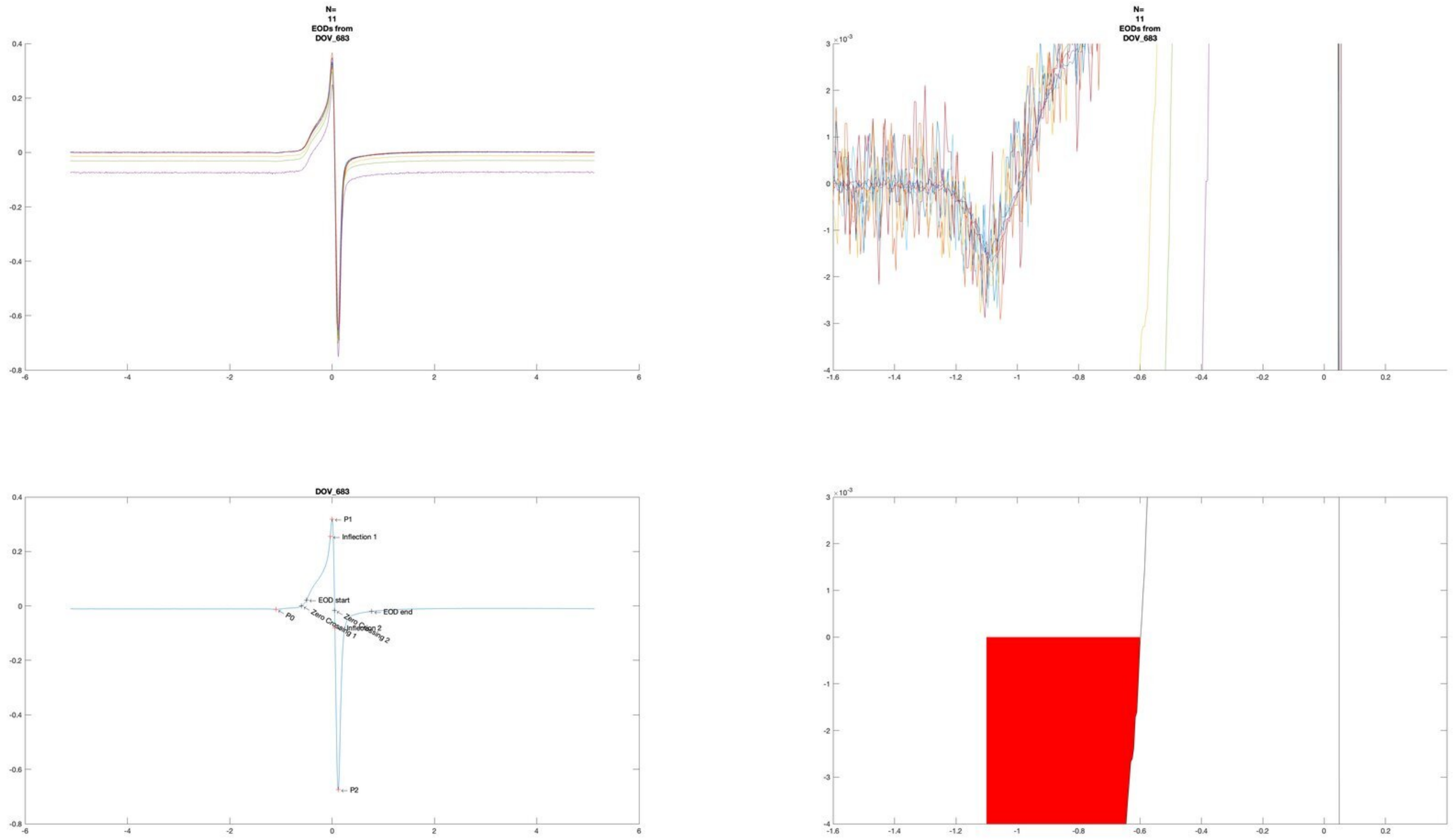
N=
11
EODs from
DOV_683
N=
11
EODs from
DOV_683
DOV_683
P1
Inflection 1
EOD start
EOD end
P0
Zero Crossing 1
Zero Crossing 2
Inflection 2
P2

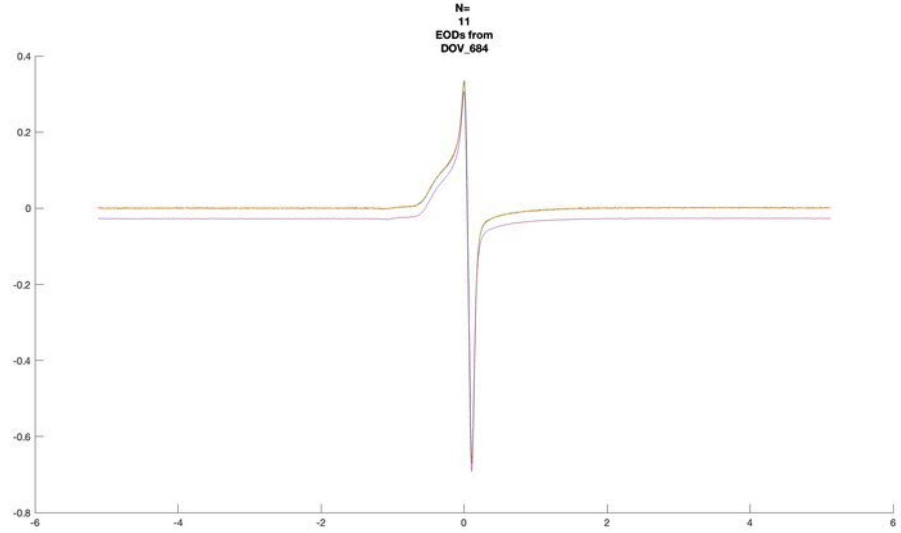
N=
11
EODs from
DOV_684

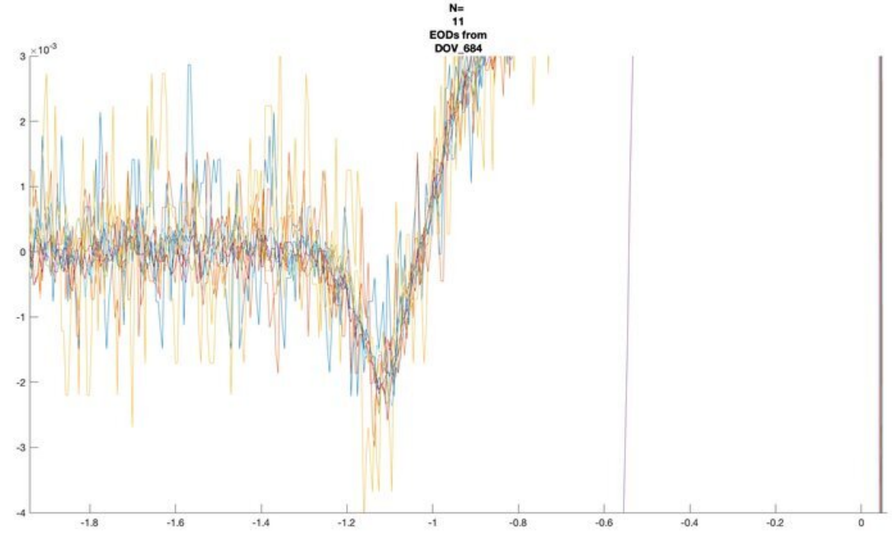
N=
11
EODs from
DOV_684

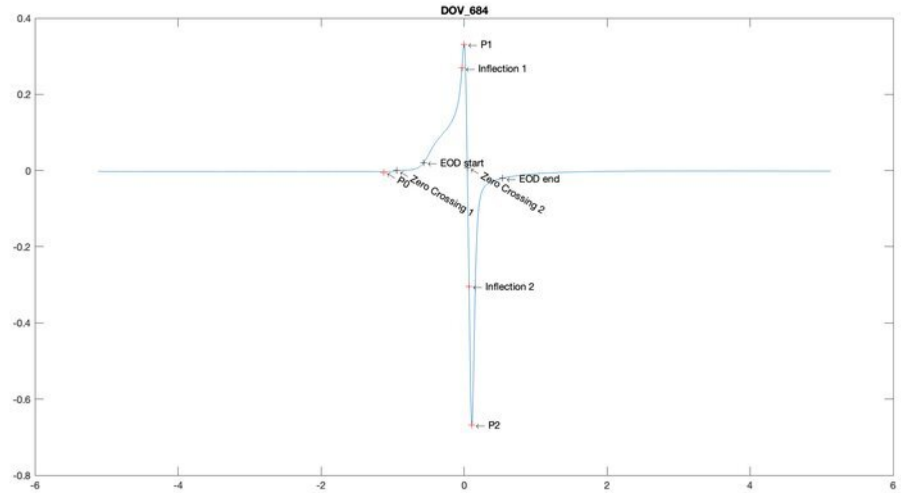
DOV_684
P1
Inflection 1
EOD start
P0
Zero Crossing 1
Zero Crossing 2
EOD end
Inflection 2
P2

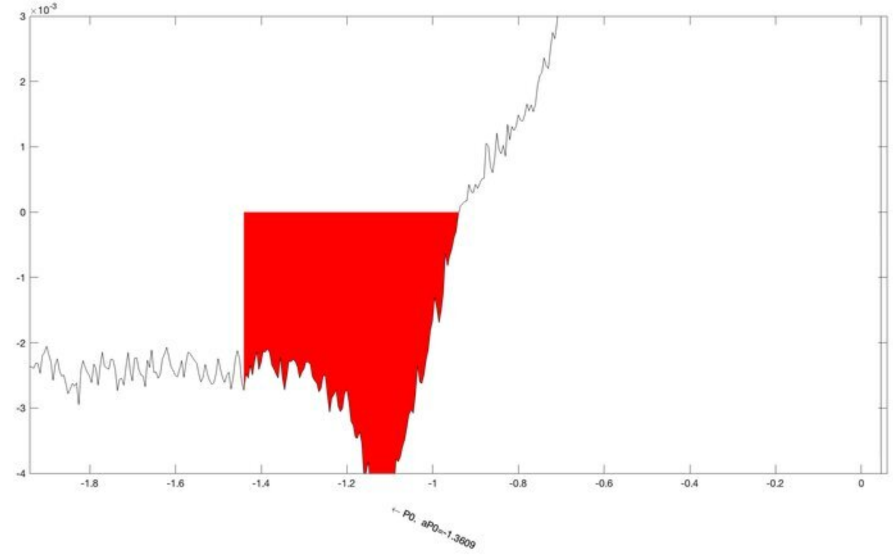
← P0, aP0=-1.3609

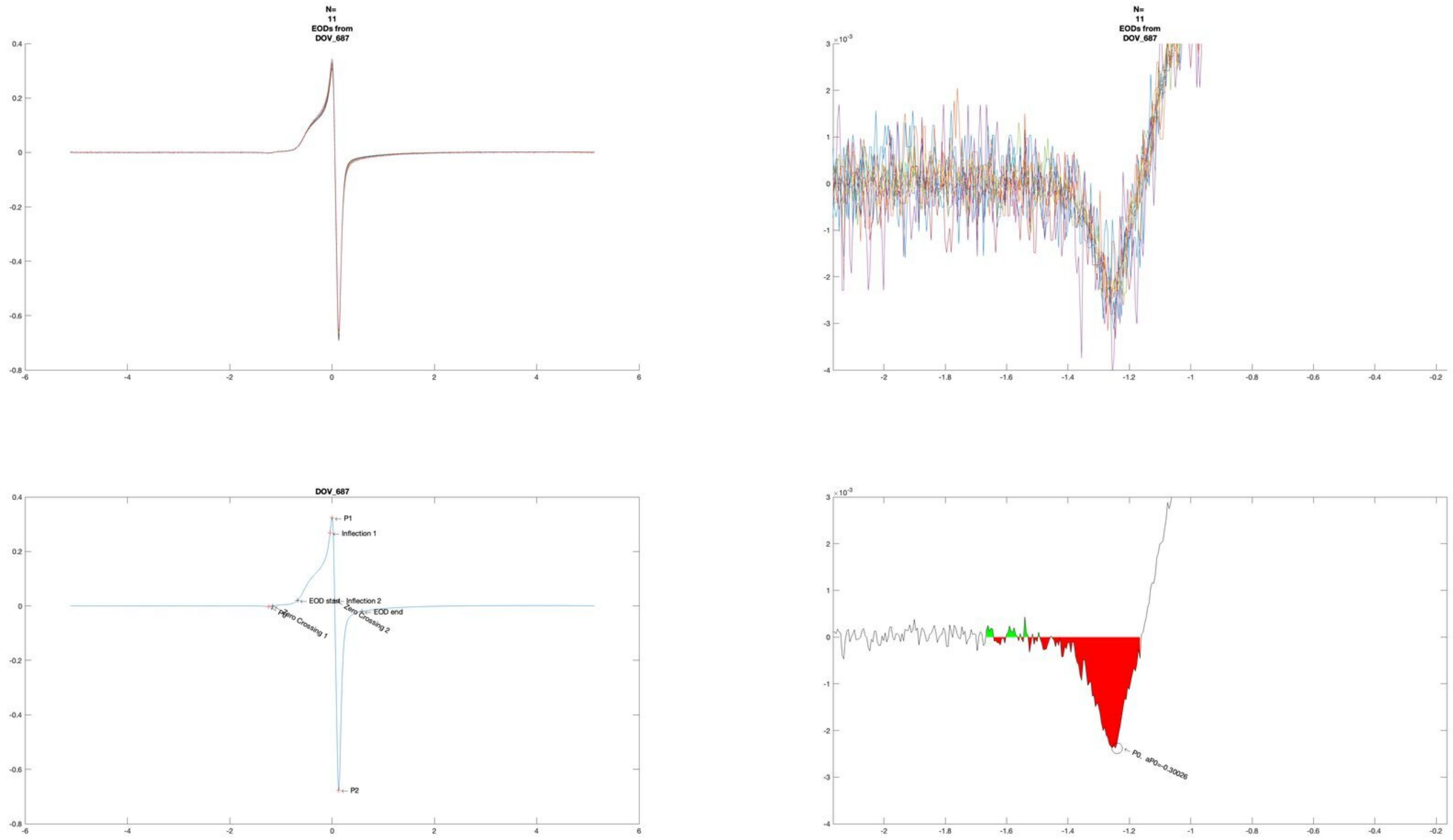
N=
11
EODs from
DOV_687
N=
11
EODs from
DOV_687
DOV_687
P1
Inflection 1
EOD start
Inflection 2
EOD end
P0
Zero Crossing 1
Zero Crossing 2
P2
P0, aP0=-0.30026

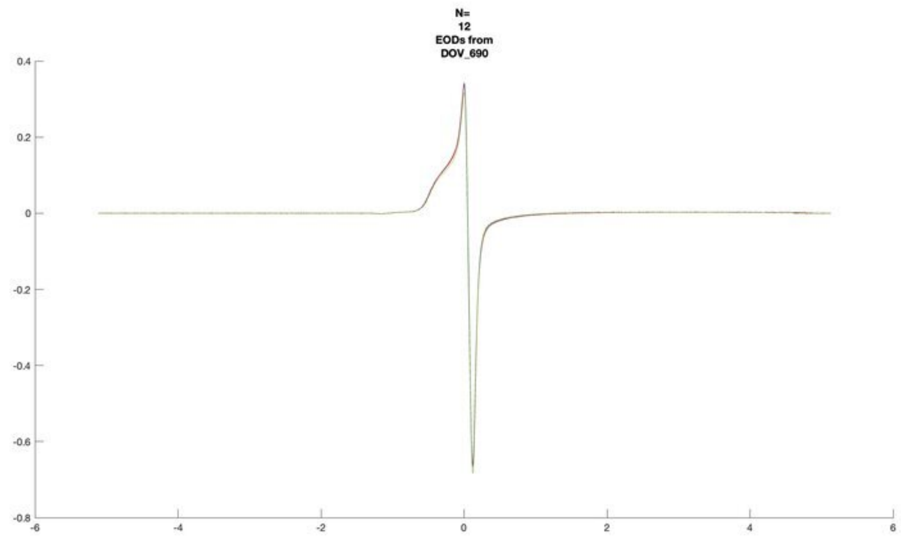
N=
12
EODs from
DOV_690

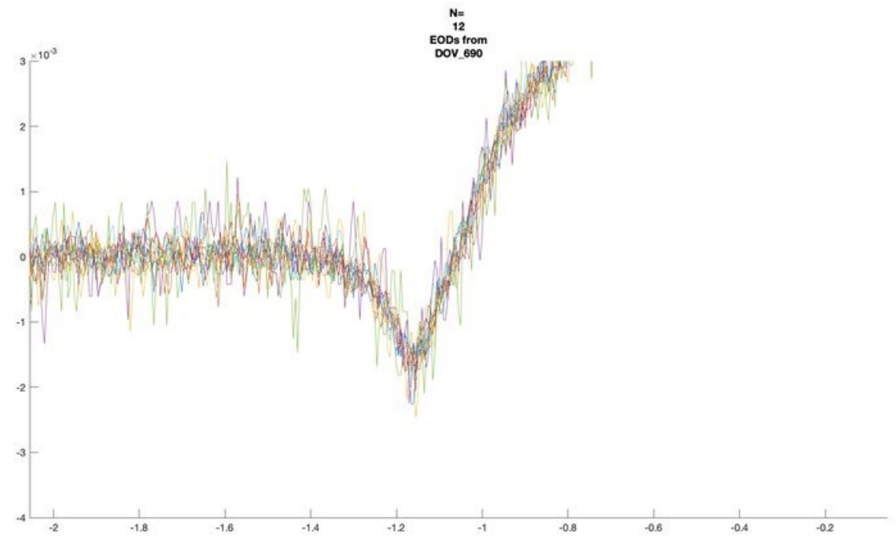
N=
12
EODs from
DOV_690
×10^-3

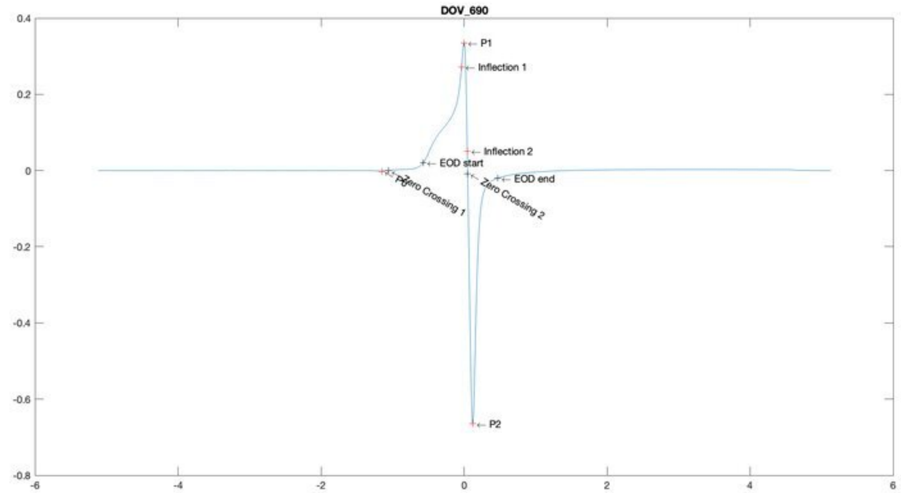
DOV_690
P1
Inflection 1
Inflection 2
EOD start
EOD end
P0
Zero Crossing 1
Zero Crossing 2
P2

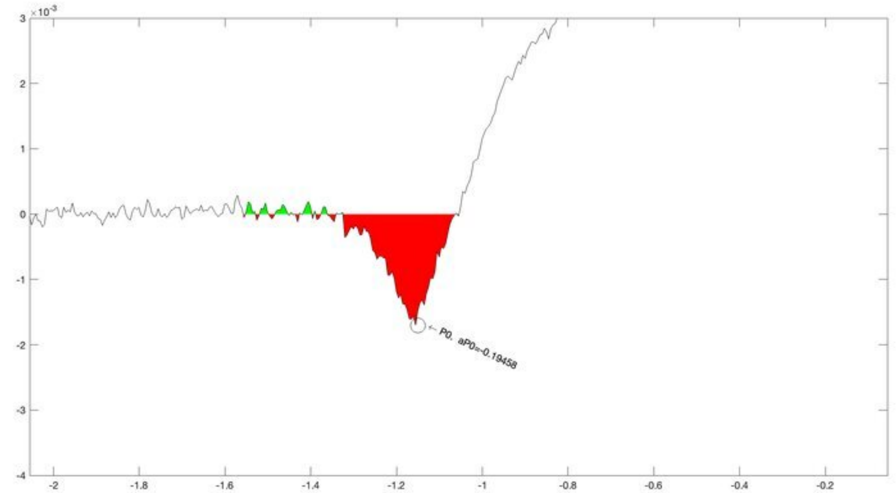
×10^-3
P0, aP0=-0.19458

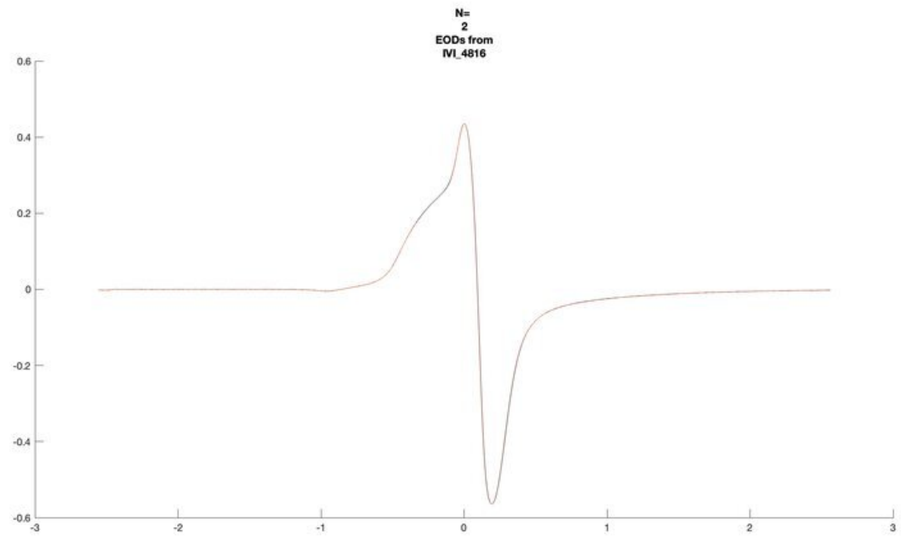
N=
2
EODs from
IVI_4816

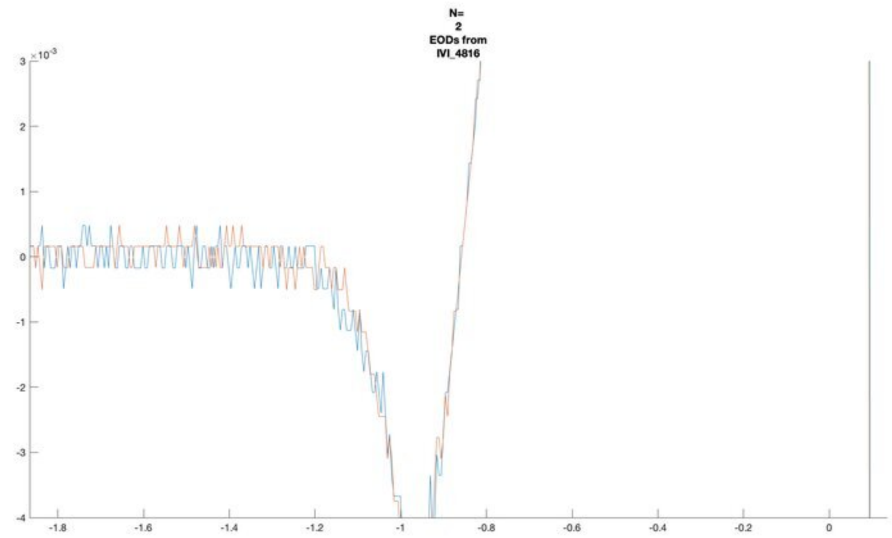
N=
2
EODs from
IVI_4816
×10^-3

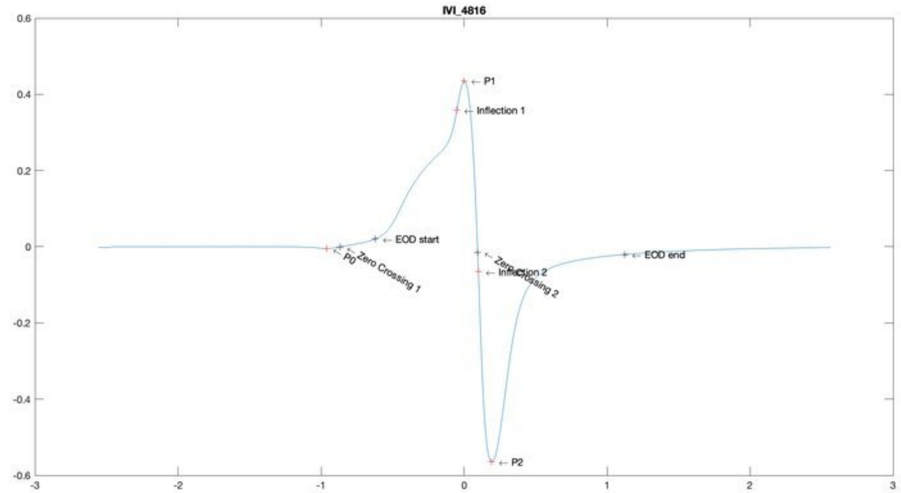
IVI_4816
P1
Inflection 1
EOD start
P0
Zero Crossing 1
Zero Crossing 2
Inflection 2
EOD end
P2

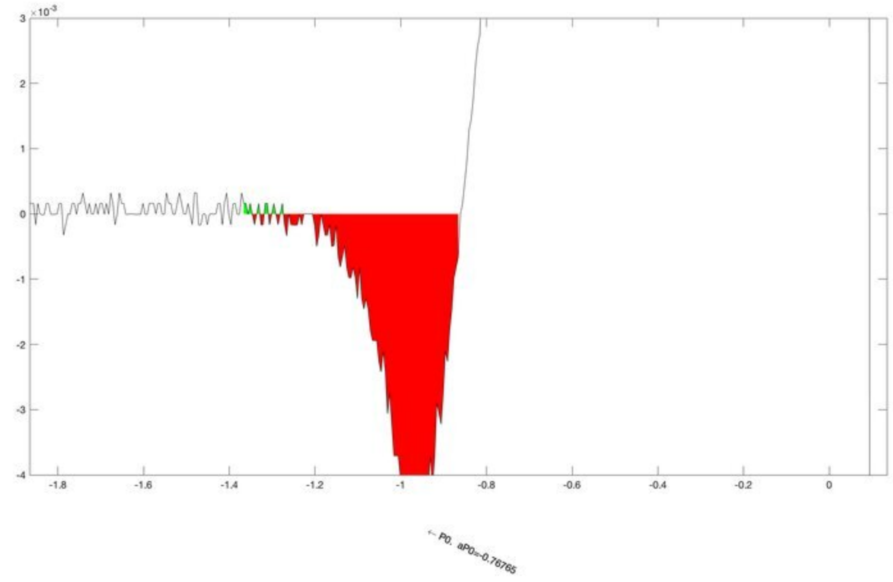
×10^-3
← P0, aP0=-0.76765

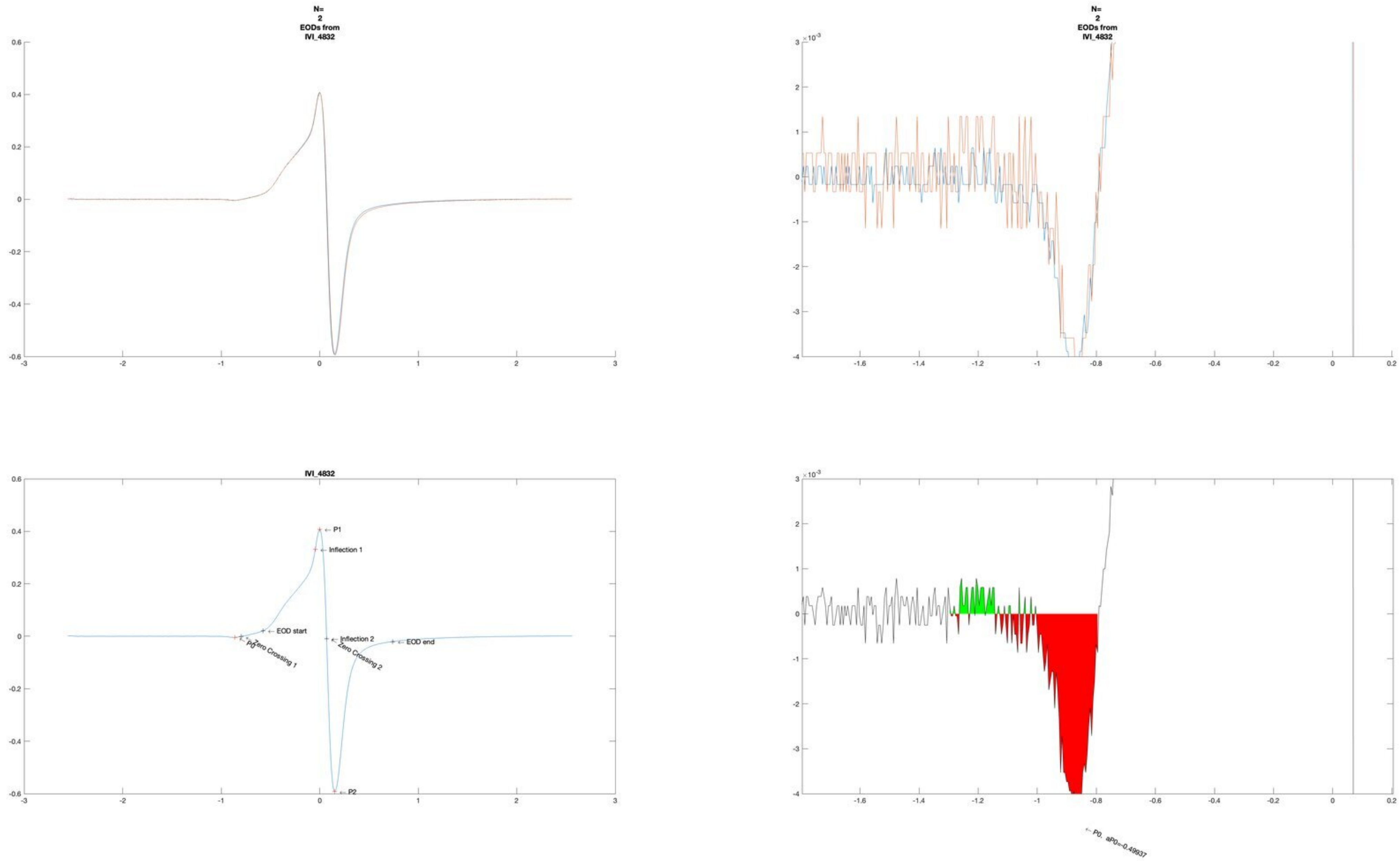
N=
2
EODs from
IVI_4832
N=
2
EODs from
IVI_4832
IVI_4832
P1
Inflection 1
EOD start
P0
Zero Crossing 1
Inflection 2
Zero Crossing 2
EOD end
P2
P0, aP0=-0.49937

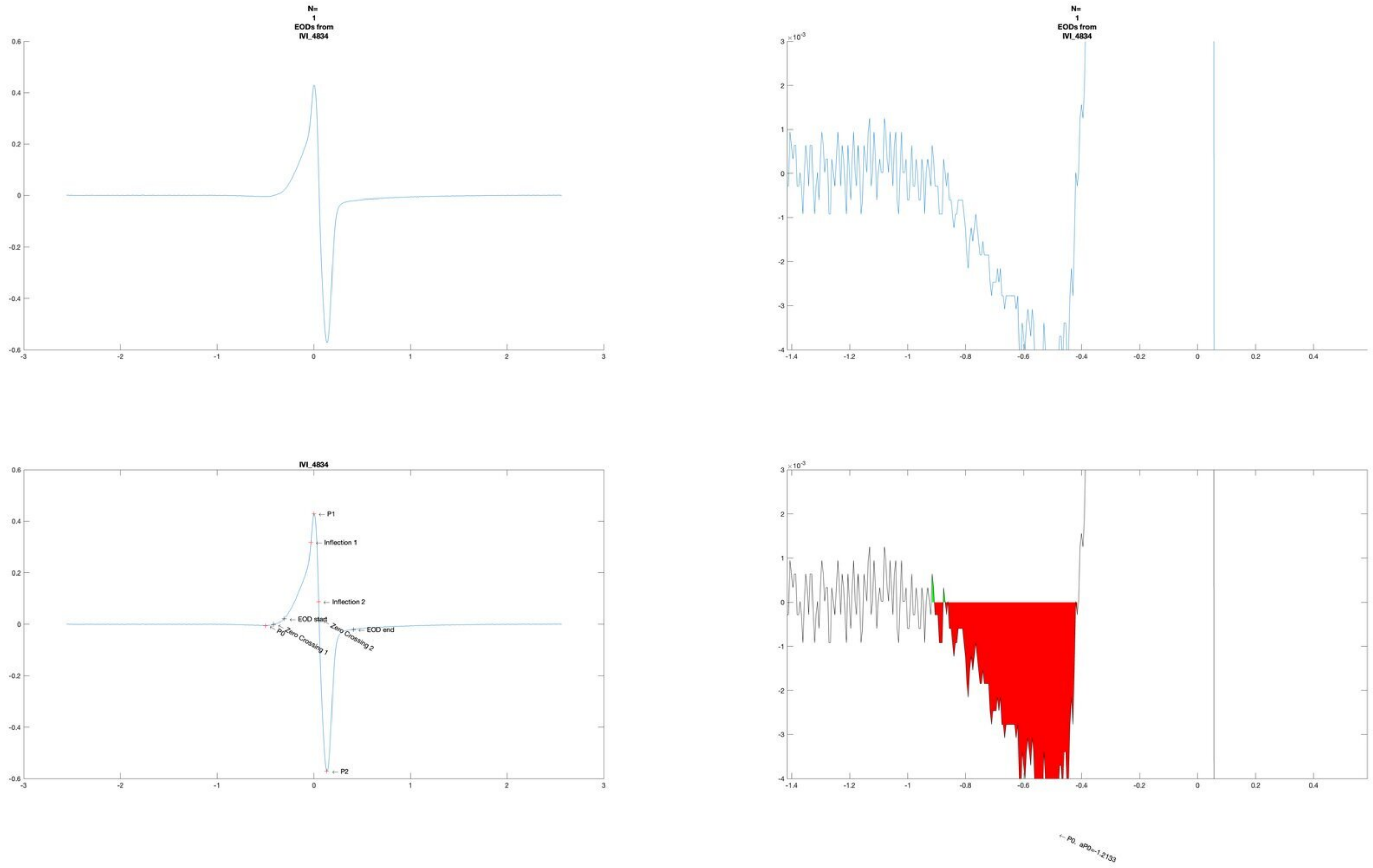
N=
1
EODs from
IVI_4834
N=
1
EODs from
IVI_4834
IVI_4834
← P1
← Inflection 1
← Inflection 2
← EOD start
← EOD end
← P0
Zero Crossing 1
Zero Crossing 2
← P2
← P0. aP0=-1.2133

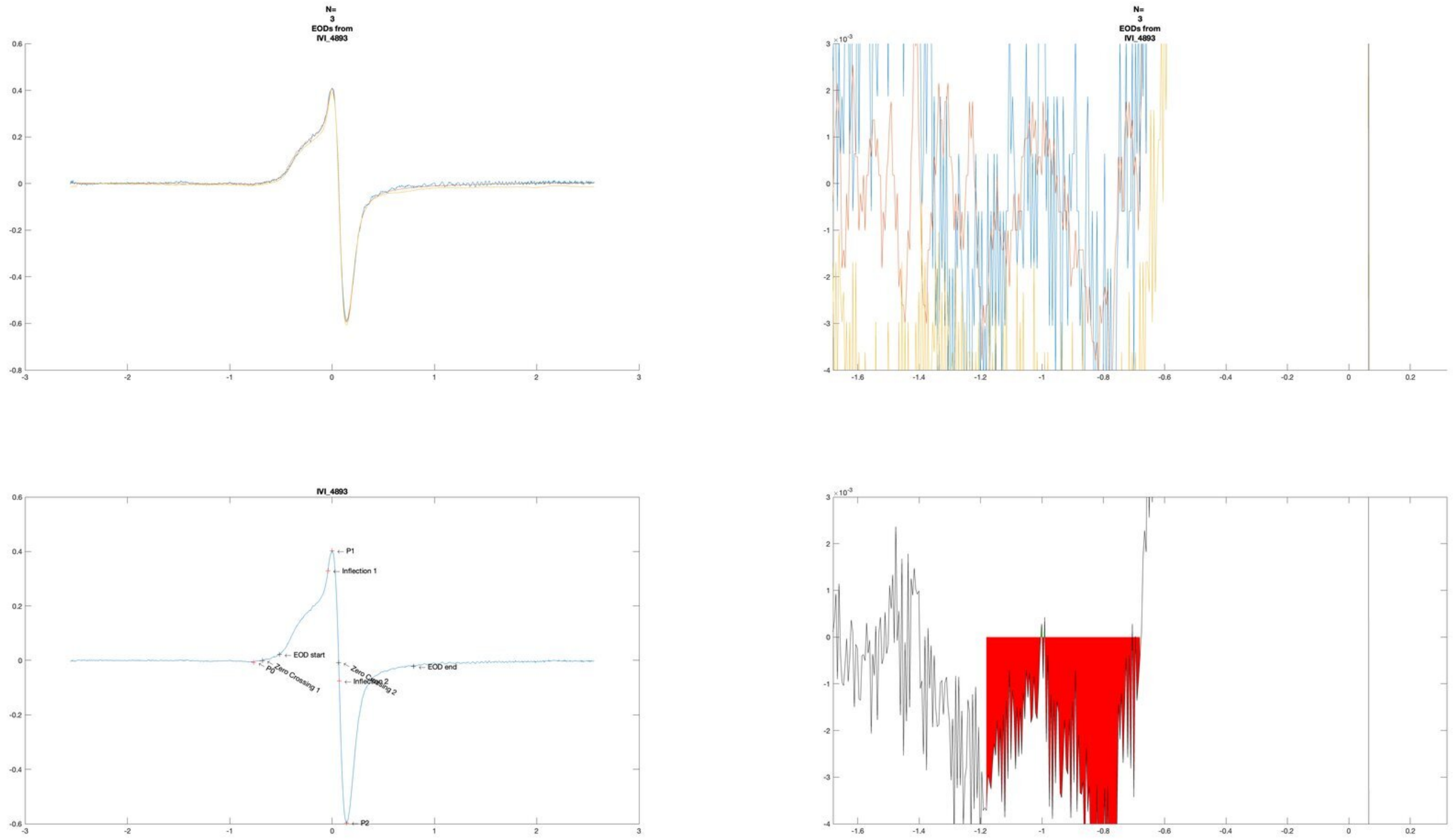
N=
3
EODs from
IVI_4893
N=
3
EODs from
IVI_4893
IVI_4893
P1
Inflection 1
EOD start
P0
Zero Crossing 1
Zero Crossing 2
Inflection 2
EOD end
P2
×10⁻³

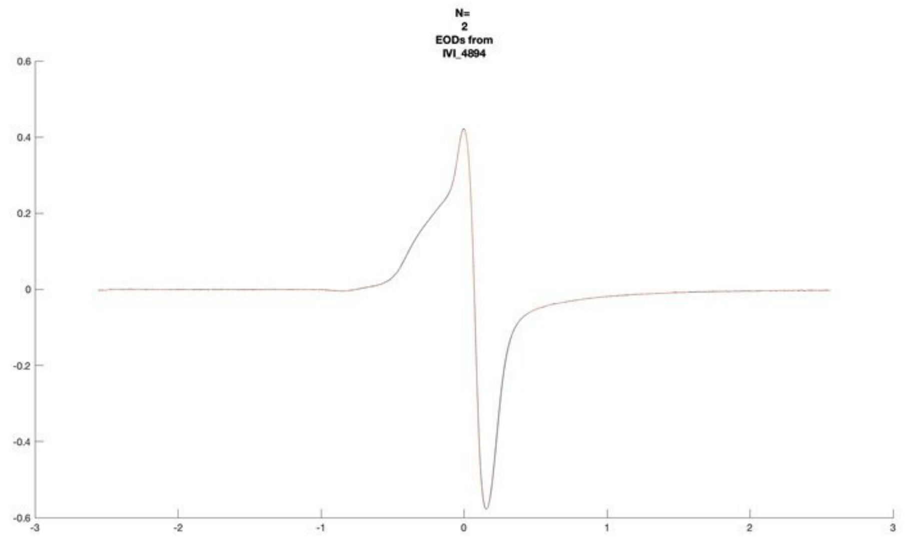
N=
2
EODs from
IVI_4894

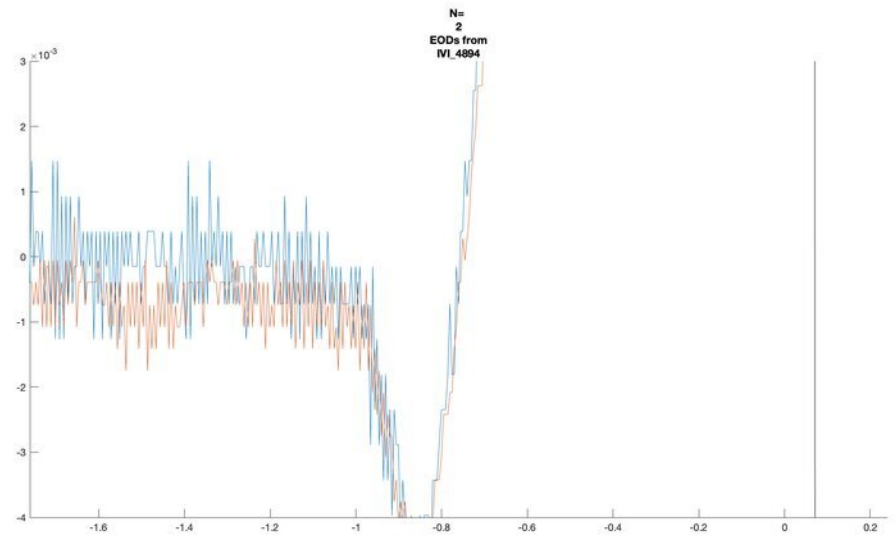
N=
2
EODs from
IVI_4894

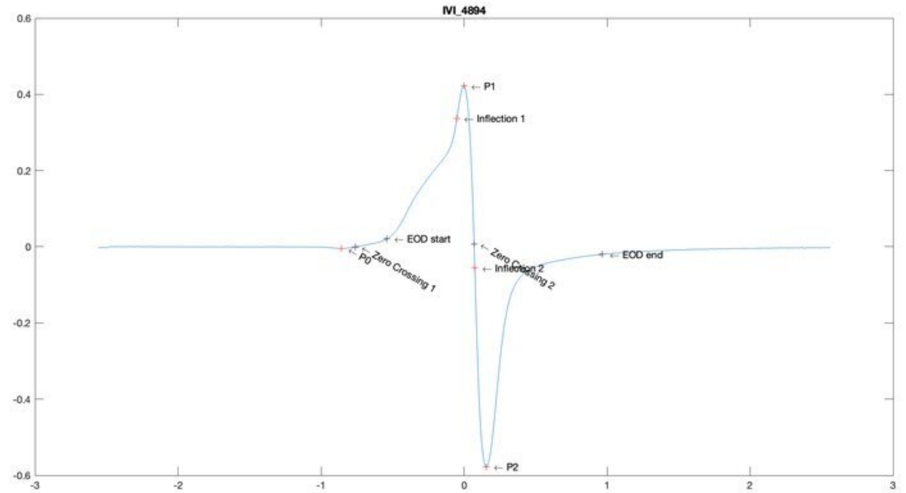
IVI_4894
P1
Inflection 1
EOD start
P0
Zero Crossing 1
Zero Crossing 2
Inflection 2
EOD end
P2

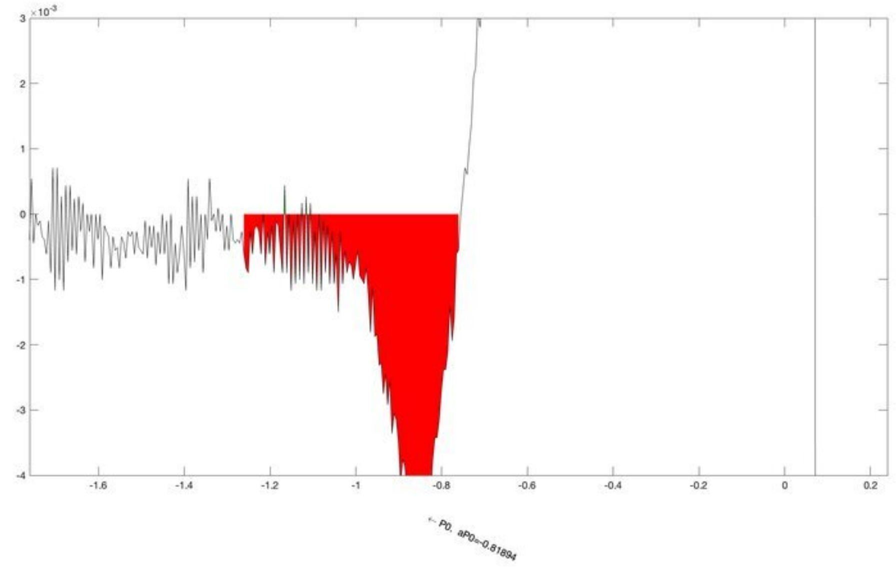
P0, aP0=-0.81894

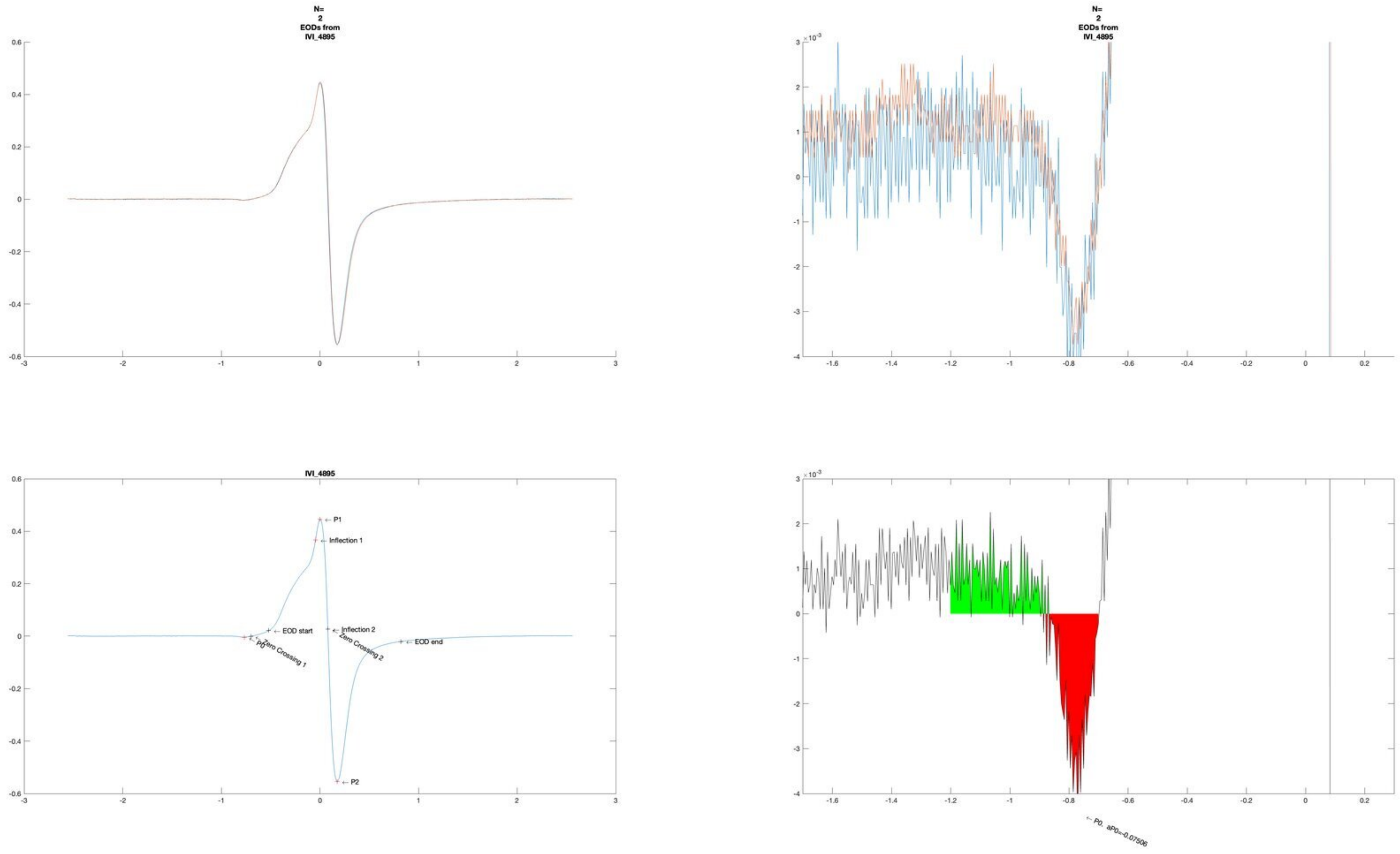
N=
2
EODs from
IVI_4895
N=
2
EODs from
IVI_4895
IVI_4895
P1
Inflection 1
EOD start
Inflection 2
EOD end
P0
Zero Crossing 1
Zero Crossing 2
P2
P0. aP0=-0.07506

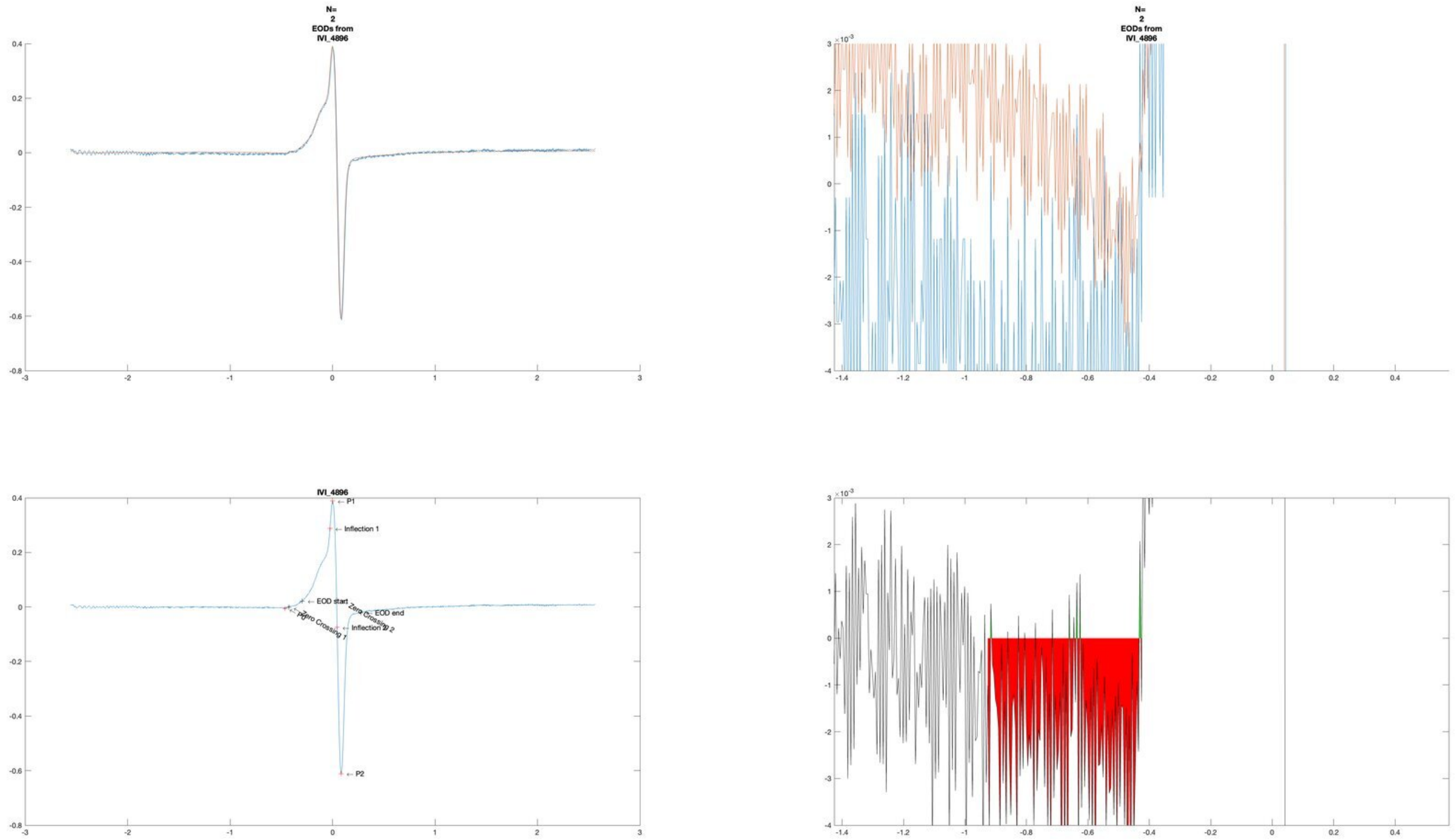
N=
2
EODs from
IVI_4896
N=
2
EODs from
IVI_4896
IVI_4896
P1
Inflection 1
EOD start
P0
Zero Crossing 1
Zero Crossing 2
Inflection 2
EOD end
P2

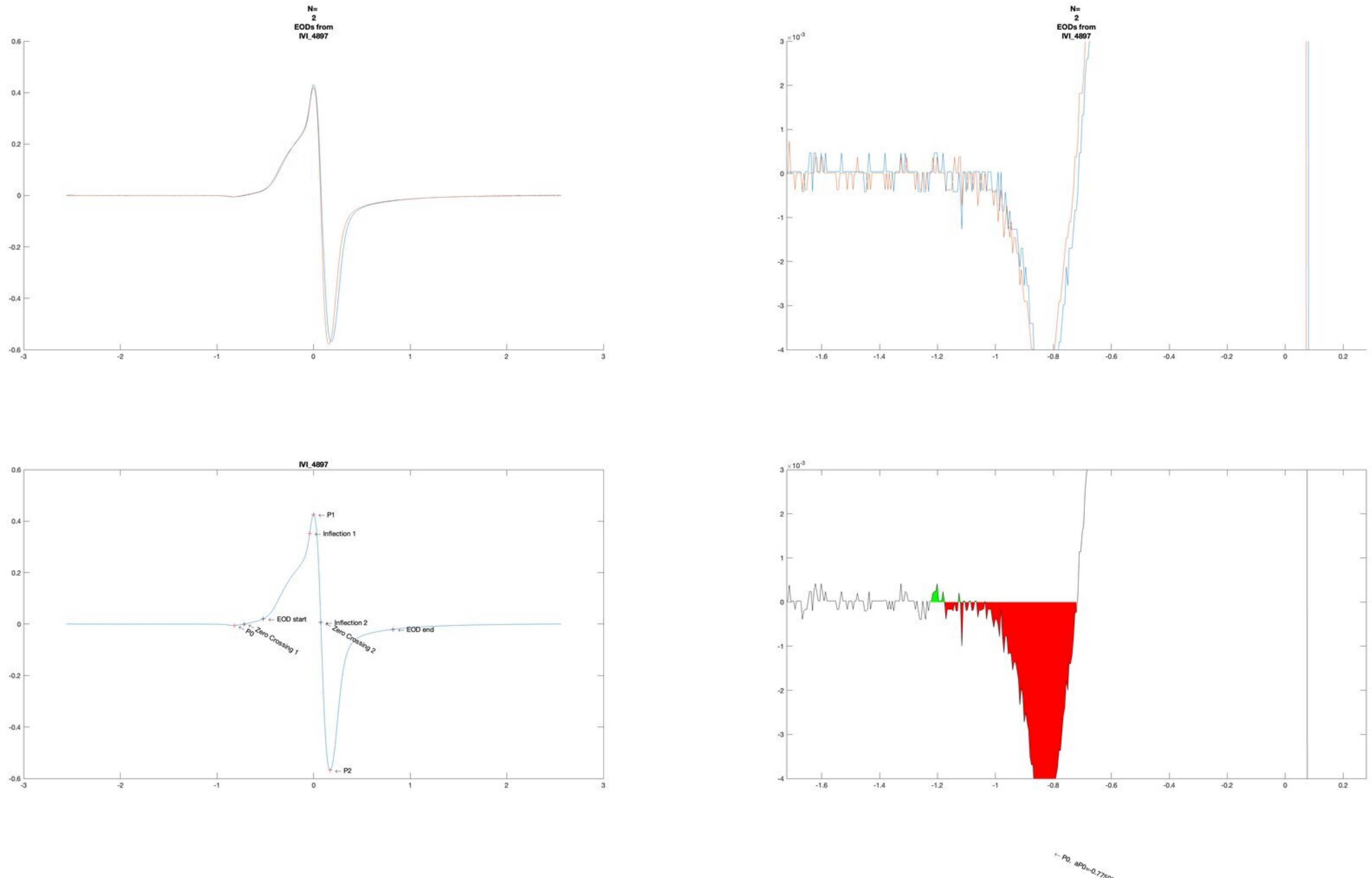
N=
2
EODs from
IVI_4897
N=
2
EODs from
IVI_4897
IVI_4897
P1
Inflection 1
EOD start
Inflection 2
EOD end
P0
Zero Crossing 1
Zero Crossing 2
P2
× 10^-3
× 10^-3

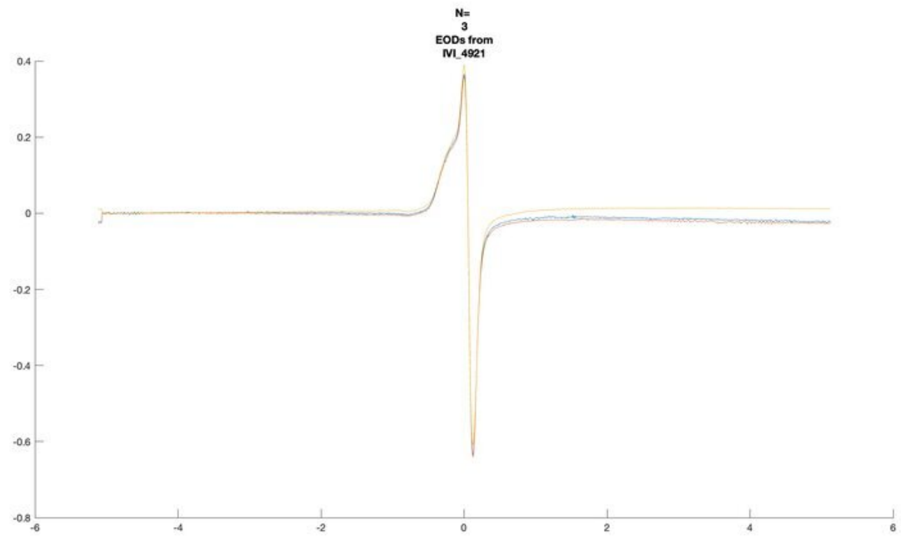
N=
3
EODs from
IVI_4921

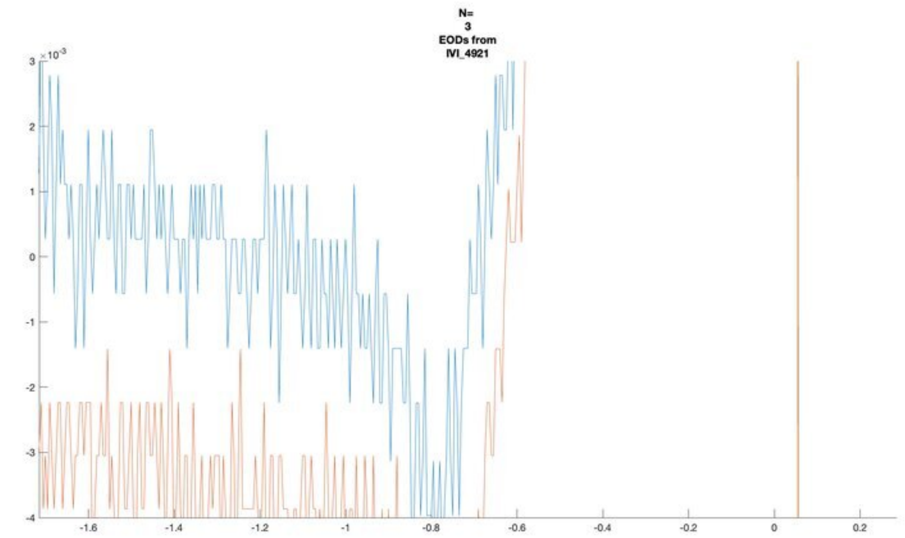
N=
3
EODs from
IVI_4921

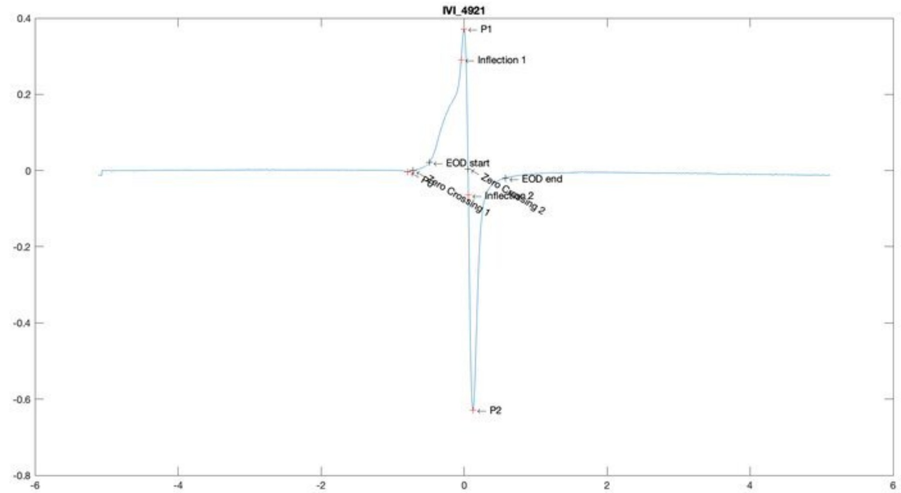
IVI_4921
P1
Inflection 1
EOD start
EOD end
Zero Crossing 1
Zero Crossing 2
Inflection 2
P2

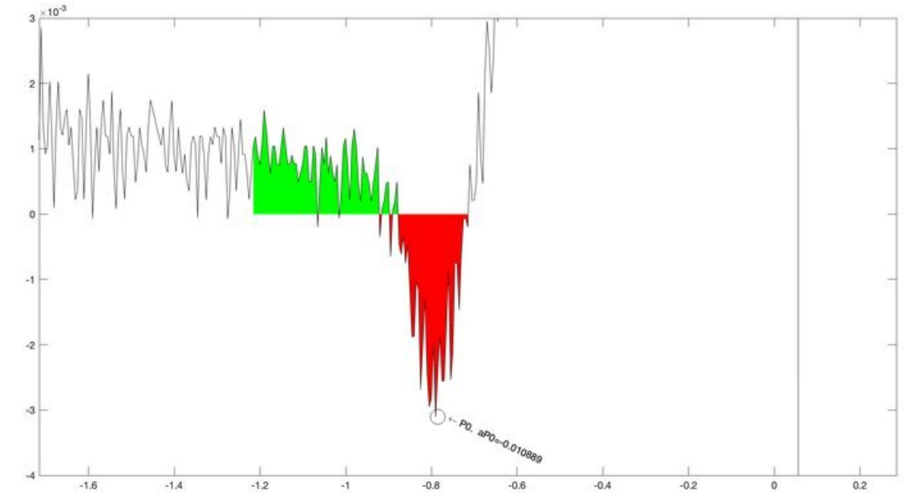
P0, aP0=-0.01089

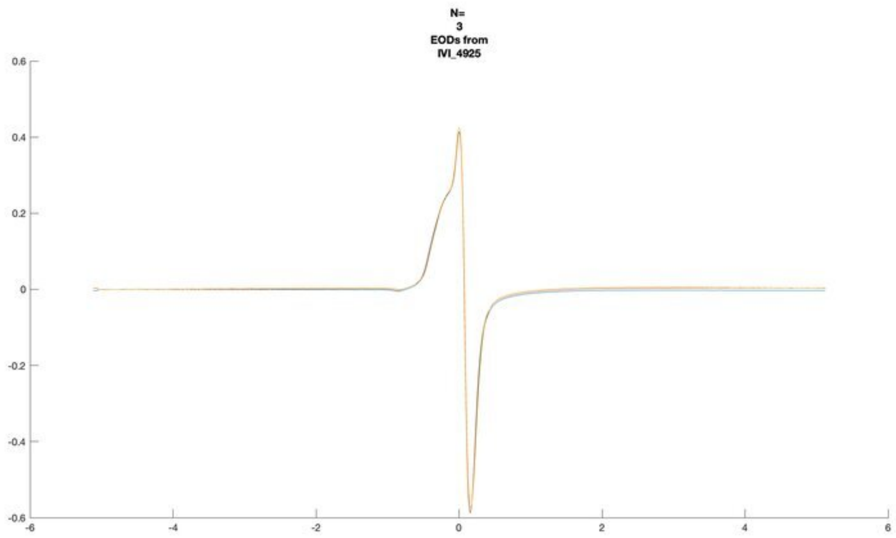
N=
3
EODs from
IVI_4925

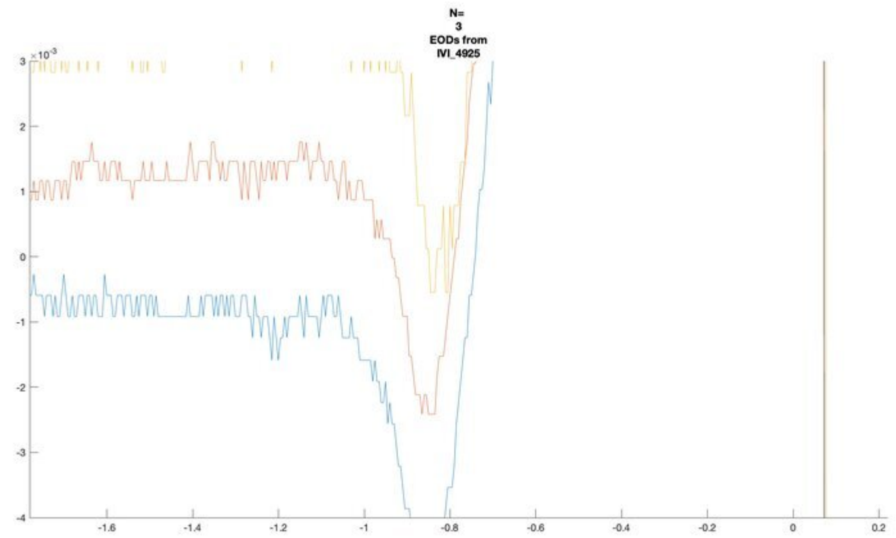
N=
3
EODs from
IVI_4925

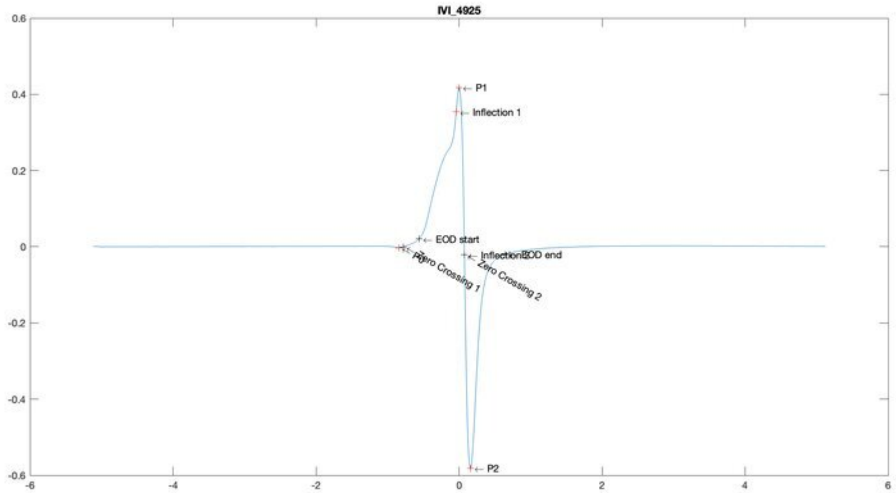
IVI_4925
P1
Inflection 1
EOD start
EOD end
Zero Crossing 1
Zero Crossing 2
P2

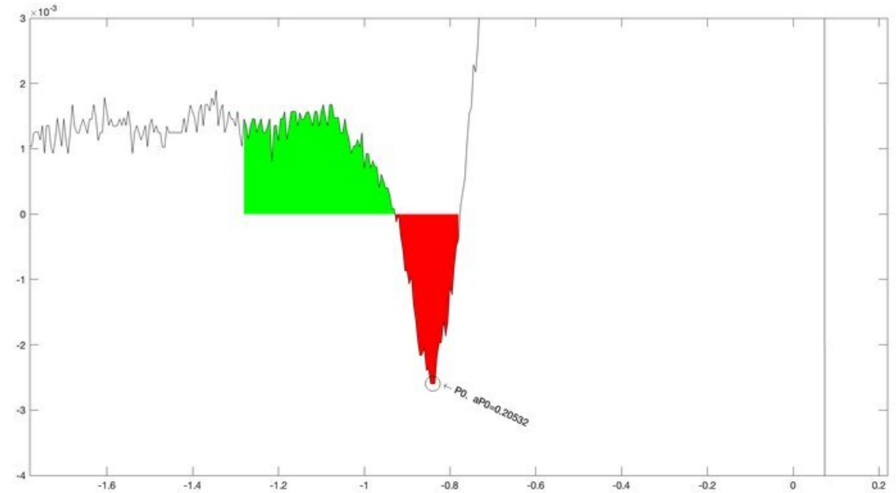
P0, aP0=0.20532

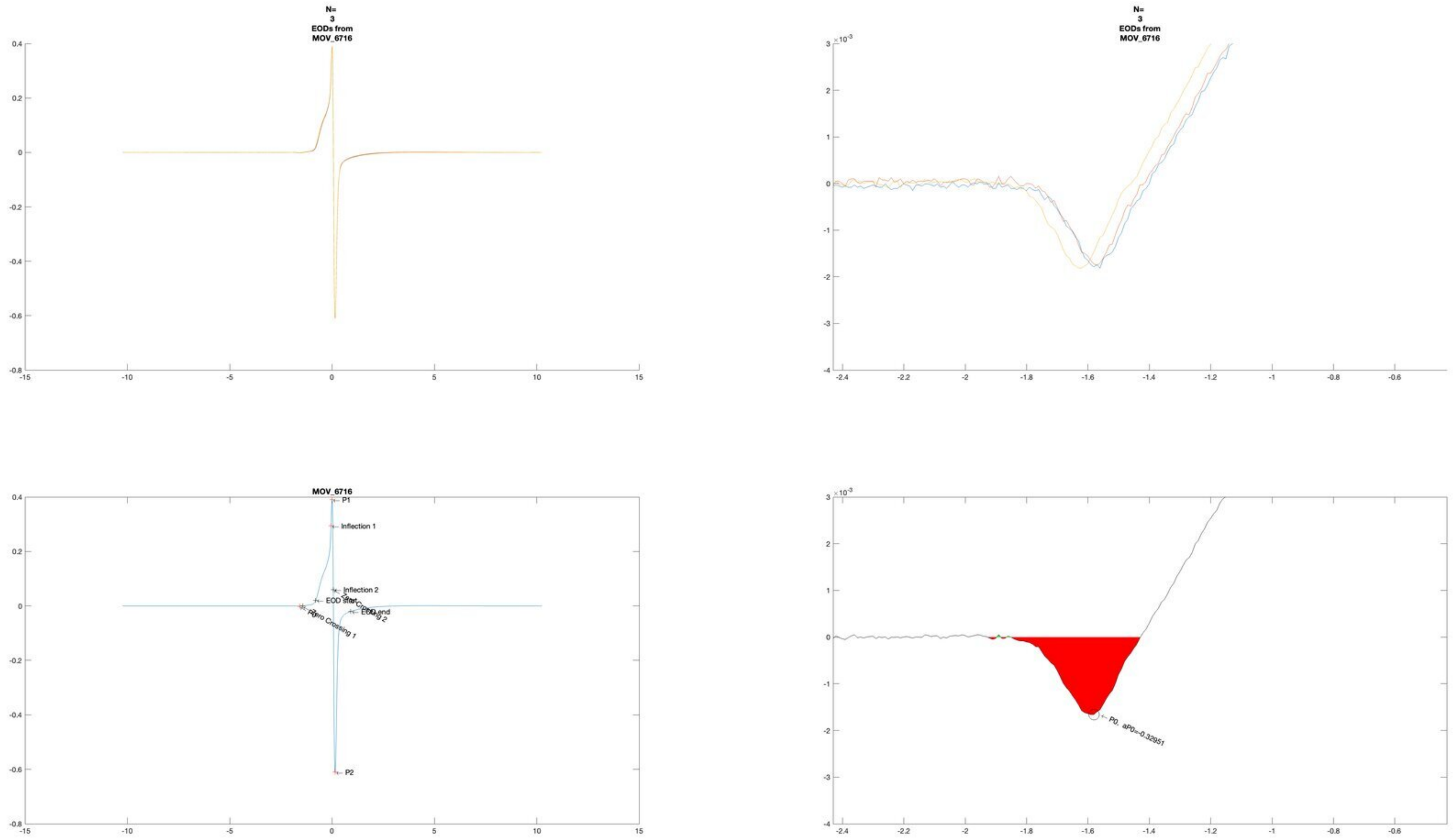

N=
3
EODs from
MOV_6716
N=
3
EODs from
MOV_6716
MOV_6716
P1
Inflection 1
Inflection 2
EOD start
EOD end
Zero Crossing 1
P0
P2
P0, aP0=-0.32951

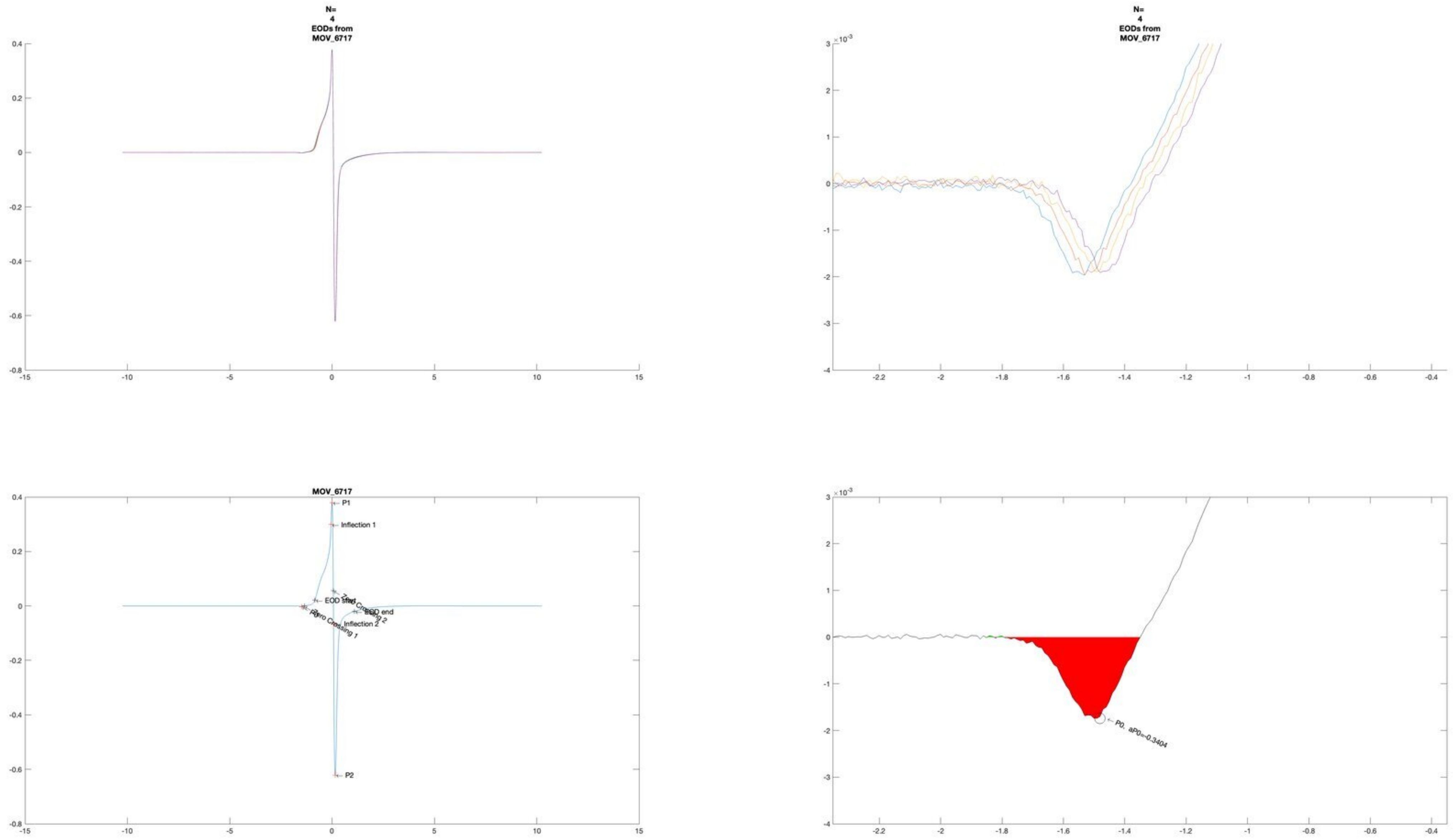
N=
4
EODs from
MOV_6717
N=
4
EODs from
MOV_6717
MOV_6717
P1
Inflection 1
EOD start
Zero Crossing 2
EOD end
Inflection 2
Zero Crossing 1
P2
P0, aP0=-0.3404

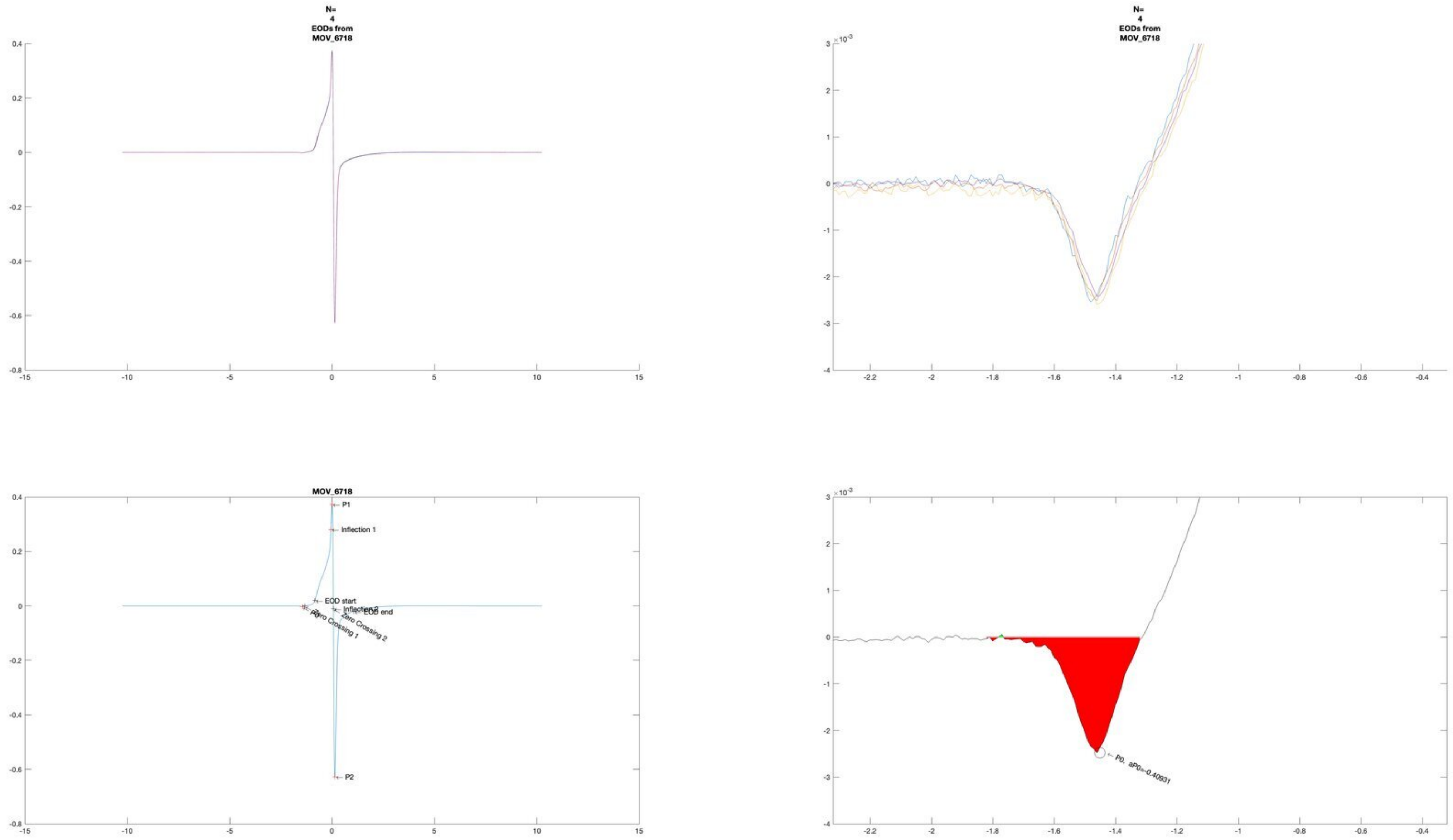
N=
4
EODs from
MOV_6718
N=
4
EODs from
MOV_6718
MOV_6718
P1
Inflection 1
EOD start
Inflection 2
EOD end
Zero Crossing 2
Zero Crossing 1
P2
P0, aP0=-0.40931

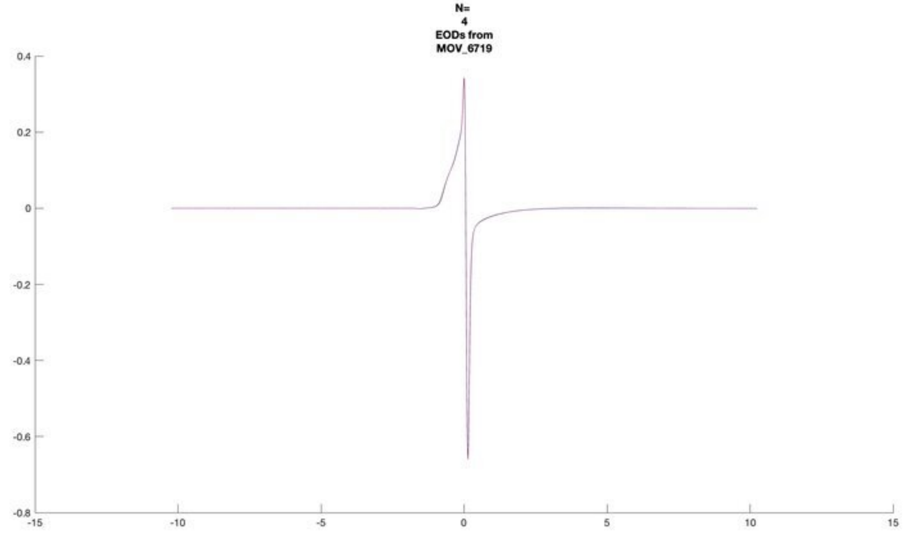
N=
4
EODs from
MOV_6719

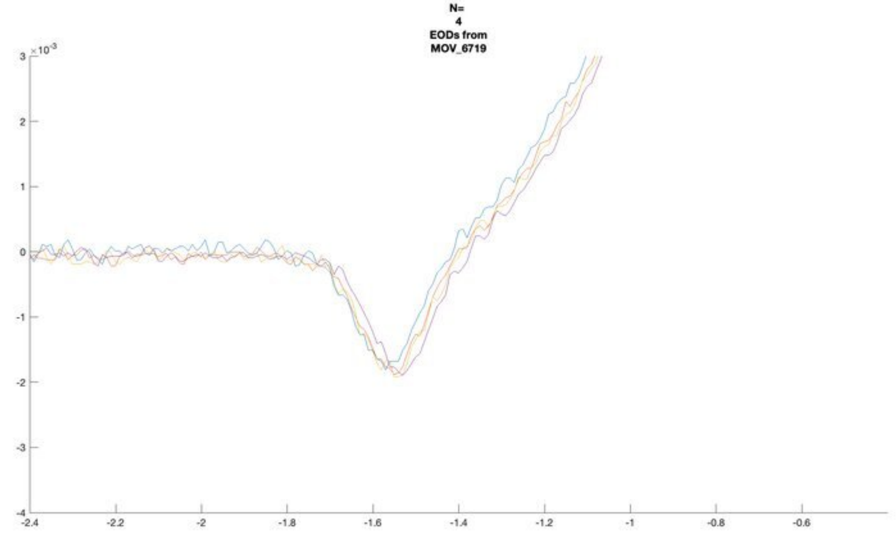
N=
4
EODs from
MOV_6719

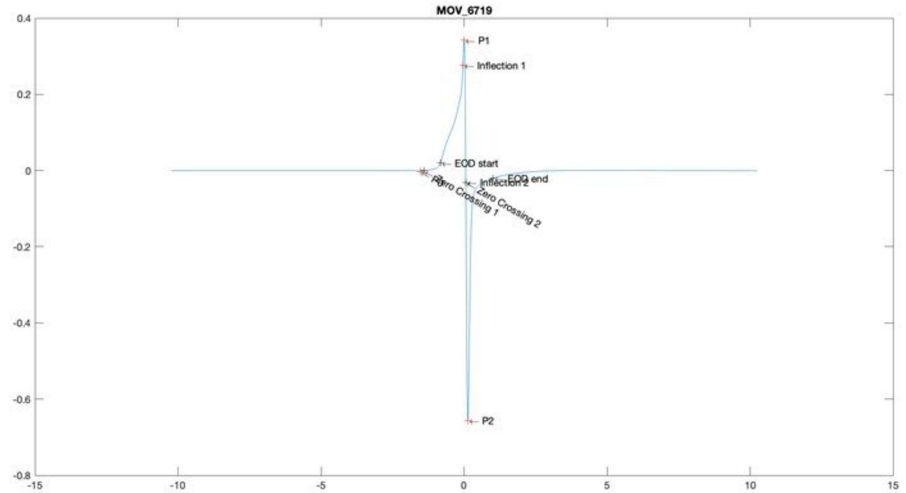
MOV_6719
P1
Inflection 1
EOD start
EOD end
Inflection 2
Zero Crossing 1
Zero Crossing 2
P2

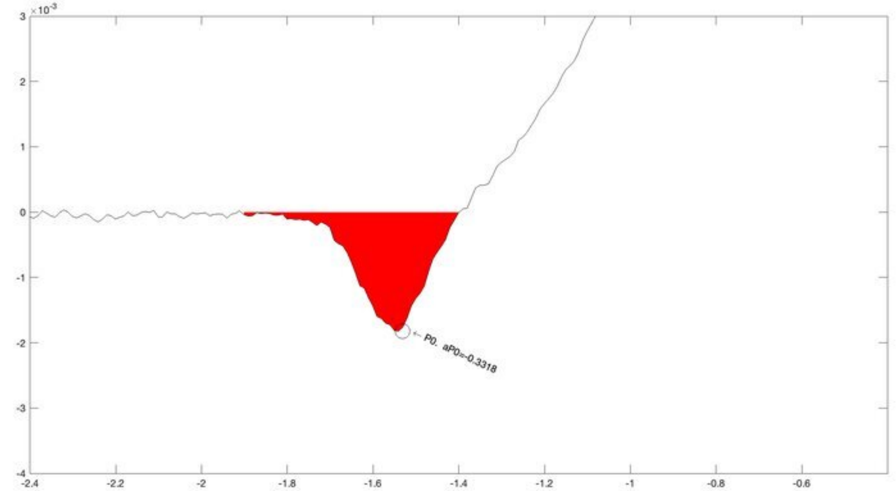
P0. aP0=-0.3318

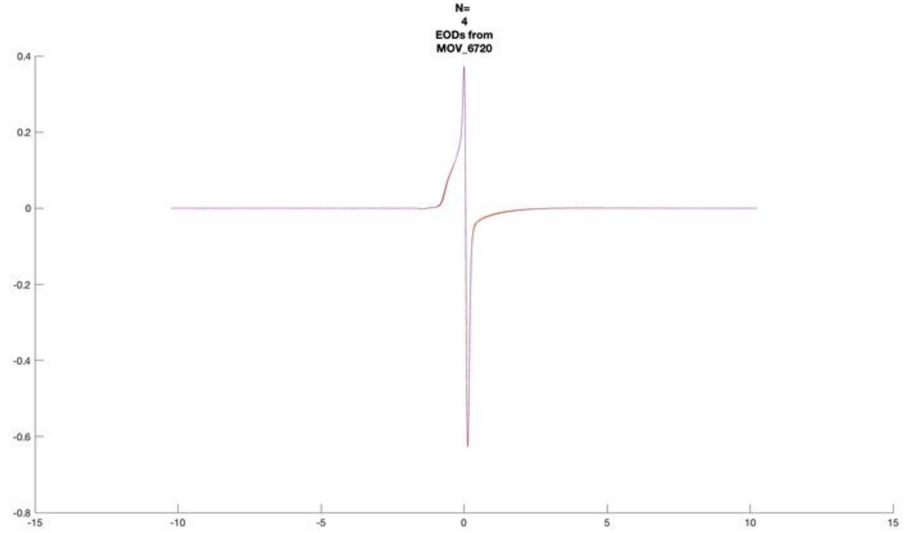
N=
4
EODs from
MOV_6720

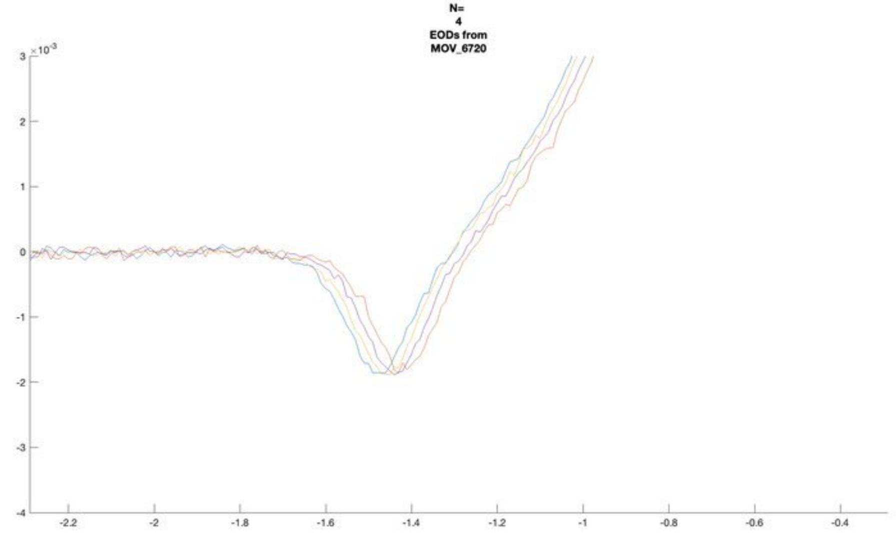
N=
4
EODs from
MOV_6720
×10^-3

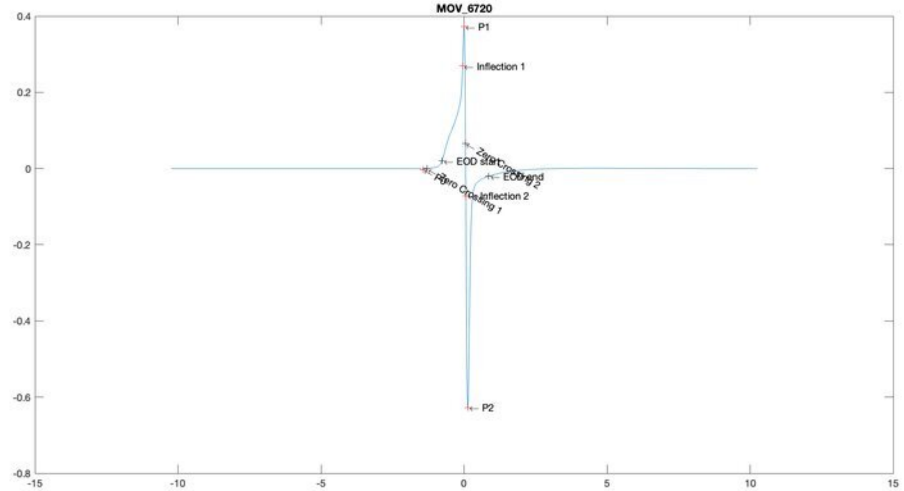
MOV_6720
P1
Inflection 1
EOD start
Zero Crossing 2
EOD end
Zero Crossing 1
Inflection 2
P2

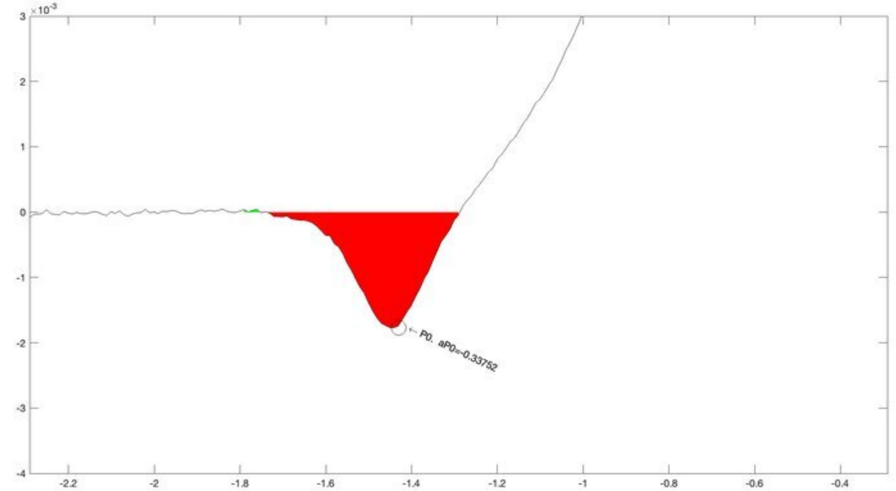
×10^-3
P0, aP0=-0.33752

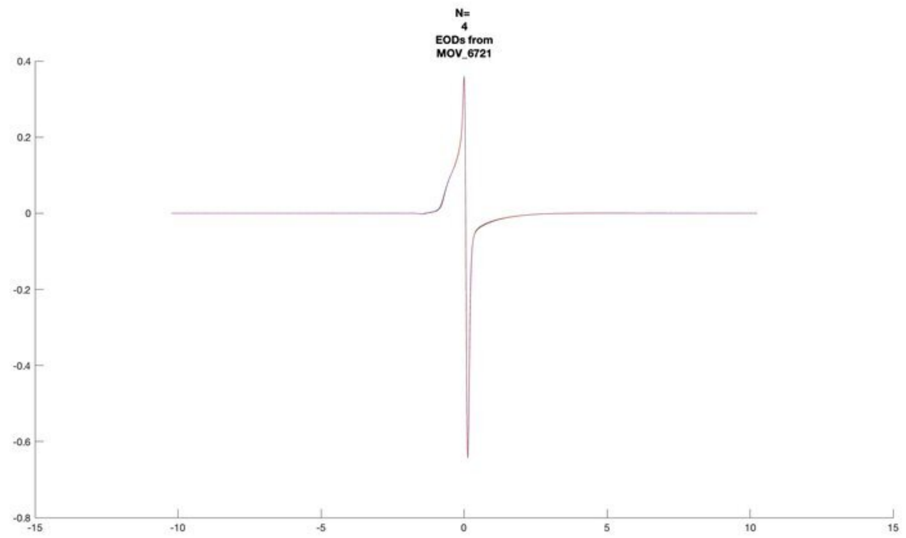
N=
4
EODs from
MOV_6721

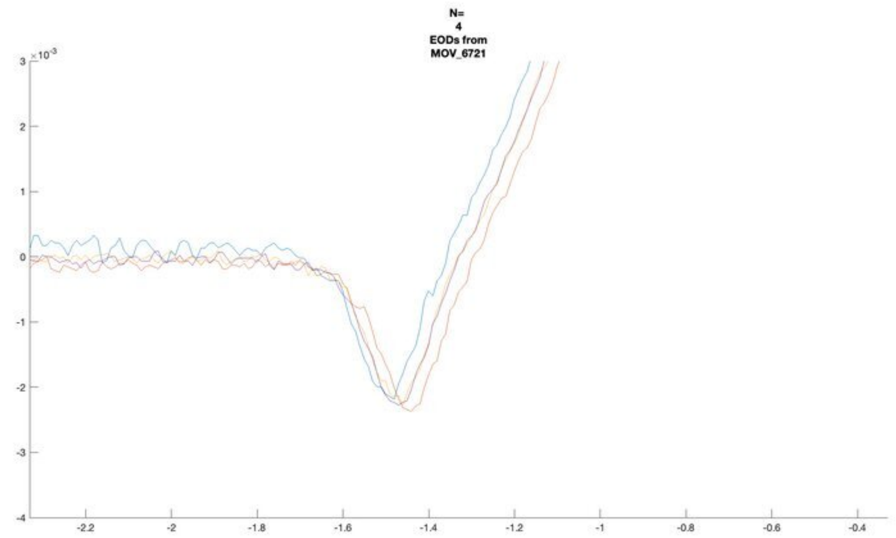
N=
4
EODs from
MOV_6721

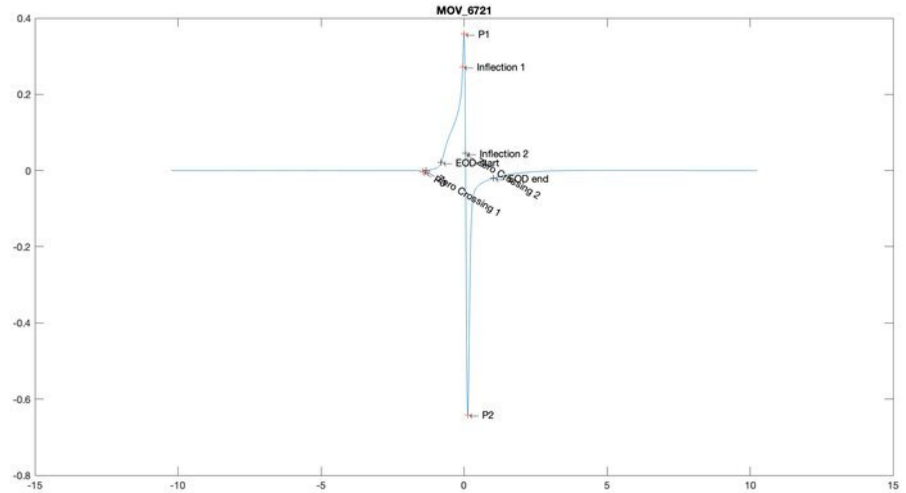
MOV_6721
P1
Inflection 1
Inflection 2
EOD start
EOD end
Zero Crossing 1
Zero Crossing 2
P2

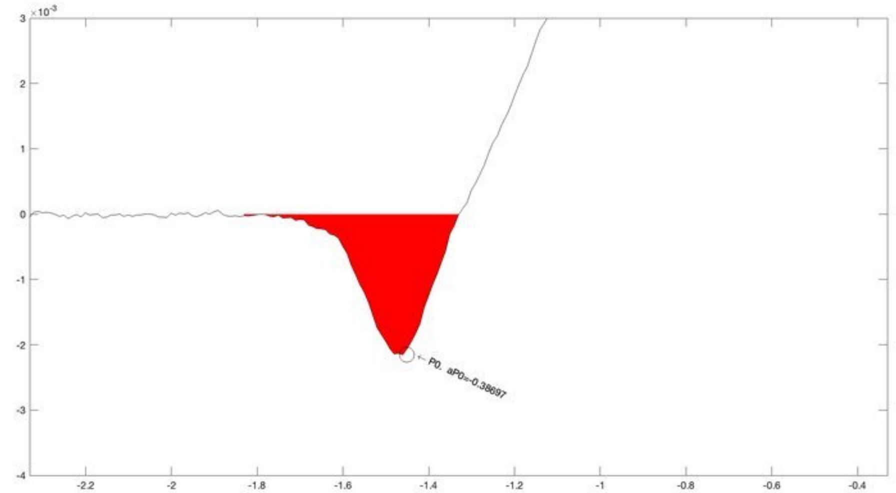
P0, aP0=-0.38697

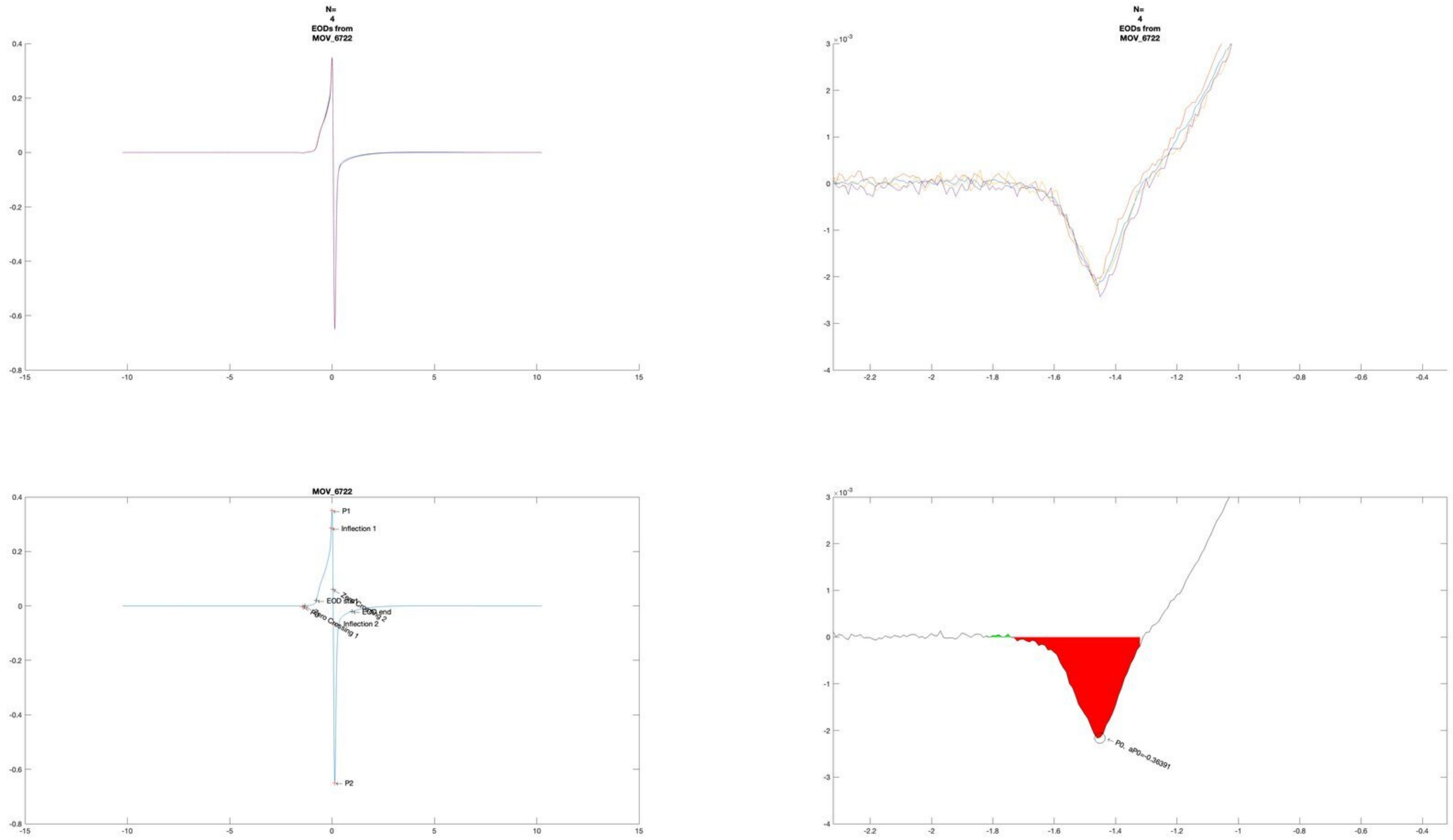
N=
4
EODs from
MOV_6722
MOV_6722
P1
Inflection 1
EOD start
Zero Crossing 1
Zero Crossing 2
EOD end
Inflection 2
P2
P0. aP0=-0.36391

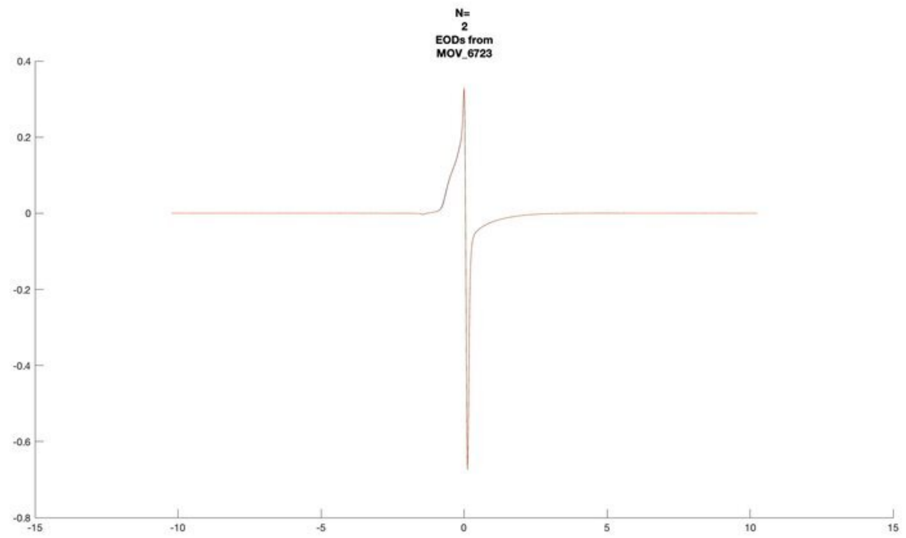
N=
2
EODs from
MOV_6723

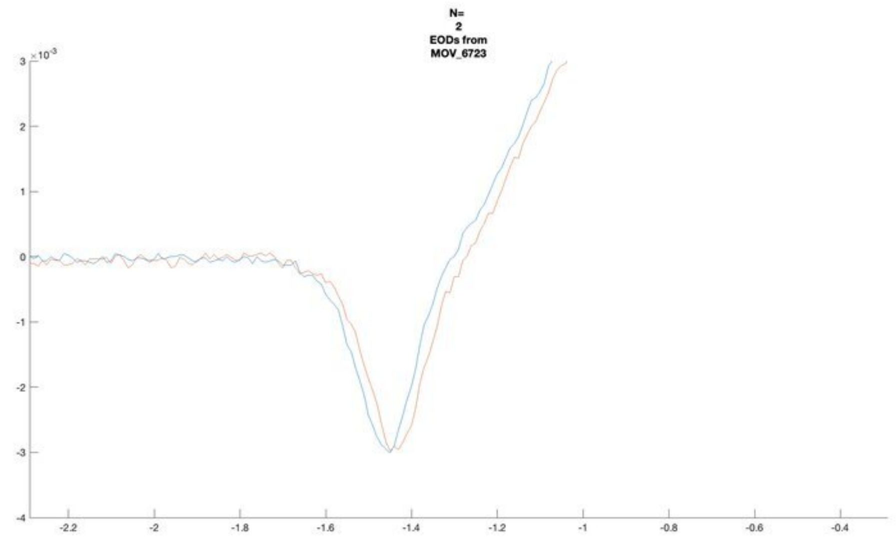
N=
2
EODs from
MOV_6723
×10⁻³

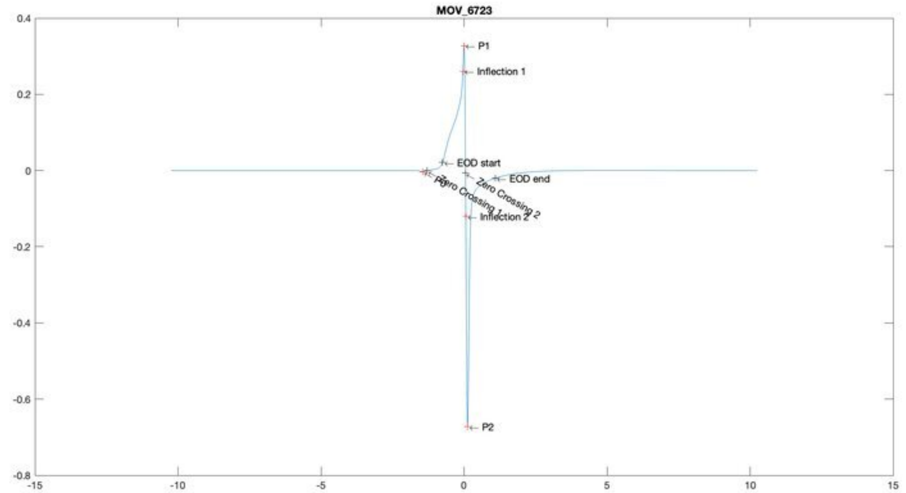
MOV_6723
P1
Inflection 1
EOD start
EOD end
P0
Zero Crossing 1
Zero Crossing 2
Inflection 2
P2

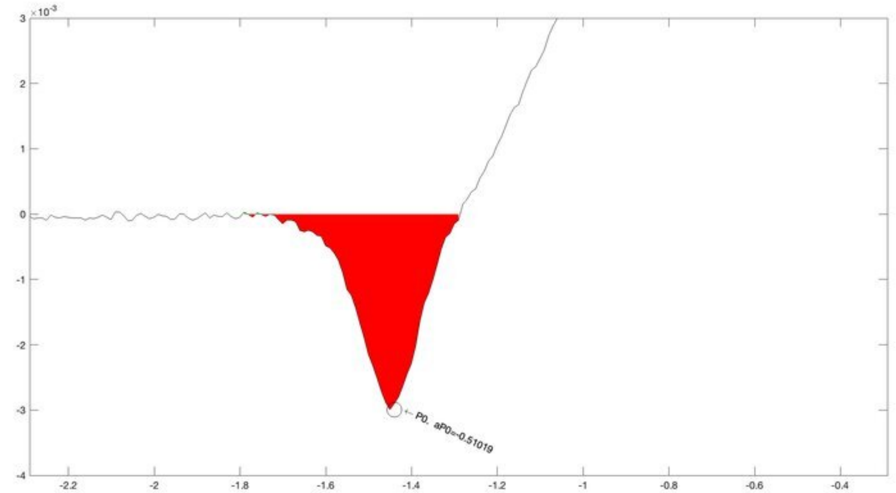
×10⁻³
P0, aP0=-0.51019

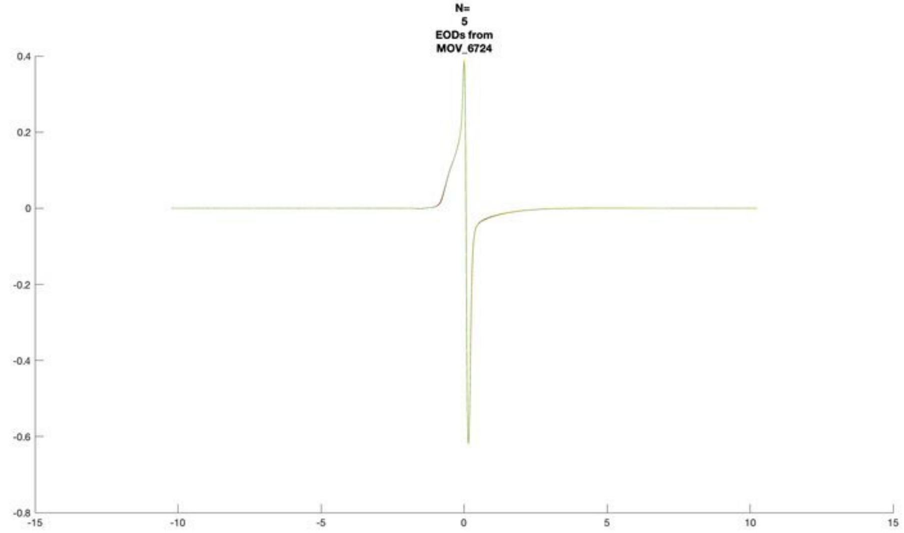
N=
5
EODs from
MOV_6724

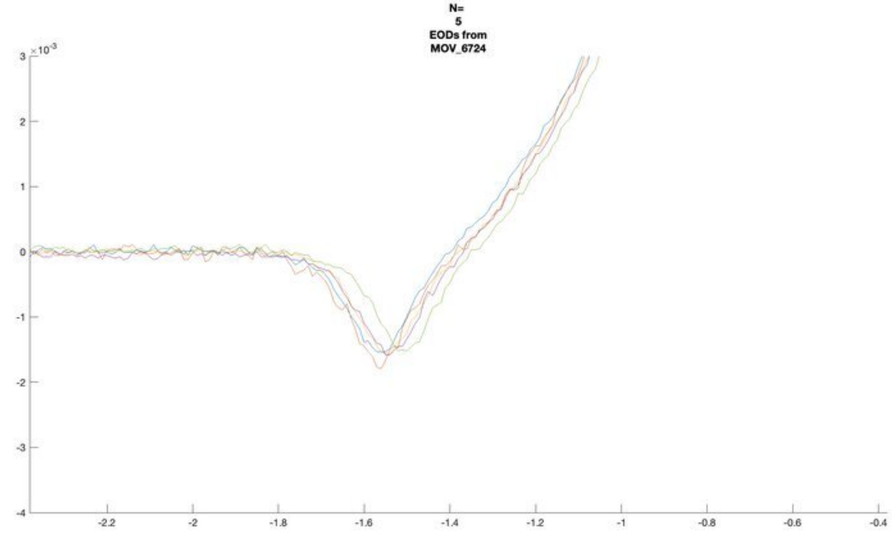
N=
5
EODs from
MOV_6724
×10^-3

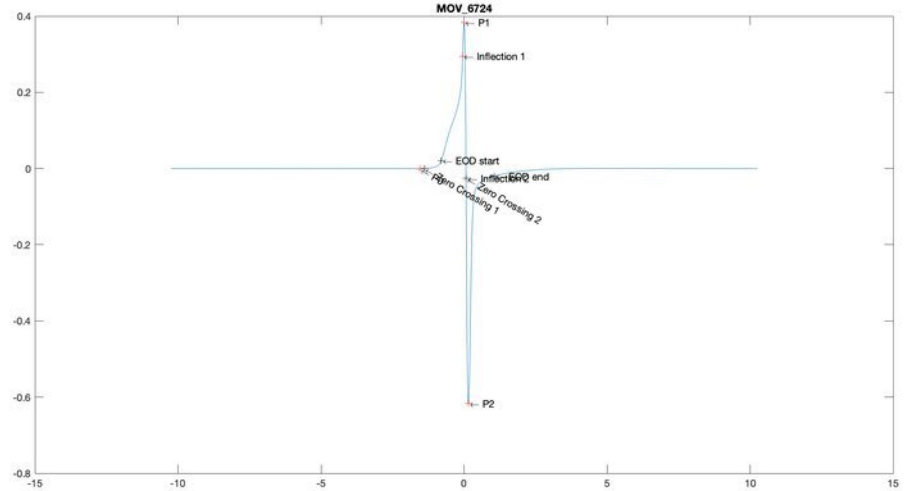
MOV_6724
P1
Inflection 1
EOD start
EOD end
Inflection 2
Zero Crossing 1
Zero Crossing 2
P0
P2

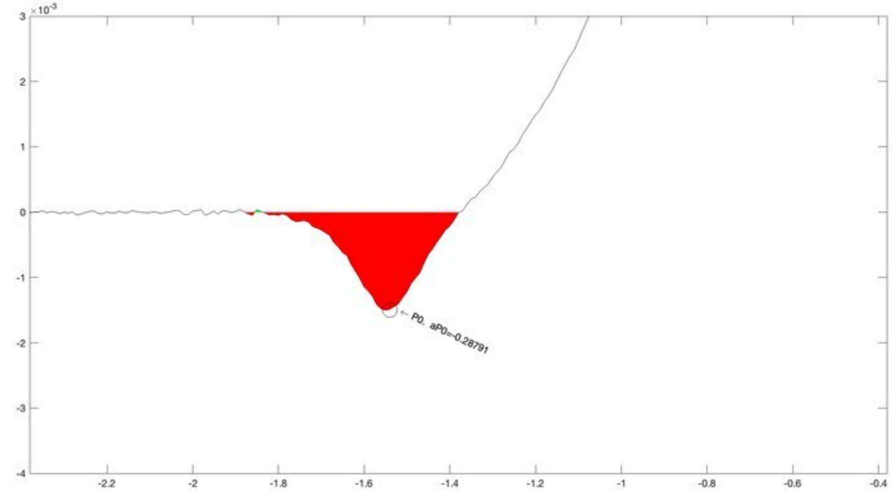
×10^-3
P0, aP0=-0.28791

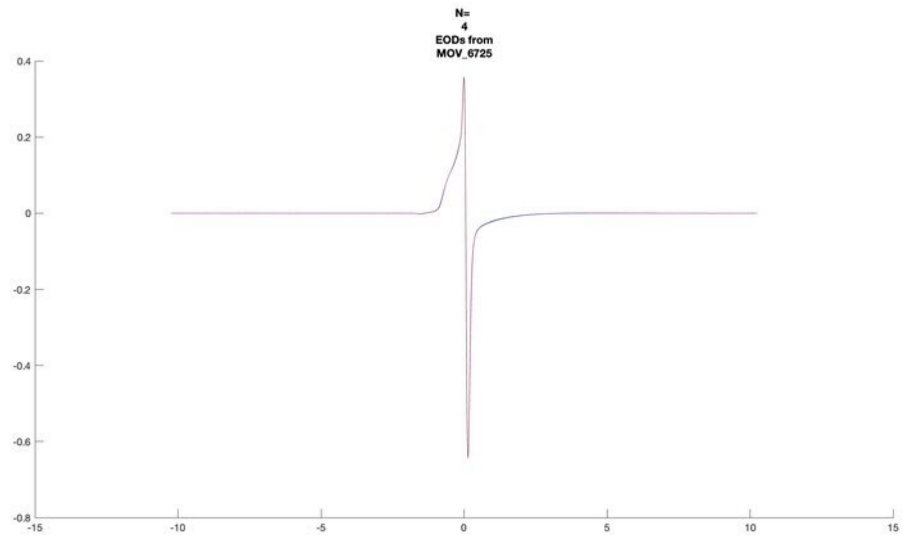
N=
4
EODs from
MOV_6725

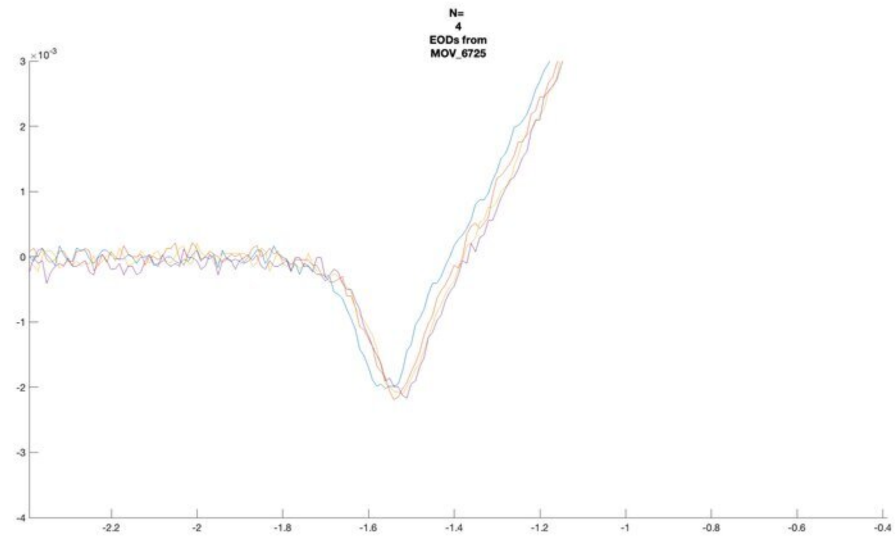
N=
4
EODs from
MOV_6725

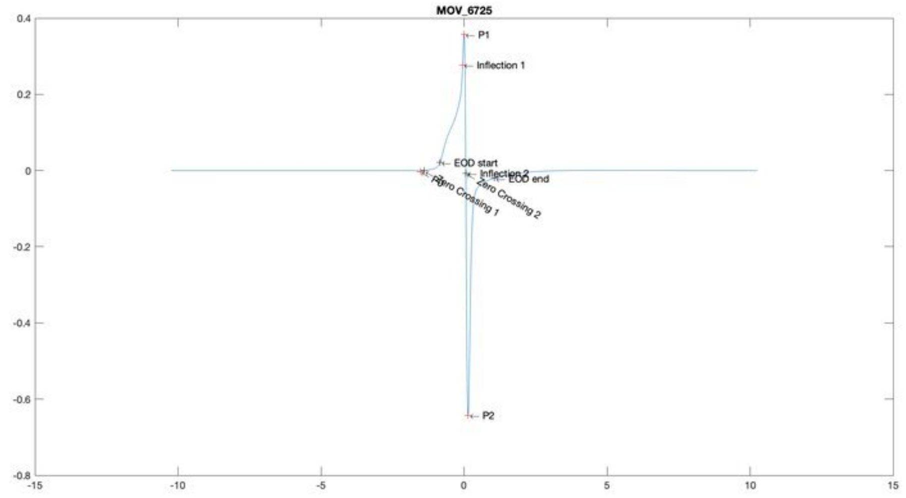
MOV_6725
P1
Inflection 1
EOD start
Inflection 2
EOD end
Zero Crossing 1
Zero Crossing 2
P2

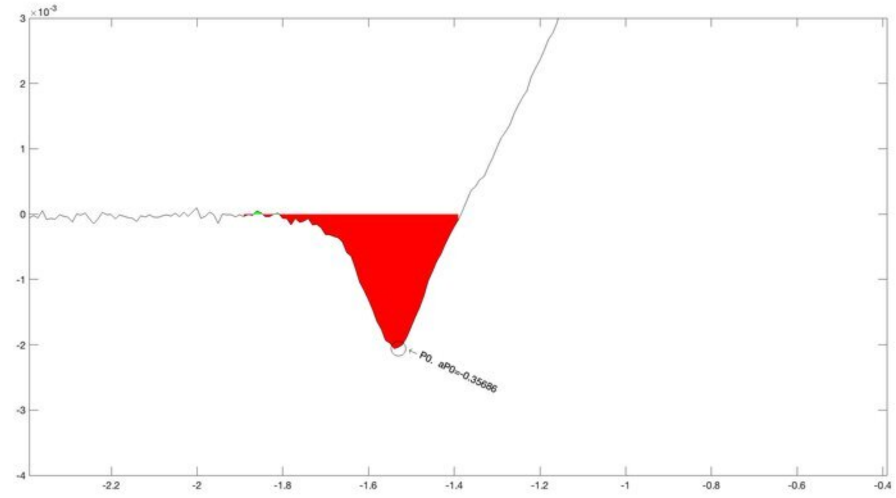
P0, aP0=-0.35686

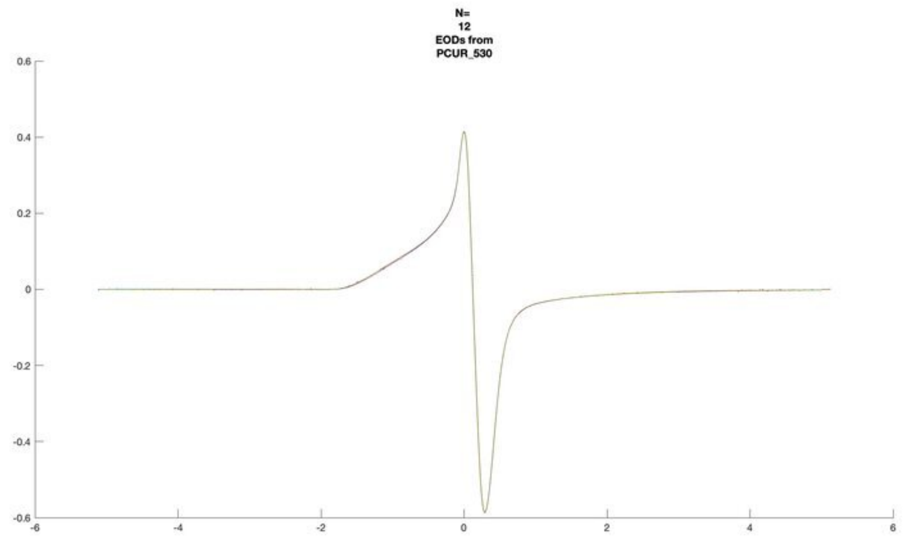
N=
12
EODs from
PCUR_530

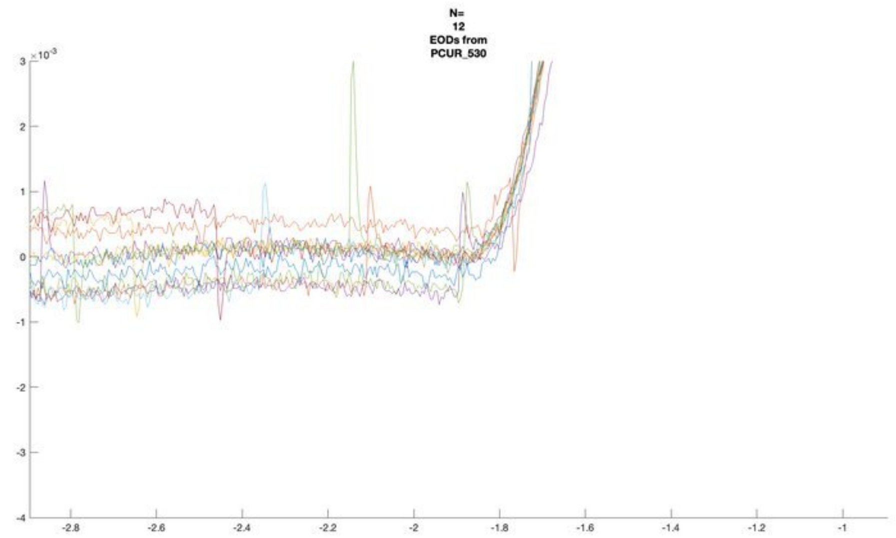
N=
12
EODs from
PCUR_530

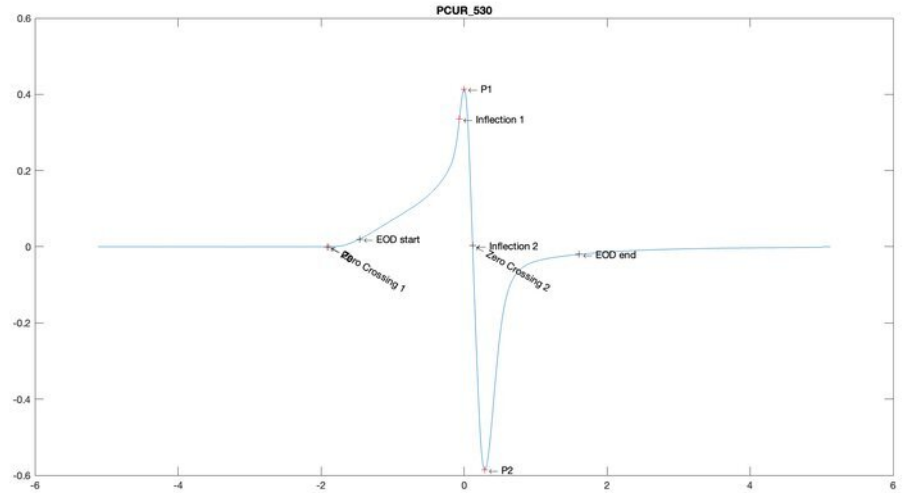
PCUR_530
P1
Inflection 1
EOD start
Inflection 2
EOD end
Zero Crossing 1
Zero Crossing 2
P2

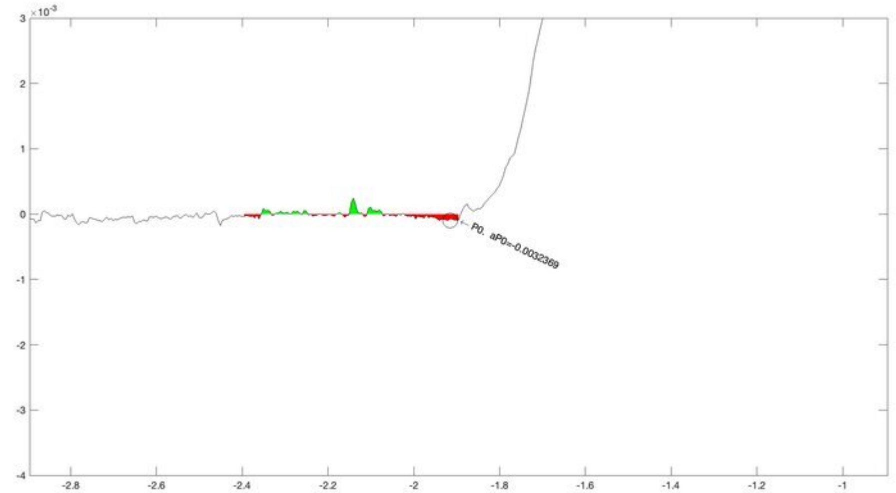
P0. aP0=-0.0032369

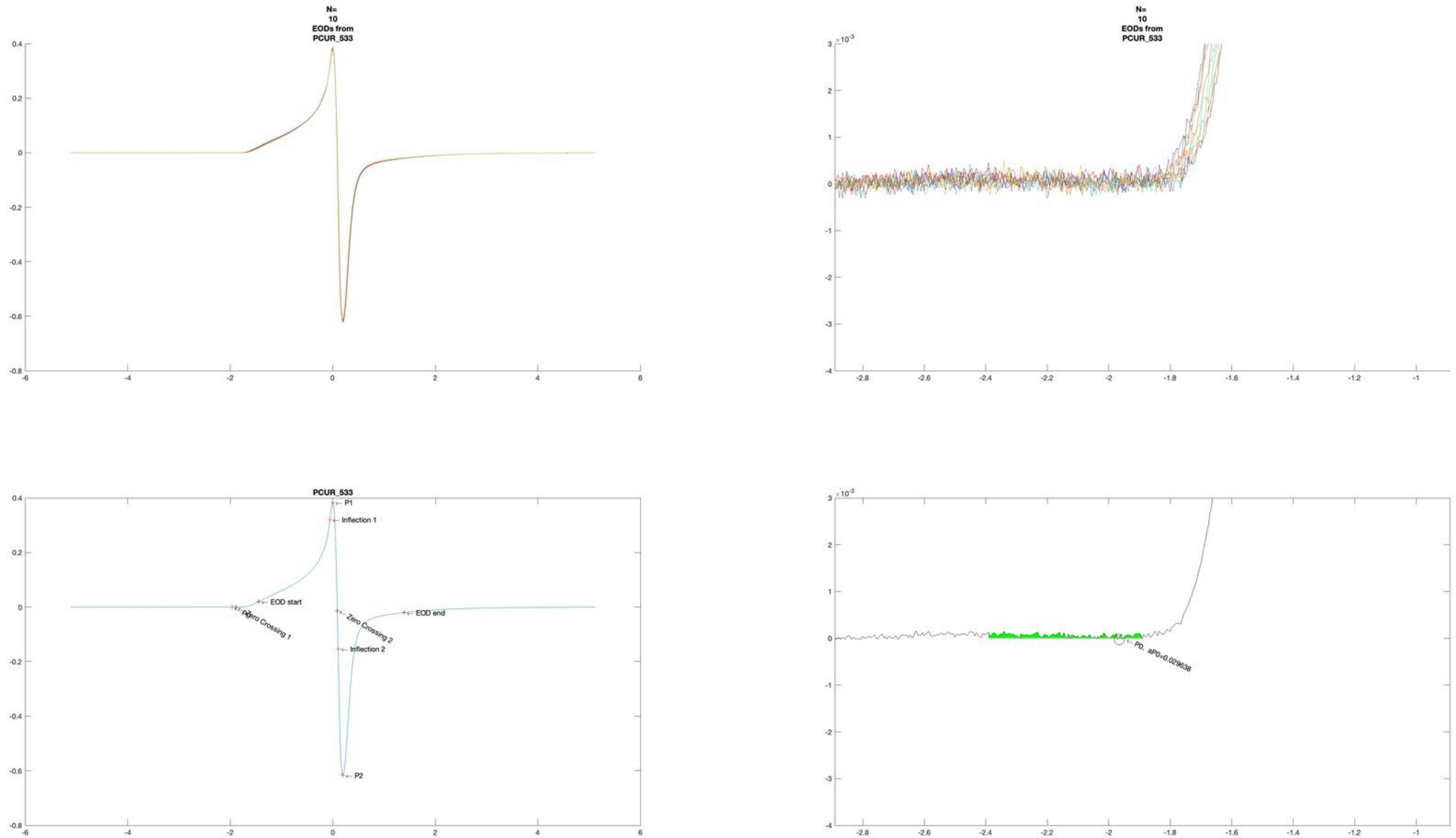

N=
10
EODs from
PCUR_533
N=
10
EODs from
PCUR_533
PCUR_533
P1
Inflection 1
EOD start
P0
Zero Crossing 1
Zero Crossing 2
EOD end
Inflection 2
P2
P0, aP0=0.029638

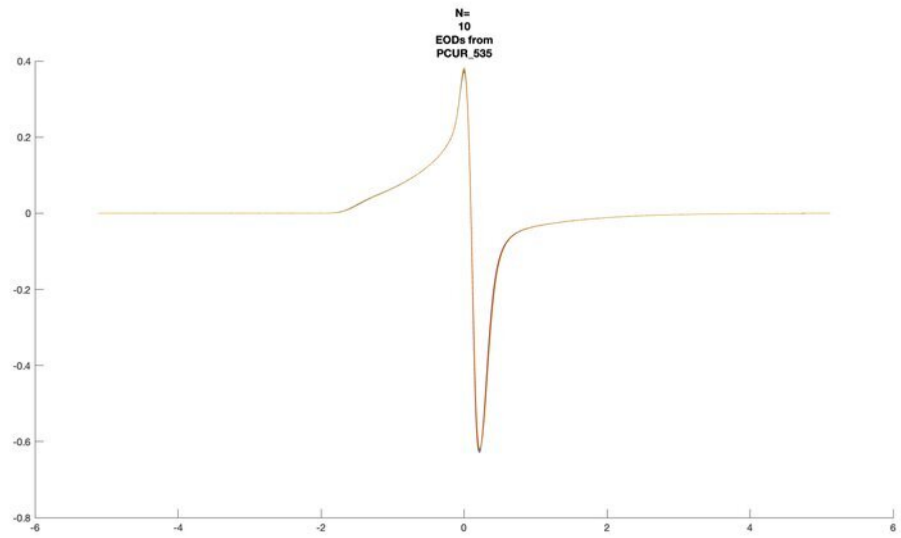
N=
10
EODs from
PCUR_535

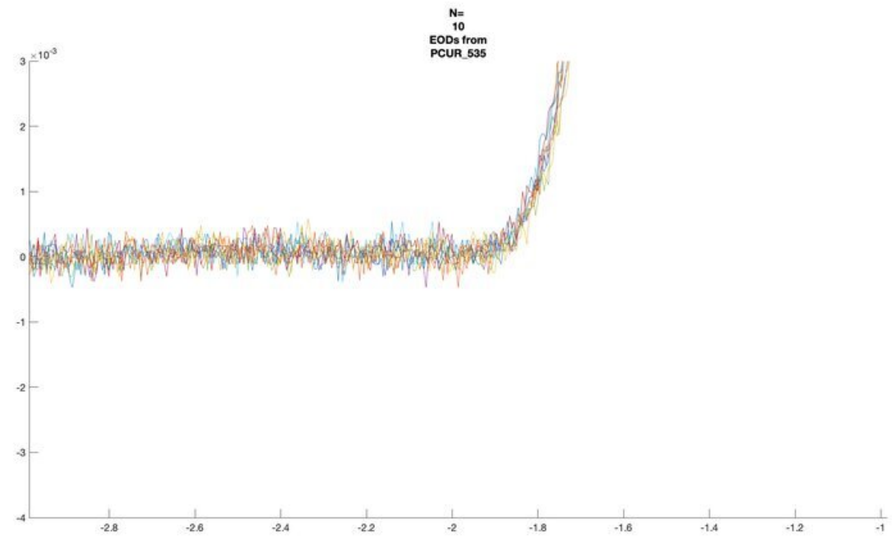
N=
10
EODs from
PCUR_535
$\times 10^{-3}$

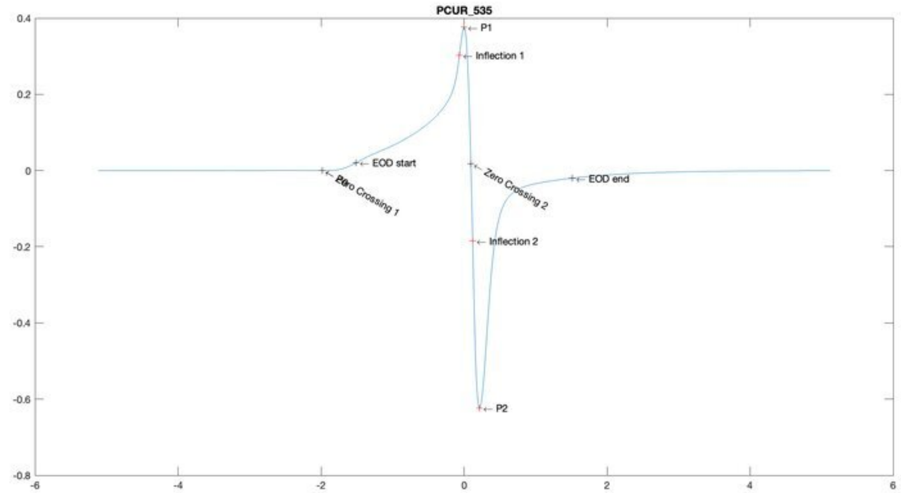
PCUR_535
P1
Inflection 1
EOD start
Zero Crossing 1
Zero Crossing 2
EOD end
Inflection 2
P2

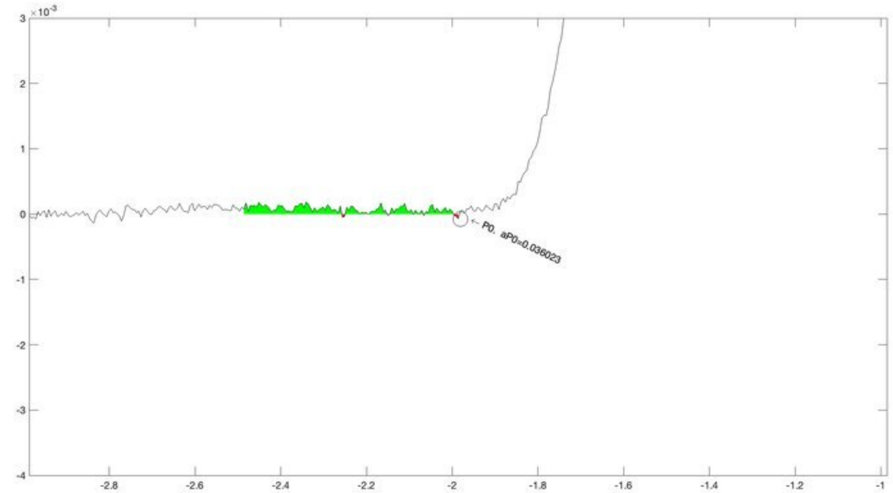
$\times 10^{-3}$
P0, aP0=0.036023

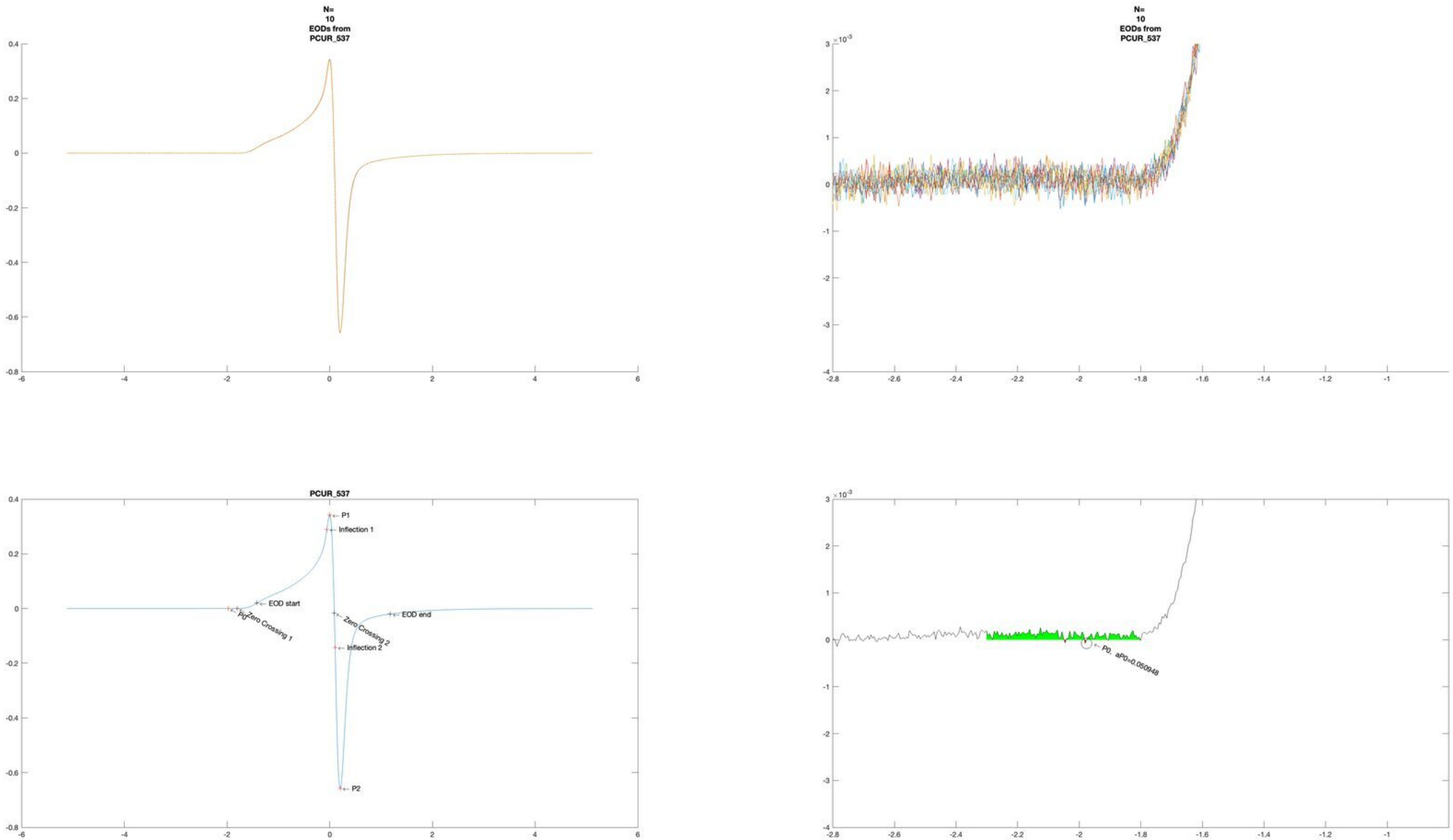
N=
10
EODs from
PCUR_537
N=
10
EODs from
PCUR_537
PCUR_537
P1
Inflection 1
EOD start
P0
Zero Crossing 1
Zero Crossing 2
EOD end
Inflection 2
P2
P0, aP0=0.050948

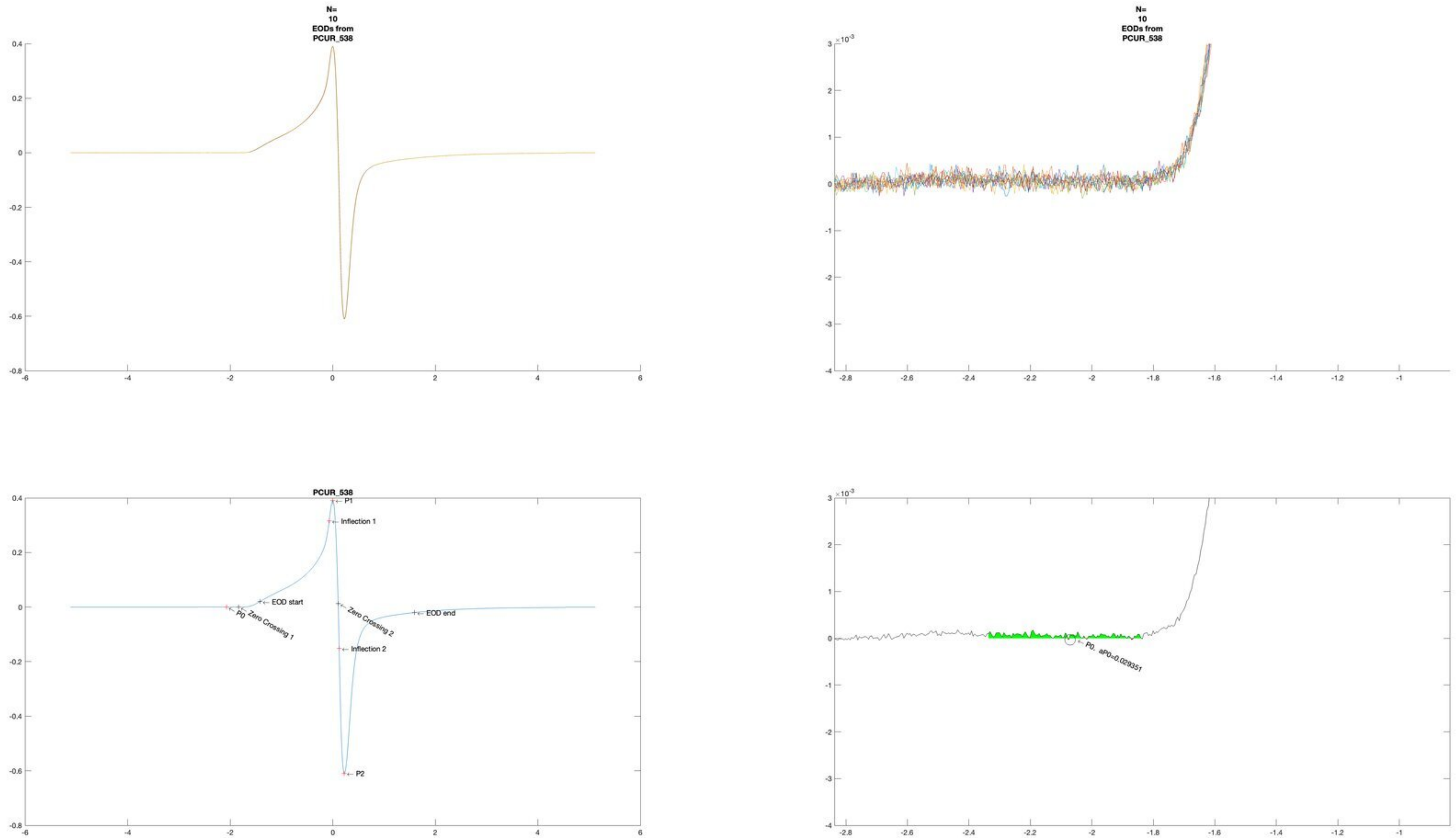
N=
10
EODs from
PCUR_538
N=
10
EODs from
PCUR_538
PCUR_538
P1
Inflection 1
EOD start
P0
Zero Crossing 1
Zero Crossing 2
EOD end
Inflection 2
P2
P0, aP0=0.029351

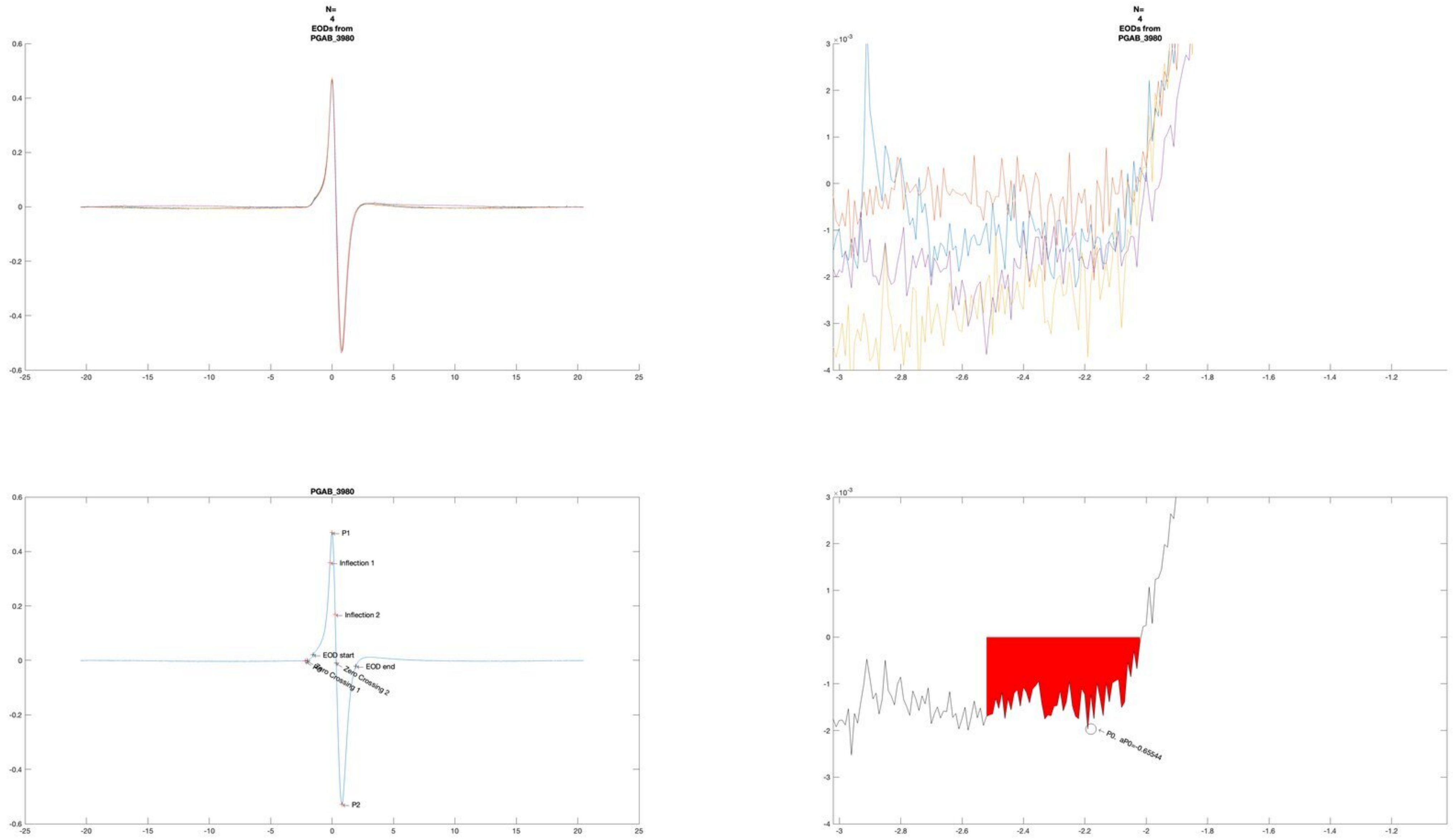
N=
4
EODs from
PGAB_3980
N=
4
EODs from
PGAB_3980
PGAB_3980
P1
Inflection 1
Inflection 2
EOD start
EOD end
Zero Crossing 1
Zero Crossing 2
P2
P0, aP0=-0.65544

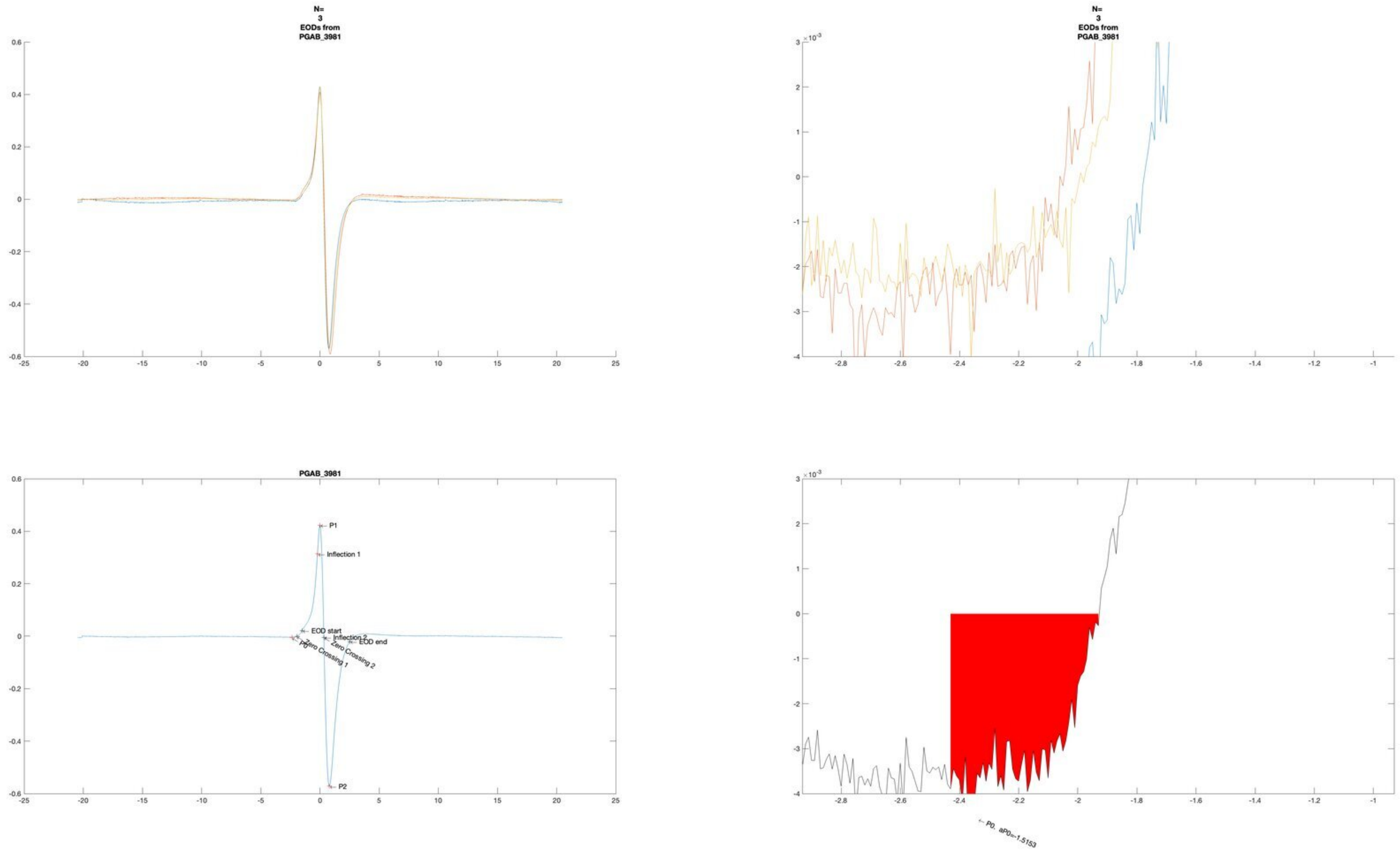
N=
3
EODs from
PGAB_3981
PGAB_3981
P1
Inflection 1
EOD start
Inflection 2
EOD end
P0
Zero Crossing 1
Zero Crossing 2
P2
← P0, aP0=-1.5153

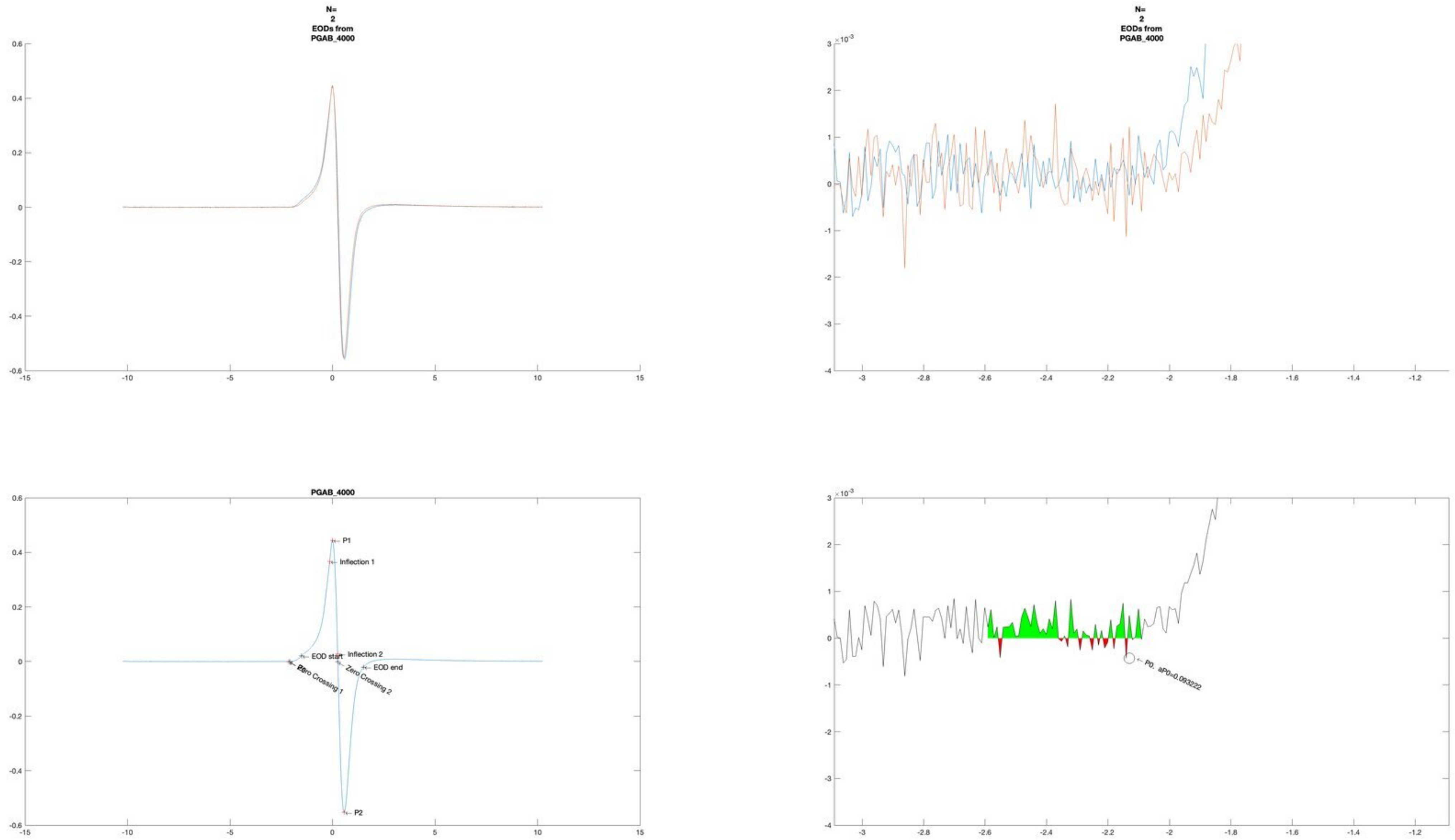
N=
2
EODs from
PGAB_4000
N=
2
EODs from
PGAB_4000
PGAB_4000
P1
Inflection 1
EOD start
Inflection 2
Zero Crossing 1
Zero Crossing 2
EOD end
P2
P0, aP0=0.093222

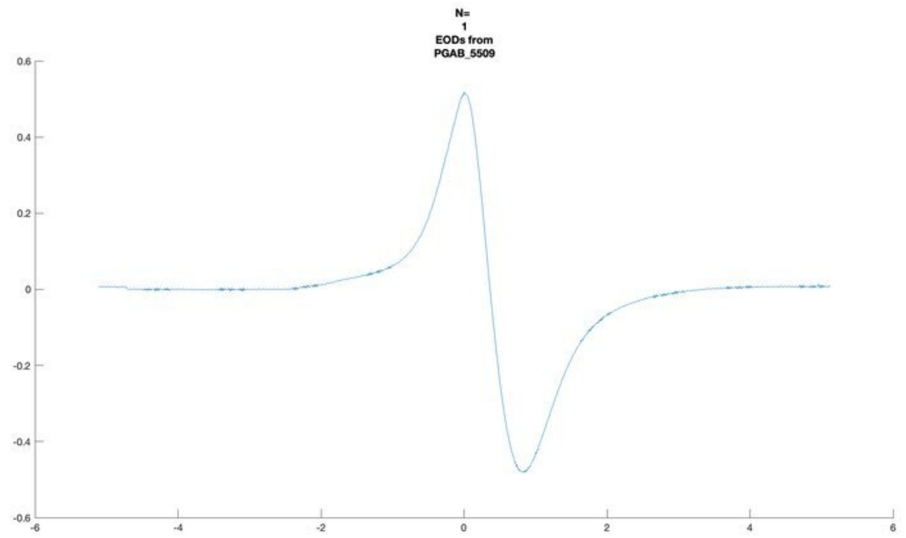
N=
1
EODs from
PGAB_5509

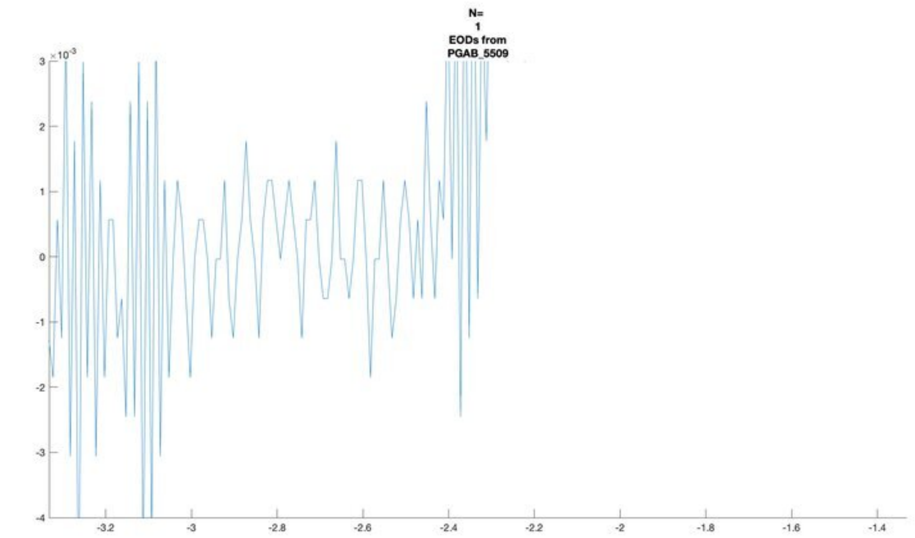
N=
1
EODs from
PGAB_5509
×10^-3

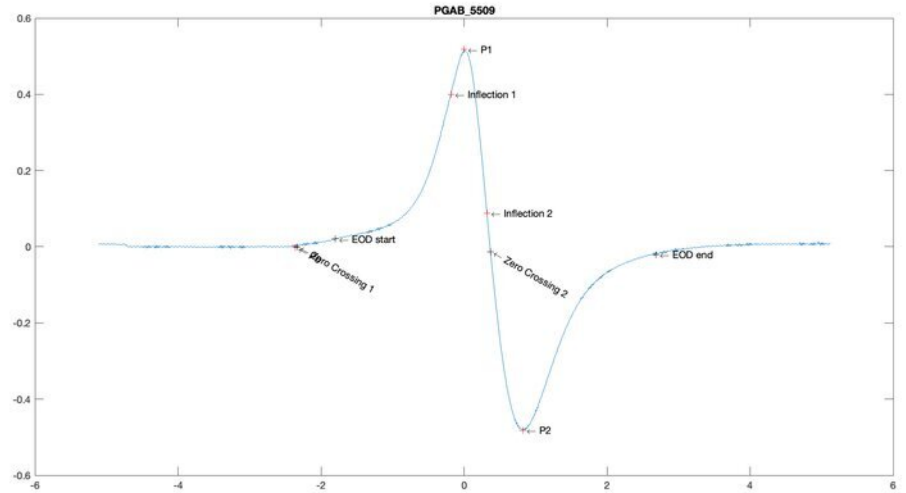
PGAB_5509
P1
Inflection 1
Inflection 2
EOD start
Zero Crossing 1
Zero Crossing 2
EOD end
P2

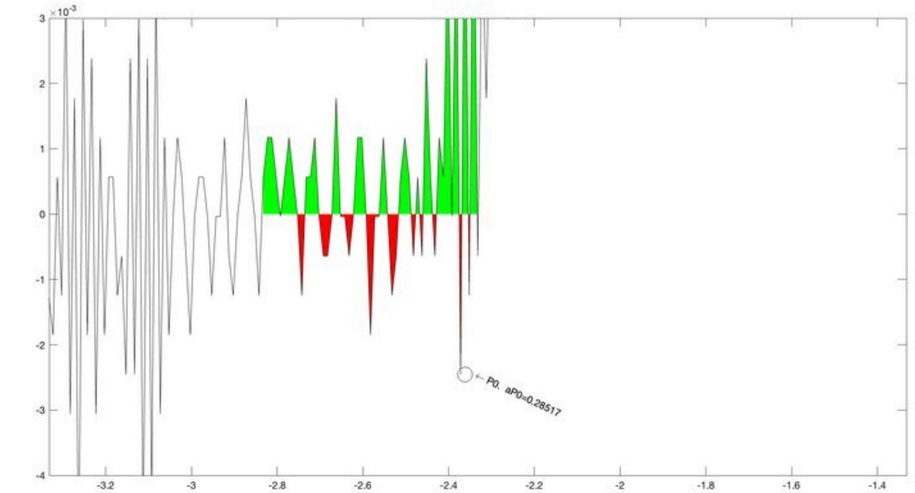
×10^-3
P0. aP0=0.28517

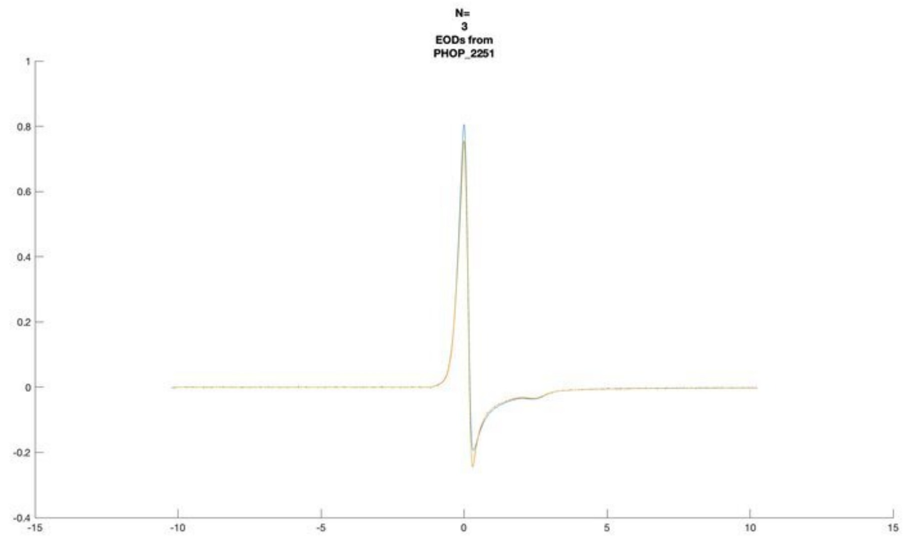
N=
3
EODs from
PHOP_2251

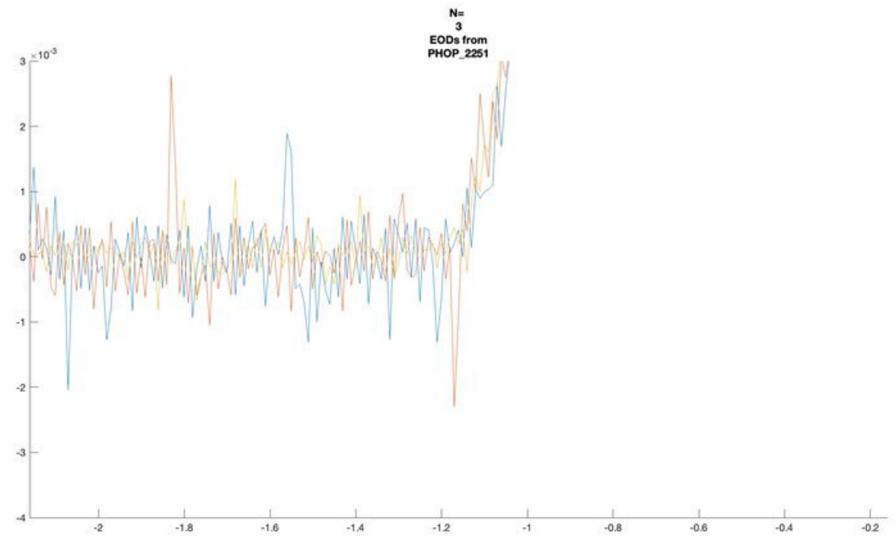
N=
3
EODs from
PHOP_2251
×10⁻³

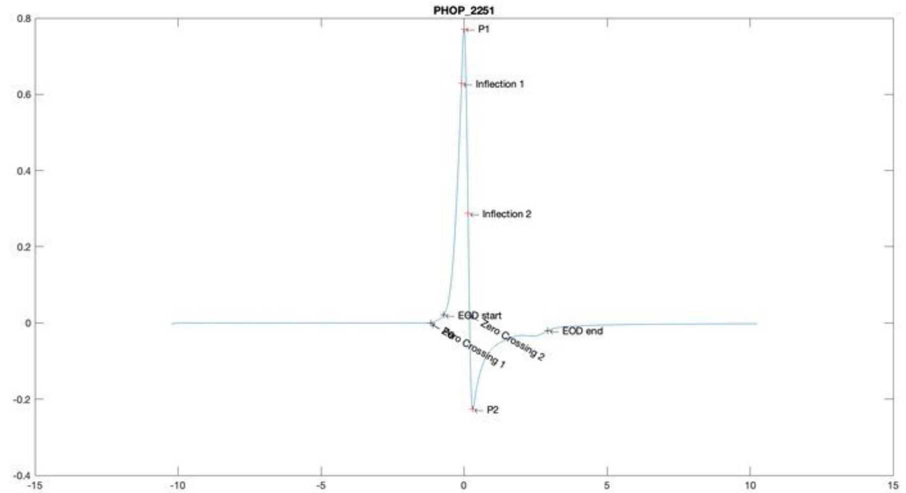
PHOP_2251
P1
Inflection 1
Inflection 2
EOD start
Zero Crossing 1
Zero Crossing 2
EOD end
P2

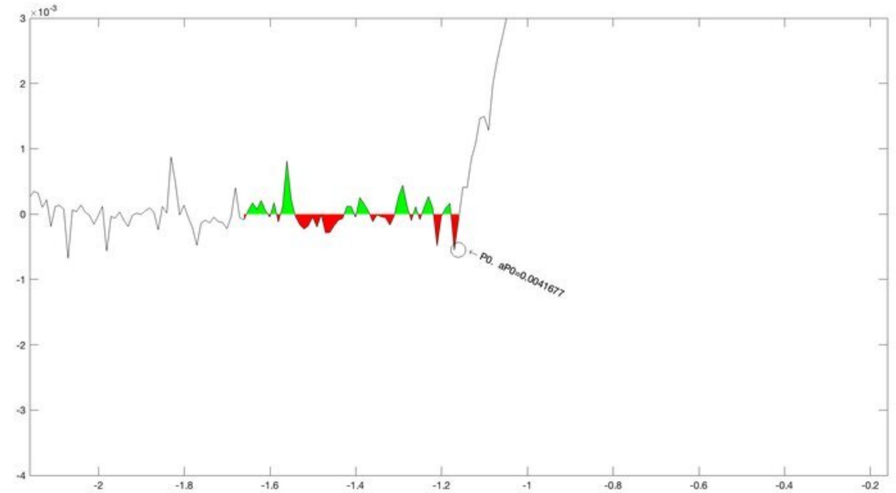
×10⁻³
P0, aP0=0.0041677

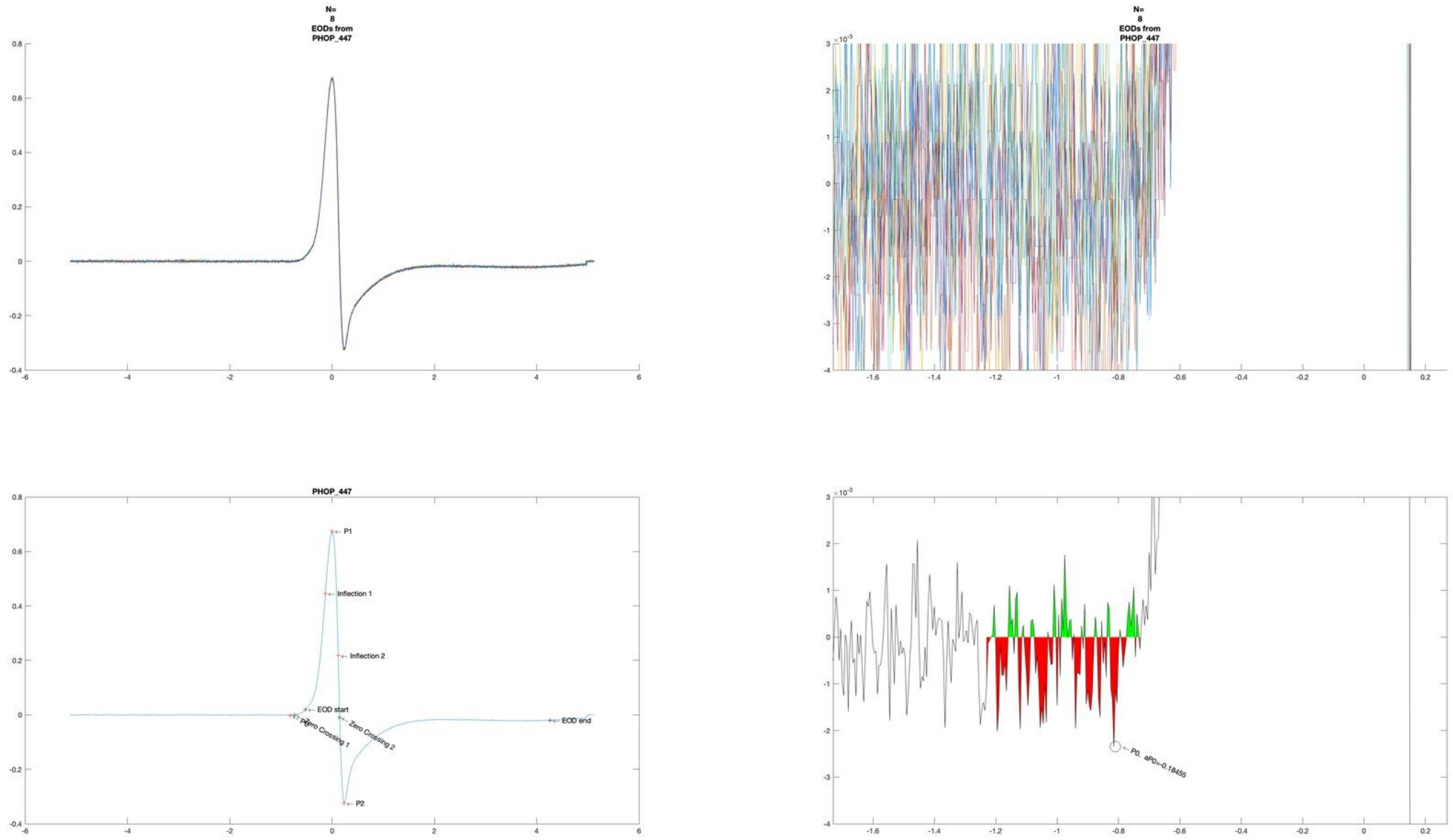
N=
8
EODs from
PHOP_447
N=
8
EODs from
PHOP_447
PHOP_447
P1
Inflection 1
Inflection 2
EOD start
Zero Crossing 1
Zero Crossing 2
P0
P2
EOD end
P0, aP0=-0.18455

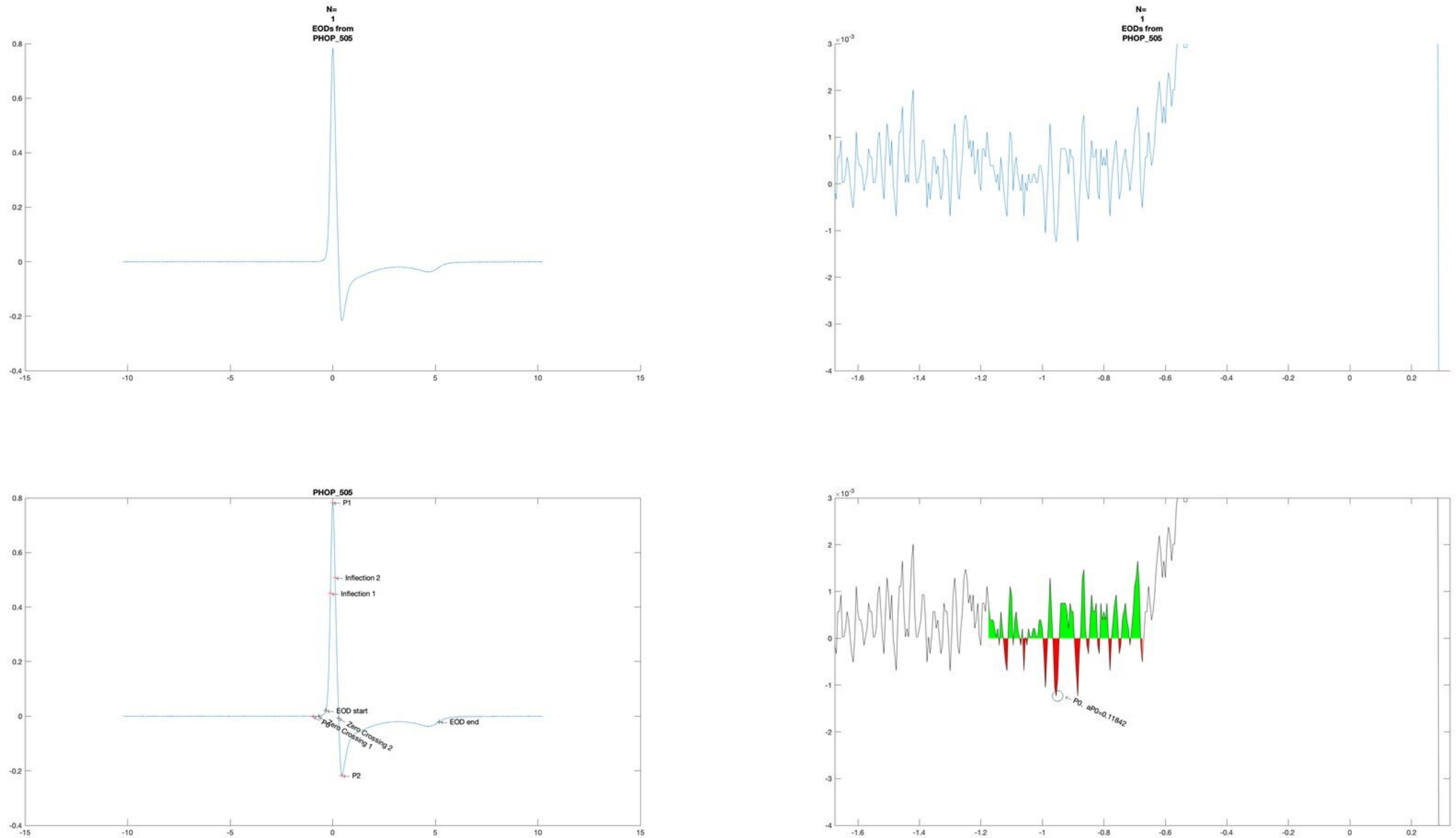

N=
1
EODs from
PHOP_505
PHOP_505
P1
Inflection 2
Inflection 1
EOD start
EOD end
Zero Crossing 2
Zero Crossing 1
P0
P2
P0. aP0=0.11842

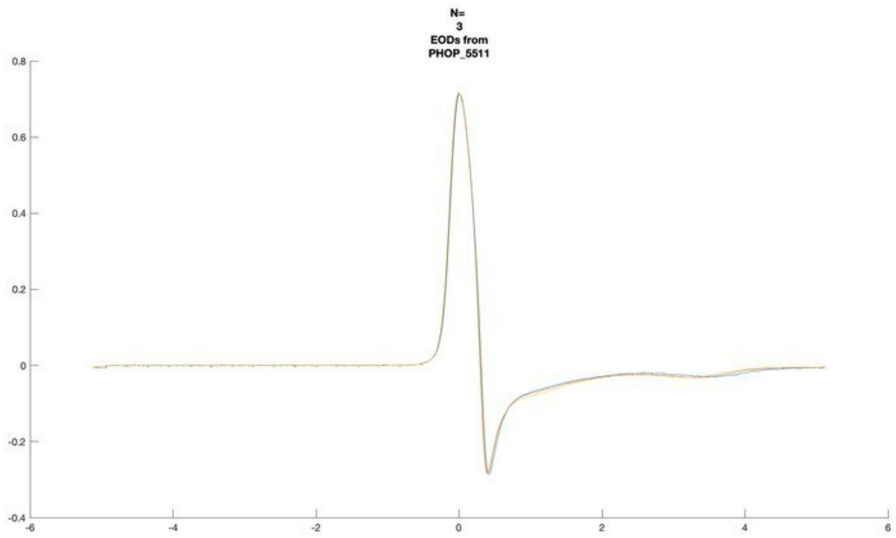
N=
3
EODs from
PHOP_5511

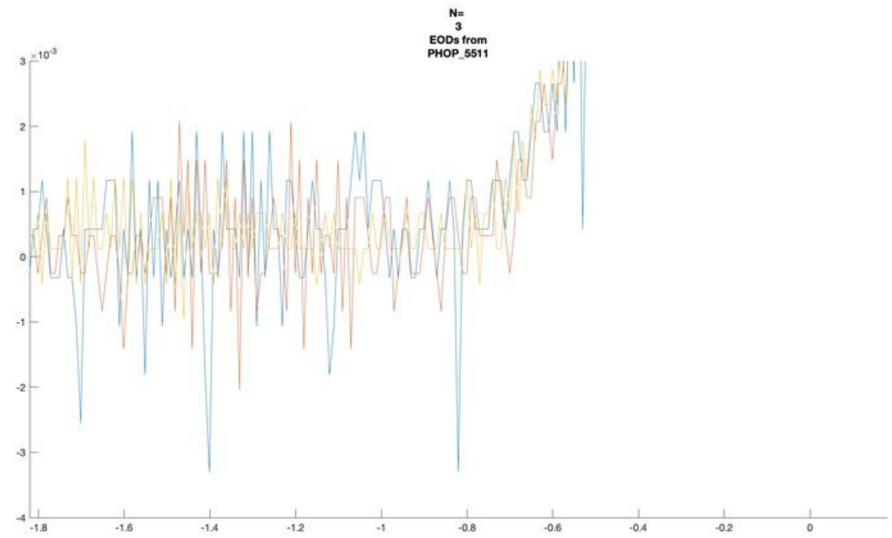
N=
3
EODs from
PHOP_5511

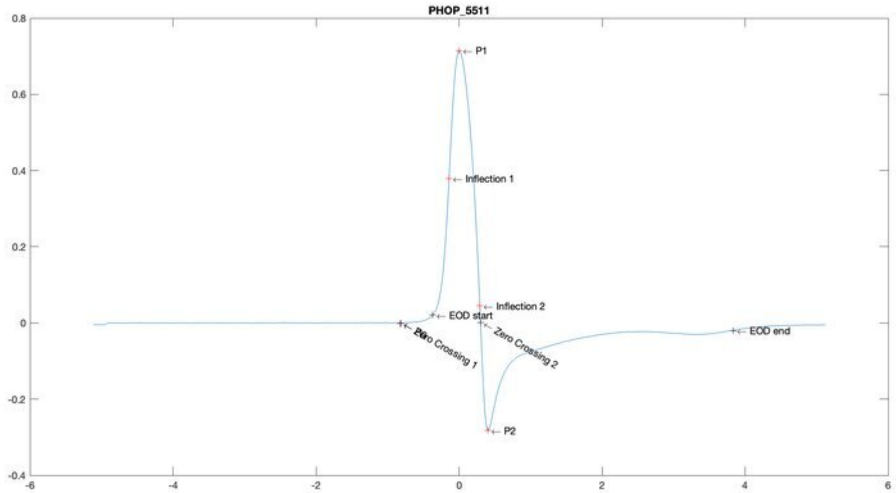
PHOP_5511
P1
Inflection 1
Inflection 2
EOD start
Zero Crossing 1
Zero Crossing 2
EOD end
P2

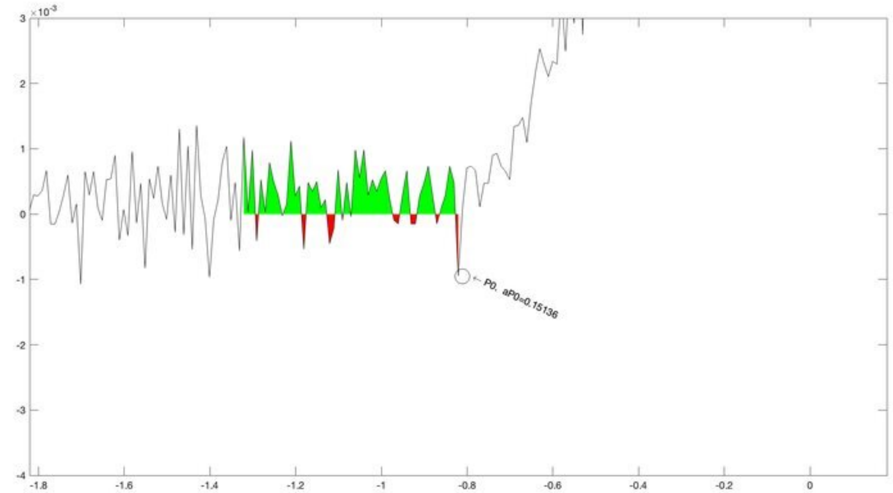
P0. aP0=0.15136

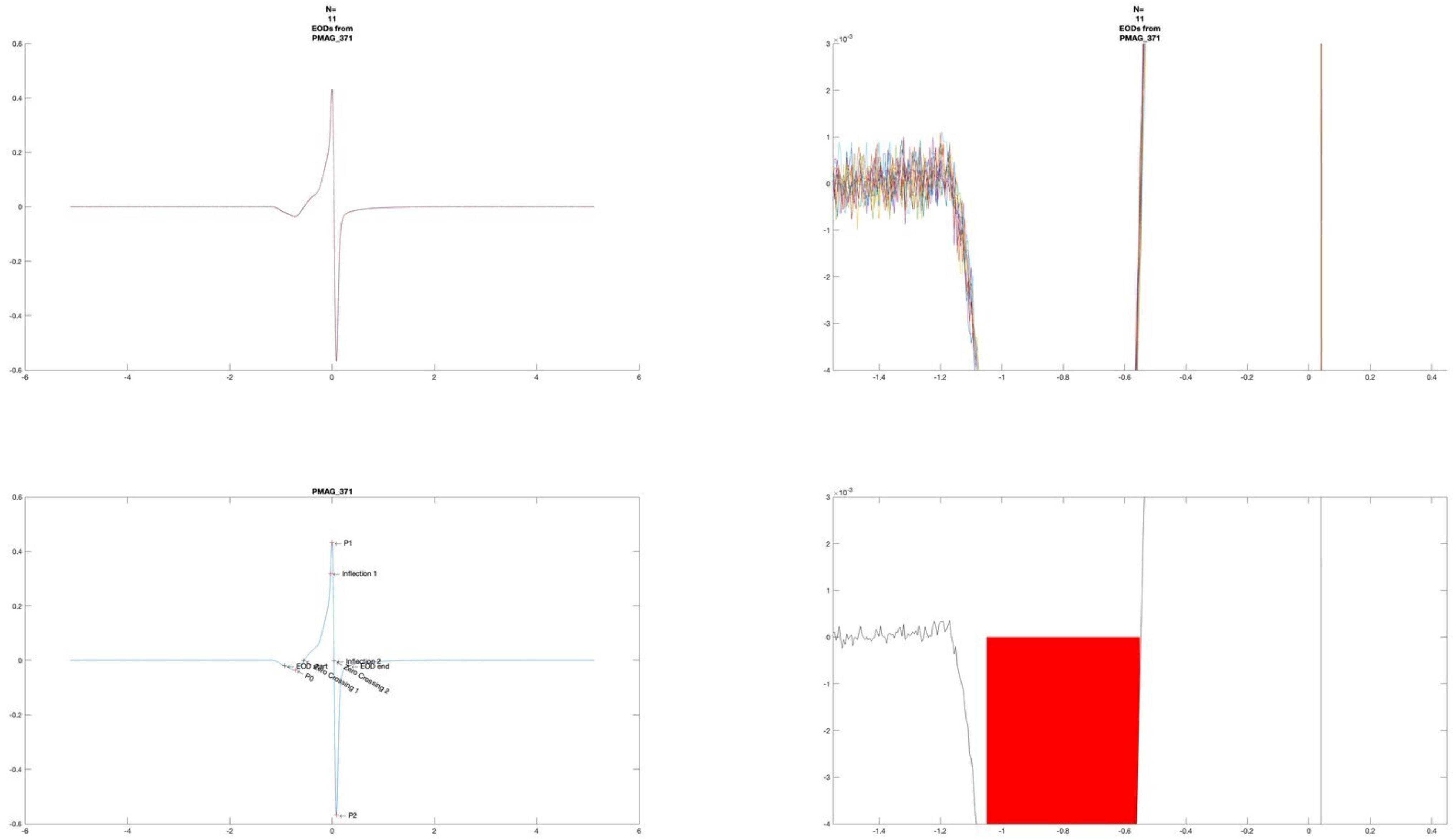
N=
11
EODs from
PMAG_371
N=
11
EODs from
PMAG_371
PMAG_371
P1
Inflection 1
Inflection 2
EOD start
EOD end
P0
Zero Crossing 1
Zero Crossing 2
P2

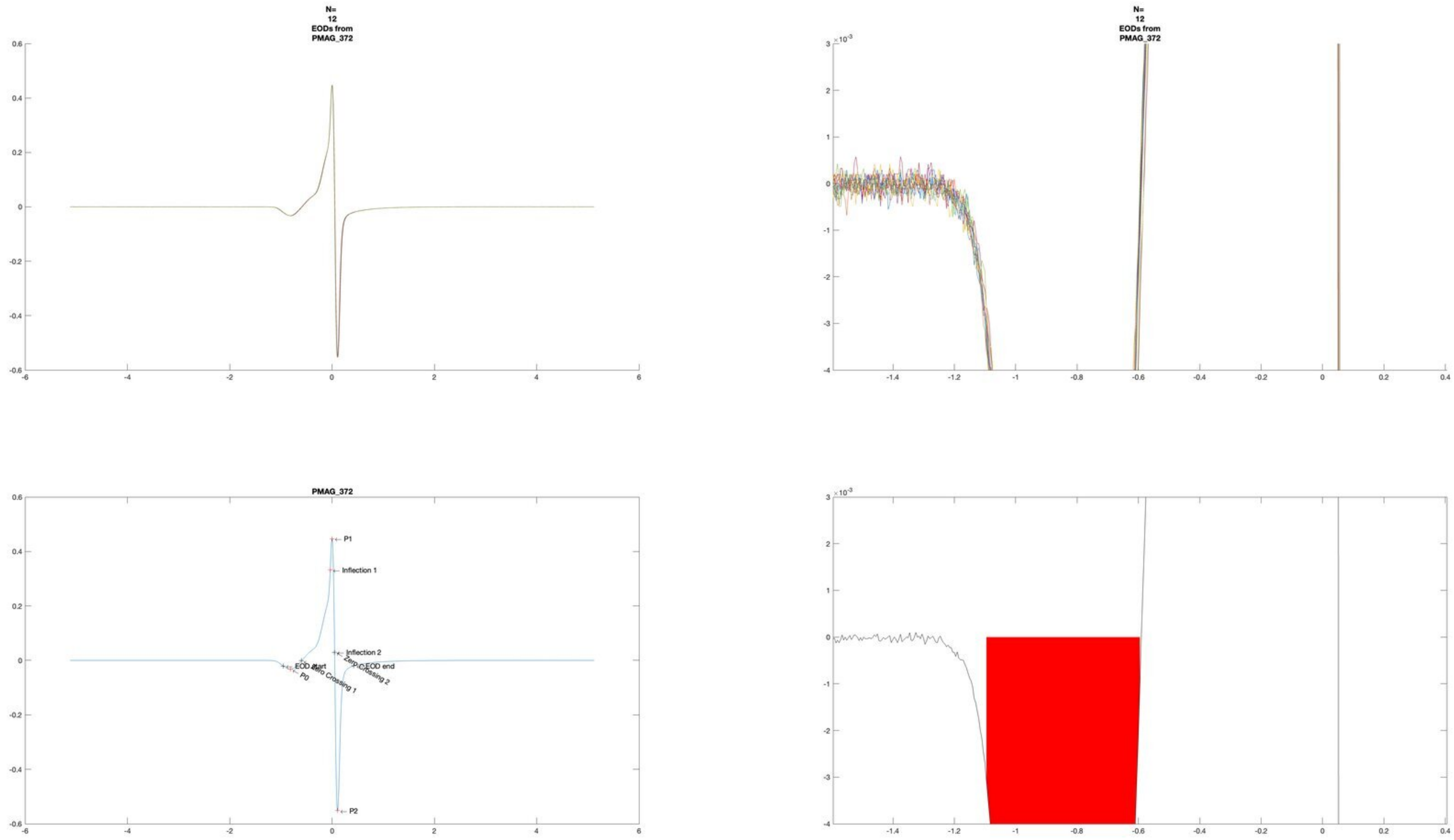
N=
12
EODs from
PMAG_372
N=
12
EODs from
PMAG_372
PMAG_372
P1
Inflection 1
Inflection 2
EOD start
Zero Crossing 1
Zero Crossing 2
EOD end
P0
P2

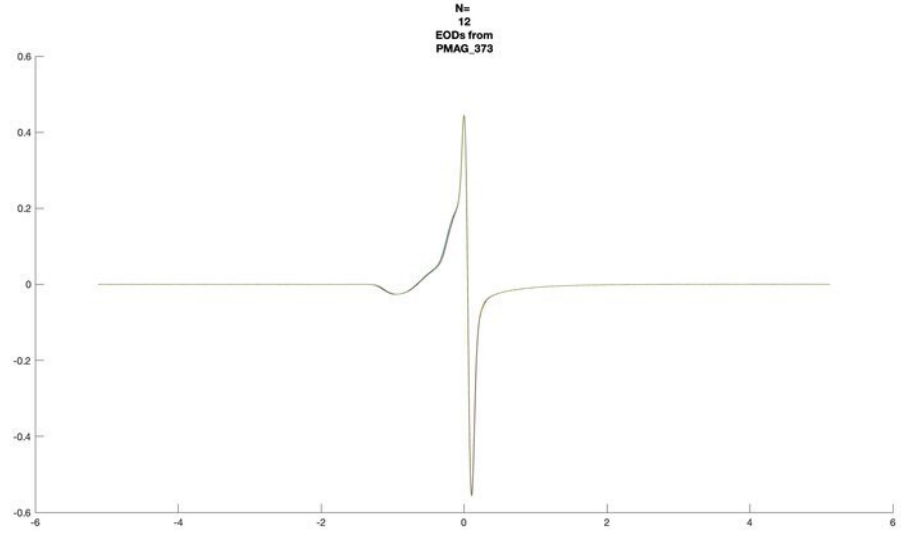
N=
12
EODs from
PMAG_373

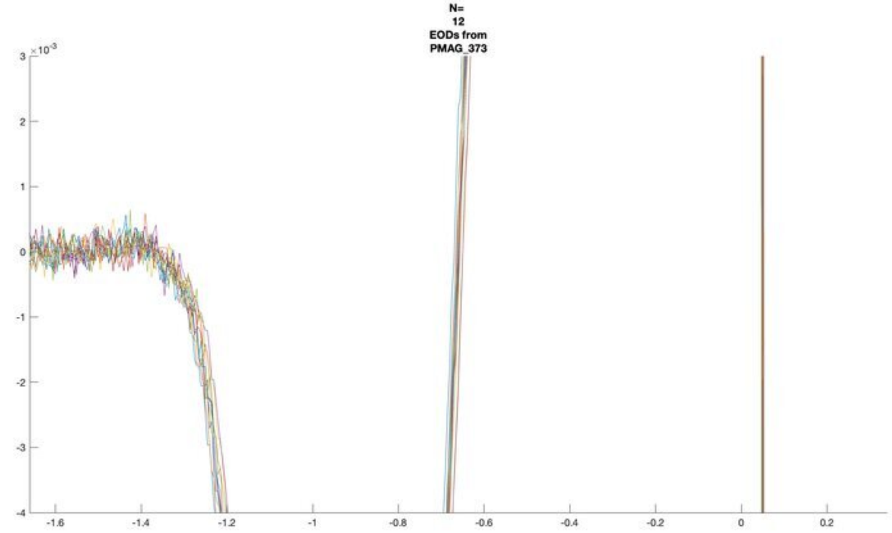
N=
12
EODs from
PMAG_373

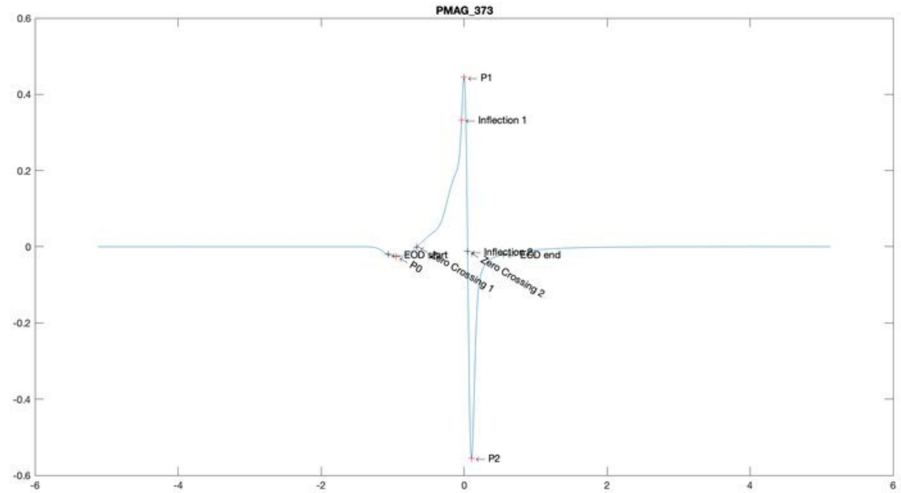
PMAG_373
P1
Inflection 1
EOD start
P0
Zero Crossing 1
Inflection 2
EOD end
Zero Crossing 2
P2

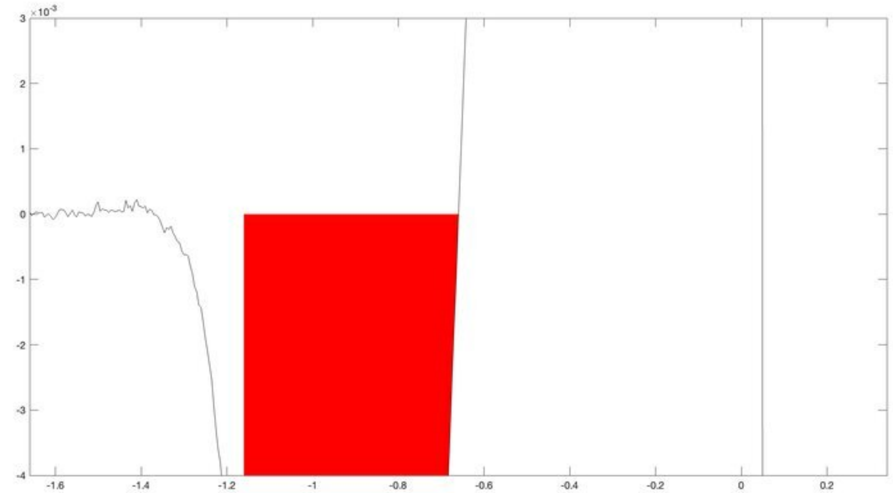

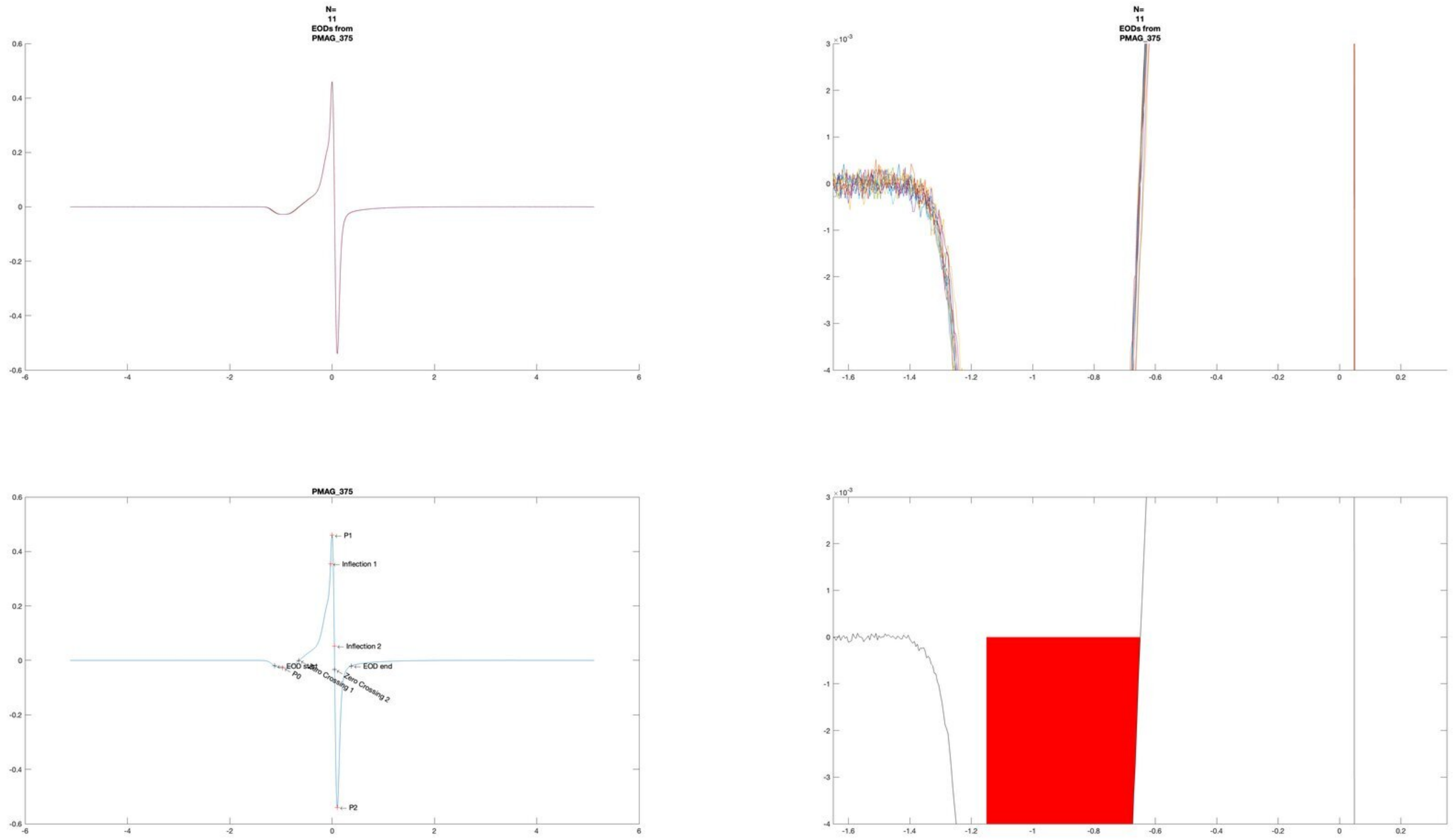
N=
11
EODs from
PMAG_375
N=
11
EODs from
PMAG_375
PMAG_375
P1
Inflection 1
Inflection 2
EOD start
EOD end
P0
Zero Crossing 1
Zero Crossing 2
P2

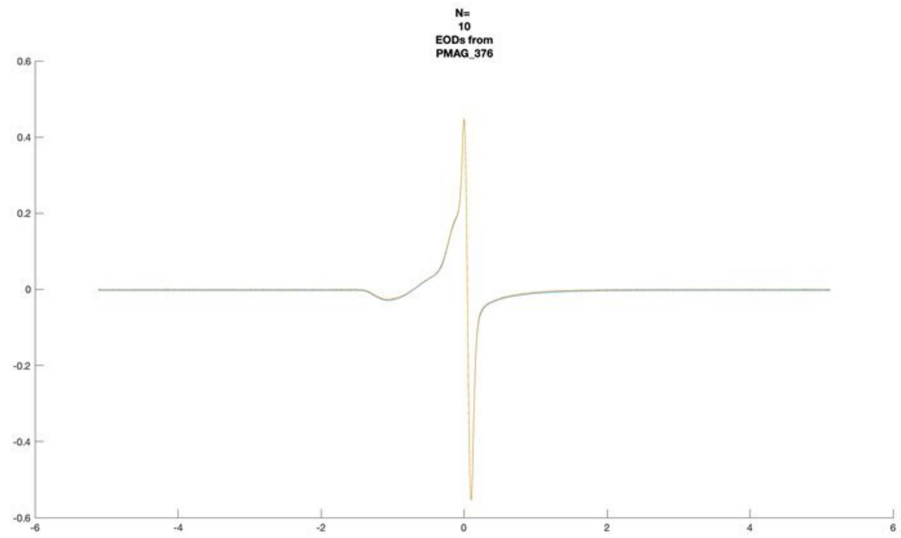
N=
10
EODs from
PMAG_376

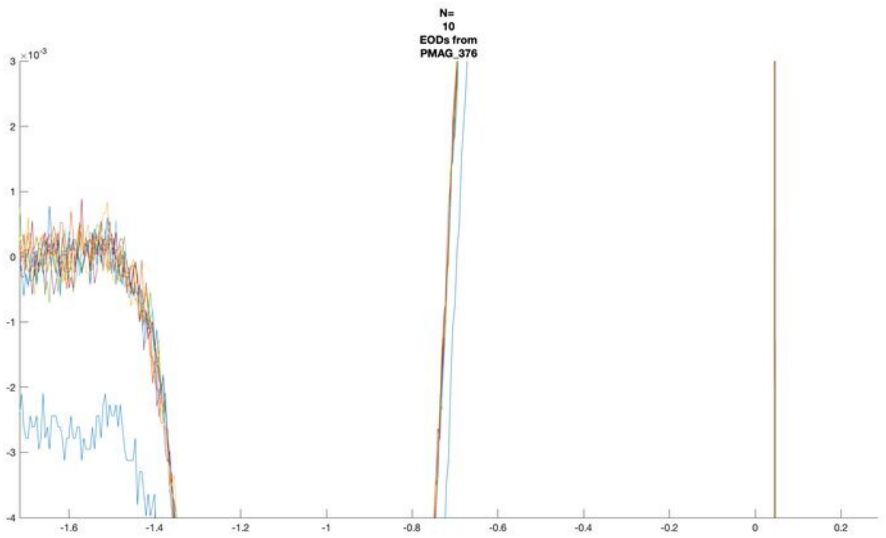
N=
10
EODs from
PMAG_376

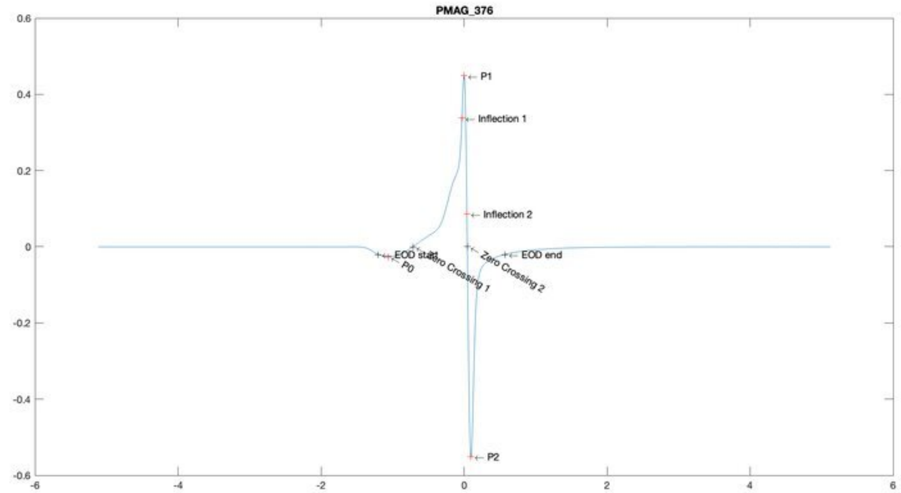
PMAG_376
P1
Inflection 1
Inflection 2
EOD start
P0
Zero Crossing 1
Zero Crossing 2
EOD end
P2

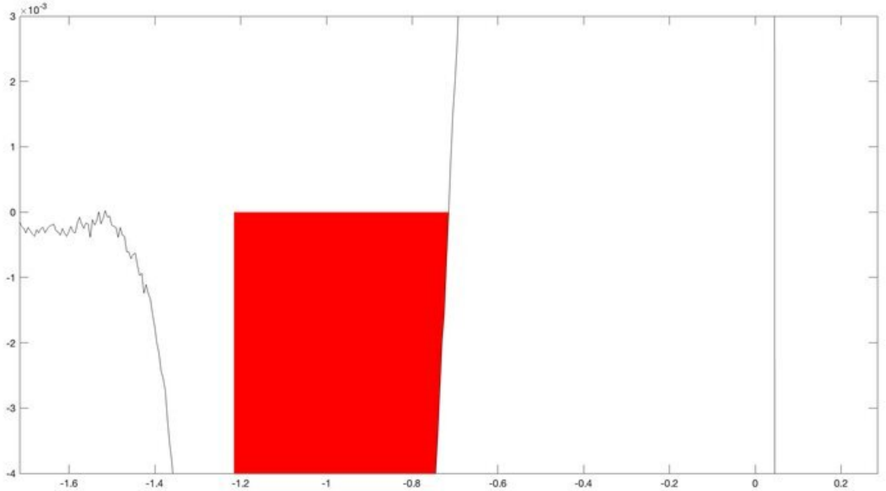

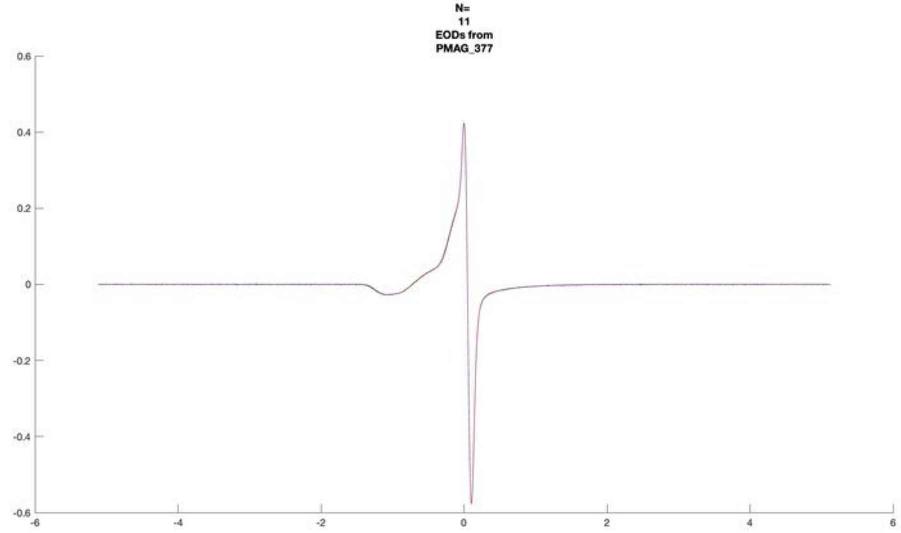
N=
11
EODs from
PMAG_377

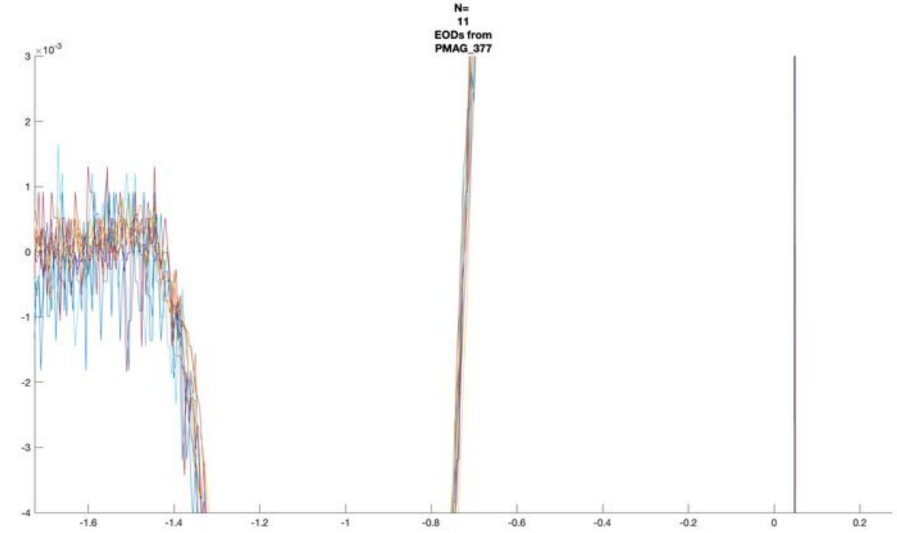
N=
11
EODs from
PMAG_377

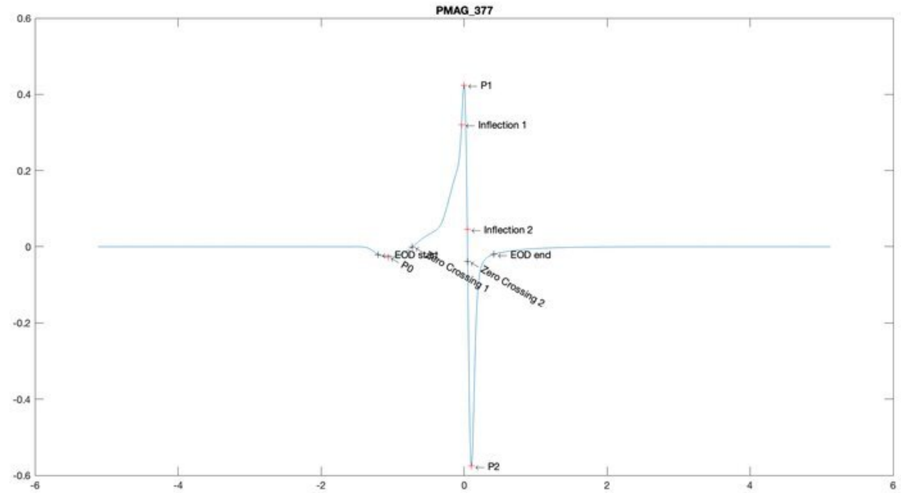
PMAG_377
P1
Inflection 1
Inflection 2
EOD start
P0
Zero Crossing 1
Zero Crossing 2
EOD end
P2

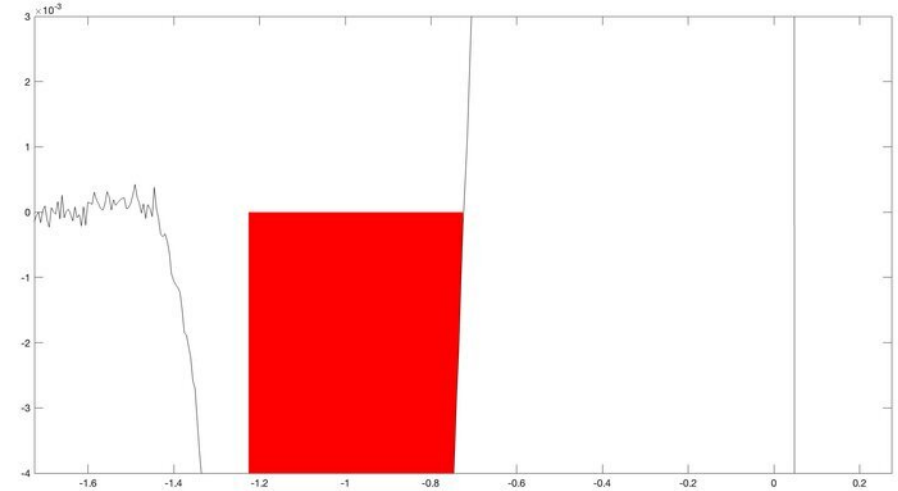

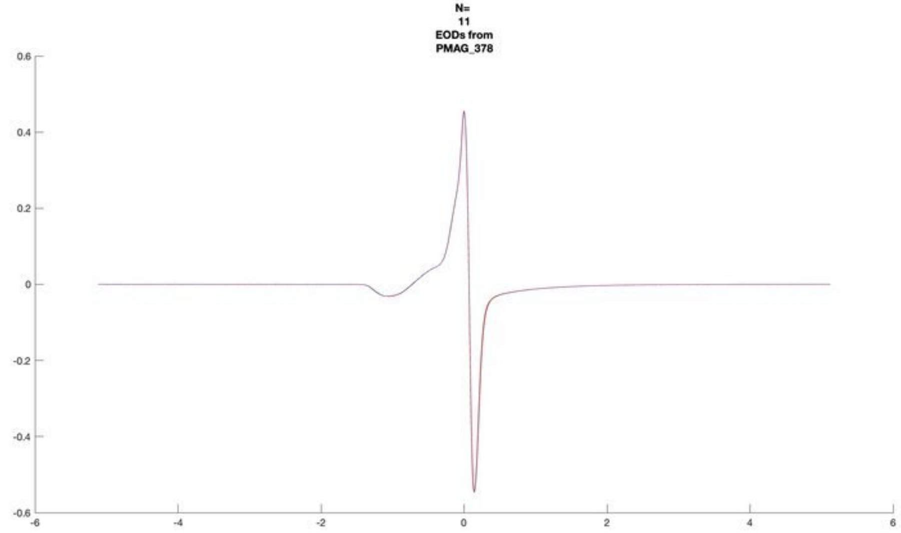
N=
11
EODs from
PMAG_378

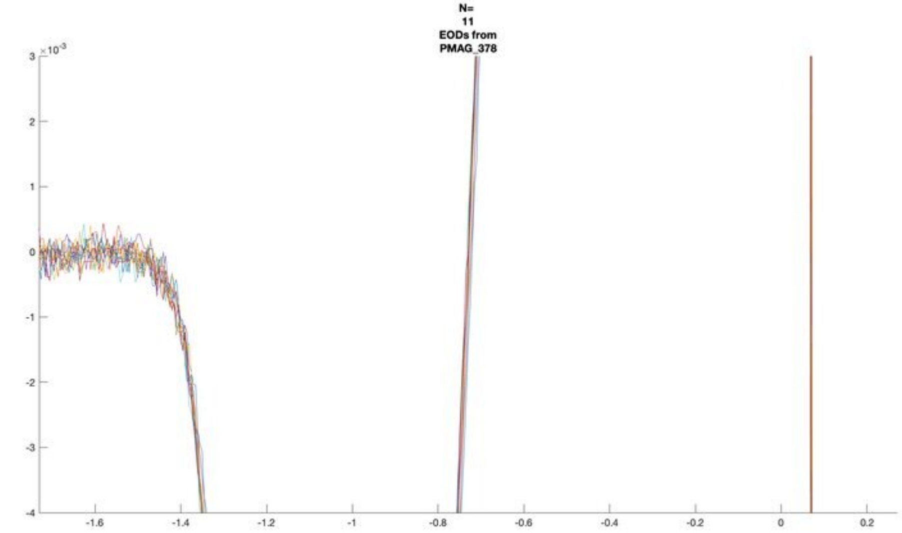
N=
11
EODs from
PMAG_378

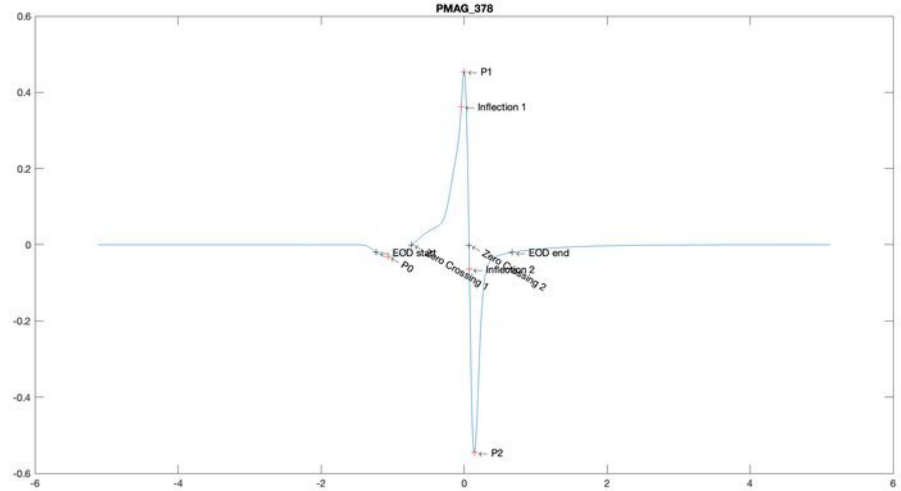
PMAG_378
P1
Inflection 1
EOD start
P0
Zero Crossing 1
Zero Crossing 2
Inflection 2
EOD end
P2

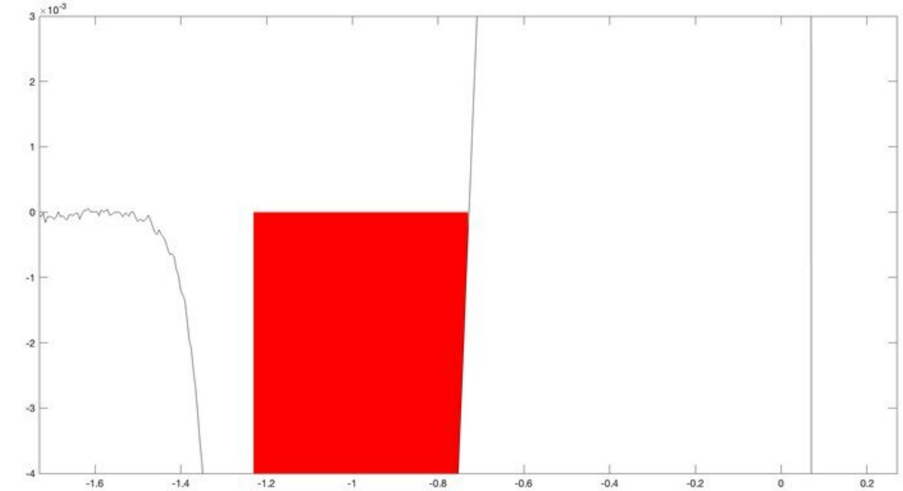

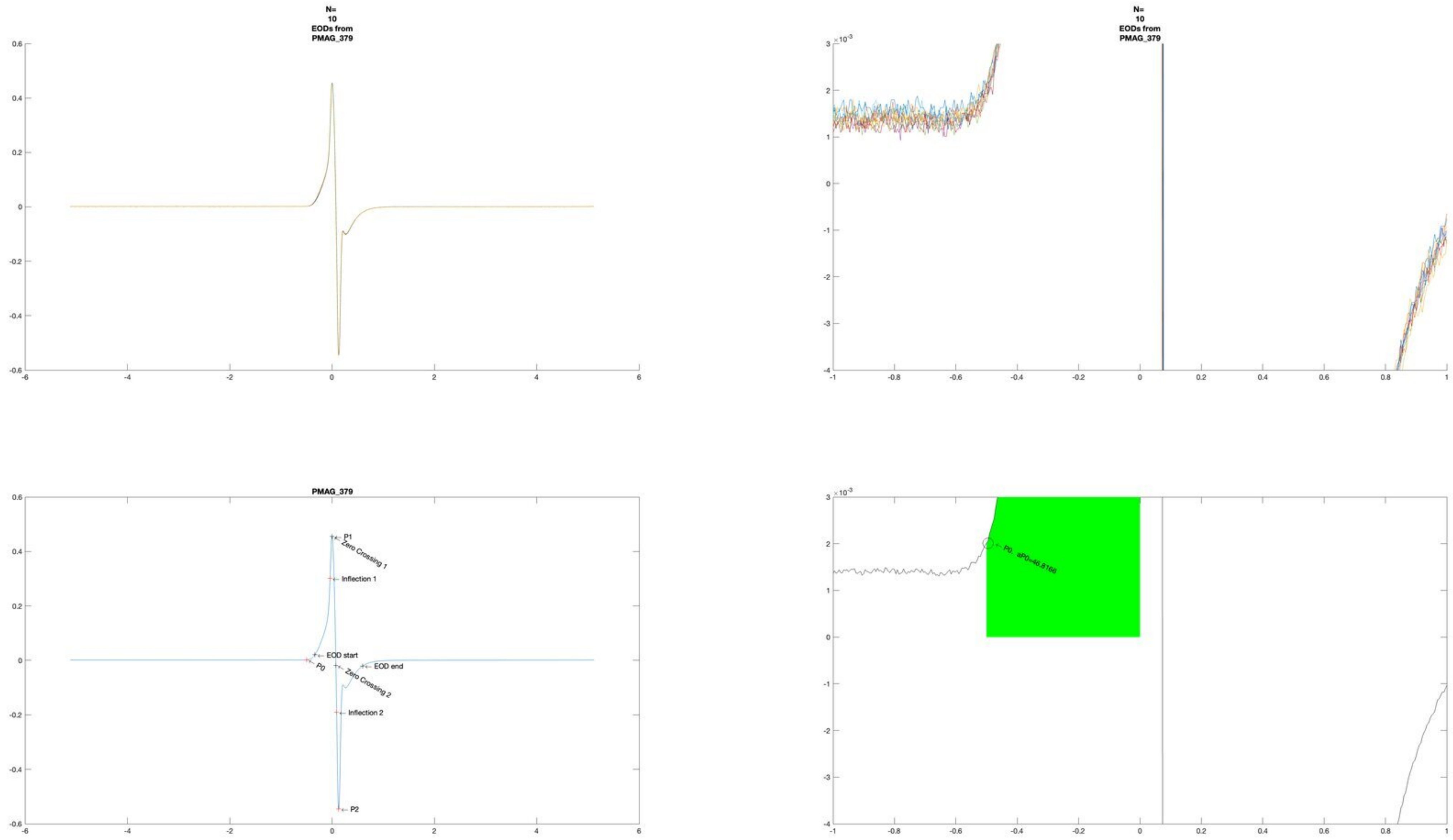

N=
10
EODs from
PMAG_379
N=
10
EODs from
PMAG_379
PMAG_379
P1
Zero Crossing 1
Inflection 1
EOD start
P0
EOD end
Zero Crossing 2
Inflection 2
P2
P0, aP0=46.8166

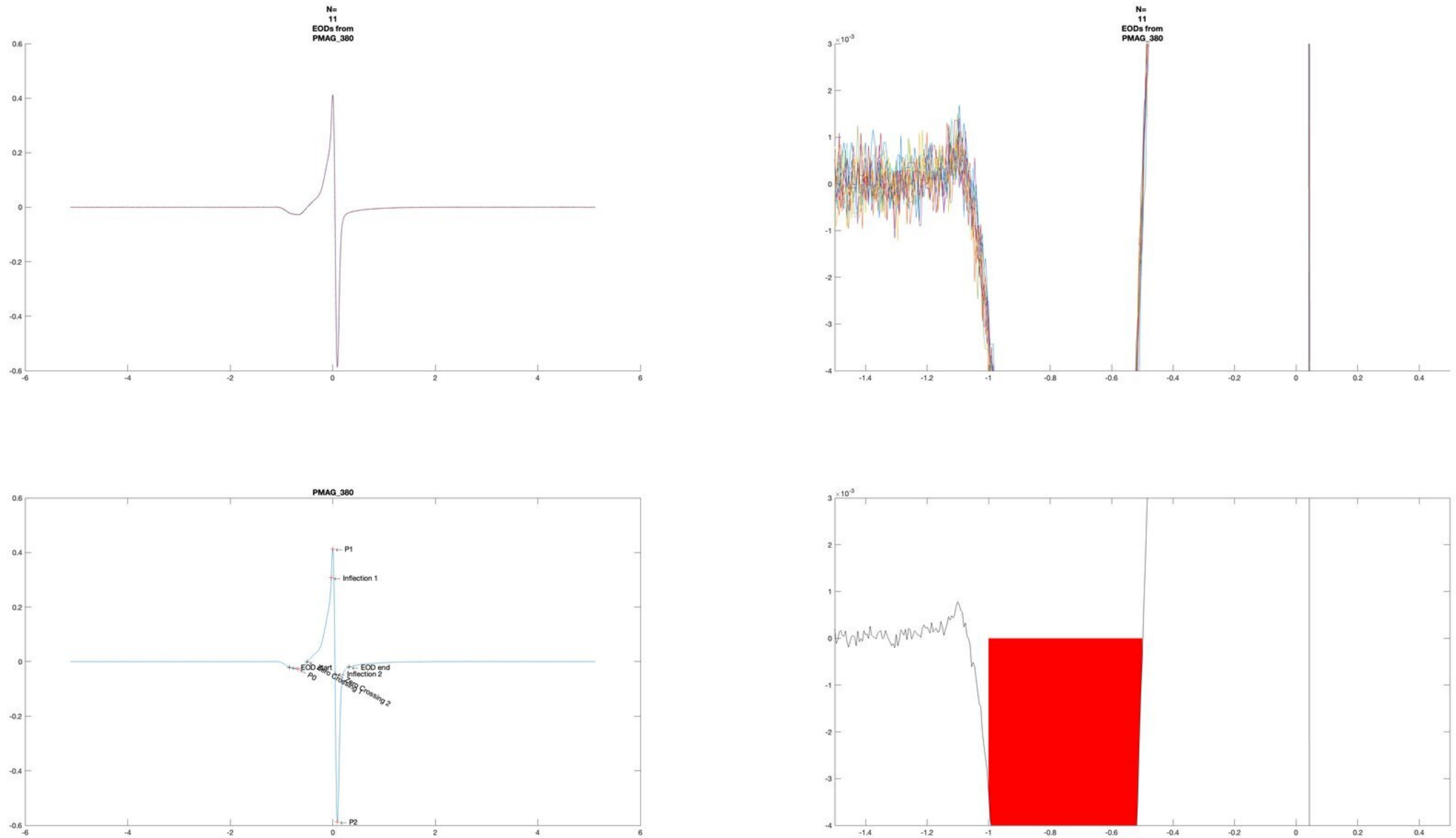
N=
11
EODs from
PMAG_380
N=
11
EODs from
PMAG_380
×10^-3
PMAG_380
P1
Inflection 1
EOD start
P0
Zero Crossing 1
EOD end
Inflection 2
Zero Crossing 2
P2
×10^-3

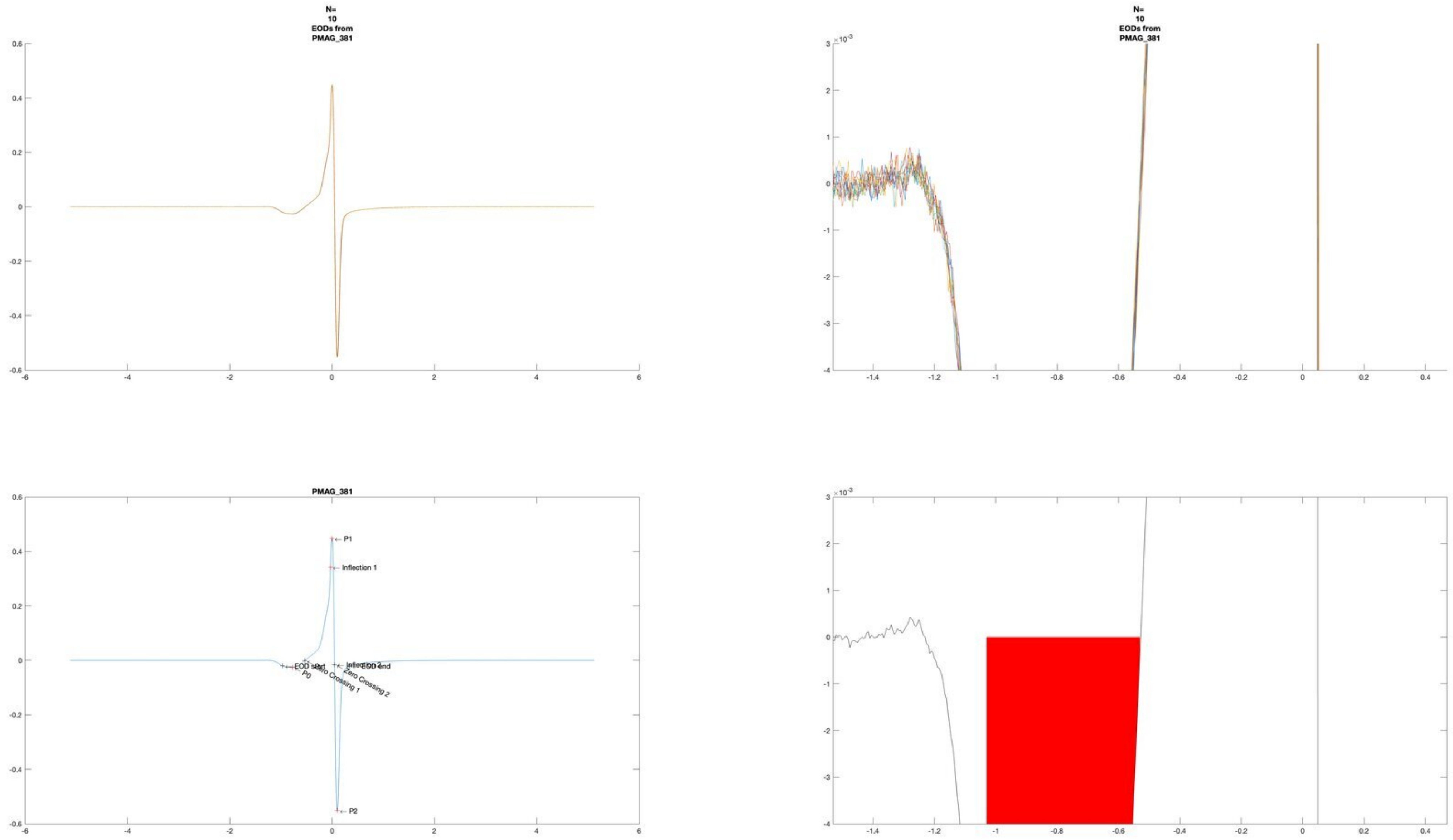
N=
10
EODs from
PMAG_381
N=
10
EODs from
PMAG_381
PMAG_381
P1
Inflection 1
EOD start
P0
Zero Crossing 1
Inflection 2
EOD end
Zero Crossing 2
P2

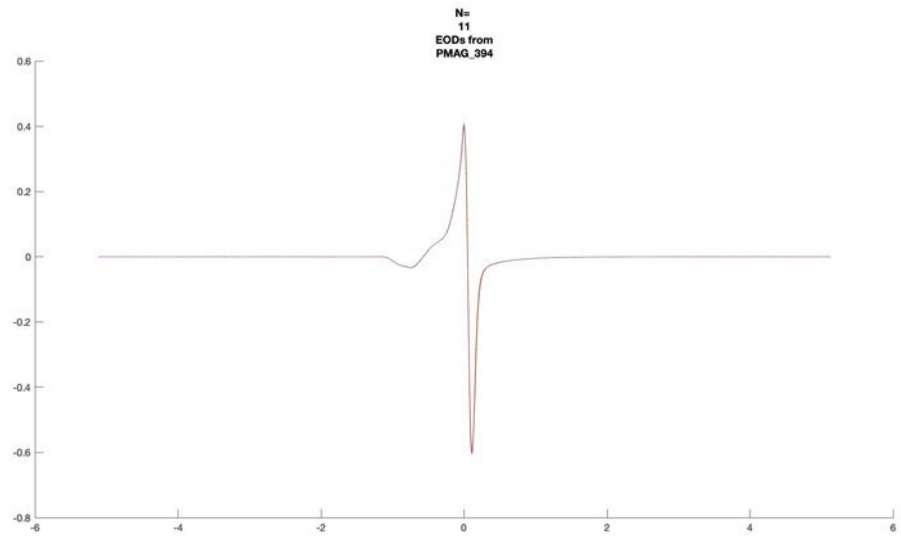
N=
11
EODs from
PMAG_394

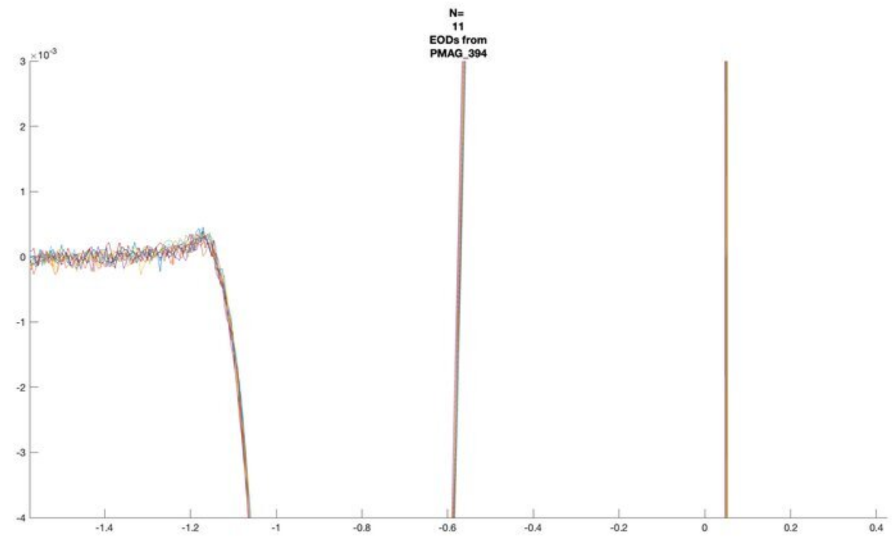
N=
11
EODs from
PMAG_394

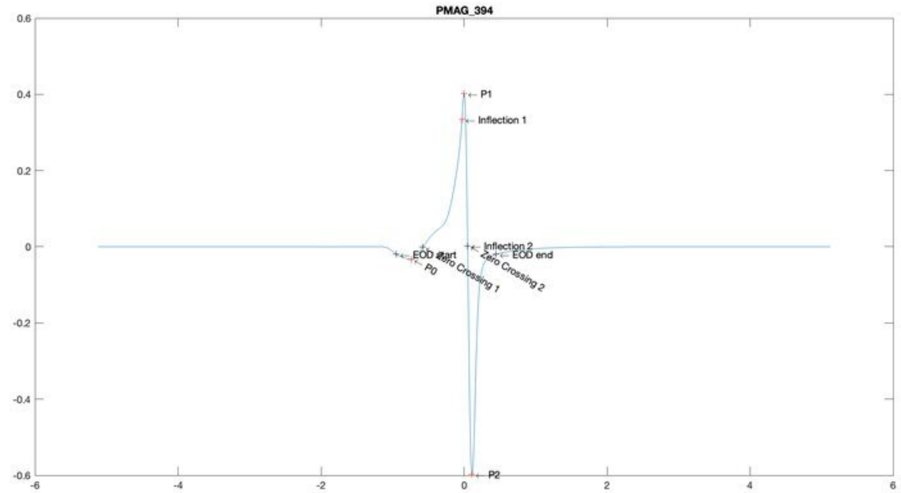
PMAG_394
P1
Inflection 1
Inflection 2
EOD start
EOD end
P0
Zero Crossing 1
Zero Crossing 2
P2

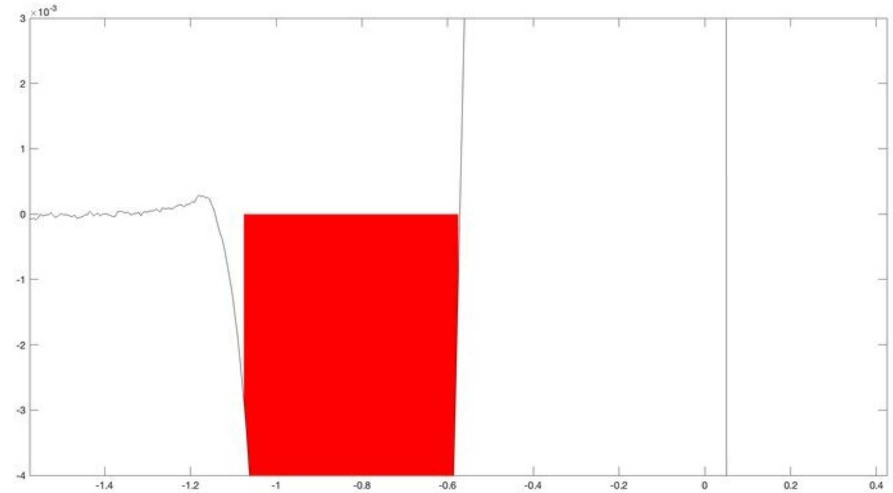

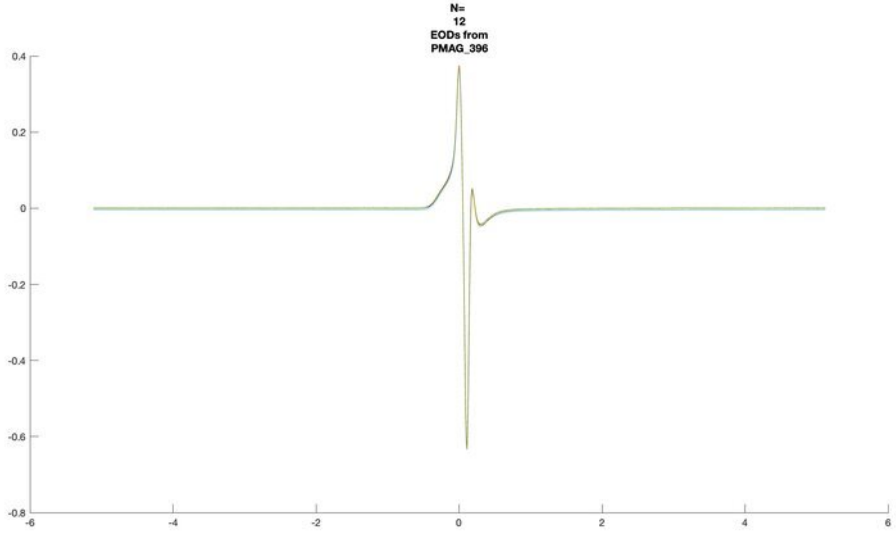
N=
12
EODs from
PMAG_396

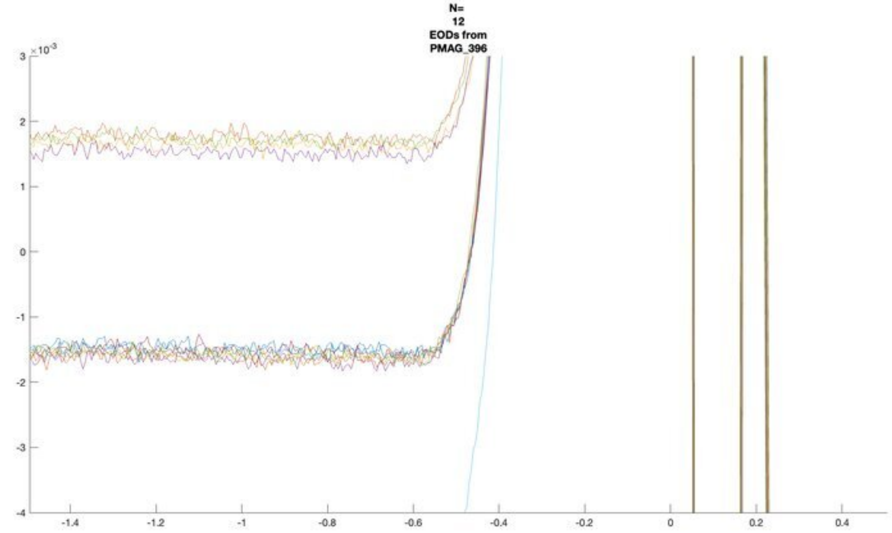
N=
12
EODs from
PMAG_396

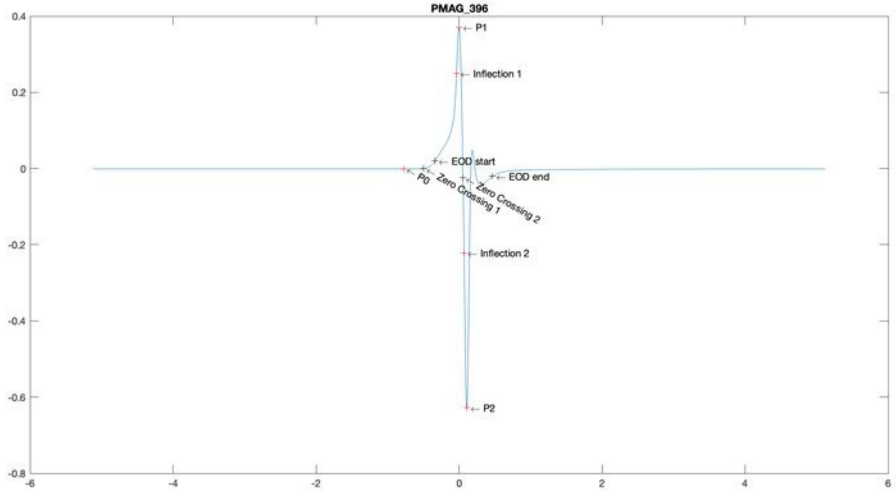
PMAG_396
P1
Inflection 1
EOD start
EOD end
P0
Zero Crossing 1
Zero Crossing 2
Inflection 2
P2

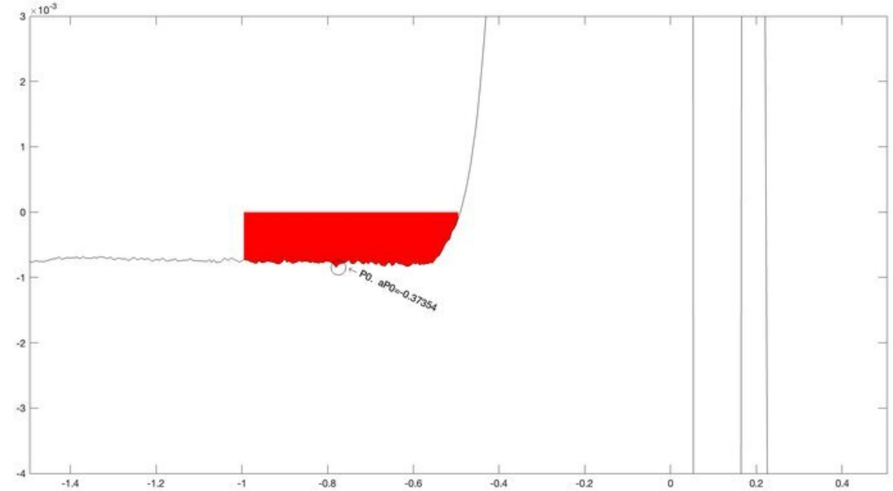
P0. aP0=-0.37354

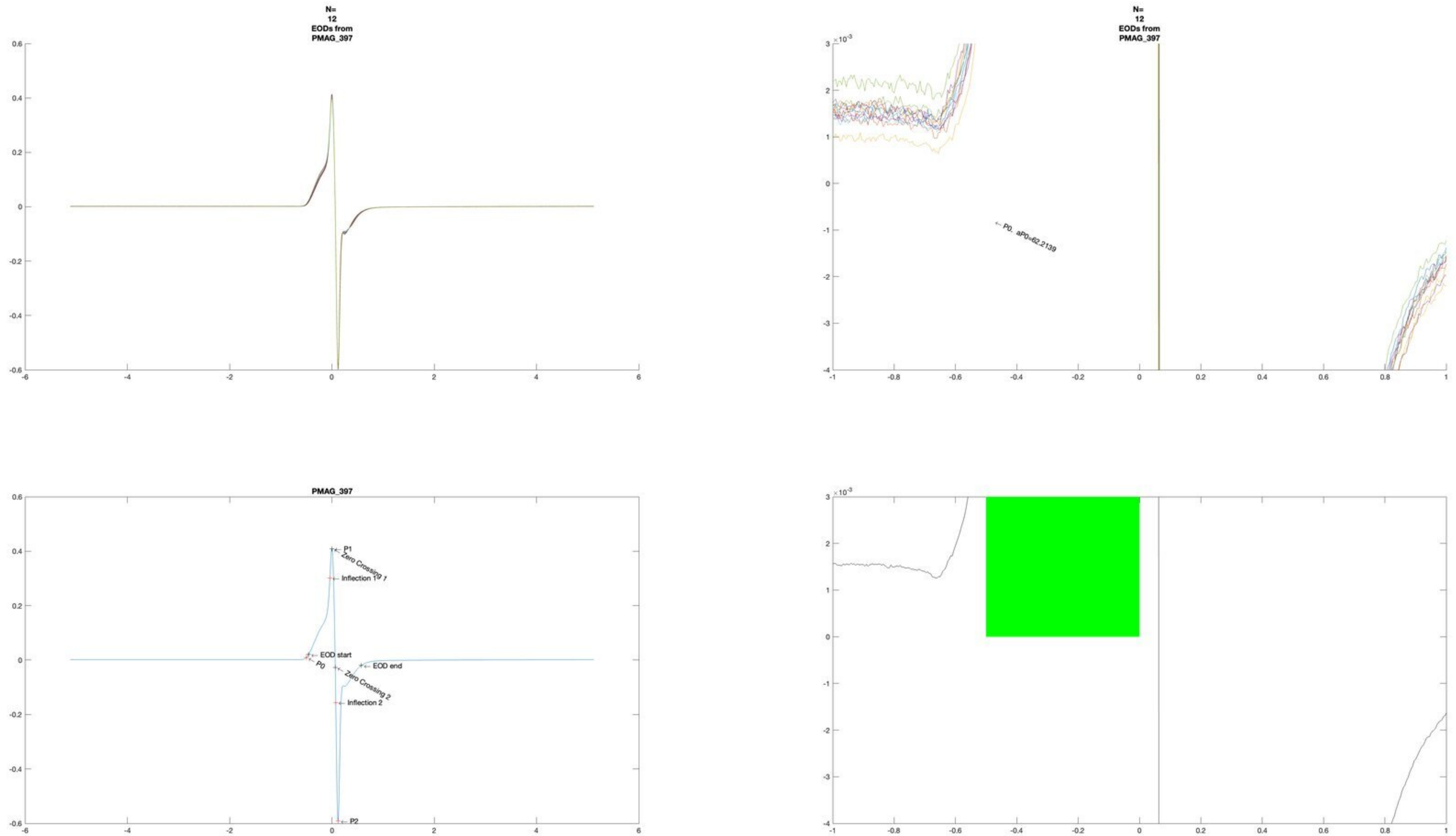

N=
12
EODs from
PMAG_397
N=
12
EODs from
PMAG_397
P0, aP0=62.2139
PMAG_397
P1
Zero Crossing 1
Inflection 1
EOD start
P0
EOD end
Zero Crossing 2
Inflection 2
P2

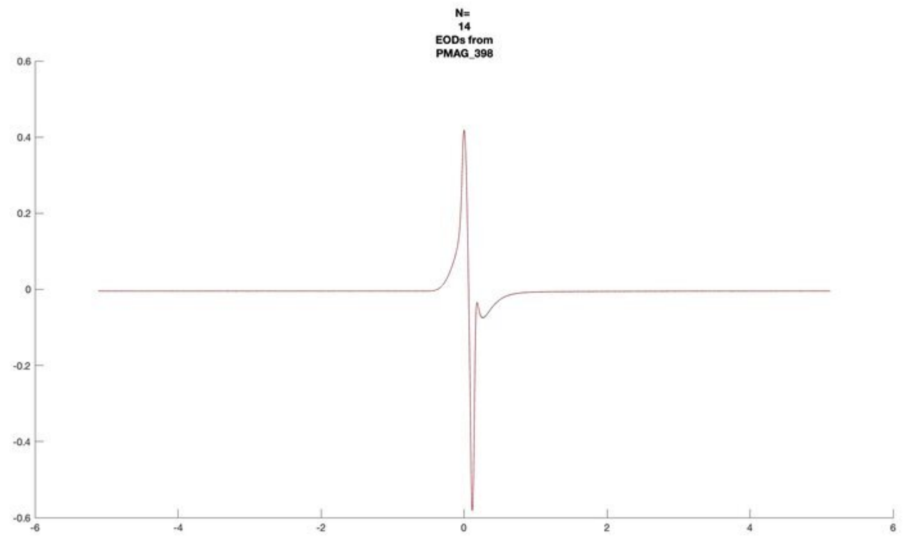
N=
14
EODs from
PMAG_398

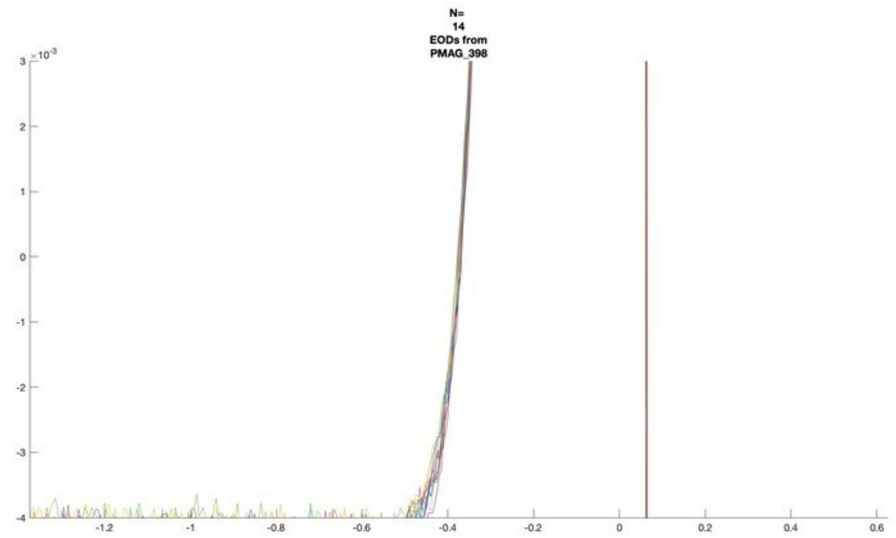
N=
14
EODs from
PMAG_398

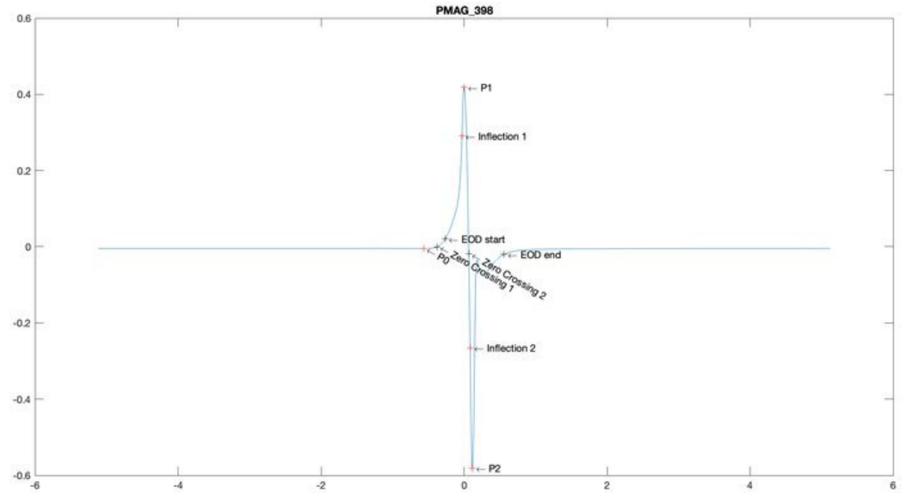
PMAG_398
P1
Inflection 1
EOD start
EOD end
P0
Zero Crossing 1
Zero Crossing 2
Inflection 2
P2

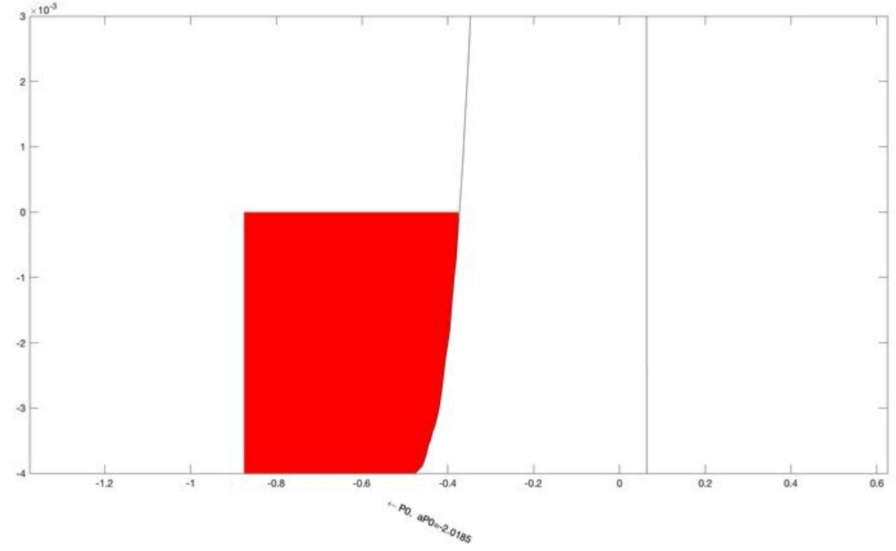
← P0. aP0=-2.0185

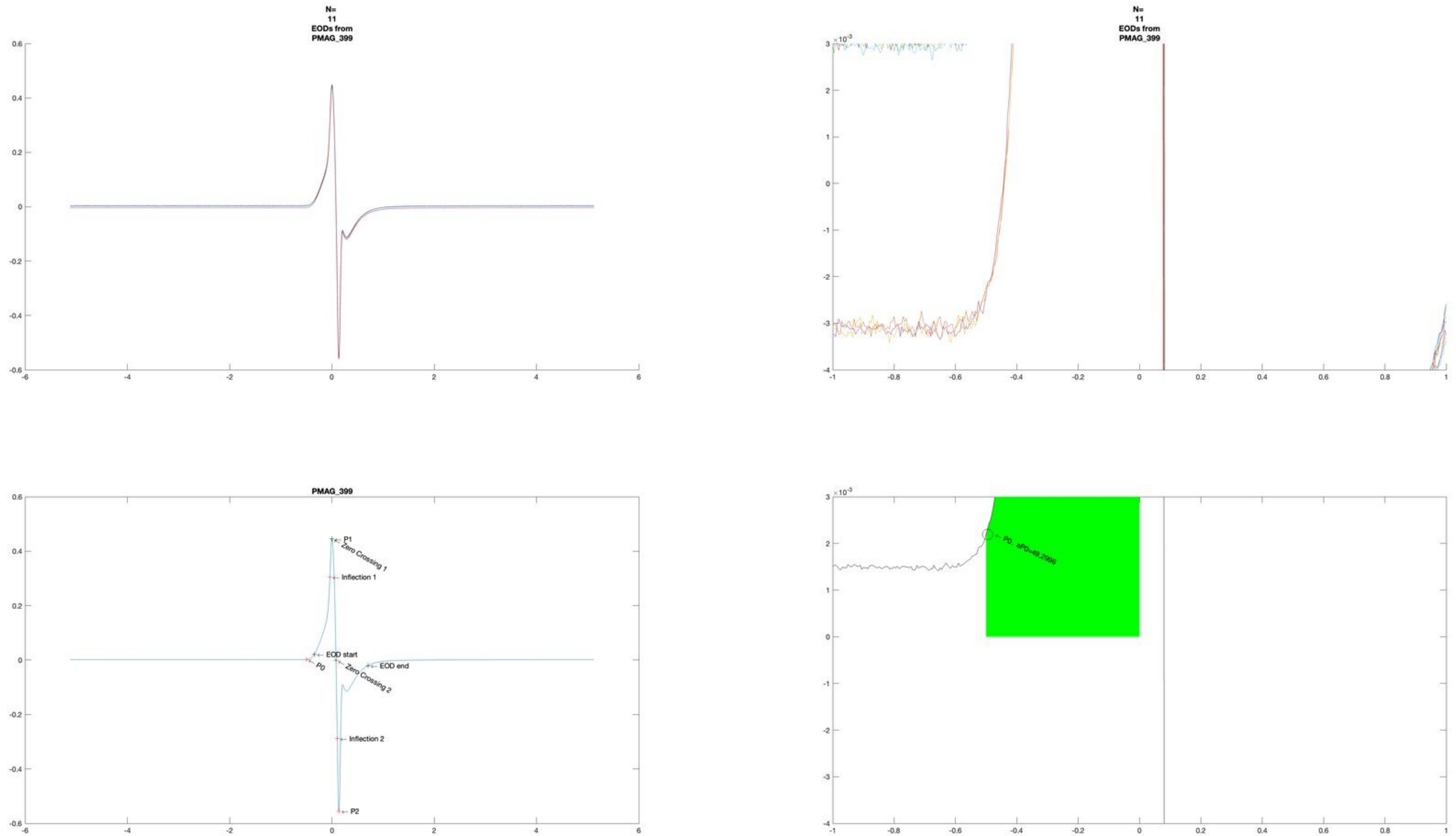
N=
11
EODs from
PMAG_399
N=
11
EODs from
PMAG_399
PMAG_399
P1
Zero Crossing 1
Inflection 1
EOD start
P0
Zero Crossing 2
EOD end
Inflection 2
P2
P0, aP0=49.2996

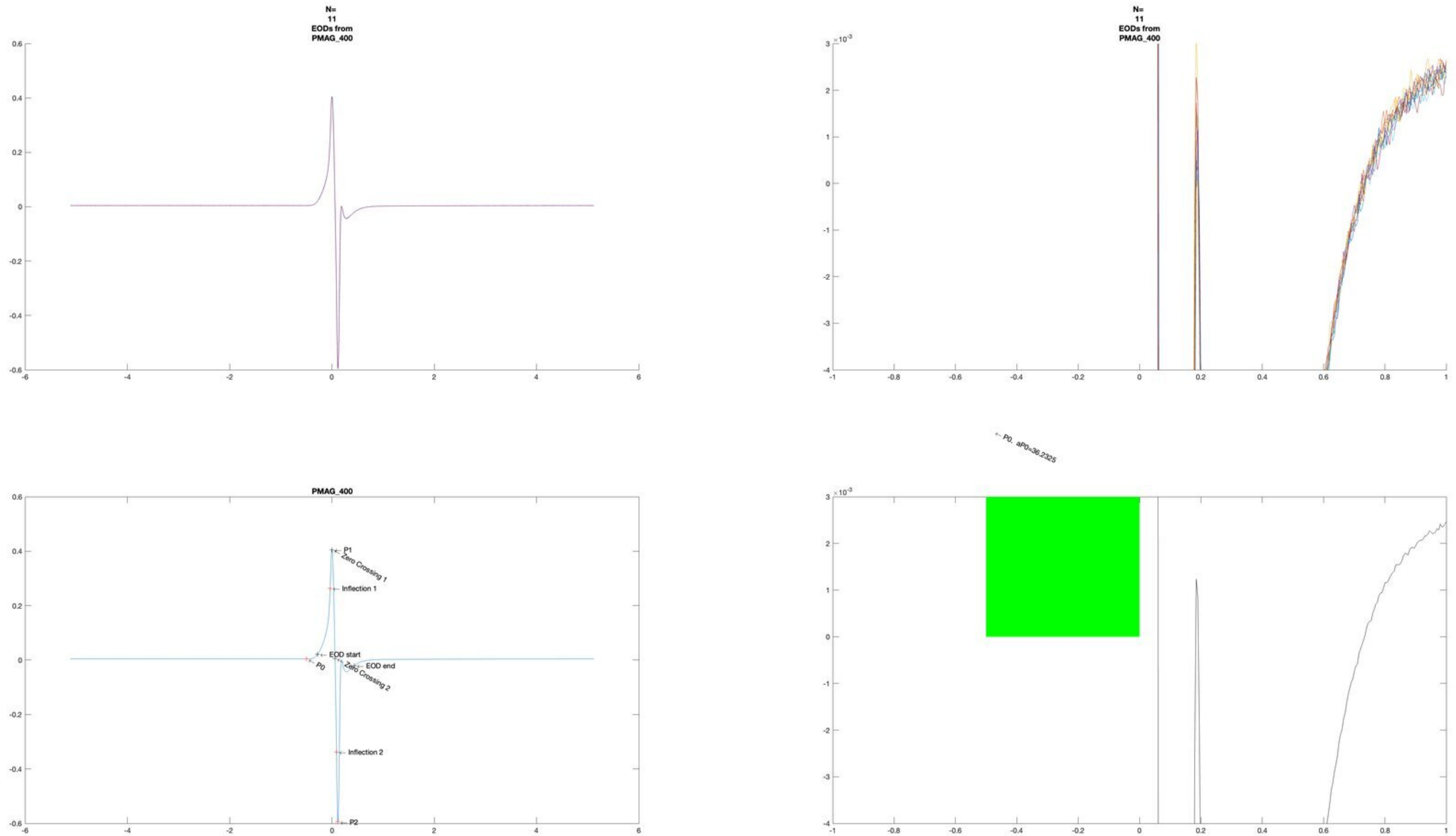
N=
11
EODs from
PMAG_400
N=
11
EODs from
PMAG_400
PMAG_400
P1
Zero Crossing 1
Inflection 1
EOD start
P0
EOD end
Zero Crossing 2
Inflection 2
P2
P0, aP0=36.2325

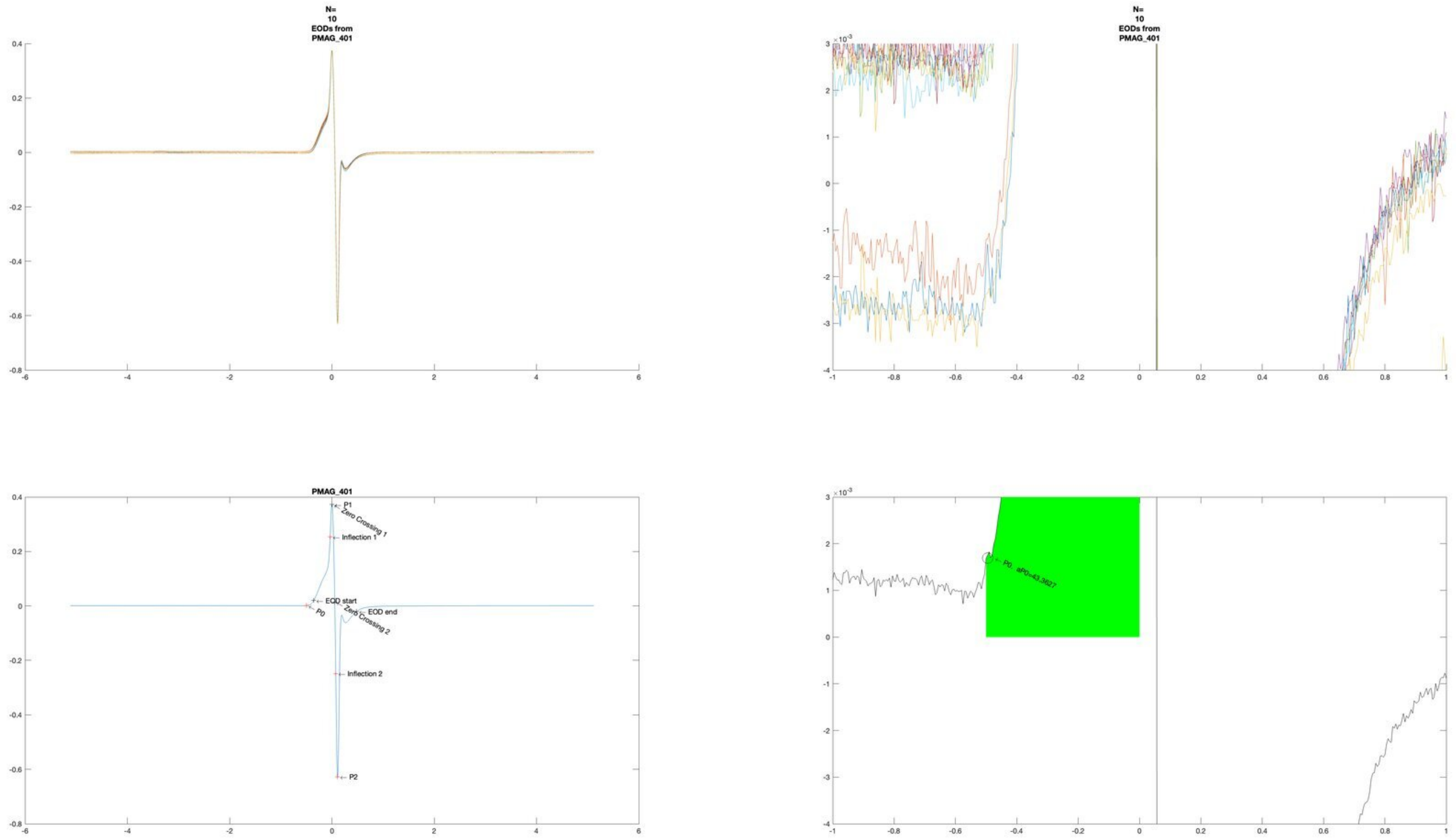

N=
10
EODs from
PMAG_401
N=
10
EODs from
PMAG_401
PMAG_401
P1
Zero Crossing 1
Inflection 1
EOD start
P0
EOD end
Zero Crossing 2
Inflection 2
P2
P0 . aP0=43.3627

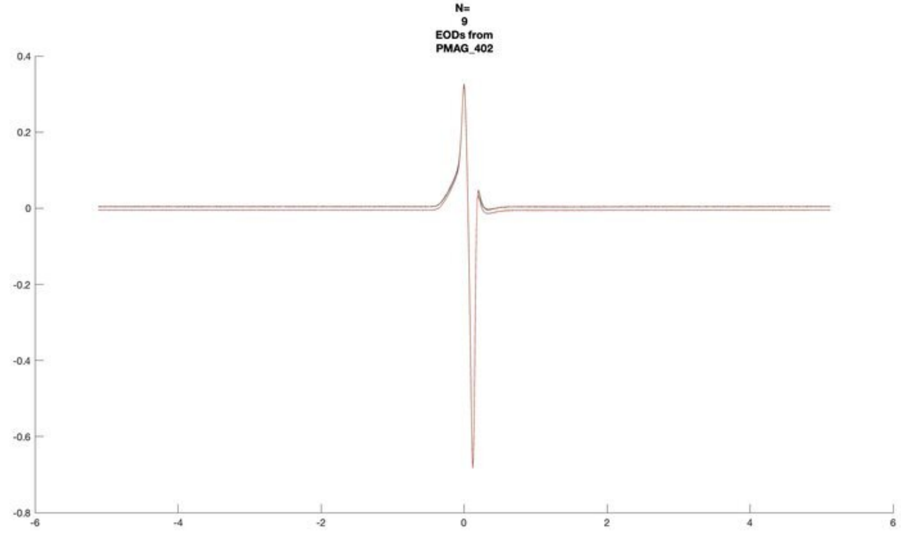
N=
9
EODs from
PMAG_402

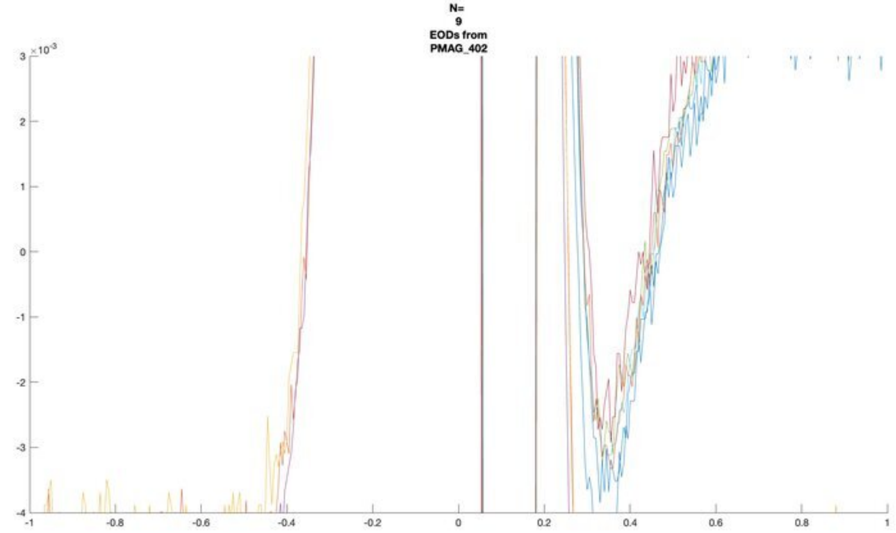
N=
9
EODs from
PMAG_402

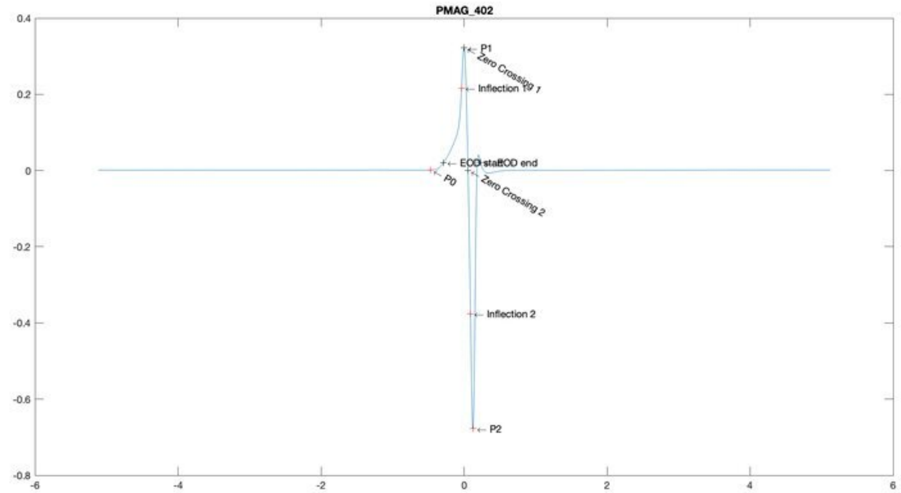
PMAG_402
P1
Zero Crossing 1
Inflection 1
EOD start
EOD end
P0
Zero Crossing 2
Inflection 2
P2

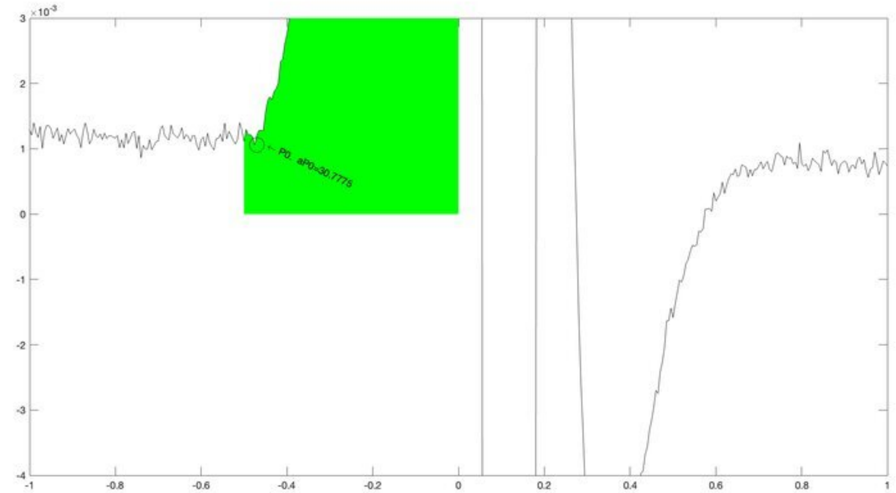
P0 aP0=30.7775

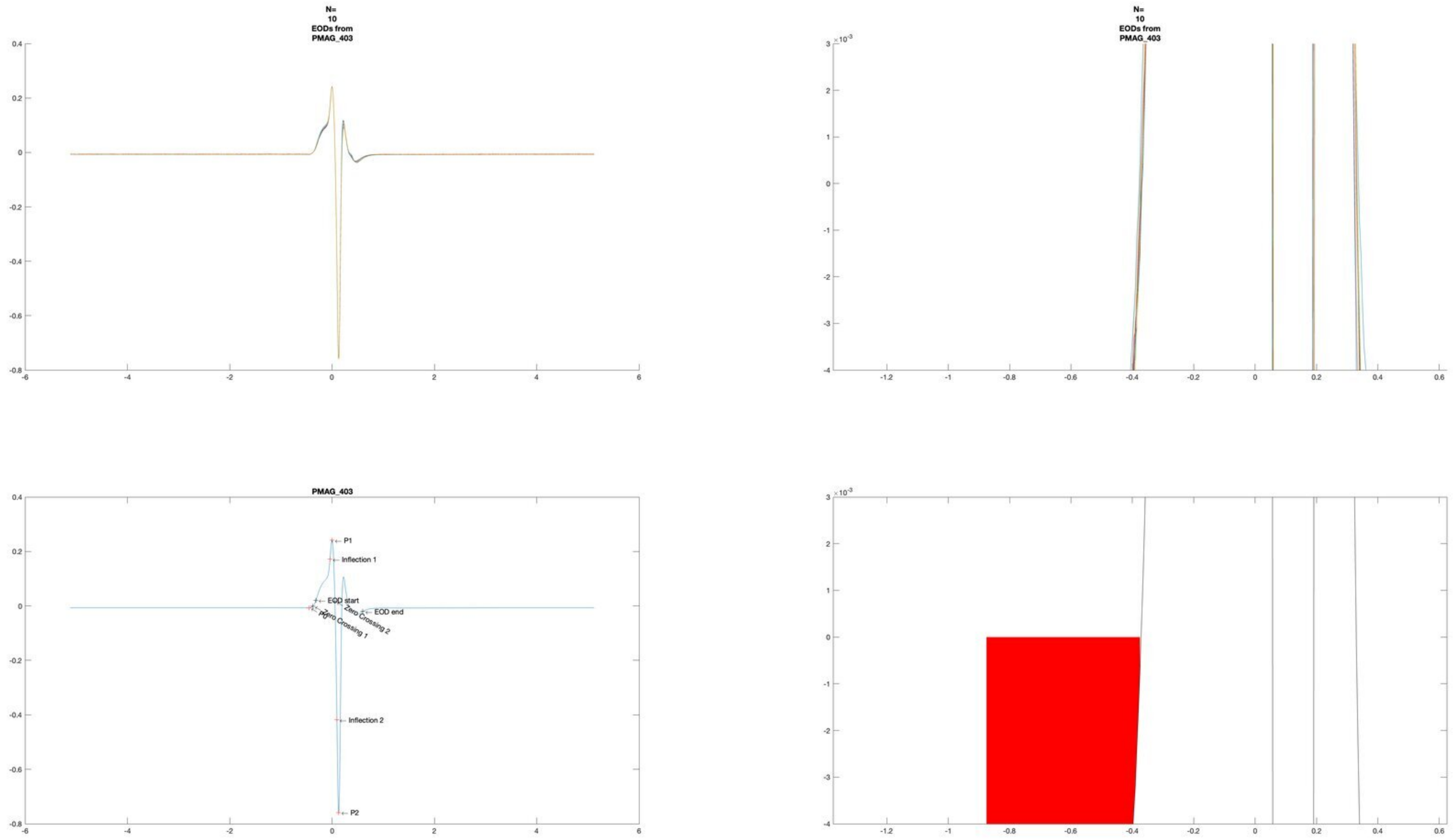
N=
10
EODs from
PMAG_403
N=
10
EODs from
PMAG_403
PMAG_403
P1
Inflection 1
EOD start
P0
Zero Crossing 1
Zero Crossing 2
EOD end
Inflection 2
P2

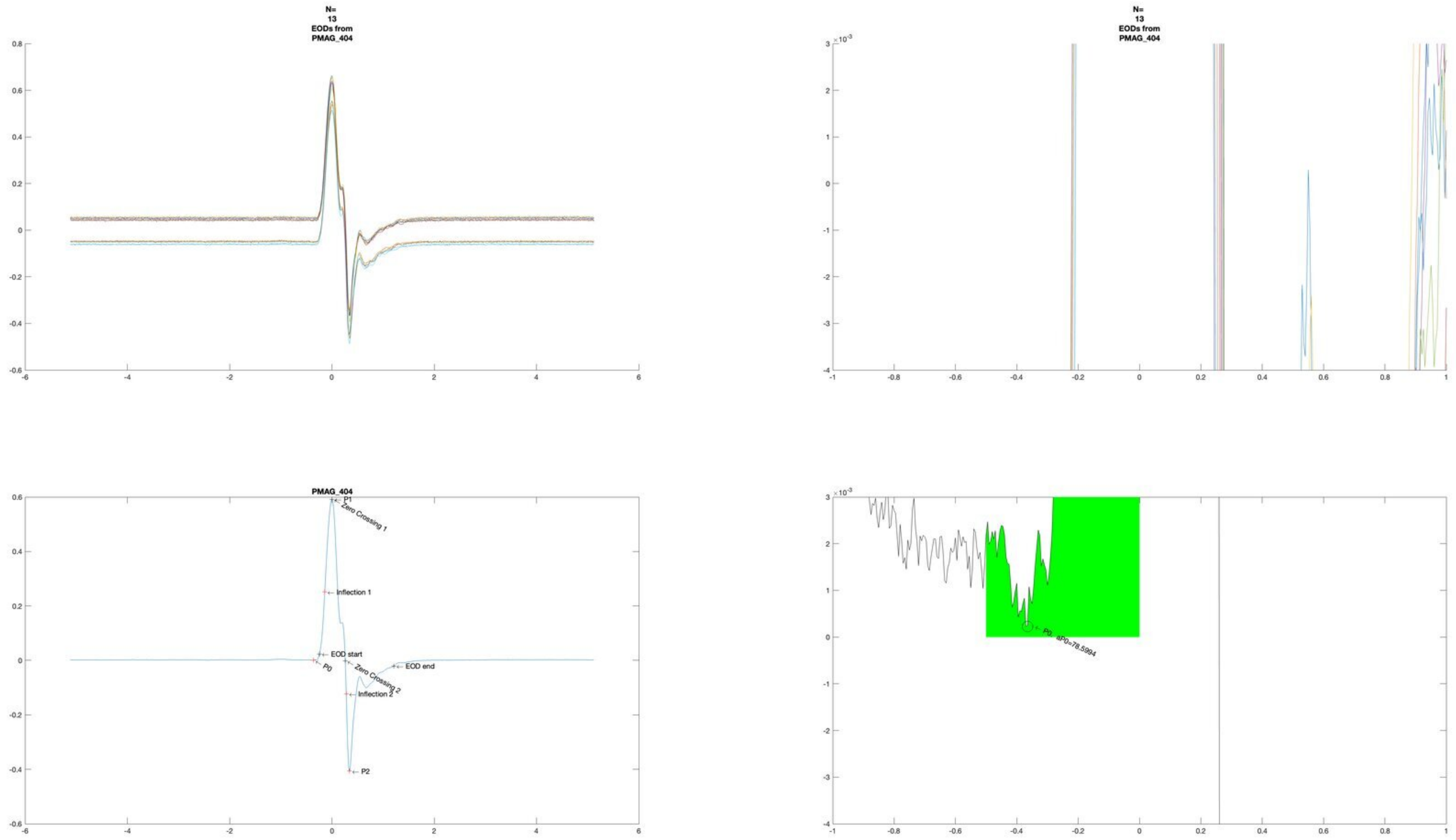
N=
13
EODs from
PMAG_404
N=
13
EODs from
PMAG_404
PMAG_404
P1
Zero Crossing 1
Inflection 1
EOD start
P0
Zero Crossing 2
EOD end
Inflection 2
P2
P0 aP0=78.5994

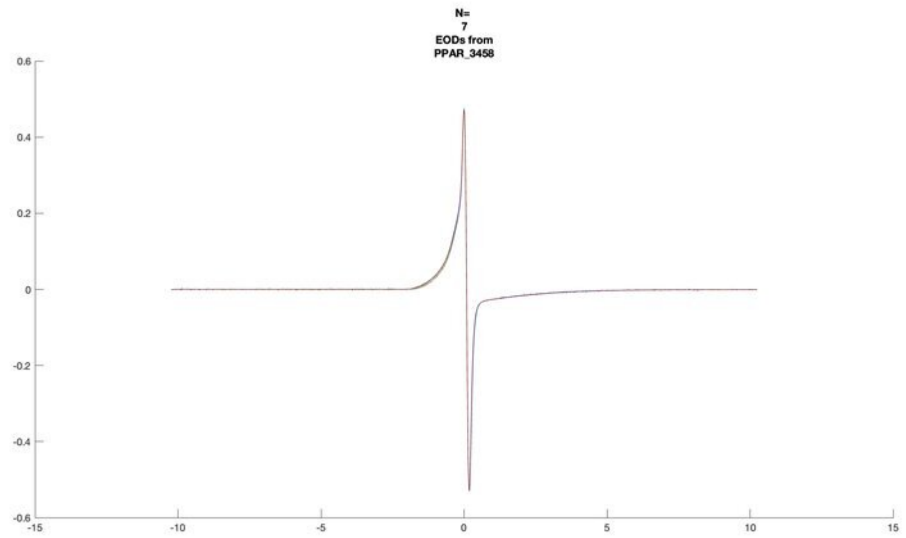
N=
7
EODs from
PPAR_3458

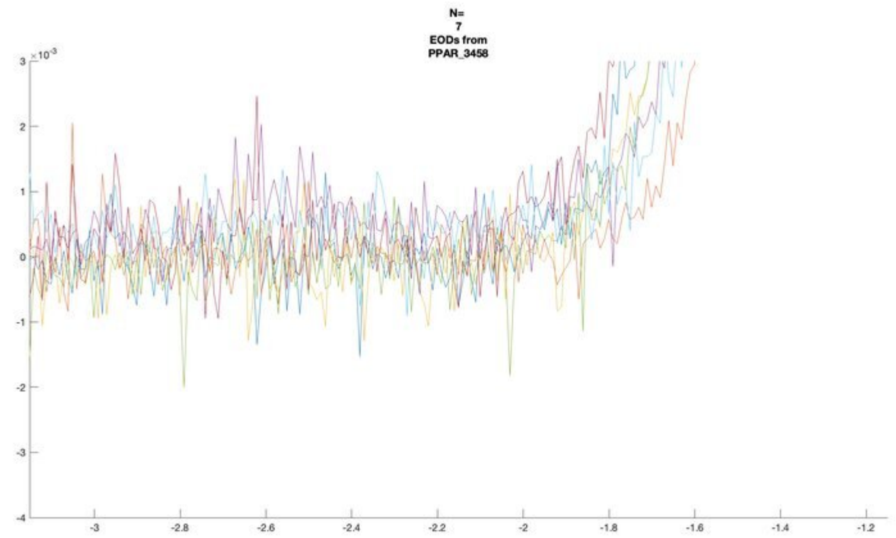
N=
7
EODs from
PPAR_3458
$\times 10^{-3}$

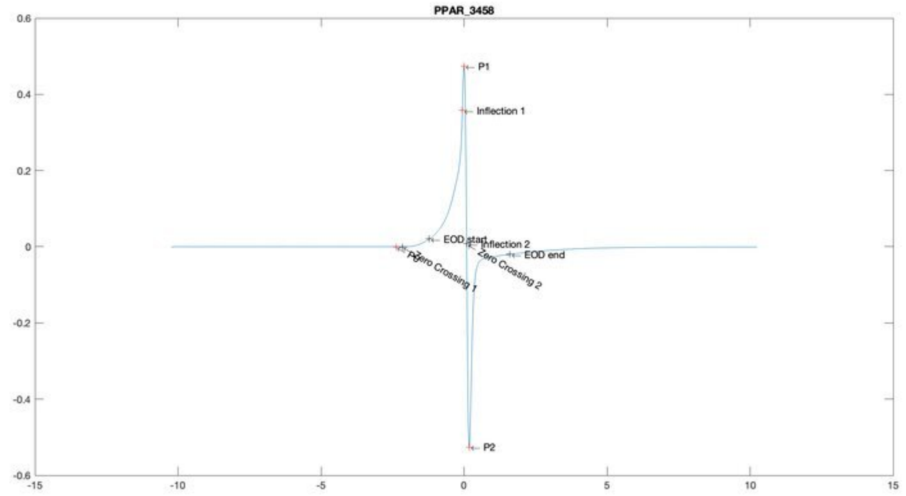
PPAR_3458
P1
Inflection 1
EOD start
Inflection 2
EOD end
Zero Crossing 1
Zero Crossing 2
P2

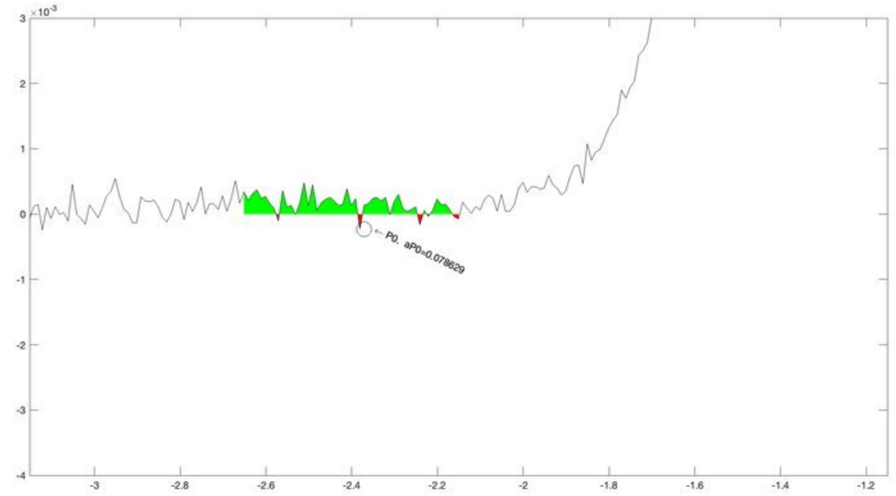
$\times 10^{-3}$
P0, aP0=0.078629

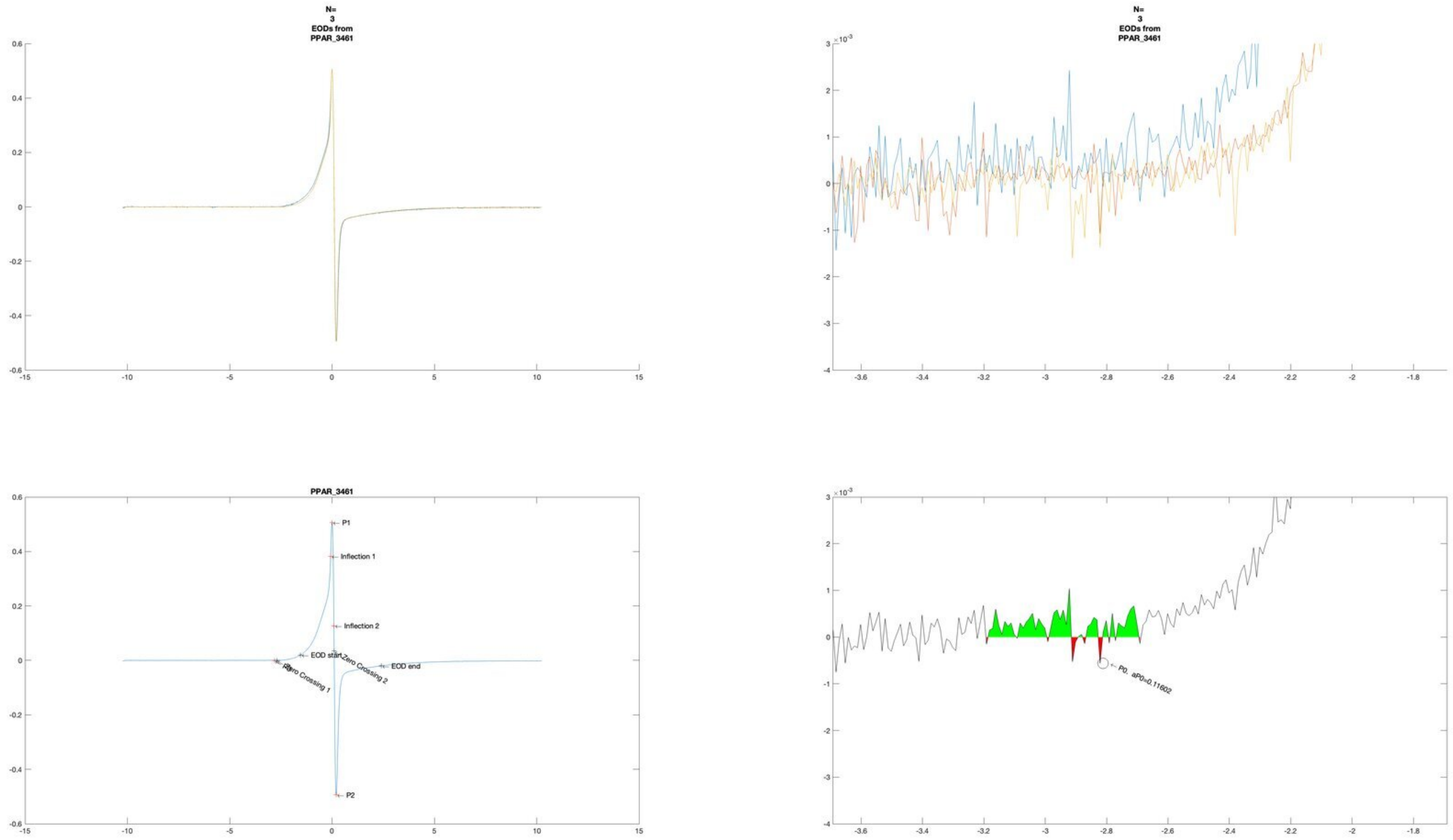
N=
3
EODs from
PPAR_3461
PPAR_3461
P1
Inflection 1
Inflection 2
EOD start
Zero Crossing 2
EOD end
Zero Crossing 1
P2
P0. aP0=0.11602

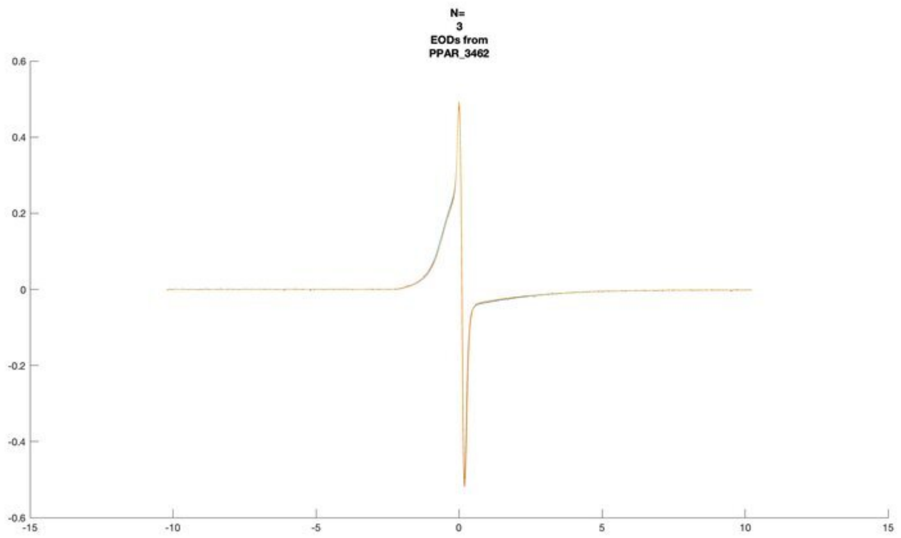
N=
3
EODs from
PPAR_3462

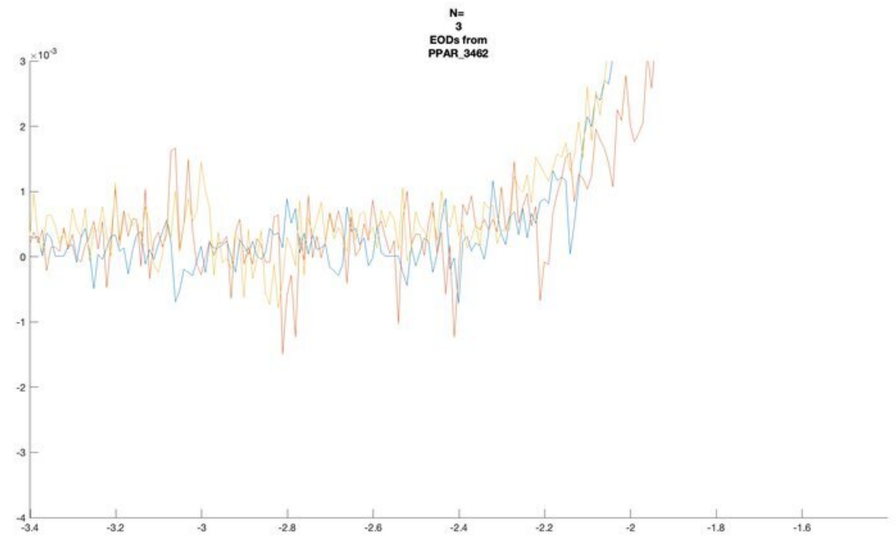
N=
3
EODs from
PPAR_3462

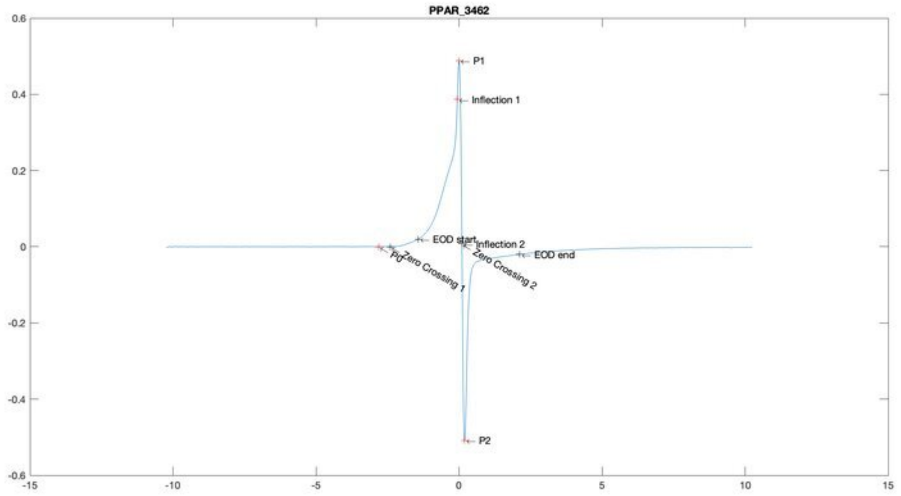
PPAR_3462
P1
Inflection 1
EOD start
Inflection 2
EOD end
P0
Zero Crossing 1
Zero Crossing 2
P2

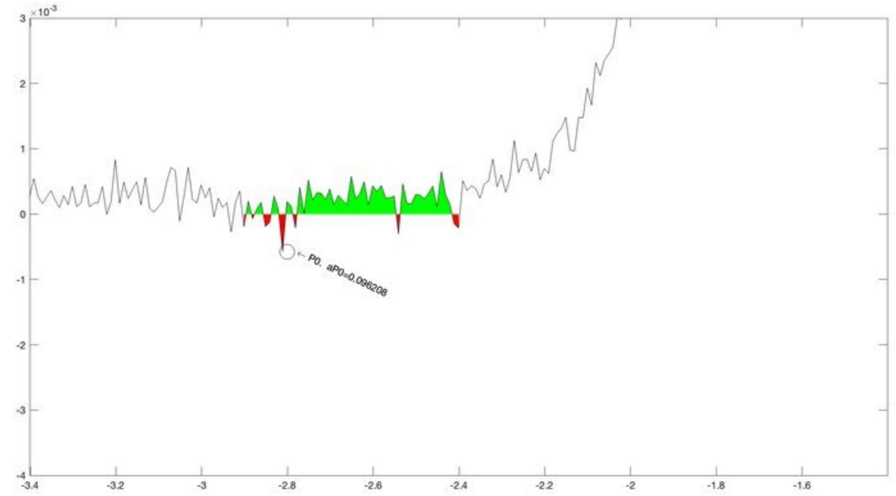
P0, aP0=0.096208

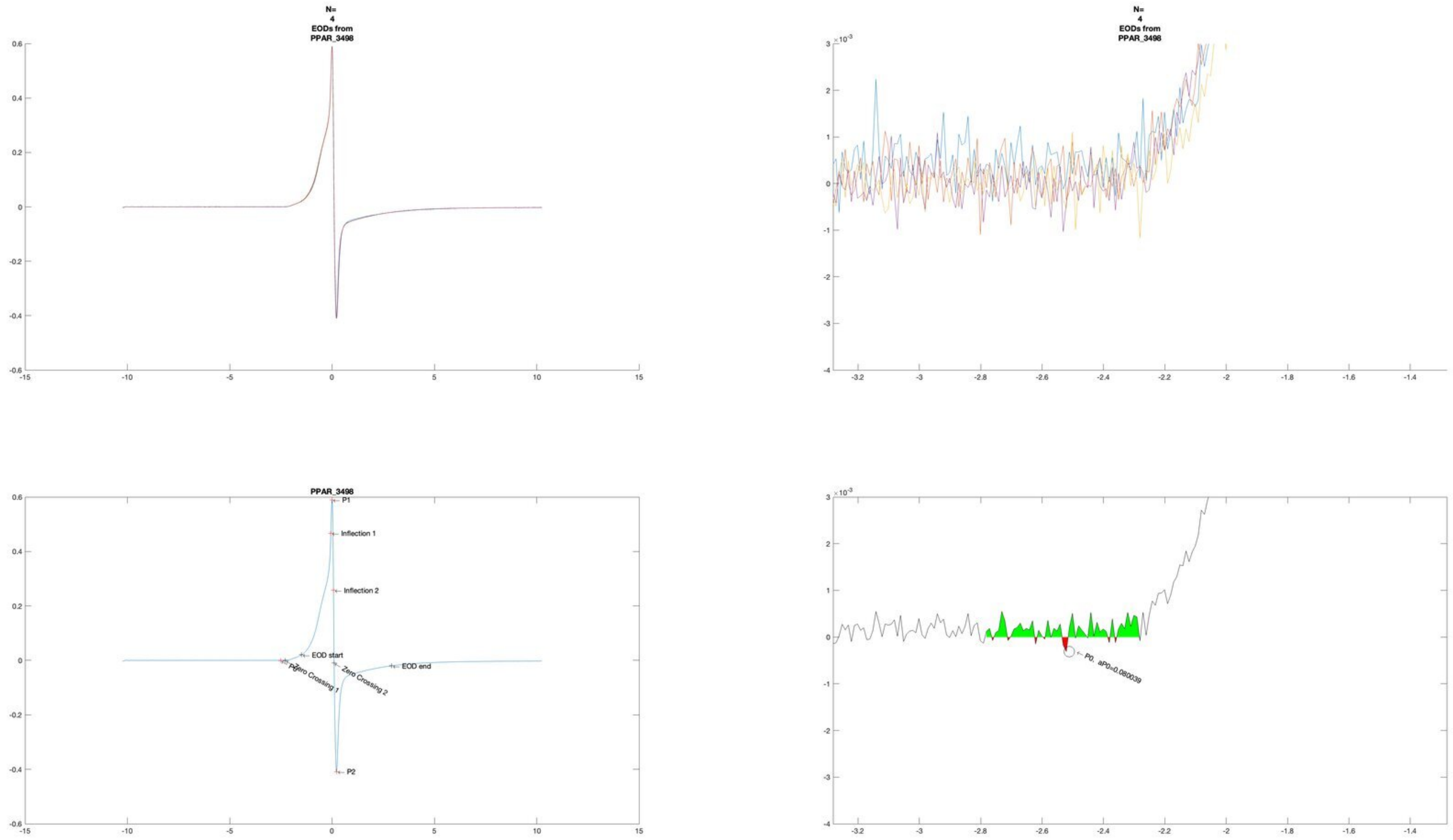
N=
4
EODs from
PPAR_3498
N=
4
EODs from
PPAR_3498
PPAR_3498
P1
Inflection 1
Inflection 2
EOD start
EOD end
Zero Crossing 1
Zero Crossing 2
P2
P0, aP0=0.080039

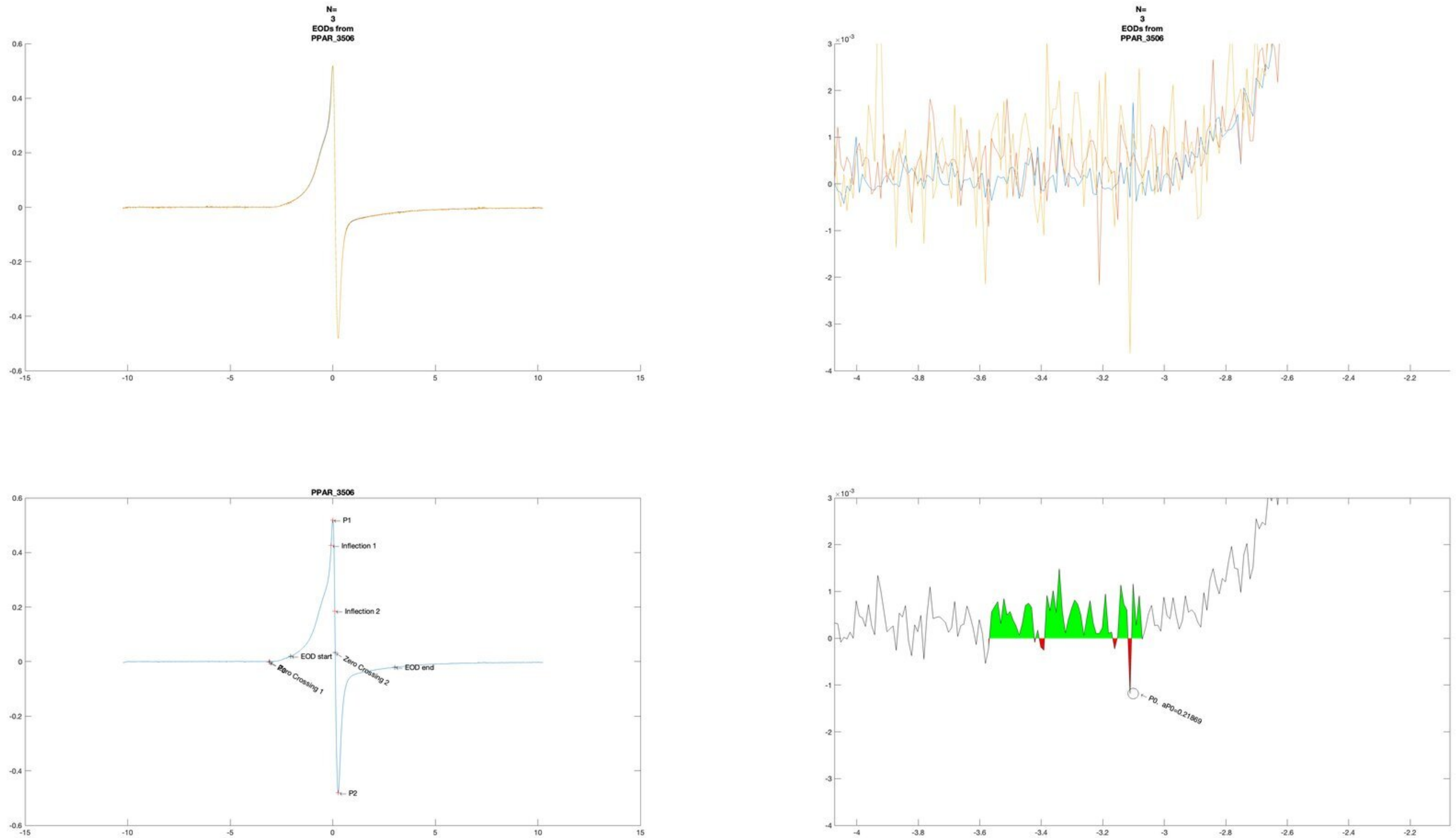
N=
3
EODs from
PPAR_3506
N=
3
EODs from
PPAR_3506
PPAR_3506
P1
Inflection 1
Inflection 2
EOD start
Zero Crossing 1
Zero Crossing 2
EOD end
P2
P0, aP0=0.21869

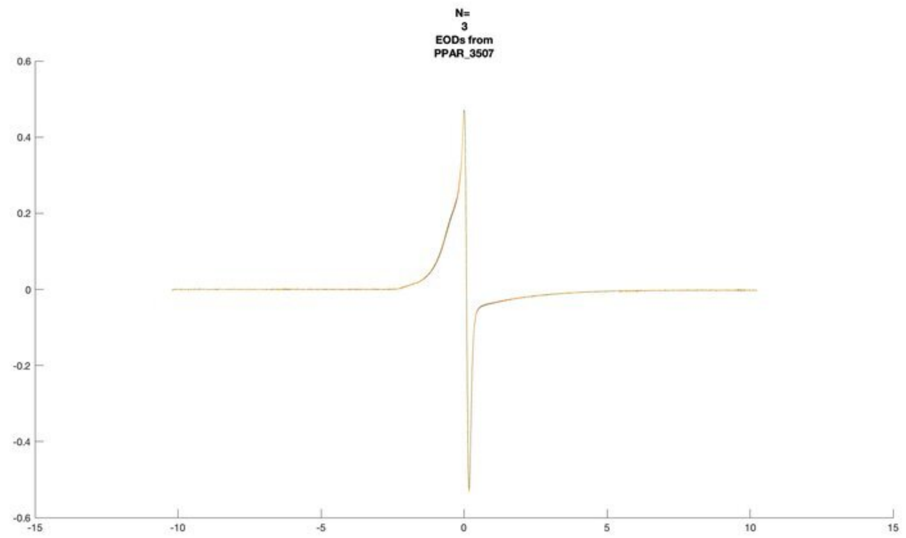
N=
3
EODs from
PPAR_3507

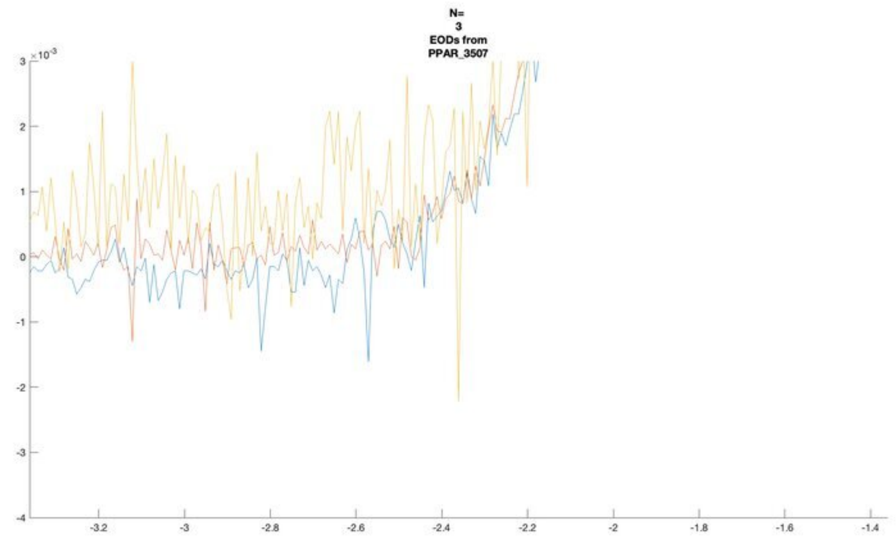
N=
3
EODs from
PPAR_3507

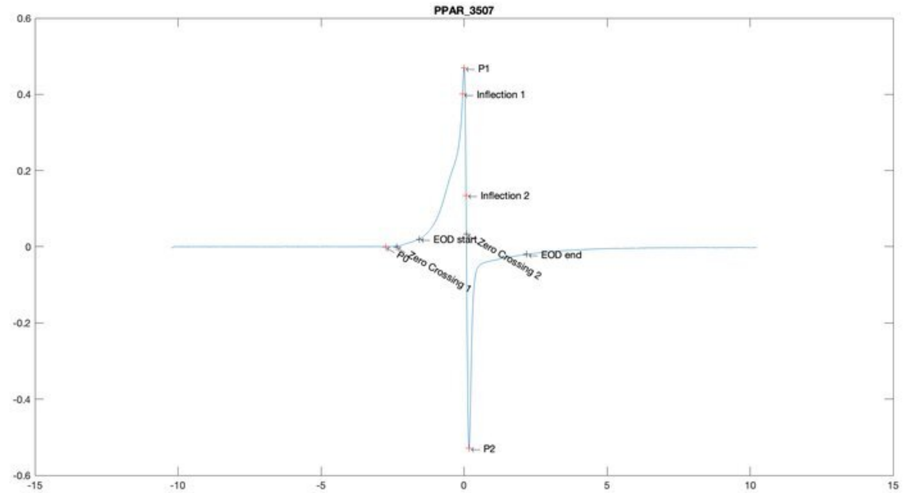
PPAR_3507
P1
Inflection 1
Inflection 2
EOD start
Zero Crossing 2
EOD end
P0
Zero Crossing 1
P2

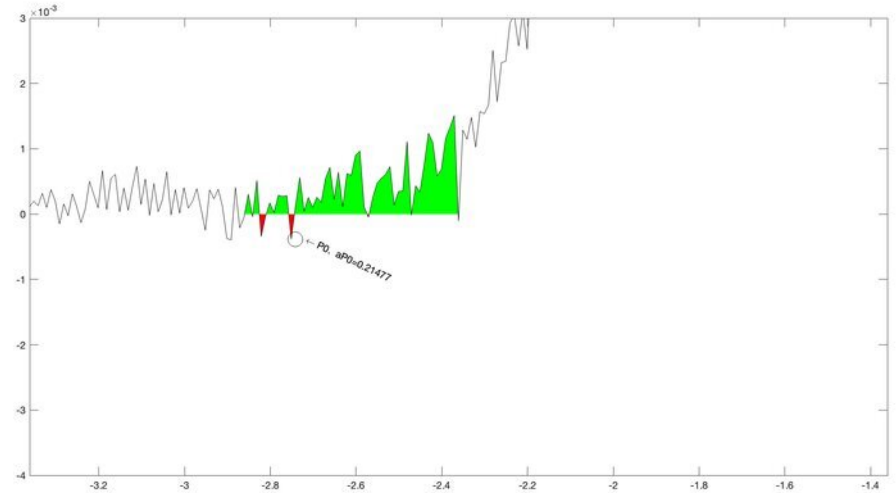
P0, aP0=0.21477

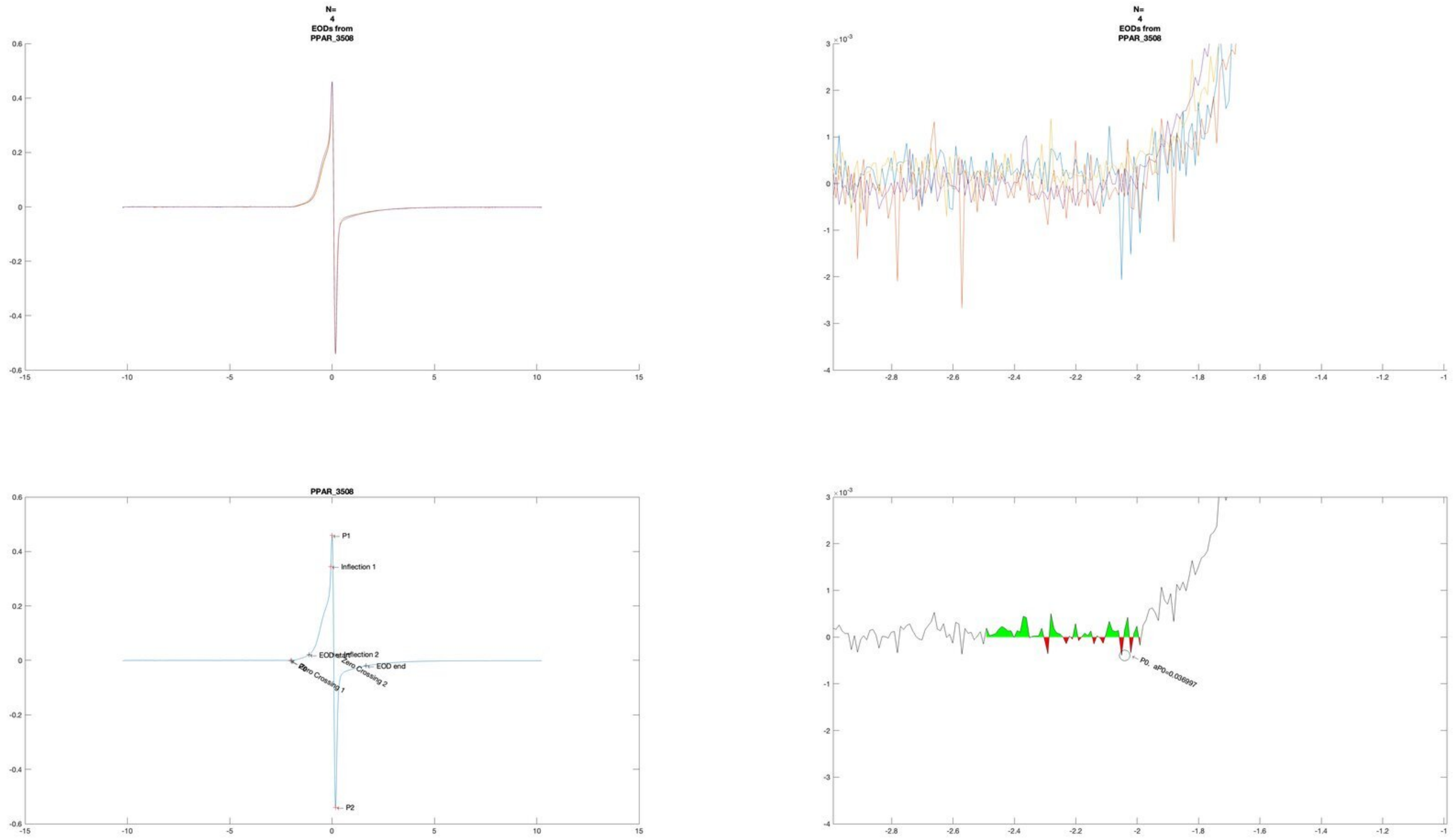
N=
4
EODs from
PPAR_3508
N=
4
EODs from
PPAR_3508
PPAR_3508
P1
Inflection 1
EOD start
Inflection 2
Zero Crossing 1
Zero Crossing 2
EOD end
P2
P0. aP0=0.036997

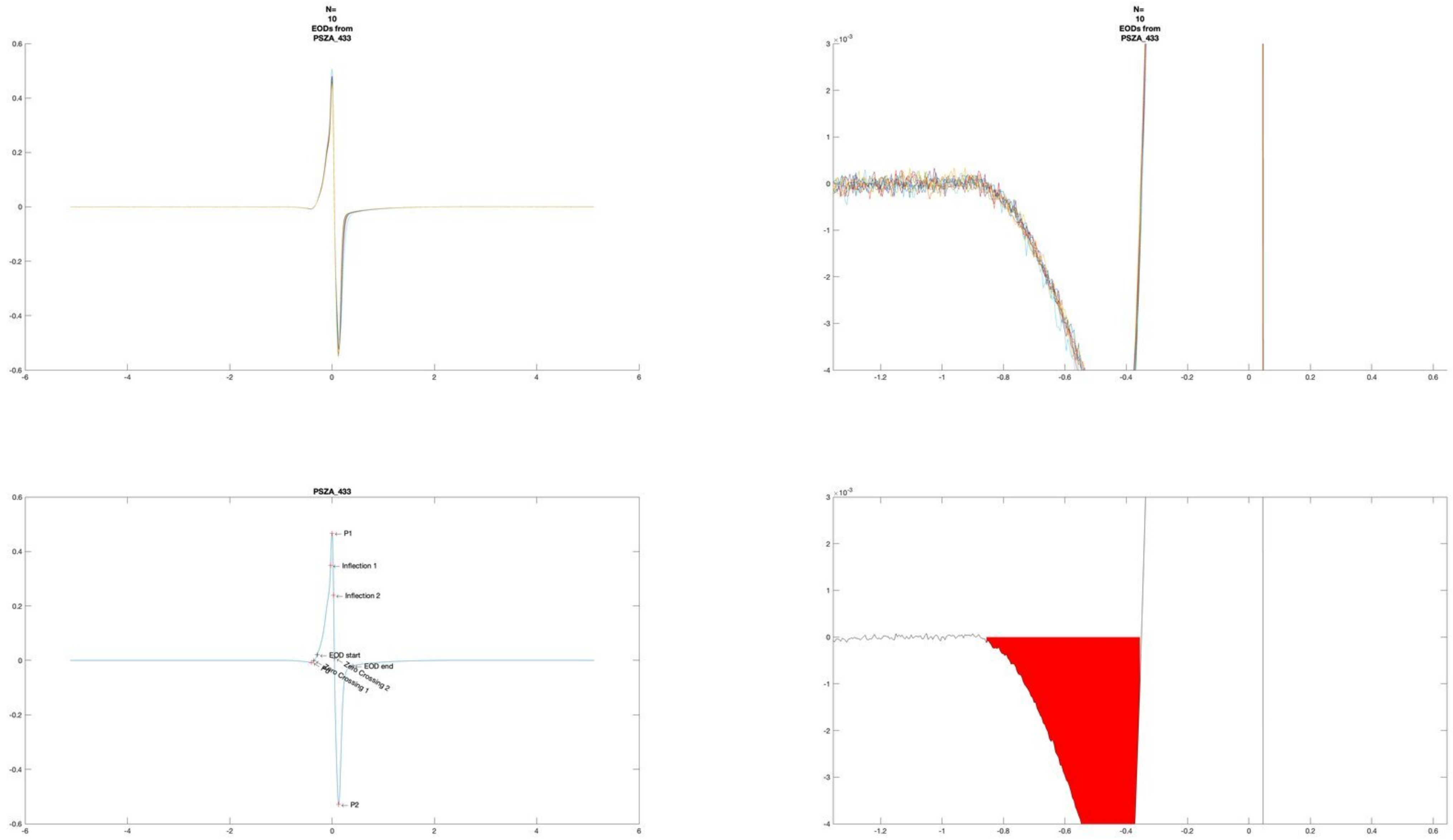
N=
10
EODs from
PSZA_433
N=
10
EODs from
PSZA_433
$\times 10^{-3}$
PSZA_433
P1
Inflection 1
Inflection 2
EOD start
EOD end
Zero Crossing 2
Zero Crossing 1
P2
$\times 10^{-3}$

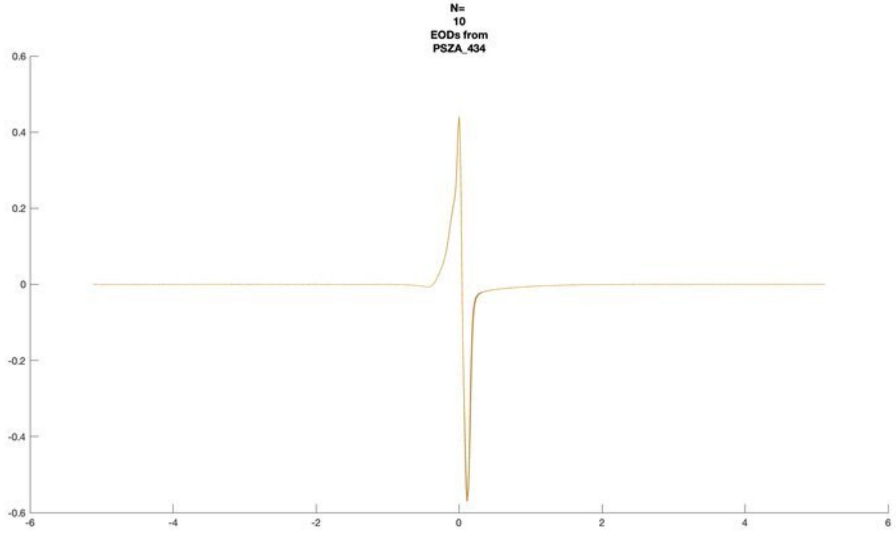
N=
10
EODs from
PSZA_434

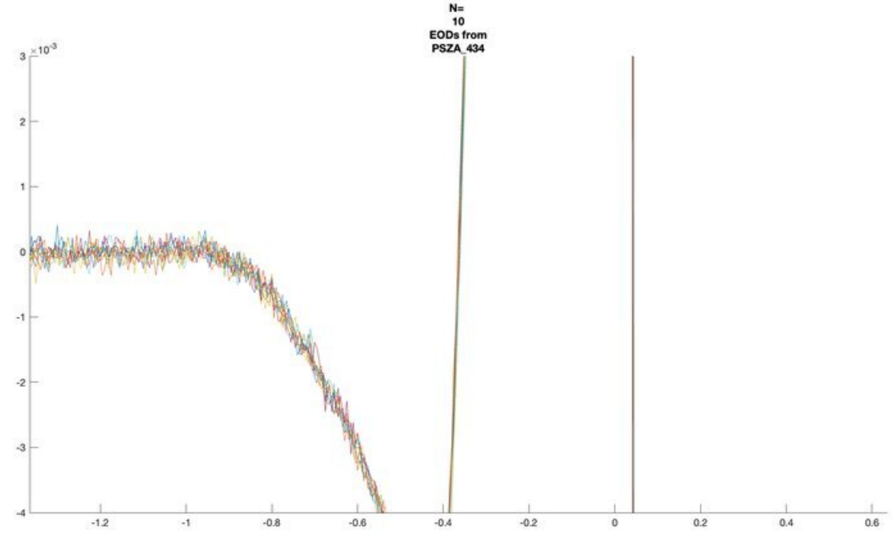
N=
10
EODs from
PSZA_434
×10^-3

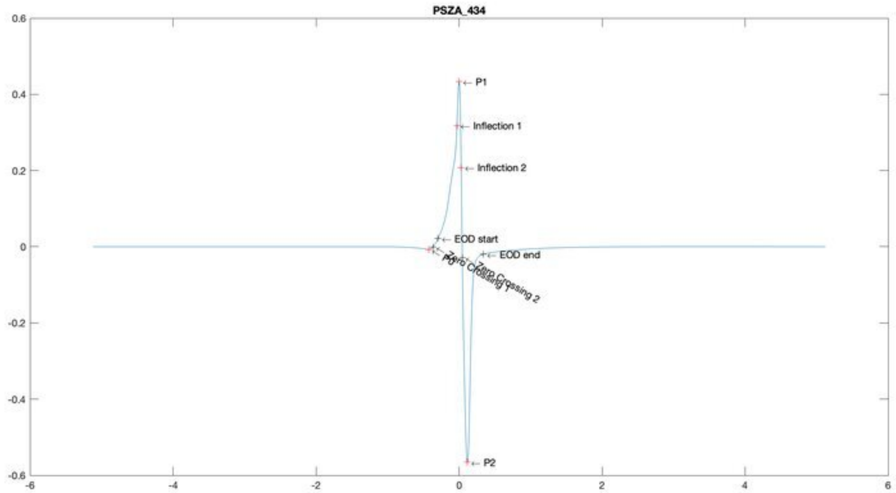
PSZA_434
P1
Inflection 1
Inflection 2
EOD start
EOD end
P0
Zero Crossing 1
Zero Crossing 2
P2

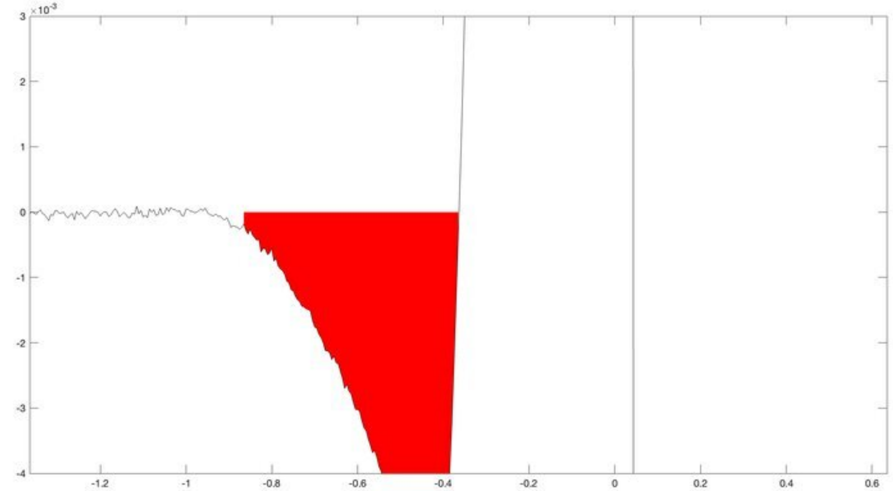
×10^-3

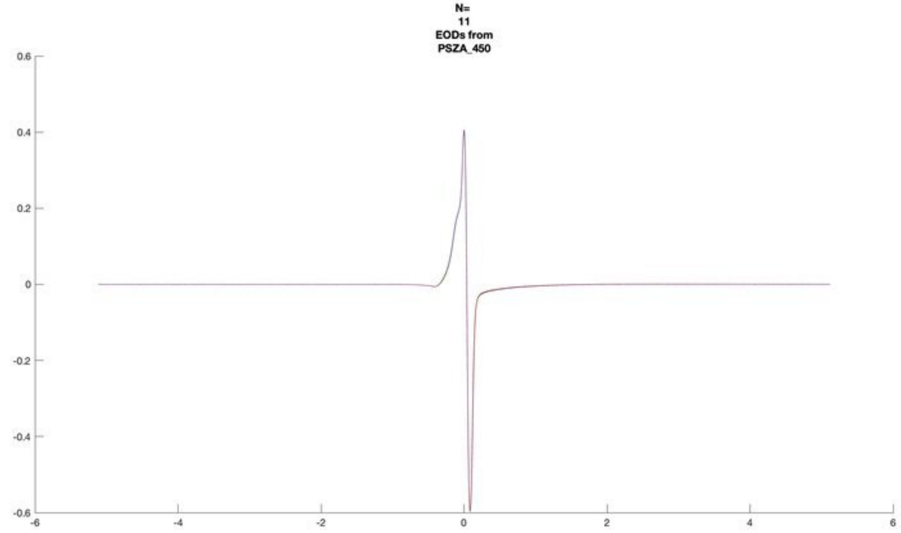
N=
11
EODs from
PSZA_450
0.6
0.4
0.2
0
-0.2
-0.4
-0.6
-6
-4
-2
0
2
4
6

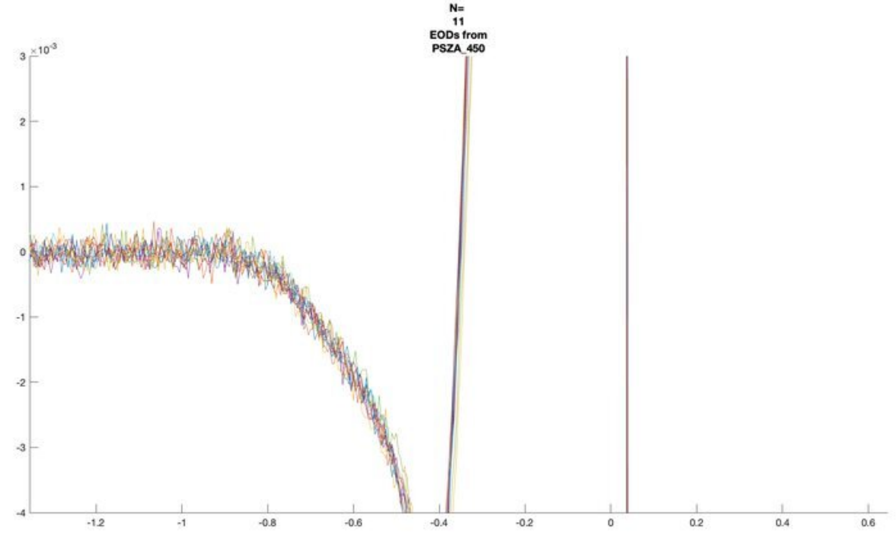
N=
11
EODs from
PSZA_450
×10^-3
3
2
1
0
-1
-2
-3
-4
-1.2
-1
-0.8
-0.6
-0.4
-0.2
0
0.2
0.4
0.6

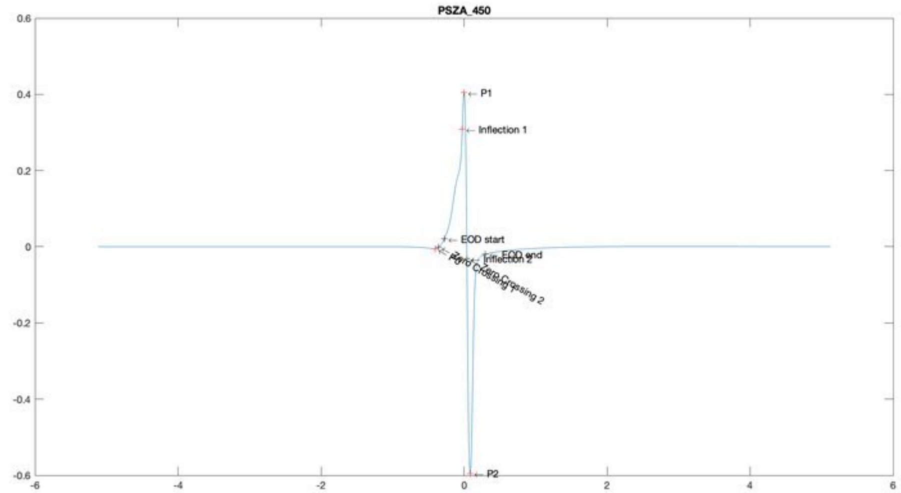
PSZA_450
P1
Inflection 1
EOD start
EOD end
Inflection 2
Zero Crossing 2
P2
0.6
0.4
0.2
0
-0.2
-0.4
-0.6
-6
-4
-2
0
2
4
6

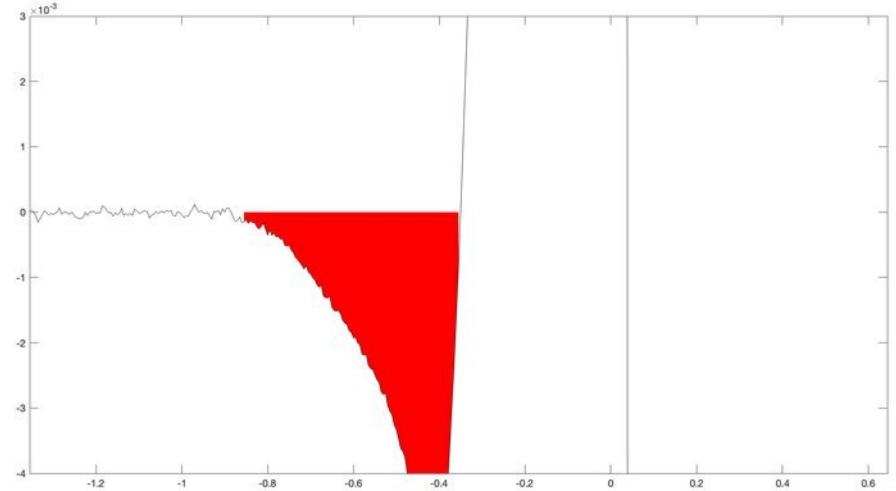
×10^-3
3
2
1
0
-1
-2
-3
-4
-1.2
-1
-0.8
-0.6
-0.4
-0.2
0
0.2
0.4
0.6

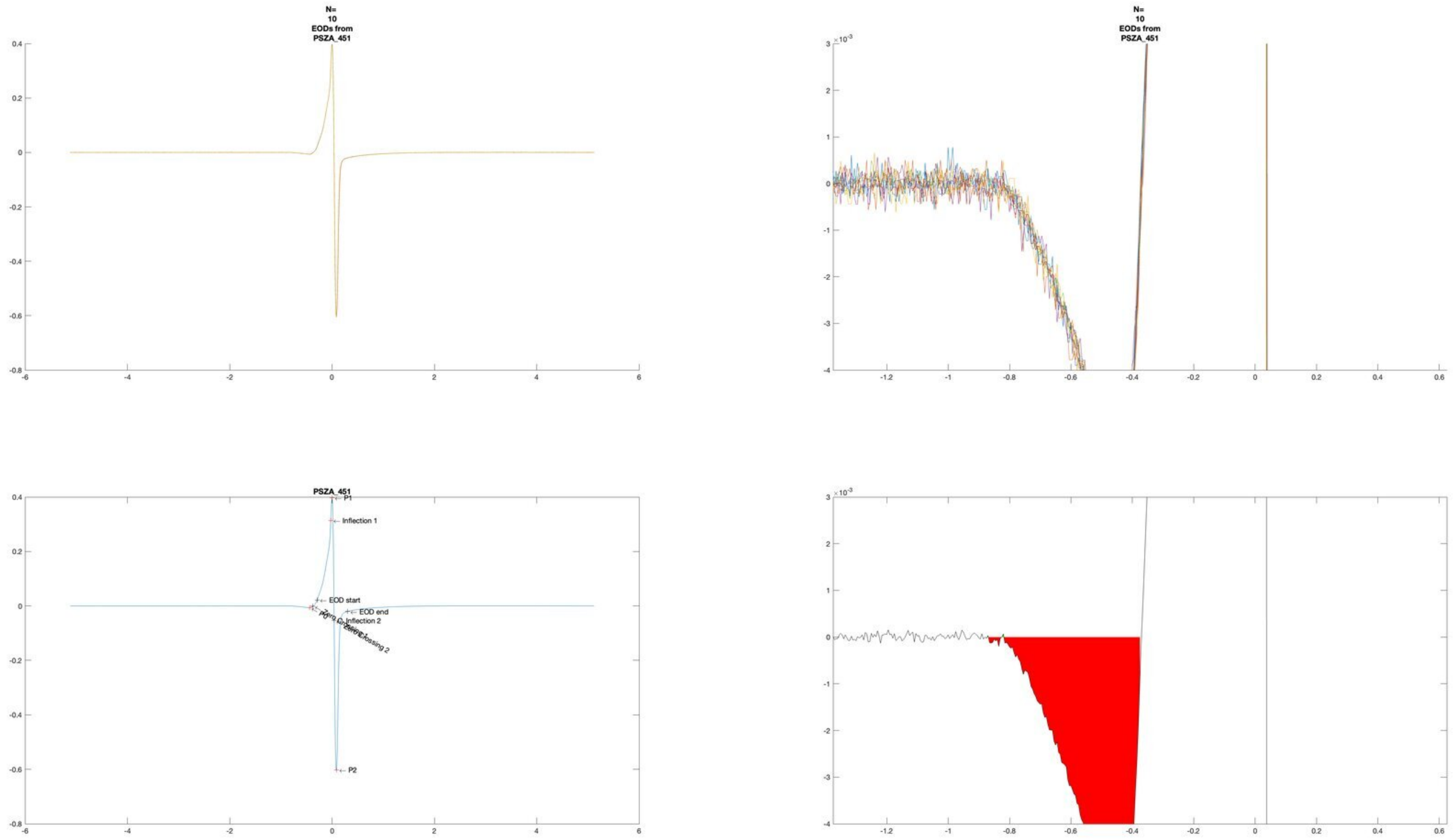
N=
10
EODs from
PSZA_451
N=
10
EODs from
PSZA_451
PSZA_451
P1
Inflection 1
EOD start
EOD end
Inflection 2
P0
P2

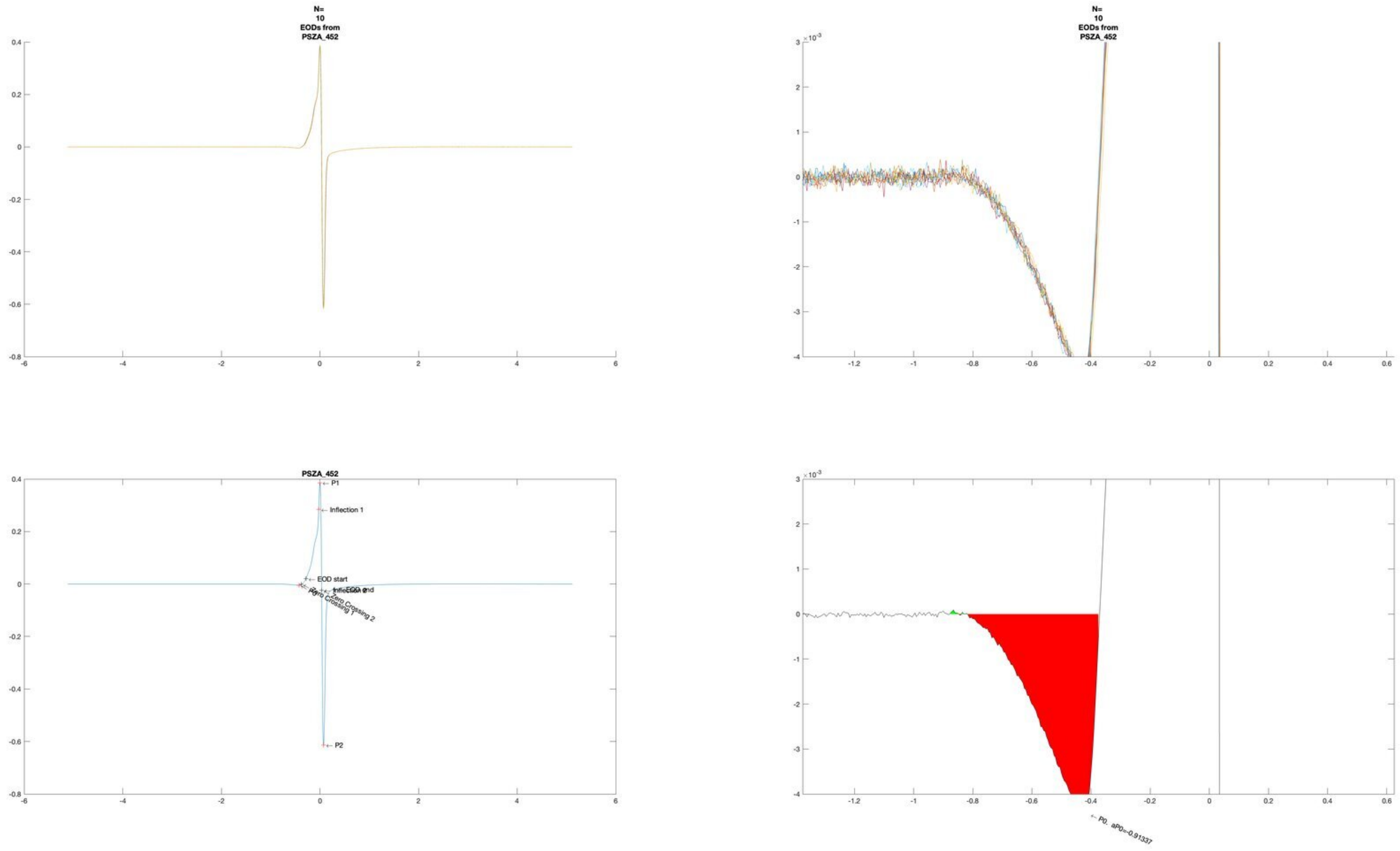

N=
10
EODs from
PSZA_452
PSZA_452
P1
Inflection 1
EOD start
EOD end
Inflection 2
Zero Crossing 1
Zero Crossing 2
P0
P2
P0, aP0=-0.91337

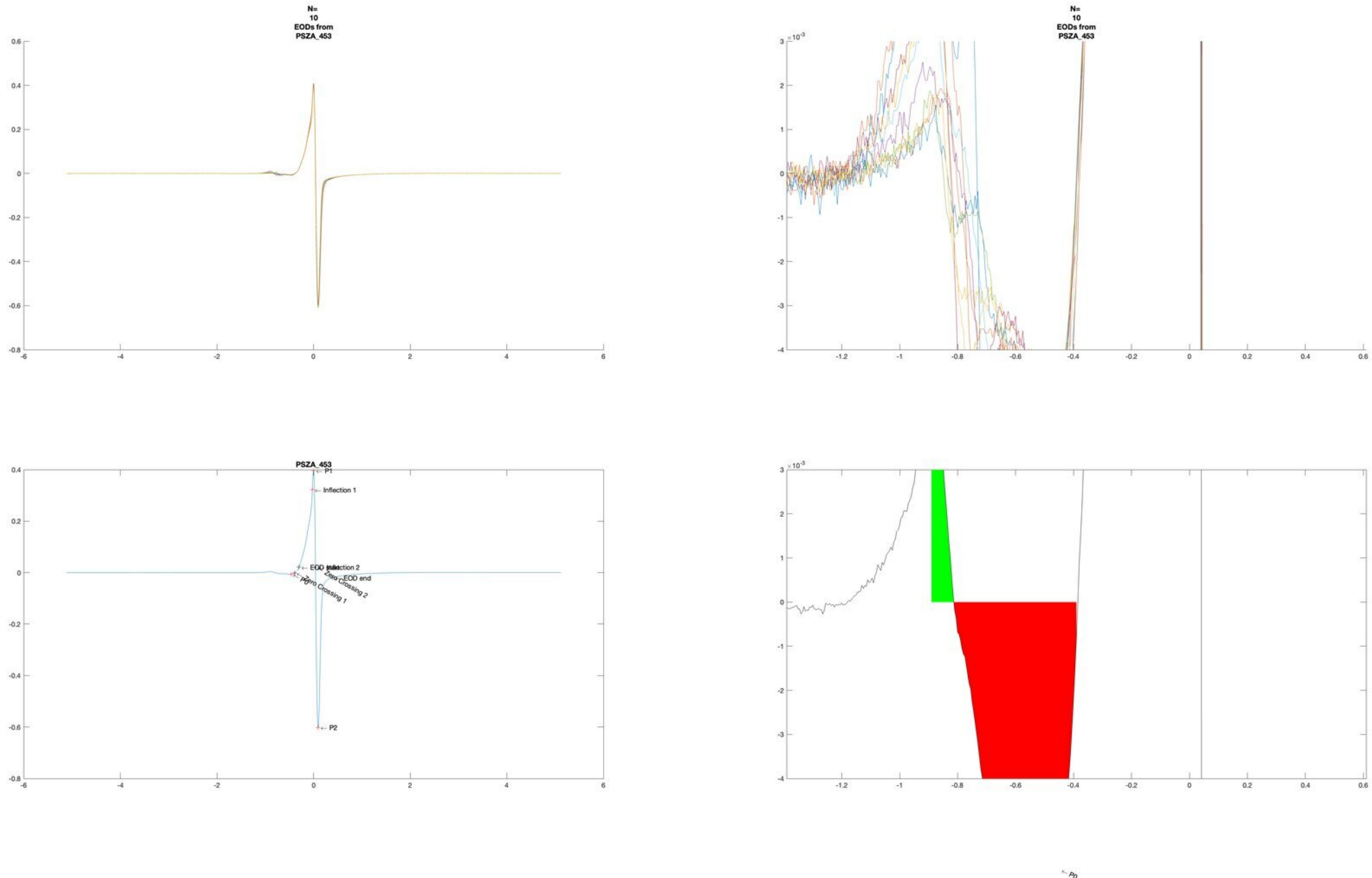
N=
10
EODs from
PSZA_453
N=
10
EODs from
PSZA_453
PSZA_453
P1
Inflection 1
Inflection 2
EOD end
Zero Crossing 1
Zero Crossing 2
P0
P2

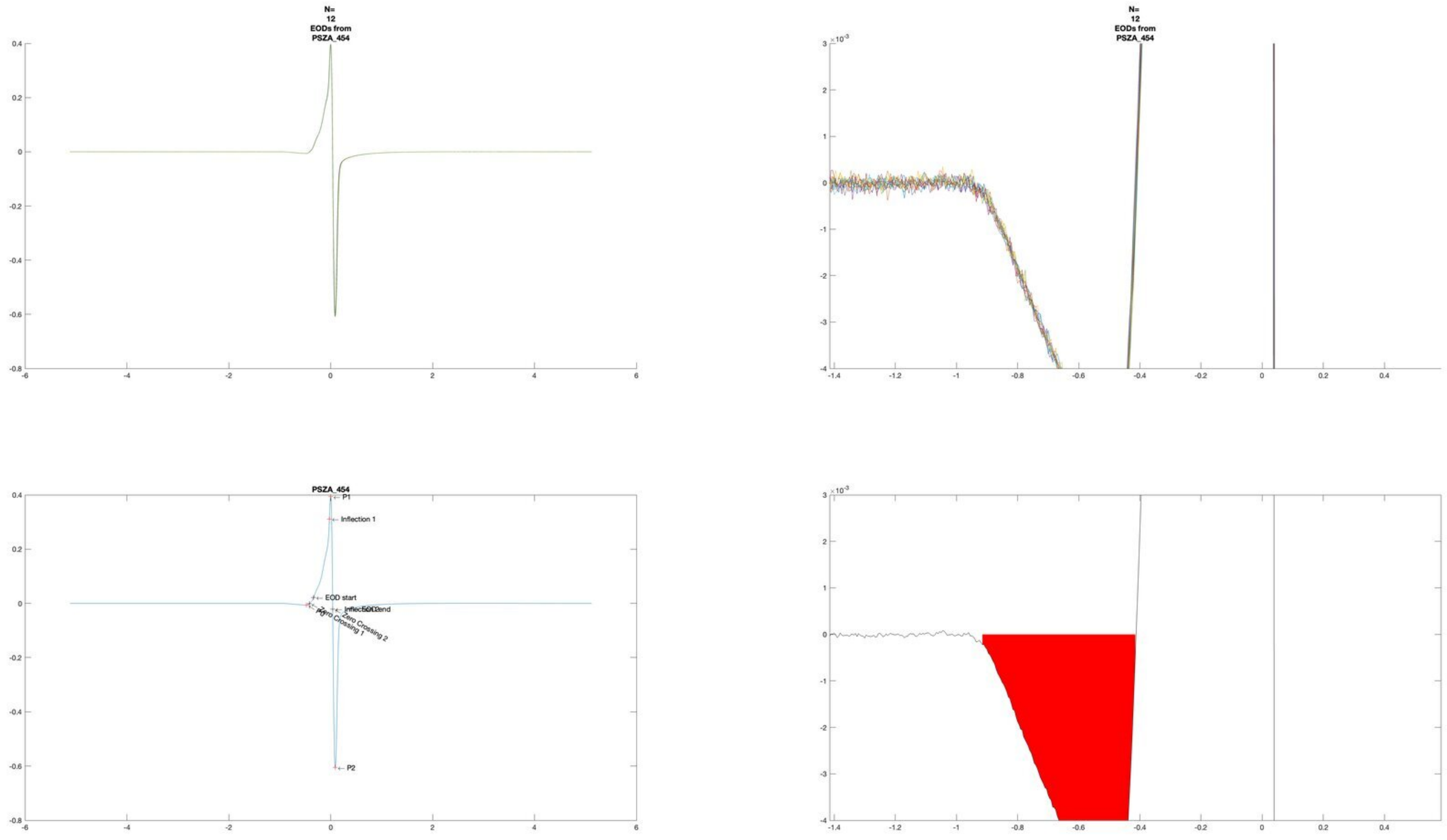
N=
12
EODs from
PSZA_454
N=
12
EODs from
PSZA_454
PSZA_454
P1
Inflection 1
EOD start
Zero Crossing 1
Zero Crossing 2
P2

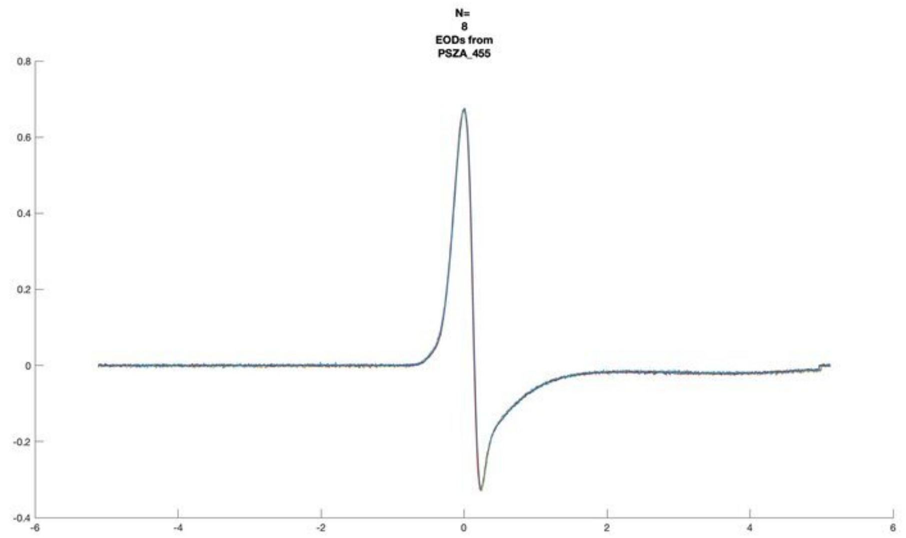
N=
8
EODs from
PSZA_455

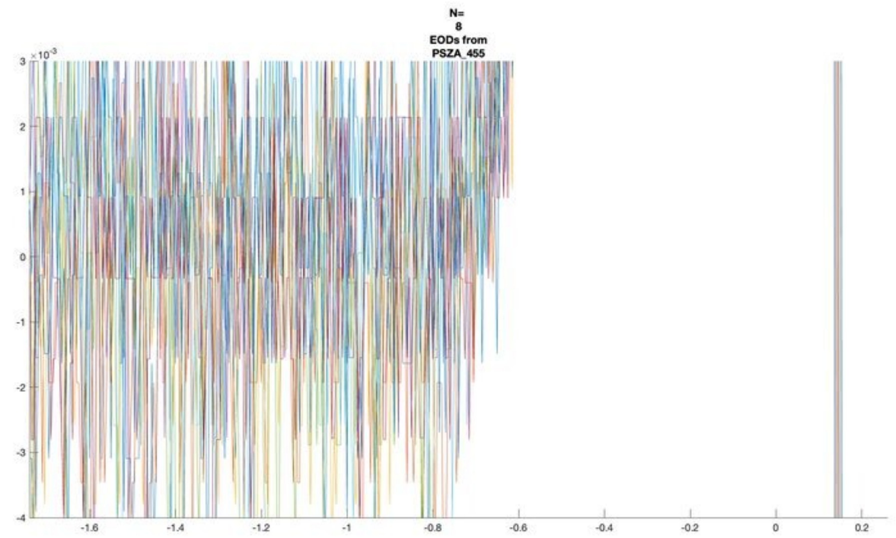
N=
8
EODs from
PSZA_455

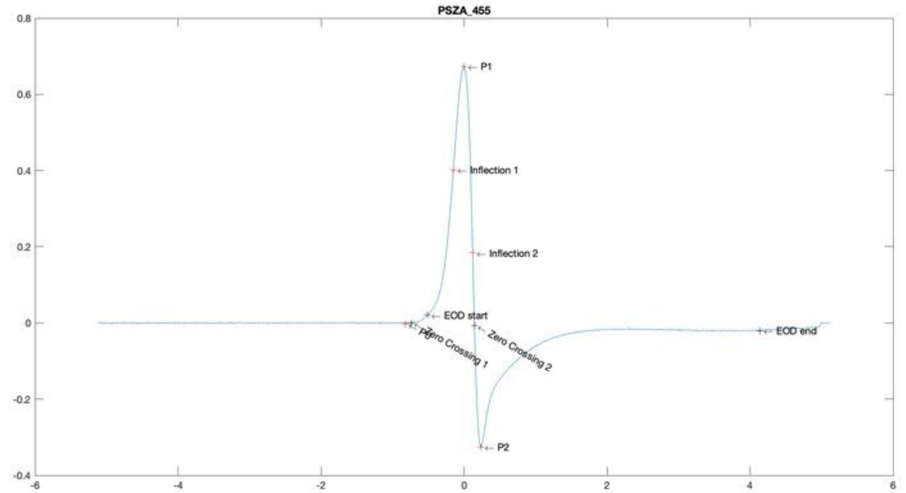
PSZA_455
P1
Inflection 1
Inflection 2
EOD start
Zero Crossing 1
Zero Crossing 2
EOD end
P2

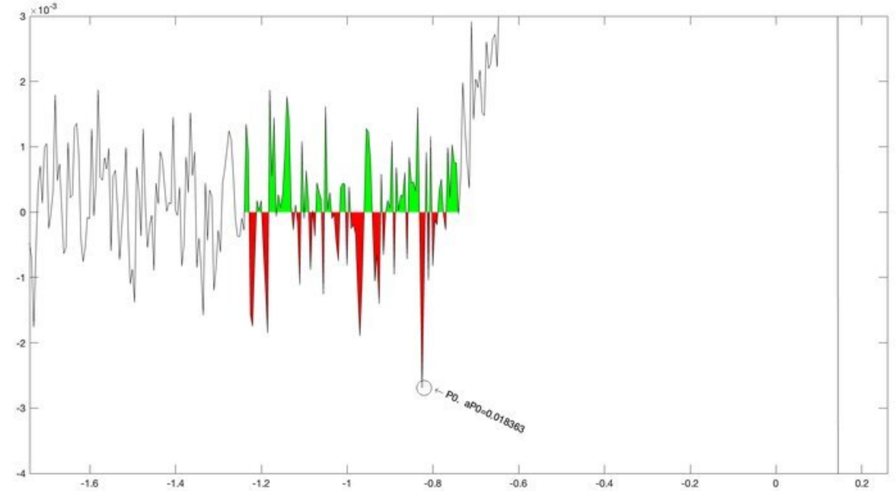
P0, aP0=0.018363

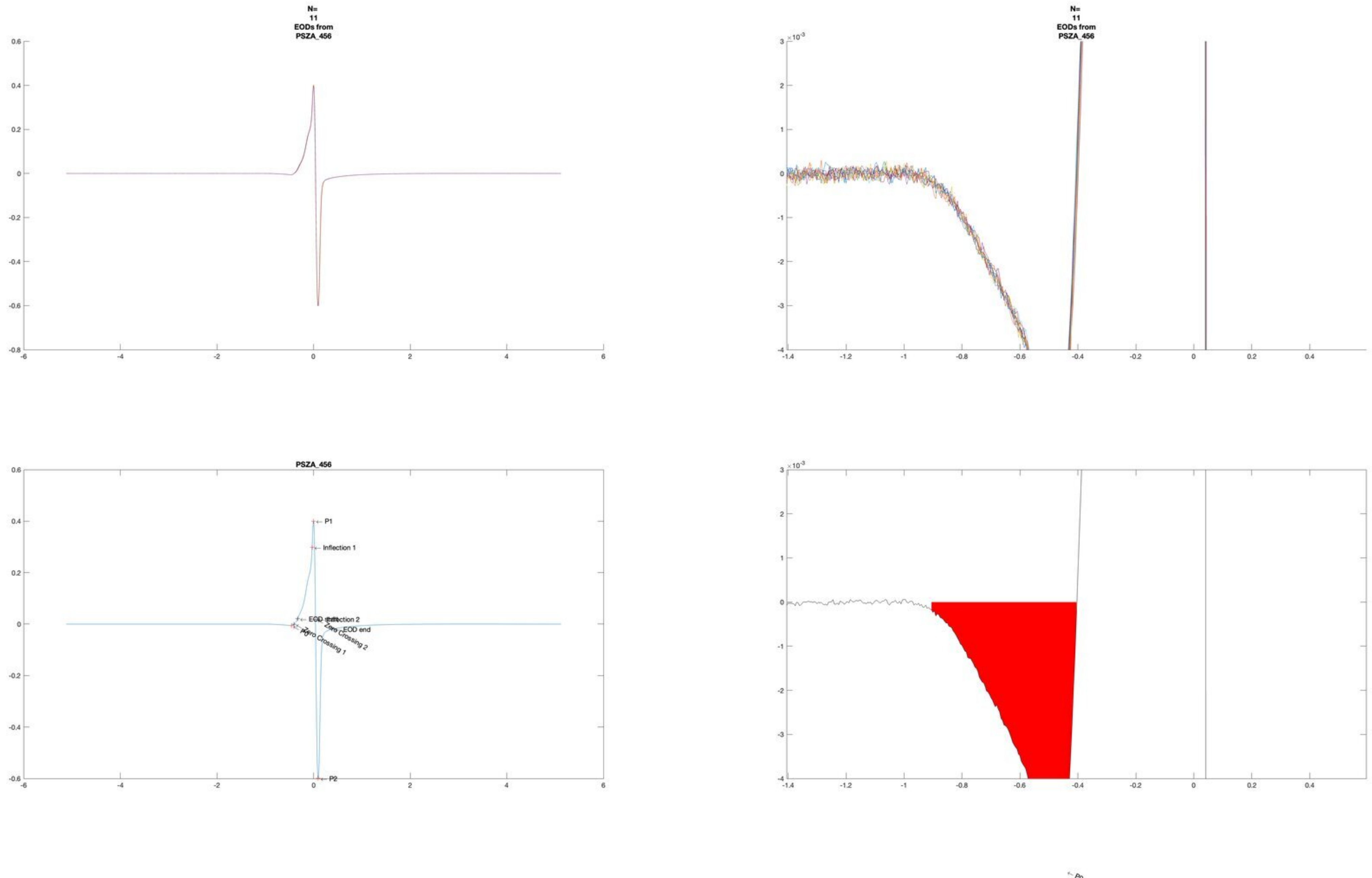

N=
11
EODs from
PSZA_456
N=
11
EODs from
PSZA_456
PSZA_456
P1
Inflection 1
Inflection 2
EOD end
Zero Crossing 2
Zero Crossing 1
P2

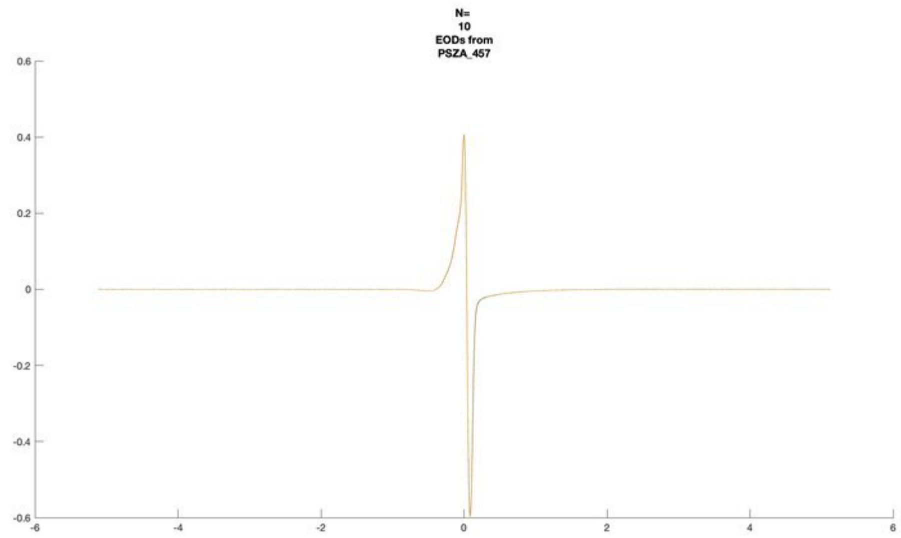
N=
10
EODs from
PSZA_457

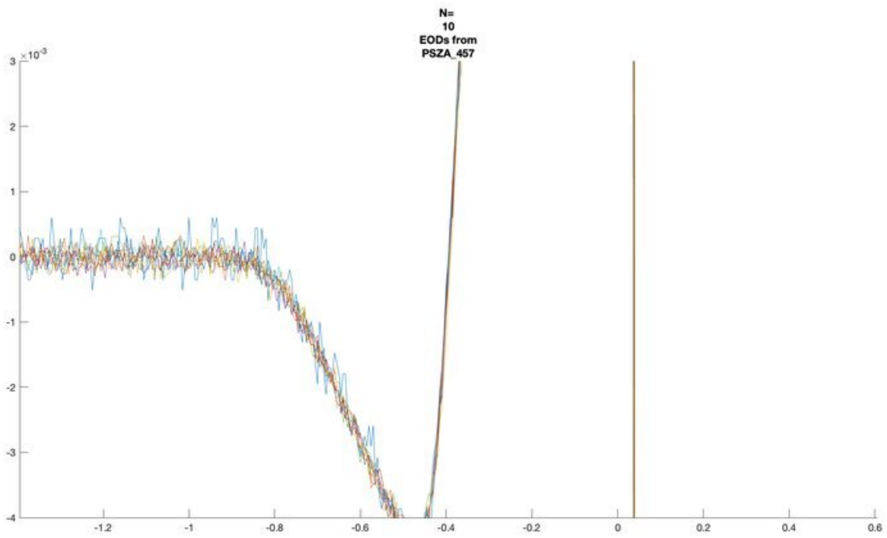
N=
10
EODs from
PSZA_457
×10^-3

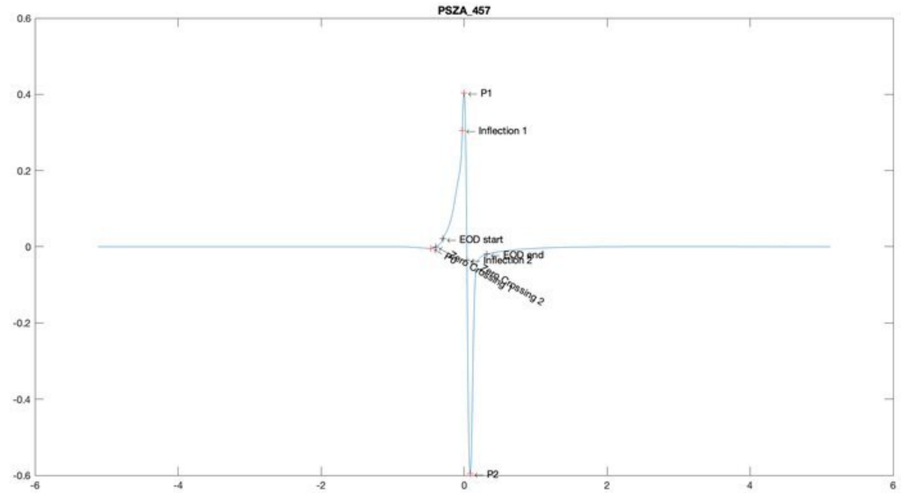
PSZA_457
P1
Inflection 1
EOD start
EOD end
Inflection 2
P0
Zero Crossing 1
Zero Crossing 2
P2

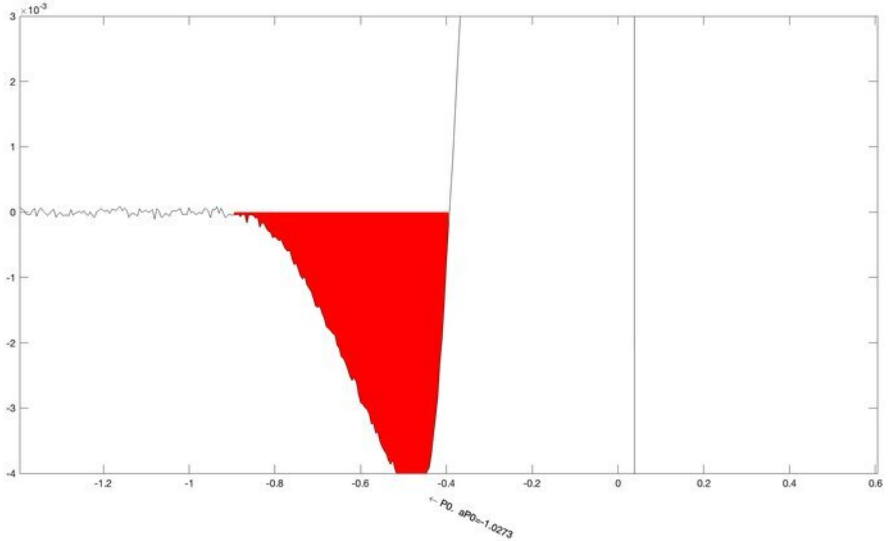
×10^-3
P0, aP0=-1.0273

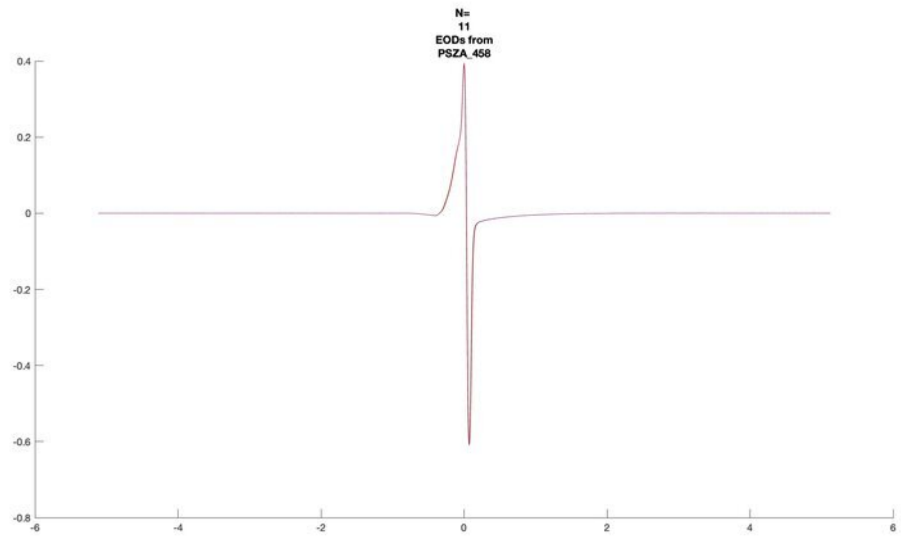
N=
11
EODs from
PSZA_458

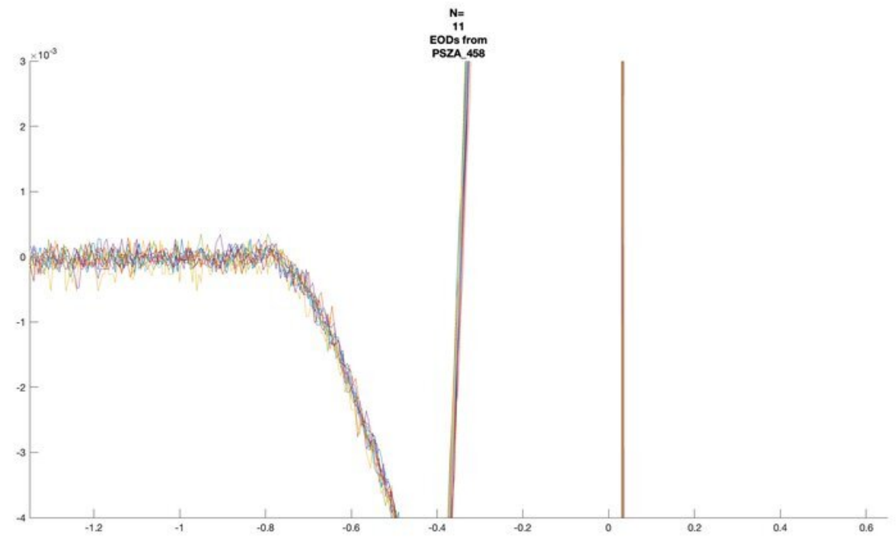
N=
11
EODs from
PSZA_458
×10^-3

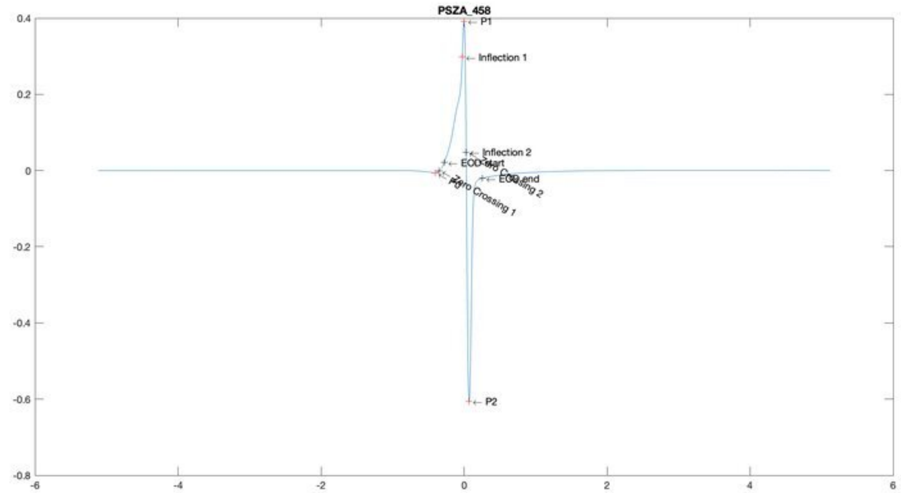
PSZA_458
P1
Inflection 1
Inflection 2
EOD start
EOD end
Zero Crossing 1
P2

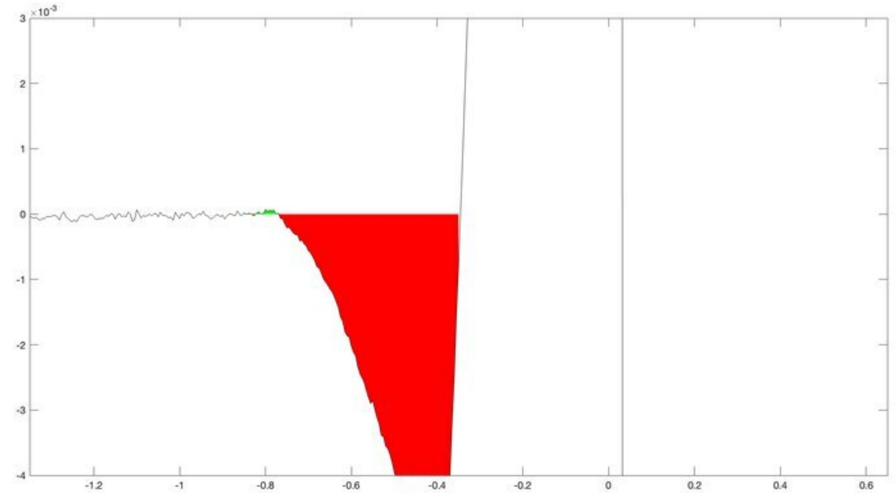
×10^-3

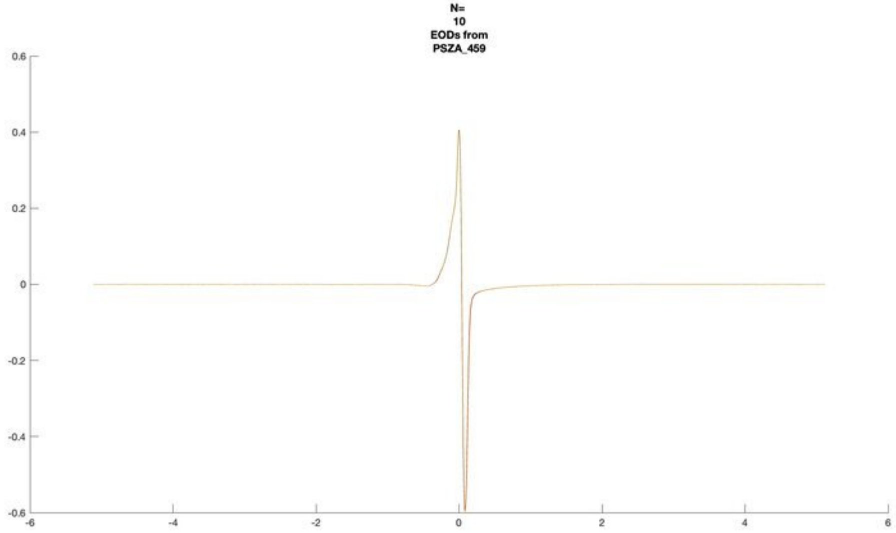
N=
10
EODs from
PSZA_459

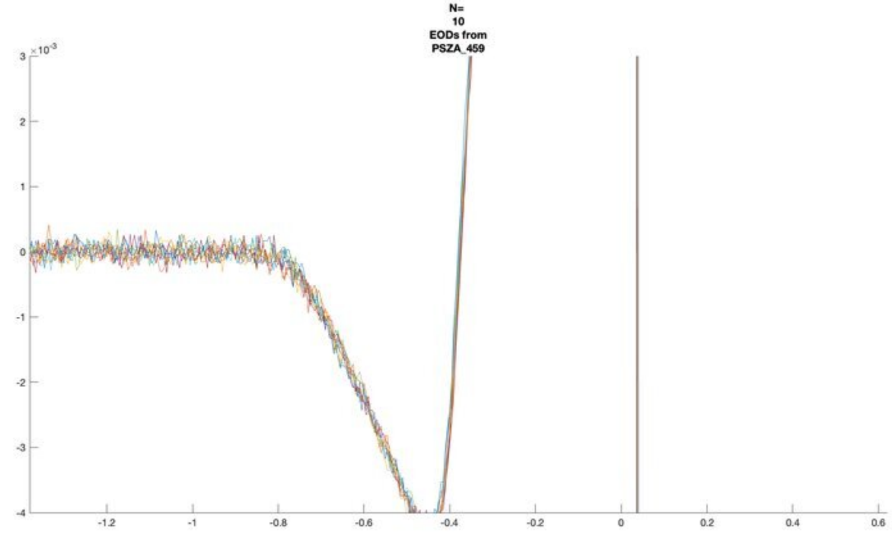
N=
10
EODs from
PSZA_459
×10^-3

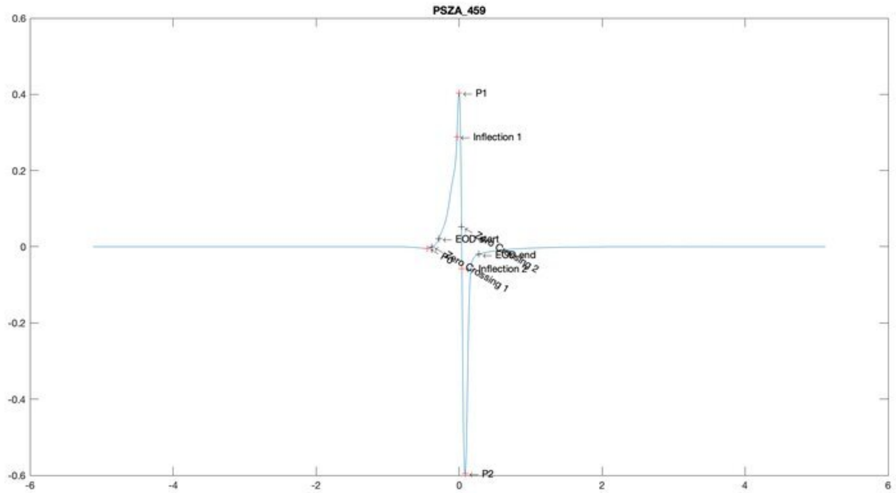
PSZA_459
P1
Inflection 1
EOD start
EOD end
Inflection 2
P0
P2

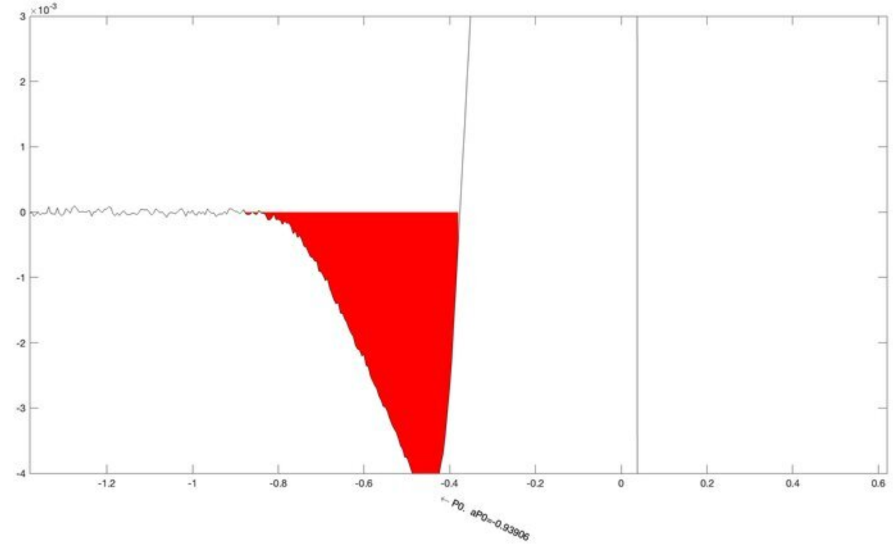
×10^-3
P0, aP0=-0.93906

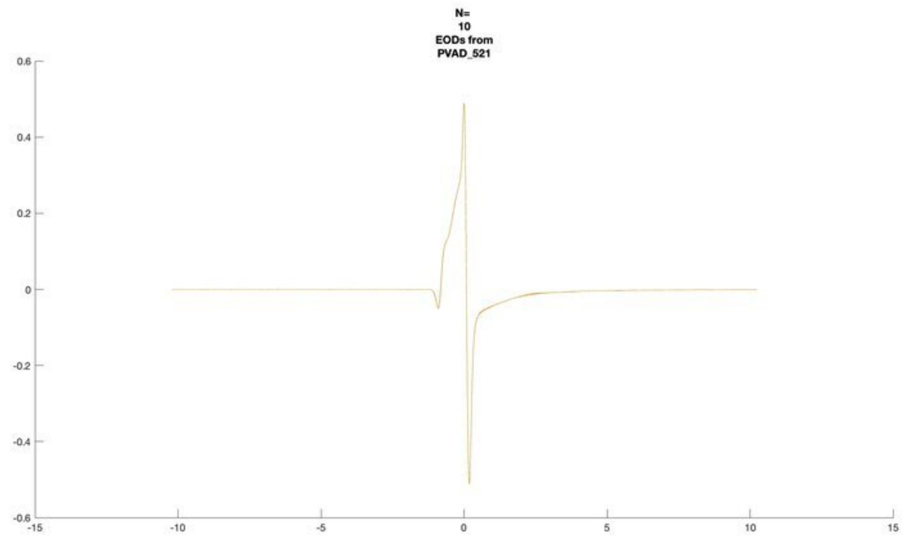
N=
10
EODs from
PVAD_521

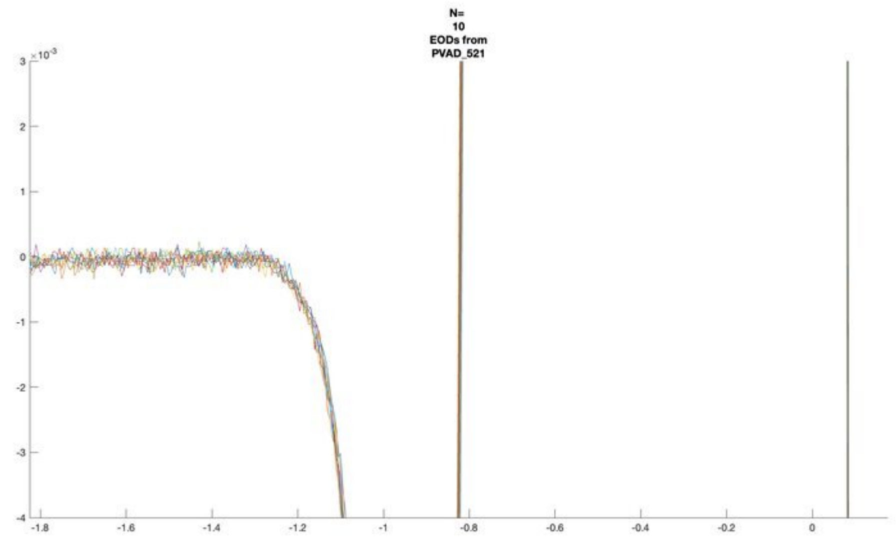
N=
10
EODs from
PVAD_521

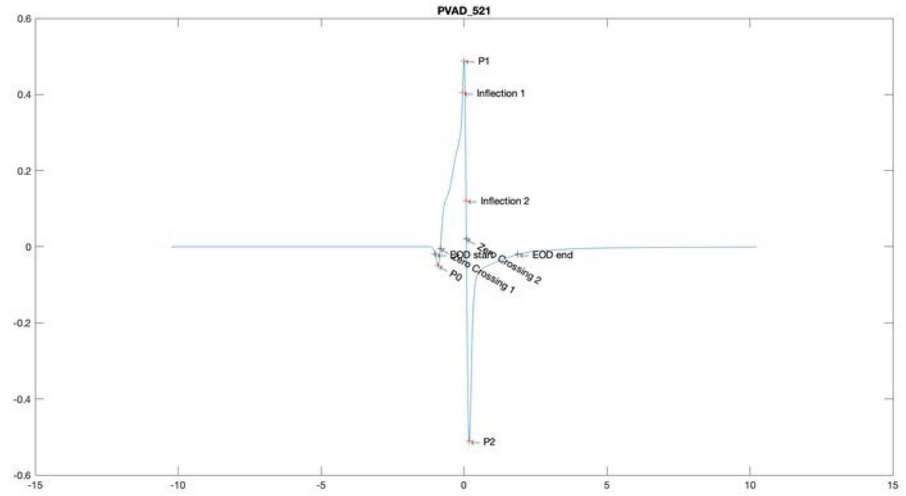
PVAD_521
P1
Inflection 1
Inflection 2
EOD start
Zero Crossing 1
Zero Crossing 2
EOD end
P0
P2

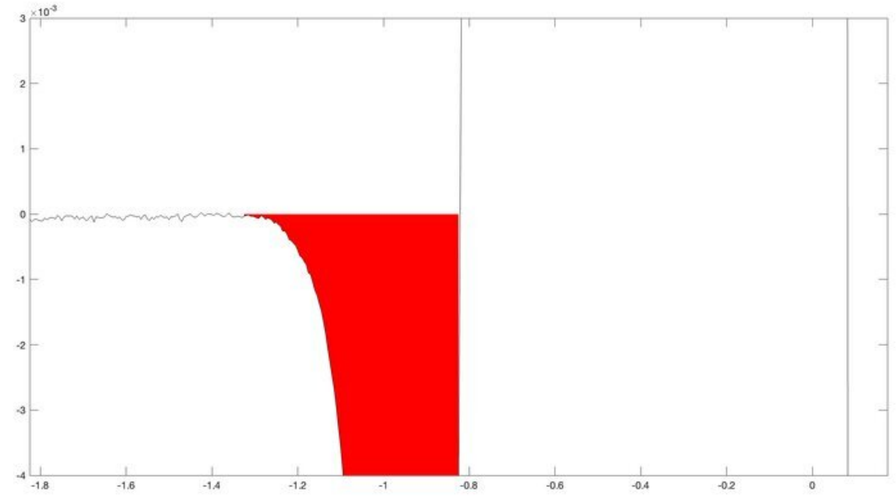

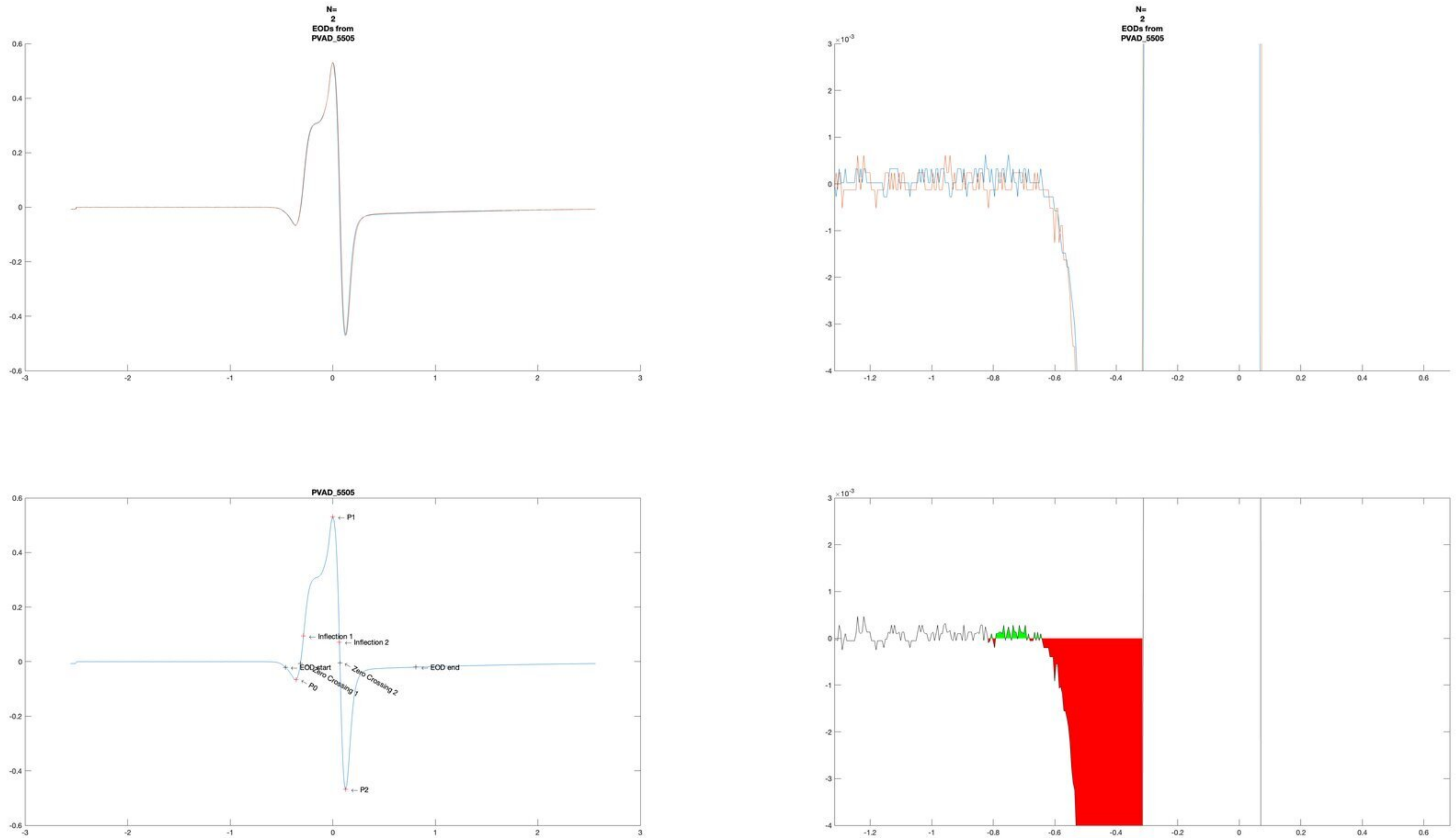
N=
2
EODs from
PVAD_5505
N=
2
EODs from
PVAD_5505
PVAD_5505
P1
Inflection 1
Inflection 2
EOD start
Zero Crossing 1
Zero Crossing 2
EOD end
P0
P2
